# Satiety, TAX-4, and OSM-9 Tune the Attraction of *C. elegans* Nematodes to Microbial Fermentation Products

**DOI:** 10.1101/2025.02.21.639594

**Authors:** Theresa Logan-Garbisch, Emily Fryer, Lara Selin Seyahi, Lucero Rogel-Hernandez, Seung Y. Rhee, Miriam B. Goodman

**Affiliations:** Department of Molecular and Cellular Physiology, Stanford University; Neuroscience Program, Stanford University; Department of Plant Biology, Carnegie Institution for Science

## Abstract

Animals are sensitive to selective pressures associated with nutrient acquisition, underscoring the evolutionary significance of chemosensation in foraging and its intersection with satiety. For the model nematode *Caenorhabditis elegans*, isoamyl alcohol (3-methyl-1-butanol) and 2-methyl-1-butanol are produced by microbial fermentation and present in bacterial food sources collected from the natural environments. Both compounds, which are structural isomers of one another, elicit strong attraction in laboratory settings. Using laboratory chemotaxis assays, we show that starvation attenuates attraction to both compounds. Well-fed *C. elegans* is largely insensitive to the biosynthetic precursors of both alcohols, with the exception of 4-methyl-2-oxovaleric acid, which is a mild repellent. *C. elegans* chemosensation relies on expression of *tax-4* cyclic nucleotide-gated (CNG) and *osm-9* transient receptor potential, vanilloid (TRPV) ion channels and animals lacking both genes are taste- and smell-blind. Animals lacking *tax-4* fail to attract isoamyl alcohol and 2-methyl-1-butanol and those lacking *osm-9* exhibit stronger attraction than the wild-type. Starvation not only attenuates attraction, but also enhances repulsion to 4-methyl-2-oxovaleric acid and uncovers repulsion in *tax-4* mutants absent in their well-fed counterparts. Collectively, these findings implicate satiety in regulating response strength, *tax-4*-dependent chemotaxis in attraction to isoamyl alcohol and 2-methyl-1-butanol, and *osm-9*-dependent chemotaxis in suppressing responses to biosynthetic precursors.

## Introduction

Nematodes co-exist with plants, animals, and microbes and are among the most abundant metazoans in the global rhizosphere (van den Hoogen et al. 2019). The nematode *Caenorhabditis elegans* eats bacteria and is attracted by small molecules synthesized via fermentation by microbes, including isoamyl alcohol (IAA) and 2-methyl-1-butanol (2M1B). Whereas IAA was identified as a *C. elegans* attractant more than 30 years ago (Bargmann et al. 1993), 2M1B was found much more recently (Worthy et al. 2018; Fryer et al. 2024). These compounds are isomers of one another and are among the most potent chemical attractants of *C. elegans* (Fryer et al. 2024). Both compounds are released by bacteria that co-occur with *C. elegans* in the wild and are present in complex mixtures of volatile organic compounds released by these bacteria (Worthy et al. 2018; Siddiqui et al. 2024). For these reasons, it is reasonable to surmise that attraction to IAA and 2M1B serve to enhance the ability of *C. elegans* to navigate toward bacterial food in their environment. Indeed, it has been proposed that attraction to IAA aids *C. elegans* in acquiring leucine, an essential amino acid that is fermented into IAA by bacteria that are known to co-occur with *C. elegans* in nature (Siddiqui et al. 2024).

Behavioral responses to chemicals are typically evaluated in the laboratory using chemotaxis assays that involve placing a population of *C. elegans* in a chemical gradient and observing whether the animals accumulate towards the chemical’s source (display attraction), distribute evenly (display indifference), or accumulate away from the chemical’s source (display repulsion). This behavior relies on chemosensory neurons that detect chemical stimuli and, similar to other animals, *C. elegans* chemosensory transduction depends on the expression of CNG channels or TRPV channels in chemosensory neurons [reviewed in (Ferkey et al. 2021)]. The *tax-4* gene encodes a pore-forming alpha subunit of CNG channels and is expressed in 10 pairs of chemosensory neurons (Komatsu et al. 1996). It was linked to chemotaxis in a forward-genetic screen reported more than 50 years ago (Ward 1973). The *osm-9* gene is also linked to chemotaxis and expressed in 10 pairs of chemosensory neurons, but it encodes a pore-forming subunit of a TRPV-like channel (Colbert et al. 1997). The two genes, *tax-4* and *osm-9*, overlap in 6 pairs of chemosensory neurons (Komatsu et al. 1996; Colbert et al. 1997). *C. elegans* mutants carrying defects in both *tax-4* and *osm-9* are unable to detect most, if not all chemical stimuli (Brissette et al. 2024; Fryer et al. 2024).

In the laboratory, animals are usually well-fed prior to chemotaxis testing and the experimental arena contains a defined chemical gradient, but no bacterial food. Natural environments, by contrast, are complex chemical landscapes in which bacterial food is present, but not spatially uniform and its availability is subject to unpredictable boom-bust cycles (Frézal and Félix 2015). It follows that *C. elegans* and other nematode species are likely to adjust their behavior depending on their satiety state. Indeed, food deprivation is known to alter a wide variety of *C. elegans* behaviors, including movement, feeding, egg-laying, and sensitivity to gases like CO_2_ [reviewed by (Baugh and Hu 2020)]. It also attenuates sensory guided-behaviors that enable *C. elegans* nematodes to navigate complex environments, including thermotaxis (Hedgecock and Russell 1975) and chemotaxis (Urushihata et al. 2016; Shimizu et al. 2019).

The fermentation pathways that yield isoamyl alcohol and 2-methyl-1-butanol in yeast have been studied for more than a century. This pathway (also known as Ehrlich degradation) begins with an amino acid, which is converted to a keto acid, decarboxylated to an aldehyde, and then reduced to a primary alcohol, which is excreted [reviewed in (Hazelwood et al. 2008)]. The starting materials for IAA and 2M1B biosynthesis are leucine and isoleucine, respectively, and there is evidence that at least one of the precursor compounds is released by bacteria that co-occur with *C. elegans* in the wild (Worthy et al. 2018; Siddiqui et al. 2024). These alcohols are also synthesized by several plants including corn (Flath et al. 1978), pineapple (Flath and Forrey 1970; Pino 2013; George et al. 2023), and banana (Myers et al. 1970), suggesting that they are common constituents of complex natural products. Both of these alcohols and acetate compounds derived from them contribute to the flavor profile of fermented foods and beverages [e.g. (Matheis et al. 2016)], suggesting that human chemosensory neurons, like those of *C. elegans*, are activated by these compounds.

Using high-throughput laboratory assays and genetic dissection, we tested the idea that *C. elegans* would preferentially detect the excreted microbial fermentation products IAA and 2M1B and that attraction to these markers of bacterial food would differ in sated and starved animals. Consistent with this expectation, we found that well-fed, sated wild-type animals are insensitive to five of the six biosynthetic precursors we tested and that sated animals are nearly 100-fold more sensitive to IAA and 2M1B than their starved counterparts. One biosynthetic precursor, the keto acid derived from isoleucine (4-methyl-2-oxovaleric acid), functioned as a mild repellent in well-fed wild-type animals. All responses were abrogated in fed mutants lacking the TAX-4 chemosensory CNG channel protein. Unexpectedly, mutants lacking the OSM-9 chemosensory TRPV channel were attracted to all six precursor compounds when well-fed. Similar to wild-type responses to IAA and 2M1B, starving *osm-9* animals reduced the attraction seen in their well-fed counterparts. These findings imply that *C. elegans* accurately detects fermentation products released to the environment and that behavioral indifference to a given compound is not equivalent to a lack of an ability to detect that compound under all conditions.

## Materials and Methods

### Chemical resources

We used serial dilution to generate solutions containing isoamyl alcohol (IAA, CAS No. 123-51-3) and 2-methyl-1-butanol (2M1B, CAS No. 137-32-6), at concentrations between 3.0 µM and 3.0 M in dimethyl sulfoxide (DMSO). The most concentrated solutions (1.0 M and 3.0 M) were generated by diluting IAA (9.2 M; MW 88.15; density 0.809 g/mL)(National Center for Biotechnology Information 2004a) and 2M1B (9.3 M; MW 88.15; density 0.815 g/mL)(National Center for Biotechnology Information 2004b), respectively, in DMSO. With this procedure, we generated 13 solutions covering 3.0 µM to 3.0 M in half-log increments.

The biosynthetic precursors of IAA are: L-leucine (I1), 4-methyl-2-oxovaleric acid (I2), isovaleraldehyde (I3). Those leading to 2M1B pathway are similarly abbreviated: L-isoleucine (M1), 3-methyl-2-oxopentanoic acid (M2), 2-methylbutyraldehyde (M3). For convenience, these compounds are referred to by these abbreviations, which are related to their position in the fermentation pathways. Acidic compounds like I1, I2, M1, and M2 generate a source of low pH in chemotaxis arenas, raising the possibility that they elicit acid avoidance (Sambongi et al. 2000) and this might overshadow responses to the compounds. To mitigate this potential confound, we dissolved all precursor compounds in ultrapure water containing HEPES (10mM), a strong pH buffer with a *p*Ka of 7.55. HEPES-buffered water is neither an attractant nor a repellent (See S1 Fig). We used IAA precursor compounds at 30mM and 2M1B precursors at 100mM. We purchased all chemicals from Sigma-Aldrich, except for anhydrous DMSO (Thermo Fisher) and stored aliquots of stock solutions at −20°C.

### Chemotaxis Assays

We performed chemotaxis assays in four-well assay plates following the workflow described by Fryer (Fryer et al. 2024) with four modifications. First, test compounds dissolved in HEPES-buffered water were placed on one end of the plate and the HEPES buffer alone on the other end of the plate served as the reference. As in our prior work (Fryer et al. 2024), test compounds that were dissolved in DMSO used DMSO as the reference. Second, we washed animals from growth plates in chemotaxis buffer, which contained (Lim et al. 2018; Fryer et al. 2024): potassium phosphate buffer (pH 6, 5 mM), MgCl_2_ (1 mM), CaCl_2_ (1 mM). Third, we delivered animals to chemotaxis arenas using a hand-held electronic repeat-dispensing micropipettor (Maestro, CAPP, Nordhausen, Germany) in place of a liquid handler. As previously described (Fryer et al. 2024), animals were suspended in a 7:3 mixture of chemotaxis buffer and iodixonal (Optiprep™, Sigma). (Iodixonal is a non-toxic polymer that increases the density of the buffer, preventing animals from settling at the bottom of holding tubes.) Fourth, immediately prior to imaging, we dispersed animals by tapping assay plates (vertical motion, keeping the plate flat) on the benchtop (2-3 times).

All of our data was collected in a blinded or masked fashion. Animal genotypes and compound identity were masked by a member of the lab who was not involved in performing the experiments. When feasible, we masked compounds in bulk prior to separation into aliquots and tested well-fed and starved animals in separate experimental sessions, and these compounds were not unmasked until we had acquired all replicates using that key.

### *C. elegans* strains and husbandry

We used four *C. elegans* strains: 1) N2 (Bristol) RRID:WB-STRAIN:WBStrain00000001; 2) GN1077 *tax-4(pr678)* III; *osm-9(ky10)* IV; 3) PR678 *tax-4(pr678)* III RRID:WB-STRAIN:WBStrain00030785; and 4) CX10 *osm-9(ky10*) IV RRID:WB-STRAIN:WBStrain00005214. The N2 strain served as the wild-type. GN1077 animals carry null alleles of *tax-4* and *osm-9* and are insensitive to most compounds (Fryer et al. 2024). Unless otherwise noted, we grew animals at 20°C and prepared age-synchronized, well-fed young-adult animals as described (Fryer et al. 2024). We starved age-synchronized animals by washing them from food-bearing growth plates ∼18 hours prior to chemotaxis testing and removing residual bacteria *via* 2-8 cycles of centrifugation and resuspension in chemotaxis buffer, stopping the process when the supernatant appeared free of debris. Animals were then transferred to 3% (w/v) agar nematode growth medium plates lacking bacteria, which reduces burrowing observed on standard 2% (w/v) agar nematode growth medium plates and enhances retrieval of the worms ∼18 hrs later. The *osm-9;tax-4* double mutants exhibit a developmental delay, taking approximately 1 day longer to reach adulthood than wild-type worms. We accounted for this in our husbandry strategy, but cannot exclude the possibility that these mutants were slightly younger than wild-type when transferred from food-bearing plates to the sterile 3% plates.

### Data analysis and statistics

We analyzed our data for chemotaxis assays as previously described (Fryer et al. 2024). In brief, we used the Our Worm Locator (OWL) software (https://github.com/Neuroplant-Resources/Neuroplant-OWL) to determine individual worm locations and to generate a *.csv file of these data to be pooled across technical and biological replicates. We excluded individual replicates containing fewer than 150 animals, an action that decreases the variance (Fryer et al. 2024). Our calculations for the mean of each replicate and the bootstrapped calculation of the differences of the means utilized the same workflow as our previous work (Fryer et al. 2024). Responses were considered significant if the 95% confidence interval of the bootstrapped mean differences excluded zero. This is equivalent to achieving *p*<0.05 in conventional hypothesis testing. We used the calculated differences of the mean position to generate dose-response curves for IAA and 2M1B (see ‘Chemical resources’ for concentrations). We fit these data to a single-site binding curve to find the effective concentration at half-maximal response 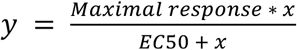 where *y* = average difference in mean position and *x* = compound concentration at the source. The correlation coefficient (*r*^2^) was calculated as follows: 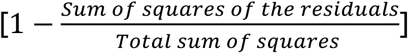.

### Code and Data availability

All code used to convert images into worm positions and estimation plots (Fryer et al. 2024) are publicly available at https://github.com/Neuroplant-Resources/Neuroplant-OWL. Code for dose-response curve fitting is available at: https://github.com/wormsenseLab. Raw (images) and processed data will be deposited in a general purpose repository.

## Results

### IAA and 2M1B attract wild-type worms

Isoamyl alcohol (IAA) and 2-methyl-1-butanol (2M1B) strongly attract well-fed, wild-type animals and were among the most effective attractants in a screen of 90 small molecules (Fryer et al. 2024). IAA and 2M1B are structural isomers of one another and are synthesized by microbes (Morgan et al. 1966; Dickinson et al. 2000) as a result of fermentation (de la Plaza et al. 2004). Figures 1A and 1B show images of typical responses of well-fed, wild-type animals to DMSO (solvent control), IAA, and 2M1B. Enlarged images reveal substantial aggregation of worms near the point sources of both alcohols, but not the solvent. We used software to determine worm position within the chemical gradient and plotted both the position of individual worms and the mean location across eight replicates in each condition (Fig. 1C, 1D). Using estimation statistics software, we calculated the difference of the mean locations (<Δ location>) for each test condition relative to control. As expected for strong attractants, the change in mean location is both statistically significant and displays a large effect size, 12.98 mm for IAA (Fig. 1C) and 11.61 mm (Fig. 1D) for 2M1B. These findings are consistent with our prior work and with the finding that IAA and 2M1B are among the strongest attractants known for wild-type *C. elegans* (Fryer et al. 2024).

**Figure 1.**
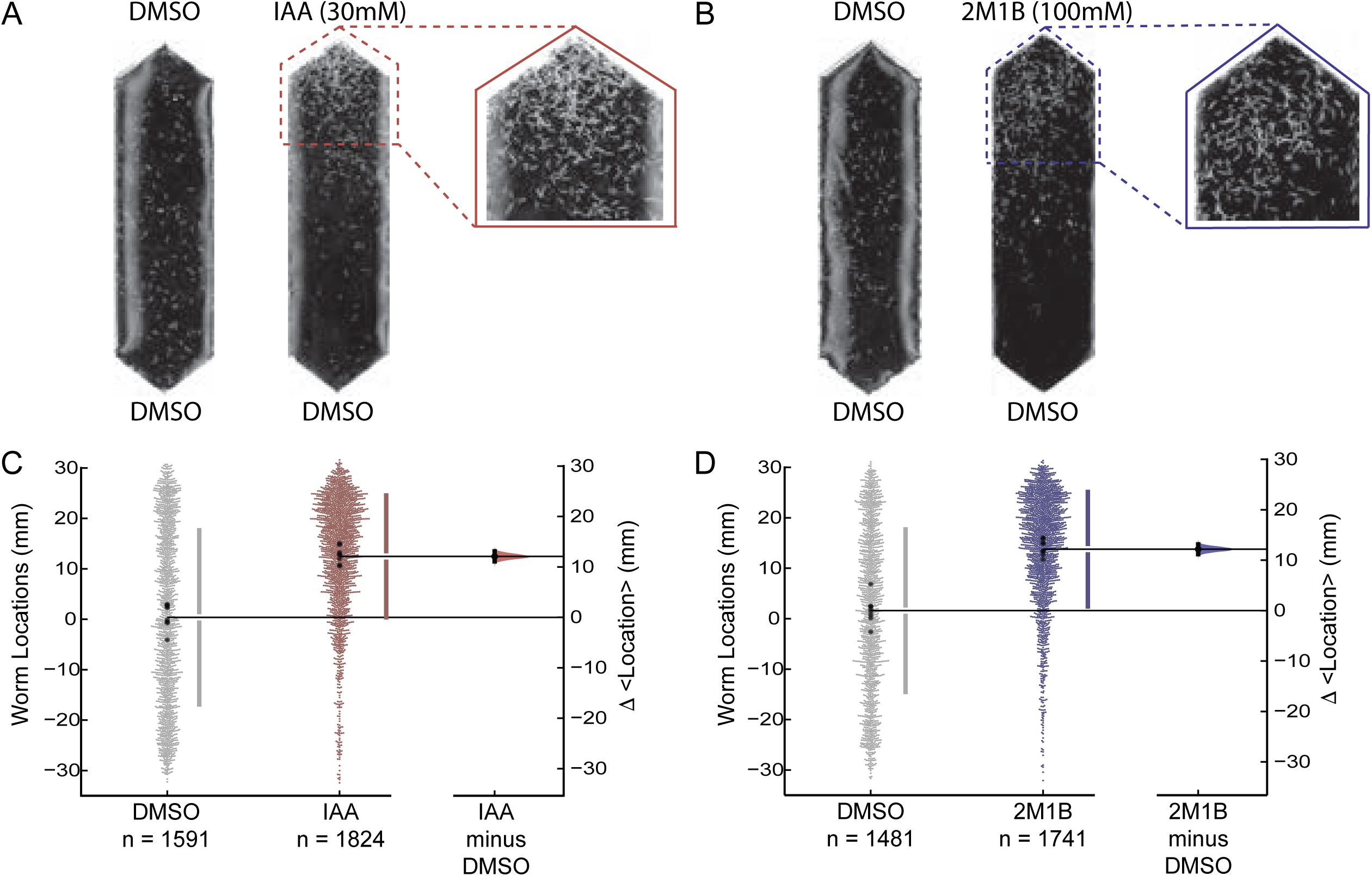
IAA and 2M1B attract well fed wild-type worms. **A, B.** Images of chemotaxis arenas comparing representative responses to the solvent control (left) and accumulation near test compounds (right). An enlarged image of the region near each test compound is also shown, IAA (**A**) and 2M1B (**B**). **C, D.** Swarm plots depicting the locations of individual worms (left), aggregated across all replicates (*N* = 8, all conditions) for solvent control (gray) and test compounds (30 mM IAA, red, **C**, and 100 mM 2M1B, blue, **D**). Each small dot (gray, red, or blue) represents a single worm and the large dots (black) show the mean position of a single replicate. The total number animals, *n*, is given below each dataset. The difference in the mean location in control vs. test conditions (<Δ location>) was computed using bootstrapping and reported as the distribution plotted on the right-hand side of each plot. The data in this figure are a subset of the data presented in Figures 2 and 3.

To learn more about the strength of the response to IAA and 2M1B, we tested wild-type worms across six orders of magnitude in source concentration (Fig 2 and Fig 3). While IAA dose-response studies have been reported previously (Bargmann et al. 1993), prior work covered only three orders of magnitude and no such information exists for 2M1B. Thus, we sought both to extend knowledge of IAA dose-response relationships and to determine dose-response relationships for 2M1B using our high throughput technique.

**Figure 2.**
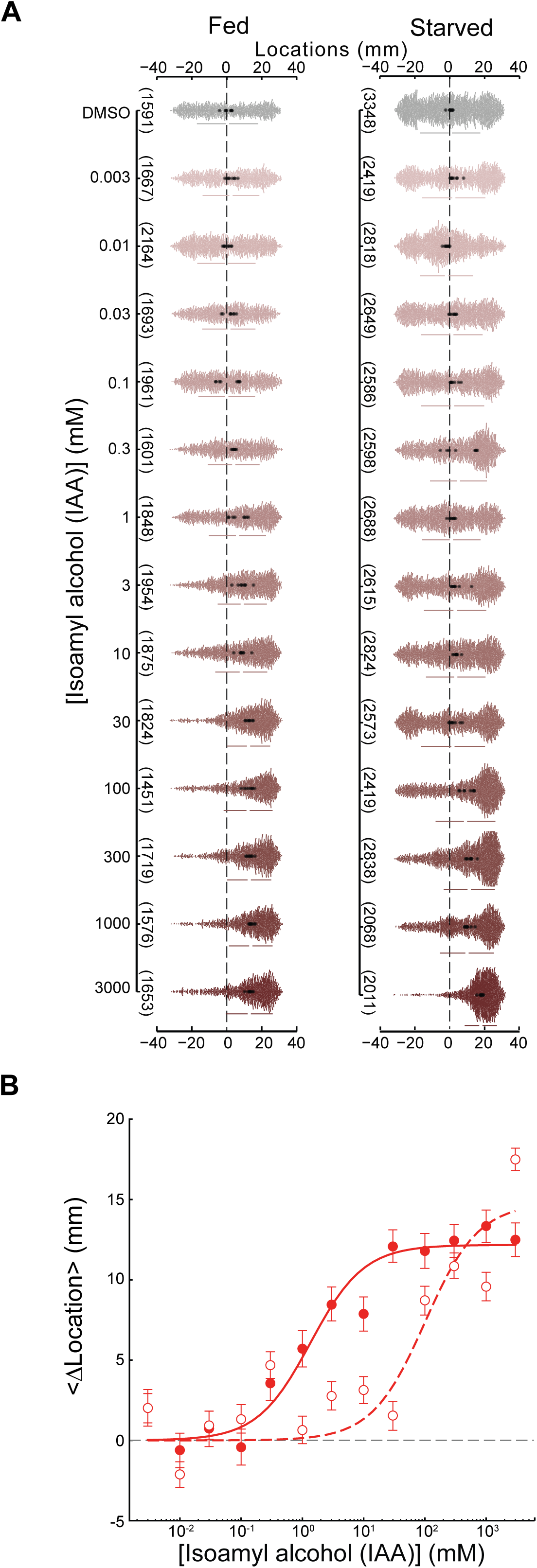
Starvation attenuates attraction to IAA by ∼75-fold. **A**. Response to [IAA] as a function of concentration at the source position (+30mm) in well-fed (left) and starved (right) worms. Swarm plots consist of the position of individual worms (small dots) pooled across 8 biological replicates (large dots are the mean location per replicate). The total number of animals tested is given in parentheses (left) and the black dashed lines indicate the initial starting position of the worms. **B**. Dose-response plot for fed (filled circles) and starved (open circles) animals. Points are <Δ location> values, and error bars are the 95% confidence interval. Smooth lines were fit to the data using the formula 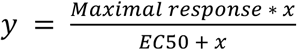 and the *EC*_50_ values were 1.36 mM and 100.97 mM for fed (solid line) and starved (dashed line) conditions, respectively. Supplemental Table Figure 2 contains the numerical values plotted in these graphs.

**Figure 3.**
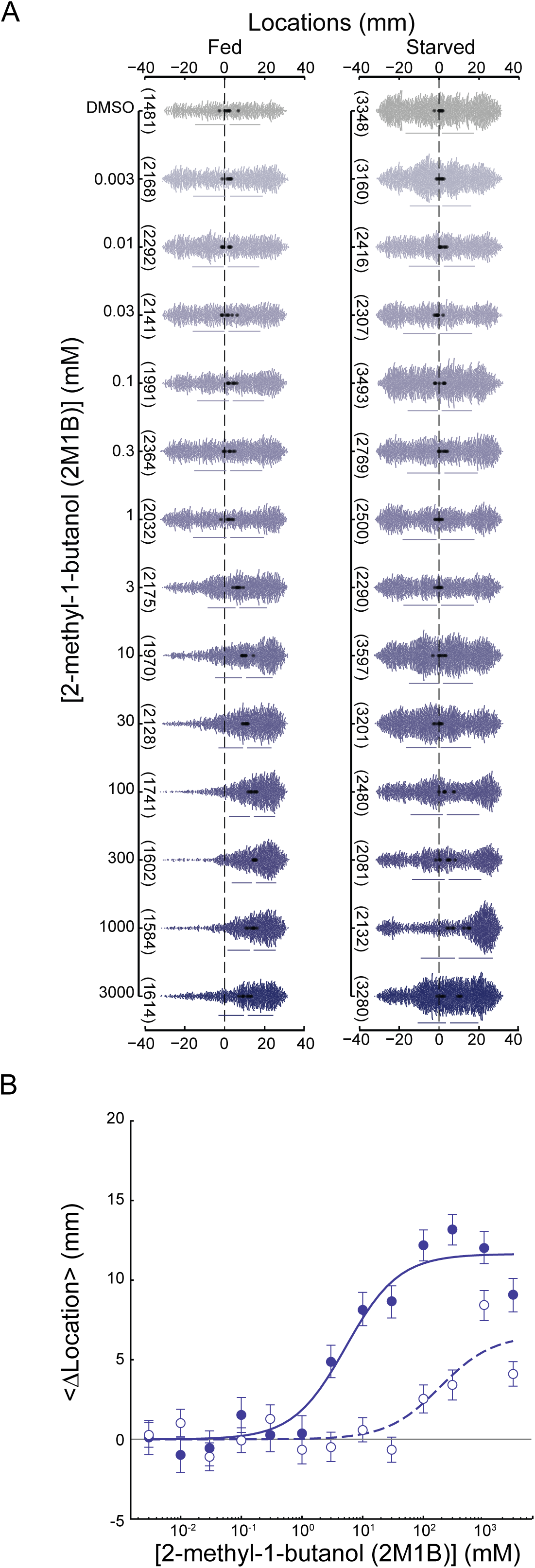
Starvation attenuates attraction to 2M1B by ∼35-fold. **A.** Response to [2M1B] as a function of concentration at the source position (+30mm) in well-fed (left) and starved (right) worms. Swarm plots consist of the position of individual worms (small dots) pooled across *N=*8 biological replicates (large dots are the mean location per replicate). The total number of animals tested is given in parentheses (left) and the black dashed lines indicate the initial starting position of the worms. The plot for the reference condition (DMSO) is replotted from Fig 2A (right), since the assays for both compounds were conducted in tandem. **B**. Dose-response plot for fed (filled circles) and starved animals (open circles) points are <Δ location> values, and error bars are the 95% confidence interval. Smooth lines were fit to the data using the formula 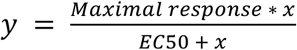 and the *EC*_50_ values were 5.29 mM and 186.93 mM for fed (solid line) and starved (dashed line) conditions, respectively. Supplemental Table Figure 3 contains the numerical values plotted in these graphs.

Given the unpredictable availability of food in natural environments for *C. elegans*, we also examined how starvation affected attraction to IAA and to 2M1B. The data include the samples shown in Fig 1.

### Starvation attenuates attraction to IAA and 2M1B

Although animals first experience compounds at the starting zone of the arena (∼30 mm from the source), we assume that the concentration at the start zone is proportional to the concentration at the source position in analyzing the concentration-dependence of their response. Thus, we use dose to refer to the concentration of each molecule deposited at the source position at the center of our ∼60mm-long assay arena.

As expected, well-fed wild-type worms show a dose-dependent attraction to IAA (Fig 2A, left, and 2B, solid lines and filled circles). Our findings are consistent with prior work (Bargmann et al. 1993), which indicates that well-fed animals placed ∼30 mm from the source can detect sub-millimolar IAA (1/10,000 dilution or 0.919 mM) and that the response saturates above a 10 mM source concentration (1/1000 dilution or 9.19 mM). Our data also show that starved wild-type worms are less sensitive to IAA (Fig 2A, right, and 2B, dashed lines and open circles) than their well-fed counterparts. This is clearly evidenced by the nearly 70-fold increase in the effective concentration for half-maximal response (*EC*_50_) (Fig 2B), even though starved animals respond in a dose-dependent fashion and eventually achieve similar strength.

In addition to exploring the chemotactic response to the expanded range of concentrations for IAA, we also investigated the behavior of both well-fed and starved worms to 2M1B across six orders of magnitude.

Similar to IAA, starvation attenuated attraction to 2M1B (Fig. 3). The effect of starvation on attraction to 2M1B was weaker than its effect on attraction to IAA, however. Specifically, the EC_50_ value for starved animals is approximately 30-fold higher than that found for well-fed animals (Fig 3). Whether well-fed or starved, wild-type worms were more strongly attracted to IAA (Fig 2) than to 2M1B (Fig 3).

### Biosynthetic precursors evoke minimal responses in well-fed wild-type animals

Structurally related compounds, such as those that comprise a microbial biosynthetic pathway might induce similar behavioral responses. Alternatively, animals might tune their sensitivity to the compounds that are excreted. The fermentation pathways studied here provide an opportunity to distinguish between these possibilities. Accordingly, we tested the chemotaxis behaviors of wild-type worms in the presence of these compounds at concentrations reflective of the saturating responses of the endpoint alcohols which still evoked responses in starved animals, although they were much weaker (30 mM for IAA, 100 mM for 2M1B).

The fermentation pathways of L-leucine and L-isoleucine to IAA and 2M1B respectively (Dickinson et al. 1997; Dickinson et al. 2000; Caspi et al. 2020) use the same biochemical reaction steps (Fig 4, below the heat map). For these experiments, we examined the behavioral response via chemotaxis to the precursor compounds as well as the end-product alcohols in 10 mM HEPES-buffered water in order to avoid the potential confound of acid-avoidance to the more acidic precursors (see Methods).

**Figure 4.**
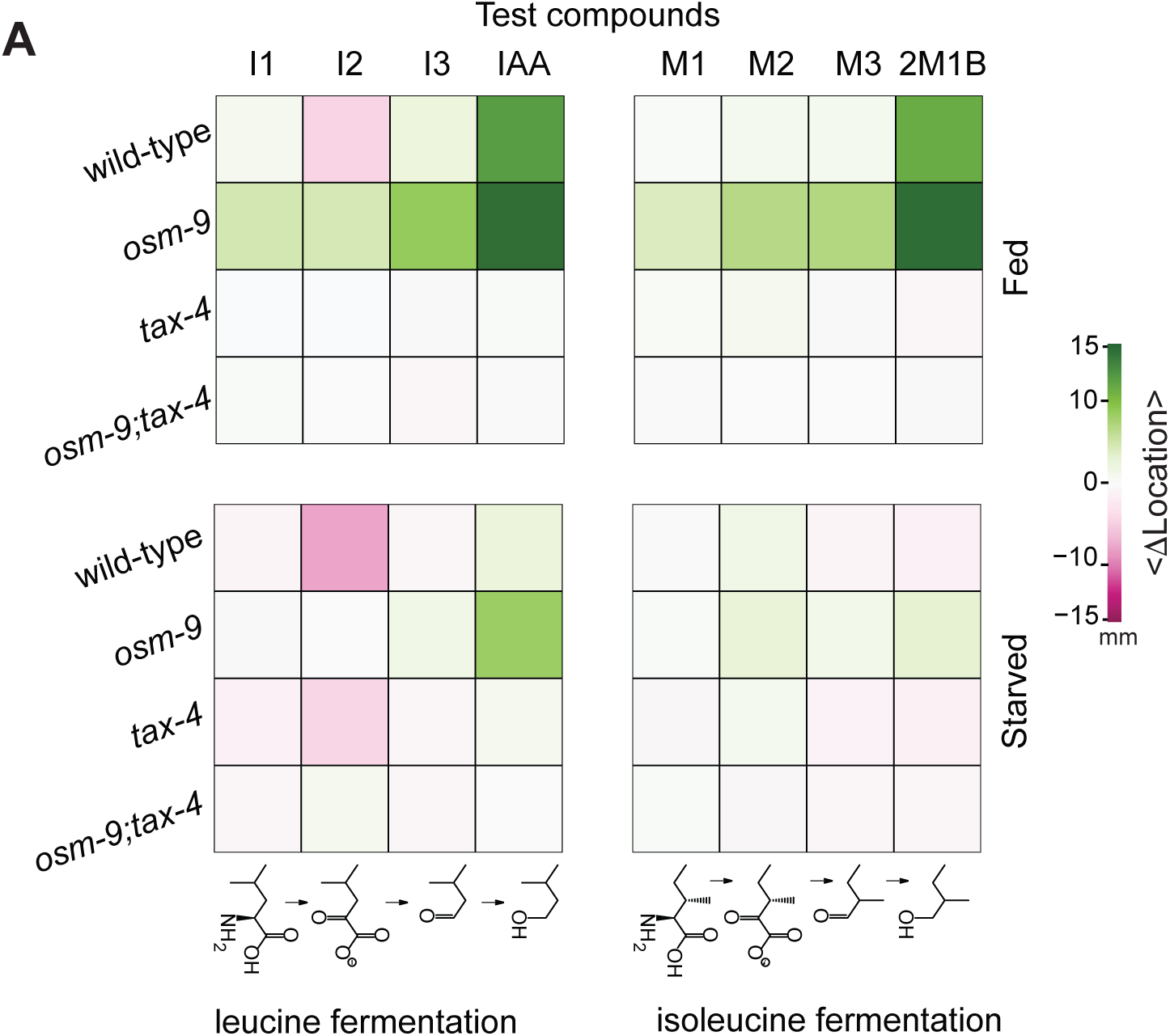
Attraction to the biosynthetic precursors of IAA and 2M1B is *tax-4*-dependent and starvation reveals an *osm-9*-dependent repulsion. A heat map representation of the response to the fermentation pathway compounds, segregated by physiological condition (fed, top; starved, bottom), fermentation pathway (isoleucine, left; leucine, right), and genotype. Green represents a change in location closer to the compound (attraction), while magenta represents a change in location farther from the test compound (repulsion), and the stronger responses for both correlate to more saturated hues. Supplemental Table Figure 4 contains the numerical values for the bootstrapped average difference of the mean position relative to responses to solvent controls represented in the heat map.

When well-fed, wild-type worms are indifferent to I1, I3, and all 2M1B precursors, but moderately repelled by I2 (<Δ location> = −4.45 mm). Both IAA and 2M1B in HEPES buffer elicit strong attraction (Fig 4, top row, S2 Fig, panel A), as found when DMSO is the solvent (Fig 1, Fig 2, and Fig 3) (Fryer et al. 2024). These findings show that responses to IAA and 2M1B are independent of the solvent used.

### Genetic dissection of chemosensory transduction

To gain insight into which chemosensory signaling pathways were involved in responding to these eight compounds, we tested mutants defective in the expression of *tax-4*, *osm-9*, or both chemosensory transduction channel genes. We were interested in seeing how the signaling pathways (if any) used for the precursors (I1-I3, and M1-M3) compared to those for the final fermentation products (IAA, 2M1B). Loss of *osm-9* alone enhanced sensitivity by more than 5-fold, on average, to all eight compounds relative to wild-type animals in the well-fed condition (Fig 4, top panels, and S2 Fig, panel B). Loss of *tax-4*, but not *osm-9*, abolished all responses in well-fed mutants (Fig 4, top panels, and S2 Fig, panel C). Additionally, the *tax-4; osm-9* double mutants are indifferent to all compounds (Fig 4, top panels, and S2 Fig, panel D), verifying that the response to these compounds depends on chemosensory transduction.

In wild-type animals, starvation attenuates attraction and enhances the repulsion to I2 (Fig 4, bottom panels and S3 Fig, panel A). For starved *osm-9* mutants, we observe a similar attenuation of attraction to all compounds, and no repulsion appears (Fig 4, bottom panels and S3 FigB). On the other hand, when starved, *tax-4* mutants weakly avoided I2 (Fig 4, bottom panels and S3 Fig, panel C), but remained indifferent to all other compounds. The starved *osm-9; tax-4* animals behaved no differently than their well-fed counterparts, displaying uniform indifference, again as expected of chemo-insensitive worms (Fig 4, bottom panels and S3 Fig, panel D). Taken together, these observations suggest that the mild repulsion to I2 in wild-type animals amplified by starvation might reflect integration of *tax-4-*dependent attraction and *osm-9*-dependent repulsion being modulated by feeding state. These findings underscore the utility of analyzing signal transduction mutants.

## Discussion

Here, we analyzed the ability of fermentation products and their biosynthetic precursors to attract *C. elegans*, focusing on two strong attractants, IAA and 2M1B, synthesized from leucine and isoleucine, respectively. Like leucine and isoleucine, IAA and 2M1B are structural isomers of one another and have identical molecular weights and similar or nearly identical physicochemical properties including solubility, density, and vapor pressure. Starvation attenuates attraction to both compounds (Fig. 2, 3) and loss of TAX-4 function eliminates this response (Fig. 4). Yet, the *EC*_50_ for attraction to IAA is nearly 4-fold less than it is for 2M1B (Fig 2, Fig 3). This is not the only example in this study in which *C. elegans* responds differently to structural isomers. We found that I2 (4-methyl-2-oxovaleric acid) weakly repels well-fed and starved animals, while its structural isomer, M2 (3-methyl-2-oxopentanoic acid), evokes no detectable response (Fig. 4). The variation in dose-response relationships between IAA and 2M1B could reflect parallel differences in the affinity of IAA and 2M1B for one or more olfactory receptor protein(s), in the presence of receptors that bind to one compound, but not the other, or in how sensory information is converted into attraction behavior. Future work measuring neural responses to both compounds or linking these compounds to their receptors will be needed to distinguish among these possibilities. Collectively, these findings suggest that despite their shared physicochemical properties, biosynthetic origins in microbial fermentation, and behavioral phenotypes, IAA may be a more salient chemical signal than 2M1B.

We found that an 18-hour starvation period dramatically attenuated the response to both IAA (Fig 2) and to 2M1B (Fig 3). What might account for this effect of starvation? At the molecular level, starvation could decrease the affinity of olfactory receptor(s) for IAA and 2M1B or modify the ensemble of receptors expressed in chemosensory neurons that detect these chemicals. There is evidence for the latter effect in the AWA neurons following a short period of food deprivation (McLachlan et al. 2022), although further study will be required to determine if similar effects are seen more broadly. At the cellular and neural circuit levels, starvation could decrease chemosensory neuron excitability, the efficiency of synaptic transmission generally, or the conversion of chemosensory signals into behavior. Starvation could also affect the behavioral strategy responsible for chemotaxis, increasing the extent of chemosensory adaptation that occurs during the 1-hour assay period.

Whereas starvation dramatically decreased attraction to IAA and 2M1B, it amplified repulsion evoked by the I2 (4-methyl-2-oxovaleric acid) precursor compound (Fig. 4); the average change in position of well-fed wild-type animals was −4.45 mm vs −7.83 mm when starved. Starvation also revealed I2-evoked repulsion in *tax-4* null animals (fed −0.08 mm, starved −4.16 mm), an effect that depended on *osm-9* expression since both well-fed and starved *tax-4;osm-9* double mutants were indifferent to I2 (Fig. 4). Together, these findings implicate *tax-4* in attraction to I2 and *osm-9* in repulsion and suggest that the balance between these two drives is altered by starvation.

All of our assays allowed animals to navigate in a pre-established chemical gradient for 1 hour. Apart from our dose-response studies, it is not known how robust the outcomes we report are to assay conditions. For instance, weak responses might reflect adaptation that occurs during the 1-hour assay. As a result, conditions that decrease sensitivity (starvation, loss of *tax-4* or *osm-9*) might arise from an increase in adaptation rather than a loss of chemosensory function. Time-lapse imaging and modification of the assay period will be needed to address this uncertainty.

Wild-type *C. elegans* are strongly attracted to IAA and 2M1B and most of their biosynthetic precursors evoked little or no chemotactic response. However, as noted above, genetic dissection and analysis of the effect of satiety suggest that indifference reflects the integration of opposing signals rather than a lack of sensitivity. From these findings, it is tempting to speculate that *C. elegans* relies on its nervous system not only to determine which chemical species are worthy of attention, but also those that it ought to ignore. This concept may account for our finding that starvation increases the *EC*_50_ for attraction to both IAA and 2M1B (Fig. 3, 4), despite the association of both compounds with bacterial foods (Chai et al. 2024; Siddiqui et al. 2024). Indeed, de-emphasizing attraction to some chemical species could enable starved animals to redirect their energy towards other, potentially more fruitful attractants. Future studies, potentially making use of a high-throughput chemotaxis platform (Fryer et al. 2024), will be needed to evaluate this model in detail.

## Acknowledgements

This research was funded by grants from the Wu Tsai Neurosciences Institute (Big Ideas, Research Accelerator to MBG, SYR, and TRC) and the National Institutes of Health (R35NS10502 to MBG) as well as a BioX Interdisciplinary Fellowship to LRH. We thank Z. Liao for technical support and members of the Goodman lab and L. E. O’Brien, L. O’Connell, and T. R. Clandinin for their insightful commentary.

## Author contributions

Conceptualization: TLG, SYR, MBG; Data curation: TLG, EF; Formal analysis: TLG, EF; Funding acquisition: LRH, SYR, MBG; Investigation: EF, LRH, TLG; Methodology: TLG; Project Administration: TLG, MBG; Software: TLG, EF, LSS; Supervision Oversight: MBG; Validation: TLG; Visualization: TLG, EF, LSS, MBG; Writing (original): TLG, MBG; Writing (review): TLG, EF, LRH, LSS, SYR, MBG.

## Abbreviations used in the manuscript

Abbreviation: Term
I1: L-leucine (CAS No. 61-90-5)
I2: 4-methyl-2-oxovaleric acid (CAS No. 816-66-0)
I3: Isovaleraldehyde (CAS No. 590-86-3)
IAA: isoamyl alcohol *aka* 3-methyl-1-butanol (CAS No. 123-51-3)
M1: L-isoleucine (CAS No. 73-32-5)
M2: 3-methyl-2-oxopentanoic acid (CAS No. 1460-34-0)
M3: 2-methylbutyraldehyde (CAS No. 96-17-3)
2M1B: 2-methyl-1-butanol (CAS No. 137-32-6)
DMSO: dimethyl sulfoxide (CAS No. 67-68-5)
HEPES: 4-(2-hydroxyethyl)-1-piperazineethanesulfonic acid (CAS No. 7365-45-9)
*EC*_50_: Effective concentration at half-maximal response
<Δ location>: Mean change in location
CNG channel: cyclic-nucleotide gated channel
TRPV channel: vanilloid transient receptor potential channel

## Supplemental Figures

Figure S1 | **Both fed and starved animals are indifferent to pH-buffered water**. Swarm plots (top) showing the position of individual worms (small dots) and the mean location derived from 16 replicates (large dots) for fed (**A**) or starved (**B**) wild-type worms. Distributions of the mean difference (bottom) between each test condition and the reference (water on both sides) are the result of bootstrapped calculations of the difference in mean location (<Δ location>) represented by the central dot, the 95% confidence interval represented by the bar, and the distribution of bootstrapped means to the right of the confidence interval. Supplemental Table Figure S1 contains the numerical values plotted in these graphs.

Figure S2 | **Responses of well-fed wild-type and mutant worms to biosynthetic precursors and fermentation products shown as swarm plots and differences of the mean position**. Swarm plots and bootstrapped differences of the mean position are color coded according to the tested compound, with gray representing the control (HEPES-buffered water), red representing the isoleucine degradation pathway (I1, I2, I3, and IAA), and blue representing the leucine degradation pathway (M1, M2, M3, and 2M1B). **A**. Wild-type animals, fed. *N* = 8 for all compounds. B. *osm-9* null animals, fed. *N* = 10 for HEPES-buffered water, I1, I3, IAA, M1, M2, and M3, *N* = 9 for I2, and *N* = 16 for 2M1B. C. *tax-4* null animals, fed. *N* = 9 for HEPES-buffered water, M2, and M3, *N* = 10 for I1, I2, I3, IAA, M1, and 2M1B. D. *osm-9; tax-4* animals, fed. *N* = 8 for all compounds. Small dots in each swarm plot (top) show the position of individual worms and large dots are the average position of a single replicate. The number of animals tested across replicates is given in parentheses. Distributions of the mean difference (bottom) are the result of bootstrapped calculations of the difference in mean location (<Δ location>) represented by the central dot, the 95% confidence interval represented by the bar, and the distribution of bootstrapped means to the right of the confidence interval. Supplemental Table Figure S2 contains the numerical values plotted in these graphs.

Figure S3 | **Responses of starved wild-type and mutant worms to biosynthetic precursors and fermentation products shown as swarm plots and differences of the mean position**. Swarm plots and bootstrapped differences of the mean position are color coded according to the tested compound, with gray representing the control (HEPES-buffered water), red representing the isoleucine degradation pathway (I1, I2, I3, and IAA), and blue representing the leucine degradation pathway (M1, M2, M3, and 2M1B). **A**. Wild-type animals, starved. *N* = 8 for all compounds. B. *osm-9* null animals, starved. *N* = 8 for all compounds. C. *tax-4* null animals, starved. *N* = 8 for all compounds. D. *osm-9; tax-4* animals, starved. *N* = 8 for all compounds. Small dots in each swarm plot (top) show the position of individual worms and large dots are the average position of a single replicate. The number of animals tested across replicates is given in parentheses. Distributions of the mean difference (bottom) are the result of bootstrapped calculations of the difference in mean location (<Δ location>) represented by the central dot, the 95% confidence interval represented by the bar, and the distribution of bootstrapped means to the right of the confidence interval. Supplemental Table Figure S3 contains the numerical values plotted in these graphs.

