## Supplementary figures and images for "Satiety, TAX-4, and OSM-9 Tune the Attraction of *C. elegans* Nematodes to Microbial Fermentation Products"

### Supplemental Figure S1

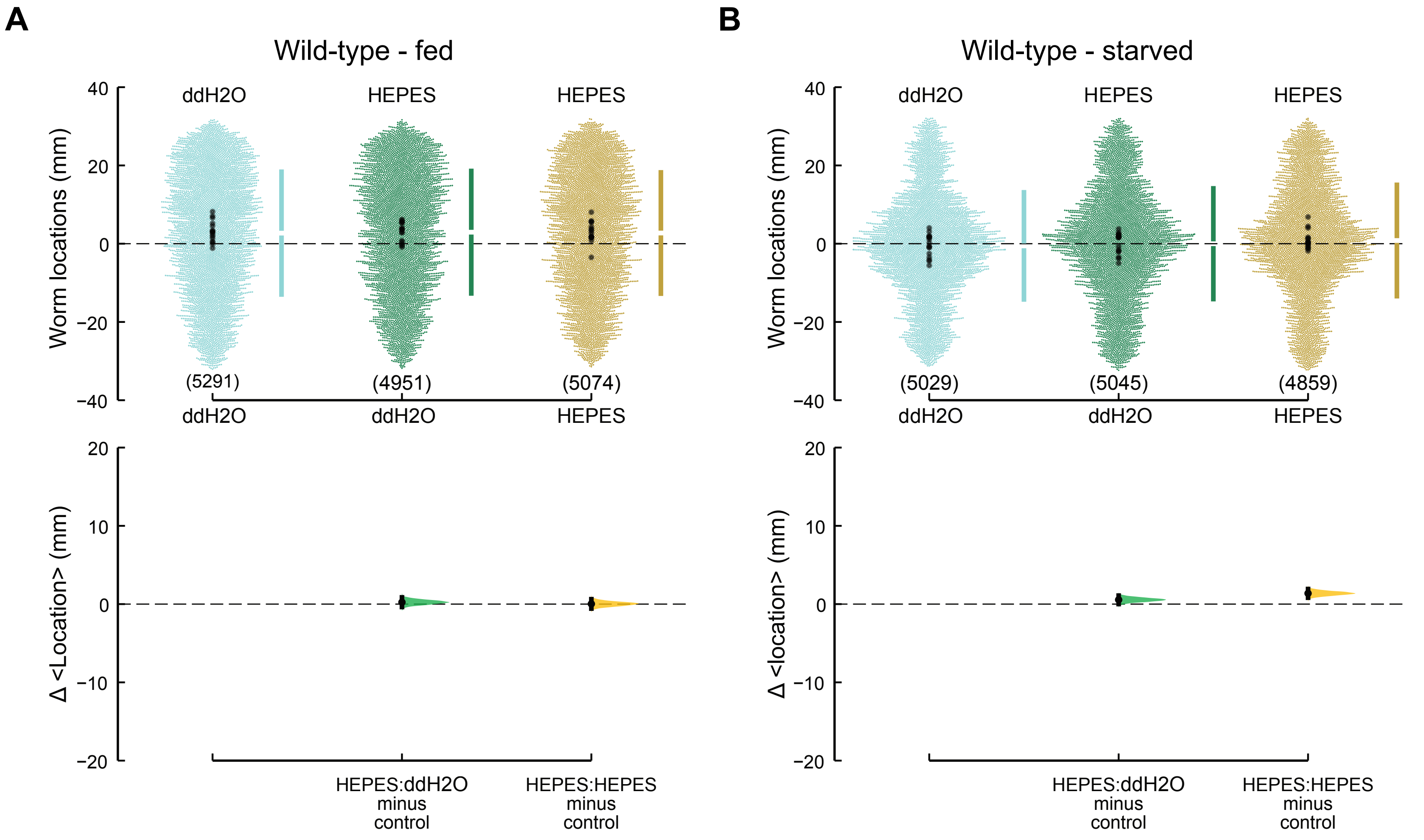

### Supplemental Figure S2

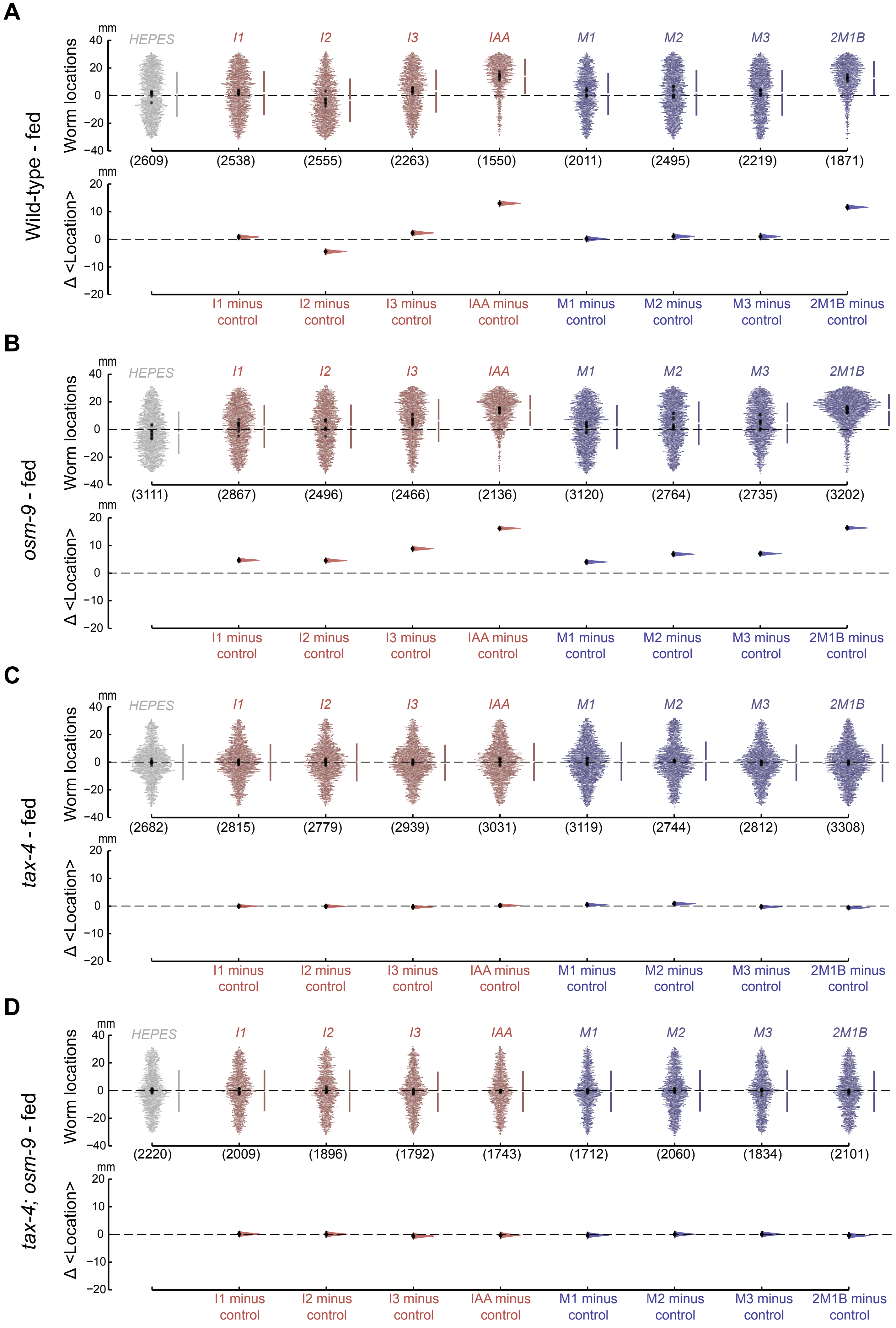

### Supplemental Figure S3

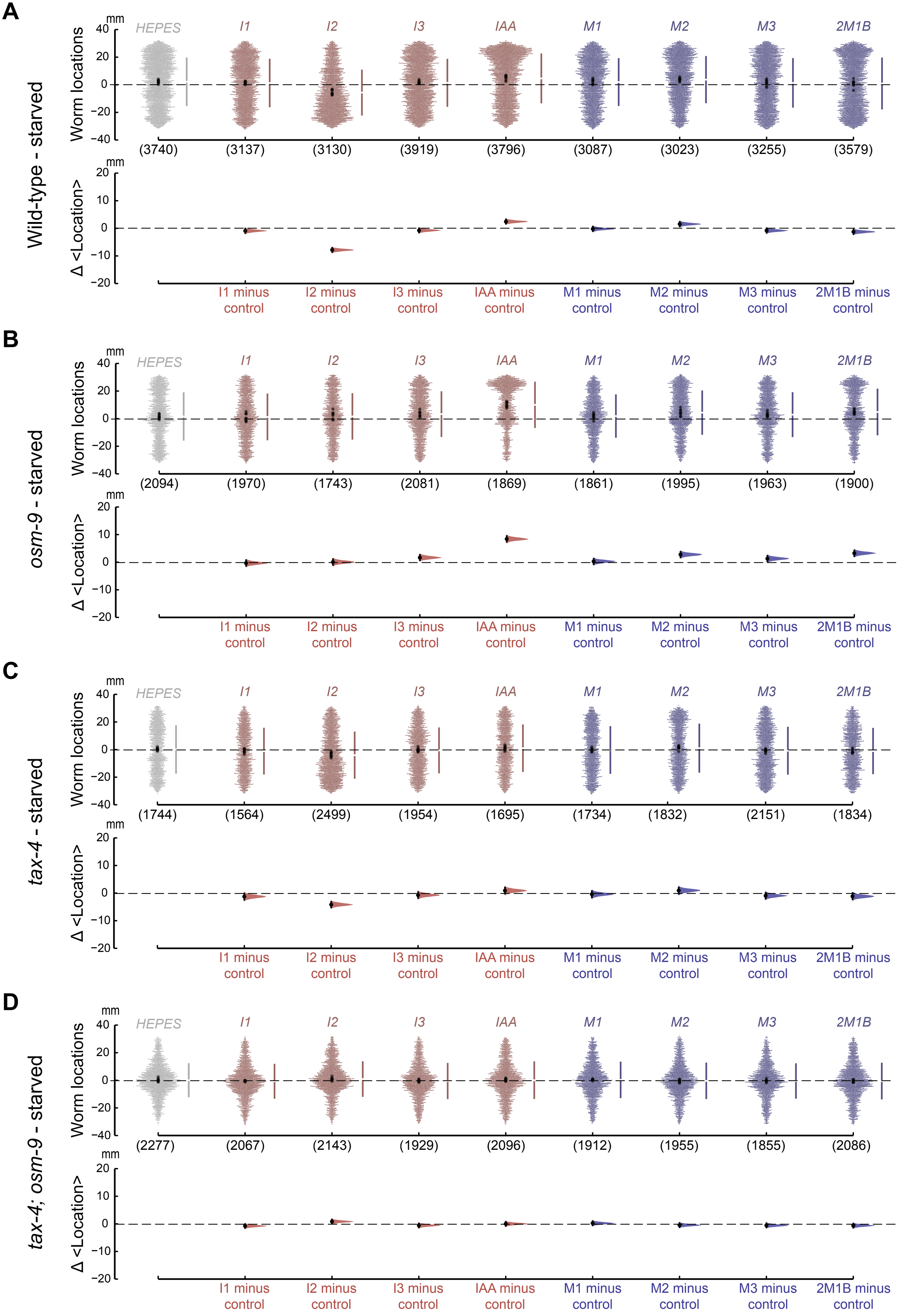
