## Supplemental to Figures 2-4 for "Satiety, TAX-4, and OSM-9 Tune the Attraction of *C. elegans* Nematodes to Microbial Fermentation Products"

FIGURE-2A-REPLICATE MEANS

| Well | Total worms | Chemotaxis index | Compound | Concentration (mM) | Physiological condition | File name | Well width | Plate ID | Mean position (mm) |
|---|---|---|---|---|---|---|---|---|---|
| 1A | 170 | 0.110 | DMSO | Solvent | Fed | IAADR02_Scan002 | 2996 | IAADR02_P01 | 2.527 |
| 3A | 160 | 0.007 | DMSO | Solvent | Fed | IAADR03_Scan002 | 3063 | IAADR03_P03 | -0.769 |
| 3A | 163 | -0.054 | DMSO | Solvent | Fed | IAADR07_Scan001 | 3071 | IAADR07_P03 | -0.523 |
| 1A | 224 | -0.317 | DMSO | Solvent | Fed | IAADR07_Scan002 | 3026 | IAADR07_P05 | -4.117 |
| 1A | 269 | 0.054 | DMSO | Solvent | Fed | IAADR08_Scan002 | 3009 | IAADR08_P01 | 2.269 |
| 3A | 220 | 0.102 | DMSO | Solvent | Fed | IAADR08_Scan002 | 3057 | IAADR08_P03 | 2.809 |
| 1A | 189 | 0.155 | DMSO | Solvent | Fed | IAADR09_Scan003 | 3034 | IAADR09_P01 | 2.934 |
| 3A | 196 | 0.006 | DMSO | Solvent | Fed | IAADR09_Scan003 | 3066 | IAADR09_P03 | -0.157 |
| 4A | 297 | 0.199 | IAA | 0.003 | Fed | IAADR12_Scan001 | 3042 | IAADR12_P04 | 4.451 |
| 4B | 206 | 0.326 | IAA | 0.003 | Fed | IAADR12_Scan001 | 3054 | IAADR12_P04 | 4.476 |
| 4C | 176 | 0.337 | IAA | 0.003 | Fed | IAADR12_Scan001 | 3048 | IAADR12_P04 | 6.333 |
| 4D | 250 | 0.177 | IAA | 0.003 | Fed | IAADR12_Scan001 | 3046 | IAADR12_P04 | 3.172 |
| 3A | 182 | -0.051 | IAA | 0.003 | Fed | IAADR13_Scan001 | 3075 | IAADR13_P03 | 0.127 |
| 3B | 174 | -0.038 | IAA | 0.003 | Fed | IAADR13_Scan001 | 3068 | IAADR13_P03 | -1.058 |
| 3D | 185 | 0.012 | IAA | 0.003 | Fed | IAADR13_Scan001 | 3069 | IAADR13_P03 | 1.096 |
| 1C | 197 | 0.078 | IAA | 0.003 | Fed | IAADR14_Scan001 | 3054 | IAADR14_P01 | 1.369 |
| 3A | 260 | 0.172 | IAA | 0.01 | Fed | IAADR12_Scan001 | 3069 | IAADR12_P03 | 2.755 |
| 3B | 233 | 0.091 | IAA | 0.01 | Fed | IAADR12_Scan001 | 3074 | IAADR12_P03 | 2.534 |
| 3C | 259 | -0.202 | IAA | 0.01 | Fed | IAADR12_Scan001 | 3079 | IAADR12_P03 | -2.406 |
| 3D | 269 | -0.004 | IAA | 0.01 | Fed | IAADR12_Scan001 | 3085 | IAADR12_P03 | 0.963 |
| 2A | 280 | 0.090 | IAA | 0.01 | Fed | IAADR13_Scan002 | 3078 | IAADR13_P06 | 1.118 |
| 2B | 306 | -0.055 | IAA | 0.01 | Fed | IAADR13_Scan002 | 3062 | IAADR13_P06 | -1.027 |
| 2C | 258 | -0.120 | IAA | 0.01 | Fed | IAADR13_Scan002 | 3080 | IAADR13_P06 | -1.368 |
| 2D | 299 | -0.103 | IAA | 0.01 | Fed | IAADR13_Scan002 | 3070 | IAADR13_P06 | -1.582 |
| 1A | 196 | 0.145 | IAA | 0.03 | Fed | IAADR12_Scan001 | 3056 | IAADR12_P01 | 2.053 |
| 1B | 240 | 0.257 | IAA | 0.03 | Fed | IAADR12_Scan001 | 3077 | IAADR12_P01 | 4.185 |
| 1C | 157 | 0.147 | IAA | 0.03 | Fed | IAADR12_Scan001 | 3062 | IAADR12_P01 | 2.431 |
| 1D | 166 | 0.189 | IAA | 0.03 | Fed | IAADR12_Scan001 | 3048 | IAADR12_P01 | 3.791 |
| 1A | 210 | 0.486 | IAA | 0.03 | Fed | IAADR13_Scan001 | 3050 | IAADR13_P01 | 5.568 |
| 1D | 155 | 0.130 | IAA | 0.03 | Fed | IAADR13_Scan001 | 3045 | IAADR13_P01 | 2.057 |
| 2A | 296 | -0.205 | IAA | 0.03 | Fed | IAADR14_Scan001 | 3055 | IAADR14_P02 | -3.067 |
| 2B | 273 | -0.134 | IAA | 0.03 | Fed | IAADR14_Scan001 | 3085 | IAADR14_P02 | -2.281 |
| 2A | 211 | 0.475 | IAA | 0.1 | Fed | IAADR12_Scan002 | 3072 | IAADR12_P06 | 7.427 |
| 2B | 251 | 0.433 | IAA | 0.1 | Fed | IAADR12_Scan002 | 3066 | IAADR12_P06 | 6.550 |
| 2C | 201 | 0.395 | IAA | 0.1 | Fed | IAADR12_Scan002 | 3073 | IAADR12_P06 | 7.088 |
| 2D | 208 | 0.385 | IAA | 0.1 | Fed | IAADR12_Scan002 | 3078 | IAADR12_P06 | 5.809 |
| 1A | 244 | -0.418 | IAA | 0.1 | Fed | IAADR13_Scan002 | 3053 | IAADR13_P05 | -6.262 |
| 1B | 308 | -0.343 | IAA | 0.1 | Fed | IAADR13_Scan002 | 3047 | IAADR13_P05 | -4.206 |
| 1C | 242 | -0.461 | IAA | 0.1 | Fed | IAADR13_Scan002 | 3043 | IAADR13_P05 | -6.307 |
| 1D | 296 | -0.231 | IAA | 0.1 | Fed | IAADR13_Scan002 | 3032 | IAADR13_P05 | -3.482 |

| | | | | | | | | | |
|---|---|---|---|---|---|---|---|---|---|
| 2A | 214 | 0.265 | IAA | 0.3 | Fed | IAADR12_Scan001 | 3081 | IAADR12_P02 | 4.458 |
| 2B | 277 | 0.333 | IAA | 0.3 | Fed | IAADR12_Scan001 | 3081 | IAADR12_P02 | 5.433 |
| 2C | 179 | 0.236 | IAA | 0.3 | Fed | IAADR12_Scan001 | 3077 | IAADR12_P02 | 4.005 |
| 2D | 219 | 0.311 | IAA | 0.3 | Fed | IAADR12_Scan001 | 3076 | IAADR12_P02 | 4.708 |
| 2A | 172 | 0.147 | IAA | 0.3 | Fed | IAADR13_Scan001 | 3071 | IAADR13_P02 | 2.598 |
| 2B | 199 | 0.205 | IAA | 0.3 | Fed | IAADR13_Scan001 | 3077 | IAADR13_P02 | 3.352 |
| 2D | 168 | 0.113 | IAA | 0.3 | Fed | IAADR13_Scan001 | 3080 | IAADR13_P02 | 3.648 |
| 1A | 173 | 0.245 | IAA | 0.3 | Fed | IAADR14_Scan001 | 3041 | IAADR14_P01 | 4.730 |
| 1A | 243 | 0.585 | IAA | 1 | Fed | IAADR12_Scan002 | 3049 | IAADR12_P05 | 9.890 |
| 1B | 205 | 0.665 | IAA | 1 | Fed | IAADR12_Scan002 | 3055 | IAADR12_P05 | 11.387 |
| 1C | 202 | 0.753 | IAA | 1 | Fed | IAADR12_Scan002 | 3047 | IAADR12_P05 | 12.306 |
| 1D | 205 | 0.638 | IAA | 1 | Fed | IAADR12_Scan002 | 3037 | IAADR12_P05 | 10.175 |
| 4A | 252 | 0.030 | IAA | 1 | Fed | IAADR13_Scan001 | 3031 | IAADR13_P04 | 1.055 |
| 4B | 281 | -0.024 | IAA | 1 | Fed | IAADR13_Scan001 | 3036 | IAADR13_P04 | 1.044 |
| 4C | 199 | 0.124 | IAA | 1 | Fed | IAADR13_Scan001 | 3056 | IAADR13_P04 | 3.487 |
| 4D | 261 | 0.292 | IAA | 1 | Fed | IAADR13_Scan001 | 3069 | IAADR13_P04 | 4.637 |
| 1B | 182 | 0.854 | IAA | 3 | Fed | IAADR08_Scan002 | 2925 | IAADR08_P01 | 15.278 |
| 3B | 161 | 0.376 | IAA | 3 | Fed | IAADR09_Scan003 | 3088 | IAADR09_P03 | 6.548 |
| 1B | 219 | 0.393 | IAA | 3 | Fed | IAADR10_Scan003 | 3023 | IAADR10_P01 | 8.221 |
| 1C | 234 | 0.010 | IAA | 3 | Fed | IAADR10_Scan003 | 3049 | IAADR10_P01 | 2.881 |
| 1D | 230 | 0.426 | IAA | 3 | Fed | IAADR10_Scan003 | 3052 | IAADR10_P01 | 8.371 |
| 1A | 264 | 0.654 | IAA | 3 | Fed | IAADR11_Scan001 | 3032 | IAADR11_P01 | 9.668 |
| 1B | 349 | 0.737 | IAA | 3 | Fed | IAADR11_Scan001 | 3029 | IAADR11_P01 | 10.462 |
| 1C | 315 | 0.772 | IAA | 3 | Fed | IAADR11_Scan001 | 3034 | IAADR11_P01 | 10.693 |
| 1C | 251 | 0.652 | IAA | 10 | Fed | IAADR08_Scan002 | 3019 | IAADR08_P01 | 9.353 |
| 3C | 155 | 0.500 | IAA | 10 | Fed | IAADR08_Scan002 | 3046 | IAADR08_P03 | 9.579 |
| 2A | 177 | 0.434 | IAA | 10 | Fed | IAADR10_Scan003 | 3046 | IAADR10_P02 | 8.258 |
| 2B | 175 | 0.369 | IAA | 10 | Fed | IAADR10_Scan003 | 3009 | IAADR10_P02 | 7.740 |
| 2D | 267 | 0.408 | IAA | 10 | Fed | IAADR10_Scan003 | 3066 | IAADR10_P02 | 8.353 |
| 4A | 258 | 0.496 | IAA | 10 | Fed | IAADR10_Scan004 | 3041 | IAADR10_P08 | 8.094 |
| 4C | 332 | 0.192 | IAA | 10 | Fed | IAADR10_Scan004 | 3049 | IAADR10_P08 | 3.980 |
| 1D | 260 | 0.883 | IAA | 10 | Fed | IAADR11_Scan001 | 2973 | IAADR11_P01 | 14.326 |
| 1D | 230 | 0.716 | IAA | 30 | Fed | IAADR08_Scan002 | 2983 | IAADR08_P01 | 12.709 |
| 3D | 158 | 0.551 | IAA | 30 | Fed | IAADR08_Scan002 | 3043 | IAADR08_P03 | 10.741 |
| 3C | 159 | 0.877 | IAA | 30 | Fed | IAADR10_Scan003 | 3066 | IAADR10_P03 | 15.041 |
| 3D | 200 | 0.684 | IAA | 30 | Fed | IAADR10_Scan003 | 3046 | IAADR10_P03 | 12.005 |
| 2A | 284 | 0.731 | IAA | 30 | Fed | IAADR11_Scan001 | 3086 | IAADR11_P02 | 10.547 |
| 2B | 293 | 0.866 | IAA | 30 | Fed | IAADR11_Scan001 | 3053 | IAADR11_P02 | 12.904 |
| 2C | 224 | 0.839 | IAA | 30 | Fed | IAADR11_Scan001 | 3053 | IAADR11_P02 | 13.234 |
| 2D | 276 | 0.858 | IAA | 30 | Fed | IAADR11_Scan001 | 3049 | IAADR11_P02 | 14.782 |
| 2A | 160 | 0.689 | IAA | 100 | Fed | IAADR07_Scan002 | 3051 | IAADR07_P06 | 12.475 |
| 2A | 221 | 0.541 | IAA | 100 | Fed | IAADR08_Scan002 | 3070 | IAADR08_P02 | 10.083 |

| | | | | | | | | | |
|---|---|---|---|---|---|---|---|---|---|
| 4A | 239 | 0.627 | IAA | 100 | Fed | IAADR08_Scan002 | 3055 | IAADR08_P04 | 11.161 |
| 4A | 159 | 0.400 | IAA | 100 | Fed | IAADR09_Scan003 | 3041 | IAADR09_P04 | 8.168 |
| 4A | 168 | 0.885 | IAA | 100 | Fed | IAADR10_Scan003 | 3046 | IAADR10_P04 | 15.920 |
| 4B | 157 | 0.708 | IAA | 100 | Fed | IAADR10_Scan003 | 3038 | IAADR10_P04 | 13.827 |
| 4C | 173 | 0.788 | IAA | 100 | Fed | IAADR10_Scan003 | 3046 | IAADR10_P04 | 14.483 |
| 4D | 174 | 0.836 | IAA | 100 | Fed | IAADR10_Scan003 | 3031 | IAADR10_P04 | 14.554 |
| 2B | 231 | 0.717 | IAA | 300 | Fed | IAADR08_Scan002 | 3034 | IAADR08_P02 | 11.282 |
| 4B | 175 | 0.675 | IAA | 300 | Fed | IAADR09_Scan003 | 3038 | IAADR09_P04 | 12.366 |
| 4B | 187 | 0.742 | IAA | 300 | Fed | IAADR09_Scan004 | 3036 | IAADR09_P08 | 12.835 |
| 1A | 241 | 0.850 | IAA | 300 | Fed | IAADR10_Scan004 | 3023 | IAADR10_P05 | 14.333 |
| 1B | 200 | 0.879 | IAA | 300 | Fed | IAADR10_Scan004 | 3046 | IAADR10_P05 | 16.090 |
| 1C | 200 | 0.813 | IAA | 300 | Fed | IAADR10_Scan004 | 3042 | IAADR10_P05 | 14.266 |
| 1D | 201 | 0.741 | IAA | 300 | Fed | IAADR10_Scan004 | 3053 | IAADR10_P05 | 13.447 |
| 3C | 284 | 0.660 | IAA | 300 | Fed | IAADR10_Scan004 | 3069 | IAADR10_P07 | 11.002 |
| 2C | 151 | 0.694 | IAA | 1000 | Fed | IAADR07_Scan002 | 3042 | IAADR07_P06 | 14.017 |
| 2C | 203 | 0.720 | IAA | 1000 | Fed | IAADR08_Scan002 | 3077 | IAADR08_P02 | 12.591 |
| 4C | 175 | 0.783 | IAA | 1000 | Fed | IAADR09_Scan004 | 3042 | IAADR09_P08 | 14.502 |
| 2A | 162 | 0.868 | IAA | 1000 | Fed | IAADR10_Scan004 | 3060 | IAADR10_P06 | 15.593 |
| 2B | 228 | 0.690 | IAA | 1000 | Fed | IAADR10_Scan004 | 3056 | IAADR10_P06 | 13.171 |
| 2C | 271 | 0.774 | IAA | 1000 | Fed | IAADR10_Scan004 | 3052 | IAADR10_P06 | 13.375 |
| 2D | 208 | 0.806 | IAA | 1000 | Fed | IAADR10_Scan004 | 3058 | IAADR10_P06 | 13.424 |
| 3D | 178 | 0.915 | IAA | 1000 | Fed | IAADR10_Scan004 | 3067 | IAADR10_P07 | 16.405 |
| 2D | 169 | 0.529 | IAA | 3000 | Fed | IAADR02_Scan002 | 3039 | IAADR02_P02 | 10.240 |
| 2D | 242 | 0.683 | IAA | 3000 | Fed | IAADR08_Scan002 | 3091 | IAADR08_P02 | 12.255 |
| 4D | 203 | 0.702 | IAA | 3000 | Fed | IAADR08_Scan002 | 3034 | IAADR08_P04 | 13.288 |
| 2D | 202 | 0.799 | IAA | 3000 | Fed | IAADR09_Scan003 | 3066 | IAADR09_P02 | 13.563 |
| 2D | 198 | 0.665 | IAA | 3000 | Fed | IAADR09_Scan004 | 3049 | IAADR09_P06 | 13.259 |
| 4D | 194 | 0.785 | IAA | 3000 | Fed | IAADR09_Scan004 | 3051 | IAADR09_P08 | 14.593 |
| 3A | 174 | 0.886 | IAA | 3000 | Fed | IAADR10_Scan004 | 3059 | IAADR10_P07 | 15.128 |
| 3B | 271 | 0.811 | IAA | 3000 | Fed | IAADR10_Scan004 | 3074 | IAADR10_P07 | 13.001 |
| 1A | 454 | 0.036 | DMSO | Solvent | Starved | DRS01_Scan002 | 3024 | DRS01_P05 | 0.740 |
| 1B | 562 | 0.085 | DMSO | Solvent | Starved | DRS01_Scan002 | 3034 | DRS01_P05 | 1.334 |
| 1C | 426 | 0.052 | DMSO | Solvent | Starved | DRS01_Scan002 | 3031 | DRS01_P05 | 1.517 |
| 1D | 519 | -0.039 | DMSO | Solvent | Starved | DRS01_Scan002 | 3034 | DRS01_P05 | 0.219 |
| 3A | 485 | 0.031 | DMSO | Solvent | Starved | DRS07_Scan002 | 3061 | DRS07_P07 | 0.080 |
| 3B | 308 | 0.080 | DMSO | Solvent | Starved | DRS07_Scan002 | 3036 | DRS07_P07 | 1.802 |
| 3C | 308 | -0.190 | DMSO | Solvent | Starved | DRS07_Scan002 | 3068 | DRS07_P07 | -2.353 |
| 3D | 286 | -0.012 | Solvent | Solvent | Starved | DRS07_Scan002 | 3068 | DRS07_P07 | 0.104 |
| 1A | 279 | 0.197 | IAA | 0.003 | Starved | DRS03_Scan003 | 3065 | DRS03_P09 | 3.929 |
| 1B | 194 | 0.096 | IAA | 0.003 | Starved | DRS03_Scan003 | 3051 | DRS03_P09 | 2.150 |
| 1C | 250 | 0.146 | IAA | 0.003 | Starved | DRS03_Scan003 | 3045 | DRS03_P09 | 4.273 |
| 1D | 299 | 0.452 | IAA | 0.003 | Starved | DRS03_Scan003 | 3047 | DRS03_P09 | 7.915 |

| | | | | | | | | | |
|---|---|---|---|---|---|---|---|---|---|
| 1A | 278 | 0.070 | IAA | 0.003 | Starved | DRS06_Scan003 | 3044 | DRS06_P09 | 1.152 |
| 1B | 331 | 0.085 | IAA | 0.003 | Starved | DRS06_Scan003 | 3042 | DRS06_P09 | 1.304 |
| 1C | 402 | 0.036 | IAA | 0.003 | Starved | DRS06_Scan003 | 3052 | DRS06_P09 | 0.901 |
| 1D | 386 | -0.036 | IAA | 0.003 | Starved | DRS06_Scan003 | 3043 | DRS06_P09 | 0.211 |
| 4A | 420 | -0.163 | IAA | 0.01 | Starved | DRS02_Scan002 | 3059 | DRS02_P08 | -1.457 |
| 4B | 461 | -0.082 | IAA | 0.01 | Starved | DRS02_Scan002 | 3056 | DRS02_P08 | -0.527 |
| 4C | 397 | -0.121 | IAA | 0.01 | Starved | DRS02_Scan002 | 3067 | DRS02_P08 | 0.038 |
| 4A | 388 | -0.321 | IAA | 0.01 | Starved | DRS05_Scan002 | 3011 | DRS05_P08 | -2.776 |
| 4B | 377 | -0.200 | IAA | 0.01 | Starved | DRS05_Scan002 | 3025 | DRS05_P08 | -2.292 |
| 4C | 293 | -0.061 | IAA | 0.01 | Starved | DRS05_Scan002 | 3027 | DRS05_P08 | -0.815 |
| 4D | 243 | -0.340 | IAA | 0.01 | Starved | DRS05_Scan002 | 3024 | DRS05_P08 | -4.244 |
| 2C | 239 | -0.077 | IAA | 0.01 | Starved | DRS07_Scan002 | 3078 | DRS07_P05 | -1.878 |
| 2A | 449 | -0.012 | IAA | 0.03 | Starved | DRS01_Scan001 | 3062 | DRS01_P02 | -0.454 |
| 2B | 400 | 0.054 | IAA | 0.03 | Starved | DRS01_Scan001 | 3060 | DRS01_P02 | 0.526 |
| 2C | 428 | 0.137 | IAA | 0.03 | Starved | DRS01_Scan001 | 3030 | DRS01_P02 | 2.581 |
| 2D | 439 | -0.003 | IAA | 0.03 | Starved | DRS01_Scan001 | 3061 | DRS01_P02 | -0.013 |
| 2A | 187 | 0.431 | IAA | 0.03 | Starved | DRS04_Scan001 | 3074 | DRS04_P02 | 3.879 |
| 2D | 165 | 0.298 | IAA | 0.03 | Starved | DRS04_Scan001 | 3068 | DRS04_P02 | 2.850 |
| 1C | 309 | 0.054 | IAA | 0.03 | Starved | DRS07_Scan001 | 3024 | DRS07_P01 | 1.865 |
| 1D | 272 | 0.198 | IAA | 0.03 | Starved | DRS07_Scan001 | 3037 | DRS07_P01 | 3.556 |
| 2A | 254 | 0.071 | IAA | 0.1 | Starved | DRS03_Scan001 | 3053 | DRS03_P02 | 1.439 |
| 2B | 226 | 0.343 | IAA | 0.1 | Starved | DRS03_Scan001 | 3063 | DRS03_P02 | 5.094 |
| 2C | 246 | 0.189 | IAA | 0.1 | Starved | DRS03_Scan001 | 3057 | DRS03_P02 | 2.769 |
| 2D | 228 | 0.349 | IAA | 0.1 | Starved | DRS03_Scan001 | 3062 | DRS03_P02 | 6.536 |
| 2A | 518 | 0.009 | IAA | 0.1 | Starved | DRS06_Scan001 | 3053 | DRS06_P02 | 0.769 |
| 2B | 448 | 0.010 | IAA | 0.1 | Starved | DRS06_Scan001 | 3039 | DRS06_P02 | 0.340 |
| 2C | 301 | 0.047 | IAA | 0.1 | Starved | DRS06_Scan001 | 3034 | DRS06_P02 | 1.119 |
| 2D | 365 | 0.009 | IAA | 0.1 | Starved | DRS06_Scan001 | 3045 | DRS06_P02 | 0.498 |
| 1A | 428 | -0.403 | IAA | 0.3 | Starved | DRS02_Scan002 | 3037 | DRS02_P05 | -5.402 |
| 1B | 516 | -0.123 | IAA | 0.3 | Starved | DRS02_Scan002 | 3038 | DRS02_P05 | -1.563 |
| 1C | 365 | -0.108 | IAA | 0.3 | Starved | DRS02_Scan002 | 3034 | DRS02_P05 | -0.378 |
| 2A | 276 | 0.915 | IAA | 0.3 | Starved | DRS05_Scan002 | 3067 | DRS05_P05 | 15.772 |
| 2B | 280 | 0.847 | IAA | 0.3 | Starved | DRS05_Scan002 | 3081 | DRS05_P05 | 14.538 |
| 2C | 259 | 0.918 | IAA | 0.3 | Starved | DRS05_Scan002 | 3079 | DRS05_P05 | 15.071 |
| 2D | 246 | 0.808 | IAA | 0.3 | Starved | DRS05_Scan002 | 3071 | DRS05_P05 | 14.984 |
| 2D | 228 | 0.209 | IAA | 0.3 | Starved | DRS07_Scan001 | 3059 | DRS07_P02 | 3.373 |
| 1A | 370 | -0.060 | IAA | 1 | Starved | DRS02_Scan003 | 3053 | DRS02_P09 | -0.002 |
| 1B | 351 | -0.138 | IAA | 1 | Starved | DRS02_Scan003 | 3055 | DRS02_P09 | -1.620 |
| 1C | 340 | 0.031 | IAA | 1 | Starved | DRS02_Scan003 | 3050 | DRS02_P09 | 1.150 |
| 1A | 441 | 0.042 | IAA | 1 | Starved | DRS05_Scan003 | 3054 | DRS05_P09 | 0.743 |
| 1B | 381 | 0.091 | IAA | 1 | Starved | DRS05_Scan003 | 3044 | DRS05_P09 | 2.061 |
| 1C | 258 | 0.145 | IAA | 1 | Starved | DRS05_Scan003 | 3047 | DRS05_P09 | 3.329 |

| | | | | | | | | | |
|---|---|---|---|---|---|---|---|---|---|
| 1D | 331 | 0.088 | IAA | 1 | Starved | DRS05_Scan003 | 3038 | DRS05_P09 | 2.315 |
| 2D | 216 | 0.173 | IAA | 1 | Starved | DRS07_Scan002 | 3065 | DRS07_P05 | 2.646 |
| 4A | 443 | 0.023 | IAA | 3 | Starved | DRS01_Scan001 | 3029 | DRS01_P04 | 1.000 |
| 4B | 367 | 0.169 | IAA | 3 | Starved | DRS01_Scan001 | 3025 | DRS01_P04 | 2.857 |
| 4C | 362 | 0.213 | IAA | 3 | Starved | DRS01_Scan001 | 3047 | DRS01_P04 | 3.060 |
| 4D | 428 | 0.140 | IAA | 3 | Starved | DRS01_Scan001 | 3043 | DRS01_P04 | 2.523 |
| 1A | 237 | 0.692 | IAA | 3 | Starved | DRS07_Scan002 | 3052 | DRS07_P06 | 12.524 |
| 1B | 199 | 0.234 | IAA | 3 | Starved | DRS07_Scan002 | 3029 | DRS07_P06 | 5.460 |
| 1C | 292 | 0.019 | IAA | 3 | Starved | DRS07_Scan002 | 3032 | DRS07_P06 | 1.092 |
| 1D | 287 | 0.075 | IAA | 3 | Starved | DRS07_Scan002 | 3034 | DRS07_P06 | 1.889 |
| 3A | 386 | 0.154 | IAA | 10 | Starved | DRS01_Scan002 | 3058 | DRS01_P07 | 2.517 |
| 3B | 368 | 0.222 | IAA | 10 | Starved | DRS01_Scan002 | 3066 | DRS01_P07 | 4.005 |
| 3C | 562 | 0.228 | IAA | 10 | Starved | DRS01_Scan002 | 3071 | DRS01_P07 | 3.720 |
| 3D | 523 | 0.215 | IAA | 10 | Starved | DRS01_Scan002 | 3071 | DRS01_P07 | 3.658 |
| 3D | 179 | 0.485 | IAA | 10 | Starved | DRS04_Scan002 | 3073 | DRS04_P07 | 6.758 |
| 3B | 250 | 0.069 | IAA | 10 | Starved | DRS07_Scan001 | 3035 | DRS07_P03 | 1.762 |
| 3C | 241 | 0.161 | IAA | 10 | Starved | DRS07_Scan001 | 3048 | DRS07_P03 | 3.625 |
| 3D | 315 | 0.265 | IAA | 10 | Starved | DRS07_Scan001 | 3052 | DRS07_P03 | 4.325 |
| 1A | 337 | 0.150 | IAA | 30 | Starved | DRS03_Scan002 | 3047 | DRS03_P06 | 3.147 |
| 1B | 241 | 0.371 | IAA | 30 | Starved | DRS03_Scan002 | 3035 | DRS03_P06 | 7.263 |
| 1C | 258 | 0.339 | IAA | 30 | Starved | DRS03_Scan002 | 3014 | DRS03_P06 | 5.990 |
| 1D | 303 | 0.090 | IAA | 30 | Starved | DRS03_Scan002 | 3086 | DRS03_P06 | 1.773 |
| 1A | 337 | 0.062 | IAA | 30 | Starved | DRS06_Scan002 | 3041 | DRS06_P06 | 0.864 |
| 1B | 264 | 0.076 | IAA | 30 | Starved | DRS06_Scan002 | 3032 | DRS06_P06 | 1.412 |
| 1C | 418 | -0.014 | IAA | 30 | Starved | DRS06_Scan002 | 3045 | DRS06_P06 | -0.075 |
| 1D | 415 | -0.050 | IAA | 30 | Starved | DRS06_Scan002 | 3047 | DRS06_P06 | -0.489 |
| 1A | 218 | 0.623 | IAA | 100 | Starved | DRS03_Scan001 | 3044 | DRS03_P01 | 14.048 |
| 1B | 200 | 0.679 | IAA | 100 | Starved | DRS03_Scan001 | 3036 | DRS03_P01 | 13.438 |
| 1C | 230 | 0.615 | IAA | 100 | Starved | DRS03_Scan001 | 3024 | DRS03_P01 | 11.883 |
| 1D | 259 | 0.717 | IAA | 100 | Starved | DRS03_Scan001 | 3043 | DRS03_P01 | 13.984 |
| 1A | 283 | 0.498 | IAA | 100 | Starved | DRS06_Scan001 | 3061 | DRS06_P01 | 8.284 |
| 1B | 373 | 0.242 | IAA | 100 | Starved | DRS06_Scan001 | 3055 | DRS06_P01 | 5.199 |
| 1C | 384 | 0.408 | IAA | 100 | Starved | DRS06_Scan001 | 3039 | DRS06_P01 | 8.399 |
| 1D | 472 | 0.326 | IAA | 100 | Starved | DRS06_Scan001 | 3052 | DRS06_P01 | 5.822 |
| 3A | 546 | 0.576 | IAA | 300 | Starved | DRS02_Scan001 | 3067 | DRS02_P03 | 9.660 |
| 3B | 447 | 0.667 | IAA | 300 | Starved | DRS02_Scan001 | 3061 | DRS02_P03 | 12.352 |
| 3C | 419 | 0.545 | IAA | 300 | Starved | DRS02_Scan001 | 3065 | DRS02_P03 | 10.177 |
| 3D | 293 | 0.634 | IAA | 300 | Starved | DRS02_Scan001 | 3056 | DRS02_P03 | 12.009 |
| 3A | 328 | 0.642 | IAA | 300 | Starved | DRS05_Scan001 | 3067 | DRS05_P03 | 11.786 |
| 3B | 235 | 0.787 | IAA | 300 | Starved | DRS05_Scan001 | 3080 | DRS05_P03 | 15.817 |
| 3C | 277 | 0.692 | IAA | 300 | Starved | DRS05_Scan001 | 3070 | DRS05_P03 | 12.525 |
| 3D | 293 | 0.448 | IAA | 300 | Starved | DRS05_Scan001 | 3079 | DRS05_P03 | 9.022 |

| | | | | | | | | | |
|---|---|---|---|---|---|---|---|---|---|
| 1A | 379 | 0.411 | IAA | 1000 | Starved | DRS01_Scan003 | 3016 | DRS01_P09 | 8.691 |
| 1B | 340 | 0.466 | IAA | 1000 | Starved | DRS01_Scan003 | 3048 | DRS01_P09 | 8.490 |
| 1C | 285 | 0.504 | IAA | 1000 | Starved | DRS01_Scan003 | 3026 | DRS01_P09 | 9.968 |
| 1D | 324 | 0.473 | IAA | 1000 | Starved | DRS01_Scan003 | 3050 | DRS01_P09 | 9.212 |
| 1A | 172 | 0.575 | IAA | 1000 | Starved | DRS04_Scan003 | 3040 | DRS04_P09 | 9.981 |
| 1B | 175 | 0.800 | IAA | 1000 | Starved | DRS04_Scan003 | 3039 | DRS04_P09 | 12.199 |
| 1D | 162 | 0.512 | IAA | 1000 | Starved | DRS04_Scan003 | 3036 | DRS04_P09 | 10.150 |
| 4A | 231 | 0.734 | IAA | 1000 | Starved | DRS07_Scan002 | 3051 | DRS07_P08 | 14.603 |
| 4A | 269 | 0.938 | IAA | 3000 | Starved | DRS03_Scan002 | 3078 | DRS03_P08 | 18.116 |
| 4B | 229 | 0.946 | IAA | 3000 | Starved | DRS03_Scan002 | 3056 | DRS03_P08 | 15.512 |
| 4C | 193 | 0.935 | IAA | 3000 | Starved | DRS03_Scan002 | 3056 | DRS03_P08 | 17.455 |
| 4D | 254 | 0.959 | IAA | 3000 | Starved | DRS03_Scan002 | 3065 | DRS03_P08 | 17.753 |
| 4A | 296 | 0.945 | IAA | 3000 | Starved | DRS06_Scan002 | 3045 | DRS06_P08 | 18.580 |
| 4B | 261 | 0.880 | IAA | 3000 | Starved | DRS06_Scan002 | 3055 | DRS06_P08 | 17.933 |
| 4C | 266 | 0.985 | IAA | 3000 | Starved | DRS06_Scan002 | 3047 | DRS06_P08 | 19.377 |
| 4D | 243 | 0.924 | IAA | 3000 | Starved | DRS06_Scan002 | 3051 | DRS06_P08 | 18.911 |

### FIGURE 2A-ISOAMYL ALCOHOL-FED-WORMPOSITIONS

| DMSO | IA D01 | IA D02 | IA D03 | IA D04 | IA D05 | IA D06 | IA D07 | IA D08 | IA D09 | IA D10 | IA D11 |
|---|---|---|---|---|---|---|---|---|---|---|---|
| 8.533 | -17.579 | 19.266 | -7.725 | -0.106 | -1.255 | -8.184 | -0.988 | 20.108 | 9.956 | -0.553 | -16.270 |
| -17.510 | -21.407 | 15.587 | 22.894 | 9.785 | -2.087 | 0.935 | 22.612 | 9.882 | 14.165 | 5.313 | -22.902 |
| 16.583 | -2.616 | -9.935 | -13.968 | 17.845 | -3.722 | 18.424 | -12.585 | 10.995 | -6.653 | -4.707 | -11.898 |
| 13.806 | 0.192 | 0.900 | 4.814 | 20.792 | -5.780 | -2.478 | 1.576 | 0.916 | 13.576 | 12.176 | 0.386 |
| 6.017 | -1.350 | -7.482 | -15.568 | -8.272 | -6.542 | -0.157 | -14.763 | 14.185 | 4.514 | 14.743 | -2.856 |
| 14.738 | -8.838 | -8.048 | -1.623 | 0.806 | -7.173 | -15.610 | -21.876 | 18.900 | 20.419 | -5.427 | -25.788 |
| -7.485 | -12.591 | -18.140 | -26.045 | -2.308 | -8.697 | 13.317 | -15.744 | 20.929 | 24.284 | 16.119 | 8.368 |
| -22.856 | 17.744 | 16.352 | 6.406 | 18.237 | -1.147 | 12.339 | 27.574 | -8.403 | -14.468 | -7.690 | -11.103 |
| 18.125 | 17.147 | 8.029 | -17.661 | 8.407 | -7.829 | -12.105 | 16.749 | 6.365 | 12.613 | -22.677 | 13.109 |
| -24.836 | -1.000 | 0.958 | 9.524 | -6.663 | 24.473 | 2.080 | 8.270 | 17.934 | 24.969 | 9.310 | -1.401 |
| -3.443 | 10.629 | 8.768 | -21.306 | -4.589 | -14.819 | 14.981 | 5.897 | 17.166 | -13.593 | -21.044 | 6.801 |
| 25.193 | -5.064 | 20.881 | -11.404 | -21.991 | 8.630 | 17.954 | 15.097 | 18.526 | 1.719 | 14.331 | 13.747 |
| 2.230 | 7.507 | 6.227 | -3.422 | 21.455 | 9.372 | 22.849 | -3.777 | 16.723 | 8.355 | -14.645 | 9.788 |
| -14.630 | 5.303 | 15.779 | 1.542 | 23.169 | 7.398 | -16.009 | 14.394 | -2.255 | 21.707 | 23.513 | 19.697 |
| -18.786 | -19.672 | -4.096 | 20.463 | 20.436 | 7.993 | 6.544 | 17.605 | 24.611 | -16.654 | -18.377 | -21.627 |
| 0.975 | 20.741 | -24.563 | 19.521 | 16.321 | 8.356 | 18.307 | 18.847 | 20.006 | 26.375 | 19.878 | 22.890 |
| 20.744 | 6.062 | -21.251 | 8.152 | 21.951 | 7.023 | -17.908 | 26.371 | 14.294 | 10.885 | 4.233 | 11.199 |
| 2.551 | 25.922 | -8.980 | -10.126 | 10.037 | 24.496 | -2.382 | 22.050 | 11.138 | 19.770 | 11.509 | 11.626 |
| -7.733 | -7.120 | -0.144 | 8.725 | -17.101 | 5.830 | 7.592 | 29.403 | -19.386 | 5.113 | 14.822 | 6.526 |
| 11.703 | 2.701 | -10.250 | -29.257 | 26.352 | 1.663 | 19.976 | 6.310 | 15.992 | -1.145 | 26.350 | 17.301 |
| -19.789 | -21.353 | 1.516 | 22.031 | 2.232 | 19.637 | 27.347 | 28.087 | 2.301 | 24.880 | 18.043 | 8.389 |
| -1.341 | -16.137 | -2.967 | -4.000 | 3.289 | 21.447 | -9.166 | 14.870 | 6.702 | 5.767 | 0.555 | 13.848 |
| -19.232 | 25.469 | 12.426 | 14.569 | 19.654 | -2.747 | 6.021 | 27.377 | 24.973 | 2.924 | 25.562 | 16.441 |
| 22.676 | -5.758 | -26.417 | -4.376 | 18.592 | 24.655 | 18.806 | 26.141 | 26.128 | 14.199 | -1.785 | 3.429 |
| -0.789 | 20.850 | -8.373 | -4.966 | 20.910 | 1.390 | 22.379 | 7.216 | 21.134 | 22.034 | -21.854 | 4.590 |
| -20.850 | -24.977 | -15.340 | 4.761 | 8.910 | -15.068 | 14.808 | 20.970 | -7.824 | 19.916 | 21.948 | 26.054 |
| 3.497 | -6.463 | -16.556 | -6.015 | 12.053 | 0.385 | 0.644 | 28.016 | 9.366 | 3.753 | 17.759 | -27.362 |
| -6.062 | 7.952 | -3.365 | -6.749 | 17.922 | 10.850 | 26.926 | 23.246 | 23.168 | 11.168 | -5.786 | 23.655 |
| -2.020 | 18.148 | -0.127 | -3.282 | -9.312 | 16.747 | 20.622 | 21.998 | 12.837 | 8.269 | 20.797 | 17.076 |
| 12.368 | 2.835 | 10.420 | -13.686 | 20.679 | -4.161 | 10.632 | 11.066 | 8.405 | 13.279 | 10.046 | 19.840 |
| -23.732 | 22.189 | 9.373 | 7.557 | -2.055 | -15.449 | 24.729 | 21.034 | 3.760 | 21.207 | 19.436 | 26.860 |
| -18.487 | -3.209 | -0.929 | -17.670 | 13.652 | 19.335 | 25.441 | 24.615 | 26.790 | 2.492 | 21.937 | 9.283 |
| 6.047 | 16.975 | 12.919 | -1.865 | 26.031 | -2.578 | 19.577 | 16.555 | 17.247 | 24.114 | -27.121 | 12.212 |

| | | | | | | | | | | | |
|---|---|---|---|---|---|---|---|---|---|---|---|
| 17.544 | -6.666 | 11.617 | 23.725 | -19.376 | 11.384 | 9.449 | 7.969 | 13.329 | 27.453 | 18.430 | 1.555 |
| -7.623 | 12.048 | 20.809 | 2.613 | 26.106 | 21.555 | 7.893 | 10.457 | -5.822 | 17.754 | 14.968 | -11.935 |
| 13.681 | 27.318 | -20.433 | -13.256 | 6.325 | 12.952 | -13.613 | 27.908 | 15.165 | 23.599 | 23.707 | 24.913 |
| -19.624 | 21.453 | 24.137 | -14.889 | 10.124 | -0.852 | 22.190 | 17.570 | 21.106 | 22.161 | 26.016 | 25.555 |
| -0.919 | -16.117 | -12.217 | 18.641 | 4.886 | 4.684 | 0.633 | 27.174 | 19.597 | 7.304 | -11.798 | -12.582 |
| 23.803 | -4.996 | -13.878 | 13.302 | 28.885 | -31.232 | 14.991 | 26.437 | 12.319 | 26.062 | 0.672 | 9.900 |
| 18.402 | -6.383 | -23.673 | 20.941 | 3.067 | 18.199 | 17.195 | 20.440 | 3.870 | 18.799 | 17.825 | 22.527 |
| -3.790 | 15.822 | 19.284 | 4.186 | 20.578 | 23.278 | 28.243 | 21.948 | -13.833 | 14.396 | 16.672 | 16.248 |
| 12.912 | 5.673 | -18.940 | 20.075 | -24.416 | -2.674 | 27.079 | 3.290 | 13.149 | 9.342 | 21.565 | 20.523 |
| -25.612 | 11.035 | 14.836 | 15.165 | -4.464 | -3.222 | 24.056 | 28.263 | 21.805 | 26.493 | 14.766 | 12.961 |
| -25.445 | -19.421 | -8.110 | 4.909 | -2.681 | 5.663 | 1.522 | 29.134 | 1.116 | 24.894 | 19.598 | 14.135 |
| -7.013 | -4.687 | 7.783 | 7.571 | 26.591 | 27.123 | 25.014 | 21.617 | -3.352 | 7.790 | 11.181 | 7.140 |
| -27.240 | 24.655 | -16.933 | -18.490 | 2.603 | -20.413 | 22.582 | 27.949 | 24.412 | 15.274 | -4.622 | 23.393 |
| 4.776 | 7.497 | 16.910 | -14.228 | 13.101 | 31.021 | 16.082 | 4.185 | 23.877 | 10.047 | 16.519 | 12.221 |
| -4.230 | 17.130 | 6.746 | -10.653 | 11.374 | 9.916 | 29.383 | 29.034 | 15.489 | 15.793 | 11.797 | 15.334 |
| 25.398 | 9.715 | -11.646 | -7.872 | 28.098 | 1.868 | 26.798 | 17.770 | 21.900 | 11.439 | -4.462 | 18.238 |
| -3.113 | 14.618 | 17.733 | 15.485 | 16.359 | -6.358 | 29.116 | 19.845 | 27.712 | -20.973 | 22.233 | 4.248 |
| 2.741 | 3.582 | -21.336 | -0.573 | 14.430 | -11.715 | 20.098 | 17.118 | 16.602 | 19.725 | 25.554 | 17.298 |
| -15.886 | -13.652 | -27.630 | -11.594 | 19.814 | 24.713 | 13.442 | 26.000 | -3.251 | 10.258 | 24.083 | 16.409 |
| 21.434 | 11.557 | -8.315 | 3.797 | 19.186 | 23.526 | 24.144 | 23.062 | 8.000 | 25.194 | -1.937 | 10.361 |
| 8.635 | 27.213 | 18.999 | 19.044 | 13.061 | -18.649 | 17.130 | -1.167 | 10.291 | 23.108 | 18.264 | 21.106 |
| 23.443 | 13.347 | 12.063 | 6.848 | 8.006 | 5.344 | -22.491 | 19.224 | 19.004 | 5.720 | 17.471 | 23.099 |
| 12.633 | -4.386 | 7.460 | 7.754 | 2.938 | -0.990 | 25.197 | 27.926 | 25.883 | 27.680 | 10.824 | 18.555 |
| -12.085 | -23.207 | 19.979 | 5.120 | 10.945 | -9.597 | 25.222 | 20.490 | 18.231 | 19.072 | 25.145 | 25.138 |
| 7.961 | 19.116 | -24.097 | 2.634 | 28.203 | 14.509 | 20.867 | -2.246 | 15.194 | 16.336 | 27.652 | 21.841 |
| 16.811 | -6.060 | 10.839 | -10.208 | 29.175 | 6.804 | 26.514 | 18.371 | 0.359 | -8.536 | 9.308 | 8.375 |
| -15.329 | -2.766 | -8.917 | 11.992 | -5.536 | 31.645 | 9.196 | 20.069 | 23.484 | 23.916 | 4.474 | 16.618 |
| -3.300 | -11.977 | 22.880 | 5.716 | -19.039 | 17.645 | -16.838 | 26.790 | -1.716 | 15.041 | 23.906 | 23.508 |
| 21.759 | 27.188 | 3.896 | -14.612 | 31.016 | 15.140 | 18.970 | 22.402 | 19.788 | 26.544 | 21.911 | -2.744 |
| -24.908 | -9.306 | -3.501 | -6.749 | 8.849 | -5.530 | 24.041 | 28.538 | 9.445 | 17.654 | 28.234 | 4.740 |
| 16.579 | 11.868 | -15.550 | -10.662 | 31.421 | 12.172 | 27.865 | 25.498 | 17.457 | 25.055 | 26.792 | 10.588 |
| -21.013 | -7.009 | 3.007 | 9.292 | -29.569 | -0.768 | 16.578 | 2.163 | 12.316 | -4.107 | 27.118 | 20.076 |
| 10.242 | 24.289 | 28.483 | 0.530 | 29.909 | 10.108 | -21.969 | 26.830 | -9.486 | 1.095 | 27.595 | 28.206 |
| 14.789 | 7.675 | 8.778 | -3.218 | 7.780 | -19.643 | 17.987 | 6.044 | 2.098 | 21.436 | 18.831 | 2.981 |
| -29.624 | -23.836 | 15.770 | -9.138 | 24.404 | 4.221 | 0.495 | 12.547 | 15.311 | 17.523 | 22.375 | 13.739 |

| | | | | | | | | | | | |
|---|---|---|---|---|---|---|---|---|---|---|---|
| 19.980 | 15.572 | -7.864 | -19.268 | 12.880 | 23.007 | 3.014 | 15.574 | 8.061 | 14.984 | 16.932 | 7.519 |
| -7.954 | -22.979 | -25.357 | -3.659 | 17.276 | 18.665 | 20.889 | 3.242 | -5.600 | 15.617 | -9.028 | -17.922 |
| 7.166 | -13.247 | -5.987 | -1.194 | 5.406 | 14.657 | -1.444 | 22.266 | -10.460 | 25.911 | 25.758 | 24.277 |
| -11.923 | 9.702 | -18.174 | 25.286 | 11.234 | 26.623 | 30.024 | 19.270 | 10.388 | 8.479 | 0.794 | 4.806 |
| -23.717 | 20.883 | -28.529 | 16.103 | 8.882 | 5.659 | 24.834 | 24.251 | -8.166 | 19.881 | 30.505 | -30.085 |
| 9.732 | -10.335 | -23.645 | -7.569 | 13.111 | -6.196 | 15.276 | 21.159 | 11.551 | -22.588 | 10.084 | -3.591 |
| 25.558 | 3.258 | -29.883 | 9.171 | 31.096 | -7.775 | 11.629 | 14.380 | 8.745 | 12.447 | 19.157 | 12.880 |
| -27.984 | -6.774 | 13.811 | -14.401 | 4.025 | -8.879 | 26.103 | 2.422 | -4.772 | 11.557 | 21.198 | 24.958 |
| 3.682 | 14.797 | -8.438 | 23.733 | -23.426 | -23.079 | 10.981 | 25.675 | -6.929 | -3.709 | 24.593 | 18.872 |
| 21.698 | -16.359 | 12.409 | -22.737 | 2.288 | 11.358 | 20.496 | 16.241 | 17.730 | 21.973 | 22.265 | 7.513 |
| 23.825 | -8.039 | 22.004 | -5.217 | -31.360 | 20.649 | -24.852 | 22.881 | 15.091 | -4.296 | 18.329 | 26.361 |
| 22.804 | 10.118 | -30.369 | -10.469 | 17.427 | 0.818 | 28.222 | 13.408 | 18.803 | 14.057 | 9.341 | 23.125 |
| 18.196 | 8.697 | 28.052 | -8.910 | -8.683 | 22.856 | 15.375 | 21.956 | 14.124 | -24.964 | 28.671 | 28.897 |
| 29.057 | -16.790 | 20.633 | 2.141 | 12.747 | 5.469 | 1.141 | 26.753 | -9.134 | 19.498 | 26.445 | 13.904 |
| -9.650 | -13.515 | 14.742 | 8.577 | 28.239 | 27.254 | 8.299 | 16.817 | 16.823 | 16.898 | 24.077 | 11.149 |
| 21.107 | 22.346 | -13.713 | 12.860 | -28.798 | 17.355 | 17.516 | 23.354 | 19.949 | 26.224 | 23.286 | -2.150 |
| -6.913 | -9.070 | -19.835 | 4.567 | 11.878 | 28.085 | 29.086 | 25.336 | -1.992 | 24.861 | 27.841 | 15.883 |
| -0.646 | 5.713 | -11.883 | 19.633 | 18.422 | -4.846 | -2.336 | 21.070 | 21.351 | -0.534 | 19.533 | 19.148 |
| -19.756 | -9.808 | 21.224 | -19.023 | 3.586 | 16.197 | 2.269 | 8.439 | -8.590 | 11.201 | 28.198 | 16.719 |
| 23.934 | 10.378 | -10.347 | 22.232 | 7.062 | 21.499 | 19.522 | 20.224 | 7.696 | -15.067 | 18.722 | 18.071 |
| -17.312 | -19.716 | -17.715 | -12.542 | 26.483 | 14.290 | 29.645 | 18.251 | 14.218 | 16.136 | 22.030 | -23.352 |
| 7.619 | 6.117 | 27.313 | -7.181 | -0.691 | 11.457 | 24.226 | 13.634 | 10.785 | 23.796 | 17.207 | 9.867 |
| -8.692 | 4.510 | 18.797 | 6.858 | 18.876 | -9.608 | 6.115 | 19.302 | 8.459 | 20.288 | 10.481 | 22.660 |
| -0.641 | 23.628 | 18.038 | -18.455 | 8.598 | -24.496 | 28.048 | 17.379 | 17.869 | 12.341 | 17.509 | 21.051 |
| 22.071 | 6.966 | -9.115 | 10.436 | 21.188 | 10.877 | 20.775 | -8.997 | 10.269 | 15.642 | 26.309 | 25.100 |
| 14.736 | 13.325 | -18.866 | 20.576 | 17.173 | 25.016 | -0.051 | 23.325 | 18.969 | -16.784 | 18.460 | 6.211 |
| -6.758 | -7.533 | -15.791 | 13.883 | 2.331 | -14.949 | -2.709 | 9.397 | 13.166 | 4.325 | 14.000 | 29.786 |
| 29.434 | 18.813 | 23.499 | 21.857 | -11.845 | 14.155 | 12.348 | 14.538 | 24.715 | 15.452 | 20.236 | 5.002 |
| 17.368 | 15.878 | 21.637 | 16.637 | -21.374 | -10.616 | 7.809 | 6.457 | 11.685 | 22.996 | 3.952 | 13.730 |
| -2.375 | 5.294 | -2.291 | -4.715 | -16.101 | -17.049 | -3.789 | 14.985 | 9.715 | 25.605 | 23.905 | 11.078 |
| 25.647 | 0.980 | -7.108 | -20.681 | 16.182 | 26.486 | 5.316 | 13.930 | 18.559 | 13.206 | 22.665 | 20.051 |
| 27.227 | -6.864 | 11.136 | -7.192 | 15.217 | 19.593 | 8.470 | 22.780 | 13.577 | 16.213 | -2.739 | 9.895 |
| 6.510 | -8.756 | 20.414 | 7.767 | 7.813 | 27.440 | 31.112 | 8.442 | 21.268 | -0.542 | 18.091 | 25.239 |
| 2.014 | 24.318 | 6.524 | -11.077 | -2.240 | 17.341 | 3.700 | 24.292 | 3.240 | -17.496 | -14.938 | 16.131 |
| 13.501 | -23.029 | -10.157 | 18.182 | -5.100 | 3.389 | 5.678 | 21.136 | -24.314 | 15.267 | 5.757 | 17.524 |

| | | | | | | | | | | | |
|---|---|---|---|---|---|---|---|---|---|---|---|
| 0.118 | -29.601 | 2.270 | 10.684 | 0.273 | -5.036 | 27.096 | 18.120 | 9.378 | 8.631 | 2.184 | 12.101 |
| 17.064 | 24.711 | 13.186 | 3.713 | 20.526 | 17.775 | 30.476 | 12.849 | 7.963 | 21.320 | 19.631 | 28.915 |
| 7.833 | -18.610 | 31.101 | -20.340 | 9.424 | -9.114 | 8.498 | 11.793 | 17.949 | 23.745 | -22.780 | 23.626 |
| 27.294 | -19.753 | 17.511 | -17.308 | 2.491 | -21.095 | 7.657 | 6.715 | 6.897 | 11.891 | 20.922 | -2.971 |
| 19.369 | -12.065 | 23.245 | 8.442 | 8.695 | 16.682 | -0.883 | -17.561 | 20.172 | 31.621 | -7.149 | 11.347 |
| -19.583 | -26.487 | -24.570 | 9.855 | 29.007 | -15.043 | 24.325 | 23.722 | 7.124 | 13.038 | -1.097 | 8.627 |
| 22.056 | -17.099 | -6.211 | 18.402 | 13.442 | 10.587 | 18.881 | 11.030 | 10.945 | 18.209 | -20.772 | 19.194 |
| 16.010 | 23.105 | -12.041 | 15.513 | 18.633 | -23.788 | 10.493 | -2.173 | 5.433 | -24.370 | 14.384 | 5.283 |
| 16.403 | 20.393 | -25.625 | -6.017 | 14.187 | 29.351 | 17.967 | 2.097 | 19.877 | 20.926 | 10.618 | 22.620 |
| 7.166 | 16.446 | 14.763 | 5.541 | 11.186 | 28.532 | 9.948 | 19.246 | 31.047 | -8.941 | 24.666 | 14.889 |
| 7.712 | 6.379 | 10.390 | -9.825 | -1.817 | 23.652 | 15.320 | 24.972 | 17.004 | 17.621 | 2.608 | -6.791 |
| -9.940 | 10.045 | 8.341 | 0.208 | -12.920 | -6.662 | 24.086 | 16.658 | 16.108 | 9.094 | 16.431 | 25.587 |
| -23.999 | 13.179 | 24.379 | -23.639 | -5.964 | 7.406 | 13.169 | 14.987 | -18.254 | 12.231 | 17.593 | 9.018 |
| -1.545 | 21.879 | 22.348 | 3.125 | -20.977 | -20.564 | -1.900 | 19.894 | 13.119 | 15.447 | 23.036 | 26.318 |
| -25.996 | 19.331 | 26.837 | -17.078 | -16.215 | -10.791 | 8.549 | 26.880 | 14.371 | 9.839 | 20.080 | -19.654 |
| 25.925 | -21.869 | -14.567 | -15.238 | 23.855 | -16.341 | -12.591 | 25.860 | 10.653 | 16.940 | 4.859 | 24.105 |
| -20.571 | 15.873 | 14.306 | -8.428 | -9.819 | 16.897 | 28.157 | 15.589 | 19.389 | 21.655 | 6.680 | -16.100 |
| 27.841 | 22.554 | 19.844 | 4.826 | 1.277 | -5.495 | -17.156 | -3.538 | 8.657 | 10.892 | 13.401 | 16.454 |
| -9.804 | -3.016 | -12.484 | -19.999 | 18.733 | 9.181 | 16.603 | 21.709 | 20.855 | 23.919 | 22.316 | -6.367 |
| 25.967 | 17.145 | 15.458 | -2.812 | -5.433 | 3.233 | -4.853 | 17.877 | 16.973 | 12.775 | 19.835 | 20.095 |
| -14.554 | -19.878 | -22.488 | 11.661 | 5.693 | -7.272 | -24.911 | 25.259 | 10.435 | -13.506 | -9.855 | 10.152 |
| -6.822 | 27.067 | 3.215 | -0.650 | -2.494 | -19.981 | 20.093 | 10.585 | -12.965 | 24.567 | 15.634 | 21.619 |
| 10.814 | 11.130 | 8.230 | 12.391 | 4.378 | 17.945 | -7.396 | 0.655 | 12.495 | 17.676 | 10.817 | -25.461 |
| -11.105 | 20.730 | 21.765 | -1.291 | 22.608 | -1.787 | 17.556 | 6.320 | 6.355 | 9.477 | 23.613 | 11.637 |
| 12.367 | -4.810 | 24.179 | -4.630 | 18.482 | -4.046 | 29.798 | 7.857 | 14.152 | 18.967 | 16.212 | 22.199 |
| -26.915 | -9.459 | -25.867 | -2.664 | 25.818 | 6.177 | -0.716 | 15.047 | 11.269 | 12.899 | 20.526 | 24.498 |
| -17.062 | 23.528 | -19.447 | 10.517 | 3.778 | -14.395 | 19.023 | 24.243 | 11.956 | 26.110 | 24.292 | -29.640 |
| 3.035 | 13.073 | 28.797 | 7.312 | 14.094 | -0.381 | 11.644 | -0.664 | 18.397 | 14.354 | 18.988 | 28.079 |
| 19.302 | 9.898 | 9.325 | 13.733 | -4.242 | 24.512 | 10.739 | 22.064 | 1.750 | 22.133 | 26.422 | 26.874 |
| 0.523 | -19.235 | 7.394 | -5.930 | -1.383 | -7.023 | 21.197 | 2.205 | 17.331 | 17.640 | 12.110 | 11.139 |
| 26.012 | -0.071 | -10.595 | 3.969 | 6.873 | 14.575 | -3.953 | 6.580 | 12.942 | 22.884 | -24.756 | 29.045 |
| -23.062 | -21.285 | 26.903 | 4.585 | -6.611 | 25.226 | 28.782 | 23.244 | 9.141 | 19.941 | 24.940 | -1.667 |
| 0.594 | 18.581 | 25.596 | 2.508 | -5.544 | 16.134 | 27.464 | 5.852 | -12.531 | 19.584 | 5.057 | 16.031 |
| 21.462 | 12.072 | 3.518 | 24.678 | -24.379 | -12.654 | 0.754 | 20.684 | 0.606 | 12.418 | 23.152 | 12.505 |
| 15.894 | -11.202 | 5.981 | 20.695 | 6.862 | 19.816 | -8.287 | 18.859 | 16.845 | 27.099 | 18.262 | 19.011 |

| | | | | | | | | | | | |
|---|---|---|---|---|---|---|---|---|---|---|---|
| 26.957 | -18.331 | 28.325 | 16.829 | 2.542 | 3.851 | 6.366 | 27.265 | 16.065 | 22.936 | 23.746 | 25.241 |
| 13.661 | -11.767 | -7.717 | 19.752 | 20.003 | -22.577 | -6.064 | 21.554 | -7.795 | 7.930 | -14.899 | 11.709 |
| -20.162 | 27.222 | 7.937 | 22.388 | 16.805 | 12.240 | 25.981 | 25.093 | 7.137 | 16.668 | 0.606 | 24.721 |
| -5.416 | 0.566 | -17.617 | -6.970 | 10.752 | 10.315 | 21.798 | 15.630 | 19.905 | 20.740 | 14.307 | 13.349 |
| -18.651 | 18.278 | 9.886 | 0.207 | 21.782 | -1.534 | 17.778 | 11.798 | -24.931 | 21.372 | 4.911 | 9.225 |
| 9.493 | -18.844 | 13.989 | 22.198 | -2.667 | 22.634 | 9.187 | 8.615 | 10.973 | 19.207 | 5.906 | 18.584 |
| -10.038 | 22.197 | 29.900 | 18.561 | 23.047 | -19.434 | 22.861 | -11.601 | 6.607 | 25.602 | 4.307 | 9.901 |
| 21.348 | 19.628 | -31.236 | -20.153 | -3.455 | 8.222 | 19.968 | 13.721 | 22.331 | 9.422 | 24.094 | 19.935 |
| -3.270 | 16.810 | 5.857 | -0.921 | -1.172 | -25.483 | 14.841 | 2.951 | 18.789 | 13.275 | 7.153 | 27.493 |
| -13.365 | -6.399 | 0.405 | 2.090 | 13.092 | 17.526 | 26.766 | 25.429 | -13.492 | 19.106 | 12.846 | 18.105 |
| 10.480 | 20.743 | -11.231 | 23.716 | 12.082 | 3.093 | 24.119 | 16.698 | -16.126 | 11.575 | 17.720 | -4.998 |
| 1.323 | -18.450 | -28.918 | 12.420 | 25.099 | 19.102 | -2.143 | 7.506 | 19.156 | 10.913 | 15.173 | 16.275 |
| 23.606 | 11.878 | -1.917 | -5.754 | 7.423 | -4.588 | 4.295 | 26.863 | 7.518 | 8.777 | 22.530 | 23.008 |
| 10.999 | 23.816 | 29.252 | -6.607 | -7.035 | 23.508 | -7.485 | 22.281 | 18.902 | 24.006 | -7.164 | 19.995 |
| 10.279 | 16.927 | -0.541 | 6.198 | -6.047 | -17.358 | 5.149 | 24.294 | 19.870 | 22.442 | -2.760 | -26.226 |
| -12.643 | 7.951 | 18.245 | 27.928 | -3.531 | 16.707 | -6.385 | 0.973 | 12.351 | -25.016 | 13.764 | 17.484 |
| 20.791 | -9.556 | 14.835 | -21.122 | -8.509 | 5.455 | 10.384 | 9.307 | 21.130 | 5.414 | -3.966 | 15.207 |
| 8.429 | 3.210 | 26.761 | -0.890 | 13.112 | 24.740 | 6.692 | 22.799 | 20.250 | 8.140 | 25.007 | 25.600 |
| -22.032 | -5.455 | -25.737 | -3.143 | 12.227 | 6.396 | 11.846 | 14.310 | -9.231 | 25.041 | 17.231 | 23.769 |
| 5.504 | -3.324 | -22.264 | -17.899 | 11.451 | 0.825 | 28.607 | 15.552 | 21.511 | 22.607 | 13.203 | 5.928 |
| -0.985 | -10.180 | 15.225 | 8.372 | 22.326 | -9.958 | -7.606 | 16.982 | 22.072 | 26.627 | 5.398 | 10.672 |
| -13.963 | -7.031 | 8.186 | 11.543 | 4.800 | 2.906 | 16.451 | 12.179 | 15.153 | 20.593 | 22.594 | 9.199 |
| -0.957 | -8.207 | 23.793 | 9.249 | -4.085 | -6.753 | 28.067 | 13.699 | -7.666 | -5.733 | 22.021 | -12.003 |
| 13.923 | 21.632 | 17.912 | 16.635 | -5.080 | 17.881 | -1.219 | 28.700 | 19.292 | 26.375 | 22.654 | 11.731 |
| 15.781 | 18.423 | -2.232 | -0.052 | 1.951 | -15.199 | -27.120 | 11.095 | 12.517 | -0.993 | 16.021 | 12.416 |
| -0.190 | -9.607 | -29.772 | 18.453 | -7.154 | -0.590 | 6.958 | 4.258 | -14.260 | 17.608 | -2.919 | 3.873 |
| 3.373 | 29.446 | 27.527 | -1.545 | 8.029 | 23.443 | 8.014 | 8.180 | 13.699 | 23.452 | 16.107 | 23.887 |
| -22.989 | -16.950 | -9.298 | 5.323 | -0.609 | 0.215 | 11.042 | 24.940 | 13.221 | 6.113 | 20.834 | 22.645 |
| -5.126 | 30.779 | 12.729 | 10.268 | 7.290 | 11.135 | -6.268 | 14.410 | -4.700 | 27.114 | 19.077 | -9.180 |
| -15.887 | 17.711 | 17.510 | -4.348 | 25.195 | 8.946 | 25.259 | 10.087 | -13.743 | -1.027 | -13.863 | -18.166 |
| -20.660 | -12.581 | -21.668 | -22.809 | 2.112 | -12.952 | 15.257 | 14.770 | 11.401 | 16.320 | 14.803 | 25.464 |
| 23.426 | 15.250 | 8.884 | 6.165 | -9.833 | -1.585 | 21.033 | 3.333 | 5.430 | 20.171 | 19.610 | 20.247 |
| -11.624 | -21.694 | -8.526 | 9.663 | -11.485 | 26.864 | -2.517 | 8.160 | 9.648 | 21.330 | 20.034 | 11.863 |
| -3.839 | -10.293 | 13.208 | -8.338 | 19.447 | 7.257 | 4.258 | 17.225 | 19.504 | 24.640 | 22.873 | 21.444 |
| -17.544 | 22.575 | -1.476 | 21.307 | 5.555 | 4.858 | -0.965 | 19.351 | 25.012 | 22.155 | 20.817 | 12.423 |

| | | | | | | | | | | | |
|---|---|---|---|---|---|---|---|---|---|---|---|
| -1.256 | 21.237 | 16.982 | 1.245 | 11.566 | 9.933 | -10.013 | 26.429 | 14.640 | 0.728 | 15.814 | 25.748 |
| -18.509 | -24.219 | 25.788 | 8.618 | 9.632 | 12.654 | 23.388 | 18.377 | 12.389 | 28.219 | -8.662 | -25.552 |
| -21.592 | -7.952 | -0.295 | 24.060 | -6.343 | 14.040 | -13.584 | 25.495 | 16.415 | 10.078 | 7.952 | 11.213 |
| -2.953 | 27.740 | -4.485 | -7.172 | 16.333 | 17.550 | 25.944 | 7.320 | -7.127 | -2.947 | -2.129 | 5.659 |
| 9.794 | 12.658 | 22.366 | -16.658 | 17.514 | -17.642 | 24.705 | 15.770 | 13.968 | 26.108 | 16.667 | 24.771 |
| 7.586 | -12.025 | -13.240 | -9.538 | -22.383 | 10.598 | -12.209 | -10.285 | -13.560 | 20.387 | 4.181 | 4.805 |
| -20.858 | -28.095 | 13.604 | -7.697 | 3.898 | -6.893 | -0.165 | -1.449 | 18.826 | 14.344 | 26.592 | 23.985 |
| 14.884 | 20.951 | 10.065 | -11.993 | -2.090 | 3.527 | -2.521 | 8.105 | 25.563 | 27.062 | 25.744 | 19.561 |
| 3.241 | 11.807 | -25.371 | 26.569 | -21.605 | 9.454 | 6.629 | 0.153 | 3.943 | 22.331 | 14.413 | 12.950 |
| 16.192 | 24.610 | 7.454 | 19.382 | 20.540 | -3.884 | -4.001 | 2.537 | 7.167 | 8.912 | -18.911 | 16.628 |
| -0.290 | 16.638 | 11.604 | 16.163 | -4.300 | -8.545 | 8.411 | 2.397 | 1.090 | 16.870 | 0.518 | 16.561 |
| -7.399 | 13.809 | 0.327 | -5.983 | 10.944 | -14.009 | 0.954 | -8.757 | 11.033 | 21.227 | 7.641 | 20.005 |
| 10.546 | 1.878 | 8.920 | -2.050 | 1.423 | 11.522 | 21.556 | -17.616 | 17.106 | 25.348 | 27.490 | -3.619 |
| 3.058 | 16.047 | 24.663 | -2.939 | 1.998 | 13.502 | -9.263 | -10.525 | 5.371 | 23.120 | 10.103 | 23.012 |
| -6.103 | -13.707 | 26.054 | 5.606 | 7.293 | 21.651 | 5.058 | -6.342 | 9.319 | 26.904 | -27.956 | 15.510 |
| 1.690 | 26.832 | -5.172 | -10.952 | 23.195 | -10.478 | -5.932 | 3.558 | 5.954 | 6.694 | -18.946 | 18.165 |
| -7.799 | -20.368 | 5.793 | 6.675 | -9.915 | 24.844 | 5.851 | -15.141 | -1.918 | 8.294 | -4.846 | 20.525 |
| -2.919 | 17.083 | 3.462 | 20.225 | 21.066 | -16.774 | 20.171 | 4.610 | 15.830 | 24.718 | 28.597 | 7.373 |
| 6.718 | 23.153 | 23.155 | -13.426 | 18.918 | 8.389 | 2.397 | 19.357 | 28.062 | 21.164 | 24.963 | 13.513 |
| 4.415 | -17.325 | 26.871 | 3.788 | 13.053 | -18.085 | 16.828 | -16.523 | 10.305 | 1.976 | -18.144 | 6.250 |
| 0.768 | 7.139 | 9.493 | 5.224 | 23.553 | 25.423 | -10.246 | 1.765 | 8.290 | 23.471 | -15.989 | 19.672 |
| 13.000 | -7.880 | 22.458 | 14.987 | 4.280 | -17.018 | 14.354 | 13.202 | 27.196 | 16.000 | 18.974 | 23.645 |
| -4.726 | 21.081 | -12.735 | 11.411 | 24.170 | 0.283 | 22.009 | -13.128 | -0.779 | 17.507 | 13.069 | 3.048 |
| -26.745 | -10.433 | 16.361 | 10.738 | 16.107 | -0.176 | 23.908 | -16.800 | 2.416 | -21.630 | 6.382 | -25.982 |
| -25.615 | 24.283 | -3.577 | 5.714 | 14.879 | 7.580 | 12.267 | -18.239 | 20.017 | 22.127 | 25.170 | 6.564 |
| 24.056 | -14.860 | -23.518 | -5.248 | 18.664 | 3.832 | -6.446 | 25.779 | -7.025 | 25.262 | 27.976 | 17.179 |
| 8.972 | 25.928 | 17.221 | -6.167 | -6.121 | -4.612 | 7.881 | 9.050 | 12.253 | 12.655 | -10.224 | 27.615 |
| 25.898 | 14.364 | 23.880 | 7.450 | -21.739 | -20.265 | -4.589 | 20.082 | -2.008 | 9.066 | -8.436 | -2.621 |
| 21.682 | 9.744 | 22.704 | -2.011 | 7.682 | -20.704 | 25.413 | -26.181 | 16.507 | 7.223 | 16.261 | 18.114 |
| 12.223 | -25.623 | 9.138 | -16.360 | 15.958 | 15.901 | 19.455 | -22.049 | 17.970 | 17.911 | 25.797 | -1.407 |
| 5.680 | 28.829 | 20.589 | -20.340 | 21.806 | 14.182 | 10.403 | 1.111 | -11.789 | 26.226 | 17.578 | -3.812 |
| 6.416 | 19.528 | -5.307 | -6.822 | 12.673 | -5.250 | 8.419 | 25.949 | 8.936 | 21.676 | 27.280 | 25.758 |
| 10.249 | 12.678 | 4.787 | 10.394 | 10.941 | 10.914 | 6.333 | 15.001 | 7.791 | 19.875 | -8.430 | 9.535 |
| -7.747 | 22.164 | 3.823 | -17.052 | -6.346 | -18.902 | 20.130 | 11.374 | -2.563 | 23.970 | 23.920 | 9.543 |
| -6.571 | -6.583 | 7.554 | 3.857 | 24.196 | -17.631 | 22.259 | 20.322 | 9.563 | 3.276 | 3.984 | -3.336 |

| | | | | | | | | | | | |
|---|---|---|---|---|---|---|---|---|---|---|---|
| -14.555 | -18.330 | -22.736 | -6.475 | -14.183 | 12.381 | 10.596 | 14.887 | 17.170 | -6.142 | 13.563 | -6.246 |
| 21.799 | 11.542 | -12.955 | -10.139 | -3.583 | 20.758 | 9.309 | -10.111 | 4.334 | -0.305 | -17.973 | 24.461 |
| -13.656 | 16.647 | 11.775 | -5.667 | 10.406 | 4.526 | 8.094 | 24.761 | 24.951 | -10.929 | 3.572 | 10.634 |
| -26.178 | -27.732 | 13.574 | 17.340 | -8.104 | -8.407 | 3.790 | 10.449 | 9.049 | 16.609 | -22.594 | -17.842 |
| -11.119 | 8.173 | -17.915 | 0.841 | 7.380 | -23.769 | -11.027 | 21.895 | 25.741 | 20.120 | -0.423 | 14.021 |
| 22.708 | 17.661 | 25.287 | 11.640 | -2.323 | 5.346 | -4.029 | 3.540 | 9.725 | 8.510 | 27.913 | 4.716 |
| -28.552 | 19.574 | 1.700 | -8.866 | 5.101 | -9.901 | 9.724 | 25.942 | 5.861 | 0.684 | 25.154 | 22.739 |
| -5.528 | -19.660 | -15.693 | 22.147 | -22.213 | -1.724 | -13.529 | -8.079 | 3.991 | 17.096 | 20.791 | 0.601 |
| -27.940 | 15.070 | 19.317 | 13.099 | 5.903 | 10.739 | 10.080 | 2.111 | 14.106 | 19.908 | 26.692 | 1.726 |
| -8.721 | 17.270 | 17.114 | 24.574 | -1.126 | 20.288 | -14.711 | -1.307 | 6.640 | 10.153 | 13.179 | 12.252 |
| 19.324 | -7.716 | 15.788 | -11.016 | 16.109 | 6.078 | -17.034 | 24.768 | -15.558 | -6.041 | 26.066 | 19.012 |
| -1.055 | -5.340 | 5.391 | -21.062 | 6.193 | -9.623 | 12.207 | 6.477 | -17.282 | 22.897 | 2.384 | 22.029 |
| 4.241 | 18.282 | -24.914 | 15.334 | -9.401 | -0.973 | 22.175 | 26.913 | 12.047 | -9.660 | 5.234 | -1.752 |
| 1.997 | 28.987 | 7.979 | 5.691 | 7.410 | 14.018 | 4.100 | 13.551 | -7.373 | 4.051 | 19.602 | 7.421 |
| 12.407 | 18.571 | -0.794 | 18.256 | 8.862 | 5.842 | 18.973 | 22.859 | 7.577 | -6.327 | 14.888 | 24.043 |
| 7.283 | 25.076 | -22.310 | -0.032 | 19.156 | -9.364 | 20.305 | 28.174 | 11.370 | 3.748 | 23.651 | 18.009 |
| 5.047 | 22.089 | 10.461 | -4.206 | -8.320 | 0.563 | 25.659 | 9.041 | 15.513 | -11.630 | 12.649 | 20.046 |
| 21.788 | 16.382 | -14.848 | 8.979 | 15.496 | 19.002 | 23.092 | 6.405 | 0.608 | -0.698 | 29.667 | -0.651 |
| 12.888 | 21.008 | 19.793 | 14.261 | 18.252 | 11.586 | 3.033 | -23.596 | 17.325 | -11.787 | 22.209 | 16.636 |
| 20.553 | -24.234 | -16.409 | -13.180 | 5.873 | -12.159 | 13.695 | 18.554 | -0.675 | -1.192 | -22.539 | 17.657 |
| 17.639 | 22.918 | 24.452 | -2.069 | -8.989 | 2.059 | -2.281 | -25.455 | 3.550 | -22.561 | 24.226 | 9.087 |
| -28.609 | 10.751 | 14.986 | 22.394 | -10.974 | 23.551 | 5.994 | -1.918 | 4.423 | -3.506 | 16.927 | 1.129 |
| 26.398 | 5.268 | 25.964 | -6.725 | 23.595 | -5.852 | -15.246 | -10.027 | 5.490 | 9.545 | 12.168 | 14.924 |
| -4.058 | 2.339 | 11.368 | -10.757 | 12.879 | 2.867 | 7.252 | 18.758 | -0.768 | 16.913 | 25.309 | -7.198 |
| 2.442 | -23.144 | 16.820 | 17.019 | -14.132 | 20.938 | 12.607 | -3.488 | 15.981 | -9.494 | 28.533 | 9.480 |
| -18.167 | 27.972 | 23.139 | 19.498 | 17.176 | 23.046 | 8.730 | 24.381 | -4.893 | -8.344 | 21.501 | -9.567 |
| 9.906 | 9.090 | -7.792 | 11.250 | -10.041 | 7.768 | 5.471 | 25.677 | -0.516 | 6.469 | 23.270 | 5.709 |
| 23.295 | 7.798 | 12.896 | -3.793 | 2.945 | 16.392 | -3.130 | 27.965 | -2.320 | 17.391 | 25.788 | -6.621 |
| -5.037 | 3.416 | -20.851 | 22.726 | 9.137 | 13.943 | -24.591 | -27.035 | 14.276 | -7.777 | 19.732 | 10.731 |
| -30.723 | 11.500 | 16.419 | 10.191 | 25.361 | 12.279 | 15.197 | 18.509 | 13.616 | 16.668 | 18.346 | 12.620 |
| 28.961 | 6.763 | -7.650 | 1.575 | -7.727 | 24.510 | -12.970 | 26.465 | 15.109 | -6.050 | 30.602 | 11.519 |
| -17.231 | -11.608 | 25.007 | 21.304 | -11.509 | 18.376 | 18.067 | -12.121 | 6.056 | 22.836 | 16.754 | 10.537 |
| 16.138 | 20.253 | -0.750 | 19.136 | 21.639 | -0.680 | 14.332 | 22.511 | 16.738 | 27.029 | -5.407 | -4.684 |
| 17.699 | -3.487 | -3.418 | 12.437 | 5.468 | -14.178 | -13.693 | -19.528 | 16.127 | -10.712 | 23.858 | 11.529 |
| -14.868 | 13.364 | 4.150 | -10.472 | -2.542 | -6.049 | -15.066 | 19.702 | -10.829 | 21.306 | -29.803 | 24.098 |

| | | | | | | | | | | | |
|---|---|---|---|---|---|---|---|---|---|---|---|
| -30.053 | -10.120 | -23.276 | -0.045 | 10.798 | -12.225 | -7.351 | 2.010 | 14.888 | 11.287 | 27.556 | 13.673 |
| -11.597 | -21.391 | -1.471 | 9.197 | -4.971 | 22.501 | 20.209 | 18.154 | 5.640 | 23.840 | 0.822 | 13.130 |
| -14.001 | -4.649 | -3.162 | -5.159 | 12.836 | 21.578 | -16.171 | -24.164 | -12.341 | -24.351 | -24.662 | 23.774 |
| 21.837 | 20.319 | -18.815 | 24.209 | 13.740 | 20.121 | -0.658 | 18.837 | 3.632 | 1.832 | 22.242 | 25.233 |
| 3.974 | 27.839 | 15.211 | 15.972 | 17.648 | 16.765 | 3.194 | 23.977 | -3.033 | 25.315 | 7.727 | 19.773 |
| -29.687 | 18.660 | 10.682 | 16.801 | -3.393 | 8.764 | 11.061 | 12.447 | -20.307 | 27.778 | 19.809 | 6.793 |
| -24.895 | 16.414 | -12.069 | 14.084 | -12.587 | 4.100 | 21.762 | -14.725 | 5.202 | -27.740 | 8.402 | 20.632 |
| 30.737 | 23.789 | -9.038 | -2.115 | 10.131 | 3.586 | 25.186 | 30.336 | -23.617 | 11.527 | 10.876 | 24.407 |
| -19.451 | -9.432 | -8.447 | 5.576 | 24.349 | 5.994 | 5.030 | -5.157 | -5.559 | 10.644 | 3.444 | 2.511 |
| 11.287 | -15.987 | 5.143 | -1.329 | 17.220 | 10.530 | 21.365 | 29.212 | -17.958 | 27.316 | 26.227 | 8.990 |
| 21.147 | 25.003 | -7.636 | 25.299 | 0.285 | -9.830 | 19.447 | 20.081 | 18.668 | 15.787 | 21.706 | 22.809 |
| -30.398 | -8.419 | -13.880 | -0.566 | 2.902 | 18.373 | 8.119 | 25.657 | 10.997 | 28.112 | 28.409 | 9.623 |
| 6.665 | -7.247 | -6.398 | -3.405 | -2.613 | 22.843 | 9.301 | 27.029 | 23.359 | 7.250 | -4.114 | 20.718 |
| -0.037 | 9.957 | -16.035 | 8.198 | 18.988 | 19.360 | 21.203 | 16.675 | 10.344 | -2.967 | 17.349 | 11.884 |
| 14.449 | -11.617 | -16.998 | -10.131 | 0.636 | -25.286 | 5.446 | -28.633 | 11.501 | 21.426 | 6.806 | 7.402 |
| -1.902 | 7.465 | -0.493 | -6.132 | -10.696 | 5.219 | 12.689 | 14.459 | 12.251 | -25.358 | 27.752 | 24.566 |
| -3.078 | 17.812 | -11.489 | -10.680 | -25.674 | 23.681 | 24.406 | 28.096 | 25.299 | 12.653 | 24.324 | -25.837 |
| -18.833 | -4.011 | 5.112 | -8.211 | 22.651 | 25.668 | 17.093 | 27.595 | 17.906 | 10.712 | 26.332 | 16.942 |
| -20.754 | -18.516 | -11.707 | 9.967 | -5.145 | 20.255 | 19.083 | -13.389 | 23.070 | 24.460 | 19.132 | -15.512 |
| -26.349 | 20.623 | -13.512 | -2.560 | 8.975 | 13.214 | -6.636 | -32.338 | 20.785 | -11.675 | 23.127 | 15.974 |
| -10.562 | -14.977 | -19.799 | 26.074 | -16.643 | -1.655 | 1.856 | 30.260 | -14.822 | 27.352 | 18.590 | 3.881 |
| -23.238 | -22.328 | -21.130 | 20.827 | 22.870 | 12.234 | 14.713 | 29.195 | 13.019 | -8.343 | 29.777 | 23.991 |
| 18.905 | 8.870 | 6.215 | 7.351 | -28.244 | 5.000 | 10.608 | -9.912 | -5.624 | -22.799 | -31.876 | 20.412 |
| -12.286 | 6.969 | -19.198 | 6.707 | -3.553 | 20.838 | 7.887 | 26.407 | -21.897 | 14.046 | -25.963 | 25.068 |
| 15.641 | 22.313 | 23.808 | -20.377 | 24.838 | -11.167 | 27.162 | 4.893 | -0.816 | 24.488 | 13.397 | 22.953 |
| 7.937 | -1.285 | 7.707 | 9.486 | 23.154 | 18.816 | 17.749 | 25.289 | 26.466 | 28.961 | 23.446 | 24.637 |
| -0.383 | -10.747 | -3.472 | 2.290 | 16.298 | -27.375 | -6.147 | -20.187 | 23.680 | 29.522 | 16.624 | 9.874 |
| 10.836 | 16.419 | -0.387 | 9.097 | 3.423 | 7.688 | 14.250 | 12.418 | 4.570 | 27.732 | -19.988 | 25.636 |
| -11.484 | -3.010 | 0.100 | 0.426 | 19.980 | 11.738 | 9.784 | 11.067 | 25.368 | -29.017 | 14.910 | 13.594 |
| -21.112 | -0.141 | 13.056 | -20.940 | -17.605 | 26.622 | 18.505 | -25.658 | 12.098 | 25.379 | -24.985 | 22.931 |
| 25.589 | 10.769 | 19.443 | 16.873 | 0.072 | 14.049 | 24.714 | 19.632 | 12.815 | -15.495 | 29.115 | 16.544 |
| 23.959 | -5.273 | 4.429 | 20.667 | -3.881 | 5.693 | -1.630 | 24.710 | -0.249 | 16.033 | 13.084 | 23.586 |
| -8.665 | -8.272 | 9.827 | 0.489 | 18.494 | 0.975 | 5.020 | 29.320 | 4.678 | 27.833 | 24.575 | 19.181 |
| -3.772 | -13.972 | 17.725 | 24.275 | 10.367 | 19.675 | -0.347 | 25.255 | 20.362 | 26.528 | 22.231 | 25.090 |
| 4.367 | 5.462 | 25.142 | 3.851 | -6.114 | 9.844 | 13.572 | -30.250 | 14.407 | 13.207 | 23.374 | -5.237 |

| | | | | | | | | | | | |
|---|---|---|---|---|---|---|---|---|---|---|---|
| 9.469 | -19.820 | -0.840 | -5.495 | -12.500 | 19.013 | 27.362 | 7.842 | 27.046 | 24.221 | -14.550 | 9.229 |
| 12.960 | 20.593 | -3.010 | 15.776 | 27.315 | -3.224 | 14.370 | -23.112 | -23.225 | -7.286 | 21.259 | 1.064 |
| -26.040 | 23.490 | 8.182 | -20.408 | 4.227 | 8.303 | 12.845 | -10.320 | 20.848 | 23.151 | 17.550 | 5.446 |
| 4.924 | 17.796 | 11.745 | -2.858 | 24.396 | 2.799 | -8.535 | 18.973 | 24.741 | -12.875 | 7.695 | 26.388 |
| -18.134 | 12.308 | -24.990 | 10.772 | 21.071 | 25.837 | 26.761 | -13.202 | 23.519 | 24.257 | 19.367 | 21.223 |
| -8.510 | 0.530 | 13.860 | -9.052 | -4.366 | 21.153 | 20.890 | 8.343 | 15.492 | 23.108 | 18.567 | 25.953 |
| -28.864 | 9.220 | -1.937 | 11.472 | -1.732 | -5.689 | 25.943 | 2.141 | 5.725 | 17.368 | 10.635 | 15.598 |
| -24.107 | 14.954 | -8.699 | 19.478 | 17.467 | 10.433 | 9.306 | 23.651 | -9.480 | 13.813 | 10.048 | 28.266 |
| 19.087 | -24.796 | 20.476 | -10.195 | 5.727 | 11.268 | 15.564 | -7.847 | 27.947 | 27.601 | 26.126 | 18.593 |
| -10.534 | 11.384 | 19.233 | -19.914 | 25.843 | -14.466 | 28.884 | 27.766 | -27.940 | 27.566 | 25.178 | 14.032 |
| 5.394 | -13.514 | -7.237 | 2.625 | -6.796 | -6.608 | 3.498 | 6.779 | 10.448 | 30.773 | 15.893 | -2.914 |
| -15.296 | 22.337 | 8.780 | -0.784 | -3.113 | 13.007 | -3.285 | 26.737 | 8.133 | 21.168 | 25.474 | 12.729 |
| -1.486 | -3.186 | 21.480 | 17.622 | 7.863 | -10.146 | 21.859 | -21.681 | 24.807 | 19.715 | 13.287 | -13.571 |
| 22.420 | 16.798 | -26.524 | 14.701 | -3.699 | 17.004 | 11.173 | 16.704 | -29.358 | 18.380 | -28.923 | 21.068 |
| -16.425 | 22.012 | 23.956 | 24.908 | -8.922 | 8.050 | 16.599 | 24.714 | -6.843 | 24.051 | -28.533 | 23.421 |
| 3.099 | -6.487 | -12.709 | 7.541 | 0.428 | -5.877 | -4.946 | 20.251 | 20.688 | -26.079 | 23.395 | -4.889 |
| -26.074 | 20.233 | 2.078 | -1.328 | -29.954 | 17.828 | 29.123 | 8.653 | 17.772 | 29.809 | 24.267 | 18.272 |
| -25.400 | 21.079 | 15.220 | 11.678 | 22.332 | -21.608 | 23.679 | 25.341 | 12.157 | 5.352 | 11.605 | 14.999 |
| -18.492 | 20.108 | -13.982 | 25.847 | 8.709 | -2.584 | 1.909 | 10.129 | 23.795 | -31.754 | 19.402 | 10.435 |
| 15.050 | 16.910 | 9.890 | 8.568 | 26.564 | -20.291 | 12.576 | 4.032 | 22.683 | -22.617 | 23.089 | 8.073 |
| 19.978 | 8.771 | -18.791 | 19.403 | 24.521 | 28.333 | 19.864 | 6.771 | 20.657 | -19.240 | 9.033 | 19.986 |
| -6.932 | -3.079 | 22.282 | 13.820 | 21.683 | -0.356 | 10.412 | -5.998 | 18.882 | -17.182 | 8.056 | -11.661 |
| -13.686 | 22.427 | -9.295 | 2.714 | 26.068 | 11.162 | 21.190 | 0.601 | 21.694 | 28.841 | 26.053 | 30.080 |
| 21.637 | -5.105 | 11.040 | 0.905 | 4.840 | 22.454 | 5.909 | 0.240 | 25.429 | 0.380 | 17.156 | 8.680 |
| -0.276 | 2.038 | -1.956 | 15.852 | 16.798 | 13.818 | 5.479 | 20.138 | 16.677 | 22.760 | 3.549 | 21.109 |
| -9.739 | -6.762 | -12.272 | 7.899 | 11.673 | -17.606 | 26.474 | 25.860 | -12.027 | 11.229 | 20.595 | 12.383 |
| -11.224 | 11.691 | -15.036 | 11.705 | 19.096 | 21.035 | 7.078 | 8.944 | -27.486 | 24.639 | 13.768 | 26.883 |
| 16.380 | -1.715 | 23.578 | -7.284 | -1.322 | 7.518 | 29.786 | 4.156 | -5.386 | 30.788 | 24.848 | 28.984 |
| -16.383 | 20.925 | 26.217 | 13.458 | -2.738 | 23.882 | 4.943 | 22.270 | 19.950 | -30.808 | 10.455 | 7.993 |
| -22.586 | 10.418 | 8.513 | 9.932 | 20.488 | 9.584 | 3.878 | -12.890 | 7.516 | 28.321 | 6.620 | 6.675 |
| -14.286 | 5.354 | -2.799 | 6.198 | -7.647 | 26.015 | -2.628 | -26.760 | 30.166 | 19.951 | 12.950 | 13.469 |
| 13.434 | 22.188 | 21.956 | -2.621 | -20.248 | 1.547 | 8.543 | 23.539 | 23.959 | 17.009 | 24.050 | 21.505 |
| 8.931 | 10.234 | -16.297 | 3.819 | 15.309 | -9.017 | 17.503 | -1.701 | 17.749 | 0.146 | 20.362 | 20.976 |
| 15.438 | -6.409 | -20.626 | -8.541 | 22.872 | 11.724 | -1.971 | -7.865 | 23.099 | 23.901 | 26.693 | -25.475 |
| -21.534 | 4.488 | 2.887 | 23.233 | 18.187 | -2.808 | 22.264 | -2.750 | -11.843 | 24.398 | -19.003 | 29.781 |

| | | | | | | | | | | | |
|---|---|---|---|---|---|---|---|---|---|---|---|
| -0.909 | 12.655 | -7.308 | -0.838 | -27.983 | -11.122 | 28.393 | 20.361 | 16.164 | 5.011 | -5.059 | 19.664 |
| -5.349 | -0.930 | 7.280 | 8.266 | 4.165 | 16.572 | 15.025 | 26.612 | 22.915 | 20.847 | -28.565 | 14.044 |
| 7.200 | -3.659 | -8.172 | 13.533 | -4.561 | 19.524 | 24.405 | -21.521 | 10.980 | 12.371 | 23.373 | 18.430 |
| 10.406 | 10.796 | -14.316 | 27.842 | -5.630 | 14.068 | 8.464 | 4.895 | 29.397 | 27.824 | 4.116 | -6.823 |
| 6.697 | -6.585 | 20.278 | 15.918 | -3.352 | 27.301 | 12.285 | 11.400 | 19.853 | 16.858 | -12.411 | 11.168 |
| 5.182 | 21.668 | -21.774 | -6.758 | -1.513 | -18.452 | 2.033 | 2.545 | -31.360 | 20.431 | 22.587 | -7.584 |
| 1.176 | 16.767 | -22.975 | 21.164 | 23.927 | 22.590 | 22.126 | -1.107 | 24.160 | 17.708 | 0.956 | 4.772 |
| -3.136 | 22.610 | -4.844 | 18.504 | 9.371 | -20.074 | 20.162 | 15.146 | -3.961 | 12.679 | 20.932 | -0.199 |
| -17.771 | 5.911 | -6.093 | -7.400 | -2.059 | -6.388 | 10.625 | -0.097 | 14.103 | 27.632 | 15.642 | 28.813 |
| -6.406 | 2.553 | 27.309 | 4.188 | 25.903 | 19.251 | 23.570 | 23.327 | -3.453 | 21.256 | 19.042 | 10.339 |
| 8.081 | -10.092 | 3.157 | 4.399 | -8.969 | -10.324 | 5.904 | 10.193 | 8.463 | 19.940 | -29.015 | 26.300 |
| 3.694 | 26.724 | 11.710 | -11.335 | 16.248 | 24.153 | 15.970 | 4.906 | -13.692 | 24.686 | 23.912 | 20.464 |
| -0.243 | -0.243 | 19.528 | 2.857 | 14.088 | 9.340 | 16.124 | 18.201 | -7.166 | -7.689 | 25.250 | 11.287 |
| -1.507 | 9.858 | 22.222 | 18.321 | -28.985 | 24.963 | 26.435 | 4.278 | 12.176 | 23.841 | 13.944 | 27.519 |
| 8.516 | -2.021 | 5.217 | 23.060 | -0.319 | 4.830 | 17.894 | -2.154 | 2.556 | 23.003 | 23.418 | 6.951 |
| 5.916 | 16.101 | -23.034 | 12.982 | 29.857 | 4.214 | -8.466 | 13.109 | 21.958 | 18.066 | 0.975 | 31.408 |
| 16.514 | 18.381 | -25.927 | 7.989 | -1.758 | -12.688 | 15.538 | 17.428 | -6.553 | 21.742 | 10.610 | 28.297 |
| 17.823 | 9.115 | -18.645 | -4.526 | 22.416 | -8.862 | 31.230 | 6.673 | -18.897 | 29.297 | 2.530 | 29.931 |
| 7.063 | -27.149 | -13.205 | 5.671 | 10.738 | 16.880 | 4.774 | -2.349 | 29.403 | 11.633 | 8.354 | 13.906 |
| -9.261 | 4.860 | 16.992 | 17.573 | -3.161 | 13.332 | -28.877 | -11.409 | 25.151 | 24.997 | -0.939 | -12.881 |
| -20.372 | 16.991 | 10.006 | -19.081 | -22.866 | 1.777 | 17.773 | -1.756 | 16.494 | 6.568 | 1.875 | -7.477 |
| -12.203 | 7.747 | 27.935 | 20.022 | -1.475 | 6.910 | 25.423 | -8.887 | 20.645 | 24.431 | 12.667 | 25.101 |
| 5.473 | 15.066 | -3.715 | 17.159 | 6.233 | -9.390 | 19.789 | -10.572 | 27.689 | 7.986 | 3.089 | 16.388 |
| -2.923 | 0.552 | 4.845 | -10.121 | -23.196 | 12.011 | 30.606 | 5.397 | 24.429 | -10.717 | 12.110 | 26.900 |
| -17.331 | -11.425 | -19.238 | 8.503 | 28.815 | 6.240 | 23.747 | -4.842 | 23.474 | 19.775 | 20.711 | 8.502 |
| -26.068 | -20.543 | 8.287 | -17.297 | 21.069 | -11.852 | 9.343 | 11.673 | 26.006 | -11.145 | 15.640 | 5.403 |
| 9.512 | -19.343 | 11.601 | -5.297 | 24.266 | 26.605 | -6.850 | -9.481 | 16.086 | 26.648 | 26.072 | 14.773 |
| -19.237 | 5.327 | -15.110 | 18.450 | 30.622 | -2.478 | -15.390 | 12.069 | -2.391 | 16.080 | -14.518 | -7.808 |
| 20.323 | 6.718 | -17.134 | -22.240 | 17.269 | 8.610 | 21.453 | 16.604 | 21.600 | 6.498 | 1.746 | 29.540 |
| -12.557 | 1.424 | 26.983 | 24.877 | -20.617 | 5.295 | 16.499 | 15.395 | 11.894 | 8.553 | 17.925 | 28.460 |
| 26.246 | 19.488 | -12.034 | 16.005 | 25.495 | 17.587 | 28.148 | -22.078 | 8.719 | 25.447 | 2.892 | -27.525 |
| -2.436 | -8.730 | 21.029 | -14.451 | 23.403 | 24.870 | 11.952 | -3.792 | 20.978 | 11.519 | -11.535 | -14.133 |
| -0.953 | 12.268 | 12.767 | -7.862 | -0.716 | -6.027 | 22.309 | -21.447 | 16.708 | 7.512 | -17.504 | -9.036 |
| 20.926 | 21.668 | -27.497 | 21.733 | -13.134 | 11.817 | 20.776 | -11.051 | 15.761 | 21.478 | 25.340 | 15.075 |
| 14.686 | 10.799 | 20.516 | -13.432 | 22.725 | -2.064 | 4.588 | 8.881 | -30.264 | 5.678 | -7.732 | 11.589 |

| | | | | | | | | | | | |
|---|---|---|---|---|---|---|---|---|---|---|---|
| -0.376 | 7.692 | -8.746 | 19.322 | 18.739 | -3.322 | 26.411 | 1.379 | 25.599 | -10.268 | 9.847 | 26.858 |
| -5.238 | 4.057 | -17.250 | 11.088 | -1.536 | -7.014 | 24.843 | 8.504 | -4.269 | -2.120 | -4.558 | 1.567 |
| 22.973 | 11.592 | 14.296 | 20.664 | 6.070 | -17.065 | 18.733 | -7.803 | 10.123 | 16.104 | 22.628 | 17.681 |
| -25.015 | 23.237 | 8.195 | -9.645 | -10.427 | 13.484 | 25.859 | -0.453 | 21.644 | -26.183 | 19.313 | 16.483 |
| 16.523 | 20.821 | 24.324 | -17.352 | 28.568 | 15.546 | 1.156 | -4.726 | 16.031 | -0.043 | 13.200 | 8.345 |
| 23.816 | -11.795 | 27.065 | 20.108 | 22.667 | 24.030 | 22.610 | 11.449 | 4.139 | 4.390 | 25.495 | 25.877 |
| 4.695 | -10.459 | 9.305 | 4.855 | 18.687 | 9.919 | 15.546 | 22.418 | 23.918 | 17.415 | 20.467 | -0.157 |
| 9.475 | 19.588 | -5.717 | 18.435 | 8.506 | 3.477 | 12.986 | 4.236 | 15.181 | 12.950 | 5.503 | 11.128 |
| -16.253 | 0.884 | 22.991 | -5.570 | 16.875 | 14.110 | 29.206 | 23.140 | 16.875 | 18.181 | 14.980 | 14.804 |
| 26.104 | 17.830 | -26.402 | -17.634 | 30.645 | -4.515 | 17.607 | 5.843 | 11.965 | 21.136 | -5.029 | -1.201 |
| 16.383 | 21.752 | -20.578 | -0.690 | -13.156 | -12.914 | 8.093 | 25.600 | 13.725 | 18.762 | 17.481 | 3.803 |
| -14.813 | 26.679 | 6.829 | 22.467 | 19.611 | 6.783 | 27.544 | 11.070 | -22.396 | 24.400 | 14.928 | 16.638 |
| 22.432 | 2.955 | -8.808 | 3.325 | -7.185 | 25.576 | 26.182 | 6.730 | 23.080 | 22.663 | 22.158 | 26.411 |
| 28.491 | 18.850 | 18.438 | 15.724 | -19.958 | 16.908 | 5.813 | 15.217 | 24.827 | 21.829 | -7.745 | 21.939 |
| 9.631 | -9.889 | 23.806 | -14.385 | 7.460 | 4.840 | -4.488 | 8.929 | 2.804 | -24.413 | -12.989 | 25.604 |
| 26.097 | 17.925 | 15.644 | -16.466 | 6.684 | -16.810 | 24.877 | 20.092 | 12.006 | 2.927 | 25.002 | 19.276 |
| -25.188 | 16.699 | 11.066 | -12.519 | 27.416 | 12.516 | 19.810 | -1.886 | 19.374 | -3.254 | -25.690 | 9.424 |
| 13.179 | 28.599 | -13.641 | 12.827 | -3.517 | 18.339 | 21.564 | -22.369 | -18.013 | 21.648 | -1.870 | 26.271 |
| -14.587 | 9.884 | 22.553 | 20.231 | 14.348 | 15.071 | 28.036 | 25.227 | -20.399 | -3.913 | 12.160 | 6.469 |
| -18.480 | -16.456 | -15.910 | -4.975 | 25.551 | -24.511 | 11.181 | 21.004 | 19.061 | -4.542 | -13.527 | 23.804 |
| 27.911 | 6.086 | -9.322 | 6.320 | -15.859 | 19.244 | 9.133 | 18.321 | -1.720 | 22.850 | -11.587 | 26.813 |
| -13.086 | -6.835 | -5.649 | 1.233 | 6.314 | 9.373 | 28.388 | -7.645 | 26.595 | 11.983 | -4.063 | -24.019 |
| -27.884 | 19.858 | -10.960 | -14.105 | -18.141 | 20.059 | 22.610 | -3.724 | -13.968 | 20.514 | -7.320 | 20.278 |
| -19.223 | 15.614 | -9.928 | -6.153 | 7.816 | -13.628 | 18.306 | 11.806 | -16.070 | 10.183 | 10.711 | 21.576 |
| 13.364 | -13.994 | -30.150 | 15.904 | 5.351 | 2.947 | -7.280 | 25.930 | 24.644 | 2.282 | 12.686 | 13.787 |
| -30.315 | 20.758 | 5.307 | 16.744 | -22.479 | 5.207 | -5.672 | 5.281 | -9.458 | 23.625 | -19.585 | 23.360 |
| -24.859 | 2.377 | 24.314 | 21.465 | 20.826 | -4.806 | 19.758 | 21.409 | 20.334 | -20.867 | 14.017 | 1.140 |
| -13.719 | 25.659 | -20.514 | -25.564 | 11.440 | -15.818 | -4.843 | -6.852 | 26.790 | 19.142 | 3.923 | 26.517 |
| -22.196 | -10.951 | -14.947 | 7.787 | -29.668 | 7.147 | 12.266 | 23.308 | 23.329 | 13.894 | -2.800 | -9.879 |
| 1.932 | 22.778 | 15.136 | -15.066 | 22.246 | 9.909 | 11.389 | 11.215 | 0.250 | 14.327 | 6.609 | -15.721 |
| 13.951 | -16.729 | -23.932 | 9.788 | -27.790 | -6.393 | 21.435 | 6.727 | 18.483 | 5.455 | -6.499 | 20.279 |
| -4.585 | 11.434 | 20.338 | -18.939 | 23.138 | 24.411 | 17.065 | -21.860 | 17.814 | 19.146 | -5.301 | 16.577 |
| -19.318 | 7.786 | -13.155 | 6.709 | -18.042 | 14.268 | -0.415 | 2.501 | -20.673 | -2.845 | -14.862 | -5.145 |
| -24.581 | -15.182 | 8.378 | -6.561 | 6.446 | -22.489 | 10.419 | 25.858 | -1.172 | -16.283 | 10.989 | 9.187 |
| -29.529 | 8.427 | 30.368 | 22.186 | 19.086 | 21.648 | 27.237 | -2.655 | 24.345 | -22.534 | 16.657 | 22.534 |

| | | | | | | | | | | | |
|---|---|---|---|---|---|---|---|---|---|---|---|
| -11.023 | -12.459 | 25.597 | 10.072 | -3.335 | -7.681 | -5.843 | 27.971 | 6.051 | -23.938 | 12.857 | 16.230 |
| 29.242 | 17.318 | -4.900 | 4.592 | 14.597 | 26.000 | 25.059 | -2.380 | -26.427 | 22.447 | 14.520 | -0.818 |
| -27.966 | 1.125 | -28.483 | 16.308 | 12.232 | -5.243 | 19.056 | 16.286 | 19.217 | 23.609 | -0.140 | -23.883 |
| 4.529 | -18.949 | 27.166 | -17.840 | 26.664 | 28.211 | 16.523 | -9.228 | 25.399 | 20.669 | 24.261 | -24.202 |
| -1.847 | 13.068 | 24.099 | 4.110 | 9.577 | 11.539 | 24.178 | 12.278 | 24.475 | 1.417 | 12.428 | 13.944 |
| 28.570 | -9.147 | 9.600 | -12.604 | 17.617 | -14.037 | 7.221 | 25.268 | 22.277 | 14.246 | 20.285 | 20.516 |
| 25.585 | 18.354 | 28.008 | -10.227 | 25.006 | 16.379 | -22.049 | 27.178 | 18.728 | 19.566 | 25.245 | 17.738 |
| -15.525 | -6.221 | 20.529 | 19.264 | 11.660 | -0.132 | 11.167 | -1.795 | 23.759 | 5.285 | 22.610 | 24.737 |
| 30.528 | 6.383 | 13.404 | 23.077 | 9.167 | -9.047 | 17.468 | 24.583 | 0.249 | 6.167 | 12.206 | 4.034 |
| -20.331 | -10.028 | -5.019 | -16.609 | 6.605 | 13.709 | 18.262 | -19.266 | -10.028 | 22.855 | 16.598 | -7.738 |
| -3.738 | -2.582 | -6.049 | 10.294 | 18.972 | 10.996 | 22.047 | 6.207 | 21.440 | 16.680 | -8.825 | -19.382 |
| 26.955 | -11.318 | -15.257 | -6.364 | 17.221 | -2.144 | 3.447 | -14.864 | -18.971 | 20.391 | 26.294 | 17.668 |
| 21.509 | 30.643 | 16.950 | -7.439 | 22.938 | -7.086 | 8.514 | 21.465 | -19.263 | 0.069 | -19.414 | 11.979 |
| -24.478 | 20.115 | -18.352 | 3.994 | 9.342 | -16.655 | 20.188 | 18.651 | 24.149 | 7.434 | 5.323 | 20.271 |
| 27.265 | 2.311 | -10.151 | -4.592 | 0.628 | 1.854 | -24.922 | 7.090 | -14.209 | 20.055 | 25.243 | 21.026 |
| -4.176 | 12.082 | 26.774 | -0.766 | 12.957 | -11.614 | 13.090 | 3.300 | 19.493 | 18.433 | 17.871 | 9.421 |
| -30.126 | -19.625 | -4.550 | -8.622 | 16.338 | -18.488 | -20.903 | 26.511 | 3.584 | 13.004 | 20.488 | 13.055 |
| -23.258 | 6.373 | 8.560 | 16.405 | 17.655 | 23.341 | 20.704 | 25.184 | -19.407 | 16.747 | 24.000 | 25.290 |
| 29.914 | -13.045 | 22.748 | 7.265 | 24.159 | -14.950 | -3.837 | -5.495 | 18.356 | 25.348 | 7.253 | 11.487 |
| -23.965 | 23.442 | -15.948 | 11.775 | -7.531 | 10.162 | -5.167 | 26.414 | 13.862 | 21.904 | 2.713 | 18.288 |
| 26.563 | -20.743 | -17.348 | 17.653 | -0.233 | 13.448 | 9.666 | 19.739 | 12.450 | 24.144 | 12.382 | 24.556 |
| -0.678 | -25.108 | -28.762 | 24.101 | 18.089 | -4.473 | 22.795 | 14.288 | 15.015 | 11.834 | -19.725 | 21.871 |
| -17.484 | 29.846 | 12.694 | -11.203 | 2.317 | 21.334 | 14.995 | -4.504 | -24.186 | 22.326 | 13.364 | 9.029 |
| 18.617 | 13.904 | 31.674 | 5.945 | 5.330 | 25.385 | 11.829 | -13.376 | -9.703 | 16.361 | 19.283 | 8.498 |
| -18.199 | -13.582 | 3.549 | 10.343 | -12.086 | -5.948 | -0.920 | 26.665 | 6.846 | 19.831 | 10.213 | 17.959 |
| -26.434 | 2.496 | 1.365 | 2.705 | 18.963 | 5.230 | 25.503 | -29.048 | -16.401 | 5.821 | 25.663 | 11.477 |
| -31.669 | 3.287 | -12.919 | 21.904 | 0.876 | -6.867 | 26.059 | 25.013 | 19.411 | -20.198 | -15.349 | 24.252 |
| 24.690 | 18.321 | -8.226 | 11.619 | 23.393 | 16.252 | 4.893 | -2.238 | 25.041 | 26.060 | 9.892 | 25.296 |
| 27.401 | 4.721 | -12.101 | 15.266 | -0.745 | 0.140 | 6.111 | -18.939 | 16.370 | 21.443 | 26.373 | 10.889 |
| -13.537 | 6.174 | -6.154 | -20.970 | 16.176 | 4.226 | -10.200 | 21.506 | 23.725 | 25.078 | 22.494 | 11.794 |
| -3.419 | -14.215 | 7.248 | 5.765 | 12.796 | 23.898 | 14.398 | -14.986 | 3.756 | 18.097 | -1.361 | -2.824 |
| -24.241 | 7.552 | -9.474 | -7.149 | 25.029 | 3.105 | 17.549 | 26.380 | 20.674 | 25.704 | 26.846 | -1.309 |
| -26.919 | -4.871 | 26.747 | -8.247 | 10.505 | 12.453 | -13.951 | 2.559 | 13.695 | -27.344 | 16.502 | 26.195 |
| -28.525 | 6.434 | -19.027 | -12.706 | 7.834 | -0.845 | 20.427 | -4.071 | 23.469 | 21.262 | 20.677 | 22.916 |
| 20.462 | -16.539 | 26.598 | 9.034 | 0.476 | 9.495 | 18.896 | 29.502 | 26.191 | 9.297 | 23.316 | 16.802 |

| | | | | | | | | | | | |
|---|---|---|---|---|---|---|---|---|---|---|---|
| -26.597 | -9.766 | 16.445 | -9.655 | -24.189 | -4.982 | 1.170 | 15.789 | 22.777 | 1.779 | 4.525 | -13.829 |
| 26.992 | 17.850 | 9.029 | -11.719 | -14.451 | -9.760 | 11.817 | 22.277 | -2.817 | 22.553 | -20.235 | -12.418 |
| -12.851 | 4.243 | -16.176 | -14.843 | 7.375 | 18.840 | 8.757 | 18.067 | -3.319 | 13.883 | 27.679 | 9.238 |
| 23.465 | 11.047 | -22.043 | 3.409 | 13.383 | -14.459 | 9.614 | -25.671 | 26.683 | 9.525 | 14.986 | 27.742 |
| 17.810 | -3.455 | -4.862 | -20.603 | 26.390 | 2.099 | -10.788 | 19.315 | 19.224 | 24.977 | 8.063 | 23.946 |
| 28.573 | 19.349 | 7.148 | 9.706 | -4.671 | -8.889 | 7.247 | -7.718 | 16.701 | 24.332 | 1.090 | 20.932 |
| 0.102 | 9.898 | 2.791 | 22.599 | 18.181 | 22.136 | -26.693 | -14.474 | 1.208 | 21.441 | 25.652 | 18.495 |
| 9.137 | -20.603 | -8.035 | -8.526 | -7.833 | 11.487 | 25.717 | 21.618 | -27.390 | -21.371 | -6.514 | -21.817 |
| -2.960 | -20.174 | -22.859 | -5.368 | 13.592 | -19.485 | -5.960 | 26.284 | 14.118 | 15.727 | -19.071 | 1.509 |
| -28.242 | -28.980 | 8.746 | 20.757 | 17.360 | 16.519 | 6.220 | -15.109 | -5.251 | 18.096 | 19.627 | 21.914 |
| 21.305 | 0.407 | -29.223 | -22.992 | 23.503 | -18.710 | -18.145 | -2.480 | 23.090 | 13.361 | -26.807 | -21.925 |
| -19.423 | 26.656 | 28.600 | 10.798 | -6.306 | -7.186 | -1.211 | -24.772 | -26.276 | 3.581 | 24.122 | 16.688 |
| -11.630 | -16.624 | 22.007 | 4.022 | -6.003 | 21.426 | 12.231 | -4.581 | 24.895 | 6.922 | 16.805 | 15.473 |
| -13.592 | -18.316 | -11.793 | -10.397 | 19.140 | 1.071 | -2.948 | 16.856 | 22.327 | 2.099 | 23.044 | 21.153 |
| 24.388 | 15.962 | 26.461 | 23.348 | -21.260 | -10.989 | 18.763 | 15.598 | -12.856 | -18.258 | -25.918 | 3.532 |
| -17.274 | 20.374 | -10.189 | -0.029 | 25.994 | -0.197 | -0.021 | 30.342 | -24.548 | 28.056 | 21.219 | 22.418 |
| 25.940 | -10.141 | -19.939 | 4.976 | -9.610 | 17.667 | 2.604 | 25.629 | 10.183 | -5.322 | 0.241 | -19.864 |
| 25.487 | 11.359 | -7.153 | 2.239 | -0.456 | 7.905 | 11.457 | 23.564 | 0.324 | 23.115 | 28.204 | 27.065 |
| -16.234 | 15.152 | 17.047 | -4.372 | -15.255 | -20.151 | 17.610 | 14.463 | 23.770 | 21.007 | 8.561 | 24.502 |
| 23.753 | 17.951 | 0.382 | 3.353 | -12.627 | 25.312 | 3.572 | 13.509 | 20.408 | 2.090 | 25.317 | 11.372 |
| 14.198 | 3.411 | 5.657 | -2.589 | 18.389 | 4.605 | 23.226 | 27.494 | 24.749 | 19.278 | 23.963 | 1.375 |
| -23.075 | 19.772 | -4.257 | -13.971 | 10.456 | 1.764 | 25.038 | 20.352 | 13.742 | -1.089 | 21.731 | 18.312 |
| -28.086 | 23.648 | 11.163 | -7.131 | 22.870 | 13.672 | -17.780 | 9.533 | 26.205 | 21.375 | 24.417 | -7.898 |
| 27.647 | 25.365 | -9.790 | -15.560 | 12.725 | 10.693 | 13.726 | -7.269 | 4.781 | 17.295 | 4.942 | 23.150 |
| 17.907 | -6.857 | -21.041 | -17.939 | 0.068 | -11.631 | 4.288 | 24.346 | 16.245 | 23.965 | 29.018 | -1.230 |
| -11.540 | -0.910 | 24.967 | -3.784 | 19.811 | -9.416 | 7.594 | 28.990 | -13.663 | 3.742 | 9.374 | 22.111 |
| -25.516 | 21.661 | 28.729 | -7.371 | -5.229 | -20.775 | -10.337 | -5.567 | 25.724 | 2.781 | -24.308 | 4.672 |
| -27.537 | -8.234 | 22.065 | 1.334 | 17.588 | 21.958 | -5.535 | 25.330 | 11.638 | 9.880 | 25.293 | 28.646 |
| -16.415 | -1.420 | 22.708 | -0.440 | 6.660 | 16.222 | -5.181 | 2.251 | 23.829 | 17.489 | 26.483 | 20.607 |
| 24.287 | -25.503 | 18.238 | 10.496 | 14.322 | 0.866 | -11.318 | 23.514 | -12.183 | 21.751 | -7.428 | 0.740 |
| -0.346 | 6.670 | 16.394 | -10.201 | 3.341 | 11.224 | 20.839 | -0.835 | 24.561 | -0.218 | 9.657 | 14.178 |
| 12.492 | 14.935 | -19.841 | 1.251 | -0.542 | -13.092 | 13.723 | 21.280 | 10.631 | 11.058 | 21.150 | 7.667 |
| 18.881 | 13.955 | -6.930 | -2.163 | 21.949 | 9.128 | 22.337 | -7.304 | 26.278 | 19.710 | 5.559 | 27.227 |
| -26.195 | 8.212 | 4.795 | 2.073 | 2.347 | 0.766 | 5.679 | 25.898 | 29.347 | 17.582 | 16.850 | 18.782 |
| 14.800 | -23.706 | 11.786 | 2.989 | -2.892 | 15.015 | -6.364 | 17.849 | 21.858 | 10.609 | 12.584 | 24.175 |

| | | | | | | | | | | | |
|---|---|---|---|---|---|---|---|---|---|---|---|
| 25.569 | 1.025 | -11.998 | -22.057 | 15.725 | 3.234 | 18.716 | 26.586 | 14.157 | 18.308 | 25.167 | 21.861 |
| -12.292 | -0.808 | -12.702 | -11.022 | 4.590 | -6.775 | 20.311 | 12.113 | -0.492 | 2.059 | 4.893 | 23.602 |
| -6.371 | 26.177 | -2.271 | 25.961 | 11.350 | -3.155 | -0.237 | -11.538 | 24.463 | 27.790 | 15.763 | 24.682 |
| 22.831 | -20.776 | -1.266 | -12.688 | -1.665 | -15.152 | -2.344 | -14.200 | -3.430 | 17.583 | 9.948 | 26.620 |
| 21.358 | -0.177 | -18.411 | -8.404 | 9.399 | 16.138 | 19.196 | 31.261 | 14.812 | 16.845 | 15.214 | 14.579 |
| -11.895 | 21.549 | 21.793 | 8.478 | 10.838 | 13.280 | -9.742 | 20.593 | 3.700 | 29.821 | 11.669 | -24.274 |
| 24.856 | 24.632 | 20.144 | 4.309 | 6.703 | 26.833 | 18.233 | -8.619 | -6.730 | 13.629 | -19.085 | 20.350 |
| -16.480 | 10.294 | 16.476 | 1.727 | 3.292 | 14.168 | 9.177 | -17.279 | 21.059 | 20.559 | 8.634 | 7.317 |
| -17.485 | -18.462 | 27.518 | -6.557 | 2.353 | 2.650 | 11.272 | 11.590 | 25.582 | 12.731 | 13.675 | 22.654 |
| -1.641 | 0.593 | -16.736 | -7.661 | -6.407 | -7.622 | 11.660 | 19.701 | 0.277 | 23.827 | 19.468 | 5.575 |
| 16.796 | 6.193 | -5.606 | -3.075 | -8.065 | -9.530 | 10.113 | -29.997 | 23.994 | 29.167 | 21.433 | 2.603 |
| -17.132 | 6.628 | 22.791 | -25.479 | -2.189 | -14.506 | 10.905 | -15.935 | 1.079 | 25.699 | -6.757 | 19.874 |
| -24.478 | 1.915 | -0.911 | 24.658 | -8.489 | 10.923 | 8.234 | 21.424 | 22.472 | 13.966 | 28.790 | 21.294 |
| 5.438 | -1.843 | 18.505 | -5.095 | -19.006 | -1.806 | 2.325 | -5.323 | 24.253 | 31.136 | -6.147 | 9.384 |
| -27.052 | 16.748 | 26.520 | -16.508 | -5.955 | -18.992 | -0.980 | 30.079 | -6.439 | 23.059 | -23.647 | 5.011 |
| -17.937 | -4.817 | 2.017 | -23.217 | -7.695 | 24.601 | -0.058 | -15.221 | -22.505 | 21.386 | 18.070 | 23.555 |
| -22.526 | 14.269 | -0.002 | -2.258 | 7.157 | 17.962 | 23.572 | 27.481 | -1.312 | 5.319 | 28.031 | 7.433 |
| 17.088 | 23.666 | 8.865 | -5.332 | -11.142 | 7.248 | 19.900 | 16.896 | -3.587 | 10.355 | 24.034 | 18.999 |
| 16.411 | 26.194 | -25.777 | 5.608 | 5.025 | 11.075 | 25.850 | 28.671 | 17.273 | 26.985 | 20.135 | 29.518 |
| -26.460 | -0.312 | 22.036 | -8.199 | -5.573 | -8.460 | 24.783 | 9.966 | 7.172 | 25.995 | 29.831 | -21.149 |
| -13.653 | 7.062 | 16.811 | 9.868 | 7.334 | 3.808 | 7.973 | 4.536 | 25.288 | 18.947 | 26.275 | 25.535 |
| 19.453 | 2.454 | 25.640 | 11.942 | -5.738 | -12.547 | 16.817 | 30.346 | 13.799 | 4.872 | 27.188 | 24.362 |
| -6.012 | 19.240 | -3.740 | 25.840 | 0.844 | -23.246 | 14.537 | 31.027 | 4.241 | 27.346 | -25.253 | 27.421 |
| -13.754 | 8.766 | 12.715 | -18.912 | -14.190 | 6.116 | 3.289 | -25.382 | 11.256 | 16.503 | 17.035 | 9.431 |
| 11.845 | -3.084 | 6.451 | -12.110 | 17.335 | -6.431 | 8.614 | 18.600 | 11.990 | 15.945 | -10.846 | 6.828 |
| -12.101 | -5.780 | -14.452 | -24.481 | 9.754 | -1.008 | 1.347 | 11.365 | 16.943 | 21.534 | 30.917 | 26.076 |
| 17.499 | 21.439 | -12.579 | 25.025 | -26.106 | -4.920 | -8.730 | 26.076 | -1.706 | 8.613 | -6.467 | 17.476 |
| -16.155 | 8.654 | 17.072 | 2.359 | 15.066 | 2.434 | 24.237 | 5.971 | -0.921 | 5.253 | 24.743 | 15.275 |
| -1.432 | -1.651 | -16.311 | -22.532 | 21.024 | 19.351 | 18.267 | -0.103 | 16.516 | 6.706 | 30.471 | 27.734 |
| 16.274 | -24.997 | -11.527 | 11.482 | 23.437 | -3.151 | 22.468 | 11.055 | -9.066 | 30.010 | 24.449 | 14.562 |
| 24.447 | 19.844 | 25.106 | -16.192 | 6.034 | 0.149 | 1.968 | 29.297 | 20.702 | 25.727 | 19.903 | 21.565 |
| -23.311 | 1.952 | 9.089 | -2.755 | 4.270 | 14.531 | 4.499 | 22.785 | 21.763 | 7.078 | 12.620 | -27.783 |
| 7.562 | 13.352 | 7.401 | 3.115 | 17.351 | -6.929 | -13.780 | -16.069 | 15.000 | 23.890 | 23.688 | 3.618 |
| 6.985 | 20.781 | -8.160 | 17.069 | 2.442 | 21.674 | 12.119 | 23.566 | 5.311 | 30.926 | 29.355 | 30.312 |
| 1.829 | -18.779 | 6.229 | -1.704 | 10.874 | 10.129 | 20.060 | 2.693 | -6.767 | 15.614 | 30.032 | 25.660 |

| | | | | | | | | | | | |
|---|---|---|---|---|---|---|---|---|---|---|---|
| -8.329 | -12.314 | 2.591 | 17.442 | -2.392 | -1.550 | 24.980 | 26.459 | 0.897 | 20.122 | 25.906 | 0.215 |
| -9.526 | 9.455 | 1.841 | 23.141 | -7.863 | 4.847 | 25.991 | -26.805 | 22.916 | 16.988 | 8.726 | 12.684 |
| 15.516 | 3.722 | -1.331 | 4.621 | -12.566 | 5.868 | -2.556 | -16.508 | 21.825 | -1.642 | 22.666 | -1.866 |
| -10.051 | 5.011 | -9.143 | 6.637 | 1.029 | -2.687 | 13.579 | -20.122 | -5.684 | 18.078 | 28.665 | 23.382 |
| -24.598 | 10.505 | -4.821 | -3.764 | -1.124 | -7.257 | 3.005 | 13.749 | 30.509 | 13.918 | 2.280 | 17.753 |
| 19.526 | 4.660 | 7.065 | 7.670 | 9.814 | -3.677 | 24.263 | 11.506 | -9.753 | 22.987 | 1.420 | 18.381 |
| -0.987 | -12.956 | 15.920 | 13.801 | -11.416 | 5.426 | 26.664 | 18.780 | 14.298 | -0.184 | 30.053 | 13.194 |
| -4.897 | 15.813 | 3.514 | -17.897 | 25.591 | -24.317 | 6.444 | 28.804 | -4.557 | -0.508 | 9.405 | 25.225 |
| -1.945 | 13.511 | -11.673 | 0.571 | -14.652 | 9.440 | 16.322 | -5.454 | 25.283 | 8.014 | -28.434 | 19.403 |
| 22.475 | -16.082 | 14.587 | 5.354 | 7.601 | 6.857 | 23.795 | 21.980 | 19.564 | 22.028 | 20.342 | 0.234 |
| -17.219 | -19.382 | -12.207 | 3.706 | -6.643 | 20.071 | 25.590 | 23.923 | -8.272 | 28.269 | 28.120 | 20.175 |
| -20.200 | -14.814 | 22.816 | -2.402 | -9.418 | 1.134 | 15.807 | 27.134 | 10.126 | 8.768 | 9.872 | 18.667 |
| -6.954 | -16.938 | -9.217 | 4.134 | 17.057 | -6.917 | 5.340 | 10.536 | -4.448 | 18.481 | 16.728 | 24.803 |
| -10.758 | -16.142 | -13.847 | -10.173 | 5.148 | -11.081 | 10.681 | 17.431 | 23.416 | 20.895 | -27.291 | 14.511 |
| -14.604 | -22.464 | -20.259 | -2.927 | -18.937 | -16.100 | 24.796 | -4.321 | -22.346 | 16.376 | -12.687 | 22.335 |
| 21.528 | 19.985 | -18.328 | 16.993 | 3.036 | 0.471 | 21.938 | -0.596 | 9.469 | 20.416 | 28.998 | 26.577 |
| -6.218 | 20.446 | -11.328 | 9.062 | -4.243 | -8.877 | 3.604 | -10.949 | 20.984 | 0.372 | 22.753 | 28.144 |
| 7.421 | 16.843 | 15.404 | -9.646 | 18.502 | -6.768 | 4.140 | -12.169 | 4.752 | 5.838 | 26.730 | 30.468 |
| -25.404 | 17.380 | 10.537 | 17.866 | 10.648 | -14.816 | 0.074 | 22.437 | -13.107 | 11.130 | 7.660 | 17.514 |
| 8.967 | -8.475 | -10.093 | -5.480 | 25.492 | 3.478 | 15.102 | 19.221 | 16.109 | 15.127 | 26.293 | 27.232 |
| -14.422 | -22.161 | 11.771 | -15.231 | 22.357 | -5.639 | -5.027 | 22.050 | 15.232 | 21.372 | 14.296 | 21.984 |
| -18.895 | 1.503 | 20.015 | 12.811 | -3.859 | 4.797 | 24.346 | 1.286 | -1.296 | 22.196 | -22.702 | 12.646 |
| 2.835 | -7.397 | 21.808 | 19.668 | 20.557 | 17.108 | 18.053 | -19.742 | 26.909 | 23.538 | 22.552 | 19.277 |
| -11.871 | -22.149 | 20.804 | 4.628 | -0.936 | -5.875 | 4.990 | 11.622 | 10.240 | 16.830 | 19.666 | -7.548 |
| -22.368 | 23.044 | -1.097 | -21.990 | 0.725 | -2.326 | 9.842 | 26.674 | 0.229 | 14.034 | -12.334 | 1.422 |
| 21.407 | 0.480 | 25.054 | 16.526 | 7.986 | 10.805 | 21.505 | 9.902 | 13.699 | 13.199 | 27.325 | 5.916 |
| -4.619 | 13.603 | -2.561 | 14.583 | 12.952 | -4.145 | 10.721 | 28.241 | 24.541 | 20.370 | 23.443 | -12.979 |
| -13.182 | 21.801 | -10.653 | 9.723 | -2.670 | 8.818 | 25.659 | 12.091 | -28.124 | 24.630 | 16.392 | 25.145 |
| 17.197 | 5.257 | 4.050 | 3.865 | -8.976 | -21.059 | 6.545 | 23.867 | 2.424 | 14.826 | 27.643 | 16.356 |
| -24.721 | -7.777 | 14.468 | -10.965 | 24.710 | 14.278 | 13.388 | -2.113 | -12.381 | 18.187 | 10.984 | 27.377 |
| -2.411 | -4.674 | 3.395 | 11.083 | -12.555 | 6.218 | 8.585 | -5.768 | -22.248 | 22.753 | 20.010 | -1.934 |
| 16.120 | -10.128 | -23.219 | -12.554 | 20.102 | 22.706 | 26.906 | -20.575 | 19.560 | 8.341 | 12.065 | 22.402 |
| 26.471 | -15.706 | -8.535 | -13.460 | 14.838 | -1.126 | 4.606 | 18.385 | -20.238 | 12.036 | -10.399 | 28.326 |
| -20.637 | 11.137 | 2.617 | 1.675 | -6.239 | 11.943 | 27.523 | 28.119 | -2.504 | 19.279 | 28.370 | 23.582 |
| -15.933 | 22.339 | -14.239 | 15.982 | 17.996 | -12.241 | 16.917 | -1.413 | 9.626 | 14.504 | 10.260 | 21.493 |

| | | | | | | | | | | | |
|---|---|---|---|---|---|---|---|---|---|---|---|
| -0.086 | 21.165 | 4.029 | 1.351 | 14.169 | -0.356 | 11.088 | 23.135 | 27.983 | 9.542 | 25.690 | 15.495 |
| 24.967 | -6.941 | -5.144 | -8.854 | 7.028 | -7.178 | 2.104 | 1.168 | 15.846 | 18.121 | 20.845 | 0.853 |
| -19.993 | 14.002 | -9.609 | 14.614 | 23.369 | -19.430 | 7.303 | 10.323 | 21.510 | 22.934 | 13.742 | 11.034 |
| -23.770 | 3.087 | -13.461 | -19.143 | 13.919 | -16.143 | -5.889 | 27.107 | 4.321 | 8.605 | 2.404 | 2.690 |
| -4.070 | -22.602 | -18.906 | 11.177 | -18.680 | 25.344 | 16.530 | -2.948 | 19.054 | 20.596 | 14.446 | 26.114 |
| -2.406 | 6.215 | -6.528 | 25.078 | 17.678 | 3.466 | 7.606 | 8.315 | 9.640 | 24.093 | -2.330 | 24.988 |
| 15.236 | 5.580 | 3.458 | -14.391 | 5.697 | 15.801 | 28.268 | 10.779 | -13.732 | 13.690 | 17.193 | 13.130 |
| -6.148 | 17.899 | 6.554 | 18.326 | -22.973 | -2.794 | 5.826 | 6.413 | 27.381 | 18.646 | 12.355 | 10.175 |
| -1.002 | 8.471 | 4.242 | -9.940 | 20.420 | -4.160 | 29.002 | -13.295 | 10.575 | 25.397 | 29.552 | 26.717 |
| -21.344 | 15.729 | 24.071 | 8.940 | -5.232 | -23.392 | 20.609 | 2.197 | -0.073 | 22.575 | 16.102 | 28.711 |
| -2.917 | 12.552 | 0.246 | 6.098 | 3.797 | -12.696 | 9.697 | -1.399 | -23.317 | -26.297 | 21.527 | 7.211 |
| -23.675 | -1.169 | -7.478 | -12.470 | 10.338 | 9.303 | 19.931 | -0.695 | -13.406 | 9.406 | 8.857 | 22.576 |
| -18.716 | 1.786 | -8.782 | 16.360 | 19.175 | 11.325 | -8.004 | 10.180 | 26.053 | 22.114 | -8.799 | 23.683 |
| -22.995 | -8.583 | -12.115 | 2.662 | -4.636 | -5.733 | 16.473 | -12.660 | 3.843 | 21.439 | 0.394 | -3.463 |
| -8.096 | 20.426 | -17.626 | -9.286 | 10.988 | -7.100 | 26.588 | 3.462 | 20.651 | 25.273 | 8.012 | 15.300 |
| -24.210 | -9.365 | -4.510 | 26.394 | -5.731 | 19.267 | -16.588 | 15.684 | 12.302 | 23.577 | -19.633 | 2.980 |
| 22.292 | -10.347 | -6.950 | 15.583 | 3.017 | 17.426 | 22.898 | -14.065 | 2.300 | 25.456 | 6.546 | -5.681 |
| -5.315 | 26.058 | -22.520 | 17.201 | 22.740 | -0.411 | 7.688 | 16.033 | 6.901 | 25.925 | 11.468 | 25.200 |
| 17.980 | 9.281 | -3.159 | -7.125 | 26.076 | 14.625 | -1.398 | 21.021 | -10.583 | 23.375 | 24.501 | -10.361 |
| -24.599 | 0.996 | -14.603 | 9.449 | 6.875 | -8.396 | 30.179 | 14.777 | -5.051 | 22.240 | 28.520 | 16.353 |
| -15.073 | -6.324 | 23.657 | 7.851 | 21.551 | 12.429 | 0.880 | 21.880 | 27.818 | -9.484 | 21.037 | 22.112 |
| -8.363 | -11.788 | -0.004 | -0.431 | 18.888 | -11.431 | -12.838 | 24.980 | -14.206 | -6.635 | 27.054 | -1.887 |
| -3.748 | -16.715 | 7.800 | -17.206 | 4.074 | -19.102 | 5.399 | 0.856 | -4.452 | -3.412 | 22.540 | 7.386 |
| 27.261 | 0.585 | 4.534 | -29.307 | 21.936 | -3.589 | 25.045 | -1.020 | -0.615 | -5.258 | 20.184 | 21.572 |
| -11.122 | -2.325 | -20.112 | 28.258 | -13.408 | 15.366 | 12.562 | 4.574 | 27.038 | 13.069 | 21.781 | 26.810 |
| -27.187 | -2.928 | 11.883 | 25.340 | 9.204 | 7.960 | 8.950 | 19.810 | 3.325 | 25.375 | 23.521 | 19.329 |
| -15.850 | -14.424 | -8.578 | 9.719 | -0.212 | -5.258 | 19.416 | -18.481 | 25.438 | 21.385 | 5.798 | 15.799 |
| -17.338 | 5.367 | 6.073 | 6.223 | -28.516 | 11.954 | 28.089 | -9.400 | -2.466 | 24.663 | -9.905 | 4.788 |
| -18.596 | 6.801 | -16.502 | 8.249 | 2.147 | -11.253 | 8.004 | 1.766 | -25.559 | 13.532 | 18.900 | 20.449 |
| 26.272 | 24.899 | 20.409 | -5.775 | 11.063 | 10.901 | -22.715 | 15.845 | 8.144 | 11.539 | 24.222 | 18.058 |
| -16.163 | 21.923 | 4.741 | 24.867 | -14.446 | -22.005 | 17.453 | -9.171 | -7.487 | -10.204 | 4.646 | -17.618 |
| -3.802 | 10.688 | -18.835 | 10.458 | 3.036 | 14.539 | -5.554 | -15.382 | 22.714 | 22.508 | 10.472 | -1.916 |
| -27.366 | -12.627 | -2.542 | 22.876 | 22.295 | 9.817 | 14.531 | 13.256 | 7.295 | 17.145 | 24.993 | 18.381 |
| -10.807 | 16.792 | -21.927 | 13.517 | 4.197 | 19.448 | 20.611 | -10.383 | -23.367 | -11.304 | 6.367 | 26.231 |
| -26.157 | 7.745 | -15.184 | 18.845 | 18.652 | 13.416 | 25.145 | 14.406 | -5.431 | 25.761 | -23.106 | 14.324 |

| | | | | | | | | | | | |
|---|---|---|---|---|---|---|---|---|---|---|---|
| 24.660 | 3.487 | 10.574 | -7.147 | 11.939 | 0.621 | 7.625 | 23.236 | 1.002 | -4.885 | 20.524 | 13.055 |
| 19.779 | 10.209 | -12.061 | 6.283 | 15.080 | -7.965 | 14.844 | 12.359 | -20.554 | 24.095 | 8.343 | 7.098 |
| -12.416 | 19.631 | -4.530 | -0.886 | 20.465 | 6.390 | 17.363 | 11.094 | 25.739 | 21.620 | 27.864 | 23.748 |
| 21.670 | 23.655 | 10.664 | 13.107 | 24.203 | 10.102 | 24.291 | 22.935 | 26.148 | 10.575 | 25.925 | 19.416 |
| -10.471 | -7.014 | 28.757 | 4.342 | -21.195 | 22.140 | 2.448 | -14.989 | -10.570 | 16.267 | -3.668 | -6.577 |
| 6.884 | 6.496 | 12.411 | 14.819 | -10.216 | 3.733 | 27.876 | 9.598 | 17.439 | -2.996 | -21.508 | 22.222 |
| -24.760 | -7.642 | 3.825 | -9.096 | -1.053 | 23.558 | 16.079 | -17.135 | 22.913 | -18.751 | 26.618 | 4.074 |
| -7.191 | -17.533 | 17.509 | 1.998 | 19.640 | -2.347 | -5.258 | -0.348 | 21.441 | 26.805 | 1.650 | 23.585 |
| 21.601 | 15.926 | 14.006 | -0.940 | 21.583 | -12.504 | 8.745 | -6.977 | 22.223 | 24.689 | 19.477 | 20.060 |
| -27.624 | 14.968 | -5.675 | 18.286 | 28.207 | 7.152 | 0.678 | 2.438 | -5.857 | 18.755 | 16.915 | 22.821 |
| 27.029 | 21.137 | -11.460 | 13.992 | 27.375 | -2.282 | 26.832 | 0.386 | 9.924 | 13.071 | -11.813 | 9.513 |
| -12.178 | 20.213 | 2.335 | 21.805 | 13.973 | 18.096 | 18.847 | 3.306 | 16.994 | -2.796 | 4.206 | -5.687 |
| 18.562 | 1.253 | 21.000 | 7.962 | -23.817 | -19.073 | -10.488 | -9.845 | 1.110 | 19.502 | 15.130 | -2.045 |
| -20.002 | -15.979 | -24.707 | -1.580 | -5.116 | 8.088 | 27.684 | 0.958 | 24.406 | -4.919 | -1.451 | 21.112 |
| -15.106 | 8.307 | -1.737 | 10.335 | 5.358 | -6.435 | 17.567 | 9.144 | -4.250 | 15.352 | 14.792 | 18.684 |
| 29.000 | -7.367 | -19.682 | -9.698 | 13.077 | -8.119 | 2.823 | -1.431 | -10.316 | 11.109 | 27.277 | -25.453 |
| 25.173 | 28.523 | -21.652 | 8.526 | 0.770 | 9.694 | -18.233 | -9.894 | -17.026 | 9.954 | 8.092 | 0.318 |
| -8.567 | 27.844 | 7.235 | 17.113 | 22.001 | -4.567 | 18.148 | -19.062 | 9.134 | 26.128 | 25.646 | 6.612 |
| -14.407 | -14.356 | -18.212 | -8.297 | 28.512 | -8.961 | 6.961 | 14.130 | 11.827 | -11.249 | -10.144 | 3.635 |
| -29.113 | 10.887 | 26.610 | -12.635 | -9.050 | 21.072 | 8.548 | 22.599 | 1.337 | 17.186 | 12.470 | -20.862 |
| 14.308 | 2.234 | 19.738 | 20.835 | 10.103 | 14.594 | 24.340 | -19.712 | 23.477 | 0.905 | 11.437 | 9.610 |
| -18.622 | -6.644 | -12.792 | -5.764 | 16.630 | 20.260 | 14.983 | 7.655 | 22.519 | 23.884 | 14.066 | 11.720 |
| -21.910 | 17.738 | -6.586 | -9.574 | 3.950 | 13.617 | 5.900 | 17.036 | 2.081 | 11.690 | 20.778 | -15.819 |
| -20.363 | 11.408 | 21.276 | -3.778 | 27.167 | 23.207 | 10.778 | -0.600 | 24.227 | 15.027 | 0.688 | -21.161 |
| 1.760 | 6.147 | 13.220 | 16.466 | -11.804 | -2.569 | 26.438 | 24.241 | 21.149 | 0.135 | 20.185 | 18.403 |
| -23.550 | 9.853 | -7.607 | 0.538 | -0.797 | 11.936 | 22.435 | -9.619 | 16.359 | -4.218 | -0.555 | 19.838 |
| 7.711 | -26.245 | -18.044 | 16.955 | -5.029 | -12.138 | 21.839 | 13.451 | 25.735 | 6.026 | -27.158 | 14.414 |
| -8.302 | -18.139 | 10.073 | 9.961 | 12.281 | 7.025 | 6.941 | 6.102 | 14.903 | 22.051 | 7.246 | 12.766 |
| -12.781 | 15.791 | -15.495 | -17.346 | 13.772 | 10.473 | 11.552 | -23.405 | -1.137 | -4.472 | 22.614 | 8.558 |
| -23.175 | -4.382 | -16.325 | 19.603 | -0.176 | 7.786 | 23.021 | 23.840 | 24.778 | 23.052 | 9.744 | 2.586 |
| 4.468 | 19.982 | -2.288 | 3.994 | 30.660 | 21.485 | 19.958 | 17.184 | 9.793 | -11.588 | 8.386 | 25.351 |
| -12.255 | -11.035 | 18.778 | -8.716 | 27.612 | -2.001 | -6.065 | -16.599 | 8.164 | 16.328 | 24.831 | 21.110 |
| -29.505 | 29.644 | -0.849 | -15.834 | 15.769 | -12.547 | 14.689 | -4.942 | -20.366 | 25.447 | 18.078 | 13.870 |
| -4.856 | 17.208 | -9.988 | 20.674 | 6.990 | -8.467 | 17.440 | 27.515 | 15.751 | 2.188 | 12.617 | 7.997 |
| -12.917 | -17.313 | -13.124 | 15.933 | -1.572 | 5.106 | 28.000 | -9.123 | 4.399 | 2.613 | -24.539 | -2.192 |

| | | | | | | | | | | | |
|---|---|---|---|---|---|---|---|---|---|---|---|
| 27.026 | -19.141 | -19.378 | 13.805 | 1.793 | 15.799 | 13.225 | 26.188 | 8.838 | 7.675 | -21.569 | -2.864 |
| -21.396 | -9.235 | -5.161 | 21.948 | 7.605 | -4.534 | 10.545 | -10.486 | 18.193 | 24.333 | 17.161 | 6.601 |
| -27.524 | -11.463 | -21.041 | 2.573 | -0.558 | -7.672 | 20.080 | 23.008 | 25.396 | -3.560 | -5.527 | 17.279 |
| -9.132 | 14.269 | -25.238 | -18.858 | 28.184 | -20.859 | 24.565 | -6.584 | 16.007 | 18.404 | 2.713 | 14.669 |
| 27.880 | 6.171 | 6.337 | 18.786 | 27.308 | -5.509 | 16.023 | -13.849 | 26.572 | 26.879 | -2.774 | 20.494 |
| 22.375 | -14.218 | 27.269 | -13.301 | 3.180 | 19.200 | 13.367 | 16.080 | 14.519 | -2.722 | -10.199 | 21.447 |
| 17.125 | 24.961 | -13.393 | 9.694 | 24.549 | -13.202 | 6.941 | -7.809 | -15.556 | 15.403 | 17.653 | 5.005 |
| -15.843 | 3.866 | -21.741 | -19.232 | 0.513 | 12.122 | 23.539 | 27.322 | 24.003 | 16.283 | 15.567 | 9.441 |
| 30.720 | 17.989 | 5.389 | 20.750 | -2.887 | 11.232 | 21.228 | -19.921 | 10.451 | 21.432 | 8.537 | 2.502 |
| -29.349 | 22.225 | -19.992 | 20.030 | 3.676 | -0.546 | 12.025 | 26.510 | -13.406 | 6.957 | -11.263 | -3.506 |
| -5.002 | -23.549 | 26.235 | 1.324 | -7.910 | -10.743 | 14.624 | -7.234 | 25.205 | 4.770 | 0.542 | 18.245 |
| -13.456 | -14.819 | 10.861 | -11.009 | 14.124 | -11.363 | -7.170 | -19.588 | 13.186 | -26.885 | 21.611 | 13.074 |
| -3.700 | -23.982 | 13.328 | -1.666 | 20.917 | -5.257 | 25.150 | -6.331 | -2.417 | 11.682 | 3.537 | 19.535 |
| 16.536 | 18.743 | -19.606 | -11.254 | 2.649 | -3.122 | 10.994 | 28.073 | -16.055 | 10.453 | 4.377 | 4.082 |
| -14.517 | 12.548 | 8.620 | 16.846 | 8.822 | 22.669 | 26.513 | 15.711 | 25.068 | 9.234 | 22.043 | 23.033 |
| -23.570 | 10.216 | 17.551 | 22.781 | 28.528 | 14.222 | -4.546 | -7.776 | -3.546 | -2.235 | 15.867 | -5.217 |
| 15.917 | -17.070 | -11.887 | -13.716 | 30.011 | 12.463 | 26.699 | -8.615 | 23.606 | 23.780 | 9.740 | 10.391 |
| 3.930 | 13.925 | -13.599 | 3.392 | 21.718 | 8.662 | 10.128 | 12.752 | 26.281 | 22.826 | 23.191 | 6.447 |
| -21.731 | 23.554 | -6.255 | 20.431 | -2.056 | 19.847 | -1.674 | 9.642 | 1.415 | 19.446 | 6.360 | 2.508 |
| 1.601 | -8.717 | 9.794 | 13.249 | 23.749 | -18.499 | 23.390 | -10.542 | 23.187 | 25.062 | -5.675 | -1.677 |
| 29.281 | 8.610 | 22.217 | -20.032 | -21.454 | -4.540 | 18.436 | 29.406 | 27.706 | -7.933 | 12.677 | 21.643 |
| 2.422 | 19.098 | -26.867 | -10.240 | 21.863 | 3.789 | 1.264 | 27.842 | -5.346 | 18.754 | 3.859 | 20.817 |
| -27.921 | -16.295 | -18.367 | 21.398 | 25.591 | 9.088 | 9.549 | -0.186 | -17.159 | 12.599 | 8.022 | -2.554 |
| -13.267 | -22.800 | -14.810 | -12.056 | 26.257 | 4.716 | 25.091 | -9.593 | 7.305 | 17.762 | -2.040 | 23.298 |
| -7.490 | 20.076 | -31.417 | -21.702 | 16.381 | 12.885 | 21.945 | 26.242 | -7.333 | 16.490 | -24.435 | 5.753 |
| -26.215 | 16.539 | 20.370 | -1.956 | 5.669 | 11.887 | 6.322 | -13.074 | 12.965 | 11.333 | -6.822 | 24.399 |
| -22.275 | 12.786 | -5.093 | 4.559 | 6.893 | -19.196 | 17.733 | -10.754 | -23.413 | -5.066 | -12.585 | 22.178 |
| -22.900 | 17.576 | -18.845 | 11.419 | 17.700 | -6.530 | 13.735 | -6.617 | -1.990 | 12.815 | 1.654 | 19.313 |
| -14.009 | 10.749 | -1.682 | 12.495 | 11.021 | 14.165 | 23.567 | -20.172 | 24.662 | 6.646 | 13.788 | 15.572 |
| 27.026 | 20.931 | -21.131 | 23.021 | 27.712 | 5.540 | 22.107 | 1.682 | 22.187 | -1.183 | -1.548 | 12.408 |
| -10.635 | -8.487 | -17.270 | -1.101 | 25.559 | 23.546 | 24.835 | 0.852 | 17.612 | 8.507 | -19.392 | 23.652 |
| -27.211 | 16.713 | -13.080 | 8.793 | 21.709 | 6.583 | 1.471 | 27.371 | 26.793 | 24.402 | 12.339 | 17.162 |
| -12.549 | 14.601 | -5.449 | 7.924 | 15.242 | 28.189 | 24.567 | 26.746 | 13.675 | -3.995 | 21.850 | 26.262 |
| -19.340 | -13.649 | 5.593 | 5.630 | 2.405 | 8.199 | 23.135 | -23.015 | 4.677 | 11.758 | -0.999 | 20.817 |
| -27.158 | 22.949 | 25.015 | 7.042 | 16.591 | -24.337 | 14.883 | -8.668 | 0.719 | 22.528 | -1.354 | -3.383 |

| | | | | | | | | | | | |
|---|---|---|---|---|---|---|---|---|---|---|---|
| 2.676 | 8.025 | 21.098 | 1.833 | 14.658 | -4.119 | 10.342 | 23.082 | 2.284 | 3.040 | 2.422 | 16.636 |
| 23.282 | 17.175 | 28.215 | -12.447 | 24.856 | 10.651 | 20.989 | -15.777 | 20.383 | -5.695 | 11.663 | 25.224 |
| -22.111 | 13.195 | -6.439 | 16.725 | 17.350 | -5.839 | 19.210 | 15.183 | -6.404 | 19.915 | -2.987 | 11.540 |
| 3.891 | 16.112 | 6.679 | 20.891 | -23.951 | 11.723 | 12.959 | 6.551 | -8.704 | -2.706 | 22.505 | 12.781 |
| -18.981 | -9.561 | 13.052 | 22.052 | 24.932 | 18.085 | -1.429 | 2.953 | -2.540 | 8.665 | -5.110 | 16.181 |
| -0.469 | 29.824 | -2.894 | 5.233 | 28.176 | 15.757 | -2.764 | 4.619 | -28.910 | 26.438 | 0.259 | 2.529 |
| -26.515 | -9.494 | -14.021 | -4.417 | -16.798 | 24.089 | 1.525 | -10.858 | -26.667 | 29.563 | 20.224 | 21.198 |
| -17.724 | 22.214 | -24.889 | -22.080 | 4.879 | -1.578 | 21.902 | -3.808 | 15.665 | 24.821 | 12.172 | 27.016 |
| -1.624 | 13.057 | -20.067 | 7.914 | -29.522 | 9.843 | 4.009 | -9.931 | -18.003 | -16.290 | 11.491 | 9.807 |
| 9.409 | -18.459 | -17.378 | 4.705 | 26.042 | 21.795 | 15.000 | 25.795 | 26.733 | -21.970 | 15.971 | 22.483 |
| -11.821 | 19.065 | -15.153 | 26.213 | -6.191 | -4.672 | 11.156 | -17.124 | -0.023 | 10.166 | -27.231 | 13.578 |
| -28.548 | -27.784 | 22.797 | 20.006 | 22.694 | 2.716 | 24.564 | -13.091 | -22.295 | 12.392 | 23.579 | 18.547 |
| -23.689 | 25.896 | -3.800 | 6.407 | 21.118 | 14.974 | -13.364 | 1.135 | -4.681 | 9.387 | -2.558 | 25.029 |
| -21.716 | 14.280 | -16.003 | -0.859 | 20.566 | -5.895 | 6.277 | -3.733 | 22.022 | 2.758 | 4.288 | 22.830 |
| -27.734 | 27.947 | 24.152 | -7.174 | -25.523 | 19.415 | 22.784 | 29.371 | -0.396 | 15.533 | 25.714 | 25.529 |
| -9.184 | 15.350 | 9.128 | -13.063 | 24.430 | 21.694 | -18.753 | -8.449 | 25.780 | -7.468 | 27.417 | 1.096 |
| 16.966 | -27.947 | -14.487 | 11.263 | 27.026 | -9.724 | 18.247 | -4.425 | 12.521 | 27.366 | 27.905 | 21.733 |
| -8.446 | 10.725 | 14.817 | 16.146 | 23.057 | 1.503 | 2.830 | 24.341 | 15.800 | 23.535 | -5.379 | 23.942 |
| 13.034 | 23.923 | -22.044 | -14.668 | -20.736 | 6.657 | 19.143 | 27.553 | 20.144 | 0.390 | -24.575 | 23.543 |
| -6.262 | 19.063 | -13.047 | -21.390 | -1.004 | -13.978 | -7.243 | 26.027 | 18.259 | 20.184 | -7.098 | 4.006 |
| 4.752 | 25.457 | 26.299 | 11.879 | 26.091 | 19.624 | -9.031 | 2.077 | -24.938 | 12.647 | 21.367 | -6.086 |
| 13.585 | 22.594 | -23.103 | 25.108 | 27.207 | 19.985 | -5.949 | -6.957 | -3.369 | 22.264 | 24.315 | -7.029 |
| -22.953 | -16.796 | 14.216 | 20.882 | 25.885 | 6.473 | -14.640 | -11.002 | -23.010 | 10.409 | 25.377 | 9.655 |
| -23.884 | 18.511 | -8.214 | -5.060 | 18.893 | 7.727 | -4.774 | 26.830 | 26.481 | 17.013 | 11.177 | 16.580 |
| -19.273 | 7.620 | -9.528 | 5.878 | 26.088 | 27.069 | 2.641 | 17.221 | -14.002 | 25.384 | 27.307 | 17.913 |
| 2.581 | 19.448 | -10.105 | 21.536 | -16.024 | 15.948 | 11.834 | -15.037 | 23.032 | -7.904 | 28.191 | 1.377 |
| -9.851 | 20.120 | -20.424 | -6.752 | -19.919 | 9.940 | -6.088 | 9.077 | -0.399 | 21.060 | 22.185 | 20.699 |
| -28.505 | 10.126 | -31.226 | -4.503 | -20.211 | 6.663 | 7.544 | -7.604 | 11.214 | 19.976 | 17.053 | 10.688 |
| 25.215 | 23.286 | -1.706 | 17.624 | 16.820 | -9.017 | -7.465 | 29.419 | 26.862 | 29.276 | 26.008 | 6.684 |
| 27.695 | 11.277 | -6.815 | 19.305 | -13.617 | 23.813 | -6.745 | 4.140 | 23.922 | -13.454 | 5.021 | 19.477 |
| -25.819 | 17.838 | 1.077 | -3.196 | -17.892 | 21.043 | 6.167 | -4.445 | -5.180 | 6.236 | 26.379 | 8.909 |
| -12.842 | -5.644 | 7.529 | -7.535 | -25.585 | 0.892 | -9.718 | 12.801 | 2.766 | 27.453 | 13.524 | -25.931 |
| -3.733 | 10.453 | 11.842 | 26.565 | -21.377 | 21.405 | 8.726 | 25.350 | -29.538 | 14.698 | 8.524 | 23.127 |
| 3.525 | 21.264 | 16.649 | 21.271 | 11.417 | -14.058 | -9.041 | 10.268 | 29.837 | 19.068 | -3.708 | 18.531 |
| 6.008 | 6.296 | -6.358 | 18.466 | 19.641 | 26.455 | 17.131 | 14.522 | -18.898 | 0.801 | 0.019 | -0.471 |

| | | | | | | | | | | | |
|---|---|---|---|---|---|---|---|---|---|---|---|
| 22.140 | 20.504 | 17.652 | -2.358 | -25.741 | -9.083 | 15.274 | -5.052 | 15.512 | 22.727 | -5.276 | 22.211 |
| -15.727 | 17.453 | 5.088 | -17.801 | 0.951 | -10.635 | 24.252 | -9.242 | -15.412 | -6.016 | 24.771 | 26.827 |
| -6.070 | 22.954 | 20.869 | 3.801 | -10.719 | 1.998 | 16.126 | -16.801 | 22.088 | -9.279 | 27.623 | 18.555 |
| 21.083 | 12.217 | 27.934 | 8.903 | 19.745 | 6.870 | 6.705 | 19.091 | 13.982 | -2.890 | 29.160 | -5.091 |
| -9.939 | 5.499 | -17.656 | -16.182 | 20.332 | 14.980 | 3.105 | 30.077 | 28.396 | 21.871 | 19.086 | 8.651 |
| -22.082 | 23.765 | -30.841 | -11.481 | 19.835 | 19.646 | 13.696 | -0.504 | 15.551 | 9.026 | 27.474 | 22.785 |
| 23.902 | -11.244 | -9.164 | -5.926 | 8.107 | 22.908 | 17.318 | -6.343 | -8.624 | 2.995 | 15.553 | 18.482 |
| -24.351 | -4.846 | 17.853 | 23.690 | 11.290 | 7.710 | -4.866 | -18.854 | 20.064 | 20.761 | 10.371 | 23.369 |
| 15.188 | 22.498 | -13.545 | 13.981 | -7.135 | 20.957 | 23.675 | -8.538 | 12.254 | -0.633 | 24.886 | 21.379 |
| -13.850 | -13.889 | -10.186 | 2.019 | 11.012 | -5.432 | 12.199 | 6.516 | 30.119 | 29.724 | 21.903 | -0.793 |
| 5.272 | -16.165 | -10.900 | -12.045 | -1.296 | 1.673 | 22.940 | -26.440 | -1.042 | 19.075 | 7.894 | 12.562 |
| 20.560 | 21.516 | 0.857 | 30.006 | 24.371 | 0.448 | 23.380 | -5.134 | 5.727 | 10.894 | 25.964 | 2.234 |
| -19.222 | 16.037 | -3.628 | -8.617 | 7.340 | 8.519 | 14.426 | -15.586 | 17.711 | 30.471 | 12.958 | 9.817 |
| -27.148 | 16.633 | -23.012 | -16.164 | 11.059 | -4.497 | -2.409 | -8.604 | 16.370 | 16.987 | -2.597 | 7.868 |
| -20.899 | 2.709 | -17.033 | 27.425 | 1.690 | 7.014 | 12.226 | 10.222 | 27.867 | 20.027 | 16.537 | 30.153 |
| 25.795 | 15.887 | -20.495 | 20.328 | 17.816 | -20.300 | 10.387 | 26.176 | 25.714 | 16.360 | -19.676 | 16.048 |
| -1.367 | 17.731 | 14.803 | 16.832 | -12.095 | -3.218 | -6.812 | 4.560 | -4.573 | -12.940 | 0.875 | 16.993 |
| -11.508 | 13.031 | -29.857 | 21.086 | -9.019 | 18.701 | -13.318 | 16.584 | 29.210 | 22.430 | 29.565 | 5.072 |
| 8.422 | 18.967 | 7.556 | -5.556 | -5.946 | 22.815 | 19.707 | 13.243 | 12.031 | 19.385 | 28.671 | -5.485 |
| 19.678 | 15.619 | 21.487 | -11.112 | -0.758 | 6.026 | -15.844 | 17.889 | -18.558 | 23.423 | 17.616 | 21.623 |
| -18.473 | 7.809 | 1.061 | 9.739 | 6.380 | 25.022 | 2.086 | -15.330 | 5.107 | -3.302 | 23.432 | 29.479 |
| 23.564 | -17.310 | 26.411 | 13.486 | -12.622 | -13.691 | 25.608 | 27.852 | 19.488 | 0.554 | 13.065 | 22.489 |
| 20.497 | -16.538 | -0.411 | 29.528 | 0.542 | 22.687 | -22.298 | -8.174 | 23.540 | 21.110 | -8.148 | 8.441 |
| -5.923 | 21.140 | -9.429 | 12.532 | 18.259 | -3.179 | 19.309 | -3.820 | -28.346 | 24.376 | 24.376 | -6.486 |
| -19.608 | 18.892 | -8.478 | -12.023 | 9.497 | -12.137 | 27.059 | -19.870 | 27.369 | 21.752 | 25.666 | 13.226 |
| -22.333 | -15.964 | 6.458 | 14.179 | 0.958 | 8.820 | 20.057 | -12.728 | 15.237 | 5.251 | -6.232 | 24.000 |
| -26.074 | 26.142 | -10.911 | -13.853 | -10.845 | 4.066 | 6.019 | 24.715 | 26.230 | 24.532 | 18.316 | 5.541 |
| 24.849 | -23.618 | 27.022 | -4.911 | 22.980 | 12.785 | 25.674 | 23.876 | 10.909 | 6.542 | 28.166 | 25.422 |
| 2.888 | 0.552 | -19.172 | 16.359 | 5.176 | 17.286 | 16.254 | 4.108 | 25.429 | 13.463 | 29.210 | 19.452 |
| -16.068 | 4.824 | -26.457 | -24.813 | -4.722 | 8.024 | 26.734 | 25.843 | 3.030 | 15.824 | 11.240 | 12.495 |
| 13.799 | 8.114 | -14.677 | -7.641 | -3.648 | 2.444 | 9.674 | 13.754 | -4.667 | 17.104 | 26.630 | 28.868 |
| 18.110 | 10.200 | -15.755 | 1.874 | 22.369 | 13.516 | 10.381 | -14.753 | 24.385 | 7.093 | -5.892 | 7.502 |
| -8.159 | 11.414 | -9.327 | -12.824 | 16.168 | -7.658 | -14.826 | -15.738 | 10.840 | 18.780 | 18.600 | 24.951 |
| -21.862 | -0.837 | 11.663 | -31.038 | -20.759 | -2.337 | 1.778 | 22.599 | -24.662 | 23.443 | -3.792 | 26.152 |
| 10.627 | 1.307 | -12.627 | -15.409 | 10.203 | 0.722 | 20.887 | -18.236 | -1.248 | 29.231 | 21.646 | 22.223 |

| | | | | | | | | | | | |
|---|---|---|---|---|---|---|---|---|---|---|---|
| 18.023 | 2.814 | -18.563 | 12.431 | -1.481 | -17.083 | 10.815 | -4.456 | -24.095 | 16.101 | 24.661 | 16.924 |
| 19.144 | 14.263 | 28.662 | 22.689 | 3.175 | 21.147 | 14.167 | -7.295 | 28.481 | 19.960 | 2.313 | -3.824 |
| -20.886 | -1.510 | 22.226 | -2.716 | -10.881 | 3.254 | 13.457 | -16.723 | -15.274 | 23.939 | 25.454 | 22.594 |
| 5.772 | -22.868 | -25.935 | -7.720 | 17.968 | 14.835 | -19.165 | 16.364 | 29.552 | 22.041 | 19.121 | 23.212 |
| -25.127 | -6.605 | -5.456 | -17.841 | -26.359 | -1.139 | 12.613 | 28.111 | 25.602 | 22.997 | 11.730 | 30.588 |
| 17.391 | -2.435 | 13.324 | -6.697 | -11.669 | 6.653 | 6.445 | -26.588 | 12.762 | 14.850 | 23.089 | 20.055 |
| -0.425 | -10.606 | 0.224 | 5.106 | -2.291 | -14.223 | 4.846 | -9.515 | -0.380 | 25.985 | -4.613 | 7.144 |
| -16.288 | 19.802 | -7.612 | -21.925 | 12.585 | 19.822 | 25.126 | 23.409 | 1.772 | 5.861 | -2.874 | 16.085 |
| 0.284 | 13.116 | -1.619 | 20.499 | 6.761 | 7.535 | 15.269 | -2.586 | 25.869 | -0.413 | 26.886 | 25.942 |
| -15.409 | 5.514 | 8.192 | -20.977 | 2.221 | -18.136 | 2.920 | 12.973 | 23.232 | 12.146 | -17.723 | 27.306 |
| -22.759 | 27.485 | -10.302 | 19.164 | 10.934 | 10.647 | 8.305 | 20.604 | 2.127 | -2.773 | 20.915 | 26.561 |
| 8.012 | -3.152 | -2.302 | 17.336 | -17.934 | 25.082 | -14.270 | 8.828 | 0.392 | -24.759 | 23.338 | 8.621 |
| 24.009 | 13.693 | -4.750 | 18.314 | -5.743 | 23.692 | 26.417 | 4.503 | -4.104 | 24.743 | 24.406 | 9.038 |
| 16.334 | 2.521 | -12.939 | 5.554 | 3.007 | 17.292 | -10.350 | 5.819 | -6.178 | 9.128 | 27.444 | 1.835 |
| -6.157 | 20.827 | 27.473 | 19.796 | 15.219 | -0.190 | -25.064 | -13.956 | 22.472 | 21.227 | 21.787 | 23.936 |
| -10.385 | -11.442 | -11.988 | 14.608 | -20.331 | -4.095 | 24.184 | -13.139 | 12.002 | 22.732 | -12.394 | 14.447 |
| -7.122 | -19.565 | -20.727 | -4.589 | 7.576 | 12.388 | -7.645 | -6.091 | -25.959 | -5.778 | 16.688 | 6.770 |
| 1.988 | 24.613 | -23.732 | 8.721 | 1.168 | -10.152 | -0.003 | 25.524 | 28.338 | 12.249 | -4.448 | 3.587 |
| -1.732 | 12.352 | -24.920 | 27.724 | 11.855 | 8.371 | 11.295 | 6.316 | 4.787 | 25.660 | 18.989 | -15.179 |
| 13.391 | -3.602 | -1.212 | -3.119 | 4.615 | 5.591 | -16.744 | 26.066 | 27.380 | 22.043 | 16.163 | 25.571 |
| 6.584 | -0.077 | -11.012 | -17.482 | -24.466 | 21.208 | 24.784 | 8.266 | -14.677 | 20.288 | 17.243 | 19.261 |
| -18.086 | 3.875 | -21.284 | 12.316 | 20.741 | -8.398 | 23.303 | -3.249 | 25.670 | 12.954 | -7.048 | 16.074 |
| -8.155 | -11.968 | 5.932 | -27.889 | 15.042 | 11.894 | 3.605 | -10.777 | 7.686 | 3.849 | 8.525 | -0.688 |
| 24.629 | -1.557 | -13.818 | 19.960 | 13.813 | -14.157 | 22.387 | -6.500 | -5.596 | -22.102 | -11.616 | 15.474 |
| -17.489 | -23.520 | 27.251 | -9.314 | 8.421 | 19.718 | 27.381 | 20.887 | -19.982 | 21.504 | -1.833 | 14.379 |
| 5.269 | -5.485 | -5.378 | 26.896 | 4.443 | 15.805 | 3.948 | 28.442 | 5.526 | -5.297 | 22.145 | 11.317 |
| -18.814 | 21.252 | -11.269 | 20.054 | 21.541 | -6.662 | 4.582 | 13.609 | 0.285 | 8.809 | 3.150 | 27.592 |
| 25.200 | 14.866 | -14.997 | 15.650 | 10.397 | 7.705 | 14.612 | -4.814 | 26.687 | 24.749 | 15.228 | 28.446 |
| -22.791 | -4.718 | -17.305 | 28.688 | 17.072 | -2.343 | 25.379 | 21.197 | -1.304 | 23.121 | -4.892 | 26.370 |
| -17.152 | -1.897 | 0.464 | 16.948 | 20.115 | 14.173 | -14.151 | 10.559 | 20.806 | 19.584 | 16.294 | 24.848 |
| 9.924 | -8.624 | 5.847 | 8.722 | -21.838 | 16.267 | 26.263 | 6.034 | -23.205 | 24.189 | -15.305 | 12.763 |
| -7.821 | -3.928 | -2.441 | -19.563 | 6.043 | 6.678 | -22.356 | -8.071 | 19.686 | 4.057 | 21.217 | 18.189 |
| 10.758 | 17.305 | 2.569 | -21.168 | 1.589 | 12.037 | 18.646 | -23.090 | 23.596 | 3.364 | 10.836 | 21.195 |
| 26.422 | 23.127 | -25.287 | 9.718 | 20.974 | 20.560 | 8.821 | -27.312 | -13.798 | 19.110 | 2.351 | 5.115 |
| -1.170 | 8.831 | -19.918 | 11.343 | -1.432 | -14.762 | 28.400 | 2.660 | 14.580 | 14.041 | 7.116 | 5.980 |

| | | | | | | | | | | | |
|---|---|---|---|---|---|---|---|---|---|---|---|
| 25.764 | 6.365 | 2.602 | -5.315 | -23.404 | 14.659 | 21.751 | 27.850 | 10.904 | 7.442 | -26.391 | 13.390 |
| -10.142 | 24.499 | -12.209 | 5.997 | -8.291 | 5.525 | 6.192 | 27.205 | -2.054 | 22.658 | 25.445 | 1.880 |
| 9.524 | 27.065 | -13.079 | 22.880 | -13.369 | 9.005 | 27.465 | 9.091 | 4.316 | 25.275 | 15.728 | 16.995 |
| 12.845 | -19.811 | -5.607 | 17.511 | -14.030 | -1.500 | 24.059 | 4.134 | 3.769 | 14.405 | 14.674 | 21.626 |
| -10.915 | 14.821 | -25.137 | 13.419 | 22.870 | -5.793 | 19.900 | -16.960 | -5.922 | 13.154 | 3.366 | 6.659 |
| -20.632 | 10.934 | 4.907 | -2.910 | -17.157 | 10.043 | 15.880 | 6.603 | -2.927 | 17.367 | -13.363 | 27.855 |
| -1.487 | 25.352 | -4.122 | 25.096 | 6.245 | 24.070 | 3.482 | -15.537 | -5.191 | 20.059 | -19.053 | 26.159 |
| 27.372 | 27.685 | -1.835 | -9.593 | 15.875 | -11.618 | 22.796 | 24.042 | 23.775 | 24.745 | 11.346 | 7.936 |
| 13.916 | -0.179 | -15.620 | 10.653 | -17.713 | 6.092 | 5.067 | -25.134 | 25.713 | -4.269 | -7.884 | -0.495 |
| -15.070 | -21.857 | -25.287 | -22.304 | 8.748 | -8.763 | -5.372 | -24.518 | 13.896 | 7.543 | 6.168 | 25.173 |
| -11.699 | -11.790 | -1.078 | -13.842 | -3.548 | -3.370 | 10.764 | 19.060 | 16.131 | 16.554 | 4.828 | 2.937 |
| -5.429 | 11.468 | -7.379 | -25.549 | 28.561 | -6.440 | 9.574 | 1.126 | 26.787 | 18.459 | 0.256 | 12.712 |
| 7.577 | 5.496 | -10.173 | -8.619 | 6.976 | 3.582 | 18.303 | 9.382 | 20.405 | 13.968 | 24.330 | 22.239 |
| 23.214 | -19.874 | -6.093 | 6.723 | -6.858 | -19.397 | -2.182 | -1.667 | -5.798 | -21.368 | -19.712 | -2.505 |
| -0.148 | -21.566 | 14.538 | 8.822 | 4.036 | 10.765 | 25.141 | -13.135 | 23.343 | -10.076 | 10.816 | 26.632 |
| 22.017 | -0.401 | -4.962 | 12.655 | 13.365 | 22.016 | 23.671 | 26.150 | -17.437 | 25.188 | -6.429 | 20.148 |
| 19.369 | -24.343 | 5.265 | 3.661 | -4.976 | 19.386 | 4.468 | 0.160 | 23.658 | -26.758 | -26.014 | 5.501 |
| -24.077 | 5.933 | -12.187 | -7.547 | 19.561 | 16.340 | 26.206 | -8.123 | 3.545 | -4.479 | -22.929 | 10.452 |
| 24.537 | -11.045 | -0.621 | -0.275 | 22.977 | -2.321 | 5.756 | 2.309 | -4.402 | 22.597 | -17.927 | -6.436 |
| -19.927 | 4.674 | 7.833 | 15.671 | 24.238 | -13.453 | 25.874 | -9.470 | -12.561 | 19.935 | 7.083 | 14.871 |
| 25.414 | 12.200 | 16.357 | -4.161 | 9.058 | 4.634 | 15.136 | 21.472 | 1.747 | -22.342 | -4.610 | 23.751 |
| 19.335 | 6.788 | 20.839 | -5.821 | -5.641 | 7.914 | 12.315 | 10.507 | 3.720 | 23.347 | -19.853 | 23.423 |
| -18.238 | -20.848 | -2.343 | 3.459 | 5.053 | 21.087 | 20.989 | 13.084 | -13.421 | 24.861 | 26.511 | 26.303 |
| 26.898 | 28.241 | 4.202 | -15.792 | 13.323 | 13.847 | 27.542 | 24.082 | 20.655 | 20.452 | 17.318 | 21.571 |
| -25.533 | -1.348 | 10.536 | -3.129 | -6.715 | -1.240 | 10.908 | -8.186 | 25.549 | -3.778 | -17.105 | 4.388 |
| -3.407 | -9.277 | 19.799 | -10.188 | 6.752 | -6.769 | -3.468 | 8.433 | 22.366 | 19.747 | 4.013 | 6.319 |
| 0.285 | -22.449 | -13.626 | 24.425 | 5.729 | 25.890 | 20.012 | 20.839 | 21.486 | 8.085 | 16.328 | 27.049 |
| 19.843 | 9.323 | 7.417 | 23.400 | -29.468 | -5.235 | -8.491 | 5.911 | 17.978 | -19.727 | -8.057 | -0.787 |
| -1.578 | -11.445 | 0.040 | 21.867 | 7.545 | 27.122 | 6.054 | -26.981 | -10.327 | 21.765 | -18.175 | -7.151 |
| -21.684 | -18.735 | 17.211 | 4.491 | -28.979 | 3.131 | 21.859 | 11.278 | -6.600 | 5.327 | 7.204 | 19.488 |
| 16.943 | -7.193 | -19.529 | 23.070 | -8.715 | -20.359 | -11.283 | -4.834 | -10.846 | 20.635 | -23.556 | 12.309 |
| -23.554 | 20.173 | 22.956 | 5.346 | 10.359 | 24.773 | 7.923 | -10.433 | 24.714 | 11.904 | 19.089 | 5.695 |
| -20.141 | 1.040 | -11.539 | 24.979 | 6.646 | -2.438 | 14.064 | 25.228 | 13.563 | 4.690 | 4.479 | 17.528 |
| 26.760 | 15.623 | -20.104 | -24.658 | -14.206 | 16.236 | 22.529 | 21.530 | -1.897 | 13.291 | -22.251 | 20.164 |
| -14.833 | 22.199 | 5.577 | 0.966 | 17.040 | -7.985 | 19.441 | -7.979 | 2.363 | -8.925 | 6.053 | 18.851 |

| | | | | | | | | | | | |
|---|---|---|---|---|---|---|---|---|---|---|---|
| -0.043 | 10.609 | 25.123 | 4.905 | 4.115 | -14.383 | 12.341 | -11.541 | -24.982 | -2.381 | 21.416 | 20.809 |
| 23.212 | -21.828 | 11.766 | -15.875 | 14.522 | 2.143 | 24.184 | 1.907 | 0.493 | 25.597 | -2.870 | -2.826 |
| -23.961 | 23.765 | -4.168 | 17.258 | 4.923 | -9.015 | -23.482 | -4.739 | 6.589 | 23.866 | -15.187 | 22.277 |
| -7.630 | -20.208 | -6.474 | 5.438 | 2.311 | 9.134 | 23.377 | 3.868 | -18.425 | -5.387 | -9.365 | 11.831 |
| 2.423 | 28.109 | -5.863 | -22.482 | -30.299 | 23.577 | 21.956 | 26.068 | 19.689 | 8.872 | -7.635 | 17.991 |
| -16.384 | 14.369 | -4.943 | 0.633 | 29.702 | -10.226 | 24.833 | 20.455 | -9.617 | 18.815 | -23.500 | 19.373 |
| 28.279 | 8.557 | 20.461 | -14.669 | 9.744 | -25.649 | 6.779 | 17.021 | 9.741 | 21.262 | -12.981 | 25.216 |
| 27.491 | -10.138 | -21.119 | -8.686 | 30.111 | -4.469 | 11.128 | -13.370 | -2.403 | -11.365 | 20.780 | 6.064 |
| -21.187 | 0.002 | -25.607 | 12.239 | 28.951 | -4.995 | 23.169 | -23.441 | -3.611 | 20.647 | 20.538 | 20.105 |
| -24.589 | -23.741 | 22.695 | -24.604 | 25.273 | 0.036 | 19.302 | -8.314 | -10.914 | -6.413 | 16.875 | 25.108 |
| 6.667 | 21.381 | 2.425 | 13.812 | 16.079 | 15.617 | -8.452 | -17.174 | 13.161 | 15.131 | 13.012 | 12.871 |
| -0.710 | -9.024 | -4.482 | -3.456 | 11.492 | -21.508 | 12.829 | -9.032 | 14.355 | 13.756 | 21.982 | -4.238 |
| 29.236 | 22.188 | -13.116 | 1.631 | -4.322 | 14.866 | -2.042 | 9.670 | 15.963 | -27.703 | 21.254 | 11.595 |
| 26.521 | 23.797 | 26.051 | 19.470 | 27.750 | 6.838 | 19.333 | -7.139 | -2.782 | 3.441 | 24.920 | 18.189 |
| -1.668 | 18.706 | 10.527 | 18.357 | 13.840 | -8.108 | -16.422 | -13.601 | 1.619 | 12.292 | 11.434 | 25.480 |
| -15.493 | 7.384 | -27.156 | 25.157 | 13.329 | 21.310 | 17.311 | -21.621 | -18.206 | 19.680 | 18.250 | 16.392 |
| -24.031 | -19.199 | 17.585 | 5.218 | -2.776 | 24.859 | 27.877 | 25.101 | 25.466 | 16.401 | 12.025 | 9.680 |
| -6.580 | 26.441 | 14.952 | 23.326 | 10.599 | -19.019 | 20.288 | -1.099 | 5.626 | 16.973 | -2.096 | 21.096 |
| 8.709 | 14.753 | -6.364 | -24.037 | 17.070 | 14.079 | 18.076 | -13.987 | 19.881 | 11.472 | 25.574 | 6.585 |
| 30.350 | -10.978 | 25.045 | -26.342 | 12.024 | -13.109 | 5.311 | -10.082 | 9.996 | 24.847 | 24.687 | 23.742 |
| 24.433 | 25.621 | -7.160 | 18.254 | 22.346 | -11.345 | 14.910 | -8.425 | 8.670 | 10.970 | 21.488 | 22.398 |
| 22.324 | 19.709 | -9.321 | 13.891 | 19.403 | 0.986 | 11.265 | 23.085 | 21.211 | 5.446 | 26.646 | 19.039 |
| -11.674 | -15.506 | 2.400 | 26.069 | 11.357 | 19.596 | 13.907 | 3.063 | 1.251 | 17.190 | 23.424 | 26.157 |
| -21.392 | 9.009 | -4.553 | 4.145 | 9.676 | 7.044 | -7.328 | -10.328 | 20.593 | 21.778 | 11.187 | 21.722 |
| -10.874 | -19.932 | 19.299 | 23.561 | 21.043 | 16.919 | 22.432 | -7.394 | 14.732 | 26.792 | 9.325 | 7.567 |
| 25.774 | -16.021 | -19.100 | 2.907 | 24.079 | 4.582 | 6.043 | -20.099 | 13.398 | 1.526 | 25.123 | 15.939 |
| -2.496 | 1.884 | 8.707 | 15.425 | 29.466 | 9.302 | 23.196 | 7.183 | -14.878 | 19.354 | 21.969 | 9.913 |
| -1.767 | -28.827 | -13.439 | 14.849 | 4.662 | -1.683 | 25.065 | 9.459 | 0.733 | 25.773 | 12.426 | 22.664 |
| 25.418 | -21.930 | -24.028 | 2.489 | 3.960 | 14.584 | 4.439 | -17.762 | 25.675 | 20.072 | 20.585 | 20.084 |
| -15.280 | -8.972 | -17.835 | -18.302 | 16.733 | 15.097 | 10.100 | -22.911 | 15.582 | 24.995 | 22.405 | 12.554 |
| -10.424 | 18.780 | 21.617 | 5.783 | -6.067 | 10.576 | 23.681 | -24.142 | -6.331 | 11.367 | -0.225 | -1.053 |
| 30.105 | -9.642 | -27.161 | 9.470 | 29.795 | 20.400 | 15.160 | -24.699 | 5.430 | 15.568 | 24.026 | -5.715 |
| 11.621 | -25.540 | -0.515 | 2.243 | 12.795 | 16.558 | 21.771 | -13.859 | -12.143 | 17.305 | 15.921 | 22.277 |
| -22.069 | 14.234 | 13.360 | 4.765 | -1.355 | -7.078 | 15.941 | 2.044 | 11.735 | 16.261 | 25.396 | 17.217 |
| -14.696 | -5.748 | -8.292 | 9.337 | 28.800 | -1.097 | 22.219 | -8.637 | -2.527 | 17.724 | 27.166 | 20.305 |

| | | | | | | | | | | | |
|---|---|---|---|---|---|---|---|---|---|---|---|
| 21.162 | 4.876 | 9.851 | 5.128 | 12.844 | -9.878 | 22.136 | -2.865 | 4.806 | 2.148 | 15.602 | 5.247 |
| 15.448 | -0.242 | 5.150 | 11.863 | -0.127 | 13.845 | 3.212 | -10.088 | 0.939 | 26.451 | 26.287 | 23.623 |
| -10.136 | 17.343 | 25.910 | -14.658 | 0.779 | -3.496 | 9.779 | -9.300 | -11.019 | 2.716 | 25.100 | 26.850 |
| 26.933 | -1.221 | 14.325 | 28.112 | 6.115 | 7.787 | 14.188 | 10.051 | 27.347 | 24.872 | 1.317 | -0.052 |
| -6.685 | -13.512 | -10.871 | 0.239 | -1.113 | -5.791 | 10.024 | 2.007 | 22.107 | 27.760 | 21.490 | -2.472 |
| -25.032 | -19.246 | 0.700 | -23.665 | 23.572 | -18.547 | 25.839 | 11.352 | 19.852 | 20.512 | 19.409 | 23.378 |
| -2.587 | 19.351 | -26.497 | -1.877 | 22.221 | 24.347 | 11.853 | 8.525 | -1.295 | 18.403 | 16.728 | 25.012 |
| 27.989 | 11.010 | 12.003 | -27.458 | -20.374 | -10.210 | 17.946 | 25.157 | -8.088 | 21.996 | 24.536 | 20.221 |
| -1.271 | 12.420 | -2.415 | 12.491 | 14.853 | 12.774 | 14.742 | -3.917 | 25.588 | 9.997 | 11.505 | 18.534 |
| -16.840 | 8.973 | 7.367 | 9.760 | 17.335 | 17.462 | 9.127 | 12.296 | -4.012 | -4.607 | 18.982 | 6.339 |
| 23.162 | 21.423 | 12.914 | 1.156 | 2.147 | -26.876 | 4.154 | 21.357 | 18.196 | 15.950 | -0.638 | 22.645 |
| 25.478 | -18.614 | 8.757 | 16.888 | 26.649 | 1.379 | 27.045 | 4.829 | -18.016 | 23.875 | 24.647 | 14.700 |
| 14.256 | -11.450 | -26.969 | -3.671 | 5.624 | 18.284 | 23.982 | -12.241 | -11.364 | 1.145 | 25.980 | -3.467 |
| -9.939 | -13.702 | 11.745 | 4.558 | -1.337 | 22.091 | 7.617 | 13.819 | 12.586 | -1.227 | 23.143 | 16.814 |
| -10.553 | -14.548 | -19.040 | 20.873 | -7.497 | 16.029 | 5.049 | 0.531 | -19.760 | -5.658 | 24.307 | 20.828 |
| -7.037 | -14.904 | -26.037 | 19.913 | 21.677 | -7.206 | -8.114 | 3.581 | 24.394 | 15.141 | 14.635 | 22.012 |
| 26.199 | 16.680 | 21.773 | 22.325 | 19.262 | -2.772 | 1.403 | 10.991 | 18.986 | 26.176 | 28.246 | -0.099 |
| 24.964 | -27.587 | -4.695 | -27.535 | 7.714 | 20.905 | 4.594 | -3.466 | 14.324 | 3.955 | 13.362 | 18.905 |
| 14.130 | 24.477 | -23.234 | 22.904 | 18.413 | 14.332 | 20.565 | 8.321 | 27.938 | -6.563 | 4.451 | 0.097 |
| -27.829 | 22.966 | -5.757 | 24.634 | 20.809 | 24.290 | 21.162 | -19.497 | 16.608 | 22.587 | 17.184 | 6.449 |
| -23.877 | 31.111 | -8.528 | 23.891 | 3.538 | 6.037 | 12.609 | -8.085 | -3.933 | 12.679 | -0.141 | -0.345 |
| -11.469 | 14.400 | 1.110 | 12.216 | 17.341 | -3.305 | 9.874 | 22.711 | -15.618 | 28.834 | 26.891 | 11.671 |
| -12.277 | 26.494 | 23.095 | 2.763 | -19.774 | 19.647 | 8.750 | 21.197 | 26.270 | 2.613 | 18.087 | 23.728 |
| -4.597 | 10.368 | 11.772 | 0.077 | 25.467 | -9.543 | 22.651 | 21.929 | 17.470 | -11.862 | 28.443 | 14.836 |
| -25.059 | 2.141 | 8.849 | 17.776 | 28.317 | 9.278 | 5.127 | -19.260 | -7.266 | 19.816 | 12.078 | -0.259 |
| -6.079 | 26.047 | 18.819 | -4.395 | 18.382 | 18.254 | 19.162 | -5.894 | 24.000 | 3.136 | 27.440 | 14.383 |
| 23.878 | -17.352 | -4.342 | 9.823 | 12.667 | 16.256 | 21.844 | -10.855 | -28.470 | 21.815 | 24.862 | 19.991 |
| 27.042 | 8.789 | -11.292 | -18.234 | 23.814 | 15.005 | 8.679 | 2.895 | -18.409 | 21.103 | 20.348 | 22.123 |
| 3.730 | -18.296 | -17.216 | 21.958 | -25.910 | 26.191 | 14.279 | 9.486 | 20.335 | 16.163 | 13.957 | 17.739 |
| 9.319 | -20.596 | 27.138 | 27.565 | 19.705 | -3.113 | 15.811 | -22.058 | 16.046 | 17.705 | 16.862 | 15.454 |
| 8.598 | 21.244 | -1.734 | 28.127 | 20.018 | -11.610 | 11.562 | -9.711 | -7.843 | 29.065 | 19.140 | 11.551 |
| -29.553 | 17.734 | 6.954 | -9.441 | 14.722 | 8.115 | -24.495 | 10.831 | 22.049 | 10.986 | 16.319 | 19.737 |
| -21.452 | 24.374 | -5.199 | -13.296 | 5.921 | 21.662 | 15.193 | -16.155 | -5.920 | 5.421 | 24.710 | 16.154 |
| 29.849 | 13.915 | -9.246 | 22.672 | -3.668 | -14.058 | 3.224 | 18.611 | 12.698 | 24.811 | 23.594 | -2.163 |
| -30.363 | -17.749 | 10.147 | -28.143 | -5.495 | 14.123 | 7.042 | 19.260 | -22.489 | 18.000 | 14.237 | 16.037 |

| | | | | | | | | | | | |
|---|---|---|---|---|---|---|---|---|---|---|---|
| -25.993 | 1.027 | 15.135 | 24.980 | -7.323 | -16.744 | 18.498 | 4.483 | 18.427 | 18.359 | -11.218 | 25.940 |
| 25.447 | 15.144 | 18.575 | -20.174 | 27.058 | 9.943 | 5.255 | 17.941 | -7.528 | 0.387 | 28.086 | 18.412 |
| 3.892 | 27.527 | -12.121 | 5.507 | -9.865 | -0.257 | -8.904 | 22.175 | 28.644 | 22.585 | 17.049 | 14.642 |
| 22.680 | 21.997 | -14.989 | 20.272 | -18.856 | 17.898 | -25.384 | -0.172 | 23.386 | 27.946 | 28.758 | 22.177 |
| -12.581 | 7.928 | 15.994 | 13.131 | -4.558 | 16.350 | 10.400 | 3.925 | -13.638 | 25.006 | 11.219 | 1.666 |
| -4.714 | -13.468 | -24.744 | 22.039 | -9.016 | -20.194 | 11.858 | -7.994 | 24.724 | 17.148 | 8.940 | 22.393 |
| 30.482 | 13.193 | 14.144 | 6.634 | -24.246 | 19.553 | 20.892 | 26.020 | 6.097 | 4.887 | 26.356 | 8.765 |
| -9.386 | -29.136 | -19.481 | 24.037 | 20.379 | 26.869 | 14.458 | 23.600 | 3.975 | -0.810 | 13.722 | 21.176 |
| -19.022 | -19.531 | -29.066 | -4.971 | 25.891 | 17.862 | -0.387 | 26.959 | -7.157 | 8.032 | 24.928 | 13.548 |
| 27.486 | 16.596 | 16.773 | 14.455 | 26.648 | 24.341 | 11.034 | 17.030 | 8.519 | 3.074 | 18.239 | 1.373 |
| 13.686 | 15.710 | 23.773 | -27.143 | 6.045 | -3.107 | 0.786 | 19.007 | -4.188 | 1.572 | 14.361 | 19.350 |
| -11.602 | -3.118 | -5.405 | -25.896 | -27.464 | 24.711 | 12.500 | 10.953 | 27.057 | 14.827 | 9.307 | 16.441 |
| 19.513 | -15.419 | 13.003 | 25.513 | -17.174 | -4.838 | 9.432 | -11.675 | 22.777 | 0.247 | -0.760 | 24.061 |
| 7.515 | -16.941 | 26.925 | 6.588 | 6.756 | -2.877 | 13.954 | -8.150 | 18.546 | 27.544 | 27.703 | 4.734 |
| -10.578 | -9.399 | 5.446 | 5.214 | -4.069 | 6.861 | -13.122 | 14.451 | -11.148 | 5.666 | 25.310 | 6.933 |
| 29.423 | 27.350 | -8.769 | 10.902 | 23.657 | -20.704 | -21.555 | -26.828 | 15.907 | 7.777 | 19.045 | 22.464 |
| 6.128 | -21.855 | 19.020 | 6.126 | -18.128 | 18.692 | 8.902 | 9.850 | 25.938 | 4.309 | 22.322 | -5.533 |
| -8.428 | 25.094 | 19.789 | -28.816 | 24.941 | 14.121 | 22.716 | 23.275 | -14.258 | 16.627 | 14.588 | 26.496 |
| 23.261 | 12.996 | 8.861 | 25.320 | 21.107 | 19.940 | 4.104 | -6.757 | 5.099 | 7.116 | 18.076 | 17.765 |
| 17.193 | -25.457 | -4.815 | 5.266 | 22.738 | 13.001 | -12.672 | -16.005 | 24.493 | 15.635 | 23.864 | 19.082 |
| -14.971 | 25.439 | -23.182 | 28.064 | 9.012 | -19.229 | 9.378 | 9.400 | 27.805 | 29.337 | 21.923 | 8.548 |
| 21.286 | 22.383 | -5.425 | 14.483 | -13.596 | 20.335 | -21.047 | -19.492 | 1.205 | 18.192 | 30.620 | 13.394 |
| 29.915 | 17.342 | -10.633 | 20.933 | -2.069 | 16.067 | -22.818 | 25.700 | -27.998 | -0.902 | 20.771 | 25.606 |
| 12.869 | 20.814 | -15.077 | -24.559 | -18.804 | 6.777 | 25.849 | 1.477 | -3.306 | -2.010 | 11.982 | 16.339 |
| -14.143 | -17.502 | 4.243 | 22.319 | -9.008 | -15.702 | -8.464 | 27.031 | 30.417 | 12.356 | 28.666 | 12.055 |
| 24.278 | -24.997 | -22.347 | -30.158 | 12.827 | 19.712 | 21.845 | -8.638 | 16.677 | 14.994 | 29.173 | 26.145 |
| 4.697 | 9.235 | 12.902 | 7.870 | -17.021 | 21.424 | 16.197 | -2.026 | -5.234 | -32.419 | 12.414 | 6.935 |
| -10.404 | 15.243 | 26.407 | 3.733 | 21.566 | 18.022 | -7.001 | 24.329 | 8.599 | 7.992 | 10.898 | -0.777 |
| 8.795 | -15.288 | 14.796 | 6.340 | 10.787 | 3.359 | 16.160 | 28.721 | 23.255 | 21.877 | 15.506 | 13.299 |
| 19.821 | 18.688 | -22.930 | -20.827 | -25.758 | 9.588 | -21.309 | -24.457 | -14.120 | 13.388 | 25.328 | 7.966 |
| 14.105 | 13.883 | -6.229 | 18.979 | 20.424 | -20.008 | 10.235 | 26.679 | 25.673 | 8.964 | 19.306 | 18.310 |
| 13.592 | -11.955 | 21.226 | 7.603 | -16.082 | 3.451 | -13.992 | -27.865 | 11.087 | 20.553 | 28.023 | 19.601 |
| 26.571 | 24.419 | -3.855 | 2.297 | 24.903 | 10.137 | -18.353 | 28.498 | 1.966 | 2.426 | 17.900 | 28.788 |
| 5.664 | 19.521 | -7.429 | 10.132 | 10.383 | -17.575 | 25.162 | 5.273 | 26.827 | 29.180 | 18.585 | 22.484 |
| 10.257 | 6.807 | 17.739 | 27.424 | -10.668 | -20.995 | -10.697 | 26.977 | 27.417 | 7.054 | 27.104 | 3.282 |

| | | | | | | | | | | | |
|---|---|---|---|---|---|---|---|---|---|---|---|
| -21.021 | 12.300 | 18.267 | -23.873 | 11.471 | -14.072 | 9.180 | 20.818 | 12.485 | 17.755 | 16.532 | 21.349 |
| 21.321 | -10.616 | -5.510 | 27.906 | 3.718 | -0.388 | -20.050 | 10.233 | -11.946 | 22.749 | 17.695 | 24.280 |
| 24.953 | -17.250 | 15.127 | 3.241 | -16.487 | 8.327 | 24.212 | 0.094 | 23.805 | 10.663 | 22.638 | 11.815 |
| 20.478 | 23.711 | -11.941 | 13.745 | 2.183 | -1.828 | 14.471 | -20.968 | 0.300 | 26.768 | 8.534 | 29.117 |
| -22.057 | -6.153 | -7.873 | 29.061 | -25.236 | -13.621 | -21.472 | 11.114 | 28.598 | 16.389 | 25.069 | 16.391 |
| 16.007 | -20.589 | 13.830 | -26.080 | -15.452 | -26.233 | 20.927 | 21.636 | 12.010 | 19.142 | 19.781 | 24.952 |
| 3.347 | 22.808 | -13.438 | 26.641 | 0.175 | -9.344 | 16.807 | 1.883 | -27.297 | 8.467 | 23.450 | 26.001 |
| 11.971 | -17.799 | 25.724 | 20.172 | 9.577 | -14.361 | 24.880 | -5.014 | 14.932 | 3.301 | -17.871 | -9.998 |
| -27.071 | 21.363 | -26.500 | 17.816 | -15.891 | -1.485 | -2.905 | 24.519 | 7.397 | 12.194 | 4.050 | 14.725 |
| 22.868 | -15.996 | -24.543 | 2.618 | 10.793 | 2.486 | 18.628 | -6.193 | 29.772 | 28.734 | 15.425 | 0.887 |
| 0.003 | -26.441 | -4.465 | 0.823 | -23.999 | -7.428 | -22.892 | 21.207 | 30.604 | 13.718 | 13.465 | -0.448 |
| -6.878 | 19.059 | 11.691 | 23.497 | -15.648 | 15.668 | -19.763 | 16.720 | 13.350 | 20.453 | 10.248 | 27.897 |
| 16.492 | 15.536 | 8.930 | 11.215 | -18.676 | -6.561 | 8.771 | -10.668 | 17.427 | -3.464 | 25.943 | 2.944 |
| 22.509 | -20.569 | 16.468 | -2.784 | -13.947 | -2.295 | 27.358 | 25.029 | 27.812 | -5.631 | 17.442 | 17.266 |
| 23.946 | -11.481 | -15.669 | -27.727 | -6.661 | -3.227 | 20.310 | 25.804 | 19.792 | 8.200 | 27.625 | 27.416 |
| 19.189 | 17.851 | 25.286 | 4.759 | 0.532 | -4.759 | 7.159 | 21.769 | 4.035 | 6.135 | 27.990 | 25.848 |
| -5.214 | -16.522 | -17.342 | -1.129 | -21.882 | 13.682 | 24.496 | 28.643 | 23.431 | 11.189 | 11.666 | 10.426 |
| -12.546 | -8.082 | 19.716 | 26.175 | -3.469 | -5.381 | 10.734 | 13.204 | -2.843 | 15.581 | 26.669 | 13.359 |
| 10.019 | 21.140 | -1.992 | 8.242 | -25.801 | -6.250 | -6.459 | 16.034 | -6.581 | 12.327 | 22.581 | 29.865 |
| 15.073 | 15.578 | 22.046 | 8.597 | -19.657 | -27.446 | -1.413 | 24.095 | 5.522 | 14.920 | 8.247 | 23.434 |
| 27.796 | 10.058 | 8.912 | -11.928 | -6.619 | 21.086 | -22.552 | 12.287 | -9.214 | 21.186 | 3.311 | 25.056 |
| -9.330 | 7.270 | -18.273 | -19.649 | -14.966 | 26.454 | -13.099 | 25.552 | 5.110 | 13.814 | 9.403 | 28.915 |
| 15.880 | 20.039 | -14.495 | 7.929 | -22.647 | 2.286 | 23.774 | 5.723 | 26.799 | 13.369 | 11.001 | 18.487 |
| -7.735 | 25.878 | -23.784 | 1.516 | -15.768 | -23.214 | 26.031 | 26.736 | -15.198 | 6.645 | 18.130 | 20.051 |
| -17.514 | 14.861 | 30.065 | 21.124 | -25.658 | -19.458 | -0.590 | -0.729 | 15.271 | 11.111 | 6.741 | -9.342 |
| -13.455 | 14.278 | 11.248 | 2.142 | 6.181 | 9.415 | 13.174 | 29.286 | 14.887 | 19.859 | 20.582 | -0.357 |
| 12.195 | -4.878 | 21.395 | 10.788 | -5.662 | -9.604 | 22.556 | 20.373 | -2.829 | 7.247 | 13.480 | 14.812 |
| 11.282 | 8.061 | 22.326 | 23.621 | -17.910 | 24.981 | -17.902 | 27.079 | 12.744 | 11.678 | 8.467 | 15.606 |
| 26.339 | -10.943 | -5.932 | 9.474 | -17.019 | -21.437 | 24.716 | 22.331 | 30.198 | -1.105 | 7.833 | 13.393 |
| -22.203 | 24.562 | -7.108 | 9.030 | -14.385 | 13.571 | -25.711 | 28.323 | 27.586 | -2.994 | 10.289 | 21.396 |
| 7.348 | 6.533 | -23.986 | 27.798 | 3.937 | 4.319 | 14.423 | -3.297 | 20.850 | 5.575 | 27.131 | 27.593 |
| -24.416 | -24.145 | -27.699 | 19.551 | -19.528 | -0.453 | 22.504 | 10.478 | -4.900 | -24.173 | 25.068 | 16.704 |
| -15.863 | -14.578 | 21.157 | -19.946 | 2.157 | 12.053 | -15.973 | 10.962 | 16.659 | 3.932 | 27.799 | -4.196 |
| 23.361 | -6.459 | -18.314 | 23.606 | -16.588 | 26.473 | 29.462 | -14.812 | 28.651 | 13.883 | 14.351 | 26.808 |
| 17.640 | -24.467 | -11.901 | 18.925 | 20.671 | 18.961 | -18.294 | 9.451 | 31.097 | -4.839 | 3.375 | 13.013 |

| | | | | | | | | | | | |
|---|---|---|---|---|---|---|---|---|---|---|---|
| -12.031 | 16.267 | 21.386 | 13.432 | -24.369 | 21.353 | 27.753 | 24.354 | -7.081 | 12.622 | 6.137 | 25.421 |
| -27.199 | 20.473 | 18.187 | 10.473 | -10.386 | 8.596 | 30.860 | 15.770 | 27.751 | 28.670 | 23.523 | 30.966 |
| 20.342 | 17.442 | -18.966 | 14.618 | 17.180 | 25.445 | 24.805 | 9.201 | 14.858 | 8.532 | 5.292 | 5.006 |
| -25.691 | 18.406 | -5.319 | 4.305 | -5.952 | 15.939 | -16.612 | -13.559 | -10.933 | 22.156 | 24.530 | 9.029 |
| 11.556 | -21.849 | -29.185 | 2.586 | 12.254 | 3.866 | 21.972 | 5.345 | 19.897 | 19.605 | 18.970 | 18.389 |
| 2.263 | 15.304 | 11.794 | 11.575 | 17.502 | 9.727 | -27.803 | -15.596 | 12.305 | 1.995 | 7.924 | 17.902 |
| -1.796 | 15.251 | 19.811 | 21.008 | -28.711 | -13.400 | -19.527 | -23.634 | 24.217 | 7.826 | 26.808 | -4.326 |
| 7.538 | 8.749 | 10.059 | 7.302 | -14.674 | -13.060 | 7.334 | 16.973 | 26.727 | -29.994 | 3.593 | 26.688 |
| 23.305 | -9.443 | -16.303 | 6.246 | -24.611 | 23.753 | -18.559 | 27.283 | 11.243 | -30.751 | 11.969 | 23.224 |
| 25.916 | -4.421 | -11.226 | 3.199 | -16.880 | 22.793 | 13.042 | 11.343 | -12.673 | 20.548 | 17.171 | 25.168 |
| 4.113 | 2.721 | -18.794 | 25.426 | -20.336 | -6.183 | -28.595 | 27.714 | 24.697 | 15.347 | 22.544 | 22.049 |
| 16.029 | -17.370 | -2.639 | 4.720 | -21.609 | 29.900 | 23.617 | 25.283 | 25.534 | 21.532 | 13.244 | 25.904 |
| 19.226 | -8.190 | -24.545 | 16.959 | -17.964 | 19.394 | 29.946 | 23.986 | -4.815 | 2.890 | 20.004 | 8.774 |
| 7.980 | 17.377 | -7.978 | 16.159 | -14.461 | 5.360 | 26.615 | -12.077 | -13.471 | 4.587 | 25.317 | 21.579 |
| -17.733 | 6.279 | -10.182 | 1.714 | -7.036 | -10.831 | -22.072 | 14.731 | 19.967 | 1.159 | 1.800 | 24.056 |
| -21.313 | 20.590 | -0.459 | 12.530 | 12.968 | 7.978 | -16.923 | 7.050 | 15.051 | 12.405 | 22.494 | 2.683 |
| 5.480 | -22.639 | 14.613 | -19.550 | -19.077 | 10.873 | 25.628 | 30.314 | 10.727 | 11.528 | 23.952 | 27.775 |
| -4.726 | 4.003 | 22.526 | -2.684 | -1.053 | 14.677 | 12.838 | 21.809 | 16.543 | 6.947 | 13.434 | 24.151 |
| -10.147 | 6.536 | -17.530 | -12.607 | -9.419 | 0.502 | 19.151 | -12.152 | 11.912 | 23.585 | 20.605 | 28.390 |
| -4.087 | -2.646 | 20.239 | 0.827 | 27.243 | 1.379 | 28.414 | -3.592 | 29.382 | 14.630 | 6.979 | 15.503 |
| 12.724 | -5.413 | 26.218 | -7.877 | -25.736 | 9.552 | 10.189 | 9.741 | 23.881 | 8.644 | 4.955 | 4.540 |
| 3.033 | 23.116 | -9.049 | 2.415 | -7.439 | 26.673 | -14.246 | 12.080 | -7.624 | 28.021 | 17.252 | 1.539 |
| -16.403 | 0.529 | 24.787 | 0.217 | -12.036 | 18.262 | -24.354 | 29.466 | 25.094 | 21.160 | 18.333 | 20.125 |
| 25.606 | 25.268 | -4.280 | -27.386 | -11.876 | -3.023 | -29.542 | -10.781 | 22.876 | -5.151 | 5.814 | 26.140 |
| -8.716 | 15.998 | 14.974 | 14.914 | -20.719 | 11.799 | 19.878 | 30.955 | 3.373 | -19.182 | 15.714 | 29.932 |
| 17.827 | -20.501 | 29.175 | 4.447 | -23.193 | 27.643 | -16.275 | 19.599 | 14.182 | 14.391 | 20.719 | 22.599 |
| 24.951 | 2.244 | 5.807 | 12.118 | -22.449 | -6.273 | 8.099 | 8.199 | 3.960 | 14.233 | 11.583 | 12.589 |
| 8.239 | 26.399 | -24.266 | 5.554 | -24.892 | 25.932 | 7.568 | 12.802 | 28.297 | 10.094 | 25.189 | 21.500 |
| 2.700 | -24.565 | -13.410 | 6.461 | 1.828 | 2.161 | -19.931 | 28.116 | 2.933 | 26.742 | 2.514 | 10.217 |
| -5.806 | -21.590 | 13.643 | -23.140 | -8.806 | 15.389 | -19.059 | 25.683 | -0.227 | 25.094 | 19.895 | 0.048 |
| 20.778 | 14.225 | -7.154 | 11.651 | -13.017 | 28.526 | -16.707 | -2.047 | -8.278 | 13.486 | 9.087 | 14.886 |
| -21.280 | 21.375 | 21.624 | 0.662 | -18.138 | 19.946 | 25.544 | 11.862 | 29.578 | 21.656 | 4.646 | 28.440 |
| 4.279 | -8.268 | -6.677 | 17.990 | -1.733 | -12.794 | 16.803 | 23.501 | 29.001 | -0.214 | -13.547 | 20.649 |
| 5.153 | 8.029 | -11.403 | 1.149 | 18.336 | 1.336 | -18.184 | 29.016 | 19.421 | 18.520 | -18.331 | 19.426 |
| 18.429 | 5.531 | 17.233 | 7.498 | -9.125 | -20.371 | 22.086 | 30.489 | -4.581 | 23.321 | -16.324 | 29.322 |

| | | | | | | | | | | | |
|---|---|---|---|---|---|---|---|---|---|---|---|
| -19.418 | 11.697 | 27.950 | 16.646 | -23.222 | -12.373 | 20.238 | 7.010 | -10.302 | 1.635 | -22.259 | 28.080 |
| -13.057 | -0.951 | -18.901 | -5.200 | -19.751 | -15.452 | -11.264 | -5.056 | 16.479 | -0.572 | -19.334 | 21.826 |
| 20.035 | 1.703 | -25.658 | 0.372 | -29.036 | 7.396 | -0.669 | 2.298 | 28.693 | 5.976 | -15.648 | 7.824 |
| -0.080 | 27.667 | -10.346 | -11.619 | 18.976 | -19.978 | 28.470 | 17.348 | -1.648 | -3.801 | -16.796 | 15.717 |
| 2.684 | 21.381 | -16.041 | 11.440 | 27.927 | 16.164 | 27.689 | 21.005 | 26.454 | 25.340 | 12.961 | 17.546 |
| -4.921 | 4.865 | -18.401 | -21.599 | -25.125 | 11.891 | 31.524 | -6.163 | 7.993 | 4.851 | 16.517 | 23.116 |
| 21.407 | -2.751 | 23.379 | 9.762 | -17.334 | -4.345 | -22.393 | -8.599 | 3.196 | 4.194 | 24.241 | 28.325 |
| 18.563 | 26.281 | -19.673 | 16.998 | -16.456 | 10.520 | 5.501 | 7.391 | -1.072 | 24.226 | 21.507 | 6.424 |
| 11.507 | -3.938 | -21.070 | -0.814 | -22.614 | 6.896 | -14.827 | 29.469 | 18.191 | 10.000 | 7.323 | 20.120 |
| 22.145 | -24.655 | 19.977 | 5.852 | 16.482 | -10.209 | 17.846 | 26.647 | -7.568 | 19.069 | 17.981 | -14.747 |
| 4.984 | 13.199 | -25.993 | -1.801 | 0.138 | -19.822 | 7.857 | 21.345 | -12.229 | 22.584 | 12.699 | 0.324 |
| -7.014 | 6.973 | 15.053 | -23.503 | -20.025 | -15.062 | 8.606 | -3.314 | 23.758 | 13.563 | -13.806 | -4.929 |
| -12.708 | -3.268 | -1.761 | 8.578 | 17.277 | -23.394 | 23.479 | 25.087 | -3.295 | 11.124 | 14.242 | -0.123 |
| -3.061 | -0.548 | 25.658 | -3.604 | 29.842 | -16.072 | 13.354 | 5.267 | 20.656 | 6.709 | 23.578 | 9.662 |
| -23.366 | -4.475 | 4.642 | 24.297 | -7.984 | -20.660 | -23.612 | -1.138 | 7.286 | 5.312 | 24.249 | 16.528 |
| -22.180 | -16.578 | 19.286 | 3.437 | -17.085 | 2.408 | 24.396 | -29.062 | 23.782 | 20.039 | 14.994 | 11.737 |
| -24.753 | 14.960 | -19.114 | 7.789 | 4.040 | 18.484 | 15.078 | 22.372 | 2.895 | 10.570 | -16.398 | 18.099 |
| 18.037 | 8.743 | 24.004 | 20.658 | -9.705 | 11.548 | -18.009 | 21.750 | 13.255 | 12.283 | 17.931 | 8.552 |
| -9.110 | -3.188 | 21.280 | 17.043 | -6.814 | 6.641 | 21.294 | 10.876 | 21.893 | -12.221 | 26.453 | 21.267 |
| 16.900 | -1.823 | 27.012 | 15.376 | -23.081 | -13.047 | 26.973 | 28.665 | 25.686 | 0.780 | 23.853 | 10.523 |
| 24.428 | -13.107 | 9.599 | 4.639 | -15.868 | 23.487 | -13.438 | 3.285 | 25.684 | 18.859 | -3.196 | -7.044 |
| -10.658 | -13.485 | -15.543 | -24.842 | 5.992 | -22.277 | 26.419 | -11.081 | 12.237 | 24.721 | 26.329 | 27.629 |
| -2.452 | 30.327 | -3.810 | 6.454 | 24.029 | -15.349 | -10.675 | -27.839 | -1.867 | 1.426 | 23.214 | 7.208 |
| -19.789 | 20.957 | -5.007 | -3.611 | 18.964 | -1.377 | -18.820 | 7.715 | 22.706 | -2.336 | 22.349 | 24.464 |
| -22.658 | -17.492 | -11.468 | -14.053 | 15.802 | 27.643 | 5.841 | -0.728 | -7.111 | 26.748 | 21.903 | 20.440 |
| -0.292 | -19.648 | -8.190 | -21.910 | 7.950 | 19.920 | -8.827 | 25.401 | -14.833 | 9.947 | 25.156 | 14.788 |
| 7.795 | 14.952 | 2.487 | 17.921 | -29.308 | 21.953 | -21.346 | 2.112 | -11.374 | 26.788 | 20.550 | 1.244 |
| -0.904 | -16.924 | 24.735 | 10.901 | -13.022 | -19.341 | -16.027 | 27.904 | 8.253 | 0.848 | -21.353 | 2.920 |
| -17.125 | -24.768 | -12.717 | 6.557 | 4.452 | 14.877 | 24.181 | -0.381 | -9.531 | -6.298 | 18.121 | 4.233 |
| -24.021 | 6.476 | -7.779 | -5.961 | -17.458 | 13.206 | -13.171 | 2.863 | -2.510 | 19.180 | 24.205 | 22.226 |
| 6.418 | 19.159 | 3.885 | -7.920 | -20.873 | -22.095 | 16.879 | -4.597 | 7.505 | 11.588 | 21.062 | 27.744 |
| -9.462 | 24.657 | -24.350 | -0.105 | 25.643 | 22.522 | 13.835 | 24.778 | -22.844 | 7.050 | 18.770 | 17.203 |
| -21.423 | 24.275 | 20.271 | 15.553 | -25.455 | -10.334 | -14.792 | 9.556 | 18.017 | 4.757 | 24.965 | -10.963 |
| 5.427 | -3.915 | -19.104 | -14.879 | -10.674 | -11.464 | 23.694 | 8.662 | -5.664 | 25.494 | 21.510 | 6.827 |
| -16.011 | 13.542 | 13.847 | -22.127 | -18.859 | -9.022 | -30.714 | 26.275 | 7.672 | 8.128 | -1.296 | 19.496 |

| | | | | | | | | | | | |
|---|---|---|---|---|---|---|---|---|---|---|---|
| 1.111 | -24.223 | -13.912 | 6.649 | -7.977 | -14.373 | -14.145 | -22.359 | 21.692 | 16.326 | 13.668 | 11.892 |
| -11.014 | -10.211 | -14.955 | -4.312 | -14.511 | 9.908 | -19.668 | 19.659 | 22.065 | 23.209 | 26.982 | 5.027 |
| -4.315 | -15.000 | 17.805 | -8.572 | 30.051 | 2.393 | 17.933 | -6.864 | 16.416 | -8.844 | 26.109 | 21.435 |
| -22.266 | 24.356 | -25.576 | 20.227 | 7.359 | -3.261 | -23.259 | 7.213 | -0.400 | 13.429 | 14.062 | 9.560 |
| -21.377 | -18.548 | 24.553 | -9.270 | -25.634 | 9.276 | -1.227 | -18.795 | 24.423 | 14.784 | 25.274 | 19.210 |
| -18.961 | -5.595 | -20.631 | -6.984 | 2.817 | -6.528 | 21.088 | 25.631 | 27.054 | 24.442 | -20.245 | 16.892 |
| -8.858 | 12.286 | -0.695 | 4.853 | -3.212 | 22.627 | -15.722 | -8.807 | -8.584 | 18.036 | 19.533 | 23.205 |
| -16.212 | 6.811 | -14.460 | -14.470 | -14.193 | -27.079 | -20.676 | 26.992 | -0.086 | 11.316 | 3.397 | 17.371 |
| -25.866 | 14.126 | 15.425 | 12.521 | -15.407 | 4.395 | 13.778 | -11.859 | 19.221 | 17.018 | 22.526 | 14.087 |
| 4.941 | -6.123 | 4.410 | -17.664 | -19.616 | -11.628 | 14.464 | -9.811 | 25.140 | 13.391 | 21.815 | 22.188 |
| 10.181 | -20.983 | 25.525 | -24.294 | 4.430 | -2.653 | 29.691 | 27.928 | 18.489 | -5.165 | 18.013 | 23.869 |
| 11.163 | 23.217 | 7.499 | -7.393 | -26.435 | 6.448 | -16.914 | 21.150 | 23.028 | -2.140 | 22.998 | 27.407 |
| 23.811 | -23.033 | -23.367 | -20.741 | -0.361 | -12.095 | -11.624 | 8.859 | -10.411 | 10.931 | 11.859 | 12.936 |
| 19.053 | 8.255 | 24.128 | -26.228 | -18.236 | 9.913 | -23.538 | 0.945 | -19.028 | -10.110 | 15.986 | -8.425 |
| -13.800 | -29.658 | -24.294 | 2.242 | -16.150 | 3.171 | 16.744 | -4.465 | 19.028 | 22.881 | 13.882 | 20.409 |
| -6.910 | 10.890 | 8.464 | 15.312 | 25.582 | 25.414 | 3.339 | 20.329 | -7.347 | 3.378 | 21.543 | 25.093 |
| -16.227 | 28.707 | -5.927 | -5.804 | 28.221 | 9.175 | 22.522 | 2.953 | 6.301 | 21.065 | -7.972 | 22.210 |
| 13.934 | -17.042 | 22.262 | 15.607 | 9.059 | 0.688 | -24.411 | 20.702 | 26.309 | -9.416 | 29.592 | 15.456 |
| -6.132 | 24.763 | -13.726 | 7.758 | -11.197 | -4.247 | -13.235 | -10.346 | 3.036 | 4.596 | 27.881 | 18.124 |
| -24.931 | -4.437 | 19.839 | -1.557 | -13.756 | -6.222 | 5.124 | 26.555 | 6.747 | 8.405 | -19.750 | 25.006 |
| -20.309 | -19.114 | 22.799 | 6.900 | -10.260 | -7.901 | 21.431 | 24.687 | 16.900 | 20.511 | 14.524 | 4.194 |
| -1.365 | 8.882 | 6.713 | 1.567 | -14.166 | -13.391 | -8.719 | -5.489 | 22.671 | 5.443 | 23.944 | 27.600 |
| 6.148 | -15.006 | -1.498 | -18.930 | -12.064 | -5.526 | -21.587 | 15.511 | -9.272 | 9.308 | 19.887 | 20.152 |
| -4.584 | 28.960 | -21.625 | -21.399 | -20.240 | 11.923 | -29.261 | 19.757 | 7.461 | 16.048 | 21.598 | 26.579 |
| -9.851 | 7.666 | -2.814 | -0.389 | -15.540 | -20.321 | 27.479 | 26.039 | 19.940 | -1.417 | -0.568 | 23.835 |
| 22.047 | -4.806 | 17.223 | -14.313 | -6.249 | 24.713 | 28.887 | 12.796 | -25.127 | 7.734 | 15.581 | 16.398 |
| 14.256 | 20.116 | 13.647 | 9.925 | 5.470 | -2.666 | -22.473 | 0.587 | 17.413 | 6.304 | 18.607 | 25.319 |
| -2.849 | -11.747 | 0.902 | 6.440 | -7.895 | 7.299 | 28.178 | 19.112 | 5.194 | 1.929 | 30.097 | 23.630 |
| -8.207 | -7.086 | -10.604 | 3.545 | -9.695 | -8.766 | 4.095 | 22.675 | 26.433 | -0.627 | 28.543 | 11.525 |
| -22.067 | 10.811 | 21.635 | 11.616 | -6.791 | -10.203 | -8.440 | 8.461 | 6.307 | -7.299 | -10.337 | 25.914 |
| 19.602 | 22.805 | 0.160 | 5.169 | 10.603 | 8.817 | 25.345 | 5.092 | -27.074 | 4.237 | 19.761 | 20.380 |
| -1.221 | 13.136 | -8.390 | 2.183 | -24.316 | 12.899 | 22.499 | 20.197 | 2.064 | -26.052 | 20.521 | 23.743 |
| -10.058 | 21.262 | -21.893 | 13.484 | 0.740 | -17.601 | -7.924 | 25.821 | 25.794 | 25.280 | 22.834 | 24.782 |
| 13.498 | -26.027 | -22.080 | 8.509 | 6.595 | -0.398 | -23.924 | 11.263 | -1.045 | 14.399 | 18.500 | -2.104 |
| 14.659 | -31.503 | 8.103 | -19.043 | -12.643 | -6.235 | -20.763 | -0.403 | 21.152 | 23.337 | 27.545 | 10.076 |

| | | | | | | | | | | | |
|---|---|---|---|---|---|---|---|---|---|---|---|
| -7.113 | 3.367 | -9.037 | 14.612 | 11.541 | 11.401 | -14.271 | -17.863 | -26.730 | 13.711 | 14.259 | 17.343 |
| -10.656 | -19.177 | -17.428 | -10.035 | -22.483 | -15.761 | -13.316 | 23.267 | 25.628 | 12.561 | 29.182 | 20.371 |
| -8.301 | -4.384 | 19.530 | 20.674 | 10.036 | 14.270 | -11.677 | 23.818 | -15.504 | 8.545 | 17.980 | 20.933 |
| -15.321 | -6.612 | -0.760 | -12.090 | -21.908 | 8.061 | 5.850 | 8.263 | 16.492 | 12.245 | 8.569 | 22.496 |
| 7.777 | -15.951 | -19.033 | -7.438 | 3.036 | 2.600 | -19.332 | 15.729 | 14.906 | 13.314 | 14.480 | 5.474 |
| -13.701 | 7.755 | -7.861 | 17.848 | -13.886 | 6.586 | 10.115 | 1.064 | 18.143 | 3.096 | 19.912 | -6.028 |
| 18.613 | 26.469 | -24.176 | 16.863 | -23.001 | 26.410 | 12.888 | -24.826 | -21.985 | 1.397 | 23.966 | 2.589 |
| -24.450 | 20.925 | 19.098 | 4.708 | -15.678 | 15.116 | -16.653 | 6.930 | 4.186 | -23.030 | 8.977 | 19.146 |
| -6.111 | -2.511 | 6.460 | -17.386 | 4.311 | 4.752 | 28.295 | -6.666 | -13.541 | 23.895 | 25.577 | 12.882 |
| -8.468 | 0.038 | -20.745 | 29.506 | -7.870 | -2.442 | -13.759 | 19.014 | 5.688 | 5.703 | 17.057 | 25.356 |
| -2.596 | -3.980 | 8.819 | -8.154 | -8.534 | 13.443 | -4.540 | 11.744 | -1.039 | -3.617 | -5.568 | 23.229 |
| 25.042 | -18.383 | -2.976 | 2.909 | -3.863 | -8.353 | -26.182 | 21.384 | -5.846 | -25.864 | 26.585 | 7.488 |
| 27.465 | -5.445 | -26.039 | 23.931 | 5.024 | -16.876 | 21.662 | 24.485 | 20.669 | 8.082 | 22.458 | 17.879 |
| 14.059 | -30.597 | -4.635 | -7.499 | -10.385 | -0.151 | -8.672 | 19.489 | -15.031 | -2.228 | -0.420 | 18.917 |
| 22.272 | -7.950 | 0.473 | -24.699 | 16.677 | -14.170 | 18.335 | -5.117 | -14.310 | 19.480 | 30.014 | 22.691 |
| -11.925 | -1.222 | 25.044 | 16.087 | 10.403 | -0.260 | -8.124 | -4.355 | 20.943 | 23.617 | 24.895 | 15.722 |
| 25.768 | -19.266 | 17.742 | 4.299 | -5.350 | -12.077 | -20.145 | 1.827 | 19.084 | 14.892 | 22.913 | 18.225 |
| 14.527 | 21.793 | 19.496 | 1.267 | -20.349 | 7.874 | -15.596 | 11.949 | -11.191 | -4.952 | -13.629 | 20.952 |
| 19.275 | -7.180 | 18.605 | 15.196 | 25.371 | 3.371 | -4.058 | -0.730 | 5.297 | 3.345 | 20.351 | 19.429 |
| -4.351 | 27.422 | -15.384 | 19.407 | -25.267 | -1.384 | 23.068 | 21.097 | -13.245 | 9.157 | 13.822 | 12.083 |
| 21.178 | -20.111 | 17.317 | -4.766 | 6.309 | 5.911 | -9.713 | -1.057 | 7.529 | -25.226 | 14.518 | 19.143 |
| 27.701 | 19.429 | -10.649 | 12.808 | -13.088 | 11.578 | -28.025 | 21.701 | 15.095 | 22.338 | -3.413 | 11.506 |
| 19.440 | 1.827 | -16.255 | -21.565 | -5.974 | 1.596 | 25.365 | 1.521 | 19.816 | 10.367 | 12.700 | 10.199 |
| 20.809 | 0.239 | 20.264 | 7.483 | -21.178 | -2.842 | 13.011 | 20.362 | -15.868 | 12.847 | -0.937 | 8.313 |
| -7.913 | 9.960 | 23.922 | 30.069 | -17.340 | 21.020 | 9.593 | 3.082 | 23.008 | 16.235 | -22.377 | 6.615 |
| 9.112 | 25.790 | 19.203 | 22.968 | -2.795 | -5.401 | 1.632 | -0.170 | 21.395 | 7.868 | 17.390 | 4.453 |
| -19.154 | -4.137 | 6.716 | 15.352 | -7.470 | -3.338 | 7.695 | 3.544 | 11.036 | -6.407 | 17.923 | 1.235 |
| -16.500 | -9.588 | 17.057 | -12.566 | -9.180 | -15.679 | -22.438 | -18.211 | 23.923 | 21.160 | -5.579 | -2.545 |
| 15.592 | 24.809 | -4.880 | 28.143 | 1.156 | 5.453 | -3.063 | 10.579 | 15.740 | 15.714 | 27.957 | -5.246 |
| -21.704 | 8.213 | -12.852 | -4.781 | 9.259 | 12.863 | -5.759 | 1.844 | 12.119 | 3.641 | -3.170 | 9.210 |
| 22.526 | 8.947 | 23.923 | 5.805 | -11.031 | 25.792 | 17.948 | 9.393 | -14.871 | 22.793 | 22.011 | 22.712 |
| 0.379 | -5.920 | 12.963 | 16.181 | -4.449 | -0.554 | 11.888 | -15.532 | 6.653 | 10.273 | 26.492 | 21.284 |
| 9.765 | -10.134 | 7.453 | 10.041 | -0.917 | -4.784 | 11.142 | 21.731 | 26.355 | 8.209 | 11.007 | 18.049 |
| 24.899 | 6.610 | -18.108 | 3.493 | -13.509 | 19.251 | 22.498 | 22.843 | 17.713 | -7.904 | -16.667 | 20.324 |
| -20.763 | -19.933 | 21.904 | 0.134 | -1.390 | 12.361 | -24.554 | 24.995 | -6.447 | 20.038 | 14.974 | 24.390 |

| | | | | | | | | | | | |
|---|---|---|---|---|---|---|---|---|---|---|---|
| -18.372 | 9.855 | 24.664 | 27.462 | 7.636 | 10.561 | 6.706 | 10.543 | 25.330 | 6.228 | 28.388 | 12.341 |
| -24.640 | -4.543 | 23.978 | -19.734 | -3.826 | -25.794 | -0.533 | -10.449 | 15.497 | 5.458 | 17.080 | 25.213 |
| 13.216 | -6.885 | 27.630 | -10.241 | -18.254 | -1.971 | -12.113 | -11.230 | 22.984 | -16.778 | 23.704 | 19.806 |
| -6.649 | -5.642 | -22.262 | 28.981 | 24.518 | -5.299 | 10.211 | 25.916 | 21.960 | 17.033 | 11.520 | 15.847 |
| 26.013 | -24.992 | 6.575 | -24.947 | -26.511 | 9.667 | 7.471 | 3.752 | 23.965 | 17.706 | 2.988 | 12.017 |
| 29.133 | 3.989 | -0.829 | 17.992 | 7.081 | 11.867 | 8.814 | 12.526 | 11.194 | 3.365 | -1.649 | -2.690 |
| -29.743 | -23.142 | 21.871 | 5.051 | -8.278 | 1.878 | -17.692 | 27.107 | 25.995 | 21.851 | 8.392 | 24.460 |
| 22.271 | -13.617 | 17.084 | 23.634 | -17.632 | -2.661 | 13.568 | 0.726 | 19.623 | 24.710 | 20.195 | 21.920 |
| -16.922 | -5.303 | -28.987 | 0.540 | -22.451 | -14.128 | 4.226 | 21.885 | 14.804 | 15.583 | 27.319 | 19.152 |
| 18.733 | -19.820 | -18.786 | -26.583 | 5.388 | 3.005 | -27.336 | 25.567 | -17.870 | 1.568 | 25.152 | -13.875 |
| 8.965 | 27.175 | -23.601 | 27.234 | 0.415 | -7.939 | -11.233 | 8.197 | 18.484 | 9.490 | 21.180 | 18.563 |
| 25.660 | 16.417 | -22.420 | -6.916 | -23.032 | -14.632 | -12.405 | 4.049 | 5.276 | 25.237 | 24.250 | 15.714 |
| 22.868 | -14.378 | 3.956 | 24.986 | -11.719 | -13.711 | 9.558 | 23.346 | 12.929 | 0.325 | 22.494 | 9.446 |
| 28.818 | -7.160 | 28.159 | 19.829 | -23.717 | 1.469 | 11.986 | 3.077 | 15.986 | 6.184 | -20.469 | 24.786 |
| 26.512 | -2.193 | 12.340 | 25.629 | -3.044 | 15.755 | 1.972 | 20.208 | -16.539 | 10.687 | 27.741 | 23.846 |
| 24.755 | -22.614 | -14.387 | -8.607 | 1.252 | 0.297 | 7.550 | 12.670 | 1.703 | -0.426 | 4.141 | 20.535 |
| 17.542 | -26.642 | 17.500 | -18.860 | 13.062 | -24.117 | -21.564 | 26.827 | -1.177 | 9.069 | 3.381 | 13.518 |
| 9.470 | -7.949 | -16.372 | -1.765 | 4.736 | 20.623 | 22.438 | 18.554 | 21.073 | 18.516 | 25.838 | 19.340 |
| 19.818 | -18.917 | -9.307 | 6.633 | -17.631 | -26.397 | -5.649 | 22.623 | -5.482 | 23.443 | -8.822 | -12.089 |
| 24.082 | 16.397 | 25.420 | -17.053 | -7.789 | 7.101 | -10.957 | -20.739 | 23.932 | 7.286 | 15.565 | 15.140 |
| -17.501 | 13.029 | -17.228 | -18.128 | -13.817 | 2.239 | -19.429 | 14.356 | -19.880 | 5.721 | 21.810 | 11.178 |
| 23.178 | 24.264 | -6.705 | -6.500 | -19.811 | -1.617 | -7.812 | 19.754 | 18.301 | 26.252 | 16.667 | 21.654 |
| -23.972 | -16.489 | 26.527 | -4.427 | -22.246 | -25.902 | -4.944 | 18.972 | 7.309 | 10.078 | 23.541 | 22.983 |
| -26.903 | -1.326 | -5.582 | 10.250 | -6.495 | 5.042 | -3.610 | 23.581 | 17.508 | 21.683 | 17.357 | 27.100 |
| 8.225 | -15.546 | -26.894 | 4.866 | -11.916 | 6.626 | -2.245 | 4.114 | 11.979 | 8.335 | 5.211 | 15.924 |
| 23.702 | -3.214 | -19.336 | 27.249 | 3.326 | 16.335 | -23.943 | -26.594 | -3.426 | 27.275 | -7.919 | -3.193 |
| 22.092 | 7.609 | 16.494 | -6.310 | -16.249 | 0.780 | 6.325 | 23.589 | -10.971 | 1.831 | 6.403 | 23.397 |
| -10.570 | -11.299 | 13.470 | 9.133 | 27.483 | 8.161 | 22.209 | 19.156 | 23.114 | 18.467 | 12.367 | 22.358 |
| 9.888 | -5.435 | 5.242 | -3.309 | -14.631 | -4.233 | -4.481 | 22.123 | 20.493 | -4.532 | 22.784 | 10.147 |
| -14.283 | 15.571 | 23.767 | -30.316 | -10.075 | 3.470 | -25.513 | 14.285 | 7.057 | 16.342 | 2.805 | 4.658 |
| -30.243 | 10.621 | -24.265 | 20.103 | 10.576 | 19.253 | -14.971 | 7.543 | 8.315 | 3.200 | 19.700 | 16.293 |
| 25.855 | -1.641 | -29.706 | -20.873 | -5.213 | 17.823 | 13.840 | 12.098 | 23.694 | 10.653 | 24.295 | 21.177 |
| -9.335 | 9.420 | -20.436 | 26.169 | -25.603 | -16.131 | 0.182 | 9.110 | 21.915 | -1.329 | -10.937 | 19.556 |
| -25.796 | 20.390 | 15.578 | 17.077 | -20.124 | -7.700 | -16.263 | 21.788 | 13.085 | 19.712 | -5.253 | 8.721 |
| 19.284 | 0.270 | -18.796 | -17.392 | 7.727 | -6.617 | 4.673 | 26.034 | 21.051 | 7.833 | 18.465 | 12.942 |

| | | | | | | | | | | | |
|---|---|---|---|---|---|---|---|---|---|---|---|
| -26.655 | -19.758 | 14.077 | -22.785 | 3.842 | 5.050 | -20.341 | 14.269 | 29.856 | 14.153 | 12.921 | -20.106 |
| 0.074 | -2.490 | 7.559 | -29.943 | 5.595 | 24.659 | -1.440 | 25.228 | 7.049 | 3.485 | 17.739 | 7.015 |
| -14.620 | -17.656 | 18.041 | 18.107 | 2.205 | 19.588 | -22.246 | -10.597 | -16.744 | 12.781 | 27.380 | 25.961 |
| 26.301 | 12.536 | 23.808 | 23.335 | -15.477 | 21.400 | 14.213 | 11.387 | 22.854 | 8.009 | -24.287 | 13.886 |
| 24.428 | -12.460 | -19.198 | 5.966 | -1.367 | 24.152 | 10.581 | 23.703 | -0.910 | -0.627 | -20.491 | 28.872 |
| 29.296 | 8.358 | -18.344 | 3.135 | -9.290 | -24.500 | -5.864 | 30.235 | 6.562 | 7.271 | 20.227 | -7.315 |
| 9.633 | -25.081 | 17.621 | -29.417 | 7.093 | 26.095 | 2.844 | 17.117 | -14.386 | 23.797 | 19.146 | -5.198 |
| -25.930 | -15.811 | -14.346 | 25.325 | -2.499 | 11.083 | 6.359 | -4.257 | -3.650 | 19.641 | 22.406 | 3.579 |
| -23.555 | 10.389 | -29.043 | 23.439 | 0.373 | -3.597 | -3.041 | 4.919 | 6.271 | 2.098 | 24.779 | 24.112 |
| 14.997 | -20.710 | -11.741 | 13.478 | -12.465 | 18.239 | 13.709 | 19.005 | -6.174 | 1.251 | -8.976 | 28.176 |
| -24.125 | 14.451 | 9.551 | 10.352 | 0.980 | 1.428 | -22.934 | 9.156 | 14.245 | 5.403 | -12.206 | 21.300 |
| -10.063 | -22.072 | -5.198 | 11.702 | -15.080 | 19.489 | -4.907 | 25.336 | 13.146 | 8.206 | 22.238 | 11.437 |
| 9.398 | 15.282 | 24.660 | -28.776 | -4.096 | -9.344 | 11.470 | 21.158 | 24.536 | 13.563 | 19.353 | 15.575 |
| -3.463 | -8.459 | -17.591 | 14.394 | -4.512 | 5.521 | 3.110 | 0.878 | 0.775 | 19.262 | 3.981 | 9.102 |
| -30.542 | -6.678 | -29.967 | -1.478 | -3.431 | 9.618 | -20.392 | -0.546 | -7.353 | 15.532 | 13.121 | 25.496 |
| -10.492 | -15.218 | -2.022 | 23.335 | 1.647 | 26.183 | -10.373 | 11.552 | 22.939 | 25.461 | 4.999 | 12.874 |
| -2.654 | 9.571 | -19.465 | 7.596 | -16.555 | -2.368 | -21.652 | -1.997 | 12.608 | 23.478 | 23.794 | -0.578 |
| 21.808 | 17.514 | -31.650 | -15.501 | 8.712 | 19.361 | -5.458 | -5.778 | 7.107 | 21.056 | 15.520 | 22.518 |
| 22.980 | -15.285 | -12.524 | 19.975 | 7.125 | 22.913 | 9.745 | 6.153 | 17.703 | 12.341 | 24.438 | 24.531 |
| 17.828 | 1.957 | -23.252 | -25.149 | -18.792 | 20.222 | -1.889 | 24.509 | 9.634 | 24.006 | 15.211 | 26.412 |
| 18.357 | -16.961 | 9.856 | 10.428 | -0.759 | 23.816 | -13.823 | 12.471 | 21.368 | 9.803 | -0.749 | 5.575 |
| 27.688 | 15.035 | -22.431 | 23.783 | 6.177 | -4.637 | 7.562 | -8.917 | 16.510 | 4.330 | -4.098 | 3.283 |
| -5.732 | -0.178 | -5.509 | -26.016 | -10.295 | -13.175 | -20.422 | 18.630 | 30.032 | 3.335 | 19.897 | 29.275 |
| -4.636 | -8.630 | 30.672 | -20.480 | 25.409 | -9.666 | -10.206 | 8.493 | 4.288 | 18.309 | -0.826 | 9.634 |
| 16.328 | -0.768 | -3.153 | 0.143 | -14.675 | -20.985 | 3.111 | 6.672 | -11.361 | 7.484 | -1.568 | 6.757 |
| -22.826 | -9.406 | 19.869 | 5.781 | -21.063 | -25.797 | -8.062 | -2.973 | 15.347 | 26.707 | 23.080 | 28.020 |
| -7.068 | -6.320 | 28.472 | 25.646 | -17.855 | -17.140 | 2.620 | 19.828 | -28.966 | 25.702 | 26.045 | 17.384 |
| -19.083 | -8.678 | -8.468 | 23.687 | -15.445 | 24.470 | 1.217 | 29.119 | 22.724 | 18.341 | 6.300 | 7.864 |
| 19.692 | 3.901 | 0.356 | 14.274 | 7.610 | -7.244 | 10.092 | 4.049 | 14.294 | 26.235 | -0.909 | 14.953 |
| -0.193 | 18.502 | 10.379 | -11.578 | -0.150 | 5.645 | 22.515 | 13.423 | -6.929 | 16.600 | 16.626 | -2.434 |
| 5.689 | 5.019 | -1.484 | -9.700 | -16.465 | 12.958 | -5.403 | 20.451 | 17.687 | 15.516 | 22.387 | 19.825 |
| 24.130 | -9.380 | -21.989 | -22.192 | -23.755 | -4.974 | 8.123 | -2.593 | 9.772 | 19.761 | 18.609 | 0.952 |
| 16.666 | 23.043 | -3.868 | 18.001 | 24.505 | 8.680 | 15.530 | 28.950 | 14.005 | 17.272 | 9.471 | 25.309 |
| -3.128 | 6.315 | -18.725 | -18.457 | 23.469 | -0.835 | -0.899 | 21.571 | 23.835 | 8.433 | 18.214 | 21.019 |
| -14.720 | -11.974 | 7.666 | 6.061 | 22.118 | 17.987 | -10.439 | 25.582 | 18.556 | 18.629 | 12.105 | -10.854 |

| | | | | | | | | | | | |
|---|---|---|---|---|---|---|---|---|---|---|---|
| -9.201 | 21.676 | -0.923 | 5.150 | 20.039 | -26.758 | -3.071 | 14.796 | 12.572 | 6.577 | 22.719 | 8.895 |
| 26.533 | -9.754 | 30.289 | 17.479 | 21.392 | 5.577 | 20.907 | 26.356 | 17.071 | 28.564 | 10.920 | 27.248 |
| 25.236 | -3.173 | -12.339 | -24.476 | 19.004 | 25.333 | 5.212 | 0.274 | -2.629 | 24.767 | 19.301 | 18.878 |
| -29.811 | 4.823 | 17.294 | 6.700 | 20.801 | 29.340 | -6.955 | 9.581 | 4.252 | 10.138 | 21.973 | 0.890 |
| -28.156 | -19.164 | -11.563 | -7.478 | 0.589 | -9.407 | -4.217 | 5.419 | 12.164 | 5.441 | 3.254 | 27.476 |
| -16.590 | 1.068 | -13.016 | -8.873 | -0.210 | 3.817 | -14.640 | 13.457 | 19.565 | 13.539 | 6.414 | 24.149 |
| 13.389 | 10.358 | 15.510 | -13.486 | 17.911 | 7.998 | 24.867 | 3.701 | 11.303 | 12.155 | 24.425 | 14.815 |
| -5.222 | -4.157 | -2.000 | -10.547 | -17.558 | 25.976 | -18.670 | 14.536 | 23.089 | 12.866 | 20.196 | 5.476 |
| 19.370 | 10.888 | 11.707 | -16.073 | -1.281 | -21.058 | 19.788 | 6.560 | -6.096 | 7.148 | 14.881 | 28.306 |
| -8.215 | 2.800 | 14.465 | -5.513 | -3.856 | -6.657 | 0.613 | 11.108 | 20.660 | 22.688 | 20.444 | 24.079 |
| -11.631 | -0.389 | 7.563 | 12.291 | -4.824 | 14.873 | -12.284 | 19.438 | 17.659 | 17.934 | 25.220 | 26.553 |
| 22.094 | 13.825 | 23.740 | 13.965 | -8.900 | -29.794 | -25.402 | -22.088 | 30.145 | 8.097 | 19.198 | -1.006 |
| -20.356 | 23.705 | 16.645 | 20.989 | -18.450 | -28.919 | 24.385 | 25.825 | -11.386 | -3.945 | 5.039 | -2.748 |
| 9.666 | -10.625 | -14.926 | -16.540 | 2.573 | 26.068 | 17.349 | 23.815 | 24.604 | 12.702 | 21.820 | 20.206 |
| 0.068 | 4.083 | -24.761 | -21.019 | -19.471 | 1.492 | -24.310 | 9.113 | 22.276 | 27.948 | -2.413 | 28.831 |
| 4.727 | 5.354 | -20.353 | 6.027 | -23.165 | -9.535 | -23.514 | -1.580 | 10.625 | 9.371 | 3.794 | 25.347 |
| -6.864 | -10.092 | 5.463 | 8.652 | 25.242 | 10.015 | 25.583 | -2.890 | -22.548 | -1.828 | -17.634 | 23.446 |
| -11.574 | -25.962 | -19.166 | -15.473 | 11.905 | -10.939 | 2.104 | 26.592 | 29.309 | -2.938 | 23.951 | 9.976 |
| -20.001 | -12.899 | -3.302 | 19.865 | 26.072 | -1.045 | 26.968 | 16.114 | 7.592 | -1.141 | 26.647 | 30.359 |
| -18.271 | -8.210 | 21.684 | -25.074 | 1.341 | -13.464 | 10.428 | 30.128 | 23.947 | 23.679 | -3.279 | 15.735 |
| 25.511 | -5.277 | 16.838 | -6.513 | 19.456 | -18.810 | 7.121 | -0.469 | 4.257 | 24.599 | 22.955 | 10.679 |
| 21.034 | 9.329 | -7.332 | -14.575 | -0.121 | 8.189 | -20.996 | 30.845 | 28.967 | 8.209 | 2.480 | 14.132 |
| -13.728 | 7.159 | 22.869 | 2.546 | -20.541 | 22.403 | 19.824 | -21.935 | 5.760 | 21.759 | 25.494 | 26.858 |
| 18.071 | -0.841 | 0.756 | 15.507 | -23.636 | -10.432 | 5.349 | 22.858 | 12.400 | 27.080 | 28.265 | 11.328 |
| -10.497 | 0.757 | -18.069 | 24.969 | 26.166 | 27.758 | -4.361 | 15.139 | 8.335 | 6.684 | 25.045 | 3.167 |
| -28.252 | 23.705 | -17.565 | 14.335 | 8.836 | 19.993 | 15.165 | 24.798 | -18.465 | 28.284 | 22.142 | 18.240 |
| 10.984 | -20.779 | -24.276 | -5.850 | -1.638 | 6.585 | 21.172 | 31.240 | 15.688 | 14.392 | 24.608 | 22.958 |
| 27.989 | 2.246 | 17.450 | -14.699 | -15.010 | -20.086 | -25.120 | -11.939 | 10.442 | 16.647 | 17.844 | 30.647 |
| -7.791 | -13.143 | 10.481 | 8.028 | 10.602 | 31.232 | 22.367 | 12.939 | -1.162 | 0.384 | 23.322 | 24.458 |
| 0.563 | -2.250 | 20.781 | -4.110 | -4.991 | -6.725 | -26.257 | 13.834 | 18.793 | 20.959 | 8.717 | -20.336 |
| -13.098 | 11.693 | -16.480 | 6.531 | -26.443 | -7.475 | -2.528 | 18.819 | -4.188 | 15.415 | 18.943 | 5.239 |
| -22.803 | 13.205 | -20.546 | 22.040 | -9.124 | 5.543 | 17.558 | 26.933 | -22.506 | 17.384 | 9.787 | -15.422 |
| -21.184 | 25.387 | -21.099 | -18.811 | 3.858 | 10.584 | -15.904 | 7.368 | 6.766 | 18.715 | 20.819 | 29.580 |
| -27.279 | -5.368 | 15.673 | 18.433 | -25.782 | 25.531 | 18.399 | 29.386 | 7.719 | 25.646 | 17.758 | 28.655 |
| 22.103 | -16.535 | 28.399 | 25.529 | -18.311 | -8.206 | -16.955 | 4.940 | -11.904 | 8.902 | 24.019 | 27.365 |

| | | | | | | | | | | | |
|---|---|---|---|---|---|---|---|---|---|---|---|
| -11.133 | 12.232 | 7.701 | -5.145 | -14.129 | -32.432 | -10.392 | 17.876 | -15.397 | 10.429 | 22.697 | 10.408 |
| 26.134 | 5.185 | 1.279 | 21.431 | -23.045 | -9.587 | 19.119 | 27.975 | 22.564 | 21.046 | -5.650 | 8.560 |
| -19.249 | -8.007 | 22.504 | 19.559 | -17.553 | 15.257 | -12.510 | 16.466 | 5.871 | 17.550 | -5.896 | 15.197 |
| 26.983 | 4.376 | -26.326 | -12.388 | -3.781 | 8.026 | -8.259 | -1.152 | -3.943 | 29.035 | 28.219 | 25.759 |
| 11.682 | -0.136 | 0.102 | -3.060 | -16.127 | -15.322 | 22.877 | 8.918 | 29.782 | 13.066 | 19.884 | -10.121 |
| 16.075 | 6.647 | 23.306 | -16.367 | 15.864 | -0.260 | 28.678 | 5.532 | 23.783 | 7.346 | 9.198 | 6.396 |
| -5.923 | -6.173 | 3.097 | -1.939 | -2.906 | 14.053 | 0.984 | 19.204 | -11.245 | 28.411 | 15.710 | 7.136 |
| 8.109 | -23.412 | 14.181 | -11.198 | -25.456 | 24.543 | -15.853 | -3.812 | -4.486 | 12.355 | 7.935 | 24.065 |
| 6.699 | -4.736 | 4.245 | -26.402 | 6.251 | 6.766 | 18.425 | 25.012 | 30.817 | 21.483 | 24.737 | 20.884 |
| 24.529 | -20.477 | -5.165 | -19.327 | -11.254 | 26.482 | 5.255 | 12.000 | 20.565 | 13.429 | 21.694 | 18.512 |
| 21.960 | 27.200 | 18.324 | -18.379 | 19.185 | -17.973 | 6.111 | 26.223 | 18.006 | 5.421 | 13.481 | 26.796 |
| 19.851 | -8.687 | -16.376 | -25.080 | -1.555 | 27.285 | 11.141 | 4.178 | 9.615 | 18.864 | 18.581 | -4.576 |
| -10.110 | 6.659 | -10.221 | -19.762 | -0.084 | -3.993 | -22.698 | -1.425 | -0.385 | 20.872 | 26.737 | 23.167 |
| 21.430 | 13.324 | 17.418 | 19.674 | -26.077 | -8.123 | 27.811 | 22.408 | -19.404 | 24.447 | 26.014 | 4.066 |
| 12.474 | 5.727 | -15.356 | -12.638 | 2.236 | -24.577 | 12.825 | 10.399 | 16.909 | 18.601 | -10.904 | 28.947 |
| 25.571 | -29.537 | 26.376 | -22.548 | -19.620 | -1.655 | 4.352 | -0.453 | 7.249 | 29.459 | 23.318 | 24.257 |
| 4.173 | -13.482 | 15.335 | 13.282 | -8.400 | 16.210 | 7.348 | 28.141 | -6.189 | 2.712 | 4.338 | 21.334 |
| -6.605 | -17.774 | -28.388 | -26.164 | -27.388 | 1.002 | -2.008 | 0.674 | -5.888 | -8.759 | 18.137 | 8.950 |
| 16.687 | 11.885 | -0.344 | 23.371 | 6.646 | 23.836 | -3.483 | 27.128 | 14.203 | 29.935 | 27.159 | 23.048 |
| -0.741 | 17.256 | -23.358 | 26.430 | -22.333 | 17.772 | 1.448 | -4.153 | 21.896 | -31.362 | 6.311 | 10.132 |
| -10.708 | 20.330 | 28.696 | 14.337 | -24.906 | 5.452 | 10.712 | 30.037 | -21.292 | 30.886 | 28.518 | 3.391 |
| 7.331 | 25.467 | 10.820 | 5.254 | -5.406 | -1.179 | -18.911 | 19.766 | 11.273 | 6.765 | 21.587 | 22.596 |
| -5.055 | 3.594 | 21.053 | -28.227 | -9.392 | -5.114 | 20.146 | 8.568 | -5.422 | 11.500 | 15.031 | 27.843 |
| -15.938 | 15.050 | -6.098 | -8.386 | -18.841 | -12.051 | -4.697 | -18.586 | -6.959 | 1.685 | 25.443 | 17.438 |
| 15.535 | -8.577 | -6.900 | -11.830 | 28.399 | 14.578 | 14.691 | 22.111 | 19.237 | 26.329 | 22.871 | 24.707 |
| 21.383 | 7.330 | 22.692 | -5.632 | 2.988 | 13.040 | -14.943 | 10.555 | 12.833 | 24.901 | 3.796 | 24.226 |
| -25.846 | -6.025 | -16.550 | 27.385 | 7.936 | -6.456 | 25.282 | 17.809 | 8.590 | 10.326 | 27.788 | 19.047 |
| 19.401 | -30.256 | -4.368 | -14.605 | 5.521 | 25.901 | 21.678 | 29.931 | 17.900 | 23.705 | 19.265 | 26.817 |
| 4.053 | -2.266 | -14.441 | -21.905 | 3.217 | -3.109 | 15.247 | 4.818 | -1.084 | 16.682 | 21.618 | 20.110 |
| -22.986 | 20.431 | -15.669 | -28.224 | -21.500 | -7.358 | -4.001 | 2.021 | 10.178 | -11.878 | 28.839 | 28.795 |
| -14.549 | -13.864 | 5.766 | 13.317 | -16.644 | -3.903 | 7.660 | 30.725 | -3.051 | 8.713 | 9.664 | 4.618 |
| 17.369 | -22.479 | 0.206 | -11.541 | -6.273 | -24.350 | 23.111 | 7.605 | 3.485 | 30.066 | 3.844 | 6.783 |
| 13.094 | 25.376 | -1.934 | -20.073 | -25.053 | 22.609 | -15.358 | 13.975 | 20.124 | 17.469 | 23.897 | 17.927 |
| 4.954 | -27.787 | -13.128 | -29.342 | -6.812 | 20.085 | 14.119 | -2.558 | -9.752 | 14.450 | 22.816 | 11.398 |
| -10.569 | 30.807 | -8.879 | -3.984 | 1.977 | 14.892 | -7.196 | 15.848 | 0.086 | 27.537 | 26.721 | -28.427 |

| | | | | | | | | | | | |
|---|---|---|---|---|---|---|---|---|---|---|---|
| 25.328 | 12.571 | 0.826 | -15.813 | -16.060 | 22.075 | -17.126 | 9.085 | 13.040 | 15.433 | 27.308 | 23.981 |
| 11.233 | -28.372 | -11.094 | -18.238 | -5.142 | -1.705 | -21.642 | 7.146 | -0.703 | 16.138 | 22.872 | 2.174 |
| 19.719 | -6.169 | 6.538 | -19.482 | -21.724 | 28.393 | -0.113 | -0.608 | 20.979 | 5.542 | 18.445 | 14.037 |
| -7.843 | 10.068 | 11.034 | 4.449 | -14.442 | 4.605 | 25.807 | 18.154 | 17.789 | 26.424 | 20.120 | 27.705 |
| -8.402 | 8.053 | -7.665 | -25.151 | 19.739 | -16.790 | -21.284 | -6.854 | 15.909 | 6.484 | -21.949 | 11.050 |
| 8.393 | -17.322 | -22.308 | -24.095 | 21.471 | 7.898 | -5.979 | 11.797 | 4.513 | 13.338 | 25.829 | 17.247 |
| 20.158 | -30.802 | 25.261 | 12.919 | -18.676 | 20.238 | 21.176 | 14.110 | 0.718 | -1.841 | 13.947 | 21.635 |
| 4.553 | -17.763 | 9.139 | 20.709 | -22.482 | -0.784 | 15.847 | -2.212 | 23.980 | 17.835 | 21.417 | 25.475 |
| -17.279 | 5.631 | -13.738 | 0.232 | -27.405 | 23.393 | -17.370 | 6.104 | -12.938 | 29.479 | 17.654 | 8.183 |
| 2.267 | -18.364 | -17.254 | -26.565 | 16.300 | -3.598 | 23.198 | -1.120 | 7.425 | 15.825 | 27.544 | 15.768 |
| -19.742 | -9.332 | 20.281 | -25.367 | -9.914 | -6.709 | -13.224 | -2.814 | -6.330 | 28.349 | 22.016 | -14.148 |
| 16.199 | -27.609 | 14.704 | 9.306 | -6.743 | -21.914 | -1.758 | 10.803 | 24.784 | -5.059 | 5.483 | -0.183 |
| 8.558 | 3.723 | -18.995 | -27.412 | 7.757 | 18.668 | 20.000 | 10.138 | -1.634 | 22.772 | 5.128 | 23.482 |
| -21.765 | -25.679 | 18.596 | -23.017 | -16.876 | 21.006 | -22.277 | 7.848 | 25.458 | 21.605 | 20.599 | 6.596 |
| 23.570 | -4.670 | -21.120 | -21.946 | 15.376 | 23.285 | 17.751 | 0.684 | -3.535 | 10.444 | 14.668 | 20.444 |
| 12.048 | 16.969 | 8.210 | 18.034 | 0.658 | -5.786 | 15.482 | -4.753 | 11.594 | 23.730 | 19.183 | 3.817 |
| -3.549 | -27.471 | 10.687 | -3.275 | -8.451 | -5.259 | -0.639 | 13.224 | -16.280 | 18.825 | 18.742 | 18.737 |
| -13.873 | 20.479 | -7.480 | -9.672 | -23.163 | 13.495 | -14.038 | -22.546 | 26.608 | 12.025 | 21.168 | 22.327 |
| -11.266 | -12.126 | 0.761 | -15.458 | 25.786 | -19.209 | 30.235 | 2.499 | 23.426 | 18.209 | 0.284 | 26.039 |
| -8.175 | 12.115 | -25.515 | -2.523 | -14.550 | 16.483 | 26.190 | 12.157 | 20.930 | 9.544 | 16.023 | 8.064 |
| 13.995 | 19.865 | -4.212 | 21.824 | -4.729 | -27.823 | 22.687 | -1.989 | 8.189 | 29.392 | 16.947 | 21.917 |
| -0.380 | -4.034 | 18.217 | -2.130 | -9.838 | 7.146 | -21.687 | 4.626 | 28.812 | 25.330 | 9.797 | 24.888 |
| 1.856 | 24.223 | -15.192 | -0.554 | 2.652 | 23.809 | -25.599 | -18.972 | -1.599 | 13.629 | -3.317 | -14.152 |
| 0.552 | 7.420 | 16.119 | -29.809 | 13.511 | 6.280 | 24.856 | 7.236 | -19.060 | 21.954 | 22.440 | 20.114 |
| 11.781 | -13.783 | -3.439 | -20.442 | 4.771 | 3.937 | 10.151 | 8.519 | 4.254 | 11.077 | -10.054 | -14.177 |
| -1.105 | -23.381 | -10.199 | -27.320 | -6.553 | -18.735 | -16.286 | 1.317 | 8.073 | 1.381 | 11.090 | 22.563 |
| -24.911 | 30.842 | 25.348 | -5.476 | -22.850 | 8.085 | 23.734 | 15.126 | 18.096 | -0.928 | -7.662 | 11.916 |
| -8.878 | -27.924 | -18.762 | -13.923 | -20.456 | -3.783 | 27.119 | 9.598 | 19.682 | -6.827 | 13.345 | 23.474 |
| -25.786 | 20.784 | -20.787 | 27.500 | -14.342 | 9.330 | 14.444 | -17.418 | -0.932 | 22.798 | 5.691 | 23.904 |
| -10.228 | 1.202 | -20.284 | 1.301 | -10.887 | 5.030 | -18.414 | 16.055 | 6.530 | 4.807 | 27.726 | 20.252 |
| -24.451 | -20.293 | 13.063 | 8.330 | -11.541 | -27.738 | 12.707 | -3.937 | 21.822 | -17.429 | -3.707 | -7.720 |
| 21.441 | 6.240 | 15.160 | 9.044 | -27.068 | 25.654 | 4.291 | -1.207 | -8.416 | 6.506 | -4.993 | 21.223 |
| 20.971 | 12.290 | -23.445 | 20.927 | 28.534 | -12.858 | 28.796 | 1.947 | -28.285 | 27.854 | 20.396 | 0.584 |
| -17.388 | -19.778 | -7.044 | -3.617 | -28.804 | 22.134 | -14.355 | 15.763 | 22.654 | 15.845 | -15.667 | 24.674 |
| 12.376 | -6.020 | 7.839 | -26.024 | -25.862 | 10.767 | 11.385 | 5.088 | 15.590 | 14.895 | 19.833 | 5.831 |

| | | | | | | | | | | | |
|---|---|---|---|---|---|---|---|---|---|---|---|
| -15.412 | -11.094 | 17.029 | 18.060 | 23.717 | 23.133 | -7.240 | -3.361 | 4.763 | 8.611 | 10.040 | 16.440 |
| -2.638 | -27.017 | 17.293 | -9.290 | -17.252 | 13.624 | 6.696 | 11.847 | -22.553 | 26.510 | 26.931 | 7.362 |
| -26.513 | -14.485 | -1.556 | -4.703 | -15.240 | 14.540 | -14.464 | 25.061 | -27.763 | 18.758 | 18.444 | 6.845 |
| 11.168 | -5.339 | 18.819 | -29.896 | -19.603 | -1.464 | -11.709 | -14.391 | -0.504 | 0.627 | 16.648 | 2.454 |
| -6.490 | 11.963 | 15.834 | -19.867 | 5.153 | -8.312 | 25.436 | 26.519 | -20.557 | 20.833 | 25.638 | 16.993 |
| -7.125 | 25.126 | 7.354 | -26.610 | -24.543 | -6.631 | -9.637 | -2.563 | 10.541 | 12.454 | 9.763 | 19.008 |
| 9.964 | 8.294 | -18.235 | 7.039 | -18.778 | -15.032 | -17.973 | 6.709 | 16.486 | 23.919 | 26.461 | 5.510 |
| 19.409 | -7.851 | 0.832 | -8.043 | 25.980 | 9.549 | -17.068 | 1.459 | 9.678 | 14.654 | 20.586 | 16.051 |
| 8.682 | -21.243 | -20.906 | 3.497 | -20.508 | 20.305 | -14.027 | 11.002 | -3.131 | 26.477 | 18.955 | 22.192 |
| 18.889 | -24.127 | -24.154 | -8.594 | -4.099 | -4.000 | -23.562 | 2.376 | 17.862 | 13.802 | -4.797 | 17.519 |
| -26.762 | -24.697 | 6.506 | -27.451 | -29.982 | -12.586 | -17.496 | -1.538 | 16.701 | -0.745 | 24.086 | -5.192 |
| 24.642 | 7.276 | -14.682 | 18.533 | 11.669 | 0.677 | -16.065 | 12.363 | -9.968 | 6.474 | 21.064 | 1.046 |
| 23.429 | -2.658 | -2.686 | 18.258 | -24.470 | 24.907 | -2.641 | 24.763 | -22.205 | -2.155 | -3.423 | 11.829 |
| 5.699 | 1.673 | 18.118 | -10.471 | 20.087 | -2.719 | 21.654 | 3.466 | 0.964 | 3.639 | 4.519 | 17.246 |
| -5.200 | -11.031 | 10.346 | -13.529 | 2.216 | -11.969 | 29.604 | -10.239 | 5.127 | 24.831 | -8.438 | 24.753 |
| -10.529 | 12.281 | 3.664 | 5.648 | 7.848 | -4.956 | 13.626 | -22.420 | 11.881 | 25.829 | 3.725 | 14.397 |
| -12.453 | 3.928 | -11.900 | -15.967 | -18.847 | 18.873 | -7.590 | 27.437 | 11.540 | -4.236 | 16.463 | 20.053 |
| 15.328 | 7.363 | 5.622 | -21.502 | -20.753 | 16.315 | -23.216 | -8.099 | 3.747 | 12.214 | 24.665 | -3.266 |
| 9.852 | 12.249 | 0.362 | -7.046 | 7.331 | -3.095 | -18.885 | 25.364 | -0.589 | 24.345 | 14.489 | 17.214 |
| 26.457 | -6.878 | 15.296 | -30.468 | 2.773 | 12.589 | 10.130 | 8.888 | 27.430 | 17.684 | 21.335 | -8.763 |
| 27.673 | -7.881 | 3.293 | 23.639 | -1.746 | -15.945 | -10.120 | 5.223 | -7.725 | 16.465 | 26.691 | 22.550 |
| 8.538 | 27.385 | -0.670 | -23.958 | 31.300 | -18.028 | -25.942 | 20.130 | -11.501 | 28.405 | -1.286 | 15.931 |
| -26.561 | -20.672 | 13.530 | 19.008 | -7.004 | -2.394 | -19.435 | -0.488 | 25.455 | 14.120 | 19.755 | 10.275 |
| 14.922 | 8.642 | -8.922 | -12.205 | 30.621 | 11.441 | -21.550 | 24.829 | -10.170 | 18.908 | 8.240 | 14.531 |
| 19.593 | 14.804 | -9.385 | -9.709 | -15.357 | 4.124 | 24.602 | -8.590 | 20.895 | 0.796 | -12.518 | -1.730 |
| -19.964 | -22.983 | 9.971 | -5.208 | -10.321 | -15.258 | 11.459 | 4.698 | 10.124 | 21.701 | 20.721 | -13.312 |
| 1.819 | -18.211 | -21.354 | 7.905 | -25.424 | -1.623 | -11.693 | 16.161 | -12.493 | 5.741 | 5.926 | -6.244 |
| -19.595 | -5.215 | 2.224 | -25.265 | 25.085 | 10.389 | 29.848 | -2.427 | 10.855 | 20.067 | 22.916 | 18.540 |
| -22.642 | -23.530 | 16.331 | -3.556 | 15.930 | -19.543 | 29.042 | -10.622 | 19.196 | 4.901 | 19.276 | 13.808 |
| 9.397 | -28.944 | -1.256 | -0.613 | -8.999 | -14.450 | 16.493 | 7.369 | 19.953 | -0.893 | 9.654 | 23.436 |
| -12.386 | -15.983 | 5.012 | 7.163 | -9.744 | 10.667 | -6.531 | 25.491 | 11.841 | 26.822 | 25.297 | 25.303 |
| -5.510 | 5.108 | -4.936 | -9.147 | 14.129 | -23.091 | 13.113 | -16.502 | 0.161 | 16.102 | 16.456 | 26.330 |
| 2.195 | 20.991 | 0.570 | 6.375 | -28.180 | -3.195 | -16.489 | 13.711 | 9.341 | -0.625 | 6.183 | -22.461 |
| 18.550 | -8.468 | 21.885 | -14.895 | -9.175 | 14.700 | -17.243 | 8.705 | 9.892 | -3.719 | 8.703 | 16.867 |
| 10.011 | -5.128 | -3.430 | 11.056 | -10.822 | -14.136 | -19.772 | 7.878 | 4.856 | 18.394 | 25.672 | -1.458 |

| | | | | | | | | | | | |
|---|---|---|---|---|---|---|---|---|---|---|---|
| -18.706 | 15.183 | 8.359 | -14.312 | 31.472 | 7.782 | 5.782 | 3.763 | 10.861 | 17.365 | 20.297 | -4.830 |
| -20.486 | -5.845 | 12.487 | -7.736 | 14.018 | 14.070 | 23.573 | 24.314 | 2.554 | 5.303 | 19.092 | -8.731 |
| 23.338 | 9.799 | 12.882 | 4.352 | -8.318 | 22.827 | -17.293 | 17.304 | 26.992 | -1.460 | 13.195 | -3.260 |
| 21.239 | -27.907 | -2.754 | 27.303 | 28.911 | 18.950 | -12.138 | -10.909 | -8.567 | 4.640 | 24.848 | 10.746 |
| -14.206 | -6.781 | 9.927 | 23.432 | -5.611 | 18.252 | -0.696 | -6.493 | 17.440 | 19.148 | 22.322 | -7.698 |
| -6.981 | 20.674 | 10.541 | -27.400 | -23.410 | 3.294 | -20.304 | 2.815 | 23.671 | 15.008 | 21.049 | 24.592 |
| 13.844 | -4.115 | 18.113 | 2.323 | -20.222 | -3.582 | 19.992 | 4.298 | 11.426 | 23.140 | -2.277 | 11.348 |
| -3.158 | 12.911 | 14.154 | -15.876 | -12.691 | -6.263 | 23.211 | 16.106 | 4.158 | 0.731 | 12.097 | 17.980 |
| -11.839 | 23.085 | -16.320 | 21.875 | 31.003 | -17.718 | -29.346 | 15.273 | -2.157 | 9.037 | 18.743 | 3.931 |
| -1.118 | 23.244 | -5.649 | -4.699 | 18.919 | 5.398 | 17.154 | 5.578 | 19.973 | 23.771 | -10.078 | 18.656 |
| 3.904 | 9.508 | -11.310 | -8.211 | -6.744 | -2.594 | 3.745 | 20.194 | -22.346 | -0.773 | 18.527 | 12.409 |
| 0.762 | -16.209 | 4.882 | -25.048 | 5.441 | -24.168 | 20.526 | 21.500 | 24.017 | 19.800 | -0.130 | -25.012 |
| 22.417 | 20.864 | 5.854 | 7.686 | -19.342 | -9.319 | -13.848 | 13.190 | 15.394 | 14.201 | -6.004 | 22.634 |
| 17.358 | -25.039 | 1.119 | -20.613 | -5.172 | -17.140 | 29.628 | 17.440 | 22.822 | 3.417 | 23.304 | 5.546 |
| -24.099 | -20.354 | 0.105 | -23.261 | 18.995 | -7.871 | 6.451 | 3.406 | -17.473 | 20.470 | -7.587 | -0.843 |
| -19.635 | -12.297 | -13.478 | -11.804 | 4.213 | -18.673 | -23.631 | -12.025 | 21.121 | 3.950 | 13.823 | 26.040 |
| -17.085 | -4.931 | 24.458 | -20.279 | -13.785 | -13.463 | 21.611 | 14.575 | 18.708 | 27.151 | 20.888 | 27.605 |
| 6.057 | -8.780 | -24.982 | 0.891 | -22.270 | -26.421 | 18.054 | 3.014 | 5.887 | 6.006 | 14.828 | 5.336 |
| 14.265 | -6.685 | -7.461 | 15.646 | -24.145 | 6.323 | 15.305 | 13.653 | 15.299 | 12.775 | 17.722 | 19.073 |
| 11.393 | 6.815 | 21.136 | 4.784 | -4.774 | 7.476 | 26.984 | 9.218 | 20.212 | 14.512 | 15.143 | 14.932 |
| 3.302 | 4.646 | 17.760 | -5.563 | -20.441 | -5.039 | 22.479 | 20.505 | 5.080 | 15.903 | 11.027 | 20.197 |
| 19.266 | -21.439 | 11.255 | 9.422 | 13.236 | 12.871 | -10.946 | 12.499 | 18.782 | 18.269 | 1.312 | 23.919 |
| 29.831 | 5.793 | 19.097 | -10.656 | 4.546 | -23.887 | -21.912 | 23.324 | -23.552 | 26.195 | 10.398 | 17.883 |
| -12.023 | 7.322 | -15.588 | -22.715 | 19.390 | -9.403 | 27.958 | -2.129 | 9.686 | 9.008 | 0.309 | 16.715 |
| 9.998 | -12.131 | 9.300 | 11.992 | -15.433 | -8.518 | 16.051 | -6.975 | -18.843 | 6.651 | 11.221 | 21.142 |
| 20.160 | -15.972 | -5.768 | 2.392 | 30.556 | 3.806 | 19.923 | -3.923 | 19.544 | -1.399 | 14.378 | 25.933 |
| -11.688 | -0.993 | -1.106 | -5.185 | -22.941 | 2.479 | -26.247 | 24.653 | -6.893 | 24.394 | 22.110 | 27.788 |
| 1.285 | 24.297 | 19.220 | -6.801 | 11.302 | -1.423 | 25.469 | 3.490 | -2.124 | 10.725 | 23.637 | 27.192 |
| 13.033 | 22.490 | 17.403 | -3.828 | 7.843 | 2.012 | 22.838 | -5.819 | 11.249 | 5.100 | -0.871 | 28.286 |
| -5.253 | -9.160 | -15.283 | -9.148 | -17.003 | 20.492 | -19.652 | 14.276 | 10.411 | 3.794 | 15.789 | 29.462 |
| 9.648 | 21.238 | 2.266 | -1.835 | -18.743 | -3.961 | -28.093 | 8.467 | 9.868 | 16.761 | -5.530 | 21.748 |
| 25.914 | 12.111 | -13.979 | -23.566 | -29.400 | -11.845 | -4.627 | 2.134 | -18.304 | 8.220 | 8.525 | 7.218 |
| -25.180 | -4.297 | 12.507 | 17.278 | -21.982 | 9.585 | 5.518 | -0.790 | 2.208 | 19.476 | 24.944 | 13.150 |
| 25.288 | 10.793 | -19.949 | 8.911 | -22.843 | -10.748 | -23.340 | -15.792 | 21.705 | 21.758 | 10.881 | 28.152 |
| 22.164 | -8.677 | 19.610 | 5.886 | -27.629 | 22.365 | 27.838 | -11.316 | 25.335 | 0.717 | 26.390 | 27.197 |

| | | | | | | | | | | | |
|---|---|---|---|---|---|---|---|---|---|---|---|
| -9.053 | -13.676 | 10.720 | -18.009 | -17.478 | 1.404 | -11.740 | 18.571 | 19.911 | 19.000 | 1.876 | 17.570 |
| 30.384 | 16.052 | -23.246 | 2.095 | 3.904 | -2.146 | 26.832 | -13.933 | -15.897 | 14.108 | 25.558 | 22.425 |
| 27.260 | -25.209 | 7.895 | 11.970 | -4.122 | -7.707 | -12.680 | 17.409 | 15.238 | 9.717 | 23.140 | 24.472 |
| -12.453 | 6.583 | 15.983 | 0.203 | -18.103 | 22.650 | -11.047 | 19.823 | 9.071 | 15.932 | 16.513 | 25.971 |
| 3.770 | 19.657 | -1.967 | -10.293 | -0.329 | 2.175 | 6.862 | -6.767 | -8.466 | 26.180 | -23.350 | 5.904 |
| 28.200 | 9.113 | -17.565 | -6.311 | -28.109 | -16.565 | 17.169 | 4.660 | 11.728 | 5.618 | 16.491 | 26.786 |
| 10.511 | -11.832 | 3.649 | -8.603 | -19.179 | -24.392 | 26.074 | 18.180 | -16.928 | 16.518 | 11.021 | 23.732 |
| -4.880 | -21.514 | 4.095 | -7.742 | 28.036 | -14.467 | -26.642 | 1.158 | 22.745 | 22.836 | 19.717 | 24.864 |
| 18.743 | 21.510 | 17.847 | -14.834 | -9.304 | 25.898 | -20.851 | -0.135 | 9.999 | 17.364 | 20.751 | -3.903 |
| 21.088 | -18.474 | 8.418 | 2.800 | 29.804 | 13.645 | -7.813 | 11.087 | 15.496 | 6.440 | 15.455 | 14.260 |
| 23.059 | -21.623 | -13.878 | -16.477 | 19.374 | 8.622 | -9.952 | 14.051 | 2.167 | -0.255 | 21.640 | 7.220 |
| 20.341 | 1.574 | -9.579 | -13.197 | 27.699 | 2.501 | -27.479 | 20.383 | -25.003 | -14.419 | 19.363 | 8.761 |
| 18.218 | -0.495 | 0.218 | -2.759 | -6.819 | 10.206 | -8.454 | -9.101 | 17.847 | 23.287 | -5.737 | 19.933 |
| -23.242 | 19.643 | -12.996 | -28.966 | -14.416 | -5.765 | -28.832 | 21.907 | 24.213 | -10.523 | 9.510 | 0.164 |
| -22.431 | 16.550 | -7.366 | -4.436 | -7.976 | 15.586 | -15.790 | 23.942 | 7.401 | 0.917 | 14.273 | 29.696 |
| -10.992 | -24.924 | 11.389 | -9.783 | -6.029 | -14.976 | 22.744 | 19.134 | -4.429 | 22.110 | -6.033 | 5.408 |
| 7.324 | 8.202 | -11.147 | -15.927 | -17.242 | 1.356 | 10.563 | 23.656 | 24.823 | 12.007 | 15.653 | 30.225 |
| 25.996 | 9.680 | 19.232 | 9.917 | -23.465 | 14.317 | -6.476 | 3.930 | 4.807 | 20.331 | -3.453 | -8.752 |
| -3.755 | 6.411 | -25.127 | 6.958 | 8.165 | 21.701 | -19.327 | 16.650 | 2.110 | 7.037 | 18.578 | 16.062 |
| 11.007 | 11.202 | -5.187 | 6.003 | -24.367 | 16.657 | 2.009 | 19.880 | 15.808 | -2.173 | 21.002 | 25.323 |
| 22.415 | -8.566 | -17.583 | 13.665 | -19.804 | 22.184 | -7.785 | 8.451 | 6.517 | 24.550 | 17.460 | 23.615 |
| 0.717 | 19.349 | -11.513 | -23.072 | -20.972 | 0.611 | 20.525 | -6.000 | -27.057 | -2.792 | -2.325 | 12.955 |
| -5.511 | 1.795 | 24.633 | -27.846 | 9.413 | -16.428 | -25.508 | 6.873 | 23.265 | 14.224 | 28.487 | 21.413 |
| 21.592 | -7.318 | 0.943 | -14.704 | -1.491 | 26.698 | -19.027 | 9.370 | 10.122 | -0.281 | 21.757 | -1.561 |
| -28.017 | -14.763 | -2.517 | -20.103 | -15.847 | 1.945 | -21.125 | 13.364 | 20.503 | 12.836 | 12.456 | 9.437 |
| -26.870 | 15.275 | 25.065 | -9.034 | 21.258 | 2.519 | -1.919 | 0.992 | -7.879 | 20.487 | -1.930 | 22.687 |
| -1.460 | 9.570 | -14.841 | 7.879 | -7.655 | 3.188 | -11.556 | 17.279 | -15.798 | -21.337 | 22.575 | 11.616 |
| 17.780 | 22.146 | 8.320 | 3.739 | -23.904 | -5.144 | -16.449 | 8.236 | -18.935 | 20.975 | -2.218 | 16.078 |
| 15.403 | -11.236 | 16.218 | 9.328 | 22.380 | -19.207 | 23.996 | 19.055 | 26.531 | 12.142 | 23.821 | -11.853 |
| 12.424 | -20.298 | -4.549 | 22.355 | -19.676 | 18.501 | -21.035 | -17.556 | 7.853 | 9.362 | 19.594 | 29.653 |
| -15.041 | -3.806 | 25.296 | 11.705 | -5.050 | -26.972 | -21.909 | 11.310 | -7.017 | 8.500 | 4.897 | 20.013 |
| 1.417 | -9.739 | -12.056 | -24.624 | -21.933 | 9.576 | 15.324 | 9.122 | 23.986 | -5.757 | -23.286 | 15.014 |
| 28.344 | -12.331 | 3.583 | 14.337 | -4.734 | -18.274 | -19.456 | 23.571 | 22.188 | 1.463 | 11.037 | 18.786 |
| 16.194 | 11.609 | -24.619 | -0.601 | -23.212 | 6.317 | 14.588 | 12.038 | 5.829 | 16.503 | -1.878 | -3.466 |
| 7.899 | -2.485 | 11.873 | -7.394 | -14.600 | 23.496 | 28.586 | 18.043 | -9.804 | 14.486 | 24.928 | 9.000 |

| | | | | | | | | | | | |
|---|---|---|---|---|---|---|---|---|---|---|---|
| 9.702 | 19.376 | -1.927 | -10.126 | 6.773 | 3.078 | 3.340 | 7.078 | 27.315 | 10.849 | 12.682 | 6.282 |
| 13.775 | 6.828 | -17.211 | -6.694 | -13.090 | -5.927 | -22.224 | 1.958 | -11.153 | -12.424 | 15.817 | 22.213 |
| -20.683 | -7.810 | -25.044 | -1.947 | -9.199 | 27.158 | -26.716 | -1.030 | -3.006 | 24.931 | 19.309 | 28.577 |
| -8.235 | 8.845 | 13.919 | 8.063 | -15.738 | 10.582 | 0.049 | 24.825 | -14.334 | 9.796 | 11.560 | 24.777 |
| -4.854 | -11.952 | -23.155 | 17.927 | -2.714 | 30.207 | -27.090 | 13.279 | -0.114 | 22.910 | 28.072 | 17.282 |
| -5.799 | 21.866 | 9.405 | -2.822 | -26.323 | 25.776 | 26.778 | 15.182 | 22.706 | 18.474 | -8.324 | 30.215 |
| -9.068 | -17.947 | 4.171 | -15.009 | -18.210 | -13.844 | -11.967 | -8.196 | -13.880 | 21.983 | 25.469 | 12.233 |
| 14.908 | 4.651 | 14.475 | -22.320 | -4.965 | -6.436 | -11.698 | 20.137 | -5.476 | 17.060 | 8.011 | 1.711 |
| 26.008 | -6.215 | 8.487 | 25.358 | -11.806 | -11.728 | -23.175 | 7.995 | -21.937 | 7.547 | 22.803 | 9.617 |
| -23.347 | -1.098 | -3.155 | 6.545 | 7.856 | -24.701 | 16.878 | 11.731 | 27.827 | 25.588 | 11.980 | 28.290 |
| 7.058 | -14.517 | 24.523 | -3.909 | 3.648 | 28.050 | 0.620 | 27.384 | 9.788 | 1.504 | 28.730 | 11.554 |
| 14.013 | -4.868 | 2.712 | 2.163 | -24.797 | 20.384 | -12.087 | 9.849 | 12.763 | 0.464 | 14.640 | 10.941 |
| -1.127 | 26.597 | 19.909 | 11.189 | -3.683 | 18.819 | -16.962 | 2.568 | 11.399 | 19.660 | 25.293 | 13.854 |
| -6.141 | 12.948 | 27.252 | -13.442 | -17.011 | 15.668 | -8.648 | -3.551 | 23.509 | 23.841 | 24.084 | 12.433 |
| 9.389 | 5.855 | -22.114 | 4.903 | -2.002 | 15.046 | -4.585 | 13.779 | -3.569 | -16.599 | 19.463 | 9.295 |
| 13.436 | 4.907 | -19.708 | -21.136 | 24.425 | 3.229 | 0.729 | 25.235 | -11.603 | 16.216 | 14.103 | 0.401 |
| 20.222 | 16.603 | 15.542 | -22.810 | -8.374 | 20.593 | 11.991 | 5.188 | 20.497 | 4.101 | 17.327 | 18.223 |
| 4.419 | 0.835 | -23.353 | -9.111 | 7.205 | 21.546 | -4.212 | -8.261 | -4.724 | 13.061 | 15.599 | 23.309 |
| 22.681 | -27.820 | 4.890 | -10.572 | -18.893 | 25.602 | 21.339 | 18.168 | -15.932 | 22.920 | 9.941 | 20.052 |
| 25.852 | -21.035 | 3.902 | -15.651 | -0.259 | 23.625 | 2.478 | 15.856 | 21.648 | 25.272 | -5.174 | 23.751 |
| 12.548 | -20.429 | 8.344 | -4.589 | -9.369 | -8.993 | -11.001 | 0.771 | 19.240 | 7.122 | 23.205 | 29.585 |
| 2.889 | 27.986 | 19.790 | 0.216 | -6.870 | -3.265 | 13.700 | 16.944 | 17.455 | 3.720 | 29.628 | 10.255 |
| -11.627 | 11.386 | 17.059 | 11.844 | 12.151 | 4.885 | -24.283 | 14.053 | 7.798 | 2.069 | 28.748 | 8.151 |
| 4.862 | -9.770 | 21.604 | -3.839 | 16.685 | 28.505 | 16.349 | -3.119 | -14.630 | -16.354 | 5.254 | 22.546 |
| 19.951 | -15.836 | -6.511 | -17.329 | -17.184 | -7.738 | -14.637 | 19.419 | 25.192 | 2.770 | 10.490 | 15.652 |
| 7.937 | 0.036 | 24.985 | -7.982 | 19.450 | 21.212 | -5.648 | 15.192 | 21.661 | 18.497 | 21.786 | -0.952 |
| -13.912 | 28.306 | 3.380 | 1.068 | -20.169 | 26.462 | -21.783 | 10.731 | -13.231 | 5.590 | 25.177 | 11.819 |
| 25.391 | 19.123 | 23.186 | -9.493 | 18.867 | 8.083 | -23.291 | -2.148 | 8.824 | 16.743 | -4.907 | -5.567 |
| -3.218 | 16.838 | -8.613 | 27.539 | -11.392 | 24.521 | -12.329 | 24.709 | -7.955 | 9.269 | 19.005 | -12.075 |
| 3.903 | -4.135 | -18.073 | -5.054 | 26.880 | -12.066 | -14.636 | 12.506 | 26.969 | 23.999 | 17.100 | 18.447 |
| 9.021 | -21.927 | -26.573 | 7.099 | -18.322 | 25.231 | 18.285 | 20.376 | 6.553 | 13.980 | 3.036 | 28.513 |
| -10.200 | -11.745 | 11.003 | 18.173 | -8.596 | -22.110 | 22.981 | 28.029 | 17.866 | 1.339 | 3.904 | 21.296 |
| -19.893 | 3.868 | 3.827 | -18.407 | -12.580 | 20.294 | 20.166 | 6.694 | 10.560 | 9.628 | 15.530 | -25.461 |
| 22.762 | -20.725 | -16.318 | 9.809 | 16.127 | -27.452 | 24.625 | 18.514 | -9.813 | 25.672 | 11.211 | -1.662 |
| 1.284 | 27.447 | 16.201 | 4.178 | 6.926 | 28.979 | -5.387 | 4.759 | -15.626 | 10.614 | 28.214 | 19.841 |

| | | | | | | | | | | | |
|---|---|---|---|---|---|---|---|---|---|---|---|
| -1.741 | -19.859 | 7.760 | 4.717 | -2.107 | -13.349 | 1.325 | -0.898 | 14.311 | 5.320 | -16.077 | 23.644 |
| 7.239 | 4.720 | 5.875 | 11.962 | 0.496 | -17.771 | -10.170 | 26.834 | 22.584 | 25.268 | 20.178 | 1.849 |
| -17.602 | -19.715 | -6.566 | 13.510 | -8.275 | 20.301 | 19.962 | 3.845 | -22.907 | 21.635 | 27.306 | 16.469 |
| 4.677 | 3.024 | -22.066 | 20.047 | -27.025 | 22.828 | 5.588 | 23.158 | -1.381 | 17.434 | 13.660 | 30.190 |
| -0.877 | 19.296 | -27.484 | -3.536 | -25.889 | 26.716 | -4.152 | 0.268 | 5.792 | 14.649 | 0.895 | 24.913 |
| 25.067 | 28.835 | -20.642 | 16.898 | -20.753 | 19.153 | -8.864 | 15.159 | 16.464 | 3.121 | 14.564 | 7.496 |
| -12.617 | -9.835 | 19.018 | 21.327 | 2.122 | -18.368 | 12.545 | 26.589 | 4.787 | 6.700 | 26.742 | 16.844 |
| -23.475 | 9.706 | 12.074 | 2.334 | -6.042 | 23.663 | 1.256 | 10.772 | -11.497 | 2.339 | 21.945 | 10.735 |
| 13.423 | -5.964 | 26.118 | 8.535 | -15.333 | -17.612 | -4.341 | 8.718 | 13.980 | -7.843 | 9.067 | -5.247 |
| -7.288 | -20.775 | 17.195 | 0.711 | -0.620 | -4.930 | 5.287 | 6.860 | 22.171 | 15.412 | 29.474 | 19.647 |
| -8.852 | 6.719 | -2.269 | -15.705 | 3.031 | -15.123 | 24.516 | 18.421 | 27.182 | 7.562 | 13.495 | 6.822 |
| 22.857 | -19.586 | -24.656 | 13.989 | -11.440 | -20.881 | 7.664 | 27.199 | 9.859 | 11.375 | 12.580 | -24.789 |
| 4.903 | -15.227 | -19.514 | -1.189 | -18.584 | 26.451 | 23.709 | 19.206 | 21.279 | 19.806 | 13.313 | 25.110 |
| -10.974 | 25.949 | -28.735 | -11.345 | 11.558 | -8.900 | -20.867 | 10.011 | 13.156 | 24.732 | 22.478 | 15.638 |
| -2.510 | 28.515 | -12.538 | -6.794 | -14.006 | 22.548 | -6.050 | -2.094 | -5.723 | 20.978 | 11.109 | 22.829 |
| 15.244 | 23.749 | 9.783 | -19.269 | -22.587 | 26.926 | -12.428 | 20.405 | -4.410 | 16.136 | 19.384 | 24.620 |
| -6.476 | 1.635 | 6.038 | -9.696 | 7.999 | 15.760 | -8.878 | 27.956 | -14.481 | 22.031 | 27.794 | 5.980 |
| -17.901 | 24.808 | 24.863 | -24.403 | -13.611 | -17.022 | 11.951 | 11.731 | 19.113 | -10.707 | 27.143 | 7.293 |
| -1.473 | -10.043 | -0.106 | 11.356 | -20.247 | 27.776 | 2.941 | -2.398 | -7.643 | 10.713 | 17.472 | 27.650 |
| -0.758 | 8.095 | -21.307 | 0.231 | 0.561 | -25.045 | 17.250 | 7.414 | 16.182 | 28.142 | 24.993 | -0.762 |
| -5.280 | 27.564 | 4.257 | -9.106 | -8.737 | 20.956 | 9.332 | 1.941 | -15.853 | 6.552 | 23.452 | 5.140 |
| 7.422 | -17.240 | -7.222 | -5.257 | -3.422 | 17.342 | -17.975 | 17.024 | 10.600 | 25.856 | 9.470 | -4.699 |
| -0.039 | 28.452 | 13.179 | -11.607 | -5.311 | 12.866 | 1.690 | 22.946 | 22.919 | 7.036 | 17.885 | 25.975 |
| -4.326 | 7.722 | 25.623 | -3.005 | -2.863 | -3.439 | -19.848 | 9.075 | -22.986 | 18.570 | 16.972 | 23.537 |
| -8.333 | -20.592 | 20.409 | 12.045 | -25.966 | -0.423 | -3.649 | 2.598 | 15.209 | 17.239 | 4.635 | 9.525 |
| -3.303 | 29.538 | 18.415 | 1.611 | 14.562 | -13.999 | 6.607 | -7.316 | 28.738 | 21.384 | 16.478 | 21.192 |
| 8.029 | -2.271 | 12.395 | 6.248 | -9.509 | 20.583 | -0.311 | -16.641 | 27.072 | 11.679 | 28.423 | 26.719 |
| -2.294 | -30.756 | -15.269 | -8.031 | -17.157 | -29.589 | -6.853 | 6.558 | -18.032 | 10.688 | 11.416 | -15.297 |
| 12.318 | 30.591 | -11.730 | -7.400 | -6.515 | 13.793 | -4.894 | 13.358 | -4.418 | 25.313 | 24.264 | 21.410 |
| -2.809 | -14.560 | -2.847 | -21.691 | -10.433 | 25.571 | 23.739 | 8.300 | 2.032 | 16.693 | 7.575 | 17.674 |
| -9.525 | 20.441 | -12.750 | -23.313 | 1.333 | 27.126 | 10.429 | 0.681 | -13.063 | -17.460 | 21.592 | -22.222 |
| 11.490 | -18.287 | -13.280 | 13.427 | -14.136 | 26.643 | -2.236 | -1.789 | 11.736 | 12.961 | 15.262 | 24.067 |
| 11.986 | -9.036 | -29.349 | -14.474 | -18.849 | -28.565 | -19.250 | 5.802 | 6.765 | -5.488 | 22.074 | 12.969 |
| -3.235 | 0.408 | -6.821 | 21.473 | -1.490 | 16.265 | 10.803 | 12.183 | -14.288 | 18.384 | 23.749 | 10.989 |
| -17.616 | -21.126 | -10.550 | -11.268 | -5.483 | 26.872 | -17.177 | 24.093 | 29.133 | 0.725 | 2.300 | 24.575 |

| | | | | | | | | | | | |
|---|---|---|---|---|---|---|---|---|---|---|---|
| 3.337 | -22.883 | 24.635 | -12.031 | -9.784 | 24.470 | -5.915 | 23.308 | -12.272 | 26.459 | 19.545 | 19.383 |
| 3.711 | -26.413 | -16.286 | -25.641 | -23.550 | 17.376 | 17.863 | 9.620 | 13.030 | 13.896 | 0.041 | 14.396 |
| -23.869 | 22.279 | 21.357 | 10.773 | -14.870 | 1.805 | -19.803 | 20.594 | -3.498 | -2.201 | 28.449 | 5.865 |
| 4.047 | -31.720 | 28.085 | 26.045 | 0.089 | -4.284 | -21.962 | 14.865 | 4.181 | 21.699 | -5.814 | 9.842 |
| -19.404 | 4.438 | -25.305 | 9.453 | 2.361 | 20.162 | -16.251 | -0.060 | 18.693 | 27.464 | 17.161 | 16.148 |
| -7.274 | 18.763 | 23.542 | 16.151 | 1.572 | -6.172 | 18.654 | 18.558 | -5.202 | -0.604 | 6.924 | 19.860 |
| -11.063 | -29.427 | -21.856 | -20.262 | 2.921 | 25.612 | 14.907 | 22.354 | 26.640 | 19.174 | 12.763 | 18.201 |
| 17.474 | -16.588 | -9.307 | -20.795 | 1.858 | 11.336 | 12.510 | -5.073 | 27.216 | 25.911 | 26.947 | 26.351 |
| 3.057 | 12.753 | -20.411 | -7.095 | 11.773 | 19.031 | -7.299 | 10.311 | -24.447 | 16.106 | 23.021 | 25.245 |
| -25.621 | -17.523 | 23.919 | 2.798 | -19.764 | 18.563 | 15.612 | 20.419 | -26.553 | 28.074 | 25.183 | 22.155 |
| -22.694 | 24.956 | -1.768 | 8.247 | -5.223 | -12.610 | 6.953 | 3.627 | -7.467 | 20.623 | -11.418 | 7.649 |
| -9.196 | 27.286 | 16.287 | 6.742 | -24.372 | -6.873 | 6.258 | 15.866 | -10.151 | 5.079 | 9.544 | 25.870 |
| -16.905 | 8.023 | 5.906 | -16.212 | 6.037 | 21.729 | -24.272 | 8.365 | 25.354 | 13.497 | 0.874 | 25.889 |
| 14.674 | 19.616 | -18.903 | -15.315 | 18.947 | -18.502 | 22.066 | 28.518 | 17.461 | 11.227 | 19.622 | 17.018 |
| -24.817 | -19.142 | -27.344 | -21.302 | 2.493 | -2.848 | -22.143 | 1.498 | 7.290 | 17.634 | 13.584 | -11.912 |
| 23.451 | 0.854 | -24.628 | 7.104 | -11.418 | -3.506 | -12.759 | 27.725 | 24.311 | 15.037 | 7.192 | 18.235 |
| 9.945 | 9.417 | -19.944 | 4.175 | 24.955 | 20.083 | 23.765 | 12.193 | 28.572 | 19.373 | 21.480 | 19.284 |
| 18.258 | -20.238 | -14.559 | 17.708 | -7.394 | -12.857 | -7.838 | -3.955 | 12.270 | 17.944 | 7.299 | 23.662 |
| 22.811 | 13.289 | 28.030 | -6.280 | 14.939 | -13.971 | -15.543 | 11.814 | 2.056 | 11.503 | -6.743 | 12.484 |
| 11.111 | -9.577 | 30.393 | 2.023 | 12.025 | -12.466 | -10.477 | 7.324 | -11.975 | 27.933 | 23.826 | 24.798 |
| -13.531 | 18.607 | -22.056 | 3.182 | -12.727 | -10.518 | -12.028 | 22.530 | 22.039 | 26.289 | 8.001 | 25.714 |
| 14.732 | -18.874 | 7.675 | 16.332 | -14.935 | -3.959 | -24.278 | 14.899 | 15.608 | 20.083 | 25.187 | 23.480 |
| -6.175 | -17.885 | -10.133 | -16.518 | -10.121 | 19.469 | -6.187 | 21.613 | 26.220 | -31.531 | 15.927 | 9.377 |
| 18.956 | 11.972 | -14.754 | -12.987 | -20.552 | 2.065 | 1.796 | 15.419 | 16.539 | 0.096 | 16.755 | 23.958 |
| 12.003 | 23.706 | -18.510 | -10.756 | -23.498 | -1.396 | -21.086 | 11.367 | 13.872 | 16.461 | 23.774 | 24.892 |
| -10.045 | 24.615 | -15.616 | -16.418 | 21.218 | -24.171 | -11.239 | 3.816 | -3.417 | 18.520 | 21.277 | -9.638 |
| 20.611 | 13.958 | 14.680 | 0.880 | 20.024 | -26.131 | 24.804 | 16.726 | 8.320 | 15.518 | 22.027 | 19.599 |
| -1.528 | 21.156 | 17.664 | -19.191 | -23.355 | -7.079 | 15.983 | 13.810 | -16.090 | 22.823 | 27.377 | 15.813 |
| -21.602 | 17.128 | 26.204 | 4.146 | -16.433 | -21.013 | 11.614 | 8.887 | 2.531 | 9.269 | 15.661 | 22.662 |
| -26.978 | -13.926 | -28.852 | -13.393 | -11.034 | -17.201 | -0.667 | -1.503 | -9.424 | 28.864 | 24.610 | 25.738 |
| -21.462 | -0.498 | -30.586 | -21.272 | -14.766 | 1.574 | 22.942 | 26.764 | 8.690 | 13.958 | 19.298 | 16.827 |
| -11.857 | 17.434 | -9.184 | -22.172 | 1.314 | 20.982 | -15.313 | 18.484 | 18.819 | 21.917 | 3.220 | 13.308 |
| -10.623 | 26.684 | -19.335 | -12.188 | 3.188 | -6.684 | -21.218 | 9.823 | -4.423 | 10.748 | 13.388 | 5.256 |
| 4.939 | 28.226 | -4.171 | 14.812 | -24.142 | -9.615 | -19.339 | 4.914 | 21.271 | 19.937 | 25.321 | -0.519 |
| -2.322 | 12.332 | -6.491 | -9.013 | -6.292 | -24.273 | -22.955 | 23.657 | 20.241 | 11.638 | 17.027 | -22.689 |

| | | | | | | | | | | | |
|---|---|---|---|---|---|---|---|---|---|---|---|
| -15.787 | -17.019 | 0.618 | 16.769 | 4.011 | 22.473 | 25.164 | 8.869 | -3.034 | 9.962 | 14.124 | -20.088 |
| -25.052 | -12.552 | 16.178 | -10.482 | 0.360 | 8.966 | 21.946 | 17.249 | 24.909 | 0.917 | 22.948 | -21.309 |
| 15.101 | -9.539 | 29.724 | 7.962 | -12.344 | 4.308 | 12.570 | 6.174 | -20.171 | 17.455 | 20.475 | 16.734 |
| -21.977 | -24.581 | 15.010 | 17.685 | -25.139 | 26.172 | 4.479 | 12.011 | 19.486 | 14.886 | 10.875 | 9.628 |
| -9.922 | -25.133 | -12.377 | -6.830 | 21.649 | -12.616 | -1.363 | 5.733 | 3.629 | 24.382 | 19.754 | 14.721 |
| 20.837 | -9.128 | -2.479 | -0.609 | -23.199 | -17.199 | 11.844 | -3.544 | -6.704 | 25.943 | 23.066 | 10.392 |
| -24.591 | 23.605 | 27.077 | -2.610 | 20.070 | 2.839 | -5.445 | 24.005 | -9.932 | 10.536 | 14.092 | 12.579 |
| -4.850 | -19.420 | 18.917 | -19.322 | -11.097 | -10.658 | -17.341 | 15.454 | -5.427 | -1.389 | 7.741 | 20.259 |
| -7.721 | -11.766 | 11.161 | -1.369 | -15.647 | 25.330 | 15.929 | 29.686 | 29.500 | 20.181 | 26.673 | 16.158 |
| -28.841 | -14.695 | -12.513 | 22.961 | -7.767 | 13.811 | 25.755 | 25.627 | 28.567 | 14.449 | -1.179 | -6.462 |
| 14.105 | 19.133 | -26.714 | -3.897 | -12.636 | 22.928 | -13.481 | 17.837 | 10.797 | -14.571 | 5.640 | 18.034 |
| -26.311 | 3.341 | 3.102 | -15.390 | 4.947 | 16.716 | 10.500 | 12.984 | -3.398 | 28.384 | 22.169 | 23.066 |
| 11.192 | -15.173 | 1.746 | -6.084 | 11.495 | -15.654 | 9.670 | 2.617 | 23.531 | 18.166 | 20.770 | 17.682 |
| 22.237 | 26.767 | 19.386 | 24.073 | -22.722 | 26.691 | -10.009 | 1.294 | 16.183 | 20.934 | 10.056 | 12.784 |
| 16.186 | -12.811 | -3.635 | -23.834 | 7.105 | -19.080 | -26.824 | 7.964 | -25.001 | 7.333 | 9.119 | 19.267 |
| 23.819 | -19.127 | 15.993 | -2.621 | -17.527 | 23.050 | 25.016 | 27.465 | -11.508 | 29.183 | 13.991 | 13.885 |
| -1.537 | -8.277 | -15.376 | -18.827 | -3.909 | -16.822 | 27.362 | 28.597 | 22.820 | 22.004 | 4.035 | 0.679 |
| 18.465 | -22.663 | -28.802 | 1.827 | -13.859 | -26.510 | 26.802 | 10.930 | 24.526 | 18.592 | 14.674 | 15.449 |
| -23.597 | 22.687 | 3.466 | -10.190 | -22.064 | 22.537 | -6.723 | 14.277 | -22.146 | 22.805 | | 8.090 |
| 28.824 | -11.233 | -19.433 | 14.620 | -6.983 | -6.657 | 21.095 | 15.324 | 14.904 | -4.809 | | 10.547 |
| 10.355 | -11.777 | 6.338 | 9.954 | 13.684 | 1.967 | -19.931 | 31.034 | -13.529 | -0.880 | | 25.365 |
| 28.413 | -19.741 | -22.812 | -2.227 | 7.575 | -6.089 | -17.887 | 9.633 | -15.918 | 15.008 | | -15.039 |
| 16.577 | -2.001 | -5.668 | -11.600 | -13.080 | 0.500 | -5.978 | 8.769 | -12.415 | 24.790 | | 2.220 |
| 17.055 | -4.793 | 24.867 | -0.225 | -11.063 | -9.349 | 22.375 | 5.226 | -8.910 | 19.857 | | 11.096 |
| 0.239 | 24.338 | -24.712 | -13.605 | 1.799 | 21.296 | -19.039 | 10.770 | 11.104 | 19.079 | | -16.664 |
| 1.805 | 20.919 | -15.493 | -12.124 | -20.778 | 4.649 | -18.649 | 22.732 | -24.273 | 8.648 | | 21.717 |
| -0.271 | -13.092 | 23.647 | -0.895 | -4.712 | -0.879 | 11.419 | 11.777 | 0.593 | -0.260 | | 8.301 |
| -16.849 | -15.657 | -0.750 | 19.144 | -9.122 | 22.194 | -10.091 | -4.921 | 17.091 | 0.517 | | -15.269 |
| 21.662 | -10.817 | 22.254 | -21.260 | -15.881 | 4.983 | 17.731 | 25.751 | 8.590 | 15.966 | | 20.534 |
| -20.368 | 2.252 | 7.453 | -5.283 | -17.093 | 0.880 | 28.199 | 15.042 | -4.485 | 10.569 | | 6.577 |
| -7.607 | -21.146 | 30.735 | 11.597 | 0.323 | -1.047 | 7.361 | 18.730 | -1.500 | 27.334 | | -4.691 |
| 22.266 | 10.746 | 8.892 | -26.337 | -7.681 | 10.713 | 3.520 | 6.832 | -11.722 | 10.320 | | -3.780 |
| -15.210 | 11.338 | -5.972 | -19.283 | -0.270 | 24.818 | -23.015 | -2.083 | 25.025 | 8.601 | | 11.950 |
| -10.800 | 18.150 | 5.315 | 1.438 | -3.295 | 23.016 | 23.490 | 16.202 | 21.817 | 24.953 | | -2.831 |
| 30.140 | 8.615 | -9.119 | 26.724 | -24.039 | 22.120 | -5.996 | 1.459 | 16.196 | 11.307 | | 8.920 |

| | | | | | | | | | | | |
|---|---|---|---|---|---|---|---|---|---|---|---|
| 14.525 | -7.430 | 0.276 | -1.193 | -20.226 | 13.221 | -12.265 | 22.190 | 30.168 | 27.945 | | 23.671 |
| -20.280 | -19.222 | -11.783 | -16.249 | 4.266 | -10.478 | 26.974 | 29.270 | 12.498 | 20.175 | | 23.243 |
| -20.596 | 0.233 | 17.181 | 18.599 | -8.090 | -19.706 | -19.040 | 6.299 | -10.338 | 6.982 | | 20.793 |
| -24.736 | 17.895 | -5.149 | 12.598 | 14.415 | 23.700 | -10.891 | 8.559 | -14.064 | 4.291 | | 18.015 |
| -27.814 | 12.792 | -20.292 | -6.165 | -2.188 | -6.996 | 2.994 | 1.370 | 22.879 | 29.019 | | 15.900 |
| -30.658 | -0.434 | -29.939 | 5.365 | 8.587 | -23.569 | -13.221 | -4.927 | -14.526 | 15.358 | | -10.566 |
| -29.801 | -4.602 | 26.017 | -27.646 | -18.871 | -18.527 | -1.325 | 23.733 | 28.364 | 12.662 | | 21.217 |
| -25.675 | 11.640 | -26.177 | -2.967 | -9.587 | 0.257 | 21.195 | 20.640 | -6.385 | 17.240 | | -2.455 |
| 20.947 | -1.436 | 6.086 | -4.985 | -26.974 | 21.156 | 28.070 | 25.364 | -21.344 | 20.877 | | -3.109 |
| -22.020 | -25.614 | -19.818 | -16.116 | -12.593 | 19.141 | -12.356 | 27.471 | 8.952 | 8.753 | | 8.153 |
| 22.511 | 10.416 | -22.848 | -13.647 | -15.269 | 21.857 | 6.059 | -28.109 | 6.754 | 23.726 | | -0.865 |
| 26.534 | 8.624 | -18.988 | 8.427 | 27.014 | 2.632 | 3.960 | 26.381 | -10.419 | 14.715 | | 9.384 |
| -14.089 | -3.363 | 29.744 | -22.129 | -11.847 | 12.409 | 21.671 | 19.678 | 26.181 | 7.335 | | 18.401 |
| -4.400 | 9.446 | -27.701 | 0.665 | -14.448 | 22.836 | 16.397 | 14.398 | 9.567 | 19.414 | | 10.863 |
| 23.178 | 6.048 | -25.389 | 14.341 | 5.448 | 17.615 | 18.033 | 6.520 | -8.702 | 22.246 | | -23.633 |
| 25.092 | -12.026 | -24.143 | -7.047 | -0.801 | -23.167 | 20.707 | 16.593 | -18.423 | -4.094 | | 26.121 |
| 23.997 | 15.221 | 9.958 | -13.931 | -6.238 | 1.833 | -4.793 | 9.333 | 18.699 | 20.282 | | 14.844 |
| 30.367 | -1.873 | -8.313 | 0.487 | 25.865 | 5.412 | -18.524 | 23.726 | 7.763 | 4.628 | | 22.879 |
| 3.591 | -0.128 | -13.709 | 6.159 | -24.892 | -5.635 | 30.627 | 8.169 | 11.849 | 3.321 | | 0.031 |
| -8.636 | 3.928 | -23.360 | -3.730 | -19.083 | -23.806 | 29.145 | 0.952 | -12.616 | 13.628 | | -1.865 |
| -32.225 | 0.709 | -30.207 | -7.438 | -17.672 | -26.751 | 13.105 | 21.719 | -5.944 | -2.725 | | 12.799 |
| -15.679 | 23.178 | -5.399 | -13.026 | -9.488 | -6.383 | 22.288 | -1.966 | -7.153 | 25.973 | | -6.298 |
| 23.740 | 1.731 | 10.358 | -22.786 | 9.891 | -11.564 | 22.997 | 24.750 | -23.716 | 19.088 | | 24.746 |
| -1.433 | 7.046 | -8.710 | 16.479 | -5.701 | -1.216 | 8.184 | 3.379 | 3.212 | 18.524 | | 5.500 |
| 29.860 | 20.264 | -9.568 | 9.266 | -19.914 | -2.762 | -10.703 | -3.086 | -8.555 | 12.317 | | 16.760 |
| 15.033 | 6.136 | 9.376 | 5.102 | -18.483 | 19.339 | -13.405 | 12.887 | 16.965 | 1.148 | | 10.854 |
| 0.940 | -6.326 | -15.527 | 2.978 | 14.297 | 23.062 | -25.501 | 5.184 | -20.577 | 6.308 | | -5.148 |
| -13.664 | 4.444 | -13.039 | 1.492 | -21.817 | 21.639 | 29.735 | -7.397 | -10.757 | 27.939 | | 18.382 |
| -18.720 | -17.023 | -16.908 | 8.920 | -1.348 | 20.075 | 21.893 | -1.274 | -6.249 | -3.980 | | 22.287 |
| 25.407 | -25.825 | -15.037 | -26.243 | -12.651 | 17.444 | 30.315 | 13.608 | -21.115 | 15.470 | | 26.545 |
| -30.420 | 25.523 | -7.887 | 22.278 | -11.720 | -14.056 | -19.483 | 2.479 | 7.129 | 22.019 | | -0.435 |
| 7.589 | 24.886 | -1.695 | -25.386 | -23.629 | 22.972 | -31.227 | -3.596 | 10.957 | 24.256 | | 13.716 |
| -9.162 | 11.584 | -13.404 | -5.774 | 18.845 | 2.432 | 11.604 | 21.825 | -8.999 | 22.742 | | -3.901 |
| -21.702 | -22.181 | -20.092 | -12.500 | -19.240 | 29.177 | 30.830 | -2.662 | -29.308 | 12.561 | | 7.889 |
| -26.501 | 26.413 | -21.415 | -26.793 | -13.774 | 31.340 | -3.018 | -8.704 | 8.245 | -2.608 | | 6.523 |

| | | | | | | | | | | | |
|---|---|---|---|---|---|---|---|---|---|---|---|
| -8.727 | 19.571 | -28.709 | 10.248 | -2.301 | 6.381 | -21.565 | 11.716 | -1.919 | -3.815 | | 25.454 |
| 27.376 | -11.727 | 2.614 | 7.705 | 10.325 | 21.731 | 25.934 | 6.984 | -8.201 | 19.380 | | 28.657 |
| -27.090 | -20.283 | 11.781 | -18.282 | -7.245 | 28.638 | 7.871 | 9.981 | 5.164 | 17.987 | | 23.965 |
| -10.487 | 21.236 | 15.633 | -24.637 | 2.960 | 22.427 | 24.447 | 25.616 | 22.583 | 13.803 | | 25.259 |
| 5.594 | 3.223 | 9.431 | -19.416 | -1.920 | -6.138 | -20.177 | 12.254 | -9.765 | 20.578 | | 13.199 |
| -15.446 | 5.845 | 7.861 | -21.053 | -20.400 | 4.652 | 29.230 | 0.225 | -19.253 | 16.548 | | 19.246 |
| 1.408 | -11.035 | -27.817 | -5.841 | -27.620 | 4.058 | -12.665 | 3.097 | 9.899 | -4.969 | | -12.810 |
| -13.026 | 2.962 | 17.825 | 5.169 | -10.704 | -3.704 | -4.115 | -6.658 | -22.470 | 21.506 | | -10.725 |
| 2.752 | -7.216 | 2.142 | -13.563 | 20.012 | 15.566 | 27.493 | -8.714 | 11.735 | 22.491 | | -9.389 |
| 8.770 | -0.592 | -2.896 | -16.550 | -22.633 | 18.236 | 19.491 | 14.908 | -8.285 | 9.071 | | 26.122 |
| -22.665 | 27.626 | 10.979 | -2.859 | -20.724 | -10.247 | -17.420 | 14.152 | -22.859 | 10.194 | | 16.257 |
| 18.833 | 2.134 | 17.348 | 3.788 | 13.274 | -8.881 | 12.855 | 22.985 | 23.472 | 3.473 | | 24.628 |
| 20.940 | 12.254 | -11.123 | -24.122 | -24.138 | -12.439 | -15.596 | 0.979 | 2.555 | 26.134 | | 21.958 |
| 9.239 | 25.443 | 0.422 | 26.833 | -4.470 | -1.140 | 21.383 | -6.426 | -13.543 | 14.374 | | -8.744 |
| 3.429 | -24.677 | -14.407 | -17.772 | -10.228 | 27.041 | -19.079 | 1.816 | -2.964 | 23.232 | | 14.520 |
| 1.462 | -1.895 | -25.363 | 9.017 | 25.679 | 24.542 | 16.347 | -1.865 | -16.490 | 22.269 | | -3.636 |
| -3.196 | 18.714 | -1.923 | -9.031 | 4.275 | 20.364 | 18.780 | 4.535 | 5.412 | 4.975 | | 27.563 |
| 15.756 | -4.326 | 15.495 | -3.801 | -17.314 | 0.604 | 25.175 | -9.894 | -1.641 | 9.322 | | -4.863 |
| 20.538 | -26.794 | 11.857 | -4.256 | 9.102 | -23.206 | 12.900 | 7.127 | -29.353 | 25.082 | | 18.019 |
| 18.656 | 21.045 | 27.383 | -16.789 | -12.157 | 23.948 | 29.176 | -3.004 | 13.546 | 5.289 | | 12.467 |
| -28.875 | -4.024 | -6.337 | 26.490 | -25.780 | 19.578 | 26.809 | 5.703 | 8.732 | 19.606 | | 28.382 |
| -17.434 | -0.743 | 5.672 | -24.913 | 3.792 | 28.248 | 7.090 | 15.578 | 25.385 | 13.758 | | 18.370 |
| 8.074 | 8.449 | 9.944 | 0.325 | 27.661 | 15.154 | 27.739 | 17.180 | 12.053 | -3.112 | | 21.740 |
| -12.915 | 15.743 | -6.610 | -14.180 | -25.157 | -6.373 | 21.271 | 26.349 | 15.397 | 24.312 | | 5.731 |
| -4.067 | -11.579 | -7.823 | 21.441 | 10.889 | 20.652 | 16.123 | 2.621 | 10.203 | 8.671 | | 24.800 |
| -0.889 | -6.067 | 18.915 | 6.674 | -18.022 | 17.913 | -19.727 | 12.048 | -11.190 | 5.381 | | 10.944 |
| -15.930 | 19.492 | -22.743 | -26.218 | 14.739 | -3.195 | -4.745 | 3.437 | -6.308 | 11.271 | | 19.423 |
| -21.955 | -15.021 | 23.280 | 20.793 | 4.970 | -4.481 | -13.487 | -0.093 | 3.712 | 2.113 | | 1.290 |
| -17.292 | 3.540 | -19.458 | -16.050 | -7.853 | 25.640 | -27.799 | 28.115 | 20.543 | 19.410 | | -5.843 |
| 10.501 | -4.323 | 21.680 | 8.739 | -20.605 | -3.015 | 26.853 | 25.144 | 13.700 | 6.235 | | 3.186 |
| -8.094 | -24.267 | -2.483 | -6.512 | 6.745 | 27.607 | -3.959 | 0.579 | -4.390 | 16.448 | | 8.576 |
| 11.538 | 20.420 | -20.144 | 26.177 | 28.213 | -19.296 | 1.974 | 9.576 | -20.169 | 12.968 | | 28.140 |
| -18.464 | -10.148 | 5.432 | -29.551 | -8.872 | -2.237 | -10.122 | 28.351 | 19.793 | 22.883 | | 23.359 |
| 20.867 | 22.582 | -10.943 | -19.230 | -14.729 | 6.311 | 15.478 | 15.748 | 4.517 | -16.863 | | -8.315 |
| 13.301 | 6.636 | -25.242 | -21.141 | -17.956 | 18.300 | -26.878 | 23.421 | 25.803 | 4.056 | | 28.829 |

| | | | | | | | | | | | |
|---|---|---|---|---|---|---|---|---|---|---|---|
| 8.162 | 15.489 | 15.886 | -12.869 | -5.642 | 22.610 | 20.410 | 18.443 | 18.103 | 23.521 | | 16.437 |
| 16.074 | 9.943 | 4.629 | 3.902 | -12.166 | 16.986 | -28.119 | 3.826 | 7.942 | 16.727 | | -22.406 |
| 0.451 | 28.641 | -18.602 | -23.158 | -11.081 | -1.572 | 27.558 | 10.906 | 24.529 | 19.164 | | 22.390 |
| -14.171 | 20.230 | 27.535 | -9.946 | -18.487 | -20.544 | 24.867 | 5.599 | -5.640 | 10.262 | | 13.734 |
| -15.605 | -15.377 | 16.718 | -31.355 | -6.805 | -24.589 | 23.538 | 24.578 | -12.411 | 9.270 | | 10.582 |
| 18.019 | 17.712 | -9.532 | 21.583 | -8.170 | 12.862 | 20.789 | -0.788 | 17.044 | 17.439 | | 16.962 |
| -9.735 | -23.444 | 24.910 | 23.420 | 10.970 | 22.873 | 11.237 | 2.305 | -9.324 | 14.470 | | 16.038 |
| -10.954 | -28.557 | -6.563 | -10.454 | -9.039 | 28.539 | -19.048 | 1.277 | -7.127 | 3.615 | | 21.647 |
| -1.279 | 23.810 | -17.570 | -22.848 | -10.020 | 7.112 | 24.341 | 25.865 | 6.679 | 1.811 | | 20.467 |
| -21.974 | 9.019 | -10.955 | -3.381 | -3.537 | -3.117 | 1.245 | -2.275 | 3.496 | 12.179 | | -9.117 |
| 7.612 | 3.738 | 4.722 | 2.912 | -21.330 | -10.075 | -11.645 | 28.790 | -16.654 | -3.246 | | 28.612 |
| -1.950 | 1.641 | -13.492 | 6.553 | 3.072 | -9.071 | -22.790 | 10.595 | -14.721 | 2.090 | | -5.005 |
| -13.331 | -24.048 | 15.085 | -7.465 | 13.419 | 8.211 | -21.754 | 9.588 | -19.507 | 2.412 | | -14.673 |
| 12.458 | 16.919 | -25.514 | -17.757 | 5.099 | 11.081 | 23.366 | 7.819 | 23.813 | -22.497 | | 7.261 |
| -24.501 | -3.773 | -2.720 | 18.131 | -11.327 | -17.430 | 6.920 | 6.203 | 9.470 | 4.310 | | 2.638 |
| 8.211 | 8.500 | 10.981 | 5.850 | -6.725 | 9.038 | 10.181 | 5.703 | 15.445 | 9.182 | | 18.573 |
| 6.583 | 3.167 | -9.352 | 27.438 | -3.149 | 3.553 | 19.005 | 4.573 | -27.335 | -3.294 | | 20.372 |
| 9.160 | -6.192 | -23.015 | 7.388 | -8.815 | 7.871 | 5.192 | -7.911 | -5.212 | 21.871 | | 13.887 |
| -5.963 | 30.662 | 22.549 | -21.582 | -19.661 | -0.265 | 6.349 | -9.760 | -10.748 | 17.894 | | 25.652 |
| -11.916 | -9.929 | -4.953 | 3.826 | -10.172 | -23.131 | 26.238 | 10.241 | 22.674 | 1.139 | | 30.365 |
| -4.096 | -17.253 | 7.453 | 0.748 | -14.920 | 16.414 | 14.174 | 19.083 | -21.720 | 20.309 | | 16.416 |
| 21.113 | -18.632 | 4.104 | 22.323 | -29.146 | 4.834 | 24.402 | -23.779 | 3.659 | 24.432 | | 14.592 |
| -2.876 | -10.504 | -21.787 | 14.602 | -17.829 | -25.017 | 18.669 | 26.246 | -2.400 | 24.020 | | -9.990 |
| 0.304 | 8.279 | -27.682 | -7.398 | -16.312 | 13.503 | 3.048 | 10.333 | -3.648 | 9.594 | | -22.192 |
| 3.161 | 31.271 | 23.905 | -8.185 | 31.194 | 2.100 | 25.031 | -11.106 | -17.080 | -3.692 | | -8.972 |
| 26.411 | 4.144 | 3.892 | -23.691 | -24.819 | 7.411 | -6.408 | 27.674 | -11.662 | 8.895 | | 23.934 |
| 17.229 | -23.874 | -11.384 | -18.610 | -23.988 | 8.602 | 11.603 | -0.999 | 7.313 | 6.899 | | 22.903 |
| 18.669 | 12.686 | 4.485 | 19.980 | -0.984 | 3.580 | 0.717 | 12.033 | 14.522 | 10.233 | | 21.501 |
| -3.703 | -13.941 | 10.013 | -12.228 | -7.486 | -2.329 | 19.962 | 1.048 | -7.778 | 19.143 | | -4.836 |
| -9.424 | 25.353 | -7.194 | -16.711 | 3.834 | -8.396 | -13.610 | 2.064 | -18.133 | 12.408 | | 16.246 |
| -12.231 | -15.583 | 25.961 | -25.825 | 15.532 | -20.548 | -25.239 | 1.336 | -19.046 | -26.766 | | 13.383 |
| -13.037 | 18.602 | 26.800 | 2.236 | -8.768 | -7.414 | -26.820 | 7.191 | 22.393 | 17.080 | | 24.764 |
| 2.016 | 7.647 | -2.660 | -8.227 | 24.271 | 19.420 | -17.317 | -3.301 | 16.942 | 15.205 | | 20.522 |
| 7.039 | -22.883 | 20.130 | 26.429 | -29.552 | 16.867 | 2.593 | 15.692 | 19.927 | 23.321 | | 11.015 |
| -26.692 | 21.794 | 5.224 | 5.775 | -19.597 | 11.399 | -24.572 | 14.372 | 7.287 | -11.045 | | 15.176 |

| | | | | | | | | | | | |
|---|---|---|---|---|---|---|---|---|---|---|---|
| 7.476 | -5.364 | -6.906 | -15.116 | -9.907 | 4.708 | 26.484 | 25.098 | -2.984 | 3.967 | | 18.979 |
| 0.517 | -14.636 | 0.003 | -29.733 | 9.386 | 26.745 | -8.931 | -2.034 | -9.441 | 9.606 | | 27.902 |
| -3.754 | -26.985 | -9.562 | 30.467 | -21.272 | 1.905 | -23.827 | 23.366 | 14.643 | 22.552 | | 9.875 |
| -14.388 | -5.718 | -0.872 | 9.903 | 12.985 | -7.190 | -16.164 | 9.370 | 10.494 | -0.546 | | -29.460 |
| -1.515 | -24.693 | -24.158 | 3.680 | -12.714 | -24.029 | 25.555 | -4.901 | 3.783 | 16.612 | | 23.679 |
| 12.757 | 11.493 | -2.044 | 8.318 | -5.395 | 20.969 | 18.897 | 5.901 | 2.413 | 24.914 | | 4.337 |
| 3.871 | 30.353 | 21.994 | -11.863 | 23.128 | -11.644 | -0.910 | 17.366 | -12.176 | 8.531 | | -5.961 |
| -12.043 | 3.638 | -3.950 | -23.105 | 8.128 | 1.076 | -9.559 | 16.200 | 25.121 | 5.986 | | 13.792 |
| 7.486 | 6.956 | -12.748 | 9.312 | -29.417 | 4.002 | 20.150 | 8.051 | 4.325 | 5.603 | | 9.119 |
| 24.045 | -25.159 | 4.002 | -3.177 | -15.288 | 9.860 | 8.837 | -10.837 | -24.344 | 5.170 | | 28.179 |
| 9.597 | 13.548 | -8.251 | -10.829 | -7.436 | -12.311 | 15.496 | -5.714 | 15.889 | 14.436 | | 25.487 |
| 22.644 | -6.483 | -23.042 | -16.599 | -25.120 | 3.335 | -20.145 | 15.066 | -11.673 | 23.600 | | 30.530 |
| 14.941 | 1.042 | 10.751 | -15.208 | -9.490 | -9.898 | 7.414 | 13.713 | -20.106 | 21.452 | | 16.561 |
| 13.209 | 29.968 | -9.637 | 14.317 | 29.113 | 11.841 | -22.937 | 21.636 | 3.630 | 8.033 | | -1.273 |
| 7.348 | -9.955 | 12.253 | -3.566 | 2.962 | -2.619 | 19.258 | 18.880 | 26.104 | 15.660 | | -11.416 |
| 0.964 | -0.397 | -3.532 | 9.764 | -20.313 | -18.131 | 18.079 | 25.823 | 18.977 | 10.270 | | 5.424 |
| 13.841 | -29.441 | 1.477 | 25.100 | -18.714 | -3.070 | 15.337 | -0.752 | -12.992 | 7.040 | | 0.513 |
| -1.979 | -23.986 | 13.675 | 4.837 | -4.459 | 9.562 | 17.246 | 24.338 | 8.032 | 12.928 | | 25.758 |
| | -24.592 | 17.843 | 18.624 | -13.487 | -22.074 | 20.556 | 19.612 | 16.741 | -6.181 | | 10.587 |
| | 7.827 | -1.629 | -11.718 | 0.035 | 16.613 | 15.608 | 11.614 | -23.417 | 18.504 | | -0.474 |
| | -12.338 | 2.885 | -21.929 | 1.255 | 23.387 | -0.393 | 6.995 | 9.546 | 24.269 | | -3.004 |
| | -23.146 | 8.720 | -17.292 | -19.753 | -8.197 | 11.632 | 10.624 | -20.691 | 14.846 | | 16.696 |
| | -26.818 | 9.977 | -9.024 | -21.934 | -12.135 | 12.875 | 22.455 | 10.552 | 0.988 | | 23.961 |
| | 1.651 | -24.409 | -20.387 | -14.758 | 19.487 | -5.948 | 23.969 | 0.647 | 15.338 | | 18.406 |
| | -4.350 | 0.849 | 1.851 | -7.724 | 12.492 | 0.479 | -21.815 | -10.709 | 12.196 | | 22.127 |
| | -22.098 | -26.062 | -12.489 | -16.446 | -4.856 | -19.128 | 27.624 | 23.563 | 21.840 | | 20.865 |
| | -25.028 | 11.973 | 8.463 | -23.718 | 5.665 | -10.526 | 17.723 | 10.241 | 13.941 | | 8.082 |
| | 4.430 | -19.163 | -13.709 | 27.012 | 14.896 | 1.494 | 20.370 | 24.054 | 8.511 | | 8.792 |
| | 26.580 | -5.374 | 17.523 | -26.111 | | 2.670 | 19.325 | -14.523 | -8.800 | | -3.090 |
| | -3.113 | 3.743 | -4.115 | -20.480 | | 17.310 | 10.347 | 3.145 | 25.852 | | 23.261 |
| | -10.512 | -4.348 | 7.777 | 22.291 | | 19.362 | 18.811 | -17.815 | -23.583 | | -1.595 |
| | 17.758 | -8.650 | 9.099 | 14.899 | | 8.512 | 5.449 | 16.395 | 23.151 | | 6.346 |
| | 22.358 | -9.179 | -27.145 | -17.566 | | -4.865 | 15.083 | 23.178 | -2.226 | | 23.836 |
| | -23.862 | -16.827 | 20.997 | -27.811 | | -3.767 | 16.874 | 14.607 | 16.284 | | 18.640 |
| | 7.355 | -12.164 | 20.197 | 4.137 | | -2.181 | 9.843 | 7.921 | 9.432 | | 24.467 |

| | | | | | | | | | | | |
|---|---|---|---|---|---|---|---|---|---|---|---|
| | 5.959 | -10.126 | -2.423 | -1.587 | | 0.355 | 6.706 | 8.632 | 27.417 | | 9.537 |
| | -19.467 | 11.364 | -19.509 | -19.896 | | 4.119 | 8.461 | 16.195 | 7.453 | | 16.710 |
| | 10.154 | -9.413 | 28.838 | -23.216 | | -0.190 | 15.826 | 15.168 | 8.532 | | -7.967 |
| | 28.345 | -8.320 | 15.857 | 26.877 | | -18.511 | 1.654 | -3.060 | 11.890 | | 12.783 |
| | -26.049 | -7.992 | 14.772 | -13.604 | | 9.977 | 12.791 | -1.105 | 16.182 | | 25.203 |
| | -24.147 | -12.213 | 0.609 | -24.200 | | 19.281 | 7.796 | 13.967 | 21.728 | | -4.244 |
| | -4.160 | 8.473 | 21.774 | 17.168 | | -23.644 | 17.566 | 24.536 | 15.192 | | 26.474 |
| | 22.862 | 15.879 | -25.827 | -14.979 | | 2.496 | 15.052 | -0.655 | -5.054 | | 25.020 |
| | 2.609 | -8.118 | -24.417 | 27.431 | | -19.789 | 6.403 | 11.252 | 19.930 | | 0.205 |
| | 26.493 | -12.628 | 18.174 | -27.136 | | 11.696 | -7.797 | 0.148 | 25.950 | | 11.652 |
| | -11.057 | -2.892 | -8.748 | -25.142 | | -7.351 | 4.447 | 13.226 | 15.815 | | 17.252 |
| | -6.471 | -1.859 | -3.933 | -18.245 | | 1.337 | 21.141 | 17.904 | 11.128 | | 9.870 |
| | 16.111 | -6.862 | 1.910 | -13.244 | | 19.933 | 21.920 | 22.914 | 18.224 | | 8.159 |
| | 12.132 | -20.977 | 6.779 | -15.123 | | -16.881 | 7.484 | 9.387 | -2.044 | | -21.168 |
| | -21.492 | -20.216 | -20.779 | -21.771 | | 23.750 | 19.560 | 7.584 | 21.661 | | 6.272 |
| | -17.845 | -15.851 | -10.510 | -20.576 | | 11.029 | -1.279 | 16.387 | 1.308 | | 28.979 |
| | 2.210 | -9.448 | 17.442 | -8.989 | | 9.515 | 10.289 | 2.030 | 9.405 | | 13.826 |
| | 22.164 | -6.154 | -29.261 | -6.061 | | 20.618 | 16.351 | 18.718 | 22.622 | | 17.564 |
| | 11.182 | -24.911 | 0.429 | 6.549 | | 5.217 | 3.360 | 0.544 | 24.576 | | 0.456 |
| | -18.854 | 7.637 | 3.925 | -9.724 | | -0.701 | 0.627 | 10.411 | 17.933 | | -12.867 |
| | 24.398 | -7.959 | 22.541 | -6.311 | | 24.666 | 13.498 | -20.829 | 22.590 | | 20.440 |
| | -23.311 | -18.284 | -14.917 | -7.418 | | -16.762 | -20.583 | 21.534 | 12.267 | | -12.392 |
| | 20.982 | -0.563 | 2.671 | 21.250 | | -24.180 | 23.413 | 25.526 | 25.849 | | 11.340 |
| | -0.916 | 22.399 | -7.111 | -24.928 | | 15.296 | -16.036 | 13.605 | 24.220 | | 6.421 |
| | 25.796 | 13.275 | 2.093 | 6.941 | | -2.516 | 17.316 | 24.893 | 20.115 | | 27.894 |
| | 20.574 | -11.701 | 24.108 | -16.908 | | -7.744 | 6.737 | 3.861 | -0.773 | | 12.366 |
| | -9.835 | 4.309 | -25.476 | -22.686 | | 14.890 | -13.217 | 20.459 | 27.925 | | -20.689 |
| | -10.670 | 3.991 | 6.551 | -17.806 | | -13.113 | 20.867 | 1.485 | 14.065 | | 1.200 |
| | -6.223 | 3.017 | -17.660 | -11.374 | | 16.803 | -2.664 | -24.262 | 21.123 | | 8.706 |
| | -3.922 | 22.850 | -4.201 | -16.051 | | -4.216 | 15.916 | -13.406 | 22.375 | | 5.584 |
| | -1.492 | -20.315 | -3.314 | 17.031 | | -13.785 | -4.112 | 10.607 | 14.301 | | 21.292 |
| | 7.555 | -26.621 | -13.588 | -4.077 | | -12.399 | 10.929 | 6.629 | 0.610 | | 15.089 |
| | -18.213 | -15.597 | -18.153 | -0.442 | | -26.186 | 13.119 | 25.729 | 20.786 | | 25.578 |
| | 20.605 | -6.765 | 28.092 | 17.714 | | -18.671 | 11.709 | 4.221 | 27.482 | | -3.320 |
| | 15.686 | -11.037 | 7.649 | -25.802 | | 1.840 | 7.740 | 9.981 | 18.205 | | 9.512 |

| | | | | | | | | | | | |
|---|---|---|---|---|---|---|---|---|---|---|---|
| | 2.885 | 13.308 | -9.036 | -11.647 | | 3.651 | 1.942 | 21.084 | -16.871 | | 22.501 |
| | -6.983 | -20.931 | 11.954 | -15.429 | | -11.455 | 1.101 | 4.511 | 25.128 | | 6.963 |
| | 3.719 | 23.777 | -11.342 | -12.046 | | -3.740 | -0.650 | 3.596 | 17.174 | | 21.346 |
| | 26.106 | 25.278 | 27.314 | -17.541 | | -15.048 | 13.895 | -14.918 | 27.372 | | 27.210 |
| | -3.614 | -22.360 | 25.248 | -13.870 | | 11.401 | 5.514 | 13.110 | 26.012 | | 10.336 |
| | -8.469 | -9.189 | 26.248 | -3.216 | | 19.095 | 3.090 | 7.277 | 23.881 | | 16.226 |
| | 1.739 | -0.529 | -0.823 | -16.882 | | 28.524 | 2.003 | 19.418 | 0.704 | | 24.267 |
| | -5.979 | 12.068 | 8.316 | -10.001 | | 16.768 | 12.058 | 3.920 | 18.446 | | 14.516 |
| | -8.948 | 16.636 | 14.773 | -5.105 | | 8.796 | 23.128 | 2.915 | 13.409 | | -1.963 |
| | 18.377 | 25.781 | 8.908 | 10.554 | | -24.880 | 6.802 | 13.768 | 14.313 | | 23.202 |
| | 6.554 | 27.511 | 12.193 | -18.055 | | 24.175 | 24.620 | 21.229 | 2.652 | | -3.916 |
| | 5.644 | 7.534 | -19.136 | -14.349 | | -21.825 | 22.040 | 0.319 | 29.631 | | 7.386 |
| | 14.941 | -3.073 | 5.060 | -9.787 | | 5.713 | 20.921 | 22.455 | 16.869 | | 27.898 |
| | -3.271 | -11.597 | -10.008 | 16.756 | | -15.978 | 11.005 | 17.549 | 19.609 | | 26.944 |
| | 3.971 | 22.645 | -8.532 | 9.637 | | 27.376 | -7.654 | 25.736 | 20.163 | | 19.259 |
| | -0.804 | 5.573 | -17.715 | -6.071 | | 23.097 | 9.113 | -8.901 | 11.553 | | 13.143 |
| | -24.525 | 5.004 | -11.938 | -4.757 | | -19.776 | 2.889 | 5.914 | 5.239 | | 15.142 |
| | -12.771 | -18.275 | -1.970 | -8.832 | | 26.140 | 13.239 | 19.204 | 15.887 | | 9.414 |
| | 16.322 | -16.318 | 17.326 | -11.170 | | 3.282 | 7.394 | 24.187 | 19.938 | | 10.232 |
| | -7.728 | -17.036 | 22.344 | -12.453 | | 3.960 | 8.296 | 21.498 | 4.200 | | 6.180 |
| | 5.250 | -20.810 | -4.318 | -4.842 | | 27.656 | 24.256 | 4.127 | 24.171 | | 22.319 |
| | 0.112 | -8.342 | 0.871 | -13.790 | | 20.626 | 12.862 | -0.418 | 12.499 | | 5.173 |
| | 18.499 | 12.383 | -15.193 | 9.639 | | 6.114 | 4.509 | 7.355 | 15.860 | | 8.255 |
| | -16.406 | -0.863 | -19.190 | 4.666 | | 8.582 | -3.245 | 20.100 | 6.616 | | 19.811 |
| | | 17.180 | -23.905 | 10.546 | | 11.425 | 14.835 | 22.497 | 18.315 | | 25.589 |
| | | -4.173 | -6.957 | 3.056 | | 13.802 | 11.140 | 3.531 | 19.695 | | 26.603 |
| | | -6.974 | 4.818 | -9.692 | | 5.134 | -1.756 | 5.395 | 5.544 | | 6.728 |
| | | -0.142 | -0.029 | 21.580 | | 6.869 | 4.241 | 17.171 | 29.395 | | 11.272 |
| | | 11.060 | -26.710 | -24.483 | | -22.161 | 19.787 | 15.323 | -6.108 | | -0.524 |
| | | 4.951 | -26.108 | 22.331 | | 21.454 | 11.497 | 16.708 | 7.148 | | -5.085 |
| | | 27.800 | 8.840 | 9.206 | | -4.617 | 6.214 | 13.374 | 9.835 | | 16.667 |
| | | -4.883 | -1.012 | 6.934 | | -24.299 | 10.279 | 14.419 | 28.399 | | -13.344 |
| | | 8.691 | 9.845 | -23.309 | | 28.763 | 1.460 | 24.922 | 17.974 | | -26.592 |
| | | -9.472 | -17.932 | 25.725 | | 11.032 | 17.614 | 5.997 | 25.030 | | 14.908 |
| | | 14.181 | -5.938 | 5.389 | | 0.163 | 14.786 | 0.111 | 23.202 | | 20.114 |

| | | | | | | | | | | | |
|---|---|---|---|---|---|---|---|---|---|---|---|
| | | -17.285 | -1.039 | 6.594 | | -3.502 | -4.746 | 15.689 | 26.779 | | 21.561 |
| | | 2.960 | 6.121 | -4.996 | | -21.080 | 2.087 | 12.220 | 12.535 | | -3.397 |
| | | -0.597 | 3.917 | 8.899 | | -6.596 | 24.204 | 0.570 | 16.547 | | 14.463 |
| | | -15.224 | 16.220 | 7.347 | | -1.653 | -3.663 | 18.432 | 1.807 | | 2.597 |
| | | 12.053 | -10.596 | 5.515 | | 19.482 | 3.075 | 20.540 | 21.717 | | -23.612 |
| | | -13.341 | -5.104 | 10.608 | | -19.117 | 15.074 | -0.928 | -4.480 | | 18.912 |
| | | -1.030 | 17.176 | -18.148 | | -11.457 | 6.649 | 14.982 | 13.475 | | 4.196 |
| | | 6.655 | 8.463 | -20.948 | | -17.045 | 27.283 | 21.545 | -2.296 | | 25.381 |
| | | 25.406 | 12.853 | 12.763 | | 26.889 | -8.323 | 28.538 | -1.900 | | 20.901 |
| | | 5.825 | -26.241 | -6.922 | | -13.964 | 13.858 | 11.508 | 23.197 | | 16.527 |
| | | 4.436 | -11.797 | 14.428 | | -23.261 | 17.146 | 19.844 | 24.686 | | 23.396 |
| | | -23.419 | -21.718 | 1.767 | | -9.978 | 21.836 | -15.044 | 17.195 | | -2.829 |
| | | 12.812 | 23.696 | 7.688 | | -2.902 | 6.070 | 13.094 | 14.930 | | 10.129 |
| | | -11.087 | 7.642 | -4.220 | | 21.679 | 24.143 | -9.458 | 16.177 | | -8.294 |
| | | 16.041 | -20.761 | 11.823 | | -19.095 | 13.532 | 3.656 | 27.271 | | -11.387 |
| | | 1.789 | | -2.288 | | 4.531 | 2.356 | -1.230 | 7.776 | | 19.875 |
| | | 14.731 | | -24.200 | | 25.761 | 22.922 | 9.807 | 18.711 | | 24.590 |
| | | -6.163 | | -5.479 | | -21.673 | 17.368 | 27.805 | 25.768 | | 5.157 |
| | | -28.275 | | 4.052 | | 6.762 | 5.678 | 16.290 | 17.203 | | 11.911 |
| | | 20.019 | | -8.572 | | -3.685 | 16.211 | 12.850 | 14.718 | | 20.050 |
| | | 20.654 | | -0.866 | | -12.996 | 1.646 | 28.558 | 26.003 | | 6.609 |
| | | -18.495 | | -2.366 | | 8.071 | -9.094 | 27.290 | 4.857 | | 23.327 |
| | | -20.762 | | -15.473 | | -20.919 | 14.339 | 0.915 | 13.048 | | -0.800 |
| | | 13.359 | | -13.019 | | 19.358 | 12.664 | 18.603 | 15.701 | | 19.493 |
| | | -16.952 | | -17.276 | | 27.934 | 9.733 | 19.904 | 28.758 | | 16.852 |
| | | 9.605 | | 21.668 | | -23.735 | 9.293 | 7.071 | 10.213 | | 26.205 |
| | | -11.575 | | 2.879 | | 5.348 | 3.681 | 24.210 | 13.543 | | 25.217 |
| | | -16.680 | | -18.521 | | -12.237 | 26.138 | 29.476 | 3.871 | | 7.482 |
| | | 17.226 | | -8.099 | | 12.550 | 6.895 | -21.044 | 20.109 | | 3.605 |
| | | 9.068 | | 23.487 | | 29.311 | 7.324 | 20.360 | 8.586 | | 20.408 |
| | | -14.060 | | -22.806 | | 26.820 | 10.078 | 27.841 | 30.380 | | 16.393 |
| | | 23.311 | | -7.117 | | 15.950 | 19.646 | 17.735 | 15.271 | | 4.534 |
| | | 14.934 | | 12.127 | | -25.377 | 16.135 | 22.096 | 18.016 | | 14.242 |
| | | -20.065 | | 3.822 | | 9.422 | 18.557 | 16.284 | 7.690 | | 21.621 |
| | | 26.607 | | 6.909 | | 4.649 | 4.504 | 6.913 | 28.308 | | 12.570 |

| | | | | | | | | | | | |
|---|---|---|---|---|---|---|---|---|---|---|---|
| | | -2.602 | | -15.579 | | 18.380 | 3.342 | -12.090 | 20.096 | | 11.418 |
| | | 2.290 | | -21.530 | | 8.292 | 2.598 | 19.519 | -2.756 | | -3.976 |
| | | 19.755 | | 0.412 | | 26.078 | 5.896 | 18.781 | 29.140 | | 4.335 |
| | | -26.627 | | -0.967 | | 9.992 | 10.811 | 5.780 | 23.670 | | 12.557 |
| | | 27.174 | | 9.233 | | 10.585 | -2.635 | 24.377 | 12.433 | | -12.632 |
| | | -9.698 | | -12.201 | | -21.855 | 15.079 | 19.862 | 25.415 | | 0.997 |
| | | -26.065 | | -9.667 | | -7.848 | 23.548 | 26.745 | 7.568 | | |
| | | 1.061 | | -16.638 | | -20.740 | 0.990 | 15.241 | 24.103 | | |
| | | -1.552 | | 2.079 | | 28.993 | 20.657 | 28.409 | 8.929 | | |
| | | -5.409 | | 8.960 | | 25.304 | -1.389 | 0.799 | 21.031 | | |
| | | 2.020 | | 21.004 | | 11.000 | 9.208 | 19.095 | 10.776 | | |
| | | -25.422 | | 13.938 | | 27.282 | 11.005 | 20.621 | 23.052 | | |
| | | 8.362 | | -13.325 | | -24.016 | 1.140 | 16.356 | 19.913 | | |
| | | 11.965 | | 4.535 | | 4.872 | 21.936 | 5.036 | 24.687 | | |
| | | -17.072 | | -3.134 | | -4.555 | 10.048 | 22.360 | 16.695 | | |
| | | -8.758 | | 3.398 | | 18.165 | 24.384 | 19.845 | 26.263 | | |
| | | -9.550 | | -1.785 | | 10.554 | 18.999 | 16.732 | 14.162 | | |
| | | -13.099 | | 0.374 | | 26.757 | 15.876 | -1.085 | 20.689 | | |
| | | -18.783 | | 8.086 | | 4.293 | 19.927 | 25.957 | 19.315 | | |
| | | -1.998 | | 20.016 | | 16.364 | 25.160 | 18.098 | 29.575 | | |
| | | -12.412 | | -10.371 | | -26.366 | 8.711 | 8.112 | 26.976 | | |
| | | 27.778 | | -23.119 | | 0.238 | 25.389 | 13.629 | 26.118 | | |
| | | -19.300 | | -24.868 | | 13.226 | 11.952 | 28.265 | 19.928 | | |
| | | -4.550 | | 28.401 | | 19.771 | 3.910 | 7.421 | 22.539 | | |
| | | -16.760 | | 20.322 | | 13.768 | 20.912 | -1.678 | 18.349 | | |
| | | 12.449 | | -20.422 | | -4.631 | 0.454 | 29.223 | 20.441 | | |
| | | -21.155 | | 16.854 | | -25.505 | 1.077 | 20.535 | 28.211 | | |
| | | -26.816 | | -12.354 | | 20.851 | 23.116 | 20.060 | 26.705 | | |
| | | -18.401 | | -11.212 | | 29.479 | 17.256 | 27.594 | 21.701 | | |
| | | 11.612 | | -21.518 | | 25.201 | 24.352 | 26.200 | 9.056 | | |
| | | -2.687 | | -23.862 | | 30.824 | 8.300 | 17.713 | 23.499 | | |
| | | -5.704 | | 0.145 | | 26.474 | 2.638 | 19.489 | 12.698 | | |
| | | -9.153 | | 9.528 | | 13.003 | -2.179 | 11.585 | 11.175 | | |
| | | -32.104 | | -13.532 | | -2.651 | -1.405 | 21.242 | 9.983 | | |
| | | -17.383 | | 26.349 | | -23.117 | 14.690 | 21.785 | -3.680 | | |

| | | | | | | | | | |
|---|---|---|---|---|---|---|---|---|---|
| | | -15.791 | | -4.021 | | 7.877 | 26.356 | 25.237 | 25.703 |
| | | 20.976 | | 10.891 | | 23.750 | -3.419 | 28.876 | 20.318 |
| | | 10.109 | | -18.517 | | 28.407 | 1.558 | -1.083 | 8.503 |
| | | -25.215 | | -9.852 | | 25.727 | 23.826 | 18.057 | 24.028 |
| | | 7.323 | | 27.923 | | 15.166 | 11.381 | 26.675 | 16.381 |
| | | -19.521 | | -23.239 | | 11.073 | 18.643 | 3.197 | 15.320 |
| | | 28.830 | | 11.963 | | -10.703 | 15.543 | 10.019 | 9.429 |
| | | -6.455 | | -15.680 | | 13.776 | 19.467 | 22.060 | 10.938 |
| | | -6.890 | | -14.630 | | 21.279 | 14.346 | 11.006 | 10.066 |
| | | -20.790 | | -18.254 | | -7.183 | 9.101 | 15.792 | 27.034 |
| | | 11.503 | | 3.786 | | 2.347 | 13.848 | 28.763 | 21.601 |
| | | -20.488 | | -18.830 | | 23.156 | 10.109 | 13.526 | 14.792 |
| | | -14.290 | | -10.385 | | -27.004 | 14.853 | 7.957 | 23.922 |
| | | -2.838 | | 6.550 | | 12.445 | 9.303 | 15.031 | 0.164 |
| | | 21.454 | | -11.794 | | 22.346 | 23.005 | 29.390 | 22.552 |
| | | 20.058 | | -3.985 | | 17.624 | 13.974 | 27.585 | 3.408 |
| | | -11.246 | | 8.294 | | 26.028 | 18.168 | 30.272 | -26.529 |
| | | -16.301 | | 19.110 | | 19.715 | 10.669 | 26.652 | 28.112 |
| | | -17.878 | | -14.003 | | 23.702 | -5.128 | 25.943 | 24.863 |
| | | 5.965 | | 15.906 | | 16.988 | 22.492 | 26.334 | 18.274 |
| | | -13.104 | | -0.049 | | 23.352 | 13.118 | 1.207 | 22.026 |
| | | 20.758 | | -15.074 | | 1.572 | 16.666 | 22.000 | -5.095 |
| | | -20.479 | | -4.006 | | 18.426 | 25.326 | 12.319 | 11.386 |
| | | -21.666 | | 2.737 | | 24.933 | 24.103 | 21.284 | 12.114 |
| | | -9.815 | | -24.546 | | 6.838 | 20.980 | 20.556 | 24.862 |
| | | -18.867 | | -23.325 | | 26.256 | -1.454 | 15.155 | 14.667 |
| | | 31.037 | | 9.998 | | -19.237 | 8.765 | 18.250 | 22.839 |
| | | 27.743 | | 3.516 | | 1.169 | -7.149 | 13.951 | 13.740 |
| | | -5.784 | | -3.057 | | -30.632 | 11.028 | -5.291 | 9.401 |
| | | 16.852 | | -13.196 | | 22.193 | 12.485 | 28.499 | 0.741 |
| | | 26.176 | | 8.267 | | 24.198 | -4.210 | 10.327 | 26.953 |
| | | -3.356 | | -20.597 | | 19.201 | 9.010 | 2.527 | 21.989 |
| | | 8.906 | | 22.727 | | 1.522 | -30.791 | 16.690 | 20.610 |
| | | -23.741 | | 27.208 | | 0.310 | 17.626 | 14.085 | 24.532 |
| | | -10.622 | | 21.484 | | 23.699 | 9.466 | 28.222 | 19.757 |

| | | | | | | | | | |
|---|---|---|---|---|---|---|---|---|---|
| | | -21.750 | | -18.565 | | -20.292 | 7.597 | 1.625 | 24.832 |
| | | 18.940 | | 11.053 | | -21.518 | 4.550 | 11.903 | 15.180 |
| | | 14.961 | | -17.984 | | 27.657 | 27.275 | 12.483 | 13.739 |
| | | 11.283 | | -10.148 | | 14.392 | 12.933 | 29.306 | 21.890 |
| | | -29.940 | | -13.178 | | 18.847 | 13.784 | -0.744 | 17.853 |
| | | -23.387 | | -14.707 | | -10.413 | 18.828 | 22.733 | 26.045 |
| | | 15.445 | | 13.082 | | 25.713 | 16.020 | 30.181 | 11.666 |
| | | 14.178 | | -20.099 | | 19.459 | 15.530 | 23.303 | 23.203 |
| | | -15.321 | | 0.238 | | 27.741 | 10.504 | 21.068 | 9.144 |
| | | -26.103 | | -8.355 | | 24.562 | -6.903 | 5.513 | -8.720 |
| | | -2.033 | | 17.449 | | 25.442 | 9.775 | 4.889 | 25.561 |
| | | -18.167 | | -18.740 | | 18.526 | 22.664 | 26.955 | 24.892 |
| | | -7.226 | | 1.530 | | -23.681 | -4.123 | 13.142 | 13.940 |
| | | -21.704 | | 2.774 | | -29.821 | 15.369 | 28.385 | -8.884 |
| | | -23.678 | | 14.188 | | 22.861 | 11.998 | 19.373 | 21.266 |
| | | 22.855 | | -7.558 | | 7.217 | 25.769 | 16.176 | 26.189 |
| | | -0.727 | | -15.637 | | -19.928 | 14.686 | 2.071 | 13.686 |
| | | -18.250 | | 30.105 | | -26.488 | 9.044 | 26.271 | 17.457 |
| | | -22.018 | | 5.783 | | 1.222 | -1.906 | 21.871 | 21.948 |
| | | 18.630 | | -21.610 | | 21.007 | 0.058 | 4.284 | 14.859 |
| | | -4.005 | | -16.831 | | -18.891 | 27.137 | 24.556 | 9.748 |
| | | -24.908 | | -26.358 | | 11.583 | 15.856 | 14.443 | 22.692 |
| | | 14.984 | | 6.849 | | 27.145 | -2.235 | 5.049 | 19.725 |
| | | 20.106 | | -12.619 | | -24.692 | 18.358 | 24.398 | 25.158 |
| | | 5.952 | | -23.204 | | 21.719 | 31.099 | 17.330 | 15.072 |
| | | 25.904 | | -2.311 | | 20.557 | 25.363 | 18.535 | -4.303 |
| | | -19.893 | | -20.842 | | -18.596 | 5.512 | 9.629 | 24.289 |
| | | -3.131 | | 16.915 | | 11.525 | -8.802 | 7.118 | 1.848 |
| | | 27.611 | | -14.209 | | -25.678 | 7.987 | 20.323 | 21.582 |
| | | 28.293 | | -8.991 | | 16.187 | 17.551 | 27.236 | 18.559 |
| | | 20.701 | | -24.241 | | -27.109 | 23.056 | 23.564 | 13.370 |
| | | -2.169 | | 8.184 | | -22.017 | 21.526 | -7.596 | 9.954 |
| | | 15.664 | | -11.925 | | 14.195 | 22.128 | 26.593 | 14.699 |
| | | -9.320 | | -24.765 | | -23.691 | 18.363 | 18.635 | 17.897 |
| | | -6.406 | | -18.080 | | 8.325 | -11.388 | -0.208 | 19.333 |

| | | | | | | | | | |
|---|---|---|---|---|---|---|---|---|---|
| | | -21.073 | | 0.204 | | -23.094 | 16.812 | 2.778 | 15.468 |
| | | -23.029 | | -10.484 | | -0.115 | -17.891 | 29.004 | -7.135 |
| | | 21.635 | | -13.511 | | 23.490 | -8.885 | 1.952 | 20.870 |
| | | -22.108 | | -6.730 | | 11.430 | 23.151 | 25.100 | 19.099 |
| | | 9.218 | | -7.668 | | 14.157 | 4.371 | 12.639 | 14.394 |
| | | -19.464 | | 28.156 | | -21.361 | 0.944 | 15.645 | 10.640 |
| | | -25.687 | | 23.676 | | 22.630 | 7.733 | 17.682 | |
| | | 11.882 | | -19.485 | | 21.502 | -3.037 | 3.257 | |
| | | 19.779 | | -16.867 | | 25.623 | 15.165 | 8.734 | |
| | | 26.851 | | 16.264 | | -8.963 | 13.529 | 26.247 | |
| | | -11.515 | | -26.764 | | 23.146 | -7.411 | 10.510 | |
| | | 0.138 | | -18.931 | | 18.781 | 18.811 | 22.172 | |
| | | 26.109 | | 3.554 | | -2.119 | 27.779 | 20.737 | |
| | | -24.725 | | -12.625 | | 16.854 | 10.559 | -0.247 | |
| | | 2.044 | | 15.351 | | 9.468 | 16.402 | 24.996 | |
| | | 11.147 | | 19.238 | | -9.649 | 9.553 | 20.977 | |
| | | -18.527 | | 4.121 | | -10.392 | 13.830 | 23.047 | |
| | | -26.809 | | -5.049 | | 2.919 | 7.591 | 3.013 | |
| | | 19.871 | | -7.462 | | 18.564 | -15.718 | 9.273 | |
| | | 4.993 | | 5.269 | | 2.391 | 11.288 | 15.954 | |
| | | -8.956 | | 23.707 | | -5.872 | 4.614 | 17.591 | |
| | | -12.718 | | -2.982 | | 1.002 | 8.550 | 19.347 | |
| | | -0.272 | | -15.047 | | -8.987 | 5.623 | 11.564 | |
| | | -10.280 | | 26.736 | | -0.195 | 25.354 | 6.121 | |
| | | 13.799 | | 20.476 | | 9.373 | 14.508 | 16.185 | |
| | | 19.084 | | -15.493 | | 5.086 | 13.371 | 13.356 | |
| | | 13.126 | | -21.840 | | 0.770 | 16.849 | 27.641 | |
| | | -17.339 | | -13.472 | | -25.482 | 4.169 | 11.065 | |
| | | -2.850 | | -8.101 | | -22.503 | 23.704 | 10.051 | |
| | | -26.556 | | -27.905 | | -23.267 | -9.458 | 26.647 | |
| | | -2.052 | | -2.668 | | | 19.347 | 20.534 | |
| | | -10.324 | | -18.177 | | | 12.551 | 18.822 | |
| | | -7.784 | | 29.497 | | | 10.400 | 25.571 | |
| | | -21.712 | | 5.391 | | | -5.870 | 21.983 | |
| | | 6.619 | | -22.807 | | | 14.673 | 26.319 | |

| | | | | | | | | |
|---|---|---|---|---|---|---|---|---|
| | | 7.439 | | -5.999 | | | -3.845 | 9.148 |
| | | 5.066 | | -8.882 | | | 9.434 | 16.846 |
| | | 26.137 | | -16.363 | | | 13.446 | 17.462 |
| | | 19.306 | | -15.688 | | | 17.838 | 7.263 |
| | | -4.062 | | -4.677 | | | 26.711 | 9.127 |
| | | 23.047 | | -14.490 | | | 15.820 | 5.420 |
| | | -1.869 | | -11.255 | | | 7.134 | 14.561 |
| | | -24.278 | | -15.846 | | | -17.090 | -2.894 |
| | | 2.402 | | -9.268 | | | -5.312 | -7.477 |
| | | 16.670 | | -25.614 | | | -9.115 | 3.625 |
| | | -18.643 | | 5.295 | | | 29.416 | 8.964 |
| | | 2.310 | | -7.564 | | | 11.787 | 21.357 |
| | | -12.787 | | 18.066 | | | -6.554 | 7.989 |
| | | 6.269 | | -0.617 | | | 15.276 | 3.889 |
| | | 16.433 | | -20.282 | | | -2.013 | 12.858 |
| | | 18.502 | | 7.159 | | | 18.281 | 23.021 |
| | | -22.218 | | 17.045 | | | 6.526 | 20.489 |
| | | -23.567 | | 8.056 | | | -0.598 | 13.450 |
| | | 15.625 | | -18.027 | | | 13.863 | 23.521 |
| | | 8.328 | | 20.980 | | | 0.004 | 21.634 |
| | | -1.905 | | -20.860 | | | 29.001 | 8.890 |
| | | -20.987 | | -6.685 | | | 16.747 | 5.854 |
| | | -2.526 | | -10.742 | | | 13.191 | |
| | | -17.748 | | -22.621 | | | 7.493 | |
| | | -19.343 | | -21.440 | | | 17.583 | |
| | | -14.341 | | 6.766 | | | 15.735 | |
| | | -12.616 | | -19.994 | | | 14.380 | |
| | | -16.798 | | 5.606 | | | 2.517 | |
| | | 17.541 | | 20.549 | | | 25.200 | |
| | | 20.341 | | 16.368 | | | 14.855 | |
| | | -0.727 | | -7.567 | | | 19.338 | |
| | | -6.586 | | 9.832 | | | 23.924 | |
| | | -23.189 | | 3.180 | | | -2.019 | |
| | | 0.222 | | -22.565 | | | 18.242 | |
| | | -25.965 | | -9.813 | | | 0.166 | |

| | | | | | | | |
|---|---|---|---|---|---|---|---|
| | | 0.933 | | -18.370 | | | -2.849 |
| | | 21.638 | | 9.192 | | | 10.385 |
| | | -8.969 | | -15.416 | | | -0.771 |
| | | -0.577 | | 14.248 | | | 9.111 |
| | | 19.502 | | -6.145 | | | 22.756 |
| | | -1.412 | | -24.186 | | | 12.576 |
| | | 1.698 | | -20.462 | | | 6.130 |
| | | -19.132 | | 9.731 | | | 2.740 |
| | | -14.673 | | 26.236 | | | 5.329 |
| | | 16.782 | | -1.507 | | | 0.887 |
| | | -4.617 | | -13.917 | | | 11.673 |
| | | -6.925 | | -26.690 | | | 16.325 |
| | | 8.286 | | 10.026 | | | 13.909 |
| | | 10.733 | | 7.177 | | | 9.056 |
| | | -12.774 | | -24.096 | | | 17.227 |
| | | -11.476 | | -5.011 | | | 2.147 |
| | | -13.984 | | -12.338 | | | 15.577 |
| | | -22.776 | | -23.085 | | | 7.926 |
| | | -10.827 | | -9.460 | | | 6.363 |
| | | -19.589 | | 1.737 | | | 0.055 |
| | | -17.421 | | -22.288 | | | 27.225 |
| | | 21.724 | | -21.841 | | | 19.993 |
| | | -7.632 | | -21.382 | | | 12.416 |
| | | -16.101 | | -26.331 | | | 0.838 |
| | | -24.955 | | -28.639 | | | 18.144 |
| | | -20.037 | | 15.936 | | | 5.617 |
| | | -1.819 | | 5.896 | | | 25.801 |
| | | -5.021 | | 24.574 | | | 10.018 |
| | | -10.198 | | -20.311 | | | 14.992 |
| | | 5.784 | | -22.746 | | | 14.431 |
| | | 9.465 | | 1.086 | | | -0.853 |
| | | -18.099 | | -1.924 | | | 16.077 |
| | | -0.645 | | -13.513 | | | 21.084 |
| | | 24.742 | | -23.369 | | | 0.788 |
| | | 7.724 | | -24.434 | | | 24.844 |

| | | | | | | | |
|---|---|---|---|---|---|---|---|
| | | -17.685 | | -6.320 | | | -1.409 |
| | | 17.365 | | -0.368 | | | 18.492 |
| | | 10.952 | | -12.950 | | | 14.180 |
| | | -26.969 | | -4.636 | | | 12.379 |
| | | -6.455 | | -10.955 | | | 21.337 |
| | | -5.566 | | -24.075 | | | 16.591 |
| | | -13.663 | | 2.918 | | | 17.527 |
| | | 0.797 | | 26.783 | | | -17.144 |
| | | 11.409 | | 10.186 | | | 15.678 |
| | | 8.876 | | -25.309 | | | -8.839 |
| | | 20.098 | | -24.592 | | | 24.180 |
| | | -1.189 | | 0.838 | | | 11.915 |
| | | -24.405 | | -24.332 | | | 11.657 |
| | | 15.656 | | -17.991 | | | 21.909 |
| | | 1.268 | | 14.413 | | | -24.367 |
| | | -10.310 | | -8.863 | | | 23.289 |
| | | -7.817 | | 19.550 | | | 3.562 |
| | | 1.784 | | -8.682 | | | 13.172 |
| | | -2.861 | | -21.494 | | | 24.508 |
| | | -27.439 | | -24.490 | | | -2.805 |
| | | 5.563 | | 5.393 | | | 10.386 |
| | | -3.103 | | -17.761 | | | -7.563 |
| | | 10.091 | | -8.261 | | | 11.655 |
| | | -11.798 | | -18.569 | | | 0.621 |
| | | 2.260 | | -10.181 | | | -6.806 |
| | | 19.419 | | -4.194 | | | 22.796 |
| | | -4.371 | | 7.395 | | | 16.403 |
| | | -1.579 | | 1.298 | | | 13.871 |
| | | -3.880 | | -16.395 | | | 11.957 |
| | | -23.906 | | 21.763 | | | 16.847 |
| | | -9.162 | | -17.583 | | | -10.287 |
| | | 4.175 | | 13.133 | | | |
| | | -22.440 | | 23.301 | | | |
| | | -15.500 | | -6.502 | | | |
| | | 9.036 | | 7.891 | | | |

| | | | | |
|---|---|---|---|---|
| | | -5.206 | | -16.993 |
| | | 12.305 | | -13.652 |
| | | 4.006 | | -17.672 |
| | | -13.841 | | |
| | | 22.722 | | |
| | | 14.150 | | |
| | | -22.009 | | |
| | | -19.101 | | |
| | | 8.039 | | |
| | | -20.020 | | |
| | | 21.055 | | |
| | | -0.072 | | |
| | | -11.559 | | |
| | | -17.502 | | |
| | | -10.553 | | |
| | | 10.509 | | |
| | | -14.664 | | |
| | | -20.946 | | |
| | | -8.108 | | |
| | | -18.625 | | |
| | | -7.573 | | |
| | | 25.228 | | |
| | | 15.761 | | |
| | | 13.186 | | |
| | | 9.022 | | |
| | | 12.365 | | |
| | | 8.365 | | |
| | | 28.591 | | |
| | | 27.118 | | |
| | | -6.872 | | |
| | | -26.344 | | |
| | | 9.728 | | |
| | | -27.513 | | |
| | | 20.765 | | |
| | | 25.117 | | |

| | | |
|---|---|---|
| | | 7.839 |
| | | 27.406 |
| | | 21.306 |
| | | -18.399 |
| | | 6.767 |
| | | 8.652 |
| | | -20.004 |
| | | 16.970 |
| | | 1.995 |
| | | -20.841 |
| | | -4.030 |
| | | 13.594 |
| | | -20.597 |
| | | 17.622 |
| | | -19.793 |
| | | -16.925 |
| | | -28.188 |
| | | -13.246 |
| | | 21.432 |
| | | 22.320 |
| | | 15.545 |
| | | 1.002 |
| | | -18.138 |
| | | 6.620 |
| | | 23.801 |
| | | -19.013 |
| | | -23.065 |
| | | 0.260 |
| | | -28.402 |
| | | -12.363 |
| | | -9.453 |
| | | -19.388 |
| | | 17.682 |
| | | 15.341 |
| | | 7.138 |

| | | |
|---|---|---|
| | | 13.266 |
| | | -21.793 |
| | | -6.261 |
| | | -10.263 |
| | | -16.018 |
| | | 22.901 |
| | | -11.141 |
| | | 27.738 |
| | | 7.992 |
| | | 19.230 |
| | | 3.219 |
| | | -9.266 |
| | | -15.678 |
| | | -16.802 |
| | | 13.107 |
| | | 10.298 |
| | | -4.992 |
| | | -21.087 |
| | | -27.237 |
| | | -9.831 |
| | | -29.130 |
| | | -17.115 |
| | | -23.550 |
| | | 12.162 |
| | | 25.403 |
| | | -15.096 |
| | | -25.163 |
| | | -6.032 |
| | | 13.086 |
| | | -13.847 |
| | | 16.959 |
| | | -3.325 |
| | | -0.776 |
| | | 7.091 |
| | | -19.901 |

| | | |
|---|---|---|
| | | 10.320 |
| | | -21.065 |
| | | 20.198 |
| | | -20.545 |
| | | 18.909 |
| | | 24.706 |
| | | 16.669 |
| | | -1.947 |
| | | -16.823 |
| | | 12.859 |
| | | 26.879 |
| | | -11.575 |
| | | -18.944 |
| | | -21.028 |
| | | -21.649 |
| | | -30.415 |
| | | -27.485 |
| | | -14.611 |
| | | 26.269 |
| | | 18.236 |
| | | -21.541 |
| | | 28.007 |
| | | 13.522 |
| | | 20.711 |
| | | 12.538 |
| | | -6.934 |
| | | -19.929 |
| | | 24.417 |
| | | -18.280 |
| | | -23.443 |
| | | -24.001 |
| | | 3.319 |
| | | 14.944 |
| | | 16.694 |
| | | -7.305 |

| | | |
|---|---|---|
| | | -24.461 |
| | | -26.227 |
| | | -15.352 |
| | | 27.949 |
| | | -2.744 |
| | | 17.314 |
| | | 21.241 |
| | | 25.367 |
| | | -18.501 |
| | | -28.404 |
| | | 20.178 |
| | | -7.805 |
| | | -21.643 |
| | | 13.497 |
| | | 27.201 |
| | | 21.074 |
| | | -23.115 |
| | | -18.068 |
| | | 19.069 |
| | | -23.799 |
| | | 19.133 |
| | | 13.851 |
| | | -6.870 |
| | | 24.095 |
| | | 20.050 |
| | | 6.734 |
| | | 0.859 |
| | | 0.332 |
| | | -25.953 |
| | | -25.269 |
| | | -10.353 |
| | | 20.828 |
| | | 25.384 |
| | | 2.180 |
| | | -2.667 |

| | | |
|---|---|---|
| | | -19.975 |
| | | -2.896 |
| | | -8.253 |
| | | -22.850 |
| | | 7.057 |
| | | -15.979 |
| | | -23.688 |
| | | -6.620 |
| | | 17.724 |
| | | 7.318 |
| | | 0.126 |
| | | -2.991 |
| | | -16.742 |
| | | 6.891 |
| | | 22.107 |
| | | 16.970 |
| | | 15.723 |
| | | -25.128 |
| | | 1.321 |
| | | -23.583 |
| | | 19.966 |
| | | 3.344 |
| | | 14.976 |
| | | 4.169 |
| | | -14.665 |
| | | 14.578 |
| | | -13.214 |
| | | 0.945 |
| | | -18.447 |
| | | -19.955 |
| | | -24.579 |

| IA D12 | IA D13 |
|---|---|
| -5.067 | 0.100 |
| 24.996 | -8.727 |
| 16.474 | 15.544 |
| 8.324 | -5.568 |
| 16.092 | -7.310 |
| -3.110 | -14.418 |
| 0.728 | 21.135 |
| 17.181 | -25.608 |
| 18.454 | 17.644 |
| -4.411 | 10.427 |
| -22.851 | 24.120 |
| 21.447 | 5.053 |
| -5.412 | 1.221 |
| -13.088 | 8.519 |
| 24.118 | -21.232 |
| -8.211 | 3.963 |
| -21.390 | 23.538 |
| 13.875 | 18.961 |
| 15.469 | 21.728 |
| 23.358 | 2.830 |
| 22.112 | 5.877 |
| 16.728 | -15.174 |
| 20.573 | 1.105 |
| -22.578 | 24.626 |
| -25.838 | 16.455 |
| 25.479 | 25.517 |
| -13.716 | 21.333 |
| 1.535 | 17.355 |
| 26.601 | -11.624 |
| 20.938 | 26.438 |
| 14.995 | 6.542 |
| 8.150 | 24.177 |
| 25.716 | 21.958 |

| | |
|---|---|
| 22.512 | 23.037 |
| -6.462 | -26.417 |
| 28.170 | 23.746 |
| 9.154 | 17.715 |
| 24.556 | 10.900 |
| 15.000 | 24.803 |
| 11.565 | 20.079 |
| 23.268 | 26.464 |
| 22.265 | 18.495 |
| 27.981 | 27.910 |
| 25.840 | -2.239 |
| -27.698 | 4.459 |
| 29.546 | 27.307 |
| 13.691 | 21.549 |
| -24.764 | 27.864 |
| 21.002 | 23.599 |
| 6.361 | -17.262 |
| 16.938 | 22.079 |
| 25.341 | -6.681 |
| 28.033 | 28.453 |
| 29.094 | 19.866 |
| 12.895 | 21.170 |
| 20.817 | 24.923 |
| 10.742 | 12.376 |
| 30.170 | 24.237 |
| 22.189 | 11.495 |
| 22.878 | -24.448 |
| 28.822 | 21.453 |
| 26.372 | 26.047 |
| -14.214 | -5.846 |
| 24.552 | -28.796 |
| 13.337 | 17.385 |
| 6.343 | -28.115 |
| 27.583 | 22.912 |
| 29.302 | 16.733 |

| | |
|---|---|
| 28.536 | -11.816 |
| 18.351 | -29.989 |
| 22.815 | 27.031 |
| 25.413 | 16.045 |
| 16.504 | -22.889 |
| 23.860 | 24.838 |
| 18.114 | 22.699 |
| 9.938 | 25.869 |
| 30.428 | -9.238 |
| 18.725 | -6.786 |
| -15.887 | 20.101 |
| -26.676 | 0.796 |
| 26.842 | -3.139 |
| -17.109 | -29.023 |
| 14.077 | 27.824 |
| 27.746 | 20.137 |
| 28.382 | 9.117 |
| 18.368 | -0.320 |
| 21.597 | 17.641 |
| 26.334 | 12.944 |
| 9.569 | 10.407 |
| -5.251 | 24.671 |
| 13.475 | 22.398 |
| 30.166 | 18.713 |
| 28.945 | -21.379 |
| 28.030 | 30.841 |
| 20.085 | -18.752 |
| -11.941 | 21.156 |
| 18.031 | 24.785 |
| 21.242 | 23.717 |
| 29.028 | 20.904 |
| 8.073 | 18.655 |
| 18.264 | 13.452 |
| 29.408 | 1.816 |
| 18.005 | 26.736 |

| | |
|---|---|
| 10.860 | 21.430 |
| 16.028 | 27.703 |
| 29.038 | 15.771 |
| 26.605 | 11.923 |
| 28.108 | 18.488 |
| 10.234 | 20.015 |
| 23.915 | -19.461 |
| 6.231 | 28.478 |
| 25.460 | -10.759 |
| 27.675 | -17.242 |
| -5.295 | 17.115 |
| 9.693 | 22.303 |
| 16.733 | 26.766 |
| 26.717 | 4.567 |
| 21.623 | 14.676 |
| 28.001 | 18.289 |
| -3.730 | 28.277 |
| 8.444 | 24.825 |
| -0.018 | 16.259 |
| 15.102 | 23.955 |
| 25.550 | 3.083 |
| 16.067 | -14.961 |
| 23.225 | 27.673 |
| 27.745 | 7.241 |
| 17.420 | 0.325 |
| 4.878 | 22.273 |
| 8.179 | 3.418 |
| 4.564 | 17.625 |
| 22.092 | -11.491 |
| 26.446 | 20.020 |
| 13.449 | 21.245 |
| 18.908 | 24.396 |
| 10.352 | 23.261 |
| 9.886 | 3.582 |
| 3.261 | 25.812 |

| | |
|---|---|
| 20.091 | -20.387 |
| -2.396 | 21.461 |
| 18.170 | 17.479 |
| 7.392 | 22.031 |
| 16.799 | -16.875 |
| 21.564 | -10.360 |
| 11.124 | 24.066 |
| 23.738 | 19.694 |
| 16.969 | 1.393 |
| 14.418 | -10.415 |
| -22.014 | 16.188 |
| 6.562 | 1.015 |
| -1.660 | 20.626 |
| 10.135 | 12.122 |
| 25.030 | 10.329 |
| -2.983 | 17.405 |
| -4.507 | 9.607 |
| 14.516 | -1.419 |
| 19.163 | 19.767 |
| 15.897 | 16.661 |
| 17.954 | 8.780 |
| -9.512 | -17.784 |
| 20.659 | 6.227 |
| 16.077 | 12.714 |
| 18.510 | -9.819 |
| 8.112 | 23.408 |
| 15.899 | -18.577 |
| 23.212 | 8.741 |
| 25.478 | 20.460 |
| 6.795 | 13.090 |
| 19.229 | -12.583 |
| 20.147 | 24.444 |
| 22.339 | -12.160 |
| 1.187 | 16.828 |
| 10.591 | -0.523 |

| | |
|---|---|
| 10.526 | 18.290 |
| 15.048 | 15.649 |
| 19.200 | 2.168 |
| -16.251 | 14.284 |
| 18.284 | 0.421 |
| 21.351 | 20.540 |
| 24.626 | 19.773 |
| 17.038 | 21.292 |
| 20.582 | 7.814 |
| 24.024 | 17.539 |
| 23.621 | 25.127 |
| 13.107 | 15.083 |
| 5.608 | 21.588 |
| -8.728 | 24.084 |
| 10.631 | -8.592 |
| 12.838 | 22.191 |
| 13.947 | 11.864 |
| -7.909 | 1.264 |
| 11.328 | 9.730 |
| 26.471 | 21.036 |
| -0.270 | 18.155 |
| 5.667 | 9.849 |
| 23.025 | 25.876 |
| 11.752 | -1.561 |
| 5.275 | 8.816 |
| -14.277 | -6.671 |
| 15.386 | 22.111 |
| 26.733 | -3.493 |
| 16.424 | 24.256 |
| 19.524 | 14.070 |
| 8.153 | 10.807 |
| 21.490 | -5.225 |
| -6.266 | 20.827 |
| 14.158 | 18.246 |
| 18.181 | 10.340 |

| | |
|---|---|
| 20.105 | 12.667 |
| 19.461 | 3.685 |
| 28.797 | 14.825 |
| 11.271 | 25.377 |
| -30.224 | 17.572 |
| 25.509 | 22.679 |
| 17.654 | -8.303 |
| 18.915 | 5.703 |
| 18.514 | 2.061 |
| 27.416 | -3.245 |
| 24.282 | -9.173 |
| 26.387 | -8.765 |
| 22.220 | 27.518 |
| 11.500 | 23.851 |
| 28.955 | -1.836 |
| -4.144 | 20.779 |
| -5.365 | 14.724 |
| 16.116 | 21.446 |
| 22.781 | 20.228 |
| 16.878 | 11.412 |
| 21.030 | 12.627 |
| -21.027 | 26.454 |
| 25.241 | -25.145 |
| 0.062 | 14.358 |
| 29.618 | 25.316 |
| 14.195 | 25.962 |
| 9.934 | 18.351 |
| 23.303 | -6.258 |
| 20.658 | 22.185 |
| 26.526 | 24.366 |
| 2.437 | 28.361 |
| 14.388 | -3.684 |
| 15.133 | 10.677 |
| 23.851 | -5.064 |
| 21.375 | 1.947 |

| | |
|---|---|
| -4.505 | 18.799 |
| 18.829 | 21.641 |
| 13.189 | 11.737 |
| -26.160 | 9.713 |
| 13.746 | 27.873 |
| 22.356 | 20.235 |
| -3.499 | 13.466 |
| 16.953 | -30.492 |
| 29.330 | -13.709 |
| 20.137 | 22.488 |
| 13.350 | 19.715 |
| 8.288 | 16.678 |
| 21.868 | 23.129 |
| 2.953 | 7.195 |
| -31.843 | 25.243 |
| 26.302 | 28.930 |
| 21.339 | 2.893 |
| -7.026 | -12.176 |
| 12.262 | 25.589 |
| -2.467 | 17.351 |
| 16.286 | 23.706 |
| 18.614 | 10.572 |
| 10.459 | 24.356 |
| 1.046 | 29.808 |
| 13.642 | 12.779 |
| 28.087 | 18.807 |
| 11.435 | 12.147 |
| 25.772 | 15.916 |
| 22.031 | 9.451 |
| 17.159 | -22.561 |
| -23.012 | 21.140 |
| 24.882 | 14.857 |
| 16.715 | 20.283 |
| 20.047 | 22.647 |
| -9.972 | 17.470 |

| | |
|---|---|
| 21.286 | 29.123 |
| 14.083 | 2.908 |
| 14.608 | -30.281 |
| 16.236 | -16.541 |
| 26.013 | 28.356 |
| -29.929 | 30.064 |
| 12.782 | 20.384 |
| 22.176 | 11.184 |
| 27.785 | 5.933 |
| 26.962 | -11.715 |
| 17.618 | 27.589 |
| 5.223 | 20.993 |
| 3.007 | 24.385 |
| 18.849 | -31.530 |
| 3.648 | 23.857 |
| -11.799 | 19.165 |
| 25.337 | 17.288 |
| 10.268 | 27.720 |
| 19.511 | 26.550 |
| 15.765 | 23.489 |
| -6.935 | 20.877 |
| 23.764 | 27.448 |
| 9.943 | 23.709 |
| -1.794 | -3.511 |
| 16.838 | -30.545 |
| 12.154 | 22.633 |
| 22.351 | 17.580 |
| -14.232 | 14.033 |
| 5.381 | 24.422 |
| -19.784 | 16.023 |
| 12.833 | 26.052 |
| 10.718 | 11.954 |
| 0.578 | 21.974 |
| 18.175 | 3.657 |
| 25.294 | 28.662 |

| | |
|---|---|
| 25.846 | -12.547 |
| 16.014 | 29.202 |
| 12.987 | 25.242 |
| 18.663 | 13.405 |
| 22.891 | 20.345 |
| 10.022 | 21.821 |
| 11.318 | 16.274 |
| 5.735 | 1.394 |
| -10.408 | 13.803 |
| 21.038 | 20.011 |
| 15.727 | 25.269 |
| 4.629 | 24.065 |
| 25.825 | 8.796 |
| 21.231 | 17.668 |
| 18.263 | 19.464 |
| 24.890 | -28.153 |
| 20.653 | 12.912 |
| 23.189 | 11.918 |
| 13.895 | 27.632 |
| 9.037 | 23.332 |
| 8.057 | 17.234 |
| -15.530 | 7.637 |
| 24.195 | -19.491 |
| 19.135 | 18.709 |
| 17.181 | 19.765 |
| -12.998 | 17.764 |
| 22.362 | 13.991 |
| 15.071 | 24.294 |
| -8.294 | 22.251 |
| 16.659 | 12.688 |
| -3.457 | 26.309 |
| -2.413 | 16.240 |
| 20.443 | 24.268 |
| -2.663 | -7.365 |
| -1.498 | 14.489 |

| | |
|---|---|
| 21.560 | 28.502 |
| 17.847 | 13.490 |
| 20.994 | 19.125 |
| 2.076 | 19.401 |
| 20.567 | 12.589 |
| 19.988 | 20.714 |
| 14.988 | 27.023 |
| 14.758 | 25.775 |
| 14.501 | 7.517 |
| 23.690 | 14.499 |
| 11.543 | 22.402 |
| 10.690 | 12.680 |
| -0.981 | -9.702 |
| 17.815 | -1.194 |
| 16.642 | 16.172 |
| 5.891 | 21.501 |
| -25.253 | 0.963 |
| 16.942 | 11.576 |
| 25.595 | -8.143 |
| -4.268 | 26.355 |
| 19.856 | 25.959 |
| 20.511 | 22.698 |
| 3.611 | 24.630 |
| 20.161 | 12.828 |
| 24.407 | -22.392 |
| 11.743 | 16.855 |
| 17.184 | -12.075 |
| 18.113 | 12.374 |
| 25.456 | 10.326 |
| -5.586 | 19.699 |
| -22.776 | 14.250 |
| -22.440 | 9.582 |
| -4.308 | 15.849 |
| 9.028 | 18.475 |
| -12.246 | 21.648 |

| | |
|---|---|
| -13.871 | 13.691 |
| 18.486 | 4.700 |
| 23.249 | -5.334 |
| -1.825 | 24.931 |
| 20.293 | 10.506 |
| 24.944 | 1.689 |
| 5.465 | -26.426 |
| 6.582 | -8.467 |
| 23.217 | 9.145 |
| 3.385 | -0.302 |
| 21.213 | 15.924 |
| 21.959 | -19.804 |
| 12.986 | 7.776 |
| 27.631 | 1.157 |
| 25.569 | -18.911 |
| 13.937 | 23.999 |
| 28.418 | -20.263 |
| 9.312 | 23.008 |
| 26.146 | 14.734 |
| 26.519 | 8.341 |
| 22.194 | -10.279 |
| 22.980 | 9.701 |
| 12.619 | 15.746 |
| 17.679 | 4.999 |
| 27.095 | 15.795 |
| 26.146 | -10.161 |
| -3.198 | 18.885 |
| 17.543 | 2.828 |
| 23.607 | -4.557 |
| 6.460 | 25.350 |
| 14.177 | 6.569 |
| 27.694 | 20.281 |
| 1.014 | 23.769 |
| 18.495 | 8.602 |
| 3.077 | 3.025 |

| | |
|---|---|
| 20.436 | -13.559 |
| 29.300 | 24.424 |
| 24.888 | 21.715 |
| 13.478 | 20.086 |
| 23.783 | 24.385 |
| 21.352 | 7.486 |
| 6.264 | 26.213 |
| 24.222 | 12.630 |
| 26.162 | 8.287 |
| -28.716 | 2.094 |
| 19.127 | 16.458 |
| 22.125 | 14.849 |
| 21.351 | 21.053 |
| 31.848 | -25.765 |
| 27.508 | 14.940 |
| 23.306 | -15.017 |
| 26.189 | -6.061 |
| -0.835 | 24.732 |
| -5.503 | 23.656 |
| 14.307 | 24.359 |
| 15.535 | 19.728 |
| 27.002 | 16.327 |
| 24.684 | 13.820 |
| 16.757 | 26.634 |
| 30.516 | 9.864 |
| 22.490 | 21.930 |
| 14.695 | 21.141 |
| -0.243 | -18.456 |
| 25.667 | 25.913 |
| -4.678 | 11.295 |
| 30.071 | 25.502 |
| 19.515 | 19.536 |
| 23.328 | 4.638 |
| 21.722 | -8.853 |
| 6.366 | 25.769 |

| | |
|---|---|
| 3.430 | 18.494 |
| 18.123 | 21.231 |
| -2.888 | -20.519 |
| 24.405 | 17.266 |
| 30.522 | 2.951 |
| 29.896 | 25.375 |
| 13.926 | 22.788 |
| 8.886 | -5.519 |
| 20.323 | -16.788 |
| 24.641 | -14.283 |
| 6.112 | 19.215 |
| -5.576 | -6.919 |
| 13.309 | 26.501 |
| 11.432 | 27.170 |
| -1.224 | -4.868 |
| -25.718 | 21.224 |
| -2.455 | 19.766 |
| 17.478 | 18.433 |
| 24.630 | 3.641 |
| 20.345 | 29.159 |
| 18.547 | 23.873 |
| 13.608 | 27.979 |
| 29.382 | -12.804 |
| 21.832 | 23.300 |
| 19.336 | -17.503 |
| 8.916 | 24.662 |
| 25.817 | 14.370 |
| 25.877 | 26.537 |
| 13.297 | 22.241 |
| 26.935 | 12.785 |
| 12.346 | 0.550 |
| 9.941 | 7.906 |
| 14.494 | 27.656 |
| 17.715 | 6.759 |
| 28.006 | 12.063 |

| | |
|---|---|
| 24.364 | -1.513 |
| -26.426 | 24.793 |
| 22.592 | 29.724 |
| 25.799 | 26.105 |
| 25.113 | 10.158 |
| -13.834 | 13.262 |
| 3.211 | 23.919 |
| 7.672 | 22.382 |
| -1.529 | 15.952 |
| 21.260 | 21.894 |
| 16.308 | 11.713 |
| 19.315 | 21.487 |
| -2.631 | 25.992 |
| -6.129 | 28.492 |
| 22.713 | 15.233 |
| 0.604 | 8.018 |
| 25.137 | 9.825 |
| 5.516 | 19.713 |
| 23.989 | 29.223 |
| 27.180 | -22.425 |
| 7.782 | 20.817 |
| 26.472 | 13.496 |
| 25.812 | 24.895 |
| 26.084 | 29.771 |
| 17.591 | 7.809 |
| 19.121 | 15.976 |
| 20.022 | -25.253 |
| 18.301 | 30.438 |
| 23.122 | 26.144 |
| 14.143 | 23.781 |
| 24.808 | 26.866 |
| 22.267 | 7.643 |
| 10.730 | 0.900 |
| 18.000 | 7.463 |
| 21.388 | 27.401 |

| | |
|---|---|
| 23.010 | 20.987 |
| 16.374 | 20.145 |
| 12.565 | 24.158 |
| 2.474 | 22.637 |
| 9.126 | 14.399 |
| -24.071 | 11.556 |
| 21.011 | 5.033 |
| 18.182 | -6.942 |
| 19.046 | -12.693 |
| 21.156 | 8.068 |
| 24.416 | 12.981 |
| 8.301 | 23.393 |
| 21.706 | 14.174 |
| 13.707 | 14.749 |
| 16.604 | 24.185 |
| 19.939 | 18.464 |
| 26.059 | 28.295 |
| 8.463 | 28.831 |
| -11.365 | 24.828 |
| 25.198 | 21.300 |
| 16.423 | 11.663 |
| 15.564 | 25.998 |
| 23.554 | 22.329 |
| 24.304 | 10.845 |
| 12.555 | 27.066 |
| 26.221 | -10.436 |
| 27.103 | 20.017 |
| 6.262 | 8.653 |
| 9.055 | 24.949 |
| 22.186 | 16.334 |
| 25.268 | -8.535 |
| -14.400 | 21.405 |
| 11.854 | 26.726 |
| 27.775 | 25.943 |
| 23.470 | -16.950 |

| | |
|---|---|
| 25.594 | 21.741 |
| 10.327 | 25.305 |
| 13.152 | 7.850 |
| 26.567 | 16.135 |
| 7.214 | 8.504 |
| 24.792 | 23.722 |
| 13.574 | 15.428 |
| 27.504 | 26.803 |
| 16.769 | 22.338 |
| 24.003 | 20.288 |
| 10.616 | 17.420 |
| 8.878 | 21.063 |
| 23.159 | 12.028 |
| 25.791 | 27.506 |
| 26.924 | 20.908 |
| 20.729 | 3.861 |
| 1.659 | 15.459 |
| 8.530 | -7.510 |
| 24.837 | 11.662 |
| 27.416 | 27.011 |
| 26.616 | -6.192 |
| -1.339 | 2.035 |
| 10.168 | 26.507 |
| 24.445 | 8.388 |
| 0.072 | 23.976 |
| 19.420 | 25.051 |
| 9.508 | 7.207 |
| 16.407 | -23.385 |
| 19.398 | 13.702 |
| 15.934 | 16.787 |
| 17.067 | 21.362 |
| 18.782 | 20.156 |
| 3.763 | 22.032 |
| 11.552 | 2.580 |
| 27.807 | -1.681 |

| | |
|---|---|
| 30.390 | 15.944 |
| 14.364 | 23.931 |
| 22.639 | 15.455 |
| 20.433 | 8.982 |
| 12.400 | 1.837 |
| 21.892 | -0.442 |
| 18.112 | -18.156 |
| 21.587 | 2.577 |
| 11.939 | 16.129 |
| 28.666 | -2.107 |
| 17.519 | 22.580 |
| 27.557 | -17.119 |
| 19.972 | 12.538 |
| 8.413 | 5.275 |
| 17.331 | 2.345 |
| -20.405 | 8.453 |
| 24.480 | 15.131 |
| 14.207 | -15.053 |
| 18.468 | 10.431 |
| -10.831 | 8.031 |
| 16.385 | 4.655 |
| 19.078 | 0.535 |
| -0.398 | 11.346 |
| 23.082 | 6.005 |
| 29.215 | 14.811 |
| 25.848 | -4.122 |
| 27.218 | 10.086 |
| 28.556 | 8.237 |
| 24.059 | -24.095 |
| -13.680 | -3.656 |
| 9.792 | 17.261 |
| 17.411 | -5.886 |
| 28.095 | 23.080 |
| 24.792 | 19.871 |
| 8.736 | 10.078 |

| | |
|---|---|
| 26.100 | 18.915 |
| 18.643 | 9.373 |
| 26.722 | 3.948 |
| 10.356 | 3.077 |
| -4.833 | 4.663 |
| 15.918 | 2.593 |
| 19.240 | -26.939 |
| 12.963 | 22.281 |
| 8.243 | 17.888 |
| -10.130 | 20.813 |
| 9.624 | 13.115 |
| 17.461 | 5.455 |
| 22.189 | 21.930 |
| 20.669 | -8.662 |
| 26.700 | 24.295 |
| 15.341 | -20.568 |
| 11.501 | 25.184 |
| -21.512 | 2.169 |
| 11.973 | 4.162 |
| 16.201 | 21.677 |
| 22.669 | 10.376 |
| 23.757 | 23.923 |
| 26.123 | 6.538 |
| 24.372 | 19.804 |
| 5.857 | 12.582 |
| 17.711 | 25.842 |
| 4.961 | 2.355 |
| 24.975 | 14.167 |
| 19.752 | 9.955 |
| 19.482 | 23.988 |
| 17.162 | 24.759 |
| 21.101 | 22.665 |
| 26.399 | 20.974 |
| 4.856 | 14.673 |
| 4.201 | 9.280 |

| | |
|---|---|
| 6.986 | -0.704 |
| 16.062 | 19.897 |
| 22.386 | -27.137 |
| 2.512 | 17.744 |
| -3.322 | 25.995 |
| 12.263 | 20.372 |
| 21.855 | 24.421 |
| 10.999 | 12.856 |
| 24.068 | 0.507 |
| 0.943 | 18.713 |
| 5.312 | 15.444 |
| 25.395 | 9.379 |
| 2.732 | 24.848 |
| 23.343 | 12.037 |
| 24.184 | 21.323 |
| 16.111 | 17.255 |
| 22.040 | 28.468 |
| -0.994 | 26.820 |
| 17.868 | 24.230 |
| 13.131 | -1.910 |
| 7.639 | 13.199 |
| 23.612 | 20.293 |
| 20.535 | 28.019 |
| 3.936 | 20.175 |
| -11.248 | 25.098 |
| 5.777 | 9.220 |
| 0.054 | 16.034 |
| -12.369 | 14.499 |
| 25.679 | 18.593 |
| 24.547 | 29.494 |
| 12.801 | 1.533 |
| 12.115 | 17.203 |
| 17.785 | 24.608 |
| 14.539 | 10.248 |
| 20.784 | -7.281 |

| | |
|---|---|
| 15.153 | 15.772 |
| 23.304 | 25.861 |
| 18.056 | -1.338 |
| 3.530 | 22.957 |
| 19.735 | 24.234 |
| 15.341 | 14.349 |
| 21.200 | 22.262 |
| 23.755 | 18.857 |
| 7.300 | 19.377 |
| 12.798 | 23.104 |
| 7.990 | 16.489 |
| 24.631 | 14.651 |
| 11.470 | 11.499 |
| 25.486 | 19.606 |
| 15.868 | 18.213 |
| 13.307 | 22.390 |
| 23.921 | 28.425 |
| -2.053 | 30.292 |
| 20.075 | 18.488 |
| 10.228 | 7.809 |
| 18.563 | 9.628 |
| 17.703 | 1.014 |
| 22.459 | 14.357 |
| 25.242 | 21.907 |
| 27.766 | 25.668 |
| -2.890 | 26.902 |
| 7.202 | 21.224 |
| 4.219 | 27.597 |
| 8.079 | 24.089 |
| 28.533 | 19.602 |
| 25.865 | 26.074 |
| 6.821 | 24.860 |
| 21.859 | 1.291 |
| 16.208 | 15.971 |
| 29.133 | 16.769 |

| | |
|---|---|
| 24.198 | 19.587 |
| 17.599 | -14.494 |
| 11.687 | 25.990 |
| 19.932 | -0.553 |
| 23.507 | 10.015 |
| 1.871 | 27.327 |
| -6.899 | 5.595 |
| 27.079 | 16.997 |
| -8.369 | 14.864 |
| -9.307 | 27.796 |
| 12.215 | 12.086 |
| 15.391 | 21.235 |
| 24.316 | 22.779 |
| 18.657 | 12.589 |
| 28.726 | 9.456 |
| 24.622 | 0.740 |
| 25.190 | -2.542 |
| 23.530 | 17.703 |
| 6.499 | 19.342 |
| 27.652 | 10.080 |
| 26.199 | 20.935 |
| 16.056 | 26.621 |
| 25.142 | -13.661 |
| -23.450 | 28.715 |
| -19.657 | 15.933 |
| 20.242 | -7.377 |
| 27.062 | 23.368 |
| 28.343 | 10.067 |
| 0.195 | 23.265 |
| 17.515 | 13.773 |
| 18.095 | 18.002 |
| 21.440 | 23.712 |
| 31.070 | 21.386 |
| 17.366 | 15.216 |
| 4.003 | 24.715 |

| | |
|---|---|
| 9.203 | 16.582 |
| 30.362 | 14.160 |
| 29.372 | 26.606 |
| 28.365 | 24.357 |
| 4.479 | 25.468 |
| 1.952 | 27.081 |
| 29.090 | 22.397 |
| 23.957 | 16.652 |
| 31.754 | 10.215 |
| 1.837 | 3.487 |
| -7.544 | 20.325 |
| 9.567 | 22.347 |
| 25.788 | 6.283 |
| 30.829 | 15.656 |
| 22.373 | 23.394 |
| 14.888 | 18.290 |
| 30.268 | 14.771 |
| -8.924 | 20.310 |
| 1.985 | 11.870 |
| 17.600 | -5.338 |
| 15.401 | -3.735 |
| 20.770 | 1.398 |
| -13.504 | -2.327 |
| 25.080 | 14.093 |
| 2.786 | 20.548 |
| 14.223 | 13.384 |
| 26.445 | 8.718 |
| 28.086 | 19.926 |
| 18.740 | 8.008 |
| 13.213 | 1.817 |
| 5.177 | -0.628 |
| 10.553 | 21.290 |
| -5.784 | 5.973 |
| 29.665 | -0.747 |
| 11.146 | 23.154 |

| | |
|---|---|
| 25.756 | -5.338 |
| -11.609 | 24.524 |
| 21.382 | 0.798 |
| 9.742 | -7.637 |
| -0.879 | 25.043 |
| 17.648 | -18.277 |
| 7.999 | -24.949 |
| -7.534 | 15.265 |
| 3.764 | -11.115 |
| 26.724 | 18.508 |
| 14.330 | 16.546 |
| -8.798 | -4.067 |
| 20.360 | 17.167 |
| 4.760 | -6.764 |
| 9.379 | 19.253 |
| 8.778 | -17.505 |
| -10.081 | -14.633 |
| 22.537 | 8.245 |
| -10.380 | -20.874 |
| 29.655 | 16.408 |
| 8.104 | 21.974 |
| 10.911 | -6.522 |
| 28.046 | 22.723 |
| 21.428 | 2.181 |
| 25.562 | 18.351 |
| 2.605 | -25.518 |
| 27.501 | 24.793 |
| 11.124 | 13.697 |
| -7.027 | 10.896 |
| 9.316 | 11.904 |
| -2.742 | 12.381 |
| 22.099 | 19.091 |
| -10.257 | 23.043 |
| 26.948 | 12.614 |
| 12.319 | 9.930 |

| | |
|---|---|
| 28.663 | 12.759 |
| -7.447 | 23.165 |
| 16.487 | 15.322 |
| -4.568 | 27.132 |
| 15.445 | 17.574 |
| 10.271 | -8.910 |
| -13.795 | 23.189 |
| 21.718 | 7.478 |
| 27.061 | 18.592 |
| -16.005 | 25.661 |
| 22.569 | 26.242 |
| 9.827 | 16.414 |
| 20.185 | -5.671 |
| 28.158 | -19.543 |
| 16.777 | 12.437 |
| 18.406 | -1.738 |
| 25.609 | 15.540 |
| 24.771 | 8.892 |
| -11.629 | -8.111 |
| 23.705 | -4.159 |
| 3.708 | 19.518 |
| 27.351 | 21.957 |
| 8.250 | 8.419 |
| -23.646 | -26.173 |
| 28.409 | 23.790 |
| 8.954 | 10.203 |
| 12.145 | 15.988 |
| 22.362 | 3.588 |
| 26.529 | -21.181 |
| 27.308 | 27.050 |
| 0.955 | 12.202 |
| 25.840 | 22.354 |
| 15.568 | 20.868 |
| 9.640 | 24.450 |
| 10.482 | 20.408 |

| | |
|---|---|
| -7.091 | 19.785 |
| 17.351 | 22.942 |
| 18.797 | 18.657 |
| 25.217 | 13.281 |
| 24.072 | 10.374 |
| -0.244 | 22.114 |
| 21.491 | 15.041 |
| 10.795 | 24.207 |
| 22.861 | 25.005 |
| 11.805 | 20.096 |
| 18.426 | 23.486 |
| 21.284 | -13.563 |
| -9.012 | -19.433 |
| -6.399 | 25.884 |
| 4.829 | 12.272 |
| 17.329 | 22.959 |
| 11.729 | 20.329 |
| 6.660 | 8.056 |
| -20.027 | 16.597 |
| 17.351 | -19.747 |
| 19.789 | 15.065 |
| -8.353 | 7.283 |
| 19.304 | 27.222 |
| 6.845 | 20.536 |
| 17.253 | 24.912 |
| 12.240 | 31.043 |
| 12.977 | 30.122 |
| 13.635 | -27.084 |
| -0.603 | 17.540 |
| -1.231 | 18.314 |
| 22.301 | 22.092 |
| -4.446 | 20.822 |
| 7.792 | 28.777 |
| 14.167 | 22.644 |
| 16.798 | -2.649 |

| | |
|---|---|
| 20.753 | 26.585 |
| 12.077 | 18.177 |
| -5.782 | -10.789 |
| 25.017 | -23.370 |
| -8.015 | -25.472 |
| -12.907 | 10.822 |
| 17.698 | 31.862 |
| 10.735 | 15.794 |
| 9.382 | 12.133 |
| -20.417 | 24.639 |
| -24.395 | 31.307 |
| 14.299 | -6.850 |
| 3.147 | 30.013 |
| 23.354 | 27.393 |
| -7.958 | 26.403 |
| 0.991 | 21.113 |
| 9.178 | 16.491 |
| 3.695 | 31.196 |
| 14.653 | 5.426 |
| -12.024 | 13.672 |
| 9.581 | 29.427 |
| 19.318 | 25.858 |
| 22.367 | -6.760 |
| 26.073 | 15.482 |
| 0.722 | 23.714 |
| 13.606 | 17.737 |
| 18.523 | 16.317 |
| 7.522 | 29.341 |
| 23.205 | 19.717 |
| 16.654 | 14.278 |
| 25.621 | 24.025 |
| 19.983 | 21.528 |
| -3.052 | 18.954 |
| 17.625 | -13.488 |
| 16.948 | 27.904 |

| | |
|---|---|
| 6.604 | -5.238 |
| 5.792 | 16.418 |
| 22.259 | 26.565 |
| 15.615 | 17.517 |
| 15.694 | 25.066 |
| 24.758 | 11.017 |
| 10.261 | 15.585 |
| 20.008 | 30.132 |
| 7.293 | 16.060 |
| 9.505 | 29.287 |
| 3.929 | 24.760 |
| 25.496 | 15.176 |
| 27.297 | -12.503 |
| 20.885 | 28.715 |
| -19.571 | 10.791 |
| 11.609 | 27.223 |
| 26.005 | 25.747 |
| 19.325 | 19.146 |
| 26.771 | 12.915 |
| 1.282 | -20.095 |
| 21.874 | 28.458 |
| 3.624 | 26.918 |
| 24.358 | 20.774 |
| 10.661 | 24.753 |
| -0.069 | 11.274 |
| 9.838 | 19.567 |
| 5.059 | 25.716 |
| 7.147 | 21.801 |
| 21.366 | 15.926 |
| 17.616 | 14.104 |
| 27.723 | -25.338 |
| 18.165 | 23.483 |
| 26.162 | 4.477 |
| 14.411 | 17.936 |
| 24.093 | 12.111 |

| | |
|---|---|
| 11.167 | 20.726 |
| 25.183 | -14.991 |
| 13.183 | 24.721 |
| 16.713 | -4.818 |
| 6.059 | 13.115 |
| 15.859 | 25.478 |
| -24.417 | 15.534 |
| 24.731 | 26.871 |
| 15.259 | 14.523 |
| 12.524 | 6.528 |
| 14.089 | -25.616 |
| 3.384 | 20.192 |
| 6.263 | 16.725 |
| 2.674 | 8.402 |
| 7.298 | 24.843 |
| 25.998 | -18.414 |
| 19.031 | 10.893 |
| 9.749 | -0.605 |
| 17.961 | 1.913 |
| -5.139 | 22.794 |
| 11.678 | 20.825 |
| 19.513 | 17.485 |
| 16.919 | 0.205 |
| 18.880 | 12.509 |
| 6.622 | 1.031 |
| 14.309 | 14.983 |
| 4.406 | 9.263 |
| 26.783 | 18.185 |
| 15.353 | 21.355 |
| 28.910 | 19.514 |
| 5.875 | 24.954 |
| -1.507 | 20.981 |
| 18.112 | 18.301 |
| 28.457 | -9.723 |
| -7.401 | 19.665 |

| | |
|---|---|
| 21.615 | 11.264 |
| 19.442 | 21.236 |
| -5.334 | 19.030 |
| 10.177 | -5.344 |
| -14.086 | 25.076 |
| 19.869 | 5.649 |
| 21.505 | 17.715 |
| 22.878 | 6.392 |
| 24.894 | 8.923 |
| 21.985 | 4.363 |
| 18.986 | 25.474 |
| 20.540 | 26.556 |
| 29.424 | 18.242 |
| 14.113 | -7.662 |
| 30.067 | 24.325 |
| 20.339 | 22.509 |
| 15.045 | -0.356 |
| -2.061 | 3.101 |
| 25.487 | 8.820 |
| -1.492 | 11.186 |
| 5.384 | -12.481 |
| 18.513 | 23.032 |
| 21.166 | -1.696 |
| 3.437 | 19.232 |
| 9.718 | -2.472 |
| 26.949 | 9.764 |
| 0.559 | 21.426 |
| 12.897 | 25.808 |
| 17.311 | 14.904 |
| 16.389 | 21.793 |
| 30.477 | 19.561 |
| 29.778 | -19.776 |
| 28.852 | 24.028 |
| 0.881 | 26.746 |
| 16.530 | 17.103 |

| | |
|---|---|
| 26.217 | 11.611 |
| 7.954 | 28.247 |
| 27.942 | 18.072 |
| 27.287 | -12.514 |
| 24.509 | 10.730 |
| 21.892 | -7.393 |
| 29.264 | 9.798 |
| -3.974 | 20.252 |
| -9.939 | 25.639 |
| 10.206 | -25.816 |
| 23.874 | 16.298 |
| 16.459 | 19.302 |
| -3.669 | 12.438 |
| 5.565 | 23.766 |
| 10.970 | 26.844 |
| 28.996 | 24.717 |
| -0.381 | 26.164 |
| 22.005 | 21.182 |
| 21.582 | 17.009 |
| 9.276 | 7.821 |
| -13.004 | 21.892 |
| 7.457 | 18.113 |
| -12.639 | 17.361 |
| 20.092 | 29.704 |
| 14.319 | 28.637 |
| 24.578 | 27.481 |
| 15.259 | 25.981 |
| 23.540 | 24.661 |
| 11.941 | 14.913 |
| 26.860 | 30.089 |
| 18.858 | 10.653 |
| 28.348 | 1.702 |
| 27.556 | 16.199 |
| 23.148 | 30.502 |
| 22.694 | 2.735 |

| | |
|---|---|
| 17.319 | 24.422 |
| -8.097 | 20.403 |
| 13.338 | 30.368 |
| -15.993 | 23.546 |
| 21.327 | 30.739 |
| 28.407 | -9.038 |
| 18.341 | 21.197 |
| -4.061 | 22.547 |
| 30.236 | 16.256 |
| 27.136 | 25.173 |
| 5.939 | 19.518 |
| 16.360 | 15.753 |
| 22.216 | 11.287 |
| 26.106 | 22.480 |
| 2.989 | 16.846 |
| 15.394 | 29.134 |
| 23.428 | 31.084 |
| 18.737 | 23.089 |
| 13.768 | 13.419 |
| 8.751 | 1.128 |
| 22.933 | 24.785 |
| -9.989 | 28.937 |
| 27.627 | 8.506 |
| -12.354 | 26.213 |
| -14.011 | 13.554 |
| -3.680 | 30.326 |
| 15.533 | 10.684 |
| 21.186 | 21.902 |
| -10.882 | 12.402 |
| 19.816 | 27.722 |
| 0.904 | 21.474 |
| 17.643 | 18.367 |
| 17.957 | 1.388 |
| 7.038 | -25.750 |
| -16.662 | 16.454 |

| | |
|---|---|
| 9.748 | -14.616 |
| 2.543 | 11.896 |
| 19.113 | 26.032 |
| 23.241 | 14.692 |
| 13.692 | 24.397 |
| 17.174 | 30.560 |
| 26.062 | 23.287 |
| 17.598 | 22.555 |
| 12.817 | 0.268 |
| 18.410 | 13.270 |
| 9.685 | 8.829 |
| 11.164 | 20.835 |
| 15.950 | 16.926 |
| 24.759 | 27.725 |
| 23.439 | -25.412 |
| 27.128 | 19.087 |
| 8.251 | 24.347 |
| 20.526 | 22.491 |
| 16.845 | 13.900 |
| 11.616 | 26.869 |
| 25.720 | 15.136 |
| 19.570 | -2.207 |
| -20.799 | 29.007 |
| 23.775 | 18.387 |
| 17.764 | 25.768 |
| 16.315 | 19.046 |
| 8.767 | 29.042 |
| 5.249 | 12.186 |
| 26.055 | 14.595 |
| 20.270 | 21.135 |
| 22.665 | 7.096 |
| 23.772 | 24.699 |
| 5.478 | 12.773 |
| 17.840 | -28.641 |
| 13.816 | 25.623 |

| | |
|---|---|
| -10.682 | 22.831 |
| 22.096 | 10.576 |
| 0.124 | 19.496 |
| 18.519 | 20.740 |
| 24.365 | 21.641 |
| 9.572 | 10.142 |
| 26.432 | 13.572 |
| 20.840 | -25.156 |
| 0.758 | 24.928 |
| 12.847 | 28.015 |
| 5.086 | 22.903 |
| 14.528 | 10.831 |
| -26.033 | 20.513 |
| 26.009 | 15.010 |
| 22.793 | 18.786 |
| 8.117 | -19.753 |
| 16.428 | 21.425 |
| -11.208 | 16.496 |
| 25.023 | 26.816 |
| 5.068 | 23.792 |
| 8.457 | 20.511 |
| 8.865 | -1.579 |
| 19.656 | 25.451 |
| 22.058 | 22.336 |
| 11.133 | 15.782 |
| 16.523 | 14.939 |
| 9.139 | 21.318 |
| 0.034 | -14.959 |
| -3.452 | 19.548 |
| 13.889 | 11.352 |
| -7.422 | -9.940 |
| 14.679 | -0.384 |
| 8.672 | 21.885 |
| 6.388 | 10.224 |
| -2.048 | 23.564 |

| | |
|---|---|
| 10.148 | -22.311 |
| -24.192 | -20.559 |
| 15.031 | 25.660 |
| 10.405 | 19.330 |
| 17.704 | 25.061 |
| 19.115 | 24.186 |
| 4.164 | -1.906 |
| 5.501 | -11.009 |
| 25.124 | 12.430 |
| 12.069 | 13.806 |
| -0.228 | 22.587 |
| 22.408 | -1.305 |
| 13.280 | 2.668 |
| -15.082 | 15.712 |
| 17.119 | 10.068 |
| 9.298 | 5.117 |
| 0.161 | 14.226 |
| 15.323 | 3.453 |
| 13.815 | 5.162 |
| 10.164 | -1.369 |
| -6.366 | 19.803 |
| 4.905 | 13.243 |
| 19.195 | 17.696 |
| 13.533 | 21.118 |
| 19.801 | 8.044 |
| 21.849 | -25.532 |
| 16.206 | 1.778 |
| 15.411 | 16.901 |
| -10.037 | 10.068 |
| 13.360 | -9.170 |
| 18.179 | 22.780 |
| 14.025 | 22.638 |
| 9.344 | 11.807 |
| 8.116 | 26.620 |
| 24.403 | 17.379 |

| | |
|---|---|
| 6.182 | 20.778 |
| -1.068 | 14.433 |
| 23.112 | 13.409 |
| -6.878 | 26.071 |
| 17.976 | 10.385 |
| -7.923 | 23.226 |
| 2.131 | 14.334 |
| 12.821 | 14.905 |
| 1.559 | 12.535 |
| 9.926 | 24.829 |
| 13.835 | 27.862 |
| -8.509 | 22.692 |
| 24.861 | 27.382 |
| 7.651 | 8.985 |
| 21.664 | 28.394 |
| 2.166 | 15.097 |
| 23.298 | 17.665 |
| 20.297 | -23.302 |
| 20.643 | 26.159 |
| 15.276 | 21.968 |
| 8.814 | 21.849 |
| 22.580 | 21.165 |
| 15.941 | 23.258 |
| 10.004 | 1.342 |
| 11.662 | 29.199 |
| -4.656 | 23.702 |
| 18.514 | 24.336 |
| -25.151 | 12.850 |
| 4.009 | 13.864 |
| 7.083 | 17.609 |
| 26.517 | -7.234 |
| 20.750 | 26.012 |
| 19.677 | 17.482 |
| 27.404 | 27.037 |
| 3.160 | 14.941 |

| | |
|---|---|
| -2.143 | 29.456 |
| 18.885 | 12.427 |
| 23.300 | 25.826 |
| 25.314 | 27.499 |
| 19.457 | 19.141 |
| 9.545 | 10.785 |
| -3.102 | 19.908 |
| 2.920 | 16.818 |
| 19.761 | 0.523 |
| 26.107 | 25.043 |
| 12.499 | -12.353 |
| 20.575 | 29.974 |
| 30.213 | 21.813 |
| 23.819 | 23.957 |
| 15.342 | -0.563 |
| 28.096 | 8.411 |
| 20.919 | 29.214 |
| 12.353 | 22.542 |
| 5.311 | 18.082 |
| 22.636 | 21.981 |
| 18.362 | 12.165 |
| 9.127 | 10.211 |
| -0.671 | 13.348 |
| 23.533 | 19.603 |
| 3.530 | 31.577 |
| 29.234 | 24.872 |
| 10.173 | 19.873 |
| -9.284 | 16.025 |
| 20.949 | 23.487 |
| 16.176 | 4.263 |
| 28.326 | 3.198 |
| 22.295 | 12.012 |
| 13.485 | 7.049 |
| 18.358 | 24.421 |
| 31.303 | 26.853 |

| | |
|---|---|
| 23.336 | 30.563 |
| 13.358 | 16.334 |
| 29.796 | 14.238 |
| 29.358 | 20.937 |
| -4.266 | 15.118 |
| 22.552 | 18.073 |
| -7.912 | 0.961 |
| 1.301 | -0.043 |
| 19.464 | 23.688 |
| -0.421 | 20.395 |
| 18.002 | 16.582 |
| 28.126 | 15.759 |
| 14.698 | 7.314 |
| 8.771 | 11.665 |
| 5.375 | 13.641 |
| 7.382 | 26.694 |
| 10.317 | 18.001 |
| 17.757 | 14.641 |
| 0.742 | 29.445 |
| -2.480 | 27.353 |
| 19.326 | 23.940 |
| 23.387 | 7.467 |
| 14.993 | 28.197 |
| 24.818 | 10.006 |
| -5.221 | 22.781 |
| 26.935 | 20.914 |
| 7.332 | 28.774 |
| 23.085 | 12.321 |
| 29.391 | 6.924 |
| 8.813 | 9.172 |
| 16.798 | -1.203 |
| 17.806 | 25.814 |
| 12.598 | 18.424 |
| 25.207 | 16.988 |
| 22.063 | 0.181 |

| | |
|---|---|
| -13.885 | 27.335 |
| 23.807 | 13.706 |
| 20.527 | -22.318 |
| 2.062 | 9.890 |
| 5.771 | 13.955 |
| 7.216 | 5.094 |
| 18.253 | 16.291 |
| 17.315 | 15.634 |
| 14.632 | 0.691 |
| -2.309 | 22.138 |
| 21.978 | 12.163 |
| 29.539 | 26.841 |
| 26.537 | 19.578 |
| 21.158 | 8.780 |
| 10.689 | 18.602 |
| 19.989 | -10.613 |
| 12.716 | 10.026 |
| 17.450 | 10.948 |
| 7.891 | 7.210 |
| 11.702 | 21.622 |
| 3.931 | 20.441 |
| 25.723 | 15.103 |
| 19.959 | 22.936 |
| 26.823 | 24.429 |
| 6.907 | 25.458 |
| 21.314 | 1.872 |
| -17.909 | 4.214 |
| 22.944 | 0.972 |
| 22.031 | 18.159 |
| 24.677 | 23.245 |
| -0.136 | 10.988 |
| 3.934 | 8.942 |
| 26.348 | 7.960 |
| -13.680 | 15.990 |
| 20.177 | -0.858 |

| | |
|---|---|
| 21.087 | 12.325 |
| 15.380 | 0.464 |
| 26.114 | 4.638 |
| 16.038 | 3.855 |
| 9.184 | 6.916 |
| 14.124 | 1.094 |
| -2.697 | 21.726 |
| 25.727 | -11.875 |
| 20.173 | -13.239 |
| 21.857 | 13.880 |
| 20.853 | 24.093 |
| 13.380 | 24.825 |
| -5.383 | 16.451 |
| 19.614 | 17.558 |
| 14.008 | 24.030 |
| 23.506 | 14.757 |
| 21.356 | 3.418 |
| 14.828 | 10.505 |
| 22.723 | 21.351 |
| 13.882 | 19.216 |
| -0.251 | 20.619 |
| 19.645 | 16.247 |
| 12.720 | 10.585 |
| 23.825 | -1.140 |
| 25.664 | 9.434 |
| 5.018 | 22.616 |
| -0.783 | 15.401 |
| 11.705 | 14.311 |
| 15.773 | 5.192 |
| -14.147 | -4.915 |
| 8.419 | 11.905 |
| 18.841 | 21.367 |
| -6.623 | 24.109 |
| 13.546 | 15.493 |
| -0.021 | 19.915 |

| | |
|---|---|
| 22.132 | 11.531 |
| 15.524 | 6.431 |
| 12.012 | 19.204 |
| 9.886 | 17.690 |
| 13.225 | 20.671 |
| 17.762 | 26.107 |
| 25.116 | 18.549 |
| 9.558 | 16.714 |
| 21.055 | -16.519 |
| 25.698 | 22.717 |
| 10.400 | 10.541 |
| 26.019 | 23.449 |
| 13.166 | 5.418 |
| 12.451 | 14.666 |
| 5.258 | 7.824 |
| 23.275 | 22.664 |
| 20.753 | 25.548 |
| 17.958 | -2.586 |
| 19.872 | 8.777 |
| -1.490 | 19.548 |
| 0.817 | 4.424 |
| 21.448 | 9.383 |
| 20.381 | 14.475 |
| 16.193 | 15.050 |
| -3.620 | 5.453 |
| 6.281 | 10.786 |
| 27.310 | -7.491 |
| 25.163 | 17.702 |
| 11.238 | 27.032 |
| 16.489 | 20.356 |
| 20.919 | 11.906 |
| 14.112 | 12.904 |
| -1.068 | 1.282 |
| 12.442 | 12.451 |
| 10.031 | 9.086 |

| | |
|---|---|
| 18.904 | 22.711 |
| 7.557 | 4.827 |
| 26.706 | 13.870 |
| 12.041 | 16.521 |
| 23.792 | -6.299 |
| 13.783 | -14.842 |
| 7.445 | 17.769 |
| 10.829 | 8.842 |
| 17.238 | 24.101 |
| 18.694 | 14.135 |
| 28.659 | 9.630 |
| 16.323 | 27.923 |
| -8.878 | 17.560 |
| 20.827 | 19.213 |
| 26.387 | 11.471 |
| -1.359 | 26.352 |
| 27.649 | 11.018 |
| 29.002 | 19.029 |
| 26.779 | 13.355 |
| 23.212 | -9.003 |
| -15.163 | 16.289 |
| 13.337 | 14.923 |
| 0.062 | 5.923 |
| 26.153 | 12.819 |
| 14.189 | -3.707 |
| 22.127 | 28.693 |
| 11.298 | 25.148 |
| 5.272 | 13.291 |
| 10.181 | 18.731 |
| 9.138 | 11.133 |
| 28.970 | 7.484 |
| 18.145 | 6.575 |
| -11.778 | 24.681 |
| -0.875 | 26.076 |
| 13.910 | 9.620 |

| | |
|---|---|
| 13.375 | 22.462 |
| 15.301 | 17.558 |
| 6.691 | 26.561 |
| 29.450 | 14.931 |
| 2.787 | 27.834 |
| -1.193 | 12.128 |
| 20.267 | 23.732 |
| 10.428 | 5.710 |
| 16.902 | 29.155 |
| 5.240 | 10.472 |
| 28.358 | -2.339 |
| -4.938 | 28.685 |
| 17.561 | 8.904 |
| 10.681 | 25.740 |
| 30.779 | 11.275 |
| -1.281 | 0.297 |
| 12.325 | 26.244 |
| -0.168 | 23.128 |
| 14.284 | 25.181 |
| 15.104 | 30.360 |
| 10.176 | 15.615 |
| 18.782 | -0.401 |
| 23.020 | 14.996 |
| 20.969 | 14.793 |
| 27.451 | 15.524 |
| 29.126 | -4.476 |
| 9.290 | 18.774 |
| -6.654 | 19.258 |
| 16.102 | 2.237 |
| -2.264 | 10.033 |
| 22.799 | 17.066 |
| 17.603 | 5.730 |
| 28.703 | 27.868 |
| 24.180 | 17.343 |
| 4.621 | 21.848 |

| | |
|---|---|
| 26.022 | 16.336 |
| 15.462 | -2.867 |
| 9.818 | 12.032 |
| 27.524 | 8.944 |
| 23.376 | 7.393 |
| 21.403 | 13.952 |
| 29.672 | 29.271 |
| 13.399 | 27.647 |
| 24.818 | -6.854 |
| 19.560 | 15.085 |
| 16.991 | 29.463 |
| 25.557 | 11.109 |
| 21.878 | 17.884 |
| 22.571 | 13.260 |
| 18.486 | 28.639 |
| 23.786 | 4.833 |
| 26.481 | 9.581 |
| 6.002 | -10.199 |
| 19.696 | 23.351 |
| 30.097 | 15.454 |
| 25.049 | 6.298 |
| 12.012 | 18.579 |
| 28.533 | 25.564 |
| 25.939 | -3.913 |
| 18.106 | 25.220 |
| 29.102 | 0.882 |
| 17.205 | 15.731 |
| 24.425 | 5.853 |
| 20.419 | 13.672 |
| 17.089 | -5.987 |
| 11.546 | -4.847 |
| 28.087 | 27.242 |
| 9.183 | 10.441 |
| 21.951 | 3.161 |
| -2.405 | 7.703 |

| | |
|---|---|
| 27.180 | 22.646 |
| 23.790 | 13.712 |
| 16.012 | 8.714 |
| 18.523 | 0.224 |
| 26.298 | 27.744 |
| 10.621 | 12.781 |
| 23.731 | 18.764 |
| 20.318 | 6.626 |
| 24.903 | 12.065 |
| 17.854 | 28.488 |
| 22.047 | 13.480 |
| 15.668 | 5.451 |
| 16.617 | 8.157 |
| 25.281 | 10.197 |
| 27.944 | 22.343 |
| 23.904 | 9.408 |
| 20.084 | -0.799 |
| 25.345 | 26.764 |
| -21.314 | 3.208 |
| 15.263 | -6.207 |
| 21.623 | -1.250 |
| 19.662 | 11.011 |
| 10.383 | 9.838 |
| 10.753 | 2.106 |
| 23.558 | 23.683 |
| 24.942 | -4.982 |
| 22.082 | 27.938 |
| 4.481 | 9.049 |
| 12.674 | 16.563 |
| 25.621 | 22.032 |
| 14.858 | 29.045 |
| -22.847 | 18.109 |
| 22.592 | 12.394 |
| 25.702 | 5.068 |
| 23.864 | -20.413 |

| | |
|---|---|
| 24.843 | 0.902 |
| 13.149 | 16.980 |
| 11.771 | 25.599 |
| | 23.703 |
| | -22.363 |
| | -2.881 |
| | -17.122 |
| | 26.913 |
| | 14.812 |
| | 7.432 |
| | 4.913 |
| | -0.362 |
| | -1.364 |
| | 18.026 |
| | -0.356 |
| | 13.818 |
| | 4.174 |
| | 28.324 |
| | 22.695 |
| | 20.799 |
| | 18.363 |
| | 3.511 |
| | 26.627 |
| | 16.442 |
| | 14.330 |
| | 27.625 |
| | 10.766 |
| | 24.796 |
| | -0.352 |
| | 0.326 |
| | 9.536 |
| | 24.131 |
| | 12.621 |
| | 27.086 |
| | 22.661 |

| | |
|---|---|
| | 24.972 |
| | 23.629 |
| | 26.590 |
| | 15.730 |
| | -3.694 |
| | 14.140 |
| | 21.243 |
| | 16.990 |
| | 19.280 |
| | 14.700 |
| | 12.270 |
| | 1.839 |
| | 21.836 |
| | 2.419 |
| | 25.878 |
| | 23.329 |
| | 17.075 |
| | 21.513 |
| | 13.012 |
| | 24.108 |
| | 15.128 |
| | -14.324 |
| | 17.699 |
| | 6.738 |
| | -11.676 |
| | 13.493 |
| | 13.358 |
| | 16.330 |
| | -5.731 |
| | 22.027 |
| | 23.521 |
| | 18.088 |
| | 3.066 |
| | 24.131 |
| | 14.215 |

| | |
|---|---|
| | 7.626 |
| | 14.801 |
| | 5.645 |
| | 24.491 |
| | -0.449 |
| | 14.352 |
| | 10.853 |
| | -6.449 |
| | -18.581 |
| | -25.171 |

FIGURE 2A-ISOAMYL ALCOHOL-STARVED-WORMPOSITIONS

| DMSO | IA D01 | IA D02 | IA D03 | IA D04 | IA D05 | IA D06 | IA D07 | IA D08 | IA D09 | IA D10 | IA D11 | IA D12 | IA D13 |
|---|---|---|---|---|---|---|---|---|---|---|---|---|---|
| -19.166 | -13.322 | 24.337 | 14.157 | 8.951 | -17.667 | 8.244 | 20.895 | -12.499 | 20.140 | 24.406 | 5.914 | 23.331 | 22.184 |
| 14.502 | -11.766 | 1.633 | 5.479 | 24.367 | -4.823 | 19.026 | -5.646 | -12.353 | -22.127 | 23.664 | 14.488 | 19.766 | 18.201 |
| 12.773 | -12.019 | 9.566 | -4.577 | 9.545 | -3.592 | 0.150 | -4.682 | -10.451 | -18.738 | 23.432 | 23.385 | 12.146 | 20.493 |
| -7.103 | -12.541 | -2.787 | -16.315 | -2.725 | -4.511 | 19.050 | 16.505 | -4.761 | -22.924 | 3.350 | 21.270 | 23.620 | 17.859 |
| -11.388 | 8.877 | -17.067 | 23.281 | -1.604 | -0.621 | -20.122 | 1.912 | 18.841 | -7.174 | 5.412 | -1.104 | 21.263 | 14.766 |
| 11.600 | 0.360 | -13.706 | 4.278 | -20.286 | -5.918 | -1.006 | -3.519 | 13.824 | -3.698 | 7.312 | 8.451 | 22.548 | 23.632 |
| 22.252 | -0.089 | -18.698 | 22.001 | -11.804 | -17.087 | -25.254 | 25.967 | 22.324 | 8.420 | 16.046 | 21.870 | 0.427 | 22.926 |
| -24.718 | 13.932 | 0.908 | 2.941 | 25.431 | 21.460 | -9.810 | -0.765 | 6.323 | 7.688 | 26.674 | 0.299 | -13.587 | 18.977 |
| 8.394 | 12.597 | -9.074 | -19.263 | 6.847 | -20.925 | 18.340 | 18.733 | 21.101 | 24.633 | -4.664 | 17.621 | 20.732 | 16.180 |
| 7.777 | -5.715 | -12.342 | 16.699 | -0.044 | -18.132 | -15.983 | 0.864 | 2.825 | -14.310 | 23.837 | -7.866 | -20.861 | 16.834 |
| -16.058 | -25.302 | 19.753 | 14.289 | -5.345 | 20.933 | 18.483 | -16.400 | 13.539 | 20.095 | 19.766 | 20.693 | 8.723 | 10.096 |
| 3.221 | -2.356 | -7.025 | 9.689 | 22.912 | -25.809 | 16.960 | 17.408 | 12.724 | 1.142 | 22.838 | -6.189 | 24.361 | 18.810 |
| -22.610 | 5.440 | 2.996 | 11.415 | -18.449 | 19.250 | -0.764 | 19.941 | 25.440 | -16.095 | 18.925 | 19.875 | -12.308 | 24.481 |
| -23.343 | -7.632 | -25.173 | -3.717 | 0.109 | 13.466 | 5.214 | -10.463 | 12.935 | 17.331 | 27.285 | 6.357 | 22.848 | 26.188 |
| 13.480 | 23.704 | 5.072 | -7.466 | 19.681 | 10.578 | 4.696 | -18.885 | -14.892 | 24.798 | -2.953 | 5.707 | 19.949 | 19.668 |
| -8.228 | -0.427 | -23.524 | 24.149 | -26.198 | -7.829 | 2.135 | -2.582 | 11.234 | -18.471 | 21.040 | 2.712 | -4.330 | 22.540 |
| 19.387 | 4.356 | -6.534 | 25.344 | -10.055 | -19.792 | 20.975 | -10.994 | 7.070 | 0.207 | 21.467 | 24.269 | 25.083 | 13.347 |
| -8.803 | 25.855 | 0.484 | -24.108 | 7.258 | -25.294 | 9.167 | 24.533 | -0.800 | -8.544 | 12.421 | 22.457 | 23.452 | 21.272 |
| 1.553 | -13.454 | 12.745 | -18.361 | -25.676 | -3.583 | 13.691 | 14.098 | -12.341 | 5.884 | 26.540 | -4.631 | -13.013 | 21.807 |
| 21.370 | -16.846 | 3.470 | -21.290 | 25.596 | -4.062 | 0.271 | -13.617 | 24.651 | 21.116 | 19.628 | -7.364 | 24.640 | 18.181 |
| -21.369 | 24.509 | 16.935 | -9.047 | -5.908 | 8.734 | -8.939 | 20.126 | 21.968 | -8.153 | 25.117 | 0.868 | 22.761 | 9.461 |
| 24.131 | -8.077 | -23.288 | -10.450 | 16.807 | -2.406 | 22.297 | 22.195 | 1.194 | -20.146 | 7.714 | -18.379 | 21.569 | -0.024 |
| 6.295 | -20.415 | 3.961 | 24.602 | -1.896 | -15.097 | -18.580 | -24.251 | -23.885 | 24.664 | 25.841 | 12.481 | 3.372 | 25.772 |
| -7.858 | 18.889 | -0.932 | 12.878 | -26.866 | -2.841 | 6.322 | 25.271 | -13.336 | 8.956 | -18.138 | 9.092 | 26.336 | 18.716 |
| -6.343 | -15.470 | 16.565 | 15.696 | -11.881 | -6.194 | -5.276 | -18.108 | -8.127 | -22.120 | -19.359 | -21.641 | 21.206 | 21.710 |
| -15.490 | -23.526 | -1.113 | 6.710 | 20.668 | -11.231 | -7.893 | 19.445 | -10.083 | -24.546 | -20.061 | 17.919 | 2.862 | 17.595 |
| -22.113 | 23.504 | 17.398 | -4.691 | 12.182 | 23.468 | 27.390 | 10.762 | 5.473 | -12.912 | 21.591 | -2.010 | -0.744 | 24.882 |
| 14.226 | -5.778 | 4.501 | 23.498 | -8.120 | -14.135 | -26.994 | -22.436 | 23.848 | 23.504 | 20.830 | -16.053 | 22.822 | 14.422 |
| 15.328 | 4.343 | -11.297 | -23.154 | -11.289 | 16.265 | -12.917 | 18.711 | 21.097 | -1.085 | -21.640 | -5.891 | 25.189 | 19.062 |
| 22.101 | 25.635 | 1.179 | 19.379 | 8.987 | 6.561 | 23.102 | -21.532 | 16.900 | 2.427 | 20.142 | 16.349 | 23.954 | 22.884 |
| 17.633 | -23.202 | -19.901 | -25.596 | 24.743 | -2.374 | 4.528 | -18.241 | 6.831 | -7.817 | -8.745 | 6.684 | 20.740 | 2.362 |
| -3.490 | 13.046 | 10.047 | 7.988 | 21.366 | 4.566 | -1.977 | 9.988 | 9.820 | 23.642 | 17.660 | -12.822 | -13.369 | 25.332 |
| 25.265 | 23.029 | 15.772 | 26.130 | 1.943 | 24.589 | -15.495 | 0.365 | 3.836 | -0.179 | 22.708 | 19.132 | -13.867 | 22.144 |
| -2.834 | -5.378 | -3.740 | -7.621 | -25.766 | -7.745 | 11.964 | -2.047 | 22.290 | -25.014 | 16.822 | 18.392 | -10.133 | 27.516 |
| 8.429 | -9.239 | 19.485 | 25.013 | -15.520 | -21.654 | 27.733 | 11.654 | -4.441 | -25.894 | 18.973 | 26.160 | 26.557 | 26.649 |

| | | | | | | | | | | | | | |
|---|---|---|---|---|---|---|---|---|---|---|---|---|---|
| 23.071 | 16.758 | -4.703 | 13.544 | -6.925 | -9.300 | 5.068 | -13.253 | -12.052 | -21.399 | -5.373 | 3.598 | -10.449 | 17.989 |
| 1.862 | 18.982 | -12.185 | -12.732 | -1.981 | -26.686 | 25.601 | 21.511 | 26.237 | 14.719 | 20.462 | 25.882 | 24.647 | 19.870 |
| -10.683 | 19.727 | -10.958 | -23.185 | -17.711 | -17.308 | 18.185 | 25.788 | 16.936 | -26.773 | 5.997 | -27.091 | 22.288 | 15.900 |
| -3.638 | -2.490 | -26.475 | 20.963 | -10.755 | 24.005 | -0.507 | -0.390 | -11.435 | 19.292 | 26.985 | 7.576 | 23.440 | -0.491 |
| 0.698 | 15.174 | -16.256 | 11.062 | -13.511 | -7.732 | 6.901 | 14.212 | 10.339 | 6.703 | -10.636 | 2.189 | -0.468 | 3.752 |
| -18.658 | -15.316 | 24.089 | -11.614 | -5.338 | 10.237 | -4.929 | 15.535 | -1.669 | 26.667 | 27.739 | 0.802 | -6.124 | 24.556 |
| -19.672 | 13.522 | 5.815 | -16.223 | 3.455 | -18.383 | 24.094 | -1.073 | 12.834 | 22.198 | 24.372 | 18.453 | 7.357 | 23.149 |
| 8.010 | -24.861 | 17.987 | 9.519 | -6.141 | -0.453 | -27.440 | -22.447 | -22.773 | -1.291 | 25.178 | 21.851 | -6.826 | 24.302 |
| 22.141 | 23.462 | -3.079 | -7.732 | -21.067 | -22.643 | 21.350 | -25.492 | -19.780 | 15.626 | -16.091 | 21.231 | 25.748 | 15.620 |
| -23.892 | -7.670 | 17.468 | 7.263 | -1.165 | -7.598 | -0.540 | 22.314 | 21.550 | 8.143 | 23.099 | 24.392 | 16.195 | 19.322 |
| 18.070 | 24.977 | 21.636 | -17.774 | -24.639 | 11.643 | -12.379 | 13.678 | -24.673 | 25.086 | 23.563 | 0.371 | 27.492 | 6.796 |
| -8.830 | 10.401 | 13.074 | 3.415 | -20.119 | 5.566 | 8.422 | -12.250 | 23.940 | 23.083 | 18.081 | 1.562 | 10.425 | 17.105 |
| -1.095 | -23.788 | 11.925 | 12.653 | -26.213 | -20.958 | -9.072 | 26.183 | -2.196 | 19.560 | 27.429 | 14.936 | 17.611 | 20.949 |
| -1.795 | 5.109 | -21.695 | -26.821 | -27.905 | -15.520 | -15.998 | -20.320 | -3.938 | -3.272 | 26.145 | 12.625 | 13.819 | 23.579 |
| -5.422 | -25.700 | -27.721 | 3.972 | 21.938 | -10.566 | 16.470 | -11.988 | 11.760 | 4.001 | 26.479 | 23.985 | 18.848 | 17.282 |
| -26.112 | 21.792 | -13.528 | 19.229 | -22.698 | -11.151 | -8.329 | 6.580 | 4.653 | 12.886 | 22.073 | 4.897 | -6.222 | 28.388 |
| -25.693 | -5.198 | -19.353 | 6.298 | -10.211 | -19.578 | 8.164 | 17.784 | 20.553 | 25.373 | 29.157 | 7.142 | 24.236 | -2.319 |
| 22.934 | 25.694 | -12.989 | -9.082 | -26.484 | -25.219 | 12.268 | 11.285 | -7.064 | 26.557 | 12.791 | -0.791 | 22.960 | 18.062 |
| 20.991 | -24.915 | 18.984 | -20.197 | 6.602 | -6.025 | -20.412 | 19.426 | 23.201 | 21.459 | 21.049 | 15.296 | 27.372 | 20.246 |
| -10.223 | -3.482 | -11.154 | -8.119 | -2.748 | -13.763 | 1.298 | 10.534 | 26.635 | -20.640 | 15.190 | 6.687 | 26.200 | 14.709 |
| -21.293 | 13.509 | 9.595 | 8.404 | -28.130 | -24.623 | 1.816 | 27.302 | 9.750 | 6.844 | 17.234 | -26.728 | 28.921 | 25.325 |
| -18.092 | -19.884 | 3.920 | -8.865 | 15.147 | -9.528 | -24.934 | -25.766 | 5.114 | 26.471 | 29.462 | -6.909 | 8.804 | 26.430 |
| 24.164 | -21.984 | 23.751 | -22.286 | -4.879 | -2.333 | 13.231 | 22.496 | -0.316 | 23.345 | 23.996 | 13.346 | -13.149 | 22.927 |
| 4.151 | 27.471 | -28.957 | -26.941 | -7.810 | -14.741 | 17.999 | 5.600 | -9.456 | 0.924 | 21.929 | 25.325 | 19.903 | 24.123 |
| -13.514 | -25.879 | 1.291 | 27.092 | 28.778 | -12.238 | 10.393 | 23.147 | -26.342 | -26.070 | 22.732 | -11.575 | 1.531 | 19.782 |
| 4.734 | 23.851 | -6.765 | -15.405 | 0.831 | -11.065 | 0.659 | -3.338 | 17.841 | 19.514 | 27.726 | -27.829 | -19.380 | 18.102 |
| 3.095 | 3.269 | -20.557 | -20.444 | -18.169 | 25.343 | 9.403 | 24.300 | 19.218 | -16.357 | -13.867 | 19.798 | 15.503 | 20.691 |
| -12.307 | 26.122 | -16.560 | 13.338 | 23.601 | -2.747 | -25.500 | 2.099 | -13.457 | 8.920 | 25.461 | 7.704 | 16.833 | 14.986 |
| -6.019 | 20.688 | 29.712 | 16.952 | 8.117 | 18.613 | 17.016 | 20.373 | 4.502 | -18.500 | -6.001 | 24.314 | 6.278 | 7.060 |
| -6.640 | 8.408 | -12.964 | -22.406 | 28.015 | -23.494 | -20.947 | 18.766 | 12.109 | 23.460 | -8.885 | 0.136 | -1.597 | 18.738 |
| -23.609 | 5.044 | -21.315 | -11.482 | 13.661 | 6.708 | 25.194 | -26.535 | 21.738 | 21.082 | 29.584 | 26.662 | -22.747 | 16.642 |
| -1.256 | -26.120 | -23.420 | 20.453 | -25.530 | 21.727 | 26.889 | 10.791 | 12.631 | 25.576 | 17.033 | 3.915 | 3.012 | 24.280 |
| 16.680 | -4.396 | 11.753 | 23.558 | 4.574 | -19.528 | -24.217 | 15.224 | 3.689 | 22.387 | 24.864 | 11.333 | -24.014 | 15.969 |
| 22.362 | 28.273 | 0.707 | -9.694 | 19.612 | -9.221 | 17.171 | -21.492 | -26.737 | 11.905 | 23.827 | 25.516 | 25.018 | -25.843 |
| 0.400 | -22.349 | -6.631 | 25.148 | -13.848 | 2.617 | -13.781 | -22.479 | 9.997 | -9.851 | 22.506 | 27.986 | 20.512 | 19.628 |
| 21.111 | 12.798 | 15.568 | 22.004 | 26.839 | 12.344 | 23.891 | -19.673 | -1.668 | -13.238 | 27.285 | 19.437 | -13.841 | 25.554 |

| | | | | | | | | | | | | | |
|---|---|---|---|---|---|---|---|---|---|---|---|---|---|
| -20.449 | 4.763 | 15.892 | -14.979 | 26.257 | -6.700 | -21.247 | 17.535 | 7.298 | 23.796 | 25.716 | 16.355 | 14.583 | 5.499 |
| 7.138 | -21.931 | -8.399 | -24.151 | 17.621 | -10.209 | -0.440 | -14.598 | -25.418 | -13.723 | 28.683 | -12.120 | 10.860 | 14.204 |
| 8.006 | 25.135 | 20.924 | -23.259 | -28.310 | -15.215 | 27.517 | -18.467 | 14.262 | -8.126 | -11.138 | -25.868 | 24.230 | 20.983 |
| -21.382 | -29.061 | -9.642 | 13.573 | -21.798 | -27.049 | 21.346 | 20.579 | 7.834 | 7.082 | 16.573 | 22.744 | -22.316 | 16.181 |
| -23.900 | 27.075 | 24.900 | 0.503 | 23.673 | -12.853 | -3.388 | -13.163 | -5.901 | 22.680 | 19.268 | 8.692 | 28.460 | 24.933 |
| 20.646 | 3.283 | -16.864 | -10.261 | -21.001 | 14.367 | 0.290 | 8.439 | 20.498 | 19.275 | 10.165 | 7.299 | -7.126 | 18.618 |
| 27.719 | 19.270 | -0.200 | 11.487 | 15.964 | 20.616 | -8.218 | 4.358 | 28.140 | 26.132 | 26.121 | 14.495 | 14.089 | 20.004 |
| 19.759 | 22.381 | -13.877 | 17.253 | 7.997 | 10.456 | -20.027 | -23.691 | 9.287 | 18.601 | 15.571 | -11.617 | 27.971 | 13.079 |
| -7.599 | -7.864 | -22.611 | -19.184 | 10.539 | 9.356 | 16.114 | 14.499 | 5.683 | 20.647 | 29.082 | 23.747 | 19.082 | 20.636 |
| -19.983 | 5.803 | -2.485 | -26.228 | 29.083 | -17.303 | -11.932 | 13.322 | 21.251 | 25.457 | 27.236 | 1.718 | 20.662 | 15.105 |
| 15.150 | -2.909 | 14.849 | 21.233 | -4.952 | -23.320 | -0.071 | 12.748 | 24.451 | 26.711 | 27.614 | -1.663 | 28.476 | 28.006 |
| 14.513 | 26.407 | -27.588 | -21.218 | 26.526 | -8.757 | 25.555 | 24.354 | -27.289 | 22.469 | -17.600 | 26.984 | -5.220 | 3.640 |
| 25.326 | 20.863 | 22.800 | 10.973 | 25.769 | -15.986 | -27.019 | -15.918 | -13.589 | 25.029 | 21.056 | 21.465 | -10.155 | 2.890 |
| 18.824 | 23.337 | -6.425 | 18.793 | 21.789 | -22.056 | -9.836 | 5.175 | 27.383 | -17.564 | -7.724 | 16.779 | 5.273 | 23.997 |
| 18.096 | -27.632 | 29.881 | 9.731 | -29.278 | -5.186 | -3.751 | -23.367 | -16.526 | -23.625 | 18.175 | 25.572 | 19.526 | 26.266 |
| -18.855 | 26.042 | -9.644 | -25.033 | 1.726 | -5.840 | -14.018 | -0.859 | -13.178 | 12.281 | 26.089 | 24.307 | 22.574 | 25.449 |
| -3.645 | -13.008 | 23.249 | 8.649 | 18.186 | 24.322 | 9.165 | 25.677 | -12.109 | 30.117 | 22.729 | -10.724 | 27.128 | 28.514 |
| 0.410 | 20.544 | 6.703 | -3.107 | 19.289 | -14.042 | -4.341 | -9.268 | 7.481 | 29.018 | 23.378 | 19.697 | -8.439 | 17.541 |
| -0.494 | 6.052 | 17.430 | 0.692 | 7.029 | 21.319 | 12.838 | 2.019 | 26.234 | 22.977 | 13.223 | 6.867 | 25.285 | 18.946 |
| -5.237 | -1.492 | -17.819 | -23.589 | -30.515 | 15.661 | -14.389 | -4.525 | 8.617 | 20.231 | 12.306 | -6.476 | 26.021 | 11.467 |
| -18.305 | 25.536 | -19.880 | 7.498 | -25.564 | -1.435 | 20.473 | -22.193 | 6.373 | 26.074 | 25.666 | -7.130 | 12.778 | 0.327 |
| -24.333 | -8.268 | 10.279 | 27.584 | 23.015 | -2.388 | 13.370 | 12.122 | 23.645 | 21.791 | 27.339 | 21.882 | 23.082 | 14.773 |
| -23.251 | 27.122 | 20.513 | 24.872 | -9.278 | -2.682 | -16.365 | 24.979 | 3.559 | 18.605 | -28.220 | 24.797 | 10.947 | 25.395 |
| 17.187 | 21.966 | -13.322 | -6.415 | 14.336 | 15.932 | 16.166 | -0.228 | 10.073 | -25.658 | 29.147 | 10.113 | 21.660 | 21.516 |
| 5.789 | 22.854 | 5.394 | -5.234 | -31.431 | -22.158 | -3.701 | 23.177 | 15.222 | -17.660 | 22.068 | 16.131 | 16.380 | 14.487 |
| 4.701 | 23.705 | 24.806 | -25.164 | 15.644 | -18.066 | 21.268 | -6.279 | -16.024 | -29.635 | 19.903 | 11.848 | 30.166 | 18.589 |
| -12.825 | -27.693 | 13.867 | -27.484 | 6.872 | -21.115 | 17.439 | 20.828 | 19.737 | -10.914 | 25.130 | -2.351 | -15.375 | 19.503 |
| 24.730 | 17.577 | 26.256 | -3.525 | 30.756 | 17.124 | -18.259 | 6.258 | -19.289 | -25.385 | 18.745 | 21.413 | -9.136 | 21.224 |
| 4.103 | 16.585 | 22.474 | -1.621 | 29.566 | 21.616 | -2.295 | -7.347 | -3.321 | 23.130 | 16.489 | 3.809 | 4.684 | 19.173 |
| 0.926 | -18.954 | -13.853 | -24.354 | 24.548 | -15.951 | -7.088 | 20.303 | 22.966 | -20.077 | 26.240 | -3.432 | 19.504 | 15.985 |
| 14.018 | 17.852 | -9.777 | -15.219 | -27.634 | -9.866 | -23.464 | -1.109 | 26.878 | -24.652 | -5.446 | 23.668 | 27.326 | 14.905 |
| 12.703 | 22.876 | -4.747 | -23.762 | 18.836 | -23.376 | 1.061 | -27.492 | 13.568 | -12.343 | 25.323 | 20.550 | 7.670 | 25.788 |
| 7.443 | 25.923 | 15.274 | 23.577 | 20.443 | 5.755 | -4.725 | -24.441 | -25.705 | 27.043 | 21.373 | -17.705 | 10.112 | 24.930 |
| 27.860 | 4.623 | -27.874 | 13.688 | -10.470 | -23.682 | -12.943 | 3.979 | -15.883 | 21.539 | 23.531 | 20.475 | -3.634 | 5.391 |
| 2.570 | -26.113 | -15.213 | -20.912 | -25.327 | -25.124 | 7.834 | -26.588 | 25.323 | 3.908 | 20.405 | 17.086 | 1.141 | 29.428 |
| 0.232 | -21.881 | -9.042 | 16.105 | 6.050 | -3.251 | 23.123 | -25.176 | -1.228 | 23.756 | -16.217 | 10.787 | -19.283 | 30.970 |

| | | | | | | | | | | | | | |
|---|---|---|---|---|---|---|---|---|---|---|---|---|---|
| 22.605 | -10.660 | -4.185 | 4.682 | -6.635 | -12.836 | -17.237 | 21.868 | 28.629 | 9.960 | 26.536 | 7.117 | 23.810 | 20.834 |
| 15.919 | 21.340 | -1.581 | -10.355 | 23.809 | -5.065 | -5.301 | 9.366 | 12.665 | -19.945 | -31.094 | 22.049 | 26.552 | 7.239 |
| -17.668 | 25.740 | -10.843 | 1.312 | 28.072 | -7.731 | 18.894 | -7.901 | 9.339 | -25.651 | 18.899 | 8.423 | 9.045 | 25.656 |
| -6.701 | 29.447 | 4.795 | -0.908 | -30.203 | -18.085 | -13.781 | 26.131 | -22.918 | -27.461 | 23.124 | -19.062 | 17.023 | 11.865 |
| -26.636 | 23.013 | 21.794 | 3.338 | 19.289 | -8.920 | -3.711 | 19.520 | -24.091 | 21.893 | 27.246 | 16.496 | -8.265 | 19.588 |
| -21.283 | 21.833 | -18.376 | -17.540 | -15.287 | -20.251 | -27.561 | -11.817 | -17.384 | 20.163 | 29.127 | 15.762 | 18.973 | 29.154 |
| -25.448 | -24.566 | -9.492 | -25.699 | 21.761 | -2.665 | -29.768 | -6.302 | -19.594 | -16.361 | 18.969 | 13.588 | 22.481 | 28.790 |
| 16.390 | 31.203 | -17.654 | -12.460 | -28.851 | -5.994 | 26.124 | -21.563 | 29.445 | 27.296 | -24.937 | 0.227 | 21.038 | 19.156 |
| 12.053 | 19.153 | -24.599 | 26.225 | -6.450 | -21.579 | 12.362 | -8.645 | 14.995 | -12.389 | 18.300 | 22.101 | 15.897 | 25.219 |
| 18.795 | 19.726 | 17.057 | 8.143 | -15.804 | -18.919 | 27.479 | -16.646 | 20.721 | -28.254 | -7.338 | 16.271 | -16.841 | 21.716 |
| -5.101 | 11.249 | 31.997 | -26.498 | -13.195 | 4.307 | 14.626 | -3.751 | 21.129 | -21.982 | -6.737 | 11.500 | 29.830 | 30.776 |
| -13.546 | 6.869 | -11.335 | -3.801 | 14.672 | -1.417 | -2.681 | -14.279 | -0.036 | 25.212 | 21.956 | 9.647 | 19.819 | -31.985 |
| -0.065 | -4.420 | 23.764 | 24.581 | -25.335 | -15.238 | 20.340 | 13.243 | 26.176 | -16.142 | -4.412 | -5.614 | -27.283 | 24.103 |
| -19.328 | -14.505 | -2.588 | -0.649 | 25.852 | -12.643 | -2.981 | 3.061 | 5.982 | -17.411 | 14.118 | 23.531 | -20.051 | 27.874 |
| -27.332 | 27.341 | -9.529 | 4.574 | 15.828 | 14.072 | -5.819 | 23.590 | 7.293 | 26.780 | 23.063 | 23.781 | -13.126 | 24.603 |
| 18.096 | 31.232 | 30.200 | -27.119 | -8.916 | 1.582 | -0.609 | -1.421 | 2.672 | 30.643 | 11.638 | 6.317 | 12.079 | 31.440 |
| 19.348 | 28.158 | -12.083 | 16.994 | -7.588 | -22.352 | 8.324 | 21.679 | -3.768 | -24.254 | 27.326 | 23.137 | 17.318 | 19.883 |
| 2.014 | 26.561 | -7.638 | -23.632 | -22.956 | -16.197 | 21.068 | -10.020 | -11.865 | 24.681 | 12.841 | 7.533 | 18.270 | 18.482 |
| 20.489 | -6.940 | 20.391 | -12.350 | -25.971 | 10.876 | 15.178 | -13.011 | 20.056 | 20.725 | 26.247 | 19.241 | -21.683 | 30.619 |
| 13.288 | 19.246 | 6.695 | -20.053 | -18.884 | -3.643 | -19.213 | -9.660 | 22.527 | -16.159 | 19.205 | 8.470 | 8.396 | 15.189 |
| -0.654 | 12.099 | -3.354 | -20.897 | -28.067 | 0.104 | -6.824 | -27.821 | 23.640 | 29.341 | 21.685 | 24.182 | 25.035 | 27.668 |
| 14.857 | 15.189 | -9.907 | -25.024 | 20.206 | -9.213 | -27.808 | 15.286 | 18.174 | -18.282 | 20.052 | -4.168 | 23.548 | 18.901 |
| 12.720 | -20.690 | -29.416 | -8.428 | -14.604 | -23.812 | -15.932 | -8.966 | 11.674 | 22.736 | 27.716 | -28.932 | -4.538 | 4.129 |
| 1.101 | 26.984 | -18.593 | 27.372 | -24.055 | -30.114 | -28.439 | -19.145 | 8.064 | 17.890 | -4.837 | -14.080 | 28.372 | 21.170 |
| 24.105 | -27.955 | -3.117 | 12.577 | 13.895 | 16.992 | -17.162 | -0.809 | 3.819 | 28.075 | 24.640 | 25.109 | -16.309 | -31.569 |
| -15.298 | 6.681 | -23.185 | -26.464 | -18.233 | -16.427 | -26.390 | 14.073 | 27.607 | -7.459 | 3.802 | 19.718 | 1.085 | 27.467 |
| 20.780 | 9.720 | 30.607 | -11.634 | -19.289 | -14.629 | -1.829 | 2.109 | 16.410 | 23.996 | -3.903 | 12.987 | -23.623 | 24.252 |
| 2.518 | 23.553 | 16.711 | 2.039 | 15.718 | 1.432 | -2.921 | 1.615 | 21.717 | 19.371 | -25.073 | 27.477 | 10.384 | 31.750 |
| -1.230 | 3.217 | 28.537 | -5.029 | -25.453 | -17.530 | -3.714 | 18.041 | 12.939 | -22.818 | 22.907 | 26.546 | -29.000 | 28.155 |
| -2.614 | -4.395 | 11.899 | -26.745 | -18.147 | 4.025 | -8.540 | 9.063 | -3.645 | -5.705 | 14.013 | 9.488 | 0.572 | -6.719 |
| -25.278 | 26.723 | -21.872 | 9.165 | 3.033 | -23.172 | -3.267 | -11.225 | -27.931 | -21.465 | 29.163 | -15.400 | -3.539 | 19.365 |
| 19.956 | -18.116 | 17.919 | -28.103 | -27.347 | -23.392 | 16.001 | -0.617 | -21.859 | -29.375 | 24.204 | -18.900 | 22.312 | 22.954 |
| 26.747 | -4.874 | -15.982 | -9.373 | 12.653 | -0.958 | -1.497 | -19.831 | -12.322 | -9.515 | 21.911 | -27.034 | -8.863 | 27.300 |
| 0.210 | 23.257 | 11.644 | -24.482 | -24.004 | -19.134 | -13.284 | -3.635 | 29.980 | -2.686 | 17.596 | 20.935 | 11.985 | 20.048 |
| 29.185 | 24.039 | -8.945 | 25.905 | 14.903 | 9.912 | 5.055 | -28.895 | 25.558 | 19.872 | 25.168 | 20.686 | 26.205 | 14.755 |
| -22.949 | -25.140 | -11.286 | 2.405 | 4.585 | -4.470 | 16.391 | 17.045 | -24.179 | -20.934 | 22.684 | -0.261 | -6.422 | 19.328 |

| | | | | | | | | | | | | | |
|---|---|---|---|---|---|---|---|---|---|---|---|---|---|
| 16.554 | 27.268 | -6.612 | -7.080 | 26.515 | -17.523 | 28.249 | -2.861 | -2.417 | 20.412 | 24.030 | 28.330 | 15.138 | 9.605 |
| -17.537 | 2.407 | 20.128 | -15.678 | 19.361 | -22.073 | -2.802 | 26.419 | -6.364 | -16.971 | -4.983 | -4.671 | 2.736 | 25.355 |
| -3.702 | -10.014 | 25.654 | -29.168 | 21.595 | 13.851 | -2.323 | -2.178 | -7.937 | -18.887 | 19.147 | 17.152 | -0.879 | 21.326 |
| -19.120 | -28.948 | 16.536 | 27.888 | 23.090 | -1.209 | 0.291 | 1.323 | -24.767 | 24.373 | 20.223 | 27.801 | 4.413 | 26.018 |
| 19.040 | -27.463 | 11.222 | 20.191 | 2.628 | -22.598 | -8.830 | 28.382 | -5.686 | 6.759 | 29.224 | 17.606 | 0.089 | 15.409 |
| -28.591 | 9.775 | -21.673 | -30.275 | -8.220 | 12.527 | -21.288 | -5.950 | 23.891 | -8.454 | 26.990 | 29.924 | -27.178 | 23.950 |
| 11.795 | -12.661 | 23.284 | 22.992 | 24.061 | -15.064 | -31.870 | 18.288 | -26.371 | 13.594 | 21.941 | 12.466 | 21.849 | 25.004 |
| -5.591 | 17.662 | -8.887 | -25.526 | 19.791 | -22.976 | -16.322 | 17.788 | -13.606 | 23.527 | 1.837 | 8.190 | -7.039 | 15.916 |
| -20.884 | 13.421 | -16.896 | -18.021 | 18.755 | -5.336 | 20.540 | 3.279 | 1.642 | -26.280 | 15.053 | 13.766 | -9.085 | 26.293 |
| -21.699 | 21.922 | -14.235 | -20.183 | -10.898 | -19.665 | -1.407 | 14.563 | 28.752 | -18.658 | -0.041 | 20.174 | -20.807 | 9.418 |
| 25.205 | -17.996 | -5.161 | 19.351 | 22.419 | -3.523 | 6.459 | 25.027 | -19.586 | -21.996 | 18.537 | 18.452 | -25.525 | 22.118 |
| 21.111 | 25.786 | 2.411 | 21.168 | 17.088 | 31.317 | 4.178 | 15.353 | -8.078 | 31.531 | 20.865 | 15.538 | 21.873 | 25.394 |
| -23.731 | 8.740 | -22.179 | 24.358 | 3.469 | -9.044 | -23.964 | -9.186 | -16.083 | 20.801 | 27.495 | 2.086 | 3.310 | 17.367 |
| -29.164 | 28.779 | -24.030 | -2.342 | 14.594 | 15.490 | -13.140 | -11.710 | 20.346 | -19.062 | 23.337 | -6.528 | -5.402 | 30.805 |
| -17.910 | 5.975 | -20.712 | 14.487 | 8.450 | 12.680 | -2.795 | -18.809 | -5.018 | 25.445 | 24.722 | 23.099 | 29.339 | 16.031 |
| 8.152 | -15.715 | -30.220 | 12.553 | -7.975 | -21.163 | -0.229 | -23.620 | 14.024 | 23.923 | 24.101 | 22.433 | 10.723 | 20.014 |
| 1.829 | -6.756 | -15.516 | 30.374 | 24.158 | -21.510 | 28.932 | -24.844 | 25.483 | 0.772 | 17.795 | 21.172 | -9.461 | 27.068 |
| -0.602 | 25.010 | 24.831 | -29.963 | -6.766 | -25.351 | -21.430 | 0.897 | 20.012 | -16.879 | 13.402 | 2.687 | 22.504 | 24.675 |
| 4.695 | 17.950 | -9.536 | -21.071 | 1.373 | -21.898 | -8.843 | -20.902 | -18.790 | -0.689 | 25.892 | 1.265 | -5.349 | 13.777 |
| 27.846 | -23.927 | 0.512 | -17.804 | 19.325 | -11.390 | 11.566 | -26.932 | -27.194 | -4.357 | 23.634 | -6.111 | 24.427 | 29.346 |
| 0.515 | 21.846 | -25.589 | 3.947 | 12.318 | -8.377 | -15.667 | 22.261 | 1.171 | 21.428 | 8.651 | 26.330 | -0.719 | 22.682 |
| -2.003 | -6.071 | 12.955 | 10.295 | -8.799 | -23.955 | -0.545 | 4.609 | 0.013 | 18.665 | 4.068 | 24.256 | 2.873 | 28.720 |
| 14.433 | -5.028 | 5.278 | -29.349 | 17.641 | 26.484 | 17.625 | -3.495 | -30.452 | 9.405 | 25.904 | 19.016 | -23.311 | 23.264 |
| -19.735 | 8.576 | 6.002 | -25.980 | -0.785 | 20.929 | 3.220 | -19.417 | 7.686 | 30.743 | 25.289 | 12.769 | 28.411 | 26.557 |
| -4.154 | -5.558 | -11.960 | -12.134 | 22.678 | -12.489 | -2.825 | -28.484 | 15.056 | -9.229 | 27.385 | 28.777 | 12.196 | 18.691 |
| 3.707 | 28.001 | -16.146 | -7.515 | -7.945 | -30.111 | 2.803 | 17.908 | 18.244 | 20.010 | 20.589 | -23.291 | -27.586 | 24.181 |
| -26.008 | 22.554 | 1.105 | -24.080 | -22.819 | 23.724 | 2.449 | -4.881 | 5.899 | 24.855 | -18.877 | 10.055 | 18.828 | 20.636 |
| 26.887 | -25.195 | -17.132 | 18.271 | 1.936 | 5.842 | -11.723 | 29.078 | -3.415 | 23.154 | 12.591 | 26.346 | 26.036 | 17.352 |
| 0.899 | -23.535 | -14.808 | 16.686 | 22.158 | -4.479 | 25.412 | -19.158 | 9.699 | -3.400 | -2.773 | 18.441 | 24.239 | 12.151 |
| -17.833 | -16.095 | 14.376 | -19.759 | 0.906 | -22.948 | -10.611 | 17.393 | -26.105 | 13.378 | 27.567 | 29.395 | -16.445 | 29.944 |
| 28.485 | -8.324 | -11.658 | -19.102 | 4.929 | -12.913 | -6.344 | 9.106 | -10.077 | -18.772 | 25.452 | 24.257 | 3.974 | 25.716 |
| 12.074 | 2.120 | -9.758 | 20.179 | 15.720 | -24.255 | -1.982 | 14.348 | 29.656 | 7.061 | 22.999 | 16.945 | 13.950 | 22.190 |
| 4.005 | -11.639 | -21.820 | -8.072 | 10.673 | 23.218 | 2.058 | -11.127 | 28.711 | 11.276 | 12.140 | 4.251 | 27.623 | 28.815 |
| -1.517 | -2.985 | -20.126 | 12.551 | -23.295 | -8.416 | -27.489 | -16.358 | 7.981 | -0.051 | 26.676 | 21.820 | 0.221 | 29.522 |
| 1.594 | -17.052 | 23.512 | -6.663 | 25.315 | -2.664 | -18.667 | -5.841 | -13.737 | 27.341 | 20.251 | 16.567 | 23.784 | 16.199 |
| -27.230 | 21.865 | 10.096 | -30.671 | 24.173 | -25.054 | 10.869 | 16.433 | 3.646 | -23.035 | 15.678 | 2.493 | 30.618 | 28.178 |

| | | | | | | | | | | | | | |
|---|---|---|---|---|---|---|---|---|---|---|---|---|---|
| 23.980 | 8.854 | 6.491 | 28.702 | 6.130 | -26.278 | -3.370 | 16.866 | 21.466 | 19.193 | 1.681 | 24.566 | 21.969 | 18.123 |
| 5.378 | 6.816 | 0.510 | 2.351 | 13.403 | -9.342 | -4.538 | -12.983 | 15.985 | -23.755 | 19.652 | 20.280 | 15.964 | 23.172 |
| -21.248 | 17.813 | -2.216 | -20.860 | -19.076 | -15.918 | -5.749 | 10.089 | 18.278 | 22.387 | 20.926 | 19.122 | 14.975 | 26.821 |
| 27.568 | -24.133 | -5.187 | -15.363 | 9.451 | -19.438 | -9.164 | -23.219 | 5.871 | -12.814 | 9.408 | 13.409 | -4.414 | 27.887 |
| 0.444 | 1.887 | 18.500 | 29.694 | 14.126 | 4.799 | -21.539 | 9.338 | 0.829 | 10.617 | 11.292 | 27.126 | 16.456 | 2.609 |
| -2.517 | -6.093 | -6.308 | -21.856 | -0.614 | -7.010 | 18.817 | 20.702 | 26.214 | -28.300 | -4.820 | 6.721 | -20.112 | 13.551 |
| 29.324 | 8.741 | -23.261 | 11.505 | -3.541 | -17.307 | -7.164 | 28.722 | -6.925 | 15.960 | 8.135 | 7.272 | 2.241 | 19.242 |
| -12.245 | -14.116 | -29.727 | -2.719 | 6.670 | -3.037 | -16.080 | 19.156 | 31.161 | 7.141 | 22.999 | -25.549 | 25.142 | 26.090 |
| 25.420 | 0.931 | -2.528 | -22.252 | 11.995 | -16.656 | 25.533 | -28.867 | -21.880 | 24.879 | 22.125 | 10.516 | -3.802 | -4.308 |
| -0.738 | -2.665 | -16.558 | -10.940 | 23.579 | -23.127 | 17.340 | 29.831 | 13.649 | -25.507 | 19.226 | 10.228 | 14.603 | 15.270 |
| 26.886 | -3.047 | -3.723 | 5.090 | 18.672 | -0.957 | 22.790 | -1.782 | -18.828 | 21.859 | 2.321 | 3.195 | -4.569 | 23.117 |
| -29.766 | -24.743 | -4.225 | -25.714 | -26.903 | -22.319 | 0.120 | 12.452 | -30.260 | -17.818 | 23.126 | 21.368 | 15.999 | 22.258 |
| -3.824 | 18.605 | 12.702 | 13.094 | 18.139 | 24.538 | 13.455 | 13.120 | -5.148 | -27.424 | 21.152 | -9.764 | -0.962 | 21.019 |
| -0.025 | 13.698 | -5.574 | 3.753 | 21.520 | 20.711 | -7.173 | -29.830 | -9.849 | -16.429 | 17.570 | 30.590 | 13.129 | 13.215 |
| -28.920 | -17.418 | 25.400 | -9.721 | 3.430 | -21.937 | 3.080 | -1.147 | -1.074 | -7.478 | -24.409 | 25.338 | 18.020 | 27.817 |
| -25.548 | 20.165 | -3.400 | -29.938 | 8.229 | -15.369 | 1.265 | 23.008 | 12.365 | -20.199 | 19.507 | 20.032 | 27.400 | 21.284 |
| -18.203 | 21.935 | -5.191 | -8.827 | -8.217 | -11.822 | -12.395 | 5.883 | 1.928 | 16.965 | 0.447 | 21.136 | 20.473 | 25.907 |
| -7.334 | 10.381 | -7.587 | 19.164 | -18.511 | -24.458 | -6.108 | -11.775 | -18.200 | 2.599 | 26.442 | -7.644 | 22.377 | -2.748 |
| 6.100 | 13.148 | -12.080 | 1.416 | 26.475 | -8.077 | 20.256 | 24.526 | -29.235 | -23.453 | 25.623 | 13.699 | 26.389 | 26.542 |
| 4.423 | 21.489 | 17.243 | 6.593 | 7.325 | -19.991 | -30.378 | -10.674 | -2.232 | -22.103 | 3.813 | 9.515 | 3.763 | 20.288 |
| 9.132 | -22.249 | -23.739 | -16.441 | -19.358 | -21.367 | -18.964 | -18.838 | -31.231 | -11.349 | 24.591 | -29.628 | 23.111 | 18.346 |
| -24.082 | -1.288 | 29.398 | 14.571 | 4.919 | 19.706 | -0.747 | 16.470 | 18.000 | 25.891 | 20.214 | 26.156 | 10.912 | 24.256 |
| 27.531 | 22.786 | -20.082 | 30.881 | -15.966 | -10.922 | -4.478 | -3.649 | 23.189 | 24.168 | 2.246 | 16.691 | 30.950 | 24.398 |
| -26.757 | 1.883 | 18.139 | -15.260 | -4.219 | 5.861 | 19.668 | -28.125 | 11.835 | -29.098 | -14.616 | 29.442 | -10.151 | 15.349 |
| -8.648 | -18.439 | 9.844 | 24.874 | 13.961 | -15.164 | 8.737 | -9.212 | -8.556 | 22.765 | 2.959 | 25.257 | 28.679 | 22.654 |
| 3.539 | 6.073 | -6.976 | 18.259 | 18.197 | -14.127 | -18.687 | -11.012 | -13.941 | 23.589 | 9.934 | 15.785 | 29.006 | 25.610 |
| -1.564 | 4.828 | 19.578 | 15.841 | 24.793 | -19.149 | -8.251 | 14.865 | -24.156 | 28.275 | 25.522 | 7.002 | 7.077 | 17.008 |
| -5.343 | 3.418 | 11.884 | 8.204 | 9.060 | 21.619 | -15.745 | 30.577 | -30.691 | -19.812 | -17.184 | 17.220 | 14.642 | 22.136 |
| 16.141 | -11.258 | -4.818 | -15.923 | -4.470 | -2.488 | 17.111 | -23.858 | 0.918 | -22.304 | -16.733 | 30.933 | 22.416 | 13.951 |
| -9.189 | 2.462 | -24.560 | -30.580 | -15.774 | -29.159 | -9.739 | -3.378 | 18.355 | 26.325 | -18.818 | 18.108 | 2.075 | 19.019 |
| 17.399 | 0.154 | 9.263 | -10.189 | 4.214 | -0.852 | 29.300 | 23.569 | -29.421 | 19.985 | -15.961 | 15.250 | 21.791 | 18.580 |
| -18.296 | -22.898 | -3.174 | -10.541 | 15.593 | 0.754 | -20.850 | 18.178 | 26.748 | 18.336 | -17.556 | 9.509 | 15.553 | 25.500 |
| 25.636 | 16.777 | -6.652 | 23.259 | 3.593 | 13.074 | -2.810 | -4.752 | 28.825 | 25.233 | -2.417 | 19.859 | -9.153 | 23.818 |
| 20.847 | 4.166 | 26.236 | -14.172 | -10.960 | -21.028 | -26.694 | 30.152 | 27.762 | 16.144 | -16.483 | -4.916 | -14.237 | 22.931 |
| -14.386 | 2.696 | -15.090 | 12.392 | 8.701 | 11.803 | -22.315 | 22.271 | -28.201 | 12.719 | 19.971 | 12.718 | -3.105 | 16.841 |
| -20.422 | -2.552 | 20.697 | -5.016 | -4.347 | 9.328 | -4.076 | 26.409 | -9.516 | 14.922 | 16.162 | 19.260 | 18.243 | 12.153 |

| | | | | | | | | | | | | | |
|---|---|---|---|---|---|---|---|---|---|---|---|---|---|
| 22.947 | -3.715 | -20.488 | -4.458 | 26.289 | 6.370 | 4.012 | 24.263 | -23.683 | -7.817 | 21.824 | -0.047 | 21.452 | 16.523 |
| -13.446 | -11.617 | -28.328 | 27.715 | -26.723 | -9.225 | 2.634 | 14.839 | 11.041 | -10.958 | -10.359 | 23.977 | 25.899 | 24.299 |
| -27.532 | -12.875 | 17.473 | 11.371 | -19.228 | -19.725 | -8.377 | -10.081 | 9.712 | 24.274 | -8.861 | 10.384 | 24.913 | 7.074 |
| 14.681 | -5.928 | 22.239 | -12.413 | 23.216 | -18.246 | -1.226 | -12.979 | -6.297 | -20.194 | 20.939 | -9.300 | 20.640 | 4.074 |
| 13.798 | 25.458 | -16.991 | -29.551 | 1.461 | -23.478 | 16.662 | 20.285 | -20.889 | -23.886 | 10.150 | 0.490 | -14.285 | 20.587 |
| -3.456 | -23.745 | 13.084 | 2.808 | -2.625 | -26.839 | 12.868 | -26.643 | 22.529 | -16.385 | 22.807 | 28.392 | 27.306 | 21.488 |
| -25.802 | 7.792 | -10.159 | 19.623 | 0.355 | 4.040 | -20.969 | -30.622 | 7.332 | -21.228 | -6.899 | 20.979 | 5.538 | 7.275 |
| -19.315 | -4.652 | 15.001 | 1.028 | -8.570 | -25.513 | -16.768 | 28.914 | -3.610 | 10.580 | 1.904 | 10.041 | 23.210 | 15.585 |
| 28.690 | 24.523 | -6.467 | -16.289 | 21.525 | -5.497 | -26.927 | -2.179 | 12.470 | -11.844 | -12.269 | 7.451 | -25.669 | 8.675 |
| 29.769 | -14.482 | 4.426 | 1.586 | 24.482 | -28.715 | -9.720 | 5.532 | -0.929 | -26.407 | 3.700 | 6.337 | -10.117 | 5.313 |
| -29.780 | -17.146 | 22.098 | 9.169 | 16.699 | -21.865 | 5.800 | 28.010 | -18.047 | 9.893 | 11.280 | 3.061 | 7.156 | 19.952 |
| 15.791 | 27.071 | -5.864 | 13.666 | -2.288 | -10.569 | -1.886 | 25.282 | -22.178 | -27.955 | 20.162 | 27.805 | 18.062 | 25.384 |
| 30.452 | 5.529 | 9.396 | -20.095 | -7.135 | -14.886 | -4.954 | -5.516 | 28.653 | 18.226 | 24.904 | 26.651 | 21.204 | 10.887 |
| -23.122 | 3.431 | -9.519 | 5.295 | 20.338 | -8.423 | 6.559 | -15.666 | -16.302 | -19.403 | 13.819 | 0.964 | 13.269 | 15.320 |
| 23.322 | 2.363 | -19.226 | 24.501 | -20.552 | -1.102 | -6.482 | -23.392 | -21.596 | 22.695 | 18.968 | 16.712 | -0.521 | 7.092 |
| 3.782 | -1.336 | -3.672 | -0.468 | 8.662 | -15.862 | 7.254 | 19.409 | -29.790 | 28.514 | 11.256 | 17.642 | 9.707 | 25.657 |
| -17.929 | 8.416 | 17.399 | -14.335 | 10.585 | 1.614 | -9.237 | 2.545 | 25.384 | 5.909 | 4.811 | 10.930 | 17.471 | 20.532 |
| -6.144 | 16.165 | -8.636 | -22.640 | -10.000 | 0.627 | 29.219 | 14.568 | 26.341 | 20.711 | -2.122 | -2.271 | 1.098 | 11.090 |
| 26.748 | 0.206 | -0.741 | -17.735 | -0.463 | -13.580 | -3.014 | -16.401 | 12.835 | 7.123 | -11.258 | -12.926 | -8.477 | 19.908 |
| 1.903 | -14.418 | -10.020 | 31.078 | -14.815 | -18.377 | -0.561 | -27.033 | 22.863 | -5.929 | -4.290 | -27.518 | 22.099 | 21.747 |
| -1.752 | -16.486 | -8.329 | 21.241 | -23.069 | -0.058 | 2.663 | -8.074 | -2.023 | 19.465 | 15.375 | 13.807 | -6.518 | 20.759 |
| -26.137 | 12.783 | -12.356 | -16.722 | 22.750 | 15.564 | 13.117 | -22.147 | 4.339 | -17.198 | -2.471 | -24.492 | 29.649 | 18.713 |
| -9.420 | -12.785 | 21.467 | -19.592 | -4.028 | -6.471 | -2.563 | 14.891 | 18.988 | 23.987 | 14.729 | 14.446 | 25.220 | 21.551 |
| 12.754 | -2.145 | -1.358 | 11.931 | -21.346 | -3.976 | 10.795 | 13.284 | -26.296 | -29.139 | 19.597 | 23.958 | 20.207 | -1.451 |
| -2.992 | 16.957 | 4.121 | -28.218 | -20.061 | -20.228 | -20.046 | -11.683 | -0.563 | -18.783 | -25.986 | 9.627 | 19.519 | 24.023 |
| 29.084 | -26.008 | -0.049 | 15.086 | 11.713 | -21.262 | -6.940 | 9.312 | 22.383 | -14.936 | -8.341 | 8.171 | 28.218 | 12.737 |
| -22.046 | 14.621 | 5.034 | 24.710 | 3.607 | -27.630 | -10.438 | -29.607 | -10.825 | -21.189 | 7.550 | 24.255 | 24.068 | 20.622 |
| -4.849 | 1.696 | 22.753 | 22.700 | 1.787 | 3.776 | -5.398 | 7.697 | 13.527 | -25.631 | -10.606 | 5.696 | -17.437 | 5.551 |
| 21.644 | 9.040 | -27.123 | -31.009 | -15.674 | -13.425 | -3.899 | -30.704 | 2.944 | -23.574 | 23.696 | -9.984 | 15.113 | 12.010 |
| 20.969 | 24.414 | -20.007 | 20.016 | -1.423 | 24.748 | -17.620 | 22.553 | 27.086 | -19.391 | 22.668 | -13.883 | -19.589 | 23.484 |
| -17.166 | -4.398 | -3.734 | 6.448 | -11.983 | -18.281 | 9.208 | -4.062 | 30.588 | -24.207 | 26.811 | 25.176 | 21.851 | 2.965 |
| 1.870 | -11.678 | -5.258 | -21.710 | -10.211 | 3.208 | -23.072 | 17.663 | 28.405 | 11.101 | 20.468 | -12.235 | 15.692 | 22.890 |
| 16.554 | 4.570 | -23.719 | -22.205 | -4.344 | -5.913 | 26.831 | 0.694 | 11.207 | 6.048 | 13.992 | 30.056 | 2.099 | 24.861 |
| -18.850 | 16.218 | 3.110 | -25.789 | 11.358 | -15.672 | 20.026 | 16.026 | 18.604 | 18.785 | 21.915 | 29.495 | 6.115 | 19.097 |
| 23.101 | 10.321 | 21.023 | -15.929 | -10.895 | -25.762 | 5.575 | 5.975 | 15.860 | -11.121 | 16.064 | 12.413 | 23.340 | 21.952 |
| 9.936 | -13.446 | -2.631 | 12.277 | -3.395 | 1.793 | 3.483 | 1.799 | 12.084 | -22.585 | 5.680 | 9.319 | 9.913 | 20.047 |

| | | | | | | | | | | | | | |
|---|---|---|---|---|---|---|---|---|---|---|---|---|---|
| -24.689 | 24.770 | 7.596 | -12.282 | 7.297 | -16.366 | 9.981 | -25.123 | -10.361 | -21.419 | 24.862 | 13.613 | 25.750 | 9.705 |
| -26.050 | 4.246 | -8.818 | -0.695 | 7.870 | -18.084 | 4.486 | -9.214 | -25.798 | -6.975 | 22.947 | 15.776 | 23.752 | 5.164 |
| 9.199 | 6.638 | -24.324 | -10.324 | -7.985 | -20.920 | 3.951 | 5.282 | 9.259 | -24.404 | 12.455 | -27.083 | -0.913 | -4.006 |
| 24.368 | -1.922 | -7.522 | 14.979 | -21.352 | -26.021 | -6.622 | -12.347 | -19.447 | 7.723 | 6.383 | 19.570 | 2.014 | 20.648 |
| -20.892 | 14.086 | -20.363 | -30.853 | -21.733 | -12.304 | -4.054 | -25.655 | 6.342 | -11.944 | 15.866 | 15.154 | 28.248 | -1.918 |
| 20.709 | 23.109 | -24.971 | 13.192 | -9.038 | 0.851 | -8.378 | -10.831 | 26.381 | 28.369 | 23.425 | 13.194 | 12.146 | 25.635 |
| 6.915 | 16.984 | 25.098 | 10.982 | -2.636 | 14.769 | 27.881 | -20.799 | -15.445 | -22.872 | 25.087 | 26.310 | -20.536 | 14.638 |
| 18.643 | 4.233 | 24.207 | -23.894 | 2.415 | 6.832 | -2.606 | -3.320 | -24.010 | -4.130 | 24.212 | 27.491 | 25.094 | -24.415 |
| -14.837 | -16.751 | -10.099 | 17.986 | 3.669 | -9.395 | 12.363 | 23.571 | 27.290 | -7.691 | -21.210 | 7.832 | 19.674 | 23.110 |
| 29.304 | -10.195 | 26.642 | -27.260 | -10.009 | -13.259 | -8.981 | 20.429 | 8.422 | -9.255 | 21.451 | 23.992 | 29.455 | 3.369 |
| 21.257 | -9.427 | -11.700 | -14.126 | 4.833 | -24.269 | 16.265 | -11.610 | 22.938 | -15.539 | 17.490 | 18.462 | 16.954 | 19.296 |
| 15.986 | -13.543 | -17.218 | 24.097 | -26.658 | -26.255 | 2.834 | 12.721 | 7.736 | 19.403 | 25.716 | 22.923 | -3.058 | 15.155 |
| -7.371 | 11.089 | -12.236 | 30.196 | 23.092 | -10.028 | -11.029 | 14.738 | -29.890 | 8.317 | 23.436 | 14.172 | 15.282 | 18.730 |
| -29.186 | 14.449 | 9.685 | -12.479 | 14.566 | 11.153 | -1.907 | -4.227 | 28.245 | 16.208 | 24.893 | 30.892 | 18.695 | 2.359 |
| 14.067 | -8.037 | -4.589 | 26.608 | -1.436 | 1.360 | -3.427 | -18.473 | -9.730 | 6.663 | 26.217 | -1.640 | 26.872 | -0.237 |
| -18.343 | 8.699 | -6.416 | 27.155 | 12.612 | 6.093 | 11.783 | -25.858 | 25.384 | -25.064 | 22.331 | -1.119 | 19.076 | 16.296 |
| 14.503 | -13.044 | -0.933 | 5.687 | -23.318 | 18.866 | 24.999 | 15.923 | -3.677 | 27.442 | 20.999 | -18.860 | -10.316 | 11.560 |
| -9.330 | 12.611 | -21.958 | 0.309 | 8.946 | 22.428 | 3.924 | 0.342 | -4.707 | -19.474 | 19.689 | 26.970 | 10.380 | -4.704 |
| 0.147 | 2.635 | -7.440 | 15.029 | 4.171 | -16.420 | -6.142 | -2.466 | 23.570 | 21.146 | 26.896 | 29.993 | 14.010 | 15.574 |
| -28.315 | -8.644 | 0.918 | 18.596 | 24.010 | -25.320 | -2.816 | 11.464 | -28.703 | 7.252 | 10.362 | -9.030 | 11.619 | 14.976 |
| -0.431 | 24.879 | 11.415 | -25.894 | 19.593 | 3.588 | -7.086 | 9.809 | -28.176 | -18.014 | 8.265 | 12.014 | -11.496 | 20.774 |
| 31.273 | 18.016 | -11.753 | -9.274 | 1.562 | 21.730 | -10.992 | -11.341 | 15.005 | -21.288 | -22.338 | -25.974 | -10.258 | 16.351 |
| -2.232 | 0.644 | -9.296 | 21.549 | -8.423 | 7.154 | 16.220 | 17.415 | -25.418 | 4.884 | 22.005 | 28.411 | 13.202 | -1.176 |
| 23.792 | 8.851 | -2.312 | -21.092 | 18.620 | -3.435 | -24.296 | -29.697 | -8.916 | 24.167 | 17.305 | 10.378 | -16.025 | 9.711 |
| 10.292 | 18.820 | 5.793 | 14.167 | -0.821 | 13.248 | -8.756 | 10.201 | -18.844 | -6.453 | 27.205 | 22.651 | 1.530 | 14.070 |
| 5.279 | 16.434 | -17.967 | 12.016 | -25.997 | -7.530 | -25.132 | -5.329 | -29.338 | 16.456 | 17.718 | 11.768 | 28.269 | 17.151 |
| -29.269 | 23.110 | 8.280 | 29.567 | -21.462 | -12.287 | -3.986 | 7.542 | 12.258 | 1.769 | 19.254 | 15.861 | 16.567 | 12.679 |
| 28.528 | 16.808 | 1.858 | -24.420 | 21.798 | 5.892 | -26.989 | -8.838 | -6.484 | 8.636 | 14.641 | 7.973 | 15.920 | 15.900 |
| 0.341 | -8.938 | 7.860 | -16.922 | 11.353 | -26.502 | 18.978 | -22.927 | 23.208 | -0.394 | 28.787 | 28.348 | 22.487 | 13.118 |
| -21.420 | -23.757 | -14.302 | 26.376 | -0.264 | -6.941 | 0.052 | -22.023 | 10.730 | -18.534 | 18.830 | 20.702 | 24.722 | 13.613 |
| 15.252 | -24.376 | 6.611 | -5.288 | -16.678 | -8.853 | -0.646 | -20.682 | -1.984 | 22.867 | 18.265 | 20.108 | 3.840 | 16.308 |
| 7.667 | -18.416 | -1.542 | 24.339 | 0.771 | -17.789 | 5.097 | 15.225 | -10.877 | -21.636 | -9.921 | 26.419 | 22.728 | 10.504 |
| -4.960 | -6.708 | -8.687 | 27.654 | 17.287 | -24.356 | 22.460 | -26.090 | -26.106 | 19.554 | -9.460 | 21.821 | 21.209 | 12.133 |
| 23.230 | -16.164 | 0.166 | -30.309 | 13.308 | 3.108 | 13.043 | 23.398 | 26.592 | 25.446 | 25.139 | 18.598 | 19.110 | 9.622 |
| -13.136 | -21.946 | -4.817 | -18.885 | 4.063 | 2.166 | 1.441 | -0.336 | -10.010 | 20.722 | 22.965 | 13.440 | 14.502 | 21.243 |
| -14.938 | -2.036 | -10.736 | -29.603 | -1.580 | -2.725 | 4.499 | 21.792 | -22.836 | 11.814 | 7.232 | 19.222 | -20.871 | -1.763 |

| | | | | | | | | | | | | | |
|---|---|---|---|---|---|---|---|---|---|---|---|---|---|
| -27.215 | 4.897 | -18.775 | 30.180 | -9.367 | -19.138 | -9.576 | 24.622 | 18.548 | -4.615 | 23.649 | 5.088 | 26.314 | 25.880 |
| -12.718 | 1.012 | -23.250 | 20.104 | 0.537 | 14.056 | -19.994 | -7.299 | -7.239 | 24.743 | 26.348 | -14.955 | -8.270 | 22.296 |
| -24.783 | -16.848 | 13.876 | 28.317 | 6.467 | 2.730 | -4.702 | 18.684 | -28.915 | 18.221 | 1.030 | 22.591 | 24.441 | 21.385 |
| -2.494 | -11.513 | 9.212 | -0.328 | -12.245 | -9.870 | -2.679 | 9.181 | -11.505 | -1.043 | 22.127 | -17.423 | 10.653 | 12.159 |
| 0.823 | 3.072 | 1.385 | -14.274 | 16.607 | 15.097 | -4.080 | 23.935 | -8.570 | 5.922 | 28.260 | 27.909 | 2.737 | 8.238 |
| 13.728 | -16.246 | -19.537 | -17.295 | 7.351 | 22.418 | -25.404 | -26.592 | -23.619 | 26.668 | 16.403 | 29.882 | 9.065 | 6.263 |
| -7.022 | -18.518 | 3.373 | -4.611 | -2.099 | -9.248 | 3.867 | 9.716 | 11.150 | 0.942 | 25.184 | 13.812 | 16.937 | 22.816 |
| -3.715 | 11.159 | -3.863 | -28.944 | 3.270 | -17.623 | 17.703 | 7.067 | 22.722 | 12.403 | 17.704 | 2.448 | 18.004 | 20.201 |
| 14.178 | 24.591 | 18.876 | 13.148 | -17.455 | -6.795 | 1.722 | -2.120 | -1.396 | 9.021 | -7.722 | 16.971 | -5.542 | 13.151 |
| 30.466 | -5.844 | 14.455 | -15.029 | 21.275 | -18.625 | -3.180 | 26.893 | 27.611 | -25.623 | 15.654 | 19.462 | 19.634 | 14.033 |
| 21.495 | 8.446 | 1.476 | -23.334 | 18.550 | -7.545 | -2.302 | 1.037 | -19.427 | 16.717 | 24.530 | 10.288 | 28.301 | 16.290 |
| -4.593 | -10.533 | 0.321 | 13.619 | 1.565 | 17.619 | -6.952 | -25.337 | -27.907 | -9.671 | 27.036 | 24.590 | 11.578 | 22.684 |
| -1.188 | 1.624 | -26.138 | -21.198 | 26.030 | -12.203 | 20.335 | 1.890 | 0.050 | 21.188 | 17.684 | 29.418 | 27.696 | 6.276 |
| -29.181 | -14.545 | 26.629 | 7.583 | 2.776 | -11.356 | -14.910 | -26.392 | -27.011 | 9.535 | 10.985 | 9.668 | 1.743 | 9.884 |
| 23.835 | -4.734 | -6.279 | 3.319 | -26.095 | -0.837 | -8.152 | 2.958 | -3.782 | 21.755 | 18.572 | 20.769 | -9.095 | 26.860 |
| -0.724 | -13.320 | -6.911 | 11.568 | 19.344 | 2.841 | -1.017 | 12.475 | -5.221 | 2.286 | 22.358 | -11.979 | 1.202 | 15.850 |
| 1.112 | 25.650 | -13.390 | -20.205 | -1.232 | -13.755 | 0.721 | -22.410 | 6.302 | 0.480 | 20.594 | 25.900 | 7.762 | 4.568 |
| 17.878 | 1.301 | -14.524 | -1.124 | 20.755 | 13.526 | -8.980 | -27.104 | -10.885 | 10.208 | -21.535 | 0.134 | -1.552 | 20.849 |
| 5.634 | -3.171 | -7.859 | 18.285 | -27.522 | -26.608 | 3.550 | 18.230 | -25.266 | -3.985 | 20.021 | 12.794 | 15.110 | 18.206 |
| -5.023 | -1.914 | -11.746 | 24.973 | -25.092 | -20.724 | -6.597 | -22.870 | 22.222 | 26.373 | 12.466 | 14.453 | 3.889 | 22.043 |
| -20.207 | -8.552 | -22.618 | -16.696 | 26.816 | -24.267 | 6.927 | -3.666 | -28.262 | -3.407 | 28.738 | 7.704 | -9.081 | 19.036 |
| 26.511 | 12.887 | -5.337 | 11.754 | -24.615 | 13.708 | -15.187 | 8.881 | 18.553 | -24.773 | 13.819 | -16.125 | -13.398 | 17.086 |
| -9.283 | -0.379 | 22.474 | -5.442 | -26.688 | -9.059 | -18.544 | 29.587 | 3.373 | 12.751 | 30.052 | 23.095 | 17.729 | 17.688 |
| 9.090 | 13.378 | -20.652 | -21.889 | 24.710 | 23.799 | -1.580 | -19.108 | 9.860 | 13.914 | 19.873 | 24.456 | 20.707 | 27.989 |
| -0.021 | -7.331 | 11.163 | 29.256 | 26.197 | 13.276 | 16.297 | 10.645 | 7.434 | -5.604 | 15.261 | 11.422 | 19.053 | 20.009 |
| -19.510 | -18.792 | 3.916 | 21.225 | 16.699 | -5.956 | 21.670 | 4.376 | -19.190 | 2.838 | 27.553 | 4.700 | 17.039 | 28.837 |
| 6.593 | 6.569 | -8.700 | 10.018 | 11.008 | -9.656 | 11.175 | -9.893 | 5.255 | 10.436 | 26.643 | 26.685 | 7.204 | 22.611 |
| -28.038 | 23.604 | 21.518 | 12.261 | -29.316 | 24.224 | -14.667 | -4.968 | -9.231 | -16.091 | 22.710 | 20.926 | 5.600 | 21.155 |
| 25.780 | 19.539 | 8.401 | 26.162 | -0.949 | -0.060 | -9.724 | 21.282 | 11.063 | 4.773 | 17.974 | 27.954 | -5.449 | 16.815 |
| -10.764 | 0.059 | 13.287 | -18.143 | -7.300 | -14.582 | 23.565 | -2.765 | -23.783 | 22.955 | 26.480 | 25.290 | -2.199 | 12.080 |
| 21.810 | -6.068 | -14.699 | 2.225 | -23.913 | -13.417 | 12.098 | -9.474 | 6.282 | 20.613 | 6.023 | -0.761 | 25.931 | 15.028 |
| 29.115 | 11.532 | -0.162 | -13.796 | -6.726 | -16.582 | 7.480 | -0.797 | -16.884 | 17.520 | 29.620 | 28.344 | -1.633 | 12.876 |
| 3.345 | 12.919 | 17.920 | -15.256 | 24.107 | -3.312 | 10.112 | -23.687 | 17.226 | 21.743 | 19.798 | 14.271 | -15.445 | 18.377 |
| -25.013 | 7.055 | -2.498 | -21.455 | -12.343 | -11.640 | -4.295 | -16.910 | 25.413 | 16.057 | 22.084 | -4.916 | 8.276 | 12.366 |
| -2.626 | 6.124 | -3.238 | -27.854 | 21.924 | -24.458 | -13.999 | -15.157 | 13.632 | 15.265 | -6.642 | 8.609 | 25.219 | 21.023 |
| -17.546 | -19.894 | -7.707 | -21.965 | 6.728 | -10.134 | -12.772 | 17.843 | 25.857 | -20.376 | 24.523 | 6.901 | 23.882 | 20.463 |

| | | | | | | | | | | | | | |
|---|---|---|---|---|---|---|---|---|---|---|---|---|---|
| -27.236 | -23.546 | 7.206 | -6.309 | -24.282 | -12.105 | -2.139 | -21.326 | -22.820 | -21.825 | 20.470 | 29.656 | 14.772 | 17.160 |
| 12.863 | -3.118 | 22.027 | 19.307 | 28.066 | 15.577 | -26.374 | 21.758 | 24.192 | 23.317 | 9.311 | 11.537 | 24.297 | 23.550 |
| -22.200 | -0.447 | -8.233 | 14.626 | 25.191 | -2.395 | 12.956 | 3.238 | -3.650 | -3.385 | 25.552 | 22.082 | 22.551 | 27.407 |
| -19.020 | -19.795 | 14.416 | -23.002 | 8.090 | -18.605 | -1.617 | -10.484 | 24.753 | 11.034 | 18.552 | 2.836 | 15.882 | 16.325 |
| -26.581 | -4.275 | -4.005 | -28.560 | 1.165 | 1.861 | -8.952 | -11.662 | 21.838 | 24.901 | 23.707 | 25.429 | 19.297 | 18.070 |
| 19.684 | 19.514 | -19.505 | -24.899 | 24.125 | -6.402 | 3.261 | 21.259 | 18.173 | -18.800 | 27.506 | 24.649 | 20.805 | 20.546 |
| 18.249 | 2.099 | 11.938 | 16.507 | -6.798 | -20.214 | -7.308 | 26.717 | 16.435 | -20.840 | 29.782 | 19.602 | 9.876 | -4.807 |
| -21.864 | -4.702 | 9.291 | -3.904 | 4.483 | -21.980 | 3.945 | -25.117 | -8.694 | 9.434 | 24.373 | -6.137 | 25.594 | 19.418 |
| -6.034 | 14.881 | -11.264 | -16.843 | 11.863 | 21.109 | 23.160 | 10.693 | 19.117 | 12.721 | 10.852 | 15.877 | -17.948 | 14.129 |
| 5.294 | 0.956 | 11.190 | 5.007 | 22.867 | -15.401 | 7.722 | -1.984 | 22.937 | -5.587 | 28.353 | 7.852 | -7.299 | 10.607 |
| -22.915 | 15.989 | -13.929 | 19.093 | 27.852 | -6.859 | -3.490 | 24.437 | 8.958 | 22.169 | 26.783 | 10.321 | 16.724 | 11.793 |
| 20.816 | -24.259 | 2.119 | -2.424 | -27.339 | -21.159 | 15.921 | 5.061 | -16.620 | 13.148 | 20.472 | 23.176 | 0.648 | 9.706 |
| -20.254 | -2.343 | -7.578 | -7.383 | 14.075 | 3.623 | 25.641 | 25.334 | 19.504 | 5.136 | 13.332 | 26.780 | -22.387 | 18.821 |
| -18.438 | -13.482 | -22.984 | 19.780 | -24.844 | -24.088 | -6.266 | 29.122 | -6.833 | -0.774 | 21.336 | 26.118 | 18.541 | 15.218 |
| 22.029 | 17.221 | 16.690 | -19.082 | 25.406 | 6.063 | -9.296 | 0.299 | -22.970 | -16.705 | 9.045 | 21.382 | -25.209 | 26.552 |
| 16.217 | 13.167 | -0.713 | -24.494 | 28.699 | 4.079 | -11.318 | -24.064 | -25.427 | 7.451 | 15.576 | 7.398 | -23.646 | 16.538 |
| -23.745 | -0.721 | -4.979 | -14.161 | 15.921 | -11.203 | 24.511 | -0.642 | 25.739 | 5.343 | -16.211 | 18.441 | 21.403 | 13.620 |
| -24.971 | 24.515 | 3.753 | -20.827 | 1.654 | -18.303 | -17.134 | 19.739 | -10.729 | -13.379 | 27.977 | 28.915 | -16.129 | 22.655 |
| 2.930 | 14.509 | -9.663 | 29.015 | -8.522 | -20.499 | 12.895 | 7.199 | -23.921 | 9.380 | -7.863 | 15.590 | -3.146 | 10.469 |
| -27.012 | 3.201 | -19.086 | 23.810 | 9.813 | -5.351 | 15.494 | 3.678 | -26.543 | 18.616 | 4.930 | 19.860 | 18.253 | 22.195 |
| 24.660 | 27.095 | 4.101 | 28.563 | -30.556 | -7.917 | 4.884 | 13.001 | -18.267 | 1.537 | 25.714 | 12.441 | 10.593 | 10.982 |
| -1.714 | 26.658 | -12.003 | -10.458 | 25.894 | -13.722 | 21.615 | 27.579 | 8.285 | -2.668 | 9.135 | 9.465 | 21.978 | 28.862 |
| -16.977 | 28.011 | -26.496 | 2.774 | 7.722 | -14.751 | -6.660 | -27.476 | -23.346 | -14.766 | 21.200 | 0.359 | 2.396 | -3.537 |
| -5.320 | -24.897 | 19.995 | -26.310 | 6.698 | -22.152 | 6.435 | 17.445 | 0.146 | 5.811 | 13.033 | 16.488 | -2.265 | 24.489 |
| -9.725 | 26.404 | 5.315 | 20.502 | -26.554 | -7.238 | -11.605 | -6.854 | -24.915 | -0.571 | 9.355 | 24.944 | -11.597 | 13.689 |
| -12.784 | -26.753 | -24.714 | -5.910 | 24.704 | -21.706 | 17.035 | -17.571 | 20.412 | -5.245 | 26.599 | 22.776 | -12.349 | 27.295 |
| -24.171 | 17.049 | 0.156 | 17.165 | -6.072 | 5.294 | -4.784 | -26.861 | -14.228 | 25.758 | 18.609 | 5.852 | 23.447 | 19.984 |
| 5.298 | 14.614 | -1.780 | 18.350 | -8.975 | 6.306 | -6.047 | 23.278 | -20.191 | -21.457 | 17.632 | 5.478 | 26.727 | 4.192 |
| -21.613 | 21.192 | -7.706 | -26.743 | 29.012 | -6.127 | -16.277 | 11.767 | -19.174 | -9.574 | 26.190 | 14.501 | 13.227 | 16.173 |
| 23.770 | 0.034 | -5.178 | -25.718 | 21.798 | -16.500 | 18.687 | -11.308 | 15.697 | 18.944 | 21.014 | 21.883 | -10.230 | 7.377 |
| -12.138 | 18.613 | -9.377 | -17.598 | 7.340 | -20.350 | 10.146 | 12.396 | 23.859 | 17.660 | 22.761 | 19.274 | -22.720 | 15.323 |
| -24.733 | 14.428 | -10.451 | 16.099 | 29.662 | 1.740 | -6.284 | -18.709 | 5.870 | -24.636 | 19.894 | -6.706 | 14.126 | 9.268 |
| 28.692 | -14.328 | 17.364 | 11.992 | -26.728 | -1.503 | -11.558 | 20.055 | 18.127 | 21.260 | 20.419 | 16.783 | 7.343 | 13.331 |
| -23.512 | -17.936 | -5.889 | -25.249 | 19.826 | -5.054 | -10.997 | -11.733 | -1.941 | -0.883 | 7.958 | 15.138 | 14.545 | 16.996 |
| 8.614 | -21.898 | -23.140 | -16.046 | 17.158 | -11.554 | -24.788 | -15.680 | 7.317 | 17.274 | 16.973 | 8.263 | 12.082 | 20.913 |
| 22.884 | -10.123 | -25.223 | -6.603 | 8.473 | 17.301 | -10.614 | -14.422 | 24.505 | 8.983 | 24.096 | -5.265 | -11.484 | 15.392 |

| | | | | | | | | | | | | | |
|---|---|---|---|---|---|---|---|---|---|---|---|---|---|
| 17.347 | 2.283 | 11.973 | 26.053 | 19.373 | -25.914 | -16.429 | 7.939 | 23.353 | -2.251 | 16.389 | 28.902 | -20.806 | 20.364 |
| -22.140 | 19.681 | -4.631 | 14.169 | 13.236 | -0.807 | 5.217 | 14.448 | 14.082 | 9.195 | 13.864 | 18.611 | -15.359 | 22.630 |
| -9.813 | -19.661 | 5.537 | 23.501 | 23.553 | -17.667 | -19.317 | 18.904 | 4.868 | -24.772 | 21.041 | 5.643 | -8.608 | 16.444 |
| 0.968 | 29.236 | -3.470 | 27.361 | 16.437 | -23.792 | 14.694 | 27.570 | 4.272 | 26.222 | 5.773 | -3.378 | -9.460 | 2.287 |
| -2.720 | -21.777 | 8.058 | 8.639 | 11.635 | 12.524 | -13.805 | 17.811 | -10.990 | 9.366 | 27.436 | -20.580 | -23.208 | 13.659 |
| -25.256 | -28.109 | 4.837 | 22.439 | 6.111 | -1.820 | 6.530 | -2.376 | -25.736 | -11.737 | -6.238 | 24.276 | 9.620 | 10.849 |
| 17.799 | -19.266 | 6.888 | 13.728 | 0.731 | -9.856 | 10.385 | 5.874 | -12.758 | -22.233 | 12.675 | 15.444 | -21.610 | 25.391 |
| 13.037 | -26.068 | 3.633 | -23.180 | -27.449 | 18.602 | 1.803 | -9.866 | -6.534 | 25.369 | 18.559 | 14.064 | 22.405 | 28.113 |
| -19.542 | -29.506 | 11.306 | -23.944 | 12.297 | -3.298 | -12.324 | -12.603 | -9.605 | -16.959 | 22.831 | 21.247 | 8.046 | 12.978 |
| 21.650 | 5.213 | 9.558 | 13.292 | 16.683 | -11.753 | 8.602 | -6.356 | 11.005 | -22.272 | 27.416 | 5.251 | 3.975 | 23.547 |
| -6.445 | 12.165 | 1.760 | -8.943 | -20.601 | -2.597 | -13.886 | 9.967 | -12.235 | -2.925 | 23.588 | 25.793 | 22.780 | 17.301 |
| -26.348 | 12.804 | 0.973 | -11.697 | 9.649 | -5.343 | 14.885 | -21.669 | 25.228 | 22.695 | -20.644 | 1.363 | 20.049 | 19.588 |
| 11.265 | -26.550 | 2.988 | 24.120 | -15.034 | 6.870 | -14.731 | -19.350 | -9.261 | -24.147 | 13.045 | 8.786 | -16.473 | 5.151 |
| -14.618 | 27.295 | 13.165 | 22.926 | 28.213 | 10.235 | -13.192 | 24.645 | -14.716 | 20.630 | 26.280 | 9.084 | 6.707 | 8.730 |
| 1.823 | 23.229 | -19.416 | 8.317 | 26.479 | 4.001 | -23.084 | 24.088 | -13.989 | 25.750 | 25.164 | -4.340 | 9.169 | 11.756 |
| 6.538 | 20.726 | -10.419 | 26.589 | -32.056 | -10.867 | 12.708 | -4.179 | -8.252 | -9.070 | -20.824 | 19.810 | -18.042 | 18.019 |
| -25.484 | -2.304 | -7.142 | -27.205 | 19.978 | -16.560 | 5.943 | -20.015 | -9.915 | 8.352 | 19.924 | 2.372 | -4.113 | 13.827 |
| -25.963 | 9.030 | -4.997 | -21.162 | 13.954 | 1.168 | 23.894 | 1.407 | 22.819 | 15.302 | 24.099 | 17.118 | -2.009 | 27.800 |
| 27.634 | -17.546 | 4.392 | 25.746 | 23.781 | -15.295 | -10.058 | -26.736 | 12.295 | -24.972 | -3.524 | -28.221 | -11.288 | 16.201 |
| 2.426 | -19.582 | -6.250 | 11.157 | 10.321 | -3.307 | -16.482 | 2.710 | 14.183 | 20.752 | -6.420 | 11.209 | -9.529 | 10.328 |
| 14.400 | 3.562 | -11.137 | 15.001 | 16.001 | -4.498 | 8.891 | -10.530 | 16.321 | 18.965 | 15.076 | -19.099 | 10.643 | 24.635 |
| 7.638 | -17.056 | -12.200 | -17.684 | 20.611 | 16.987 | 18.169 | 11.782 | 20.098 | 12.911 | 11.235 | 25.107 | 16.664 | -5.850 |
| 3.035 | -7.539 | 11.801 | 9.052 | 17.195 | 14.630 | 5.006 | -25.332 | -0.594 | 21.172 | -19.732 | -11.472 | 19.172 | 22.828 |
| 15.159 | 28.935 | -11.723 | 24.715 | 22.798 | -7.893 | -0.888 | -4.031 | 9.582 | 14.772 | 22.363 | -18.609 | -10.830 | 14.775 |
| -13.562 | -16.310 | 9.200 | -14.105 | 13.829 | -9.167 | 6.626 | 16.539 | 24.834 | 12.729 | 21.686 | 14.650 | 21.560 | 13.150 |
| 20.883 | 31.109 | -13.353 | 21.878 | 15.942 | -24.249 | 15.935 | 15.364 | 11.515 | 10.120 | 8.141 | -0.422 | -2.580 | 16.309 |
| -20.881 | 12.875 | -9.458 | -0.378 | 22.529 | 5.512 | 0.082 | -12.555 | 2.157 | 18.390 | 26.804 | 21.431 | -20.650 | 9.385 |
| -24.512 | 6.546 | -2.809 | -24.776 | -29.735 | 0.184 | -2.653 | 23.922 | 22.253 | -4.401 | 20.891 | 10.034 | -17.383 | 20.806 |
| 25.893 | 30.209 | -11.022 | 2.719 | -27.142 | -14.118 | -8.154 | 17.609 | 17.789 | -7.772 | 21.903 | 19.373 | -17.043 | 14.957 |
| 20.294 | 17.237 | -1.197 | -23.200 | 31.067 | 3.984 | 13.398 | 17.141 | 12.235 | -23.526 | 6.970 | -8.345 | 19.583 | 22.069 |
| 8.315 | 19.827 | 6.346 | -16.112 | 7.064 | 9.472 | -12.569 | 9.203 | 11.676 | -25.103 | 7.631 | 22.659 | 18.933 | 13.142 |
| 0.716 | -17.814 | -15.566 | 6.729 | 11.785 | -11.149 | 12.230 | 21.886 | -10.149 | 3.117 | 24.325 | 6.833 | -24.699 | 19.932 |
| 8.949 | 25.432 | 24.625 | -1.560 | 23.483 | 24.156 | -3.184 | -13.801 | 16.367 | 25.065 | 22.710 | 20.400 | 11.239 | 12.838 |
| 23.575 | 30.250 | 17.078 | -26.707 | 7.442 | 3.201 | -19.276 | 11.186 | 2.615 | 20.788 | 19.960 | 13.215 | 21.278 | 15.569 |
| -10.558 | -20.560 | 21.635 | -16.981 | -25.228 | -2.004 | 1.816 | 12.423 | 24.109 | 12.630 | 17.133 | 10.409 | 25.378 | 25.345 |
| 12.547 | 4.983 | -11.399 | -22.089 | 24.708 | 19.721 | -0.648 | -25.898 | 1.817 | 24.822 | -15.186 | 18.342 | -8.146 | 28.228 |

| | | | | | | | | | | | | | |
|---|---|---|---|---|---|---|---|---|---|---|---|---|---|
| 26.991 | 0.792 | -18.134 | 13.067 | 15.842 | -17.499 | 7.727 | -8.429 | 19.314 | -4.601 | 8.906 | 15.794 | 8.415 | 20.290 |
| -11.624 | 10.380 | -22.431 | -27.611 | -23.495 | 20.576 | -1.121 | -23.199 | 18.143 | 25.950 | 13.138 | 24.868 | 22.725 | 17.045 |
| -25.588 | 3.686 | -19.098 | -14.235 | 19.050 | 5.952 | 18.995 | 19.293 | -10.989 | 13.856 | 14.496 | 16.187 | 21.767 | 13.416 |
| 25.828 | -21.445 | -12.533 | -25.805 | -29.537 | 19.110 | 8.492 | -15.641 | 25.400 | 9.106 | 23.640 | 5.811 | 2.924 | 23.037 |
| -17.090 | 28.939 | 10.924 | 19.324 | -11.599 | 15.256 | -19.928 | 25.993 | -9.618 | -16.844 | 19.565 | 18.845 | -6.289 | 9.873 |
| 5.536 | -11.069 | 25.306 | 5.398 | -20.067 | -6.629 | -2.001 | 10.029 | 24.783 | 27.569 | 12.971 | 27.149 | 12.316 | 14.227 |
| 3.508 | -25.177 | -14.784 | 21.480 | 29.653 | 12.147 | -6.438 | -10.773 | 18.447 | 23.772 | -1.017 | 23.335 | -25.072 | 12.998 |
| 7.701 | 6.676 | 12.507 | -19.134 | 26.188 | 2.609 | -13.230 | -22.150 | 16.358 | 28.360 | -20.669 | 0.979 | 13.762 | 11.286 |
| 6.671 | 18.973 | 0.976 | 23.653 | 19.741 | -7.282 | -9.697 | 25.362 | 8.643 | 20.158 | 17.649 | 3.702 | -13.446 | 21.263 |
| 22.625 | 9.060 | -11.018 | -2.056 | 5.647 | -18.003 | 19.220 | 13.624 | -6.259 | 24.113 | 7.064 | 21.348 | 22.965 | 15.085 |
| -14.961 | 17.348 | -11.758 | 17.657 | 18.428 | -0.096 | -1.941 | 14.545 | 17.191 | 20.828 | 21.694 | 25.733 | 25.576 | 15.578 |
| -19.144 | -9.359 | -22.816 | 11.130 | -6.803 | -9.273 | -20.478 | 20.483 | 14.930 | 24.673 | 9.936 | 13.027 | 4.759 | 18.252 |
| 4.469 | 12.520 | -13.461 | 14.906 | -19.322 | -8.039 | 0.825 | 16.261 | 0.022 | 29.656 | 9.035 | 12.141 | 1.449 | 25.119 |
| -8.290 | -7.614 | -24.268 | 14.189 | 16.337 | -4.201 | 7.289 | -22.984 | 25.123 | 19.087 | 7.798 | 4.148 | 18.510 | 12.828 |
| 1.605 | 29.716 | 0.615 | -11.965 | 7.178 | 0.297 | -1.913 | 7.808 | -17.864 | -18.073 | 0.072 | -6.437 | 21.358 | 15.266 |
| -26.443 | 26.946 | 7.447 | 16.877 | 25.989 | -13.274 | 8.109 | -12.257 | -5.650 | -14.475 | 9.051 | -7.734 | -3.171 | 21.094 |
| 22.567 | -20.611 | -12.722 | 3.171 | -22.590 | -7.365 | 0.413 | 7.173 | 7.506 | 22.315 | -13.399 | 11.448 | -26.461 | 21.522 |
| -22.437 | -14.275 | 17.741 | -7.229 | -22.366 | 10.745 | 1.926 | -0.800 | 20.379 | 19.962 | -14.232 | -12.049 | 9.341 | 22.085 |
| -7.738 | -3.526 | -12.223 | -15.889 | 22.463 | -22.066 | -0.390 | 0.882 | 26.012 | 4.264 | -24.799 | -4.479 | -8.464 | -0.390 |
| 14.668 | 25.914 | -10.378 | 25.111 | 28.181 | 18.541 | 2.993 | -10.798 | 12.893 | 11.336 | -14.525 | -16.589 | 20.907 | 25.889 |
| 20.643 | -8.373 | -0.422 | -26.726 | 8.878 | -24.119 | -24.428 | -7.209 | 23.167 | -30.328 | -21.275 | 22.091 | 22.678 | 27.716 |
| -3.428 | 17.930 | -20.875 | -17.295 | -5.913 | 22.375 | -10.212 | -9.288 | 15.833 | 20.544 | -7.903 | 7.130 | -9.475 | 13.719 |
| 23.356 | 4.163 | -18.636 | -25.097 | 4.530 | -16.307 | -7.939 | -2.069 | -20.134 | 25.513 | -13.426 | 1.671 | 19.920 | 24.454 |
| 6.697 | 18.102 | 0.292 | -14.343 | -24.837 | -3.176 | -0.824 | -24.915 | -23.822 | 27.606 | 21.990 | 0.330 | -15.269 | 21.977 |
| 4.102 | -10.630 | -15.708 | 4.965 | 21.297 | -18.459 | -2.554 | -24.196 | 23.749 | -21.422 | -16.025 | 26.130 | -13.293 | 16.941 |
| 18.609 | 26.649 | 25.041 | 11.065 | 10.830 | 1.605 | -11.160 | -8.212 | -14.885 | 17.402 | 10.779 | 25.594 | 18.510 | 14.585 |
| -14.806 | -22.622 | 18.608 | -11.514 | -11.302 | -3.650 | -17.852 | 24.197 | 7.633 | 21.719 | -16.212 | 18.486 | -0.073 | 23.674 |
| 22.084 | -20.106 | 0.145 | 22.478 | 21.851 | 3.422 | -17.249 | 21.157 | 10.277 | 23.380 | 18.600 | 16.064 | 23.592 | 9.092 |
| 19.862 | 16.784 | 7.381 | -22.132 | 23.273 | -21.779 | 22.880 | -14.626 | -7.525 | 20.791 | -15.254 | 11.697 | 20.764 | 8.657 |
| -24.859 | 28.756 | -11.641 | 7.769 | 28.454 | 10.716 | -15.889 | -21.910 | -9.314 | 9.424 | -15.901 | -5.188 | 24.798 | 19.507 |
| 4.959 | 20.689 | 5.985 | 22.610 | 7.763 | 0.552 | -25.660 | -10.871 | -3.318 | -14.811 | 19.011 | 11.057 | 7.870 | 15.455 |
| 20.596 | 25.694 | -6.268 | 25.633 | 22.586 | -12.897 | 18.533 | -1.116 | -15.654 | -26.095 | -16.217 | -12.766 | 19.864 | 22.999 |
| 22.072 | 16.026 | -10.003 | 8.787 | 25.830 | 12.496 | 8.117 | -8.586 | 13.860 | -29.788 | 1.796 | -7.005 | 17.586 | 25.641 |
| -16.463 | 21.536 | -1.518 | 2.500 | 17.049 | 0.898 | -2.053 | -18.910 | 9.037 | 25.111 | 8.346 | 21.284 | 10.472 | 10.179 |
| 25.589 | 9.890 | -9.060 | 15.316 | -12.799 | 25.587 | -9.910 | 11.686 | -0.487 | -30.619 | 21.710 | 8.329 | 8.860 | 21.024 |
| 24.854 | -18.564 | 3.493 | 9.808 | 28.638 | -6.146 | 5.979 | -23.400 | -4.415 | 23.616 | 0.448 | -26.611 | -11.561 | 23.788 |

| | | | | | | | | | | | | | |
|---|---|---|---|---|---|---|---|---|---|---|---|---|---|
| -13.996 | 7.544 | 6.518 | 19.648 | 23.885 | -16.127 | -9.033 | -15.832 | 24.558 | 21.346 | 24.026 | -20.161 | -18.163 | 16.768 |
| -17.313 | -9.396 | -14.456 | -25.097 | -9.940 | -24.600 | -1.108 | 22.362 | -10.186 | 29.563 | 15.735 | 26.983 | 20.270 | 14.680 |
| -22.786 | 0.093 | -6.873 | -20.827 | 23.142 | -23.402 | 18.064 | 18.736 | 17.671 | -14.036 | 7.247 | 19.864 | -23.640 | 11.322 |
| 12.024 | -23.994 | -0.767 | -1.270 | 20.917 | 2.530 | 14.807 | 4.741 | -25.638 | 22.173 | -15.923 | 13.390 | 23.624 | 14.914 |
| 4.472 | 11.149 | -20.754 | -15.683 | 16.555 | -18.861 | -9.522 | 1.821 | 2.337 | 15.963 | -12.945 | 16.913 | 12.751 | -1.093 |
| -12.280 | 24.435 | 18.600 | -22.006 | 27.823 | -21.330 | 25.719 | -22.299 | 27.938 | 19.302 | 9.070 | 22.920 | 25.533 | 14.407 |
| 0.893 | 9.428 | -1.877 | 15.913 | -10.182 | 0.652 | -6.369 | -9.672 | 26.415 | -23.034 | 4.717 | 17.540 | 0.646 | 9.564 |
| -19.694 | 19.984 | 6.475 | -0.123 | 3.964 | 2.251 | 2.190 | -12.376 | 25.232 | -30.299 | 21.820 | 25.493 | 19.026 | 13.956 |
| -18.893 | 2.481 | -1.481 | 4.614 | -9.464 | -8.987 | -5.379 | 11.239 | 24.046 | 24.735 | 26.146 | -3.074 | 15.792 | 18.019 |
| 3.233 | -6.399 | -0.652 | 18.667 | -15.220 | -25.457 | -26.777 | -19.844 | 10.477 | 27.891 | -24.935 | -7.799 | 17.500 | 16.088 |
| 25.629 | -22.921 | -9.674 | -19.802 | 20.226 | 10.242 | 8.323 | -21.195 | 0.796 | 24.068 | 14.589 | -0.315 | 8.012 | 10.881 |
| 7.605 | -7.413 | -0.440 | 5.228 | 26.246 | -7.459 | -3.641 | -12.137 | 27.470 | 17.007 | 19.431 | -20.236 | 14.760 | 17.303 |
| 8.644 | 13.534 | -7.629 | 11.376 | 17.868 | 25.471 | -7.306 | -23.522 | 3.393 | -26.944 | 17.271 | 21.801 | -6.376 | 14.259 |
| -21.766 | -9.608 | -8.506 | -7.344 | -26.590 | -18.633 | -22.072 | -19.405 | -5.539 | 21.722 | 22.218 | 1.008 | 17.806 | 5.998 |
| 14.938 | 12.979 | -21.712 | 21.339 | -8.028 | 14.711 | -19.631 | -24.378 | 21.867 | 6.867 | -18.608 | 0.624 | 19.602 | 15.475 |
| 10.192 | -12.664 | -18.908 | 19.933 | -22.851 | -6.706 | 10.716 | 1.602 | 17.622 | 27.183 | 21.597 | 7.525 | 27.719 | 4.389 |
| 12.031 | -3.647 | -1.781 | -3.842 | -27.080 | 1.205 | 8.931 | 12.256 | 5.301 | 24.443 | 17.815 | -3.102 | -9.865 | 19.132 |
| 0.165 | -20.974 | -8.936 | 4.096 | 17.742 | 4.403 | -24.403 | 3.061 | -18.619 | -18.412 | 3.940 | -27.750 | 22.404 | 27.027 |
| -20.552 | 19.786 | -17.536 | 17.872 | 10.056 | -3.280 | 3.992 | -10.005 | -20.696 | 12.031 | 6.302 | 23.912 | -3.012 | 22.766 |
| -2.186 | 3.894 | -1.639 | 8.304 | 5.902 | -15.711 | -7.618 | 15.608 | 19.192 | 23.893 | 6.816 | 18.578 | 26.757 | 17.077 |
| 5.381 | 16.465 | -5.001 | 13.236 | -17.853 | -14.308 | -11.332 | 21.573 | 21.466 | -19.636 | 23.024 | -19.061 | 5.982 | 12.780 |
| 23.621 | 4.559 | -3.377 | 1.499 | 22.199 | 26.033 | 22.460 | -9.001 | 9.819 | 17.988 | 2.269 | 15.034 | 22.870 | 13.793 |
| 15.671 | -8.831 | -0.421 | 5.236 | 14.477 | 7.852 | 6.732 | -17.640 | -0.907 | -4.218 | 26.331 | -1.326 | -17.614 | 13.348 |
| -21.480 | 13.225 | 2.314 | -15.798 | -8.066 | -10.000 | -14.637 | 19.326 | 18.622 | -4.920 | 17.567 | 26.684 | 21.214 | 23.799 |
| -10.381 | 19.596 | 0.404 | 1.078 | 16.951 | -24.692 | -8.123 | 23.453 | -4.493 | -13.812 | 1.861 | -14.514 | 28.219 | -0.808 |
| -3.025 | -21.335 | 10.747 | -7.594 | 1.322 | -8.354 | 19.721 | -22.385 | 8.388 | 22.934 | 17.016 | -3.020 | 20.044 | 11.716 |
| -23.782 | 2.303 | 18.988 | -1.543 | 5.297 | -0.799 | -12.559 | 3.965 | -2.454 | -25.216 | -5.575 | 26.740 | 8.070 | 18.143 |
| 1.065 | -4.613 | 13.422 | -2.910 | -2.348 | -25.874 | -0.252 | -3.193 | 17.285 | 26.370 | 26.884 | 15.258 | -1.324 | -5.624 |
| 4.944 | 3.249 | 12.485 | 17.801 | -11.376 | -23.714 | -4.202 | 7.639 | 12.397 | 25.202 | 1.491 | 6.778 | -27.200 | 9.128 |
| 0.565 | 11.736 | -2.828 | 10.102 | 2.108 | 18.809 | -2.606 | -17.566 | -28.052 | 18.317 | 23.593 | -1.779 | 27.482 | 23.188 |
| -22.527 | -16.014 | -5.312 | -0.477 | -1.818 | 2.627 | 6.317 | 25.374 | 23.346 | 22.329 | -13.353 | 8.098 | 23.396 | 14.438 |
| 9.065 | 4.421 | -19.493 | 6.660 | -18.072 | -1.354 | -15.498 | 21.865 | -10.690 | 20.508 | 18.508 | -9.688 | 21.689 | 18.376 |
| 14.334 | 10.485 | -6.341 | -1.134 | 18.325 | -16.968 | -0.570 | 5.077 | -16.847 | -19.275 | 3.912 | 20.860 | 24.379 | 10.534 |
| -0.014 | -11.509 | 4.248 | -11.751 | 9.204 | 5.364 | -13.104 | 13.267 | 22.752 | 8.434 | 6.558 | 19.471 | 21.019 | 20.771 |
| 15.824 | -8.028 | 16.543 | -25.459 | -16.701 | -18.224 | 3.651 | -13.402 | -10.130 | -11.255 | -20.994 | 3.457 | 4.398 | 6.640 |
| -21.075 | 9.454 | -4.239 | 26.580 | 22.145 | -22.726 | 2.176 | -21.893 | -0.861 | 30.066 | 21.453 | 23.099 | 24.839 | 21.803 |

| | | | | | | | | | | | | | |
|---|---|---|---|---|---|---|---|---|---|---|---|---|---|
| 4.226 | 3.111 | -14.783 | -6.651 | -18.849 | -17.672 | -11.233 | -2.379 | 16.581 | -20.248 | 24.330 | 22.449 | 19.328 | 6.154 |
| -8.958 | -14.623 | 18.413 | -5.599 | 22.897 | -4.193 | -26.252 | 17.944 | 7.494 | -16.532 | 28.660 | 16.180 | -23.554 | 20.113 |
| -18.835 | 19.434 | -15.887 | 4.062 | -3.681 | -13.311 | -13.609 | -22.273 | -7.946 | 7.865 | -5.246 | 17.204 | 13.646 | 16.844 |
| 25.107 | 2.161 | -20.761 | 1.463 | 23.787 | -8.736 | -4.014 | -19.647 | 26.059 | -29.756 | -25.922 | 14.658 | 12.438 | 23.402 |
| 19.261 | 21.916 | 8.887 | -3.853 | 7.834 | -15.866 | -6.734 | 21.578 | 1.730 | 31.192 | 27.019 | -0.186 | 0.936 | 12.050 |
| -20.260 | 20.850 | -22.157 | -22.685 | 11.420 | 7.474 | -2.469 | 24.101 | 26.923 | 24.040 | 27.874 | 17.644 | 4.793 | 15.830 |
| 15.240 | -15.867 | -24.666 | 12.783 | 3.902 | 2.398 | -10.219 | 5.673 | 7.024 | 25.846 | 26.172 | 5.326 | -0.663 | 14.306 |
| 2.897 | -3.756 | -3.798 | -24.315 | 12.390 | -22.276 | 5.662 | -23.083 | 24.666 | -17.026 | 18.670 | 4.482 | 19.931 | 19.749 |
| -12.829 | 9.398 | 1.971 | -11.656 | -2.409 | -0.728 | -0.841 | 10.441 | 29.183 | 23.475 | 23.565 | 14.281 | -14.199 | 20.696 |
| -22.410 | -3.396 | -2.707 | 25.642 | 22.860 | 8.750 | -23.346 | 1.415 | 8.117 | 22.298 | 21.266 | 24.332 | -15.594 | 14.883 |
| 23.579 | -20.045 | -23.638 | 6.392 | 5.159 | 5.173 | -20.433 | 24.551 | 22.840 | 28.413 | 22.153 | 21.299 | -5.469 | 18.984 |
| 22.700 | 11.062 | 25.269 | -17.251 | -0.959 | -3.147 | 9.826 | -12.947 | 21.590 | -12.056 | 24.786 | 7.781 | 10.516 | 4.019 |
| 13.943 | -1.755 | 5.709 | -21.222 | 21.055 | 12.191 | -5.767 | -13.924 | 13.072 | 29.177 | 27.514 | 13.622 | -7.037 | 17.353 |
| -21.749 | 3.610 | 16.399 | -15.571 | 14.608 | 9.078 | -10.893 | 19.066 | 25.721 | 20.982 | 22.392 | 11.397 | -28.125 | 16.662 |
| 15.996 | 1.621 | -24.367 | 3.557 | 0.877 | 0.530 | 26.373 | -18.346 | -24.017 | 18.494 | 0.477 | 4.976 | 8.575 | 11.880 |
| -24.266 | 20.615 | -23.031 | -0.744 | -2.970 | -2.448 | 23.115 | 12.343 | 16.755 | 26.857 | 12.508 | 10.874 | 20.700 | 20.974 |
| 25.193 | 17.185 | 18.963 | -7.993 | -12.270 | 7.539 | 4.224 | 23.793 | -15.308 | 10.156 | -6.187 | -7.177 | 26.639 | 4.933 |
| 8.047 | 6.654 | -7.182 | 9.353 | 11.376 | 19.371 | -0.379 | 16.432 | -4.648 | -17.882 | 7.524 | 9.004 | 13.966 | 12.762 |
| -4.106 | -3.457 | 4.503 | -14.616 | 22.982 | -17.483 | 1.904 | -23.027 | 21.612 | 24.713 | -2.642 | -14.140 | -2.243 | 7.872 |
| 22.711 | -14.365 | 9.809 | -12.073 | 22.195 | -11.168 | -22.188 | 24.472 | -18.505 | 23.313 | 16.536 | 2.268 | 25.651 | 6.672 |
| -2.024 | -1.671 | -16.675 | 26.261 | 6.011 | -9.718 | 4.646 | -2.338 | -20.248 | 15.913 | -15.474 | 15.536 | 24.187 | 17.543 |
| 2.429 | 7.946 | -25.335 | 19.852 | 17.803 | 2.172 | 8.051 | 22.167 | -24.373 | 26.672 | 20.237 | 19.349 | 21.999 | 14.892 |
| 16.343 | 6.085 | 21.723 | 2.940 | -11.900 | -21.788 | -16.287 | 14.321 | 28.409 | 12.656 | 15.778 | 11.524 | -4.825 | 12.551 |
| -6.626 | -16.584 | 7.976 | -10.420 | -0.418 | 24.507 | -2.456 | 3.666 | 4.391 | 28.858 | -22.927 | -2.027 | -2.550 | 11.436 |
| -8.397 | 13.071 | 2.678 | 11.001 | 23.987 | -3.321 | 2.720 | -22.250 | -24.997 | 16.334 | 15.112 | -18.822 | -11.762 | 10.823 |
| -19.662 | 11.748 | -0.821 | -23.371 | -14.423 | -6.350 | 23.103 | 9.567 | 19.748 | 5.796 | 18.152 | 18.440 | 22.754 | 20.698 |
| -20.225 | 0.110 | -8.685 | -25.839 | 3.393 | -24.810 | -11.968 | -9.453 | -11.809 | 20.432 | 6.268 | 8.244 | 15.866 | 13.094 |
| 19.235 | -4.635 | 17.199 | 21.578 | 18.217 | 17.492 | 2.053 | 15.146 | 26.223 | 7.203 | 18.962 | -20.192 | 1.831 | -18.356 |
| -26.148 | -2.820 | -24.284 | 1.430 | -1.622 | 16.835 | 26.341 | 11.497 | 6.252 | -17.231 | 26.946 | -6.089 | 21.008 | -17.079 |
| 21.859 | 2.865 | 9.801 | -24.123 | -4.046 | -24.250 | -21.054 | -16.887 | 26.899 | -20.876 | 29.088 | -0.133 | 11.092 | 9.960 |
| 10.563 | 20.104 | 11.128 | 24.152 | 13.343 | -1.628 | -0.735 | 3.350 | -23.914 | -1.557 | -1.138 | -4.968 | 8.092 | -16.529 |
| 3.424 | -18.238 | -3.548 | -21.360 | 8.714 | -15.912 | -6.876 | -24.140 | -1.336 | 22.131 | 25.236 | -19.528 | 26.297 | 23.827 |
| -13.434 | -9.811 | -24.120 | 22.074 | -4.616 | -6.931 | -12.059 | -26.677 | -19.638 | -16.197 | 17.805 | 25.565 | -17.038 | 8.806 |
| 23.325 | 19.926 | -4.251 | 23.482 | -18.329 | -10.604 | -28.822 | -20.216 | -7.678 | 28.067 | 20.152 | 2.767 | 23.458 | 3.968 |
| -0.986 | 2.207 | -14.750 | 27.327 | 22.717 | -7.982 | -2.042 | -18.959 | 20.712 | 30.157 | 19.759 | -11.802 | 3.716 | -2.789 |
| -9.555 | 11.372 | 10.853 | -19.674 | 11.607 | 27.132 | 6.622 | 15.554 | 23.006 | 17.355 | 11.066 | 9.328 | -22.289 | -14.183 |

| | | | | | | | | | | | | | |
|---|---|---|---|---|---|---|---|---|---|---|---|---|---|
| 15.369 | 22.421 | -13.074 | 9.793 | -21.610 | -5.741 | 27.349 | 24.202 | -14.981 | 10.835 | -2.802 | 14.149 | 18.653 | 1.954 |
| 10.226 | 3.034 | 14.302 | -18.324 | 10.999 | 3.284 | 26.639 | 18.134 | 4.589 | 7.399 | 24.865 | -7.410 | 25.188 | 16.059 |
| -22.679 | 13.856 | 3.923 | -26.854 | 10.618 | 0.999 | -28.659 | 19.104 | -16.513 | 18.447 | 28.140 | -17.740 | 30.995 | 25.183 |
| -6.535 | 20.198 | -10.382 | 26.447 | 8.953 | -0.449 | 18.054 | 4.974 | -20.738 | 21.389 | -24.095 | 5.838 | 28.787 | 22.716 |
| 0.509 | -15.652 | 17.008 | -2.536 | 8.308 | -8.970 | -11.875 | 18.140 | 28.852 | 22.867 | 5.949 | 12.907 | 29.881 | 22.217 |
| 24.565 | -7.233 | -0.114 | -24.852 | -2.609 | -23.153 | -0.938 | -0.394 | 10.225 | 14.319 | -11.359 | -2.233 | 20.985 | 19.466 |
| -21.963 | -6.753 | -2.249 | -23.574 | 13.501 | 26.232 | -2.052 | 12.021 | 5.311 | -28.208 | 25.523 | 17.431 | 14.264 | 21.716 |
| 23.322 | -14.344 | 17.392 | 13.616 | -18.953 | 22.898 | -15.647 | 13.713 | 27.820 | -14.798 | 23.120 | 16.428 | 19.469 | 0.731 |
| 20.636 | -23.968 | -12.686 | 23.519 | -17.099 | 16.082 | -9.078 | 0.234 | 21.674 | 18.119 | 20.098 | 19.393 | 26.177 | 7.477 |
| 3.289 | 21.272 | 28.324 | 24.715 | 23.133 | -10.519 | 4.105 | 28.260 | 29.385 | 13.688 | 27.004 | 11.672 | 14.841 | 10.856 |
| -20.581 | 16.094 | -23.168 | 27.864 | 10.290 | 1.739 | -27.220 | -23.940 | -19.961 | 8.138 | 16.834 | 10.392 | -1.744 | 20.421 |
| -25.758 | 25.381 | -25.883 | -16.526 | 21.670 | -13.137 | -23.129 | -1.910 | 26.196 | -16.816 | 22.422 | -8.860 | 25.686 | 21.116 |
| 19.266 | -11.762 | 2.914 | 28.213 | -2.656 | 25.573 | -29.126 | 22.140 | 13.893 | 19.985 | 23.921 | -11.067 | 24.929 | 10.709 |
| -4.000 | -21.305 | 1.272 | 19.473 | 25.501 | -15.479 | 8.727 | 12.679 | -30.034 | 3.918 | 1.722 | -20.858 | 21.504 | 19.698 |
| -10.391 | 22.399 | -3.295 | 27.003 | -22.195 | -15.159 | 4.526 | 25.509 | 30.339 | 27.430 | 19.232 | -24.747 | 4.958 | 3.191 |
| -24.848 | -0.927 | -2.741 | -17.195 | 5.787 | 15.081 | -0.817 | -4.573 | 1.091 | -17.495 | 18.006 | 22.946 | 10.420 | 25.754 |
| -23.514 | -4.624 | 22.268 | 14.014 | -9.254 | 17.310 | -14.982 | 24.760 | 19.542 | 25.079 | -22.228 | 10.025 | 17.965 | 17.345 |
| 5.351 | 1.152 | 17.653 | -8.852 | -26.424 | -6.177 | -17.836 | 11.911 | 16.607 | -23.545 | -23.624 | 8.477 | -21.194 | 6.041 |
| 4.834 | -7.551 | 4.972 | 7.954 | -16.188 | -11.334 | 19.329 | 10.806 | 7.208 | 16.248 | 25.864 | 23.384 | 6.516 | 21.330 |
| 27.275 | 1.709 | -8.469 | 4.192 | 2.608 | 5.696 | -29.734 | -18.272 | 5.860 | 25.524 | 27.130 | 16.653 | -14.140 | 3.947 |
| -26.842 | -9.118 | -11.869 | -23.487 | -14.581 | -3.878 | 26.554 | 1.222 | 11.136 | 21.754 | 12.391 | 4.664 | 20.721 | 22.974 |
| 24.503 | -9.956 | 27.623 | -27.584 | 20.805 | -18.784 | -1.815 | -6.351 | 20.303 | -8.859 | 21.628 | -13.851 | -13.178 | 24.605 |
| 4.484 | 21.436 | 0.628 | -17.805 | 17.446 | 27.287 | -6.036 | -15.810 | 2.403 | -14.956 | 30.380 | 18.759 | 26.740 | 14.102 |
| 20.490 | -4.558 | -0.113 | 6.701 | -11.565 | 23.628 | -16.310 | -3.526 | -18.751 | -6.451 | -5.999 | -17.236 | 13.686 | 26.603 |
| -24.323 | -20.394 | 2.800 | 13.828 | 17.738 | -0.074 | -4.527 | 17.059 | 8.513 | 14.956 | 24.535 | -18.745 | 31.357 | 22.413 |
| -19.005 | 1.951 | 15.832 | 18.529 | -6.495 | 8.766 | -21.432 | -5.836 | -14.454 | -8.056 | 7.027 | -14.569 | 27.589 | 23.417 |
| 27.458 | 20.355 | 15.079 | 16.082 | 7.636 | 19.781 | -18.977 | 4.034 | -18.288 | 28.594 | -22.693 | 14.834 | 18.130 | 16.721 |
| 11.308 | 25.980 | -24.292 | -20.640 | 13.184 | -27.376 | 11.350 | -19.602 | 18.148 | -10.275 | 17.122 | 1.795 | 29.132 | -2.093 |
| 23.923 | 17.958 | 6.587 | -22.423 | 11.159 | -1.725 | 7.039 | 26.307 | 21.878 | -18.646 | 29.747 | -5.827 | 9.519 | 21.456 |
| -10.061 | -6.513 | -4.399 | 24.138 | 7.386 | -5.519 | 23.056 | -25.874 | -3.233 | 9.817 | 22.765 | -21.409 | -16.937 | 20.616 |
| -8.946 | -5.806 | -19.107 | -1.398 | -25.163 | -21.092 | -14.248 | 7.336 | -29.933 | 18.916 | -11.755 | 14.157 | 12.468 | 7.078 |
| -27.525 | -3.577 | 6.940 | 28.839 | 6.653 | -20.066 | -25.103 | 22.127 | 20.875 | 2.313 | 15.643 | 10.607 | 30.835 | 14.523 |
| 0.663 | 7.342 | 3.078 | 18.656 | 1.250 | 22.080 | 24.204 | 2.669 | 19.024 | 21.686 | -21.542 | 22.585 | 24.943 | 26.917 |
| 21.485 | 21.570 | 8.199 | -0.297 | 5.864 | 13.598 | -11.088 | 0.417 | 14.011 | 12.051 | 25.690 | 16.984 | -17.676 | 15.300 |
| 11.899 | -1.276 | -1.034 | -17.714 | -17.680 | 2.761 | 0.501 | -13.578 | -2.235 | 11.520 | 31.071 | 9.609 | -22.634 | 18.752 |
| -5.579 | 17.638 | 6.176 | -11.205 | -26.944 | -18.009 | -25.747 | -5.010 | -26.397 | -5.471 | 24.110 | 11.359 | 18.747 | 10.352 |

| | | | | | | | | | | | | | |
|---|---|---|---|---|---|---|---|---|---|---|---|---|---|
| 14.565 | -2.807 | 28.591 | -25.334 | 15.117 | 12.374 | -11.818 | 7.860 | -0.390 | 23.446 | 5.480 | 12.468 | 14.159 | 19.657 |
| -2.578 | -18.492 | 8.694 | 24.916 | 9.468 | -7.386 | -1.363 | -28.874 | -8.816 | 11.562 | 19.613 | 4.549 | 17.802 | 10.915 |
| -26.532 | 5.914 | 2.144 | -8.896 | 14.284 | -9.985 | 17.976 | -15.099 | 25.316 | 27.253 | -1.361 | -10.130 | 20.028 | 25.645 |
| -4.243 | -10.770 | 29.929 | -15.430 | -21.444 | -17.086 | 2.566 | 0.696 | 19.743 | 22.686 | 28.807 | 7.695 | 15.075 | 21.445 |
| -0.859 | -9.993 | -16.160 | 13.180 | 26.215 | -2.512 | 15.567 | 23.801 | -16.617 | 15.415 | 25.596 | -11.917 | 22.059 | 5.502 |
| 13.788 | -18.054 | -9.943 | -21.718 | 15.592 | 1.968 | -16.832 | 18.214 | 30.011 | 21.901 | 22.006 | -21.960 | 27.653 | 17.613 |
| -24.319 | -26.544 | -13.899 | 9.156 | -8.556 | -26.338 | -17.233 | 3.398 | 26.777 | 21.223 | 26.080 | -7.881 | 23.186 | 25.294 |
| -6.426 | 8.557 | 24.853 | -17.091 | -0.758 | 4.964 | -30.943 | 22.025 | -3.525 | 3.651 | -8.783 | 12.918 | -18.544 | 14.681 |
| 27.365 | 18.820 | 4.849 | 24.027 | 3.382 | -9.534 | -25.532 | -1.950 | 5.732 | -6.535 | 20.788 | 24.568 | 21.354 | -1.471 |
| 24.452 | 15.548 | -3.471 | 29.212 | -23.904 | -5.204 | -13.849 | 27.711 | 16.479 | -20.883 | -21.556 | 5.199 | 24.123 | 9.464 |
| 25.653 | 6.093 | -7.256 | -4.325 | -18.421 | -8.387 | 14.935 | -17.256 | 24.040 | 15.639 | 1.164 | 10.088 | 3.210 | 7.241 |
| 18.655 | -7.479 | -24.593 | -1.185 | 28.332 | -3.399 | 0.732 | 20.402 | -14.752 | 26.231 | 27.965 | 25.235 | 2.640 | 24.141 |
| 5.348 | 12.876 | -4.493 | 25.857 | -15.675 | -25.845 | 28.351 | 15.343 | 6.395 | -24.018 | 19.372 | 21.229 | 16.028 | 27.659 |
| 24.058 | 28.561 | -6.659 | -9.205 | -19.321 | 17.253 | 2.033 | 14.182 | 18.926 | -23.716 | 21.014 | 20.515 | 22.621 | 22.912 |
| -17.862 | 16.880 | -5.257 | -10.331 | 4.973 | 11.405 | -20.494 | -3.158 | 23.180 | 25.513 | 15.800 | -3.634 | 24.790 | 20.082 |
| 23.359 | -28.571 | -1.236 | 3.705 | 5.315 | -0.023 | 19.318 | -7.791 | -7.984 | 12.430 | 17.593 | 9.154 | 8.681 | 15.924 |
| -1.287 | 7.095 | -5.554 | -14.271 | -7.786 | 3.114 | 6.733 | -9.994 | -23.507 | -3.614 | 24.185 | 2.938 | 27.222 | 17.088 |
| -25.180 | 24.387 | -10.453 | -23.724 | 6.318 | 10.539 | -10.743 | 18.807 | 7.801 | -9.032 | 18.939 | -5.877 | -0.488 | 26.728 |
| -25.923 | -23.553 | 3.325 | 27.207 | -0.969 | 8.907 | 26.955 | 3.002 | 28.312 | 13.197 | 27.311 | 22.652 | -27.049 | 19.360 |
| 2.372 | -2.253 | -22.005 | 25.653 | 7.063 | 0.075 | -4.026 | 28.467 | -11.909 | -23.117 | -15.044 | 24.798 | 29.704 | 19.169 |
| -9.529 | 28.087 | 0.304 | 24.689 | -8.455 | -20.736 | -15.908 | 2.733 | 25.540 | 14.342 | 21.566 | 10.783 | 28.536 | 20.491 |
| 18.628 | 23.135 | -15.477 | -27.594 | 13.700 | -15.268 | -6.766 | -25.182 | 21.931 | 18.968 | -28.794 | 21.775 | 18.780 | 13.400 |
| -4.005 | 23.978 | 19.544 | -20.278 | -9.563 | 28.883 | -1.453 | 16.837 | 4.866 | -0.541 | 16.667 | 13.664 | 23.262 | 17.899 |
| 22.869 | 4.031 | 25.820 | 3.944 | 7.628 | 8.091 | -22.555 | 9.121 | -2.447 | -23.390 | 18.077 | 18.367 | -7.599 | 19.121 |
| 1.843 | 25.225 | 23.521 | 22.338 | 4.851 | 2.670 | -7.801 | -3.828 | 3.081 | 10.790 | 7.179 | 16.541 | -7.120 | 11.528 |
| 4.138 | 0.334 | 4.667 | 17.633 | -5.762 | -3.960 | -4.728 | 6.032 | -0.431 | 18.007 | 30.090 | 24.666 | 27.365 | 28.739 |
| 26.327 | -7.421 | -7.453 | -25.842 | 6.766 | 19.581 | -3.058 | 23.901 | 0.577 | 19.728 | 12.033 | 18.915 | 13.280 | 30.430 |
| 2.155 | -25.081 | -20.134 | 7.257 | -16.766 | 5.902 | 1.888 | -13.951 | -1.419 | -2.854 | 0.827 | 24.159 | 22.971 | 28.045 |
| -28.153 | -8.335 | 18.076 | 23.863 | -21.427 | -7.229 | -5.585 | 0.019 | -5.137 | -23.787 | 22.100 | 7.418 | 19.534 | 24.272 |
| 25.620 | 6.331 | -1.444 | 21.306 | 11.593 | -19.418 | -30.683 | 22.735 | 18.079 | 8.156 | 25.771 | -17.798 | 16.231 | 10.399 |
| 0.713 | -13.926 | -5.216 | -16.176 | -15.077 | -7.584 | -3.827 | 28.526 | 17.475 | 6.057 | 20.272 | 22.476 | 8.529 | 24.727 |
| -16.827 | 3.739 | 27.475 | 20.256 | -14.403 | 17.908 | 9.775 | -14.628 | 16.574 | 10.296 | 23.267 | 12.432 | 15.833 | 23.742 |
| 7.292 | 16.415 | 1.461 | 13.271 | -4.744 | 9.360 | 3.466 | -18.631 | -6.114 | 21.265 | 18.853 | 25.859 | -7.739 | 18.096 |
| -24.230 | 25.896 | -28.222 | -8.356 | 13.392 | -3.333 | -22.711 | -21.967 | -16.290 | 23.531 | 28.668 | 21.894 | 5.508 | 19.923 |
| -25.030 | -5.508 | -18.352 | -9.372 | 24.053 | -10.968 | -5.818 | -13.161 | 13.306 | -12.097 | 19.409 | 25.329 | 23.564 | 23.288 |
| -1.096 | -9.871 | -6.309 | -24.262 | 7.118 | -13.042 | 12.456 | -2.862 | -31.648 | 4.289 | 25.566 | 16.076 | 9.771 | 7.998 |

| | | | | | | | | | | | | | |
|---|---|---|---|---|---|---|---|---|---|---|---|---|---|
| -26.388 | -5.856 | -7.955 | -21.180 | 25.489 | -2.251 | 7.564 | -7.743 | 11.263 | 20.168 | 26.799 | 23.816 | -30.853 | 19.318 |
| 27.398 | 14.566 | -12.150 | 2.879 | 24.727 | 19.073 | -3.406 | -9.006 | -25.073 | 9.737 | 20.061 | 18.170 | 19.773 | 21.747 |
| -23.622 | -6.945 | 7.555 | -28.428 | -1.633 | 20.777 | 11.086 | -15.054 | -2.363 | 9.015 | 20.897 | -21.808 | 6.746 | 22.432 |
| -9.815 | 9.742 | 2.504 | -18.051 | 19.765 | 1.010 | -10.322 | 26.887 | -27.881 | 15.847 | 13.234 | 24.497 | 2.099 | 6.786 |
| 16.106 | -27.380 | -14.899 | -15.406 | 6.320 | -25.300 | 6.180 | -2.476 | 26.124 | -19.900 | 30.201 | -2.147 | 7.451 | 28.838 |
| -16.064 | 26.714 | 14.920 | -26.550 | -28.994 | -23.431 | 8.717 | -26.294 | 7.276 | 15.221 | 24.530 | 21.087 | 26.852 | 30.085 |
| -22.788 | -22.263 | -10.722 | 17.543 | 27.702 | 15.413 | -28.721 | 21.310 | 0.224 | 5.902 | 22.238 | 16.444 | 13.622 | 16.446 |
| -11.764 | 0.062 | 18.375 | 23.336 | -1.963 | 6.575 | -29.000 | -17.570 | -15.143 | 12.288 | 15.773 | -12.056 | -13.864 | 23.149 |
| -4.588 | 24.595 | 9.462 | -29.815 | -27.455 | -26.685 | -9.027 | 2.992 | 3.683 | -21.849 | 4.836 | 5.649 | 20.025 | 21.053 |
| 1.695 | -5.973 | -7.436 | -19.055 | 13.328 | -17.634 | 3.316 | 22.686 | 23.291 | 4.871 | 20.807 | -1.599 | -10.906 | 21.843 |
| 7.068 | -6.521 | 30.991 | 12.193 | -25.954 | -17.421 | -15.174 | 16.656 | -0.709 | -17.032 | 28.549 | 26.534 | -24.426 | 15.225 |
| 24.896 | 29.810 | -2.624 | -14.600 | 29.496 | -29.192 | -0.685 | -4.106 | -26.211 | -4.302 | 23.889 | -5.382 | 27.951 | 23.671 |
| 2.128 | 31.266 | 22.049 | 16.757 | 4.906 | -28.517 | -6.393 | -21.385 | 17.029 | 23.332 | -1.568 | 14.566 | 15.809 | 17.594 |
| 3.752 | 8.291 | 0.205 | -30.413 | 24.977 | -2.971 | 25.184 | -26.595 | 8.175 | 6.825 | 26.001 | -22.536 | 8.346 | 18.929 |
| 0.015 | 17.871 | -0.546 | 4.669 | 18.966 | -24.443 | 6.168 | 7.191 | 16.339 | 5.458 | 19.237 | 19.759 | 21.324 | 18.040 |
| -14.928 | 9.712 | -10.224 | 19.227 | 23.779 | 22.289 | 3.469 | -27.118 | 28.376 | -22.609 | 28.704 | 19.118 | -3.370 | 16.991 |
| -17.177 | -3.447 | 1.950 | 28.268 | -23.447 | -17.288 | 26.770 | -9.283 | 29.375 | -25.395 | 16.696 | 23.157 | 11.130 | 19.744 |
| 5.775 | -7.880 | -25.703 | 22.682 | -11.194 | -16.930 | 21.812 | 13.174 | 6.133 | 24.771 | 6.244 | 21.752 | -7.632 | 25.366 |
| 21.948 | -21.148 | -21.313 | 15.625 | 16.864 | -6.677 | 5.425 | -17.912 | -18.776 | -1.807 | 20.925 | 20.295 | 25.547 | 24.009 |
| 12.657 | 17.919 | 3.248 | -16.989 | 19.150 | 19.893 | 14.764 | 15.239 | 18.807 | -20.140 | -22.305 | 4.214 | 15.971 | 28.816 |
| 26.722 | -1.666 | 6.337 | 1.694 | -9.665 | 27.919 | -20.836 | -4.225 | -31.203 | 4.700 | 24.837 | 21.312 | -4.295 | 14.472 |
| -6.959 | 21.231 | -1.095 | -16.366 | 10.608 | -25.444 | 19.855 | 25.762 | 15.414 | 12.537 | 19.774 | 26.935 | -11.327 | 30.784 |
| -27.015 | 5.766 | -1.563 | -20.629 | -27.363 | -27.307 | 1.258 | 20.059 | 8.755 | 18.106 | -13.460 | 23.872 | -6.182 | 13.274 |
| -8.354 | 28.998 | 28.728 | -12.522 | -20.543 | 9.270 | -25.818 | -12.930 | 14.052 | -24.947 | -19.082 | 22.440 | 16.496 | 17.875 |
| -23.949 | 23.849 | -5.328 | 2.275 | 13.952 | 2.536 | -10.206 | 30.741 | -6.796 | 23.705 | 24.597 | 8.589 | 10.014 | 15.144 |
| -24.781 | 3.520 | 9.233 | -2.498 | 15.514 | -28.110 | -5.595 | -4.645 | 23.501 | 13.540 | 18.765 | 12.862 | 23.115 | 27.492 |
| -27.727 | -16.278 | -5.841 | 7.444 | -16.905 | -17.631 | -26.820 | -8.387 | 12.149 | 2.891 | 25.421 | 25.588 | 22.179 | 23.553 |
| 18.587 | -19.394 | -16.573 | 11.069 | -5.310 | -20.214 | -5.041 | -25.340 | 16.912 | -26.220 | 29.447 | 11.244 | 15.254 | 21.848 |
| 26.008 | -12.484 | -13.676 | 6.290 | 21.082 | 7.618 | 1.885 | 17.172 | 24.889 | 21.062 | -28.087 | 0.063 | 24.907 | 16.590 |
| -20.621 | 28.016 | 17.171 | -6.161 | 27.006 | -7.563 | -7.530 | 29.738 | 0.143 | -8.312 | -18.137 | 14.806 | 27.619 | 10.226 |
| 14.361 | 23.964 | -16.982 | 20.207 | 26.533 | -14.901 | 1.384 | -13.795 | -11.334 | -21.937 | -20.513 | 13.021 | 19.729 | 15.955 |
| 24.695 | 10.160 | -19.272 | 16.880 | 5.490 | -0.602 | -2.476 | 22.350 | -20.081 | 17.074 | 21.509 | -0.968 | 10.100 | 17.761 |
| 19.956 | 31.540 | -17.746 | 15.224 | -3.279 | 15.903 | -19.016 | 14.914 | -27.637 | -7.570 | 14.434 | -4.332 | 19.142 | 20.238 |
| 10.377 | -30.324 | 7.096 | -25.825 | 22.363 | 10.187 | 15.365 | -24.085 | -29.160 | -1.378 | 23.542 | 24.097 | 17.728 | 16.177 |
| -15.237 | 26.449 | 5.251 | -26.789 | 24.037 | 8.599 | -13.168 | -19.976 | -17.407 | 4.037 | 22.723 | -24.213 | 6.691 | 22.469 |
| -16.188 | 7.938 | -7.434 | 20.638 | 14.401 | 13.502 | -2.141 | 26.146 | 10.862 | 8.593 | 9.141 | 15.893 | 5.549 | 24.311 |

| | | | | | | | | | | | | | |
|---|---|---|---|---|---|---|---|---|---|---|---|---|---|
| 1.048 | 7.034 | -8.408 | 14.097 | -7.621 | -25.180 | -15.937 | 19.533 | 7.644 | -22.274 | 19.450 | -18.262 | -29.153 | 25.259 |
| 0.114 | 22.614 | 0.664 | 31.290 | -26.721 | -1.862 | -4.014 | -25.637 | 20.246 | -26.534 | -1.842 | 21.361 | 26.012 | 19.890 |
| -20.055 | 11.376 | -1.898 | -0.255 | -6.616 | 5.981 | -15.191 | -30.612 | -3.359 | 7.951 | 27.454 | 11.052 | 18.535 | -4.717 |
| -2.223 | -13.685 | 13.335 | -22.948 | 21.537 | -26.194 | 3.658 | 9.737 | 17.319 | -10.491 | 21.039 | 16.657 | -18.088 | 18.588 |
| 18.956 | 17.448 | -6.861 | -25.072 | -5.198 | -17.567 | -6.823 | 8.592 | 13.905 | 2.109 | 8.724 | 22.258 | 22.967 | 23.861 |
| -28.985 | -5.528 | -24.057 | 7.563 | 22.033 | -16.516 | -12.412 | 17.349 | 30.066 | 25.975 | 18.372 | -23.213 | 21.362 | 15.055 |
| -25.002 | -6.854 | 29.160 | 9.556 | -22.710 | 29.168 | 18.038 | -22.975 | -22.103 | 21.027 | 19.566 | 10.270 | 22.334 | 27.037 |
| 23.080 | 21.199 | -14.919 | -30.841 | 7.647 | -13.992 | -3.275 | 7.687 | 24.734 | 7.061 | -4.776 | 13.098 | 14.743 | 16.335 |
| 11.945 | -28.529 | -31.045 | 6.057 | -11.032 | -0.439 | -21.592 | 28.384 | 21.879 | 24.949 | 14.684 | 23.696 | 12.677 | 6.913 |
| 2.649 | 18.804 | -12.944 | 1.038 | 14.978 | -21.221 | -8.020 | -20.463 | -18.816 | 10.172 | 24.536 | 12.202 | -8.416 | 28.889 |
| -14.385 | 20.294 | -4.852 | -4.801 | -6.085 | 22.030 | 4.917 | 20.987 | 6.023 | 10.705 | 23.064 | 2.467 | -16.892 | 8.737 |
| 15.294 | -7.648 | -3.153 | -19.873 | 13.152 | -26.678 | 17.046 | -7.847 | -25.142 | -2.996 | -4.009 | 18.199 | 27.509 | 24.633 |
| -0.879 | 22.546 | 13.875 | 19.579 | 20.538 | -11.817 | 19.789 | 27.022 | 17.687 | 5.199 | 1.106 | 19.653 | 27.083 | 22.133 |
| 3.125 | 31.322 | 5.357 | 15.086 | 4.579 | 3.221 | -15.823 | -18.336 | 26.440 | 2.813 | 5.275 | 20.322 | -19.061 | 17.636 |
| 5.334 | -6.006 | 25.577 | 18.093 | -14.908 | -0.959 | -22.617 | 4.169 | 28.220 | 22.010 | 22.130 | 8.222 | 16.370 | 26.724 |
| -17.795 | -13.544 | 30.424 | -14.510 | 19.500 | 25.371 | -20.521 | -7.595 | -3.826 | 14.731 | -1.469 | 17.288 | 17.012 | 20.918 |
| 7.979 | 18.888 | 1.184 | 17.183 | -9.387 | -13.534 | -11.475 | 30.969 | -17.579 | -3.409 | 5.795 | -15.302 | -14.653 | 24.486 |
| -26.470 | -26.091 | -6.966 | -16.802 | 18.226 | 6.522 | -17.168 | 12.367 | 23.765 | -19.503 | 7.864 | 23.278 | 25.277 | 14.486 |
| 25.753 | 28.046 | 13.340 | -21.848 | -25.480 | -14.947 | 0.478 | -25.747 | 21.350 | -1.398 | 21.145 | 19.636 | 10.207 | 16.565 |
| 1.660 | -16.321 | -21.017 | -23.558 | 16.413 | 8.448 | -23.567 | -0.247 | 7.850 | 19.019 | 22.583 | 14.810 | -28.009 | 17.378 |
| -9.542 | 19.483 | 23.752 | 27.618 | 25.231 | -18.371 | 4.755 | -4.955 | 4.555 | -25.085 | 18.696 | 24.988 | 24.409 | 16.061 |
| -22.188 | -5.372 | 6.541 | 14.643 | -16.684 | -24.507 | 24.076 | 31.733 | -29.395 | -21.206 | 16.322 | 16.349 | -27.549 | 27.257 |
| -7.303 | 16.296 | -31.178 | 12.704 | 29.366 | 11.505 | 20.974 | 6.119 | -13.524 | 23.765 | -13.707 | 14.011 | 18.482 | 17.553 |
| 3.707 | 3.015 | 0.978 | -4.227 | -10.043 | -22.953 | -22.045 | 29.356 | 27.915 | -26.962 | 20.531 | -24.644 | -20.011 | 3.168 |
| 6.722 | 30.510 | -4.268 | -0.452 | 7.102 | -2.085 | 19.396 | -4.211 | 22.481 | 17.481 | 9.323 | 24.484 | 21.466 | 22.503 |
| -19.101 | -3.235 | -17.473 | 28.852 | -3.025 | -29.242 | -7.438 | 20.414 | -4.941 | -23.712 | 8.212 | 28.052 | 23.387 | 18.299 |
| -1.941 | 6.409 | 28.095 | 2.949 | 9.970 | -20.159 | 23.682 | 6.615 | 13.814 | 27.224 | 14.334 | 22.879 | 14.769 | 27.685 |
| -6.188 | 29.908 | 2.159 | 30.548 | -23.346 | -9.088 | 2.449 | -2.571 | -2.334 | 5.738 | 21.157 | 19.513 | -28.580 | 25.997 |
| -28.502 | -6.550 | 3.747 | 8.526 | -15.662 | 26.940 | 0.652 | 22.361 | 21.080 | 21.426 | -14.069 | 26.253 | 19.529 | 28.068 |
| 26.536 | 16.770 | -23.614 | -29.438 | 16.116 | -2.264 | 21.478 | -18.536 | -11.900 | 26.054 | 22.980 | 23.843 | 12.726 | 19.420 |
| 1.253 | 18.676 | -1.638 | 4.980 | 9.550 | -7.396 | 25.264 | 11.572 | 26.813 | -17.921 | -4.559 | 18.610 | 8.726 | 21.926 |
| -12.185 | -9.639 | -4.889 | 14.945 | -29.514 | -13.991 | -13.114 | -26.852 | -14.011 | -24.117 | 18.444 | 5.408 | 24.723 | 20.913 |
| 28.315 | -15.264 | -8.031 | -22.605 | 20.263 | 18.017 | -22.126 | 24.722 | -7.279 | 16.328 | 10.811 | 19.645 | 22.188 | 22.589 |
| -0.306 | -23.505 | -12.521 | -28.510 | 28.359 | -17.500 | 14.564 | -22.731 | 26.354 | 17.049 | 12.232 | 14.025 | -0.335 | 3.942 |
| -15.383 | 26.708 | 19.631 | -30.299 | 19.863 | 14.715 | -6.523 | 19.655 | 23.716 | 14.828 | 15.936 | 21.333 | 20.196 | 8.641 |
| 5.570 | 27.608 | -0.592 | 30.165 | 6.849 | 9.967 | 8.272 | 27.137 | -0.677 | -14.144 | -12.197 | 18.803 | 26.443 | 24.672 |

| | | | | | | | | | | | | | |
|---|---|---|---|---|---|---|---|---|---|---|---|---|---|
| 16.991 | 28.653 | -1.265 | -8.020 | -8.651 | 6.071 | 7.474 | 7.162 | 20.340 | -15.798 | 9.169 | 24.059 | 21.368 | 11.974 |
| 14.759 | 29.378 | -6.906 | 24.290 | -27.719 | -5.472 | -16.505 | 9.393 | 13.432 | 9.311 | 2.820 | 24.389 | 10.371 | 26.725 |
| 4.588 | -1.720 | 9.478 | 23.115 | 22.612 | -21.992 | 25.427 | -24.439 | -9.312 | 21.914 | 9.483 | 17.244 | -24.300 | 26.145 |
| -11.649 | 25.996 | -3.169 | -12.415 | 23.200 | -19.407 | -7.554 | 1.271 | 9.174 | 3.830 | -9.247 | 17.832 | 20.524 | 27.324 |
| -8.872 | -28.880 | -17.669 | 26.888 | -25.706 | 21.815 | -15.730 | 28.362 | -26.493 | 28.182 | 14.873 | 25.564 | 9.497 | 22.041 |
| -19.561 | -4.410 | 1.232 | -19.454 | 22.116 | -13.302 | -12.146 | 20.522 | 20.956 | 7.780 | 19.082 | -28.423 | -2.646 | 15.976 |
| 20.681 | 5.161 | 7.932 | -7.431 | -1.297 | 28.572 | -3.324 | 0.104 | -28.053 | 28.983 | 14.164 | -15.099 | 25.199 | 9.495 |
| -11.238 | 19.014 | 13.991 | 17.364 | -20.366 | 31.113 | -7.920 | -7.173 | -9.023 | 15.775 | 21.008 | 27.691 | 24.151 | 24.566 |
| -28.255 | -17.291 | -24.765 | -31.876 | -7.175 | -8.598 | 22.143 | -9.539 | 25.387 | -9.712 | 14.596 | 20.994 | 6.607 | 18.944 |
| 21.983 | 22.633 | 22.002 | 22.333 | -2.417 | 13.964 | -3.074 | 3.998 | 21.371 | 7.353 | 21.472 | 24.268 | 21.459 | 21.991 |
| 5.027 | 29.157 | 3.546 | 14.864 | 11.366 | 31.903 | 8.753 | -26.268 | 17.222 | -3.433 | 16.452 | -9.241 | 15.804 | 21.516 |
| 24.820 | 16.376 | 7.537 | -15.666 | 4.137 | -27.100 | -26.571 | -26.721 | 27.718 | 27.590 | 15.717 | 16.031 | 2.280 | 17.096 |
| 21.474 | -4.175 | -20.546 | 6.375 | -5.716 | 23.487 | 14.372 | -27.165 | -27.472 | 13.473 | 23.346 | -27.584 | 4.154 | -3.269 |
| -25.748 | 27.419 | 19.544 | 21.011 | 23.462 | 6.664 | 1.281 | 9.576 | -5.338 | 2.398 | 26.009 | 14.980 | 10.599 | 15.704 |
| -4.446 | -7.672 | 0.158 | -11.866 | 9.431 | -4.550 | -0.650 | 27.036 | -17.882 | 10.741 | 18.184 | 27.523 | 22.607 | 12.096 |
| 8.485 | 19.599 | -2.581 | 25.403 | -29.841 | 7.033 | -17.644 | 19.662 | -1.467 | -22.311 | 20.101 | -3.972 | 20.089 | 20.202 |
| 5.835 | -10.696 | -2.087 | 18.754 | 7.626 | 16.433 | -0.950 | 25.981 | -4.150 | 14.465 | 15.665 | 1.039 | 17.709 | 19.075 |
| -9.442 | -1.235 | 20.692 | -13.006 | 14.167 | -6.364 | -10.859 | -27.610 | 23.547 | 22.242 | 11.974 | 20.177 | 7.470 | -5.739 |
| 23.834 | 22.791 | 17.268 | -11.245 | -28.023 | -15.932 | 17.647 | -24.925 | -22.908 | -22.633 | 0.888 | 10.179 | 19.118 | 23.214 |
| -0.329 | -21.251 | 8.556 | -1.426 | 27.032 | 19.152 | -5.766 | -25.370 | -23.851 | 23.592 | -8.150 | 25.694 | 16.787 | 16.120 |
| -23.320 | 6.671 | 23.959 | -26.259 | -26.722 | -5.722 | -11.541 | -28.218 | -18.525 | 16.060 | 8.040 | 18.225 | 24.253 | 14.822 |
| 18.244 | -17.961 | -2.939 | 4.143 | 12.736 | -11.426 | 7.451 | 8.160 | -24.736 | 27.847 | -2.868 | 14.200 | 26.356 | 21.151 |
| -2.421 | 22.878 | 5.107 | 29.279 | 23.184 | -11.953 | -13.108 | 24.249 | -6.495 | 23.355 | -3.894 | -28.197 | 12.637 | 15.562 |
| 20.146 | -7.816 | -5.231 | -21.419 | 24.064 | 17.896 | -3.238 | 25.392 | 26.394 | 14.785 | 7.130 | 27.926 | -8.310 | 13.697 |
| -3.014 | -19.418 | 11.763 | 9.971 | -2.492 | -2.955 | -14.835 | -17.672 | 14.573 | -18.042 | 17.393 | 22.329 | -2.472 | 24.084 |
| -24.392 | 18.764 | -6.143 | -25.400 | -1.396 | -20.276 | 3.481 | 11.566 | -26.595 | 20.910 | -2.396 | 15.984 | -20.657 | 18.415 |
| -23.566 | 1.642 | -7.899 | 10.796 | 22.100 | -22.533 | 27.276 | 21.262 | 25.633 | 26.570 | 21.698 | 23.996 | 9.551 | 23.232 |
| 16.153 | -1.211 | -8.436 | -18.133 | 7.627 | 15.733 | 5.824 | 16.327 | 11.534 | 26.166 | 1.289 | 9.227 | 13.754 | 25.547 |
| 19.475 | 27.307 | -13.925 | 13.898 | -8.435 | 12.922 | 19.283 | -16.533 | 12.034 | 16.012 | 24.765 | 26.340 | -4.906 | 25.642 |
| 17.901 | -2.181 | -17.266 | -6.881 | 20.115 | 28.594 | 12.497 | 12.229 | -24.679 | 30.815 | 5.124 | 14.723 | 25.635 | 22.560 |
| 11.148 | 23.237 | -1.130 | -20.268 | -23.556 | -16.462 | 1.725 | 0.049 | 17.535 | 25.247 | 22.612 | -28.619 | 23.806 | 20.427 |
| -20.427 | 4.708 | 3.267 | 19.368 | 22.073 | -21.011 | 2.485 | -24.282 | -3.256 | -9.230 | 16.949 | 12.863 | 2.680 | 17.495 |
| -7.274 | 1.075 | -21.978 | 7.831 | 14.873 | -25.671 | -20.512 | 13.269 | 7.894 | -9.010 | 15.989 | -1.782 | -6.508 | 22.735 |
| 22.194 | -1.513 | -5.865 | 5.046 | -8.391 | 26.292 | 18.417 | -14.299 | -1.117 | 23.882 | 17.740 | 22.081 | -25.568 | 19.020 |
| -30.482 | 22.395 | 17.040 | -26.995 | 24.702 | -27.902 | -0.164 | -0.588 | 26.845 | 18.030 | 23.958 | 24.682 | -12.570 | 18.464 |
| -18.925 | 5.565 | -21.253 | 21.419 | -26.403 | -31.091 | 0.760 | -21.317 | -7.736 | -29.441 | 26.297 | 7.974 | 11.020 | 0.369 |

| | | | | | | | | | | | | | |
|---|---|---|---|---|---|---|---|---|---|---|---|---|---|
| 24.092 | -19.493 | 5.058 | 11.241 | -28.279 | 18.375 | -2.749 | 28.614 | 5.466 | 14.093 | 24.665 | 27.167 | 14.458 | 19.765 |
| 20.902 | -21.818 | 2.127 | 0.441 | -10.131 | 17.183 | 2.843 | 26.668 | 14.805 | 25.288 | -21.949 | 23.579 | 22.981 | 9.726 |
| -25.262 | 21.637 | 0.566 | -18.766 | -5.654 | -10.060 | -1.492 | 11.280 | 13.345 | -2.416 | -2.516 | 14.831 | 1.030 | 17.163 |
| -29.360 | 27.014 | -2.719 | -5.584 | 8.096 | -28.946 | -6.559 | -17.730 | 8.515 | -1.399 | -6.559 | 7.273 | -20.168 | 7.451 |
| 3.580 | -23.903 | 24.889 | -23.881 | -4.780 | 2.287 | 6.142 | -25.196 | -6.840 | 9.812 | 21.872 | -23.846 | 15.389 | 5.274 |
| 16.048 | -6.875 | 9.260 | -11.428 | 8.819 | -20.687 | 3.764 | 16.248 | 23.218 | -28.016 | -18.298 | 18.131 | 24.315 | -17.831 |
| 5.267 | -9.835 | -5.646 | -10.123 | 8.317 | 4.266 | 26.402 | -23.477 | 9.592 | 21.200 | -7.167 | 20.401 | -4.725 | -18.148 |
| -3.255 | 3.552 | 3.462 | 8.585 | 23.369 | 7.235 | 20.506 | 25.526 | -2.321 | 26.660 | 28.160 | 27.864 | 8.054 | -20.595 |
| 8.658 | -2.942 | -12.737 | -3.858 | -10.147 | 1.360 | -13.008 | 29.042 | 3.860 | 19.353 | -23.633 | 11.073 | -1.940 | 24.263 |
| 29.035 | -4.202 | 15.360 | 26.664 | -23.685 | -18.307 | 15.136 | 24.210 | 21.977 | 24.413 | 16.619 | 21.450 | 10.648 | 8.809 |
| 29.612 | 18.043 | -17.334 | -17.927 | 9.404 | 8.061 | -9.553 | 19.136 | -27.410 | -22.498 | -11.977 | 21.224 | 6.134 | 21.489 |
| 2.435 | -3.702 | 8.222 | 14.495 | 27.272 | -25.130 | -12.269 | -27.510 | -26.004 | 10.046 | 21.495 | 5.871 | 19.967 | 5.939 |
| -28.366 | 26.380 | -11.686 | -2.436 | 14.027 | -8.117 | 6.432 | 21.947 | 15.540 | 24.060 | 20.800 | -7.614 | -3.336 | 13.258 |
| 27.742 | -9.867 | -4.285 | -8.070 | 8.445 | 30.749 | -11.012 | 0.623 | -24.924 | -21.040 | -11.353 | 21.841 | 1.778 | 16.119 |
| 26.074 | -12.611 | -23.077 | 17.403 | 22.943 | -16.383 | 3.570 | 18.275 | 2.818 | 23.575 | 7.250 | 25.832 | 15.095 | 8.090 |
| -0.834 | -1.677 | 17.442 | 1.835 | 19.481 | 9.731 | -24.099 | 15.344 | -23.679 | 20.222 | 19.454 | 19.149 | -9.206 | 6.572 |
| -27.258 | 21.552 | -6.066 | 19.247 | -13.417 | -4.456 | -4.250 | 14.810 | 5.381 | -3.582 | 10.793 | 14.148 | -5.569 | 24.053 |
| 24.249 | -8.078 | -6.608 | 0.005 | -20.433 | -8.883 | 2.659 | 16.483 | -5.928 | 19.260 | 20.053 | 27.275 | 21.674 | 5.832 |
| -19.791 | -22.802 | -9.201 | 5.981 | -17.544 | 23.254 | -25.527 | 6.188 | 16.139 | -10.090 | 25.521 | 28.134 | -14.222 | 12.490 |
| -29.837 | -14.706 | -4.829 | -24.844 | 25.003 | 19.311 | -18.314 | -3.126 | -10.074 | -15.847 | -5.036 | 16.285 | 10.168 | 19.422 |
| -31.371 | 19.126 | 29.940 | 26.556 | 11.898 | -12.733 | -14.829 | -0.397 | 17.518 | 21.589 | -23.257 | -27.308 | 1.617 | 18.979 |
| -4.067 | -3.585 | 5.425 | 8.102 | -6.159 | 17.831 | -3.975 | 18.586 | -12.287 | 20.830 | 23.444 | 22.968 | -23.348 | 23.298 |
| 26.762 | -4.850 | 7.670 | -20.385 | 19.936 | -3.604 | -7.101 | 8.685 | -12.895 | -18.235 | 27.525 | 6.671 | -21.312 | 16.808 |
| 16.693 | 6.850 | 6.155 | -12.470 | -14.059 | -9.749 | 4.672 | 4.760 | -5.279 | 23.175 | -10.479 | 25.126 | 3.378 | 22.358 |
| 4.596 | 21.969 | 26.990 | -23.280 | 20.668 | -12.079 | 20.387 | -16.109 | -14.571 | 25.422 | 6.189 | 13.636 | -12.502 | 20.683 |
| 3.023 | -20.336 | -26.198 | 22.739 | 24.200 | -28.556 | 12.343 | 17.754 | 25.413 | -14.709 | -18.160 | 14.902 | -0.992 | 14.969 |
| 1.561 | 24.783 | 4.035 | -9.338 | 22.727 | 10.820 | -12.513 | -4.963 | 7.627 | 24.438 | 19.880 | 30.028 | -14.853 | 21.255 |
| -0.533 | 17.613 | -7.409 | -30.130 | -2.010 | 9.001 | 14.209 | 23.576 | -24.420 | 30.700 | 19.454 | 13.358 | -17.171 | 13.607 |
| 5.788 | 13.141 | 14.963 | 29.358 | 0.717 | 5.139 | -24.117 | -1.804 | -19.095 | 27.402 | 18.922 | 0.771 | -13.410 | 3.491 |
| -6.011 | 14.218 | -8.170 | 23.627 | 0.267 | -16.805 | 3.374 | 20.634 | -16.293 | 1.869 | 7.275 | 29.631 | 1.137 | -0.649 |
| 10.055 | -25.372 | 3.581 | 20.229 | 19.218 | -25.191 | 13.587 | 22.871 | -23.419 | 25.303 | 22.158 | 27.821 | -22.345 | 17.921 |
| 14.610 | 18.202 | -4.557 | -0.512 | -2.665 | -3.213 | -5.112 | 17.076 | -21.733 | 26.113 | 17.389 | 16.938 | 9.846 | 16.625 |
| 20.912 | 12.715 | 28.865 | 17.524 | 8.381 | 8.467 | 11.400 | -21.339 | 2.232 | 24.007 | -8.329 | 16.411 | -15.921 | 12.184 |
| -19.590 | -11.810 | 0.851 | 5.786 | -24.738 | 7.402 | 11.944 | 15.179 | -26.516 | 18.610 | 24.575 | 15.721 | -21.072 | 10.147 |
| 12.813 | -1.408 | 11.238 | -19.721 | -9.870 | -10.063 | -21.882 | 25.308 | 10.584 | 21.025 | 20.143 | 1.704 | -2.605 | 0.439 |
| 30.736 | -2.248 | -15.282 | 27.572 | -26.304 | 16.325 | 11.017 | 7.056 | -5.419 | 7.556 | 16.261 | 12.249 | -19.032 | 20.006 |

| | | | | | | | | | | | | | |
|---|---|---|---|---|---|---|---|---|---|---|---|---|---|
| -1.365 | 9.327 | -17.598 | 23.171 | 6.504 | -27.223 | 12.658 | -13.343 | 19.036 | 26.756 | 29.507 | 11.163 | 21.636 | 26.169 |
| -29.031 | 21.877 | -2.226 | -27.518 | 10.743 | 11.930 | 9.298 | 24.907 | -25.847 | -26.665 | 8.220 | 18.964 | 14.645 | 21.142 |
| -2.351 | 16.310 | -10.267 | 10.562 | -11.057 | -0.251 | 19.740 | -21.726 | 16.948 | 24.753 | 28.621 | 10.359 | 23.448 | -3.334 |
| -16.131 | 19.016 | -14.201 | -28.215 | -14.889 | -6.116 | 0.038 | 28.460 | 11.665 | -10.326 | -22.750 | 28.923 | 25.643 | 24.565 |
| 24.506 | 8.710 | -24.477 | -6.489 | 13.891 | -12.511 | -11.808 | -23.513 | -18.494 | 20.435 | 23.688 | 17.628 | 13.979 | 26.428 |
| 4.051 | 25.041 | 8.162 | 19.609 | -9.195 | -21.722 | 18.029 | -17.298 | 21.796 | 23.893 | 19.513 | 23.617 | -5.435 | 11.837 |
| 3.072 | 19.950 | -22.179 | -24.764 | -13.474 | 6.212 | 25.807 | 9.814 | 24.194 | 18.113 | 26.711 | 21.792 | 2.439 | 20.023 |
| 19.755 | 6.709 | -1.733 | 13.793 | 0.891 | -20.772 | 6.956 | 26.077 | -26.088 | -19.343 | 16.660 | 30.733 | 15.291 | 24.962 |
| -13.311 | -0.659 | -20.588 | 8.891 | 21.955 | 24.817 | -24.624 | -15.615 | -22.578 | 16.312 | 20.551 | 26.255 | 7.389 | 26.256 |
| 30.176 | -0.267 | 1.709 | -26.650 | 2.096 | -10.630 | -0.778 | 22.546 | -4.260 | 26.003 | 21.576 | 23.956 | -15.303 | 24.195 |
| 6.135 | 9.623 | -2.856 | 9.879 | 4.734 | -13.568 | -20.052 | -9.789 | -24.091 | 19.664 | 24.235 | 24.333 | -6.014 | 23.009 |
| 1.616 | 25.064 | 4.878 | -29.414 | 14.992 | 26.998 | 18.165 | -27.232 | 15.515 | 23.096 | 22.590 | 28.481 | 26.051 | 21.179 |
| -15.514 | -3.400 | 14.346 | 3.899 | 13.719 | 22.542 | -0.966 | 25.309 | 7.488 | 29.391 | 15.166 | 25.578 | 8.913 | 6.882 |
| -17.688 | -13.206 | -1.662 | -24.028 | -6.547 | 26.099 | -16.932 | -23.229 | -2.091 | 24.393 | 22.974 | 13.094 | 22.461 | 23.409 |
| -22.550 | 19.745 | 20.073 | -4.797 | 9.832 | -29.876 | 1.502 | -25.921 | 8.680 | -19.008 | 25.382 | 20.272 | -12.223 | 16.321 |
| -28.540 | 3.712 | 16.696 | -29.119 | -25.599 | 28.410 | -7.553 | 12.801 | 10.095 | 18.770 | 12.311 | 19.568 | 4.731 | 9.761 |
| 17.973 | 22.975 | -8.987 | -14.128 | 3.760 | 23.151 | -18.436 | -24.136 | 12.641 | 21.035 | 28.040 | 24.720 | -15.131 | 26.270 |
| -26.132 | 11.892 | -1.319 | 25.606 | -11.668 | 14.447 | 17.699 | 7.733 | -14.523 | -24.636 | 13.136 | 30.329 | -7.600 | 13.063 |
| -23.675 | 4.167 | -4.368 | 16.840 | -2.422 | 23.874 | -8.436 | -12.637 | 20.282 | 25.362 | 10.122 | 23.902 | -0.404 | 16.573 |
| 23.267 | 13.278 | -2.489 | 6.893 | -1.102 | 3.168 | 17.072 | -8.417 | 11.375 | -28.607 | 30.053 | 25.471 | 2.839 | 13.795 |
| 2.205 | -21.266 | 20.416 | -17.814 | -7.404 | -17.738 | -1.380 | 19.201 | -5.737 | 21.712 | 11.557 | -28.597 | 13.300 | 17.818 |
| -21.601 | -16.147 | 9.926 | 28.772 | -3.869 | 9.924 | 21.705 | 10.155 | 0.497 | -17.833 | -4.774 | 29.150 | 3.912 | 15.679 |
| 4.346 | 19.970 | 20.005 | -10.632 | 0.136 | -10.812 | 2.962 | -23.812 | 17.800 | 21.000 | -24.371 | 30.030 | 10.905 | 23.316 |
| 3.758 | -17.725 | 2.578 | 2.295 | -3.084 | 20.290 | 0.672 | 25.644 | -12.684 | -9.556 | 27.064 | 17.137 | 24.408 | 24.340 |
| -20.715 | 21.605 | 5.236 | -5.900 | 6.350 | -4.571 | -8.018 | 16.813 | -13.507 | 28.012 | 29.283 | 22.668 | 8.228 | 17.184 |
| 1.253 | -24.871 | -6.570 | 26.815 | -4.922 | -9.006 | 12.966 | -15.832 | 10.684 | 10.911 | 19.272 | 27.932 | -8.295 | 18.860 |
| -9.712 | 9.666 | 17.625 | -14.559 | 24.791 | 7.839 | -1.856 | 26.361 | -7.866 | -10.106 | 21.188 | 24.224 | -1.370 | 21.077 |
| 19.168 | -1.810 | -2.519 | -12.318 | 15.646 | 2.260 | -23.369 | 17.961 | 7.487 | -2.885 | -6.519 | 17.825 | 15.982 | 1.463 |
| -5.643 | -20.952 | -0.463 | -13.081 | 8.911 | 23.585 | -25.541 | 5.303 | 14.827 | 23.875 | -19.856 | 12.406 | 27.610 | 13.503 |
| 17.579 | 11.339 | -14.497 | 20.754 | 13.741 | 26.083 | 22.738 | -6.884 | -5.297 | 14.425 | -0.216 | 15.699 | 17.882 | 29.064 |
| -19.932 | 11.934 | -19.345 | -27.952 | 19.283 | 17.879 | 10.883 | -22.045 | -7.842 | -21.650 | 6.728 | 14.993 | 20.325 | 22.211 |
| 16.357 | -7.549 | -25.404 | 13.592 | 8.152 | 1.403 | -17.063 | -22.829 | -7.481 | 19.717 | 17.330 | 20.650 | 1.800 | 25.407 |
| 8.192 | 0.576 | 1.848 | 25.002 | 7.823 | -24.913 | 21.204 | -25.883 | -13.511 | 21.901 | 7.239 | 29.686 | 25.631 | 5.233 |
| -3.489 | -10.152 | -12.713 | 9.426 | 10.841 | -23.319 | -6.914 | 22.379 | -12.344 | -2.222 | 28.499 | 14.326 | -22.315 | 9.887 |
| 22.640 | 18.605 | -1.569 | 24.384 | -9.276 | -29.639 | 3.587 | -12.136 | 5.173 | 20.343 | 4.385 | 30.631 | 23.584 | 14.862 |
| -29.224 | -19.783 | -3.253 | -7.438 | 14.327 | 24.737 | 14.566 | 23.307 | 22.050 | 8.062 | 22.073 | 27.013 | -21.403 | 17.218 |

| | | | | | | | | | | | | | |
|---|---|---|---|---|---|---|---|---|---|---|---|---|---|
| 27.713 | 13.599 | -19.479 | 28.389 | 15.782 | -16.351 | -4.226 | 24.780 | -10.790 | 24.639 | 30.118 | 16.471 | 22.782 | 2.949 |
| 19.956 | 1.113 | 21.616 | -23.674 | 22.612 | 15.132 | -8.480 | 4.546 | 9.939 | -2.982 | 17.631 | 24.952 | 1.152 | 7.410 |
| 10.101 | 9.350 | -4.334 | -3.875 | 8.974 | -21.593 | 2.369 | 21.214 | -2.420 | 16.664 | -0.161 | 28.863 | -3.827 | 20.822 |
| -13.702 | 17.827 | -6.493 | 21.544 | 10.162 | 5.479 | -11.988 | -5.308 | 12.850 | -28.530 | 17.116 | 19.652 | 26.851 | 15.070 |
| -22.372 | 23.912 | -25.751 | -9.727 | 19.410 | -7.802 | 21.368 | 15.800 | 23.999 | 23.900 | 18.523 | 23.888 | 25.143 | 19.723 |
| -28.379 | 5.949 | 9.214 | 8.090 | 18.880 | -24.439 | -2.743 | 6.821 | -1.306 | 7.594 | 9.957 | 27.810 | 5.446 | -19.103 |
| -25.600 | 26.448 | -1.295 | -4.550 | -8.952 | -24.952 | -3.104 | 10.875 | 7.999 | 17.607 | 30.640 | 19.250 | 9.276 | 14.581 |
| -1.311 | 21.140 | 24.607 | 18.955 | 17.194 | 20.133 | 15.682 | -25.533 | -14.032 | 28.892 | 12.510 | 13.369 | 21.669 | 23.312 |
| -24.285 | 3.147 | 1.399 | 22.159 | 22.168 | -2.053 | 9.365 | 19.779 | -2.751 | -12.667 | 19.052 | 0.288 | 22.311 | 0.846 |
| 8.942 | 19.277 | -18.306 | 15.902 | 25.645 | -27.535 | 8.842 | 21.964 | 21.414 | -10.076 | 25.111 | 26.888 | -0.688 | 25.129 |
| 4.825 | 9.831 | -10.097 | 5.990 | 16.530 | -3.704 | 2.948 | 25.447 | 15.251 | 23.125 | 17.922 | 25.666 | 26.417 | 29.577 |
| 2.238 | 8.558 | 11.521 | -12.379 | 20.835 | 8.992 | -5.944 | 26.640 | 19.405 | 7.938 | 20.432 | 30.043 | 4.684 | 28.567 |
| -10.746 | -2.935 | 2.100 | -11.295 | 25.078 | 15.890 | 13.185 | -2.991 | 5.181 | 1.570 | 27.840 | 26.320 | 15.576 | 15.574 |
| 17.293 | -20.789 | -10.994 | 24.434 | -21.683 | 6.784 | -24.808 | 6.206 | 22.689 | 18.106 | 23.052 | 19.974 | 22.826 | 24.633 |
| 26.786 | 25.387 | 3.340 | -24.815 | 7.003 | -2.801 | 13.597 | 4.242 | 23.622 | 23.373 | 13.364 | 24.395 | 9.628 | 21.811 |
| -24.479 | 21.418 | -5.330 | -14.830 | -2.325 | 3.985 | 5.312 | -13.264 | 9.428 | 15.932 | 24.578 | 24.827 | 19.043 | 17.280 |
| 8.488 | 26.519 | -17.804 | 27.270 | 22.247 | -6.470 | -0.709 | -18.711 | 16.334 | 12.478 | 19.903 | 16.658 | 17.287 | 16.111 |
| 19.784 | 25.901 | -7.501 | 9.656 | -5.197 | -11.776 | -13.100 | 15.262 | -23.009 | -14.055 | 12.346 | 12.277 | -8.152 | 25.461 |
| 0.150 | -12.189 | 18.612 | 8.102 | 25.968 | -23.330 | -4.832 | 22.839 | -25.177 | 19.546 | 28.978 | 18.722 | 23.727 | 26.554 |
| 10.003 | 14.588 | 2.670 | -5.212 | 11.194 | -28.932 | -4.043 | 12.161 | 20.333 | -22.700 | 22.026 | -9.304 | 2.893 | 17.246 |
| 16.070 | 19.810 | -19.578 | 10.483 | -3.290 | -0.539 | -12.676 | -12.681 | 25.963 | -24.606 | 26.291 | 4.989 | 27.164 | 21.426 |
| -8.852 | 2.037 | 1.250 | -19.651 | -8.338 | -20.934 | 5.941 | 20.463 | 24.973 | 28.163 | 7.912 | 21.800 | -2.973 | 8.271 |
| 17.890 | 4.292 | -6.945 | 11.690 | 16.235 | 24.286 | 17.524 | 19.502 | 1.060 | 2.805 | -24.278 | 13.218 | 11.239 | 30.883 |
| 21.934 | 7.748 | 18.082 | -10.620 | 15.376 | 14.797 | 11.969 | 9.150 | -21.515 | -0.677 | 22.646 | 15.314 | 24.861 | 11.746 |
| 20.294 | -8.309 | -13.532 | -28.613 | 6.841 | -26.507 | 2.491 | 10.061 | -1.571 | 22.728 | 14.216 | 28.353 | -10.633 | 18.777 |
| -7.597 | 3.981 | -4.343 | 15.763 | 2.410 | -12.954 | -6.536 | -14.306 | 15.019 | 20.198 | 18.571 | 10.532 | 18.118 | 13.632 |
| -28.316 | -17.875 | -22.367 | 24.548 | -5.250 | -6.069 | -19.853 | -23.412 | -0.072 | 19.885 | 9.608 | 6.224 | 20.112 | 23.774 |
| 16.953 | 19.220 | -25.025 | 21.799 | -24.595 | -19.952 | 3.543 | -24.975 | -15.601 | 7.153 | 24.481 | 4.744 | -2.030 | 20.326 |
| -6.156 | 20.276 | 17.306 | 4.883 | 13.860 | -17.451 | -7.461 | -4.522 | -2.223 | -1.663 | 15.661 | 21.637 | 2.228 | 8.835 |
| -10.281 | 20.850 | -15.515 | -1.780 | 2.575 | 22.000 | -17.986 | 16.217 | -7.459 | 16.457 | 19.791 | 23.037 | 26.983 | 17.796 |
| 8.028 | 25.590 | 15.429 | 16.850 | 1.618 | 11.357 | 11.278 | 26.234 | 22.741 | 18.902 | 18.809 | 26.706 | 18.927 | 18.472 |
| 6.373 | 4.368 | -0.089 | 4.042 | 11.875 | 21.204 | 5.253 | -18.432 | -3.933 | 17.086 | 27.288 | 28.697 | 3.019 | 11.804 |
| -21.207 | -25.770 | 16.182 | 12.457 | -15.566 | 9.479 | 13.068 | 2.698 | 17.526 | -0.671 | 20.543 | 13.823 | -19.829 | 21.682 |
| -16.589 | 12.806 | -18.966 | 5.736 | 27.352 | -26.099 | 6.773 | 13.723 | -27.062 | 18.065 | 14.396 | 20.226 | -26.523 | 22.937 |
| -6.978 | 23.245 | 14.521 | 7.533 | 21.930 | 16.779 | -5.422 | 7.220 | 13.188 | 5.023 | 17.736 | 19.186 | 23.896 | 20.244 |
| -20.644 | 19.650 | 1.992 | -12.682 | -7.763 | 2.600 | 17.016 | -15.278 | -11.512 | 3.937 | 21.280 | 29.812 | -22.148 | 18.922 |

| | | | | | | | | | | | | | |
|---|---|---|---|---|---|---|---|---|---|---|---|---|---|
| -28.963 | 10.957 | 11.198 | -1.879 | 8.356 | -2.210 | -8.624 | -7.889 | 12.365 | 10.660 | 28.457 | -8.394 | -21.604 | 25.745 |
| 28.989 | 15.451 | 1.548 | -27.540 | 19.773 | -19.835 | -7.877 | -23.206 | 21.252 | 10.384 | 16.287 | 27.937 | 21.835 | 8.617 |
| 7.087 | 18.620 | 10.595 | 7.052 | -4.472 | 10.473 | 20.966 | 5.146 | 19.162 | 12.654 | 25.456 | 25.668 | 21.319 | 18.311 |
| -14.758 | 11.832 | 22.251 | -10.542 | -1.431 | 5.786 | 0.633 | 19.273 | -12.272 | -2.164 | 31.022 | 21.135 | -13.103 | 15.953 |
| 19.441 | 26.808 | 19.404 | -5.910 | -1.071 | -9.325 | -17.426 | -0.926 | -13.263 | 12.073 | 15.535 | 17.538 | 28.465 | 30.469 |
| -19.036 | -23.814 | -8.195 | -23.080 | -6.329 | 12.960 | 7.307 | 3.185 | -20.737 | 20.392 | 21.708 | -9.036 | 2.814 | 21.588 |
| 22.611 | 28.028 | -10.025 | -1.478 | -27.636 | -4.855 | -0.593 | 0.825 | -24.414 | 22.285 | 12.025 | 26.093 | -5.545 | 22.672 |
| -25.699 | 19.324 | 7.849 | 26.007 | 24.810 | -22.546 | -3.443 | 6.930 | 8.034 | 3.861 | 29.685 | 22.456 | 26.968 | 14.247 |
| 26.251 | 12.388 | -14.592 | 26.393 | -7.264 | 24.768 | 3.574 | 4.032 | 7.348 | -4.019 | 24.033 | 24.883 | 5.458 | 23.933 |
| -3.562 | 15.866 | 6.471 | -19.828 | 25.656 | -10.575 | -3.118 | -0.171 | 15.883 | 1.579 | 23.099 | -28.202 | 15.515 | -1.752 |
| -8.199 | -7.289 | 6.072 | 18.772 | 15.038 | -25.469 | -13.999 | -5.468 | 19.972 | -5.985 | -10.545 | 27.103 | 24.765 | 23.710 |
| 1.066 | 15.274 | -3.405 | 21.516 | 26.539 | 20.490 | 17.895 | 8.120 | 14.440 | 2.803 | 20.242 | -28.703 | 1.722 | 0.790 |
| -26.708 | 21.242 | -20.454 | -14.233 | -7.563 | -8.774 | 11.883 | -3.972 | -4.531 | -26.546 | 28.879 | 30.363 | -12.032 | 18.468 |
| 4.087 | -23.114 | 9.847 | -27.604 | 26.887 | -27.739 | 0.627 | 16.344 | -21.841 | 0.547 | 11.361 | 22.628 | 27.899 | 21.472 |
| -5.698 | 17.531 | 4.421 | -19.513 | 18.208 | 5.180 | -21.312 | 8.544 | -8.533 | -23.982 | -18.364 | 20.147 | 22.418 | 19.106 |
| 4.817 | 24.139 | 2.619 | 15.978 | -2.081 | 6.393 | 12.822 | 0.412 | -15.634 | -22.417 | 19.674 | 16.674 | 11.795 | 26.219 |
| -4.102 | 20.464 | 0.961 | 5.107 | -5.792 | -6.508 | 19.194 | 3.698 | 9.725 | -18.706 | 18.411 | 29.507 | -17.828 | 29.734 |
| 8.950 | 22.886 | -1.929 | 9.202 | -14.120 | -18.745 | 15.428 | 20.499 | 5.324 | 21.788 | -9.255 | 24.116 | 23.898 | 27.965 |
| 18.185 | 21.997 | -4.185 | -17.671 | 8.711 | 7.724 | -10.508 | 7.026 | 8.752 | -12.964 | 9.621 | 19.346 | -16.194 | 15.895 |
| -2.484 | -21.104 | -6.079 | -22.637 | 19.691 | -1.797 | -10.023 | -3.172 | 23.768 | -0.715 | 16.890 | 23.396 | 9.959 | 17.088 |
| -11.422 | 23.506 | 13.762 | 19.014 | -14.627 | 4.549 | -4.425 | 3.570 | -2.297 | 0.149 | 18.770 | 25.609 | 26.850 | 1.369 |
| -18.157 | 25.887 | 4.905 | -26.845 | 17.066 | -24.520 | 25.696 | 14.247 | 6.970 | 9.324 | 25.046 | 27.632 | 11.234 | 13.881 |
| 28.489 | 17.596 | -4.530 | 22.910 | 24.597 | -21.060 | -1.704 | 13.185 | 11.807 | -23.838 | 29.719 | 28.151 | -12.209 | 20.894 |
| 27.126 | 16.600 | 7.993 | -20.306 | -26.110 | 26.261 | -4.825 | 16.323 | -1.886 | 12.001 | 21.555 | 18.581 | 23.810 | 17.944 |
| -0.830 | 16.111 | -7.809 | 3.934 | 16.284 | -10.646 | -14.984 | 7.821 | -19.974 | 24.899 | 5.977 | 30.055 | 28.877 | 24.401 |
| 23.802 | 2.996 | -15.401 | 16.495 | 11.552 | -14.295 | 24.764 | -3.926 | -17.672 | -14.466 | 22.441 | 13.275 | 20.660 | 4.815 |
| 6.749 | 26.866 | -25.539 | -25.566 | 18.106 | -11.463 | -21.844 | 6.395 | 10.535 | -3.245 | 20.293 | 26.480 | 14.277 | 21.612 |
| 27.815 | -22.279 | 18.446 | -19.861 | 14.321 | 17.048 | 1.197 | -7.206 | 26.070 | -19.019 | 24.625 | 11.104 | -19.671 | 19.318 |
| 25.434 | -10.913 | -17.535 | 11.649 | -6.389 | 21.441 | 7.357 | -12.451 | 13.395 | -20.907 | 27.246 | 13.667 | 11.903 | 20.237 |
| 18.621 | 13.094 | -1.805 | -11.415 | -15.055 | 16.542 | -10.824 | 12.731 | 3.397 | 8.195 | 17.558 | -14.014 | 25.974 | 25.474 |
| 29.007 | 27.035 | -1.410 | 6.491 | 25.186 | 14.174 | 6.879 | -8.503 | -24.314 | -17.683 | 15.290 | 18.885 | -4.219 | 27.133 |
| -28.421 | 22.287 | 5.255 | 3.805 | 9.459 | 16.031 | -17.674 | 17.216 | -23.706 | 2.969 | 11.677 | 25.782 | 19.124 | 27.735 |
| 14.627 | 8.924 | -21.911 | -5.800 | 26.365 | 5.747 | -23.464 | -22.987 | -26.029 | -20.410 | 19.505 | -29.082 | 24.387 | 17.883 |
| 24.033 | 3.499 | 7.867 | 16.862 | 30.122 | 8.471 | -15.579 | 21.485 | 15.189 | -18.361 | 28.552 | 29.546 | -25.030 | 23.888 |
| 22.128 | 28.171 | -7.194 | -24.092 | -17.727 | 28.226 | -7.017 | -5.220 | -2.833 | 23.360 | 8.216 | 19.959 | -19.089 | 19.592 |
| 9.700 | 12.999 | 5.761 | 4.609 | -26.995 | -5.912 | 2.106 | 1.437 | 21.721 | 0.362 | 20.481 | 22.868 | -14.801 | 23.138 |

| | | | | | | | | | | | | | |
|---|---|---|---|---|---|---|---|---|---|---|---|---|---|
| 6.419 | 7.869 | -3.987 | 3.500 | 25.188 | -27.006 | -20.816 | 13.855 | 8.026 | 9.767 | 9.089 | 16.912 | 22.360 | 20.776 |
| 3.286 | 27.790 | -26.801 | 19.951 | 13.143 | -3.056 | 27.828 | 4.699 | -26.724 | 6.307 | 6.767 | -26.368 | 11.235 | -2.627 |
| 20.983 | -15.365 | 11.442 | 8.313 | -4.441 | 17.400 | -5.229 | 12.548 | 6.304 | 22.157 | 25.340 | 26.799 | -21.751 | -2.176 |
| -11.277 | 22.777 | -1.475 | -25.855 | -8.741 | -14.446 | 8.775 | 8.969 | 0.846 | 16.816 | 29.442 | -0.344 | 26.554 | 27.397 |
| -27.068 | 16.742 | -23.531 | 22.905 | 20.642 | 7.583 | 0.305 | -21.643 | -5.232 | -7.763 | 17.947 | 24.888 | 28.356 | 11.786 |
| -27.978 | -18.549 | 1.459 | -21.179 | -10.814 | -15.366 | 11.738 | 18.648 | -17.555 | 0.911 | 23.110 | 20.082 | 3.676 | 25.883 |
| 22.846 | 0.003 | 4.652 | 16.477 | -19.409 | -4.299 | 7.121 | -7.811 | 20.585 | 11.931 | 13.651 | 28.284 | 14.841 | 17.372 |
| -0.526 | -24.134 | 8.085 | -8.023 | 21.595 | -17.018 | -4.457 | 15.541 | -9.737 | -11.142 | -24.566 | 13.510 | 10.475 | 26.533 |
| -27.611 | -10.554 | 10.864 | -10.893 | -25.232 | -26.258 | 3.209 | 23.910 | 18.340 | -9.087 | 25.652 | 14.375 | -7.194 | 29.025 |
| 24.508 | 28.359 | 18.390 | -15.052 | -18.700 | -15.606 | 22.511 | -19.935 | 4.572 | 12.070 | 18.280 | 16.285 | 5.210 | 15.976 |
| -4.384 | 9.255 | -7.603 | 5.133 | -30.147 | 28.334 | -17.927 | 18.193 | 23.044 | -9.439 | 8.619 | 3.952 | 20.041 | 22.908 |
| -20.725 | 29.005 | -9.155 | 21.897 | 28.309 | 6.359 | 13.669 | 11.557 | 8.280 | -2.846 | 21.136 | -18.321 | 11.844 | 13.847 |
| -26.276 | 25.834 | -2.492 | -14.058 | 8.411 | -3.937 | -6.240 | 17.163 | 20.388 | 3.055 | 22.379 | 26.585 | 26.283 | 30.225 |
| -18.965 | -23.201 | 3.038 | 19.002 | 11.201 | 7.034 | 16.242 | 25.787 | -10.481 | 10.020 | -4.711 | 22.744 | 18.501 | 27.056 |
| 0.779 | 17.374 | 5.348 | -20.429 | -10.859 | 3.088 | 22.716 | 2.962 | 14.509 | -0.724 | -21.096 | 21.010 | 12.940 | 25.583 |
| 25.825 | 13.048 | -21.678 | 11.810 | 17.164 | -10.408 | -9.852 | -3.292 | 2.698 | 0.614 | 24.870 | -13.212 | 28.980 | 21.557 |
| -24.335 | 20.689 | -4.287 | -17.119 | 25.185 | -26.845 | -13.555 | 18.124 | 27.146 | 23.328 | 26.912 | 16.920 | 20.490 | -1.054 |
| -23.107 | 15.062 | -16.355 | 7.479 | -7.920 | 20.905 | 19.782 | 21.950 | 21.216 | -10.474 | 28.969 | 24.349 | 24.519 | 25.245 |
| -9.480 | 26.695 | 13.401 | -13.902 | 19.874 | 18.216 | 16.505 | -12.810 | 15.538 | 20.650 | 26.015 | 28.862 | 27.792 | 17.161 |
| -1.171 | -17.690 | -9.853 | -5.180 | 30.019 | -14.514 | 19.246 | 1.330 | 3.819 | -19.840 | 20.473 | 23.544 | 22.578 | 19.632 |
| -15.802 | -19.856 | 25.243 | -21.880 | 5.004 | 12.138 | -7.338 | 6.559 | 19.135 | -1.453 | 14.504 | 20.106 | -15.554 | 13.759 |
| -19.573 | -24.029 | 16.647 | -20.686 | -2.865 | -6.442 | 21.282 | 15.170 | -19.789 | -20.768 | 18.403 | 7.673 | 23.377 | 18.039 |
| 21.625 | -27.836 | -11.613 | -16.588 | 8.373 | 13.935 | -3.163 | 1.933 | -20.648 | 9.888 | 15.999 | -9.246 | 19.821 | 22.710 |
| -3.130 | 21.269 | 3.759 | -20.065 | 30.998 | -12.372 | -5.166 | 10.416 | 23.051 | -13.645 | 26.658 | -9.661 | -26.842 | 23.744 |
| -6.674 | -25.144 | 5.715 | 3.485 | 28.500 | -15.419 | 23.946 | 20.741 | -22.248 | -17.125 | 18.633 | 8.917 | 31.030 | 21.878 |
| 4.505 | 7.483 | -1.662 | 17.031 | 11.285 | 8.824 | -11.919 | -8.695 | -26.573 | -12.522 | 26.962 | 20.680 | 30.698 | 13.634 |
| -25.296 | 28.836 | 6.852 | 18.674 | 22.580 | -7.659 | 11.549 | 3.552 | 22.008 | -11.168 | 22.408 | 23.614 | 25.324 | 4.708 |
| 20.377 | -11.372 | -16.655 | 8.638 | 18.670 | 16.270 | -5.450 | 0.578 | 2.704 | 17.687 | 19.530 | -11.628 | 24.270 | 28.646 |
| -10.348 | 25.420 | 13.897 | 5.548 | 25.220 | 20.245 | -0.470 | 25.389 | 20.403 | 23.864 | 10.725 | 27.306 | -13.730 | 23.187 |
| 27.208 | -20.131 | -3.129 | 2.028 | -13.908 | 26.051 | 28.656 | -21.953 | 9.608 | 7.457 | 5.445 | 17.217 | -14.414 | 16.444 |
| 28.209 | -22.672 | -10.732 | 20.795 | -23.820 | -24.946 | 10.132 | 1.176 | -14.664 | 0.023 | 26.234 | 20.135 | 21.562 | -6.075 |
| -2.459 | 16.538 | 5.389 | 19.650 | 27.780 | -1.621 | -4.983 | 4.453 | 19.723 | -24.992 | 6.704 | 15.824 | 18.072 | 15.760 |
| 10.696 | 11.093 | 0.149 | -6.166 | 26.283 | -9.630 | -6.078 | 24.119 | 13.481 | 20.012 | 22.790 | 28.307 | 19.091 | 27.721 |
| -11.922 | 23.702 | -9.468 | -12.555 | 12.436 | -4.895 | -6.790 | 27.821 | -19.126 | -18.340 | -26.592 | 4.508 | 1.109 | 21.078 |
| -26.361 | 22.452 | 3.856 | 14.633 | -8.583 | -15.333 | 4.278 | 10.207 | -0.042 | -26.907 | 25.585 | 26.881 | 30.512 | 16.998 |
| 14.247 | 12.175 | -7.306 | 10.782 | 22.432 | -22.925 | -12.886 | -3.998 | 6.501 | 15.680 | 22.038 | 25.950 | -1.277 | 13.191 |

| | | | | | | | | | | | | | |
|---|---|---|---|---|---|---|---|---|---|---|---|---|---|
| 26.360 | 19.722 | 22.961 | -5.334 | 28.213 | -2.606 | 16.457 | 1.539 | 25.558 | -24.849 | 12.363 | 23.368 | 6.072 | 24.338 |
| 18.148 | -9.657 | 4.463 | -22.685 | 28.548 | 9.925 | 3.416 | -10.455 | -6.460 | 7.479 | 9.232 | 5.058 | 20.412 | 15.502 |
| 29.052 | -21.637 | -24.001 | 17.885 | -10.145 | -0.917 | -21.725 | 23.507 | 22.723 | -24.881 | 24.063 | 24.987 | 28.984 | 21.307 |
| 9.109 | 11.683 | 14.579 | 11.456 | 29.755 | -26.738 | -2.012 | 16.322 | -20.235 | 17.877 | 14.948 | 22.213 | 9.506 | 15.186 |
| 9.860 | 19.453 | 6.076 | -24.432 | 8.405 | 1.334 | -11.498 | 20.496 | 0.897 | 2.199 | 27.842 | 20.380 | 22.413 | 8.051 |
| -14.034 | 21.256 | -10.950 | 23.525 | 27.565 | 17.521 | 20.478 | -22.000 | -18.615 | -19.768 | 10.651 | 18.588 | 15.477 | 16.469 |
| 6.365 | 18.923 | 13.060 | 3.224 | 24.190 | -1.459 | 4.949 | 26.549 | 26.198 | 21.555 | 22.366 | -4.907 | 23.074 | 18.760 |
| 14.504 | -25.004 | -3.841 | 26.058 | 3.862 | -4.343 | -1.048 | 4.916 | -10.058 | 8.596 | 19.060 | 5.097 | 14.155 | 25.600 |
| 5.502 | 29.217 | -18.965 | -2.080 | 26.141 | 6.961 | -7.251 | 16.185 | 12.522 | -12.783 | 17.055 | 3.458 | -6.818 | 24.186 |
| -6.930 | 26.738 | -7.410 | 1.131 | 5.598 | -17.222 | -0.230 | 13.390 | 18.975 | 21.089 | 21.704 | -2.461 | 10.505 | 23.284 |
| 7.069 | -0.919 | 1.660 | 15.734 | 19.577 | -24.186 | 15.249 | -9.309 | -0.265 | 19.801 | 26.642 | 21.660 | 2.031 | 25.059 |
| 4.752 | 25.838 | -2.639 | -24.989 | 11.161 | -16.405 | 10.860 | 21.014 | -22.025 | 16.060 | -5.323 | -25.750 | 26.444 | 16.425 |
| -25.543 | 17.207 | 0.944 | 8.509 | 28.197 | -18.963 | 6.576 | -24.162 | -11.805 | -12.428 | 23.877 | 24.916 | 0.464 | 17.246 |
| -26.488 | 29.874 | -4.273 | 15.222 | 19.252 | -2.208 | -22.401 | -15.364 | 24.375 | 12.129 | 12.534 | 5.921 | 27.494 | 21.660 |
| -13.831 | -28.441 | -7.082 | 13.327 | 22.413 | 12.674 | 5.180 | -19.964 | 24.916 | 22.186 | 25.588 | 26.328 | 19.914 | 27.501 |
| 2.575 | 23.111 | -9.173 | -24.647 | 24.306 | 25.160 | -27.337 | 17.104 | 28.192 | 23.796 | 26.149 | 22.106 | 4.093 | 13.875 |
| 0.502 | -24.217 | -10.236 | 25.671 | -17.934 | 20.607 | -16.271 | -23.453 | 18.373 | 22.988 | 11.794 | -5.009 | -5.250 | 14.559 |
| -4.620 | -17.765 | -4.457 | 22.351 | 27.000 | 4.171 | 25.222 | 9.188 | 11.959 | -18.452 | 23.183 | -0.910 | 29.001 | 15.996 |
| 27.835 | -5.878 | 23.358 | -2.919 | -8.485 | -12.027 | 6.063 | 27.430 | -11.340 | -1.019 | 25.002 | 25.498 | 25.300 | 18.607 |
| -10.082 | 25.421 | 7.754 | 5.739 | 27.941 | -12.578 | -4.072 | 21.998 | 11.226 | -12.616 | 19.696 | -12.216 | 7.322 | 23.626 |
| 25.577 | 10.126 | -9.819 | -21.730 | -6.629 | 23.180 | 19.888 | -18.145 | -28.356 | -19.633 | 14.557 | 21.732 | -8.014 | 16.911 |
| 17.541 | 20.221 | -16.440 | 16.473 | 8.104 | 2.560 | 8.718 | 0.078 | 20.507 | -10.377 | 21.309 | 11.253 | 27.205 | 18.514 |
| 8.736 | 18.376 | 1.455 | 12.497 | 18.430 | -5.608 | -11.416 | -25.265 | 10.225 | 21.651 | 11.798 | 10.421 | 0.331 | 12.237 |
| 21.699 | -18.852 | 17.775 | 24.022 | 20.489 | -26.236 | -10.934 | 9.853 | 22.033 | 24.488 | 16.670 | -8.092 | 24.048 | 18.978 |
| -7.982 | -2.904 | 16.292 | 26.768 | -28.014 | 17.976 | -19.415 | 12.697 | -0.508 | 5.601 | 9.017 | -27.362 | 23.096 | 17.919 |
| -16.227 | -19.510 | 25.161 | -7.016 | 8.236 | -16.137 | -26.378 | 1.243 | 22.994 | -21.350 | -0.919 | 5.474 | 10.839 | -22.164 |
| -17.231 | 22.794 | -0.588 | 5.516 | 14.346 | -3.157 | 9.943 | -1.238 | 7.034 | -12.583 | 15.737 | -24.144 | 8.586 | 20.740 |
| 14.616 | 31.381 | 11.522 | -11.010 | -22.088 | 5.241 | -20.668 | -9.334 | 27.572 | 10.069 | 20.334 | 3.135 | -10.072 | 22.366 |
| -5.221 | 10.506 | -7.018 | 9.785 | -29.302 | 11.305 | 21.841 | 3.399 | 19.882 | 19.981 | 9.519 | 23.510 | 19.633 | 16.502 |
| -13.370 | 7.419 | 23.585 | -6.623 | 25.767 | 16.137 | -16.164 | 17.777 | -6.267 | 17.756 | -1.978 | 22.995 | 18.837 | 14.685 |
| 23.743 | 27.440 | -8.428 | -16.860 | 6.823 | 7.479 | 30.956 | 28.169 | -4.044 | 16.560 | 18.585 | 27.637 | -11.864 | 21.155 |
| -8.820 | 15.762 | 20.521 | 20.198 | -5.599 | -4.676 | -30.120 | 26.489 | -14.587 | 12.133 | 9.105 | 26.817 | 28.769 | 25.726 |
| 5.642 | 23.576 | 15.957 | 19.394 | -14.431 | 2.743 | 16.364 | -27.546 | 19.246 | -23.531 | -9.578 | 26.383 | 26.465 | 19.918 |
| -19.686 | -28.740 | 14.102 | 2.504 | 6.034 | -5.310 | -0.676 | 19.320 | 14.224 | -27.093 | 9.397 | -18.556 | 27.315 | 24.345 |
| -27.386 | -21.835 | 1.458 | -26.442 | -16.273 | -11.568 | -10.411 | 11.615 | -28.421 | -3.106 | 19.743 | 23.598 | 25.178 | 9.666 |
| 22.785 | 17.318 | -9.608 | -22.874 | 14.593 | -5.834 | -29.229 | 14.187 | 15.443 | 24.038 | -1.724 | 20.641 | 24.378 | 24.906 |

| | | | | | | | | | | | | | |
|---|---|---|---|---|---|---|---|---|---|---|---|---|---|
| 17.441 | 30.795 | -19.181 | -7.245 | -13.054 | 11.819 | 25.686 | 7.674 | -15.482 | 0.454 | -19.372 | 0.913 | 28.084 | 23.439 |
| -1.085 | 23.930 | -7.378 | 10.569 | -19.643 | -6.432 | 5.169 | -16.068 | 29.050 | 24.756 | -7.810 | 15.771 | 15.803 | 15.822 |
| -4.297 | -19.246 | -10.261 | 22.721 | -7.130 | -2.556 | -23.061 | 23.772 | 16.089 | 22.148 | -3.233 | 8.235 | -17.751 | 19.209 |
| -5.916 | 31.732 | -11.140 | 26.084 | 13.653 | -15.114 | 12.164 | 15.970 | 1.175 | -8.311 | -24.289 | 17.994 | 21.659 | 20.295 |
| -23.820 | 7.997 | 7.674 | 7.455 | -29.098 | -22.742 | 21.124 | 27.043 | -19.921 | 26.050 | -9.172 | 8.757 | 22.379 | 27.041 |
| 11.806 | 26.055 | 0.771 | -21.542 | 20.506 | -25.253 | 6.600 | -20.750 | 10.406 | 22.735 | -9.630 | 1.387 | 14.783 | 21.707 |
| -1.939 | 18.964 | -6.691 | 27.859 | 24.444 | 4.497 | 28.930 | 3.893 | -2.696 | -25.089 | 19.989 | 12.507 | 25.950 | 16.804 |
| 2.374 | 16.640 | 4.292 | -24.487 | 14.114 | 20.502 | 11.310 | 13.714 | 20.105 | -19.472 | 23.950 | -6.198 | 21.331 | 22.964 |
| -15.919 | 27.845 | 3.240 | 18.406 | 18.926 | 2.365 | -8.606 | 21.120 | -29.481 | 16.702 | -8.241 | 7.923 | 18.148 | 18.526 |
| 18.441 | 27.077 | -2.539 | 8.470 | 19.758 | -5.875 | 5.811 | 16.415 | -0.432 | -3.218 | 12.855 | -1.804 | -3.447 | 24.405 |
| -25.810 | -18.170 | -11.928 | 12.549 | 10.395 | 22.347 | -24.946 | 4.309 | 9.261 | -7.199 | 20.611 | 24.976 | 2.618 | 21.944 |
| -6.239 | -20.015 | -25.530 | -9.154 | 7.743 | 9.658 | 28.047 | 20.165 | -5.052 | -13.012 | 9.656 | 16.986 | -25.004 | 26.214 |
| 16.880 | 22.493 | 1.312 | 21.260 | 25.737 | -9.898 | 12.911 | -9.154 | -28.029 | 14.786 | -19.844 | 9.768 | 16.149 | 14.508 |
| -18.842 | 9.532 | -9.885 | -12.190 | 7.095 | 6.450 | 21.071 | -0.914 | 24.975 | -28.366 | -6.002 | -9.936 | 9.755 | 26.669 |
| 27.743 | 15.022 | -3.716 | -16.483 | -15.617 | 0.425 | -19.128 | -21.592 | 10.683 | 15.927 | 24.189 | -6.627 | 25.338 | 22.379 |
| -26.608 | 29.775 | -5.752 | 2.529 | 11.432 | -17.551 | 14.600 | -21.027 | 28.370 | -18.381 | 20.599 | 9.311 | 28.605 | 17.735 |
| -5.322 | -10.726 | -18.938 | 1.909 | -18.331 | -4.918 | -17.772 | 16.662 | -22.440 | -22.376 | 19.419 | -2.758 | 15.479 | 16.420 |
| 19.291 | 17.696 | -10.683 | -19.994 | 24.071 | -7.103 | 11.931 | 22.150 | 22.725 | -27.640 | 19.050 | 10.990 | 12.283 | 21.352 |
| 10.741 | 10.116 | 19.422 | -26.005 | -14.656 | -0.685 | -22.246 | 11.496 | -15.468 | 20.346 | -1.178 | 5.039 | -5.148 | 19.602 |
| 27.326 | 29.886 | -1.527 | 7.120 | -0.292 | -6.256 | -7.626 | 19.702 | 27.223 | 13.350 | -6.750 | -8.063 | 23.465 | 15.771 |
| -6.512 | 9.223 | -3.271 | -21.256 | -2.164 | -9.228 | -15.327 | -28.759 | -1.022 | -27.033 | 21.616 | 23.861 | 26.054 | 23.301 |
| -10.013 | 3.977 | -18.142 | -3.760 | 19.215 | -19.227 | 8.582 | 25.078 | 23.661 | 19.543 | 24.371 | -1.308 | 16.575 | 25.166 |
| 24.272 | 21.322 | 3.095 | 6.505 | 10.219 | 8.714 | -12.118 | 12.314 | -3.654 | -17.625 | 8.068 | 12.800 | 8.365 | -11.805 |
| -14.126 | 18.666 | 4.535 | -22.774 | 15.380 | -16.345 | -21.790 | 0.755 | -21.166 | -19.359 | 18.303 | -19.770 | 13.996 | 26.370 |
| 26.394 | 28.224 | 11.727 | -12.769 | -14.181 | 23.295 | 21.836 | -26.358 | 27.946 | 22.526 | 13.460 | 1.821 | 28.048 | 19.193 |
| 22.039 | 14.446 | -14.960 | 21.766 | 3.532 | 22.412 | 29.011 | 3.345 | 21.847 | 12.009 | -25.711 | 0.537 | 12.020 | 17.959 |
| -24.623 | 24.226 | 0.425 | 20.713 | 21.381 | 3.085 | -12.883 | 28.324 | 14.598 | 24.995 | 23.051 | 17.226 | 18.499 | 24.782 |
| 25.260 | 26.360 | -19.601 | -23.479 | 18.374 | -3.058 | 18.716 | 17.488 | 11.197 | 22.004 | 18.411 | -8.866 | -9.700 | 22.855 |
| 25.994 | -6.018 | 27.140 | 24.948 | 16.235 | 5.057 | -26.378 | -23.386 | 4.731 | 14.966 | -2.628 | -3.450 | 17.657 | 18.427 |
| 20.581 | -3.478 | 9.478 | -6.295 | 21.206 | -14.974 | 20.275 | -25.132 | 24.488 | 1.077 | 22.195 | 25.645 | 8.664 | 17.071 |
| 14.147 | 25.754 | 2.259 | 24.542 | 11.295 | -22.121 | 15.741 | 26.476 | 2.650 | -13.044 | 26.439 | 2.828 | 20.329 | 12.055 |
| -1.414 | 11.337 | -2.370 | 14.713 | -17.262 | -10.438 | 25.043 | 19.589 | -27.686 | -24.370 | 14.626 | -24.454 | -22.176 | 21.626 |
| -3.654 | -28.800 | -8.521 | -24.694 | -18.563 | 1.324 | -4.210 | 10.653 | -28.640 | 17.414 | -1.087 | -27.300 | 21.957 | 15.893 |
| -25.270 | 12.066 | 8.092 | -0.127 | 17.590 | 0.773 | -20.017 | -21.316 | 27.326 | 16.132 | 19.363 | 1.358 | -4.891 | 20.398 |
| 16.775 | 23.570 | 6.923 | -9.651 | 10.015 | 0.127 | -29.384 | 15.368 | 9.289 | 23.598 | 25.102 | 20.588 | 25.762 | 23.115 |
| 23.243 | 16.233 | -4.662 | 18.192 | 16.614 | 24.106 | 9.253 | -14.516 | -20.840 | -22.038 | -10.097 | 6.001 | -0.063 | 10.314 |

| | | | | | | | | | | | | | |
|---|---|---|---|---|---|---|---|---|---|---|---|---|---|
| 25.349 | 24.784 | -5.945 | 19.111 | 23.824 | -3.697 | -22.325 | -5.469 | 12.112 | -27.917 | 22.438 | 26.512 | 24.414 | 21.039 |
| -9.377 | -23.037 | 16.902 | 17.101 | -0.306 | -5.577 | 22.908 | 17.553 | 4.126 | 18.753 | 18.315 | 24.822 | 20.727 | 5.606 |
| 12.906 | -7.846 | 25.230 | 7.986 | -21.186 | 7.661 | -7.417 | 2.222 | -22.437 | 12.886 | 1.931 | 9.758 | 19.479 | 23.984 |
| 2.027 | 30.185 | 9.043 | -8.992 | 12.139 | 12.274 | 4.207 | 27.454 | -12.387 | 26.690 | 11.988 | 6.671 | 21.652 | 12.102 |
| -4.092 | 18.476 | 23.101 | 20.831 | -5.083 | 15.917 | 26.709 | 16.922 | 1.739 | 18.203 | -1.854 | 13.761 | -26.882 | 6.991 |
| 5.786 | 16.807 | 12.997 | 11.690 | 8.756 | 21.878 | 23.419 | -6.287 | -23.349 | 25.548 | -14.930 | -25.119 | 10.283 | 12.812 |
| 8.642 | 7.097 | -10.304 | -20.442 | -20.925 | -11.282 | -7.561 | 14.954 | 5.806 | -20.199 | 20.075 | -3.012 | 22.925 | 9.010 |
| -18.956 | 20.079 | -1.427 | 28.325 | 24.008 | 2.520 | -8.945 | -28.157 | -18.507 | 21.188 | 4.402 | 9.195 | 17.587 | -2.358 |
| -20.255 | 6.800 | -8.052 | 18.335 | 16.673 | 15.128 | -9.605 | 24.333 | 9.907 | 19.467 | 24.500 | -15.210 | -19.246 | 15.087 |
| 26.420 | 3.195 | -9.784 | 29.565 | 3.991 | 9.070 | 8.627 | -21.028 | -25.830 | 23.054 | 16.293 | 4.369 | 10.085 | -6.743 |
| -16.215 | -1.390 | 19.027 | -6.521 | -20.983 | -4.730 | -11.864 | -2.157 | 18.912 | 20.987 | 25.737 | 23.439 | -11.075 | 19.221 |
| 18.686 | 25.100 | -25.814 | 1.890 | 17.896 | 5.163 | -13.811 | 22.222 | -15.279 | -25.496 | 1.466 | -0.076 | -16.081 | 1.481 |
| 22.717 | 25.759 | 3.344 | 16.747 | 12.652 | 4.528 | 24.318 | -22.856 | 24.659 | 20.098 | 11.214 | 12.853 | -27.243 | 20.575 |
| 19.925 | 25.315 | 0.996 | 7.500 | 7.654 | 20.889 | -22.551 | 28.283 | -0.316 | -16.417 | 6.806 | 21.996 | 6.492 | 19.375 |
| 4.941 | 12.243 | 2.001 | -13.933 | -0.519 | 14.517 | -1.284 | 6.874 | 28.852 | -20.401 | 21.892 | 15.047 | 4.171 | 19.865 |
| 15.784 | -4.921 | -6.718 | -28.372 | -23.107 | 21.464 | 23.772 | -7.335 | 8.206 | 25.740 | 27.800 | 9.972 | 24.734 | 22.113 |
| 17.661 | -15.613 | 14.413 | 14.865 | -10.219 | 2.925 | -2.619 | 19.345 | -6.032 | 22.133 | 13.186 | -6.404 | -22.671 | 17.698 |
| 16.768 | -3.302 | -0.141 | -0.912 | -4.591 | -16.068 | 16.203 | 3.987 | -8.084 | 17.105 | 12.390 | -2.097 | -18.085 | 21.447 |
| 7.102 | 11.693 | -3.854 | -4.604 | -24.146 | 15.687 | 27.707 | 23.813 | 23.791 | -24.343 | -13.607 | 22.289 | 14.954 | 25.665 |
| -24.569 | 8.820 | 6.565 | 8.329 | -12.062 | -5.837 | -4.483 | -0.896 | 15.907 | 20.117 | 24.244 | 1.174 | 19.472 | 22.490 |
| 9.542 | -28.797 | 13.746 | 12.150 | 0.251 | -1.835 | 17.433 | 26.408 | 22.770 | -26.749 | -25.658 | 18.111 | 27.051 | 15.185 |
| -15.782 | -6.812 | 10.466 | -20.344 | 9.040 | 11.628 | 14.382 | 20.787 | -3.809 | 26.201 | 20.855 | 25.777 | 13.467 | -16.268 |
| -6.355 | -5.824 | -4.845 | 19.564 | 9.953 | -22.862 | -6.929 | -15.239 | -7.282 | 24.000 | -26.310 | -3.002 | 20.684 | -25.238 |
| 10.416 | 4.722 | -6.385 | -25.940 | -3.611 | -8.429 | 3.812 | 9.883 | 29.961 | 29.195 | 13.483 | -3.674 | 2.377 | 24.647 |
| -4.360 | 23.669 | 5.331 | 10.896 | -8.609 | -5.261 | -11.774 | -11.428 | 28.833 | 22.712 | -0.652 | 2.444 | 15.011 | 20.545 |
| -25.694 | -8.245 | 26.354 | 10.186 | -5.386 | -6.222 | -22.093 | 30.175 | 26.645 | -0.518 | 27.111 | -8.084 | 21.418 | 18.145 |
| 8.110 | 16.429 | -3.147 | 17.959 | 18.916 | 18.711 | -13.857 | -28.059 | 20.172 | 19.566 | 1.559 | -10.309 | -12.260 | 6.152 |
| -23.166 | 21.465 | 1.014 | -0.614 | 9.424 | 22.499 | -16.035 | 14.733 | 18.399 | -24.809 | 17.334 | 19.716 | 23.612 | 12.496 |
| -25.910 | 1.703 | -1.645 | -18.700 | 21.873 | -15.429 | 14.955 | 18.745 | 17.707 | 23.470 | 4.008 | 3.575 | 17.189 | 17.050 |
| 1.607 | 18.069 | -18.658 | -21.125 | -9.716 | -19.039 | 14.026 | -24.702 | -24.020 | -20.004 | -26.709 | 20.910 | 22.378 | 21.407 |
| 0.881 | 8.908 | 9.137 | -10.573 | -1.480 | 5.190 | -3.806 | -29.718 | 25.544 | 21.600 | 23.540 | 25.053 | 9.073 | 20.299 |
| 9.221 | 27.256 | 2.030 | -13.976 | 17.171 | 2.488 | -22.901 | 21.499 | 27.802 | 24.294 | 18.558 | -2.072 | 18.042 | 19.131 |
| 20.629 | 20.341 | -26.188 | -27.811 | -8.846 | -7.858 | -21.136 | 18.990 | 13.371 | -26.825 | 19.874 | -22.864 | 16.846 | 22.817 |
| 13.987 | -3.601 | 26.805 | 28.885 | 13.354 | 15.875 | 12.395 | 1.185 | 6.664 | 24.312 | -3.398 | 20.446 | 24.232 | 13.499 |
| 10.047 | 16.990 | -7.030 | 19.103 | -5.374 | -17.719 | 19.567 | -6.135 | -18.945 | -20.572 | -17.933 | 4.485 | 26.635 | 19.718 |
| 22.155 | 13.944 | 3.282 | 4.864 | -4.727 | -5.363 | 7.853 | -8.340 | -27.590 | -22.295 | 12.554 | -20.001 | 25.230 | 17.224 |

| | | | | | | | | | | | | | |
|---|---|---|---|---|---|---|---|---|---|---|---|---|---|
| 6.782 | -14.807 | 22.275 | -29.533 | -0.104 | 6.356 | 0.224 | 20.217 | 20.603 | 19.751 | 23.351 | 3.045 | 22.604 | 15.286 |
| -12.202 | -27.831 | -9.865 | 13.292 | 2.231 | -4.259 | 20.417 | 25.422 | 8.488 | -21.564 | 13.243 | 1.380 | 15.731 | 16.539 |
| -23.813 | -2.523 | 10.803 | -5.134 | 8.960 | 17.698 | 4.404 | 22.646 | -14.999 | -24.615 | 5.007 | 1.990 | -12.345 | 19.657 |
| 20.993 | 3.485 | 28.786 | -28.599 | -15.383 | -25.479 | -5.350 | 15.926 | -0.803 | -25.155 | -12.760 | 23.667 | 18.749 | 26.336 |
| -19.742 | 20.214 | -0.331 | -17.520 | 11.535 | 14.217 | -6.212 | -13.722 | 25.921 | -17.297 | -15.831 | 5.273 | 24.570 | 11.666 |
| 18.173 | 7.595 | 14.785 | -4.337 | 5.981 | -21.129 | 23.158 | -21.427 | 10.780 | -27.172 | 22.163 | -4.951 | 16.319 | 27.417 |
| 14.917 | 20.416 | -1.030 | 24.748 | 3.092 | 14.521 | 3.891 | -24.590 | 16.758 | -28.253 | 28.195 | 24.607 | 11.232 | 25.132 |
| 23.704 | 7.948 | -13.505 | 11.744 | 19.467 | -15.498 | -25.983 | 1.657 | -21.620 | 28.859 | 19.104 | 16.130 | -18.262 | 19.728 |
| 17.343 | -23.584 | -1.465 | 2.479 | 22.422 | -14.494 | -2.363 | -10.451 | -6.439 | 17.212 | 20.089 | 10.360 | 21.773 | -3.392 |
| 5.093 | 1.420 | 3.449 | -11.532 | 5.551 | -4.729 | -21.881 | 30.835 | 8.552 | -7.855 | 14.523 | 0.185 | 21.423 | 8.171 |
| 16.087 | 14.352 | 17.744 | 16.539 | 10.574 | -8.754 | 21.061 | -7.180 | 4.214 | -15.049 | -1.694 | -10.797 | 17.886 | 21.397 |
| 10.580 | 11.165 | 24.968 | -4.019 | -16.759 | 18.777 | 10.761 | -4.869 | -0.348 | -23.544 | 15.776 | -3.467 | 13.639 | 18.295 |
| 15.787 | 2.970 | -18.311 | 28.246 | -13.126 | 10.005 | 13.760 | 17.965 | -5.016 | 23.293 | 13.957 | 1.202 | -23.105 | 9.046 |
| 8.273 | -4.568 | -14.509 | -18.782 | 1.291 | -7.089 | -20.471 | -4.117 | 25.117 | -27.608 | 20.847 | -9.800 | 16.715 | 15.718 |
| -7.776 | 8.632 | -6.592 | 22.664 | -6.504 | 19.228 | 24.499 | 23.882 | 24.227 | 28.738 | 21.633 | -9.364 | -0.278 | 26.363 |
| 16.608 | 26.823 | -22.892 | 23.935 | 21.105 | -6.756 | -25.972 | -6.595 | 19.543 | 21.817 | -23.882 | -5.306 | 9.320 | 13.872 |
| -13.104 | -4.422 | -9.050 | 19.840 | 18.497 | -23.833 | 11.498 | 6.622 | 17.403 | -22.286 | -3.531 | -3.650 | 13.672 | 22.745 |
| 12.610 | -5.391 | 11.695 | -27.281 | -18.782 | -6.123 | -19.853 | 13.433 | 9.100 | -2.943 | 12.167 | -10.610 | -0.882 | 21.835 |
| 20.052 | -0.067 | -20.377 | 25.369 | -10.321 | 21.811 | -6.565 | 7.560 | 7.872 | 2.260 | 28.589 | -25.076 | 15.708 | 17.520 |
| 0.679 | 4.310 | 0.770 | -27.870 | 12.007 | -10.661 | -27.671 | 20.151 | -23.873 | 25.525 | -11.859 | 0.794 | -22.200 | 23.997 |
| -23.400 | -2.429 | 7.724 | 11.858 | -8.998 | 7.570 | -1.043 | -10.887 | 28.122 | 13.811 | 9.597 | -2.320 | -24.410 | 20.132 |
| -2.568 | 6.028 | 18.016 | 27.584 | -7.196 | -19.005 | -3.520 | -0.675 | -5.316 | 29.811 | 26.437 | -23.773 | -8.637 | 19.234 |
| 2.134 | 24.746 | 1.284 | -25.523 | 13.508 | 23.127 | -7.653 | 22.615 | 2.780 | 27.365 | 21.948 | 22.212 | 22.306 | 25.722 |
| 13.465 | -6.903 | 17.236 | 26.268 | 20.525 | 20.212 | 4.814 | 10.042 | -11.408 | 26.851 | 21.076 | 4.027 | 7.722 | 27.042 |
| 8.479 | 13.535 | -9.066 | 27.905 | 22.305 | -17.107 | 8.756 | 15.041 | 18.128 | 16.610 | 6.607 | 16.702 | -8.003 | 27.461 |
| 24.361 | -20.079 | -10.484 | 19.603 | -7.812 | -0.107 | -10.814 | -13.281 | -1.829 | 18.073 | 17.664 | 8.274 | -11.220 | 17.490 |
| -21.428 | 20.695 | 17.552 | -2.565 | 19.836 | -3.850 | 6.185 | 26.653 | -23.131 | -18.155 | 21.451 | 24.870 | 4.177 | 23.218 |
| 20.145 | -9.336 | -0.406 | 17.966 | 25.743 | -5.134 | -2.720 | 17.758 | 28.679 | -9.039 | 13.805 | 24.056 | -14.851 | 20.446 |
| 19.215 | 0.931 | -26.891 | 5.751 | -16.578 | -11.277 | 20.789 | 15.756 | -24.235 | -30.241 | -29.401 | 11.794 | -7.423 | 27.897 |
| -14.019 | -4.001 | 26.720 | -24.963 | 7.991 | -5.554 | -3.835 | 26.846 | -16.037 | -22.844 | 10.756 | 22.730 | 1.257 | 26.020 |
| -18.925 | 20.628 | -3.809 | 8.953 | -21.876 | 15.337 | -12.609 | 23.657 | -27.535 | 15.336 | -28.775 | 17.308 | -1.804 | 24.957 |
| -5.242 | -14.662 | 6.662 | -28.777 | 26.556 | -19.754 | 9.548 | 16.355 | 12.094 | 20.850 | 12.954 | -0.862 | 11.270 | 9.543 |
| -14.670 | -4.500 | -5.112 | -27.100 | 6.652 | -22.203 | 6.969 | 17.526 | -0.221 | 22.594 | 12.048 | -2.722 | -21.644 | 10.727 |
| -3.109 | 20.227 | 17.123 | -29.824 | -20.880 | 25.518 | -8.246 | -7.815 | 0.826 | 18.570 | 18.322 | 13.760 | -1.481 | 16.270 |
| 6.696 | 10.111 | 4.286 | 24.217 | -0.148 | -10.338 | 18.829 | -25.843 | -14.787 | -22.295 | -1.281 | 8.909 | 19.669 | 14.596 |
| 5.143 | 4.754 | -15.397 | 15.845 | -1.646 | -8.852 | -19.315 | 24.073 | -12.742 | 19.511 | 22.502 | 1.454 | -9.144 | 21.522 |

| | | | | | | | | | | | | | |
|---|---|---|---|---|---|---|---|---|---|---|---|---|---|
| 14.421 | 3.090 | -7.454 | 12.125 | -14.411 | -8.401 | 22.601 | 28.814 | -28.864 | 23.794 | 6.447 | -12.275 | -2.032 | 15.657 |
| 7.336 | 0.285 | -12.548 | 21.625 | 20.641 | 7.814 | 15.547 | 6.063 | 16.177 | 30.261 | 27.646 | -9.641 | 1.495 | 21.890 |
| -1.018 | 11.357 | -23.244 | -16.422 | 24.793 | 22.139 | -16.963 | 7.863 | -16.492 | -22.793 | -1.129 | -0.333 | -11.086 | 19.357 |
| -5.070 | 24.070 | -19.507 | -11.485 | 25.247 | 21.383 | -5.108 | 25.360 | -25.329 | 13.666 | 25.385 | 10.391 | 22.959 | 27.025 |
| 15.488 | 11.752 | -4.744 | -3.487 | -23.422 | 5.915 | -5.643 | 4.956 | 29.062 | 23.639 | 28.515 | 11.964 | -14.643 | 21.127 |
| 11.903 | -16.730 | -17.672 | -15.183 | -18.496 | -24.337 | 2.414 | -2.747 | 15.564 | 22.510 | 14.238 | 25.066 | -22.782 | 14.546 |
| 7.557 | 25.224 | -14.217 | 19.162 | 18.225 | 15.014 | -4.900 | -31.166 | 15.029 | 24.774 | -30.254 | 23.227 | -12.127 | 13.480 |
| 16.484 | 0.512 | 29.254 | 18.744 | -10.964 | 3.244 | -13.387 | 21.055 | -26.718 | -19.933 | 20.031 | 10.830 | 17.513 | 26.242 |
| 11.362 | 10.799 | -13.031 | 16.792 | -21.032 | -1.610 | 17.691 | 26.944 | 21.807 | 20.047 | 21.780 | -7.063 | 13.907 | 25.372 |
| -17.530 | 6.173 | 18.707 | 9.919 | -22.550 | 1.930 | -4.030 | -18.438 | 14.707 | 21.032 | -23.599 | 22.871 | 5.726 | 18.809 |
| -12.045 | 12.098 | 12.056 | 11.627 | -13.533 | -12.140 | -25.791 | -30.516 | 13.916 | -21.099 | 16.503 | -8.841 | -7.004 | 22.476 |
| -23.223 | 4.045 | -2.380 | 23.607 | 12.848 | 0.274 | 18.723 | -16.026 | 26.579 | 21.994 | 22.808 | 24.175 | 16.686 | -27.097 |
| 11.361 | 21.405 | -11.611 | 25.746 | -7.448 | 0.978 | -27.393 | -20.301 | 12.745 | -15.428 | 11.202 | 16.886 | 21.755 | 16.326 |
| 10.713 | -8.982 | -0.686 | -22.573 | 12.113 | -13.057 | 7.931 | 1.536 | -27.901 | 25.257 | 21.084 | 18.632 | 24.319 | 28.772 |
| 9.898 | -16.976 | -5.457 | -15.368 | -4.129 | 2.398 | -5.695 | 20.971 | 20.874 | 23.193 | -26.944 | 9.691 | -17.548 | 15.249 |
| 9.364 | 19.211 | 13.344 | 24.105 | 2.431 | -0.386 | 24.688 | 30.268 | -17.908 | -26.085 | 24.941 | -0.903 | 25.228 | 28.036 |
| -5.742 | 10.988 | -3.483 | 0.803 | -26.549 | -7.705 | 27.633 | -29.529 | -31.952 | 26.279 | 11.895 | 9.325 | 18.538 | 23.446 |
| 7.934 | 22.491 | 18.439 | -27.999 | -19.282 | -6.295 | 9.924 | -28.883 | -3.587 | 13.634 | 6.849 | -5.469 | 20.977 | 16.383 |
| -3.607 | 0.342 | -2.606 | -19.722 | 15.794 | -4.013 | -18.772 | -4.420 | 6.407 | 22.355 | 22.901 | -18.823 | -3.663 | 24.224 |
| 21.241 | 25.589 | 1.360 | -21.279 | 21.942 | -15.055 | 8.570 | -23.889 | 3.760 | 14.864 | 13.806 | 23.631 | 22.463 | 28.987 |
| 20.962 | 2.332 | -6.627 | -16.690 | 12.326 | 21.629 | -26.050 | 16.241 | -17.185 | -23.008 | 30.355 | 11.359 | 20.182 | 26.274 |
| 15.384 | -4.016 | 11.729 | -26.825 | -23.286 | -2.923 | 7.906 | -26.554 | 24.743 | 17.448 | 22.251 | 4.195 | 16.764 | 24.320 |
| -4.288 | 24.401 | 9.248 | -18.930 | 25.502 | -20.050 | 1.659 | -8.235 | 11.523 | -29.554 | 18.245 | -5.931 | 25.532 | 22.689 |
| -11.957 | 20.049 | 0.687 | -28.780 | 18.302 | 12.675 | 0.479 | -0.253 | 15.030 | 18.661 | 12.789 | 25.391 | 24.765 | -17.517 |
| -18.931 | 8.384 | -4.679 | -29.567 | -25.349 | -14.688 | 18.586 | -19.230 | 1.933 | 21.633 | 14.468 | 2.945 | 19.495 | 14.600 |
| 8.886 | 22.936 | -5.698 | 18.711 | -18.494 | -3.171 | 25.529 | 17.183 | -14.281 | 24.337 | 27.053 | -9.425 | 2.793 | 12.181 |
| -14.496 | 9.525 | 8.950 | 30.472 | 20.880 | 20.665 | 20.617 | 26.335 | 20.374 | 20.889 | 13.051 | 24.606 | -1.093 | 29.365 |
| -8.031 | 2.759 | -27.276 | 4.180 | -2.619 | -4.705 | 2.838 | -0.647 | 5.448 | -25.195 | 26.560 | -10.089 | 17.421 | 28.813 |
| -20.311 | -24.475 | 21.261 | -20.535 | 15.194 | 22.355 | -12.001 | 24.580 | 23.897 | 27.305 | 13.859 | 20.174 | 14.241 | 12.806 |
| -26.197 | -9.261 | -1.221 | 20.462 | 7.567 | 10.241 | -17.786 | 21.306 | -20.071 | -24.553 | 27.608 | 7.866 | 9.364 | 27.717 |
| 20.495 | 23.952 | -19.899 | 16.981 | -13.318 | 13.966 | -0.763 | 6.845 | 30.232 | -9.133 | -15.825 | 0.025 | 1.353 | 21.300 |
| 19.708 | 5.353 | 3.032 | 30.165 | 26.879 | -9.023 | 0.412 | 0.804 | -4.720 | 30.729 | -28.796 | -0.673 | -23.474 | 25.490 |
| 25.818 | 22.961 | -12.126 | -3.245 | 4.498 | 12.708 | 16.093 | 22.503 | 0.796 | -16.024 | -29.582 | -26.603 | 10.285 | 19.332 |
| 3.262 | -15.244 | 12.025 | -13.576 | -17.727 | 25.121 | -23.494 | -23.220 | -16.305 | -4.103 | -5.507 | 22.758 | -5.515 | 17.516 |
| 24.225 | -3.524 | 18.309 | 7.795 | 16.383 | -8.702 | 26.085 | -29.093 | 26.321 | 19.339 | -9.114 | -17.855 | -6.812 | 11.998 |
| -18.118 | 24.103 | -7.475 | -27.552 | -0.416 | -15.544 | 5.896 | -7.256 | 27.412 | -23.364 | 12.323 | 26.866 | 15.723 | 23.807 |

| | | | | | | | | | | | | | |
|---|---|---|---|---|---|---|---|---|---|---|---|---|---|
| 10.938 | 23.078 | 3.415 | 13.027 | 9.166 | 24.603 | -20.869 | 18.654 | 28.629 | -22.242 | 6.713 | 25.894 | 20.300 | -11.063 |
| 6.275 | -21.177 | -8.543 | 9.898 | 10.425 | 10.793 | 14.152 | 11.511 | 21.289 | 23.080 | 7.594 | -4.614 | 23.248 | 30.085 |
| -13.191 | 0.883 | 22.020 | 9.062 | -9.102 | -2.303 | 20.473 | 4.800 | -20.917 | 17.223 | 27.513 | 1.412 | 17.599 | 20.597 |
| -23.603 | 4.951 | 1.238 | -18.009 | 18.127 | 21.809 | 5.139 | -22.125 | 24.539 | -22.004 | 20.869 | 19.156 | 25.643 | 13.280 |
| 7.556 | -24.517 | -1.840 | 12.761 | 14.530 | -20.427 | 5.405 | -0.398 | 29.531 | -24.490 | -0.774 | -12.864 | 9.950 | 14.778 |
| 2.239 | 21.616 | -6.270 | -30.141 | 12.028 | 0.297 | 10.550 | -13.974 | 18.778 | 17.826 | -30.034 | 4.264 | 6.447 | 10.497 |
| 21.073 | -25.045 | -15.926 | -28.137 | -27.087 | -17.930 | 1.895 | 1.190 | -14.703 | -16.013 | 13.932 | 19.531 | 21.054 | 28.007 |
| 25.409 | 16.890 | -2.414 | -15.973 | 1.132 | 19.960 | -21.836 | 22.931 | 17.329 | 18.059 | 16.341 | 18.144 | 22.706 | 20.190 |
| -21.828 | -26.229 | -21.830 | 17.973 | -4.854 | 7.529 | -16.693 | 14.388 | -30.840 | -4.331 | 19.944 | 5.411 | 22.300 | -11.039 |
| 23.757 | -1.584 | -13.313 | 19.242 | 24.909 | -6.856 | 5.428 | -24.045 | 30.326 | -23.223 | 14.597 | -1.278 | 14.868 | 28.375 |
| -26.255 | 17.942 | 5.615 | -10.950 | -20.143 | -7.689 | 19.651 | -29.551 | 3.253 | -15.802 | 31.334 | -15.208 | -8.970 | 15.554 |
| 18.458 | -15.306 | 2.167 | -8.999 | 23.088 | 15.751 | 23.713 | 21.578 | -10.461 | -23.705 | 11.839 | 25.276 | -9.618 | 29.159 |
| 6.528 | 18.866 | -6.753 | -23.127 | -1.610 | 2.275 | 5.823 | -19.243 | 22.148 | 24.273 | 17.993 | 11.295 | 26.469 | 26.726 |
| 25.686 | 25.236 | 18.053 | 0.308 | 22.077 | -25.515 | -12.916 | 0.387 | 9.243 | -25.562 | 30.798 | 4.707 | -6.774 | 14.048 |
| -5.567 | 24.329 | -4.048 | 27.143 | 21.884 | -1.417 | -14.600 | -0.286 | -1.067 | 10.905 | 17.293 | 21.121 | -18.946 | 25.947 |
| 22.214 | 26.988 | 18.542 | 17.066 | -14.340 | 11.184 | -4.135 | 14.041 | -3.361 | 7.152 | 25.436 | 19.148 | 22.153 | 27.463 |
| 14.758 | 21.967 | 2.689 | 21.069 | -18.828 | 6.149 | 1.351 | -20.727 | -18.343 | 18.102 | -8.153 | 13.101 | -3.558 | 16.555 |
| 8.600 | 5.105 | -20.101 | -11.995 | -21.057 | 1.296 | -10.789 | 15.767 | 1.891 | 23.769 | -28.998 | -0.243 | 28.151 | 18.540 |
| 5.729 | -20.260 | -21.046 | 20.781 | -0.893 | -2.192 | 22.277 | 15.286 | 3.493 | -9.477 | 21.478 | -18.260 | -12.630 | 20.002 |
| -3.027 | 22.490 | -10.016 | -2.132 | 6.476 | -5.645 | 20.716 | -23.119 | -0.351 | 27.027 | 11.291 | -10.850 | 10.724 | 16.088 |
| 13.000 | -24.978 | -0.105 | -26.007 | 25.708 | -20.994 | 19.869 | -3.498 | -9.474 | 9.005 | -2.769 | 22.583 | 27.349 | 17.039 |
| -10.458 | -20.785 | 7.685 | 15.198 | 18.763 | -27.189 | 11.987 | 23.297 | 15.780 | -26.447 | 26.784 | 23.116 | 13.993 | 25.031 |
| 20.542 | -23.260 | 27.489 | 5.750 | -22.592 | -3.579 | 20.775 | -8.395 | 2.773 | 13.903 | -30.594 | 5.366 | 18.110 | 23.395 |
| 21.510 | 6.967 | 21.313 | -28.527 | 14.377 | -11.279 | 8.688 | -25.483 | -13.120 | -20.243 | -27.867 | 9.836 | -3.736 | 28.229 |
| -24.690 | -5.105 | 11.746 | 14.405 | -0.559 | 11.348 | 18.119 | 24.956 | 23.222 | -27.790 | 4.173 | -0.720 | 21.904 | 23.974 |
| -7.494 | -20.924 | -7.320 | -4.246 | -12.191 | 0.818 | 15.199 | 8.005 | 12.795 | -21.887 | 29.276 | -11.971 | 20.255 | 17.270 |
| 15.457 | 4.524 | 1.257 | -18.057 | -15.545 | 24.254 | 3.817 | -0.598 | 11.129 | -16.196 | 25.565 | -17.120 | 25.281 | 15.706 |
| 0.907 | 11.228 | -11.921 | 29.688 | 27.039 | -24.578 | -0.714 | -3.766 | 6.394 | -14.216 | 30.273 | 7.597 | -25.782 | 11.963 |
| 14.075 | 19.376 | 19.940 | -19.448 | -24.495 | -26.497 | 16.742 | -22.584 | 26.596 | -4.374 | -18.541 | -21.438 | -5.525 | 0.421 |
| -22.502 | 20.092 | 18.619 | -17.452 | -13.040 | 22.968 | 9.856 | 29.060 | 22.794 | -12.125 | 17.232 | 21.925 | 21.090 | 21.475 |
| 18.576 | -13.060 | -9.185 | 10.077 | 22.570 | 1.419 | -3.248 | 11.646 | -5.804 | -19.856 | 31.375 | 14.007 | 4.259 | 25.982 |
| 23.502 | 18.798 | 4.473 | 22.984 | 0.938 | -7.089 | 23.333 | 19.487 | -8.764 | -24.381 | 27.714 | 27.168 | 14.190 | 12.908 |
| -17.624 | -24.630 | 13.416 | 11.615 | 19.290 | -20.347 | 4.688 | -26.790 | 13.887 | -1.346 | 13.707 | 19.099 | -11.818 | 16.058 |
| -3.546 | -25.721 | -2.298 | 1.976 | -23.814 | 26.493 | 10.734 | 27.336 | 27.725 | 26.045 | 23.358 | -9.369 | 19.173 | 18.997 |
| -27.634 | -1.434 | -18.680 | -3.361 | -23.407 | -25.079 | 12.922 | 5.251 | -2.238 | -19.095 | -25.159 | -2.896 | 16.912 | 21.067 |
| -26.110 | -17.444 | -9.693 | 10.639 | 11.613 | -8.092 | 6.027 | -16.602 | 28.280 | -14.945 | -26.354 | 17.130 | 21.348 | 28.766 |

| | | | | | | | | | | | | | |
|---|---|---|---|---|---|---|---|---|---|---|---|---|---|
| 9.419 | 21.856 | 21.793 | -16.955 | 7.033 | -14.915 | -7.756 | 26.159 | 20.431 | 6.171 | 16.375 | -16.362 | 29.304 | 23.504 |
| -19.016 | 15.386 | 1.838 | 12.688 | 25.006 | 7.738 | 6.550 | -4.796 | 24.865 | -12.936 | -14.610 | -7.906 | 15.543 | 22.930 |
| 19.759 | 14.866 | 3.200 | 3.785 | 22.051 | 7.297 | 21.737 | -20.166 | 8.472 | -6.157 | -23.876 | 24.361 | 13.100 | 18.998 |
| -26.429 | 1.129 | -8.246 | -13.013 | -24.497 | 7.445 | 11.793 | -24.862 | -16.572 | -21.133 | -20.479 | 18.066 | 20.316 | 16.516 |
| 22.385 | -12.237 | -9.160 | -11.551 | -26.973 | -7.628 | 7.672 | -25.651 | 21.580 | 24.583 | 21.785 | 0.161 | 9.248 | 26.756 |
| -15.653 | -27.713 | -20.182 | -29.385 | 27.545 | 12.936 | 3.320 | 16.683 | 28.832 | 18.022 | 18.001 | 14.798 | 23.435 | 31.412 |
| 6.109 | -4.080 | -6.357 | 22.657 | 19.145 | -21.486 | -5.586 | 0.843 | -18.232 | -22.743 | 19.709 | -2.052 | 22.149 | 31.641 |
| -21.645 | 21.442 | -4.093 | 13.630 | -10.039 | -9.994 | -12.079 | 18.059 | 15.813 | 1.389 | 29.045 | 23.437 | -19.941 | 31.042 |
| 16.893 | -20.217 | 19.318 | 24.971 | 25.530 | -11.278 | -25.038 | 6.605 | -14.886 | -27.460 | -17.209 | 20.640 | 0.930 | 27.747 |
| -27.652 | -3.122 | 13.043 | -20.826 | 16.840 | -2.381 | -14.399 | 8.782 | -18.823 | -16.307 | -30.624 | 23.083 | 28.568 | 12.205 |
| 13.949 | -24.289 | -8.156 | 1.648 | -14.157 | -7.776 | -10.493 | -19.206 | 16.542 | 22.013 | 16.806 | 21.935 | 24.705 | 30.183 |
| -1.854 | 17.455 | -11.050 | -4.192 | -19.005 | -25.328 | -17.605 | -26.039 | 1.354 | -14.620 | -25.134 | 3.067 | 27.270 | 24.872 |
| 10.243 | -25.670 | 7.921 | 21.461 | -14.519 | -8.140 | -1.596 | -25.397 | 29.624 | -10.880 | -11.233 | 26.121 | 7.136 | 14.500 |
| -5.519 | 18.994 | 8.868 | -21.767 | 0.963 | 26.030 | -8.736 | -20.581 | -28.468 | -9.145 | 16.307 | -6.906 | -8.706 | 28.831 |
| -18.913 | 6.365 | 20.945 | 16.673 | -17.141 | -11.788 | 19.068 | -26.567 | 20.368 | -23.363 | 3.791 | 25.711 | 9.006 | 24.164 |
| 20.544 | 27.429 | 10.673 | -28.508 | 15.084 | -27.060 | 24.439 | 12.090 | 0.766 | 14.146 | 14.508 | 7.957 | 16.444 | 20.745 |
| -11.995 | 1.138 | 19.549 | -12.551 | -22.477 | -24.988 | -0.669 | -0.621 | -10.895 | -9.333 | -2.191 | -25.941 | 17.778 | 15.553 |
| -10.820 | -25.041 | 9.734 | -25.455 | -27.634 | 0.077 | -17.543 | 12.606 | -23.695 | -11.764 | 20.820 | 13.614 | 29.760 | 21.817 |
| 27.763 | 28.046 | -20.038 | 20.179 | 29.925 | -13.209 | -18.702 | 22.928 | 18.763 | 25.293 | 14.109 | 21.464 | 1.350 | 16.881 |
| -26.080 | -8.711 | -27.655 | 27.154 | 20.521 | 3.977 | 7.612 | 7.888 | -30.208 | 11.702 | 29.553 | 20.404 | 12.365 | 29.495 |
| -16.637 | -28.276 | 23.717 | -19.301 | -25.645 | -15.528 | 0.294 | -16.302 | 10.090 | 0.673 | -29.196 | 27.353 | 27.348 | 15.155 |
| 25.290 | 4.608 | -8.266 | 1.000 | 28.477 | 29.676 | -25.668 | 16.941 | -10.067 | -20.087 | 22.942 | 18.846 | 29.180 | 11.825 |
| -21.605 | 16.385 | -8.872 | 31.236 | 7.172 | 26.371 | -4.837 | 24.996 | 21.946 | 18.650 | 27.618 | 1.614 | 19.539 | 23.407 |
| 8.903 | 18.083 | -11.451 | 19.282 | -2.883 | -14.472 | -11.980 | 10.783 | -29.555 | 23.322 | 20.263 | 15.655 | 16.073 | 17.988 |
| 21.856 | -25.604 | -27.391 | -18.381 | 22.753 | -16.878 | 19.310 | -22.535 | -11.947 | 27.309 | 24.353 | 27.029 | 5.823 | 29.998 |
| -7.135 | 20.252 | -2.028 | 9.101 | 23.970 | 1.204 | -23.938 | 19.094 | -7.691 | 11.796 | 17.340 | 23.847 | 3.131 | 7.665 |
| -27.698 | -16.598 | 10.053 | 17.295 | 8.380 | -21.471 | -20.779 | 13.576 | -21.250 | 7.595 | 21.683 | -16.016 | 4.199 | 7.896 |
| 25.162 | 28.612 | -5.999 | -22.419 | -11.918 | -26.355 | 23.231 | -26.818 | -28.968 | -26.207 | 0.080 | 25.030 | 11.840 | 23.514 |
| -3.164 | 25.489 | -20.052 | -7.914 | -16.625 | -7.464 | -9.735 | -0.241 | 17.754 | 22.271 | 10.226 | -12.380 | -9.743 | 24.444 |
| -9.342 | 18.945 | -7.517 | -14.498 | 27.242 | 10.149 | -10.420 | 11.491 | 26.340 | 17.833 | 6.024 | 6.482 | 28.904 | 26.590 |
| -2.330 | -19.319 | 10.264 | 23.292 | 18.688 | -5.955 | 24.923 | 9.934 | 5.080 | -24.078 | 24.792 | -8.680 | 26.147 | 25.677 |
| -16.249 | -28.143 | -6.890 | 2.749 | 0.505 | -21.219 | 26.547 | 16.825 | 1.804 | 23.408 | -13.551 | 22.606 | 20.459 | 28.029 |
| -20.323 | 7.083 | -7.828 | 15.133 | -3.674 | -5.279 | 16.925 | 18.021 | 30.672 | 15.393 | 3.279 | 9.632 | 22.941 | 18.639 |
| -3.947 | 5.330 | -12.262 | 30.660 | 15.167 | 25.774 | 11.603 | 9.088 | -11.062 | 3.292 | -12.917 | 19.993 | 0.794 | 30.478 |
| 18.789 | -22.517 | -27.173 | 29.188 | -26.696 | 14.361 | -17.164 | 4.788 | 27.184 | -24.867 | -21.085 | 18.486 | -22.688 | 20.285 |
| 21.306 | -0.867 | 12.049 | 10.649 | -21.278 | -23.567 | 2.084 | 3.594 | 15.247 | 24.452 | 23.301 | 23.695 | 23.721 | 16.822 |

| | | | | | | | | | | | | | |
|---|---|---|---|---|---|---|---|---|---|---|---|---|---|
| -20.447 | -1.475 | 15.631 | 14.701 | -24.864 | 21.103 | -19.986 | -19.175 | 25.402 | -10.765 | -29.427 | -8.179 | 19.835 | 15.557 |
| -27.273 | -3.017 | 6.812 | -24.262 | -21.868 | -25.753 | -15.042 | 20.828 | 6.433 | -3.166 | -21.709 | 15.449 | 14.829 | 23.677 |
| -18.957 | 29.407 | 20.635 | 13.387 | -10.775 | -2.679 | -14.279 | 19.782 | 24.416 | -18.849 | 20.727 | 8.436 | 20.978 | 27.669 |
| -24.742 | 28.810 | -4.381 | 11.076 | -1.222 | 26.149 | 3.473 | 20.639 | 16.620 | -3.337 | -26.436 | 23.070 | 1.670 | 14.483 |
| -15.459 | 2.247 | -9.432 | 24.034 | -17.613 | -7.766 | 0.656 | 16.095 | 25.945 | 17.818 | -3.451 | 26.343 | 18.983 | -3.054 |
| 28.190 | -1.140 | -13.524 | 8.215 | 20.760 | -10.518 | 26.814 | 5.928 | -12.455 | -21.762 | 19.830 | 20.562 | 8.834 | 16.891 |
| -8.535 | -9.519 | -6.520 | -22.391 | -19.258 | 17.462 | -1.579 | 4.906 | -5.524 | 8.857 | -14.138 | 27.233 | -11.955 | 12.171 |
| -26.594 | -6.344 | 15.364 | 9.669 | -0.953 | 2.023 | -6.019 | -17.627 | -28.534 | -26.288 | 28.153 | -16.860 | 21.521 | 15.718 |
| -0.589 | 30.464 | -16.960 | -7.769 | -15.246 | 26.661 | -14.538 | 7.522 | -12.969 | 19.682 | 22.706 | 21.589 | 28.467 | 22.082 |
| -15.895 | 22.320 | 21.514 | -28.100 | -21.385 | -12.346 | 6.475 | -15.677 | -0.453 | -15.871 | 23.684 | -18.170 | 15.183 | 9.874 |
| -10.809 | -19.623 | -16.337 | 21.552 | 3.456 | -8.320 | 25.474 | -23.517 | 4.467 | -10.658 | 26.467 | 19.085 | 20.638 | 25.350 |
| 22.841 | 0.327 | 11.616 | 24.875 | 1.617 | 10.333 | -8.786 | -21.230 | 21.525 | 12.182 | 14.756 | 14.701 | 19.738 | 22.853 |
| 20.458 | -23.052 | -20.860 | -27.331 | -30.171 | -18.633 | -2.764 | 5.775 | 26.375 | -1.336 | 12.577 | -1.446 | 19.161 | 16.680 |
| -5.081 | -11.335 | -7.037 | 27.270 | 24.392 | -26.743 | -15.830 | -1.585 | -22.497 | 11.375 | 13.267 | -24.248 | 26.552 | 20.692 |
| -21.895 | -27.657 | 19.561 | 0.019 | 8.458 | 11.537 | -3.436 | -24.585 | 28.121 | -2.833 | 9.817 | -8.202 | -2.977 | 15.875 |
| -13.743 | -29.743 | 6.210 | -25.479 | -2.830 | 5.154 | -10.573 | 18.524 | 17.006 | -18.640 | 25.776 | 13.138 | 10.971 | 13.654 |
| 27.662 | -12.800 | -3.794 | -10.691 | 28.876 | -10.843 | -18.929 | -4.500 | 14.042 | -20.438 | 24.910 | 23.124 | 22.392 | 18.816 |
| 25.396 | 21.301 | 18.490 | 27.240 | -28.765 | -17.213 | 23.630 | 13.894 | 11.874 | -0.726 | -22.607 | 21.638 | 24.438 | 18.311 |
| -1.586 | -3.626 | -26.083 | 13.361 | -29.086 | -20.967 | 13.076 | -21.744 | -29.512 | 21.831 | 10.792 | 27.412 | 25.468 | 29.923 |
| 19.337 | 26.498 | -14.789 | -18.453 | -26.753 | 18.142 | -16.475 | 7.658 | -25.037 | 9.352 | 20.429 | 21.279 | 13.987 | 26.481 |
| 15.336 | 14.888 | 20.410 | -19.998 | -10.185 | -4.593 | -10.672 | -8.497 | 16.833 | -2.600 | -21.272 | 26.109 | -14.775 | 16.835 |
| 12.673 | 10.998 | 21.287 | -1.062 | -21.720 | 25.079 | -8.455 | 17.228 | -7.797 | -1.467 | 2.737 | 0.048 | -1.759 | 29.017 |
| 3.930 | -30.676 | 16.080 | -6.761 | 28.031 | -15.994 | -14.859 | -15.921 | -15.925 | 6.365 | 11.949 | -7.870 | 24.860 | 25.983 |
| -20.704 | 28.375 | -4.926 | -14.077 | -8.708 | -18.694 | 18.434 | 5.077 | 14.788 | -3.022 | 26.630 | 7.795 | 9.119 | 25.413 |
| 8.779 | 23.046 | 30.031 | -17.837 | -25.979 | 16.519 | -19.271 | 8.607 | 22.816 | 19.723 | 27.705 | -23.330 | -15.706 | 23.404 |
| 17.877 | -3.379 | 24.395 | -12.856 | 17.458 | 12.215 | 17.057 | 16.358 | -1.734 | -5.495 | 1.703 | -13.204 | 2.167 | 19.785 |
| 9.886 | -19.815 | -14.954 | -26.158 | -9.414 | 18.484 | 13.459 | 8.045 | -20.263 | 11.494 | -6.444 | 27.064 | -16.486 | 24.488 |
| -7.652 | -18.487 | -10.576 | 6.610 | -27.544 | -23.116 | 8.632 | 19.443 | -26.363 | 21.421 | 24.600 | 25.336 | 29.471 | 13.383 |
| 23.704 | -28.220 | -2.598 | -1.903 | -20.252 | -29.704 | 25.621 | 12.381 | 12.345 | -24.011 | 8.243 | -5.100 | 16.321 | 22.904 |
| -20.036 | 0.632 | 1.227 | 17.907 | -22.223 | -9.914 | 20.107 | -5.393 | 5.269 | 11.085 | 20.400 | 23.771 | 28.741 | 29.129 |
| 2.441 | 31.138 | -6.210 | -8.911 | -23.815 | -1.134 | 4.338 | -18.474 | 23.611 | 23.890 | -9.237 | 21.869 | 25.952 | 27.847 |
| -15.423 | -17.904 | -16.047 | 18.943 | 2.613 | -8.638 | -21.233 | -0.776 | 5.945 | 19.987 | -21.150 | 17.724 | 11.960 | 17.190 |
| 25.904 | -20.640 | 29.636 | 18.401 | 22.298 | -8.293 | -21.969 | -19.409 | 4.766 | 19.333 | -19.823 | 7.112 | 11.167 | 14.445 |
| -8.181 | -3.964 | -3.686 | 24.628 | 11.898 | 9.976 | -5.098 | -17.317 | 3.099 | 21.996 | -13.120 | -13.876 | -4.320 | 21.682 |
| -9.646 | 20.049 | -18.179 | -6.109 | -5.871 | -3.825 | -5.652 | 4.824 | 25.128 | -7.691 | 21.661 | -0.794 | 24.787 | 26.810 |
| 22.826 | -4.605 | -10.274 | 14.483 | 12.797 | -17.158 | -1.407 | 24.958 | -12.042 | -24.971 | 25.511 | 6.259 | 23.367 | 6.970 |

| | | | | | | | | | | | | | |
|---|---|---|---|---|---|---|---|---|---|---|---|---|---|
| 1.764 | 27.440 | 7.963 | -17.645 | 16.269 | 14.155 | -15.701 | 14.285 | 21.068 | 22.893 | 16.393 | 22.175 | 29.684 | 25.649 |
| -28.098 | 18.543 | 28.555 | 5.162 | 26.169 | -19.911 | -6.032 | -24.610 | -3.846 | 3.767 | -2.005 | 15.084 | 20.168 | 13.451 |
| -14.572 | 25.861 | 10.298 | -20.783 | 13.772 | 23.136 | -27.585 | 21.742 | -23.662 | -1.103 | 4.682 | 20.863 | 30.527 | 10.563 |
| -29.711 | -26.259 | -17.711 | -8.082 | 11.373 | -29.605 | -19.031 | -10.754 | 11.467 | 8.668 | 11.268 | 1.721 | 6.369 | 1.244 |
| 27.896 | -16.769 | -12.737 | 0.600 | 28.685 | 20.765 | 14.025 | 23.513 | -24.601 | 18.381 | 26.725 | -25.783 | 27.101 | 21.176 |
| 24.341 | 5.773 | 26.614 | -9.797 | 28.029 | -8.679 | 13.445 | -21.291 | -7.218 | 25.870 | 15.960 | 23.738 | 19.130 | 28.972 |
| -8.803 | 13.192 | -6.027 | -15.840 | 18.838 | -29.083 | 5.876 | 22.001 | -25.044 | 21.468 | 24.851 | 25.464 | 21.483 | 20.277 |
| 30.466 | 26.986 | -4.247 | -22.955 | 20.041 | -19.200 | 22.765 | -16.293 | 27.994 | -23.188 | -6.746 | 23.030 | -16.509 | 23.928 |
| -11.187 | 20.530 | 9.277 | -0.565 | 2.484 | -6.383 | 3.621 | -23.864 | -29.523 | -6.093 | 2.155 | 26.344 | 18.167 | 12.923 |
| -4.001 | -17.213 | -4.670 | 21.691 | -9.169 | 12.163 | -22.631 | 13.229 | 10.372 | -18.683 | 10.413 | 9.146 | 29.117 | 16.661 |
| -5.041 | -19.430 | -16.833 | 13.639 | -16.864 | 25.810 | 8.070 | 20.837 | 0.686 | -0.328 | 15.049 | 29.199 | 24.972 | 8.499 |
| 16.219 | -15.718 | -8.293 | 23.204 | -21.665 | 5.034 | -23.889 | 3.655 | -26.056 | -20.031 | 11.872 | 10.781 | -26.715 | 21.914 |
| -29.151 | -18.282 | -8.833 | 15.162 | 6.294 | -11.870 | -24.570 | -22.889 | 20.396 | -24.929 | 4.720 | 24.122 | -13.486 | 28.288 |
| 19.555 | 20.023 | 4.043 | -18.119 | 23.562 | 19.126 | -26.207 | -19.356 | -11.530 | 16.445 | 20.057 | 19.841 | -12.552 | 17.152 |
| 21.305 | 28.423 | 19.006 | 23.458 | 4.425 | -17.337 | 23.756 | -6.392 | -23.848 | 8.290 | -25.499 | 4.252 | -29.042 | 12.767 |
| 26.836 | -19.839 | -13.476 | -2.497 | -12.000 | -4.508 | 25.421 | -24.083 | 13.342 | -22.307 | 26.249 | 1.117 | -21.684 | 28.437 |
| 22.219 | 29.991 | -9.095 | -26.097 | -21.343 | 15.953 | 20.774 | 2.147 | 10.013 | 17.615 | 17.435 | 27.839 | 27.766 | 26.587 |
| -22.022 | 16.131 | 28.924 | 9.226 | 24.861 | 17.669 | -1.907 | -9.890 | -19.807 | -8.510 | -5.805 | 1.436 | 13.705 | 14.396 |
| -30.763 | -20.828 | -25.858 | -17.165 | -3.756 | 29.685 | 18.461 | -21.803 | 18.648 | -17.800 | 14.916 | -21.209 | -18.276 | 7.689 |
| -23.877 | 30.262 | -19.540 | -3.336 | -7.856 | -6.896 | -2.441 | -22.383 | -10.080 | 7.348 | 14.268 | 8.500 | 21.917 | 18.231 |
| 9.299 | -18.239 | -0.608 | 17.539 | 14.080 | -28.564 | 15.483 | 5.014 | -23.001 | -16.982 | 7.534 | 20.542 | 23.209 | 19.450 |
| -7.862 | -19.718 | -15.035 | -2.133 | -15.701 | -15.483 | -13.250 | 17.347 | 28.317 | -11.993 | 23.982 | 15.911 | 15.312 | 19.953 |
| -16.744 | -31.405 | -12.264 | 27.136 | -22.470 | 14.716 | -2.862 | -9.520 | 21.097 | -22.743 | 25.413 | 21.593 | 19.295 | 10.234 |
| -26.443 | -4.247 | -3.259 | 12.137 | 3.360 | -5.607 | -3.578 | -17.454 | -0.235 | 24.238 | 0.393 | 19.156 | 10.177 | 13.267 |
| -2.673 | 5.496 | 2.157 | 13.260 | -13.293 | 9.091 | 13.328 | -11.869 | 6.980 | 23.857 | 15.766 | 12.640 | 29.755 | 23.742 |
| -18.778 | 27.622 | -5.248 | 8.176 | -23.716 | -9.224 | 24.490 | -24.794 | -29.911 | 18.475 | 8.312 | 25.023 | 11.264 | 12.501 |
| 6.530 | 20.107 | -28.323 | 16.404 | -16.032 | 2.196 | 17.016 | 1.233 | 22.978 | 20.249 | -20.115 | 7.525 | 14.187 | 19.257 |
| -7.492 | -23.079 | -19.814 | -17.746 | -19.301 | 24.184 | -26.066 | 6.141 | 13.853 | -10.165 | -8.213 | 22.228 | 27.198 | 14.281 |
| 8.080 | -27.820 | 10.613 | -3.864 | -31.064 | -12.642 | 21.591 | 18.552 | 25.793 | -20.304 | -24.033 | 28.222 | 24.859 | 7.593 |
| 24.527 | -17.766 | 19.248 | -11.597 | 22.705 | 20.169 | 28.389 | 7.449 | 2.499 | -15.662 | 17.400 | 5.457 | 23.822 | 15.764 |
| 10.541 | 19.138 | 2.779 | 3.360 | 19.340 | -3.456 | -8.626 | -1.935 | -24.367 | 20.907 | -13.252 | -15.498 | 20.196 | 13.571 |
| -27.221 | 0.850 | 14.079 | 0.279 | -14.434 | 3.323 | 24.987 | 9.278 | -17.848 | -24.514 | -22.128 | 26.301 | 17.844 | 15.333 |
| 17.514 | -3.005 | 9.465 | 28.409 | -17.675 | 19.125 | 17.304 | 15.702 | -5.967 | -12.863 | 9.686 | 15.687 | -20.610 | 14.702 |
| 29.091 | -15.135 | -7.395 | 23.976 | 2.464 | -21.687 | -6.872 | -18.180 | 17.413 | -21.028 | -21.437 | -2.333 | 10.852 | 24.542 |
| 14.519 | -1.602 | -2.327 | -13.328 | -12.441 | -12.391 | -27.578 | 25.192 | 19.752 | 11.726 | 6.216 | 17.103 | -3.913 | 23.347 |
| -21.453 | -0.176 | 19.113 | 13.888 | -1.390 | 0.938 | -16.160 | -17.827 | -3.056 | -18.370 | 5.426 | 23.821 | 16.645 | 21.020 |

| | | | | | | | | | | | | | |
|---|---|---|---|---|---|---|---|---|---|---|---|---|---|
| 21.631 | 29.313 | -19.494 | -0.371 | 24.067 | 25.706 | -20.938 | -10.167 | -7.528 | -25.333 | 19.321 | 14.622 | -2.799 | 3.000 |
| -24.094 | -28.040 | 14.612 | 2.047 | 30.886 | -22.262 | -2.843 | 25.594 | 23.884 | -5.886 | 16.037 | -2.537 | -10.732 | 18.684 |
| -5.552 | 19.418 | -6.631 | -15.579 | 13.324 | -15.900 | -12.672 | -12.750 | -13.875 | -22.612 | 17.542 | -13.290 | -15.835 | 15.881 |
| 22.031 | 22.282 | -10.386 | -22.855 | 28.055 | 27.122 | -1.121 | -19.500 | -16.252 | 2.208 | -21.695 | 21.624 | 26.121 | 17.910 |
| -6.837 | 0.914 | -12.194 | 14.792 | -6.058 | 12.149 | 18.297 | 20.542 | -4.809 | 22.284 | -19.166 | 27.524 | 22.773 | 26.021 |
| 0.177 | -29.807 | -2.632 | 26.693 | 0.076 | -24.724 | 23.451 | 15.177 | -24.893 | 25.859 | -25.080 | 18.401 | -2.220 | 20.348 |
| -4.603 | -5.715 | 16.191 | 26.153 | 19.949 | 6.938 | -20.161 | -19.064 | 13.708 | -24.277 | -7.390 | 26.799 | 5.794 | 21.315 |
| -3.059 | 28.594 | 21.967 | 27.474 | 11.309 | 10.634 | -23.113 | -3.779 | 10.166 | -19.927 | -16.772 | 21.017 | 24.648 | 19.408 |
| 28.411 | -29.077 | 7.631 | -24.248 | -9.283 | -4.269 | 30.299 | 20.765 | -17.023 | 19.622 | -24.092 | -3.315 | 19.509 | 17.427 |
| -19.125 | -26.539 | -18.047 | 3.881 | 8.499 | 14.404 | 13.308 | 4.146 | 6.754 | 19.295 | 1.873 | 22.641 | 13.649 | 27.391 |
| 3.948 | 28.341 | -21.773 | 17.577 | -21.253 | 4.205 | 17.666 | -13.871 | 22.217 | 24.094 | 8.523 | -14.822 | -11.385 | 14.912 |
| -1.739 | -0.950 | -15.377 | -19.520 | -26.064 | -6.222 | -1.204 | -26.036 | 21.691 | 0.074 | -24.590 | -23.657 | 27.470 | 25.842 |
| 0.452 | -18.318 | -2.450 | 9.991 | -14.809 | -22.542 | -25.449 | 0.262 | 7.386 | 1.451 | -13.706 | 15.442 | 26.736 | 13.133 |
| 12.725 | 18.608 | -22.920 | 25.785 | 6.500 | -19.088 | 6.135 | 3.352 | -12.218 | -2.862 | -22.148 | -21.535 | 5.600 | 22.368 |
| 16.074 | 4.120 | -9.942 | 20.730 | -26.436 | 1.171 | -28.552 | 15.873 | -27.398 | -22.523 | -11.014 | 18.715 | 9.351 | 21.637 |
| 17.404 | 19.433 | 21.518 | 18.114 | -23.260 | 15.527 | 25.559 | 16.803 | -1.121 | -1.721 | 25.470 | -20.290 | 17.816 | 24.443 |
| 24.863 | 10.984 | 5.945 | 8.854 | 31.373 | 10.700 | -21.304 | 12.785 | -18.992 | -1.006 | -22.785 | -12.622 | -24.926 | 19.913 |
| 17.733 | 6.092 | -7.069 | 28.200 | -22.055 | 23.696 | -12.524 | 2.612 | 19.920 | 25.241 | 16.737 | 24.196 | 24.596 | 22.831 |
| 28.779 | -28.476 | -5.475 | 5.812 | 29.658 | -18.380 | -7.604 | 7.121 | -10.240 | 5.768 | 10.256 | 5.534 | 14.888 | 20.516 |
| 2.275 | -20.745 | -24.903 | -10.304 | 17.303 | 21.134 | 21.435 | -3.361 | -15.694 | 23.128 | 18.453 | -2.071 | -27.661 | -19.417 |
| -25.628 | -20.180 | 4.780 | -21.835 | -5.003 | -9.474 | 11.614 | 19.344 | 12.733 | 0.661 | -12.020 | 26.425 | 22.476 | 26.641 |
| -27.943 | -22.984 | -14.447 | 7.652 | 23.751 | -24.635 | -6.608 | 25.236 | 1.387 | 5.377 | 23.242 | 14.865 | 17.123 | 16.457 |
| -29.046 | -29.179 | 2.308 | 21.390 | 30.923 | -15.072 | 3.620 | 12.626 | -4.377 | 21.442 | 15.964 | 8.800 | 7.569 | 15.610 |
| 19.028 | -1.971 | 20.732 | 20.280 | 12.340 | 3.074 | -16.675 | -18.963 | -8.233 | 20.146 | 20.245 | 28.700 | -6.450 | 18.233 |
| 10.764 | -24.270 | -16.407 | -15.095 | -26.911 | 14.388 | 20.356 | 3.238 | -7.488 | -23.957 | 23.994 | 7.424 | 19.748 | 19.177 |
| -11.978 | -18.514 | -5.581 | -5.928 | -16.513 | -9.515 | -29.049 | -14.954 | -22.052 | -27.369 | -16.732 | 17.429 | 23.280 | 7.527 |
| -1.123 | -19.360 | -2.738 | 2.502 | 14.174 | -23.926 | 2.155 | 27.437 | 18.529 | 14.152 | -18.892 | -22.362 | 13.000 | 25.035 |
| -22.793 | -27.951 | -6.494 | 8.370 | 27.347 | 22.338 | 10.738 | 20.547 | -9.463 | 22.865 | -11.699 | 19.654 | 21.593 | -17.321 |
| -24.438 | 15.747 | 5.698 | -11.688 | -20.138 | 25.300 | -9.204 | 7.894 | -2.012 | 17.909 | -19.704 | -16.895 | -1.380 | 20.810 |
| 12.085 | -0.478 | -14.302 | -19.558 | -14.124 | -27.363 | -3.711 | 6.509 | 24.890 | 6.374 | 11.035 | 21.496 | 0.054 | 19.914 |
| 27.465 | 4.022 | 24.059 | -25.714 | 16.717 | 3.717 | 27.533 | -0.170 | 9.067 | -11.305 | 5.713 | -9.910 | 28.112 | 22.906 |
| 11.730 | 20.571 | 3.027 | 9.994 | -1.210 | 0.645 | 9.355 | 16.328 | -27.971 | -13.204 | 4.041 | 18.886 | 4.223 | 25.996 |
| 21.360 | -23.950 | -12.855 | 23.730 | -8.622 | -1.186 | -10.078 | -10.289 | -5.689 | -24.675 | 21.723 | -16.052 | -5.390 | 18.574 |
| 23.914 | -0.170 | 6.933 | 4.739 | 19.205 | 13.287 | 23.315 | 11.050 | 22.399 | -23.350 | 19.986 | 10.377 | 20.296 | 24.118 |
| 15.518 | 6.253 | 7.651 | 0.276 | 7.117 | 18.728 | 26.174 | 25.192 | 19.383 | 27.606 | -26.404 | 16.854 | 25.435 | 22.565 |
| -7.086 | -21.863 | 2.761 | -25.160 | -1.728 | -14.697 | -27.680 | -1.710 | 7.104 | -19.708 | -18.603 | -8.343 | 6.115 | 13.345 |

| | | | | | | | | | | | | | |
|---|---|---|---|---|---|---|---|---|---|---|---|---|---|
| 22.053 | -28.444 | -17.968 | -8.069 | -5.704 | 10.476 | 29.219 | 2.238 | -0.557 | 23.787 | 25.217 | 15.250 | -5.982 | 17.637 |
| -18.652 | 15.143 | 21.494 | 24.072 | -17.830 | -2.381 | 24.862 | -21.373 | 21.033 | 20.900 | 14.627 | -5.327 | 16.598 | 19.600 |
| -21.786 | 20.131 | 21.818 | -25.253 | 14.508 | -7.663 | -14.259 | -7.163 | 27.393 | -26.956 | 18.342 | 13.370 | -25.939 | 6.394 |
| -19.060 | 26.725 | 18.704 | 18.268 | -3.560 | 13.418 | -27.634 | -5.847 | -13.746 | 5.982 | -19.350 | -19.807 | -15.983 | -1.055 |
| -22.868 | 19.011 | 8.983 | -22.148 | -6.682 | -28.705 | 26.086 | -6.711 | 1.969 | 5.115 | 23.353 | 3.862 | -8.234 | 15.091 |
| 8.935 | 4.674 | -4.190 | -3.918 | -19.068 | 6.256 | 12.920 | -23.814 | -24.681 | -1.158 | 24.112 | -2.444 | -11.765 | 13.856 |
| 23.135 | -25.511 | 16.995 | -17.668 | -22.338 | -3.543 | 28.346 | 25.488 | 14.998 | -15.594 | 25.644 | 19.522 | 22.235 | 13.082 |
| -14.552 | 21.878 | -5.802 | 26.986 | 29.570 | 0.935 | 20.403 | 19.962 | -19.324 | 24.627 | -8.960 | -23.022 | 26.828 | 16.495 |
| -7.017 | 16.500 | -23.556 | 22.294 | 26.958 | -19.770 | -7.942 | 1.807 | -21.644 | -19.071 | 26.147 | 26.125 | 18.059 | 15.096 |
| -13.201 | 17.725 | 2.728 | -23.958 | 21.512 | 20.501 | -25.054 | 12.598 | 25.076 | -26.785 | 6.676 | 17.652 | 21.031 | 12.221 |
| -19.172 | -17.741 | -0.439 | 5.858 | 20.439 | 12.974 | 29.132 | -26.795 | 20.462 | -19.539 | 20.349 | 19.872 | 4.874 | 21.574 |
| 19.476 | -5.295 | 7.291 | -21.357 | -24.829 | 27.657 | 6.055 | 22.625 | 18.270 | -25.323 | 22.855 | 16.190 | -11.155 | 8.767 |
| -14.416 | 21.406 | 20.496 | 11.434 | 15.466 | -19.034 | -22.154 | 8.033 | 11.450 | -13.047 | 18.836 | 11.106 | 24.126 | 19.265 |
| -24.871 | -20.430 | -1.248 | -7.977 | 24.881 | 24.739 | -17.142 | 25.830 | 8.157 | -28.197 | 12.946 | 4.787 | 12.376 | 14.292 |
| -6.773 | 12.534 | 26.257 | 18.812 | 6.665 | -2.224 | -29.134 | -7.470 | 23.182 | -0.667 | -25.293 | 26.635 | 25.683 | 10.327 |
| 29.425 | 27.271 | -9.955 | 17.421 | -25.520 | -3.104 | 7.648 | -12.773 | -1.544 | 13.415 | 28.032 | 29.263 | 27.238 | 13.219 |
| 28.016 | 22.519 | -17.246 | 7.110 | 10.213 | -1.502 | 19.311 | 15.895 | 6.742 | 4.486 | 15.805 | 14.640 | 17.335 | 19.228 |
| 24.162 | -20.964 | 12.251 | 2.227 | -8.260 | -26.946 | 13.391 | 15.267 | 18.465 | 18.001 | -22.925 | 28.005 | -17.710 | 4.183 |
| -25.886 | -5.113 | -23.007 | 25.310 | -20.692 | 9.627 | -0.840 | -22.458 | 17.802 | 2.625 | 22.231 | 30.909 | 14.821 | -14.919 |
| -1.875 | -25.610 | 5.849 | -7.536 | 16.635 | 16.839 | 26.043 | 13.954 | -5.559 | 2.091 | 9.876 | -1.333 | -20.441 | -17.299 |
| -10.318 | 26.753 | -0.157 | 26.629 | 28.072 | -8.675 | -6.645 | 19.671 | 26.730 | -7.045 | 23.755 | 23.250 | 17.914 | -21.268 |
| 26.288 | 7.978 | -11.662 | 0.840 | 0.213 | -18.519 | 20.339 | 18.000 | -27.096 | 26.818 | 19.432 | 1.531 | -16.783 | 10.090 |
| 3.648 | 0.486 | -16.471 | 24.635 | 21.161 | 15.720 | -11.782 | -19.202 | -22.963 | -18.837 | 23.300 | -5.270 | -13.562 | 15.882 |
| -0.757 | -19.607 | -6.621 | 17.943 | -11.407 | 8.379 | -22.278 | -12.375 | 17.500 | 17.779 | 27.438 | 26.678 | 0.813 | -24.594 |
| 9.564 | 21.422 | 1.812 | 13.925 | -23.743 | 13.826 | 25.140 | 18.368 | 7.690 | 0.519 | 25.395 | 23.961 | 22.976 | 21.441 |
| -2.761 | -20.773 | -11.500 | 21.617 | -20.090 | -9.171 | -3.104 | 17.078 | 12.995 | 28.951 | -3.957 | 17.315 | 25.152 | 21.793 |
| 15.581 | 13.448 | -4.795 | 20.299 | 18.358 | -25.447 | 1.835 | -0.742 | 5.664 | 10.009 | -11.141 | 19.128 | 20.512 | 23.528 |
| -12.787 | 23.339 | 13.094 | -3.863 | -7.749 | 4.443 | 27.754 | -10.250 | 4.741 | -2.734 | 4.469 | 9.798 | -18.716 | 23.509 |
| -28.155 | -21.539 | 18.777 | -15.295 | -18.253 | 21.513 | 22.609 | 28.459 | -18.285 | -23.273 | 14.327 | 18.405 | 16.769 | 0.229 |
| 11.006 | 17.242 | -17.463 | -4.995 | 29.053 | -9.443 | -29.790 | -18.494 | 21.383 | 16.324 | -8.579 | -8.726 | 7.052 | 22.271 |
| 8.333 | 12.182 | 21.791 | 23.494 | -5.510 | 1.589 | 28.774 | 22.360 | 4.326 | 16.784 | -3.223 | 10.111 | 23.282 | 21.425 |
| 28.976 | 22.881 | 20.636 | 3.796 | 29.777 | -4.182 | -7.385 | -13.610 | -12.798 | -14.090 | 25.192 | 20.512 | 18.381 | 25.410 |
| 0.107 | 22.325 | -8.957 | 11.732 | 1.368 | 14.404 | -23.935 | -5.215 | -1.657 | -20.164 | 10.171 | -0.362 | 17.822 | -10.127 |
| 9.100 | 8.107 | -7.335 | 25.768 | 14.517 | -13.607 | -10.618 | 21.935 | -11.611 | -21.857 | 16.470 | 3.980 | -5.543 | 20.295 |
| -0.730 | 11.997 | -6.571 | -14.456 | 9.372 | -16.563 | -20.096 | 5.752 | 21.590 | 21.029 | 23.958 | -6.498 | 8.328 | 19.682 |
| -4.356 | -26.729 | -3.297 | -5.750 | -26.578 | 21.872 | -2.464 | 0.307 | 19.280 | -26.984 | 27.604 | 27.509 | 13.145 | 24.772 |

| | | | | | | | | | | | | | |
|---|---|---|---|---|---|---|---|---|---|---|---|---|---|
| -22.373 | -6.047 | 12.287 | -18.488 | 10.617 | 20.557 | -19.284 | 9.513 | -3.390 | 8.711 | 21.740 | 14.164 | 6.226 | 26.096 |
| -13.454 | 21.470 | 22.123 | 17.643 | -7.386 | 25.950 | 23.350 | -18.784 | 18.552 | 3.195 | 18.304 | 2.683 | -19.624 | 18.977 |
| 27.601 | -16.976 | 3.266 | 22.446 | -27.625 | -16.137 | 11.077 | 2.643 | -1.943 | -24.762 | 26.478 | -5.226 | 17.293 | 20.907 |
| -19.755 | -10.542 | -8.788 | 23.862 | -18.881 | 23.950 | 12.300 | 27.723 | -18.768 | -21.292 | -11.219 | -9.118 | 20.572 | 22.249 |
| 19.807 | 9.052 | -22.509 | -4.715 | -24.493 | 21.456 | -22.594 | -1.067 | 12.563 | -22.870 | -23.202 | 25.556 | 17.548 | 25.563 |
| 3.531 | 24.436 | -17.633 | 8.924 | 0.326 | -25.776 | -21.415 | -14.837 | 18.341 | -2.500 | 19.971 | 20.766 | 10.823 | 19.191 |
| -10.542 | -22.946 | 1.355 | -14.899 | -9.295 | -21.307 | 19.505 | 19.161 | 25.361 | 20.026 | -4.890 | 8.578 | 26.728 | 17.039 |
| -1.777 | -6.906 | -18.816 | 7.651 | 13.599 | -1.902 | -23.556 | 16.272 | 11.770 | 5.955 | -14.792 | 11.258 | 25.335 | 18.590 |
| -20.862 | -13.336 | -1.467 | -6.325 | 6.785 | 25.648 | -25.514 | 4.857 | 18.974 | -4.109 | -13.390 | 26.601 | 4.404 | 26.625 |
| 2.645 | 1.691 | -24.854 | 25.858 | -4.477 | 20.968 | 20.680 | -2.685 | 20.154 | 15.166 | 20.636 | 17.219 | 19.900 | 23.787 |
| 11.808 | -6.067 | 8.573 | 24.730 | -22.548 | 15.540 | -16.125 | 13.815 | 17.467 | 24.750 | 28.640 | 5.852 | -15.275 | 15.675 |
| -3.697 | 25.108 | -13.564 | -3.981 | -14.143 | 13.680 | 26.631 | -14.167 | 21.790 | -11.164 | -15.636 | 28.593 | 11.920 | 14.841 |
| 8.552 | -15.436 | 2.866 | -1.855 | -26.726 | 18.812 | -29.350 | -17.319 | 16.763 | 23.058 | -21.127 | 28.813 | 13.272 | 16.566 |
| 13.839 | -24.130 | -16.287 | 14.410 | -15.306 | -16.166 | 18.761 | 17.667 | 25.759 | 30.278 | -7.169 | 23.360 | 12.788 | 2.602 |
| 22.388 | 19.502 | -5.970 | 8.507 | 2.691 | -11.866 | -12.095 | 22.321 | 21.183 | 29.702 | -28.310 | 27.673 | 3.408 | 25.093 |
| -19.576 | -9.819 | -15.579 | 16.782 | 27.608 | -3.209 | -25.843 | -26.391 | -13.191 | 19.451 | 1.411 | 18.647 | 18.692 | 26.780 |
| -28.214 | 0.980 | -5.595 | 23.534 | -19.697 | -5.416 | -2.734 | 26.342 | -4.746 | -2.307 | 18.965 | -3.719 | 8.553 | 20.742 |
| -2.213 | 22.948 | -7.441 | 22.589 | -17.415 | -2.269 | -24.918 | 12.687 | -19.864 | 3.477 | 15.372 | 27.181 | -10.814 | 19.306 |
| -4.014 | -1.826 | -9.315 | 3.403 | -25.861 | -9.743 | -27.709 | 17.052 | -25.151 | -18.195 | 21.237 | 9.535 | 26.571 | 22.242 |
| -23.498 | 24.998 | 5.823 | 11.052 | -28.310 | -4.666 | -30.075 | -17.713 | -3.143 | 15.002 | 24.294 | 9.246 | -4.946 | 11.086 |
| -26.332 | -9.736 | 3.611 | 24.693 | 29.240 | -5.747 | 26.014 | -1.225 | -8.041 | 7.393 | 23.107 | 21.244 | 9.855 | 24.494 |
| -26.754 | 3.295 | 4.877 | -1.455 | 14.017 | -13.688 | 22.742 | -9.024 | 6.812 | -6.158 | -10.491 | 20.589 | 25.231 | 27.386 |
| 11.366 | -26.003 | 15.812 | 11.806 | -1.699 | 0.645 | 27.952 | -7.774 | -23.296 | -7.118 | -5.153 | 15.273 | 5.301 | 27.642 |
| 23.301 | 20.563 | 0.722 | 19.057 | 12.109 | 5.565 | -23.523 | 19.100 | 11.552 | -26.610 | 19.963 | 7.672 | 13.129 | 21.539 |
| 18.220 | 15.555 | 1.443 | 8.262 | -18.631 | -14.029 | 18.686 | 22.944 | 23.117 | -0.130 | -28.053 | -20.407 | -5.201 | 27.347 |
| 30.451 | 24.620 | -8.207 | -19.854 | 26.954 | -22.612 | -28.803 | 4.413 | 16.814 | 18.351 | 9.461 | 24.898 | 14.540 | 18.808 |
| -25.060 | 17.752 | 19.127 | 14.786 | 27.739 | 19.519 | 22.391 | -13.618 | -1.434 | 29.904 | 18.352 | 17.208 | 18.990 | 25.254 |
| -5.528 | -19.660 | -6.438 | 24.869 | -24.048 | 4.094 | -30.256 | 20.890 | -5.586 | 23.851 | 24.827 | 16.417 | -1.343 | 22.863 |
| 5.143 | -3.128 | -23.766 | 21.376 | -7.560 | -18.231 | 6.993 | -10.484 | -12.165 | 2.165 | -18.499 | 25.911 | 9.516 | 26.191 |
| 19.984 | -20.890 | 24.756 | 23.566 | 2.839 | 16.109 | 6.708 | 9.528 | -19.812 | 28.724 | -25.136 | 8.664 | 19.747 | 24.141 |
| 13.802 | -1.502 | 18.841 | 4.531 | 17.708 | -11.742 | -9.213 | -21.387 | -14.240 | 21.998 | -20.728 | -7.610 | 15.807 | 17.450 |
| -14.051 | 7.821 | 9.817 | -21.011 | 16.348 | 1.281 | -25.125 | -19.331 | -24.698 | 30.739 | 17.719 | 11.411 | -7.533 | 11.965 |
| -20.842 | 2.092 | 1.487 | 24.358 | 23.330 | 10.020 | 28.844 | -2.367 | 15.967 | 29.469 | 20.345 | -30.151 | 18.901 | 19.807 |
| 25.877 | 14.822 | -20.610 | -17.694 | -3.052 | 2.708 | 20.376 | -11.281 | 9.042 | 11.516 | 27.702 | 26.667 | 20.564 | 19.472 |
| 15.924 | 19.103 | 6.143 | -7.066 | -13.536 | -21.919 | -27.488 | -5.700 | 6.209 | -0.521 | -29.283 | -31.705 | -1.424 | 27.370 |
| -13.324 | 0.856 | 0.020 | -18.720 | 24.438 | 20.172 | 23.693 | -2.437 | 11.257 | -30.775 | 4.585 | 25.534 | -16.556 | 16.510 |

| | | | | | | | | | | | | | |
|---|---|---|---|---|---|---|---|---|---|---|---|---|---|
| -15.711 | 9.325 | -17.002 | 18.107 | -23.374 | 20.753 | 19.102 | -0.479 | 12.927 | -1.440 | -12.714 | 18.576 | 7.188 | 26.145 |
| 24.516 | 4.589 | 25.173 | -5.513 | 21.045 | -10.806 | -23.261 | 22.982 | 23.309 | -6.256 | 15.012 | 20.102 | -17.866 | 27.797 |
| 28.843 | -15.093 | 7.994 | 25.422 | 1.642 | 12.664 | -24.704 | 11.276 | 1.893 | 5.879 | 19.038 | -5.861 | 11.479 | 15.523 |
| -27.822 | -22.163 | 6.266 | -11.709 | -0.348 | -17.609 | -30.497 | -1.974 | 0.306 | 26.530 | -24.561 | -19.511 | 15.808 | 25.104 |
| 7.269 | -18.321 | -5.734 | 6.361 | -21.662 | 16.334 | 26.894 | -26.129 | 21.440 | 20.827 | -14.543 | 15.827 | 14.382 | 20.173 |
| 21.433 | 20.938 | 26.355 | -16.682 | 26.272 | 9.258 | 17.041 | 0.468 | -5.451 | -30.033 | -5.553 | 17.889 | 12.563 | 27.104 |
| 9.157 | 10.359 | -19.237 | 26.562 | -9.722 | -1.097 | -5.782 | -29.834 | -22.735 | 2.479 | 29.180 | 20.389 | 10.729 | 23.320 |
| -8.659 | -15.800 | 1.835 | 25.576 | -19.272 | 11.906 | -22.883 | 12.446 | 15.097 | 27.368 | 23.911 | -23.028 | 13.249 | 17.722 |
| 18.220 | 18.373 | -22.957 | 2.321 | -6.781 | -19.720 | 27.512 | -19.585 | 13.304 | -30.485 | 21.895 | 23.469 | 8.586 | 19.419 |
| 17.089 | -7.131 | -2.160 | 14.352 | -27.004 | 1.320 | -20.038 | -4.006 | 8.904 | 17.820 | 13.199 | 24.792 | -6.667 | 16.474 |
| -11.579 | 24.648 | -7.318 | 8.245 | -5.140 | -6.578 | -11.938 | -16.808 | 4.795 | 4.315 | -26.804 | 26.561 | -11.396 | 17.251 |
| -2.387 | -24.144 | 5.548 | -26.437 | -25.069 | -11.161 | 22.274 | 16.346 | 23.616 | 19.688 | 11.803 | 9.499 | -16.555 | 29.050 |
| -21.966 | 22.609 | -24.961 | 4.663 | -17.564 | -16.921 | -25.063 | -9.622 | -6.845 | 2.762 | 24.499 | 19.673 | 17.494 | 25.201 |
| 21.464 | -20.171 | 4.323 | -13.588 | -11.797 | -16.025 | 6.651 | 13.972 | 16.951 | 20.733 | 20.634 | 20.192 | -4.489 | 20.312 |
| -19.789 | 19.249 | 22.867 | -19.759 | 9.095 | 3.096 | -2.065 | -27.150 | 13.850 | -8.989 | -22.514 | 24.217 | -18.225 | 23.211 |
| 8.668 | 1.236 | 3.885 | 19.111 | -3.320 | 14.375 | 25.325 | -10.992 | -13.105 | -1.022 | 15.652 | 27.259 | -20.020 | 16.124 |
| -20.450 | -19.127 | -15.124 | -25.466 | -21.483 | 24.349 | 26.831 | 4.728 | 9.545 | -4.002 | -30.346 | 20.921 | 7.497 | 11.464 |
| -24.471 | 0.281 | -19.644 | 21.144 | -8.392 | 23.870 | -29.138 | -12.139 | -14.853 | 30.804 | 16.346 | 18.541 | 10.514 | 26.491 |
| 2.372 | -14.643 | -24.775 | 8.912 | 26.061 | 11.367 | -11.057 | 20.714 | 3.356 | -20.096 | -27.756 | 25.304 | 6.754 | 14.670 |
| -0.285 | 23.525 | -6.162 | -2.840 | -1.062 | 21.344 | -23.447 | 18.641 | 15.961 | -24.891 | 22.654 | -15.539 | 15.185 | 28.570 |
| 23.692 | 24.229 | -15.761 | -20.610 | 24.250 | 17.520 | -2.338 | 13.080 | 3.705 | 0.674 | 5.002 | 26.203 | 9.569 | -4.218 |
| 29.603 | -11.621 | -4.517 | -13.842 | -2.355 | 3.744 | -16.518 | 19.624 | 0.675 | -2.412 | -13.283 | 19.876 | -5.214 | 20.424 |
| 13.099 | 7.354 | -10.010 | -25.021 | -26.186 | 15.022 | 22.796 | 15.734 | 1.649 | -6.987 | -19.610 | 21.130 | -9.587 | 18.232 |
| 7.495 | 2.948 | -20.886 | 10.811 | 10.521 | -0.904 | 12.074 | -26.760 | -13.358 | 5.697 | 28.059 | 22.074 | -11.272 | 23.415 |
| 4.168 | 26.898 | 0.672 | 5.077 | 5.668 | 25.695 | 3.192 | -0.784 | -18.306 | 21.868 | 27.240 | -11.464 | -19.712 | 20.934 |
| -7.217 | 25.676 | -0.268 | -26.218 | -19.132 | 2.143 | 27.520 | -5.156 | 15.644 | 16.694 | -18.773 | 28.699 | 5.126 | 27.352 |
| 1.310 | -5.587 | 11.005 | 19.829 | 19.422 | 7.796 | -20.769 | -21.568 | 21.566 | 22.619 | -27.565 | -10.800 | 11.905 | 15.579 |
| -3.400 | -0.938 | -4.061 | 21.176 | 7.402 | -3.742 | 29.131 | 9.679 | 9.327 | -4.085 | 25.247 | 23.353 | 2.914 | 24.994 |
| 1.573 | -17.153 | -5.290 | -24.581 | 13.285 | 24.031 | 12.706 | 10.409 | 21.086 | 8.958 | -20.952 | 6.161 | -23.507 | 29.168 |
| 8.832 | -6.242 | -18.764 | -15.889 | -6.502 | 20.524 | -4.103 | 7.430 | -5.152 | 6.669 | 15.260 | 20.636 | 8.822 | 26.775 |
| 19.138 | 8.331 | -9.475 | -21.402 | -15.940 | 14.506 | 2.175 | 5.541 | 5.578 | 28.925 | -22.285 | 1.226 | -4.445 | 29.666 |
| 26.441 | 26.863 | -15.097 | 3.960 | -23.246 | -22.237 | 21.532 | -29.043 | 7.598 | -6.120 | -23.453 | -29.801 | -17.940 | 25.051 |
| -24.133 | 20.085 | 24.919 | 16.539 | 27.806 | 1.098 | -27.722 | 2.568 | 22.242 | 25.345 | -5.141 | 28.348 | -22.845 | 18.475 |
| 22.180 | -20.903 | -17.439 | -18.455 | 7.883 | -13.545 | 28.541 | -8.199 | -1.124 | 5.600 | 26.742 | 25.355 | 8.217 | 28.198 |
| 0.020 | 18.349 | 0.601 | 12.837 | -16.887 | 21.065 | 14.292 | -15.057 | -12.400 | -30.358 | -8.474 | 14.312 | 21.502 | 12.988 |
| 24.896 | -6.889 | 0.159 | 24.682 | 3.122 | 19.894 | 1.429 | 16.709 | -14.296 | 14.656 | 18.253 | 17.562 | 18.588 | 17.626 |

| | | | | | | | | | | | | | |
|---|---|---|---|---|---|---|---|---|---|---|---|---|---|
| 10.088 | 15.391 | -12.342 | -16.292 | -11.795 | 12.548 | -1.265 | 14.758 | 17.223 | -3.124 | -8.087 | 2.962 | 24.513 | 24.502 |
| -16.547 | 6.801 | 1.967 | -8.726 | 16.497 | -2.102 | 19.978 | 21.925 | 4.012 | -0.530 | 30.437 | -3.975 | 0.487 | 7.340 |
| 21.052 | 9.592 | -3.343 | -23.660 | 17.355 | 13.815 | 22.971 | -22.245 | 22.572 | 8.056 | 21.964 | 20.268 | -5.434 | 16.309 |
| -1.601 | 18.767 | -2.850 | 27.652 | -20.489 | 20.755 | -21.292 | -29.829 | -10.578 | -15.158 | 19.711 | 28.290 | -11.281 | 30.237 |
| 5.533 | 5.275 | -9.794 | -27.436 | -4.021 | 16.413 | 5.673 | 1.278 | 8.563 | -21.235 | 17.363 | 4.629 | -1.811 | 26.914 |
| 19.319 | 21.435 | -24.682 | -15.275 | 23.295 | 18.458 | 19.528 | 22.978 | -4.586 | 26.336 | -30.930 | 10.654 | -2.825 | 23.127 |
| -1.132 | 26.851 | -14.018 | 22.250 | 13.918 | 21.724 | 4.597 | -30.899 | 24.195 | 4.213 | 15.537 | 18.909 | 3.863 | 22.080 |
| -13.385 | -24.715 | -0.518 | -3.472 | 8.330 | 15.466 | 3.978 | 16.247 | 6.345 | 22.865 | 23.612 | 25.994 | -4.959 | -3.604 |
| -6.948 | 24.491 | -8.408 | -28.135 | -17.693 | 22.779 | -29.508 | -10.653 | -2.221 | 0.778 | 11.149 | 27.223 | -1.238 | 18.479 |
| 15.810 | -13.136 | -7.468 | 23.768 | -2.010 | 22.153 | -28.139 | 24.322 | -13.612 | 5.135 | 24.741 | 23.834 | 24.879 | 14.534 |
| -6.401 | 14.445 | -16.413 | 26.674 | -7.156 | 26.266 | -22.955 | 21.122 | -26.943 | 16.796 | -9.376 | 23.356 | 17.981 | 17.195 |
| 16.477 | 18.345 | -5.739 | -10.614 | -22.247 | 3.879 | -26.970 | -12.805 | -17.787 | -27.927 | 27.692 | -8.803 | -3.642 | 23.272 |
| -19.317 | -2.956 | -27.951 | -26.115 | -12.669 | 21.350 | 21.356 | -5.343 | 1.139 | -1.328 | 6.017 | 8.854 | -9.025 | 15.621 |
| 7.346 | -3.788 | 5.412 | 9.959 | -3.677 | 27.107 | 28.796 | -9.866 | 5.759 | -2.824 | 24.521 | 10.326 | 0.266 | 15.190 |
| 12.168 | 27.460 | -4.920 | -16.369 | -27.735 | 18.122 | 24.160 | 10.296 | 22.732 | -13.636 | 26.011 | 27.655 | 0.732 | 19.532 |
| -22.760 | -22.925 | -9.308 | 21.232 | -18.872 | 21.225 | 6.812 | -30.534 | 23.583 | 24.147 | 9.618 | 28.503 | -2.042 | 29.013 |
| 27.899 | 3.649 | -27.023 | 14.956 | -21.453 | 28.198 | -24.922 | 21.831 | 9.623 | -5.822 | 28.984 | 7.655 | -4.011 | 27.511 |
| -3.469 | 28.278 | -3.395 | 24.499 | 21.393 | 19.368 | -0.534 | 28.909 | -25.861 | 19.788 | 28.443 | 11.698 | 0.198 | 13.619 |
| 11.292 | 23.906 | -2.896 | 6.090 | 17.974 | 22.751 | 6.579 | 2.217 | 11.485 | 0.908 | 16.796 | -6.698 | 14.366 | 25.964 |
| 10.342 | -25.354 | -9.044 | -26.136 | -17.096 | 17.842 | 5.220 | -3.389 | -23.072 | -25.239 | -28.499 | 27.067 | 25.842 | 15.120 |
| -13.375 | -10.061 | 3.715 | -27.451 | -23.142 | 22.147 | 15.780 | 9.079 | -25.156 | 21.702 | 18.407 | -6.757 | 4.933 | 17.861 |
| -17.624 | 21.713 | 2.691 | -5.452 | -24.071 | -0.103 | -4.235 | -3.700 | 16.577 | 22.603 | -29.325 | 28.216 | 23.688 | 22.381 |
| -26.845 | 0.780 | -3.791 | 11.412 | 21.998 | 16.773 | -8.613 | 25.691 | 20.819 | -2.353 | 1.231 | 19.071 | 5.458 | 24.442 |
| 23.266 | -11.873 | 19.628 | 17.359 | -8.324 | 25.716 | 26.448 | 18.040 | 14.091 | 4.229 | 31.631 | 22.731 | 18.203 | 26.364 |
| 14.832 | -22.464 | -4.088 | -3.333 | 0.877 | -5.093 | 25.903 | 23.045 | -21.651 | 6.248 | 0.188 | 8.451 | -2.118 | 29.996 |
| 19.965 | -17.133 | -21.582 | 24.234 | 26.013 | 17.865 | 18.234 | 11.438 | 23.509 | -29.016 | 30.015 | 27.088 | 24.807 | 5.329 |
| -9.538 | 26.250 | -6.119 | -28.499 | 16.667 | 18.967 | 11.669 | 1.142 | 13.680 | -7.871 | 18.910 | 14.469 | 23.053 | -6.723 |
| -7.640 | 27.756 | -0.329 | -8.672 | 5.864 | 15.702 | 4.616 | 13.639 | 8.754 | -18.039 | 14.131 | 22.019 | -5.461 | 22.904 |
| 22.939 | -0.135 | -23.231 | 25.183 | -1.008 | 8.871 | -21.892 | 22.152 | 3.502 | 0.753 | -11.643 | 26.102 | 26.196 | 20.216 |
| 19.174 | 12.254 | -1.720 | 28.683 | -9.883 | 28.517 | -9.750 | 23.619 | -19.884 | -4.131 | -28.995 | 25.660 | 0.516 | 28.980 |
| -24.888 | 20.653 | -4.903 | 4.733 | 7.742 | 25.430 | 15.864 | -1.273 | -1.806 | -3.008 | 16.565 | 2.991 | -12.876 | 15.614 |
| -3.623 | -25.091 | 24.654 | -24.227 | -19.360 | 19.835 | -25.891 | 7.094 | 17.818 | -1.386 | -18.915 | -25.763 | -5.512 | -2.683 |
| 11.317 | 0.578 | 17.374 | 15.165 | 23.735 | 22.385 | -6.424 | -8.437 | -3.304 | -9.321 | 24.253 | 15.184 | 27.808 | 29.947 |
| -9.484 | 25.134 | 17.731 | 14.344 | -23.636 | 16.466 | 25.098 | 9.678 | 26.484 | -19.734 | 21.386 | 25.501 | -24.160 | 17.991 |
| 25.175 | 15.367 | -2.644 | 1.516 | -7.352 | 14.426 | 0.975 | -25.452 | 17.199 | 18.004 | 4.019 | 19.541 | 28.386 | 29.323 |
| 21.052 | -25.642 | -22.128 | 29.638 | 2.200 | 18.527 | -19.937 | -5.760 | 2.412 | 0.175 | -30.104 | 27.457 | 25.715 | 12.152 |

| | | | | | | | | | | | | | |
|---|---|---|---|---|---|---|---|---|---|---|---|---|---|
| 21.623 | 1.243 | -3.182 | 0.101 | 1.577 | 21.387 | 10.352 | 26.479 | 14.194 | 27.290 | 19.667 | 25.964 | 13.577 | 23.021 |
| 13.507 | -16.378 | -26.267 | -15.483 | 18.910 | 23.011 | -3.448 | -20.312 | -9.886 | 20.108 | 26.366 | 2.512 | 24.443 | 23.935 |
| -16.994 | -8.109 | -28.255 | -4.362 | -11.763 | 10.560 | -1.091 | 8.335 | -13.825 | -16.313 | 20.746 | -14.851 | 3.178 | 16.959 |
| 18.424 | -10.820 | -0.867 | -28.247 | 6.793 | 21.433 | -2.014 | 14.340 | 27.668 | -6.334 | 12.308 | 24.248 | 23.717 | 25.874 |
| -18.755 | -17.341 | -17.840 | -21.611 | 17.669 | 18.898 | 1.892 | 2.640 | 21.449 | 21.729 | -22.792 | 24.902 | -4.971 | 25.470 |
| 9.232 | 4.429 | -8.618 | 24.638 | 4.803 | 13.407 | -16.708 | -27.465 | -26.268 | 0.832 | 13.869 | 20.359 | 14.514 | 19.358 |
| 14.616 | 6.067 | -12.535 | 9.240 | -1.766 | 26.651 | -9.875 | 3.533 | 6.022 | -29.097 | -24.874 | 9.489 | 17.674 | 12.586 |
| -21.995 | -20.801 | -23.543 | -8.025 | 6.006 | 17.605 | -18.428 | 23.842 | 26.517 | -22.473 | 22.961 | 23.843 | 3.269 | -9.420 |
| 25.241 | 5.637 | 9.972 | 21.782 | 11.338 | 9.262 | -1.456 | 16.414 | 24.715 | -5.876 | -18.327 | 20.131 | 17.117 | 20.727 |
| -11.292 | -1.446 | -26.522 | 23.338 | -5.774 | 20.267 | -9.514 | 28.043 | 10.672 | 16.392 | 2.005 | -4.557 | 27.768 | 21.446 |
| -4.611 | 29.469 | 9.538 | 11.220 | -20.254 | 23.515 | 21.785 | 29.539 | -9.494 | -11.691 | -28.751 | 23.465 | 2.777 | 2.241 |
| -13.811 | 25.522 | 1.809 | -14.910 | 20.495 | 11.550 | 5.553 | 21.124 | -11.869 | -1.862 | 28.571 | 9.016 | 24.978 | 8.012 |
| -8.179 | -27.776 | -12.521 | 20.525 | -10.158 | 24.291 | 18.539 | -0.276 | -4.768 | 27.931 | -23.206 | -7.877 | 24.443 | 5.908 |
| 8.535 | -20.350 | -23.081 | -22.510 | 22.612 | 29.254 | -4.203 | -4.129 | 12.328 | 12.653 | 29.265 | 17.489 | 22.801 | 15.476 |
| -17.736 | -8.675 | 1.140 | 29.889 | -1.179 | 21.853 | -15.202 | -21.965 | -18.972 | 9.552 | 3.094 | 23.130 | 0.098 | 22.672 |
| -15.517 | -25.640 | -1.879 | -11.031 | -24.855 | 18.159 | 27.427 | -22.829 | 8.717 | -1.152 | -14.668 | 25.795 | 9.232 | 4.806 |
| -4.213 | -7.454 | -1.350 | -23.997 | 21.445 | 18.833 | 23.256 | 23.966 | 10.068 | 16.153 | 31.405 | -14.162 | 26.153 | 21.272 |
| 4.856 | -3.940 | -5.490 | 4.761 | -3.903 | 19.503 | -3.023 | 15.960 | -11.511 | 10.597 | 20.063 | 14.986 | 24.907 | 15.977 |
| 23.747 | 16.004 | -4.597 | 13.347 | -14.459 | 19.945 | -20.769 | 5.389 | 26.156 | -25.483 | -13.817 | 23.060 | 1.857 | 15.304 |
| 20.144 | -15.589 | -18.933 | -21.739 | -11.411 | 8.831 | 6.301 | -23.740 | 13.737 | 9.598 | 17.608 | 6.894 | 21.643 | 25.970 |
| 24.854 | -2.208 | 27.937 | -26.577 | 14.729 | 23.969 | -8.033 | 31.380 | 2.547 | 6.608 | -27.909 | 2.436 | 2.234 | 23.262 |
| 19.701 | -5.048 | -3.086 | -27.504 | -2.911 | 22.348 | 12.303 | 1.369 | 10.613 | 4.623 | 18.743 | -17.174 | -0.640 | 26.867 |
| -6.234 | 6.182 | 3.178 | 24.157 | 23.055 | 17.636 | -18.126 | 17.385 | 19.692 | -22.893 | -3.561 | -0.061 | 29.940 | 1.169 |
| -9.055 | -2.360 | -6.378 | -24.559 | 24.045 | 15.866 | 1.859 | 12.444 | -12.644 | -6.389 | 16.460 | 11.328 | 26.554 | 27.932 |
| 0.936 | -28.339 | 27.640 | -29.484 | -18.383 | 0.189 | -16.032 | -28.810 | -4.314 | -21.277 | 24.722 | -2.844 | 23.251 | 25.263 |
| 3.808 | 22.117 | -25.270 | 22.001 | 11.172 | 19.424 | 10.080 | 30.289 | 9.270 | -1.783 | 25.201 | 9.536 | 23.973 | 29.873 |
| 26.356 | -23.779 | -11.869 | -18.321 | -5.463 | 17.114 | -5.721 | 2.562 | -8.999 | 25.726 | -11.175 | 7.263 | 25.223 | 19.155 |
| -21.266 | -23.018 | 30.151 | 22.850 | -6.689 | 18.594 | 27.166 | -1.349 | -3.712 | -9.103 | -26.819 | -1.264 | 25.526 | 13.973 |
| 14.743 | 10.729 | -21.960 | -17.699 | -17.267 | 19.296 | 18.954 | 4.465 | -8.385 | -13.144 | 27.731 | -20.488 | 26.173 | 14.564 |
| -24.473 | -21.151 | 0.338 | 23.723 | 3.879 | 14.471 | 12.580 | 6.078 | -17.858 | -6.397 | 12.908 | 21.140 | 27.133 | 7.633 |
| 11.523 | 23.142 | -20.367 | -29.811 | -5.821 | 25.979 | -20.195 | 16.429 | -24.259 | -25.125 | 9.605 | -11.820 | 28.079 | 17.796 |
| 19.128 | 25.285 | 16.808 | 9.455 | -0.185 | 20.862 | 4.593 | 22.045 | -26.410 | 14.421 | -18.047 | 18.630 | 4.569 | 9.332 |
| 6.787 | 24.358 | -0.375 | -15.853 | -1.276 | 17.817 | 3.955 | -18.945 | 13.851 | 19.469 | 30.436 | 13.403 | -6.937 | 0.770 |
| -17.066 | 19.868 | -4.651 | -30.203 | -26.530 | 17.509 | 7.928 | -7.329 | 26.952 | 4.156 | 18.881 | 1.828 | 25.414 | 27.419 |
| 2.360 | -28.942 | -16.820 | 28.455 | -27.264 | 18.480 | 5.844 | 19.138 | 7.593 | 0.913 | -8.038 | 10.166 | 20.276 | 30.182 |
| 13.505 | -1.242 | -6.668 | -25.451 | 23.765 | 15.690 | -4.602 | -16.442 | 2.818 | -3.934 | -10.842 | 12.303 | -7.910 | 8.435 |

| | | | | | | | | | | | | | |
|---|---|---|---|---|---|---|---|---|---|---|---|---|---|
| 13.753 | -0.624 | -20.992 | -19.988 | -18.939 | 31.683 | 8.625 | -12.067 | -6.350 | -20.868 | 21.370 | 9.098 | 30.638 | 16.458 |
| 1.241 | -30.404 | -26.605 | -16.744 | 16.171 | -5.340 | -13.764 | -0.328 | 24.189 | -2.633 | 14.729 | 7.369 | 24.052 | 26.562 |
| -22.301 | -19.405 | -21.003 | -20.513 | 7.567 | -3.781 | -2.919 | -27.179 | -14.159 | 12.888 | 19.638 | 2.228 | 20.013 | 25.175 |
| 24.328 | -2.792 | -17.154 | 8.584 | 5.031 | 14.815 | 25.507 | -25.134 | 12.831 | 4.640 | 25.120 | -20.809 | 18.483 | 26.069 |
| 4.337 | -20.845 | -13.981 | 1.981 | -16.299 | 17.727 | -27.345 | 10.051 | 10.061 | -18.451 | 8.579 | 1.353 | 24.788 | 23.113 |
| -11.440 | 9.044 | -3.616 | -1.437 | 20.451 | 25.687 | -1.984 | -26.481 | 10.901 | -26.322 | 29.732 | -1.407 | 21.528 | 28.566 |
| -4.985 | 0.193 | 17.602 | -2.266 | -17.614 | 23.055 | -19.053 | 6.917 | -12.552 | 2.899 | 23.402 | 16.172 | 25.745 | 27.183 |
| -21.398 | -4.468 | -23.229 | -23.500 | 3.590 | 26.295 | 20.587 | 13.103 | -20.276 | -0.611 | -24.515 | -19.675 | -7.585 | 19.360 |
| 0.571 | 13.886 | -5.257 | 6.053 | 5.702 | 18.328 | -14.832 | -2.178 | -28.774 | 8.269 | 29.007 | -11.687 | 23.963 | 8.436 |
| 2.813 | 27.897 | 18.959 | 15.096 | 14.124 | 11.225 | -11.104 | 8.405 | -26.799 | 5.533 | 13.403 | 6.958 | 29.875 | 14.369 |
| 8.396 | -27.212 | 28.859 | 24.063 | 20.707 | 19.562 | -12.959 | 22.109 | 23.302 | -14.352 | 20.116 | 13.372 | 4.818 | 24.341 |
| -23.363 | 0.301 | -20.983 | 6.908 | 8.842 | 7.704 | -3.861 | 11.183 | 14.768 | -3.597 | -27.987 | 9.091 | -30.755 | 27.712 |
| 17.571 | -6.971 | -13.969 | -15.591 | 26.374 | 30.141 | 12.675 | -0.902 | -28.270 | 8.989 | -4.622 | 9.541 | 18.606 | 21.414 |
| -24.621 | 11.483 | -27.983 | -5.582 | -2.192 | 12.794 | -16.269 | 27.971 | 21.393 | -6.801 | 16.852 | -6.754 | 20.641 | 14.537 |
| 23.704 | 18.228 | -6.686 | -26.558 | 25.976 | 22.010 | 1.887 | -21.993 | 19.237 | -6.113 | 27.369 | 6.097 | 16.852 | 26.374 |
| -10.735 | 24.172 | -11.228 | 28.367 | -10.823 | 18.541 | -7.221 | 20.942 | 16.427 | 6.797 | 15.393 | 5.153 | 31.253 | 15.431 |
| -3.682 | -24.003 | 1.555 | 4.561 | 0.015 | 27.368 | 7.923 | 17.766 | 9.780 | 5.120 | -20.786 | 14.281 | 28.407 | 11.436 |
| -9.683 | -9.627 | -5.849 | -8.868 | -5.200 | 24.072 | 11.701 | -22.990 | 4.946 | 0.742 | 4.101 | -21.362 | 23.797 | 18.671 |
| 5.169 | 19.261 | -28.026 | 26.799 | 26.892 | 23.633 | 1.122 | 26.864 | -18.006 | -5.015 | -23.738 | 2.719 | 5.103 | 28.652 |
| 14.420 | 17.479 | 20.787 | 1.427 | -12.835 | 26.134 | 3.278 | -17.241 | -29.280 | 5.953 | 17.265 | -12.876 | 26.650 | 22.159 |
| -18.895 | 31.330 | -3.431 | 5.437 | 23.413 | 22.035 | -17.666 | 24.596 | 22.306 | 22.780 | -24.557 | 3.359 | 22.189 | 27.455 |
| 19.189 | 8.478 | -20.956 | 20.470 | 11.992 | 20.278 | 3.594 | 25.459 | 14.260 | 6.986 | 25.802 | 21.318 | -2.274 | 5.566 |
| 2.505 | 2.560 | -2.200 | -22.497 | 6.310 | 7.471 | 9.317 | 19.681 | 27.829 | -19.394 | -28.272 | 6.906 | 20.793 | 10.457 |
| 18.403 | -30.184 | -13.674 | -20.816 | 17.823 | 19.439 | 12.535 | 0.531 | 28.591 | 19.258 | 21.806 | 12.815 | 5.708 | 15.331 |
| 5.955 | 5.749 | -10.708 | -27.084 | -26.551 | 20.931 | 5.394 | -7.854 | 1.325 | 25.953 | -7.728 | 12.384 | -6.408 | 17.404 |
| -3.619 | 26.512 | -19.594 | -10.423 | 22.573 | 17.175 | 23.092 | -3.992 | 20.859 | -20.747 | -21.457 | -8.835 | 18.365 | 20.206 |
| 3.027 | -0.766 | -2.458 | -19.765 | 17.154 | 26.100 | 9.916 | -6.522 | -16.356 | 16.462 | 18.740 | 3.891 | 28.894 | 22.908 |
| 16.178 | 25.643 | -17.683 | 0.999 | 3.953 | 13.221 | -5.177 | -15.223 | 24.265 | -25.339 | 23.777 | 11.100 | 5.430 | 14.564 |
| 9.272 | 15.739 | -20.269 | -2.886 | -7.553 | 24.081 | 10.998 | 2.288 | 27.262 | -3.823 | 12.997 | -23.212 | 28.277 | 28.212 |
| -11.747 | -1.795 | -28.275 | 12.040 | 12.650 | 29.982 | -20.467 | 18.347 | 17.913 | -9.931 | 28.620 | 10.013 | 29.847 | 19.283 |
| -4.813 | -10.870 | 27.519 | -0.752 | -5.622 | 15.261 | 2.186 | 27.658 | 6.479 | -22.180 | 11.104 | 22.374 | -1.309 | 14.007 |
| 17.881 | -18.485 | -26.894 | 29.571 | -1.211 | 13.495 | 6.762 | -16.698 | 23.080 | -6.847 | 15.137 | 11.547 | 5.637 | 26.611 |
| 10.208 | -20.686 | 2.284 | -18.754 | 8.331 | 14.931 | 2.594 | -8.277 | -9.113 | 0.107 | 28.018 | -14.665 | -5.364 | 11.694 |
| -4.502 | 9.931 | 23.009 | 21.583 | -1.900 | -5.375 | -6.070 | -1.922 | 26.802 | -1.583 | 19.026 | 17.172 | -1.940 | 27.336 |
| -1.006 | -5.221 | -8.189 | -14.891 | -23.508 | 18.755 | 18.859 | 28.909 | 16.165 | 3.815 | 10.312 | -26.308 | 28.654 | 21.154 |
| 14.543 | 1.865 | 28.807 | -12.217 | 20.998 | 27.623 | -18.569 | 1.337 | 18.879 | 2.504 | 6.716 | -6.163 | 11.822 | 23.761 |

| | | | | | | | | | | | | | |
|---|---|---|---|---|---|---|---|---|---|---|---|---|---|
| 5.708 | -27.547 | 5.244 | 21.185 | -25.500 | 20.799 | 12.216 | 14.925 | -3.347 | -0.884 | -10.290 | -7.891 | 27.468 | -2.304 |
| 5.315 | 23.511 | -6.614 | 17.392 | 24.212 | 30.803 | 23.854 | 24.476 | 3.165 | -23.003 | -22.355 | 24.992 | -29.204 | 12.802 |
| -3.805 | -23.614 | 31.617 | 2.792 | 0.729 | 12.204 | 5.977 | -21.503 | 28.699 | 1.469 | -14.630 | 24.026 | 25.870 | 24.433 |
| -0.066 | -24.182 | -20.434 | -8.496 | 26.049 | 22.379 | 2.152 | 16.470 | 21.428 | -3.265 | 27.184 | 11.976 | 26.294 | 14.453 |
| -7.816 | -30.013 | 16.647 | 10.281 | -0.889 | 18.226 | 0.371 | -4.123 | 23.793 | -2.314 | 19.742 | 16.283 | 6.654 | 19.671 |
| -10.407 | -18.758 | -22.710 | 13.570 | -16.729 | 22.899 | -3.028 | 19.218 | -11.181 | 13.063 | -24.011 | 24.687 | 28.758 | 9.207 |
| -6.983 | 26.976 | 17.552 | -20.389 | -3.737 | 19.912 | -16.862 | 30.156 | -18.953 | -4.890 | -5.631 | -14.322 | 12.879 | 25.619 |
| 21.610 | 19.112 | -13.814 | -15.163 | 7.416 | 27.114 | 8.226 | 23.065 | -22.978 | -8.064 | 17.029 | 9.334 | 24.427 | 21.940 |
| 8.025 | 27.735 | -7.304 | -23.850 | 1.413 | -23.749 | 21.928 | 11.508 | 25.092 | -26.449 | 12.546 | 1.469 | -3.675 | 17.919 |
| 20.662 | 25.510 | -29.209 | -0.735 | 17.781 | -3.505 | -18.002 | -16.010 | 19.863 | 14.611 | -9.466 | 20.091 | 9.562 | 23.116 |
| 9.015 | 26.096 | -5.784 | -12.668 | 15.685 | 20.369 | 3.068 | 9.386 | -18.207 | -25.046 | 24.487 | 7.364 | 0.052 | 15.008 |
| 10.285 | -20.839 | 1.416 | 16.035 | 9.285 | -21.801 | -6.753 | 14.460 | 12.759 | -0.940 | 1.578 | -3.423 | 9.939 | 17.258 |
| -22.656 | 28.957 | 13.597 | 14.995 | -24.350 | 14.590 | 24.371 | 1.689 | -0.171 | -6.497 | -4.093 | 19.443 | 6.266 | 20.959 |
| -8.196 | 1.260 | 30.558 | 24.753 | -1.987 | 17.938 | -12.888 | 20.978 | 0.504 | -19.790 | -4.565 | 11.480 | 22.251 | -18.993 |
| 6.869 | -20.047 | -4.172 | -24.845 | 21.722 | 18.476 | -7.726 | -13.523 | 9.618 | -17.745 | -22.069 | 4.622 | 6.711 | 18.284 |
| 17.321 | 16.539 | 17.868 | 7.105 | 12.427 | 28.216 | -8.461 | 16.871 | 1.112 | -23.191 | 21.674 | 5.388 | 20.279 | 2.700 |
| 24.456 | -28.105 | -8.808 | 3.828 | -5.733 | 24.274 | -24.877 | -3.271 | 16.348 | 7.280 | 22.581 | 26.645 | 2.099 | 19.135 |
| -4.256 | -0.572 | -10.722 | 22.248 | 26.315 | 1.158 | -17.482 | 19.815 | 13.693 | -3.190 | -16.610 | 9.229 | 1.392 | 24.762 |
| 13.810 | -23.047 | -19.379 | 12.518 | 16.235 | 21.925 | 9.485 | -16.389 | 6.155 | -21.712 | 28.948 | 26.072 | 3.754 | 20.426 |
| 19.327 | 18.621 | -8.639 | -4.667 | -20.785 | 17.278 | -22.177 | -9.702 | -8.822 | 13.682 | 25.243 | 24.166 | 28.286 | 22.678 |
| -5.210 | -31.479 | 3.721 | 13.864 | -22.702 | 11.838 | 3.947 | 17.444 | -16.100 | 18.462 | 18.461 | 27.077 | -1.030 | 17.616 |
| 1.538 | -19.173 | 18.117 | 18.731 | 23.476 | 20.020 | 0.004 | -12.244 | 28.630 | -3.428 | -27.557 | 26.641 | -1.063 | 27.109 |
| -19.346 | 20.311 | -1.023 | -12.038 | -23.801 | 25.753 | -4.860 | 15.898 | 18.044 | -4.908 | 27.967 | 23.050 | -1.360 | 19.797 |
| -12.731 | 15.187 | -20.259 | -11.144 | 6.222 | 19.130 | 23.303 | -25.120 | -2.793 | -9.882 | 21.089 | -22.594 | -2.498 | 6.329 |
| -23.413 | -1.510 | 13.751 | -30.039 | 14.395 | 20.501 | -3.977 | -29.315 | 24.282 | -5.607 | -12.529 | 13.625 | 23.053 | 17.546 |
| -6.633 | 28.781 | -28.414 | 11.768 | -3.594 | 19.584 | -7.731 | 13.655 | 15.717 | 11.922 | 6.200 | -12.631 | 1.263 | 15.247 |
| 24.951 | -26.261 | -8.222 | 31.227 | -11.586 | 25.028 | -2.426 | -4.245 | 11.524 | 0.418 | 1.712 | 21.358 | 2.528 | 25.087 |
| 16.283 | 19.143 | -27.513 | -2.822 | 24.077 | 13.858 | -2.984 | 29.142 | 4.111 | 7.583 | 20.238 | -15.264 | 1.468 | 18.164 |
| 18.224 | -19.060 | -21.243 | -29.327 | 20.995 | 16.717 | 2.569 | 13.079 | -29.366 | 9.326 | -29.136 | 27.004 | -1.758 | 19.018 |
| 12.399 | 28.190 | 18.825 | 9.611 | -4.136 | 20.914 | -22.378 | 28.130 | 1.501 | -1.675 | 26.099 | 3.020 | -0.493 | 21.499 |
| -24.520 | 27.288 | 7.632 | -22.204 | 0.575 | 23.698 | 12.508 | -0.107 | -7.454 | -0.772 | 12.814 | 10.179 | -4.557 | 23.367 |
| 19.943 | 16.016 | -1.223 | 3.455 | -10.401 | 9.674 | -6.509 | -0.549 | 20.491 | 2.057 | 22.445 | 5.118 | 0.542 | 22.063 |
| -3.298 | -26.445 | -11.670 | -7.076 | -2.592 | 15.574 | 10.740 | -2.300 | 18.563 | -5.560 | 26.651 | 23.904 | 27.641 | 14.904 |
| -19.985 | -20.574 | 5.619 | 2.729 | -21.930 | 17.449 | 6.737 | 3.387 | -3.512 | 5.098 | 15.216 | 20.252 | -23.454 | 27.515 |
| -2.344 | 10.875 | -14.360 | -8.687 | -17.840 | 20.074 | 22.518 | -26.908 | 7.884 | -2.683 | 9.099 | 26.388 | -0.867 | 13.834 |
| -24.047 | -1.183 | -12.726 | 8.039 | 0.614 | 23.031 | 3.011 | -6.764 | 6.591 | -3.297 | 27.177 | -6.375 | 0.966 | 18.381 |

| | | | | | | | | | | | | | |
|---|---|---|---|---|---|---|---|---|---|---|---|---|---|
| -3.984 | 29.209 | -13.744 | -16.702 | -12.804 | 11.149 | 21.303 | -18.938 | -18.450 | -22.962 | 21.245 | 20.609 | 27.023 | 19.403 |
| 23.343 | 25.704 | -15.397 | 30.096 | -11.506 | 10.653 | 14.744 | 8.631 | 9.470 | -8.513 | 11.238 | -2.414 | 0.198 | 23.782 |
| 2.028 | -20.825 | 3.447 | 12.864 | -24.114 | 22.471 | 12.203 | -11.552 | -2.105 | -19.415 | 6.532 | 28.371 | -0.430 | 21.217 |
| 22.664 | -14.491 | -7.168 | 25.800 | 10.502 | 17.026 | -1.986 | 10.309 | 25.062 | 17.647 | 8.174 | 21.069 | 4.292 | 26.452 |
| -5.794 | 12.975 | 28.921 | 13.982 | 12.447 | 18.784 | -10.373 | 19.260 | 2.485 | -9.457 | 20.064 | 8.636 | -1.399 | 13.701 |
| 26.562 | 4.094 | -10.815 | -7.711 | -19.594 | 15.573 | -0.302 | 0.942 | 23.230 | -10.771 | -9.784 | 24.150 | 21.684 | 19.680 |
| -13.586 | -30.208 | 3.688 | -14.597 | 8.249 | 13.490 | -19.648 | -21.747 | -19.971 | 12.768 | 10.887 | 23.043 | 4.999 | 14.938 |
| -15.260 | -24.880 | -5.123 | 4.507 | 6.525 | 8.946 | 1.533 | -1.424 | -21.280 | 19.913 | 10.061 | -15.391 | -23.183 | 23.586 |
| 14.047 | 24.822 | -17.965 | -10.006 | 21.633 | 28.676 | -6.660 | 2.229 | 5.691 | -25.017 | -11.191 | 22.191 | -0.900 | 25.827 |
| -9.677 | 19.168 | 19.654 | 17.210 | 1.140 | 24.071 | -22.662 | 27.953 | 27.452 | 20.686 | -11.967 | -29.621 | -1.022 | -1.470 |
| -19.970 | 20.252 | 15.804 | -24.102 | -21.167 | 21.649 | 8.591 | 16.792 | 26.128 | -3.157 | 5.983 | 22.650 | -3.745 | 25.076 |
| 11.574 | -16.981 | 26.790 | -0.389 | 9.538 | 14.457 | -15.259 | -12.439 | -21.927 | 17.626 | 27.705 | 28.148 | 1.005 | 17.373 |
| -13.883 | -20.353 | 5.059 | 3.521 | 21.898 | 18.511 | 3.955 | 28.325 | 22.010 | 9.017 | -9.035 | 26.470 | 1.851 | -5.469 |
| -3.485 | 1.084 | 25.121 | -3.993 | 16.649 | 7.385 | -24.488 | -6.524 | 1.809 | 21.389 | -27.623 | 28.333 | -14.009 | 22.517 |
| -25.285 | -29.366 | 2.604 | 5.966 | -0.793 | 10.527 | 13.432 | -15.803 | -3.861 | -0.730 | 24.264 | -25.202 | 0.255 | 21.197 |
| -20.342 | 2.122 | 25.459 | -19.972 | -4.551 | 16.397 | -15.475 | -4.589 | 7.982 | 5.470 | -25.885 | -14.639 | -0.383 | 26.025 |
| 2.926 | 17.438 | 6.865 | -21.081 | 20.885 | 21.927 | -16.117 | -23.946 | 19.548 | -25.349 | 17.819 | 16.972 | 20.804 | -18.584 |
| -8.109 | 3.284 | -6.684 | 18.050 | -23.510 | 26.653 | 4.333 | 22.532 | -0.384 | 3.744 | -12.813 | 26.912 | -3.986 | 20.206 |
| -10.206 | -19.719 | 1.859 | -27.971 | -21.467 | 12.403 | 14.405 | 14.751 | -7.120 | 6.606 | -25.207 | -28.624 | 10.254 | 21.719 |
| 21.352 | -3.124 | -24.294 | -9.444 | -25.436 | 18.905 | -2.875 | -0.566 | 18.103 | 17.325 | 15.736 | 16.947 | 11.151 | -9.342 |
| 9.917 | 29.554 | -21.254 | -25.912 | -2.115 | 8.305 | 7.469 | -22.598 | -24.497 | 25.126 | -11.514 | 27.429 | 1.919 | -20.780 |
| -24.159 | 24.847 | 9.864 | -14.084 | 25.159 | 19.451 | -12.716 | 13.304 | -9.405 | 23.957 | 22.291 | 26.525 | 9.594 | 10.619 |
| 24.062 | -28.502 | 15.734 | 1.408 | -24.417 | 18.320 | -4.540 | 19.378 | -20.527 | -2.786 | -8.435 | 23.668 | -11.615 | 5.828 |
| -7.483 | -30.194 | -10.411 | -20.371 | 3.250 | 20.817 | -8.550 | 9.997 | 22.480 | -24.142 | -16.917 | 23.915 | 8.996 | 22.618 |
| 6.768 | -19.135 | -25.417 | -17.220 | 24.106 | 22.725 | -12.119 | -11.965 | -20.217 | -24.569 | 19.380 | 20.024 | -24.874 | -7.470 |
| 23.416 | -27.270 | -13.047 | 30.576 | 10.094 | 22.148 | -14.960 | -26.848 | -23.661 | 2.768 | -19.259 | -16.334 | -2.082 | 12.181 |
| -21.362 | 21.365 | 7.206 | 22.485 | 3.922 | 27.921 | -24.384 | 5.123 | 17.720 | 17.213 | -25.099 | 21.960 | -0.540 | -11.953 |
| -20.807 | -4.937 | 3.112 | 15.367 | 22.968 | 15.919 | 9.753 | -0.863 | -12.105 | 24.645 | 18.162 | 28.684 | -21.374 | 21.820 |
| 26.210 | -21.507 | -26.058 | -16.574 | 20.484 | 11.494 | 5.975 | 21.976 | -12.866 | 22.841 | -13.054 | 26.082 | 10.429 | 17.106 |
| -20.361 | -29.713 | 6.035 | -26.450 | -15.491 | 19.846 | 2.703 | 0.195 | 9.747 | -16.789 | 21.213 | 22.477 | -3.962 | 18.272 |
| -25.682 | 16.176 | -20.366 | 6.089 | -22.733 | 12.499 | 24.188 | -20.717 | -9.891 | -10.086 | 10.321 | -22.591 | 1.420 | 12.365 |
| 9.047 | 5.821 | 14.035 | -8.144 | 11.991 | 11.131 | 17.022 | -9.998 | 9.211 | 20.721 | 11.908 | 20.624 | -0.877 | 11.945 |
| -8.479 | 4.104 | -2.439 | 8.840 | -24.116 | 18.910 | 20.346 | 18.034 | -22.703 | 4.893 | -3.110 | 24.131 | -0.040 | 22.468 |
| -0.103 | 18.725 | -4.885 | -17.995 | 21.579 | 4.698 | 21.149 | 14.200 | 29.203 | 14.827 | -6.163 | 16.610 | 7.124 | 24.310 |
| -24.976 | -25.258 | -5.221 | -13.252 | 15.922 | 16.006 | -9.322 | 12.540 | -18.772 | -15.826 | -16.583 | 18.741 | 0.194 | 19.754 |
| 20.809 | -2.530 | 3.672 | 20.307 | 26.022 | 21.751 | -7.719 | 4.694 | 16.094 | 23.778 | 16.543 | 6.638 | 12.912 | 21.417 |

| | | | | | | | | | | | | | |
|---|---|---|---|---|---|---|---|---|---|---|---|---|---|
| 13.925 | 27.516 | -19.139 | 27.473 | 20.991 | 8.401 | -5.997 | -5.226 | 21.981 | 19.406 | 13.602 | 22.878 | 6.792 | 19.217 |
| -22.761 | -5.001 | -22.102 | -14.772 | -15.518 | 22.728 | 8.565 | -12.224 | 15.756 | 21.647 | -11.197 | 3.463 | -0.338 | 25.246 |
| -26.593 | -1.010 | 11.354 | 23.620 | -17.585 | 15.303 | 24.979 | -19.767 | 25.265 | 4.997 | 18.154 | -15.546 | 20.936 | 3.646 |
| 1.096 | 20.332 | -8.513 | 21.469 | -17.827 | 20.098 | -24.310 | 11.379 | -29.747 | -18.482 | 10.221 | 21.752 | 8.922 | 14.976 |
| 19.455 | 6.736 | -20.117 | 9.760 | -13.429 | 8.889 | -16.918 | 19.596 | 27.631 | -21.628 | -21.102 | 26.786 | 10.472 | 22.823 |
| -7.250 | -27.901 | 4.221 | -18.449 | 12.288 | 20.971 | -0.020 | -8.615 | -15.413 | -1.657 | 14.579 | 25.506 | 10.999 | 10.960 |
| -15.767 | -14.278 | -9.845 | 1.996 | -19.654 | 12.738 | 3.399 | 13.252 | 13.576 | -0.720 | -1.088 | 23.790 | 0.464 | 18.699 |
| -17.369 | -21.236 | -4.317 | -29.912 | 10.515 | 23.996 | 15.062 | 14.992 | -23.690 | 0.679 | 25.212 | 16.275 | 0.805 | 14.040 |
| 5.440 | 1.468 | 18.730 | -2.344 | -5.425 | 10.852 | 17.537 | -0.977 | 10.384 | -6.899 | -3.389 | -12.033 | 2.175 | 20.841 |
| -20.680 | 16.351 | 0.717 | -20.722 | -11.261 | 19.180 | -8.280 | 25.182 | 12.372 | 8.517 | -22.509 | 26.050 | 24.701 | 13.608 |
| -21.985 | 18.468 | -5.874 | 16.383 | 1.458 | 10.338 | -3.043 | 10.511 | -13.150 | 1.775 | 25.972 | 27.596 | 0.059 | 2.992 |
| -19.831 | 8.670 | -11.010 | 21.167 | 8.502 | 0.002 | 21.340 | 15.655 | -17.794 | 25.849 | 20.362 | 20.778 | 6.601 | 25.940 |
| -2.982 | -2.448 | 29.204 | 17.996 | -25.554 | 14.962 | -15.523 | -21.818 | 17.124 | 25.309 | 19.149 | 19.688 | 1.134 | 20.243 |
| -11.458 | 3.711 | 1.538 | 2.564 | -6.741 | 19.952 | -20.318 | 17.951 | -2.773 | 11.859 | 26.765 | 24.948 | 11.143 | 22.465 |
| 26.953 | 16.512 | -18.460 | -26.793 | 16.281 | 7.364 | 17.805 | -15.879 | -22.151 | -27.596 | 23.168 | 28.258 | 4.429 | 21.131 |
| 15.166 | 0.780 | -3.513 | 6.613 | 6.813 | 20.334 | -5.647 | -6.753 | 28.388 | 4.956 | -24.087 | -6.688 | -9.883 | 21.934 |
| -12.719 | -18.576 | -24.653 | 5.855 | 24.565 | 1.777 | 5.694 | 14.650 | 11.667 | -19.990 | 24.122 | 21.337 | 13.709 | 19.658 |
| 28.272 | -5.491 | 5.145 | -14.105 | 22.593 | -1.366 | -4.657 | 13.656 | -25.114 | 27.399 | 11.017 | 24.603 | 6.660 | 19.012 |
| -23.502 | -6.259 | 23.018 | 28.453 | 12.916 | 21.311 | 9.612 | -14.882 | 25.048 | 8.458 | 9.151 | 25.566 | -1.463 | 25.369 |
| -0.082 | -23.867 | 12.392 | -24.871 | 3.680 | 9.617 | 15.316 | 18.725 | 23.155 | -26.553 | 24.457 | 30.704 | 16.971 | 13.550 |
| -8.183 | -3.534 | 15.974 | 21.828 | 19.860 | 25.731 | -12.291 | 8.780 | -16.467 | 20.438 | 16.406 | 27.803 | 2.043 | 20.558 |
| -14.440 | -27.322 | -10.200 | 4.855 | 9.711 | 13.586 | -21.075 | -25.715 | 21.594 | 26.779 | 23.604 | 15.829 | -0.598 | 24.282 |
| 23.639 | 7.059 | -26.215 | -12.321 | -15.188 | 16.358 | -2.087 | 27.171 | 15.537 | -25.549 | -19.591 | 11.063 | 14.608 | 22.583 |
| 13.047 | 8.826 | -8.920 | 20.264 | -22.110 | 17.667 | 16.250 | 7.890 | -18.518 | 21.634 | -8.395 | 25.995 | 8.114 | 21.588 |
| -18.923 | -18.788 | -4.854 | -26.424 | 15.441 | 10.278 | -19.524 | -7.429 | 27.248 | -22.320 | -16.155 | 30.769 | 0.324 | 19.647 |
| 18.651 | -27.910 | -13.976 | 14.926 | -16.213 | 16.842 | 13.977 | 13.193 | -7.015 | 1.062 | 23.000 | 19.169 | 11.204 | 15.486 |
| -7.100 | 10.583 | 18.111 | 9.587 | -24.290 | 20.803 | 19.044 | -6.369 | -19.193 | -21.006 | 12.715 | 20.374 | 27.295 | 18.428 |
| 5.243 | 20.679 | -21.203 | 9.265 | -24.700 | 5.908 | -23.476 | -3.373 | 17.370 | -6.066 | -13.618 | 31.314 | 22.499 | 19.233 |
| 2.422 | 23.391 | -27.553 | -1.327 | 0.494 | 11.019 | 26.547 | 15.052 | -16.979 | -24.656 | -14.426 | 17.103 | -1.250 | 26.283 |
| 7.986 | 11.481 | 1.940 | -4.582 | -11.638 | 23.807 | -18.118 | -24.028 | 10.177 | -4.254 | -23.810 | 20.970 | 14.238 | 16.462 |
| -1.336 | 6.249 | -22.339 | 22.739 | 24.061 | -1.854 | 6.884 | 22.776 | 1.767 | 22.078 | 26.606 | 5.693 | -2.219 | 11.408 |
| 22.800 | 18.534 | -25.065 | 18.690 | -23.506 | 13.662 | -0.225 | -11.689 | -7.809 | -8.438 | 22.245 | 18.755 | -1.145 | 14.087 |
| 13.559 | 3.333 | -0.423 | -13.044 | 25.514 | 21.437 | 24.672 | 9.874 | -9.952 | 16.985 | 4.835 | 26.274 | 27.422 | -13.213 |
| -9.762 | 1.200 | -20.477 | 16.981 | -14.748 | -8.879 | 12.440 | 7.208 | -10.604 | -1.941 | 10.235 | 25.344 | -0.416 | 24.482 |
| -25.942 | 27.101 | 7.024 | 25.011 | 22.753 | 16.096 | -17.651 | 16.852 | 30.452 | -19.322 | -1.960 | 25.081 | -1.649 | 20.682 |
| -16.590 | -22.827 | -21.026 | 11.903 | 11.138 | 16.861 | 22.672 | -18.689 | -28.617 | -7.051 | 19.193 | 29.490 | 11.267 | 21.154 |

| | | | | | | | | | | | | | |
|---|---|---|---|---|---|---|---|---|---|---|---|---|---|
| 28.912 | 28.290 | -0.279 | -3.737 | 20.095 | -19.754 | 23.940 | -21.740 | 8.784 | 29.488 | -7.139 | 27.738 | -1.711 | 23.127 |
| 26.071 | 22.019 | -4.216 | -18.932 | -26.379 | 20.961 | -7.321 | -18.258 | -26.408 | 23.920 | -19.441 | 16.577 | 26.565 | 12.708 |
| 12.959 | -22.407 | 9.834 | 8.533 | -18.409 | 14.315 | 21.286 | -12.146 | -27.310 | 4.998 | 21.820 | 6.537 | 22.603 | 25.470 |
| -3.969 | 12.175 | -6.596 | 6.889 | -2.829 | 23.990 | 5.118 | -20.231 | -3.324 | 25.203 | -9.712 | 24.209 | 10.781 | 19.798 |
| -8.220 | -20.701 | 0.087 | 5.464 | -20.286 | 5.313 | 23.316 | 14.368 | -16.283 | -28.375 | 7.128 | 5.696 | 26.874 | 12.772 |
| 10.681 | 25.619 | -17.015 | -12.396 | 1.253 | 13.419 | 8.619 | 5.861 | 18.898 | 6.230 | 13.311 | 21.152 | 23.743 | 24.142 |
| -24.075 | 17.197 | 14.151 | 28.598 | -26.793 | 18.907 | 22.884 | -23.993 | -11.264 | 7.298 | -16.940 | -24.299 | 25.087 | 20.128 |
| 24.887 | 19.394 | -24.045 | 15.134 | -22.482 | 3.172 | -19.855 | -25.917 | -14.417 | 23.298 | 21.528 | 10.007 | 27.945 | -0.795 |
| 7.421 | 4.430 | -5.959 | -14.138 | -22.017 | 10.399 | -4.675 | 22.215 | 20.644 | 9.460 | 22.644 | 23.704 | 3.877 | 14.156 |
| 23.954 | -23.499 | -10.359 | -8.420 | 1.317 | 19.182 | -4.244 | 3.261 | 29.011 | 4.714 | -7.111 | 16.955 | 28.301 | 6.444 |
| 18.666 | -0.718 | -0.993 | -3.212 | -25.845 | 16.555 | 21.283 | -23.500 | 21.188 | 28.573 | -9.824 | 16.037 | 18.828 | 12.274 |
| -24.938 | -21.130 | 16.914 | 2.411 | 23.279 | 17.956 | 3.355 | 2.940 | -1.846 | -18.664 | -15.168 | 25.688 | 17.583 | 24.414 |
| 15.306 | -21.770 | -5.052 | -20.203 | 21.700 | 14.078 | 19.664 | 19.745 | 12.152 | 19.028 | -19.784 | 24.687 | 23.750 | 12.742 |
| 1.944 | 21.730 | -7.734 | -6.054 | -12.678 | 13.274 | 24.442 | 23.721 | -4.944 | 17.872 | 5.565 | 22.323 | 27.665 | 26.020 |
| -22.537 | -18.782 | 8.330 | 12.195 | -23.299 | 12.998 | 27.303 | 7.186 | -23.272 | -27.830 | 4.183 | 24.946 | 9.646 | 19.249 |
| -0.576 | -22.295 | -21.630 | -2.199 | -28.679 | 21.027 | 25.371 | 4.694 | 22.067 | -28.624 | 14.877 | 13.296 | -18.180 | 21.917 |
| -19.968 | 24.933 | 23.292 | 22.698 | 25.082 | 4.993 | -0.486 | -23.870 | -4.482 | -29.165 | 21.379 | 17.747 | 20.629 | 16.726 |
| 26.151 | 3.143 | 25.191 | 21.479 | 18.651 | 11.794 | 21.948 | 11.343 | 16.238 | 12.579 | -6.051 | 12.552 | -2.433 | 24.981 |
| -2.700 | -17.826 | 1.680 | -22.853 | 23.264 | 6.099 | -16.108 | 19.101 | 2.449 | 20.295 | -10.706 | 28.826 | 24.409 | 14.941 |
| -9.381 | -19.024 | -8.441 | 25.706 | -7.137 | 18.573 | 2.049 | -7.596 | -6.012 | -26.312 | 17.084 | 29.233 | 28.515 | 26.617 |
| -26.917 | 4.278 | -13.736 | 24.124 | -19.647 | 12.113 | -21.606 | 6.241 | 13.511 | 7.015 | 18.388 | -7.832 | -17.403 | 11.333 |
| 21.465 | 23.124 | 19.892 | -27.136 | 1.058 | 23.532 | 15.370 | -21.223 | -21.879 | 25.485 | -10.197 | -8.606 | 14.130 | 23.343 |
| 1.699 | -23.129 | -3.770 | -16.283 | 12.719 | 11.548 | -23.462 | -6.615 | 23.868 | 28.222 | 25.229 | 19.701 | 23.624 | 20.249 |
| -13.167 | 27.267 | -11.758 | 4.044 | -15.996 | 22.120 | 16.223 | -10.972 | 18.482 | -4.385 | 25.874 | 16.059 | 27.609 | 27.708 |
| -14.827 | -24.387 | -20.735 | 20.672 | -27.784 | 10.627 | 26.082 | 24.180 | 9.701 | -22.562 | 12.239 | 22.008 | 26.423 | 22.647 |
| 16.240 | 26.379 | 7.597 | 20.064 | 19.139 | 20.290 | 28.894 | -16.773 | 24.464 | -27.403 | 24.346 | 27.448 | 21.550 | 14.077 |
| -20.927 | 4.840 | 14.938 | -16.943 | -18.697 | 4.357 | 0.936 | 0.203 | 23.188 | -9.475 | 11.636 | -3.668 | 26.845 | 18.449 |
| -11.988 | 2.353 | 12.179 | 14.665 | -28.884 | 12.829 | -13.047 | 17.102 | 20.434 | 21.415 | -6.377 | 10.192 | 23.187 | 17.892 |
| 14.069 | 21.366 | -4.681 | 15.207 | 17.520 | 8.756 | 1.713 | 23.034 | 10.777 | 17.144 | 1.836 | -4.568 | 16.843 | 26.836 |
| -3.409 | 9.084 | 17.378 | -19.878 | 21.216 | 7.519 | 28.107 | -9.228 | -7.062 | -22.780 | 14.707 | 23.032 | 24.685 | 22.587 |
| 25.960 | -23.515 | -3.534 | -19.057 | 16.726 | 3.584 | -25.722 | 14.129 | -20.728 | 24.359 | 8.024 | -7.900 | 27.250 | 16.125 |
| 22.982 | 24.337 | 7.058 | -7.978 | 7.656 | 1.904 | 22.465 | -12.804 | 26.944 | 27.632 | 2.268 | -10.660 | 3.564 | 13.218 |
| 10.131 | -16.939 | 5.439 | 3.174 | 23.974 | 9.360 | 17.982 | -21.519 | -10.888 | 28.499 | -5.118 | 17.672 | 31.313 | 17.241 |
| 29.368 | -11.212 | 3.630 | -1.643 | -8.525 | 8.075 | -4.645 | 12.975 | 11.441 | -30.064 | 9.716 | 20.961 | 27.822 | 14.256 |
| 10.848 | -26.665 | 10.397 | 17.322 | -13.685 | 6.234 | 17.132 | 10.715 | -12.676 | 3.612 | 18.440 | -11.049 | 21.451 | 20.516 |
| 3.178 | 25.503 | -18.675 | -21.509 | -24.551 | 4.392 | -0.952 | -20.692 | 13.063 | -2.402 | 16.022 | 21.584 | 6.237 | 18.272 |

| | | | | | | | | | | | | | |
|---|---|---|---|---|---|---|---|---|---|---|---|---|---|
| -10.096 | 18.855 | -5.697 | 22.351 | -9.742 | 11.595 | -3.026 | 15.897 | 6.265 | -0.044 | 15.117 | 22.494 | 29.948 | 15.514 |
| -5.459 | -24.118 | -6.777 | 8.913 | 27.898 | -8.859 | -22.721 | 5.374 | 28.977 | 8.110 | -24.213 | 22.511 | 28.331 | 5.958 |
| 17.704 | -25.063 | 12.064 | 16.007 | 26.491 | 4.500 | 28.781 | 8.949 | 16.810 | -21.825 | 22.260 | 7.344 | 19.546 | 24.322 |
| 20.094 | 15.997 | 2.781 | 18.354 | -20.633 | 20.457 | 7.530 | -24.542 | 28.048 | 29.555 | 21.687 | 12.568 | 23.013 | 18.497 |
| -22.905 | 13.605 | 6.211 | -14.706 | -14.745 | 13.134 | 10.361 | -2.405 | 25.534 | -4.464 | 26.775 | 7.658 | 17.712 | 26.108 |
| 26.699 | -12.872 | -26.385 | -5.509 | 7.579 | 19.476 | 20.642 | 15.368 | -13.948 | 28.781 | 17.781 | 19.991 | 20.302 | 27.351 |
| 16.700 | 27.022 | -3.092 | -9.108 | 15.793 | 20.847 | -9.950 | 25.279 | 7.299 | 15.056 | -7.214 | 24.307 | 20.072 | 21.975 |
| 21.569 | 18.432 | -11.558 | -25.060 | 22.936 | 14.851 | -12.106 | 2.938 | -19.328 | 19.072 | -11.223 | 1.444 | -0.449 | 23.242 |
| 28.575 | 0.520 | -2.576 | 24.007 | -5.173 | -3.228 | -24.485 | -10.254 | -10.530 | 26.563 | -20.137 | 15.504 | 24.456 | 9.521 |
| 24.127 | -14.300 | 21.276 | 1.909 | -9.751 | -5.153 | -19.756 | 24.688 | 27.259 | 31.053 | 3.800 | 19.179 | 30.361 | 16.256 |
| -17.404 | -21.817 | 14.088 | 16.126 | -27.800 | 8.730 | -22.361 | -1.384 | 17.921 | 27.062 | -18.812 | 19.902 | -1.042 | 6.376 |
| 9.954 | 6.618 | -13.042 | -6.625 | 20.266 | -4.949 | 5.763 | -1.964 | 26.591 | 9.349 | -14.679 | 23.079 | 23.555 | 21.238 |
| -24.213 | 5.051 | -25.056 | 21.068 | 24.514 | 21.745 | 26.030 | -5.198 | 24.953 | 14.416 | 24.805 | 17.231 | 30.801 | 12.512 |
| 0.113 | -25.784 | -6.738 | -12.282 | -26.131 | 18.584 | 27.548 | -22.142 | -15.911 | -2.277 | 20.322 | -3.088 | 22.289 | 28.573 |
| -24.870 | 15.924 | 2.220 | -26.214 | 13.979 | 23.964 | 0.621 | -23.853 | 4.393 | 30.433 | -17.670 | 24.429 | 7.002 | 25.663 |
| 11.525 | -24.791 | -8.780 | -3.992 | -23.345 | 13.903 | 22.783 | 18.881 | -28.074 | -23.030 | 12.001 | -27.080 | 24.814 | 16.076 |
| -4.155 | 10.827 | -1.661 | -21.884 | -20.600 | -0.719 | 9.768 | -6.585 | -11.749 | -29.923 | 9.738 | 16.443 | 27.583 | 14.186 |
| -27.321 | 14.843 | -9.505 | 21.860 | -2.593 | 20.935 | -15.732 | 8.651 | -5.611 | 24.153 | -22.561 | 21.060 | 26.341 | 22.365 |
| 10.337 | 3.052 | -5.275 | 19.498 | -12.441 | 17.816 | -19.211 | 10.028 | 5.399 | -10.888 | -13.791 | -2.123 | 22.225 | 17.351 |
| -7.096 | 17.337 | -10.311 | 9.163 | -22.719 | 23.108 | 20.278 | 18.367 | -8.241 | -7.756 | 23.384 | -0.951 | 15.419 | 27.337 |
| -28.109 | -26.464 | 11.360 | -15.574 | -5.238 | 6.958 | 10.846 | 23.837 | -23.364 | 28.555 | 20.563 | -15.600 | 26.652 | 19.135 |
| -15.372 | -22.411 | 3.792 | 3.508 | -8.621 | 15.181 | 26.245 | 22.343 | 11.825 | -28.947 | 18.597 | 15.582 | -2.895 | 20.952 |
| 1.340 | 25.409 | 7.675 | 26.514 | -28.446 | 20.326 | -6.715 | 11.354 | 26.263 | 21.623 | 18.190 | 24.802 | 4.308 | 23.313 |
| -2.242 | 3.085 | -3.207 | -13.162 | -19.423 | 20.765 | -24.953 | 20.121 | -1.720 | -16.528 | 26.444 | 21.340 | 3.633 | 20.964 |
| 14.506 | -2.143 | 20.743 | 20.143 | 18.082 | 25.405 | 0.863 | 1.098 | -20.820 | -0.221 | 17.417 | 11.213 | 28.191 | 17.710 |
| 11.014 | 14.510 | 3.025 | -9.192 | -25.333 | 15.335 | -12.352 | 7.397 | -3.403 | 18.768 | 3.897 | 26.205 | 26.187 | 22.328 |
| -10.347 | 25.046 | -14.696 | -28.159 | 27.810 | 10.324 | -15.737 | 24.847 | 1.492 | -31.435 | -23.558 | -29.557 | 30.347 | 12.033 |
| -17.915 | 26.212 | -1.241 | -11.184 | -0.396 | 14.213 | -28.174 | 24.284 | 23.493 | -28.977 | 20.077 | 27.429 | 28.454 | 15.610 |
| 17.345 | -20.002 | 1.368 | 25.731 | -16.851 | 17.879 | 29.332 | 21.408 | -18.054 | 17.956 | 14.660 | 27.872 | -1.339 | 8.386 |
| 20.647 | 24.586 | 3.818 | -4.827 | -24.056 | 2.287 | -6.125 | -5.345 | 8.604 | -5.600 | 21.345 | 14.378 | 23.161 | 30.544 |
| -17.102 | 1.450 | -2.347 | -24.555 | -11.665 | 2.766 | -20.469 | 23.299 | 24.125 | -20.356 | 13.348 | 18.558 | -2.724 | 18.656 |
| 29.603 | -12.081 | -20.225 | 16.539 | 10.999 | 13.731 | -22.252 | 18.290 | -10.546 | -30.711 | 25.482 | 8.439 | 26.368 | 15.570 |
| 18.100 | -18.508 | 0.661 | 15.605 | -25.527 | 4.471 | -19.323 | 19.386 | 11.689 | 29.707 | 4.603 | -7.676 | 22.729 | 25.691 |
| 15.351 | 24.241 | -12.745 | -17.193 | -4.138 | 25.251 | 17.253 | 2.443 | -29.551 | 17.022 | 15.487 | 20.700 | 23.523 | -0.463 |
| -26.021 | 2.731 | 5.205 | 26.635 | -22.625 | 1.944 | -18.531 | 17.945 | 28.195 | 26.892 | 1.223 | -14.168 | 24.244 | 17.100 |
| -8.240 | -19.173 | -4.613 | 21.203 | 29.376 | 18.658 | -28.034 | 11.683 | 14.770 | -28.833 | 16.142 | -25.538 | -2.791 | 24.053 |

| | | | | | | | | | | | | | |
|---|---|---|---|---|---|---|---|---|---|---|---|---|---|
| 25.080 | 16.957 | -18.450 | -28.525 | 10.103 | -1.955 | -27.047 | 18.817 | 5.501 | -11.282 | -18.803 | -4.759 | 3.716 | 14.031 |
| 18.625 | -15.281 | -4.058 | -27.336 | 18.860 | 14.285 | 10.518 | 6.181 | 20.258 | -0.989 | -2.039 | 20.454 | -1.892 | 19.875 |
| -11.863 | 23.203 | -6.048 | -8.307 | -30.525 | 0.803 | 19.326 | 17.623 | -6.771 | -0.290 | 20.100 | 28.559 | 29.286 | 22.192 |
| -14.684 | 16.535 | -6.546 | -18.250 | 18.353 | 19.048 | 17.946 | -0.805 | 11.738 | -3.155 | 23.541 | 17.167 | 28.040 | 24.820 |
| 20.615 | 13.745 | -7.087 | -3.948 | -27.261 | -3.234 | 14.802 | 20.968 | 13.782 | -23.347 | 25.596 | 23.255 | 4.536 | 16.438 |
| 17.167 | 5.003 | 23.656 | -16.286 | -18.990 | 25.284 | 26.274 | -24.968 | 17.784 | 7.450 | 24.567 | 25.777 | -2.337 | 11.987 |
| -22.529 | -18.428 | -1.022 | 4.126 | 19.468 | 21.907 | -11.626 | -16.871 | 16.156 | 19.590 | 16.806 | 18.693 | 19.445 | 26.338 |
| -4.373 | -10.617 | -6.224 | 24.580 | -30.066 | -15.883 | 28.821 | 16.579 | -22.961 | 2.532 | 21.437 | 11.489 | 0.889 | 17.330 |
| -7.234 | -5.376 | 18.397 | -27.395 | 27.433 | 20.153 | -26.628 | 10.927 | -4.802 | 28.071 | -25.966 | 12.669 | 11.288 | 15.758 |
| 28.898 | -1.925 | 0.078 | -22.425 | 26.264 | 18.072 | -25.527 | 23.748 | 25.872 | 15.431 | 21.043 | -23.673 | -1.739 | 22.700 |
| 10.954 | 22.650 | 13.205 | -23.436 | 16.163 | 19.271 | 18.754 | 14.370 | 21.968 | -2.136 | 26.252 | 28.344 | 3.841 | 18.044 |
| -5.612 | 16.467 | -10.555 | 19.787 | -25.372 | 11.485 | -23.239 | -24.949 | 3.980 | -6.640 | 19.848 | 22.774 | -1.871 | 22.130 |
| 12.707 | 19.829 | 2.678 | -0.569 | -12.543 | 16.626 | 29.634 | 21.913 | -22.162 | -28.962 | 0.877 | 18.351 | 0.601 | 27.591 |
| 1.243 | -11.371 | -1.744 | 19.211 | -13.782 | 3.475 | -7.625 | 18.262 | 10.473 | 6.643 | 22.828 | 6.705 | 23.102 | 23.181 |
| 21.497 | 2.281 | -24.242 | -19.227 | 15.194 | 24.070 | 21.632 | 23.971 | -14.217 | 20.990 | 8.656 | 24.645 | 29.508 | 24.399 |
| -17.498 | 7.923 | -13.811 | 15.375 | 20.540 | 20.439 | -8.328 | -20.279 | 8.804 | -1.012 | -17.539 | 21.438 | 28.005 | 13.308 |
| -4.740 | -1.004 | 2.087 | -6.068 | 2.836 | 22.785 | 25.824 | 10.828 | -8.509 | -14.566 | -9.248 | -23.147 | 29.127 | 12.141 |
| -29.129 | -16.374 | -14.724 | 25.584 | 21.661 | 16.002 | 23.368 | 11.618 | 12.516 | -15.556 | 22.296 | 14.173 | 1.215 | 19.376 |
| 9.525 | 11.021 | 4.541 | -17.285 | -26.246 | -2.959 | 0.084 | 21.430 | 8.084 | -24.497 | -15.692 | 10.994 | 19.554 | 22.801 |
| 22.950 | 18.735 | 3.516 | -20.024 | 10.073 | 25.538 | -20.644 | -15.756 | -29.301 | 2.938 | -26.927 | 15.501 | -18.149 | -1.581 |
| -27.857 | 18.153 | 17.048 | -14.115 | 1.517 | -27.321 | 23.793 | 26.292 | 20.226 | 5.366 | 21.383 | 4.470 | 17.141 | 7.310 |
| 30.258 | 19.805 | -11.589 | 5.644 | -17.572 | 14.400 | 13.522 | 21.117 | -5.511 | -3.314 | -2.683 | 20.732 | 0.862 | 26.404 |
| 6.128 | -17.086 | -7.355 | -11.202 | -2.834 | 27.743 | 28.770 | 25.978 | 9.390 | 1.426 | 18.389 | 6.292 | 23.616 | 19.041 |
| -9.018 | 4.103 | 5.239 | 23.017 | -21.275 | 13.658 | 25.654 | 7.992 | -7.011 | -29.312 | 24.643 | 16.996 | 6.520 | 16.920 |
| -20.880 | 24.316 | 6.372 | 26.258 | -23.214 | 25.964 | 19.862 | 20.355 | -21.451 | 28.024 | 12.422 | 25.742 | 1.594 | 25.735 |
| -18.885 | 2.491 | 19.926 | 19.718 | 5.572 | 12.811 | 5.858 | -5.890 | 17.699 | 27.383 | 23.665 | -19.500 | -3.556 | 20.941 |
| 16.639 | 22.895 | -8.047 | -8.925 | -15.226 | 22.062 | -29.056 | 26.878 | 7.021 | 0.849 | 19.934 | 14.009 | -4.366 | 30.047 |
| 21.078 | -15.099 | -1.100 | -26.284 | -17.486 | 15.322 | -26.011 | -16.120 | -23.408 | 18.759 | -23.104 | 7.371 | 21.938 | 29.637 |
| -21.595 | -23.488 | -6.512 | 20.979 | -7.763 | 17.110 | -23.201 | 20.497 | 23.156 | -0.729 | 1.089 | 24.383 | 9.455 | 28.935 |
| 5.128 | 1.633 | -13.483 | -21.488 | -22.298 | 20.713 | -23.703 | 25.109 | -8.115 | -18.868 | 11.149 | 23.888 | 14.622 | 15.994 |
| -14.817 | -12.724 | 21.005 | 13.045 | 8.591 | 16.323 | 28.348 | 18.178 | 1.937 | 24.285 | 9.465 | 17.161 | -4.479 | 15.290 |
| 13.532 | 26.241 | 6.072 | -11.515 | -23.291 | 15.038 | -21.102 | 22.256 | -19.070 | 14.524 | 25.748 | 26.893 | -4.299 | 13.841 |
| -29.800 | -8.005 | -9.921 | 5.491 | -29.115 | 15.481 | 25.017 | 25.648 | 28.608 | 16.408 | 8.106 | 26.193 | 3.052 | -0.617 |
| -6.295 | 12.530 | 7.670 | -19.240 | 17.679 | 14.215 | 5.134 | 23.787 | -1.051 | -18.680 | 14.476 | 5.897 | 6.100 | 27.980 |
| 23.812 | 13.132 | -12.067 | -13.601 | -12.752 | 17.756 | 24.553 | 26.857 | 0.777 | 3.049 | -24.316 | 15.023 | 26.002 | 26.993 |
| -12.115 | 14.176 | 0.761 | 11.710 | -16.020 | 19.589 | -22.566 | 8.856 | -16.053 | -18.741 | -6.568 | 14.411 | 18.597 | 25.066 |

| | | | | | | | | | | | | | |
|---|---|---|---|---|---|---|---|---|---|---|---|---|---|
| -15.353 | 0.439 | -14.201 | 3.150 | 28.520 | 9.877 | 25.656 | -25.911 | -13.582 | 4.567 | 23.165 | 26.675 | 19.675 | 17.296 |
| -16.420 | 17.050 | -15.514 | 24.484 | -0.641 | 21.145 | 31.331 | -26.216 | 4.015 | -5.562 | 28.832 | 5.289 | 27.006 | 30.867 |
| 27.447 | -25.182 | 11.892 | 26.617 | -9.452 | 21.746 | 10.663 | 25.363 | -11.816 | -10.726 | -19.956 | 9.679 | 23.241 | 4.751 |
| 19.774 | -2.666 | -2.623 | -24.813 | 14.028 | 18.935 | 6.166 | 27.789 | 20.791 | -26.693 | 26.907 | 15.707 | 4.420 | 11.637 |
| 29.120 | -26.229 | 23.019 | -4.371 | -20.434 | 24.726 | -30.434 | 21.020 | -2.826 | -17.508 | 16.757 | 22.179 | 6.791 | 21.594 |
| -18.647 | 17.479 | 0.267 | 11.428 | 8.953 | 12.020 | -5.396 | 17.959 | -9.560 | 5.482 | 30.099 | 21.438 | 19.308 | 20.526 |
| 30.573 | 8.327 | 6.008 | -0.191 | 19.565 | 22.876 | 27.500 | 19.094 | -6.042 | -9.642 | 27.646 | 9.402 | -9.305 | 29.225 |
| 7.194 | 6.977 | -7.841 | 23.519 | 27.857 | 20.344 | 7.991 | 13.220 | -16.594 | -26.954 | -2.878 | 27.093 | -8.153 | 8.573 |
| -13.766 | 26.168 | -7.320 | -18.541 | 10.520 | 24.085 | -7.854 | -4.925 | 8.561 | 9.808 | -6.191 | 4.749 | 10.194 | 13.719 |
| 3.993 | 8.979 | -3.772 | 8.796 | -21.408 | 2.573 | -28.727 | 26.889 | 24.369 | -18.495 | -10.844 | 1.922 | 9.143 | 30.292 |
| 17.901 | -20.557 | -4.383 | -5.247 | -26.756 | 15.067 | 28.589 | 15.402 | -17.725 | -26.367 | 24.851 | 7.516 | -0.658 | 26.941 |
| -4.668 | 16.067 | -3.692 | 10.073 | -9.941 | 27.444 | 23.109 | -18.481 | 20.222 | 6.936 | -15.544 | 17.525 | 21.764 | 26.530 |
| 25.709 | 12.366 | -18.939 | 13.534 | 29.240 | 7.982 | 20.530 | 11.987 | -3.296 | 3.085 | 10.463 | -20.096 | 11.285 | 27.780 |
| -28.051 | 13.421 | 10.877 | -8.271 | 18.774 | -1.772 | 21.344 | -26.962 | -25.216 | 26.938 | 8.744 | 25.861 | 12.099 | 24.052 |
| -28.950 | 25.660 | -0.100 | 15.282 | -24.959 | 4.657 | 24.342 | 18.847 | 7.824 | -23.018 | 23.211 | 9.828 | 2.148 | 24.922 |
| -16.020 | 4.563 | -12.425 | -23.506 | 24.820 | 20.670 | 5.295 | 25.398 | 29.091 | 25.888 | -16.024 | 21.288 | -2.281 | 19.338 |
| 6.269 | -11.546 | 16.733 | 24.606 | 13.360 | 14.634 | 27.312 | 29.084 | 4.918 | 14.128 | 25.831 | 9.351 | -16.842 | 15.845 |
| 27.411 | -9.650 | -23.471 | 0.138 | -28.907 | 21.872 | -15.815 | 24.343 | 3.584 | 23.278 | 20.341 | 23.324 | 21.707 | 16.638 |
| -30.961 | 20.799 | -22.412 | 22.767 | 17.605 | 18.161 | -12.736 | -27.290 | 27.600 | 2.716 | -27.460 | 7.011 | 2.079 | 10.365 |
| -3.830 | 14.856 | -19.751 | 2.187 | -13.342 | 16.764 | 4.650 | 17.301 | 23.047 | 21.560 | 17.012 | 10.391 | 14.630 | 21.274 |
| -14.592 | 22.185 | -20.806 | 10.760 | -14.714 | 13.618 | 25.566 | -0.589 | 1.866 | -11.563 | -25.662 | -24.401 | 1.659 | 28.970 |
| -2.669 | -24.244 | -3.418 | -17.810 | -30.075 | 19.163 | 29.696 | 7.040 | -12.417 | -7.903 | 28.881 | -25.116 | 4.030 | 29.603 |
| 18.323 | 27.393 | 25.268 | -19.064 | -3.745 | 4.846 | 20.195 | 14.458 | 25.013 | -4.935 | 22.435 | 3.862 | 4.646 | 18.061 |
| 3.377 | -12.492 | -12.163 | -9.702 | 26.488 | 9.005 | -20.327 | 4.978 | -17.024 | 14.916 | 17.485 | 6.901 | 2.478 | 23.574 |
| -4.481 | 16.007 | -25.320 | -13.975 | -4.341 | 28.862 | -24.420 | 18.227 | -21.152 | -19.560 | 6.019 | 20.131 | -0.977 | 15.550 |
| -29.752 | -7.442 | -0.405 | -5.613 | -18.689 | 25.775 | -31.475 | 26.822 | 22.439 | 23.964 | 7.697 | -0.300 | -4.791 | 20.484 |
| -12.238 | 23.975 | -13.246 | 21.403 | 30.207 | 22.278 | 24.606 | 22.082 | 28.093 | 16.075 | 13.529 | 11.638 | 1.180 | 19.882 |
| 28.824 | 17.814 | -17.821 | 18.440 | -8.981 | 17.214 | -25.048 | 17.698 | -11.905 | -26.932 | -13.606 | 25.215 | -20.710 | 28.932 |
| 15.051 | 6.906 | 0.631 | 23.393 | 7.642 | -3.749 | -14.937 | 18.448 | 13.849 | 27.522 | 20.897 | 22.430 | 24.535 | 18.822 |
| 24.618 | 14.753 | 16.194 | -21.643 | 25.286 | 21.213 | -26.134 | -2.245 | 1.678 | -20.665 | -8.268 | 18.230 | 10.764 | 18.484 |
| 17.333 | -3.702 | -21.787 | -24.574 | -6.926 | 15.830 | 5.364 | 16.711 | -12.883 | -24.201 | 11.308 | 24.252 | 3.314 | 24.552 |
| 5.572 | -7.773 | -1.306 | -25.963 | 15.557 | 6.380 | -9.632 | 15.796 | -23.639 | -1.187 | -21.563 | -19.542 | 0.777 | 19.551 |
| 18.742 | -19.850 | -3.084 | -0.001 | -0.137 | 20.107 | -13.141 | 27.216 | -7.104 | -23.642 | 4.533 | 14.436 | -25.524 | 27.842 |
| 12.017 | -24.872 | 22.449 | -11.845 | 18.166 | 19.169 | 9.799 | -18.666 | -16.063 | 3.622 | 9.737 | 6.849 | 12.580 | 23.123 |
| -26.228 | -3.710 | -13.966 | -19.661 | 29.679 | 17.935 | 23.756 | 25.688 | 9.863 | 2.817 | 8.509 | -0.828 | 19.394 | 25.553 |
| 20.610 | -25.664 | -16.264 | 19.918 | -28.295 | 8.236 | 3.301 | 22.950 | 2.427 | -2.463 | 22.097 | 24.625 | 22.586 | 16.546 |

| | | | | | | | | | | | | | |
|---|---|---|---|---|---|---|---|---|---|---|---|---|---|
| -13.838 | 25.141 | 1.944 | -1.200 | 1.748 | 19.691 | -23.051 | 18.212 | 11.338 | -5.635 | 23.431 | 20.241 | 4.233 | 15.050 |
| 10.957 | -1.930 | -4.157 | -2.293 | -0.652 | 17.569 | 26.199 | 29.994 | 19.902 | -18.695 | 19.527 | 17.138 | -3.517 | 14.281 |
| 0.637 | -27.483 | 21.330 | -23.751 | -2.870 | 20.679 | -20.755 | 17.057 | 18.842 | 23.561 | -0.467 | -25.487 | 3.220 | 20.926 |
| -19.409 | 24.328 | -20.305 | 2.948 | -23.800 | 15.201 | -22.738 | 19.515 | -12.888 | 21.922 | -8.484 | 6.310 | -5.108 | 23.832 |
| -4.463 | -5.023 | 6.644 | 10.988 | -26.181 | 24.694 | -25.369 | 18.631 | -9.558 | 24.791 | 18.515 | 25.615 | 0.726 | 21.308 |
| -9.440 | 17.531 | 9.305 | -25.246 | -16.058 | 22.496 | 27.062 | 23.559 | -15.252 | -3.682 | 25.067 | 11.667 | 8.409 | 25.018 |
| 6.179 | -2.981 | -9.459 | 8.523 | -4.850 | 18.458 | -20.272 | 22.805 | 17.078 | 26.118 | 28.975 | 15.945 | 24.501 | 22.071 |
| -8.185 | 18.327 | 8.700 | -10.636 | -18.611 | -8.766 | -27.918 | 15.016 | 12.372 | -21.268 | 30.983 | 22.814 | 0.432 | 15.930 |
| 2.135 | -19.942 | -22.514 | 6.194 | 20.930 | 21.515 | 8.471 | -5.895 | -11.939 | -26.113 | 26.994 | 19.335 | 22.098 | 22.976 |
| -5.896 | 22.981 | 3.095 | -21.673 | 21.918 | 25.679 | -6.158 | -12.947 | 15.514 | 25.487 | 29.635 | -11.492 | 1.354 | 3.140 |
| -3.926 | 19.736 | 5.060 | -9.927 | -22.654 | 13.162 | 22.171 | -21.100 | 2.767 | -20.101 | 17.423 | 24.271 | 5.534 | 14.631 |
| 26.722 | -18.208 | -1.529 | -14.148 | 26.320 | -5.600 | -30.533 | 15.635 | 26.045 | -25.440 | 5.685 | -6.992 | 1.311 | 28.518 |
| -20.992 | -25.366 | 21.037 | -16.441 | 25.475 | -3.770 | 29.802 | 8.302 | 11.175 | 19.290 | 26.789 | -19.148 | 19.521 | 25.911 |
| -7.300 | -2.336 | -5.129 | 12.497 | -9.390 | 20.578 | 11.519 | -28.091 | 17.484 | 17.800 | 15.280 | -8.350 | 22.910 | 22.184 |
| -29.846 | -22.821 | 5.829 | -5.679 | 18.982 | 14.840 | -4.362 | 17.503 | 18.041 | 0.400 | 2.545 | 8.630 | -3.863 | 12.894 |
| 11.485 | -26.374 | -3.786 | -4.683 | -14.278 | 17.837 | -14.023 | -15.833 | 10.477 | 2.489 | 21.617 | 19.894 | -4.583 | 27.775 |
| 14.477 | 27.298 | 4.399 | -7.567 | 27.012 | 15.369 | -29.908 | 30.981 | -5.903 | 0.911 | 6.730 | 12.952 | -4.958 | 16.448 |
| 22.805 | 19.960 | 5.496 | 16.549 | 2.295 | 5.632 | 29.154 | 16.267 | -7.340 | -7.603 | -9.278 | 25.430 | 2.067 | 20.992 |
| 2.800 | -5.961 | -11.587 | -11.081 | -12.200 | -5.640 | -4.298 | 20.514 | 20.492 | 21.339 | 29.114 | 2.560 | -5.459 | 19.220 |
| -24.998 | 23.560 | -19.165 | -3.159 | 6.314 | 13.839 | -23.089 | 18.104 | 21.554 | -26.571 | 6.764 | -25.390 | 10.238 | 18.712 |
| 29.398 | 29.249 | -3.116 | -24.032 | -21.856 | 16.526 | -22.501 | 18.585 | -24.263 | -21.308 | 18.187 | 3.994 | 22.805 | 13.870 |
| 17.902 | 19.624 | -21.934 | 4.064 | -17.517 | 10.400 | -7.283 | 22.636 | -17.372 | -0.091 | -13.911 | -6.483 | 5.126 | 21.614 |
| 1.852 | 28.178 | 24.944 | -9.265 | -15.174 | -9.474 | -21.987 | 16.744 | 6.677 | -0.691 | 28.291 | -7.903 | 12.833 | 20.746 |
| -16.587 | -28.048 | -2.725 | -22.547 | 11.533 | 23.383 | 27.213 | 17.296 | 24.894 | -5.804 | -26.368 | -4.948 | 26.022 | 8.010 |
| 21.245 | 20.223 | 2.431 | -15.369 | 18.009 | 9.723 | -26.472 | 25.344 | -6.493 | 10.163 | 22.586 | 23.645 | -2.813 | 20.406 |
| 4.297 | -6.421 | -21.156 | 2.700 | 28.223 | -29.801 | -19.187 | -16.366 | 12.155 | 17.747 | 9.217 | 25.006 | 26.589 | 23.735 |
| 17.209 | -25.032 | -24.004 | 12.486 | 23.801 | 25.629 | -20.461 | 21.542 | -20.497 | -0.467 | 13.708 | -10.294 | 16.860 | 21.963 |
| 2.257 | -20.759 | 21.036 | -25.002 | 14.537 | 24.886 | 6.791 | 29.610 | 19.612 | -24.117 | 24.975 | 4.276 | 23.094 | 15.616 |
| -9.215 | 2.826 | -5.434 | 10.905 | -2.065 | 21.612 | -12.922 | 25.775 | -7.692 | 26.831 | -5.836 | 20.343 | 8.031 | 23.307 |
| -15.051 | 25.286 | -22.589 | 9.761 | -16.880 | 15.200 | 24.745 | 22.225 | 7.375 | 6.823 | 17.485 | 25.190 | 25.509 | 11.391 |
| -19.389 | 27.671 | 24.169 | -9.165 | 19.526 | 13.324 | 7.460 | 12.844 | -11.405 | -25.105 | -26.668 | 24.576 | -2.759 | 26.115 |
| 13.923 | -4.914 | -25.615 | 3.299 | -20.145 | 7.133 | 22.167 | 17.123 | -23.727 | -8.227 | -4.912 | 20.708 | -6.948 | 12.752 |
| -13.645 | -6.607 | -23.943 | 14.368 | 13.165 | 14.165 | -10.448 | 20.317 | 22.964 | 26.095 | 21.322 | 25.823 | -7.356 | 24.579 |
| 1.531 | 29.006 | -3.905 | -1.400 | -20.969 | -31.101 | 26.545 | 14.582 | -10.582 | -16.719 | -30.680 | 27.065 | 23.586 | 25.609 |
| 24.549 | -28.583 | -23.241 | -7.867 | -28.272 | 19.941 | -26.861 | 23.481 | 27.020 | 11.327 | 25.632 | 2.943 | 26.324 | 23.737 |
| -17.464 | 22.136 | -3.519 | 11.542 | -8.483 | 14.543 | -28.900 | 16.647 | 5.412 | -14.518 | 27.802 | 8.784 | 7.263 | 23.238 |

| | | | | | | | | | | | | | |
|---|---|---|---|---|---|---|---|---|---|---|---|---|---|
| -19.105 | 5.863 | 2.155 | 8.550 | 28.922 | -4.497 | -28.355 | 18.560 | -28.365 | -6.002 | 18.618 | 23.552 | -25.398 | 17.154 |
| 3.526 | -10.591 | -6.600 | -1.458 | -19.210 | 16.701 | 30.180 | 17.734 | 13.368 | -13.554 | 19.372 | 17.376 | 5.173 | 21.225 |
| -18.810 | -21.706 | -9.550 | -22.870 | -9.571 | 18.956 | 24.171 | 22.044 | -12.713 | -0.134 | 15.211 | -28.307 | 18.734 | 15.475 |
| -19.903 | -26.725 | -10.988 | 7.022 | -2.964 | 24.841 | -4.565 | -30.176 | 15.444 | -22.592 | 26.843 | 25.163 | -4.308 | 10.267 |
| -4.473 | 14.495 | -1.145 | 4.442 | 10.120 | 12.229 | -25.927 | 18.841 | -3.907 | 4.814 | -13.766 | 22.298 | 26.813 | 23.161 |
| -8.415 | -9.762 | -7.639 | -4.915 | 7.062 | 30.811 | 29.385 | 14.395 | 17.041 | 15.748 | 31.034 | 17.845 | 4.023 | 27.149 |
| -21.935 | 18.906 | -24.977 | 9.358 | -2.471 | 15.436 | -20.051 | 20.171 | -0.449 | -8.829 | 10.441 | -8.592 | -15.854 | 19.184 |
| -29.704 | 16.596 | 3.476 | -4.388 | -21.244 | 22.187 | 11.209 | 29.037 | 16.013 | -12.194 | 28.477 | 9.338 | 21.741 | 24.962 |
| 0.701 | 19.999 | -2.321 | 10.041 | 10.588 | 19.043 | -9.825 | 23.607 | 25.927 | -16.515 | -21.059 | 27.170 | 23.049 | 25.843 |
| 19.697 | -5.620 | -14.589 | 8.645 | -13.979 | 31.571 | 7.280 | -9.440 | 6.068 | 4.170 | -24.628 | 24.056 | 6.935 | 25.415 |
| -17.096 | 21.170 | -11.561 | 3.056 | 4.110 | -3.921 | 25.235 | 20.674 | -26.968 | 12.204 | 25.938 | 27.333 | 26.072 | 17.127 |
| 17.779 | -23.386 | 4.190 | 14.133 | 22.781 | 30.743 | 16.713 | 27.692 | 5.163 | -15.761 | -15.106 | 27.664 | 28.595 | 19.567 |
| -5.939 | 23.623 | -17.989 | 1.554 | -17.803 | 30.176 | -16.977 | 17.636 | -11.840 | 4.320 | 11.539 | 11.877 | 22.123 | 24.516 |
| -7.903 | 4.349 | -22.987 | 22.175 | 11.327 | 20.053 | 23.406 | 29.551 | 4.299 | 21.738 | -28.775 | 26.490 | -2.799 | 20.815 |
| 28.497 | 24.084 | 6.758 | 14.650 | 17.229 | 7.044 | 28.673 | -4.655 | -13.958 | 1.669 | 26.465 | 23.951 | 22.734 | 24.600 |
| -7.247 | 22.804 | -3.464 | -22.197 | 3.216 | 17.689 | -26.377 | 22.982 | -24.834 | 9.078 | 22.295 | 2.282 | 23.219 | 18.182 |
| 29.399 | 10.361 | -24.710 | -14.007 | -16.106 | 11.157 | -7.331 | 19.473 | 22.503 | -12.344 | 20.551 | -17.441 | 3.256 | 26.541 |
| 21.811 | -5.231 | -1.127 | -2.256 | -21.839 | 21.791 | -19.814 | 13.498 | 11.540 | -20.918 | 16.972 | 22.268 | 25.296 | 25.964 |
| -19.660 | -13.200 | 8.443 | 7.931 | 24.172 | 13.582 | 28.011 | 17.397 | -18.537 | 15.360 | 6.325 | 28.227 | -7.760 | 21.381 |
| -28.542 | 10.875 | -9.501 | -0.288 | -0.872 | 25.632 | 23.763 | -1.012 | 8.759 | 8.076 | 21.681 | 26.102 | 24.391 | 16.782 |
| 19.211 | 12.562 | -12.889 | -5.507 | 5.445 | 20.963 | 7.109 | 20.626 | 7.892 | -21.654 | -12.957 | 20.189 | 14.754 | 25.473 |
| -21.480 | -29.626 | -6.045 | -1.483 | 28.833 | 16.106 | 18.605 | -0.715 | 6.698 | -16.137 | 29.301 | 17.921 | 27.223 | 19.207 |
| -25.428 | 0.655 | -22.304 | 26.651 | 0.631 | 22.319 | -20.344 | 19.376 | 3.617 | -0.717 | -21.914 | 27.711 | 29.885 | 23.886 |
| -27.067 | -19.852 | -11.051 | -6.990 | 15.230 | 17.149 | -8.468 | 29.103 | 23.523 | -1.840 | -3.110 | 24.029 | 28.443 | 21.960 |
| -5.805 | 29.673 | -27.889 | 9.556 | -14.196 | 18.808 | 12.984 | 13.761 | -0.543 | -9.339 | 29.885 | 22.057 | -5.218 | 22.691 |
| -22.953 | -21.638 | -18.266 | 7.095 | -2.993 | 29.856 | 8.246 | 22.360 | -13.243 | -4.039 | 29.034 | 25.267 | 22.774 | 13.421 |
| 5.822 | -13.860 | 9.319 | 8.296 | -0.873 | 20.226 | 1.396 | -27.148 | 13.926 | -2.535 | 13.886 | -29.468 | 0.315 | 25.200 |
| 24.391 | 20.890 | 7.822 | -5.096 | -25.472 | 17.664 | -9.589 | 16.958 | -21.519 | 10.863 | 9.725 | 27.382 | 27.603 | 15.794 |
| 28.251 | 2.930 | -13.334 | 25.795 | -20.523 | 15.492 | -24.716 | 1.411 | -3.678 | 1.811 | 2.875 | -8.481 | 0.803 | 18.184 |
| 17.235 | -18.626 | -26.411 | 5.460 | 21.780 | 13.833 | 12.434 | 21.373 | -6.147 | -7.847 | 23.699 | 21.493 | 27.065 | 23.212 |
| -3.086 | 11.888 | 4.332 | -7.294 | 17.986 | 19.316 | 3.977 | 19.002 | -27.717 | 24.445 | 24.745 | 25.597 | -12.918 | 21.357 |
| -14.629 | -2.168 | -23.464 | -3.091 | -16.643 | 30.600 | 21.210 | 17.905 | 9.315 | 3.341 | 21.965 | 19.995 | 26.672 | 16.419 |
| 20.212 | -4.731 | -10.218 | 4.311 | -26.051 | 15.708 | -23.301 | 18.373 | -8.061 | -3.820 | 21.116 | 11.803 | 29.809 | 24.428 |
| 14.711 | -29.278 | 1.028 | -4.740 | -10.247 | 16.391 | -7.142 | 21.211 | 21.240 | 21.637 | 27.855 | 20.295 | 17.497 | 18.887 |
| 11.145 | 30.723 | -27.535 | 10.132 | -11.745 | -7.685 | 10.346 | 20.634 | 12.044 | 6.114 | 19.250 | -21.706 | 18.988 | 23.781 |
| -4.871 | 4.184 | -3.190 | 8.061 | -26.761 | -0.390 | -28.409 | 22.722 | -12.475 | -6.331 | 13.119 | 18.232 | 16.580 | 22.785 |

| | | | | | | | | | | | | | |
|---|---|---|---|---|---|---|---|---|---|---|---|---|---|
| -3.414 | -24.851 | 29.355 | 7.151 | 7.029 | 25.259 | -15.707 | 13.545 | -13.978 | 12.785 | 0.785 | 28.890 | 3.945 | 21.978 |
| -25.289 | 12.454 | 7.224 | 13.815 | 5.541 | 14.892 | 28.276 | 27.588 | 23.731 | -2.070 | -29.057 | 23.015 | 27.478 | 14.360 |
| 13.432 | 25.239 | -20.450 | 2.186 | 16.452 | 24.039 | -7.679 | 13.229 | 0.630 | -10.505 | 6.758 | -26.475 | 24.973 | -2.877 |
| -6.836 | -15.165 | 13.576 | 11.152 | -27.731 | 17.717 | -24.093 | 21.997 | 18.250 | 21.729 | 14.413 | 26.947 | 3.602 | 7.476 |
| -11.119 | 24.500 | -11.668 | -4.307 | 20.297 | -5.720 | 27.588 | 17.470 | 10.230 | -3.396 | 2.878 | 28.799 | 29.349 | -21.594 |
| 1.896 | 18.367 | 0.788 | 21.387 | -5.808 | 16.236 | 2.344 | 14.788 | -0.600 | -5.534 | -3.868 | 6.869 | 23.321 | -15.364 |
| -23.795 | -8.040 | -27.736 | 4.509 | 16.960 | -1.812 | -13.108 | 28.462 | -1.087 | -7.130 | -29.703 | 30.234 | 1.072 | 16.778 |
| 26.109 | -30.369 | -13.076 | 11.664 | 28.177 | 21.211 | 20.444 | 22.386 | -26.622 | 10.349 | 24.516 | -6.928 | 30.296 | 19.305 |
| 8.498 | 28.499 | -6.827 | 3.873 | 15.520 | 23.096 | 5.465 | 20.765 | 24.957 | 3.535 | 24.050 | 16.564 | 21.959 | 12.007 |
| -19.056 | -22.099 | 29.955 | 20.955 | -19.406 | 18.014 | -23.092 | 15.634 | 14.346 | -16.422 | 30.045 | 7.788 | 18.668 | 18.750 |
| -20.585 | 3.055 | 15.802 | 14.462 | 26.686 | 21.850 | 26.873 | 12.467 | -20.232 | 13.685 | -27.952 | -9.961 | 27.439 | 17.849 |
| -0.387 | 5.445 | 21.851 | 6.746 | 25.093 | 22.322 | 21.526 | 26.759 | -3.808 | 0.175 | 17.237 | 19.196 | 25.661 | 11.586 |
| -16.771 | 1.094 | -18.990 | 20.918 | -28.310 | 19.506 | 5.801 | 15.015 | -6.235 | 1.637 | -2.543 | 9.652 | 24.396 | 15.529 |
| -10.532 | 26.741 | 12.945 | 21.314 | -28.980 | 13.531 | -13.423 | 19.020 | 19.908 | -0.276 | 29.076 | 24.936 | 21.165 | 19.934 |
| 27.756 | 0.131 | -21.067 | -28.264 | -19.770 | 9.563 | -28.026 | 18.113 | -6.959 | 17.692 | 18.114 | 27.098 | 24.802 | -24.448 |
| 17.401 | 6.849 | -3.987 | 15.703 | -6.845 | -6.584 | -26.236 | 11.990 | 21.578 | -11.247 | 21.227 | 17.378 | -3.349 | 9.613 |
| 6.653 | -6.625 | -25.955 | 8.077 | -12.482 | 26.337 | 28.518 | 8.866 | 16.613 | 6.544 | -19.120 | 26.335 | 12.853 | 24.091 |
| -28.264 | 31.025 | 7.424 | 14.315 | 27.909 | 20.725 | -8.444 | 19.543 | -5.701 | 5.505 | 22.618 | 23.293 | -6.380 | 13.835 |
| -10.969 | -13.102 | -6.784 | -9.140 | 5.935 | 16.304 | 21.105 | -2.606 | -26.931 | -17.181 | 6.349 | 20.393 | -4.487 | 6.227 |
| -28.835 | -4.612 | 8.641 | 20.217 | 22.437 | 13.957 | -14.471 | 19.303 | 26.097 | -5.955 | -21.574 | 19.895 | -6.597 | 18.796 |
| -21.508 | 27.851 | -7.761 | 6.593 | -11.515 | 20.921 | 12.094 | -2.380 | 7.723 | -21.074 | -5.745 | 19.274 | -5.975 | 21.846 |
| -22.749 | 2.424 | 20.708 | 3.629 | 9.448 | 18.704 | 4.031 | 15.334 | 1.985 | -4.193 | 30.773 | 6.192 | 6.137 | 20.051 |
| 9.469 | 21.052 | -27.634 | 24.396 | 25.611 | 12.822 | -22.027 | 16.303 | -2.653 | 5.252 | 25.537 | 23.976 | 5.789 | 19.346 |
| 18.062 | -21.468 | 21.118 | 22.932 | 11.838 | 10.904 | 10.717 | 17.921 | 11.753 | -18.198 | -26.689 | 9.486 | 2.566 | 3.373 |
| 5.731 | -15.489 | 7.681 | 16.893 | 1.822 | 5.862 | -19.174 | 11.546 | 6.065 | 6.443 | 15.375 | 21.855 | -4.025 | 7.125 |
| -20.739 | 11.775 | -4.732 | 13.873 | -24.941 | -3.424 | 7.100 | 16.798 | 21.802 | -16.991 | 0.858 | 27.359 | 22.462 | 22.226 |
| 26.763 | -5.970 | -12.532 | -8.487 | -26.187 | 22.741 | 0.811 | 15.478 | 25.363 | 20.842 | 29.918 | 19.615 | 29.046 | 17.327 |
| 7.317 | -16.598 | -15.932 | 5.130 | -11.016 | 16.686 | 24.424 | 17.396 | 0.437 | -9.805 | 21.398 | 18.801 | 2.137 | 20.432 |
| -4.855 | 0.452 | 23.149 | 18.143 | 2.228 | 15.425 | 5.880 | 7.006 | 7.770 | -8.651 | 1.664 | -13.969 | 25.999 | -19.666 |
| 1.471 | 27.234 | -20.878 | -30.222 | 22.872 | 21.876 | -26.010 | 24.357 | 12.925 | -9.402 | -27.560 | 26.582 | -4.332 | 22.363 |
| -0.109 | -26.552 | -21.731 | 18.570 | 1.309 | 15.806 | -1.511 | 17.985 | 19.778 | -22.527 | 28.454 | 28.691 | -4.727 | 20.969 |
| -18.493 | -2.955 | 23.996 | 4.698 | 19.962 | 6.239 | -13.858 | -1.016 | 25.861 | -8.231 | 27.598 | 9.194 | 23.935 | 13.293 |
| 8.642 | -3.795 | 25.932 | 21.612 | -15.437 | -11.391 | 9.744 | 20.070 | 4.512 | -15.468 | 18.932 | 31.532 | 22.078 | 6.500 |
| -9.083 | -18.621 | 20.226 | -5.831 | -13.183 | 16.668 | 6.770 | 16.835 | 24.604 | 0.453 | 2.521 | 19.711 | -2.270 | 16.731 |
| -10.039 | 19.711 | -13.432 | 4.946 | 15.483 | 20.334 | -4.385 | 20.321 | -0.016 | 17.471 | -6.155 | 11.922 | 26.435 | 21.226 |
| -17.943 | -15.891 | 25.470 | 29.995 | 11.781 | 12.608 | 22.697 | -7.703 | 9.598 | -5.427 | 18.341 | 25.704 | 21.022 | 11.483 |

| | | | | | | | | | | | | | |
|---|---|---|---|---|---|---|---|---|---|---|---|---|---|
| -27.693 | 28.329 | -29.007 | 1.052 | -13.755 | 20.189 | 15.592 | 14.331 | -21.970 | 13.174 | -20.477 | 20.643 | 3.204 | 16.009 |
| -13.071 | -16.713 | 5.836 | 0.532 | 26.065 | 25.996 | 27.522 | -1.415 | -22.718 | -23.764 | -7.577 | 15.834 | 1.640 | 19.311 |
| -7.970 | -18.022 | -14.700 | -4.122 | -12.531 | 22.363 | 2.662 | 8.387 | 1.433 | 26.532 | -16.807 | -6.737 | 2.221 | 24.937 |
| -16.227 | -19.062 | -12.239 | 5.292 | 23.828 | 24.531 | 11.307 | -7.024 | 11.665 | -0.390 | 13.907 | 26.424 | 19.967 | 13.583 |
| 25.079 | -24.063 | 12.824 | 9.339 | 14.879 | 28.012 | -3.115 | -5.106 | 10.039 | -13.962 | 4.554 | -8.989 | 22.060 | 26.150 |
| 16.228 | 29.337 | -12.582 | 7.849 | 6.190 | 15.399 | -11.713 | -12.670 | -1.358 | -27.268 | 12.993 | 24.710 | -1.170 | 10.297 |
| -22.226 | 4.131 | 4.538 | 8.401 | 2.172 | 25.287 | 3.823 | 16.539 | 14.223 | 27.205 | 26.022 | 23.294 | 1.594 | 16.836 |
| 23.564 | 7.090 | -9.525 | 2.363 | 0.014 | 19.309 | -3.553 | 13.486 | 11.110 | -23.017 | 14.456 | 25.422 | 10.658 | 18.654 |
| -10.819 | 3.155 | 1.152 | -4.216 | -5.688 | -24.194 | -24.314 | 21.650 | 14.959 | 14.710 | -17.834 | 21.228 | 20.664 | 15.602 |
| -24.053 | -17.439 | -23.105 | -1.639 | 28.142 | 18.063 | -8.362 | 15.899 | 10.522 | -2.597 | 20.273 | 18.233 | 14.832 | 23.184 |
| 9.638 | 25.673 | -17.843 | -1.255 | -7.860 | 15.623 | -21.093 | 20.528 | 12.349 | -10.859 | -13.296 | 11.621 | 16.895 | 20.195 |
| 21.506 | -15.051 | -26.638 | -16.089 | -18.112 | 21.065 | 22.139 | 18.347 | 22.486 | 6.849 | -11.988 | 21.391 | -0.037 | 2.636 |
| -0.500 | 15.903 | -8.677 | 3.948 | 9.391 | 26.611 | 6.214 | 23.860 | 20.587 | 7.373 | 6.693 | 20.057 | 19.923 | 26.156 |
| 21.235 | 30.530 | 31.087 | 6.286 | 26.376 | 15.503 | -26.658 | 19.317 | -9.854 | -2.036 | 25.134 | 30.767 | 26.359 | 15.916 |
| -11.563 | -23.622 | -27.870 | 26.339 | -9.770 | 24.546 | -12.418 | 7.807 | 25.888 | -6.077 | -2.941 | 22.900 | 24.877 | 11.857 |
| 10.274 | 26.834 | -5.744 | 2.651 | 8.378 | 22.305 | 3.450 | 17.597 | 2.956 | -26.967 | 7.222 | 19.464 | -3.149 | 12.269 |
| 8.145 | -0.954 | 8.832 | -17.907 | -15.764 | 23.037 | -2.446 | -2.356 | -17.956 | -18.319 | 29.457 | -25.456 | 0.708 | 25.421 |
| -24.960 | -12.996 | 20.310 | 0.709 | 24.140 | 20.459 | 10.740 | -0.559 | -14.477 | 20.639 | 22.390 | 17.887 | 27.799 | 4.549 |
| 2.144 | -16.071 | -24.028 | 29.445 | 10.166 | 23.923 | 5.672 | 16.784 | 23.148 | 18.644 | -20.650 | 25.907 | 9.158 | 20.898 |
| 24.330 | -8.258 | -9.743 | 6.262 | -9.058 | 19.246 | -3.819 | 19.274 | -9.080 | 9.306 | -7.093 | 25.086 | 8.916 | 14.708 |
| -7.282 | 26.165 | -2.134 | 16.518 | 18.421 | 15.152 | 8.458 | 10.000 | 23.721 | 26.159 | -8.391 | 28.541 | 25.570 | 15.591 |
| 18.649 | -30.230 | -7.767 | 21.049 | 5.549 | 21.200 | 25.014 | -14.887 | -5.845 | 21.809 | -2.824 | 19.674 | 23.155 | 27.065 |
| 17.628 | 26.582 | 18.841 | -2.134 | -5.104 | 12.316 | 26.871 | 21.450 | -9.340 | -24.722 | -25.874 | -12.728 | 2.187 | 19.321 |
| 4.484 | -29.537 | -29.922 | -0.885 | -0.621 | 26.302 | 16.718 | 6.589 | 20.962 | 22.251 | 27.342 | -0.623 | 26.361 | 20.145 |
| 19.260 | 28.737 | -2.182 | 0.342 | 27.187 | 16.325 | -2.817 | -2.706 | -23.109 | -1.242 | 14.475 | 27.681 | 21.293 | 17.751 |
| 27.785 | 18.084 | 6.732 | -0.689 | 22.586 | 18.334 | 9.681 | 23.040 | 0.488 | 25.568 | 24.992 | 19.078 | 21.547 | 18.636 |
| -27.834 | -26.697 | 29.079 | 2.372 | 11.132 | 24.370 | -1.009 | 10.928 | -2.765 | -2.894 | 16.271 | 24.526 | 6.949 | 24.936 |
| 23.805 | -10.625 | 21.128 | 5.154 | 16.141 | -16.436 | -23.233 | 7.539 | -25.021 | 10.836 | -26.547 | 14.569 | -17.579 | 27.785 |
| 1.252 | -17.391 | -12.927 | -1.134 | -8.397 | 11.747 | 10.880 | -15.417 | 13.349 | 4.677 | 0.302 | 24.103 | -14.474 | 18.893 |
| -3.065 | -8.351 | -0.692 | 8.817 | -14.158 | 27.419 | -9.192 | 6.357 | 10.544 | 17.051 | 18.682 | 25.510 | -13.477 | 12.873 |
| -25.966 | 30.409 | -19.865 | 12.190 | 19.009 | 17.869 | -0.472 | -6.705 | -11.160 | -22.636 | -15.939 | -27.538 | -8.287 | 19.629 |
| -1.774 | 27.240 | 9.998 | 1.217 | 7.881 | -8.863 | -9.824 | -1.960 | 11.543 | -2.328 | 24.792 | 26.392 | 2.412 | 9.073 |
| -25.757 | 14.235 | -5.305 | -29.150 | 0.043 | 22.992 | 13.020 | 9.825 | 18.107 | 2.936 | -12.615 | 7.628 | -7.150 | 7.261 |
| 9.178 | -2.140 | -19.204 | -4.460 | -1.783 | 21.690 | 26.044 | 16.745 | 15.453 | -5.893 | -0.171 | 28.815 | 14.212 | 28.718 |
| -11.871 | -30.367 | 18.287 | 28.002 | 21.154 | 14.637 | 8.390 | 4.535 | 4.356 | 21.102 | 28.151 | 18.118 | 3.208 | -1.729 |
| -19.419 | -21.506 | 31.806 | -6.718 | -24.387 | 17.206 | 1.360 | -6.649 | 16.298 | -10.736 | 19.955 | 22.085 | 13.455 | 10.223 |

| | | | | | | | | | | | | | |
|---|---|---|---|---|---|---|---|---|---|---|---|---|---|
| 3.605 | 20.710 | 15.715 | 13.837 | -21.650 | 11.309 | -1.524 | 11.481 | 21.466 | 11.858 | 22.456 | -4.906 | -16.893 | 15.454 |
| -16.000 | 19.247 | -13.399 | 2.043 | -12.318 | 8.999 | 26.415 | -13.019 | 19.337 | -7.472 | 18.733 | 19.888 | -21.945 | 22.235 |
| 5.683 | -22.158 | 0.939 | 7.378 | 3.880 | 5.159 | 21.183 | 17.883 | 7.563 | -3.388 | 23.361 | 10.916 | -25.229 | 12.490 |
| 3.138 | 22.119 | -19.328 | 3.334 | -15.736 | 14.420 | -10.627 | 11.407 | 11.893 | 26.731 | -26.544 | -3.435 | -1.853 | 25.596 |
| -27.144 | 0.727 | 2.624 | 0.838 | -20.595 | 24.493 | 25.207 | -0.370 | -0.713 | -18.584 | 29.146 | -30.463 | 18.903 | 16.449 |
| 19.871 | -29.067 | -27.404 | 4.536 | -14.525 | 19.247 | 19.539 | 2.605 | -1.165 | -23.397 | 25.828 | 16.186 | 20.410 | 23.339 |
| 0.721 | -5.191 | -8.515 | -0.707 | 11.886 | 5.127 | 6.432 | 22.033 | 5.794 | -8.573 | -21.883 | 13.743 | 14.342 | 14.840 |
| 6.992 | 8.543 | 14.660 | 22.746 | -17.447 | 15.378 | 2.180 | 23.040 | -1.855 | -12.605 | 18.011 | 22.355 | 18.340 | 10.525 |
| 27.280 | 3.319 | -19.797 | 7.472 | 10.203 | 18.520 | 3.896 | -13.549 | 1.784 | -25.707 | 20.847 | 25.372 | -0.303 | 18.462 |
| -26.524 | 27.578 | -24.905 | 7.085 | 23.542 | 17.582 | 0.485 | 12.782 | -6.363 | 25.885 | 9.533 | 21.310 | 7.311 | 28.734 |
| 7.825 | -21.723 | -9.933 | 3.979 | -8.024 | 25.868 | 2.585 | -9.244 | -19.354 | -7.614 | 26.297 | 30.160 | 16.395 | 26.925 |
| -9.991 | 9.281 | -11.822 | -3.615 | 27.992 | 24.678 | 13.593 | 17.937 | 7.173 | -21.330 | -23.281 | 27.336 | 4.420 | 21.979 |
| 22.108 | 11.333 | -22.927 | -12.311 | 17.870 | 6.680 | -25.486 | -16.035 | -2.327 | 16.826 | -21.859 | 6.847 | -13.341 | 29.460 |
| -11.248 | -3.407 | 21.531 | 1.158 | -15.606 | 1.339 | -2.438 | 25.528 | -11.067 | -0.505 | 17.110 | -6.236 | 20.658 | 25.054 |
| -3.004 | -23.645 | 20.520 | -19.315 | 19.992 | 5.042 | 11.043 | 22.179 | 0.720 | 18.304 | 7.239 | 29.168 | 22.707 | 27.422 |
| 9.308 | -26.863 | 6.603 | -0.902 | 18.651 | 18.645 | 18.438 | -24.188 | -0.868 | -1.807 | 21.578 | 19.473 | 15.386 | 17.931 |
| -14.089 | -27.170 | -20.163 | 6.831 | 8.175 | 6.457 | 26.196 | 6.918 | 23.222 | -17.486 | 29.730 | 10.807 | 3.242 | 15.133 |
| -20.721 | 19.829 | -13.779 | 23.566 | -22.397 | 20.681 | 12.805 | -24.798 | 1.572 | -3.912 | 24.403 | 22.834 | 24.476 | 14.360 |
| -5.947 | -18.526 | 30.249 | 1.253 | -8.231 | 5.774 | -10.048 | -16.124 | 16.871 | 22.590 | 28.455 | 25.972 | 22.973 | 7.781 |
| -22.886 | -29.870 | -21.891 | 26.415 | -18.935 | -23.622 | 7.004 | -23.865 | 20.095 | 4.762 | 17.472 | 29.466 | 16.522 | 24.897 |
| 9.914 | 29.329 | -9.043 | -1.945 | -21.329 | -19.014 | -17.968 | -24.232 | 22.183 | 8.809 | 12.090 | -26.441 | 13.989 | 21.233 |
| 24.063 | -27.543 | 17.498 | 2.061 | 25.198 | 16.039 | -4.842 | 14.921 | 18.333 | -8.904 | -0.213 | 18.187 | 20.718 | 15.720 |
| 0.293 | 6.592 | -20.291 | 6.928 | 22.136 | 11.567 | -10.442 | -19.130 | -5.338 | 14.233 | 22.552 | -7.139 | 25.019 | 1.503 |
| -10.305 | 26.006 | -6.826 | 6.542 | 6.646 | 9.276 | 4.766 | 26.457 | -1.245 | 21.702 | 23.012 | 9.513 | 25.818 | 17.074 |
| -24.465 | 22.091 | 2.806 | 16.667 | -4.401 | 22.401 | 10.563 | 25.636 | -14.752 | 11.990 | 20.156 | 15.979 | -26.358 | 26.739 |
| 4.232 | 9.818 | 11.647 | 15.057 | 13.364 | 24.201 | 25.647 | 3.868 | -21.100 | -1.770 | 16.271 | 4.588 | 11.880 | 16.579 |
| -6.665 | -22.521 | -28.652 | -10.746 | -0.901 | 13.218 | -0.244 | 23.913 | -3.805 | -3.541 | 25.152 | 13.222 | -17.169 | 27.256 |
| -17.267 | 14.198 | -0.395 | 4.226 | -15.341 | 1.635 | 1.219 | -0.784 | -23.352 | 19.274 | 10.298 | 3.136 | 26.271 | 18.847 |
| 21.761 | -28.140 | 4.208 | -21.934 | 8.951 | 5.569 | -20.598 | -23.668 | 10.874 | -0.731 | -25.510 | 9.771 | 23.242 | 18.150 |
| 14.301 | -23.042 | 19.607 | 1.043 | 17.285 | 4.832 | 13.474 | 20.120 | 11.660 | -11.922 | -22.994 | 24.925 | 23.833 | 20.774 |
| -25.403 | -25.210 | 9.135 | -0.320 | -13.655 | 7.799 | 11.551 | 22.746 | -2.950 | -19.035 | 21.530 | 5.288 | 24.682 | 26.151 |
| -12.081 | -1.066 | 15.768 | -19.595 | 10.191 | 15.982 | -18.307 | -0.193 | 24.228 | -5.630 | 22.156 | 26.644 | 22.721 | 24.094 |
| -16.396 | -17.830 | 14.002 | 2.301 | -10.792 | -2.611 | -2.250 | 0.650 | 8.916 | -22.728 | 26.535 | 21.954 | -1.863 | 25.597 |
| -19.702 | 24.663 | 9.639 | -18.376 | 12.053 | 1.991 | -5.070 | 22.179 | 17.374 | 25.837 | -19.076 | -10.733 | 16.432 | 19.945 |
| 12.789 | 23.233 | -1.232 | 1.578 | 19.276 | 16.393 | -15.290 | -1.285 | 1.214 | -18.580 | 26.925 | 17.583 | 15.428 | 12.802 |
| -14.528 | -22.493 | -7.153 | -22.214 | 20.522 | 9.143 | 21.272 | 24.801 | 5.765 | -2.016 | 24.952 | 6.158 | 18.421 | 28.317 |

| | | | | | | | | | | | | | |
|---|---|---|---|---|---|---|---|---|---|---|---|---|---|
| -3.893 | -30.165 | -14.733 | 7.168 | 10.638 | 7.366 | -10.336 | -0.703 | -1.496 | 28.263 | -17.771 | 23.973 | 27.377 | 23.535 |
| -13.171 | 26.313 | -8.518 | -0.969 | -25.130 | 10.688 | -10.886 | 4.010 | 4.101 | -11.825 | -18.373 | 10.186 | -26.378 | 17.129 |
| -19.324 | 15.660 | -12.545 | 3.492 | -23.406 | 19.820 | -1.295 | -19.220 | 1.415 | -16.212 | -14.730 | 10.898 | 21.145 | 25.757 |
| -9.874 | 17.825 | 10.369 | 8.968 | 11.819 | 17.295 | -15.833 | 8.227 | -27.254 | 2.809 | 22.724 | -24.985 | 20.455 | 17.772 |
| -19.953 | -27.842 | -2.876 | 2.582 | 6.143 | 25.010 | 13.323 | 20.187 | 2.605 | -2.852 | -8.156 | 20.550 | -4.059 | 19.730 |
| 15.798 | -4.107 | 26.828 | 4.778 | -21.118 | 18.971 | -9.006 | -21.860 | 2.041 | -15.445 | 7.115 | 8.113 | -26.288 | 12.609 |
| 17.243 | 28.592 | 18.069 | -2.359 | 27.140 | 0.236 | -11.557 | 21.316 | -20.433 | 17.338 | -23.105 | 12.820 | 24.495 | 24.565 |
| 1.879 | -0.715 | 8.891 | 7.347 | -8.451 | 17.761 | 6.899 | 25.677 | -0.420 | -28.637 | 23.564 | 21.391 | 21.177 | 26.765 |
| 24.196 | 11.090 | 4.164 | 5.152 | 24.013 | 16.015 | -23.890 | -10.273 | 19.404 | -8.775 | 16.744 | 24.731 | 11.196 | 21.025 |
| 24.932 | -20.826 | 0.152 | 16.140 | -26.375 | 10.416 | 2.086 | 7.761 | -2.939 | 26.564 | 15.340 | 26.359 | 27.610 | 26.200 |
| -7.523 | 2.077 | -29.089 | 0.939 | 12.288 | 11.247 | -3.536 | 24.559 | 4.625 | -2.219 | 19.479 | 27.409 | -15.798 | 30.478 |
| -24.137 | -17.390 | 3.564 | 4.141 | -19.050 | 24.513 | -2.842 | -20.889 | 1.277 | -23.543 | 27.826 | 27.938 | 25.330 | 10.311 |
| 26.790 | -29.365 | -16.977 | -7.187 | -15.887 | 0.616 | 12.490 | -14.018 | 9.105 | -5.373 | 21.518 | 23.827 | 12.353 | 25.422 |
| 5.314 | -23.298 | 7.617 | -6.657 | -4.547 | 19.327 | 2.427 | -24.319 | -3.482 | 12.093 | -7.911 | 16.563 | 22.611 | 1.514 |
| -19.204 | -1.744 | -24.852 | -0.172 | -12.731 | 17.016 | 24.414 | 17.645 | 0.608 | -6.741 | 16.103 | 21.021 | 23.655 | 24.012 |
| -23.161 | 29.307 | 20.845 | -1.799 | 25.094 | 19.869 | 7.998 | 5.300 | -12.355 | -13.504 | 18.551 | -6.860 | -9.875 | 16.567 |
| -3.835 | 3.309 | 6.986 | -9.152 | 15.930 | 22.199 | -7.277 | 27.139 | 15.600 | -24.714 | -11.290 | 3.284 | 27.154 | 20.725 |
| -3.234 | -4.286 | -25.731 | -9.916 | 20.202 | 21.603 | 20.878 | 25.787 | -13.503 | -26.402 | 23.592 | 15.982 | 13.638 | 22.573 |
| -18.809 | 0.176 | 12.857 | 5.230 | 12.654 | 12.805 | -9.383 | -22.944 | 4.492 | 8.575 | 3.150 | 21.640 | 20.233 | 19.503 |
| 18.912 | -25.752 | -5.058 | 15.194 | -22.516 | 17.476 | 19.528 | 27.466 | 14.286 | -26.056 | 22.728 | 19.952 | 24.141 | 22.129 |
| 4.069 | 27.197 | -1.734 | -0.565 | 23.183 | 4.815 | -15.065 | 13.619 | 17.670 | 17.705 | 24.900 | 3.979 | 25.877 | 24.432 |
| 14.031 | -5.703 | -11.434 | -1.589 | 24.170 | 4.474 | -1.226 | 19.016 | -3.584 | -12.759 | 27.106 | 7.559 | -18.548 | 31.490 |
| -11.726 | -27.282 | 26.104 | 6.367 | 8.415 | 12.147 | -10.656 | -9.869 | 3.823 | 22.989 | 20.611 | 13.264 | 25.210 | 23.691 |
| -20.303 | 7.233 | 13.413 | 22.553 | -20.817 | 15.497 | 10.060 | -8.862 | 4.144 | 27.241 | 19.714 | 24.663 | -26.004 | 29.690 |
| 6.311 | -16.401 | -25.045 | 8.365 | -22.852 | 3.493 | 20.833 | -23.228 | 20.362 | 26.076 | -18.233 | 22.782 | 27.984 | 20.996 |
| -2.287 | -17.117 | 5.253 | -1.629 | 8.808 | 18.145 | -4.228 | 12.714 | 4.385 | 28.837 | 23.733 | 25.300 | 21.387 | 17.704 |
| 8.521 | 19.416 | -7.996 | -3.136 | 2.714 | 25.386 | 1.879 | 28.386 | 6.038 | 24.194 | -15.146 | 10.531 | 12.285 | 28.918 |
| 25.353 | 8.087 | -12.943 | 13.274 | -13.215 | 14.855 | -2.499 | -5.473 | 0.115 | -24.786 | 2.054 | 14.631 | 26.492 | 20.330 |
| 9.831 | 5.669 | -0.461 | 4.524 | 5.346 | 16.155 | -20.048 | 26.702 | -21.859 | 19.363 | 11.753 | 27.248 | 14.893 | 30.870 |
| 11.782 | 13.069 | -22.445 | 2.681 | 21.650 | 10.629 | 16.180 | 15.873 | 22.132 | -12.389 | 28.088 | 18.032 | 28.971 | 22.586 |
| -10.509 | -1.915 | -25.617 | -17.571 | 22.486 | 7.545 | 9.021 | -0.583 | 22.582 | -28.998 | 24.713 | 16.517 | 27.217 | 18.277 |
| 7.129 | 20.570 | -4.352 | -19.032 | -18.374 | 25.143 | -8.022 | 8.303 | 22.409 | 30.790 | 26.849 | 3.120 | 18.056 | 21.839 |
| 4.607 | -25.356 | 8.741 | 6.067 | 20.290 | 21.288 | -6.810 | 8.993 | 23.829 | 21.945 | 0.348 | 11.850 | 15.636 | 25.796 |
| -25.500 | -22.330 | -23.421 | 12.356 | -10.588 | 14.996 | -14.871 | -27.443 | 26.016 | 26.437 | 26.022 | 19.802 | -26.009 | 24.829 |
| 20.983 | -12.287 | 29.670 | 14.351 | 23.015 | 17.255 | -17.382 | 24.673 | 12.510 | -1.887 | -18.400 | 7.425 | 27.756 | 18.949 |
| 20.202 | 1.021 | 5.948 | -19.056 | 19.171 | 24.236 | -3.108 | -11.524 | 20.485 | -23.204 | 5.063 | 24.141 | 23.924 | 15.776 |

| | | | | | | | | | | | | | |
|---|---|---|---|---|---|---|---|---|---|---|---|---|---|
| 5.443 | 27.964 | -3.547 | 0.721 | 9.425 | 5.218 | -8.028 | 25.304 | 4.920 | -22.369 | 20.904 | -4.234 | 16.028 | 31.677 |
| 17.988 | -21.759 | -8.919 | -0.932 | -6.329 | 24.746 | -0.002 | 17.427 | 5.749 | 15.603 | 17.291 | 22.015 | 22.897 | 30.228 |
| -12.703 | 29.049 | -27.283 | 6.997 | 19.397 | 20.138 | -6.106 | 26.644 | 5.281 | 7.151 | -14.370 | 1.207 | 18.311 | 12.359 |
| 22.249 | 19.997 | -17.674 | 2.319 | -9.971 | 13.221 | 17.411 | 24.049 | 5.454 | -1.033 | -16.110 | -13.977 | 26.089 | 20.895 |
| -6.794 | 18.746 | -11.261 | -19.251 | 16.948 | 22.099 | 8.152 | -10.167 | 1.815 | 3.994 | 11.541 | 17.024 | 10.584 | 22.802 |
| 18.577 | 12.542 | 8.975 | 4.882 | -10.531 | 5.975 | -10.502 | 27.448 | -7.271 | -27.816 | 18.347 | 5.360 | 20.787 | 19.042 |
| 9.862 | 23.187 | -3.936 | -22.879 | 24.299 | -1.315 | -1.806 | -20.937 | -18.414 | 27.356 | 19.927 | 15.257 | 13.492 | 16.018 |
| 5.980 | -12.772 | 14.923 | -0.368 | -16.763 | 20.884 | -0.588 | 28.596 | 3.638 | 23.856 | 5.484 | 13.608 | 29.253 | 27.272 |
| 16.792 | -29.452 | 2.448 | 10.594 | -1.002 | 13.695 | -1.161 | -23.799 | 23.475 | 13.647 | -24.384 | 9.340 | 22.318 | 23.602 |
| 14.536 | -20.075 | -10.262 | 1.075 | 12.800 | 6.515 | -23.834 | 22.687 | -11.966 | -26.486 | 14.700 | 4.714 | 19.685 | 20.165 |
| -21.467 | 3.364 | -19.358 | 13.292 | 3.961 | 18.564 | 20.871 | 22.080 | 21.106 | -5.281 | 23.469 | 24.357 | 13.097 | 14.437 |
| 11.646 | -14.012 | -4.413 | 3.438 | 12.042 | 17.257 | 26.398 | -6.803 | 26.932 | -13.884 | 21.615 | 8.542 | 28.498 | 28.578 |
| -24.693 | -26.043 | 22.944 | -17.871 | 5.397 | 22.541 | -2.262 | -27.083 | 18.864 | -2.198 | 13.165 | 20.625 | 20.354 | 21.398 |
| 22.473 | 26.594 | -1.659 | 25.134 | -12.405 | 7.512 | 2.090 | -29.344 | 12.136 | -5.709 | 4.385 | 2.626 | -1.263 | 29.080 |
| 0.008 | 17.015 | -12.898 | 23.013 | -20.134 | 15.607 | -8.674 | 27.854 | -24.449 | -24.256 | 1.290 | -6.215 | 23.601 | 23.631 |
| 9.042 | -14.905 | -27.470 | 17.214 | -20.782 | 17.926 | -21.108 | 19.799 | 31.005 | -30.631 | -14.541 | 16.314 | 7.284 | 17.708 |
| -16.417 | 20.849 | 0.608 | -14.902 | 5.839 | 10.655 | -16.440 | -10.419 | 28.698 | -28.154 | -23.685 | 0.942 | -25.761 | 25.711 |
| 1.248 | 10.868 | 1.518 | -1.515 | -6.832 | 6.913 | -0.206 | -28.465 | -31.459 | 25.071 | -4.622 | 5.794 | -24.706 | 12.835 |
| 5.021 | -3.311 | -10.117 | 10.377 | 10.656 | 23.895 | -5.205 | -24.022 | -5.667 | 14.310 | 16.491 | 4.086 | 20.385 | 30.392 |
| -19.425 | 15.120 | -16.844 | -26.369 | 13.128 | 12.021 | 23.649 | 22.415 | -23.335 | -27.660 | 12.310 | 10.227 | 14.628 | 20.111 |
| 25.511 | -2.012 | -8.495 | -7.606 | -3.958 | 21.467 | 25.572 | 24.422 | -5.384 | 20.868 | 11.459 | 7.130 | 29.175 | 27.754 |
| 5.971 | -24.531 | 18.405 | 4.596 | -12.600 | 20.270 | -23.731 | 27.130 | 3.604 | -1.599 | -11.632 | -2.099 | 19.266 | 22.486 |
| -1.599 | 10.242 | 0.995 | 5.237 | 1.222 | -5.619 | 25.229 | 29.030 | 28.452 | -24.554 | -3.610 | -11.251 | -19.858 | 25.803 |
| 22.294 | 21.312 | 12.532 | 0.207 | -19.952 | 17.429 | -18.678 | 16.423 | 25.750 | -16.080 | -10.730 | 9.440 | -26.199 | 16.573 |
| -21.917 | -14.916 | -19.908 | -9.528 | -6.015 | 17.115 | 0.439 | 30.487 | 23.743 | 22.240 | -12.425 | 23.100 | 28.045 | 19.317 |
| -13.588 | 2.206 | -0.796 | 1.547 | 9.811 | 14.518 | 17.077 | -29.311 | 22.547 | 12.848 | 14.401 | 24.813 | 24.758 | 28.423 |
| -17.537 | 16.571 | 0.253 | 20.192 | 10.839 | 9.449 | -3.116 | 19.030 | 29.343 | 1.697 | -15.146 | 26.449 | -25.638 | 18.976 |
| -19.703 | 14.791 | 14.459 | 7.413 | -9.672 | 19.293 | -21.372 | 22.315 | 19.635 | 5.499 | 23.864 | -6.586 | 19.729 | 26.504 |
| 21.685 | 0.866 | 9.197 | 3.891 | -10.427 | 26.750 | 10.013 | -2.513 | 29.003 | 25.349 | -5.670 | 19.956 | 27.590 | 12.343 |
| -12.136 | -27.347 | -1.529 | 5.546 | 20.918 | 5.294 | -0.673 | 29.323 | 29.456 | -22.260 | -4.876 | 4.391 | 21.560 | 30.338 |
| -11.021 | 27.750 | -6.524 | 1.156 | -0.671 | 16.581 | 20.645 | 25.827 | 26.525 | 17.259 | 15.132 | -7.423 | 30.222 | 21.576 |
| 16.740 | 23.446 | -20.062 | -19.573 | 22.949 | 15.245 | -10.800 | -5.692 | 24.866 | 4.982 | -19.235 | 11.973 | 26.691 | 20.421 |
| -15.426 | 25.983 | -13.564 | -4.453 | 23.845 | 7.753 | -14.566 | -17.841 | -5.303 | -28.363 | -10.776 | 1.045 | 25.514 | 24.884 |
| 9.077 | 16.373 | -23.367 | -3.478 | 12.024 | 13.349 | -21.149 | 30.707 | 17.669 | -19.238 | 23.359 | 5.758 | 29.033 | 26.968 |
| -0.414 | -24.053 | 3.227 | 6.059 | 10.485 | 11.676 | -16.928 | -27.789 | 26.895 | 22.698 | -1.136 | 22.196 | 19.434 | 10.408 |
| 23.848 | -16.625 | -8.191 | 3.364 | -17.613 | 9.178 | 18.700 | 24.821 | 24.272 | -31.301 | 24.242 | -3.774 | 21.585 | 29.494 |

| | | | | | | | | | | | | | |
|---|---|---|---|---|---|---|---|---|---|---|---|---|---|
| 14.052 | -22.365 | 2.165 | 0.311 | -10.305 | 23.705 | 23.605 | 27.592 | 9.065 | -26.123 | 18.231 | 1.957 | 26.069 | 30.342 |
| 1.283 | -11.147 | 1.328 | -4.264 | 16.066 | 16.702 | -0.850 | 23.458 | 19.981 | 26.857 | -3.985 | 15.393 | -11.048 | 1.788 |
| 19.014 | 25.364 | -11.850 | -3.885 | -23.662 | 21.339 | 26.158 | 31.018 | 26.046 | -26.996 | 8.436 | -15.993 | 22.502 | 14.757 |
| -7.756 | -5.486 | 3.654 | -4.440 | 7.082 | 14.238 | -9.128 | 7.351 | 26.671 | 8.750 | 1.615 | -4.404 | 25.168 | 19.490 |
| 4.297 | -12.694 | -5.867 | -1.764 | 13.954 | 24.970 | -10.454 | 16.999 | 25.160 | -14.373 | -16.038 | 6.769 | 24.433 | 24.228 |
| 15.587 | -22.566 | -7.734 | -20.944 | 26.594 | 15.741 | -17.153 | 28.178 | -0.059 | -22.214 | -20.319 | 5.222 | 20.401 | 20.803 |
| -3.979 | -1.210 | 8.665 | 0.739 | 23.695 | 15.065 | 7.765 | -21.221 | 21.902 | -9.976 | 6.982 | 17.018 | 16.986 | 26.200 |
| -19.253 | 23.140 | -13.072 | 19.552 | -25.607 | 11.201 | -3.760 | -28.226 | -2.018 | 17.266 | 13.163 | 21.227 | 30.810 | 23.423 |
| -20.133 | 6.640 | 23.878 | -3.024 | 25.289 | 3.737 | -8.577 | 30.499 | 28.932 | 7.226 | -7.746 | 18.276 | 18.758 | 25.670 |
| -9.378 | -5.268 | 5.122 | 21.842 | 26.471 | 20.176 | 19.243 | -31.373 | -1.360 | -28.758 | 19.237 | -10.996 | 22.875 | 28.552 |
| -11.425 | -1.194 | 0.172 | 10.696 | 14.300 | 5.119 | 18.032 | 23.145 | 4.303 | 26.198 | -5.688 | 16.559 | 18.289 | -28.151 |
| 25.414 | 15.518 | -1.784 | -1.676 | 22.951 | -3.768 | 5.261 | -26.097 | -18.227 | 28.095 | -9.301 | 8.695 | 21.421 | 2.948 |
| -4.746 | -16.035 | 5.835 | -5.433 | -21.625 | 18.899 | 3.677 | -20.714 | -3.761 | 3.968 | -16.881 | 9.211 | 20.856 | 27.102 |
| 22.695 | -20.767 | 4.253 | -1.365 | -0.500 | 9.608 | -18.618 | 30.368 | -15.300 | 7.959 | 10.489 | 9.855 | 14.980 | 21.226 |
| 0.768 | 23.956 | -4.730 | 3.408 | 3.054 | 25.683 | -4.156 | -30.778 | 2.880 | -30.384 | 19.736 | -5.540 | -1.164 | 18.186 |
| -8.333 | -24.997 | -6.484 | 5.562 | -0.924 | 12.185 | 24.340 | 28.780 | 16.524 | 3.120 | 0.581 | -3.787 | -2.885 | 27.572 |
| 19.921 | 21.732 | 26.812 | -5.453 | -10.500 | 19.815 | 0.457 | 15.370 | 0.587 | -2.151 | -20.173 | 11.671 | -25.392 | 18.644 |
| -7.573 | 26.678 | -6.065 | 22.290 | 10.914 | 21.725 | -3.583 | -4.466 | 27.142 | -19.351 | -13.668 | 21.126 | 14.534 | 15.302 |
| 8.770 | -6.105 | 16.082 | -7.581 | 15.910 | 1.409 | 22.952 | -27.487 | 18.979 | -10.474 | -9.136 | 3.386 | 11.638 | 25.564 |
| 18.240 | 11.127 | -3.626 | -4.030 | -13.167 | 18.561 | 21.346 | 18.125 | -5.540 | -0.192 | 18.660 | 14.100 | 20.438 | 21.988 |
| 9.432 | 1.646 | -19.276 | 17.018 | -22.281 | 16.383 | -0.939 | -28.182 | -28.394 | 18.496 | -19.913 | -9.225 | 17.208 | 27.953 |
| -7.508 | -5.542 | 8.371 | 7.682 | 21.799 | 7.652 | -23.837 | 21.873 | 9.094 | 10.190 | 2.940 | 27.083 | 27.301 | 8.711 |
| -1.529 | 24.755 | -0.504 | -1.018 | 24.588 | 16.022 | 16.441 | 29.785 | 28.432 | -12.825 | -8.589 | 23.504 | 8.521 | 26.984 |
| -18.516 | -11.977 | -9.103 | -4.667 | 18.348 | 11.463 | 2.360 | 26.233 | -3.518 | -27.531 | -12.659 | 5.782 | 25.901 | 22.542 |
| 14.787 | -22.263 | 13.282 | 4.534 | 0.681 | 17.943 | -7.588 | -28.157 | 25.637 | -1.118 | -14.906 | 19.566 | -15.595 | 24.082 |
| -0.539 | 19.798 | -23.477 | 1.048 | 2.160 | 8.863 | 20.016 | 27.879 | 23.950 | -4.518 | 18.828 | 4.121 | 25.530 | 19.705 |
| -23.836 | -9.922 | -14.976 | 27.823 | 5.116 | 20.713 | 28.464 | -12.495 | 3.283 | 4.581 | -19.347 | -2.926 | 20.558 | 19.459 |
| -11.860 | 14.537 | 9.963 | 6.801 | 20.848 | 16.992 | -21.751 | 26.616 | 26.072 | -2.405 | 20.590 | 18.842 | 26.282 | 26.254 |
| -21.762 | -11.742 | 3.876 | 1.166 | 25.845 | 5.883 | 5.593 | 28.030 | 23.010 | -24.598 | 19.776 | 10.337 | 21.844 | 24.130 |
| 23.721 | -6.315 | -6.141 | 6.046 | 19.655 | 23.998 | 10.954 | 26.079 | 28.239 | 28.396 | 26.344 | -4.251 | -25.283 | 10.698 |
| 12.471 | 16.466 | 18.152 | -9.804 | -23.377 | 14.560 | 7.552 | 25.130 | 3.688 | 7.529 | 13.010 | 12.145 | 25.232 | 15.461 |
| -20.254 | -23.361 | -3.062 | 0.642 | -25.784 | 27.519 | 17.835 | 29.112 | -3.139 | -6.582 | 12.167 | 0.620 | 21.691 | 22.987 |
| -1.889 | 11.542 | 6.235 | 26.701 | -18.812 | -26.961 | 8.934 | -27.936 | 8.922 | 2.196 | 7.187 | 11.381 | 17.599 | 25.879 |
| -10.043 | 5.820 | 21.905 | 12.840 | 6.381 | 9.848 | 5.154 | 23.775 | -3.735 | -25.574 | -21.261 | 13.582 | 21.238 | 26.915 |
| -8.718 | -1.964 | -8.289 | -0.849 | -8.743 | 19.095 | -22.030 | 20.045 | -26.642 | -17.992 | -17.125 | 19.557 | 19.370 | 15.761 |
| -18.026 | 25.481 | 16.679 | -13.784 | 11.830 | 16.951 | 4.466 | 28.155 | 22.038 | -23.145 | 5.191 | 12.587 | 23.005 | 22.856 |

| | | | | | | | | | | | | | |
|---|---|---|---|---|---|---|---|---|---|---|---|---|---|
| 0.153 | -21.435 | -0.790 | 22.263 | 9.958 | 21.064 | 0.581 | 29.421 | 21.477 | -11.329 | 1.811 | -15.925 | 17.294 | 21.017 |
| -16.756 | 18.206 | -3.467 | -24.037 | -26.223 | 19.333 | 19.358 | 23.331 | 16.169 | 25.758 | 3.109 | 4.605 | 18.267 | -20.907 |
| -19.443 | -22.064 | -15.674 | 16.386 | -26.941 | 16.522 | -4.594 | -28.996 | 1.980 | -14.332 | 24.947 | 8.164 | 5.834 | 24.951 |
| -18.889 | 17.571 | -22.746 | 8.221 | -27.459 | 25.367 | -19.335 | 27.019 | 7.235 | 23.614 | 16.890 | 25.307 | 23.576 | 15.148 |
| -11.064 | -5.696 | -15.077 | 1.514 | 21.510 | 12.711 | -0.146 | -22.878 | 19.439 | 11.148 | -1.706 | 10.307 | 21.465 | 17.405 |
| 2.055 | 18.535 | 1.055 | 6.625 | 2.192 | 28.492 | 3.377 | 25.296 | -0.838 | 8.429 | 22.819 | 3.621 | 20.222 | 21.937 |
| 0.911 | 3.942 | 8.296 | 11.864 | 25.496 | 14.101 | -19.791 | -18.493 | -2.779 | -1.476 | 25.444 | 10.830 | 28.884 | 26.959 |
| 3.561 | -3.335 | 19.489 | -2.357 | 27.597 | -2.837 | -28.579 | 25.729 | 23.937 | -20.593 | 18.892 | -10.502 | 15.876 | 24.259 |
| 1.369 | 23.569 | 11.224 | 1.022 | 15.107 | 14.904 | 16.877 | -27.660 | 4.480 | 22.476 | -14.034 | 18.985 | 19.955 | 26.470 |
| 4.453 | 10.788 | -1.658 | -11.046 | 23.276 | 23.177 | 18.205 | -22.329 | 19.738 | 15.203 | -7.155 | -16.812 | 26.970 | 23.897 |
| -1.827 | -26.478 | -5.799 | -6.212 | -22.615 | 9.930 | -6.690 | 26.583 | 1.456 | 18.394 | 14.225 | -0.044 | 22.486 | 27.503 |
| 22.129 | -25.665 | -9.995 | 25.387 | -26.276 | -0.451 | 22.731 | -27.972 | 26.980 | -6.628 | 22.100 | -4.521 | 21.067 | 26.553 |
| 18.064 | -4.870 | -13.022 | -31.143 | 4.050 | 17.028 | -24.415 | -25.485 | -2.153 | -23.253 | 15.916 | -1.506 | 24.023 | 14.928 |
| -24.326 | 4.589 | -4.351 | 15.833 | -6.291 | 6.023 | 23.414 | 9.295 | -10.544 | 21.175 | 11.077 | 25.769 | 15.898 | 25.196 |
| 9.874 | -0.428 | 2.785 | 27.210 | -23.851 | 26.249 | 24.261 | -9.510 | 23.258 | -28.816 | 9.449 | 15.321 | 28.601 | 23.254 |
| 9.389 | -10.636 | -6.629 | 3.203 | 27.287 | 13.226 | 18.915 | -25.647 | 13.472 | -18.130 | -20.739 | -8.194 | 14.800 | 9.165 |
| -0.824 | -12.441 | 15.721 | 22.505 | 25.122 | 14.289 | -1.914 | 1.051 | -18.329 | -29.186 | 19.759 | 11.672 | 28.133 | 24.508 |
| -19.382 | 23.773 | 13.420 | -0.160 | -28.324 | 17.790 | 26.618 | 22.904 | 15.109 | -3.736 | 23.090 | 4.103 | 25.067 | 21.803 |
| 7.119 | 22.463 | 2.009 | 14.643 | -17.856 | 15.338 | -15.688 | 12.596 | 0.438 | -24.213 | 17.429 | 17.283 | 23.732 | 22.580 |
| 23.361 | 17.343 | -3.920 | -1.841 | -1.504 | 9.416 | 30.092 | -19.977 | 23.029 | -1.011 | 14.940 | 16.883 | 6.247 | 20.724 |
| 14.452 | 16.474 | -12.124 | 4.826 | 2.645 | 24.961 | 12.643 | 28.370 | 25.101 | 21.509 | 13.877 | -2.754 | 26.683 | 25.294 |
| -6.017 | 3.927 | -7.975 | -21.686 | -21.135 | 22.367 | -24.944 | -16.465 | 20.017 | -28.524 | 24.373 | 16.187 | 8.221 | -10.407 |
| 7.269 | -1.795 | -1.041 | 6.090 | 14.712 | 18.915 | 4.467 | 25.949 | -6.393 | 28.505 | -19.665 | 25.280 | 20.326 | 22.091 |
| -7.497 | -23.267 | 7.841 | 6.894 | 21.186 | 18.392 | 26.284 | 6.711 | -17.774 | 5.931 | 18.886 | 23.395 | 13.391 | 18.698 |
| 19.507 | 14.723 | -19.852 | 3.741 | -12.781 | 22.913 | -9.998 | 21.034 | -8.605 | 22.366 | 22.515 | -6.873 | 10.502 | 23.758 |
| -4.624 | -10.955 | 26.494 | 17.818 | 26.549 | 23.714 | 19.170 | 11.751 | 17.311 | -22.771 | 19.955 | 7.399 | 21.236 | -19.876 |
| -15.707 | -19.962 | 7.310 | 10.619 | 16.978 | 29.141 | 5.781 | 25.144 | -18.786 | 5.336 | 23.549 | -3.008 | 22.731 | 15.959 |
| -10.532 | 24.382 | -9.202 | -0.243 | -7.717 | 22.031 | 16.289 | 23.624 | 0.793 | 0.195 | 21.069 | 6.272 | 27.640 | 13.636 |
| -0.225 | -22.189 | 25.002 | 20.397 | -1.043 | 20.967 | -13.163 | -5.074 | 6.150 | 25.577 | 25.049 | 11.305 | 6.101 | 24.500 |
| 15.781 | -24.258 | -2.166 | 19.956 | 4.452 | 28.338 | 27.582 | -21.331 | 23.502 | -4.724 | -0.963 | -2.478 | 29.199 | -6.866 |
| -25.343 | -17.896 | -14.000 | 15.789 | 12.450 | 15.737 | -16.781 | -16.722 | -0.451 | -6.357 | 2.018 | -23.603 | 28.640 | -8.035 |
| 24.555 | 5.765 | 11.498 | 5.144 | -4.105 | 14.422 | -27.503 | -8.687 | -6.985 | 27.068 | 22.244 | 3.365 | 6.883 | 22.320 |
| 11.598 | -19.440 | 7.640 | -15.785 | 29.264 | 4.177 | 26.884 | 13.472 | -17.709 | -1.324 | 10.002 | 21.612 | 24.946 | |
| 7.400 | -25.912 | 14.672 | -1.113 | 27.250 | 26.910 | 23.215 | 22.184 | 0.420 | 11.229 | -1.530 | 8.990 | -0.940 | |
| 18.481 | 3.705 | -5.588 | 29.521 | 24.911 | 17.428 | -6.706 | 10.307 | 0.022 | 8.941 | -11.439 | 18.464 | 15.993 | |
| 12.272 | 13.156 | 13.648 | 16.169 | -23.668 | 21.720 | 12.089 | -19.262 | -3.813 | -12.713 | -8.840 | 4.824 | 21.654 | |

| | | | | | | | | | | | | |
|---|---|---|---|---|---|---|---|---|---|---|---|---|
| 13.904 | -15.933 | -5.040 | -31.399 | 23.604 | 19.774 | -7.828 | -25.392 | 1.139 | 27.594 | 21.448 | 2.324 | 19.519 |
| -19.483 | 8.416 | -7.641 | -4.211 | -18.955 | 6.775 | 19.532 | 10.290 | 2.558 | -26.784 | 0.191 | 5.235 | 23.533 |
| -22.868 | -23.383 | -3.122 | 27.308 | -26.071 | 24.717 | 29.728 | 19.460 | 4.571 | -18.931 | 5.066 | 8.454 | 26.006 |
| 12.810 | -24.954 | -20.577 | 4.188 | 26.816 | 22.193 | 14.686 | -24.294 | -5.277 | -28.903 | 23.427 | 21.003 | 13.794 |
| -21.759 | 16.254 | -0.469 | -24.553 | -6.221 | 24.546 | 29.993 | 26.406 | 9.513 | 24.053 | -22.910 | -0.604 | -4.304 |
| 11.518 | 10.797 | 20.709 | -2.712 | -9.182 | 9.255 | 2.581 | 14.156 | 18.620 | 5.221 | 6.415 | -10.870 | 4.653 |
| -0.334 | 9.926 | 1.676 | 2.261 | 10.406 | 14.348 | -30.005 | 12.296 | -1.643 | -21.871 | 26.766 | -7.382 | -19.875 |
| 21.587 | -15.740 | -19.247 | 14.269 | -28.971 | 12.474 | -20.415 | 25.332 | -22.545 | -23.077 | -0.213 | 7.032 | -1.773 |
| 25.295 | 15.305 | 2.335 | 8.306 | -0.427 | 23.770 | 27.944 | 23.967 | -24.833 | -9.403 | 24.125 | -5.348 | 17.531 |
| -21.038 | -21.707 | -11.068 | -18.063 | -24.356 | 16.111 | 7.551 | 22.018 | 6.342 | 6.656 | -26.888 | -1.652 | 23.202 |
| 27.847 | 18.478 | -21.112 | 17.709 | 21.957 | 17.586 | 18.032 | 5.641 | 8.603 | 22.528 | 8.426 | 3.925 | 18.888 |
| -21.989 | -9.032 | 8.362 | 23.623 | 12.005 | 24.573 | 19.120 | 1.782 | 12.387 | 28.438 | 17.044 | -4.020 | 21.820 |
| 7.980 | -19.915 | -17.749 | 16.318 | -2.631 | -6.148 | -13.094 | 19.341 | -23.826 | -7.547 | -15.416 | 18.574 | 24.631 |
| -23.750 | 9.086 | -24.566 | -25.267 | 19.780 | 18.414 | 19.802 | 20.848 | -2.933 | 16.884 | 17.818 | 15.722 | -6.358 |
| -11.354 | -17.506 | -8.432 | -2.394 | -11.836 | -30.609 | 5.532 | -11.129 | 19.204 | -6.563 | -23.456 | 12.923 | -4.693 |
| 10.312 | 8.448 | -1.129 | -16.118 | -7.788 | 21.813 | 10.580 | -19.857 | 16.475 | 9.812 | -25.686 | 4.258 | -2.970 |
| 9.738 | -24.220 | -6.285 | 1.273 | -21.654 | 30.553 | 13.005 | -23.680 | -20.312 | 20.887 | -27.082 | -4.702 | -25.092 |
| 22.480 | -23.400 | 19.394 | -0.650 | -23.187 | 15.757 | 28.191 | 23.382 | 1.318 | -21.220 | -0.024 | 13.518 | 21.763 |
| -9.236 | -6.497 | -5.578 | -2.014 | 14.446 | 9.586 | 17.981 | -7.232 | 4.106 | -22.507 | 12.363 | 15.343 | 1.646 |
| 25.223 | 4.561 | 17.779 | -2.606 | -28.561 | 20.695 | 6.492 | -20.348 | -5.565 | -27.545 | -11.715 | 23.528 | 9.120 |
| -25.895 | -1.200 | -14.794 | -6.744 | -12.418 | 15.219 | -4.834 | -12.775 | 12.089 | 15.798 | 20.051 | 21.499 | 20.927 |
| -26.509 | 4.989 | -13.913 | -1.377 | 8.319 | 26.315 | 26.561 | 9.512 | -7.071 | -28.198 | -0.807 | 19.506 | 22.118 |
| 2.717 | -10.924 | 12.733 | 3.336 | 26.565 | -8.756 | -30.277 | 1.490 | -20.306 | 11.071 | -22.506 | 5.888 | 25.106 |
| -3.169 | -24.592 | -2.873 | -18.839 | 20.713 | 14.093 | 23.434 | 16.593 | 9.235 | 13.935 | 22.674 | 0.124 | 23.628 |
| -10.696 | 3.607 | 9.826 | -2.894 | -30.067 | 24.863 | 23.812 | 21.544 | 6.755 | -16.914 | 26.151 | 15.346 | 24.482 |
| -25.413 | -12.781 | 25.598 | 19.328 | 24.455 | 20.619 | 28.342 | -14.896 | 15.989 | 19.415 | 28.864 | 4.170 | -5.593 |
| 9.492 | 7.931 | 7.091 | -0.477 | 2.924 | 23.273 | 15.642 | 9.802 | -5.926 | 2.567 | -11.778 | -0.720 | 23.014 |
| 24.000 | 12.649 | 16.423 | 24.195 | 11.095 | 19.209 | -17.945 | -11.784 | 18.059 | 26.608 | -14.019 | 20.426 | 17.072 |
| 23.373 | -24.071 | -23.745 | 2.249 | -26.136 | 21.825 | 11.727 | 24.464 | 4.539 | 22.250 | 15.244 | 3.751 | 25.260 |
| 21.215 | -1.738 | -12.638 | -1.505 | -18.113 | 15.185 | 19.625 | -17.118 | -0.085 | -4.747 | 19.952 | 8.392 | 20.032 |
| -0.057 | 3.146 | -10.026 | 3.803 | -14.841 | 19.614 | -21.669 | 0.928 | -21.683 | 25.401 | 8.956 | 19.225 | 12.046 |
| -15.734 | -2.164 | -22.419 | -0.872 | 16.328 | 21.368 | 17.901 | -7.279 | -3.837 | 20.629 | 3.100 | 12.345 | -9.737 |
| -9.107 | 17.907 | -1.919 | 3.417 | 20.969 | 27.316 | 22.412 | -24.893 | 23.596 | -23.930 | 18.596 | 20.937 | 18.999 |
| 3.768 | 21.322 | -14.436 | 1.543 | 11.548 | 18.661 | -20.345 | 21.729 | 24.756 | -22.437 | 16.401 | 6.425 | 22.441 |
| -16.142 | 3.771 | -22.550 | -0.913 | 17.391 | 26.053 | 21.217 | 23.729 | 12.061 | -22.913 | 15.770 | 16.950 | 19.226 |
| 13.394 | 10.784 | -24.744 | 0.867 | 4.112 | 19.432 | 16.759 | 24.821 | -7.113 | -2.926 | 16.784 | 21.746 | 7.065 |

| | | | | | | | | | | | | |
|---|---|---|---|---|---|---|---|---|---|---|---|---|
| 4.843 | 25.526 | 11.382 | 5.552 | -11.656 | 20.769 | 29.001 | -21.715 | -8.050 | -26.088 | -21.211 | 14.858 | 17.947 |
| 22.703 | -21.496 | 4.377 | 26.469 | 24.874 | 21.030 | -10.305 | 25.580 | 3.451 | -27.374 | -10.217 | 13.680 | 23.524 |
| 27.334 | 20.734 | -8.797 | 3.514 | 28.679 | 18.927 | 4.235 | -9.498 | -19.891 | 18.008 | 27.308 | 6.209 | 7.730 |
| -16.541 | 5.431 | 4.912 | 17.367 | -24.991 | 28.938 | 26.409 | -5.482 | 1.642 | -8.338 | -15.626 | 24.336 | 16.129 |
| 8.543 | -7.025 | -7.556 | 0.408 | -21.850 | 19.588 | 9.653 | -23.066 | 4.568 | -24.135 | 24.980 | 23.693 | 3.359 |
| 23.796 | 20.470 | -0.188 | 4.550 | -27.104 | -17.793 | 4.612 | -6.426 | 0.856 | -26.312 | 13.441 | 5.020 | 8.505 |
| 0.256 | -5.887 | -4.051 | 13.121 | -23.887 | -0.233 | 21.701 | 0.580 | -1.488 | 15.250 | 20.583 | 8.801 | 1.018 |
| 5.764 | 26.121 | -5.303 | 21.470 | 22.719 | 20.517 | 12.405 | -18.515 | -24.293 | 4.151 | 5.412 | 8.266 | 14.311 |
| -21.999 | 2.341 | 5.992 | -24.105 | 19.134 | 22.160 | -12.258 | 24.710 | -6.574 | -20.368 | 29.356 | -4.794 | -4.575 |
| -12.925 | 10.612 | 18.782 | -6.722 | 26.124 | 21.616 | 13.166 | 20.784 | 17.131 | 10.266 | 22.820 | 25.056 | 16.920 |
| 1.945 | 3.534 | 14.593 | -7.849 | -27.584 | 3.268 | 7.362 | 19.284 | -2.978 | -10.517 | 17.810 | 19.903 | 18.705 |
| -20.669 | -1.829 | -6.572 | -9.485 | 15.405 | -9.425 | -11.049 | 17.924 | 27.888 | -25.485 | 27.761 | 13.180 | 24.668 |
| 15.869 | 6.540 | -16.025 | 20.856 | 20.861 | 20.202 | -19.632 | -4.870 | 14.326 | 27.563 | -1.086 | -6.755 | 23.228 |
| -2.728 | -0.981 | -14.302 | -22.505 | 6.646 | 17.397 | 28.505 | -18.756 | -15.189 | 17.349 | 28.654 | 26.441 | 16.295 |
| -8.324 | 24.631 | 3.105 | 0.283 | 2.470 | 15.079 | 21.831 | 7.619 | 12.951 | 20.511 | 21.357 | 22.585 | 24.240 |
| -6.142 | 17.592 | -7.442 | 10.875 | 21.360 | 26.646 | -27.244 | 20.038 | -7.986 | 23.291 | 18.743 | 22.153 | 19.934 |
| 22.517 | -21.730 | -5.869 | 6.690 | 29.626 | 25.068 | 21.463 | -3.424 | -0.627 | 25.733 | -16.728 | 21.359 | 22.846 |
| 18.666 | -5.994 | 8.942 | 21.203 | 14.600 | 23.316 | 25.708 | 1.699 | 0.789 | 14.296 | -28.842 | 22.794 | |
| -18.740 | 26.105 | -1.540 | -5.727 | 11.633 | 15.907 | 22.594 | 17.756 | 4.281 | 7.918 | 26.797 | 25.699 | |
| 17.171 | 20.081 | -1.990 | 19.457 | 9.707 | 13.605 | -19.897 | 7.248 | -19.150 | 2.815 | -9.884 | 25.080 | |
| 13.123 | 25.559 | 8.554 | -4.971 | 18.144 | 15.284 | -26.075 | 20.709 | -12.044 | 25.189 | -22.690 | 22.638 | |
| -19.310 | -2.358 | -0.667 | -8.930 | 20.830 | 16.989 | 17.438 | 9.333 | 10.692 | -24.580 | 11.752 | 4.362 | |
| -25.314 | -6.945 | 2.012 | -4.349 | 23.970 | 23.368 | -9.504 | -12.676 | 3.056 | -8.856 | 8.431 | 16.691 | |
| 18.387 | -24.692 | 21.786 | -6.207 | -19.503 | 22.269 | 6.017 | -1.913 | -4.870 | 18.776 | 23.119 | 28.860 | |
| -3.000 | 24.110 | -20.219 | -23.472 | -28.566 | 18.589 | -5.530 | -16.715 | -11.957 | 8.368 | -26.615 | 23.497 | |
| -0.765 | -10.771 | 23.365 | 4.784 | 27.057 | 20.936 | -21.362 | -3.566 | -21.024 | 11.581 | -10.409 | 18.337 | |
| 14.312 | -17.697 | -3.250 | -1.186 | 16.094 | 15.278 | 9.351 | -26.424 | 5.748 | -18.453 | 19.753 | -0.006 | |
| -27.222 | -26.731 | 8.339 | 23.540 | 30.972 | -2.599 | 26.368 | -21.728 | 27.906 | 27.012 | 25.677 | 25.881 | |
| -17.655 | -9.240 | -10.896 | 14.488 | -23.972 | 6.393 | 23.407 | -14.796 | 16.276 | 7.331 | 16.657 | 21.033 | |
| 9.480 | 21.397 | 18.066 | 11.247 | 1.026 | 28.312 | 11.648 | -15.364 | 8.268 | -6.330 | -7.529 | -5.560 | |
| 2.675 | -8.220 | -4.154 | 8.456 | -26.386 | 23.214 | 25.286 | 4.262 | 2.534 | -19.586 | 4.242 | 27.056 | |
| 4.406 | -1.431 | 17.431 | -7.049 | 22.900 | 13.817 | 10.414 | -10.936 | -13.181 | 24.028 | -15.554 | 23.974 | |
| 21.706 | -15.847 | -10.424 | -0.266 | -22.075 | 22.273 | 20.427 | -18.333 | -23.820 | -0.002 | -19.345 | 25.471 | |
| 8.810 | 8.135 | -25.888 | -4.639 | -9.627 | -18.987 | 23.315 | -26.375 | 22.040 | -22.117 | 20.780 | 26.152 | |
| -17.087 | 25.804 | -4.588 | 22.127 | -27.690 | 22.101 | 0.309 | 26.610 | -1.283 | 21.836 | 26.801 | 22.321 | |
| -3.923 | 8.711 | 4.439 | -23.121 | -28.678 | 12.833 | 15.957 | 2.768 | -2.788 | -2.146 | 4.917 | 10.459 | |

| | | | | | | | | | | | |
|---|---|---|---|---|---|---|---|---|---|---|---|
| -13.229 | -11.284 | 18.370 | 15.210 | -4.389 | 20.494 | 23.208 | -24.240 | 5.837 | -6.856 | 17.927 | -8.046 |
| -14.895 | 22.193 | -4.255 | -3.668 | -3.432 | 19.129 | -26.771 | -19.964 | 18.340 | -16.710 | 15.501 | 20.998 |
| 22.555 | 25.085 | 10.458 | 7.774 | 7.180 | 19.634 | 12.886 | 27.375 | 22.690 | -23.090 | 21.038 | 18.531 |
| 28.681 | -21.702 | -3.472 | -9.520 | 21.173 | 0.332 | -27.555 | -18.115 | -4.116 | -15.398 | -16.579 | 26.946 |
| 23.190 | 3.182 | 8.092 | 11.353 | -10.877 | -0.102 | 25.190 | -3.244 | -28.471 | 22.970 | 27.442 | 19.071 |
| 26.695 | 9.189 | -7.964 | 20.126 | 30.252 | 17.497 | -4.636 | -7.256 | -21.917 | 12.371 | 24.723 | 24.374 |
| 13.715 | -20.428 | -17.684 | -14.567 | -19.607 | 12.019 | 20.589 | -26.545 | -7.726 | -0.386 | -14.269 | 22.939 |
| 1.398 | 18.614 | 17.912 | -24.798 | -29.705 | 23.750 | -7.472 | -19.536 | -8.397 | -25.087 | 19.590 | 29.542 |
| 25.466 | -3.676 | -2.678 | 18.361 | -9.523 | 7.363 | -23.759 | 15.839 | 19.502 | -3.008 | 0.014 | 21.834 |
| 12.469 | 21.596 | -2.069 | 1.651 | 19.838 | -16.797 | -1.061 | -23.629 | -10.616 | 12.663 | -12.268 | 20.576 |
| -1.221 | 19.491 | 0.108 | -7.234 | 29.071 | 18.730 | -19.218 | 8.425 | 4.750 | -14.585 | 22.628 | 23.793 |
| -29.481 | -1.014 | -10.726 | -25.522 | -22.599 | 5.318 | 21.056 | -14.456 | 26.845 | 2.346 | 16.602 | 29.782 |
| 7.591 | -26.724 | 8.472 | 24.888 | 12.153 | 11.171 | -12.385 | -15.113 | -27.622 | 8.076 | 2.310 | 29.100 |
| -11.851 | -5.134 | -3.154 | 5.754 | -11.753 | 10.690 | -4.841 | 20.850 | 22.219 | 23.395 | -17.247 | 18.115 |
| -6.522 | 11.568 | -6.630 | -4.084 | 20.490 | 18.039 | 7.405 | -23.914 | -6.360 | 19.660 | 21.213 | -5.038 |
| -20.036 | 0.268 | -24.338 | 11.058 | 17.543 | 25.296 | -11.762 | -9.144 | 27.938 | 6.251 | 11.097 | -19.565 |
| 6.540 | 23.508 | -23.761 | 7.467 | 28.387 | 21.132 | 25.088 | 25.746 | 26.822 | 27.101 | 12.655 | 26.177 |
| 4.655 | 1.689 | -16.207 | -24.417 | 9.339 | 17.379 | -16.546 | -27.831 | 12.581 | -26.566 | -2.483 | 20.435 |
| 17.389 | 1.939 | -14.649 | 19.721 | 29.830 | 0.308 | -9.044 | -18.494 | -3.669 | -16.022 | 1.127 | 28.296 |
| 21.996 | 17.547 | 6.479 | -6.213 | 24.061 | 14.085 | 20.933 | 16.435 | -28.384 | 24.521 | 30.396 | 13.905 |
| -28.397 | -17.156 | 7.099 | 20.420 | 11.576 | 20.058 | -22.373 | 14.629 | 21.992 | -1.310 | 22.183 | 18.643 |
| 14.318 | 3.562 | 3.873 | -4.069 | 16.055 | 16.264 | -10.345 | 28.310 | 24.682 | 20.238 | 12.066 | 22.365 |
| 16.546 | 5.262 | -2.196 | -16.244 | 8.137 | 3.787 | -12.682 | -26.552 | 14.281 | 22.485 | 15.455 | 23.351 |
| -19.122 | -21.460 | 26.274 | 8.496 | -23.925 | 4.427 | -1.615 | -1.108 | 1.056 | 0.080 | -11.506 | 17.889 |
| -24.012 | 26.928 | 8.014 | 2.831 | 27.654 | 25.583 | 12.523 | 26.642 | 26.202 | -24.554 | 20.439 | 16.300 |
| 22.801 | -27.029 | -10.356 | 4.551 | 21.760 | 18.191 | -4.895 | 22.140 | 15.091 | 2.881 | 10.884 | 30.376 |
| -15.485 | -0.384 | 15.350 | -10.693 | 23.580 | 1.293 | 17.082 | 21.243 | -0.627 | -9.242 | 16.868 | 27.633 |
| 19.044 | 11.192 | -26.042 | -25.184 | 14.117 | 20.627 | -3.242 | 23.069 | -16.136 | -21.719 | 27.011 | 24.897 |
| -16.878 | -22.564 | -1.767 | 0.443 | -21.299 | -14.335 | -19.128 | 10.064 | 25.883 | 7.849 | -24.667 | 22.774 |
| 20.748 | 13.332 | -18.379 | 14.051 | -24.522 | 20.158 | -10.746 | -6.377 | -20.705 | -22.611 | 27.825 | 20.733 |
| -2.341 | 28.422 | 8.621 | 7.666 | -12.326 | 11.503 | 2.112 | -10.612 | 4.286 | 10.649 | -30.337 | 3.163 |
| -21.919 | -10.840 | 12.021 | -9.859 | -27.579 | 8.416 | -20.899 | 26.729 | -6.503 | 3.992 | 14.031 | 25.874 |
| 18.499 | 12.385 | -6.545 | -26.512 | 21.340 | 11.923 | 26.678 | -20.049 | -10.814 | 24.904 | -9.346 | 11.639 |
| -11.029 | 8.418 | -9.286 | 26.991 | 20.031 | 17.864 | 15.556 | 18.153 | -17.212 | -22.741 | 21.387 | 9.918 |
| 8.202 | -25.323 | 0.093 | 1.063 | -2.385 | 6.823 | 6.743 | -24.756 | 14.227 | -25.651 | -27.029 | 27.893 |
| 22.325 | 20.703 | 2.956 | -4.992 | 28.783 | 24.833 | -23.620 | -27.329 | -31.088 | -0.589 | 16.131 | 21.377 |

| | | | | | | | | | | | |
|---|---|---|---|---|---|---|---|---|---|---|---|
| 3.785 | -0.711 | 5.667 | 3.136 | -15.124 | 22.164 | -21.895 | 30.365 | 3.049 | 25.395 | 8.482 | 8.845 |
| 8.751 | -2.691 | 4.936 | -24.298 | 15.930 | -5.324 | 20.520 | 29.110 | 29.515 | 21.595 | 9.385 | 24.539 |
| -14.348 | -26.583 | 26.214 | 0.272 | -25.792 | 16.749 | -25.505 | -22.258 | 19.979 | 18.162 | 12.378 | 25.415 |
| -19.677 | 5.771 | -20.954 | 3.687 | -8.475 | 6.946 | 20.847 | -3.456 | 21.765 | -3.897 | -14.473 | 17.199 |
| 6.138 | 20.502 | -5.270 | -9.946 | -5.982 | -2.023 | 11.316 | -2.274 | 13.168 | -6.553 | -30.279 | 16.850 |
| 29.342 | -5.235 | -25.963 | -11.498 | -9.061 | 8.389 | -24.369 | -19.336 | -19.338 | 20.181 | 17.478 | 7.531 |
| 2.476 | 2.033 | -7.537 | 14.664 | 5.311 | 9.398 | 19.088 | -16.505 | -5.234 | -21.254 | 13.059 | 26.906 |
| -17.464 | 1.222 | 11.099 | -6.070 | -10.897 | 25.009 | -6.374 | 30.754 | -12.080 | 19.650 | -4.847 | 28.923 |
| 19.361 | 23.803 | 3.627 | -5.154 | -7.892 | 18.418 | 3.748 | -12.905 | 31.756 | 11.161 | 14.001 | 19.003 |
| -28.997 | -25.390 | 20.687 | 23.061 | -20.953 | 25.418 | 2.312 | -22.981 | -2.712 | 12.226 | 26.077 | 19.814 |
| -1.596 | -28.566 | -4.619 | 8.945 | -26.935 | 20.657 | 19.168 | 20.359 | 29.140 | -7.649 | 20.455 | 21.563 |
| -15.921 | 1.582 | -25.282 | -10.674 | -10.089 | 0.339 | -10.532 | 2.485 | 2.691 | -14.452 | -24.104 | 22.876 |
| 0.460 | -0.828 | 9.029 | 9.977 | -25.282 | -3.556 | -24.464 | -14.521 | 22.448 | 0.228 | 31.122 | 22.188 |
| -31.434 | -11.202 | 16.163 | 5.898 | -1.839 | 19.962 | 23.463 | 10.538 | -2.135 | -2.671 | 18.744 | 25.217 |
| 9.804 | 6.303 | -11.534 | -25.500 | -17.723 | 22.368 | -25.136 | -20.913 | -8.228 | 7.452 | 3.827 | 23.853 |
| 9.250 | 8.220 | 12.416 | 12.270 | -8.843 | 17.334 | -16.865 | 12.564 | 28.981 | 3.360 | 24.046 | -30.146 |
| -11.620 | 1.673 | 17.841 | -22.765 | -18.255 | -4.302 | -2.125 | 18.872 | 29.769 | 4.874 | 25.135 | 28.324 |
| 29.898 | -8.509 | -10.369 | -8.789 | 10.282 | 16.741 | -20.771 | -20.442 | 6.054 | -5.459 | -16.356 | 24.346 |
| -19.458 | -19.351 | 13.894 | 25.933 | 0.577 | -0.361 | -1.919 | -27.133 | 10.121 | -4.524 | 30.637 | 23.219 |
| -20.715 | -29.055 | -19.413 | -3.671 | -26.128 | 9.530 | 9.348 | -16.693 | 10.408 | -11.405 | 24.795 | 16.431 |
| -12.449 | 11.074 | 25.863 | 12.246 | 15.649 | 12.799 | -12.971 | 28.641 | -13.824 | 16.981 | 2.435 | 18.246 |
| 12.813 | 9.029 | 11.239 | -15.200 | -27.794 | 16.170 | 2.177 | 9.034 | 6.290 | 10.927 | 5.454 | 22.275 |
| 11.879 | -9.800 | -17.222 | 24.387 | -9.923 | 0.332 | -13.624 | 22.861 | 22.446 | 1.266 | 0.977 | 27.336 |
| -9.348 | -0.212 | 22.685 | 25.340 | 17.347 | 14.137 | 11.803 | -29.309 | -2.828 | 23.353 | 13.394 | 24.050 |
| -21.654 | 18.970 | 5.948 | 21.993 | -24.694 | 25.663 | -12.355 | 24.319 | 26.642 | 0.876 | -13.570 | 26.523 |
| -18.454 | -16.246 | 15.167 | 18.464 | -23.336 | 8.156 | 6.564 | 22.071 | 16.875 | -24.431 | 9.746 | 19.306 |
| 9.511 | 13.417 | -23.287 | 8.893 | 23.257 | 1.553 | 13.656 | -28.902 | 21.472 | -21.677 | 4.529 | 8.891 |
| 8.755 | 0.472 | 2.886 | 10.893 | -21.053 | 21.008 | -15.861 | -19.740 | -0.958 | 13.579 | -3.809 | 27.602 |
| 11.344 | -21.844 | -14.017 | 15.843 | -26.984 | 17.338 | 1.421 | 26.884 | 25.192 | 6.884 | -9.957 | 18.477 |
| 2.281 | -7.642 | -22.194 | 24.065 | 27.668 | 24.886 | -8.793 | 9.873 | 0.356 | -15.246 | -18.084 | -13.711 |
| 1.822 | 19.455 | -10.157 | 2.821 | 1.321 | 15.386 | -13.850 | 0.490 | -16.000 | -19.086 | 14.577 | 23.454 |
| -11.547 | -5.155 | -2.936 | 10.405 | 24.596 | 5.294 | 2.442 | 21.436 | -21.168 | -7.600 | 24.427 | 14.154 |
| -5.123 | 27.548 | -1.772 | 23.299 | 27.966 | 22.158 | -6.887 | 3.389 | 26.651 | -22.329 | -20.036 | 21.495 |
| 10.715 | -22.466 | -11.296 | -27.100 | 15.503 | 21.530 | -22.197 | 16.998 | 25.638 | 0.057 | -22.962 | 23.247 |
| 18.461 | -22.047 | -24.200 | 14.101 | -8.525 | 11.239 | -6.187 | 30.305 | 8.549 | -2.595 | 22.273 | 25.918 |
| 4.775 | 29.068 | -17.673 | 29.454 | -19.425 | 6.469 | 9.721 | -1.555 | -9.561 | -9.451 | -2.727 | 26.358 |

| | | | | | | | | | | | |
|---|---|---|---|---|---|---|---|---|---|---|---|
| 0.388 | 11.696 | 3.853 | 19.993 | 6.230 | 0.127 | -6.460 | -30.298 | 24.898 | -7.287 | 28.130 | 27.734 |
| -20.098 | -3.609 | -1.192 | 25.008 | 7.816 | -3.919 | -17.808 | -28.783 | -22.014 | 8.996 | 10.710 | -10.313 |
| -6.631 | -12.316 | -28.409 | -7.476 | 14.269 | 18.122 | -7.589 | 16.533 | -1.871 | 20.695 | -25.519 | -3.540 |
| 11.770 | -8.273 | 27.552 | -21.058 | -20.525 | 19.086 | -1.550 | -22.500 | 18.580 | -20.124 | -24.139 | 20.840 |
| 1.498 | 1.588 | -21.539 | -23.745 | -24.717 | 22.093 | -21.058 | -29.373 | 15.368 | 13.782 | 5.775 | 24.444 |
| 17.613 | -24.368 | 0.570 | 26.266 | 26.664 | 3.470 | 0.313 | 10.069 | 27.028 | 3.104 | -20.653 | 23.816 |
| -16.314 | 26.423 | -25.307 | -28.418 | 25.867 | 2.399 | -0.344 | -12.570 | 14.238 | 20.377 | 23.768 | 9.727 |
| -10.581 | -15.544 | 19.329 | 24.438 | 23.070 | 26.252 | -16.222 | 29.766 | 28.125 | 13.870 | -15.673 | 16.786 |
| 27.958 | 22.204 | 3.470 | -6.786 | -23.518 | 8.255 | -9.051 | -15.907 | -2.172 | -19.409 | -28.825 | 19.489 |
| 28.305 | 22.974 | 8.080 | -18.579 | -22.320 | 17.214 | 5.564 | -20.671 | -4.153 | 4.270 | -13.047 | 12.189 |
| -9.033 | -15.263 | -6.932 | 6.314 | -20.681 | 24.904 | -18.128 | -30.115 | -0.233 | -5.506 | 26.784 | 26.444 |
| -28.725 | -26.665 | -16.400 | -29.190 | 9.618 | 20.929 | 18.063 | 22.983 | 21.646 | 22.606 | 11.920 | 24.435 |
| -20.460 | 19.230 | -14.058 | -0.650 | -16.710 | 9.556 | 2.566 | 19.209 | -20.585 | -1.878 | -5.318 | 24.978 |
| -27.269 | 16.220 | -7.620 | 25.881 | 24.979 | 15.903 | 12.320 | 25.045 | -25.378 | -17.014 | 12.884 | 27.068 |
| 21.360 | -21.718 | 16.042 | -12.132 | -26.357 | 13.618 | -1.043 | 29.265 | 27.295 | 24.106 | 5.437 | 18.676 |
| -11.744 | -0.934 | 8.108 | -5.854 | 10.413 | 17.832 | 13.345 | 14.875 | -4.346 | -20.636 | 16.544 | -5.091 |
| -18.049 | -9.151 | 18.585 | -23.701 | -14.926 | 25.384 | -11.596 | 27.746 | -19.258 | 9.023 | 29.893 | 25.661 |
| 7.267 | -12.472 | 9.797 | 30.447 | -9.721 | 12.075 | 2.525 | -4.874 | 10.726 | -2.743 | -19.905 | 22.253 |
| -2.449 | -13.928 | -21.078 | -11.112 | -17.165 | 20.889 | -14.710 | -23.867 | -14.322 | 7.157 | -21.298 | 17.620 |
| 16.763 | 25.079 | 25.013 | 21.860 | -24.542 | 13.373 | -5.871 | 23.225 | -22.045 | -11.686 | -5.544 | 22.610 |
| 10.031 | -0.443 | 4.657 | -27.178 | 3.361 | 19.412 | 10.909 | 11.306 | -13.067 | -18.953 | 1.337 | 19.763 |
| -26.033 | -20.986 | -22.162 | -28.517 | 20.936 | 19.996 | -12.122 | 22.279 | 25.654 | 13.415 | 21.444 | 18.556 |
| 22.453 | -25.758 | -19.554 | 24.914 | -25.497 | 22.643 | -22.561 | -15.827 | 2.316 | 21.883 | -0.543 | 21.659 |
| -24.196 | -21.733 | 28.202 | -28.156 | -20.144 | 1.829 | 26.298 | 14.790 | 24.499 | 20.005 | 10.014 | 12.088 |
| -29.547 | -3.346 | -21.527 | -29.349 | 26.392 | -2.956 | -6.981 | 1.373 | -18.794 | 12.432 | 2.450 | 26.618 |
| 14.543 | 30.484 | 6.238 | -2.128 | -23.183 | 25.813 | 9.115 | 20.048 | -5.498 | 1.284 | 1.982 | -11.763 |
| -15.411 | 26.583 | -6.739 | 13.277 | -13.578 | 23.103 | 2.199 | 19.236 | -8.493 | -5.341 | -8.650 | 29.483 |
| 17.304 | 20.962 | -9.258 | 11.345 | -22.036 | 14.563 | -2.185 | 15.200 | -1.507 | 18.987 | -19.521 | 8.392 |
| -6.589 | 24.674 | 18.210 | 20.957 | 5.077 | 17.843 | -9.856 | -22.058 | 5.942 | 25.693 | 26.341 | 23.128 |
| -17.324 | -25.813 | -11.863 | -6.504 | 5.958 | 20.972 | -4.840 | -24.440 | -16.789 | 6.401 | -22.590 | 21.565 |
| -26.324 | -15.489 | -8.464 | 23.697 | 14.806 | 12.993 | 18.671 | -28.661 | 4.312 | -19.064 | 19.690 | -29.000 |
| -18.750 | -28.706 | -17.711 | 19.974 | -6.150 | 16.644 | -3.160 | -24.688 | -21.501 | 23.522 | -24.476 | 7.734 |
| -14.751 | 2.523 | 21.122 | 30.802 | 14.068 | 15.558 | 8.072 | -25.253 | -6.829 | -24.684 | 9.160 | 18.546 |
| 15.202 | 0.644 | 4.686 | 24.511 | -17.759 | 8.897 | 0.280 | -22.256 | -0.640 | -20.492 | 4.415 | -31.090 |
| 14.270 | -24.995 | 19.277 | -29.549 | -19.270 | -0.722 | 2.274 | 22.863 | 2.266 | -15.072 | 6.564 | 26.965 |
| -16.518 | -30.221 | 9.403 | 3.387 | -20.295 | 5.868 | -10.300 | 23.849 | 20.663 | 21.392 | -30.978 | 21.933 |

| | | | | | | | | | | | |
|---|---|---|---|---|---|---|---|---|---|---|---|
| 27.174 | 8.613 | -14.578 | 17.899 | 8.407 | 8.793 | 3.553 | -29.219 | -9.360 | -14.198 | 22.730 | 12.425 |
| 12.750 | 19.384 | 7.884 | -28.933 | 22.079 | -7.681 | -9.450 | -5.863 | 1.490 | -6.413 | -7.931 | -3.689 |
| 13.572 | 10.892 | 9.920 | 31.330 | 23.591 | 4.386 | 16.700 | 13.629 | -0.829 | -27.194 | -20.513 | -7.820 |
| 0.807 | -10.873 | 16.038 | 5.284 | 10.976 | 15.878 | -24.149 | -26.424 | -11.725 | -3.917 | 0.791 | -12.472 |
| 16.582 | 30.070 | -23.311 | 25.513 | 26.259 | 9.856 | -10.266 | 9.549 | -14.582 | 22.377 | 4.742 | 23.570 |
| -1.684 | 28.576 | -11.493 | 6.801 | 15.912 | 6.069 | 23.428 | 27.672 | -24.098 | 8.346 | 3.366 | 22.730 |
| 22.956 | 21.648 | -14.825 | 19.648 | 16.521 | 22.804 | 18.477 | 21.125 | -13.042 | -15.630 | 19.969 | 23.976 |
| -20.969 | -21.172 | 24.266 | 14.819 | -16.676 | 24.538 | -23.870 | 17.278 | 0.628 | -1.697 | 24.822 | 8.826 |
| 12.373 | 7.056 | -16.662 | -5.495 | -7.230 | 19.091 | -9.405 | -13.998 | 8.296 | 24.527 | -4.057 | 15.955 |
| 23.462 | 7.773 | -25.754 | 20.181 | 24.061 | 24.165 | -21.662 | 14.110 | 17.738 | 22.130 | 20.733 | 26.814 |
| -29.156 | -24.021 | -23.489 | 28.243 | -10.378 | 26.034 | 18.048 | -16.145 | -23.257 | -0.770 | 19.062 | 20.864 |
| 27.518 | -0.656 | 30.792 | -27.262 | -20.710 | 26.843 | 23.936 | -24.137 | 11.883 | -19.163 | -5.404 | 19.234 |
| 2.725 | -10.404 | -6.543 | -30.119 | -7.099 | 23.792 | 4.719 | 16.674 | -9.631 | -25.567 | 23.934 | -7.420 |
| 9.097 | 16.979 | -9.437 | 8.674 | -22.159 | 19.890 | 6.536 | 20.331 | -24.974 | -22.765 | 7.733 | 8.270 |
| -22.884 | 24.285 | 16.409 | 6.796 | -4.402 | 20.881 | -24.262 | -4.298 | -25.984 | -20.288 | -25.862 | 28.109 |
| 20.266 | 30.917 | 7.578 | 18.674 | -7.132 | 22.217 | 8.949 | -24.250 | -15.431 | 14.466 | 29.685 | 26.268 |
| 8.395 | 3.842 | 25.816 | -13.003 | 20.991 | 11.452 | 1.347 | -29.237 | 10.412 | 13.886 | 6.941 | 23.121 |
| 6.474 | -6.496 | 11.443 | -28.688 | 14.453 | -12.204 | -11.738 | -17.158 | -7.838 | 23.380 | 9.761 | 11.065 |
| 5.920 | -7.880 | 10.574 | -22.744 | -23.875 | 26.483 | 20.665 | 2.182 | -13.178 | -3.694 | 16.384 | 23.944 |
| -27.093 | -2.270 | 23.154 | 28.778 | -16.501 | 24.339 | -13.704 | -0.047 | 7.711 | -24.026 | 6.119 | 26.747 |
| 10.617 | 6.219 | 28.335 | 10.777 | -1.906 | 15.143 | 20.256 | 28.718 | 19.563 | 11.942 | 25.438 | 9.974 |
| 14.369 | -19.509 | -12.765 | -23.831 | 21.992 | 7.768 | 17.932 | 24.848 | -3.282 | -17.032 | 23.459 | 21.723 |
| -25.265 | -5.886 | 29.675 | -30.961 | 9.588 | 25.383 | 3.203 | 24.107 | 1.588 | 3.430 | -6.362 | -3.895 |
| 26.011 | 19.620 | 18.836 | -17.317 | 18.062 | 13.667 | -17.572 | 27.855 | -19.436 | -13.248 | 23.950 | 23.482 |
| 18.871 | -0.925 | -19.262 | -12.864 | -5.672 | 18.315 | 17.416 | 2.988 | -26.012 | -20.516 | 0.567 | 24.658 |
| -6.855 | -20.018 | 26.667 | 27.862 | 13.632 | 21.145 | 22.844 | 25.673 | 15.776 | 0.518 | -8.811 | 18.148 |
| -17.759 | -29.548 | -29.333 | -11.248 | -10.782 | 10.398 | 14.543 | -3.984 | -12.318 | -22.261 | 14.979 | 25.437 |
| 15.733 | 1.499 | 7.829 | 19.455 | -7.719 | 18.527 | -5.457 | -28.108 | 1.802 | 26.128 | -5.303 | 11.565 |
| 9.983 | 24.633 | -27.201 | -22.148 | -21.069 | -20.667 | -10.981 | -5.935 | -7.448 | -8.050 | 30.184 | 6.553 |
| -16.967 | 22.166 | -4.841 | -28.703 | -9.966 | 25.759 | 24.736 | -22.319 | 11.538 | 17.697 | 6.528 | 28.174 |
| 27.285 | -28.609 | -28.242 | -0.781 | -12.140 | 16.855 | -25.197 | -15.460 | 13.528 | -21.068 | -14.789 | 19.726 |
| -2.612 | 2.857 | -11.038 | -30.270 | -8.407 | 22.911 | 19.671 | -14.433 | -4.148 | -16.146 | 21.937 | 11.165 |
| -19.376 | -24.976 | 17.815 | 20.945 | 2.384 | 2.335 | -0.024 | 26.302 | -3.180 | -14.731 | 17.761 | 17.530 |
| 22.303 | -22.395 | 6.638 | -29.842 | 5.095 | 15.484 | -2.731 | -2.391 | 0.745 | -2.355 | 20.708 | 27.986 |
| 21.894 | -27.315 | -19.395 | -11.162 | -22.249 | 22.506 | -3.996 | 24.747 | 13.223 | 0.385 | -29.558 | 14.105 |
| 24.258 | 7.411 | 31.340 | -27.717 | -25.916 | -8.801 | -27.277 | 14.718 | 17.853 | 27.407 | 24.575 | 16.727 |

| | | | | | | | | | | | |
|---|---|---|---|---|---|---|---|---|---|---|---|
| -3.659 | 0.551 | 12.783 | -5.950 | -0.764 | 12.369 | -5.102 | 7.223 | 7.387 | 8.522 | -23.390 | 11.278 |
| 26.394 | 11.950 | -27.091 | -17.908 | -2.753 | 23.561 | 22.785 | 27.867 | 5.416 | 2.882 | -26.478 | 13.072 |
| 28.783 | -10.172 | 4.732 | -0.956 | -13.365 | 15.765 | -2.849 | 21.996 | 23.932 | 28.075 | 26.563 | 22.707 |
| 21.584 | -21.045 | -5.286 | -21.626 | -17.868 | 24.116 | 24.227 | 21.446 | 14.349 | 7.509 | 20.114 | -18.349 |
| -8.832 | 1.475 | -17.927 | -10.627 | -20.700 | 17.453 | 2.744 | -0.211 | -5.597 | -10.081 | -16.217 | -5.653 |
| 4.581 | -26.871 | -24.761 | -17.236 | -10.592 | 17.084 | -17.751 | -9.909 | -22.778 | -5.667 | 29.424 | 25.277 |
| -28.070 | -30.642 | 15.365 | 18.771 | -0.109 | 9.664 | 4.529 | 18.458 | -14.040 | -24.715 | 7.070 | 23.380 |
| -7.343 | 28.284 | -21.848 | 29.805 | 23.979 | 23.064 | 8.443 | -19.373 | 21.762 | -3.880 | -2.585 | 21.980 |
| -20.307 | 9.505 | 29.151 | -9.639 | -10.129 | 20.792 | -19.584 | -7.883 | 20.191 | 19.959 | 6.502 | 15.781 |
| 18.275 | -20.298 | 5.621 | -16.956 | -18.927 | 21.861 | -27.167 | 20.116 | -7.869 | -27.016 | 18.324 | 25.817 |
| 7.873 | 30.984 | 10.595 | 26.656 | 25.564 | 19.674 | -16.764 | -10.340 | -24.464 | -24.223 | 17.049 | 12.552 |
| 12.193 | -0.081 | -19.476 | 28.197 | -21.767 | 24.481 | 19.554 | -22.905 | -9.056 | -25.885 | -19.776 | 21.371 |
| 10.186 | 30.328 | -1.907 | 30.808 | -14.750 | 12.539 | 7.781 | 26.364 | 2.310 | 2.939 | 27.728 | 20.678 |
| -11.341 | -18.201 | 22.688 | 8.501 | -22.991 | 21.853 | -20.759 | 25.246 | 5.823 | -28.865 | 25.592 | -1.251 |
| -18.725 | 6.034 | 18.000 | -10.833 | 19.128 | 11.743 | 5.520 | 3.613 | 4.052 | 25.649 | -13.798 | 26.051 |
| -22.002 | -5.276 | -23.971 | 23.678 | -6.167 | 21.084 | -23.188 | 7.640 | -11.389 | 16.170 | 21.512 | -4.816 |
| -24.778 | 4.555 | 8.263 | 22.917 | -17.670 | 23.517 | 18.618 | 18.582 | -1.812 | 2.311 | 20.880 | 23.666 |
| -14.520 | -15.748 | -8.545 | 28.220 | 16.607 | 5.434 | 3.073 | 6.378 | 7.352 | -18.339 | -10.367 | 11.098 |
| 16.031 | 10.762 | -10.232 | -11.160 | 24.924 | 16.473 | -20.415 | 19.591 | -8.217 | 27.777 | -11.921 | 6.621 |
| 2.532 | 28.436 | 5.530 | -24.761 | 6.523 | 16.961 | 24.934 | -23.207 | -3.275 | 5.712 | 19.393 | 21.759 |
| -11.070 | -0.665 | -12.335 | -23.926 | -3.717 | 24.601 | 27.002 | -20.599 | 0.327 | 26.822 | 15.164 | 14.444 |
| -19.840 | -14.389 | -25.434 | -4.912 | -5.215 | 19.219 | -1.887 | 5.666 | -12.682 | 19.266 | 30.633 | 2.865 |
| -25.748 | 27.505 | -9.712 | 13.904 | 9.954 | 16.073 | 7.779 | -2.457 | 2.433 | -10.741 | 0.624 | 23.287 |
| 28.700 | -11.208 | -24.586 | 3.018 | -9.237 | 12.970 | 19.029 | 11.559 | -0.720 | -12.028 | -16.526 | 9.454 |
| -23.908 | -20.010 | -21.221 | 26.256 | -23.219 | 26.494 | 8.224 | 24.834 | -0.997 | 17.320 | 27.446 | -15.670 |
| -25.184 | 29.059 | 14.534 | 8.693 | 24.224 | 21.501 | 25.399 | 9.161 | 2.233 | -19.078 | 13.143 | 23.803 |
| 12.040 | 17.978 | -11.451 | 27.207 | -24.814 | 14.660 | -25.525 | 7.267 | -19.125 | 12.955 | 28.915 | 14.992 |
| -24.453 | -18.816 | -21.972 | 21.341 | 5.704 | 14.265 | -27.802 | 22.982 | -21.298 | -29.327 | 6.240 | 16.844 |
| 21.685 | -26.548 | -8.114 | -14.483 | -25.717 | -6.452 | -3.781 | 17.096 | -8.763 | -8.618 | -29.261 | -27.015 |
| 11.316 | -29.818 | 7.234 | -26.810 | -10.554 | 15.435 | -0.390 | -21.424 | 19.274 | 22.612 | 29.882 | -4.364 |
| -19.626 | -23.901 | -20.602 | -0.064 | -13.784 | 28.199 | 17.045 | -2.908 | 7.803 | 0.912 | 19.236 | 15.583 |
| -13.538 | 9.355 | 15.469 | 24.828 | -27.060 | 25.593 | 23.846 | -17.114 | 1.585 | -5.769 | 5.790 | 23.991 |
| -20.326 | -29.988 | -14.082 | 8.092 | -20.282 | 18.116 | 13.462 | 1.968 | 22.220 | -9.006 | -8.327 | 1.539 |
| 13.619 | 29.536 | 26.574 | -3.895 | 18.425 | 20.663 | 21.174 | -8.598 | 17.542 | -11.282 | -26.563 | 3.911 |
| -24.829 | -25.077 | 6.612 | 19.387 | -15.768 | 6.607 | -23.204 | -3.921 | 8.271 | 26.272 | 21.554 | 9.607 |
| -9.703 | 13.321 | 5.498 | 20.696 | -23.255 | -27.191 | -19.469 | 16.257 | -15.095 | -0.914 | 15.738 | -5.896 |

| | | | | | | | | | | | |
|---|---|---|---|---|---|---|---|---|---|---|---|
| -23.103 | -4.129 | -16.986 | 10.217 | 24.919 | -6.593 | 3.551 | -11.220 | 4.071 | -29.579 | -28.941 | -24.442 |
| -6.225 | -5.180 | 7.909 | -12.287 | -13.284 | 2.277 | 26.089 | 3.454 | 22.232 | 3.144 | 27.207 | -5.670 |
| 8.366 | 9.558 | 3.125 | 17.876 | -24.305 | 22.630 | 4.392 | 24.023 | 11.501 | -4.021 | 20.415 | 8.510 |
| 19.105 | 29.874 | -19.262 | 17.282 | 8.553 | 26.204 | -20.549 | 26.718 | 6.025 | -20.990 | 21.141 | -6.369 |
| 0.550 | -13.820 | -16.913 | 26.651 | -11.044 | 12.269 | -18.864 | -13.689 | 8.772 | -7.332 | 10.382 | 15.275 |
| -7.576 | 18.428 | -17.988 | 13.336 | -9.135 | 0.506 | 29.324 | -15.582 | -6.705 | 13.429 | 2.118 | 3.449 |
| 7.274 | -26.745 | 28.971 | 23.967 | 0.208 | 22.154 | -13.512 | -17.122 | 12.972 | 21.817 | 7.859 | 0.705 |
| 23.119 | -28.046 | 8.091 | 28.815 | 24.130 | 19.213 | 21.706 | 18.482 | 2.118 | 19.583 | 22.468 | -0.206 |
| -8.802 | 5.909 | 29.657 | 18.495 | 20.844 | 18.483 | 24.472 | 8.249 | 19.188 | 1.054 | 12.949 | 9.494 |
| 13.996 | -27.640 | 12.571 | 27.057 | 20.460 | 22.864 | -0.422 | 3.177 | 2.672 | 4.853 | 21.819 | 14.428 |
| -10.344 | 20.697 | 24.120 | -7.028 | -24.848 | 21.584 | 3.739 | 1.016 | 7.618 | -3.163 | 18.067 | 22.224 |
| 9.037 | 28.323 | 5.668 | -12.805 | 25.386 | 20.965 | -26.721 | -6.730 | 16.954 | -11.310 | -20.835 | 13.773 |
| 23.707 | -22.451 | -7.579 | -24.938 | -23.923 | 16.969 | 6.378 | -0.732 | 3.851 | -22.487 | 14.304 | -24.138 |
| -24.741 | -4.682 | -24.303 | 14.724 | -27.212 | 27.027 | 21.871 | 9.270 | 0.382 | -21.256 | -16.305 | 12.091 |
| -25.785 | -28.393 | -6.536 | 4.590 | -6.414 | 24.002 | 24.152 | 16.227 | -5.454 | -28.662 | -27.009 | 10.403 |
| 4.745 | -3.793 | -0.563 | -2.467 | -12.471 | 24.891 | 5.021 | -16.549 | -1.823 | 16.382 | -27.726 | 5.810 |
| -1.193 | 0.993 | 4.794 | 4.234 | -2.423 | 20.159 | 2.622 | -22.040 | 0.391 | -9.648 | 27.971 | 19.099 |
| -0.795 | -6.485 | 7.792 | -26.191 | -1.226 | 18.189 | 19.988 | 6.829 | -11.437 | -28.071 | 18.920 | -0.983 |
| -3.330 | 21.832 | -4.546 | 16.320 | -19.029 | 15.246 | 15.053 | 20.925 | 9.674 | -6.968 | 20.906 | 3.064 |
| 20.828 | 9.348 | 20.962 | -18.953 | -11.998 | 17.418 | -5.229 | -8.135 | 1.954 | 2.009 | 25.169 | 7.469 |
| -15.473 | -28.903 | 17.921 | -18.165 | -24.934 | 13.462 | 16.196 | -3.615 | 11.382 | -4.302 | 6.479 | 22.065 |
| 8.169 | -7.811 | 10.156 | -7.593 | 21.974 | 28.441 | 26.551 | -6.511 | -10.665 | -0.643 | -27.313 | 4.220 |
| 0.608 | -22.702 | -10.970 | 7.387 | 0.307 | 20.937 | 7.133 | -25.435 | -7.280 | 1.649 | 7.603 | 0.992 |
| 26.694 | -27.572 | 4.290 | -17.808 | 16.425 | 23.361 | -23.633 | 17.446 | 11.214 | -7.906 | -24.645 | 11.046 |
| 10.131 | -23.387 | -21.454 | 22.268 | 20.964 | 19.937 | -12.915 | 15.031 | 19.795 | -23.084 | 19.838 | 17.470 |
| -10.207 | -27.025 | 8.559 | 20.682 | 10.027 | 26.111 | 20.647 | 7.794 | 18.124 | -19.600 | -3.382 | -1.962 |
| -21.641 | 24.450 | -5.437 | 2.564 | 7.307 | 26.891 | 21.294 | -10.900 | -23.647 | -27.143 | -19.133 | -0.201 |
| 16.905 | -21.698 | -6.499 | -11.201 | -0.027 | 17.056 | -22.156 | 10.034 | 8.183 | 7.499 | 25.698 | 3.191 |
| 25.828 | -21.051 | -3.393 | 25.768 | -17.986 | 21.839 | 9.931 | -26.355 | 21.865 | -11.663 | -21.742 | 23.066 |
| 15.100 | -6.060 | 2.742 | 6.360 | -14.639 | 18.348 | -20.752 | -16.145 | 6.312 | 25.859 | 6.100 | 17.015 |
| -13.986 | -15.489 | 22.767 | -8.143 | 9.221 | 24.535 | 16.737 | -1.250 | -26.576 | 11.124 | -14.577 | -8.665 |
| 6.004 | 17.912 | -17.678 | -20.615 | 0.740 | 14.039 | 27.624 | -4.706 | -20.820 | 18.494 | 23.581 | 11.054 |
| -27.981 | -22.200 | 9.995 | -20.120 | 11.372 | -21.937 | -8.573 | 21.363 | -23.001 | -20.914 | 27.821 | 15.090 |
| -5.329 | 16.954 | 25.775 | 13.764 | -15.516 | 19.071 | 20.792 | 25.678 | 14.501 | -26.986 | 9.791 | 1.511 |
| -15.122 | 28.136 | -8.628 | 5.096 | 14.152 | 31.225 | 18.467 | 10.721 | 25.312 | 29.819 | -1.751 | 6.800 |
| -22.427 | 28.590 | 8.683 | -9.357 | -6.345 | 29.859 | 8.235 | 2.603 | 7.356 | 25.169 | 29.336 | 26.698 |

| | | | | | | | | | | | |
|---|---|---|---|---|---|---|---|---|---|---|---|
| 2.469 | 3.674 | 4.840 | -9.710 | 7.991 | 8.548 | 24.128 | 22.540 | 9.622 | 6.536 | 14.163 | 11.974 |
| 4.911 | 24.527 | 2.873 | -7.913 | -20.793 | 18.564 | -11.062 | 11.566 | 20.142 | -1.360 | 23.489 | 11.225 |
| 23.917 | -3.751 | -26.208 | -25.125 | 19.844 | 15.385 | 1.261 | -13.482 | -5.689 | -25.241 | 13.130 | 23.708 |
| -16.167 | 21.141 | 3.511 | 8.230 | 20.490 | 19.190 | -22.382 | 6.139 | 4.661 | 25.958 | 27.315 | 6.898 |
| -17.218 | -10.098 | -15.103 | -20.598 | -26.703 | 14.296 | 23.230 | 20.057 | 1.001 | 18.230 | -21.818 | 21.062 |
| -21.975 | 24.236 | -12.609 | -26.969 | -15.758 | 17.209 | -18.927 | -9.910 | 13.205 | 27.530 | 22.904 | 20.499 |
| 3.871 | -19.854 | -10.417 | 13.338 | 25.967 | -26.542 | 20.379 | 18.205 | 13.593 | -30.907 | 8.702 | 14.336 |
| -20.303 | -25.735 | -19.188 | -23.086 | 15.929 | 25.753 | 4.441 | 24.597 | 9.066 | -17.310 | -8.508 | -0.928 |
| 8.162 | -13.700 | -8.466 | -18.387 | 16.505 | 22.271 | -27.718 | -3.882 | 7.164 | 13.724 | 25.254 | -22.947 |
| 27.336 | -6.750 | -3.577 | -21.349 | -26.157 | -5.108 | 24.705 | 23.706 | 4.229 | 0.506 | 15.713 | 8.299 |
| -5.935 | -23.085 | -20.837 | 25.628 | 26.838 | 24.979 | 17.117 | -24.343 | -19.326 | -27.761 | 12.667 | 15.275 |
| -11.727 | 26.401 | -24.894 | -9.879 | -18.679 | 20.947 | -19.979 | -3.258 | -1.423 | 4.216 | -18.101 | 11.142 |
| 12.729 | 18.096 | 11.567 | 6.108 | 23.565 | 23.850 | -23.065 | 1.519 | 23.436 | -4.328 | -22.661 | 8.332 |
| 1.271 | -27.086 | 8.835 | 21.279 | 6.954 | 21.249 | -29.518 | 7.149 | 8.643 | -21.209 | 2.609 | -18.576 |
| -20.727 | -2.938 | -6.471 | 18.265 | -15.397 | 11.574 | 23.733 | -17.351 | -4.559 | 3.561 | 17.267 | 23.332 |
| 10.803 | -3.875 | -4.525 | 24.103 | 14.126 | -12.804 | 15.321 | 8.201 | 12.169 | 22.097 | 9.663 | 6.418 |
| -14.548 | -9.700 | -1.197 | 20.312 | -10.082 | 22.143 | -11.985 | -1.551 | 0.585 | -30.445 | -23.923 | -8.989 |
| 13.812 | -14.642 | 19.618 | 2.947 | -24.065 | 20.501 | 25.313 | -21.957 | 0.159 | -22.335 | 20.991 | 21.361 |
| -2.282 | 25.254 | 19.282 | -6.356 | -8.065 | 18.491 | -18.760 | -20.266 | 27.724 | -22.923 | -25.488 | 8.273 |
| -3.303 | -2.050 | -6.927 | 4.531 | -28.554 | 16.810 | 28.298 | -20.150 | -11.375 | -5.582 | 21.889 | 9.851 |
| 20.179 | 27.305 | -13.386 | -26.093 | 22.938 | 22.490 | 21.153 | 4.747 | -7.240 | 30.308 | 19.711 | -6.750 |
| 22.297 | -5.817 | 2.080 | -12.452 | -12.623 | 23.015 | 13.960 | -1.846 | 3.397 | 15.284 | 11.510 | -5.517 |
| 6.598 | 12.279 | -21.962 | 8.330 | 9.500 | 23.432 | -20.256 | -21.814 | -4.903 | -7.349 | 8.036 | 18.375 |
| -23.996 | -23.142 | -18.720 | 25.231 | -27.528 | 18.182 | -2.156 | -23.518 | 10.288 | 27.708 | 20.129 | 27.512 |
| 9.065 | 1.895 | -12.327 | -14.875 | -15.808 | 25.776 | -17.048 | -22.871 | 6.107 | 12.038 | 18.328 | 25.347 |
| 0.486 | -13.782 | -4.330 | 20.818 | -18.996 | 16.257 | 23.076 | 19.695 | 16.367 | -25.834 | -10.988 | -22.900 |
| -25.457 | -2.875 | 8.710 | -0.016 | -23.046 | 19.123 | 27.173 | 16.831 | -2.353 | -2.182 | 6.231 | 24.946 |
| 19.805 | -19.894 | -9.514 | -16.932 | -24.957 | 27.216 | -14.146 | 15.514 | 7.796 | -23.033 | 25.878 | 16.370 |
| 16.392 | 21.325 | 10.439 | -24.518 | 28.149 | 21.327 | 25.479 | 14.751 | 22.950 | -20.003 | 18.747 | 21.013 |
| -6.417 | -13.449 | 11.987 | -17.778 | -11.485 | 17.938 | -11.552 | 17.887 | 10.652 | 23.611 | 17.508 | 10.834 |
| 12.598 | -22.532 | 24.602 | 24.908 | 23.795 | 27.287 | 18.906 | 7.499 | 0.003 | -24.274 | -27.985 | -7.072 |
| -17.097 | 26.178 | 25.723 | -9.130 | 4.156 | 24.756 | 22.217 | 16.945 | -6.011 | 26.980 | 1.767 | 23.761 |
| -22.141 | -1.513 | 10.462 | -20.903 | -21.979 | 20.254 | 27.712 | 13.483 | 9.574 | -12.135 | 15.656 | 14.743 |
| -27.879 | -2.286 | -8.017 | 23.954 | -17.896 | 16.483 | -22.307 | -17.438 | 4.213 | 0.002 | 23.918 | 22.027 |
| -12.925 | -6.761 | -14.986 | 2.838 | 9.970 | 15.765 | 3.239 | -8.860 | -1.850 | -17.865 | 14.853 | -6.091 |
| 18.401 | -9.352 | -16.976 | 17.994 | -26.008 | 13.201 | 16.172 | -14.226 | -5.908 | -1.462 | 21.013 | 26.640 |

| | | | | | | | | | | | |
|---|---|---|---|---|---|---|---|---|---|---|---|
| 4.195 | -5.857 | -23.428 | -8.674 | 27.502 | 11.446 | 19.933 | 12.167 | -15.957 | 26.014 | -10.362 | 21.594 |
| -1.513 | -17.662 | -20.980 | -2.904 | 8.654 | 15.055 | 30.173 | -5.054 | -6.476 | 24.981 | 3.919 | 3.632 |
| -7.348 | 20.542 | 7.726 | -24.524 | -14.012 | 24.103 | 11.693 | -5.949 | -18.999 | 20.382 | -20.015 | 8.630 |
| 21.611 | -22.001 | -7.957 | 24.520 | 15.324 | 14.293 | 21.200 | 12.064 | -24.844 | 19.282 | -25.748 | 20.631 |
| -22.529 | -9.889 | -8.678 | -5.162 | -20.104 | 19.162 | 12.717 | 0.021 | 14.500 | 7.802 | 12.172 | 26.862 |
| 0.359 | -5.237 | 3.476 | 19.887 | -28.637 | 26.904 | 15.762 | 9.361 | -7.495 | -19.288 | 22.318 | 5.332 |
| -6.936 | -10.737 | 11.885 | 19.141 | 21.828 | 24.656 | 4.002 | -21.896 | -7.994 | -28.271 | 21.560 | 18.798 |
| -24.719 | -13.531 | -2.779 | 18.100 | -19.090 | 27.826 | -0.564 | 12.353 | 7.326 | -0.132 | 16.477 | 17.913 |
| -2.030 | -23.831 | -17.325 | 6.923 | -14.818 | 2.820 | -27.595 | -23.697 | -27.374 | -29.907 | 20.128 | 22.748 |
| 16.145 | 27.271 | 1.581 | 26.169 | 16.411 | 16.294 | 7.588 | 2.507 | -6.561 | -11.715 | 5.891 | 11.995 |
| -27.132 | 12.835 | -25.475 | 25.223 | 7.582 | 26.608 | 20.180 | 25.486 | -9.284 | 27.651 | -4.306 | 21.365 |
| 5.322 | 8.621 | -5.447 | -5.559 | 1.152 | 17.484 | -4.322 | 22.841 | -16.704 | -26.954 | 18.463 | 24.716 |
| -6.913 | -24.599 | -22.906 | 18.677 | -12.721 | -8.532 | 21.953 | 13.346 | 19.550 | -3.087 | 8.164 | -4.228 |
| 9.095 | 22.124 | -8.266 | -11.271 | -18.546 | 18.023 | -17.455 | 8.091 | -7.511 | 12.060 | -17.250 | 9.823 |
| 7.236 | -19.659 | -21.965 | 15.067 | -21.765 | 26.515 | -19.445 | -22.680 | 3.858 | 26.880 | 26.293 | 8.583 |
| -8.233 | 3.454 | 10.666 | -12.420 | 19.903 | 2.177 | 27.615 | 18.794 | 29.342 | -4.940 | 24.800 | 29.329 |
| -23.367 | 23.052 | -20.867 | -21.981 | -23.221 | 23.935 | -29.192 | -5.705 | 9.731 | -10.642 | 19.430 | 22.615 |
| 19.880 | -6.707 | -11.576 | 1.261 | 23.300 | 17.494 | 23.941 | -12.467 | 19.605 | 29.331 | -1.951 | 20.580 |
| -11.202 | -6.037 | 23.559 | 3.020 | 12.212 | 20.794 | 31.662 | 15.916 | 6.415 | 7.377 | 23.227 | 15.005 |
| -15.193 | -26.443 | -13.365 | 20.007 | 25.273 | 20.600 | 16.695 | -2.053 | -3.530 | 5.147 | 17.432 | 7.670 |
| 18.225 | 14.797 | 17.150 | 19.318 | 28.623 | 1.156 | -11.189 | 16.743 | -19.270 | -25.281 | 10.398 | 25.851 |
| 1.484 | -24.343 | 4.686 | -2.366 | 14.475 | -9.489 | 10.466 | 17.259 | -1.425 | 13.855 | -16.060 | 6.915 |
| 22.754 | 26.167 | 14.313 | -5.291 | 13.744 | -7.856 | 2.826 | 8.553 | 8.518 | 9.028 | -2.789 | 20.004 |
| 11.144 | 19.780 | -25.591 | 1.473 | 10.269 | 16.918 | 0.031 | -8.214 | -20.657 | 26.460 | -19.791 | 18.758 |
| -12.240 | 19.976 | 8.186 | -2.946 | 9.757 | -19.993 | -21.772 | -24.917 | -5.901 | -4.035 | 23.758 | 26.322 |
| -12.623 | 18.232 | -2.104 | -14.425 | 5.544 | -21.321 | -11.821 | 21.560 | -12.624 | 29.709 | 0.762 | 22.127 |
| 18.660 | -15.177 | -21.484 | -19.481 | -5.330 | -10.966 | 4.082 | -19.649 | 1.784 | -18.226 | 18.252 | 9.639 |
| 15.809 | -23.210 | -9.357 | 12.225 | 24.994 | 25.625 | 8.950 | 26.037 | 19.563 | -19.284 | 5.403 | 24.304 |
| -3.997 | -11.107 | 0.905 | -8.251 | -20.097 | 13.804 | 25.432 | 23.597 | 8.231 | -21.896 | 2.502 | 0.989 |
| 4.352 | -18.169 | -4.610 | 22.429 | -6.229 | -16.433 | -3.476 | 17.048 | 23.365 | -27.886 | -23.312 | 24.998 |
| 3.538 | 2.729 | -3.353 | -20.132 | 22.491 | -5.470 | -3.136 | 2.674 | 21.717 | -2.035 | -13.522 | 16.910 |
| -15.705 | -22.513 | 0.394 | 14.794 | 21.640 | 17.494 | -12.203 | 19.179 | -21.125 | 0.614 | 21.325 | 23.119 |
| 24.521 | 22.397 | -11.998 | 4.466 | 27.925 | 7.065 | -0.547 | -1.804 | 27.882 | -24.817 | -24.640 | 18.705 |
| -7.973 | 12.992 | -0.542 | 15.550 | -19.295 | 1.454 | -20.718 | -18.149 | -19.150 | 21.386 | -25.539 | -10.132 |
| 8.567 | 26.398 | -11.331 | 2.619 | 27.033 | 11.848 | 10.773 | 14.524 | 22.605 | -2.521 | -8.505 | 9.125 |
| -9.782 | -0.719 | -24.041 | -15.511 | 2.710 | -0.707 | 21.810 | -7.779 | 30.391 | 27.551 | 6.815 | -14.818 |

| | | | | | | | | | | | |
|---|---|---|---|---|---|---|---|---|---|---|---|
| 10.542 | -16.503 | -6.627 | 12.990 | 6.906 | 11.501 | -6.085 | 6.322 | 6.585 | 19.998 | 15.598 | 15.584 |
| 25.962 | 21.754 | -20.204 | 2.773 | -15.003 | 6.151 | -19.316 | -12.774 | 16.975 | 28.476 | 26.301 | 14.348 |
| 26.745 | 25.016 | 17.895 | -8.005 | -15.465 | 16.818 | 28.806 | 1.248 | 25.025 | -5.915 | 23.643 | 21.011 |
| 10.233 | 26.229 | -1.156 | 24.801 | -6.101 | 16.351 | 18.136 | -1.791 | 22.761 | -24.238 | -6.362 | 25.855 |
| 0.922 | -12.919 | -14.839 | 10.336 | -16.165 | -1.018 | 20.332 | -18.666 | -14.238 | 13.449 | 1.256 | 23.964 |
| -3.084 | 20.566 | 3.260 | -22.877 | 20.605 | 5.357 | 25.245 | 24.800 | 25.482 | -28.643 | 0.330 | 30.185 |
| 0.398 | -6.927 | 10.131 | 19.525 | -20.459 | 19.121 | 24.401 | 5.591 | -9.268 | -27.425 | 5.101 | 24.923 |
| 18.716 | -25.265 | 2.838 | 19.138 | -25.995 | 19.800 | 25.580 | -1.092 | 26.637 | -25.996 | 14.138 | 6.840 |
| 14.432 | -10.721 | -6.960 | 17.772 | 22.315 | -10.338 | -28.946 | -4.303 | -6.257 | 26.134 | 8.182 | 16.680 |
| -9.335 | -10.482 | 16.097 | 11.803 | 11.206 | -7.213 | 5.589 | -3.595 | -14.139 | 6.471 | 25.427 | 23.778 |
| -24.576 | 24.856 | -12.854 | -20.357 | 23.279 | -0.647 | 17.141 | -21.693 | -27.977 | 14.674 | 7.204 | 12.534 |
| 3.365 | -18.233 | -6.143 | 4.763 | -20.480 | 17.658 | -23.119 | -23.785 | 22.030 | 4.268 | -14.733 | -17.249 |
| -4.467 | -22.386 | -9.228 | 17.249 | -6.629 | 3.420 | 21.406 | -26.007 | -6.901 | 19.506 | -17.880 | 27.518 |
| -13.918 | 19.963 | 13.884 | -20.823 | 21.279 | 2.099 | 27.554 | -22.670 | 16.341 | -17.149 | -11.538 | 26.883 |
| 2.940 | 19.323 | 15.106 | -21.308 | -23.859 | 21.631 | -10.021 | -0.041 | 30.540 | 25.073 | -7.796 | -11.545 |
| 8.046 | -20.428 | 9.176 | -3.176 | 29.112 | 14.456 | -21.077 | -5.654 | -1.590 | 23.117 | 16.419 | 20.778 |
| 2.542 | 22.115 | -11.293 | -8.213 | 24.049 | -1.064 | 18.105 | -15.462 | -3.367 | 0.468 | 1.400 | 21.359 |
| -23.517 | -0.466 | -22.857 | -21.765 | 22.608 | 7.666 | -11.926 | 7.986 | 21.185 | -6.086 | -15.783 | 23.378 |
| 10.179 | 25.689 | -14.485 | 4.148 | -16.837 | 23.021 | 2.273 | -22.314 | 29.150 | 19.859 | 23.724 | 7.768 |
| -5.185 | -24.466 | 1.506 | -3.613 | 30.419 | -14.450 | 24.889 | -5.970 | -15.051 | 14.065 | -22.286 | 6.652 |
| 13.612 | -17.653 | -24.460 | -8.857 | -28.910 | -17.407 | -20.588 | -8.829 | 10.394 | -22.073 | 20.394 | 25.223 |
| 1.770 | 20.454 | 24.185 | 21.982 | -18.349 | -19.048 | -21.869 | 25.985 | 1.848 | -10.333 | 19.963 | 12.282 |
| 5.035 | 15.573 | -25.488 | 18.967 | 24.631 | -21.121 | 26.285 | 26.568 | -7.790 | 3.657 | 19.163 | -6.855 |
| -16.348 | -9.499 | -0.080 | 4.863 | -2.066 | -5.528 | -25.604 | 12.187 | 26.928 | -14.831 | 7.922 | 22.113 |
| -22.489 | 14.856 | -9.525 | 5.994 | -13.319 | -23.627 | -20.107 | 26.884 | -12.143 | -22.778 | 21.743 | 27.839 |
| -3.715 | 6.419 | 8.101 | 20.922 | 21.392 | -6.804 | -3.828 | 25.310 | 28.577 | -27.484 | -7.893 | 19.388 |
| -10.399 | -7.587 | 4.274 | 2.325 | 19.189 | 1.115 | 17.333 | 16.584 | -6.971 | 26.202 | -9.185 | 17.223 |
| 5.815 | 21.552 | 10.369 | 23.051 | -23.941 | -0.163 | -9.970 | 18.634 | 3.387 | 17.211 | -13.272 | -3.148 |
| 1.534 | 9.817 | -7.108 | -14.276 | 20.765 | 12.280 | 7.113 | 27.044 | 1.734 | 15.153 | -5.534 | 26.474 |
| -10.870 | 18.748 | -22.454 | -3.699 | -29.094 | 5.561 | 5.296 | -15.871 | 31.102 | 4.621 | -11.099 | -14.565 |
| 22.997 | 2.373 | 7.026 | -13.722 | 18.158 | 4.429 | -6.082 | 1.836 | 20.037 | -18.925 | 22.208 | 10.723 |
| -1.152 | 6.299 | -16.278 | -16.987 | -4.461 | 20.441 | -7.460 | 8.179 | -6.783 | -29.518 | -6.609 | 20.751 |
| -11.719 | -4.314 | -5.760 | 20.574 | -21.571 | 6.628 | 20.625 | 26.047 | -28.981 | -15.112 | 12.932 | 20.090 |
| -17.388 | 11.646 | -2.411 | 1.106 | -2.547 | -6.893 | 24.048 | 5.102 | 11.940 | -23.342 | 7.001 | 18.701 |
| 10.354 | 13.353 | 11.143 | -1.562 | -10.164 | 26.330 | -16.079 | 9.222 | 29.570 | -0.988 | 17.514 | 24.081 |
| 20.161 | -12.143 | 7.539 | 9.402 | 14.493 | 10.571 | -21.488 | -16.768 | 26.739 | 28.487 | 11.311 | -6.770 |

| | | | | | | | | | | | |
|---|---|---|---|---|---|---|---|---|---|---|---|
| 14.414 | 16.276 | -11.391 | -22.931 | 15.366 | -2.445 | 23.071 | 1.598 | 28.136 | -19.628 | -16.189 | -16.897 |
| -9.546 | -6.979 | 3.397 | -18.519 | -0.398 | 1.843 | 11.432 | 28.679 | 10.874 | -26.036 | 13.621 | 25.510 |
| -8.121 | 12.404 | 9.723 | 16.094 | 25.297 | -22.957 | 22.368 | -28.056 | 22.562 | -27.861 | -0.841 | 19.701 |
| 8.866 | 14.898 | -6.205 | 11.803 | 20.700 | 13.716 | 7.946 | 0.639 | 29.573 | -23.932 | 1.347 | 9.422 |
| -5.386 | 5.089 | -26.139 | 18.185 | -25.954 | 0.714 | -10.773 | -6.143 | 16.471 | 22.915 | 22.540 | 24.329 |
| -0.055 | -22.874 | 16.839 | 7.328 | 16.021 | 12.778 | -18.659 | -27.189 | -6.119 | 21.755 | 19.012 | 14.086 |
| 11.349 | 21.144 | -24.100 | -13.982 | 26.627 | 9.549 | -3.193 | 15.380 | 27.279 | 15.084 | 3.507 | 22.167 |
| 10.805 | 13.930 | -4.441 | 22.519 | -19.640 | -3.602 | 7.260 | -12.741 | -7.325 | -16.199 | 23.769 | 27.174 |
| -24.742 | | -12.827 | -21.967 | -9.248 | -3.113 | -6.329 | 29.306 | -6.677 | -28.957 | | -17.452 |
| -1.476 | | 18.979 | 13.931 | -16.081 | 11.846 | 25.934 | 20.023 | 20.168 | 27.444 | | 23.825 |
| -6.895 | | -13.919 | 14.783 | -29.104 | 4.528 | 1.290 | 5.293 | 25.987 | 20.911 | | -9.855 |
| -13.690 | | 1.460 | 23.203 | -17.667 | 13.453 | -4.955 | 4.044 | 26.563 | 25.903 | | 17.872 |
| -8.838 | | 5.720 | -3.462 | -28.515 | 11.284 | -1.734 | 28.780 | 12.136 | 16.513 | | 15.847 |
| -17.885 | | -4.361 | -13.614 | -20.537 | 1.604 | -19.170 | 29.860 | 25.727 | -4.544 | | -23.418 |
| -17.160 | | -24.003 | -27.294 | -21.275 | -26.481 | -12.371 | -22.066 | 19.690 | -17.164 | | 23.182 |
| -4.757 | | 0.531 | -15.283 | 28.703 | 6.467 | 4.141 | 4.707 | 2.547 | -26.032 | | 27.056 |
| 12.163 | | -5.286 | -0.733 | 24.768 | 6.968 | 20.732 | -7.750 | 28.640 | -10.384 | | 14.347 |
| 2.224 | | 26.601 | -3.567 | -24.533 | 23.009 | -20.078 | 29.945 | 21.745 | 5.723 | | 16.677 |
| -15.253 | | 20.860 | -18.312 | 19.678 | -22.331 | -4.800 | 27.416 | -23.314 | 19.581 | | 30.001 |
| -20.865 | | -17.189 | 5.005 | 28.156 | -5.732 | -15.565 | -16.667 | -11.141 | -1.550 | | 20.523 |
| 20.563 | | -18.006 | -6.219 | 6.631 | 24.004 | -2.112 | 29.256 | 28.018 | -27.855 | | 18.814 |
| 19.082 | | 4.954 | 20.180 | 16.230 | -0.929 | 10.016 | -19.989 | 0.984 | 24.693 | | 4.245 |
| -10.665 | | -11.736 | 19.885 | 5.441 | 17.456 | -18.645 | -4.063 | -28.075 | -14.941 | | 8.982 |
| 5.338 | | 2.062 | -11.977 | 7.758 | -6.636 | -5.263 | -17.113 | 23.859 | 17.071 | | 19.986 |
| 13.855 | | 9.678 | 10.874 | -13.541 | 20.350 | -12.032 | 7.370 | -10.012 | -10.064 | | 18.428 |
| -12.488 | | 18.089 | 16.313 | 21.151 | -21.878 | -8.175 | 6.176 | 25.403 | 8.596 | | 4.487 |
| -18.463 | | 3.891 | -1.722 | 8.390 | 4.115 | -10.105 | -2.562 | -20.560 | -10.633 | | 21.556 |
| -4.081 | | -8.165 | 18.162 | -1.163 | -0.036 | -16.429 | -17.146 | 26.403 | 28.982 | | 21.966 |
| 7.646 | | -4.104 | 16.006 | 30.303 | 26.004 | 16.271 | 20.507 | -5.691 | 22.555 | | 26.387 |
| 1.526 | | -10.896 | 20.517 | -9.231 | -19.368 | 5.628 | -21.537 | 25.200 | -15.832 | | -20.660 |
| 0.663 | | -16.166 | -16.980 | -18.580 | 2.106 | -11.242 | 29.732 | 27.815 | 13.583 | | 24.150 |
| -0.548 | | 8.432 | 28.106 | -25.396 | -18.100 | -7.660 | 19.699 | 9.824 | 8.170 | | 8.624 |
| 20.992 | | -18.523 | -18.458 | 29.544 | 5.096 | 24.294 | -28.926 | -10.406 | -13.869 | | 24.776 |
| -19.496 | | 18.612 | -27.257 | 9.961 | 7.954 | 13.224 | 31.609 | 2.493 | 7.198 | | 11.653 |
| -21.847 | | -20.314 | -0.116 | 20.899 | -5.190 | -6.087 | 21.669 | 18.350 | 25.514 | | 25.770 |
| 2.928 | | 18.225 | 27.530 | 15.346 | 7.198 | 15.065 | -0.137 | -1.685 | -9.250 | | 28.211 |

| | | | | | | | | | | | |
|---|---|---|---|---|---|---|---|---|---|---|---|
| 0.045 | | -12.433 | 26.289 | -24.402 | 18.026 | 11.179 | -14.967 | -28.175 | -23.166 | | 19.613 |
| -6.277 | | -14.006 | 22.018 | -15.286 | 6.077 | 20.947 | -19.244 | -5.850 | 24.397 | | 27.468 |
| -10.378 | | -8.918 | -3.703 | 24.531 | -0.041 | -2.380 | 27.826 | -4.754 | 4.673 | | 23.885 |
| -19.676 | | -3.439 | 13.439 | 23.523 | -5.573 | 1.037 | 19.480 | 14.399 | -3.373 | | 21.265 |
| -24.739 | | -2.406 | 7.863 | 22.923 | -15.915 | 6.461 | -25.156 | 23.660 | -19.265 | | 26.507 |
| -24.385 | | -9.894 | 4.134 | -18.972 | 6.417 | 10.887 | 29.089 | 7.265 | -1.587 | | 22.862 |
| 22.791 | | -27.585 | 25.495 | -2.666 | 28.828 | 10.614 | 28.732 | -27.034 | -25.966 | | 21.772 |
| 18.574 | | 4.779 | -7.739 | -22.510 | 26.024 | 10.116 | -27.067 | -2.060 | 19.439 | | -7.658 |
| 1.493 | | 3.288 | 5.071 | 10.200 | -15.457 | -14.684 | 24.095 | 21.377 | -22.063 | | -8.518 |
| -9.998 | | -18.556 | 10.696 | 3.036 | 9.976 | 12.758 | -6.930 | 27.027 | -13.944 | | 23.625 |
| 3.417 | | -15.127 | -5.338 | 29.160 | 3.571 | -6.055 | -23.849 | 23.293 | 15.317 | | 10.975 |
| 0.139 | | -13.548 | 14.634 | 27.426 | -14.017 | 13.704 | 18.339 | 8.451 | -1.170 | | 14.203 |
| -0.436 | | -8.459 | -13.018 | 26.484 | 22.523 | 25.097 | 14.659 | 25.524 | 16.708 | | 16.075 |
| 4.688 | | -19.361 | 18.499 | 25.876 | 11.595 | -8.439 | 17.262 | -26.385 | -7.652 | | 24.948 |
| -6.825 | | -10.267 | 28.269 | -14.792 | 16.246 | -14.405 | -20.821 | 3.881 | -20.845 | | -0.918 |
| 18.604 | | 5.870 | -16.270 | 18.687 | 17.146 | 17.085 | -8.167 | -9.202 | -15.631 | | 28.938 |
| 23.109 | | 1.731 | 15.911 | -18.305 | 4.852 | -7.008 | -28.466 | 6.368 | -27.117 | | 12.190 |
| -23.421 | | -28.303 | 1.666 | 14.285 | -8.250 | 13.472 | 0.402 | -3.657 | -11.923 | | 25.496 |
| 25.074 | | -12.527 | 17.173 | 8.446 | 26.733 | 2.341 | 9.795 | -9.362 | 23.891 | | -20.844 |
| -3.756 | | -16.687 | 23.948 | -18.742 | 2.591 | 11.158 | -17.608 | -22.825 | 2.777 | | 13.607 |
| 20.333 | | 2.070 | 12.078 | -21.511 | -5.517 | -9.275 | 4.961 | 2.314 | 6.290 | | 28.389 |
| 22.875 | | -5.402 | -23.755 | -23.891 | 14.169 | 16.344 | -13.600 | 2.984 | -0.222 | | 18.495 |
| 21.578 | | -25.141 | 4.468 | -25.316 | -28.565 | -4.751 | 5.650 | -25.789 | -16.322 | | 11.475 |
| 5.895 | | -3.140 | 13.049 | -21.387 | 11.571 | -0.909 | 17.036 | -20.650 | -19.497 | | 28.975 |
| -1.931 | | -8.487 | 22.971 | -27.326 | -7.417 | 2.925 | -29.430 | -5.893 | 27.609 | | 22.965 |
| 13.464 | | -24.681 | 15.463 | -19.459 | -1.690 | 9.314 | 9.268 | 22.384 | 22.681 | | 21.531 |
| 21.639 | | 1.678 | 24.444 | -12.684 | -28.875 | 19.367 | 21.829 | 3.937 | -11.182 | | 10.864 |
| 0.410 | | 9.489 | 17.597 | 13.316 | 29.636 | 18.765 | -12.897 | -23.287 | -6.194 | | -7.269 |
| 9.378 | | -7.539 | -14.340 | 16.368 | 5.537 | 7.241 | -18.578 | 16.687 | -19.340 | | 14.224 |
| -20.919 | | -10.124 | 26.704 | -20.612 | -0.451 | 12.590 | -20.398 | 6.296 | 12.881 | | 20.877 |
| 20.811 | | 3.742 | -20.237 | 29.790 | -5.771 | 19.120 | -23.193 | 22.851 | 12.165 | | 28.851 |
| -1.089 | | 20.154 | -2.654 | -14.711 | 12.407 | -0.150 | 19.989 | -23.028 | -12.131 | | 18.562 |
| -22.966 | | -6.477 | -19.382 | 26.523 | 20.050 | 1.302 | 22.770 | -2.359 | 26.104 | | 20.203 |
| -0.117 | | -17.206 | -22.877 | -24.321 | -30.424 | -16.285 | 18.382 | 15.845 | 12.544 | | -23.201 |
| -11.889 | | -15.777 | -25.914 | -18.170 | 2.738 | -19.485 | -2.165 | -21.256 | -6.770 | | 28.147 |
| 24.492 | | -6.272 | 29.135 | 20.653 | 23.214 | -18.619 | -20.954 | 7.193 | -24.240 | | -1.999 |

| | | | | | | | | | | | |
|---|---|---|---|---|---|---|---|---|---|---|---|
| 1.352 | | -18.196 | 14.086 | 28.206 | 20.907 | 21.514 | 11.018 | -10.592 | 7.307 | | 27.232 |
| -16.902 | | -2.286 | 17.763 | 26.987 | 19.659 | 14.289 | 17.347 | -6.279 | -16.677 | | 22.854 |
| -19.609 | | -9.576 | -29.904 | 22.057 | -2.371 | 19.675 | -7.517 | -23.819 | 11.235 | | 24.085 |
| 7.169 | | -21.472 | -10.103 | -21.027 | 30.512 | 23.837 | -25.021 | 2.914 | 21.480 | | 5.765 |
| 8.474 | | -21.006 | 31.077 | 27.894 | 25.914 | 11.475 | 26.385 | -9.603 | 24.250 | | -0.600 |
| 13.936 | | 7.482 | 5.415 | -8.697 | 28.798 | 22.866 | -17.930 | -5.950 | -15.264 | | 18.829 |
| 0.971 | | -9.956 | -17.152 | 26.692 | -2.593 | 7.776 | 28.046 | 18.568 | -16.622 | | -2.269 |
| -25.365 | | 15.139 | -17.989 | 11.735 | -0.487 | -7.812 | -16.612 | 19.224 | 26.347 | | 28.874 |
| -4.671 | | -10.311 | -11.659 | -28.318 | -12.302 | -4.639 | 19.533 | 2.511 | 2.343 | | 24.963 |
| -8.971 | | 23.696 | -14.041 | -25.019 | 16.948 | 21.962 | 9.742 | 9.370 | -12.228 | | 23.434 |
| 22.981 | | -19.350 | 29.493 | 4.288 | -5.670 | -0.447 | -24.554 | -20.062 | -6.823 | | 19.077 |
| -1.341 | | 26.321 | 16.606 | -21.717 | -16.790 | -7.046 | -4.448 | -21.196 | 4.435 | | 16.617 |
| -16.835 | | -4.604 | 28.415 | 21.853 | -30.202 | 18.991 | 25.555 | -19.298 | 23.606 | | 24.861 |
| 22.889 | | -7.767 | -20.838 | -27.907 | 27.177 | 3.488 | -27.081 | -24.575 | -14.374 | | 21.174 |
| 5.848 | | 19.435 | 18.782 | 24.311 | -2.372 | -8.554 | 5.249 | 0.113 | 17.308 | | 20.375 |
| 17.845 | | -8.738 | -24.878 | 18.207 | 26.568 | -2.471 | 21.628 | -8.321 | -22.478 | | 23.520 |
| 3.350 | | 6.965 | -30.021 | 12.766 | -0.880 | -25.547 | 12.280 | 12.386 | 21.974 | | 14.614 |
| 26.905 | | 28.229 | 18.682 | -18.887 | 12.102 | 1.806 | -5.020 | -3.434 | -4.716 | | -5.286 |
| -26.327 | | -0.645 | 15.943 | -24.090 | -4.487 | 24.865 | -5.936 | 11.573 | -9.204 | | 2.078 |
| 8.985 | | -27.191 | 11.555 | 22.251 | -17.539 | 15.188 | 20.287 | -18.624 | 14.735 | | 15.672 |
| 3.976 | | -16.023 | 0.562 | -4.365 | -28.149 | -9.204 | 21.821 | -0.243 | 11.020 | | 27.380 |
| -24.521 | | -15.586 | 27.394 | -26.083 | -29.562 | -15.133 | -26.831 | -7.870 | 20.374 | | -16.291 |
| -10.428 | | -29.278 | 17.866 | -28.426 | 7.028 | -3.112 | -24.589 | 0.870 | -23.274 | | 20.198 |
| -11.809 | | 13.388 | 30.601 | 26.974 | 29.027 | 22.997 | -4.018 | -16.988 | -12.533 | | 26.731 |
| 22.451 | | 5.416 | 20.278 | 24.623 | 1.863 | 18.026 | 25.999 | -20.223 | 25.642 | | 19.583 |
| -25.909 | | -18.910 | -2.914 | 21.194 | 16.750 | -24.237 | 2.645 | -18.121 | -24.462 | | 12.658 |
| 5.976 | | -9.607 | 26.627 | 13.667 | 27.452 | 11.838 | -8.683 | 0.988 | -19.391 | | 18.132 |
| -7.912 | | 20.946 | -24.536 | 11.439 | -27.212 | -19.769 | -11.102 | 2.530 | -19.883 | | 24.720 |
| 21.533 | | -12.056 | 31.387 | -24.687 | -15.809 | 19.844 | 26.902 | -1.546 | -25.661 | | -3.148 |
| 6.759 | | -19.739 | -20.187 | -25.652 | 2.080 | 23.484 | 3.446 | 0.626 | 4.710 | | 19.090 |
| -16.976 | | -1.889 | -4.652 | -24.148 | 14.376 | 3.022 | -7.499 | 9.628 | -12.790 | | -18.006 |
| -16.393 | | -7.089 | -24.531 | 22.049 | 19.760 | -3.705 | -17.145 | -10.908 | 24.556 | | 10.956 |
| -18.912 | | -11.573 | -28.474 | 26.175 | 24.795 | -6.259 | 12.714 | 15.108 | 6.924 | | 16.735 |
| 27.215 | | 21.462 | -28.950 | -11.219 | -2.840 | 27.047 | 27.266 | 22.741 | 25.888 | | 16.266 |
| 24.105 | | -16.111 | 4.236 | 27.306 | 28.047 | 18.721 | 6.624 | -12.124 | -1.573 | | 22.549 |
| 3.478 | | -30.420 | 18.375 | 19.587 | -13.378 | -13.215 | 27.674 | -21.683 | -23.669 | | -2.303 |

| | | | | | | | | | | | |
|---|---|---|---|---|---|---|---|---|---|---|---|
| 22.907 | | -17.401 | 9.347 | 22.341 | 27.479 | 1.295 | -18.627 | -23.294 | 11.613 | | 27.008 |
| 11.986 | | 13.584 | -27.867 | 23.000 | -1.689 | 5.081 | 10.222 | -3.465 | 9.662 | | 8.986 |
| 1.505 | | 16.258 | -21.861 | -17.980 | 23.633 | -8.123 | 8.017 | -18.011 | -5.322 | | 23.811 |
| 18.822 | | -15.859 | -6.947 | -20.869 | 27.025 | 3.534 | 26.604 | 4.769 | -20.683 | | 2.701 |
| -0.570 | | 6.000 | 1.110 | -27.005 | 7.825 | -1.141 | 12.958 | 10.958 | -2.162 | | 6.034 |
| 13.798 | | -9.786 | -20.102 | 6.458 | -21.352 | 23.999 | 23.461 | -0.737 | 17.826 | | -3.801 |
| 11.059 | | -6.603 | -27.110 | 10.483 | -27.706 | -22.885 | -0.580 | 18.789 | 10.695 | | 24.138 |
| -2.930 | | -23.044 | -25.837 | 19.057 | -23.395 | 1.000 | -5.754 | -1.577 | -8.230 | | -12.355 |
| -21.971 | | -5.828 | -4.288 | 15.017 | 22.095 | 25.585 | 23.878 | 20.792 | -21.516 | | 3.888 |
| 20.060 | | -20.083 | -20.630 | -10.445 | -1.941 | 4.120 | -19.975 | -10.845 | 16.529 | | 18.946 |
| -27.148 | | -21.642 | 17.385 | -19.519 | 16.910 | 21.754 | 6.739 | 24.977 | -24.905 | | 13.457 |
| 15.234 | | -24.206 | 3.438 | 10.795 | -5.513 | 8.319 | -23.065 | -7.616 | 13.612 | | 21.866 |
| -9.408 | | -13.322 | 2.539 | 11.967 | 24.175 | -15.025 | -3.857 | 7.801 | -9.599 | | -3.356 |
| -8.727 | | -18.628 | 8.787 | 4.214 | -3.120 | 27.503 | -24.487 | -8.581 | 15.224 | | -11.582 |
| 17.737 | | 30.741 | -25.506 | -11.888 | 20.981 | 3.794 | -4.967 | 16.529 | 6.716 | | -9.754 |
| 2.612 | | -27.981 | 6.386 | 6.535 | 23.717 | 1.291 | -3.647 | 20.862 | -4.171 | | 21.480 |
| -7.892 | | 14.861 | -21.641 | 1.374 | 26.014 | 14.182 | 12.806 | 1.112 | -19.645 | | 17.437 |
| 9.927 | | -23.369 | 24.375 | 25.810 | 19.799 | 6.022 | -21.700 | -6.574 | -9.188 | | -12.657 |
| -23.695 | | -13.900 | -0.080 | -19.085 | 24.637 | 24.878 | -0.016 | 8.996 | -8.352 | | 19.557 |
| 7.505 | | 19.103 | 8.215 | -25.810 | 15.982 | -9.694 | 11.282 | 10.496 | -21.978 | | -1.045 |
| -27.894 | | 21.425 | -26.713 | -18.283 | 7.539 | 0.713 | -2.019 | -1.417 | 18.440 | | -12.282 |
| -1.122 | | -19.414 | 11.088 | 25.342 | -13.249 | -0.045 | -8.626 | -10.574 | -3.332 | | -1.464 |
| 29.922 | | 24.035 | 17.869 | 3.354 | -15.558 | 27.875 | -12.910 | 22.550 | 0.488 | | 14.525 |
| -23.290 | | -19.092 | -6.490 | 0.846 | 6.974 | -21.626 | -21.968 | -3.545 | -16.058 | | 8.394 |
| 29.072 | | -23.599 | 25.530 | -19.921 | 4.324 | -24.030 | -22.552 | -11.550 | 14.481 | | -9.377 |
| 5.581 | | -4.517 | -15.599 | -6.859 | 21.973 | 5.435 | -16.752 | 6.154 | -11.061 | | 3.680 |
| 7.833 | | -11.086 | 28.408 | 20.452 | -6.837 | -26.316 | -24.523 | -9.502 | -10.031 | | -0.720 |
| 22.992 | | 4.583 | 12.997 | -8.497 | -5.585 | -18.831 | -26.735 | 17.004 | 25.011 | | 22.170 |
| 18.814 | | -4.955 | 9.738 | -16.976 | -26.779 | 28.778 | -12.568 | 16.345 | 13.483 | | 5.078 |
| -27.268 | | -13.369 | -7.464 | -21.034 | -8.301 | -3.444 | -3.294 | 0.987 | 2.654 | | 5.607 |
| 20.227 | | -20.478 | 17.685 | -15.584 | -10.415 | -11.821 | 23.154 | 10.319 | -3.408 | | 0.256 |
| 11.567 | | -18.679 | 29.250 | 14.870 | 26.221 | -14.521 | -25.842 | 25.096 | -22.499 | | 24.534 |
| -28.314 | | 2.839 | -26.251 | 17.680 | -25.061 | 27.607 | 24.158 | 11.958 | -23.459 | | -5.026 |
| 5.310 | | 21.662 | 7.092 | 16.855 | 5.993 | -27.857 | 8.695 | 11.537 | 16.126 | | 22.726 |
| 20.705 | | -8.960 | 19.412 | 24.304 | -7.745 | -13.644 | -23.962 | 6.813 | 11.214 | | -14.020 |
| 16.059 | | 27.249 | -27.564 | 12.645 | 16.358 | 23.586 | 26.315 | 12.822 | -1.821 | | 9.988 |

| | | | | | | | | | | | |
|---|---|---|---|---|---|---|---|---|---|---|---|
| -0.201 | | -19.435 | 24.852 | 4.224 | -16.885 | -12.409 | 4.587 | -20.778 | -1.645 | | 6.234 |
| -26.627 | | 20.458 | -26.704 | 25.976 | 24.817 | 17.073 | 13.194 | -27.040 | 22.162 | | 11.338 |
| -23.445 | | -4.057 | -13.659 | 10.818 | 10.013 | 13.566 | 7.752 | 21.339 | 9.559 | | 2.462 |
| 29.746 | | -21.282 | -20.437 | -21.480 | -23.767 | 15.380 | 16.551 | -3.095 | 5.145 | | -2.541 |
| -2.376 | | -9.879 | 3.942 | 1.214 | 26.043 | 13.863 | 24.165 | 19.370 | 3.037 | | 5.010 |
| -4.990 | | -8.425 | 20.003 | -11.220 | 21.552 | 20.316 | -17.561 | -9.463 | 11.323 | | 12.476 |
| 17.150 | | 3.174 | 26.786 | 4.410 | 18.916 | 29.799 | -2.287 | -2.607 | 21.024 | | -3.267 |
| 30.118 | | 18.938 | -6.391 | 7.002 | 16.229 | -29.016 | 8.250 | 10.058 | -15.594 | | -0.738 |
| -3.295 | | -17.541 | 24.209 | -15.863 | -9.062 | -13.660 | -1.164 | -1.151 | -24.254 | | 25.193 |
| 4.201 | | -18.638 | 4.655 | -16.524 | 22.915 | -16.153 | 13.498 | -6.544 | 4.146 | | 23.699 |
| 17.330 | | -7.333 | 18.058 | 24.995 | 19.659 | 18.158 | -12.755 | 14.351 | -3.248 | | -4.382 |
| 21.645 | | -2.744 | -10.869 | -22.762 | -25.623 | -29.405 | 23.392 | -10.996 | -21.797 | | 13.791 |
| 19.362 | | -22.818 | 14.882 | 23.984 | -1.011 | -22.753 | 6.592 | 11.317 | -12.000 | | 1.916 |
| -1.382 | | -1.051 | -7.282 | -22.325 | 8.156 | 8.203 | 2.027 | 21.952 | -20.330 | | 0.652 |
| -25.150 | | 12.365 | -23.037 | -16.902 | 13.212 | 29.400 | -3.123 | -1.463 | 12.646 | | 5.663 |
| 5.467 | | 9.253 | 2.473 | 19.606 | 17.108 | -21.285 | -12.059 | 27.013 | 1.129 | | 12.581 |
| 29.005 | | 7.042 | 22.454 | 16.383 | -24.552 | 8.806 | -24.433 | 8.490 | -12.631 | | 3.613 |
| 5.975 | | 10.439 | -4.538 | 9.281 | -8.158 | 18.018 | 9.796 | 2.023 | 1.972 | | 0.305 |
| -1.172 | | -20.062 | -27.137 | 23.399 | -8.824 | 4.486 | -25.484 | 19.090 | | | 2.812 |
| 27.685 | | -16.440 | 10.212 | 22.273 | 23.025 | -19.670 | -2.261 | 9.518 | | | 19.476 |
| 24.448 | | 7.263 | 19.040 | -1.907 | -5.414 | 7.233 | 21.405 | -11.688 | | | 16.325 |
| 22.379 | | 11.622 | 24.501 | 11.146 | -6.728 | 30.035 | 9.326 | -25.760 | | | -0.909 |
| -4.173 | | 16.799 | 5.372 | -7.546 | -13.484 | 16.409 | -1.601 | -5.695 | | | 5.245 |
| -8.013 | | 20.615 | 20.445 | 19.080 | -16.360 | 29.790 | 1.187 | 15.426 | | | -7.395 |
| 21.317 | | 18.418 | 4.232 | 18.818 | -23.200 | 24.203 | -11.819 | -0.683 | | | 14.412 |
| -30.482 | | 8.550 | 16.991 | 6.235 | 20.597 | 21.113 | -10.081 | -6.903 | | | 25.860 |
| -19.667 | | -0.965 | 21.885 | 2.515 | 14.447 | -8.882 | -4.982 | 4.049 | | | 18.164 |
| 30.729 | | 11.857 | -12.938 | -7.156 | -14.988 | 28.681 | 23.918 | 28.145 | | | -10.044 |
| 20.028 | | -9.234 | 21.225 | 20.018 | 2.962 | -10.780 | -4.051 | 25.265 | | | -1.419 |
| -20.316 | | 5.031 | 22.233 | 18.833 | 0.169 | 28.056 | 16.480 | 6.201 | | | -13.708 |
| -1.903 | | 4.297 | 19.132 | 22.324 | -18.483 | -31.536 | -6.330 | -2.073 | | | 22.654 |
| 18.236 | | -14.012 | -3.258 | | 5.330 | -21.954 | -25.405 | 26.813 | | | 9.000 |
| 4.500 | | 17.048 | -25.162 | | -3.364 | -12.455 | -13.010 | -3.517 | | | 10.185 |
| -21.058 | | -20.124 | 18.039 | | -13.569 | -31.133 | -21.460 | -9.643 | | | 26.882 |
| -5.949 | | -11.258 | -15.768 | | -5.838 | -5.807 | 11.039 | 21.640 | | | 21.274 |
| 30.608 | | -4.858 | 12.064 | | -0.529 | -14.609 | 20.581 | 5.463 | | | 26.135 |

| | | | | | | | | | | | |
|---|---|---|---|---|---|---|---|---|---|---|---|
| -30.368 | | -9.468 | -26.504 | | -22.640 | 9.262 | -9.566 | 17.949 | | | 19.986 |
| -2.259 | | -8.350 | 18.431 | | -3.848 | 5.857 | 24.949 | -7.164 | | | 14.961 |
| 19.918 | | 25.103 | 25.499 | | 4.949 | 27.785 | 19.658 | 5.916 | | | -12.203 |
| -24.462 | | -15.695 | 3.783 | | -10.170 | -15.324 | 22.662 | 7.085 | | | 23.211 |
| 29.712 | | -11.542 | 10.614 | | -2.033 | 29.661 | 6.565 | 7.959 | | | 2.538 |
| -8.976 | | -3.473 | -10.439 | | -6.702 | 8.666 | -22.518 | 0.834 | | | 5.912 |
| 0.684 | | -18.990 | 2.577 | | 3.386 | -20.246 | 12.364 | -7.960 | | | 13.053 |
| 26.881 | | -2.551 | -5.756 | | | 15.659 | 1.157 | -16.730 | | | -5.075 |
| 13.799 | | 3.894 | 23.415 | | | -30.792 | 15.258 | 20.687 | | | -3.567 |
| -1.917 | | 16.656 | 18.792 | | | -30.039 | 8.634 | 3.669 | | | -7.297 |
| -18.036 | | 11.287 | 8.696 | | | -3.382 | -15.105 | 23.835 | | | 23.543 |
| 11.653 | | 10.000 | 25.943 | | | -15.425 | 2.342 | 2.369 | | | -2.010 |
| -25.622 | | -24.046 | 16.571 | | | -30.217 | -15.843 | -17.228 | | | 24.457 |
| -22.955 | | 3.676 | -19.291 | | | -29.461 | 20.136 | -17.779 | | | 1.461 |
| 23.786 | | -7.184 | 17.698 | | | 26.650 | 5.326 | 13.047 | | | -0.807 |
| 21.341 | | -4.495 | 26.513 | | | 18.522 | -1.573 | 25.989 | | | 24.991 |
| -3.243 | | -22.782 | 10.662 | | | 17.411 | 0.357 | 7.602 | | | 24.430 |
| 19.407 | | 9.366 | -26.039 | | | -27.912 | 3.082 | 17.104 | | | -13.436 |
| 28.346 | | -14.920 | -2.473 | | | 0.529 | -13.939 | 14.404 | | | 19.682 |
| 30.248 | | -13.317 | -5.965 | | | -29.989 | -7.644 | 28.032 | | | 12.995 |
| -18.954 | | 4.204 | 24.243 | | | 24.290 | -12.697 | -18.076 | | | 20.981 |
| -12.478 | | 0.801 | 1.266 | | | -22.407 | 12.531 | -2.272 | | | 3.624 |
| -23.321 | | 19.739 | 24.526 | | | -10.802 | -5.480 | 25.192 | | | 25.517 |
| -7.763 | | 18.174 | 19.337 | | | 28.220 | -17.842 | 18.623 | | | 24.127 |
| 19.846 | | 23.290 | 21.180 | | | 12.662 | | 21.627 | | | 27.462 |
| -13.034 | | 17.128 | -24.595 | | | 28.750 | | 11.283 | | | 28.322 |
| -29.608 | | -19.360 | 5.405 | | | 19.754 | | -10.100 | | | 23.484 |
| -14.378 | | 0.606 | 20.015 | | | -29.496 | | -1.169 | | | 26.075 |
| 21.384 | | 2.608 | 11.008 | | | -2.985 | | 28.881 | | | 17.836 |
| 7.721 | | -7.868 | -23.767 | | | -0.031 | | -19.131 | | | -6.886 |
| -9.098 | | 20.400 | -19.368 | | | 18.437 | | 24.627 | | | 13.758 |
| 11.683 | | 22.598 | 12.759 | | | -21.561 | | 19.506 | | | -24.735 |
| -5.759 | | 15.984 | 25.077 | | | -17.582 | | -11.543 | | | 20.749 |
| -4.007 | | -2.243 | -11.937 | | | 27.611 | | 22.581 | | | 7.609 |
| 11.198 | | -24.340 | -4.793 | | | 26.603 | | 25.738 | | | -6.139 |
| 8.079 | | 4.425 | 18.752 | | | 18.854 | | 11.126 | | | -8.340 |

| | | | | | | | | | | | | |
|---|---|---|---|---|---|---|---|---|---|---|---|---|
| -27.064 | | 0.048 | 4.549 | | | | 27.327 | | 6.442 | | | 9.632 |
| 26.125 | | 25.249 | 25.708 | | | | 7.755 | | 27.824 | | | 18.849 |
| -15.924 | | 13.307 | 19.530 | | | | 23.792 | | -11.245 | | | 22.760 |
| -18.404 | | 0.734 | 13.772 | | | | 20.604 | | 8.947 | | | 25.246 |
| -21.812 | | 26.338 | 21.615 | | | | -22.342 | | 21.815 | | | 22.000 |
| -5.077 | | -1.083 | -20.504 | | | | 8.297 | | 18.796 | | | 27.952 |
| -24.239 | | -10.625 | -8.684 | | | | 27.654 | | -19.219 | | | 19.726 |
| 1.275 | | -5.948 | -7.489 | | | | 23.578 | | -20.239 | | | 8.450 |
| 0.474 | | -26.919 | 17.546 | | | | 26.385 | | 22.212 | | | -18.637 |
| 26.528 | | -25.474 | -23.932 | | | | 17.799 | | 11.638 | | | -4.778 |
| 18.810 | | -4.414 | -23.435 | | | | -19.672 | | -17.025 | | | 19.351 |
| 22.369 | | -2.029 | 22.858 | | | | 22.287 | | 1.513 | | | 23.318 |
| -11.266 | | -22.100 | 14.729 | | | | -26.857 | | -18.952 | | | 4.386 |
| 9.962 | | 23.512 | 24.507 | | | | 20.772 | | 25.760 | | | -3.005 |
| 27.026 | | 5.127 | -9.658 | | | | 10.283 | | 22.766 | | | -9.143 |
| -6.104 | | 25.389 | 10.292 | | | | -26.144 | | 20.275 | | | 6.229 |
| -1.991 | | -28.399 | 13.795 | | | | -8.905 | | 22.410 | | | 20.174 |
| 28.615 | | -25.916 | 12.229 | | | | 15.216 | | 15.721 | | | 22.287 |
| 20.900 | | -27.631 | 22.202 | | | | 9.911 | | -24.340 | | | 26.804 |
| -22.570 | | -3.807 | 21.002 | | | | -27.400 | | 16.038 | | | 24.829 |
| -22.092 | | -19.703 | -8.867 | | | | -4.607 | | -7.600 | | | 14.676 |
| -10.332 | | -23.694 | -11.326 | | | | -27.828 | | 26.996 | | | -5.188 |
| 22.412 | | -11.425 | | | | | 26.123 | | 16.925 | | | -7.421 |
| 25.913 | | -25.033 | | | | | 10.645 | | -2.984 | | | 8.023 |
| 11.792 | | -17.810 | | | | | 15.065 | | 21.007 | | | 26.210 |
| -1.745 | | -7.458 | | | | | -13.947 | | 20.103 | | | 16.330 |
| -13.003 | | -27.204 | | | | | 18.773 | | 8.700 | | | 21.541 |
| -20.387 | | -17.438 | | | | | 25.816 | | 23.821 | | | -7.869 |
| -25.192 | | -26.503 | | | | | -27.099 | | 14.166 | | | 9.640 |
| -6.275 | | -24.794 | | | | | 3.845 | | -12.239 | | | 15.577 |
| 10.871 | | 17.701 | | | | | 9.382 | | 7.831 | | | 9.408 |
| -22.754 | | 14.239 | | | | | 21.019 | | -9.519 | | | -4.545 |
| 4.097 | | 4.493 | | | | | 17.170 | | 30.601 | | | 28.677 |
| 24.359 | | -5.455 | | | | | 26.040 | | -1.046 | | | 23.627 |
| 9.417 | | -11.499 | | | | | 2.264 | | 5.154 | | | -22.493 |
| -3.512 | | 29.141 | | | | | -14.339 | | 15.530 | | | -5.185 |

| | | | | | | | | | | | |
|---|---|---|---|---|---|---|---|---|---|---|---|
| -4.272 | | -0.363 | | | | -15.969 | | 11.153 | | | 9.708 |
| 6.279 | | 26.651 | | | | -0.032 | | 29.491 | | | -3.185 |
| 21.050 | | -27.980 | | | | -3.011 | | 26.281 | | | 27.857 |
| -25.696 | | -9.656 | | | | 4.403 | | -19.551 | | | -6.184 |
| -13.271 | | 29.546 | | | | 13.792 | | 13.711 | | | 21.057 |
| -12.150 | | 30.182 | | | | 10.718 | | 1.416 | | | 6.040 |
| -5.157 | | -23.957 | | | | -12.137 | | -18.172 | | | 10.290 |
| 18.587 | | -17.985 | | | | 18.454 | | -30.390 | | | -18.569 |
| 14.558 | | 30.667 | | | | 18.026 | | 30.913 | | | 22.228 |
| 25.942 | | 17.319 | | | | -17.493 | | 8.334 | | | 19.168 |
| -5.889 | | -7.568 | | | | -22.747 | | 13.988 | | | 23.417 |
| 1.535 | | 13.055 | | | | -16.697 | | 19.779 | | | 25.548 |
| 6.840 | | -7.178 | | | | 5.480 | | 6.173 | | | -1.435 |
| 11.648 | | 12.583 | | | | 0.777 | | -21.821 | | | -8.532 |
| -9.583 | | 16.192 | | | | -19.927 | | 25.332 | | | 28.981 |
| -5.015 | | -6.832 | | | | 12.489 | | 12.077 | | | -16.445 |
| -11.043 | | -22.796 | | | | -3.512 | | -26.240 | | | 21.122 |
| 27.356 | | -5.619 | | | | -2.070 | | -32.048 | | | -6.248 |
| 6.681 | | -28.655 | | | | -18.264 | | 23.976 | | | 20.538 |
| -24.911 | | 29.233 | | | | 12.253 | | 30.644 | | | 16.703 |
| 4.026 | | 5.017 | | | | -22.196 | | 23.588 | | | 21.798 |
| 2.247 | | 6.121 | | | | -9.963 | | -16.446 | | | -13.012 |
| -14.678 | | 27.584 | | | | -12.417 | | -13.476 | | | 1.880 |
| -9.739 | | 21.826 | | | | -16.280 | | 29.275 | | | 10.059 |
| 12.473 | | -22.410 | | | | -7.784 | | -30.850 | | | 24.855 |
| -8.415 | | 25.039 | | | | | | -30.126 | | | 18.872 |
| -4.425 | | 15.977 | | | | | | 30.016 | | | 12.704 |
| 22.960 | | -29.784 | | | | | | 27.112 | | | 11.555 |
| 21.821 | | -24.110 | | | | | | -5.825 | | | -5.962 |
| 4.421 | | -14.050 | | | | | | 19.153 | | | -13.927 |
| -15.522 | | -31.587 | | | | | | -17.861 | | | 22.803 |
| 0.058 | | -21.637 | | | | | | 10.480 | | | -15.604 |
| -25.478 | | -8.033 | | | | | | -17.154 | | | 7.172 |
| 0.670 | | -16.595 | | | | | | -29.153 | | | 20.985 |
| 5.268 | | 26.671 | | | | | | 23.146 | | | 18.140 |
| 26.495 | | 2.681 | | | | | | 8.714 | | | -17.015 |

| | | | | | | | | | | | |
|---|---|---|---|---|---|---|---|---|---|---|---|
| 8.895 | | -3.342 | | | | | | -28.575 | | | 12.195 |
| 2.668 | | 19.953 | | | | | | -20.850 | | | 16.461 |
| -2.583 | | -10.241 | | | | | | -9.073 | | | 11.097 |
| -10.505 | | -21.329 | | | | | | -25.650 | | | 27.614 |
| -4.872 | | -14.434 | | | | | | -22.665 | | | 25.805 |
| 12.433 | | -14.857 | | | | | | 26.790 | | | 19.099 |
| 2.309 | | 11.307 | | | | | | -19.187 | | | -9.674 |
| -3.324 | | 3.175 | | | | | | 18.766 | | | 11.479 |
| -18.610 | | -31.464 | | | | | | -11.936 | | | 9.120 |
| 0.322 | | -31.091 | | | | | | 25.460 | | | 4.810 |
| -8.028 | | 30.329 | | | | | | -16.584 | | | -12.461 |
| 10.007 | | -4.875 | | | | | | 16.619 | | | 9.791 |
| -16.998 | | 3.781 | | | | | | 27.712 | | | 18.060 |
| 23.875 | | -18.942 | | | | | | 20.918 | | | 15.698 |
| 15.982 | | 1.995 | | | | | | 14.115 | | | -6.005 |
| -6.950 | | -12.707 | | | | | | 16.606 | | | 28.633 |
| 11.212 | | 27.680 | | | | | | -2.047 | | | 26.159 |
| 5.560 | | 24.376 | | | | | | 24.709 | | | 19.287 |
| 6.033 | | -23.070 | | | | | | -9.362 | | | -17.069 |
| 6.771 | | -21.736 | | | | | | 26.663 | | | 19.846 |
| -10.016 | | -30.014 | | | | | | 22.949 | | | 25.650 |
| 14.947 | | 11.457 | | | | | | 22.272 | | | -9.644 |
| 8.010 | | 8.615 | | | | | | 12.089 | | | -8.814 |
| -14.777 | | -2.035 | | | | | | -14.739 | | | 19.287 |
| -24.956 | | -19.950 | | | | | | 8.744 | | | 12.886 |
| 13.322 | | 27.945 | | | | | | 7.626 | | | 23.176 |
| 1.621 | | -1.353 | | | | | | 22.492 | | | -5.062 |
| -3.530 | | 28.339 | | | | | | -24.729 | | | 24.675 |
| 8.892 | | 11.434 | | | | | | 23.851 | | | 22.415 |
| -12.399 | | 25.905 | | | | | | 4.692 | | | -19.320 |
| 3.886 | | -1.590 | | | | | | 11.221 | | | -20.026 |
| -22.866 | | -28.283 | | | | | | -2.199 | | | 26.818 |
| -6.633 | | 23.262 | | | | | | 24.293 | | | 22.054 |
| 19.158 | | 28.263 | | | | | | -10.295 | | | -8.999 |
| -6.360 | | -2.276 | | | | | | -13.083 | | | 19.766 |
| 6.944 | | -6.722 | | | | | | -8.959 | | | 16.454 |

| | | | | | | | | | | | |
|---|---|---|---|---|---|---|---|---|---|---|---|
| -23.551 | | -21.322 | | | | | | -21.101 | | | -20.057 |
| 12.118 | | 26.392 | | | | | | 25.541 | | | -6.856 |
| 4.616 | | -29.254 | | | | | | -10.502 | | | 14.340 |
| 18.580 | | -29.696 | | | | | | 5.358 | | | 13.089 |
| 6.455 | | -22.639 | | | | | | 1.662 | | | -22.842 |
| -7.585 | | -27.622 | | | | | | 25.901 | | | -15.832 |
| 21.992 | | -29.097 | | | | | | 9.426 | | | 23.528 |
| -4.139 | | -4.735 | | | | | | 27.863 | | | -29.271 |
| -4.741 | | 8.555 | | | | | | -14.072 | | | 17.294 |
| 2.263 | | -20.281 | | | | | | 16.268 | | | -13.465 |
| 3.678 | | 27.801 | | | | | | -12.437 | | | 16.810 |
| -15.706 | | 19.210 | | | | | | 10.846 | | | -16.608 |
| -7.979 | | 3.212 | | | | | | -28.029 | | | 22.232 |
| 13.108 | | -27.769 | | | | | | -19.253 | | | 21.855 |
| 7.777 | | -6.754 | | | | | | 6.851 | | | 10.757 |
| -22.689 | | 6.733 | | | | | | 23.533 | | | 9.137 |
| -15.173 | | -28.401 | | | | | | -5.092 | | | 21.616 |
| 1.282 | | -8.234 | | | | | | 26.358 | | | 18.742 |
| -4.031 | | 11.169 | | | | | | -21.825 | | | 26.748 |
| -1.523 | | -27.647 | | | | | | 6.846 | | | -15.131 |
| -25.851 | | -20.787 | | | | | | 2.065 | | | 9.905 |
| 2.512 | | -12.038 | | | | | | 24.620 | | | -8.827 |
| 7.492 | | -22.464 | | | | | | -22.943 | | | -4.259 |
| -18.755 | | -23.227 | | | | | | -23.291 | | | 24.609 |
| 8.428 | | 26.610 | | | | | | 12.082 | | | 16.207 |
| 5.203 | | -5.502 | | | | | | -25.319 | | | -0.818 |
| -10.435 | | -7.223 | | | | | | 4.408 | | | 24.358 |
| 19.663 | | 7.184 | | | | | | -7.695 | | | 25.376 |
| 20.866 | | 25.586 | | | | | | 25.174 | | | 20.080 |
| -1.476 | | -19.463 | | | | | | -14.443 | | | 27.924 |
| -10.665 | | -1.599 | | | | | | 3.257 | | | 18.097 |
| -9.510 | | 5.128 | | | | | | -23.600 | | | 22.837 |
| -0.317 | | 19.731 | | | | | | 19.383 | | | 6.261 |
| -14.448 | | 4.227 | | | | | | -2.518 | | | -2.129 |
| -19.932 | | -24.749 | | | | | | 5.495 | | | 25.995 |
| 8.806 | | -26.294 | | | | | | -17.293 | | | 14.881 |

| | | | | | | | | | | | |
|---|---|---|---|---|---|---|---|---|---|---|---|
| 22.014 | | -10.818 | | | | | | 17.509 | | | 21.998 |
| 2.571 | | 9.840 | | | | | | -18.708 | | | 17.395 |
| -9.209 | | 6.405 | | | | | | 11.652 | | | 25.019 |
| -13.081 | | -20.438 | | | | | | 21.505 | | | -22.991 |
| 4.771 | | -3.017 | | | | | | 2.839 | | | -7.103 |
| 25.393 | | 24.491 | | | | | | 10.962 | | | -8.096 |
| -17.720 | | -15.071 | | | | | | 19.217 | | | 28.413 |
| 9.917 | | 4.973 | | | | | | -20.663 | | | -18.848 |
| 7.611 | | -2.541 | | | | | | -21.663 | | | -11.468 |
| -6.044 | | 11.511 | | | | | | -13.484 | | | -22.865 |
| 3.682 | | 6.736 | | | | | | -4.595 | | | 25.687 |
| 20.265 | | 13.420 | | | | | | -19.525 | | | 7.341 |
| 20.557 | | 16.195 | | | | | | -2.972 | | | 18.614 |
| -7.089 | | 4.682 | | | | | | -23.044 | | | 27.323 |
| -2.251 | | 3.858 | | | | | | -0.532 | | | -3.484 |
| -10.630 | | 9.622 | | | | | | 15.072 | | | 26.756 |
| -25.655 | | 5.254 | | | | | | -2.008 | | | 21.502 |
| 9.154 | | 0.667 | | | | | | 9.489 | | | 20.686 |
| 14.078 | | 8.209 | | | | | | -15.042 | | | 1.013 |
| -24.599 | | -27.150 | | | | | | 19.648 | | | 18.681 |
| -26.691 | | -0.846 | | | | | | -3.566 | | | -1.314 |
| 4.428 | | -1.957 | | | | | | 7.523 | | | 14.829 |
| -8.204 | | -25.352 | | | | | | 16.135 | | | 16.080 |
| -14.918 | | -24.870 | | | | | | 8.962 | | | -14.479 |
| -11.982 | | 18.176 | | | | | | -16.107 | | | 8.706 |
| -4.035 | | 24.439 | | | | | | 8.057 | | | 3.525 |
| 19.937 | | 4.023 | | | | | | 10.229 | | | -4.904 |
| 3.019 | | -10.033 | | | | | | -12.096 | | | 24.415 |
| -18.237 | | -0.046 | | | | | | 24.313 | | | 21.095 |
| -13.193 | | -3.836 | | | | | | 13.309 | | | 16.743 |
| -3.333 | | -20.724 | | | | | | -0.828 | | | 22.848 |
| -1.514 | | -20.098 | | | | | | -11.451 | | | -9.823 |
| -5.252 | | 8.446 | | | | | | -6.304 | | | 19.419 |
| -18.662 | | -6.862 | | | | | | 2.096 | | | -11.431 |
| 5.720 | | -12.531 | | | | | | 25.476 | | | 3.056 |
| 1.945 | | 4.038 | | | | | | 7.448 | | | -3.598 |

| | | | | | | | | | | | |
|---|---|---|---|---|---|---|---|---|---|---|---|
| -25.216 | | -5.897 | | | | | | -12.284 | | | -2.242 |
| -15.906 | | 21.486 | | | | | | 14.111 | | | -5.596 |
| 8.624 | | 6.887 | | | | | | 21.040 | | | 22.459 |
| 9.896 | | -9.961 | | | | | | 19.644 | | | 23.273 |
| 6.513 | | 8.286 | | | | | | 18.584 | | | 14.250 |
| -1.576 | | 14.733 | | | | | | 24.112 | | | 24.161 |
| 11.846 | | 2.189 | | | | | | 2.000 | | | -6.797 |
| -24.466 | | -9.183 | | | | | | 3.706 | | | 26.123 |
| 17.518 | | -18.495 | | | | | | 2.825 | | | 25.027 |
| -6.268 | | -17.653 | | | | | | 7.085 | | | 22.515 |
| -20.850 | | -19.948 | | | | | | -11.199 | | | 20.019 |
| -28.587 | | | | | | | | -4.887 | | | 10.706 |
| -0.150 | | | | | | | | 18.848 | | | 6.365 |
| 21.545 | | | | | | | | -23.773 | | | -3.455 |
| -21.833 | | | | | | | | 1.562 | | | 12.976 |
| -15.162 | | | | | | | | 7.862 | | | 22.425 |
| 12.955 | | | | | | | | -10.274 | | | 16.460 |
| 11.113 | | | | | | | | | | | 15.112 |
| -4.753 | | | | | | | | | | | -10.131 |
| -21.434 | | | | | | | | | | | -24.363 |
| -28.538 | | | | | | | | | | | -15.054 |
| 17.486 | | | | | | | | | | | 26.935 |
| 4.467 | | | | | | | | | | | -1.323 |
| -20.846 | | | | | | | | | | | 17.064 |
| -1.534 | | | | | | | | | | | -10.803 |
| -20.293 | | | | | | | | | | | -12.824 |
| 8.552 | | | | | | | | | | | -7.533 |
| -29.732 | | | | | | | | | | | 2.187 |
| 5.473 | | | | | | | | | | | 13.917 |
| 1.971 | | | | | | | | | | | 25.481 |
| 24.586 | | | | | | | | | | | -19.917 |
| 7.891 | | | | | | | | | | | |
| 4.277 | | | | | | | | | | | |
| -27.162 | | | | | | | | | | | |
| -7.625 | | | | | | | | | | | |
| 20.259 | | | | | | | | | | | |

| |
|---|
| 21.783 |
| -15.302 |
| -28.090 |
| -0.958 |
| -8.994 |
| -11.515 |
| -13.961 |
| -21.436 |
| -13.408 |
| -6.382 |
| 23.322 |
| 19.983 |
| 0.314 |
| -2.449 |
| 14.587 |
| -21.903 |
| -20.701 |
| 0.461 |
| -29.318 |
| 25.374 |
| -11.743 |
| -7.972 |
| 20.509 |
| 13.407 |
| -10.488 |
| -31.248 |
| 7.727 |
| -30.055 |
| -20.883 |
| -13.539 |
| -30.642 |
| -3.412 |
| 29.818 |
| 13.503 |
| 10.229 |
| -20.396 |

| |
|---|
| 19.372 |
| 12.090 |
| 27.998 |
| -21.454 |
| 1.166 |
| -6.710 |
| -4.766 |
| 25.105 |
| -14.055 |
| 18.848 |
| 16.874 |
| 14.343 |
| -18.113 |
| -30.202 |
| 17.805 |
| 21.808 |
| -10.662 |
| 1.778 |
| -28.998 |
| -14.487 |
| -29.447 |
| -4.538 |
| 11.214 |
| -20.187 |
| -8.037 |
| -9.985 |
| 30.756 |
| -11.507 |
| -14.939 |
| 9.399 |
| -25.863 |
| 25.498 |
| -18.095 |
| 0.108 |
| -0.806 |
| -8.283 |

| |
|---|
| 1.518 |
| -22.454 |
| -10.093 |
| 24.367 |
| 30.295 |
| -21.783 |
| 30.737 |
| -11.207 |
| -13.069 |
| -14.426 |
| -13.660 |
| 14.240 |
| -6.940 |
| 18.045 |
| -2.910 |
| -8.629 |
| 21.234 |
| -18.085 |
| 29.629 |
| -22.745 |
| -15.437 |
| 15.252 |
| 21.898 |
| -12.211 |
| -16.859 |
| -23.875 |
| 19.309 |
| 29.034 |
| -4.836 |
| -29.714 |
| -28.666 |
| 22.625 |
| 6.989 |
| -11.350 |
| -12.399 |
| 30.141 |

| |
|---|
| -0.612 |
| -13.660 |
| -9.181 |
| -5.558 |
| -18.547 |
| -4.055 |
| -9.702 |
| 0.094 |
| 22.277 |
| 16.325 |
| -15.809 |
| -11.909 |
| -22.380 |
| 28.598 |
| -6.907 |
| -6.344 |
| -19.553 |
| 10.277 |
| 27.929 |
| -0.806 |
| -7.948 |
| -10.321 |
| -5.672 |
| -28.058 |
| -9.463 |
| -24.324 |
| -4.631 |
| -14.046 |
| 26.877 |
| 6.728 |
| -27.171 |
| -5.712 |
| 23.147 |
| -22.130 |
| -6.760 |
| -27.792 |

| |
|---|
| -20.595 |
| -16.677 |
| -2.882 |
| 25.944 |
| -8.055 |
| -18.402 |
| -9.530 |
| -23.498 |
| -6.170 |
| -1.624 |
| -6.918 |
| -25.437 |
| 10.055 |
| 2.233 |
| -21.129 |
| -10.465 |
| 7.716 |
| 0.759 |
| 5.177 |
| -13.823 |
| -5.645 |
| -26.809 |
| -7.242 |
| -1.347 |
| -10.000 |
| -16.514 |
| -20.824 |
| 9.630 |
| 7.040 |
| 13.657 |
| -6.843 |
| -9.548 |
| -6.053 |
| -15.198 |
| -23.256 |
| -19.670 |

| |
|---|
| -2.548 |
| -11.468 |
| -8.430 |
| 5.001 |
| -25.645 |
| -15.602 |
| 13.996 |
| 5.914 |
| 15.904 |
| 11.405 |
| 1.564 |
| -7.681 |
| -22.582 |
| 18.850 |
| -5.561 |
| -23.455 |
| 24.955 |
| 6.619 |
| 11.927 |
| 12.563 |
| -9.448 |
| 0.626 |
| -4.998 |
| 5.066 |
| -2.075 |
| -3.718 |
| -7.533 |
| 24.669 |
| 10.592 |
| 23.268 |
| -6.092 |
| 14.892 |
| 0.275 |
| 5.523 |
| 20.903 |
| 16.293 |

| |
|---|
| 9.772 |
| 7.243 |
| 8.075 |
| 17.187 |
| -3.121 |
| 20.208 |
| 7.776 |
| -1.640 |
| 12.492 |
| 8.354 |
| 3.970 |
| 9.140 |
| -2.913 |
| -15.285 |
| -11.493 |
| 7.310 |
| -10.602 |
| 15.504 |
| -9.747 |
| 1.249 |
| -4.052 |
| 5.859 |
| -3.360 |
| 4.573 |
| 18.828 |
| 23.558 |
| -19.618 |
| -7.584 |
| -1.444 |
| 2.409 |
| 14.193 |
| 4.315 |
| -19.759 |
| 3.727 |
| 2.099 |
| 22.335 |

| |
|---|
| 5.869 |
| -3.925 |
| 6.829 |
| -2.399 |
| -0.795 |
| -11.125 |
| -11.489 |
| -25.665 |
| 0.605 |
| -23.921 |
| 9.519 |
| 3.868 |
| -1.047 |
| -3.654 |
| -26.110 |
| 10.701 |
| -3.004 |
| 20.049 |
| -8.309 |
| 8.817 |
| -16.045 |
| -13.767 |
| -5.111 |
| -7.039 |
| -19.069 |
| -10.186 |
| -10.966 |
| -18.388 |
| 24.906 |
| -27.002 |
| -23.905 |
| 1.575 |
| 10.499 |
| 26.336 |
| 6.827 |
| -10.502 |

| |
|---|
| 3.126 |
| 25.157 |
| -7.281 |
| -1.799 |
| -21.142 |
| 5.920 |
| 0.135 |
| -17.083 |
| -13.835 |
| -20.396 |
| -16.446 |
| 1.230 |
| -6.411 |
| -11.668 |
| 0.922 |
| -4.717 |
| -22.492 |
| 1.284 |
| 0.120 |
| 20.457 |
| 17.121 |
| -3.192 |
| 12.091 |
| -5.761 |
| 14.001 |
| -7.824 |
| 12.879 |
| -14.263 |
| -15.831 |
| -5.426 |
| -8.721 |
| -21.004 |
| -27.828 |
| -23.433 |
| -1.257 |
| 17.879 |

| |
|---|
| 25.353 |
| 22.365 |
| -23.071 |
| -5.515 |
| 27.256 |
| 18.326 |
| 19.151 |
| -13.006 |
| -9.090 |
| 18.597 |
| -9.773 |
| -20.771 |
| 0.413 |
| -29.152 |
| 24.672 |
| -19.191 |
| -26.655 |
| -16.502 |
| 21.470 |
| 20.342 |
| 19.760 |
| 22.237 |
| 26.616 |
| 26.999 |
| 23.527 |
| -7.182 |
| 20.100 |
| 25.062 |
| -18.099 |
| 28.711 |
| -6.011 |
| -3.775 |
| -21.991 |
| -7.466 |
| 20.581 |
| 29.128 |

| |
|---|
| -22.347 |
| 9.448 |
| 12.360 |
| 20.400 |
| 11.554 |
| -18.526 |
| 10.112 |
| 4.063 |
| -8.624 |
| 7.342 |
| 17.510 |
| 14.321 |
| 2.718 |
| 30.795 |
| -21.740 |
| 5.860 |
| 1.899 |
| 23.386 |
| -11.288 |
| -11.853 |
| 30.640 |
| -16.956 |
| 23.576 |
| 1.836 |
| 16.868 |
| -20.093 |
| 17.313 |
| -19.682 |
| 30.217 |
| 6.630 |
| 28.339 |
| 24.142 |
| -20.851 |
| -12.243 |
| -16.280 |
| 21.985 |

| |
|---|
| -31.142 |
| 30.466 |
| 18.700 |
| 11.998 |
| 14.055 |
| -11.823 |
| 25.670 |
| 14.701 |
| 29.773 |
| -6.817 |
| -3.017 |
| 18.941 |
| 9.739 |
| 9.101 |
| 3.495 |
| 22.577 |
| 4.955 |
| 28.532 |
| 17.030 |
| 10.630 |
| -23.436 |
| -21.827 |
| -5.523 |
| -1.470 |
| -18.776 |
| -27.362 |
| -13.779 |
| 21.341 |
| -2.330 |
| -23.420 |
| 8.666 |
| 21.608 |
| -28.139 |
| -10.994 |
| 23.040 |
| 0.292 |

| |
|---|
| -27.323 |
| -29.124 |
| 25.541 |
| 24.890 |
| -4.058 |
| 17.843 |
| -20.078 |
| 23.078 |
| -19.713 |
| 14.368 |
| -22.005 |
| -18.688 |
| 10.518 |
| 8.215 |
| 6.315 |
| 8.795 |
| -15.461 |
| 4.466 |
| 20.127 |
| 18.559 |
| -20.619 |
| -25.186 |
| -0.249 |
| -8.637 |
| -27.174 |
| -3.718 |
| -25.926 |
| -2.154 |
| 13.754 |
| -4.266 |
| -15.654 |
| 0.521 |
| -23.556 |
| 8.751 |
| -0.301 |
| 1.143 |

| |
|---|
| -15.359 |
| 25.429 |
| 12.825 |
| -6.067 |
| -10.943 |
| -8.529 |
| -23.642 |
| -24.770 |
| 17.533 |
| 15.053 |
| 9.414 |
| 21.912 |
| -20.977 |
| 3.447 |
| -3.366 |
| 9.752 |
| 17.616 |
| 23.460 |
| -4.229 |
| -9.360 |
| -2.000 |
| -13.090 |
| -20.453 |
| -25.932 |
| -8.696 |
| -19.887 |
| 4.019 |
| -17.175 |
| -11.172 |
| -18.336 |
| -22.540 |
| -12.518 |
| -6.372 |
| -11.325 |
| -20.423 |
| -19.884 |

| |
|---|
| -21.520 |

### FIGURE-2B-DOSE-RESPONSE-CURVE

| Test compound | Concentration (mM) | Physiological condition | Mean difference (mm) | CI 5 | CI 95 | Error high | Error low |
|---|---|---|---|---|---|---|---|
| IAA | 0.003 | Fed | 2.017 | 0.890 | 3.164 | 1.147 | 1.127 |
| IAA | 0.01 | Fed | -0.607 | -1.690 | 0.447 | 1.054 | 1.083 |
| IAA | 0.03 | Fed | 0.743 | -0.385 | 1.828 | 1.086 | 1.128 |
| IAA | 0.1 | Fed | -0.424 | -1.532 | 0.710 | 1.134 | 1.108 |
| IAA | 0.3 | Fed | 3.557 | 2.469 | 4.677 | 1.120 | 1.088 |
| IAA | 1 | Fed | 5.710 | 4.577 | 6.839 | 1.129 | 1.133 |
| IAA | 3 | Fed | 8.452 | 7.444 | 9.553 | 1.100 | 1.009 |
| IAA | 10 | Fed | 7.880 | 6.806 | 8.933 | 1.053 | 1.074 |
| IAA | 30 | Fed | 12.077 | 11.091 | 13.109 | 1.032 | 0.986 |
| IAA | 100 | Fed | 11.796 | 10.712 | 12.883 | 1.087 | 1.084 |
| IAA | 300 | Fed | 12.442 | 11.403 | 13.455 | 1.013 | 1.039 |
| IAA | 1000 | Fed | 13.347 | 12.323 | 14.346 | 0.999 | 1.025 |
| IAA | 3000 | Fed | 12.495 | 11.457 | 13.543 | 1.049 | 1.038 |
| IAA | 0.003 | Starved | 2.018 | 1.091 | 2.918 | 0.900 | 0.928 |
| IAA | 0.01 | Starved | -2.118 | -2.918 | -1.330 | 0.788 | 0.800 |
| IAA | 0.03 | Starved | 0.930 | 0.074 | 1.821 | 0.890 | 0.857 |
| IAA | 0.1 | Starved | 1.319 | 0.459 | 2.226 | 0.908 | 0.859 |
| IAA | 0.3 | Starved | 4.685 | 3.863 | 5.517 | 0.832 | 0.822 |
| IAA | 1 | Starved | 0.644 | -0.199 | 1.512 | 0.868 | 0.843 |
| IAA | 3 | Starved | 2.765 | 1.896 | 3.657 | 0.891 | 0.869 |
| IAA | 10 | Starved | 3.139 | 2.290 | 3.974 | 0.835 | 0.849 |
| IAA | 30 | Starved | 1.548 | 0.628 | 2.442 | 0.894 | 0.919 |
| IAA | 100 | Starved | 8.723 | 7.803 | 9.600 | 0.877 | 0.920 |
| IAA | 300 | Starved | 10.851 | 10.099 | 11.673 | 0.822 | 0.752 |
| IAA | 1000 | Starved | 9.573 | 8.691 | 10.470 | 0.897 | 0.882 |
| IAA | 3000 | Starved | 17.497 | 16.798 | 18.198 | 0.702 | 0.698 |

| FIGURE-3A-REPLICATE-MEANS | | | | | | | | | |
|---|---|---|---|---|---|---|---|---|---|
| Well | Total worms | Chemotaxis index | Compound | Concentrati | Physiological condition | File name | Well width | Plate ID | Mean position (mm) |
| 1A | 177 | 0.436 | DMSO | Solvent | Fed | 2M1BDR01_Scan001 | 3048 | 2M1BDR01_P01 | 6.842 |
| 3A | 158 | -0.145 | DMSO | Solvent | Fed | 2M1BDR01_Scan001 | 3079 | 2M1BDR01_P03 | -2.628 |
| 1A | 172 | 0.103 | DMSO | Solvent | Fed | 2M1BDR03_Scan001 | 3043 | 2M1BDR03_P01 | 2.427 |
| 3A | 180 | 0.118 | DMSO | Solvent | Fed | 2M1BDR03_Scan001 | 3069 | 2M1BDR03_P03 | 1.407 |
| 1A | 267 | 0.122 | DMSO | Solvent | Fed | 2M1BDR04_Scan002 | 3050 | 2M1BDR04_P01 | 2.481 |
| 3A | 196 | -0.019 | DMSO | Solvent | Fed | 2M1BDR04_Scan002 | 3083 | 2M1BDR04_P03 | 0.103 |
| 1A | 162 | 0.034 | DMSO | Solvent | Fed | 2M1BDR13_Scan004 | 3050 | 2M1BDR13_P05 | 0.748 |
| 3A | 169 | 0.137 | Solvent | Solvent | Fed | 2M1BDR13_Scan004 | 3067 | 2M1BDR13_P07 | 1.915 |
| 4A | 204 | 0.148 | 2M1B | 0.003 | Fed | 2M1BDR11_Scan001 | 3039 | 2M1BDR11_P04 | 2.834 |
| 4B | 212 | 0.130 | 2M1B | 0.003 | Fed | 2M1BDR11_Scan001 | 3042 | 2M1BDR11_P04 | 3.040 |
| 4C | 192 | 0.102 | 2M1B | 0.003 | Fed | 2M1BDR11_Scan001 | 3050 | 2M1BDR11_P04 | 1.863 |
| 4D | 259 | 0.138 | 2M1B | 0.003 | Fed | 2M1BDR11_Scan001 | 3053 | 2M1BDR11_P04 | 2.109 |
| 2A | 312 | 0.213 | 2M1B | 0.003 | Fed | 2M1BDR12_Scan002 | 3074 | 2M1BDR12_P06 | 3.406 |
| 2B | 321 | 0.164 | 2M1B | 0.003 | Fed | 2M1BDR12_Scan002 | 3064 | 2M1BDR12_P06 | 2.705 |
| 2C | 326 | 0.075 | 2M1B | 0.003 | Fed | 2M1BDR12_Scan002 | 3079 | 2M1BDR12_P06 | 1.212 |
| 2D | 342 | -0.107 | 2M1B | 0.003 | Fed | 2M1BDR12_Scan002 | 3076 | 2M1BDR12_P06 | -1.162 |
| 1A | 246 | 0.046 | 2M1B | 0.01 | Fed | 2M1BDR11_Scan002 | 3042 | 2M1BDR11_P05 | 2.027 |
| 1B | 225 | -0.135 | 2M1B | 0.01 | Fed | 2M1BDR11_Scan002 | 3040 | 2M1BDR11_P05 | -1.164 |
| 1C | 228 | -0.137 | 2M1B | 0.01 | Fed | 2M1BDR11_Scan002 | 3023 | 2M1BDR11_P05 | -1.507 |
| 1D | 258 | 0.085 | 2M1B | 0.01 | Fed | 2M1BDR11_Scan002 | 3045 | 2M1BDR11_P05 | 2.405 |
| 4A | 331 | -0.024 | 2M1B | 0.01 | Fed | 2M1BDR12_Scan001 | 3081 | 2M1BDR12_P04 | -1.057 |
| 4B | 325 | 0.106 | 2M1B | 0.01 | Fed | 2M1BDR12_Scan001 | 3034 | 2M1BDR12_P04 | 2.539 |
| 4C | 319 | 0.182 | 2M1B | 0.01 | Fed | 2M1BDR12_Scan001 | 3075 | 2M1BDR12_P04 | 3.132 |
| 4D | 360 | -0.068 | 2M1B | 0.01 | Fed | 2M1BDR12_Scan001 | 3054 | 2M1BDR12_P04 | -0.570 |
| 2A | 215 | -0.123 | 2M1B | 0.03 | Fed | 2M1BDR11_Scan001 | 3046 | 2M1BDR11_P02 | -1.433 |
| 2B | 222 | 0.337 | 2M1B | 0.03 | Fed | 2M1BDR11_Scan001 | 3033 | 2M1BDR11_P02 | 6.283 |
| 2C | 227 | 0.035 | 2M1B | 0.03 | Fed | 2M1BDR11_Scan001 | 3038 | 2M1BDR11_P02 | 1.812 |
| 2D | 271 | 0.221 | 2M1B | 0.03 | Fed | 2M1BDR11_Scan001 | 3055 | 2M1BDR11_P02 | 3.791 |
| 1A | 332 | 0.111 | 2M1B | 0.03 | Fed | 2M1BDR12_Scan001 | 3050 | 2M1BDR12_P01 | 1.603 |
| 1B | 332 | 0.052 | 2M1B | 0.03 | Fed | 2M1BDR12_Scan001 | 3051 | 2M1BDR12_P01 | 0.624 |
| 1C | 247 | -0.051 | 2M1B | 0.03 | Fed | 2M1BDR12_Scan001 | 3048 | 2M1BDR12_P01 | -1.017 |
| 1D | 295 | -0.177 | 2M1B | 0.03 | Fed | 2M1BDR12_Scan001 | 3045 | 2M1BDR12_P01 | -1.507 |
| 3A | 265 | 0.052 | 2M1B | 0.1 | Fed | 2M1BDR11_Scan001 | 3038 | 2M1BDR11_P03 | 1.922 |
| 3B | 248 | 0.313 | 2M1B | 0.1 | Fed | 2M1BDR11_Scan001 | 3044 | 2M1BDR11_P03 | 4.892 |
| 3C | 180 | 0.132 | 2M1B | 0.1 | Fed | 2M1BDR11_Scan001 | 3048 | 2M1BDR11_P03 | 3.744 |
| 3D | 269 | 0.353 | 2M1B | 0.1 | Fed | 2M1BDR11_Scan001 | 3055 | 2M1BDR11_P03 | 6.104 |
| 1A | 210 | 0.074 | 2M1B | 0.1 | Fed | 2M1BDR12_Scan002 | 3054 | 2M1BDR12_P05 | 1.816 |

| | | | | | | | | | |
|---|---|---|---|---|---|---|---|---|---|
| 1B | 267 | 0.091 | 2M1B | 0.1 | Fed | 2M1BDR12_Scan002 | 3048 | 2M1BDR12_P05 | 2.229 |
| 1C | 232 | 0.218 | 2M1B | 0.1 | Fed | 2M1BDR12_Scan002 | 3048 | 2M1BDR12_P05 | 4.579 |
| 1D | 320 | 0.042 | 2M1B | 0.1 | Fed | 2M1BDR12_Scan002 | 3043 | 2M1BDR12_P05 | 1.461 |
| 1A | 331 | 0.098 | 2M1B | 0.3 | Fed | 2M1BDR11_Scan001 | 3021 | 2M1BDR11_P01 | 2.333 |
| 1B | 266 | 0.272 | 2M1B | 0.3 | Fed | 2M1BDR11_Scan001 | 3028 | 2M1BDR11_P01 | 5.071 |
| 1C | 254 | 0.203 | 2M1B | 0.3 | Fed | 2M1BDR11_Scan001 | 3029 | 2M1BDR11_P01 | 4.228 |
| 1D | 322 | 0.264 | 2M1B | 0.3 | Fed | 2M1BDR11_Scan001 | 3049 | 2M1BDR11_P01 | 2.789 |
| 2A | 262 | -0.009 | 2M1B | 0.3 | Fed | 2M1BDR12_Scan001 | 3069 | 2M1BDR12_P02 | -0.526 |
| 2B | 289 | -0.079 | 2M1B | 0.3 | Fed | 2M1BDR12_Scan001 | 3063 | 2M1BDR12_P02 | -0.327 |
| 2C | 303 | -0.027 | 2M1B | 0.3 | Fed | 2M1BDR12_Scan001 | 3075 | 2M1BDR12_P02 | 0.331 |
| 2D | 337 | 0.128 | 2M1B | 0.3 | Fed | 2M1BDR12_Scan001 | 3068 | 2M1BDR12_P02 | 2.463 |
| 2A | 177 | 0.208 | 2M1B | 1 | Fed | 2M1BDR11_Scan002 | 3062 | 2M1BDR11_P06 | 4.420 |
| 2B | 185 | -0.111 | 2M1B | 1 | Fed | 2M1BDR11_Scan002 | 3077 | 2M1BDR11_P06 | -1.824 |
| 2C | 247 | 0.073 | 2M1B | 1 | Fed | 2M1BDR11_Scan002 | 3050 | 2M1BDR11_P06 | 2.002 |
| 2D | 242 | 0.046 | 2M1B | 1 | Fed | 2M1BDR11_Scan002 | 3067 | 2M1BDR11_P06 | 2.073 |
| 3A | 274 | 0.228 | 2M1B | 1 | Fed | 2M1BDR12_Scan001 | 3074 | 2M1BDR12_P03 | 3.799 |
| 3B | 262 | 0.179 | 2M1B | 1 | Fed | 2M1BDR12_Scan001 | 3085 | 2M1BDR12_P03 | 2.876 |
| 3C | 361 | 0.106 | 2M1B | 1 | Fed | 2M1BDR12_Scan001 | 3064 | 2M1BDR12_P03 | 2.008 |
| 3D | 284 | 0.042 | 2M1B | 1 | Fed | 2M1BDR12_Scan001 | 3067 | 2M1BDR12_P03 | 1.265 |
| 3B | 188 | 0.232 | 2M1B | 3 | Fed | 2M1BDR01_Scan001 | 3070 | 2M1BDR01_P03 | 4.186 |
| 1B | 243 | 0.290 | 2M1B | 3 | Fed | 2M1BDR04_Scan002 | 3013 | 2M1BDR04_P01 | 6.139 |
| 3B | 198 | 0.337 | 2M1B | 3 | Fed | 2M1BDR04_Scan002 | 3058 | 2M1BDR04_P03 | 5.727 |
| 1A | 184 | 0.470 | 2M1B | 3 | Fed | 2M1BDR05_Scan001 | 3064 | 2M1BDR05_P01 | 7.555 |
| 1C | 193 | 0.532 | 2M1B | 3 | Fed | 2M1BDR05_Scan001 | 3046 | 2M1BDR05_P01 | 9.262 |
| 1A | 358 | 0.455 | 2M1B | 3 | Fed | 2M1BDR10_Scan001 | 3043 | 2M1BDR10_P01 | 7.223 |
| 1B | 436 | 0.380 | 2M1B | 3 | Fed | 2M1BDR10_Scan001 | 3054 | 2M1BDR10_P01 | 6.439 |
| 1C | 375 | 0.416 | 2M1B | 3 | Fed | 2M1BDR10_Scan001 | 3053 | 2M1BDR10_P01 | 6.458 |
| 3C | 159 | 0.617 | 2M1B | 10 | Fed | 2M1BDR01_Scan001 | 3067 | 2M1BDR01_P03 | 10.545 |
| 1C | 235 | 0.411 | 2M1B | 10 | Fed | 2M1BDR04_Scan002 | 2981 | 2M1BDR04_P01 | 8.910 |
| 2C | 168 | 0.763 | 2M1B | 10 | Fed | 2M1BDR05_Scan001 | 3056 | 2M1BDR05_P02 | 14.387 |
| 1C | 158 | 0.617 | 2M1B | 10 | Fed | 2M1BDR07_Scan001 | 3023 | 2M1BDR07_P01 | 10.479 |
| 1D | 157 | 0.474 | 2M1B | 10 | Fed | 2M1BDR07_Scan001 | 3054 | 2M1BDR07_P01 | 8.672 |
| 1D | 320 | 0.551 | 2M1B | 10 | Fed | 2M1BDR10_Scan001 | 3044 | 2M1BDR10_P01 | 9.171 |
| 2A | 362 | 0.636 | 2M1B | 10 | Fed | 2M1BDR10_Scan001 | 3070 | 2M1BDR10_P02 | 10.309 |
| 2B | 411 | 0.561 | 2M1B | 10 | Fed | 2M1BDR10_Scan001 | 3069 | 2M1BDR10_P02 | 8.764 |
| 3D | 172 | 0.654 | 2M1B | 30 | Fed | 2M1BDR01_Scan001 | 3063 | 2M1BDR01_P03 | 10.929 |
| 1D | 234 | 0.533 | 2M1B | 30 | Fed | 2M1BDR04_Scan002 | 2981 | 2M1BDR04_P01 | 9.225 |
| 3D | 167 | 0.673 | 2M1B | 30 | Fed | 2M1BDR04_Scan002 | 3052 | 2M1BDR04_P03 | 10.584 |
| 2C | 318 | 0.727 | 2M1B | 30 | Fed | 2M1BDR10_Scan001 | 3070 | 2M1BDR10_P02 | 11.623 |

| | | | | | | | | | |
|---|---|---|---|---|---|---|---|---|---|
| 2D | 277 | 0.674 | 2M1B | 30 | Fed | 2M1BDR10_Scan001 | 3069 | 2M1BDR10_P02 | 11.065 |
| 3A | 338 | 0.579 | 2M1B | 30 | Fed | 2M1BDR10_Scan001 | 3065 | 2M1BDR10_P03 | 8.989 |
| 3B | 345 | 0.663 | 2M1B | 30 | Fed | 2M1BDR10_Scan001 | 3082 | 2M1BDR10_P03 | 10.026 |
| 3C | 277 | 0.741 | 2M1B | 30 | Fed | 2M1BDR10_Scan001 | 3079 | 2M1BDR10_P03 | 11.271 |
| 4A | 164 | 0.709 | 2M1B | 100 | Fed | 2M1BDR04_Scan002 | 3053 | 2M1BDR04_P04 | 13.375 |
| 3A | 182 | 0.873 | 2M1B | 100 | Fed | 2M1BDR05_Scan001 | 3065 | 2M1BDR05_P03 | 15.106 |
| 3B | 168 | 0.802 | 2M1B | 100 | Fed | 2M1BDR05_Scan001 | 3062 | 2M1BDR05_P03 | 14.817 |
| 3C | 197 | 0.916 | 2M1B | 100 | Fed | 2M1BDR05_Scan001 | 3083 | 2M1BDR05_P03 | 15.877 |
| 4A | 268 | 0.777 | 2M1B | 100 | Fed | 2M1BDR10_Scan001 | 3017 | 2M1BDR10_P04 | 13.411 |
| 4B | 271 | 0.744 | 2M1B | 100 | Fed | 2M1BDR10_Scan001 | 3034 | 2M1BDR10_P04 | 11.740 |
| 4C | 282 | 0.786 | 2M1B | 100 | Fed | 2M1BDR10_Scan001 | 3053 | 2M1BDR10_P04 | 12.686 |
| 4D | 209 | 0.916 | 2M1B | 100 | Fed | 2M1BDR10_Scan001 | 3040 | 2M1BDR10_P04 | 16.075 |
| 2B | 183 | 0.857 | 2M1B | 300 | Fed | 2M1BDR04_Scan002 | 3017 | 2M1BDR04_P02 | 14.685 |
| 4C | 156 | 0.839 | 2M1B | 300 | Fed | 2M1BDR05_Scan001 | 3033 | 2M1BDR05_P04 | 15.347 |
| 4A | 154 | 0.900 | 2M1B | 300 | Fed | 2M1BDR08_Scan001 | 3049 | 2M1BDR08_P04 | 15.380 |
| 1B | 170 | 0.899 | 2M1B | 300 | Fed | 2M1BDR08_Scan002 | 3033 | 2M1BDR08_P05 | 15.785 |
| 1C | 183 | 0.804 | 2M1B | 300 | Fed | 2M1BDR08_Scan002 | 3027 | 2M1BDR08_P05 | 14.338 |
| 1D | 197 | 0.831 | 2M1B | 300 | Fed | 2M1BDR08_Scan002 | 3020 | 2M1BDR08_P05 | 14.842 |
| 1A | 269 | 0.884 | 2M1B | 300 | Fed | 2M1BDR10_Scan002 | 3044 | 2M1BDR10_P05 | 15.034 |
| 1B | 290 | 0.945 | 2M1B | 300 | Fed | 2M1BDR10_Scan002 | 3055 | 2M1BDR10_P05 | 14.470 |
| 2C | 186 | 0.665 | 2M1B | 1000 | Fed | 2M1BDR02_Scan001 | 3068 | 2M1BDR02_P02 | 10.778 |
| 4C | 171 | 0.745 | 2M1B | 1000 | Fed | 2M1BDR02_Scan001 | 3046 | 2M1BDR02_P04 | 11.766 |
| 2C | 158 | 0.807 | 2M1B | 1000 | Fed | 2M1BDR03_Scan001 | 3075 | 2M1BDR03_P02 | 14.469 |
| 2C | 183 | 0.791 | 2M1B | 1000 | Fed | 2M1BDR04_Scan002 | 3061 | 2M1BDR04_P02 | 14.278 |
| 3D | 189 | 0.878 | 2M1B | 1000 | Fed | 2M1BDR05_Scan001 | 3064 | 2M1BDR05_P03 | 16.138 |
| 4D | 167 | 0.811 | 2M1B | 1000 | Fed | 2M1BDR05_Scan001 | 3029 | 2M1BDR05_P04 | 14.893 |
| 1C | 280 | 0.865 | 2M1B | 1000 | Fed | 2M1BDR10_Scan002 | 3036 | 2M1BDR10_P05 | 14.401 |
| 1D | 250 | 0.864 | 2M1B | 1000 | Fed | 2M1BDR10_Scan002 | 3051 | 2M1BDR10_P05 | 13.304 |
| 4D | 170 | 0.537 | 2M1B | 3000 | Fed | 2M1BDR01_Scan001 | 3061 | 2M1BDR01_P04 | 9.386 |
| 4D | 196 | 0.598 | 2M1B | 3000 | Fed | 2M1BDR02_Scan001 | 3049 | 2M1BDR02_P04 | 8.854 |
| 2D | 163 | 0.535 | 2M1B | 3000 | Fed | 2M1BDR03_Scan001 | 3084 | 2M1BDR03_P02 | 7.224 |
| 4D | 169 | 0.575 | 2M1B | 3000 | Fed | 2M1BDR03_Scan001 | 3031 | 2M1BDR03_P04 | 11.272 |
| 2D | 188 | 0.780 | 2M1B | 3000 | Fed | 2M1BDR04_Scan002 | 3075 | 2M1BDR04_P02 | 13.321 |
| 4D | 168 | 0.500 | 2M1B | 3000 | Fed | 2M1BDR04_Scan002 | 2988 | 2M1BDR04_P04 | 9.299 |
| 2C | 289 | 0.764 | 2M1B | 3000 | Fed | 2M1BDR10_Scan002 | 3075 | 2M1BDR10_P06 | 12.055 |
| 2D | 271 | 0.764 | 2M1B | 3000 | Fed | 2M1BDR10_Scan002 | 3069 | 2M1BDR10_P06 | 13.019 |
| 1A | 454 | 0.036 | DMSO | Solvent | Starved | DRS01_Scan002 | 3024 | DRS01_P05 | 0.740 |
| 1B | 562 | 0.085 | DMSO | Solvent | Starved | DRS01_Scan002 | 3034 | DRS01_P05 | 1.334 |
| 1C | 426 | 0.052 | DMSO | Solvent | Starved | DRS01_Scan002 | 3031 | DRS01_P05 | 1.517 |

| | | | | | | | | | |
|---|---|---|---|---|---|---|---|---|---|
| 1D | 519 | -0.039 | DMSO | Solvent | Starved | DRS01_Scan002 | 3034 | DRS01_P05 | 0.219 |
| 3A | 485 | 0.031 | DMSO | Solvent | Starved | DRS07_Scan002 | 3061 | DRS07_P07 | 0.080 |
| 3B | 308 | 0.080 | DMSO | Solvent | Starved | DRS07_Scan002 | 3036 | DRS07_P07 | 1.802 |
| 3C | 308 | -0.190 | DMSO | Solvent | Starved | DRS07_Scan002 | 3068 | DRS07_P07 | -2.353 |
| 3D | 286 | -0.012 | Solvent | Solvent | Starved | DRS07_Scan002 | 3068 | DRS07_P07 | 0.104 |
| 4A | 407 | 0.056 | 2M1B | 0.003 | Starved | DRS02_Scan001 | 3053 | DRS02_P04 | 1.140 |
| 4B | 523 | 0.087 | 2M1B | 0.003 | Starved | DRS02_Scan001 | 3042 | DRS02_P04 | 1.649 |
| 4C | 450 | 0.118 | 2M1B | 0.003 | Starved | DRS02_Scan001 | 3054 | DRS02_P04 | 2.390 |
| 4A | 455 | 0.078 | 2M1B | 0.003 | Starved | DRS05_Scan001 | 3043 | DRS05_P04 | 1.627 |
| 4B | 348 | -0.031 | 2M1B | 0.003 | Starved | DRS05_Scan001 | 3052 | DRS05_P04 | 0.137 |
| 4C | 425 | -0.167 | 2M1B | 0.003 | Starved | DRS05_Scan001 | 3054 | DRS05_P04 | -1.248 |
| 4D | 301 | 0.000 | 2M1B | 0.003 | Starved | DRS05_Scan001 | 3059 | DRS05_P04 | 0.269 |
| 2C | 251 | -0.107 | 2M1B | 0.003 | Starved | DRS07_Scan001 | 3059 | DRS07_P02 | -0.836 |
| 3A | 369 | 0.045 | 2M1B | 0.01 | Starved | DRS01_Scan001 | 3054 | DRS01_P03 | 0.727 |
| 3B | 406 | 0.137 | 2M1B | 0.01 | Starved | DRS01_Scan001 | 3044 | DRS01_P03 | 2.571 |
| 3C | 423 | 0.055 | 2M1B | 0.01 | Starved | DRS01_Scan001 | 3042 | DRS01_P03 | 1.509 |
| 3D | 428 | 0.000 | 2M1B | 0.01 | Starved | DRS01_Scan001 | 3033 | DRS01_P03 | 0.219 |
| 3A | 194 | 0.186 | 2M1B | 0.01 | Starved | DRS04_Scan001 | 3063 | DRS04_P03 | 3.132 |
| 3D | 160 | 0.054 | 2M1B | 0.01 | Starved | DRS04_Scan001 | 3062 | DRS04_P03 | 0.539 |
| 2A | 205 | 0.096 | 2M1B | 0.01 | Starved | DRS07_Scan001 | 3052 | DRS07_P02 | 0.523 |
| 2B | 231 | 0.188 | 2M1B | 0.01 | Starved | DRS07_Scan001 | 3041 | DRS07_P02 | 3.839 |
| 4A | 277 | -0.008 | 2M1B | 0.03 | Starved | DRS03_Scan001 | 3062 | DRS03_P04 | -0.669 |
| 4B | 203 | -0.244 | 2M1B | 0.03 | Starved | DRS03_Scan001 | 3035 | DRS03_P04 | -2.331 |
| 4C | 213 | -0.130 | 2M1B | 0.03 | Starved | DRS03_Scan001 | 3036 | DRS03_P04 | -1.732 |
| 4D | 302 | -0.014 | 2M1B | 0.03 | Starved | DRS03_Scan001 | 3059 | DRS03_P04 | -0.368 |
| 4A | 350 | -0.011 | 2M1B | 0.03 | Starved | DRS06_Scan001 | 3062 | DRS06_P04 | -0.503 |
| 4B | 315 | 0.113 | 2M1B | 0.03 | Starved | DRS06_Scan001 | 3050 | DRS06_P04 | 2.062 |
| 4C | 287 | -0.090 | 2M1B | 0.03 | Starved | DRS06_Scan001 | 3045 | DRS06_P04 | -1.044 |
| 4D | 360 | -0.056 | 2M1B | 0.03 | Starved | DRS06_Scan001 | 3059 | DRS06_P04 | -0.994 |
| 1A | 499 | 0.125 | 2M1B | 0.1 | Starved | DRS02_Scan001 | 3010 | DRS02_P01 | 2.092 |
| 1B | 372 | 0.107 | 2M1B | 0.1 | Starved | DRS02_Scan001 | 3028 | DRS02_P01 | 2.581 |
| 1C | 350 | 0.180 | 2M1B | 0.1 | Starved | DRS02_Scan001 | 3043 | DRS02_P01 | 2.766 |
| 1D | 402 | 0.175 | 2M1B | 0.1 | Starved | DRS02_Scan001 | 3033 | DRS02_P01 | 2.778 |
| 1A | 570 | 0.014 | 2M1B | 0.1 | Starved | DRS05_Scan001 | 3053 | DRS05_P01 | 0.121 |
| 1B | 456 | -0.122 | 2M1B | 0.1 | Starved | DRS05_Scan001 | 3062 | DRS05_P01 | -1.842 |
| 1C | 433 | -0.091 | 2M1B | 0.1 | Starved | DRS05_Scan001 | 3047 | DRS05_P01 | -1.913 |
| 1D | 411 | -0.091 | 2M1B | 0.1 | Starved | DRS05_Scan001 | 3055 | DRS05_P01 | -2.192 |
| 4A | 405 | 0.060 | 2M1B | 0.3 | Starved | DRS01_Scan002 | 3044 | DRS01_P08 | 1.261 |
| 4B | 347 | 0.205 | 2M1B | 0.3 | Starved | DRS01_Scan002 | 3042 | DRS01_P08 | 4.101 |

| | | | | | | | | | |
|---|---|---|---|---|---|---|---|---|---|
| 4C | 387 | 0.109 | 2M1B | 0.3 | Starved | DRS01_Scan002 | 3045 | DRS01_P08 | 2.501 |
| 4D | 489 | 0.167 | 2M1B | 0.3 | Starved | DRS01_Scan002 | 3045 | DRS01_P08 | 2.830 |
| 4A | 207 | 0.176 | 2M1B | 0.3 | Starved | DRS04_Scan002 | 3042 | DRS04_P08 | 3.637 |
| 4A | 333 | -0.086 | 2M1B | 0.3 | Starved | DRS07_Scan001 | 3059 | DRS07_P04 | -0.090 |
| 4B | 298 | -0.077 | 2M1B | 0.3 | Starved | DRS07_Scan001 | 3049 | DRS07_P04 | -0.247 |
| 4C | 303 | 0.029 | 2M1B | 0.3 | Starved | DRS07_Scan001 | 3058 | DRS07_P04 | 0.231 |
| 2A | 291 | 0.043 | 2M1B | 1 | Starved | DRS03_Scan002 | 3058 | DRS03_P05 | 1.547 |
| 2B | 210 | -0.026 | 2M1B | 1 | Starved | DRS03_Scan002 | 3065 | DRS03_P05 | -0.363 |
| 2C | 225 | -0.024 | 2M1B | 1 | Starved | DRS03_Scan002 | 3065 | DRS03_P05 | 0.096 |
| 2D | 211 | -0.061 | 2M1B | 1 | Starved | DRS03_Scan002 | 3063 | DRS03_P05 | -1.798 |
| 2A | 427 | 0.003 | 2M1B | 1 | Starved | DRS06_Scan002 | 3080 | DRS06_P05 | -0.698 |
| 2B | 335 | -0.100 | 2M1B | 1 | Starved | DRS06_Scan002 | 3071 | DRS06_P05 | -1.924 |
| 2C | 423 | -0.023 | 2M1B | 1 | Starved | DRS06_Scan002 | 3070 | DRS06_P05 | 0.302 |
| 2D | 378 | 0.072 | 2M1B | 1 | Starved | DRS06_Scan002 | 3071 | DRS06_P05 | 1.254 |
| 3A | 340 | 0.106 | 2M1B | 3 | Starved | DRS03_Scan002 | 3083 | DRS03_P07 | 1.311 |
| 3B | 212 | -0.005 | 2M1B | 3 | Starved | DRS03_Scan002 | 3074 | DRS03_P07 | -0.651 |
| 3C | 204 | 0.043 | 2M1B | 3 | Starved | DRS03_Scan002 | 3080 | DRS03_P07 | 0.915 |
| 3D | 198 | -0.006 | 2M1B | 3 | Starved | DRS03_Scan002 | 3088 | DRS03_P07 | -0.503 |
| 3A | 277 | -0.085 | 2M1B | 3 | Starved | DRS06_Scan002 | 3081 | DRS06_P07 | -0.268 |
| 3B | 272 | -0.118 | 2M1B | 3 | Starved | DRS06_Scan002 | 3061 | DRS06_P07 | -2.076 |
| 3C | 388 | -0.012 | 2M1B | 3 | Starved | DRS06_Scan002 | 3055 | DRS06_P07 | 0.545 |
| 3D | 399 | -0.003 | 2M1B | 3 | Starved | DRS06_Scan002 | 3072 | DRS06_P07 | 0.276 |
| 2A | 552 | 0.196 | 2M1B | 10 | Starved | DRS02_Scan001 | 3059 | DRS02_P02 | 3.188 |
| 2B | 576 | 0.222 | 2M1B | 10 | Starved | DRS02_Scan001 | 3065 | DRS02_P02 | 3.230 |
| 2C | 465 | 0.146 | 2M1B | 10 | Starved | DRS02_Scan001 | 3066 | DRS02_P02 | 2.236 |
| 2D | 338 | 0.154 | 2M1B | 10 | Starved | DRS02_Scan001 | 3083 | DRS02_P02 | 1.438 |
| 2A | 442 | 0.015 | 2M1B | 10 | Starved | DRS05_Scan001 | 3087 | DRS05_P02 | 0.464 |
| 2B | 417 | -0.043 | 2M1B | 10 | Starved | DRS05_Scan001 | 3059 | DRS05_P02 | -0.263 |
| 2C | 383 | 0.030 | 2M1B | 10 | Starved | DRS05_Scan001 | 3072 | DRS05_P02 | 0.185 |
| 2D | 424 | -0.290 | 2M1B | 10 | Starved | DRS05_Scan001 | 3071 | DRS05_P02 | -3.125 |
| 3A | 525 | 0.039 | 2M1B | 30 | Starved | DRS02_Scan002 | 3081 | DRS02_P07 | 0.723 |
| 3B | 391 | 0.078 | 2M1B | 30 | Starved | DRS02_Scan002 | 3076 | DRS02_P07 | 1.582 |
| 3C | 352 | -0.144 | 2M1B | 30 | Starved | DRS02_Scan002 | 3109 | DRS02_P07 | -2.057 |
| 3A | 418 | -0.196 | 2M1B | 30 | Starved | DRS05_Scan002 | 3064 | DRS05_P07 | -2.755 |
| 3B | 502 | 0.049 | 2M1B | 30 | Starved | DRS05_Scan002 | 3074 | DRS05_P07 | 0.924 |
| 3C | 379 | -0.032 | 2M1B | 30 | Starved | DRS05_Scan002 | 3055 | DRS05_P07 | 0.203 |
| 3D | 343 | 0.020 | 2M1B | 30 | Starved | DRS05_Scan002 | 3087 | DRS05_P07 | 0.325 |
| 2B | 291 | -0.143 | 2M1B | 30 | Starved | DRS07_Scan002 | 3067 | DRS07_P05 | -0.642 |
| 1A | 481 | 0.111 | 2M1B | 100 | Starved | DRS01_Scan001 | 3051 | DRS01_P01 | 3.003 |

| | | | | | | | | | |
|---|---|---|---|---|---|---|---|---|---|
| 1B | 349 | 0.107 | 2M1B | 100 | Starved | DRS01_Scan001 | 3041 | DRS01_P01 | 2.638 |
| 1C | 359 | 0.113 | 2M1B | 100 | Starved | DRS01_Scan001 | 3052 | DRS01_P01 | 2.691 |
| 1D | 354 | 0.154 | 2M1B | 100 | Starved | DRS01_Scan001 | 3048 | DRS01_P01 | 2.841 |
| 1A | 161 | 0.382 | 2M1B | 100 | Starved | DRS04_Scan001 | 3057 | DRS04_P01 | 7.617 |
| 1D | 166 | 0.558 | 2M1B | 100 | Starved | DRS04_Scan001 | 3053 | DRS04_P01 | 7.304 |
| 1A | 277 | -0.102 | 2M1B | 100 | Starved | DRS07_Scan001 | 3017 | DRS07_P01 | -0.098 |
| 1B | 333 | 0.029 | 2M1B | 100 | Starved | DRS07_Scan001 | 3015 | DRS07_P01 | 2.590 |
| 2A | 339 | 0.256 | 2M1B | 300 | Starved | DRS01_Scan002 | 3065 | DRS01_P06 | 4.433 |
| 2B | 299 | 0.289 | 2M1B | 300 | Starved | DRS01_Scan002 | 3051 | DRS01_P06 | 5.185 |
| 2C | 285 | 0.287 | 2M1B | 300 | Starved | DRS01_Scan002 | 3065 | DRS01_P06 | 5.375 |
| 2D | 376 | 0.199 | 2M1B | 300 | Starved | DRS01_Scan002 | 3070 | DRS01_P06 | 4.207 |
| 1A | 178 | -0.046 | 2M1B | 300 | Starved | DRS04_Scan002 | 3047 | DRS04_P06 | 0.602 |
| 1B | 163 | -0.244 | 2M1B | 300 | Starved | DRS04_Scan002 | 3042 | DRS04_P06 | -1.657 |
| 1D | 174 | -0.008 | 2M1B | 300 | Starved | DRS04_Scan002 | 3051 | DRS04_P06 | 0.231 |
| 3A | 267 | 0.350 | 2M1B | 300 | Starved | DRS07_Scan001 | 3054 | DRS07_P03 | 8.091 |
| 3A | 224 | 0.656 | 2M1B | 1000 | Starved | DRS03_Scan001 | 3066 | DRS03_P03 | 12.384 |
| 3B | 179 | 0.708 | 2M1B | 1000 | Starved | DRS03_Scan001 | 3065 | DRS03_P03 | 15.152 |
| 3C | 207 | 0.763 | 2M1B | 1000 | Starved | DRS03_Scan001 | 3055 | DRS03_P03 | 15.269 |
| 3D | 216 | 0.748 | 2M1B | 1000 | Starved | DRS03_Scan001 | 3059 | DRS03_P03 | 14.360 |
| 3A | 413 | 0.276 | 2M1B | 1000 | Starved | DRS06_Scan001 | 3065 | DRS06_P03 | 4.914 |
| 3B | 262 | 0.306 | 2M1B | 1000 | Starved | DRS06_Scan001 | 3066 | DRS06_P03 | 6.593 |
| 3C | 313 | 0.369 | 2M1B | 1000 | Starved | DRS06_Scan001 | 3078 | DRS06_P03 | 7.351 |
| 3D | 318 | 0.204 | 2M1B | 1000 | Starved | DRS06_Scan001 | 3070 | DRS06_P03 | 4.073 |
| 2A | 480 | 0.562 | 2M1B | 3000 | Starved | DRS02_Scan002 | 3054 | DRS02_P06 | 9.512 |
| 2B | 398 | 0.630 | 2M1B | 3000 | Starved | DRS02_Scan002 | 3070 | DRS02_P06 | 10.955 |
| 2C | 377 | 0.634 | 2M1B | 3000 | Starved | DRS02_Scan002 | 3087 | DRS02_P06 | 10.360 |
| 1A | 390 | 0.101 | 2M1B | 3000 | Starved | DRS05_Scan002 | 3059 | DRS05_P06 | 1.589 |
| 1B | 471 | 0.165 | 2M1B | 3000 | Starved | DRS05_Scan002 | 3042 | DRS05_P06 | 2.574 |
| 1C | 353 | 0.106 | 2M1B | 3000 | Starved | DRS05_Scan002 | 3048 | DRS05_P06 | 1.618 |
| 1D | 427 | 0.008 | 2M1B | 3000 | Starved | DRS05_Scan002 | 3061 | DRS05_P06 | 0.822 |
| 2A | 384 | -0.124 | 2M1B | 3000 | Starved | DRS07_Scan002 | 3072 | DRS07_P05 | -1.040 |

**FIGURE-3A-2M1B-FED-WORM-POSITIONS (UNITS = mm)**

| DMSO | MB D01 | MB D02 | MB D03 | MB D04 | MB D05 | MB D06 | MB D07 | MB D08 | MB D09 | MB D10 | MB D11 | MB D12 | MB D13 |
|---|---|---|---|---|---|---|---|---|---|---|---|---|---|
| -3.286 | 4.440 | -5.943 | 9.727 | 11.349 | 10.543 | 6.145 | 23.385 | 18.803 | 6.829 | -13.658 | 8.333 | -15.025 | 0.183 |
| 18.615 | 25.121 | 21.754 | 3.903 | 2.756 | -8.041 | -1.251 | 16.371 | 17.334 | 1.604 | -9.545 | 25.917 | 4.488 | 22.860 |
| 16.387 | 23.785 | -8.124 | -2.305 | -18.741 | 20.129 | 20.537 | 25.001 | 3.367 | -2.788 | 24.464 | 1.879 | 5.757 | 22.308 |
| 10.590 | 3.760 | 25.825 | -9.843 | -19.142 | 21.207 | -7.976 | 19.741 | 16.990 | 5.857 | 20.623 | 20.468 | 7.702 | -2.864 |
| 1.023 | -4.731 | 17.811 | 8.655 | 15.343 | 9.787 | -20.628 | -4.186 | 2.777 | -7.114 | 18.875 | 16.462 | 13.394 | -10.190 |
| -0.849 | -24.468 | -8.764 | -5.889 | 17.620 | -23.470 | -21.450 | -8.299 | 4.870 | 6.481 | 12.591 | 11.121 | -22.869 | -19.425 |
| 24.704 | 9.382 | 19.835 | 14.651 | 7.005 | 1.815 | 6.634 | 23.348 | 7.626 | -0.235 | 17.573 | -21.870 | -0.777 | 25.188 |
| 0.115 | -5.584 | -10.434 | 0.074 | 23.682 | -17.877 | 1.386 | 21.410 | 7.916 | -1.673 | -12.216 | 25.827 | -5.345 | -21.253 |
| -6.850 | 10.481 | -10.971 | -16.816 | -3.186 | 10.876 | 12.821 | 7.065 | 20.254 | 16.828 | -13.359 | 11.578 | 21.114 | 18.647 |
| -0.028 | -15.960 | 4.428 | -7.415 | -8.902 | -24.132 | 11.799 | -11.535 | 15.716 | 5.630 | 15.338 | 12.192 | 17.576 | -7.965 |
| 24.734 | -24.929 | -17.910 | 11.326 | -0.006 | 20.102 | -5.725 | -21.439 | 1.867 | 19.654 | -21.266 | 20.331 | 16.426 | 5.542 |
| 20.508 | -8.486 | 22.808 | 25.303 | -7.487 | -13.825 | -4.071 | 7.193 | 6.920 | -11.777 | 20.052 | -0.560 | -14.684 | -1.528 |
| 21.989 | 13.906 | 20.871 | -12.870 | -22.517 | -9.045 | 24.744 | 1.802 | 16.145 | 9.478 | 1.806 | 8.967 | 12.960 | -6.507 |
| 24.988 | 0.545 | -8.386 | 12.324 | -4.819 | 15.119 | 4.153 | -18.977 | 14.131 | 8.515 | 25.831 | 14.767 | 19.224 | 3.425 |
| -7.067 | 22.655 | -15.669 | -25.009 | -1.811 | -20.214 | -7.050 | -3.835 | 19.420 | 16.846 | 14.020 | 17.789 | 20.598 | 19.468 |
| 4.793 | 4.542 | 2.744 | -15.666 | 25.237 | 23.218 | 23.594 | 19.750 | 18.170 | 25.127 | 13.300 | 21.326 | 3.390 | -4.804 |
| 17.791 | -13.103 | 0.734 | 9.313 | 19.146 | -16.697 | 8.600 | 6.261 | 5.915 | 15.118 | 5.287 | 20.614 | 12.647 | 9.965 |
| 10.651 | -6.062 | -12.098 | -16.640 | -12.728 | 15.990 | -19.124 | -22.791 | 2.451 | 19.233 | 10.100 | 25.510 | 13.467 | 3.135 |
| 14.057 | 2.641 | -17.059 | -24.050 | -25.562 | 9.525 | 19.895 | -3.825 | -1.687 | 2.426 | 21.128 | 5.219 | -7.030 | 4.344 |
| 26.397 | -18.355 | -13.753 | -9.246 | -17.098 | -24.054 | -10.313 | -9.310 | 25.114 | 1.071 | 20.179 | 22.208 | 4.587 | -0.184 |
| 16.474 | 12.822 | -24.915 | 27.213 | 22.539 | -9.802 | 23.644 | -16.371 | 24.702 | 8.212 | 23.033 | 18.755 | 20.144 | -15.444 |
| -11.278 | 10.963 | -9.576 | 16.762 | -11.385 | -0.157 | -8.719 | 12.810 | -7.278 | 21.801 | 23.826 | 1.734 | 23.476 | 18.910 |
| 3.665 | -7.798 | -6.817 | -0.889 | -23.960 | 2.293 | 4.222 | 18.097 | 25.563 | -9.177 | -6.676 | 19.336 | -6.081 | 9.032 |
| 5.481 | 25.487 | -8.368 | 23.932 | -26.887 | -14.745 | 3.713 | 6.929 | 8.718 | 8.891 | -11.976 | 19.654 | -17.078 | -16.886 |
| 19.293 | 5.130 | 3.204 | -11.233 | -25.244 | 15.014 | -8.641 | -24.516 | 18.435 | 21.415 | 24.012 | 12.281 | 22.905 | 20.092 |
| -5.947 | 19.154 | 21.566 | -22.223 | 21.821 | -6.066 | 2.219 | -3.139 | -5.146 | 15.814 | -3.450 | 26.136 | 15.212 | 2.226 |
| 7.795 | 3.426 | 18.594 | 9.253 | -10.261 | -12.565 | -14.189 | 19.958 | 20.061 | 20.736 | 12.999 | -22.716 | 26.221 | 13.407 |
| 13.582 | -16.276 | -2.897 | 27.067 | 1.023 | 24.977 | 13.565 | -2.534 | -10.805 | 22.332 | 11.153 | 7.032 | 24.558 | 1.528 |
| 24.182 | 25.215 | 25.543 | 17.949 | -15.898 | -17.861 | 26.516 | 3.831 | -17.294 | 19.376 | 24.601 | 15.894 | 16.915 | 17.263 |
| 9.435 | 21.466 | 9.785 | -19.041 | 11.358 | 9.357 | 20.193 | -5.658 | 24.176 | 10.858 | 12.259 | 12.999 | 25.615 | 22.961 |
| 28.496 | -0.463 | 5.862 | 20.352 | -17.561 | 23.348 | 19.536 | 1.256 | 5.827 | 8.343 | 8.412 | 21.679 | -2.662 | 20.968 |
| 10.606 | 17.484 | -18.864 | 15.400 | 21.211 | -25.397 | -10.682 | 11.756 | 18.741 | -15.556 | -10.422 | 22.243 | 20.615 | 17.380 |
| 27.251 | -15.284 | 20.881 | -2.472 | 10.820 | -8.113 | -9.658 | 25.860 | 1.338 | 25.800 | 18.859 | 5.896 | 12.087 | 16.919 |
| 22.083 | 4.573 | 4.877 | 23.136 | -10.089 | 20.926 | 28.258 | 18.552 | 9.075 | 7.370 | 1.448 | 18.954 | 2.923 | 12.238 |
| 11.881 | 7.056 | -25.984 | -21.657 | -2.539 | -2.132 | 2.729 | 27.694 | 0.173 | 0.352 | -0.492 | 23.166 | 13.331 | -6.104 |

| | | | | | | | | | | | | | |
|---|---|---|---|---|---|---|---|---|---|---|---|---|---|
| -21.721 | -19.542 | -12.553 | -8.314 | -23.664 | -18.930 | 16.494 | -9.381 | 23.968 | 13.805 | 24.927 | 14.179 | 17.761 | 24.917 |
| 7.629 | -2.888 | -13.451 | -0.661 | 17.781 | 2.862 | 19.064 | 4.047 | 16.670 | -4.350 | 11.843 | -12.300 | -3.988 | -8.807 |
| 23.586 | 19.501 | 10.852 | 10.677 | 19.821 | 0.141 | 25.498 | -16.451 | 25.707 | 15.232 | 10.484 | 8.551 | -13.883 | 7.894 |
| -27.826 | -17.773 | 20.744 | 21.466 | -20.920 | -22.430 | 13.141 | -10.306 | 5.551 | 22.823 | 18.858 | 12.123 | 24.277 | 23.341 |
| -2.413 | -24.475 | 16.649 | -2.641 | -27.668 | 24.382 | 26.576 | -25.514 | -26.298 | 14.807 | 19.519 | 3.775 | 1.334 | 10.143 |
| 5.412 | 9.136 | 12.464 | 22.009 | 24.541 | 4.809 | 22.352 | 5.949 | 22.940 | -1.717 | 28.323 | 25.590 | 8.440 | -4.811 |
| 28.621 | 13.281 | 27.894 | -19.265 | -1.906 | -19.975 | 20.942 | 16.471 | -11.785 | 8.571 | 26.936 | 26.728 | -28.490 | 15.375 |
| 20.444 | 8.450 | 8.880 | 5.134 | -11.976 | -5.421 | 19.743 | -6.032 | 11.143 | 17.526 | 6.790 | 19.525 | 6.043 | -9.265 |
| -19.455 | 18.249 | 18.769 | -6.562 | 15.089 | -21.581 | -18.442 | -14.624 | -21.760 | 24.233 | 17.706 | 22.215 | 5.185 | -3.243 |
| 3.866 | -10.329 | 21.426 | 1.112 | -18.127 | 16.298 | -16.494 | -7.519 | 7.117 | 20.911 | 23.778 | 23.144 | 9.101 | 16.221 |
| 16.625 | -15.441 | 14.937 | 2.272 | 17.059 | -12.424 | 27.641 | 7.231 | -7.370 | 19.105 | 22.357 | 24.477 | 25.646 | 26.793 |
| 10.269 | 11.086 | -2.251 | 26.951 | 5.522 | 25.698 | -26.988 | -9.075 | 8.760 | 6.209 | 7.564 | 16.352 | 13.158 | 20.940 |
| 29.359 | -16.751 | -11.339 | 6.542 | -23.282 | 0.306 | 28.485 | -12.101 | 10.561 | -9.604 | 18.557 | 21.043 | 4.420 | 5.884 |
| 27.407 | -9.781 | 2.425 | 21.059 | 21.740 | -2.848 | -30.055 | 14.115 | 22.076 | 15.138 | 25.302 | 17.513 | 24.616 | 8.897 |
| 26.477 | 16.484 | -21.046 | 24.881 | 24.236 | -26.727 | 17.928 | -1.614 | 10.105 | 11.824 | 15.353 | 21.751 | 25.063 | -1.333 |
| 15.576 | 20.082 | -27.725 | -27.429 | -6.883 | -23.408 | 4.569 | -0.526 | -5.790 | 21.587 | 27.160 | 22.223 | 13.299 | 11.342 |
| -4.721 | -2.184 | -4.926 | -3.645 | -5.428 | 18.812 | -26.181 | -21.143 | 21.326 | 18.456 | 26.616 | 27.128 | 7.342 | -13.672 |
| -18.691 | 11.959 | -28.113 | 28.173 | -0.360 | -3.680 | -6.742 | 20.525 | 24.153 | -12.642 | 15.731 | 11.636 | 28.689 | -0.134 |
| -13.153 | 4.102 | -21.946 | 2.182 | 26.357 | -19.496 | 27.407 | 23.707 | -14.287 | -4.328 | 16.168 | 18.073 | 18.113 | 23.524 |
| 18.503 | 8.356 | 16.555 | -25.393 | -1.538 | 21.991 | 25.835 | 29.162 | 3.519 | -28.139 | 8.051 | 22.800 | 9.931 | 15.627 |
| -25.980 | -12.524 | 15.129 | -4.555 | 2.407 | 16.402 | 16.922 | 16.894 | 0.427 | 5.144 | 13.986 | 17.687 | 27.289 | 27.910 |
| 7.619 | 21.876 | 26.863 | 28.764 | -23.507 | -18.762 | 8.979 | 27.758 | -4.209 | -0.125 | 26.524 | 18.677 | 18.828 | 18.299 |
| -8.699 | 19.507 | 20.118 | -7.761 | 13.728 | -25.834 | 21.702 | -7.149 | 11.250 | 13.701 | 20.297 | 12.616 | 14.771 | 25.458 |
| -1.083 | -19.309 | 2.351 | -13.098 | 16.907 | 22.774 | 1.384 | 24.251 | 24.556 | 1.007 | 17.016 | 21.970 | 10.385 | 22.216 |
| 22.326 | 13.001 | 19.328 | 6.902 | 21.372 | 17.973 | 5.786 | -6.672 | -7.187 | 14.955 | 23.991 | 27.885 | 19.413 | 16.586 |
| 11.463 | -12.251 | -7.628 | 26.185 | 22.532 | 25.145 | 29.306 | 3.732 | 29.844 | 9.405 | 12.721 | 24.351 | 23.012 | -2.706 |
| 6.471 | 11.013 | 24.541 | -18.665 | -9.504 | -15.508 | 25.216 | 9.962 | 3.564 | 25.467 | 21.701 | -25.405 | 12.021 | 5.511 |
| 24.498 | 25.545 | 23.933 | -17.906 | -17.751 | -27.301 | -29.933 | 18.766 | 28.094 | -27.681 | 13.671 | 19.909 | 28.115 | 17.462 |
| -9.693 | 20.088 | 2.186 | -28.982 | 16.165 | -0.151 | 21.013 | -15.913 | 18.732 | 24.955 | 18.776 | 25.746 | 20.528 | 4.227 |
| 22.927 | -8.772 | -20.569 | 21.041 | -8.776 | 6.593 | -3.005 | 6.042 | 15.821 | -3.756 | -18.759 | 26.898 | -26.485 | 23.090 |
| 3.736 | -29.840 | -3.359 | -19.999 | 21.939 | 1.720 | 17.636 | 19.191 | 16.984 | 16.445 | 14.963 | 17.297 | 11.107 | -7.547 |
| -7.488 | -5.775 | -3.894 | -8.673 | -2.359 | 2.623 | 17.008 | -5.214 | 5.395 | 11.421 | 10.608 | 10.841 | -2.885 | -9.147 |
| 27.807 | -12.337 | -9.438 | -16.341 | 20.786 | -7.493 | 11.913 | -27.023 | 23.902 | 15.056 | 22.479 | -1.276 | 5.874 | 22.496 |
| 5.786 | -18.794 | 25.362 | 17.122 | -11.700 | -1.685 | -28.244 | 25.980 | -6.898 | 10.783 | 18.763 | 20.673 | -13.701 | 21.296 |
| 5.478 | -19.125 | -27.975 | 20.728 | 1.207 | 4.701 | -4.617 | 27.785 | -3.371 | 18.817 | -19.656 | 30.584 | 8.062 | 27.192 |
| 1.452 | 12.806 | 12.060 | 1.643 | -13.757 | 26.533 | 17.841 | 22.069 | -7.809 | 19.588 | 23.069 | 25.064 | -13.058 | -12.142 |

| | | | | | | | | | | | | | |
|---|---|---|---|---|---|---|---|---|---|---|---|---|---|
| 13.444 | -7.951 | -27.435 | -24.834 | 9.239 | 21.355 | 16.636 | 28.909 | 12.452 | 18.113 | 8.664 | 7.644 | 18.826 | 13.954 |
| -24.807 | 22.992 | 17.532 | 29.926 | -8.316 | -11.712 | 27.054 | -6.350 | 27.960 | 23.856 | -8.863 | 7.841 | 22.784 | 29.038 |
| 16.403 | 16.390 | -20.827 | 12.712 | 26.957 | -4.800 | -11.308 | 11.538 | 9.269 | 23.327 | 20.850 | 22.259 | 26.895 | -5.187 |
| 25.088 | -10.298 | 26.513 | 2.711 | 2.352 | -15.029 | 25.124 | 7.754 | 13.642 | 25.029 | 19.867 | 23.957 | 29.934 | 9.784 |
| 4.384 | 4.874 | -3.504 | -0.014 | -6.545 | 3.478 | -1.882 | -15.485 | 7.666 | -18.910 | 23.806 | 12.769 | -10.876 | 11.528 |
| 2.405 | 19.295 | -28.868 | -29.178 | -23.443 | -10.780 | 0.387 | 15.862 | -5.949 | 5.699 | 9.453 | 31.054 | 19.132 | -15.232 |
| 27.085 | 26.469 | -4.255 | 29.902 | -28.955 | -24.004 | -29.490 | 8.489 | -18.752 | -8.932 | 0.135 | 23.927 | 21.941 | 15.465 |
| -13.371 | 11.404 | 25.461 | -3.192 | 21.385 | -1.729 | -31.208 | -9.513 | 17.603 | 9.531 | 26.130 | 18.141 | -17.470 | -22.316 |
| 10.325 | -14.428 | 24.154 | -29.905 | -27.507 | 25.082 | 15.550 | -7.867 | -11.663 | -8.400 | 24.543 | 30.628 | 27.466 | 12.591 |
| -0.530 | 20.685 | 22.714 | 0.375 | 28.560 | -5.493 | -23.970 | 10.716 | -15.618 | 20.385 | 8.391 | -20.851 | 25.790 | 25.685 |
| -9.741 | -15.963 | 7.102 | -21.596 | 1.351 | -25.975 | -20.488 | 19.480 | -15.224 | 10.599 | -6.733 | 1.819 | 8.791 | 20.056 |
| 19.722 | -18.595 | 18.069 | -18.948 | -18.624 | 23.019 | -23.690 | -6.805 | 12.458 | 29.188 | 20.380 | 11.929 | 1.636 | 13.020 |
| 16.658 | -27.237 | -16.505 | 24.723 | 10.779 | -7.440 | 27.906 | 16.786 | 25.202 | -25.102 | 7.992 | 20.263 | 15.090 | 26.533 |
| 1.282 | -1.720 | 26.173 | 13.993 | -19.842 | 21.638 | 28.731 | -1.520 | 27.595 | 24.438 | 29.166 | 11.059 | 15.911 | 27.877 |
| 14.070 | 14.224 | 22.819 | -19.507 | 25.206 | 13.150 | -2.570 | -13.553 | -9.416 | 9.993 | 24.224 | 26.702 | 12.902 | 6.252 |
| -5.130 | -20.753 | -26.266 | -27.970 | 12.855 | -25.243 | 13.202 | -3.519 | 10.502 | 9.084 | 18.253 | 23.568 | 19.391 | -17.477 |
| 9.008 | 24.742 | -7.822 | 0.578 | 17.646 | -3.000 | 20.797 | 22.979 | -14.616 | 21.832 | 15.890 | 31.467 | 20.923 | 28.572 |
| -4.574 | 6.937 | -9.078 | -14.519 | -2.454 | -29.070 | 29.063 | 29.133 | 17.228 | 25.300 | 0.430 | 15.080 | -1.573 | 23.630 |
| 24.546 | 16.205 | 11.302 | -2.671 | 19.661 | 27.258 | 26.286 | 7.484 | -1.424 | 18.084 | 28.615 | 28.803 | 23.822 | 10.993 |
| 16.212 | 13.908 | 14.944 | 29.461 | 10.602 | 9.229 | -4.654 | 4.847 | 22.118 | -0.702 | 28.008 | 25.958 | -18.484 | 21.759 |
| 23.778 | 8.678 | -7.056 | 3.704 | 22.929 | 5.521 | -11.793 | -12.447 | 20.707 | 19.995 | -0.711 | 8.553 | 28.561 | 9.766 |
| 18.879 | -7.967 | 5.720 | -9.781 | 6.102 | 10.264 | 8.668 | 20.764 | 18.552 | 6.730 | 23.021 | -23.418 | 19.232 | -8.165 |
| -23.597 | 16.995 | -18.730 | -4.895 | -5.349 | -27.502 | 23.448 | 27.444 | 13.627 | 15.923 | -9.634 | 21.891 | 28.005 | 14.298 |
| 23.010 | -28.865 | -27.368 | -24.050 | 21.464 | -19.857 | 10.452 | 15.448 | 28.274 | 11.714 | 20.430 | 10.142 | -21.492 | 31.194 |
| -9.380 | 2.281 | 31.524 | -12.363 | 24.516 | 25.549 | -28.549 | 4.174 | 23.837 | -21.098 | 28.932 | 24.217 | 2.796 | 30.568 |
| 20.957 | 20.251 | 30.023 | 30.481 | 11.010 | 22.632 | -25.747 | 16.248 | 27.499 | 4.301 | 28.403 | 25.068 | 28.710 | 25.237 |
| 11.436 | 21.726 | 28.327 | 15.509 | -24.861 | -9.110 | -30.975 | 9.582 | 23.221 | 30.759 | 28.771 | 21.010 | 11.942 | 27.595 |
| 7.889 | -17.205 | 32.014 | -16.123 | -10.098 | 4.969 | -14.412 | -5.111 | 24.056 | 29.863 | 24.803 | 14.511 | 13.768 | -16.189 |
| 17.006 | 14.107 | 25.564 | 25.822 | -29.145 | 27.632 | 19.712 | -8.417 | 28.503 | 11.228 | 7.320 | -1.992 | 11.641 | -2.029 |
| 17.760 | 20.395 | 13.282 | -24.937 | 8.310 | -4.828 | -5.596 | 14.685 | 5.065 | 25.198 | 9.136 | 3.324 | 21.739 | 16.602 |
| 5.267 | 7.132 | -23.591 | -26.357 | 18.311 | -14.129 | 25.839 | 5.399 | 21.062 | 28.219 | 19.336 | 31.103 | -27.198 | 21.587 |
| -29.200 | 12.678 | 4.911 | -16.639 | 28.799 | -6.171 | 7.400 | 4.197 | 26.245 | 13.194 | -7.069 | 18.898 | 27.129 | -9.186 |
| -3.576 | 7.883 | 9.545 | -25.446 | 26.744 | 24.919 | -6.615 | 21.068 | 10.165 | 16.384 | 25.795 | 17.939 | 19.305 | 13.407 |
| 18.450 | 27.890 | -21.624 | -14.395 | 14.618 | 13.127 | 24.524 | 10.736 | 26.939 | 29.117 | 23.283 | 6.980 | 5.177 | 20.195 |
| 4.045 | -19.898 | 31.093 | -31.149 | 9.903 | -20.945 | 19.234 | 22.918 | 16.243 | 10.913 | 20.865 | 9.924 | 5.790 | -28.015 |
| 5.949 | 31.333 | 3.940 | -25.533 | -18.015 | -17.991 | -5.080 | 6.165 | 26.462 | 31.546 | 9.793 | 22.325 | 25.206 | 11.529 |

| | | | | | | | | | | | | | |
|---|---|---|---|---|---|---|---|---|---|---|---|---|---|
| -8.497 | -28.474 | 26.234 | -7.469 | -22.018 | -25.869 | -1.225 | 29.414 | 19.255 | 19.514 | 13.631 | -0.047 | -21.258 | 18.039 |
| 1.789 | 17.893 | 14.594 | 0.840 | 26.134 | 17.302 | -3.117 | 25.459 | -4.670 | 17.992 | 27.645 | 23.226 | 23.194 | 24.009 |
| 29.348 | -7.827 | 9.694 | -27.755 | 0.370 | 21.927 | -11.283 | -1.954 | 23.131 | 28.053 | -3.270 | 17.476 | 6.123 | 11.261 |
| 15.152 | -23.953 | -23.321 | 14.369 | -20.630 | 19.516 | -16.298 | -10.680 | 25.104 | -10.793 | 3.252 | 21.625 | 15.751 | 24.218 |
| 22.854 | -14.731 | -28.461 | -25.994 | 20.453 | 11.211 | 23.740 | -18.538 | 16.147 | 30.071 | -10.432 | 25.826 | 11.906 | 29.722 |
| 4.598 | -2.498 | -18.647 | -12.645 | 12.892 | -10.834 | 28.119 | 19.043 | -0.895 | 22.097 | 24.736 | 11.731 | 14.985 | 16.095 |
| 18.852 | -5.840 | 19.625 | -29.394 | 27.920 | -17.245 | 17.649 | 7.699 | 18.841 | 8.325 | -12.859 | 3.116 | 19.966 | 14.053 |
| -12.155 | 30.109 | 10.257 | -28.190 | -16.641 | 20.225 | 5.887 | 19.591 | 26.257 | 18.490 | -9.494 | 18.752 | 10.799 | -5.637 |
| 0.271 | -11.822 | 11.070 | 13.248 | 24.148 | -22.426 | 8.289 | 0.125 | 10.623 | -14.568 | 26.577 | 10.660 | -12.949 | -19.512 |
| -6.329 | 6.456 | -4.395 | 25.754 | -29.442 | 6.365 | -20.468 | 10.156 | 4.279 | -7.667 | 0.592 | 11.069 | -13.739 | 25.940 |
| -22.458 | 21.200 | 27.816 | -9.154 | -18.567 | -19.509 | -3.951 | -30.004 | -7.884 | 19.018 | 21.665 | 24.417 | 27.950 | 2.157 |
| 27.223 | -14.349 | 3.383 | 14.513 | 28.797 | 1.137 | -21.732 | 5.552 | 24.806 | 29.286 | 25.604 | 17.540 | 6.487 | 27.982 |
| 26.751 | -4.772 | -1.925 | -20.306 | -2.625 | 16.378 | 3.355 | -17.602 | 23.182 | 3.518 | 27.580 | 7.187 | 16.283 | 6.349 |
| 9.327 | 3.830 | 16.753 | 4.272 | -12.153 | -27.435 | -5.589 | 3.602 | 17.971 | 10.743 | 15.477 | -22.710 | 13.292 | 30.004 |
| 22.552 | -0.269 | 23.532 | 29.888 | -23.603 | -8.509 | -25.338 | 14.443 | 21.974 | 2.170 | 14.665 | 21.552 | 14.522 | 14.374 |
| 16.895 | 21.447 | 4.577 | -19.348 | 24.914 | 11.348 | -4.482 | 21.451 | 19.357 | 14.604 | 22.843 | 23.671 | 12.329 | 19.679 |
| -23.876 | 2.377 | -8.510 | -2.887 | -9.270 | 7.149 | 22.793 | 23.009 | 11.706 | -4.886 | 21.652 | 17.121 | 7.846 | 18.643 |
| 9.581 | 18.509 | -11.022 | 0.604 | 8.590 | -19.840 | 5.911 | -6.999 | 2.495 | 8.246 | 7.842 | 8.670 | -13.567 | 22.177 |
| 18.271 | 15.975 | -14.848 | 28.147 | -5.731 | -28.880 | 27.449 | 16.901 | 3.248 | 29.848 | 20.126 | 12.328 | 23.520 | 12.030 |
| 3.841 | -22.005 | -26.651 | 11.081 | 18.446 | 22.765 | -10.607 | 17.459 | 14.818 | -30.512 | 27.418 | 26.551 | 22.246 | 11.392 |
| 7.394 | -13.835 | 25.285 | 0.947 | 13.795 | -23.351 | 3.033 | 10.255 | 17.861 | 12.440 | 26.073 | -28.088 | 3.265 | 21.050 |
| 15.999 | 16.056 | 15.811 | 9.261 | 6.662 | 19.761 | 4.703 | -22.029 | 4.408 | 21.742 | 15.056 | 15.884 | -9.957 | 4.396 |
| -6.506 | 10.832 | 1.052 | 3.769 | -11.711 | 16.847 | 24.601 | -17.491 | 13.683 | 23.879 | -6.959 | 25.242 | 6.939 | 21.503 |
| 21.254 | 7.683 | -25.823 | -4.069 | 25.800 | -20.730 | 26.242 | 26.038 | 10.422 | 6.448 | 26.954 | 19.401 | 24.493 | 11.105 |
| 8.040 | 23.383 | -0.115 | 18.769 | -16.712 | 0.998 | 7.939 | 19.952 | 22.136 | 22.963 | 14.942 | 27.313 | 5.855 | 6.123 |
| -9.357 | 29.744 | -7.198 | 5.070 | -3.097 | -10.232 | 3.821 | 11.855 | 6.738 | 30.176 | 4.972 | 25.887 | 10.871 | -4.613 |
| -15.605 | 22.515 | -28.454 | -23.487 | 15.908 | 26.038 | -2.132 | -15.510 | 9.372 | 9.223 | -9.758 | 23.197 | 14.024 | 24.343 |
| 10.778 | 23.267 | 23.546 | 24.889 | -12.717 | 5.287 | 23.103 | 14.159 | 15.351 | 16.615 | 25.773 | 24.995 | -13.133 | 4.691 |
| 15.648 | -7.537 | -17.605 | -22.467 | 5.893 | 31.330 | 25.506 | 2.023 | 7.712 | 26.011 | 24.789 | 12.591 | 28.944 | -3.062 |
| 6.083 | 12.507 | -14.662 | -10.175 | -20.440 | -15.302 | 9.417 | -1.233 | 4.843 | 19.462 | -9.440 | 18.470 | 25.580 | 15.076 |
| 12.182 | -27.050 | -24.893 | 10.182 | 30.241 | -6.431 | -4.492 | 20.308 | 21.103 | 15.878 | 26.189 | 17.776 | 21.565 | 6.403 |
| 21.745 | 10.310 | -9.578 | -17.880 | -18.190 | 29.190 | 4.327 | -4.347 | 24.603 | -19.796 | -9.285 | 22.018 | 17.736 | 28.985 |
| 3.619 | 20.707 | -11.448 | 15.822 | 13.731 | 27.483 | -13.468 | 6.530 | -10.266 | 5.250 | 14.941 | 4.249 | 15.058 | 19.220 |
| 2.330 | 2.680 | 11.990 | 11.338 | 0.939 | 8.672 | 12.217 | -26.119 | 11.669 | 2.161 | 1.208 | 14.230 | 5.018 | -4.147 |
| -1.793 | -17.025 | -17.916 | -8.636 | 21.890 | -19.242 | -17.943 | 9.961 | 22.845 | 26.730 | 15.638 | 24.228 | 27.049 | 22.350 |
| -4.683 | -27.700 | 10.637 | -17.070 | 24.665 | -5.768 | 14.344 | 14.688 | 24.972 | 18.486 | 11.336 | 20.874 | 0.926 | -19.031 |

| | | | | | | | | | | | | | |
|---|---|---|---|---|---|---|---|---|---|---|---|---|---|
| -2.104 | 27.933 | -20.648 | 7.079 | -29.500 | -21.621 | 20.180 | -8.647 | 9.911 | 7.655 | 12.354 | 10.097 | 9.810 | 3.904 |
| -20.724 | 8.219 | 22.282 | -7.842 | 18.152 | 15.091 | -1.168 | 8.345 | 14.401 | 29.140 | 22.955 | 16.187 | 7.713 | 17.390 |
| 14.554 | 18.992 | 23.206 | 5.644 | 26.823 | -17.522 | 19.332 | 27.410 | -15.436 | 24.180 | 6.414 | 13.552 | 6.049 | 10.788 |
| 26.019 | -21.905 | 27.869 | -15.018 | -13.855 | 8.345 | 24.784 | 2.763 | 21.627 | -2.045 | 10.020 | 18.160 | 10.594 | 12.666 |
| 17.171 | 17.324 | -7.739 | -22.003 | -20.503 | -5.394 | -5.432 | -1.023 | 18.174 | 25.798 | 10.685 | 11.643 | 21.897 | 2.832 |
| -18.443 | 23.933 | -11.305 | -16.190 | 17.619 | -7.459 | -26.620 | -3.974 | 21.432 | 26.158 | 25.978 | 25.933 | 4.079 | 17.644 |
| 3.346 | -21.051 | -27.125 | -7.228 | 29.657 | -13.055 | -19.702 | 4.123 | 2.667 | 21.055 | -7.879 | -3.563 | -3.382 | -10.924 |
| -5.401 | -3.418 | 18.318 | -6.323 | 15.209 | 3.167 | 2.155 | -22.299 | -0.776 | 25.831 | 21.577 | 21.751 | 15.049 | 27.173 |
| -11.165 | -17.837 | 19.549 | 22.867 | 8.374 | 0.927 | -20.238 | -5.406 | 7.069 | 13.434 | 17.910 | 5.848 | 12.806 | 0.056 |
| 23.986 | 14.577 | -12.505 | 1.484 | 7.655 | 21.396 | -0.308 | 2.054 | 1.203 | 18.402 | 19.743 | 0.265 | 23.838 | -12.136 |
| -7.198 | 5.277 | 4.058 | -5.555 | -21.598 | 11.725 | -3.542 | -0.043 | -16.336 | -25.543 | 14.614 | 3.732 | -5.559 | -3.400 |
| -23.769 | -9.610 | -26.546 | -25.180 | 13.775 | -19.305 | 8.950 | -13.785 | 19.708 | 12.140 | 18.916 | 12.527 | 27.660 | 16.565 |
| 4.420 | 22.382 | 27.121 | -9.670 | 24.111 | 12.941 | -7.424 | 0.939 | 24.060 | 21.620 | 11.666 | 22.624 | 6.304 | 9.402 |
| -25.300 | -7.256 | 25.350 | -23.511 | 12.426 | 30.203 | -10.420 | -1.823 | -12.391 | 19.972 | 5.701 | 4.370 | 25.793 | 17.599 |
| 15.170 | -8.986 | -21.795 | -0.056 | 18.341 | 22.483 | 19.712 | -21.050 | -1.378 | 15.017 | 15.746 | 25.370 | 16.482 | -0.844 |
| -8.424 | 23.593 | 15.845 | -7.347 | 15.835 | 23.008 | 1.916 | 24.244 | 0.066 | 13.745 | 17.747 | 24.583 | 20.238 | -2.107 |
| -22.381 | -13.236 | 20.697 | 24.974 | -7.039 | -8.249 | -18.246 | -8.554 | -20.556 | 18.740 | 18.668 | 20.562 | 18.988 | -0.872 |
| 7.025 | 6.529 | -14.370 | 5.749 | -17.775 | 27.195 | 20.492 | 16.519 | -23.393 | 15.188 | 10.907 | 0.700 | 17.647 | 26.216 |
| 24.485 | 17.778 | 6.318 | 15.840 | -1.265 | -29.108 | -23.067 | -9.839 | -8.491 | -24.956 | 0.365 | 21.829 | 10.982 | -7.442 |
| 14.111 | -19.209 | -24.945 | -16.998 | 28.932 | 9.111 | 2.697 | 24.893 | 5.693 | -18.785 | 16.522 | 23.488 | 8.734 | -7.750 |
| 23.750 | -2.114 | 23.445 | -6.855 | 21.954 | 7.664 | 26.372 | -6.959 | -5.868 | -8.421 | 3.157 | 25.733 | 19.591 | 5.527 |
| 8.001 | -5.381 | 26.635 | -15.162 | -18.757 | -0.525 | 3.235 | 11.469 | 21.507 | -9.688 | 24.399 | 14.239 | 14.971 | -17.572 |
| -0.558 | 20.862 | -15.236 | -8.010 | 25.310 | 25.724 | 24.842 | 26.168 | 2.946 | -0.113 | 9.836 | -24.441 | 11.706 | 1.053 |
| 17.305 | 19.988 | 20.066 | 9.987 | 5.612 | 2.704 | 12.169 | 13.752 | 26.311 | 6.480 | 13.168 | 9.160 | 23.802 | 8.081 |
| -4.161 | 11.949 | -5.501 | -15.802 | -25.675 | -4.888 | -7.446 | -27.340 | 4.433 | 22.878 | 19.472 | 19.968 | 6.796 | 20.644 |
| 3.125 | -2.542 | 21.497 | -25.191 | 17.222 | 24.041 | 19.270 | 7.060 | 17.863 | -2.214 | -0.861 | -0.734 | 24.375 | 3.729 |
| -1.538 | -6.363 | 15.351 | 2.473 | -26.752 | -25.470 | 1.186 | -6.087 | -6.604 | 16.720 | 13.871 | -17.951 | -21.732 | -0.355 |
| 15.138 | 22.674 | -27.746 | 14.661 | 22.587 | 5.066 | 8.782 | 21.475 | 26.804 | 24.423 | 16.837 | 17.663 | 3.045 | 10.966 |
| 1.813 | -26.308 | -4.787 | -3.828 | 27.938 | 15.253 | 24.442 | 11.576 | -18.030 | 15.702 | 18.984 | -8.675 | -14.385 | 12.562 |
| -21.616 | 28.011 | -9.010 | 6.107 | -8.689 | 19.242 | -11.195 | -7.514 | -22.602 | 9.075 | 20.788 | 11.042 | 20.977 | 19.282 |
| -23.719 | 2.686 | -16.615 | -9.130 | 23.738 | 13.963 | -0.653 | -4.238 | 11.891 | 7.343 | 23.317 | 8.790 | 15.998 | 10.215 |
| 6.975 | 14.948 | 19.448 | 4.616 | 19.992 | 3.457 | 5.982 | 10.484 | 23.626 | 2.253 | -11.818 | 13.735 | -12.737 | 3.062 |
| 14.250 | 14.361 | -7.280 | -6.430 | 4.055 | 5.607 | -12.982 | -11.192 | 21.768 | 18.622 | 23.916 | 24.017 | 17.886 | 16.390 |
| 17.352 | -7.339 | -3.211 | -22.693 | -9.324 | -18.322 | -5.604 | -0.184 | -11.626 | -1.459 | -8.559 | 16.522 | -7.185 | 12.224 |
| 15.360 | 25.698 | -19.551 | -4.110 | -4.228 | 13.293 | -9.256 | 1.864 | 0.579 | -15.211 | -16.747 | 5.383 | 7.141 | 9.163 |
| 21.525 | -19.999 | 20.674 | 8.217 | -0.258 | 11.271 | -14.602 | 7.928 | 23.832 | 13.819 | -24.284 | 11.471 | 10.193 | 0.474 |

| | | | | | | | | | | | | | |
|---|---|---|---|---|---|---|---|---|---|---|---|---|---|
| 8.697 | 16.479 | 2.641 | -4.497 | -18.752 | 1.951 | 16.853 | -18.079 | 20.458 | 6.135 | 10.331 | -1.241 | 6.609 | 17.256 |
| -16.838 | -2.463 | -21.293 | -20.904 | 12.860 | -8.278 | 16.444 | -5.322 | -6.028 | 9.731 | 13.669 | 9.688 | 24.669 | 14.766 |
| 5.974 | -26.802 | 26.637 | 22.366 | 9.013 | 25.150 | 19.620 | 1.113 | 18.935 | -3.955 | 21.314 | 11.643 | 21.537 | 6.147 |
| 5.793 | -4.178 | -19.892 | 3.841 | -23.046 | 17.343 | 10.362 | -18.303 | 20.933 | 24.435 | 12.044 | 13.965 | 6.829 | 22.855 |
| -12.544 | 21.872 | -5.140 | -2.671 | -7.696 | 6.501 | -7.867 | -3.525 | 17.897 | -18.212 | 19.265 | 16.196 | 16.646 | 9.843 |
| -3.112 | -14.854 | 19.938 | -25.602 | -23.886 | 18.316 | 3.916 | 4.433 | 27.124 | 15.682 | 26.825 | 8.913 | 20.001 | 4.699 |
| -11.225 | -3.531 | 4.935 | -18.996 | 28.608 | -14.194 | 16.886 | 5.042 | -7.341 | 26.051 | -16.253 | 18.682 | 9.383 | 16.359 |
| -0.681 | 24.064 | 0.182 | -10.431 | -11.779 | -19.750 | -1.185 | -24.494 | 22.309 | 2.961 | 19.857 | 10.661 | 3.943 | -22.488 |
| -26.629 | -17.537 | -11.202 | -13.801 | 16.806 | 27.299 | -11.381 | -0.555 | 3.735 | 18.158 | 21.934 | 25.174 | 21.863 | 20.028 |
| 26.562 | -4.654 | -3.674 | 19.306 | -6.435 | 30.060 | -7.939 | -24.474 | 2.020 | 11.043 | 10.110 | 23.799 | 8.185 | -7.283 |
| -13.402 | 20.909 | 1.236 | -3.558 | -22.117 | 22.087 | -10.609 | -6.649 | -10.537 | -15.470 | 17.504 | 9.007 | 10.564 | 12.850 |
| -20.277 | -18.360 | 27.142 | -7.827 | -4.181 | -22.577 | 21.240 | -15.976 | 19.877 | 25.940 | -14.638 | -10.753 | 20.456 | -23.843 |
| 25.290 | -19.131 | 15.167 | -4.954 | 20.225 | 26.408 | -25.137 | -1.789 | -8.178 | 5.666 | 19.093 | -7.279 | 12.088 | -1.200 |
| 6.562 | -8.126 | -17.041 | 6.432 | -5.773 | -9.116 | -20.993 | -19.184 | 17.567 | 25.263 | 8.134 | 7.727 | -3.470 | 21.629 |
| -8.108 | -21.315 | 21.235 | -2.602 | -13.239 | -10.829 | -16.752 | 13.427 | 5.144 | 9.928 | 23.356 | -20.089 | 9.598 | 12.240 |
| -3.462 | -2.843 | 0.680 | 9.716 | -8.207 | 28.278 | 10.738 | 20.066 | 11.314 | 13.956 | 7.367 | 15.850 | 18.818 | 11.038 |
| -18.368 | 13.852 | -6.035 | -8.259 | 19.253 | -6.595 | 24.264 | -12.837 | 0.752 | -6.922 | 11.317 | 24.256 | 7.028 | 13.118 |
| -28.106 | 18.568 | 15.684 | -13.817 | -18.687 | -3.354 | -17.384 | -22.920 | 14.022 | 19.532 | 18.926 | 17.328 | 6.642 | 1.311 |
| -2.537 | 19.996 | 13.935 | -12.621 | 24.035 | -28.892 | -0.016 | 14.852 | -7.893 | -8.072 | 15.335 | 21.672 | -18.065 | -1.593 |
| 20.272 | -25.175 | -18.187 | -6.066 | -2.846 | 18.711 | 11.677 | -25.058 | -25.956 | 9.455 | 16.172 | 25.711 | 21.792 | 5.001 |
| 2.330 | 2.103 | 6.004 | 1.960 | 16.299 | 24.987 | -23.995 | 11.289 | 10.280 | -19.077 | 27.650 | 14.444 | -0.034 | 22.163 |
| -5.180 | 5.930 | 13.162 | -2.492 | -6.076 | 0.321 | 7.989 | -0.112 | 25.431 | -17.277 | 22.898 | 19.627 | 21.571 | 8.540 |
| -15.496 | -0.209 | -5.505 | 4.365 | 15.404 | 23.101 | 0.203 | 25.455 | 28.762 | 25.324 | 28.816 | 11.706 | -3.482 | 23.199 |
| -12.167 | -3.497 | 0.403 | -3.763 | 26.231 | -5.256 | -16.278 | 8.473 | 12.304 | 12.157 | 14.564 | 16.456 | 11.605 | 19.245 |
| -28.829 | -2.795 | -20.972 | 5.146 | -17.889 | 11.847 | -7.765 | 24.202 | 24.987 | 20.228 | 21.180 | 20.157 | 21.040 | 11.087 |
| -25.544 | 0.261 | 18.025 | -9.258 | -0.908 | 6.400 | -12.106 | 21.680 | 17.345 | -7.079 | 20.497 | 27.105 | 22.182 | -2.690 |
| 29.048 | 1.578 | -6.571 | 7.676 | 26.978 | 19.888 | -21.024 | -21.903 | 23.940 | 26.609 | 23.965 | 23.355 | 12.974 | 6.394 |
| -23.716 | -2.878 | -25.647 | 15.589 | -3.201 | 13.757 | -17.500 | 23.236 | -14.848 | 6.178 | 8.831 | 21.270 | 15.388 | 17.053 |
| -6.536 | -17.915 | -8.710 | 0.739 | 22.027 | 8.958 | 16.122 | 2.598 | 16.574 | 20.286 | 14.894 | 13.770 | 24.338 | 7.385 |
| -27.059 | -23.747 | 3.364 | -14.920 | 4.548 | -19.592 | -3.906 | -6.394 | 20.870 | 16.276 | -0.808 | 28.876 | 25.216 | 20.223 |
| 20.498 | -22.798 | -3.336 | 10.131 | -14.100 | 16.909 | -10.158 | -26.176 | 5.731 | 18.330 | 17.220 | 11.930 | 4.995 | 1.187 |
| -7.348 | 2.174 | 15.384 | 9.390 | 9.067 | 15.864 | -26.946 | 2.042 | 4.866 | 22.045 | 26.091 | 0.788 | 20.256 | 13.828 |
| -16.677 | -24.451 | 11.119 | 5.562 | -17.071 | -27.168 | 26.554 | 8.273 | -2.277 | -15.634 | 17.729 | 1.908 | -2.844 | 2.157 |
| 23.102 | 4.246 | 2.246 | 17.451 | -10.698 | -25.541 | -15.679 | -23.285 | 28.277 | 23.041 | -29.512 | 26.394 | 7.040 | 23.623 |
| -20.947 | 7.845 | 5.118 | -6.639 | 0.682 | -0.642 | 21.772 | -15.554 | 25.352 | -22.024 | 15.415 | 25.223 | 21.900 | 15.070 |
| -13.393 | -10.321 | 21.537 | -25.082 | -23.453 | 13.398 | -22.365 | 15.496 | 24.570 | -24.130 | 13.259 | 23.400 | 17.201 | -1.197 |

| | | | | | | | | | | | | | |
|---|---|---|---|---|---|---|---|---|---|---|---|---|---|
| -8.521 | 23.734 | 24.540 | 11.289 | 1.799 | 18.253 | 0.197 | 26.503 | 26.871 | 27.716 | 14.085 | 24.090 | 8.613 | 8.866 |
| 18.742 | -13.049 | -0.829 | -0.976 | -8.802 | 24.754 | 20.462 | 25.038 | 5.906 | 8.881 | 27.215 | 28.250 | 0.959 | 14.640 |
| -20.962 | 19.632 | 8.050 | 1.810 | 26.757 | 4.108 | -12.906 | 16.370 | 4.149 | 25.094 | 25.951 | 22.749 | 27.423 | -16.536 |
| 4.851 | 7.891 | -6.346 | 7.371 | 18.347 | -17.656 | 17.051 | -9.752 | -24.458 | 8.017 | 18.456 | 21.886 | 4.512 | 17.658 |
| -14.341 | -17.617 | 8.991 | 3.128 | -24.638 | 12.383 | 22.662 | 22.644 | 20.605 | 24.138 | 28.766 | 18.157 | 26.787 | 4.790 |
| 14.055 | 0.358 | 22.168 | -12.573 | 12.700 | 20.569 | -21.168 | 0.610 | -11.376 | 6.875 | 20.283 | 10.170 | 14.113 | 5.406 |
| -11.848 | 1.103 | -3.553 | 24.937 | -5.271 | -13.459 | -18.073 | -0.133 | 16.774 | 25.628 | 19.501 | 17.284 | 17.560 | 20.146 |
| -11.380 | -18.261 | -9.792 | 20.047 | 10.348 | -1.325 | 9.077 | -3.505 | 22.111 | -9.883 | 23.784 | 19.445 | 5.406 | -3.819 |
| -19.134 | 19.568 | -10.677 | 9.394 | -7.139 | 8.865 | 5.573 | -13.307 | 7.696 | 7.576 | 10.471 | 20.425 | 24.888 | 23.728 |
| 27.480 | -21.880 | -4.573 | 21.494 | -20.642 | 10.885 | -16.322 | 17.024 | 8.594 | 9.334 | 17.555 | 11.714 | 22.096 | 18.354 |
| 4.648 | -23.690 | 2.355 | -0.924 | 6.154 | 27.193 | -4.555 | 7.790 | 10.716 | -2.617 | 24.813 | 26.828 | -16.283 | 5.937 |
| -25.547 | 3.806 | 10.134 | 7.589 | 26.747 | 25.885 | -27.871 | 2.861 | 5.729 | 26.541 | -7.295 | -17.763 | 20.957 | 18.372 |
| -7.762 | -2.264 | -13.024 | 6.860 | -14.752 | 12.006 | 5.487 | 19.431 | -6.256 | 23.391 | 11.068 | 8.408 | 21.993 | 24.337 |
| 15.032 | 20.931 | -25.389 | -10.186 | -2.605 | -22.815 | 8.270 | -12.194 | 26.616 | -27.225 | 18.110 | 3.034 | -14.720 | -6.619 |
| -9.864 | 19.390 | 6.345 | 23.478 | -7.599 | 14.200 | -24.740 | 25.543 | 0.394 | 26.966 | 9.881 | 15.358 | 25.744 | 23.355 |
| -26.586 | -5.877 | 6.639 | -20.068 | 13.735 | 1.459 | 19.750 | 21.650 | 17.703 | 25.726 | 20.022 | 27.760 | 29.251 | 16.800 |
| 7.078 | 18.795 | -9.675 | 3.025 | -5.877 | 18.968 | 16.521 | -26.077 | 24.219 | 22.348 | 6.905 | 11.032 | 9.881 | 13.748 |
| -6.454 | 4.835 | 8.225 | 21.826 | -18.897 | -0.922 | -7.264 | 28.037 | 22.055 | 11.957 | 11.888 | 12.532 | 12.567 | 9.563 |
| -2.854 | -9.362 | -2.214 | 4.287 | 23.000 | -3.339 | 0.117 | 26.917 | 12.407 | 3.745 | 19.188 | 29.433 | 14.979 | 14.358 |
| -19.535 | -10.170 | -4.832 | -0.664 | 15.292 | 28.933 | -6.227 | -4.078 | 28.811 | 6.900 | 29.151 | 18.240 | 8.027 | -11.764 |
| -14.297 | 6.973 | -6.391 | 8.121 | -13.110 | 20.992 | 11.276 | -7.400 | 25.321 | 17.829 | 27.558 | 15.730 | 3.514 | 23.271 |
| 6.056 | 8.839 | 16.775 | 22.565 | -4.010 | 16.575 | -8.097 | 13.713 | -14.813 | -24.525 | 9.796 | 28.940 | 16.833 | 25.110 |
| -18.214 | 2.601 | -5.382 | -24.560 | -8.417 | -20.212 | -1.809 | 25.545 | 20.879 | -0.927 | 21.971 | 19.306 | 16.185 | 12.945 |
| 11.385 | 8.088 | -13.703 | 3.357 | -0.407 | 25.410 | 3.528 | 12.846 | -9.648 | -26.497 | 20.901 | 13.749 | 26.598 | 3.749 |
| -8.041 | 11.390 | -3.119 | -12.148 | 21.868 | -10.646 | 9.562 | 9.288 | -25.465 | 4.161 | 14.406 | 27.278 | 19.894 | 22.074 |
| 8.682 | 22.388 | 5.163 | -8.872 | -1.365 | -7.966 | 4.928 | 29.025 | 21.360 | 17.298 | 7.375 | 15.147 | 17.303 | -2.684 |
| 13.228 | -1.577 | 4.565 | 22.355 | 13.078 | 28.134 | -9.672 | -10.348 | 23.430 | 14.850 | 18.014 | 21.049 | 22.445 | 20.981 |
| -8.538 | 14.352 | 12.733 | 16.255 | 13.948 | 26.053 | -2.316 | 5.381 | 11.547 | -9.367 | 24.127 | 13.150 | 9.396 | 12.215 |
| -3.520 | 20.648 | -23.360 | 22.989 | -9.616 | -3.822 | -7.093 | -5.365 | 29.612 | -28.856 | 11.260 | 17.183 | 27.247 | 15.373 |
| -2.641 | -0.314 | -9.152 | -1.694 | -24.426 | -14.998 | -27.726 | 23.802 | 26.110 | 17.752 | 13.861 | 19.181 | 13.001 | 14.401 |
| -19.513 | -10.746 | 2.453 | 7.032 | 19.759 | 3.769 | 25.527 | 16.598 | 28.301 | 10.493 | 27.081 | 23.503 | 22.960 | 5.804 |
| -11.265 | 7.920 | 25.157 | 17.430 | 0.972 | -3.155 | 22.068 | 28.650 | 12.154 | 13.220 | -1.633 | 14.184 | 5.786 | -7.936 |
| -10.162 | -12.695 | 18.562 | 2.288 | 6.474 | -4.414 | 1.521 | 19.860 | -26.327 | -15.482 | 23.728 | 29.160 | 21.392 | 28.221 |
| 10.902 | 27.504 | -0.062 | -20.356 | -1.589 | 16.193 | 8.514 | 8.886 | 25.508 | 4.327 | 13.796 | 12.823 | 19.591 | 23.383 |
| -12.715 | 1.729 | 1.998 | 19.678 | -12.832 | 4.901 | 27.069 | -29.096 | 23.011 | 19.272 | 18.653 | 22.700 | 20.416 | 6.834 |
| -19.001 | -14.770 | 5.143 | 11.621 | -17.772 | 22.596 | -12.752 | 27.802 | -14.134 | 17.311 | 9.343 | 17.895 | -7.445 | -23.640 |

| | | | | | | | | | | | | | |
|---|---|---|---|---|---|---|---|---|---|---|---|---|---|
| -15.318 | 22.391 | -9.000 | 16.514 | 23.816 | 17.705 | -5.788 | -15.509 | 21.991 | 14.215 | 21.269 | 14.039 | 25.991 | 10.527 |
| 5.175 | 17.399 | -10.826 | 2.936 | 17.012 | 19.256 | 2.232 | -16.284 | 24.211 | -26.848 | -3.587 | 21.431 | 12.334 | 7.100 |
| 12.770 | -5.475 | 11.099 | 13.939 | 0.660 | -11.129 | -9.452 | 9.748 | -20.819 | 23.896 | 25.363 | 29.651 | 25.352 | 15.984 |
| 4.091 | 15.847 | 1.704 | -10.413 | -21.212 | -2.604 | -20.673 | 3.122 | 19.333 | 18.094 | 28.705 | 18.999 | 14.784 | 5.860 |
| -23.501 | 9.821 | 1.315 | 1.070 | -11.932 | -26.853 | 24.045 | -7.555 | 30.063 | 16.854 | 1.870 | 12.119 | 23.512 | 11.201 |
| -5.949 | -8.763 | -3.086 | -15.307 | -1.715 | 24.324 | -28.664 | 19.018 | -10.380 | -1.601 | 11.857 | 17.611 | 10.941 | 19.817 |
| 18.797 | 12.191 | -10.452 | 6.799 | -18.831 | -24.720 | -5.026 | 28.935 | 24.964 | -8.860 | 19.377 | 29.025 | 16.418 | 11.749 |
| 7.865 | 13.723 | -6.871 | 7.935 | -6.950 | 2.361 | 29.519 | -11.137 | 16.097 | 20.471 | 15.827 | 1.153 | 17.381 | -14.977 |
| -13.375 | -0.997 | -1.751 | -3.409 | -3.376 | -1.194 | 11.249 | 6.300 | 30.761 | -29.675 | 22.554 | 22.497 | 30.180 | 4.965 |
| -3.470 | -0.108 | 12.851 | 0.738 | -20.430 | 5.256 | 21.026 | -8.428 | 15.847 | 28.852 | 23.614 | 12.596 | 21.568 | -2.953 |
| 11.180 | 9.267 | -13.811 | 22.164 | 2.170 | -10.932 | -2.779 | 20.570 | 26.593 | -12.286 | 27.099 | 30.728 | 14.755 | 0.717 |
| 5.100 | 16.978 | 21.042 | -7.370 | 9.757 | 18.320 | 7.371 | -27.254 | 11.834 | 15.653 | 12.219 | 23.865 | 15.859 | -4.250 |
| 15.502 | 7.961 | -10.055 | 18.038 | 4.179 | 13.772 | -16.535 | 12.003 | 18.951 | 12.892 | 20.556 | 19.754 | 18.266 | -10.180 |
| -8.487 | -6.963 | -5.124 | 2.082 | -10.851 | 10.589 | 30.688 | 29.458 | 22.055 | 7.990 | 21.939 | 10.776 | 13.161 | 9.481 |
| 6.527 | 19.277 | 7.404 | -24.012 | -12.394 | 21.235 | 12.657 | 27.893 | 13.126 | -27.696 | 11.680 | 29.394 | 7.863 | -6.921 |
| -5.734 | -7.496 | 4.490 | 3.760 | 3.670 | -16.307 | 28.529 | -4.946 | 28.100 | 7.115 | 15.999 | 22.316 | -23.309 | 15.442 |
| -20.893 | -2.492 | 2.296 | 13.680 | -20.860 | -18.984 | -20.005 | 27.544 | -8.949 | 4.721 | 19.960 | 11.849 | -6.821 | 12.986 |
| -17.375 | 18.474 | 6.277 | 11.388 | 1.706 | 18.158 | -18.608 | 25.682 | 31.366 | 28.742 | 23.404 | 15.940 | 4.827 | 24.313 |
| -27.768 | 23.725 | -12.286 | -11.490 | -23.919 | -0.499 | -11.441 | 2.414 | 29.827 | 28.284 | 9.961 | 14.695 | 14.410 | 21.804 |
| 11.499 | 16.326 | 8.140 | -4.819 | 9.780 | 17.657 | 25.191 | 26.659 | 21.573 | 14.268 | 26.332 | 27.948 | -22.366 | 4.531 |
| 18.257 | -9.595 | -20.774 | -0.313 | -10.412 | -21.947 | -25.792 | -14.324 | 28.359 | 15.095 | 27.473 | 9.096 | -0.148 | -1.559 |
| 13.409 | -24.113 | 5.120 | 19.318 | -1.911 | -11.823 | 2.707 | 26.945 | -14.557 | 15.859 | 22.283 | 18.545 | 26.005 | 22.996 |
| -2.139 | 29.097 | -5.228 | 3.305 | 3.785 | 10.863 | -21.335 | 16.769 | 7.143 | 28.926 | 27.013 | 13.045 | 25.387 | 16.523 |
| -1.253 | -14.235 | -4.372 | 7.885 | 17.483 | -10.809 | 22.350 | -6.107 | 25.210 | 24.971 | 12.664 | 23.989 | 22.565 | -8.094 |
| 8.849 | 24.371 | -15.016 | 5.595 | -16.667 | 15.988 | 21.206 | 23.464 | 9.399 | 8.830 | 14.103 | 7.236 | 12.579 | 14.295 |
| 3.178 | 12.210 | 6.948 | 20.015 | -22.534 | 10.012 | 12.943 | 2.974 | 10.483 | 28.678 | 18.491 | 15.987 | -25.119 | 15.369 |
| 4.264 | -11.398 | 2.232 | 12.625 | 16.921 | -15.697 | 8.396 | 13.496 | 3.774 | 26.405 | -0.387 | 20.136 | 17.701 | 8.451 |
| -19.475 | -21.587 | -4.539 | 29.053 | 6.789 | -1.491 | -24.768 | 27.775 | 17.807 | 23.390 | -5.383 | -3.529 | -5.783 | 23.406 |
| -16.866 | 20.742 | -25.873 | 28.129 | -23.594 | -21.081 | 18.228 | 18.925 | 19.236 | 15.870 | 17.929 | 9.665 | 7.053 | 22.344 |
| 24.934 | 1.767 | 11.534 | 15.037 | 24.032 | -8.644 | -5.503 | 17.166 | 23.973 | -13.526 | 7.479 | 24.740 | 14.606 | 30.005 |
| -20.497 | 15.676 | -22.764 | 10.388 | -2.640 | -22.121 | -6.900 | -13.594 | 1.382 | 23.311 | 17.221 | 20.885 | -7.561 | 20.938 |
| 1.045 | 8.708 | -19.886 | 8.452 | 7.379 | 9.381 | -26.830 | 29.477 | -9.024 | 18.359 | 10.441 | 24.029 | 19.002 | 9.975 |
| -4.896 | -7.921 | -7.456 | 1.307 | -0.101 | -3.075 | -18.945 | 14.092 | 29.070 | 22.111 | 16.217 | 15.439 | 28.827 | 11.819 |
| -9.211 | 24.755 | -22.302 | 26.033 | -16.684 | 18.971 | 30.898 | 7.607 | 27.078 | 5.586 | 19.815 | 22.521 | 13.004 | 25.791 |
| -24.421 | -24.532 | -6.514 | 7.354 | 24.476 | 5.619 | -24.544 | 21.645 | 10.465 | 19.049 | 4.484 | 23.242 | 17.230 | 12.520 |
| 24.093 | -8.355 | -5.219 | -19.025 | -8.020 | -5.732 | -27.747 | 5.829 | 24.982 | 18.138 | 24.877 | -18.417 | 14.805 | 11.230 |

| | | | | | | | | | | | | | |
|---|---|---|---|---|---|---|---|---|---|---|---|---|---|
| 13.161 | 29.760 | -3.153 | 21.675 | 14.394 | -5.077 | -16.386 | -4.364 | 23.049 | 11.805 | 16.945 | 14.271 | 10.739 | -9.771 |
| -14.434 | 23.078 | 8.010 | -20.844 | 11.809 | -0.580 | 15.420 | 1.873 | -4.861 | 15.242 | 12.190 | 28.319 | 22.183 | 23.257 |
| -6.301 | -4.933 | -3.883 | -10.294 | -2.331 | 1.600 | 15.179 | 29.605 | 18.281 | 22.110 | 10.188 | 17.072 | 29.684 | 3.258 |
| 6.886 | 13.990 | 4.452 | -21.678 | -7.149 | 11.697 | -24.338 | -11.411 | -19.068 | 19.396 | 17.067 | 28.855 | 25.222 | 9.038 |
| 22.382 | 6.303 | 15.202 | -0.284 | 19.589 | 6.645 | -26.738 | 12.347 | 25.734 | 26.440 | 20.393 | 6.607 | 19.939 | 10.616 |
| -0.928 | 16.491 | -25.968 | 16.043 | -1.382 | 0.121 | 15.771 | 25.739 | 8.319 | 29.686 | 18.189 | 4.911 | -8.857 | 14.478 |
| 4.499 | 8.114 | 17.367 | -2.544 | -17.631 | -3.890 | -30.319 | 26.960 | -3.652 | 2.011 | 24.345 | 14.046 | 23.826 | 8.141 |
| -17.985 | -9.533 | 27.536 | 23.068 | 7.508 | -10.158 | -27.960 | 22.636 | 10.168 | 23.287 | 9.792 | 14.411 | 7.906 | 15.880 |
| 10.406 | -26.743 | 8.497 | 10.956 | -3.477 | 14.937 | -1.637 | -12.707 | -20.481 | 21.429 | 14.446 | -0.107 | 16.351 | -19.439 |
| 11.090 | -11.982 | 5.075 | -6.314 | -9.435 | 9.804 | -7.182 | 6.562 | 27.748 | 4.687 | 26.351 | 24.337 | 13.822 | 21.705 |
| -7.157 | 12.096 | -15.712 | -27.477 | 23.064 | 19.193 | -25.937 | -19.772 | -15.277 | 24.070 | 22.553 | 22.472 | 22.662 | -18.272 |
| -17.405 | -2.940 | -2.427 | 27.625 | -17.358 | 22.163 | 16.109 | -1.544 | 2.691 | 11.531 | 19.037 | 7.866 | 17.958 | -7.637 |
| 8.946 | -4.788 | 4.330 | 17.129 | 10.021 | 8.875 | -17.614 | -24.788 | -4.416 | 18.674 | 16.746 | 20.587 | 7.421 | 18.056 |
| -16.551 | 25.550 | 3.052 | -13.398 | 25.949 | 9.364 | -14.023 | 21.493 | 23.163 | 10.699 | 21.427 | 17.625 | 12.333 | -13.845 |
| 5.141 | -10.673 | -14.725 | 28.453 | 5.671 | 2.908 | 8.397 | 30.045 | 21.755 | 22.957 | 6.958 | 27.823 | 15.779 | 15.707 |
| -3.821 | 27.955 | -22.414 | 7.244 | -13.431 | 18.298 | 4.152 | 19.154 | 19.959 | 19.797 | 18.918 | 10.057 | 14.936 | 13.136 |
| -0.274 | 8.102 | 9.384 | 29.777 | 21.722 | -14.184 | -10.075 | -22.016 | -0.252 | 21.355 | 14.532 | 6.045 | 7.977 | 17.333 |
| 15.010 | -15.686 | 23.714 | 25.743 | 12.158 | -12.576 | -18.311 | 20.877 | -21.454 | 24.885 | 12.492 | 20.034 | 19.471 | 21.071 |
| -18.071 | 25.931 | 8.014 | -12.082 | -11.733 | -18.410 | 19.569 | 4.385 | 20.696 | 14.877 | -4.614 | -6.434 | 17.097 | -8.963 |
| -11.035 | 23.605 | -9.037 | 24.339 | -24.532 | -6.580 | 23.111 | -25.663 | 8.562 | 3.282 | 11.114 | 18.120 | 5.803 | 13.561 |
| -17.108 | -13.103 | 11.013 | 5.103 | 13.897 | -21.754 | -25.024 | -10.452 | 26.593 | 26.478 | 25.748 | 20.016 | 6.852 | 15.030 |
| 24.472 | -3.436 | -13.945 | 15.756 | 8.480 | 24.414 | 5.146 | 30.669 | -21.921 | 17.506 | 23.720 | 18.717 | 23.141 | -23.347 |
| -13.969 | 20.047 | -17.700 | 3.604 | 24.643 | -9.403 | 15.415 | 13.302 | 11.266 | 6.590 | 13.845 | 16.477 | -2.393 | 29.002 |
| 18.928 | 0.461 | 0.152 | 5.960 | 18.714 | 8.394 | -11.775 | -28.707 | 24.802 | 18.724 | 5.925 | 25.449 | 2.460 | 16.846 |
| -15.227 | 30.563 | -6.364 | 13.442 | -15.279 | 16.284 | 28.717 | 16.874 | 16.805 | 24.964 | 15.470 | 10.035 | 13.019 | -1.290 |
| 9.223 | 17.005 | -20.649 | -9.572 | 11.520 | 4.814 | -20.288 | 27.519 | -4.173 | 19.392 | 6.527 | 26.367 | -1.829 | 7.509 |
| 18.453 | -3.980 | -8.461 | 11.928 | 23.701 | -5.191 | -11.434 | 23.032 | 12.316 | 24.211 | 22.915 | -26.536 | -29.074 | 6.345 |
| -7.652 | 18.674 | 14.726 | -10.923 | -23.109 | 0.232 | 23.716 | 8.048 | -3.893 | 17.918 | 21.508 | 23.761 | -4.279 | 23.020 |
| -16.817 | 4.831 | -13.062 | -3.492 | 24.226 | 13.262 | -18.860 | -23.166 | 28.113 | 23.143 | 9.899 | 25.305 | 13.900 | 27.090 |
| -20.491 | 13.776 | -1.352 | 9.287 | 21.982 | -14.558 | -21.635 | 20.024 | -4.797 | 13.459 | 18.701 | 6.207 | -23.738 | 11.363 |
| -8.619 | 26.020 | -11.433 | -5.501 | 20.461 | -2.091 | 3.387 | -5.625 | 24.588 | 24.317 | 25.384 | -25.766 | 18.052 | 3.115 |
| -22.565 | 28.605 | -29.151 | 18.325 | 17.902 | 16.776 | -19.366 | 16.800 | 4.043 | 23.360 | 21.084 | 21.998 | 23.730 | -24.807 |
| 12.568 | -11.877 | 29.060 | -26.077 | -9.146 | -19.201 | -9.378 | 4.101 | 19.152 | -7.950 | 2.971 | 11.494 | 20.668 | 18.784 |
| 9.938 | 22.836 | 8.084 | -6.893 | 16.056 | 25.456 | -24.826 | 28.948 | -22.084 | 24.066 | 23.781 | 18.877 | 0.260 | -0.746 |
| 1.487 | -6.533 | -7.200 | 30.958 | 18.531 | -1.557 | -15.998 | -4.981 | -25.669 | -11.366 | 22.099 | 24.759 | 25.203 | 19.623 |
| 17.597 | 10.759 | -19.002 | 20.008 | -17.258 | -16.224 | -14.252 | 16.957 | 7.641 | 19.265 | 11.628 | 14.983 | 15.242 | 14.260 |

| | | | | | | | | | | | | | |
|---|---|---|---|---|---|---|---|---|---|---|---|---|---|
| 21.488 | -14.260 | 3.008 | -4.170 | -28.042 | 7.008 | -8.362 | -21.612 | -0.044 | -12.215 | 16.036 | 23.781 | 26.672 | -4.838 |
| 15.412 | -0.293 | -3.142 | 1.447 | 8.967 | -7.369 | 21.760 | 27.281 | 18.122 | 0.406 | 24.577 | 14.595 | 14.723 | 12.366 |
| -2.772 | 15.555 | -9.102 | 3.731 | 20.974 | 24.430 | 15.498 | -0.725 | -26.235 | 7.757 | 19.232 | 14.429 | 23.351 | -8.496 |
| -24.823 | 12.013 | 22.138 | 5.344 | -7.401 | -15.787 | 12.027 | 6.416 | -1.638 | 9.443 | 19.736 | -25.400 | 17.036 | -20.696 |
| -0.062 | -13.607 | -20.406 | -26.827 | 19.754 | -3.044 | 23.599 | -8.989 | -17.764 | 17.918 | 20.964 | -20.375 | 12.802 | 4.516 |
| -4.793 | -23.509 | 30.283 | 9.368 | -25.321 | 12.858 | 17.117 | 21.609 | 1.051 | 23.513 | 10.219 | 10.898 | 6.167 | -9.480 |
| -9.440 | 18.325 | -15.197 | -13.069 | 7.340 | -12.390 | -25.259 | 1.929 | 24.468 | 5.019 | 21.488 | -10.259 | 22.790 | -4.528 |
| -11.291 | 20.877 | -26.143 | 10.398 | 16.945 | 3.735 | -15.131 | -14.809 | 1.973 | 12.234 | 5.749 | 12.469 | 9.391 | 21.632 |
| 3.771 | -9.250 | -27.243 | 20.998 | 28.791 | -8.791 | -24.265 | -10.856 | -22.063 | 25.723 | -2.399 | 22.410 | 9.843 | 11.064 |
| -0.155 | -19.625 | -7.258 | 14.424 | -24.217 | -14.881 | -23.200 | -21.930 | -6.222 | 0.568 | 24.442 | -2.404 | 24.080 | 1.586 |
| -16.969 | 12.240 | -0.063 | 13.760 | 0.423 | 4.139 | -16.781 | 7.842 | 15.114 | 7.647 | 14.748 | -1.297 | 14.776 | 20.239 |
| -25.504 | 25.927 | -8.987 | 21.352 | 22.535 | -11.010 | 5.010 | 24.081 | 27.926 | 21.893 | 16.131 | 25.079 | 2.302 | 2.874 |
| -14.513 | 22.458 | -12.081 | 23.494 | 18.079 | -8.285 | -7.329 | 2.996 | 26.633 | 16.864 | -11.997 | 14.833 | 24.592 | -2.226 |
| 5.110 | -1.031 | 12.792 | 29.341 | 28.287 | 0.047 | 4.162 | 17.114 | 17.819 | 11.037 | 4.686 | 23.868 | 16.471 | 27.418 |
| -22.933 | -20.754 | 20.585 | 12.766 | -22.390 | 6.578 | 6.544 | 9.817 | 16.606 | 24.550 | 25.687 | 6.246 | -18.427 | 24.697 |
| 13.750 | 26.645 | -29.342 | 23.819 | -3.997 | -0.916 | -10.129 | 22.900 | -4.175 | 26.558 | 19.505 | 12.442 | -25.132 | -6.821 |
| -6.527 | 5.285 | 22.863 | 2.061 | 26.181 | -15.169 | -16.484 | -7.793 | -6.017 | 12.556 | 4.420 | 6.499 | -15.331 | 4.632 |
| -19.766 | -22.510 | 13.293 | -6.170 | 1.931 | 13.176 | -4.236 | 13.798 | -18.044 | -22.250 | 11.870 | 0.643 | 0.185 | 18.057 |
| -3.219 | -16.669 | -23.943 | -30.383 | 19.721 | -1.240 | -18.487 | 15.370 | -12.341 | -3.519 | 9.391 | 1.560 | 22.020 | 27.147 |
| -2.135 | 9.503 | 3.135 | 15.525 | -17.776 | -4.860 | 21.405 | 16.779 | -1.628 | -1.858 | 9.907 | 22.017 | 9.765 | -7.043 |
| 8.248 | 31.670 | -4.870 | -8.654 | 29.025 | -11.859 | -20.863 | 4.936 | 23.165 | 16.427 | 15.792 | 24.836 | -24.994 | 16.114 |
| -19.611 | -4.298 | 4.131 | -12.373 | 20.732 | -3.375 | -17.590 | -3.535 | 3.684 | 20.447 | 16.596 | 16.684 | 19.250 | -2.406 |
| -0.342 | 14.794 | -12.884 | -27.414 | 18.810 | 10.062 | -4.993 | 26.876 | 0.540 | 13.090 | 1.029 | 9.842 | 2.932 | 9.186 |
| -22.694 | -22.943 | 21.446 | 27.437 | -19.531 | 3.767 | 24.305 | 12.854 | -0.880 | 26.863 | 22.873 | 23.750 | -9.301 | 26.483 |
| 15.362 | 27.796 | -1.466 | 24.460 | 9.524 | 2.558 | -19.673 | 18.679 | 11.857 | 25.390 | 22.377 | 21.769 | 18.458 | -10.069 |
| -6.326 | 12.895 | -13.334 | 26.541 | 5.982 | 7.921 | -18.791 | -6.665 | 24.218 | 22.877 | 16.132 | 17.588 | 7.286 | 14.894 |
| -7.639 | 29.035 | -16.885 | 3.354 | -24.479 | 0.465 | -15.134 | 3.215 | 23.398 | 22.663 | 12.474 | 11.160 | 20.171 | 17.478 |
| 16.874 | -1.713 | 6.887 | 30.238 | 23.938 | -9.382 | 22.870 | -5.058 | -13.229 | 3.714 | 15.339 | 21.893 | -17.826 | -12.877 |
| -22.054 | -4.883 | -25.033 | -28.279 | 0.400 | 9.552 | -26.020 | -3.602 | 20.347 | -21.733 | 22.552 | 11.402 | 4.472 | 18.978 |
| 13.284 | -25.701 | 10.951 | 25.279 | -30.686 | -15.293 | -21.358 | 28.308 | 4.809 | 21.175 | -22.196 | 5.497 | 11.178 | 6.764 |
| 9.673 | -6.863 | -5.803 | -20.234 | 4.597 | 19.726 | 3.649 | 17.603 | -10.087 | 12.689 | 21.491 | 13.458 | 25.451 | 21.090 |
| -20.339 | 24.536 | -17.851 | 25.826 | -0.384 | -19.890 | 5.439 | -5.844 | 27.647 | 4.076 | 26.143 | 16.741 | 12.449 | 12.521 |
| -16.051 | -23.628 | 18.326 | 20.330 | 23.277 | -13.922 | 14.120 | 9.781 | 26.780 | 19.023 | -13.259 | 23.364 | 15.768 | 24.957 |
| 7.325 | -8.210 | 10.243 | 25.305 | -20.421 | 1.232 | -8.658 | 12.705 | -14.026 | 21.900 | 4.542 | 21.089 | 10.115 | 5.338 |
| -2.517 | 30.203 | 3.296 | -3.670 | -6.995 | -20.852 | -13.199 | 4.912 | -3.469 | -3.393 | 25.939 | 6.266 | 15.226 | -0.589 |
| 23.015 | -17.104 | -19.111 | 21.660 | 25.391 | -23.012 | -3.670 | 6.762 | 21.188 | 4.812 | 24.203 | 5.732 | 11.558 | 17.280 |

| | | | | | | | | | | | | | |
|---|---|---|---|---|---|---|---|---|---|---|---|---|---|
| 4.981 | -4.112 | 11.741 | 27.417 | 8.439 | 14.641 | 24.729 | -24.892 | -4.515 | 26.974 | 9.829 | 11.726 | -25.445 | -9.711 |
| -8.908 | -9.542 | -30.090 | -12.347 | 1.140 | -8.771 | 21.849 | 23.106 | -5.719 | 12.032 | 23.627 | 19.212 | -19.624 | 14.472 |
| -13.579 | -24.816 | 28.236 | -5.493 | 27.649 | -10.994 | 10.556 | 10.600 | -16.261 | -21.961 | 18.162 | 7.528 | 13.812 | 7.684 |
| 20.625 | -0.763 | -1.697 | 29.528 | -29.715 | 6.355 | -17.233 | -5.874 | 10.519 | 8.892 | 13.876 | 20.085 | 5.313 | 11.752 |
| 12.279 | 21.589 | -7.785 | -16.912 | -11.799 | -6.239 | -10.277 | 16.240 | -8.835 | -15.298 | 3.263 | 17.128 | 15.239 | -18.912 |
| -26.796 | 10.968 | -11.213 | -25.363 | -23.106 | 21.446 | -20.023 | 1.772 | 23.521 | -7.697 | 22.310 | 28.214 | 0.487 | -19.569 |
| -25.616 | -2.124 | -14.533 | 26.511 | 30.144 | 0.680 | 9.035 | -6.616 | 3.366 | -7.452 | 20.864 | 18.651 | 19.200 | -12.832 |
| 14.027 | 15.339 | -17.972 | 1.153 | 6.072 | -12.331 | -10.859 | -13.688 | 17.483 | -19.276 | 16.098 | -1.896 | 20.667 | -16.250 |
| -14.241 | 20.136 | -22.483 | 23.418 | 18.821 | 18.946 | 11.849 | 20.969 | -7.657 | 10.895 | -4.192 | 23.069 | 24.655 | -17.842 |
| -1.786 | -23.114 | 31.111 | 13.015 | 17.539 | 2.004 | -0.848 | -1.194 | 9.867 | -20.433 | 20.398 | -6.900 | 17.347 | -0.806 |
| 19.460 | -4.012 | -10.720 | 18.787 | 19.276 | -18.349 | -3.105 | 0.824 | -5.361 | 25.161 | 8.345 | 26.908 | 10.100 | 3.422 |
| -3.073 | 11.507 | 27.765 | 28.934 | 22.132 | 3.749 | 7.894 | -12.115 | 24.376 | -7.233 | 24.414 | 27.193 | 25.137 | 1.956 |
| 2.629 | -22.249 | 11.069 | -23.031 | 15.413 | -10.402 | 7.356 | -17.739 | 21.535 | -8.211 | 13.618 | 18.106 | 24.329 | -1.733 |
| 23.071 | -12.986 | -20.779 | 12.562 | 17.424 | 24.231 | 24.281 | -7.895 | 8.563 | -13.567 | 19.505 | 19.325 | 23.868 | 7.231 |
| -24.028 | -7.233 | -1.275 | 17.306 | -14.731 | 5.728 | 1.424 | -3.607 | 15.918 | -7.607 | 20.892 | 5.820 | -23.037 | -0.019 |
| -20.230 | -23.118 | -3.865 | 8.947 | 21.051 | 13.834 | 19.821 | 19.294 | 25.983 | 20.487 | 17.474 | 21.174 | 18.439 | -10.699 |
| 27.862 | 19.788 | 21.818 | 19.274 | -12.670 | -13.630 | 10.624 | 26.029 | 10.927 | 18.265 | 22.791 | 11.800 | -17.312 | 11.305 |
| -5.603 | -21.676 | -28.200 | 29.392 | -16.526 | 7.111 | -3.726 | 11.613 | 1.859 | 3.809 | 14.517 | 4.506 | 20.461 | 24.844 |
| 26.382 | -19.356 | 26.094 | 5.017 | -27.359 | 1.702 | -4.122 | 16.139 | 2.599 | -15.004 | 27.206 | 24.344 | 12.062 | -11.691 |
| 24.444 | -26.057 | -25.210 | 24.114 | -5.786 | 11.199 | 18.857 | 2.695 | 18.807 | 16.775 | 16.781 | 7.337 | 17.790 | 12.045 |
| -12.684 | 26.918 | 12.459 | 11.509 | 26.666 | -16.553 | 14.921 | -11.546 | 5.222 | -17.039 | 4.614 | 17.355 | 23.387 | -5.571 |
| -7.700 | -10.086 | 29.437 | -7.226 | 5.455 | -17.914 | -2.513 | 13.690 | 3.439 | 5.171 | 11.325 | 9.146 | 21.054 | -12.133 |
| -17.461 | 6.805 | 30.093 | 19.984 | 29.447 | 24.452 | -19.621 | 5.961 | 12.557 | 23.849 | 7.425 | 12.652 | 24.920 | 15.886 |
| 3.547 | 26.853 | -11.867 | 28.453 | 23.797 | 22.193 | 8.469 | 11.885 | 23.553 | 21.844 | 25.768 | 11.403 | -2.715 | -13.936 |
| 15.433 | 19.629 | 22.165 | 27.113 | 18.058 | 18.920 | -16.470 | 12.308 | 7.623 | 1.485 | 8.733 | 15.484 | 24.290 | -18.462 |
| -16.040 | 3.581 | 6.241 | 0.350 | 13.412 | 26.816 | 2.503 | 4.899 | -6.432 | 0.205 | 13.574 | 7.292 | 11.769 | 13.711 |
| 22.548 | 7.819 | -5.817 | 14.017 | 25.920 | -15.794 | 18.077 | 18.774 | -1.381 | -12.060 | 18.103 | 8.056 | 12.992 | 24.994 |
| 0.218 | -4.474 | -14.977 | 22.863 | -19.671 | -4.658 | -21.334 | 17.924 | -4.961 | 14.835 | 23.518 | -1.202 | 21.911 | 4.566 |
| 26.573 | -22.862 | -9.579 | -4.673 | -29.636 | -23.357 | 12.409 | 17.058 | 11.947 | 18.054 | 24.814 | -5.026 | 25.311 | 16.623 |
| -0.567 | -2.947 | -25.266 | -22.859 | 8.493 | 15.557 | -18.947 | -15.684 | 15.096 | -22.608 | 28.629 | 17.805 | 23.036 | -26.789 |
| -20.751 | 23.019 | -3.740 | 22.144 | -25.292 | 14.917 | -25.096 | -9.118 | 12.898 | 4.402 | 9.290 | 16.837 | 27.123 | 2.550 |
| -25.099 | 5.403 | -6.814 | 18.122 | -27.214 | -17.702 | 4.301 | -5.819 | 16.509 | 12.021 | 22.887 | 5.588 | 14.461 | 11.141 |
| 24.089 | 23.462 | -24.202 | 14.250 | 9.609 | 20.706 | 22.953 | -21.029 | 1.470 | 10.439 | 21.709 | -2.159 | 13.659 | -15.366 |
| 11.643 | -5.852 | 18.091 | 8.188 | -13.928 | -19.768 | -14.670 | -19.412 | 5.542 | 5.225 | 21.121 | 23.244 | 20.365 | -0.211 |
| -21.300 | -24.498 | -4.810 | 10.273 | -10.854 | 12.997 | 25.658 | 0.095 | -1.457 | -15.352 | 17.001 | 29.158 | 18.077 | -3.626 |
| 25.424 | 16.215 | -19.693 | 24.993 | 27.843 | 4.056 | -10.324 | 15.272 | 0.476 | 9.481 | 15.974 | 30.542 | 26.371 | -1.285 |

| | | | | | | | | | | | | | |
|---|---|---|---|---|---|---|---|---|---|---|---|---|---|
| -8.053 | 11.505 | 2.086 | 27.981 | 5.924 | -1.210 | 20.894 | -3.668 | 7.356 | -9.598 | 14.506 | 24.145 | 25.650 | -11.199 |
| 23.145 | -5.969 | -5.291 | 19.838 | 14.785 | 25.497 | -9.414 | 13.568 | 8.305 | -11.979 | 13.750 | 0.139 | 16.257 | 9.756 |
| -26.535 | -9.204 | 25.653 | 12.140 | 4.847 | -18.647 | 17.217 | 4.513 | -11.021 | 14.393 | 5.791 | 20.584 | 3.671 | 19.244 |
| 27.182 | -23.448 | -14.204 | 4.646 | 15.619 | 10.619 | -18.138 | 27.439 | 24.093 | -22.043 | 23.248 | 21.978 | 2.015 | 26.420 |
| -0.839 | 24.308 | -11.421 | -15.575 | 18.088 | -14.222 | -20.699 | 18.313 | -24.777 | 13.104 | 13.976 | 26.242 | -26.562 | 2.950 |
| -21.339 | -21.443 | 19.833 | -1.802 | 22.960 | 25.048 | 2.793 | 7.632 | -5.325 | 22.622 | 19.621 | 16.929 | -9.031 | 5.085 |
| 5.562 | -8.602 | -24.986 | -14.638 | -22.846 | -10.280 | 16.892 | 3.139 | 23.804 | 24.474 | 7.579 | -5.654 | 27.541 | 13.829 |
| -23.176 | -9.143 | 13.192 | 22.298 | 18.852 | -16.965 | 24.078 | 10.975 | -13.058 | 8.259 | 30.244 | 21.124 | -1.303 | 9.498 |
| 1.398 | 6.303 | -21.649 | 18.757 | 16.865 | 5.147 | -11.495 | 5.478 | 19.115 | 5.169 | 17.750 | 4.486 | 12.733 | 15.908 |
| -2.588 | -25.290 | 17.754 | -8.284 | 12.178 | 14.355 | 20.654 | -3.124 | 23.531 | -0.463 | -4.731 | 22.435 | 20.598 | 17.518 |
| -11.503 | -24.339 | -7.169 | 2.418 | -1.380 | 17.578 | -5.408 | -7.469 | -13.831 | 3.659 | 29.682 | 26.682 | 27.616 | -18.868 |
| 21.615 | 23.525 | -2.583 | -24.289 | -26.556 | 11.582 | -4.062 | 16.371 | 11.877 | 16.898 | 9.120 | 20.130 | 18.526 | 25.422 |
| 6.353 | 1.280 | -3.058 | 25.178 | 16.086 | -0.053 | 17.178 | 22.761 | 16.881 | 18.402 | 28.052 | 29.056 | 16.397 | 9.094 |
| 29.255 | 0.249 | 12.803 | -24.931 | 22.149 | 10.214 | 19.646 | 17.171 | 22.702 | 1.318 | 22.919 | 22.170 | 22.482 | 8.259 |
| -26.569 | 12.557 | 1.775 | 19.006 | 7.454 | -14.939 | -24.634 | -0.729 | 3.595 | 4.450 | 24.411 | 24.667 | 5.041 | 15.497 |
| 20.236 | 11.879 | -2.936 | 22.674 | 5.727 | -7.243 | -15.978 | -9.319 | -17.802 | 9.574 | 25.998 | 30.538 | 9.648 | 19.725 |
| -29.045 | 2.988 | -24.797 | 11.737 | -21.350 | 16.208 | 18.173 | 5.043 | 23.140 | 16.233 | 12.018 | 9.666 | 25.490 | -1.206 |
| -0.459 | 5.944 | -8.727 | -20.550 | -9.237 | 21.406 | 7.546 | 20.576 | 26.091 | 11.440 | 22.186 | 1.147 | 18.148 | 5.104 |
| 24.824 | 18.266 | 24.348 | 14.221 | 30.152 | 9.131 | 6.706 | 10.743 | -0.242 | -3.007 | 29.683 | 22.919 | 14.297 | 9.451 |
| 26.046 | 17.218 | 23.935 | -25.445 | 20.688 | 8.192 | 14.224 | 8.816 | 26.877 | 20.964 | 0.120 | 29.821 | 26.220 | -25.966 |
| 3.695 | 15.580 | -7.639 | 20.541 | 3.152 | -25.161 | -20.477 | 1.072 | -6.932 | -4.949 | 17.801 | 11.772 | 25.881 | 11.232 |
| 7.533 | 20.079 | -27.905 | 15.054 | 24.130 | 25.435 | -6.587 | -16.566 | 13.024 | 15.835 | 24.696 | 15.252 | 0.726 | -1.207 |
| 16.146 | 1.048 | 22.651 | 5.857 | 11.614 | -17.564 | 27.134 | -0.192 | 23.558 | 9.110 | -19.213 | 26.563 | 7.483 | 18.601 |
| 14.887 | 0.076 | -21.995 | -18.442 | 9.971 | -26.065 | -22.011 | 6.262 | 0.939 | 3.010 | 8.965 | 20.585 | 23.588 | 18.042 |
| 21.560 | 23.693 | 6.063 | 26.128 | -3.678 | 17.290 | 10.363 | -16.586 | -21.651 | 3.523 | 13.792 | 12.659 | 11.916 | 17.112 |
| 5.668 | 0.963 | 3.704 | -3.572 | 23.504 | 26.962 | -3.557 | 18.503 | 18.664 | -2.390 | 20.972 | 13.430 | 4.631 | -15.775 |
| -11.869 | 18.392 | 10.995 | 18.369 | -27.567 | -13.462 | 15.694 | 4.190 | 24.489 | 10.798 | 26.098 | 29.534 | -1.399 | 16.215 |
| -24.558 | 3.275 | -6.179 | -20.714 | 12.443 | -5.172 | 24.791 | -9.145 | 22.922 | 23.747 | 31.260 | 25.102 | 9.430 | 27.309 |
| 13.375 | 3.071 | -0.571 | 14.836 | 8.873 | -9.849 | -7.642 | 11.767 | 20.741 | 20.260 | 4.886 | 23.445 | 14.574 | 25.494 |
| 23.265 | -23.551 | -9.620 | -21.470 | 17.004 | 23.201 | 28.190 | -3.842 | 27.565 | 16.316 | 17.682 | 7.199 | 28.007 | -22.944 |
| 25.793 | -15.536 | -20.034 | -5.234 | -27.082 | 12.913 | -22.558 | -7.192 | 9.586 | 8.640 | 7.547 | 18.178 | 18.098 | 5.122 |
| 25.358 | -22.246 | -17.292 | 23.091 | 10.844 | 12.138 | 2.854 | 16.851 | 25.492 | 6.834 | 26.999 | 13.230 | 15.946 | -11.081 |
| 19.503 | 16.322 | -3.798 | 12.644 | -14.798 | -19.168 | -28.130 | 10.125 | 24.283 | 13.659 | 27.356 | 12.584 | 16.740 | 13.104 |
| -23.032 | 0.893 | -13.594 | -25.038 | -8.367 | 11.755 | -3.514 | 11.501 | 25.542 | 22.905 | 12.537 | -4.448 | 29.970 | 24.635 |
| 1.498 | 7.778 | 25.453 | -8.918 | 18.920 | -2.936 | -17.926 | 18.804 | 13.921 | 3.445 | 24.996 | 26.532 | 20.562 | -16.302 |
| 16.528 | -4.768 | 14.743 | 2.866 | 22.436 | 26.481 | -13.947 | 23.156 | 12.262 | 6.126 | 28.463 | 22.037 | 22.910 | -4.520 |

| | | | | | | | | | | | | | |
|---|---|---|---|---|---|---|---|---|---|---|---|---|---|
| -25.543 | 6.364 | -9.593 | -6.732 | 20.726 | 19.759 | 25.583 | -25.701 | 6.870 | 2.123 | 26.092 | -2.215 | 4.718 | 8.123 |
| -28.574 | -19.528 | -11.959 | 3.612 | 12.279 | 1.406 | -12.384 | 0.923 | -22.534 | 18.104 | 22.825 | 29.172 | 27.292 | 9.494 |
| 13.074 | 0.737 | 23.976 | -14.891 | 17.264 | -27.544 | -9.953 | -7.953 | 28.935 | 14.207 | 18.494 | 15.503 | -2.813 | 19.358 |
| 29.709 | -14.865 | 13.989 | -9.533 | 25.945 | 19.760 | -0.867 | -11.969 | 0.683 | -26.304 | 6.034 | 17.656 | 14.719 | 11.106 |
| -15.833 | 5.193 | -22.058 | 5.842 | 15.867 | 21.655 | -25.700 | 17.224 | 23.523 | 8.255 | 13.570 | 19.656 | 28.051 | 16.210 |
| -19.291 | -15.915 | 26.926 | -0.618 | -9.650 | 9.030 | 20.559 | -10.242 | 28.121 | 18.199 | 21.567 | 18.152 | -0.321 | 4.747 |
| 7.437 | 19.353 | -22.882 | 6.855 | 18.080 | -13.465 | 12.716 | 2.944 | 8.949 | 6.666 | 9.094 | 3.554 | -26.776 | 12.547 |
| 20.801 | 14.522 | 2.995 | 5.690 | 8.357 | -2.112 | 1.833 | 4.394 | 22.749 | -12.709 | 29.224 | 2.205 | 9.967 | 6.388 |
| 0.841 | 7.177 | -23.528 | -10.255 | 17.544 | 4.147 | -22.569 | 5.999 | 18.111 | 12.101 | 4.644 | 12.819 | 30.034 | 24.531 |
| -0.455 | 25.241 | -0.183 | -15.700 | 13.785 | -18.100 | 17.647 | -13.978 | 9.943 | 21.034 | 30.334 | 14.831 | 23.667 | 23.327 |
| 3.100 | 12.280 | 18.189 | 2.523 | 5.999 | -18.709 | 20.042 | 14.566 | 11.036 | 3.870 | 24.134 | 25.742 | 26.878 | -3.859 |
| -7.157 | -4.721 | -9.796 | 14.136 | -4.948 | -21.079 | -1.080 | 16.406 | 27.518 | 1.895 | 17.362 | 23.151 | 24.667 | 7.557 |
| -24.860 | 24.412 | -12.127 | 9.152 | -20.595 | 3.002 | -29.327 | -1.620 | 21.445 | 22.911 | 21.905 | 20.296 | 25.663 | 17.419 |
| 23.344 | -25.933 | -13.897 | 16.203 | 22.906 | 15.542 | 28.749 | -18.666 | 24.507 | 24.030 | 7.476 | 12.518 | 17.095 | 14.588 |
| -16.541 | 18.338 | -11.322 | 21.450 | 7.032 | -6.986 | -5.390 | 6.521 | 3.655 | 5.494 | 9.621 | 11.046 | 23.535 | 6.934 |
| 26.675 | 16.602 | 20.424 | 5.637 | -2.554 | -25.403 | 19.998 | 11.234 | 23.226 | -25.630 | 12.766 | 15.124 | 28.857 | 17.900 |
| -14.472 | -0.504 | 17.974 | 12.267 | -5.505 | 17.094 | 2.737 | -25.084 | -20.653 | 28.307 | 22.941 | 17.294 | 25.995 | 8.845 |
| 11.590 | -3.254 | -7.571 | 2.627 | 5.025 | 28.371 | 23.931 | 24.147 | 18.006 | 12.853 | 11.649 | 19.079 | 14.130 | 12.163 |
| 28.923 | 20.153 | 25.603 | -6.678 | -14.873 | 5.932 | 24.342 | -1.232 | 28.099 | 19.169 | 26.146 | 22.667 | 27.506 | 27.123 |
| 24.617 | -5.502 | -24.803 | -23.777 | -11.719 | 2.685 | -28.029 | -0.048 | 19.168 | 25.398 | 26.894 | 24.467 | 22.493 | 29.852 |
| 0.955 | 18.950 | -25.496 | 19.360 | -4.993 | 18.844 | 6.919 | 24.962 | 9.047 | -2.698 | -3.174 | 16.814 | 27.629 | 24.809 |
| 5.487 | 22.623 | 23.698 | 13.287 | 21.831 | 27.851 | -10.222 | 17.385 | -11.645 | 19.633 | 13.731 | 25.993 | 30.679 | 21.271 |
| 6.812 | -22.192 | 21.224 | -3.561 | 23.704 | 23.659 | -20.824 | -19.844 | 9.645 | 17.241 | 8.295 | 22.036 | -10.158 | 17.031 |
| -9.134 | -7.016 | 4.295 | -22.713 | 20.066 | 7.657 | 19.651 | 6.114 | 14.796 | 12.671 | 0.818 | 10.832 | 22.821 | 13.613 |
| 26.183 | -12.939 | 9.543 | 10.466 | 18.896 | 19.154 | 8.391 | 7.142 | 18.112 | 22.448 | -5.824 | 7.136 | 19.285 | 9.100 |
| -21.798 | 14.263 | 2.046 | -9.370 | 14.501 | 11.087 | 26.905 | 10.395 | 26.883 | 20.969 | 27.902 | 27.737 | 14.994 | 9.929 |
| -25.268 | 15.859 | -19.284 | -10.573 | 25.423 | 12.754 | -5.771 | 17.450 | 8.241 | -13.198 | 15.089 | 10.664 | 23.459 | 9.490 |
| -19.548 | 1.298 | -20.989 | -4.450 | 4.080 | 4.433 | 26.150 | 10.936 | 22.354 | 28.338 | -29.108 | -6.001 | 11.992 | 18.496 |
| -8.005 | 11.144 | 0.917 | -7.473 | -22.556 | 5.954 | -3.389 | 26.369 | 22.015 | 23.268 | 16.890 | 24.091 | -6.920 | -17.619 |
| 11.562 | 3.940 | 10.789 | -14.048 | 19.384 | 24.177 | -12.615 | -7.227 | 22.885 | 3.142 | 18.237 | 27.513 | 20.162 | 28.374 |
| -6.789 | 18.814 | 16.306 | 15.250 | 28.760 | -15.857 | 24.775 | 7.641 | 30.059 | 7.032 | 15.602 | 10.363 | 27.250 | 21.501 |
| -23.570 | -0.575 | -7.318 | -16.204 | 27.213 | -25.733 | 10.656 | -2.129 | 14.340 | -9.735 | 24.549 | 25.245 | 4.423 | -10.041 |
| 19.365 | 25.070 | 3.749 | 3.050 | -27.789 | -4.840 | -3.827 | 4.922 | 19.803 | -23.407 | 6.112 | 3.771 | 13.559 | 17.943 |
| -3.612 | 4.028 | -5.011 | -17.297 | 28.041 | -0.534 | 26.405 | -12.602 | 26.090 | -28.737 | 22.747 | 13.031 | 7.196 | 16.781 |
| -4.476 | -12.776 | -3.345 | 19.528 | 26.078 | 21.315 | 16.835 | 13.791 | 7.736 | 11.626 | 17.481 | 0.095 | -1.538 | 12.850 |
| 6.758 | 21.168 | 0.113 | -15.181 | 8.417 | -6.172 | 30.396 | -27.990 | 29.283 | 12.814 | -5.317 | 26.642 | 9.240 | 10.751 |

| | | | | | | | | | | | | | |
|---|---|---|---|---|---|---|---|---|---|---|---|---|---|
| 28.878 | 14.116 | -25.104 | -2.288 | 24.372 | -9.118 | -13.571 | 16.396 | 22.503 | -18.100 | 15.138 | 22.403 | 4.292 | 28.890 |
| 23.041 | 10.275 | 12.732 | -1.993 | -13.475 | 2.092 | 12.811 | 25.070 | 25.131 | 27.527 | 3.785 | 8.617 | 27.573 | 19.850 |
| 0.364 | 3.333 | 9.858 | 16.667 | -19.398 | -20.714 | 29.803 | 22.158 | 19.640 | 21.063 | 25.068 | 20.626 | 23.091 | -29.855 |
| -2.160 | -23.192 | 6.349 | -0.173 | -2.950 | 9.881 | 3.997 | 24.462 | 5.277 | 25.855 | 26.787 | 19.087 | 25.396 | 7.595 |
| 27.407 | -15.537 | 20.088 | -8.850 | -20.432 | 0.780 | 11.679 | 22.926 | 21.968 | 9.627 | 10.677 | 23.187 | 23.945 | 21.991 |
| -22.340 | -2.930 | 11.781 | -23.250 | 6.572 | 11.857 | 2.927 | -16.640 | 24.021 | 21.897 | 7.104 | 18.626 | 5.928 | 24.979 |
| 21.226 | 14.924 | 21.549 | 25.517 | 9.779 | -8.382 | -6.822 | -4.040 | 11.694 | 23.428 | 20.096 | 20.405 | 19.392 | -26.308 |
| -4.367 | 27.042 | 19.824 | 18.230 | -26.898 | 4.430 | -18.051 | -20.042 | 25.812 | 12.049 | 23.918 | 21.228 | 2.952 | 3.569 |
| 4.296 | 12.713 | 24.761 | -9.594 | 8.867 | -12.744 | 25.099 | 19.786 | 27.781 | 8.616 | 27.647 | 24.423 | 4.331 | 12.611 |
| 26.807 | -24.413 | -4.135 | 22.014 | 21.519 | 13.730 | 27.436 | 3.103 | 29.512 | 24.578 | 27.114 | 6.179 | 22.266 | -27.182 |
| -9.442 | 7.069 | -2.711 | 8.142 | 6.152 | -2.852 | -25.772 | 16.502 | -23.205 | 6.623 | 22.517 | 15.993 | 20.753 | 10.734 |
| 2.654 | 21.046 | -3.082 | -17.838 | 27.234 | -20.725 | 14.584 | 21.904 | 31.305 | 22.014 | 16.521 | 25.463 | 27.086 | 1.429 |
| 24.557 | -4.556 | 9.311 | -5.678 | 4.212 | 26.047 | -16.463 | -6.157 | 26.667 | 2.342 | 26.051 | 19.573 | 17.591 | -20.868 |
| 13.420 | 15.528 | 12.067 | 12.220 | -21.144 | -5.312 | -17.272 | 9.726 | 19.469 | 17.163 | 23.749 | 23.717 | 7.501 | 22.818 |
| -5.814 | -11.560 | 20.299 | -10.954 | 7.963 | 22.161 | -21.805 | 19.340 | 16.281 | 26.455 | 14.824 | 20.471 | 26.404 | 18.823 |
| -10.938 | 10.056 | 22.874 | -0.850 | -5.929 | 24.017 | 10.971 | 11.698 | 20.982 | -9.358 | 26.253 | 3.028 | 2.501 | -25.037 |
| -0.827 | 16.318 | 2.693 | 3.867 | 16.952 | 13.377 | -1.406 | 16.150 | 13.609 | 17.390 | 15.835 | -3.513 | -8.338 | 9.356 |
| -7.088 | -18.264 | -22.621 | -7.109 | -16.151 | 28.642 | 28.029 | -6.213 | 6.029 | -6.359 | 18.495 | -8.202 | 14.754 | -19.625 |
| 3.554 | 8.172 | -9.382 | 14.269 | 8.528 | 6.221 | -18.497 | -25.808 | 22.375 | 28.344 | 10.260 | 19.762 | 8.597 | 10.623 |
| 2.973 | -0.743 | -7.827 | -3.125 | 25.945 | 17.621 | 17.032 | 21.426 | 12.470 | 13.608 | 16.772 | -0.153 | -13.447 | -0.581 |
| -5.978 | -8.234 | 23.988 | -9.518 | -9.812 | 27.894 | 19.047 | 14.269 | 24.726 | 27.228 | 25.390 | 6.739 | 24.967 | 26.031 |
| -9.470 | -17.875 | -13.642 | 17.781 | 16.233 | 23.119 | -24.138 | -3.079 | 18.583 | 6.523 | 8.718 | 12.111 | -4.227 | 2.019 |
| 7.318 | 23.143 | 27.330 | 26.068 | -23.881 | 4.275 | -6.514 | 5.769 | -9.140 | 8.832 | 4.151 | 17.835 | 15.288 | 25.010 |
| 22.059 | 18.692 | -6.939 | 20.134 | 11.927 | 20.300 | -1.674 | 4.661 | 20.607 | 24.153 | 11.375 | 13.476 | 18.629 | 12.810 |
| -4.694 | 6.054 | 20.690 | 27.179 | 9.953 | 18.397 | 6.982 | 24.757 | 6.492 | 19.238 | 0.274 | 10.793 | 19.719 | 23.827 |
| 18.146 | -2.410 | -6.428 | -15.193 | -14.373 | -15.004 | 11.902 | -2.549 | 25.882 | 22.827 | 23.566 | -23.257 | 15.805 | -4.481 |
| 7.606 | -21.549 | 17.005 | 8.431 | -15.995 | 8.732 | 2.368 | 23.259 | -4.073 | 16.936 | 22.368 | 6.183 | 21.540 | 11.940 |
| 14.679 | -18.615 | -7.689 | 15.744 | 23.840 | 21.369 | 21.049 | 7.622 | 28.997 | 29.048 | 13.882 | 8.778 | 24.055 | 12.034 |
| 24.750 | 10.629 | 10.255 | -23.613 | -18.551 | -3.347 | -26.149 | 28.816 | 13.400 | 23.024 | 21.435 | 21.427 | 3.852 | 27.661 |
| 16.182 | -19.760 | -0.766 | -5.009 | -25.475 | 29.493 | -17.554 | 26.765 | 17.565 | -14.265 | 15.728 | 25.005 | 25.987 | -10.093 |
| -6.198 | -16.616 | -24.223 | 21.805 | 9.409 | 2.386 | 15.697 | 28.346 | 23.775 | -1.327 | 24.481 | 4.877 | 17.099 | 3.905 |
| 4.134 | 8.345 | 6.138 | 4.336 | -21.017 | 1.170 | -1.935 | 5.605 | 19.772 | -31.061 | 19.626 | 22.527 | 26.538 | 23.402 |
| -2.656 | -1.428 | -2.283 | -8.384 | -1.098 | -4.257 | -23.419 | -28.880 | 21.569 | -1.006 | 8.344 | -1.969 | 18.666 | -24.605 |
| -5.303 | 14.902 | 11.462 | -0.348 | 25.703 | 2.949 | -20.826 | 16.003 | 18.946 | 18.970 | 10.381 | 26.055 | -0.915 | 1.651 |
| 8.484 | -14.294 | 14.063 | 12.054 | 17.947 | -27.842 | -20.017 | -2.973 | 22.853 | 19.923 | -9.474 | 19.746 | 13.492 | 20.063 |
| -13.524 | 14.133 | 5.269 | -15.127 | 14.969 | 16.530 | 16.619 | -9.395 | 5.341 | 24.875 | 24.480 | 23.524 | 23.023 | 0.539 |

| | | | | | | | | | | | | | |
|---|---|---|---|---|---|---|---|---|---|---|---|---|---|
| 17.586 | 11.695 | -23.448 | 21.571 | 20.649 | 18.340 | -25.016 | 26.575 | 9.518 | 7.487 | -18.653 | -16.383 | 14.096 | 26.178 |
| 0.711 | -22.885 | -13.525 | -19.469 | 9.848 | -0.831 | -18.642 | 6.285 | 19.134 | 23.225 | 23.321 | 21.972 | 25.755 | 16.642 |
| 16.337 | 28.704 | -20.509 | 19.958 | 0.937 | 29.309 | 29.381 | -18.077 | 17.215 | 14.992 | -18.193 | 11.069 | 23.310 | -3.919 |
| 20.872 | 9.362 | -10.410 | 25.083 | 24.253 | 11.119 | -9.015 | 23.787 | 26.305 | -16.047 | 6.281 | 15.175 | 21.405 | 7.432 |
| 23.070 | 25.831 | -22.169 | 23.665 | -7.300 | 23.596 | 24.132 | 22.348 | 19.752 | 22.655 | -20.730 | 25.185 | 14.472 | 4.887 |
| 14.234 | 23.668 | 0.331 | 9.231 | 15.034 | -7.453 | 10.275 | -20.498 | 8.070 | 20.485 | -10.178 | 19.849 | 0.194 | 1.251 |
| 6.625 | 19.457 | 12.624 | -13.189 | 18.848 | 19.879 | 12.285 | 8.606 | 4.624 | 27.821 | -11.516 | 5.446 | 8.899 | 17.899 |
| 23.939 | -29.136 | 5.602 | 2.414 | 4.550 | 27.677 | 4.763 | 13.283 | 9.551 | 19.601 | -17.536 | 14.261 | -17.297 | 23.588 |
| -11.206 | -4.481 | -4.961 | -17.270 | 15.176 | 17.619 | -15.132 | 25.373 | -0.822 | 25.854 | -21.691 | 23.005 | -6.892 | 20.427 |
| 2.407 | 15.299 | 2.424 | 24.134 | 15.096 | 11.580 | 7.883 | 27.985 | -23.610 | 5.857 | 1.902 | 24.008 | 20.796 | -0.430 |
| 2.781 | -12.166 | -8.189 | 14.063 | 11.614 | 9.935 | -22.133 | 29.358 | 14.410 | -27.814 | 22.120 | 14.785 | 20.481 | -20.101 |
| -25.306 | -21.368 | 12.566 | 4.976 | -9.449 | -19.890 | -17.790 | 7.906 | 21.620 | -4.483 | 12.075 | 11.671 | 16.642 | 19.898 |
| 25.384 | 16.488 | -9.738 | 25.843 | 2.608 | -26.202 | 10.363 | 29.916 | 28.872 | 17.143 | 25.113 | 22.568 | 14.062 | -17.985 |
| 4.758 | 17.058 | -17.956 | 22.954 | 14.833 | 15.333 | -14.150 | 2.954 | 12.973 | 26.265 | 23.013 | -3.346 | -8.581 | 12.517 |
| 3.228 | -16.448 | -20.495 | 2.872 | -8.497 | -16.398 | 30.120 | 28.212 | 26.576 | 6.106 | 18.737 | 25.849 | 2.465 | -2.148 |
| -17.281 | 22.100 | -13.234 | 1.209 | 7.805 | 24.629 | -22.617 | -10.673 | 24.423 | 11.184 | 24.734 | 18.566 | 6.213 | 21.057 |
| 4.099 | -0.920 | 7.239 | -27.565 | -11.060 | 3.433 | -23.991 | 6.807 | 4.005 | -30.672 | 9.924 | -3.504 | 15.361 | 9.811 |
| 4.777 | 20.177 | 6.642 | 9.865 | 17.788 | 5.593 | 24.128 | -3.021 | -8.609 | 24.612 | 2.934 | 3.129 | 20.927 | -11.011 |
| -23.746 | 11.661 | 28.212 | 16.274 | 0.960 | -6.457 | -11.412 | -27.718 | 25.728 | 18.167 | -5.983 | 23.429 | -25.795 | -7.123 |
| 5.125 | -24.783 | 25.040 | -3.288 | 23.436 | 18.658 | -16.304 | -1.152 | 22.647 | 25.680 | 10.485 | -4.625 | -17.788 | 14.731 |
| -14.070 | 16.118 | -25.706 | 28.965 | 1.855 | 11.220 | -24.949 | -3.740 | 24.783 | 18.726 | 22.474 | -13.207 | 9.181 | 3.527 |
| -15.181 | 25.701 | 0.063 | -19.325 | -0.265 | 17.520 | 22.498 | -25.015 | 21.926 | 29.047 | 24.581 | -13.570 | 5.514 | 21.443 |
| 11.219 | -29.733 | -7.099 | 14.873 | 4.246 | 15.654 | 1.760 | 23.811 | -0.565 | 14.007 | 21.388 | 21.853 | -10.319 | 18.893 |
| -9.514 | -15.861 | -6.495 | -6.627 | 7.005 | -27.852 | 27.082 | 19.313 | 28.099 | 16.365 | 19.953 | 20.125 | 1.496 | 15.632 |
| 23.372 | -8.757 | -5.585 | -25.287 | 1.604 | 23.086 | 26.120 | 21.127 | 6.787 | 28.326 | 7.062 | 27.119 | 7.277 | 3.466 |
| -2.184 | 23.895 | 3.653 | 22.599 | -13.204 | -24.966 | 19.819 | 22.409 | 21.073 | -4.150 | -23.731 | 25.053 | 5.934 | 13.120 |
| 10.873 | 17.459 | -4.018 | -8.578 | -8.493 | 29.834 | 11.275 | -20.238 | 23.770 | 21.384 | 8.865 | 11.122 | 14.934 | -18.985 |
| 15.502 | 22.328 | -1.120 | -17.325 | 20.388 | 16.975 | 27.769 | -11.224 | -7.861 | 14.166 | 18.165 | 23.949 | 13.192 | -6.539 |
| -9.110 | 19.984 | -19.606 | 1.571 | -9.303 | -6.367 | 21.096 | 24.981 | 20.273 | 4.630 | 11.400 | 22.976 | 22.685 | -16.399 |
| 19.564 | -15.088 | 17.019 | 18.807 | -21.356 | 20.854 | -18.385 | 7.494 | 22.732 | 27.059 | -26.504 | 13.670 | 20.892 | -21.951 |
| 3.475 | 11.301 | -2.373 | -10.528 | 21.126 | 28.449 | 19.476 | 14.176 | -21.219 | 18.743 | 23.106 | 22.889 | 10.760 | 8.116 |
| 25.932 | 24.706 | 3.947 | -15.399 | -26.869 | 24.575 | 27.142 | 29.199 | -25.633 | 6.755 | 15.185 | 27.706 | 23.225 | 23.977 |
| 6.456 | 17.938 | -25.631 | -14.193 | -19.160 | 12.777 | 22.464 | -24.526 | 24.262 | -3.312 | 9.877 | 26.486 | 24.230 | 22.333 |
| -2.397 | -10.605 | -7.270 | 24.306 | 13.278 | 17.653 | 25.159 | 25.166 | 21.592 | 5.624 | 24.762 | 7.785 | 15.510 | 17.754 |
| 5.626 | 8.417 | 26.267 | -26.450 | 8.835 | 21.034 | -7.803 | 19.527 | 17.020 | 21.741 | 21.630 | 28.935 | 24.932 | -5.882 |
| -22.916 | 23.264 | 9.743 | -29.721 | -4.737 | -25.830 | -28.172 | -9.703 | 8.104 | 11.025 | 8.198 | 17.853 | -6.149 | 8.972 |

| | | | | | | | | | | | | | |
|---|---|---|---|---|---|---|---|---|---|---|---|---|---|
| -15.220 | 21.834 | 2.849 | 14.274 | -14.436 | 3.638 | 7.450 | 31.018 | 20.187 | 17.414 | 11.591 | 11.872 | 10.519 | 14.086 |
| -22.031 | -26.062 | -21.998 | 20.971 | -8.349 | 2.957 | 29.502 | 29.923 | 21.210 | 16.586 | 24.169 | 5.209 | 7.004 | -0.019 |
| 14.853 | -19.199 | 28.121 | 10.186 | 1.202 | 26.737 | -6.415 | 10.739 | 17.419 | 27.864 | 17.306 | 1.267 | -2.829 | -3.612 |
| 21.126 | -8.283 | -4.532 | -5.635 | 24.610 | 10.489 | -10.287 | 15.425 | -0.334 | -0.927 | 19.079 | 24.044 | -9.677 | -16.074 |
| 28.074 | 14.633 | 12.952 | 8.730 | -12.328 | 5.502 | 18.466 | -27.647 | 25.037 | -2.164 | 5.168 | 21.984 | 4.445 | 24.926 |
| 1.029 | 19.248 | 0.741 | -5.040 | 22.575 | 0.101 | -24.239 | 28.986 | 22.255 | -3.475 | 17.728 | 16.859 | 1.286 | 13.632 |
| -29.038 | 26.228 | -14.665 | -11.717 | -16.014 | 8.722 | 21.100 | 25.205 | 8.592 | 14.425 | 26.062 | 19.659 | 25.508 | 24.388 |
| -16.321 | -4.808 | -23.252 | 27.235 | 12.961 | -17.144 | 23.763 | 13.612 | 19.538 | -14.892 | 25.795 | 10.981 | 20.034 | 21.008 |
| 10.650 | -18.455 | 19.785 | -24.620 | 2.866 | 20.865 | 13.622 | 25.827 | 1.040 | 24.727 | 15.491 | 7.059 | 13.840 | 11.276 |
| 4.333 | 19.648 | 8.774 | -23.280 | -17.695 | 22.190 | 8.561 | 18.263 | 19.388 | 7.186 | 21.708 | 13.098 | 10.014 | 21.948 |
| 12.286 | 16.038 | -10.288 | -12.348 | -23.993 | 25.996 | 6.949 | 21.026 | 18.015 | 15.770 | 18.454 | 28.220 | 23.607 | 24.968 |
| 11.627 | 22.661 | 16.745 | 29.769 | 22.860 | 28.346 | 14.805 | 20.139 | 25.929 | 27.653 | 9.664 | 19.400 | 21.951 | 11.975 |
| -24.175 | -4.651 | 1.885 | -19.771 | -22.706 | 23.057 | -23.224 | -19.054 | 22.195 | 4.245 | 25.038 | 20.191 | 17.973 | 2.418 |
| 22.695 | -5.219 | -11.564 | -6.890 | 18.788 | 5.631 | 4.313 | 29.201 | 17.981 | 18.252 | 23.402 | 17.736 | 12.085 | -11.554 |
| 10.432 | 13.272 | 5.080 | -20.840 | -6.265 | 4.569 | 5.843 | 27.297 | 19.553 | 25.807 | 26.631 | 21.609 | 21.124 | 9.097 |
| -13.105 | -11.477 | -2.543 | 17.067 | -7.306 | 16.517 | -3.167 | 18.873 | 21.378 | 10.401 | 20.803 | 7.016 | 8.176 | 6.437 |
| 2.688 | -20.880 | -9.177 | 10.098 | -18.371 | -10.203 | -20.036 | 5.050 | 25.090 | 16.209 | 8.151 | 16.011 | 27.941 | -0.493 |
| 5.106 | -20.304 | -28.049 | -29.936 | -25.374 | 3.066 | 24.854 | 7.795 | 23.491 | 24.225 | 19.022 | -28.747 | 11.830 | -13.680 |
| -6.918 | 20.752 | 26.514 | 25.600 | -14.255 | -17.873 | -17.240 | -12.883 | 11.416 | -25.902 | 17.996 | 13.551 | 24.191 | 23.458 |
| 7.507 | -6.181 | 25.912 | -4.257 | -11.472 | 1.729 | -21.971 | -24.197 | 13.856 | 11.604 | 11.063 | 10.089 | 21.112 | 25.308 |
| 14.874 | 26.308 | 14.567 | -28.421 | 14.702 | 22.025 | -19.625 | -27.351 | 4.978 | 2.816 | 19.349 | 17.373 | 21.611 | -14.274 |
| -28.815 | 22.022 | -28.992 | 18.362 | -23.828 | 26.309 | -20.333 | -3.120 | 9.998 | -9.421 | 24.857 | 21.190 | 16.312 | 27.100 |
| 1.431 | -12.038 | -4.305 | 7.602 | -24.892 | -12.577 | 21.453 | -29.908 | 9.051 | -1.505 | 9.470 | 8.463 | 12.922 | -5.952 |
| 11.640 | 17.348 | 23.108 | 22.786 | 24.843 | 24.216 | -14.385 | 25.063 | -3.446 | -0.374 | 10.050 | 30.700 | -5.579 | 9.313 |
| 0.381 | -24.636 | -20.899 | 25.920 | -0.397 | 0.237 | -9.192 | 17.052 | 21.609 | -9.770 | 15.164 | 19.897 | 28.113 | 19.245 |
| 21.291 | 16.087 | 6.703 | 0.676 | 26.086 | 19.450 | 25.001 | -10.792 | 22.602 | 9.506 | 7.995 | 22.695 | 14.383 | 17.911 |
| 29.149 | -3.114 | -0.449 | -11.523 | -4.546 | 5.422 | -16.132 | -15.966 | -8.880 | 16.570 | 21.794 | 1.179 | 25.341 | 15.204 |
| -13.046 | 9.369 | -5.533 | 20.637 | 20.728 | 3.517 | -25.040 | 7.845 | 6.638 | -7.214 | 19.778 | 29.604 | 13.499 | 8.812 |
| -15.184 | 17.596 | -13.136 | 14.606 | 12.597 | 13.465 | -17.523 | -21.849 | 24.639 | 10.740 | 17.543 | 21.171 | -1.248 | 26.749 |
| 9.025 | -14.628 | -17.261 | 28.688 | 9.044 | 26.843 | 4.527 | -27.684 | 24.926 | 23.703 | 16.974 | 18.094 | 7.133 | -2.295 |
| 7.040 | -20.450 | -3.409 | 10.836 | 22.695 | -0.225 | -2.878 | 14.087 | 13.802 | 24.051 | 27.148 | 25.383 | 27.763 | -13.942 |
| 20.659 | -23.336 | -16.673 | 21.847 | -6.191 | 11.825 | 18.035 | 29.184 | -25.217 | 6.212 | 25.160 | 29.888 | 21.376 | -10.604 |
| -12.930 | -15.690 | 9.541 | -18.601 | -5.536 | 3.235 | 22.502 | 26.833 | 17.574 | -8.901 | 19.197 | 4.925 | 28.363 | 16.708 |
| 10.545 | 11.565 | -13.820 | -1.700 | 29.199 | 28.220 | -24.993 | 15.021 | 14.598 | 4.521 | 12.248 | 10.450 | 26.538 | 27.559 |
| -29.847 | -8.720 | -18.773 | -1.266 | 1.485 | 24.558 | 7.984 | 22.514 | -14.919 | 22.833 | 8.434 | 14.788 | 9.894 | 18.294 |
| -24.657 | 5.387 | 12.194 | 24.452 | -22.479 | -9.908 | 13.019 | 13.151 | 9.618 | 11.531 | 6.792 | 12.454 | -6.575 | 22.468 |

| | | | | | | | | | | | | | |
|---|---|---|---|---|---|---|---|---|---|---|---|---|---|
| 14.133 | -0.767 | -7.413 | 27.051 | 23.772 | -21.810 | -10.402 | 9.217 | -20.801 | 18.886 | -29.327 | 20.863 | 23.211 | 20.832 |
| 25.106 | 24.318 | -12.688 | 22.699 | -13.512 | 20.571 | -27.583 | 8.544 | 17.887 | 20.440 | 29.777 | -1.446 | 11.598 | 10.401 |
| 24.068 | -9.134 | 7.403 | -24.815 | -21.464 | 26.210 | 24.946 | 6.149 | 25.994 | 13.328 | 14.485 | 31.743 | 21.519 | 19.677 |
| 30.153 | -0.037 | -17.895 | -5.871 | -3.498 | 23.325 | 25.669 | 20.731 | -6.065 | 8.710 | 12.743 | 13.210 | 5.206 | 16.940 |
| 17.471 | -26.282 | -20.471 | -22.076 | 6.577 | 3.737 | 27.158 | 8.055 | -7.002 | 10.127 | 28.465 | 16.103 | 13.232 | 11.055 |
| 23.617 | -14.954 | 4.541 | 3.279 | -7.476 | 24.711 | -11.620 | -16.217 | 24.565 | 12.132 | 20.800 | 20.349 | 27.271 | 3.631 |
| -10.654 | -6.270 | 10.576 | 9.790 | 21.447 | 2.960 | -4.821 | 21.308 | 21.055 | 14.809 | 17.495 | 29.988 | 25.199 | 12.185 |
| 3.086 | -1.407 | -1.153 | -9.151 | 17.043 | -3.974 | -19.667 | 16.813 | -1.575 | 8.290 | 13.808 | 22.978 | 19.035 | 12.085 |
| 10.865 | -8.246 | -17.283 | 19.906 | 25.418 | 13.410 | 17.075 | -25.434 | 0.506 | 11.542 | 18.786 | 28.432 | -0.004 | 24.006 |
| -11.300 | 5.114 | -11.352 | 15.420 | 27.787 | 25.947 | -17.032 | 22.245 | 24.229 | 7.493 | 6.656 | 16.764 | 7.337 | 17.634 |
| -4.524 | 15.423 | 15.653 | -22.244 | -17.329 | 7.574 | 12.510 | -17.986 | 25.338 | 5.289 | 23.626 | 15.598 | 27.071 | 10.205 |
| -4.217 | -14.653 | 1.490 | 25.089 | 21.639 | -3.926 | 20.530 | 20.051 | 21.548 | -8.783 | 18.771 | 17.493 | 29.397 | 25.700 |
| 13.147 | -25.310 | 26.834 | -17.115 | 14.307 | -17.065 | 25.304 | -9.131 | 0.925 | 24.228 | 24.199 | 5.030 | 22.180 | 11.602 |
| 0.685 | 11.722 | -1.966 | 1.616 | 17.322 | -15.569 | -19.445 | 19.140 | 23.077 | 2.495 | 29.615 | 21.300 | 28.103 | 18.610 |
| -31.759 | -20.236 | -0.034 | -14.816 | -8.060 | 8.704 | -4.028 | -18.002 | 8.105 | 19.704 | 25.245 | 9.227 | 17.271 | 16.232 |
| 9.340 | 25.082 | -19.567 | -0.364 | 20.083 | 9.128 | 24.509 | 5.972 | -8.910 | 25.295 | 14.033 | 24.278 | 23.525 | 15.999 |
| 21.895 | 19.274 | -29.562 | -7.418 | 26.195 | 8.065 | -10.612 | 14.109 | -13.022 | 8.606 | 13.071 | 18.371 | 16.175 | 12.922 |
| 12.031 | -21.624 | -2.484 | -28.025 | 0.441 | -14.821 | 21.889 | -26.754 | 19.321 | -10.465 | 15.723 | 22.051 | 9.489 | 6.157 |
| 16.462 | -24.495 | 12.034 | -10.405 | 22.001 | 9.661 | -3.065 | 25.996 | -7.591 | 21.712 | 10.629 | 5.787 | -16.393 | 15.329 |
| 17.344 | -2.999 | -11.064 | -13.875 | -24.660 | 18.511 | -18.056 | 2.651 | 24.558 | 6.603 | 14.220 | 23.444 | 25.453 | 17.975 |
| -2.302 | -18.843 | -8.832 | 29.323 | -15.056 | -9.104 | -22.326 | 2.082 | 23.326 | -0.892 | 22.879 | 19.192 | -1.859 | 21.122 |
| -13.094 | -25.446 | 8.274 | 21.646 | 29.519 | 16.851 | 9.445 | -24.222 | 20.901 | -4.539 | 29.667 | 21.161 | 26.539 | 12.819 |
| 9.071 | -9.642 | 17.525 | 29.902 | 23.562 | 21.429 | 7.228 | 3.995 | 29.000 | 18.064 | 20.872 | 10.378 | 20.421 | -8.349 |
| 27.261 | 1.867 | -3.545 | 1.361 | 30.761 | 22.457 | -13.236 | -3.587 | -1.403 | 10.751 | 23.861 | 30.739 | 17.628 | 28.189 |
| 22.178 | -1.992 | 4.277 | 26.142 | -18.163 | 22.109 | -7.414 | -26.076 | 20.313 | -3.809 | 14.939 | 29.445 | 10.840 | 7.834 |
| -0.630 | -16.685 | -20.825 | 18.789 | 6.721 | -6.120 | -1.267 | -7.167 | 11.076 | 8.056 | 4.212 | 22.131 | 27.535 | 27.940 |
| 18.930 | -15.305 | -17.451 | -13.237 | 25.706 | -18.006 | 22.015 | -11.705 | 28.540 | -3.219 | 23.486 | 9.976 | 13.743 | -22.418 |
| -10.673 | -14.615 | 25.749 | -3.526 | -0.997 | -8.736 | -8.761 | -8.403 | -13.060 | 20.575 | 14.105 | 19.653 | 3.885 | 22.204 |
| 6.975 | -10.679 | -21.476 | -7.412 | 21.630 | 9.467 | 2.379 | 16.847 | 24.700 | -0.535 | 30.153 | 4.852 | 26.178 | 27.303 |
| -9.278 | 4.783 | -18.382 | -4.162 | 10.686 | 20.505 | -13.463 | 10.835 | -25.363 | 7.581 | 24.707 | 24.468 | 12.501 | 28.827 |
| -3.198 | 6.478 | 20.241 | -20.535 | -13.377 | 2.384 | -2.923 | 7.248 | 21.608 | 14.836 | 20.781 | 30.895 | 22.212 | 13.994 |
| -14.242 | -10.847 | -8.135 | 22.612 | 24.861 | 12.039 | 0.340 | 25.366 | 9.711 | 3.738 | 8.733 | 15.893 | 6.640 | 16.001 |
| -8.032 | -0.064 | 21.666 | 28.374 | 13.573 | 11.044 | 14.303 | -15.057 | 29.860 | 5.143 | 15.885 | 18.125 | 16.846 | 14.864 |
| 11.650 | -24.894 | -0.168 | 24.703 | -22.675 | -0.253 | -8.525 | 2.483 | 25.981 | 22.715 | -32.180 | 11.824 | 28.093 | 31.253 |
| -5.649 | -0.406 | 9.429 | -0.732 | 15.447 | 14.477 | -25.018 | -11.884 | 20.247 | 2.267 | 13.428 | 19.845 | 31.401 | 26.432 |
| -22.251 | 10.953 | -5.342 | -9.698 | 18.899 | 21.735 | -19.801 | 15.351 | 22.829 | 26.432 | 18.167 | 10.230 | 15.767 | 19.347 |

| | | | | | | | | | | | | | |
|---|---|---|---|---|---|---|---|---|---|---|---|---|---|
| -18.233 | 1.127 | -0.005 | 11.934 | -4.831 | 8.978 | -6.574 | 25.868 | 8.344 | 23.398 | 26.149 | 10.841 | 29.487 | -9.530 |
| 25.353 | 16.329 | -7.374 | 9.456 | 1.503 | 14.332 | 2.775 | -2.033 | 23.945 | -6.001 | 21.338 | 25.490 | 18.854 | -29.776 |
| 20.386 | -2.048 | 24.812 | -27.116 | 7.587 | 6.640 | -25.302 | 21.576 | 19.161 | 13.250 | 28.931 | 6.614 | 7.329 | 29.724 |
| -20.213 | -6.174 | -16.143 | 10.395 | -23.385 | 23.903 | -11.400 | 20.324 | 25.871 | 12.288 | 16.970 | 27.835 | 19.742 | 24.610 |
| -25.499 | 1.402 | -19.655 | 8.052 | 18.472 | 17.413 | -18.314 | -2.424 | -29.884 | -20.891 | 22.916 | 7.958 | 24.609 | -12.206 |
| -14.189 | -4.671 | 23.104 | 4.031 | -17.534 | 17.880 | -8.187 | -21.164 | 6.627 | 7.273 | 19.951 | 30.185 | 10.660 | 28.457 |
| 18.875 | -1.130 | 2.738 | -24.058 | 17.209 | 10.562 | 20.520 | -9.593 | 22.911 | 19.442 | 22.784 | 4.236 | 25.938 | 4.899 |
| 6.026 | 11.814 | -24.138 | 20.712 | 9.800 | -10.158 | 2.682 | 22.571 | 19.836 | 22.562 | 28.263 | 1.731 | 28.837 | 31.056 |
| 10.421 | -5.902 | -10.964 | -14.805 | -10.105 | -22.294 | -14.796 | 8.210 | 4.486 | 22.050 | 9.525 | 9.253 | 13.628 | 13.694 |
| -19.205 | 15.313 | -25.847 | 20.065 | 20.211 | 7.896 | -0.625 | 7.503 | 7.916 | 10.976 | 18.021 | 23.549 | 23.171 | 29.391 |
| -8.659 | 3.089 | 5.772 | 1.316 | -12.899 | 8.644 | 10.001 | 17.643 | 22.525 | 8.117 | 29.950 | 3.685 | 1.875 | 25.643 |
| -2.065 | -9.459 | -19.260 | 27.018 | -25.489 | 6.441 | -7.938 | -5.144 | 30.372 | -8.889 | 15.101 | 19.000 | 24.986 | 8.323 |
| -25.777 | 12.796 | 27.956 | -1.112 | 22.324 | 11.934 | -2.583 | -4.699 | 28.151 | 25.300 | 25.276 | 29.970 | 26.915 | 26.496 |
| -15.922 | -1.492 | -18.586 | 28.211 | -4.871 | 15.130 | -17.532 | -8.089 | -0.413 | 22.927 | 28.688 | 27.829 | 12.421 | 11.407 |
| -7.756 | 18.576 | 19.407 | -12.704 | 26.971 | 19.157 | 18.641 | -9.353 | 29.918 | 21.256 | 14.562 | 22.116 | 7.685 | 27.277 |
| 10.013 | 13.585 | -11.687 | 12.833 | 18.255 | 21.527 | 11.620 | -9.651 | 19.001 | 4.631 | 17.028 | 10.395 | 23.888 | -19.740 |
| -5.747 | 8.376 | 4.882 | -28.002 | 21.007 | 27.253 | 1.443 | 17.077 | 8.798 | 9.927 | 10.108 | 25.925 | 21.318 | 20.304 |
| 19.642 | 7.630 | 6.077 | 25.801 | -5.843 | 16.754 | -10.571 | 8.565 | 28.804 | 1.325 | 4.093 | 20.156 | 29.889 | 19.601 |
| 25.907 | -2.434 | 22.114 | 10.978 | -6.367 | 24.279 | -21.327 | -23.524 | 25.472 | 26.371 | 18.785 | 18.608 | 28.051 | 9.667 |
| -3.941 | 4.051 | -7.618 | -23.514 | -15.460 | 17.777 | 24.474 | 21.972 | 15.253 | 22.361 | 17.025 | 25.084 | -22.226 | 26.913 |
| 13.662 | -7.927 | -8.737 | -3.049 | 23.703 | 25.312 | 10.905 | -0.063 | 25.044 | 21.525 | 27.325 | -1.132 | 30.675 | 22.825 |
| 5.831 | -3.966 | -16.345 | 24.118 | 13.998 | 5.052 | -9.927 | -21.118 | 6.716 | 12.078 | 24.700 | 5.603 | 0.294 | 18.259 |
| 7.209 | -21.922 | -21.643 | -9.207 | -1.881 | 26.039 | -1.103 | 16.723 | 5.356 | 13.592 | 13.833 | 21.107 | 20.120 | 9.976 |
| -8.714 | 6.631 | -23.675 | -20.010 | 7.388 | 9.787 | 16.170 | 0.204 | 22.870 | 15.992 | 29.801 | 16.748 | 22.803 | 14.902 |
| 2.570 | 0.099 | 14.771 | -11.854 | 6.312 | 3.704 | -7.841 | 16.198 | 26.413 | 24.980 | 9.761 | 24.927 | 28.527 | 27.813 |
| -0.768 | -26.895 | 18.184 | -20.526 | -3.689 | 15.674 | 25.590 | -17.857 | 19.035 | 18.120 | -0.542 | 26.802 | 24.283 | 23.960 |
| -9.975 | 10.726 | -29.580 | -15.374 | 22.581 | 22.748 | -21.613 | 6.065 | 29.448 | -2.712 | 28.830 | 8.203 | 8.210 | 18.378 |
| -1.849 | -25.345 | -21.125 | -0.956 | -12.768 | -23.680 | -25.841 | 4.174 | 21.461 | -0.783 | 19.512 | 22.576 | 14.709 | 16.098 |
| 19.298 | -5.071 | -28.739 | -19.080 | -9.731 | -7.667 | -4.248 | 9.976 | 13.774 | 13.781 | 9.374 | 20.292 | 27.923 | 14.800 |
| -12.024 | 6.657 | -19.599 | -7.161 | 25.053 | 0.814 | 25.965 | 14.024 | 29.852 | 27.332 | 23.606 | 18.835 | 8.971 | 25.958 |
| 26.728 | -11.850 | 16.767 | 20.938 | 15.282 | 13.373 | -18.905 | -6.386 | 0.025 | 24.452 | 17.650 | -2.735 | 17.925 | 26.723 |
| -13.719 | 9.453 | -14.530 | 26.867 | -27.796 | -25.526 | 23.535 | 15.752 | 3.973 | 16.333 | 11.236 | 26.016 | 11.852 | 12.046 |
| 1.910 | 7.399 | -28.200 | -17.731 | 21.193 | 8.545 | -11.252 | 17.454 | -24.829 | 13.252 | 3.366 | 28.336 | -27.374 | 20.624 |
| 20.691 | -9.963 | -11.486 | 19.946 | 25.073 | -6.355 | 6.838 | 22.227 | 6.725 | 20.411 | 16.817 | 22.813 | 8.616 | 16.054 |
| -14.355 | 5.087 | 20.185 | -24.918 | -5.511 | -18.834 | -20.108 | -0.680 | -1.438 | -1.396 | 19.018 | 23.481 | -5.320 | 27.742 |
| -26.119 | 3.846 | 14.097 | -9.880 | 28.019 | -19.943 | 14.399 | 11.757 | 19.243 | 20.105 | 23.226 | 13.600 | 20.209 | 21.603 |

| | | | | | | | | | | | | | |
|---|---|---|---|---|---|---|---|---|---|---|---|---|---|
| -4.770 | -11.184 | -23.453 | -14.129 | -19.528 | 19.352 | -1.383 | -25.436 | 29.084 | 6.197 | 15.486 | 3.611 | 29.047 | 28.905 |
| -2.128 | 0.174 | -25.560 | -0.655 | 0.612 | 9.624 | -4.427 | -22.346 | 11.043 | 26.403 | 24.555 | -26.244 | -10.414 | 18.842 |
| 8.190 | -20.794 | -2.361 | -14.843 | 27.798 | -5.381 | -18.472 | 24.875 | 13.184 | -23.015 | 21.334 | 4.132 | 23.001 | 27.843 |
| 7.155 | -5.215 | 27.111 | 26.232 | 7.151 | -21.729 | 15.739 | -13.763 | 23.311 | 9.726 | 12.085 | -2.048 | 9.341 | 24.693 |
| -3.527 | 15.078 | -15.365 | 11.935 | -18.956 | 25.539 | 1.544 | 0.418 | 11.692 | 7.105 | 6.216 | 18.903 | -10.028 | 22.313 |
| -0.525 | -8.641 | -1.359 | -25.116 | -24.245 | 13.020 | -17.952 | 7.549 | 24.253 | -10.000 | 16.177 | 7.726 | 29.077 | -11.728 |
| 3.490 | 9.036 | 17.519 | -15.155 | 19.031 | 20.540 | 16.487 | -0.522 | 22.801 | -16.447 | 11.048 | 25.642 | 21.923 | 8.509 |
| 1.484 | 24.008 | -23.971 | 7.306 | -14.761 | -10.733 | 10.299 | 6.500 | 18.441 | 14.624 | 8.064 | 15.515 | 15.636 | 23.853 |
| 18.066 | -9.705 | -20.162 | -1.946 | 1.453 | -16.262 | 12.788 | 20.218 | -26.377 | 22.080 | -2.030 | 12.242 | 24.928 | 19.949 |
| -12.824 | 0.974 | -22.303 | 18.592 | 20.302 | -3.635 | -2.548 | 23.784 | 7.413 | 4.909 | 23.699 | 4.344 | 6.689 | -26.612 |
| -22.795 | 6.687 | 15.036 | -22.860 | 20.752 | -27.727 | -0.636 | 10.467 | 23.978 | -14.712 | 28.341 | 11.243 | 14.306 | 14.360 |
| 1.882 | 9.342 | -16.909 | 5.267 | 27.031 | 3.657 | -5.847 | 12.290 | 8.320 | 8.870 | 12.282 | 24.963 | 20.033 | 12.740 |
| -10.241 | -0.305 | 23.724 | 7.689 | -21.166 | -24.372 | -14.044 | 1.182 | -24.092 | 17.639 | 21.201 | 10.245 | 13.035 | 24.959 |
| -1.530 | -10.384 | 6.563 | -6.829 | 18.423 | 24.276 | 25.572 | -21.294 | -11.906 | 2.574 | 14.767 | 14.822 | 17.466 | 23.751 |
| 5.164 | 21.625 | 16.983 | -19.944 | -16.764 | 22.431 | 9.928 | -4.411 | 19.754 | 12.628 | 4.124 | 19.132 | 0.614 | 15.759 |
| -22.147 | 11.422 | -8.674 | -17.897 | 24.908 | 10.946 | -8.573 | 22.168 | 6.745 | 4.539 | 25.426 | -12.373 | 19.374 | 23.232 |
| 4.342 | -5.707 | -4.998 | 14.050 | 19.174 | -19.371 | -16.373 | 27.236 | 18.407 | 19.053 | 24.549 | 22.146 | 16.051 | 2.695 |
| -25.145 | 18.838 | -21.308 | 0.943 | 13.189 | -21.936 | 23.170 | -23.211 | -25.088 | 19.474 | 14.984 | 17.994 | 26.419 | -14.977 |
| 11.608 | -22.721 | 14.928 | -2.190 | -7.483 | 26.640 | 6.372 | 21.275 | 8.355 | -5.357 | 10.516 | 23.959 | 24.850 | 26.394 |
| -2.856 | 14.643 | 2.454 | 9.372 | 7.210 | -20.413 | -21.001 | -25.144 | 24.256 | 29.603 | 13.766 | 3.371 | 13.347 | -9.294 |
| -8.611 | 25.718 | -9.982 | 12.303 | 18.653 | 11.807 | 9.348 | 5.952 | 11.259 | 18.299 | 21.917 | 15.894 | 6.833 | 16.499 |
| -20.335 | 16.904 | 3.904 | 1.245 | 11.101 | 25.862 | -17.835 | -6.838 | 24.672 | 8.677 | 16.778 | 16.634 | 25.773 | -13.206 |
| 14.123 | 12.808 | -19.989 | 3.082 | 4.493 | -0.545 | 16.196 | 7.353 | 15.042 | 2.339 | 25.674 | -2.223 | 22.723 | 2.442 |
| -5.336 | 24.076 | -11.440 | 10.550 | 23.440 | 13.475 | -12.122 | 23.475 | 19.201 | 16.109 | 23.675 | 8.307 | 13.325 | 8.812 |
| -2.265 | 3.536 | -24.565 | 4.000 | 1.033 | 5.979 | 24.335 | 20.005 | -16.185 | 15.378 | 20.860 | 19.429 | 14.502 | -18.564 |
| 2.745 | -11.638 | 5.461 | -18.186 | 19.710 | -6.776 | 15.620 | 11.187 | 11.987 | 26.099 | 10.356 | 17.939 | 15.910 | 24.822 |
| -11.232 | 11.108 | 24.871 | 7.283 | 2.303 | -19.814 | 18.303 | 13.517 | 4.574 | 23.722 | 5.706 | 6.267 | 16.640 | 23.448 |
| 18.427 | -2.903 | 20.765 | -25.049 | -8.864 | -21.333 | -25.521 | 22.419 | 14.008 | 16.847 | 22.682 | 21.935 | 17.952 | 13.838 |
| -10.956 | 20.407 | -23.541 | 23.398 | -10.919 | 23.264 | -22.218 | 19.809 | -1.793 | 5.153 | 0.441 | 15.282 | 9.253 | 22.398 |
| 6.333 | -15.299 | 14.352 | 0.071 | -21.627 | 16.561 | 29.019 | -16.162 | 9.104 | 6.987 | 24.451 | 2.390 | 7.589 | -11.652 |
| 19.954 | 9.504 | -26.346 | 22.708 | -3.628 | 9.364 | 6.594 | 26.553 | 23.484 | 28.120 | 23.085 | 12.331 | 13.576 | 8.287 |
| 7.457 | -5.820 | 19.767 | 2.981 | 16.403 | -23.315 | 26.815 | 2.486 | 6.572 | 29.040 | 7.629 | 26.896 | 21.057 | -17.934 |
| 19.038 | 25.503 | -2.619 | -2.573 | -18.071 | 26.716 | -10.540 | 12.400 | 21.454 | -18.515 | 13.342 | 24.495 | 11.905 | -17.296 |
| 4.940 | -2.182 | 18.542 | 13.346 | -11.918 | 2.258 | 4.492 | 13.250 | -12.647 | 14.120 | 18.271 | 0.048 | -0.177 | -3.817 |
| -23.985 | -4.633 | -18.112 | -12.468 | -4.276 | 29.384 | -20.399 | 27.614 | 26.689 | 21.075 | 27.034 | 21.250 | 24.395 | 10.612 |
| 22.285 | -21.393 | 1.938 | -16.858 | -24.218 | 24.122 | 12.778 | 25.577 | -6.091 | 24.961 | 15.855 | 20.216 | 14.609 | 1.533 |

| | | | | | | | | | | | | | |
|---|---|---|---|---|---|---|---|---|---|---|---|---|---|
| -10.858 | 15.053 | -11.491 | 6.955 | -1.122 | 7.996 | -15.451 | 23.193 | 11.019 | 27.477 | 11.484 | 18.934 | -20.307 | 17.530 |
| -7.369 | -18.932 | 23.003 | 17.119 | 9.540 | -3.202 | -17.370 | 14.504 | 17.513 | 10.041 | 24.880 | -4.461 | -13.003 | 16.969 |
| -25.016 | 10.599 | -15.114 | 20.399 | 22.561 | 19.273 | 23.114 | -5.423 | -25.702 | 3.983 | 22.818 | 5.590 | 21.258 | -21.574 |
| -12.452 | 18.168 | 15.434 | 4.894 | 6.105 | -10.881 | -22.193 | -15.403 | 17.930 | 18.807 | 25.598 | 2.135 | 19.734 | 14.144 |
| -6.287 | -26.731 | -12.124 | 2.294 | 19.948 | 25.081 | 25.820 | 21.469 | 3.863 | 26.310 | 19.404 | 19.417 | 24.381 | 6.270 |
| -8.609 | -0.858 | 19.481 | -7.657 | 7.109 | 20.409 | 7.272 | -0.694 | 27.723 | 1.913 | 3.544 | 23.659 | 23.534 | -15.786 |
| -24.970 | -18.186 | -17.360 | -17.885 | 12.468 | 28.012 | 29.268 | 13.227 | 0.854 | 29.344 | 18.245 | 25.374 | 15.970 | 4.923 |
| 12.110 | -24.332 | -7.444 | 23.147 | 1.176 | 19.530 | 20.784 | 5.934 | 9.479 | 24.315 | 13.752 | 22.059 | 9.017 | -9.116 |
| 5.407 | -8.212 | -23.137 | 19.713 | 5.826 | 6.348 | -28.603 | -28.790 | 5.078 | -17.102 | 25.024 | 6.856 | -5.134 | -1.610 |
| -25.075 | 9.321 | -2.157 | 18.233 | 0.646 | 3.225 | -17.999 | 22.751 | -14.932 | -21.433 | 25.403 | 15.443 | 8.049 | 11.698 |
| -0.925 | 23.330 | -7.914 | 13.917 | 19.917 | 1.275 | -22.253 | 29.345 | 7.816 | 28.662 | 24.324 | -2.469 | 16.745 | -7.530 |
| -12.933 | -20.102 | -16.688 | -22.157 | -10.450 | 15.971 | -24.838 | 16.279 | 28.157 | 0.145 | 8.963 | -3.034 | 22.171 | 12.186 |
| 3.023 | -12.355 | -8.796 | 8.262 | 13.465 | 16.863 | 11.137 | 14.632 | 14.275 | 20.696 | 17.340 | 16.225 | 23.647 | 10.363 |
| -17.516 | 13.071 | -17.738 | 21.641 | 1.555 | 14.628 | -10.029 | 19.359 | 18.015 | 22.565 | 11.473 | -10.173 | -22.116 | -15.498 |
| 16.117 | 23.021 | 0.169 | 5.869 | -16.769 | -5.036 | 8.285 | 25.205 | -7.555 | 21.381 | 19.536 | 16.684 | 20.594 | 21.247 |
| -6.597 | -27.087 | -19.601 | -2.287 | -17.143 | -14.405 | 25.030 | 14.013 | 8.425 | 29.296 | 23.640 | -2.753 | 12.682 | 0.126 |
| 19.008 | -7.896 | -4.964 | 24.283 | 17.026 | 26.264 | 21.108 | 22.186 | 18.900 | 7.944 | 16.712 | 15.879 | 17.304 | -5.660 |
| 13.329 | 17.358 | 22.411 | 20.983 | 10.455 | -9.172 | -28.656 | 31.231 | -3.008 | 25.548 | 3.887 | 14.720 | 1.278 | 9.651 |
| -16.817 | 25.121 | 7.780 | 11.848 | -0.174 | -12.844 | -14.198 | 27.370 | 6.705 | -6.008 | 0.305 | 27.774 | 12.025 | 8.918 |
| 12.759 | 6.799 | -9.719 | 10.056 | -0.913 | -28.023 | -22.958 | 25.686 | 20.892 | 15.402 | 13.305 | 21.506 | 22.610 | 2.211 |
| -5.903 | 15.501 | -17.025 | 19.799 | -7.033 | -22.501 | 22.980 | 17.304 | 4.471 | -6.781 | 18.147 | -20.419 | 13.500 | 9.238 |
| -17.279 | 11.921 | 3.118 | 23.646 | 11.574 | 7.970 | 21.923 | 18.821 | 18.780 | 24.771 | -19.938 | 22.910 | 11.382 | 7.603 |
| 23.978 | -23.538 | -6.452 | -3.168 | 25.376 | -16.647 | 7.187 | 9.941 | 17.444 | -0.073 | 11.580 | 14.002 | -2.836 | -22.061 |
| -21.030 | 4.737 | 0.389 | -5.058 | -19.935 | 18.263 | -10.584 | -25.363 | -14.991 | 11.757 | 13.242 | 27.070 | 18.869 | 6.984 |
| -20.494 | -15.221 | 8.675 | -0.435 | -3.676 | -25.710 | 2.268 | 0.734 | 3.577 | 27.144 | -19.436 | 25.200 | 17.026 | 23.054 |
| 11.071 | 20.885 | 25.690 | -7.438 | 23.690 | -20.714 | 15.458 | 9.227 | 2.212 | 9.826 | -21.061 | 19.673 | 12.885 | 17.666 |
| 4.594 | -4.832 | -7.426 | -23.963 | -5.221 | -4.582 | 26.209 | 24.617 | 12.363 | 18.062 | 1.199 | 23.008 | 3.434 | -20.484 |
| -23.903 | 12.761 | 20.427 | -5.714 | 18.631 | 23.796 | 20.746 | 17.606 | 10.703 | 24.305 | -0.496 | 12.487 | 19.629 | -26.198 |
| 7.494 | -28.831 | 7.189 | 2.936 | 20.497 | -8.071 | -5.562 | 23.881 | 0.825 | 10.231 | -18.082 | 24.659 | 22.954 | -3.705 |
| -18.431 | 10.490 | 2.028 | -1.717 | 18.050 | 29.421 | 19.737 | 5.336 | -5.821 | 28.269 | -18.971 | -20.324 | -5.424 | 23.187 |
| -10.464 | 28.507 | -7.089 | 12.188 | 5.614 | 2.217 | -0.475 | -31.061 | 23.979 | -17.122 | 21.420 | 25.505 | 15.773 | 4.069 |
| 12.616 | -12.782 | 11.204 | 1.774 | 6.789 | -18.258 | -14.536 | 29.931 | 0.595 | 8.918 | -7.589 | 2.620 | 22.382 | 7.530 |
| -5.026 | 9.486 | -4.384 | 7.026 | 15.128 | -12.757 | 16.560 | 26.572 | 21.074 | -26.116 | 18.313 | 29.187 | 14.986 | 23.124 |
| -15.330 | 1.876 | -15.389 | 26.178 | 7.147 | -26.009 | 17.511 | 6.180 | 0.193 | 1.083 | 14.300 | 12.886 | -23.206 | 14.714 |
| 23.168 | 24.682 | 12.695 | -24.756 | 7.701 | -19.599 | -21.850 | -23.757 | -20.572 | 14.594 | 19.635 | 20.423 | 10.807 | 20.845 |
| -1.925 | -19.865 | -7.941 | -20.995 | 14.875 | -9.605 | 8.278 | 18.283 | -1.348 | 25.915 | -6.686 | 1.734 | 17.900 | -3.353 |

| | | | | | | | | | | | | | |
|---|---|---|---|---|---|---|---|---|---|---|---|---|---|
| 19.102 | 8.495 | -0.349 | 16.990 | -19.333 | 31.080 | -6.426 | -27.359 | -0.790 | 0.361 | 15.982 | -15.872 | 17.098 | 14.013 |
| 25.721 | -12.707 | 8.373 | -15.030 | 1.446 | 21.576 | -18.091 | -6.286 | 10.688 | 15.963 | 9.421 | 23.646 | -25.597 | 23.399 |
| 21.639 | 2.007 | 3.311 | 21.472 | 22.386 | 20.160 | 16.287 | 22.506 | 18.195 | 25.238 | 25.646 | 21.234 | 23.495 | 0.913 |
| 12.119 | 3.630 | -9.828 | 8.670 | -4.053 | 10.373 | -7.229 | 1.923 | -24.654 | 23.845 | 23.530 | 25.552 | 22.695 | 9.284 |
| 10.380 | 9.949 | -10.642 | -9.162 | 26.215 | 22.485 | 21.178 | 18.646 | 2.434 | 27.078 | 22.729 | 3.749 | 7.055 | -27.342 |
| -14.178 | 15.737 | 2.366 | 18.363 | 3.314 | -24.464 | 26.765 | 9.241 | 1.615 | -2.587 | 8.720 | 20.085 | 10.097 | 21.403 |
| -16.725 | 6.815 | 27.416 | 24.997 | -23.300 | 26.945 | -19.513 | 20.900 | -8.515 | 30.202 | 3.683 | 27.305 | 23.939 | 15.209 |
| -11.155 | 18.715 | 5.058 | 20.580 | 25.396 | -22.412 | 16.963 | 25.385 | 13.309 | 9.984 | 17.683 | 23.169 | 25.103 | 11.854 |
| 5.555 | -2.177 | -13.183 | -11.492 | 22.670 | 0.922 | 25.004 | -22.489 | 21.288 | -11.796 | 26.596 | -1.822 | 13.619 | 25.102 |
| 12.763 | 7.480 | 13.912 | -3.876 | 20.423 | 16.869 | -18.583 | -29.153 | 21.603 | 18.466 | -2.367 | 20.310 | 23.138 | 19.043 |
| -9.298 | 22.702 | -26.350 | -17.624 | -21.274 | 4.580 | -23.984 | -3.628 | 22.748 | 28.004 | -26.092 | -7.487 | 17.080 | 14.184 |
| 7.392 | -22.777 | -6.080 | 3.751 | 13.925 | 19.164 | 22.549 | 17.466 | -11.194 | 11.902 | 23.812 | 26.125 | 15.509 | 26.657 |
| 25.978 | -20.491 | -16.778 | -22.589 | -3.385 | -9.265 | -12.949 | 24.288 | 8.452 | 6.146 | -1.787 | 16.766 | 18.859 | 6.246 |
| 11.875 | -24.723 | 23.419 | 23.047 | -0.533 | 23.026 | -21.685 | 16.932 | 10.105 | 16.080 | 17.478 | 28.376 | 6.048 | -3.390 |
| 20.737 | -0.901 | 2.464 | 15.309 | 21.057 | 25.028 | -2.295 | 10.662 | -2.052 | 29.447 | 24.936 | -13.240 | 27.081 | 25.813 |
| 22.637 | 0.864 | -7.837 | -2.975 | 16.522 | -4.962 | -9.593 | 4.714 | 23.072 | -9.451 | 7.849 | 20.432 | 25.300 | 7.475 |
| -23.172 | 22.653 | 3.879 | 23.268 | 24.543 | 0.043 | 19.542 | 25.812 | 25.776 | 30.699 | 4.441 | 28.085 | 15.709 | 22.624 |
| -21.079 | -29.135 | -11.820 | -22.170 | 10.055 | 31.567 | 26.755 | 28.998 | 9.319 | 1.829 | 18.953 | 18.131 | 14.551 | 24.379 |
| -6.926 | 10.661 | -6.725 | -24.951 | 3.401 | 15.799 | 12.998 | 23.935 | 1.628 | 27.712 | 23.825 | 21.321 | 21.200 | 16.423 |
| 23.818 | 28.903 | -14.417 | -14.691 | 15.799 | 14.545 | 23.843 | 26.736 | 19.240 | 8.222 | 14.114 | 15.116 | 16.739 | 18.018 |
| -11.410 | 12.643 | -5.122 | 8.904 | 19.533 | 9.663 | -21.102 | -2.869 | -10.514 | 17.793 | -1.516 | 20.071 | 17.836 | -3.043 |
| 16.684 | 7.112 | 10.699 | -20.766 | -26.579 | -14.283 | 10.077 | 29.589 | 3.940 | 14.220 | 21.903 | 3.234 | 26.073 | 21.421 |
| 9.337 | -7.280 | -15.353 | 24.581 | 14.575 | 23.811 | -12.897 | 28.228 | 5.406 | -22.570 | -2.931 | 10.509 | 20.882 | 20.463 |
| -12.449 | 2.549 | 18.296 | 10.078 | 7.607 | -20.812 | -26.234 | 14.937 | 6.422 | 28.917 | 26.906 | -29.837 | 18.734 | 25.256 |
| 22.156 | 20.430 | 28.290 | 16.192 | 21.019 | 0.408 | -3.849 | 12.661 | 22.336 | 1.750 | 18.669 | 11.804 | 22.474 | 19.563 |
| -18.039 | 17.463 | -26.676 | -9.290 | -9.591 | -6.307 | -8.554 | 22.749 | 8.007 | 27.035 | 10.375 | -1.710 | 20.605 | -5.002 |
| 23.139 | -22.357 | 1.971 | 8.272 | 19.723 | -7.858 | -11.020 | 25.970 | -0.633 | 24.384 | 19.727 | 19.626 | 24.621 | 25.523 |
| 8.141 | 24.343 | 16.390 | -19.459 | -20.844 | -0.915 | -20.165 | -0.652 | 22.469 | 19.709 | 21.618 | -28.411 | 28.675 | -2.381 |
| -3.949 | -4.003 | 1.212 | -24.990 | 7.425 | 8.439 | 4.177 | -28.283 | 21.696 | 26.582 | -7.585 | 22.827 | 23.375 | 26.382 |
| 23.858 | -25.417 | -7.395 | 0.810 | -17.795 | -22.104 | 6.963 | 20.647 | 25.126 | 25.187 | 24.483 | 26.735 | 16.513 | -6.644 |
| 6.998 | 11.906 | -18.349 | 27.951 | -19.608 | 24.155 | 27.379 | 6.261 | 26.329 | -11.040 | 0.131 | 15.344 | 25.251 | 8.533 |
| 20.180 | 7.842 | 9.837 | 26.377 | 22.112 | 9.195 | 19.976 | 26.473 | 26.717 | 12.992 | 25.709 | 21.374 | 12.346 | 21.859 |
| 26.414 | 25.099 | -25.349 | -3.792 | 25.338 | -3.702 | 13.483 | -16.860 | -2.006 | 21.690 | 11.645 | 24.528 | 20.097 | 23.639 |
| 27.364 | 4.892 | 21.868 | 20.437 | -11.968 | -15.462 | 26.337 | -19.978 | 4.824 | 9.529 | 23.778 | 32.150 | 10.478 | 19.081 |
| 3.059 | -2.791 | 14.816 | 12.094 | 6.279 | -24.189 | 23.296 | -27.321 | 17.001 | 27.315 | -1.291 | 4.034 | 5.829 | 16.152 |
| -19.778 | 14.944 | 4.367 | 22.571 | 3.396 | 5.192 | 26.867 | 10.645 | -4.663 | 22.081 | 1.642 | 22.007 | 29.338 | 14.897 |

| | | | | | | | | | | | | | |
|---|---|---|---|---|---|---|---|---|---|---|---|---|---|
| -25.455 | 19.423 | -8.041 | 25.239 | -3.152 | 19.208 | 5.158 | 18.311 | 27.129 | -4.921 | 28.689 | 28.630 | 19.172 | 11.065 |
| 18.561 | -6.343 | -26.144 | -14.956 | 9.246 | 16.692 | 17.260 | 24.859 | 18.126 | -30.099 | 23.110 | 5.193 | 15.622 | 24.117 |
| -15.382 | 9.753 | -10.433 | -23.201 | 14.205 | -28.589 | -13.521 | -5.575 | -4.726 | 29.938 | 9.059 | 21.504 | 11.542 | -29.022 |
| 7.017 | -12.613 | -15.909 | 15.451 | -4.848 | 30.809 | -13.231 | -20.516 | 0.097 | 21.035 | 19.578 | -4.065 | 10.314 | 26.072 |
| -28.193 | 16.653 | 25.609 | 6.944 | 26.333 | 19.709 | 28.260 | 20.099 | 14.557 | 9.225 | 3.751 | 7.369 | 4.523 | 8.980 |
| -22.045 | -5.529 | 6.147 | -10.006 | -11.429 | -18.659 | 10.316 | 17.770 | -7.025 | 11.013 | 18.559 | 23.624 | 29.071 | 22.202 |
| -8.263 | 19.838 | 24.206 | 27.106 | 19.832 | 11.276 | 21.848 | 15.730 | -20.504 | -8.846 | 25.669 | -30.710 | 21.571 | 20.966 |
| -23.454 | 30.068 | -18.079 | 6.200 | -21.172 | -9.313 | 4.031 | -8.476 | 23.969 | 26.507 | 4.833 | -1.395 | 23.277 | 12.305 |
| -18.178 | 5.300 | 20.587 | 13.562 | 10.193 | -14.244 | -20.455 | -26.384 | 23.348 | 22.656 | -3.928 | 20.501 | 22.180 | 7.342 |
| -9.939 | -18.179 | -4.922 | -1.993 | 16.578 | -16.954 | -21.270 | 12.530 | -1.124 | 3.570 | 12.194 | 21.932 | 8.231 | 28.620 |
| -14.948 | 20.382 | 13.793 | 12.877 | 13.097 | -19.670 | -27.716 | -17.038 | 28.335 | 18.059 | 15.061 | 25.063 | 14.827 | 17.706 |
| 25.135 | 21.759 | 16.679 | -4.799 | -11.779 | -13.438 | 24.953 | -0.580 | 1.937 | 16.949 | -4.559 | 28.422 | 17.830 | 23.091 |
| 15.289 | 23.423 | 7.297 | 22.444 | -0.565 | -21.717 | -4.074 | -19.994 | -6.135 | 21.973 | 5.220 | 24.517 | 26.811 | 20.846 |
| -11.434 | 8.331 | 4.504 | 9.149 | 29.529 | -6.382 | -4.625 | -15.408 | -13.102 | -5.532 | 19.577 | -7.359 | 4.452 | 29.125 |
| 22.844 | -10.508 | 0.043 | -22.275 | 25.774 | 20.626 | -22.619 | -4.622 | 26.138 | 23.079 | 25.035 | 19.495 | 16.791 | -5.854 |
| -5.180 | 4.792 | 23.637 | 23.273 | 8.412 | 1.093 | 22.586 | 5.479 | 16.930 | 20.536 | 21.245 | 10.969 | 27.869 | 14.743 |
| 18.287 | -26.761 | -7.604 | 16.000 | 28.015 | 22.899 | 10.854 | 26.937 | 29.380 | 18.682 | 26.006 | 8.136 | 10.222 | 25.898 |
| 4.433 | 13.175 | 10.774 | 9.967 | -29.595 | -1.217 | 17.908 | -18.621 | 28.966 | 14.667 | 9.035 | 27.137 | 29.882 | 28.230 |
| -17.854 | -3.044 | -3.296 | 18.504 | 4.214 | 17.222 | -27.138 | 11.025 | 20.584 | 25.234 | 23.638 | 21.574 | 23.349 | 22.687 |
| -28.851 | 12.399 | -1.043 | 26.862 | 21.114 | 9.892 | -17.391 | 8.846 | 26.507 | 13.069 | 22.932 | 23.325 | 18.309 | 29.755 |
| -21.673 | -8.537 | -7.165 | 7.564 | -6.579 | -4.518 | 2.567 | 2.846 | 24.437 | -18.809 | 5.803 | 19.552 | 14.335 | -3.102 |
| 13.496 | 22.442 | -1.825 | -6.778 | -19.404 | -12.582 | -23.863 | -18.135 | 31.386 | 3.329 | 28.217 | 29.340 | 28.736 | 13.987 |
| -24.776 | -3.881 | 17.648 | -4.083 | 19.053 | 27.508 | 16.888 | 10.108 | 27.656 | 10.384 | 4.078 | 4.762 | 17.652 | 8.532 |
| -4.736 | 25.282 | -5.045 | -20.745 | 12.148 | 7.205 | 23.338 | 4.629 | -4.261 | 19.032 | 27.043 | 30.046 | 24.083 | 17.624 |
| 29.423 | -19.975 | 10.273 | 19.498 | -15.474 | -25.092 | 15.174 | -24.557 | 7.431 | 4.317 | 19.683 | 19.410 | 11.330 | -31.847 |
| 19.431 | -21.285 | 18.833 | -28.158 | 21.445 | 26.652 | -0.183 | -0.763 | 27.803 | -2.352 | 23.336 | 21.469 | 21.514 | 21.598 |
| 11.016 | 10.185 | 7.379 | 5.566 | 19.906 | 18.382 | -7.026 | -19.934 | 13.959 | -2.967 | 23.716 | 9.958 | 24.697 | 20.958 |
| 25.288 | 20.856 | 21.866 | 29.068 | -20.586 | -7.967 | 25.805 | 8.332 | -16.286 | 0.690 | -2.213 | -9.040 | 15.816 | 15.781 |
| 2.802 | -25.531 | 26.308 | 26.349 | 24.467 | 21.269 | -11.825 | 14.344 | 2.622 | -1.563 | 29.796 | 20.005 | 19.048 | 16.235 |
| -28.199 | 15.623 | -16.104 | 12.272 | 17.917 | 25.253 | 9.183 | 23.369 | 16.422 | 11.634 | 24.616 | 12.584 | 17.968 | 19.531 |
| 24.589 | -0.805 | 24.222 | 1.152 | -5.446 | -24.495 | 26.594 | 20.741 | 22.515 | 1.067 | 15.482 | 24.491 | 27.461 | 25.987 |
| 3.294 | -27.934 | -21.590 | -20.131 | 26.311 | 11.459 | -1.051 | 7.749 | 3.871 | 15.412 | 2.533 | 28.867 | 12.873 | 28.681 |
| -24.299 | 17.966 | 9.355 | -20.918 | 10.328 | 16.505 | -12.447 | 18.265 | -0.240 | -4.030 | 24.937 | 22.689 | 25.260 | -21.122 |
| 13.436 | -2.799 | -19.931 | 11.203 | -18.664 | -28.633 | 16.719 | 24.902 | -29.045 | 5.386 | 28.330 | 23.152 | 20.982 | 27.801 |
| 9.634 | -14.975 | 13.813 | -5.474 | 6.045 | 29.005 | 10.537 | 17.757 | 25.568 | 23.771 | 14.114 | 14.418 | 20.243 | 24.211 |
| 4.783 | 8.311 | 6.568 | 9.704 | 28.067 | 12.761 | 24.504 | 6.163 | 29.970 | 8.998 | 29.605 | 10.638 | 19.076 | 29.414 |

| | | | | | | | | | | | | | |
|---|---|---|---|---|---|---|---|---|---|---|---|---|---|
| 28.252 | -4.615 | -1.382 | 21.868 | 23.460 | 6.164 | 19.142 | 16.256 | -23.027 | 10.153 | 6.493 | 29.924 | 15.364 | 14.906 |
| -16.262 | 10.936 | -12.664 | 3.543 | 11.178 | 17.758 | -22.527 | 24.097 | 26.637 | 12.217 | 17.999 | 28.482 | 10.113 | 6.810 |
| -8.537 | -1.628 | 3.665 | 6.456 | 7.788 | -1.468 | 2.700 | 4.217 | -9.384 | 27.340 | 14.889 | 1.365 | 28.965 | 30.651 |
| 30.242 | -6.767 | -8.880 | 24.947 | 26.234 | 19.042 | 29.068 | 13.660 | 23.498 | 7.528 | 25.522 | 9.420 | 22.464 | -31.560 |
| 6.489 | -21.934 | 18.080 | 28.214 | 17.774 | 20.488 | 13.156 | -9.761 | -4.669 | 24.383 | 9.139 | 16.222 | 11.360 | 19.229 |
| 26.249 | 21.115 | 2.430 | 20.728 | 20.589 | -19.085 | 20.528 | -15.891 | 18.278 | 0.181 | 30.084 | 19.152 | 26.730 | 25.507 |
| 9.656 | 4.921 | -23.594 | 16.600 | 27.326 | 26.294 | 3.927 | 12.715 | 17.671 | -6.617 | 17.769 | 25.318 | 7.996 | 9.564 |
| 24.140 | 18.135 | -17.984 | -8.318 | 5.457 | 24.704 | -5.024 | 20.972 | 10.013 | 8.413 | 16.120 | 15.892 | 29.584 | 6.382 |
| -7.039 | 0.929 | -19.866 | 23.171 | 13.715 | 6.910 | -11.949 | 17.865 | 14.156 | 27.676 | 7.863 | 26.047 | 17.104 | -8.841 |
| 23.191 | -24.257 | -20.513 | -19.328 | 21.631 | 23.525 | -21.455 | 22.647 | -6.368 | 15.463 | 20.073 | 21.713 | 25.404 | 19.903 |
| -19.260 | 5.375 | -15.893 | -14.745 | 1.998 | -12.686 | 22.052 | -11.731 | -25.857 | 11.129 | 18.973 | 24.022 | 12.437 | 11.054 |
| 17.835 | -9.468 | 8.560 | -21.290 | 6.650 | -10.100 | 19.924 | -0.199 | 26.110 | 10.243 | 26.588 | 27.147 | 22.658 | 21.907 |
| -31.386 | 25.034 | 5.716 | -6.606 | 25.523 | 7.629 | -1.742 | 23.814 | 27.621 | -0.396 | 12.703 | 5.388 | -3.299 | 25.850 |
| 19.003 | -12.106 | 15.966 | 0.236 | -18.708 | -21.890 | -14.106 | 14.345 | 5.985 | 2.099 | 10.856 | 16.916 | 1.729 | 14.738 |
| -10.573 | 22.328 | 9.016 | -7.279 | -6.557 | 10.279 | 27.508 | 13.321 | 22.494 | 26.678 | 7.706 | -20.611 | 21.108 | 28.015 |
| -28.275 | -8.141 | -7.246 | -13.458 | 22.015 | -0.364 | -0.485 | -7.652 | -2.480 | 17.016 | -6.533 | 19.159 | 28.489 | 7.275 |
| -29.295 | -22.409 | 27.573 | 20.221 | 24.101 | 23.305 | 23.646 | -6.987 | 19.435 | 22.895 | 26.906 | 25.756 | 21.957 | 19.574 |
| -24.603 | -13.402 | 10.218 | 17.343 | -22.486 | -28.100 | 19.977 | 21.965 | 17.345 | 21.469 | 28.397 | 18.645 | 18.317 | 22.704 |
| 5.956 | -27.465 | -19.152 | -4.686 | 11.870 | -20.464 | -19.140 | 7.001 | 22.311 | 8.887 | 21.462 | 19.992 | 19.260 | 13.958 |
| 22.892 | 20.161 | -27.727 | -20.587 | 8.414 | -26.267 | -11.438 | 5.982 | 18.707 | 25.479 | 6.528 | 26.368 | 29.205 | 25.401 |
| -8.499 | -17.934 | -8.313 | 9.762 | -15.547 | 2.438 | -5.816 | 22.999 | 17.475 | 10.761 | 3.538 | 16.904 | 23.456 | 27.267 |
| 10.961 | 19.517 | -0.531 | -26.921 | -1.045 | 27.741 | -25.705 | 3.068 | 21.404 | -8.609 | 3.417 | 19.327 | 29.703 | -23.049 |
| 27.253 | 14.888 | 25.188 | 24.065 | -4.693 | 5.392 | -16.558 | 11.476 | 7.904 | 14.952 | 0.249 | 25.087 | -31.528 | 30.204 |
| 25.338 | -22.026 | -3.149 | -19.706 | 19.806 | 22.409 | -7.763 | -24.224 | 16.847 | 23.818 | 8.324 | 5.960 | 23.689 | 25.959 |
| 7.971 | -9.896 | 5.551 | 22.507 | 6.422 | -18.913 | 19.049 | 22.518 | 28.004 | -13.450 | 29.377 | 26.861 | 23.336 | 17.154 |
| 30.598 | -19.224 | -22.668 | 2.278 | -2.183 | 8.280 | -2.167 | 18.773 | 25.277 | 23.357 | 23.140 | 10.939 | 29.210 | 24.925 |
| 10.090 | -1.678 | -12.740 | -4.738 | 25.623 | 20.768 | 9.088 | 23.468 | 10.508 | 9.662 | 6.873 | 9.495 | 5.095 | 13.052 |
| -2.420 | 13.657 | -18.957 | 28.803 | 8.172 | 23.234 | 7.367 | -2.756 | -5.778 | 14.220 | -1.770 | 0.130 | 26.071 | 3.615 |
| -29.843 | 9.117 | 11.033 | 26.024 | -19.022 | -27.869 | -8.908 | 25.732 | -0.368 | 11.264 | 16.618 | 12.891 | -5.928 | 26.041 |
| 22.879 | 18.525 | 30.196 | -11.866 | -6.040 | 12.943 | 16.813 | -1.343 | -26.864 | 22.372 | 24.756 | 3.124 | 4.335 | 26.776 |
| -29.547 | 12.724 | -28.257 | 16.470 | -7.708 | 25.111 | 13.976 | 22.171 | 4.760 | 15.577 | 24.202 | 14.359 | 27.723 | 22.726 |
| 14.840 | -20.414 | 13.607 | 26.944 | 29.164 | 24.243 | 4.897 | 9.607 | 6.399 | 1.740 | 2.912 | -9.651 | -5.412 | 18.528 |
| 18.344 | -24.636 | -5.224 | 18.204 | -17.128 | 21.839 | -9.332 | 17.976 | -1.856 | -9.427 | 0.422 | 24.613 | 13.745 | 21.506 |
| 15.610 | -6.095 | 5.924 | -14.622 | -29.292 | 27.225 | -20.118 | -25.547 | 14.841 | 27.488 | 6.307 | -0.762 | 11.551 | 17.634 |
| 9.270 | -25.679 | -8.363 | 20.442 | 22.847 | -27.537 | -12.288 | 3.682 | -7.083 | 18.555 | 17.633 | 11.894 | -2.899 | 18.114 |
| -20.607 | 2.081 | 27.149 | 27.801 | 20.491 | -19.797 | 24.665 | 26.128 | -21.789 | 23.820 | 27.105 | 15.400 | 24.558 | 10.718 |

| | | | | | | | | | | | | | |
|---|---|---|---|---|---|---|---|---|---|---|---|---|---|
| -25.590 | 22.948 | 18.064 | 25.624 | 13.996 | -22.851 | -0.013 | 4.929 | 3.926 | 4.894 | 12.212 | 23.372 | 26.959 | 14.234 |
| -17.024 | -0.108 | 22.623 | -16.924 | -5.143 | 21.426 | 15.023 | 20.357 | 11.031 | 13.226 | 13.292 | 19.771 | 27.427 | 12.698 |
| -27.471 | -1.749 | 14.580 | -9.185 | 14.734 | 1.020 | -6.772 | 6.594 | 1.443 | 3.865 | 10.301 | 25.430 | 25.495 | 26.600 |
| 26.103 | 25.965 | -16.766 | -17.863 | 18.742 | 19.371 | 10.012 | 23.730 | 21.928 | -2.810 | 18.253 | 9.358 | 28.137 | 24.751 |
| -8.413 | 18.972 | 30.738 | 30.717 | 16.814 | -19.174 | 21.416 | 10.032 | 23.405 | -23.044 | 22.413 | 6.600 | 20.607 | 21.983 |
| -0.662 | 27.097 | 23.791 | 8.777 | 0.913 | -11.190 | -8.562 | -23.884 | 21.327 | 17.529 | 3.262 | 4.899 | 8.515 | 23.062 |
| 14.726 | 21.482 | 20.594 | -5.617 | 18.189 | -22.385 | -13.653 | 26.727 | -26.632 | 6.709 | -2.637 | 1.198 | 13.958 | 11.141 |
| -11.606 | -3.757 | -23.341 | 16.059 | 4.996 | -19.249 | -16.864 | 27.893 | 14.326 | 12.695 | 21.497 | 10.376 | 28.530 | -1.298 |
| -26.103 | -9.760 | -13.204 | -27.454 | 14.549 | -10.443 | 24.562 | 2.183 | -0.917 | 9.409 | 20.835 | 17.399 | 5.510 | 12.000 |
| -17.776 | 15.492 | 5.864 | 26.739 | -19.849 | 20.005 | -15.071 | 19.297 | 11.411 | 22.485 | 29.485 | 25.031 | 15.827 | 18.409 |
| 21.121 | -7.416 | -9.051 | 12.808 | 30.319 | 15.858 | 17.849 | 23.076 | -21.236 | 16.165 | 27.454 | 21.883 | 22.404 | 26.608 |
| -12.726 | 10.920 | -18.504 | -20.355 | 13.467 | 21.572 | -9.823 | -6.834 | 12.209 | 19.892 | 16.830 | 20.624 | 12.920 | 24.225 |
| -18.682 | -23.208 | 2.694 | 0.020 | 2.692 | 4.414 | 0.411 | 20.286 | 25.037 | 13.441 | 15.054 | 18.812 | 14.300 | 25.667 |
| 4.258 | 13.121 | 1.025 | 21.833 | 1.047 | -7.025 | -0.891 | 26.052 | -0.021 | -0.136 | -0.921 | 16.056 | 26.703 | -18.481 |
| 1.552 | -20.604 | -22.673 | 20.280 | 17.420 | 1.560 | -18.005 | 25.356 | 17.656 | 5.936 | -16.186 | 16.504 | 16.002 | 17.866 |
| 13.150 | -8.241 | 29.189 | -0.434 | -29.399 | -4.885 | 25.695 | -26.862 | -24.007 | 9.191 | 25.466 | 11.913 | 20.345 | 8.986 |
| -27.987 | 24.576 | 9.863 | 23.268 | -20.406 | -9.187 | -11.797 | 29.052 | 26.194 | -1.496 | 11.156 | 15.234 | 11.684 | 27.985 |
| 5.887 | 25.975 | 25.035 | -18.436 | 19.995 | 24.683 | -4.750 | 9.519 | 19.158 | 18.395 | 26.489 | 12.641 | -0.317 | 13.326 |
| 10.320 | 14.769 | 30.985 | 24.830 | 10.416 | -6.727 | 0.199 | 24.205 | 16.192 | 16.617 | -17.326 | -16.110 | 23.462 | 18.858 |
| 20.632 | -19.918 | 25.445 | 24.388 | -10.826 | 14.727 | -5.430 | 19.931 | -9.221 | -2.201 | 17.629 | 24.765 | 10.418 | 23.776 |
| -5.595 | 23.636 | -25.182 | 18.061 | -12.368 | -14.248 | 22.902 | 21.491 | -8.582 | 26.434 | 9.146 | 9.107 | 20.941 | -12.669 |
| 28.580 | 3.052 | -11.802 | -20.570 | 1.246 | 3.056 | 21.107 | 26.623 | -0.630 | 6.962 | 25.985 | 17.119 | 7.233 | 3.240 |
| -4.188 | 15.881 | -0.861 | -4.383 | 12.692 | 23.641 | 16.003 | 25.221 | 20.017 | 19.756 | 10.475 | 22.459 | -0.181 | 25.045 |
| -6.860 | 18.860 | 22.283 | -23.106 | 3.324 | -22.092 | 23.186 | -3.100 | -1.761 | 10.314 | 30.982 | -2.638 | 25.489 | 11.792 |
| 26.233 | -5.206 | 19.919 | -22.282 | 26.684 | -5.173 | 20.777 | -2.496 | -7.517 | 7.635 | 26.566 | 18.541 | 23.152 | -0.349 |
| 24.102 | -24.944 | -13.345 | 14.538 | 24.564 | 18.293 | 19.863 | -17.275 | 8.455 | 24.123 | 16.352 | 24.333 | 3.628 | 23.372 |
| -9.561 | 14.175 | 13.471 | 23.652 | 10.090 | -19.684 | 17.331 | -16.004 | 15.738 | 8.697 | 21.778 | 1.941 | 24.254 | 22.404 |
| -15.532 | -18.019 | 5.736 | 8.725 | 22.830 | 2.368 | 7.820 | -1.330 | -3.500 | 14.620 | 18.585 | 7.320 | 11.312 | 9.448 |
| 7.208 | 13.376 | 14.609 | 12.292 | 2.271 | 3.946 | 15.624 | 6.536 | -0.012 | 11.265 | 24.604 | 15.202 | 13.030 | 24.315 |
| 27.445 | 12.776 | 30.907 | -10.190 | 14.815 | 18.898 | 20.760 | 25.481 | 19.119 | 18.551 | 13.537 | 4.055 | 25.197 | 10.184 |
| 13.028 | -3.923 | 3.032 | 20.496 | 20.715 | 12.088 | 22.046 | -12.816 | 12.526 | 17.587 | 23.242 | 20.416 | 18.899 | 9.910 |
| 25.969 | -21.601 | 27.936 | -21.113 | 21.229 | 6.411 | 1.615 | -28.754 | -19.488 | 0.673 | 16.675 | 17.390 | 21.988 | 19.143 |
| 22.832 | -12.669 | -7.675 | 6.706 | -18.602 | -14.110 | -0.422 | 27.140 | -18.464 | -4.406 | 7.895 | 14.161 | 17.713 | 15.410 |
| 28.612 | 25.071 | -22.727 | 18.985 | -17.984 | -16.376 | 1.337 | -0.230 | -6.362 | 20.702 | 7.318 | 0.591 | 25.622 | 0.405 |
| 21.928 | -8.732 | 16.295 | 21.460 | 27.966 | -19.631 | 23.154 | -5.086 | -12.896 | 12.979 | -16.107 | 1.197 | 20.112 | 6.260 |
| -10.876 | -1.983 | 6.201 | 11.276 | -19.725 | 20.587 | 19.818 | 22.164 | 20.982 | -2.970 | 5.124 | 2.926 | 15.374 | 14.323 |

| | | | | | | | | | | | | | |
|---|---|---|---|---|---|---|---|---|---|---|---|---|---|
| -3.042 | 4.407 | -2.820 | -19.954 | 25.900 | -14.361 | -7.294 | 13.036 | -16.812 | 23.686 | 6.011 | 21.405 | 13.976 | 16.852 |
| -12.408 | -11.391 | 2.079 | -10.970 | 12.208 | 8.612 | -7.699 | 30.059 | 21.640 | 24.172 | 0.788 | 7.297 | 22.401 | 13.733 |
| 7.574 | 9.685 | 27.406 | 6.442 | 22.602 | 4.627 | 12.772 | 23.562 | 7.170 | 19.583 | 27.115 | -1.677 | 19.258 | 8.264 |
| -8.674 | -4.095 | -18.688 | -30.479 | -22.699 | 10.371 | 19.013 | 28.386 | 22.053 | 15.022 | 9.924 | 11.620 | 9.773 | 22.059 |
| 22.625 | 18.453 | 17.481 | -28.271 | 24.287 | -0.473 | -11.947 | 27.507 | -22.232 | -1.371 | 29.878 | 4.946 | 12.507 | 6.869 |
| 16.381 | -21.828 | 4.413 | 10.784 | 12.866 | 19.349 | 15.355 | 9.009 | 1.896 | 1.997 | 0.364 | 18.759 | 13.100 | 23.118 |
| -3.715 | -6.613 | -23.416 | 16.841 | 0.578 | 16.105 | -12.624 | 21.893 | 2.609 | 16.802 | -10.528 | 17.558 | 17.585 | -16.819 |
| -24.745 | 1.404 | 24.084 | -15.158 | -20.943 | -24.620 | 24.699 | 10.708 | -5.946 | 12.470 | 17.595 | 14.285 | 24.394 | 14.959 |
| 21.552 | 8.000 | 22.025 | 12.434 | -24.492 | 0.525 | 4.528 | -24.521 | 21.942 | 25.040 | 14.549 | -12.898 | 22.303 | -18.890 |
| -22.774 | 22.013 | -0.633 | -29.988 | 11.656 | 19.930 | 23.466 | 17.711 | 24.086 | -4.253 | 24.162 | 20.018 | 8.370 | 15.752 |
| 17.930 | -7.833 | -2.328 | 15.582 | 6.638 | 17.419 | 17.976 | 24.958 | 18.452 | 9.704 | 24.961 | 25.247 | 25.329 | 20.989 |
| 12.738 | 0.551 | 16.698 | 25.084 | 1.455 | -21.530 | -19.582 | -2.625 | -2.831 | -9.573 | 21.246 | 6.182 | 19.624 | 9.677 |
| 19.856 | -21.384 | -18.589 | 18.712 | 19.955 | -8.617 | -9.494 | 23.143 | 24.384 | 8.407 | 10.919 | 14.524 | 14.847 | 9.062 |
| 26.948 | -17.884 | -7.751 | -12.603 | 27.815 | 15.159 | -8.070 | 5.532 | 25.587 | 17.948 | 18.645 | -4.733 | 11.384 | 23.653 |
| -18.746 | -17.114 | -15.278 | -18.512 | -1.461 | 20.772 | 18.215 | 30.981 | 10.661 | 3.228 | 28.716 | -4.073 | -24.383 | 17.952 |
| 3.664 | -19.437 | 8.437 | -23.731 | 16.960 | 2.783 | 11.834 | 27.203 | 0.884 | 20.046 | 27.145 | 6.099 | 18.299 | 22.297 |
| -10.498 | -6.953 | 29.354 | -19.317 | -9.996 | 0.042 | -9.709 | 29.361 | -24.626 | 10.454 | 10.060 | 25.683 | 12.763 | -16.863 |
| 8.054 | -1.618 | 28.667 | -28.786 | 25.967 | -0.859 | -22.596 | -22.899 | -8.700 | 5.218 | 7.836 | -4.235 | 22.509 | 0.521 |
| -11.560 | 20.466 | -21.973 | -14.209 | 18.009 | -13.753 | 25.787 | 19.978 | -23.448 | 22.096 | -11.194 | 21.800 | -6.288 | 19.461 |
| 11.708 | 25.011 | -0.975 | 23.602 | 2.810 | 3.936 | -10.228 | 26.108 | 10.606 | 24.770 | 19.962 | 26.798 | -8.902 | 13.669 |
| 23.726 | 13.225 | 12.607 | 27.245 | -7.980 | 9.885 | 8.162 | 26.942 | -0.199 | 16.638 | -3.310 | 14.295 | 19.530 | 8.112 |
| 22.528 | 1.797 | 21.832 | -0.130 | 21.355 | 16.877 | 4.900 | 12.855 | 12.808 | 11.446 | 15.519 | 16.070 | -4.424 | 17.845 |
| 6.024 | -23.723 | 19.735 | -22.793 | -2.324 | -14.617 | 20.510 | 28.409 | 3.011 | 7.390 | 20.627 | 20.983 | -8.057 | 0.456 |
| -7.246 | 22.400 | 18.151 | 28.868 | 22.936 | -17.683 | 0.764 | 21.368 | -12.406 | 18.845 | 17.525 | 24.090 | -2.272 | 21.665 |
| -2.322 | -3.263 | 22.485 | 5.950 | -17.975 | -23.975 | 23.969 | -27.840 | -22.849 | -2.018 | 28.022 | -14.100 | 13.731 | 0.716 |
| -15.496 | -21.519 | -18.294 | -6.667 | -5.901 | 24.574 | -19.183 | -4.725 | -18.420 | -7.137 | 12.645 | -14.911 | 11.954 | 18.320 |
| 24.648 | 8.037 | 10.054 | 20.376 | 10.979 | 22.990 | -19.991 | 9.077 | 19.769 | 13.673 | 9.235 | 13.663 | -0.478 | 14.804 |
| -0.505 | -19.718 | -3.743 | -3.656 | 15.957 | 26.328 | -15.675 | 29.018 | 11.867 | -5.679 | -8.083 | 15.833 | -3.951 | 25.547 |
| -12.454 | 26.692 | -22.159 | -26.379 | 18.512 | 6.130 | 11.872 | 6.008 | 9.705 | 7.158 | 3.367 | 17.915 | -11.518 | 24.773 |
| -14.914 | 21.207 | 18.847 | 24.617 | 5.795 | -11.364 | 10.086 | 30.754 | 23.859 | 1.426 | -1.154 | 24.844 | -5.533 | 7.699 |
| 20.916 | 1.873 | -1.628 | 4.912 | 1.187 | -4.480 | -22.593 | -8.325 | 4.469 | 15.491 | 22.097 | -13.903 | 12.806 | -20.772 |
| -7.402 | 23.814 | 26.727 | 15.328 | -9.527 | 6.867 | -2.223 | 8.085 | 8.310 | 17.540 | 23.903 | 9.287 | 22.998 | -9.411 |
| 18.536 | -25.549 | 28.398 | -18.761 | 11.520 | 21.293 | 12.442 | 16.426 | -0.068 | -9.994 | 10.957 | 8.125 | 17.911 | 10.318 |
| 13.360 | 7.238 | 17.573 | 18.383 | 14.896 | -24.919 | 15.498 | -3.724 | 20.716 | -3.288 | -1.904 | 27.059 | 21.147 | 19.021 |
| 14.335 | 22.743 | 5.156 | 6.542 | -18.819 | 0.559 | 25.039 | 30.187 | 8.962 | 14.569 | 16.287 | 14.555 | -22.784 | -19.211 |
| 4.675 | 4.814 | -15.676 | 13.379 | 13.401 | -9.249 | 21.284 | 18.856 | -0.822 | 5.062 | 21.135 | 19.511 | 22.545 | 9.499 |

| | | | | | | | | | | | | | |
|---|---|---|---|---|---|---|---|---|---|---|---|---|---|
| 25.070 | -8.032 | 1.376 | -16.585 | 4.388 | -25.895 | 17.243 | 21.353 | 21.454 | 9.390 | -6.114 | 20.686 | 23.325 | 22.542 |
| 27.411 | 26.765 | 1.958 | 16.952 | 21.305 | -4.816 | -22.704 | 0.196 | -2.134 | 15.729 | -5.259 | 18.840 | 20.342 | -4.605 |
| -19.850 | 18.523 | -14.823 | 4.263 | -11.421 | 26.447 | -26.889 | 23.369 | 14.849 | 19.671 | 8.713 | 0.679 | 3.260 | 24.054 |
| 16.267 | 13.431 | 13.892 | 21.467 | -12.434 | 9.079 | -7.312 | -19.283 | 12.998 | 3.139 | 8.690 | 21.802 | 21.648 | -20.600 |
| -8.540 | 16.408 | -21.037 | -0.395 | 24.985 | -23.254 | 9.917 | 22.129 | -7.654 | 23.996 | 26.618 | 28.232 | 11.136 | 8.843 |
| 22.701 | -6.501 | -12.159 | 17.920 | 27.875 | 8.059 | 24.184 | -5.001 | -17.288 | -4.683 | 23.651 | 27.089 | 12.660 | 26.023 |
| 1.074 | -26.779 | -21.739 | -14.430 | 23.054 | 19.166 | -27.735 | 29.426 | 7.561 | -9.396 | 19.951 | 27.667 | 9.022 | 16.444 |
| 21.584 | 14.892 | 15.841 | 19.307 | -28.130 | 10.519 | -6.033 | -23.578 | 11.890 | 3.762 | -6.183 | -0.773 | 23.097 | -15.291 |
| 10.260 | -3.117 | 23.658 | -21.237 | 19.086 | -3.316 | 26.227 | 5.801 | -8.630 | 15.689 | 15.476 | 9.426 | 3.119 | -0.197 |
| -15.691 | 22.031 | 18.144 | 24.063 | 24.412 | -21.854 | -1.860 | -15.140 | -7.382 | -10.302 | 14.376 | 23.882 | 6.633 | 23.905 |
| 3.486 | 17.108 | 11.289 | 10.891 | -20.059 | 11.478 | 12.662 | 26.854 | 4.190 | 5.196 | 18.970 | 10.827 | 18.776 | 10.518 |
| 12.419 | 24.298 | 6.750 | 26.087 | 26.325 | 26.001 | 20.060 | 7.150 | 26.934 | 6.816 | 1.364 | 12.832 | 21.152 | 11.528 |
| -4.151 | 15.060 | 20.701 | -23.160 | -25.635 | 1.711 | 22.803 | 15.991 | 20.985 | 5.920 | 23.446 | 8.446 | 25.108 | 7.562 |
| 6.822 | -17.238 | 5.346 | -4.228 | 20.117 | 12.828 | 28.618 | 14.967 | 24.332 | 8.980 | 15.042 | 10.155 | 21.450 | 8.247 |
| -1.589 | 12.043 | -7.753 | 3.922 | 2.795 | 2.829 | 26.921 | -3.547 | 6.765 | 4.434 | 22.496 | 15.601 | 18.177 | 14.989 |
| 11.024 | -23.873 | 27.223 | 5.181 | 12.494 | -22.785 | -29.066 | 28.588 | -13.909 | 20.570 | 12.140 | 16.786 | 16.670 | 28.090 |
| -20.361 | 23.798 | 18.692 | -9.906 | 14.896 | 15.189 | -18.954 | 11.009 | 1.999 | -13.788 | 20.402 | -2.738 | 27.436 | -27.347 |
| 24.492 | -19.074 | 0.320 | 18.601 | 8.553 | 24.942 | 22.363 | 29.865 | 25.014 | 1.076 | 7.575 | 19.986 | 22.750 | 27.256 |
| 12.344 | 10.118 | -5.569 | 5.104 | -7.987 | -0.444 | -13.810 | 25.425 | 25.946 | 0.688 | 25.374 | 13.753 | 19.134 | 25.959 |
| 19.075 | -29.491 | -16.987 | -8.231 | -15.820 | 16.979 | 15.053 | 7.909 | 25.642 | 15.836 | 27.361 | 26.900 | 23.604 | 16.840 |
| 2.165 | 12.692 | 26.094 | 14.516 | 11.035 | -23.466 | 26.814 | -3.820 | 0.383 | 10.279 | 23.028 | 25.614 | -4.612 | 11.355 |
| -9.737 | 16.276 | -18.351 | -4.610 | -18.564 | -9.488 | 19.347 | 19.996 | -11.090 | 5.380 | 9.777 | 16.620 | 6.539 | -3.996 |
| -22.529 | 22.298 | 1.443 | 10.743 | 23.596 | -19.330 | -10.104 | 30.527 | 23.597 | 13.026 | 24.013 | -2.028 | 5.644 | -3.595 |
| 18.108 | -21.475 | -23.172 | 4.950 | 22.243 | 12.975 | 8.504 | -2.341 | 10.922 | 25.713 | 16.484 | 28.872 | -11.564 | 4.410 |
| -0.204 | 17.785 | 16.238 | 8.406 | 15.249 | -26.799 | -1.506 | -19.214 | -4.115 | 18.176 | 4.040 | 17.882 | 17.875 | -5.378 |
| -16.846 | -2.624 | 19.591 | 17.587 | 7.316 | -25.008 | 21.349 | 8.976 | -13.172 | 4.380 | 4.603 | 23.033 | 7.693 | 25.440 |
| 12.224 | -12.438 | 14.969 | -14.673 | -17.119 | 0.003 | 18.655 | 15.522 | 17.032 | -2.382 | 24.550 | 25.985 | 12.682 | 26.936 |
| -3.000 | 11.410 | 12.970 | -4.130 | -3.393 | -22.428 | 25.569 | 26.812 | -22.957 | 22.999 | 17.309 | 10.304 | 23.428 | 16.287 |
| -12.912 | 19.994 | -15.322 | -7.746 | -20.504 | 23.325 | 24.327 | -15.301 | 19.460 | -7.103 | 26.347 | 23.520 | 10.219 | -3.265 |
| 3.946 | 14.035 | -2.073 | 11.598 | 13.280 | 3.629 | 15.150 | 25.268 | 7.070 | 14.162 | 14.448 | 17.197 | 13.411 | 12.085 |
| -4.714 | 12.206 | -4.408 | 2.537 | 0.093 | 24.439 | -5.891 | -23.501 | 22.816 | 5.999 | 6.111 | 28.655 | 24.550 | 15.687 |
| 15.948 | -28.413 | -16.372 | 4.654 | 20.526 | 10.550 | 5.455 | 8.943 | 1.864 | 21.925 | 10.678 | 26.260 | 21.070 | -15.052 |
| -22.108 | -9.974 | -11.616 | 5.658 | 17.554 | 6.625 | -21.913 | 20.691 | 24.208 | 8.657 | 18.584 | 25.145 | -13.282 | 29.474 |
| 24.389 | 28.360 | -25.318 | 2.808 | 24.283 | 7.348 | -1.385 | -19.498 | -17.498 | 12.153 | 15.916 | 22.699 | -20.935 | 15.805 |
| 6.046 | -6.809 | 20.798 | -22.246 | 8.879 | 25.257 | 9.070 | 27.296 | 19.674 | 17.526 | 19.995 | 21.324 | 24.833 | 23.612 |
| -0.244 | -18.979 | -9.574 | -12.295 | -17.693 | 0.974 | -13.381 | -3.019 | 10.928 | 25.381 | 18.283 | 5.182 | 20.853 | 13.263 |

| | | | | | | | | | | | | | |
|---|---|---|---|---|---|---|---|---|---|---|---|---|---|
| -24.900 | 22.424 | -10.538 | -6.385 | -0.886 | -5.377 | -20.167 | -25.838 | 8.928 | 24.502 | 12.814 | -1.721 | 28.237 | 24.931 |
| -16.855 | -7.791 | 18.545 | -8.616 | 16.484 | 19.153 | 29.650 | -25.079 | 5.542 | 16.645 | 17.668 | 18.864 | 12.708 | 20.337 |
| -24.270 | -3.150 | 6.217 | 3.379 | -21.022 | 18.233 | 28.070 | 7.684 | 15.755 | 3.748 | 21.664 | 29.498 | -8.204 | 12.877 |
| -14.572 | 0.261 | 15.128 | 25.351 | 1.086 | 3.767 | 11.684 | 19.957 | -20.908 | 10.611 | 20.559 | 12.107 | 10.213 | 30.704 |
| 21.998 | 19.913 | -7.153 | -6.687 | -21.978 | -24.695 | 17.043 | 6.746 | -6.497 | 23.479 | 25.541 | 22.128 | 17.359 | -28.942 |
| 7.226 | -22.204 | 20.259 | -7.472 | 21.770 | 21.230 | 4.344 | 21.122 | 19.221 | 1.217 | 20.071 | 14.824 | 8.781 | 20.031 |
| 0.228 | 16.411 | -0.536 | 2.124 | 24.760 | 25.697 | -17.294 | 24.569 | 3.864 | -19.016 | -4.339 | -4.269 | 2.490 | 24.405 |
| 24.835 | 24.192 | -2.629 | 20.519 | 19.323 | 15.934 | -3.166 | -26.751 | -3.832 | 19.056 | 7.605 | 17.316 | 11.631 | -8.593 |
| 22.484 | -28.903 | -14.965 | -8.105 | 12.379 | -25.605 | 5.999 | 4.522 | 8.480 | -12.766 | 18.446 | 9.274 | 29.668 | 24.041 |
| -19.607 | 18.757 | -5.420 | 20.092 | 8.120 | 2.934 | 25.319 | 29.184 | 13.448 | 17.531 | 15.507 | 28.184 | 25.302 | 25.937 |
| -9.805 | 4.249 | 0.413 | -24.504 | -11.086 | 5.937 | -5.797 | 25.099 | 25.322 | 8.945 | 21.742 | 25.666 | 15.333 | 14.733 |
| -8.664 | -24.633 | -12.840 | 11.484 | -26.760 | -4.755 | -9.528 | 19.389 | 7.582 | 19.875 | 20.569 | 20.125 | 21.474 | 12.117 |
| 2.257 | -17.537 | -16.301 | 21.849 | 22.502 | 4.992 | 6.753 | 10.685 | 2.640 | 11.109 | 2.272 | 23.798 | 22.929 | 19.523 |
| 4.338 | -16.497 | -18.635 | -8.654 | -22.212 | 0.058 | -11.019 | 8.458 | 9.898 | 22.061 | 23.842 | 31.059 | -7.617 | 27.478 |
| 5.801 | -22.426 | 22.132 | 2.064 | 21.273 | 12.832 | 19.889 | -5.476 | -14.983 | 7.420 | 8.250 | 22.426 | 18.163 | 29.902 |
| 16.722 | -20.310 | -6.578 | -25.486 | -9.148 | 18.033 | 8.188 | -0.951 | -24.229 | 18.511 | 14.858 | 11.169 | 17.127 | 19.494 |
| -3.884 | 16.377 | -21.792 | 10.794 | 19.563 | 14.391 | -21.285 | 12.972 | 7.670 | 6.830 | 21.266 | 3.348 | 28.743 | 13.298 |
| 11.939 | -19.916 | -23.173 | -21.178 | -16.670 | 8.137 | 16.996 | 0.797 | 0.787 | 8.796 | -22.905 | 30.655 | 16.996 | 5.228 |
| 9.517 | 26.016 | -7.611 | -3.842 | -10.051 | 15.365 | -23.484 | -22.510 | 2.902 | -5.652 | 16.425 | 21.220 | 11.615 | -25.194 |
| 0.697 | 29.137 | 10.608 | 23.116 | 13.721 | 10.766 | 4.724 | 25.926 | 16.547 | 8.161 | -5.382 | 29.937 | 5.873 | 29.096 |
| 7.855 | 14.657 | 23.246 | -23.216 | 16.128 | 6.807 | 10.037 | 23.262 | 24.243 | -3.549 | 0.886 | 27.042 | -27.030 | 9.220 |
| -21.833 | -21.376 | -21.032 | -10.876 | -13.830 | 19.441 | 23.509 | -3.220 | 0.105 | 11.033 | 24.743 | 22.029 | 19.674 | 27.862 |
| 17.465 | -24.410 | 18.681 | 5.336 | 5.232 | 7.714 | 19.857 | 15.466 | -2.558 | 3.980 | 5.060 | 28.571 | 27.964 | 11.829 |
| 20.760 | 18.940 | 12.527 | -13.812 | 22.079 | -6.664 | 22.710 | 19.770 | -14.077 | 2.232 | 22.417 | 23.052 | 13.685 | -12.663 |
| 12.344 | 12.949 | 12.029 | -1.291 | 1.605 | 25.413 | 0.507 | -2.308 | 19.619 | 12.066 | 17.120 | 23.104 | 16.800 | 18.491 |
| -10.952 | 20.388 | 16.894 | -26.897 | -0.998 | 20.313 | 18.791 | 2.866 | 18.609 | 24.978 | -0.408 | 28.943 | 24.346 | -0.264 |
| 25.304 | 18.407 | 0.674 | 6.200 | 0.248 | -22.503 | 8.544 | -24.666 | 18.338 | 23.046 | -4.162 | 16.063 | 9.552 | -24.689 |
| -21.041 | -18.320 | -12.015 | 15.038 | 20.597 | 16.322 | 16.813 | -21.587 | 7.157 | 20.386 | 2.377 | 8.378 | 19.311 | 30.354 |
| 0.743 | 23.852 | 7.109 | 8.383 | -17.289 | 14.240 | 21.483 | 17.544 | 22.610 | 27.919 | 21.220 | 2.159 | 21.149 | 24.113 |
| 9.781 | 21.602 | -1.820 | 24.531 | -8.833 | -14.229 | 18.179 | 26.705 | -11.784 | 14.954 | 11.903 | 7.311 | 21.770 | 14.656 |
| 17.682 | 2.924 | -7.025 | 20.271 | -18.818 | 12.728 | -16.640 | 10.836 | -21.896 | 11.442 | -3.563 | 30.705 | -18.089 | 11.700 |
| -21.230 | -4.808 | -22.880 | -25.461 | 0.136 | 4.681 | 26.666 | 25.202 | -10.744 | 5.587 | 20.011 | 24.884 | 6.115 | 25.139 |
| 22.258 | -31.340 | 21.317 | 17.568 | -19.608 | -23.870 | 22.646 | 23.216 | 11.702 | -10.171 | 18.101 | -7.625 | 29.323 | 0.202 |
| 14.399 | -21.275 | -18.461 | -26.721 | -6.233 | 10.173 | 29.034 | 21.146 | 24.939 | -29.141 | -8.057 | 20.301 | 7.004 | 27.368 |
| 6.312 | -23.890 | -2.022 | 21.690 | -7.174 | 7.079 | 7.745 | 0.511 | -6.764 | -2.141 | 0.245 | 23.134 | 20.296 | 20.199 |
| 1.322 | -3.845 | 14.422 | 19.445 | -7.836 | -2.606 | -22.032 | 9.848 | 8.644 | -14.973 | 24.845 | 21.145 | -4.728 | 31.244 |

| | | | | | | | | | | | | | |
|---|---|---|---|---|---|---|---|---|---|---|---|---|---|
| 9.183 | -24.501 | -13.086 | 17.081 | -11.801 | -0.464 | -24.959 | 24.069 | 27.608 | 24.263 | 13.777 | 22.491 | 14.697 | 21.755 |
| -24.195 | 17.040 | 25.238 | -21.027 | 5.010 | 17.454 | 4.124 | 12.013 | 20.537 | 19.797 | -1.020 | 6.350 | 27.265 | 7.606 |
| 11.182 | 8.461 | 17.643 | -24.019 | 17.005 | 25.481 | 11.087 | -12.763 | 13.374 | -16.398 | -2.778 | 19.484 | 26.489 | -11.310 |
| 1.973 | 27.358 | -8.576 | -21.738 | 13.170 | 15.159 | 27.467 | -8.350 | -14.784 | 7.808 | -4.811 | 8.803 | 20.784 | 6.009 |
| 23.250 | 21.749 | 11.743 | 16.543 | -11.242 | -26.113 | 16.006 | 9.089 | 18.827 | -6.024 | 22.483 | 23.409 | 14.240 | -3.680 |
| -15.233 | 0.044 | -12.193 | 4.505 | 23.784 | -6.397 | 24.539 | 20.766 | -17.712 | -4.165 | 8.073 | 10.057 | 13.121 | -6.532 |
| -7.123 | 14.537 | 8.885 | 21.948 | -2.250 | 10.322 | 19.900 | 15.974 | -12.567 | 14.682 | 2.781 | 12.044 | 28.199 | 11.450 |
| -26.835 | 22.618 | -4.976 | 26.606 | 8.221 | 3.406 | -17.416 | 7.868 | 8.680 | 9.847 | 0.871 | 21.316 | 22.976 | -5.304 |
| -5.922 | 18.087 | -11.542 | -14.210 | 18.238 | -3.141 | -2.936 | 10.054 | 21.136 | 3.186 | 16.337 | 3.239 | 9.876 | 8.363 |
| 27.955 | 18.908 | -9.953 | -14.835 | 10.867 | 15.946 | 26.716 | -1.099 | 1.408 | 17.392 | 13.933 | -8.261 | 17.643 | 21.909 |
| -8.415 | 23.989 | -2.111 | -18.662 | -22.329 | -1.651 | 4.229 | 26.384 | -16.199 | 2.004 | -13.339 | 27.520 | -3.466 | 10.969 |
| 6.401 | 25.689 | -19.142 | 24.583 | 23.749 | -22.115 | -22.279 | -2.849 | 0.621 | 27.381 | 20.486 | 28.462 | 21.252 | 25.331 |
| 12.296 | -21.767 | -20.914 | 24.018 | -1.523 | -28.517 | 21.650 | -20.190 | 19.828 | 25.807 | 23.560 | 11.152 | 6.471 | 8.794 |
| -21.205 | 16.659 | 20.636 | -25.914 | -27.380 | -25.605 | -27.448 | 22.057 | 26.083 | 14.793 | 3.042 | 16.845 | 24.708 | 27.257 |
| -25.536 | -28.334 | 1.075 | 23.439 | 16.801 | 20.641 | 2.426 | 20.657 | 17.993 | 3.045 | -3.782 | -22.467 | 19.281 | 29.046 |
| -6.013 | 21.893 | -5.516 | 18.007 | -25.011 | -11.613 | 18.637 | 25.815 | 10.880 | 11.029 | 5.127 | 5.275 | 22.531 | 14.339 |
| -24.112 | -8.625 | -7.459 | -20.128 | -7.944 | -19.562 | -31.422 | 5.376 | -7.678 | -6.676 | 18.935 | 24.070 | 30.537 | 2.068 |
| -3.347 | -12.622 | 22.942 | 12.239 | 18.755 | 21.582 | -8.637 | -7.178 | 7.474 | 20.781 | 8.669 | 7.916 | 28.220 | 24.052 |
| 10.835 | 26.420 | -19.496 | 14.855 | 15.802 | 19.674 | -7.084 | 7.381 | -1.035 | 22.425 | 0.247 | 18.438 | 18.125 | 10.022 |
| -25.991 | 30.032 | -2.678 | 5.955 | 1.860 | 18.050 | 29.212 | 3.163 | -4.159 | 21.524 | 5.430 | 4.647 | 17.364 | 23.051 |
| -10.306 | 24.294 | -24.764 | -28.139 | 25.928 | 2.620 | -4.710 | -5.282 | 26.918 | -2.447 | 12.858 | -3.473 | -6.599 | 6.623 |
| -12.573 | -3.058 | -15.306 | 24.711 | 0.025 | 25.272 | 20.549 | 9.740 | 15.017 | 23.059 | 19.416 | 28.292 | 29.336 | 18.480 |
| 7.020 | -24.793 | 15.651 | -18.289 | -6.357 | 18.644 | 5.315 | -23.336 | -1.404 | 11.542 | 7.968 | 9.866 | -2.791 | 7.913 |
| 14.232 | 20.053 | 9.897 | 18.436 | 27.289 | -27.213 | 30.134 | 14.714 | 9.252 | -0.893 | 21.043 | 29.346 | 15.953 | 24.224 |
| 5.933 | 27.498 | -1.238 | 9.112 | -27.668 | 12.379 | 23.753 | 3.871 | -16.442 | 28.294 | 24.854 | 22.890 | 10.087 | 27.965 |
| 23.523 | 20.915 | 12.683 | -26.348 | -23.651 | 28.438 | 9.713 | -20.832 | 14.070 | 24.586 | 15.525 | 28.805 | 19.539 | 19.684 |
| -6.665 | 16.147 | -12.670 | -0.946 | 26.623 | -21.001 | 18.769 | 21.592 | 22.745 | -18.147 | 7.390 | 27.641 | -1.928 | 11.932 |
| -16.430 | -17.931 | -19.354 | -10.476 | 21.216 | -29.640 | 4.467 | 9.042 | 15.554 | 18.820 | 14.447 | 19.002 | 8.413 | 21.297 |
| -24.243 | 29.829 | 2.200 | 15.712 | -10.912 | -20.560 | 29.788 | 5.420 | 28.535 | 27.381 | -0.788 | 25.651 | 29.566 | 8.342 |
| 12.639 | -14.890 | -23.218 | 27.166 | -11.196 | 11.575 | 22.691 | 7.463 | 16.880 | 26.312 | 0.770 | 6.369 | 16.148 | 27.134 |
| -16.870 | -21.985 | -1.362 | 22.682 | 24.222 | 23.851 | -13.066 | 13.548 | -26.536 | -12.103 | 8.145 | 23.388 | 8.867 | 7.362 |
| 17.133 | -31.931 | 19.102 | -20.065 | 3.764 | -5.862 | 13.989 | 22.106 | 23.738 | 9.027 | -12.364 | 22.617 | 11.162 | 25.685 |
| 9.207 | 18.897 | 10.542 | 17.291 | 4.470 | -2.233 | -7.472 | 5.359 | 21.390 | 7.226 | 5.214 | 19.717 | 30.075 | 11.153 |
| 17.351 | 21.631 | -20.586 | 3.675 | 15.121 | 7.725 | -1.836 | 1.180 | 9.938 | 26.682 | 23.520 | 24.703 | 20.709 | 24.987 |
| -13.156 | 26.993 | 18.526 | -0.738 | 25.149 | 8.813 | 10.144 | -6.888 | -1.980 | 6.468 | 4.789 | 1.555 | -3.181 | -19.869 |
| 20.586 | 22.723 | 25.641 | 23.908 | -6.748 | 5.843 | -20.195 | 22.749 | -9.573 | -2.172 | -6.675 | 12.779 | 25.758 | 26.141 |

| | | | | | | | | | | | | | |
|---|---|---|---|---|---|---|---|---|---|---|---|---|---|
| -4.648 | -20.430 | -1.200 | 29.346 | 21.125 | 30.005 | 27.086 | 3.038 | 6.133 | -5.205 | -5.146 | 11.709 | 14.591 | 22.108 |
| 15.042 | 9.070 | 5.698 | 13.114 | 3.189 | 4.691 | 5.214 | -6.681 | 18.553 | 9.816 | 19.437 | 17.895 | 24.644 | 12.045 |
| -25.079 | -15.685 | -1.821 | -22.902 | 7.120 | 6.661 | -6.395 | 6.780 | 3.338 | -1.120 | 3.409 | 26.096 | 13.709 | 20.557 |
| 23.703 | 27.664 | -0.114 | 9.333 | 15.349 | 15.929 | -25.547 | 12.991 | 11.486 | 15.999 | 9.005 | 23.071 | 17.869 | 21.026 |
| 28.438 | 22.079 | -26.959 | 19.699 | 22.359 | 22.316 | 15.016 | -1.715 | 25.391 | 14.854 | 25.695 | 27.512 | 22.341 | 9.285 |
| 19.779 | -26.770 | 11.200 | -19.930 | -19.622 | 14.814 | 27.567 | 20.144 | 17.650 | 20.616 | 7.021 | -3.036 | 29.633 | 7.922 |
| -4.565 | -8.819 | 13.699 | -28.540 | 6.579 | 26.525 | 19.423 | -12.268 | 27.852 | 18.970 | 24.580 | 13.886 | 24.109 | -7.803 |
| -5.785 | 18.924 | -1.010 | 28.407 | -26.817 | 23.602 | -13.725 | 22.585 | -0.126 | 30.230 | 16.052 | 2.916 | 22.496 | 21.882 |
| -19.346 | -6.258 | -5.226 | -8.826 | 24.997 | 10.984 | 28.782 | 11.892 | -18.232 | 14.961 | 11.158 | 23.904 | 25.977 | 17.953 |
| -30.178 | -0.645 | -20.836 | -24.658 | 27.866 | -2.801 | 7.352 | 7.569 | 26.037 | 0.824 | 18.493 | 16.590 | 19.672 | -8.405 |
| -2.678 | 21.124 | -22.396 | 17.886 | 10.964 | -15.068 | 10.111 | 3.540 | 12.789 | -8.540 | -6.028 | 7.303 | 20.326 | -21.930 |
| -6.787 | -25.115 | -27.027 | 14.002 | -5.280 | 12.808 | 5.822 | 10.752 | 27.033 | 23.716 | 13.282 | 27.022 | 15.542 | -11.313 |
| 11.135 | 24.351 | 9.478 | -4.988 | 8.630 | 8.902 | -4.867 | 12.671 | 29.393 | 21.724 | 2.984 | 16.014 | 12.675 | 18.652 |
| 8.138 | 16.982 | 23.674 | -18.551 | 17.965 | -26.558 | 16.153 | 4.592 | 22.461 | 5.516 | 24.695 | 8.161 | 18.662 | 8.995 |
| 29.176 | 18.067 | -24.613 | -5.695 | 16.898 | 19.931 | 3.689 | 5.817 | 19.473 | 27.114 | 14.127 | 2.232 | 23.041 | 6.914 |
| -12.982 | 21.528 | -14.227 | -12.159 | -26.377 | 8.881 | 11.480 | 19.988 | 2.034 | 24.298 | 11.311 | 11.607 | 10.239 | 0.934 |
| -17.242 | 24.778 | 1.351 | -22.579 | 5.627 | 24.027 | 5.150 | 16.651 | 5.645 | 0.691 | 26.040 | 23.219 | 21.488 | 15.467 |
| -23.251 | -31.570 | -15.666 | 16.716 | 20.730 | 19.455 | -18.821 | 13.984 | 21.969 | 4.128 | 2.663 | 26.477 | 5.081 | 9.931 |
| -18.156 | 28.853 | -6.048 | -1.851 | -19.121 | 13.320 | 20.813 | -4.211 | 18.882 | 13.355 | 22.957 | 12.992 | 19.919 | -17.080 |
| 22.429 | 25.271 | 10.152 | -29.231 | -9.589 | -23.149 | 4.409 | 4.594 | 4.654 | 2.276 | -2.278 | 8.947 | 23.512 | -16.595 |
| 1.731 | -1.401 | 22.951 | 26.545 | 27.533 | -4.809 | 5.490 | 25.989 | -2.224 | -7.102 | 26.305 | 0.548 | 22.157 | 0.499 |
| 13.526 | 18.488 | -28.536 | 18.421 | -9.953 | -29.619 | -1.190 | 13.095 | 27.391 | 19.513 | 16.628 | 9.873 | 19.074 | 3.847 |
| 4.046 | 20.444 | -13.113 | -20.628 | -12.201 | -10.213 | 19.797 | 24.637 | 10.834 | 21.971 | 9.559 | 25.604 | 15.821 | 9.486 |
| 0.706 | -28.228 | 8.906 | 19.525 | 24.966 | 18.748 | -30.550 | 19.189 | 2.202 | 15.753 | 21.498 | 18.453 | 24.648 | -16.107 |
| -5.287 | 22.261 | -13.953 | 9.305 | -4.426 | 6.869 | 14.544 | -19.127 | 6.388 | 24.556 | 23.808 | 10.696 | 18.181 | -18.081 |
| -11.556 | 26.659 | 19.713 | -12.944 | -25.725 | 26.534 | 9.559 | -6.197 | -0.214 | 18.333 | 14.709 | 13.436 | 25.463 | -18.570 |
| -1.724 | -25.342 | 28.610 | -30.098 | 27.142 | 10.970 | -30.175 | 3.089 | 4.829 | 28.810 | 16.470 | 23.782 | 21.177 | -16.371 |
| 4.728 | 27.370 | 6.574 | 25.329 | 15.624 | -18.016 | -20.082 | 4.588 | 14.895 | 13.992 | 1.304 | -13.856 | 10.892 | -18.760 |
| 24.176 | 17.249 | -25.647 | -16.528 | -13.992 | -24.117 | 25.542 | 25.970 | 10.479 | 8.477 | 26.170 | 17.231 | 25.262 | 22.719 |
| -9.988 | 18.103 | 14.653 | -11.113 | -16.470 | -27.192 | 8.744 | -11.360 | 22.607 | -14.068 | 5.158 | 7.485 | 9.926 | -5.603 |
| -23.381 | 8.764 | 2.525 | 13.444 | 8.946 | 2.107 | -18.212 | 3.633 | -13.314 | 2.503 | 28.402 | 9.848 | 6.436 | 17.890 |
| 1.359 | 4.406 | 20.398 | -20.069 | 17.424 | -25.581 | 27.065 | -7.155 | -1.472 | -4.036 | 11.734 | 22.739 | 16.448 | -21.016 |
| -6.949 | -12.189 | -7.869 | 12.479 | 23.638 | 22.195 | -3.219 | 26.190 | 9.429 | 9.412 | 22.922 | 2.558 | 15.371 | 14.633 |
| 26.099 | -1.846 | 12.453 | 1.302 | 3.237 | -28.633 | 6.610 | 2.235 | 21.125 | 21.446 | 17.862 | -3.801 | 7.389 | -17.204 |
| -16.748 | -18.614 | 9.666 | 10.113 | 13.260 | -4.760 | 23.593 | 22.014 | -16.381 | 0.949 | 15.740 | 8.054 | 10.542 | 13.710 |
| -3.035 | -21.910 | 23.543 | -21.422 | 2.929 | 30.363 | 5.308 | 23.003 | 17.095 | 29.724 | 24.566 | -8.512 | 19.920 | -1.641 |

| | | | | | | | | | | | | | |
|---|---|---|---|---|---|---|---|---|---|---|---|---|---|
| -11.744 | 25.717 | -20.165 | -27.125 | 24.817 | 25.511 | -8.190 | -3.150 | 24.531 | 25.445 | 7.521 | -18.645 | 12.751 | -0.843 |
| 7.792 | -19.863 | 18.594 | 13.874 | 21.893 | 6.165 | -22.601 | 2.932 | 13.945 | 20.513 | 4.492 | -9.681 | 17.437 | -4.732 |
| -7.807 | -27.083 | -25.350 | 4.425 | 1.017 | -2.422 | -5.816 | 19.317 | 11.896 | -15.403 | 6.134 | 14.203 | 26.256 | -17.055 |
| 20.030 | -15.179 | 7.612 | -22.541 | -13.017 | 5.020 | -26.733 | 17.361 | 25.876 | 4.690 | 14.143 | 7.150 | 4.918 | 9.351 |
| -5.094 | 23.431 | -18.957 | -19.449 | -10.862 | 15.487 | -28.036 | 9.395 | 19.267 | 12.827 | 19.308 | -1.714 | 20.905 | 2.510 |
| -14.775 | 22.031 | -16.389 | -27.446 | 27.560 | 14.871 | 13.886 | 26.864 | -6.639 | -23.411 | 22.946 | 9.963 | 12.991 | 12.965 |
| 26.871 | -21.659 | 1.880 | 27.070 | 22.658 | 8.841 | -17.454 | 14.981 | 28.311 | -1.260 | 14.915 | 15.786 | 16.502 | -18.984 |
| 12.674 | 19.763 | 15.953 | 15.876 | -20.516 | 7.005 | 8.038 | -2.135 | -4.517 | 25.763 | 12.661 | 17.084 | 2.449 | 8.186 |
| -9.313 | -22.808 | -24.826 | 20.188 | 11.360 | 12.310 | -23.011 | -7.498 | 25.042 | -10.374 | 21.497 | 21.370 | 21.186 | 7.354 |
| -29.824 | -24.346 | -22.164 | -10.440 | 15.504 | -29.900 | 17.636 | -16.834 | 23.978 | 27.382 | 9.432 | -0.320 | 18.787 | -2.196 |
| -14.064 | 18.068 | 13.119 | 23.375 | 21.166 | 18.777 | 2.994 | -6.360 | 28.995 | 18.117 | -6.664 | 22.419 | 5.398 | 14.116 |
| -16.429 | -2.463 | 15.004 | 14.266 | 27.549 | 22.782 | -5.035 | 24.015 | 26.876 | 14.804 | 1.969 | 20.335 | 4.010 | 4.910 |
| -0.413 | -21.526 | 13.737 | -15.904 | 5.678 | -24.132 | -13.563 | 11.474 | 9.145 | 24.598 | -13.495 | 23.157 | 1.546 | 23.769 |
| 22.552 | 23.243 | -19.951 | -22.040 | 3.474 | 29.089 | 8.574 | 1.562 | 26.466 | 22.653 | 10.830 | 22.316 | 4.496 | 10.669 |
| 16.847 | 13.342 | -8.518 | 4.388 | -18.996 | 16.622 | -3.743 | 3.614 | 1.519 | 8.640 | 18.875 | 12.996 | 17.471 | 10.039 |
| -8.449 | 27.780 | -11.814 | -18.146 | -10.560 | 9.308 | -27.576 | -4.351 | 19.652 | 26.358 | 6.116 | 0.992 | 22.758 | 9.459 |
| -0.789 | 6.533 | -25.785 | 6.871 | -24.395 | 23.991 | 18.104 | 26.228 | 7.367 | 28.079 | 28.955 | 24.794 | -6.037 | 12.190 |
| -4.948 | 9.294 | -8.863 | -2.051 | 25.309 | 25.277 | -2.536 | -3.228 | 8.036 | 27.177 | 10.725 | -9.956 | 18.659 | 12.780 |
| -1.118 | -4.232 | 10.057 | -2.988 | 29.327 | 3.997 | 3.938 | 19.353 | -11.908 | 29.059 | 17.962 | -2.668 | 2.842 | -3.701 |
| -8.068 | -17.837 | 4.963 | -20.753 | 5.251 | -0.455 | -20.199 | -2.021 | -14.932 | 11.196 | 15.560 | -1.105 | 10.181 | -7.171 |
| 3.756 | -0.090 | -23.621 | 9.688 | -23.861 | -21.899 | -21.020 | 4.285 | 15.925 | 3.223 | -3.689 | 18.404 | 21.628 | 22.913 |
| 9.981 | 15.055 | -5.329 | 7.585 | -20.216 | 13.325 | 27.546 | -0.415 | -29.792 | 25.556 | 25.323 | 25.084 | -0.686 | 0.416 |
| -3.907 | 20.161 | -8.496 | 22.528 | -7.312 | 3.137 | -8.396 | 21.622 | 21.913 | 21.835 | 12.388 | 13.331 | 12.805 | 24.825 |
| 8.032 | 5.369 | -27.967 | 10.775 | -27.205 | 20.929 | -5.093 | 1.188 | 18.452 | 19.681 | 27.698 | 19.279 | 22.322 | 19.738 |
| 25.025 | 9.391 | 10.494 | -7.742 | -5.618 | -15.032 | 20.449 | -10.156 | 16.808 | 9.842 | 8.146 | 26.706 | 17.961 | 13.536 |
| 14.686 | -19.554 | 9.360 | 24.332 | -12.670 | -23.506 | -13.245 | 24.540 | 23.921 | -4.285 | 2.800 | 8.585 | 14.352 | 21.455 |
| -8.892 | 23.749 | -14.237 | 6.302 | 4.065 | 27.311 | -15.929 | 22.394 | 3.490 | 11.882 | 29.558 | 24.672 | 19.913 | 5.575 |
| -22.449 | -25.503 | 19.542 | 20.530 | -22.029 | -29.585 | 15.765 | 2.178 | 8.546 | 9.126 | 8.708 | 10.295 | 5.007 | -6.498 |
| 7.863 | 11.065 | -21.317 | 3.011 | 8.366 | -26.913 | -20.437 | 15.063 | 26.152 | -2.242 | 4.749 | 20.217 | 19.090 | -21.253 |
| 4.382 | 25.166 | -15.568 | -12.961 | -1.845 | 12.416 | -26.929 | 7.518 | 12.135 | 11.974 | 19.958 | 11.002 | 21.540 | -2.307 |
| 5.419 | -10.792 | 15.781 | -23.300 | -15.570 | -22.791 | 7.584 | -14.096 | -4.547 | 27.911 | 10.123 | 13.147 | 10.661 | 15.117 |
| 1.538 | -24.470 | 17.935 | 19.749 | -7.451 | -27.721 | -23.447 | 22.695 | -16.176 | -11.025 | 25.052 | 15.918 | 17.822 | 15.900 |
| -8.536 | 19.517 | 7.875 | 0.919 | 19.259 | 7.430 | 24.038 | 25.092 | 25.381 | 10.503 | 27.409 | 27.369 | 5.440 | 14.483 |
| -5.669 | 1.499 | -10.088 | 2.263 | 9.846 | -29.669 | 20.599 | 10.561 | 22.698 | 6.506 | 24.233 | 8.852 | 6.155 | 9.469 |
| 11.156 | 29.098 | 10.950 | -24.285 | 5.960 | 23.182 | 11.198 | 7.566 | -15.024 | 26.214 | 22.697 | 18.684 | 4.963 | 6.992 |
| 17.325 | 26.630 | -1.397 | 18.027 | -21.702 | 5.989 | -0.894 | -6.251 | 10.709 | 2.468 | 21.543 | -6.893 | 14.305 | -1.131 |

| | | | | | | | | | | | | | |
|---|---|---|---|---|---|---|---|---|---|---|---|---|---|
| 27.864 | 4.229 | -23.182 | -21.165 | -0.030 | -21.239 | -24.884 | 14.967 | 3.317 | 8.299 | 28.725 | 21.345 | 2.953 | 23.363 |
| 1.590 | -18.422 | 18.315 | -2.043 | 16.649 | -28.489 | -5.807 | 23.869 | -4.055 | 29.259 | 25.741 | 20.137 | 22.435 | 16.697 |
| 4.412 | -23.472 | -26.976 | -14.416 | -2.134 | 27.653 | 16.461 | 12.057 | 13.215 | 23.384 | 1.350 | 25.544 | 2.659 | 20.411 |
| -4.521 | 27.856 | 3.435 | -0.918 | -7.375 | 6.008 | 20.883 | 9.671 | 28.106 | 26.382 | 24.022 | 10.752 | 11.001 | 23.978 |
| -13.005 | -9.028 | -29.680 | -2.985 | -8.182 | 19.430 | 5.449 | 8.995 | -14.374 | -5.025 | 9.276 | -3.090 | 24.012 | 9.942 |
| -5.568 | -22.624 | -19.915 | 8.212 | -10.385 | 3.755 | 15.516 | 20.065 | 13.969 | 28.666 | 20.416 | 26.136 | 15.959 | 2.210 |
| -17.900 | 8.508 | -22.576 | 11.298 | 23.215 | -19.282 | 8.057 | 1.738 | 29.264 | 12.678 | 16.772 | 23.385 | 2.473 | 8.259 |
| 0.557 | 18.075 | 23.473 | -16.553 | 7.968 | 25.257 | -2.472 | 14.240 | 29.788 | 27.675 | 24.855 | 25.974 | 20.012 | -1.034 |
| 14.656 | 7.540 | -28.789 | -0.144 | 22.903 | -27.734 | 12.869 | 21.959 | 19.496 | 22.013 | 20.107 | 8.820 | -1.597 | 9.277 |
| 2.424 | 25.284 | -1.055 | 18.661 | -16.932 | 18.036 | 17.678 | 27.269 | 12.168 | 21.034 | -15.680 | 20.749 | 2.933 | -13.203 |
| -0.487 | 5.748 | -30.513 | 7.867 | 19.792 | -25.205 | -7.665 | 19.358 | 22.322 | 7.374 | 15.550 | -0.769 | 18.994 | 13.817 |
| 14.997 | 4.114 | 6.861 | -27.766 | 17.309 | 6.927 | -4.219 | 25.731 | 9.612 | 27.593 | 24.281 | 23.179 | 2.269 | -6.267 |
| 7.752 | -0.542 | -15.456 | 22.293 | -27.437 | 10.001 | -16.955 | 3.459 | -6.469 | 28.233 | 19.632 | 15.766 | 25.261 | 6.663 |
| 15.985 | -23.149 | 2.317 | -25.366 | 10.362 | 5.517 | 23.804 | 0.846 | -22.385 | 25.605 | 12.995 | 19.325 | 7.208 | -2.463 |
| -3.115 | -23.675 | -5.984 | 23.650 | 18.560 | 7.609 | 20.125 | 4.978 | -4.688 | 20.389 | 8.291 | 24.719 | -11.868 | -9.931 |
| -3.768 | 26.359 | -23.085 | 6.450 | -4.977 | -17.984 | -25.998 | -3.321 | 28.189 | 14.315 | 0.796 | 5.565 | 20.316 | 24.655 |
| 5.273 | -0.021 | -14.342 | -22.023 | 26.476 | 26.638 | -11.460 | -6.824 | 14.222 | 11.534 | 10.582 | 21.197 | 13.935 | 16.750 |
| 3.298 | 6.963 | -16.165 | 15.989 | 7.193 | 19.434 | 7.236 | 10.318 | 30.078 | -27.314 | 12.375 | 26.171 | -13.169 | 6.338 |
| -4.483 | 5.008 | 19.472 | -16.244 | 12.968 | -27.420 | 14.814 | 26.265 | 20.538 | -4.824 | 27.297 | 10.653 | 6.430 | 17.990 |
| -11.488 | -24.360 | 28.315 | -30.423 | 8.461 | -23.438 | -17.238 | 20.774 | -3.286 | 21.512 | 20.076 | -3.613 | 21.514 | 2.621 |
| 10.855 | -5.892 | 2.803 | 27.213 | -12.031 | 1.179 | -15.036 | 21.648 | 18.342 | 2.285 | 9.255 | 18.902 | 18.290 | 8.639 |
| -20.998 | 20.723 | -8.284 | -7.461 | -21.456 | -26.242 | -13.114 | 27.276 | 13.577 | -7.210 | 13.922 | 12.426 | 14.755 | -10.206 |
| 2.573 | -14.746 | 10.703 | -10.937 | -17.661 | 7.818 | -9.708 | 17.658 | -1.567 | 23.145 | 15.015 | 22.548 | 8.868 | 25.715 |
| 9.110 | 19.403 | -18.945 | 21.112 | 0.728 | 12.412 | -4.874 | 14.454 | 22.059 | 15.765 | 16.753 | 16.806 | 13.078 | 15.579 |
| -26.947 | -4.260 | -23.198 | 24.399 | -11.319 | -28.895 | -4.422 | 4.959 | 1.222 | -12.134 | 23.254 | 15.740 | 24.609 | 1.155 |
| -8.257 | 8.799 | 12.678 | -1.059 | 2.880 | -1.838 | 23.195 | 22.832 | -6.459 | 5.005 | 15.656 | 21.700 | 11.678 | 27.312 |
| -22.462 | 5.850 | -23.756 | -20.040 | 5.282 | 18.925 | 20.482 | 4.405 | 4.899 | 17.428 | 1.032 | 8.493 | 28.123 | 5.952 |
| -17.187 | 22.863 | -21.586 | -21.872 | -10.127 | 7.205 | -1.690 | -0.119 | 16.956 | 25.565 | 22.107 | 14.021 | 25.078 | 13.224 |
| 1.335 | 18.091 | -28.501 | -5.230 | 16.604 | 10.027 | 0.347 | 25.301 | -20.138 | 26.981 | 29.559 | 27.864 | 16.469 | 16.327 |
| 5.651 | -26.399 | 10.093 | 9.482 | 8.370 | -22.247 | 16.355 | -22.437 | -18.839 | -0.778 | 7.238 | 21.151 | 20.426 | 23.106 |
| -3.659 | 6.839 | 22.097 | 19.947 | -4.328 | 14.742 | 1.526 | 12.133 | 28.020 | 13.010 | 25.215 | 11.721 | 12.646 | 20.765 |
| 22.269 | -24.539 | -1.997 | -26.741 | 23.352 | 25.476 | 2.400 | 27.784 | 16.689 | 28.135 | -18.168 | 14.726 | 6.761 | -8.546 |
| 14.590 | 10.011 | 9.120 | -17.899 | -19.625 | -9.957 | -1.979 | 5.288 | -13.142 | 18.965 | 13.865 | 19.780 | 5.455 | -2.509 |
| 3.036 | -27.330 | 3.220 | 17.485 | 8.652 | 16.797 | 8.574 | 17.071 | 26.023 | 26.593 | 18.156 | 21.647 | 15.520 | 26.334 |
| -20.933 | -8.372 | 8.203 | -29.196 | -12.480 | 4.892 | -12.828 | 3.318 | 19.135 | 26.002 | 26.571 | 12.630 | 13.622 | 17.181 |
| -11.154 | 11.206 | -7.709 | 3.066 | 15.534 | 27.421 | 9.767 | 14.424 | 7.761 | 14.309 | 27.042 | 0.297 | 25.829 | 9.332 |

| | | | | | | | | | | | | | |
|---|---|---|---|---|---|---|---|---|---|---|---|---|---|
| -4.445 | 22.200 | 2.151 | 22.649 | 7.259 | 6.504 | 1.446 | 26.300 | 25.171 | 23.259 | 8.624 | 19.964 | 23.109 | -1.169 |
| 0.909 | 13.407 | 4.545 | 8.017 | -15.034 | -25.764 | 2.076 | 18.957 | 15.359 | -25.145 | 24.791 | 25.704 | 18.478 | 16.471 |
| -5.332 | -24.958 | 24.050 | 19.604 | -5.836 | 1.513 | -4.126 | -2.659 | 11.651 | 10.696 | 12.490 | 25.146 | 22.086 | 19.338 |
| -3.438 | 17.411 | -6.640 | -22.735 | 15.079 | -8.966 | 2.907 | 2.242 | 22.426 | 1.593 | 15.430 | 17.788 | -3.845 | 24.537 |
| 4.520 | 2.896 | -12.703 | 0.698 | 14.466 | -16.989 | -22.916 | 7.014 | 20.116 | 25.121 | 13.905 | -0.686 | 10.233 | 20.958 |
| 0.113 | -15.386 | 6.074 | 23.339 | 24.902 | -20.340 | -23.775 | 13.136 | 20.315 | 21.881 | -1.990 | 23.604 | 3.330 | 18.433 |
| -5.518 | -12.088 | -30.841 | 20.881 | -1.310 | 28.233 | -4.878 | 4.964 | 27.665 | 15.866 | 13.282 | 29.464 | 21.260 | -0.532 |
| 15.328 | 7.071 | -10.763 | -24.385 | 8.714 | 14.377 | -16.228 | 6.500 | 10.944 | 19.437 | 11.446 | 29.864 | 26.147 | 0.357 |
| 8.489 | -24.145 | 8.052 | -5.689 | -6.094 | 26.786 | -21.591 | 4.368 | -10.677 | 19.040 | 6.298 | -3.886 | 21.823 | 10.632 |
| -17.085 | -13.059 | 13.413 | 4.247 | -27.001 | 9.379 | -16.431 | 19.619 | -25.317 | 26.333 | -0.466 | 11.273 | -0.519 | -0.090 |
| -16.573 | 25.430 | -29.977 | 10.281 | 13.287 | 5.670 | -17.477 | 23.981 | 17.799 | 15.390 | -26.408 | 16.096 | 22.500 | 24.806 |
| -1.653 | 11.621 | 18.671 | 26.130 | 16.399 | 20.982 | -4.473 | -4.114 | 19.366 | 19.845 | 7.463 | -4.516 | 18.630 | 27.749 |
| -16.352 | 8.610 | -7.517 | 27.335 | -0.210 | -18.987 | -0.745 | -8.879 | 9.145 | -26.042 | 4.731 | 18.515 | 17.383 | -7.511 |
| 3.295 | 0.009 | 12.473 | 27.708 | -19.302 | 18.152 | 5.856 | -3.536 | 28.198 | 20.630 | 20.766 | 0.397 | 26.400 | 21.567 |
| 1.492 | 26.853 | -29.828 | 19.743 | 2.552 | -10.805 | 3.690 | 6.999 | 16.078 | 24.514 | 9.435 | 12.468 | 18.703 | 12.617 |
| -0.004 | 24.954 | -8.697 | -8.632 | 7.112 | -26.641 | -4.546 | 0.417 | -2.401 | 1.405 | 1.832 | -0.836 | 14.735 | 27.575 |
| 3.737 | -0.442 | 15.885 | 27.839 | 24.347 | -0.113 | -7.102 | 29.798 | 1.467 | 3.131 | 12.313 | 27.563 | 5.036 | 21.478 |
| 7.871 | -17.047 | 26.841 | 18.325 | 4.425 | 21.524 | -5.259 | 26.663 | 19.380 | 2.073 | 18.343 | 18.593 | 25.294 | 16.875 |
| 14.899 | 25.077 | -27.213 | -4.712 | 9.727 | 14.653 | -7.565 | 2.957 | 4.141 | 18.683 | 17.352 | 27.734 | 7.756 | 19.079 |
| -13.920 | 20.980 | 24.581 | -9.810 | -7.166 | 28.054 | 16.590 | -2.900 | 3.213 | 9.222 | 13.226 | 26.123 | 27.194 | 8.535 |
| 1.283 | -23.241 | -14.207 | 26.565 | -0.396 | 23.226 | 13.518 | -17.867 | 25.427 | 17.678 | 20.216 | 9.295 | 15.730 | 7.148 |
| -4.594 | 13.333 | -10.404 | 24.763 | -14.343 | -25.436 | 6.160 | 18.033 | 26.134 | 6.376 | 15.653 | -0.096 | 28.210 | 26.237 |
| 18.765 | -9.484 | -6.836 | 11.711 | -18.621 | -9.061 | 3.187 | -3.602 | 23.835 | 0.748 | -20.216 | 14.602 | 11.157 | 15.897 |
| 19.358 | -22.303 | 13.763 | 7.955 | 24.624 | -3.971 | -10.126 | -4.717 | 28.148 | 3.923 | 8.974 | -25.765 | 9.943 | 14.623 |
| 0.620 | 21.755 | -15.544 | -21.067 | -9.345 | -18.759 | 19.061 | 11.019 | 25.828 | 25.121 | 5.515 | -0.897 | 21.527 | 9.613 |
| -4.011 | 16.334 | 23.596 | 13.444 | 5.576 | 7.512 | -26.846 | -7.364 | 20.242 | 3.251 | 7.838 | 17.031 | 23.980 | 25.240 |
| -22.075 | 10.974 | 21.885 | 1.554 | -13.274 | -0.398 | 24.354 | 14.849 | -6.975 | 2.493 | -16.018 | 18.095 | 15.157 | 16.257 |
| 11.737 | -11.109 | 12.932 | 26.525 | 1.225 | 16.761 | -24.709 | -16.273 | 11.605 | 26.552 | 27.757 | 26.833 | 26.560 | -16.158 |
| -1.170 | 25.946 | 14.815 | 11.491 | 15.782 | -20.890 | 15.014 | 8.314 | 7.943 | 6.481 | 14.318 | 1.491 | -28.699 | 29.037 |
| -9.071 | 15.508 | -2.094 | -7.193 | 14.707 | -22.515 | 11.503 | -6.688 | 9.618 | 22.476 | 4.539 | 2.898 | 9.065 | 17.204 |
| -21.114 | -18.331 | -29.312 | 6.857 | -20.778 | 22.520 | -25.803 | 1.652 | 18.644 | 18.658 | 22.114 | 8.599 | 15.666 | 0.699 |
| 22.996 | -9.065 | -5.244 | -3.168 | -10.948 | 22.111 | -24.350 | 10.240 | 3.638 | 21.239 | -4.469 | 23.044 | 16.509 | -7.804 |
| -5.553 | 22.657 | 14.511 | 21.369 | -24.322 | 19.285 | -7.050 | 4.611 | 0.470 | 20.705 | 15.694 | 21.725 | 10.405 | -11.795 |
| -9.417 | 18.911 | -30.514 | 10.906 | 9.600 | -23.099 | 8.169 | 18.619 | 8.871 | 0.341 | -7.195 | 24.630 | 22.561 | 28.680 |
| -10.179 | -6.151 | -27.026 | 12.217 | -23.515 | -8.736 | -4.367 | 30.116 | 15.665 | -13.733 | 13.973 | 9.392 | 17.232 | -4.086 |
| 14.786 | 4.418 | -28.779 | 19.262 | 11.149 | 18.506 | 25.132 | -5.047 | 12.223 | 9.403 | 16.732 | 27.279 | 7.097 | 15.578 |

| | | | | | | | | | | | | | |
|---|---|---|---|---|---|---|---|---|---|---|---|---|---|
| 9.010 | -7.612 | 25.380 | 8.406 | -23.027 | -15.349 | 25.542 | 8.977 | 5.409 | 9.216 | 17.625 | 20.039 | 21.853 | 4.419 |
| -12.747 | -0.390 | -3.340 | -4.867 | 16.259 | -7.521 | 22.485 | 17.011 | 24.294 | 7.215 | 22.275 | 3.280 | 19.030 | 25.294 |
| -2.008 | 3.857 | -14.986 | -5.742 | 12.069 | -25.136 | -9.443 | -2.979 | 26.694 | 4.856 | 4.780 | 23.575 | 9.207 | 8.942 |
| 11.802 | 16.858 | 23.160 | -21.712 | -5.475 | 23.388 | 15.523 | 12.537 | -26.567 | 21.110 | 3.874 | -1.651 | 16.050 | 20.774 |
| -5.325 | 11.420 | -9.460 | 13.709 | -17.356 | 16.371 | 17.615 | 19.601 | -7.108 | 13.234 | 5.694 | 19.766 | 3.630 | 28.336 |
| -0.712 | -3.039 | 3.220 | 3.334 | -8.601 | 26.140 | -3.516 | -5.840 | 6.196 | -11.923 | -2.539 | 12.363 | 26.246 | 17.230 |
| -8.791 | 2.979 | 5.567 | -12.843 | -15.360 | 20.581 | 8.208 | 29.680 | 15.339 | 22.325 | 29.620 | 21.876 | 5.686 | 18.732 |
| 21.932 | -9.035 | 11.353 | 20.557 | -16.651 | -24.725 | -17.465 | 10.406 | 27.192 | 21.523 | 8.342 | 7.618 | -16.434 | 13.201 |
| -17.876 | 2.406 | -8.405 | 16.739 | 1.764 | 9.693 | -11.599 | -1.476 | 18.605 | 17.986 | 25.557 | 19.366 | 29.596 | 25.068 |
| -10.309 | -26.075 | 16.467 | -3.181 | -4.928 | -22.333 | -28.369 | 16.940 | 13.301 | 24.543 | 9.653 | 20.351 | 24.452 | 22.211 |
| 13.041 | 17.339 | -19.617 | -1.768 | 14.775 | -25.772 | 14.687 | 14.399 | 12.402 | 3.555 | -1.136 | 9.658 | 14.475 | 12.104 |
| 23.189 | 8.610 | 6.183 | -8.843 | 12.385 | 22.117 | -0.163 | 1.484 | -2.654 | 1.667 | 22.920 | 22.587 | 23.642 | 23.784 |
| -19.202 | 23.885 | 27.242 | -28.164 | 9.436 | -10.932 | 17.047 | 15.719 | 5.946 | -0.880 | 24.985 | 23.036 | 5.026 | -8.930 |
| -21.304 | 18.528 | -0.220 | 11.911 | -7.708 | 4.991 | -23.526 | 16.113 | -4.654 | -17.630 | 5.365 | 11.660 | 29.293 | 18.526 |
| 1.199 | -1.795 | 8.712 | -11.704 | -4.319 | -15.536 | 26.208 | 11.653 | 14.142 | 15.264 | 28.038 | 15.832 | 22.511 | 17.928 |
| 9.356 | 4.064 | 9.464 | -19.922 | 1.528 | 2.882 | -8.007 | 2.983 | 15.538 | -9.507 | 3.293 | 29.847 | 19.426 | 5.982 |
| -3.692 | 24.674 | -23.913 | -13.754 | 9.360 | -22.861 | -23.808 | -2.007 | 25.631 | 23.231 | 27.445 | 25.207 | 21.064 | 13.983 |
| 8.412 | 7.184 | 23.609 | 9.793 | -13.008 | 15.706 | 21.977 | 5.174 | 23.416 | -11.640 | -16.559 | 2.466 | -21.676 | 27.740 |
| -1.836 | -8.235 | -27.438 | 15.579 | -22.133 | 23.938 | -1.123 | 15.155 | 21.508 | 20.739 | 18.241 | 20.986 | 22.450 | 7.421 |
| -24.163 | -9.016 | 26.260 | 1.685 | 10.626 | 22.371 | 21.020 | 27.602 | 16.192 | 9.579 | 26.398 | 16.185 | 17.595 | 15.223 |
| -6.916 | -14.423 | 11.566 | -2.485 | -15.743 | 2.873 | 0.723 | 28.976 | 19.586 | 5.883 | 21.026 | 31.377 | 20.589 | 2.690 |
| -23.176 | -25.093 | 4.639 | -24.834 | 20.014 | 10.619 | -8.375 | -10.906 | -3.362 | 7.376 | -6.155 | 29.178 | 19.059 | 8.768 |
| 7.990 | 13.342 | 2.340 | 10.732 | -3.269 | 20.068 | 18.373 | 5.401 | 4.659 | 3.490 | 6.351 | 14.106 | -12.706 | 19.808 |
| -7.642 | 15.967 | 16.210 | -26.568 | 7.147 | 9.155 | -3.447 | -1.230 | 3.272 | 10.075 | 7.381 | 28.900 | 30.646 | 12.083 |
| -22.203 | 10.190 | -21.559 | 24.458 | -10.300 | 11.215 | -17.464 | 19.184 | 9.901 | 2.538 | -2.870 | 22.343 | 2.731 | 20.928 |
| 22.955 | -21.359 | -28.997 | -9.409 | -18.469 | 1.898 | 8.079 | 8.203 | 12.147 | 17.334 | 19.431 | 21.688 | 21.194 | 29.381 |
| -18.483 | 12.145 | -10.320 | 12.408 | 1.019 | -0.355 | -3.895 | -0.134 | 26.972 | 22.758 | 17.599 | 30.016 | 13.230 | 30.141 |
| -23.828 | 7.045 | 0.131 | 2.715 | -7.949 | -14.151 | 21.409 | 14.830 | -4.518 | 16.492 | 16.542 | 20.101 | 18.210 | 19.101 |
| -11.455 | -20.302 | 15.539 | -0.185 | -1.317 | -1.144 | -23.404 | -13.003 | 22.778 | 22.155 | 3.606 | 6.616 | 10.302 | 18.009 |
| -3.780 | -4.590 | -5.937 | 14.863 | 24.149 | -2.437 | -18.769 | -20.977 | 18.638 | 17.431 | -27.165 | 17.814 | 0.280 | 23.878 |
| -0.769 | -23.433 | -4.139 | 20.842 | -8.829 | -20.050 | -5.630 | -24.886 | 17.862 | -12.283 | 24.135 | 18.448 | 28.108 | 6.415 |
| 5.899 | -3.843 | -19.992 | -4.900 | 0.504 | -16.370 | -11.291 | 26.478 | 8.860 | 14.421 | 21.576 | 26.186 | 11.251 | 8.427 |
| -20.768 | -6.018 | 6.006 | 5.954 | -8.281 | -1.477 | 26.977 | -4.712 | 20.709 | 13.044 | 13.177 | 28.285 | 14.651 | 12.629 |
| 23.224 | 8.014 | 20.130 | 8.398 | 13.561 | -19.047 | 9.135 | -6.644 | 1.356 | -1.889 | 4.395 | 29.285 | 24.542 | -21.587 |
| -21.139 | -3.569 | 8.684 | -8.844 | 19.000 | 21.961 | 21.103 | -18.793 | 23.860 | -20.379 | 11.983 | 14.697 | 28.918 | 9.677 |
| -25.096 | 12.136 | 16.037 | 11.580 | 21.434 | 8.124 | -24.672 | 3.948 | 12.501 | 1.047 | 13.940 | 30.769 | 9.272 | 23.472 |

| | | | | | | | | | | | | | |
|---|---|---|---|---|---|---|---|---|---|---|---|---|---|
| -19.386 | 15.814 | 12.302 | -24.679 | -8.973 | 9.284 | -9.751 | 21.574 | 12.886 | 2.360 | 10.580 | 15.734 | -3.663 | 3.983 |
| 14.124 | -16.563 | 7.194 | -11.126 | -8.818 | 19.446 | -19.537 | -22.843 | 0.199 | 4.784 | 0.102 | 11.658 | 30.298 | 19.832 |
| 25.112 | 0.580 | -2.031 | 9.109 | 9.956 | -18.616 | 4.690 | 14.775 | 17.694 | 0.266 | 20.019 | 6.538 | 27.015 | 4.671 |
| -18.608 | 18.950 | 0.978 | -13.245 | -11.343 | 22.927 | -3.446 | 8.153 | 3.363 | 17.350 | -20.330 | 9.402 | 26.386 | 11.150 |
| 13.284 | -0.954 | -29.673 | 26.571 | -7.654 | 20.613 | 12.052 | 31.064 | 23.687 | -0.697 | 24.765 | 25.534 | 25.569 | 16.518 |
| -5.495 | 20.337 | 16.542 | -0.660 | -20.426 | -8.229 | 25.105 | 13.697 | 22.191 | 18.298 | 20.883 | 27.704 | 19.601 | 13.912 |
| 15.869 | 2.802 | -12.441 | -27.479 | -7.020 | 4.107 | -12.520 | -28.430 | -2.287 | 7.891 | 14.982 | -1.301 | 21.681 | 9.672 |
| 6.330 | -15.767 | 4.978 | -17.875 | 19.808 | -9.182 | -18.155 | -17.835 | 4.689 | 5.333 | 7.332 | -5.485 | -19.089 | 0.282 |
| -14.203 | 9.576 | 0.327 | -19.433 | 11.637 | -16.967 | 11.211 | 10.471 | 9.339 | 6.504 | 3.548 | 23.393 | 14.872 | 27.161 |
| 18.798 | -16.974 | -0.824 | 5.197 | -17.329 | -10.603 | 15.060 | 7.460 | 10.177 | 20.538 | 1.050 | 21.504 | 9.190 | 22.243 |
| 15.935 | 19.536 | 13.831 | -26.379 | -4.088 | 24.615 | -20.680 | 5.820 | 12.303 | 7.377 | -6.492 | 28.406 | 23.443 | 9.303 |
| -25.360 | -19.322 | 15.368 | 15.244 | 6.370 | 21.721 | -27.357 | 29.788 | 17.694 | 4.978 | 19.330 | 14.509 | -29.956 | 28.666 |
| 21.512 | -21.899 | -6.603 | 2.616 | 18.325 | 4.061 | 8.483 | 23.416 | 22.305 | 10.360 | 14.627 | 27.996 | 31.928 | 22.871 |
| 13.437 | -25.719 | 10.642 | -17.062 | -3.065 | 24.315 | 28.151 | 13.864 | 18.649 | 19.742 | 4.090 | 28.866 | 13.914 | 16.172 |
| 12.848 | 18.941 | -5.570 | 16.407 | 0.668 | 11.697 | -25.984 | -5.970 | 16.366 | 17.597 | 17.754 | 17.444 | 17.121 | 0.372 |
| -24.213 | -17.728 | -16.706 | 10.399 | -8.232 | 6.615 | -0.334 | 19.741 | 17.882 | 14.712 | 24.142 | 21.091 | -0.732 | 1.498 |
| -21.619 | 8.302 | 25.150 | 7.745 | 4.184 | 14.683 | -29.334 | 15.518 | -0.720 | -17.091 | 25.444 | -1.412 | 19.964 | 13.934 |
| 19.760 | 20.426 | 11.780 | 8.808 | -9.874 | 16.853 | 20.825 | -1.151 | 10.129 | -1.127 | -4.150 | 4.366 | 26.601 | 24.971 |
| 14.774 | 1.676 | -25.385 | 8.125 | 26.381 | -3.699 | -12.129 | 21.912 | 10.800 | 19.087 | 10.285 | 28.473 | 18.302 | 7.679 |
| 3.743 | -13.192 | 6.261 | 20.307 | 13.805 | 23.191 | 5.534 | -9.389 | -13.676 | 21.827 | 13.620 | 10.544 | 16.827 | 27.933 |
| 15.193 | 22.775 | 1.008 | 17.123 | -6.152 | -23.182 | 25.595 | 30.606 | -5.551 | -0.563 | 22.428 | 17.118 | 1.165 | 29.744 |
| 13.133 | 5.784 | 19.674 | 18.069 | 17.647 | -22.042 | 11.681 | 16.547 | 24.254 | 11.493 | 21.106 | 21.057 | 22.384 | 26.615 |
| -10.214 | -17.067 | -19.815 | 0.922 | -16.272 | -25.587 | 0.380 | 4.218 | 20.311 | 7.529 | 15.683 | 12.866 | 29.501 | 11.137 |
| 23.010 | 11.497 | 8.166 | 22.848 | -7.791 | 4.066 | 27.108 | 14.525 | 3.582 | 0.119 | 13.973 | 8.675 | 25.971 | 0.984 |
| 26.861 | 4.406 | -7.258 | -2.560 | 27.290 | 22.304 | -27.715 | 15.982 | 18.479 | 24.545 | 23.905 | 11.108 | 12.270 | 20.071 |
| -21.294 | 12.956 | -0.302 | -4.781 | 3.770 | 11.991 | 22.572 | -21.482 | 11.957 | 23.080 | 1.684 | 23.012 | 8.439 | 13.410 |
| -25.308 | -15.949 | -16.393 | 12.950 | 23.701 | -24.876 | 18.362 | -17.171 | -9.397 | 9.365 | 19.978 | 22.203 | 27.217 | 22.514 |
| -17.654 | 23.932 | -8.782 | 4.178 | 28.106 | 5.020 | 7.906 | 19.314 | -22.479 | 27.894 | 3.771 | 22.745 | 7.162 | 11.069 |
| -11.381 | -8.421 | 3.675 | 5.977 | 9.273 | 22.718 | -19.802 | -2.428 | 18.844 | -4.633 | 18.308 | 17.972 | 29.998 | 19.564 |
| -19.311 | 1.677 | 2.664 | 26.109 | -26.686 | -16.680 | -29.521 | 17.325 | -7.434 | -3.110 | 25.588 | 6.385 | 5.978 | 18.485 |
| 11.690 | 6.330 | 15.448 | -24.647 | -17.663 | 14.573 | -12.141 | -6.850 | 10.368 | 21.210 | 5.206 | 12.342 | 10.647 | 8.874 |
| 8.398 | 15.563 | -26.919 | -22.291 | 20.212 | 23.479 | -18.605 | 7.665 | 7.930 | 19.911 | 2.587 | 5.203 | 18.447 | 30.330 |
| 9.189 | -24.685 | 23.159 | 23.072 | 0.245 | -17.254 | 14.268 | 20.619 | 20.227 | 7.818 | -1.021 | 25.069 | -4.145 | 4.719 |
| 19.544 | -26.442 | 16.396 | 7.913 | -4.373 | -24.797 | -26.815 | -1.262 | 15.313 | 23.548 | -17.462 | 28.695 | 15.261 | -4.652 |
| -2.515 | 20.188 | -8.981 | -5.638 | 26.664 | 27.914 | 9.093 | 10.484 | 7.706 | 26.449 | -4.790 | 7.027 | 20.957 | 10.140 |
| 29.038 | 1.098 | -21.464 | 26.251 | 5.592 | 1.296 | -15.542 | -21.975 | -23.042 | 24.253 | 16.642 | 24.762 | 19.819 | 21.868 |

| | | | | | | | | | | | | | |
|---|---|---|---|---|---|---|---|---|---|---|---|---|---|
| -1.495 | -19.903 | -5.252 | 2.839 | 5.061 | 24.359 | 7.626 | 13.605 | -27.447 | 22.343 | 22.726 | 18.060 | 23.118 | 8.609 |
| 28.014 | 25.104 | -11.343 | -11.414 | -18.443 | -17.966 | -6.027 | 0.600 | 24.484 | 9.064 | 15.244 | -2.157 | 9.382 | 14.714 |
| 21.076 | -22.963 | -6.952 | -9.881 | -8.664 | 19.084 | 20.542 | -4.895 | 14.172 | 24.985 | 21.140 | 26.160 | 3.577 | 12.689 |
| 8.495 | 25.728 | 6.313 | -18.459 | 8.330 | 13.288 | 24.492 | 28.255 | 8.682 | 14.095 | -0.836 | 4.676 | 9.053 | 25.871 |
| 14.926 | -10.910 | 25.148 | 24.239 | 28.626 | -16.673 | 27.059 | 18.643 | 19.060 | 25.566 | -10.998 | 16.478 | 27.665 | -2.962 |
| -23.180 | 17.485 | 10.335 | 4.490 | 21.393 | 12.524 | 30.494 | 28.995 | 2.277 | 21.896 | 8.630 | 20.727 | 24.656 | -9.544 |
| 31.011 | -19.331 | 21.167 | 1.640 | -22.378 | 19.790 | -14.257 | 2.971 | 17.285 | 27.529 | 13.299 | 11.587 | 11.891 | 24.724 |
| 28.674 | 3.448 | 20.201 | 13.089 | -6.984 | -27.414 | 23.623 | -9.863 | 17.559 | 12.393 | 12.800 | 3.592 | 14.319 | 19.196 |
| 27.722 | 19.728 | 8.590 | -2.471 | 6.529 | 27.513 | 7.010 | -17.744 | 26.375 | 1.298 | 10.260 | 20.179 | 20.031 | 9.133 |
| 26.144 | 14.395 | -27.532 | 10.962 | 5.582 | -17.979 | 24.847 | 14.047 | 23.161 | 21.077 | 16.868 | 26.849 | 29.486 | 22.675 |
| -2.189 | 13.119 | -12.432 | 17.616 | 23.093 | -24.755 | 21.511 | 13.633 | -3.414 | 23.210 | 15.214 | 23.508 | 21.675 | 5.843 |
| -20.746 | 12.252 | -26.643 | -5.547 | 15.894 | 25.784 | 22.267 | 11.067 | 10.208 | 7.360 | -9.047 | 20.554 | 16.454 | 2.940 |
| -22.248 | 23.088 | 22.302 | 4.939 | -1.114 | -19.591 | -1.727 | 9.027 | 24.013 | 21.912 | -11.619 | 6.864 | 25.294 | 7.265 |
| 11.124 | 4.317 | -1.136 | -3.858 | 11.752 | -6.274 | 11.438 | 7.372 | 25.140 | 10.018 | 17.787 | 18.846 | 5.783 | 4.658 |
| 26.002 | 20.636 | -7.133 | -7.420 | 7.046 | -16.676 | -30.066 | -15.093 | 14.802 | 16.525 | -5.227 | 19.719 | 0.028 | -12.234 |
| -29.280 | 6.652 | -24.877 | -17.127 | 3.046 | 23.850 | -11.199 | 14.572 | -5.739 | 15.105 | 11.556 | 26.050 | 26.744 | 25.361 |
| -14.378 | 11.495 | 17.015 | -16.047 | 10.269 | 3.717 | -25.073 | 7.736 | 20.904 | -1.712 | 16.599 | 7.834 | 10.407 | 14.665 |
| -12.084 | 19.958 | 0.927 | 2.858 | 4.748 | -17.254 | 21.989 | 21.139 | 25.761 | -1.785 | 18.502 | -1.416 | 20.571 | 25.377 |
| 8.751 | -23.328 | -6.384 | -12.603 | -2.465 | -2.908 | -9.998 | 17.161 | 7.579 | 22.319 | 4.771 | 7.232 | 18.229 | 18.221 |
| 11.514 | 18.880 | -9.640 | 18.023 | -4.429 | -29.214 | 28.081 | 18.361 | -10.562 | 24.281 | 13.937 | 5.479 | 19.610 | 10.766 |
| 16.341 | -13.075 | 4.099 | -14.878 | -20.922 | -10.346 | -17.748 | -2.872 | -6.891 | 26.436 | 23.280 | 9.766 | 17.291 | 20.785 |
| 21.534 | 23.300 | -25.041 | -1.314 | -5.893 | 21.754 | -27.967 | 21.612 | -23.739 | 3.726 | 17.146 | 8.952 | 25.388 | 15.138 |
| -2.699 | 15.780 | 4.707 | -8.509 | -19.710 | 11.494 | 23.761 | 11.395 | -6.403 | 29.114 | 12.664 | 17.662 | 24.443 | -8.400 |
| 12.575 | 10.603 | -1.998 | -9.404 | 24.999 | -28.388 | 28.909 | -6.094 | 6.821 | 18.333 | 15.921 | 27.525 | 9.000 | 23.480 |
| 11.768 | 13.482 | -8.784 | -5.032 | 17.995 | 16.856 | 28.512 | 4.813 | 22.918 | 20.778 | -8.597 | 7.518 | 15.093 | 12.667 |
| -25.786 | 17.237 | 10.482 | -5.537 | 9.785 | 1.961 | 12.873 | 26.060 | 4.910 | 12.949 | -11.071 | 26.691 | 5.380 | 2.248 |
| 10.171 | 1.949 | 17.034 | -18.183 | -16.701 | 9.134 | -27.399 | -1.989 | 28.536 | 15.760 | -12.467 | 21.839 | 28.370 | 21.494 |
| -6.823 | -20.557 | -6.871 | 15.377 | -15.158 | -23.341 | 26.568 | -10.742 | -24.680 | 20.026 | 0.714 | 23.761 | -6.213 | 27.605 |
| -25.051 | -18.364 | -7.545 | 0.966 | -23.944 | -4.491 | -12.843 | 9.536 | 2.425 | 12.409 | -4.559 | 12.669 | -2.248 | 16.627 |
| -29.089 | -25.685 | 3.190 | 14.342 | 20.951 | -25.183 | -28.490 | 15.957 | 16.207 | 22.228 | -1.546 | 19.758 | 1.616 | 7.968 |
| -10.528 | -0.487 | -20.419 | -1.263 | 18.443 | 27.168 | 30.304 | -14.625 | 12.876 | 7.551 | 7.131 | 17.992 | 22.870 | 10.360 |
| -8.069 | -20.035 | 20.521 | 15.884 | -28.121 | -19.706 | -10.742 | 26.079 | 19.836 | 1.876 | 11.993 | 18.333 | 22.248 | 4.728 |
| -13.680 | 25.479 | -12.601 | -12.249 | 29.666 | -10.802 | 16.546 | 23.942 | 22.360 | 24.681 | 18.606 | 23.951 | 19.647 | 26.394 |
| -20.185 | 16.519 | 11.645 | 18.801 | -7.626 | 12.192 | -7.069 | 20.586 | -28.412 | 10.842 | 7.569 | 20.836 | 10.071 | 20.015 |
| -6.052 | -19.242 | -3.981 | 2.928 | -18.617 | 29.954 | 8.759 | 21.558 | -6.814 | 23.961 | 6.796 | 17.748 | 4.972 | 17.819 |
| -28.168 | 12.153 | -2.283 | 11.588 | -10.380 | -17.821 | 9.722 | -1.700 | 19.213 | 15.018 | 5.495 | 10.767 | 21.500 | -17.799 |

| | | | | | | | | | | | | | |
|---|---|---|---|---|---|---|---|---|---|---|---|---|---|
| -13.917 | -29.300 | 14.681 | 1.415 | -26.982 | -0.389 | 7.545 | -13.440 | 7.949 | 28.477 | 25.233 | 19.634 | 28.787 | 13.185 |
| 26.288 | 7.484 | 18.607 | -1.929 | 19.510 | -12.983 | -28.562 | 2.174 | -4.235 | 20.403 | 17.934 | 25.629 | 17.413 | 23.195 |
| 16.085 | 21.008 | 2.998 | 6.437 | 22.652 | 19.076 | 19.657 | -8.598 | 26.494 | 23.053 | 1.393 | 14.690 | 24.884 | 2.395 |
| 22.416 | 6.394 | -13.812 | -5.782 | 8.140 | 22.738 | 17.750 | -16.715 | -17.103 | 25.811 | 14.395 | 16.090 | 14.016 | -11.878 |
| 4.318 | 22.440 | -22.609 | 5.779 | -0.454 | 1.060 | 29.487 | 7.215 | 2.876 | 22.207 | 2.198 | -2.505 | 28.899 | 9.081 |
| -17.483 | 5.222 | 20.965 | 14.884 | 30.136 | -4.939 | 16.595 | 18.694 | 10.567 | 2.688 | 4.642 | 6.470 | 21.557 | -27.345 |
| -4.464 | 2.321 | 19.594 | -0.147 | 11.599 | 24.397 | 11.303 | 5.106 | 22.745 | 24.611 | 2.796 | 25.246 | 1.304 | -7.171 |
| 12.049 | 26.604 | -18.445 | -18.270 | -22.348 | -3.267 | -11.713 | -26.723 | -25.830 | -2.924 | 20.764 | 15.462 | 24.967 | 12.783 |
| 24.016 | 14.971 | 7.337 | -16.165 | -24.169 | -12.319 | 20.817 | 8.664 | 8.836 | 13.731 | 10.682 | 9.312 | 14.633 | 15.515 |
| -26.980 | -2.939 | 1.266 | -2.399 | 14.311 | 17.591 | -29.455 | 4.360 | 19.534 | 29.690 | 23.456 | 4.611 | 17.002 | 5.373 |
| -25.803 | 23.485 | -12.127 | 15.910 | 6.750 | -29.252 | 24.618 | -6.565 | 28.241 | -1.833 | 13.424 | 17.437 | 5.080 | 19.081 |
| 14.963 | 1.359 | -9.479 | -14.923 | 0.628 | -23.006 | -22.968 | -1.386 | 2.609 | 7.580 | 14.653 | 4.984 | 20.039 | 11.800 |
| 19.362 | -7.176 | 18.621 | -0.083 | 4.655 | 24.001 | 27.287 | -10.336 | 11.635 | 16.621 | -8.940 | 5.861 | 27.313 | 25.296 |
| -6.128 | -30.035 | -7.242 | 19.740 | 15.148 | -20.850 | -20.140 | -4.904 | 17.583 | 12.415 | 8.124 | 7.548 | 29.111 | 6.732 |
| -4.560 | 19.586 | -21.410 | 11.406 | 29.244 | -21.804 | -15.344 | 6.237 | 28.829 | 24.004 | 19.720 | 22.088 | 9.615 | 18.560 |
| 23.304 | 20.473 | 4.445 | -7.613 | 13.955 | -30.191 | -17.905 | 24.337 | -2.959 | -0.602 | 2.255 | 21.111 | 25.503 | 17.876 |
| 24.107 | 6.739 | -11.332 | 24.392 | -7.116 | -3.416 | 25.857 | 15.678 | 14.972 | 21.757 | 9.249 | 9.900 | 15.663 | 19.204 |
| 25.289 | -9.224 | 9.064 | 15.050 | 30.841 | 8.389 | -23.468 | 15.014 | 24.867 | 29.150 | 13.609 | 16.757 | 27.516 | 14.597 |
| 12.663 | -27.453 | 16.440 | 9.929 | 19.855 | -16.315 | 19.027 | -9.595 | 1.177 | 7.060 | 6.840 | 22.806 | 20.719 | 8.892 |
| -0.519 | 27.469 | 5.296 | 17.203 | 23.501 | 15.789 | -0.126 | 19.355 | 20.444 | -27.180 | 4.950 | 23.518 | 5.832 | 26.001 |
| -23.949 | -0.094 | 9.468 | 21.174 | -6.358 | 25.707 | 20.124 | 20.056 | 16.851 | 3.648 | 18.598 | 13.978 | 16.162 | 12.006 |
| 18.841 | 4.637 | -8.565 | -8.606 | -30.379 | -25.034 | 25.770 | 12.369 | 22.933 | -5.959 | -13.842 | 14.911 | 12.499 | 24.014 |
| -8.658 | -6.933 | 7.154 | -21.311 | 11.326 | -7.092 | 26.549 | 13.075 | 25.668 | 22.458 | 25.623 | -3.193 | 23.275 | 16.423 |
| -24.679 | -14.326 | -22.298 | -5.413 | -26.719 | 4.428 | 2.875 | 27.550 | 18.222 | 28.112 | 14.874 | 1.600 | 26.271 | 7.227 |
| -20.465 | 11.834 | 1.498 | -9.601 | -4.258 | -20.543 | -23.921 | 1.885 | 14.169 | 23.695 | -1.588 | -5.248 | 3.638 | -14.552 |
| 16.654 | -18.322 | -24.944 | 25.566 | -21.261 | -16.804 | 29.558 | 23.241 | 15.781 | 24.665 | 14.243 | -22.937 | 18.622 | 13.336 |
| 25.078 | -11.212 | -17.378 | 15.674 | 23.943 | 23.817 | 1.028 | 25.198 | 6.605 | 30.197 | 12.314 | 20.685 | 26.630 | 18.399 |
| 22.821 | 22.539 | -23.180 | 8.588 | 24.507 | -19.971 | -2.243 | 18.402 | 27.595 | 26.140 | 24.190 | 5.130 | 7.628 | 1.180 |
| -23.311 | -2.658 | -15.174 | 4.753 | 9.547 | -27.326 | 21.800 | 2.211 | 19.042 | 23.343 | 4.017 | 19.683 | 16.488 | -8.097 |
| 10.216 | -17.626 | -20.161 | 11.744 | -26.488 | 18.074 | 9.292 | 20.481 | 3.439 | 13.679 | 10.132 | 2.434 | 8.515 | 17.655 |
| -2.789 | 19.155 | 13.934 | -13.101 | 22.193 | -15.005 | 26.955 | 7.768 | 29.538 | 15.630 | 26.034 | 20.286 | 26.760 | 15.031 |
| -7.119 | 26.358 | 10.475 | -0.512 | 22.746 | -21.236 | -30.366 | 5.722 | 12.582 | 14.554 | 9.070 | 5.889 | 7.585 | -6.108 |
| -4.230 | 16.674 | 6.104 | 25.492 | 7.029 | 19.582 | -26.633 | 0.439 | 27.450 | 10.517 | 13.246 | 24.539 | 13.432 | 10.255 |
| 14.712 | 15.658 | -22.417 | -2.764 | -7.987 | -26.255 | 26.063 | -0.074 | 14.916 | -15.729 | 15.288 | -9.435 | 8.779 | 25.294 |
| 8.786 | -19.084 | -10.873 | -15.990 | -6.989 | 19.005 | 7.145 | -25.519 | 19.611 | 11.014 | 8.400 | 18.535 | 3.033 | 11.519 |
| 17.605 | 17.547 | 7.486 | -15.350 | -26.307 | -6.392 | 23.174 | 22.092 | 3.731 | 27.483 | 18.802 | -21.885 | 21.475 | 18.075 |

| | | | | | | | | | | | | | |
|---|---|---|---|---|---|---|---|---|---|---|---|---|---|
| -0.290 | -27.977 | -23.230 | 23.790 | -31.035 | -11.976 | 17.997 | 19.184 | 24.301 | -0.062 | 17.197 | 23.635 | -0.634 | 6.183 |
| -6.812 | -11.090 | 20.377 | 3.226 | 17.399 | 17.569 | 14.555 | 11.864 | 9.668 | 25.676 | 24.697 | 17.647 | 17.392 | -3.301 |
| 23.348 | -10.216 | 16.868 | -11.077 | 29.565 | 6.913 | -2.373 | 7.078 | 28.466 | 20.405 | 23.700 | 7.746 | 13.048 | 19.848 |
| 6.373 | -15.974 | -7.857 | 10.177 | -21.807 | -14.203 | -12.053 | -6.237 | -3.805 | -4.911 | -2.885 | 18.281 | 27.709 | 26.466 |
| -12.771 | -0.269 | 8.986 | -22.292 | -9.429 | -22.072 | -29.410 | 13.762 | 25.007 | -23.627 | 22.202 | -5.238 | 7.965 | 22.080 |
| -1.004 | 26.873 | 14.093 | 21.727 | 9.994 | 20.078 | 18.139 | -4.244 | 26.013 | 7.064 | 21.493 | 19.131 | 2.036 | 16.719 |
| -15.698 | 24.494 | -3.075 | -23.858 | 26.365 | 19.598 | 5.134 | 11.824 | 11.171 | 9.164 | 27.775 | 17.115 | 21.105 | -2.717 |
| -24.336 | 28.367 | -8.311 | -2.314 | -30.269 | 6.402 | 0.665 | 12.623 | 3.692 | 17.502 | 15.854 | 24.771 | 4.663 | 13.633 |
| -1.658 | -9.220 | 3.072 | -4.419 | -18.790 | -27.759 | 13.401 | 19.577 | 13.461 | -10.468 | 8.023 | 7.924 | 3.889 | 21.153 |
| -3.570 | -13.403 | -3.979 | 3.112 | 27.134 | -31.490 | 21.819 | 8.674 | -2.481 | 15.316 | 23.191 | 18.019 | 18.789 | 2.798 |
| 14.586 | 13.736 | 5.907 | 25.974 | 25.633 | 1.212 | 26.775 | -1.138 | 29.592 | 0.783 | 25.446 | 15.181 | 24.703 | 25.304 |
| 1.366 | 20.606 | 16.141 | 5.108 | 21.474 | -8.986 | -27.262 | -25.243 | 20.372 | -1.326 | -2.173 | 22.199 | 11.616 | 14.008 |
| 21.555 | -18.494 | -10.146 | -18.176 | 27.984 | 26.339 | -9.987 | 21.351 | 1.791 | 11.501 | 15.765 | 13.513 | 17.021 | 6.666 |
| -22.300 | -26.713 | -4.699 | -19.513 | -28.408 | 4.275 | 17.256 | -3.345 | 12.551 | -13.986 | 9.061 | 19.588 | 3.709 | 19.444 |
| -7.597 | 23.800 | 25.271 | 10.734 | 2.821 | -2.113 | -17.849 | 12.338 | -1.378 | -18.313 | 22.467 | 7.416 | 10.007 | 4.626 |
| -1.418 | 25.784 | -12.526 | -14.747 | 1.016 | 21.649 | 27.647 | -15.639 | -3.319 | -3.140 | 27.253 | 18.543 | 25.772 | 0.757 |
| -20.823 | -16.522 | -6.533 | 22.577 | 17.128 | 28.478 | 4.902 | -7.044 | 25.175 | 8.914 | -6.058 | 16.827 | 23.509 | 24.282 |
| 20.245 | 4.379 | -4.307 | -26.322 | 6.764 | 19.805 | 20.948 | -11.139 | 23.092 | 12.756 | 13.516 | 22.592 | 9.519 | 22.397 |
| 18.481 | -6.683 | 13.085 | 1.978 | -9.572 | 7.584 | 9.419 | 0.029 | 7.980 | 6.803 | 9.625 | 3.719 | -2.866 | 13.211 |
| 11.155 | 14.976 | 6.286 | 3.503 | 16.698 | 8.485 | 29.685 | 19.518 | 26.455 | -8.356 | 4.149 | 1.175 | -5.573 | -1.634 |
| -2.438 | 27.255 | 6.981 | 18.736 | 11.324 | 12.444 | -28.749 | 10.401 | 2.403 | -12.946 | 9.996 | -18.763 | -2.094 | -2.560 |
| 10.884 | 13.899 | -16.973 | 5.659 | 17.943 | 0.550 | 21.939 | 20.212 | 24.006 | 14.875 | 1.876 | 6.110 | 13.658 | 20.428 |
| 1.758 | 25.893 | 11.012 | 2.457 | -18.758 | -11.174 | -23.262 | 6.084 | 14.019 | 11.979 | 11.594 | 23.510 | -8.195 | -7.271 |
| -9.264 | 9.362 | -14.153 | 19.534 | -24.424 | 23.495 | 1.564 | -10.241 | 9.071 | 16.078 | 21.227 | 21.167 | 13.164 | 10.334 |
| -22.760 | -22.722 | 4.493 | 24.440 | -28.624 | 9.195 | -11.851 | -8.134 | 5.409 | 17.311 | -2.923 | 6.614 | 9.598 | 22.704 |
| -15.050 | 27.209 | -5.690 | 25.692 | -2.166 | -6.986 | 17.413 | 25.735 | -28.998 | 3.741 | 21.695 | 10.205 | 5.890 | -4.472 |
| 15.573 | 14.605 | 19.316 | 18.798 | -7.829 | -28.028 | -27.169 | -1.585 | 13.404 | 20.161 | 19.814 | 18.213 | -5.986 | 6.905 |
| -6.993 | -22.053 | 3.664 | 0.193 | 30.781 | 13.694 | 24.078 | 18.029 | 16.670 | 28.465 | 6.966 | 14.836 | 14.070 | 24.681 |
| -18.521 | 14.238 | -18.748 | -19.194 | 21.950 | 6.875 | 19.983 | 7.306 | 15.699 | 27.855 | 23.608 | -9.659 | -8.510 | 0.807 |
| 0.954 | 25.608 | 24.033 | -26.998 | -5.849 | -27.888 | -17.564 | 23.596 | -14.858 | -10.026 | 28.403 | 9.276 | 21.590 | 3.270 |
| -4.248 | 16.439 | 12.004 | -14.255 | -25.779 | 22.811 | -29.389 | -3.625 | 12.428 | 8.175 | 10.739 | 13.880 | 20.061 | 14.325 |
| -5.972 | -4.146 | 10.051 | 6.224 | 0.242 | -29.983 | -28.402 | 14.930 | 11.191 | 30.291 | -1.357 | 16.934 | 11.009 | 21.432 |
| -26.620 | -26.771 | 20.812 | 26.238 | 26.084 | -0.608 | 16.054 | 1.855 | 22.448 | 10.507 | 5.899 | 2.184 | 20.141 | 20.824 |
| 11.670 | 27.632 | 8.900 | -8.825 | 18.399 | 7.251 | 14.569 | -12.320 | 25.058 | 29.321 | 11.153 | 23.388 | 21.656 | 15.660 |
| -0.512 | -19.015 | -19.022 | 19.885 | -8.199 | -6.615 | -4.569 | -2.638 | 26.369 | -12.932 | 16.020 | 12.239 | 1.208 | -0.934 |
| 4.538 | -24.516 | 21.791 | 3.253 | 4.587 | 20.346 | 27.284 | -17.758 | 21.422 | 13.327 | 25.100 | -5.361 | 0.428 | -7.327 |

| | | | | | | | | | | | | | |
|---|---|---|---|---|---|---|---|---|---|---|---|---|---|
| 23.019 | 23.715 | -16.383 | 13.941 | -22.625 | 10.448 | -19.254 | -19.074 | 18.224 | 15.328 | 19.429 | 10.586 | 14.482 | 14.698 |
| -25.622 | -23.730 | -2.122 | 1.792 | 24.259 | -21.737 | 17.878 | -2.167 | 31.250 | 4.630 | 15.745 | 15.513 | 25.056 | 13.441 |
| 16.302 | -12.870 | 23.140 | 8.922 | 23.813 | -15.297 | 10.674 | 10.493 | 12.982 | 1.565 | 23.549 | 24.894 | 20.611 | 20.995 |
| -27.854 | -3.396 | -7.222 | 7.024 | 8.301 | 14.000 | -8.532 | -7.022 | -4.151 | 6.980 | 17.936 | 19.799 | 2.996 | 12.604 |
| 22.706 | -0.460 | 3.496 | 25.780 | -28.918 | -5.738 | 25.090 | -0.740 | 28.486 | 27.776 | 24.352 | 23.252 | 16.037 | 18.744 |
| 19.742 | 29.243 | 24.216 | -16.917 | -26.776 | -18.953 | -4.342 | -12.445 | 27.960 | 9.960 | 19.677 | 4.530 | 5.109 | 9.457 |
| -19.492 | -25.516 | -9.027 | -25.366 | 10.269 | -1.806 | -14.091 | 19.051 | 15.668 | 7.735 | -11.379 | 17.143 | -13.155 | 7.532 |
| 19.094 | 15.650 | 13.398 | -13.286 | -1.094 | -7.074 | 8.831 | -16.340 | 22.621 | 16.892 | -9.401 | 8.352 | 21.555 | -1.264 |
| 16.331 | 13.302 | -6.073 | -22.805 | 4.274 | 5.193 | 15.470 | -24.468 | 23.292 | 18.898 | 28.587 | 22.514 | -1.329 | 17.652 |
| 0.236 | 25.714 | -22.600 | -1.071 | 27.186 | 18.205 | 14.784 | 9.305 | 21.157 | 3.801 | -2.833 | 24.665 | 7.318 | 16.217 |
| 15.487 | 29.873 | -25.635 | -3.237 | 18.695 | 22.709 | -3.051 | 5.639 | 1.559 | -10.044 | 3.244 | 18.496 | -15.186 | -6.974 |
| 14.016 | 7.354 | -4.056 | -21.513 | -23.273 | -21.644 | 18.076 | 17.318 | 3.277 | -3.764 | 29.302 | 14.049 | 11.456 | 17.628 |
| -20.137 | 0.832 | -13.680 | 25.919 | -13.617 | 28.164 | 0.266 | 0.979 | 22.206 | -12.342 | 12.909 | 9.896 | 17.519 | 2.922 |
| -11.499 | 11.038 | -22.872 | 0.039 | -19.199 | 20.562 | -24.606 | -5.889 | 13.484 | 15.150 | 8.102 | 20.593 | 14.632 | -5.510 |
| 22.505 | 21.016 | 20.595 | -23.389 | 25.459 | 14.861 | 3.093 | -15.178 | 20.460 | -5.505 | 18.479 | 1.867 | 3.187 | -18.360 |
| 10.030 | -3.629 | 26.198 | 7.925 | -7.366 | -23.900 | 3.711 | 6.814 | -12.731 | 4.994 | 17.375 | -5.433 | -5.225 | 8.702 |
| -16.547 | -15.911 | -4.828 | -8.244 | -21.574 | 9.961 | 22.286 | -8.543 | 12.084 | -7.852 | 19.000 | 15.788 | 12.007 | -24.955 |
| 20.340 | -30.511 | -24.166 | 7.547 | 28.266 | 6.159 | -26.720 | -1.108 | 25.018 | 2.479 | -7.902 | -2.139 | 15.685 | 9.347 |
| 16.849 | -18.005 | 23.017 | -5.608 | 11.893 | 11.441 | 9.339 | 16.853 | 31.519 | 15.302 | -11.292 | 26.810 | 23.483 | 21.848 |
| 21.445 | 22.901 | 21.543 | -22.605 | -28.520 | -11.008 | 5.514 | -2.203 | -20.231 | -0.691 | 4.925 | 17.451 | 10.701 | 15.208 |
| 12.738 | 15.141 | 24.134 | 5.775 | 16.240 | 0.180 | 17.137 | 3.850 | 20.298 | 7.544 | 0.998 | 22.048 | 16.939 | 10.684 |
| -7.680 | -22.337 | 20.813 | -4.635 | 10.789 | 7.287 | 23.935 | -4.287 | -2.538 | 14.550 | 21.661 | 25.860 | 20.769 | 19.603 |
| -20.785 | 25.842 | -0.522 | 2.127 | -5.366 | -1.147 | 5.898 | 13.751 | 6.639 | -2.172 | 23.219 | 7.486 | 1.233 | 25.109 |
| -12.886 | -31.253 | 25.258 | -19.853 | -22.196 | -0.437 | 0.215 | 22.022 | 2.622 | -0.290 | 9.500 | 23.436 | 19.983 | 20.922 |
| 26.899 | 13.440 | -7.083 | 26.434 | -6.231 | -14.153 | 3.876 | 15.290 | 28.544 | -8.281 | 20.640 | 12.341 | 19.283 | 15.768 |
| 15.573 | 15.817 | 4.683 | -15.261 | -28.185 | 23.053 | 18.896 | -10.228 | -0.732 | 18.085 | 26.907 | 24.651 | 6.492 | 22.738 |
| -23.489 | 14.528 | 24.767 | 19.628 | 0.712 | -8.184 | 12.024 | 21.459 | 23.448 | 9.111 | 6.884 | 17.294 | 4.375 | 8.105 |
| 6.407 | 5.773 | 8.884 | -11.807 | 18.116 | -24.567 | -13.843 | -0.070 | -30.471 | -14.628 | 25.700 | 14.887 | 3.223 | -13.559 |
| -12.547 | -1.117 | -13.515 | -23.326 | -14.854 | 19.485 | -4.769 | -4.371 | 29.773 | -20.123 | 19.953 | 19.485 | -19.560 | 9.720 |
| -2.618 | 9.721 | 2.860 | -6.350 | 12.542 | -2.968 | 22.958 | 6.230 | 19.471 | -23.916 | 16.076 | -1.000 | 26.044 | -8.796 |
| 27.392 | -7.409 | -26.684 | 3.459 | 20.460 | -25.658 | -1.063 | 24.438 | 11.291 | 13.121 | 14.227 | 26.091 | 13.484 | -1.432 |
| 24.246 | 28.024 | 12.902 | -2.209 | -13.125 | 23.860 | 7.181 | 22.222 | 12.983 | -9.007 | 21.975 | 24.084 | 6.648 | 18.403 |
| 0.968 | 6.640 | 6.994 | -17.273 | -12.234 | 14.637 | 15.890 | 8.506 | 12.233 | 10.980 | 29.703 | -6.391 | 21.737 | -12.333 |
| 11.366 | -5.724 | -5.722 | -22.589 | 24.656 | -3.654 | 25.314 | -9.561 | 3.623 | 15.171 | 11.391 | 6.151 | -0.424 | 6.537 |
| -28.935 | 17.036 | -24.041 | 24.842 | 7.985 | 10.772 | -7.347 | 21.021 | -3.491 | -11.028 | 25.476 | 10.999 | -4.240 | 1.317 |
| 25.027 | -19.047 | -17.563 | 20.276 | 11.872 | 1.484 | 25.897 | -14.327 | 22.427 | 17.293 | 6.150 | 12.575 | 11.666 | -14.363 |

| | | | | | | | | | | | | | |
|---|---|---|---|---|---|---|---|---|---|---|---|---|---|
| 7.044 | -26.520 | 6.219 | -13.023 | -26.441 | 2.665 | 2.330 | 3.991 | 20.949 | -28.796 | -1.872 | 13.399 | 5.453 | 24.392 |
| 21.598 | -26.889 | 25.294 | 12.060 | -9.784 | -11.910 | 8.562 | -6.207 | -12.489 | 19.866 | -8.385 | 16.707 | 0.496 | 3.758 |
| -2.573 | 29.202 | -8.751 | -11.332 | -14.028 | 10.137 | -9.117 | 18.591 | 17.302 | 14.358 | 20.784 | 23.240 | 14.000 | 17.716 |
| 28.089 | -23.590 | -14.555 | 4.540 | -20.485 | -9.845 | 1.111 | 1.854 | 25.093 | 18.289 | 15.188 | 15.574 | 15.172 | 8.175 |
| -13.268 | 28.414 | -15.327 | -19.848 | 26.681 | -1.496 | 0.160 | -16.221 | 24.028 | 16.336 | 19.503 | -2.398 | 7.515 | 11.896 |
| 9.518 | 19.207 | 27.168 | 1.665 | -17.414 | 11.418 | -11.907 | 4.596 | 12.900 | -2.956 | 26.522 | 1.648 | 18.153 | 18.921 |
| 16.364 | 25.947 | -6.485 | -27.497 | -19.015 | 9.004 | -20.554 | -4.479 | 7.666 | 25.447 | 17.457 | 21.208 | 23.950 | 23.288 |
| -9.879 | 24.784 | 18.423 | 16.948 | 19.084 | 15.230 | -22.476 | -3.016 | 10.934 | 16.818 | 23.503 | 26.845 | 4.570 | 21.029 |
| 4.240 | -24.979 | -25.227 | 18.036 | 25.445 | 0.367 | 5.839 | 17.840 | 5.709 | -9.767 | 24.445 | 25.811 | 21.218 | 21.925 |
| 1.532 | 1.725 | 19.863 | -3.537 | 23.457 | 13.235 | 15.553 | 6.936 | 26.433 | 6.363 | 16.662 | 10.962 | -4.754 | 14.352 |
| -24.273 | 17.139 | 4.397 | 27.101 | 27.336 | -14.558 | -9.331 | 10.720 | 14.521 | -2.223 | 19.825 | 18.017 | 8.791 | 16.342 |
| -7.755 | -17.505 | 26.726 | 21.624 | 12.952 | 6.800 | 4.063 | -12.257 | -4.019 | 22.369 | 16.189 | 19.248 | 11.316 | 14.857 |
| -4.275 | 6.493 | 14.969 | 22.760 | 10.202 | -4.427 | 10.846 | 6.017 | 30.539 | 7.403 | 10.543 | 23.806 | 16.787 | 8.153 |
| -20.852 | -29.585 | -7.004 | 13.361 | 26.875 | -21.670 | -11.289 | 7.545 | -16.572 | 11.640 | -3.560 | 22.978 | 15.959 | 5.383 |
| 22.024 | -25.682 | 22.066 | -20.458 | -18.852 | 4.202 | 17.280 | -8.981 | 0.399 | 13.835 | -6.278 | 21.311 | 5.926 | 1.271 |
| 11.298 | -28.784 | -11.558 | -14.408 | 16.614 | 7.910 | -21.329 | 22.975 | 18.481 | 24.576 | 17.461 | 27.629 | 2.043 | 7.333 |
| 1.355 | 24.874 | 18.785 | 9.947 | -23.954 | -6.675 | 14.896 | 14.784 | 28.689 | 10.951 | 23.018 | 24.961 | 14.456 | -22.324 |
| -19.409 | 27.796 | -18.706 | -24.094 | 20.440 | -14.586 | 5.583 | 0.665 | 29.539 | 13.324 | 19.983 | 13.444 | 23.819 | 11.029 |
| 19.678 | 16.303 | 16.034 | -20.851 | -0.838 | -24.277 | -2.305 | -6.934 | 15.651 | -3.844 | 5.274 | 3.639 | 12.786 | 23.589 |
| 14.426 | -20.333 | -16.466 | 15.863 | 15.336 | 23.023 | 18.232 | -0.408 | 14.974 | -9.454 | -2.475 | 12.776 | 10.748 | 23.192 |
| 13.478 | 25.546 | 19.509 | 3.902 | -22.965 | 17.588 | 14.141 | 12.537 | -12.322 | -15.276 | 19.468 | 0.815 | 25.148 | 13.304 |
| 4.931 | -14.845 | -21.946 | 28.878 | 22.134 | -9.752 | 1.824 | 6.642 | -20.202 | 2.118 | 18.298 | 24.566 | 7.907 | 15.663 |
| 28.273 | -10.828 | 6.559 | 5.909 | -1.834 | 13.268 | -1.952 | -15.148 | 30.016 | 21.206 | 7.067 | 16.994 | 20.451 | 20.266 |
| 28.760 | 28.183 | 2.438 | -22.493 | -3.512 | 15.484 | -18.947 | 8.247 | 11.349 | 17.780 | 13.644 | 6.535 | 10.055 | 18.999 |
| -12.485 | 1.055 | 4.273 | 23.062 | 19.667 | 4.610 | 9.992 | 2.385 | -20.655 | 20.148 | 24.972 | 12.322 | 13.403 | 10.457 |
| -13.775 | 21.256 | -17.959 | -28.758 | -17.555 | 8.393 | 7.493 | 10.223 | 20.775 | 16.494 | 2.407 | 18.117 | 23.110 | 21.751 |
| -30.721 | 13.890 | 18.018 | -14.255 | 12.228 | -2.908 | -9.180 | 9.201 | -7.487 | 9.562 | -10.514 | 15.664 | -5.864 | 17.966 |
| -23.252 | 24.049 | -3.697 | 14.665 | 25.722 | -12.212 | 17.196 | -5.759 | 19.686 | -2.506 | 22.244 | 26.587 | 1.872 | 26.548 |
| 5.567 | 12.319 | -10.738 | -29.413 | -7.737 | -9.396 | 8.904 | 6.575 | 13.079 | 21.968 | 20.385 | 21.496 | -3.303 | 12.002 |
| -11.247 | 15.168 | -22.312 | 20.170 | -19.983 | -7.398 | -18.336 | 12.583 | -26.110 | 2.889 | 28.826 | 19.905 | 17.843 | 2.645 |
| -28.185 | -8.940 | -26.387 | -25.051 | -21.097 | -1.212 | 14.817 | 4.384 | 22.065 | -4.476 | 26.632 | -4.291 | 15.765 | 22.174 |
| 1.607 | -25.026 | 10.798 | 20.992 | -9.229 | -3.632 | 19.351 | 24.479 | -23.526 | 15.003 | 25.282 | 27.642 | -0.165 | 7.441 |
| 7.254 | 11.946 | -28.884 | -20.260 | 6.106 | -15.483 | 10.186 | 22.705 | 27.770 | 4.238 | -7.448 | 14.878 | 14.220 | 23.641 |
| -24.153 | 9.222 | 14.501 | 26.036 | -5.067 | -7.153 | -19.755 | -5.780 | 20.045 | 6.937 | 5.136 | 17.919 | 2.263 | 14.577 |
| -31.482 | 9.789 | -8.366 | 14.804 | 23.755 | -24.074 | -9.158 | -7.495 | 29.676 | -23.068 | 9.751 | 0.853 | 28.189 | 18.338 |
| 9.751 | -1.738 | -8.950 | -16.080 | -24.594 | -9.563 | -19.010 | -11.110 | -26.797 | 19.440 | 12.010 | -1.961 | 13.192 | 21.050 |

| | | | | | | | | | | | | | |
|---|---|---|---|---|---|---|---|---|---|---|---|---|---|
| -24.790 | 21.821 | -0.839 | 21.033 | 14.053 | -18.747 | -1.481 | 10.912 | 4.539 | 4.635 | 6.907 | 21.714 | 7.738 | 19.220 |
| -22.888 | -28.062 | 14.259 | 29.105 | 17.700 | -10.691 | 10.354 | -1.126 | 23.012 | 27.573 | -9.494 | 10.114 | 19.458 | 17.207 |
| 3.880 | 20.022 | 21.518 | -4.443 | -12.179 | -23.023 | -11.167 | 25.787 | 2.240 | -1.932 | 17.029 | 10.871 | 23.999 | 0.836 |
| -0.070 | -21.682 | 27.333 | -16.566 | 2.750 | -25.156 | 1.467 | 5.075 | 1.096 | 12.742 | 19.765 | 24.730 | 21.647 | -16.925 |
| -20.567 | 17.739 | -13.208 | -12.611 | 24.636 | -9.980 | -20.315 | 15.784 | 23.654 | 1.192 | 10.394 | 23.630 | 11.723 | 26.051 |
| 8.439 | -6.433 | 28.066 | -27.504 | 13.703 | -21.983 | -8.852 | -15.813 | -12.008 | -15.603 | 19.125 | 13.288 | 8.467 | -24.441 |
| 0.408 | -15.337 | -7.945 | -10.255 | -7.925 | -16.829 | 23.511 | 5.470 | 0.714 | 7.411 | 13.611 | 8.869 | 0.504 | -2.066 |
| -22.361 | -22.387 | 4.778 | 16.019 | 10.975 | -2.507 | 6.621 | 22.924 | 11.737 | -4.355 | 17.900 | 5.771 | 24.666 | -23.441 |
| -11.161 | 25.075 | 8.602 | -26.438 | 17.399 | -1.988 | 4.058 | -6.201 | -5.730 | 4.025 | 22.539 | 15.197 | 16.430 | 24.322 |
| -27.894 | 15.435 | 7.330 | 15.051 | 23.566 | 5.322 | -9.566 | 0.742 | -7.127 | 13.530 | -16.291 | 19.502 | 16.895 | -8.643 |
| 6.451 | 11.419 | 23.907 | 12.092 | -3.981 | 11.043 | 9.025 | 10.256 | 2.215 | 10.655 | 23.092 | -0.461 | 15.885 | 3.318 |
| 24.343 | 23.037 | -25.379 | 14.344 | 26.188 | 8.514 | -6.366 | 26.969 | 17.641 | 14.973 | 27.288 | 18.535 | 17.966 | 8.184 |
| 13.958 | 27.678 | 15.785 | 18.053 | 5.395 | 17.039 | 7.815 | -24.409 | 22.891 | -4.919 | 24.207 | 15.828 | 1.883 | 19.695 |
| 20.297 | -4.364 | 20.017 | -9.215 | 9.351 | -11.498 | -12.542 | 7.528 | 5.838 | -10.514 | 10.860 | 3.692 | 18.675 | 17.539 |
| 16.913 | -18.502 | -19.202 | 0.023 | -21.663 | -6.568 | -24.691 | 19.327 | 28.054 | -23.043 | -30.510 | 15.241 | 10.354 | 21.838 |
| -23.202 | 24.228 | 25.382 | 7.584 | -25.285 | 14.633 | 16.739 | -14.529 | 10.354 | 22.389 | -18.644 | 30.661 | 26.568 | -8.110 |
| -1.710 | -20.250 | 2.444 | -22.387 | 25.756 | -23.356 | -14.870 | -5.161 | -2.312 | 4.577 | 8.745 | -0.885 | 6.082 | 27.744 |
| 23.822 | 14.859 | 28.369 | -14.931 | -12.961 | -6.649 | -5.567 | 2.689 | 6.074 | 19.478 | 10.268 | 24.150 | 26.174 | 10.289 |
| 22.425 | -25.588 | 10.505 | 11.058 | -16.399 | -12.965 | -0.151 | 8.132 | -20.469 | 25.286 | 20.482 | 21.509 | 13.423 | 19.414 |
| -21.388 | 4.673 | -18.016 | -11.166 | 16.300 | -17.273 | 11.487 | -14.176 | 11.488 | 27.734 | 25.341 | 7.152 | 19.831 | 14.652 |
| 9.875 | 17.679 | 19.407 | 13.623 | -18.299 | -23.890 | 8.161 | 5.829 | 21.395 | -0.114 | 21.946 | 15.970 | 18.904 | 9.613 |
| 23.781 | 28.366 | 1.626 | 1.846 | -3.635 | -1.522 | -11.371 | -1.352 | 3.163 | 21.808 | 7.445 | 8.740 | 22.435 | 12.767 |
| 27.397 | -28.430 | 22.684 | 25.384 | -7.198 | 15.385 | -20.137 | -7.244 | 13.591 | 3.113 | 14.530 | 26.431 | 20.560 | 14.359 |
| 27.911 | -20.223 | 17.511 | 5.033 | 16.331 | -4.608 | 0.143 | 25.137 | -12.885 | -8.725 | 21.253 | 15.592 | 5.219 | 8.613 |
| 21.756 | -10.522 | -1.700 | -24.007 | -10.408 | -17.744 | 23.121 | 17.147 | 4.993 | 17.369 | 11.991 | 29.980 | 11.068 | 12.139 |
| 8.358 | -26.288 | -16.906 | 8.779 | -23.991 | -12.720 | -18.071 | -18.420 | 28.671 | 16.373 | 16.057 | 20.715 | 14.810 | 28.825 |
| 15.937 | 13.818 | -22.961 | 22.992 | 23.067 | -8.998 | 14.087 | 27.526 | 14.862 | 14.735 | 20.770 | 15.205 | 17.497 | 17.233 |
| -7.128 | -12.139 | -29.730 | -16.569 | -1.248 | -16.049 | -6.330 | 22.432 | 19.809 | 22.142 | 18.300 | 19.433 | 2.842 | 22.641 |
| -9.605 | 25.929 | -14.019 | 30.763 | -11.385 | -13.856 | 21.629 | 20.067 | 15.939 | 18.152 | 4.887 | 5.877 | 13.820 | 21.606 |
| -8.205 | 24.758 | 23.328 | 2.532 | 12.914 | -1.041 | -4.475 | 15.781 | 3.627 | -17.078 | 22.776 | 4.183 | 18.239 | 22.313 |
| 10.689 | 18.700 | 6.861 | -8.197 | 7.626 | -9.294 | -20.972 | -5.360 | 6.509 | 13.619 | -14.760 | 24.273 | 25.480 | -29.372 |
| -4.574 | 2.500 | 6.496 | -22.036 | -10.326 | -10.685 | 24.483 | 23.593 | 16.720 | 12.020 | 29.460 | -0.137 | 6.171 | 9.530 |
| -27.300 | 0.537 | -29.255 | 1.007 | -23.368 | -16.947 | -12.973 | -4.979 | 1.438 | 9.468 | -3.255 | 13.866 | 23.175 | 24.038 |
| 9.182 | -6.918 | -26.978 | 7.530 | 14.586 | 1.241 | 20.372 | 7.078 | -25.535 | 4.202 | 15.390 | 14.825 | -28.266 | 22.992 |
| 2.565 | 0.032 | -10.529 | -7.884 | -1.534 | 20.817 | 6.426 | -11.822 | 22.815 | -7.220 | 6.717 | 19.017 | 3.297 | 18.117 |
| -6.347 | 16.015 | -7.410 | -8.422 | -2.409 | 0.433 | 25.687 | 11.126 | -3.486 | 18.796 | 10.457 | -2.451 | 17.857 | 20.035 |

| | | | | | | | | | | | | | |
|---|---|---|---|---|---|---|---|---|---|---|---|---|---|
| -4.919 | 19.095 | 26.931 | 24.251 | 19.959 | -10.162 | -25.416 | 5.843 | 18.635 | 10.062 | 21.376 | 21.578 | 25.094 | 8.573 |
| 18.754 | 23.161 | -11.968 | 19.517 | 11.296 | -8.130 | 5.442 | -16.248 | 10.611 | 7.313 | 17.886 | 9.462 | 22.151 | -10.591 |
| 24.536 | -13.066 | 27.818 | 12.884 | -7.281 | 9.281 | 1.464 | -4.668 | 22.050 | 6.352 | 12.784 | 13.453 | 16.687 | 3.927 |
| 16.649 | -23.937 | -8.031 | 9.601 | 9.863 | -7.771 | -22.554 | 18.770 | 14.449 | 27.633 | 28.383 | 5.927 | 27.599 | -6.674 |
| -0.196 | -12.445 | -5.027 | 3.232 | -8.841 | -1.538 | 23.517 | 16.438 | 23.607 | -3.323 | -11.488 | 28.835 | 13.101 | 16.921 |
| -3.978 | -22.811 | -2.311 | 22.829 | 19.477 | 0.622 | 16.295 | -7.078 | 27.181 | 8.949 | -0.441 | 19.347 | 20.819 | 15.244 |
| -5.619 | 21.452 | -13.205 | 25.127 | 18.903 | -8.352 | 14.729 | 3.167 | 4.660 | 20.160 | 11.986 | 19.889 | 5.088 | -2.242 |
| 23.960 | -8.516 | 30.178 | -21.422 | 11.015 | -7.911 | 4.104 | -9.246 | -11.859 | 11.743 | -3.334 | 7.393 | 8.767 | -3.539 |
| 15.584 | 11.894 | 11.923 | 19.736 | 1.679 | 1.558 | 27.025 | 0.623 | 24.617 | 4.665 | 23.747 | 22.410 | -3.527 | 7.022 |
| 12.526 | -11.660 | -9.303 | 16.891 | 17.477 | -20.055 | -1.747 | 27.177 | -5.649 | -10.043 | -24.541 | 3.830 | 19.409 | -1.653 |
| -11.974 | 20.877 | 21.118 | -0.315 | 0.664 | 8.757 | 1.051 | 11.815 | 25.185 | 5.091 | -2.048 | 14.203 | 19.917 | 19.142 |
| -24.817 | -18.863 | 17.666 | 8.866 | 0.150 | -1.129 | -27.137 | 8.681 | 5.357 | 22.695 | 18.521 | 27.075 | 3.955 | 14.267 |
| 9.441 | 22.027 | -27.283 | -9.536 | 20.537 | -17.698 | 25.252 | 5.028 | 23.044 | 14.435 | 3.312 | 26.015 | 21.261 | 15.569 |
| -23.914 | 13.391 | 4.907 | -18.985 | -5.924 | 0.880 | -18.339 | 2.462 | -26.014 | 25.877 | 21.443 | 21.153 | 20.639 | 0.185 |
| -8.700 | 23.096 | -15.860 | -18.109 | -17.392 | 0.188 | 8.875 | 27.791 | 10.277 | 21.204 | 15.698 | 18.626 | 22.130 | 27.464 |
| -2.713 | 8.408 | -11.893 | 11.218 | 3.789 | 18.076 | 17.751 | 23.995 | 4.455 | -6.647 | 0.032 | 29.552 | 14.190 | 24.302 |
| -19.347 | -17.016 | 20.223 | -26.471 | 19.097 | -7.148 | 6.414 | -8.225 | 18.018 | -15.322 | 4.511 | 23.881 | 13.658 | 13.975 |
| 25.558 | -25.325 | -26.570 | 13.045 | 4.436 | 25.455 | 7.629 | 26.633 | 15.572 | 8.190 | 20.288 | 5.629 | -4.675 | 16.772 |
| 1.807 | 15.471 | -31.132 | -5.982 | 8.157 | 5.120 | -22.316 | -9.393 | 0.909 | 11.948 | 18.455 | 21.423 | 16.269 | 20.081 |
| -5.902 | 10.953 | 30.847 | 3.628 | -7.456 | -15.179 | -1.167 | -0.330 | -0.925 | -4.180 | 6.187 | 19.285 | 24.017 | 24.779 |
| 2.591 | 9.850 | -4.501 | -19.392 | -19.557 | -17.724 | 26.815 | 22.618 | 5.724 | 3.353 | 24.956 | 26.444 | 3.673 | 21.611 |
| 18.712 | -1.443 | -29.174 | 7.109 | -8.981 | 1.236 | -15.080 | -11.258 | 1.748 | -10.245 | 12.621 | 30.728 | 8.313 | 9.710 |
| 13.137 | 5.226 | 8.880 | 5.986 | 1.695 | -4.792 | -24.434 | 11.481 | 19.775 | 14.925 | 25.399 | 15.847 | 5.965 | 10.560 |
| -0.734 | -16.320 | 8.247 | -19.722 | 15.447 | -19.927 | 19.279 | 7.785 | -9.390 | -11.770 | 25.307 | 14.251 | 17.635 | 12.826 |
| -3.797 | -6.519 | -10.160 | 17.963 | 12.416 | 10.037 | -10.835 | 3.689 | 10.001 | -15.137 | 24.116 | 10.490 | 23.652 | -3.344 |
| 7.922 | -8.886 | 28.474 | -18.167 | -5.857 | 9.389 | -22.492 | 2.570 | 8.784 | 0.003 | 7.687 | 27.807 | 13.339 | 5.514 |
| -21.689 | -14.608 | 14.283 | 29.643 | 25.641 | -6.107 | 1.237 | 15.621 | 24.702 | 5.531 | 14.410 | 0.469 | -3.920 | -5.920 |
| -25.285 | -11.159 | 22.865 | -29.936 | 13.938 | -18.364 | -8.959 | 17.180 | -10.664 | 21.949 | 21.879 | 7.900 | 25.188 | 28.739 |
| 7.056 | 25.566 | 21.813 | 28.173 | -2.290 | 26.316 | 1.912 | 16.357 | 11.337 | 0.630 | -4.922 | 5.049 | 1.940 | 22.950 |
| 9.323 | -23.233 | 3.166 | -3.617 | 19.550 | -9.470 | 18.716 | 6.504 | 12.571 | 14.058 | 15.870 | 29.182 | 24.946 | 11.179 |
| -1.437 | 7.349 | -15.722 | 16.548 | -9.077 | 19.850 | 10.709 | 4.092 | 13.874 | -1.186 | 23.427 | 17.319 | 21.679 | -0.376 |
| 10.206 | 6.307 | -20.810 | 6.295 | 1.066 | -12.681 | 9.590 | 19.843 | -16.727 | -8.133 | 21.017 | 2.252 | 16.668 | 29.862 |
| 5.758 | -20.636 | 31.830 | -20.310 | -6.623 | 7.637 | 12.371 | -7.670 | 22.213 | 8.791 | 20.575 | 22.616 | 19.301 | 19.337 |
| 15.000 | -6.488 | 26.090 | 8.886 | 27.665 | -6.066 | -7.518 | 23.242 | 18.914 | 1.375 | 27.378 | 24.220 | 7.454 | 21.349 |
| 3.267 | 25.004 | 24.980 | -2.031 | 8.630 | -0.334 | -5.526 | -10.622 | 24.022 | -9.060 | 17.333 | 25.034 | 20.612 | 15.896 |
| 7.309 | -25.765 | -17.886 | -8.365 | 25.246 | -7.311 | -29.071 | -1.005 | 21.383 | 24.893 | 3.776 | 30.073 | 26.677 | 22.396 |

| | | | | | | | | | | | | | |
|---|---|---|---|---|---|---|---|---|---|---|---|---|---|
| -5.880 | 17.684 | -30.412 | 13.177 | 20.653 | 11.075 | -11.335 | -6.873 | -4.995 | 15.903 | 12.621 | 19.203 | 24.167 | 28.004 |
| -1.630 | 4.590 | 12.497 | 19.767 | -0.134 | -24.753 | 21.489 | 26.055 | 7.996 | -6.251 | 8.681 | 26.511 | -5.833 | 4.952 |
| 15.349 | 11.148 | -5.048 | 17.448 | 14.390 | -15.166 | 12.647 | 21.699 | 1.975 | 3.335 | 25.914 | 21.651 | 30.133 | 14.043 |
| 19.204 | 23.598 | 14.865 | -12.497 | -0.520 | 12.057 | -2.829 | -3.951 | 5.438 | 2.871 | 24.352 | 24.531 | 17.530 | 21.976 |
| -19.382 | 26.085 | -9.438 | -1.496 | 26.718 | 24.927 | 15.255 | -11.797 | -21.161 | -12.248 | 19.868 | 26.536 | -1.060 | 10.876 |
| 11.428 | -10.057 | -24.825 | 22.687 | -2.226 | -1.260 | 23.457 | -19.357 | 27.602 | -13.038 | 17.408 | 16.730 | 14.606 | -1.040 |
| | -3.696 | 10.741 | -26.288 | -14.646 | 6.260 | -21.100 | 28.004 | 25.782 | 16.275 | 27.966 | 1.716 | 22.091 | 9.820 |
| | -1.319 | -18.902 | -14.412 | -11.235 | 9.975 | -17.553 | 1.998 | -1.436 | -5.323 | -3.904 | 15.177 | -2.183 | 29.430 |
| | -8.928 | -26.786 | 24.694 | 26.209 | -8.490 | -15.691 | -4.634 | 25.037 | 6.217 | 21.127 | 3.960 | 17.970 | 28.731 |
| | 2.611 | 23.761 | 29.025 | 12.889 | 18.782 | 25.227 | -13.814 | 19.813 | 19.344 | 25.899 | 22.696 | 0.815 | 15.628 |
| | -15.624 | 24.329 | 10.589 | 22.894 | 19.576 | 9.917 | 23.307 | 26.526 | -11.932 | 15.534 | 11.101 | 18.868 | 17.040 |
| | -18.356 | 27.289 | -18.775 | 10.664 | -22.804 | 1.595 | 11.095 | 26.763 | 11.539 | -4.060 | 29.113 | 8.768 | 14.885 |
| | -2.944 | 26.391 | 23.376 | 5.302 | -12.623 | -5.896 | 8.796 | 14.204 | 18.182 | 27.354 | 18.220 | -2.094 | 26.807 |
| | -5.435 | 20.563 | -3.971 | -14.021 | -18.372 | 27.709 | 14.885 | 12.145 | 21.208 | 3.620 | 7.822 | 22.656 | 23.864 |
| | 17.476 | 5.145 | 19.564 | 9.301 | 0.259 | 13.458 | -5.932 | 10.788 | -2.892 | 7.921 | 3.197 | -26.068 | 18.495 |
| | 7.082 | 22.272 | 15.718 | 1.471 | -17.015 | -4.682 | -18.657 | 6.557 | 23.257 | 1.368 | 4.520 | 17.104 | 23.393 |
| | -6.205 | -5.942 | 6.297 | -5.201 | -26.502 | 8.606 | 7.593 | 24.532 | -7.668 | 19.961 | 24.921 | 10.760 | 20.044 |
| | -9.923 | -28.497 | 11.697 | 2.286 | 12.816 | 20.825 | 22.917 | 1.833 | 2.465 | 5.893 | 17.971 | 7.905 | 0.677 |
| | -0.129 | 19.194 | -8.773 | 6.456 | 16.528 | -21.975 | 6.388 | 8.601 | 10.885 | 26.406 | 26.398 | 6.186 | 22.141 |
| | 15.852 | 8.659 | 3.578 | -13.526 | -5.392 | -11.257 | 5.362 | 4.156 | -18.862 | 25.977 | 24.027 | 2.642 | 13.876 |
| | 5.086 | -22.815 | -25.097 | 27.038 | -13.845 | -0.904 | 22.340 | 26.015 | 12.570 | 18.119 | 7.930 | 4.005 | 14.395 |
| | -0.827 | -27.833 | -21.307 | -20.332 | 17.575 | -28.819 | 10.910 | 15.440 | -11.776 | 18.898 | 19.264 | -0.054 | 8.485 |
| | 6.277 | -7.280 | -25.540 | 24.468 | -18.983 | 16.035 | -10.196 | -4.730 | -17.169 | 21.957 | -2.615 | 24.725 | 28.153 |
| | 16.244 | 22.818 | -7.264 | -22.986 | 14.115 | 3.997 | 18.353 | 16.538 | 25.717 | 11.772 | 20.698 | 11.583 | 6.147 |
| | 19.895 | 30.110 | 11.504 | 8.597 | 12.045 | 26.791 | 19.304 | 23.448 | 12.429 | 16.842 | 0.943 | 25.514 | 28.961 |
| | 16.570 | 11.347 | -20.568 | -0.820 | 22.679 | 13.141 | 20.012 | 1.106 | -4.404 | 12.831 | 6.947 | 18.474 | 24.241 |
| | -0.133 | 7.284 | 20.584 | 25.113 | 13.053 | 12.494 | -14.484 | 8.938 | 23.793 | 8.423 | 14.306 | 10.343 | 21.888 |
| | 9.558 | 6.527 | -19.153 | 23.891 | 24.605 | 17.393 | -2.393 | 22.600 | 11.739 | 15.442 | -3.064 | 22.601 | 22.415 |
| | 3.110 | -24.000 | 12.940 | 12.243 | 26.607 | -11.316 | -0.099 | -19.590 | 0.007 | 25.041 | 15.768 | 28.224 | 18.699 |
| | 9.772 | 16.618 | -11.541 | -11.220 | -6.859 | 4.823 | -8.778 | 24.391 | 23.256 | 12.211 | -0.425 | 17.604 | -9.259 |
| | 1.792 | -15.275 | -6.825 | 20.562 | -27.689 | 0.657 | 21.167 | 2.444 | 23.178 | 5.329 | 17.085 | 14.827 | 30.280 |
| | 20.266 | 13.599 | -2.285 | 10.305 | -13.547 | -24.725 | 28.554 | 7.267 | 21.110 | 26.481 | 16.126 | 24.724 | 29.885 |
| | 24.832 | 11.835 | 5.329 | 9.118 | -13.050 | 19.955 | -9.547 | 19.885 | 13.618 | 18.619 | 4.829 | 29.459 | 23.894 |
| | -15.698 | -20.915 | 4.136 | 2.204 | 29.423 | -9.219 | 8.727 | -1.484 | 19.527 | 23.143 | 27.505 | 25.990 | 25.742 |
| | -15.073 | 27.340 | -23.146 | 11.434 | 28.107 | 7.676 | 27.650 | 26.960 | -4.534 | 16.091 | 14.752 | 19.051 | 20.724 |
| | 24.181 | 3.852 | 24.643 | -8.643 | 19.771 | -6.058 | 18.461 | 11.312 | 18.582 | 22.225 | 20.586 | 8.433 | -6.592 |

| | | | | | | | | | | | | | |
|---|---|---|---|---|---|---|---|---|---|---|---|---|---|
| | 1.848 | 4.528 | -25.906 | -22.909 | 7.682 | -17.266 | -7.911 | -9.969 | 17.456 | 2.430 | 28.216 | 11.228 | 27.223 |
| | 21.350 | 29.816 | 13.765 | -22.369 | 1.011 | -2.811 | 22.970 | 25.614 | 4.479 | 8.266 | -0.913 | 13.544 | 18.940 |
| | 13.776 | 23.413 | -27.228 | 25.719 | -11.970 | 14.417 | 6.402 | 18.789 | 1.947 | -12.691 | 9.550 | 17.470 | 0.971 |
| | 23.103 | -29.319 | 16.057 | -0.427 | -21.029 | 18.497 | 5.826 | 18.014 | 15.612 | -4.957 | 6.969 | 7.253 | 5.228 |
| | 1.475 | -10.446 | -14.466 | -5.829 | 28.800 | 5.491 | 23.941 | 10.195 | -3.199 | 14.990 | 24.103 | 14.041 | 20.983 |
| | 11.967 | 16.671 | -10.960 | -18.645 | -10.906 | -18.797 | 15.058 | 16.554 | 14.027 | 9.906 | 19.466 | 19.822 | 17.868 |
| | -20.829 | 14.556 | 21.190 | 9.571 | -26.865 | 23.122 | 2.133 | -4.886 | -10.604 | 22.173 | 10.733 | 14.429 | 28.586 |
| | -9.012 | 22.591 | 18.905 | 5.745 | -19.774 | 25.230 | 11.193 | 14.353 | 8.871 | -15.702 | 4.291 | 26.580 | 6.161 |
| | 22.625 | -18.034 | 12.506 | -1.753 | -8.728 | 29.081 | -3.548 | 16.008 | 1.586 | 14.578 | 12.442 | 20.985 | 11.748 |
| | 8.787 | 1.312 | 0.680 | 10.751 | 27.098 | -11.492 | -19.527 | 1.263 | 12.967 | 14.259 | 2.302 | 12.785 | 23.704 |
| | -10.787 | -12.312 | -10.057 | 0.009 | -22.858 | -23.482 | 8.899 | -6.737 | 9.486 | 19.302 | 16.245 | 2.983 | 29.340 |
| | -5.322 | -7.140 | -26.761 | 3.134 | 2.650 | 2.163 | 0.939 | 13.639 | 15.777 | 15.868 | 21.776 | 29.875 | 24.883 |
| | -19.026 | 18.992 | 7.557 | 19.135 | -2.176 | -16.157 | 7.609 | 11.559 | 19.884 | -22.666 | 1.010 | 24.366 | 16.997 |
| | 10.632 | 5.723 | -7.483 | -3.491 | -6.489 | -0.943 | -14.788 | 19.274 | 19.477 | -2.389 | 10.492 | 28.197 | 23.702 |
| | 21.017 | -22.389 | -16.704 | 26.977 | 14.230 | 22.487 | 17.150 | -6.283 | -0.129 | 17.316 | 22.849 | 13.585 | 14.909 |
| | -24.838 | 17.252 | 18.589 | 9.437 | 26.172 | 29.687 | -6.425 | 5.013 | -1.180 | 8.359 | 21.447 | 21.810 | 8.284 |
| | 8.714 | -8.324 | 10.305 | -5.333 | -28.295 | 21.171 | 23.410 | 20.723 | 14.279 | 18.391 | 7.820 | 7.086 | 18.677 |
| | -24.123 | -27.871 | -23.324 | 24.737 | -6.220 | 13.976 | -5.308 | 3.242 | 13.569 | 2.741 | 16.574 | 15.584 | -0.503 |
| | 0.192 | 25.575 | 17.449 | 23.531 | 3.459 | -8.026 | 25.535 | 23.803 | -9.166 | 8.919 | 15.609 | 13.956 | 22.107 |
| | -19.889 | 22.249 | -17.105 | 16.931 | 1.671 | 8.941 | 16.745 | 19.913 | 20.747 | 0.123 | 11.401 | 6.055 | -16.251 |
| | 5.815 | -11.440 | -8.803 | -6.400 | -23.568 | 27.906 | 21.159 | 15.391 | -5.131 | -4.199 | 19.957 | 23.459 | 26.229 |
| | -23.587 | -10.405 | -18.220 | -0.676 | -17.024 | 27.431 | -1.269 | -2.800 | 0.602 | 5.168 | 23.097 | 14.707 | 7.593 |
| | 7.470 | 7.316 | -17.499 | 22.110 | -20.486 | 24.079 | 22.527 | 11.037 | 9.843 | -3.716 | 25.006 | 17.906 | 7.142 |
| | -14.846 | 3.837 | -6.569 | 10.142 | 8.653 | 25.795 | -11.359 | 4.052 | 6.423 | 10.197 | 14.605 | 28.628 | -6.057 |
| | 18.030 | 17.697 | 7.448 | -7.896 | 1.427 | -13.239 | -5.914 | 24.460 | 22.243 | 17.944 | 3.833 | 18.494 | 29.477 |
| | 13.324 | -17.831 | 12.263 | 18.416 | 19.467 | 16.022 | 14.654 | 2.367 | -2.978 | 9.214 | 13.542 | 10.989 | 20.566 |
| | 21.557 | 19.338 | 2.018 | -12.640 | -10.470 | -20.858 | -4.676 | 16.713 | 24.248 | 25.054 | 24.157 | 19.034 | 28.292 |
| | 14.821 | 26.371 | 8.278 | -4.308 | 11.007 | -23.063 | 2.564 | 8.177 | 14.000 | 20.262 | 22.464 | 24.235 | 25.612 |
| | -17.537 | -22.489 | -24.485 | -5.011 | -3.029 | -6.221 | 24.935 | -3.217 | 21.522 | 24.300 | 25.484 | 5.367 | 24.180 |
| | -5.104 | 1.330 | -25.067 | 21.596 | 20.410 | 15.622 | 18.994 | -8.211 | 4.738 | 11.570 | -10.373 | 21.823 | 18.897 |
| | -9.965 | -5.963 | -2.287 | 19.508 | -4.589 | 0.347 | 6.964 | 24.488 | 25.553 | 3.223 | 27.204 | 25.636 | 13.843 |
| | -23.865 | 14.530 | -18.430 | 8.883 | 29.753 | -19.044 | 14.029 | 23.156 | 18.043 | 18.906 | 18.958 | 21.020 | 22.998 |
| | 6.820 | -26.580 | -1.934 | -5.286 | 26.976 | 19.784 | 27.391 | 11.776 | 12.984 | 19.953 | 14.917 | 6.681 | 12.226 |
| | 1.737 | -8.124 | 8.989 | -9.397 | 4.069 | 16.606 | 9.584 | 0.397 | 9.599 | 16.335 | 9.479 | -8.934 | 26.496 |
| | -15.612 | 5.868 | -20.435 | -9.010 | 23.683 | -26.158 | 6.156 | 3.787 | 20.056 | 5.041 | 22.543 | 26.700 | 7.075 |
| | -19.352 | 23.162 | 0.913 | 17.582 | -5.597 | 27.079 | -0.871 | 21.861 | 24.963 | 19.384 | 5.862 | 9.322 | 21.409 |

| | | | | | | | | | | | | | |
|---|---|---|---|---|---|---|---|---|---|---|---|---|---|
| | 23.801 | 13.838 | 19.879 | 9.928 | -10.038 | 20.132 | -6.818 | 1.779 | 3.784 | 13.636 | 5.352 | 22.444 | 20.786 |
| | 16.742 | 28.321 | 15.066 | -1.258 | 1.047 | 27.650 | -7.541 | -4.189 | 3.989 | 17.911 | 18.384 | 5.017 | -12.246 |
| | -12.930 | -6.772 | 2.868 | 28.478 | 21.219 | 23.958 | -21.369 | 19.685 | 11.116 | 23.023 | 21.062 | 2.146 | 14.941 |
| | 11.593 | -26.096 | -15.853 | 13.616 | -18.121 | 17.224 | 0.202 | 5.600 | 12.599 | 22.257 | 9.019 | 23.512 | 24.107 |
| | 15.908 | 2.892 | 5.184 | -17.320 | -21.243 | 18.661 | 23.191 | 3.058 | -4.236 | 20.850 | 10.480 | 24.224 | 16.829 |
| | 19.989 | 18.415 | -10.722 | 29.318 | 15.032 | -0.286 | 8.022 | -6.062 | 14.447 | 25.823 | 23.602 | 26.113 | 27.759 |
| | 5.666 | -4.851 | -24.023 | 25.880 | 16.819 | 25.787 | 17.372 | 9.840 | 22.056 | 15.961 | 13.035 | 21.283 | 22.737 |
| | -7.114 | -21.363 | -12.791 | 11.499 | 20.843 | 12.418 | -12.959 | -0.621 | 23.428 | 20.222 | 8.730 | 22.725 | 15.824 |
| | 25.587 | -18.131 | 24.176 | -11.750 | -16.689 | -19.063 | -15.252 | 23.238 | 24.765 | 5.854 | 18.736 | 10.556 | 3.561 |
| | 17.582 | -23.070 | -12.132 | -3.828 | -24.901 | 16.649 | 0.910 | 1.769 | 17.830 | 21.939 | 14.172 | 4.312 | -9.040 |
| | 8.616 | 15.854 | -23.144 | 26.967 | -11.736 | 28.242 | 2.152 | -2.672 | 3.877 | 21.299 | 13.448 | -3.051 | -13.047 |
| | -7.916 | 8.276 | 5.551 | 6.005 | -16.100 | -7.703 | 22.535 | 3.829 | 16.210 | 15.994 | 21.655 | 0.870 | 21.113 |
| | -17.155 | -9.732 | -17.182 | 2.978 | -3.754 | 11.101 | 15.217 | 19.040 | 18.868 | 12.745 | 14.987 | 5.606 | 25.407 |
| | -23.994 | 2.073 | -6.411 | 16.548 | 25.008 | -13.384 | 26.141 | 20.670 | 3.189 | -18.210 | 26.140 | 23.446 | -1.513 |
| | 3.026 | 22.878 | 24.737 | 12.564 | -5.529 | 23.410 | 25.105 | 5.268 | 10.748 | -1.330 | 3.876 | 6.260 | 8.200 |
| | -16.618 | -24.955 | 16.805 | 18.943 | -30.413 | -6.029 | 20.574 | 16.776 | 4.990 | 21.848 | 9.370 | 21.607 | 12.310 |
| | 12.992 | -3.823 | -13.830 | -8.783 | 18.091 | -29.433 | 7.729 | 25.050 | 16.588 | 19.999 | 25.054 | 21.130 | 4.761 |
| | 16.343 | 17.860 | -25.095 | 16.021 | 4.707 | -30.690 | -6.029 | -14.043 | -0.154 | 18.907 | 19.072 | -15.210 | 15.759 |
| | 21.824 | 17.034 | -22.425 | 25.355 | -22.355 | 13.924 | 1.654 | -8.426 | 18.117 | 23.661 | 23.279 | 5.360 | 26.084 |
| | -9.425 | 3.905 | -21.451 | 22.946 | -21.757 | -21.236 | 17.249 | 10.118 | 18.588 | 27.318 | 0.325 | 18.149 | 18.325 |
| | -14.852 | 20.892 | 9.548 | 13.537 | 23.219 | 7.948 | -0.145 | 22.132 | 1.515 | 6.818 | 21.976 | 18.655 | 14.016 |
| | 24.286 | 16.028 | -1.038 | -6.821 | 18.641 | 17.338 | 26.318 | -11.881 | 19.108 | 2.706 | 7.850 | 1.465 | 26.857 |
| | 15.018 | -10.635 | 11.610 | -8.843 | 28.759 | 2.569 | 22.923 | 17.615 | 9.635 | 22.834 | 15.106 | 17.014 | 24.825 |
| | 0.746 | 10.927 | -6.871 | -21.171 | -3.438 | -17.351 | -2.428 | 1.046 | 8.912 | 13.981 | 16.962 | 18.101 | 10.534 |
| | -28.281 | 25.100 | 22.237 | -7.810 | -12.593 | -14.643 | 21.275 | 6.694 | 22.264 | 17.062 | 12.104 | 24.414 | 20.521 |
| | 26.779 | -23.749 | -22.165 | 10.220 | 27.176 | 22.382 | 8.782 | 25.697 | 25.868 | 14.780 | 4.015 | 21.559 | 23.574 |
| | -23.780 | 25.525 | 19.104 | -11.292 | 22.944 | -23.882 | 6.213 | -10.786 | 8.563 | 16.010 | 0.995 | 20.282 | -9.406 |
| | 17.789 | -9.258 | 14.237 | 7.672 | 7.561 | -16.567 | -3.191 | 16.326 | 25.231 | 15.184 | 3.029 | 25.182 | -0.419 |
| | 10.030 | 21.424 | 21.421 | 23.862 | -4.644 | -18.681 | -4.005 | 8.227 | 24.694 | 21.141 | 24.750 | 16.849 | 7.308 |
| | 1.635 | -2.944 | 17.993 | 8.407 | -32.049 | 28.696 | 9.631 | 20.347 | 7.735 | 18.753 | 20.264 | 6.803 | 17.868 |
| | 23.562 | -5.779 | -20.969 | -5.213 | -6.212 | 25.644 | 19.669 | -1.095 | -0.553 | 27.853 | 8.160 | -19.681 | 18.313 |
| | 22.824 | 19.981 | 3.506 | -9.590 | 21.316 | -3.930 | 16.216 | 1.094 | 28.351 | 20.865 | 9.119 | 1.088 | 16.233 |
| | -22.289 | 13.679 | -16.809 | -27.296 | 1.393 | -4.946 | 29.927 | 14.327 | 14.648 | 27.258 | 24.019 | 18.841 | 24.538 |
| | 10.661 | 2.024 | -6.093 | -16.264 | 25.350 | 19.367 | 23.908 | 15.713 | 2.106 | 26.024 | 16.261 | 15.171 | 14.589 |
| | -6.124 | -17.918 | -10.148 | -2.538 | 23.266 | -6.542 | -14.156 | 2.228 | 13.248 | 8.549 | 1.310 | 5.153 | 10.416 |
| | 20.665 | 13.113 | -11.929 | 22.711 | 29.154 | -20.968 | 12.236 | 9.654 | -5.937 | 28.849 | 5.212 | 20.814 | 7.738 |

| | | | | | | | | | | | | | |
|---|---|---|---|---|---|---|---|---|---|---|---|---|---|
| | 17.058 | 3.089 | 14.773 | 8.626 | -22.870 | 20.798 | 21.457 | 17.284 | 10.196 | 19.887 | 21.835 | 22.470 | 19.580 |
| | -18.942 | -9.877 | -13.427 | 1.443 | -9.721 | 0.560 | 25.180 | 8.277 | 25.120 | 15.136 | 0.321 | | 0.556 |
| | 23.432 | -23.167 | 6.304 | -4.688 | -12.605 | 28.814 | -13.174 | 26.394 | 28.969 | 24.338 | -3.463 | | 25.195 |
| | 26.417 | -4.873 | -1.505 | -3.545 | -28.560 | -11.745 | 3.538 | -6.362 | 18.314 | 17.755 | 22.877 | | 23.908 |
| | 4.692 | 21.569 | 7.221 | 24.859 | -9.006 | -12.274 | 8.945 | -21.080 | 9.220 | 14.140 | 5.067 | | 20.130 |
| | -18.381 | 6.165 | 0.059 | 5.185 | 0.230 | -29.277 | -9.736 | 4.557 | -13.330 | 26.811 | 0.443 | | 17.803 |
| | 24.402 | 12.417 | -19.694 | 12.021 | -25.128 | -9.194 | -2.381 | -0.305 | 17.277 | 9.964 | 20.569 | | 14.145 |
| | 12.359 | -18.601 | 17.917 | -29.614 | 30.844 | 26.891 | 17.648 | 6.298 | -0.889 | 13.701 | 16.458 | | 7.919 |
| | 9.297 | 10.750 | 20.153 | 21.904 | 1.377 | -10.002 | -1.509 | 24.663 | -16.745 | 14.534 | 9.387 | | -0.716 |
| | 2.706 | 16.940 | 15.262 | -6.673 | 4.742 | 13.838 | 6.245 | -0.736 | 7.469 | 25.989 | 4.196 | | 24.742 |
| | 22.270 | -0.527 | 16.966 | 16.432 | -29.144 | 10.670 | 8.139 | 7.353 | -5.228 | -8.656 | 9.795 | | 19.157 |
| | -13.722 | -6.664 | 6.957 | -8.664 | 20.052 | -29.778 | -12.150 | -1.682 | 6.566 | 19.733 | 7.355 | | 16.214 |
| | 19.263 | 15.554 | -2.649 | 28.267 | 7.022 | 17.016 | -9.106 | 19.792 | 27.877 | 16.313 | 12.925 | | 1.844 |
| | -12.622 | 4.355 | -6.402 | 27.368 | 26.897 | -0.396 | -5.192 | 9.506 | 19.600 | 24.572 | 6.621 | | 25.263 |
| | 6.002 | 18.755 | -1.204 | -9.776 | 9.516 | 17.859 | 2.917 | 14.618 | 17.047 | 7.604 | 10.361 | | 22.825 |
| | -8.644 | -6.972 | 13.747 | 22.865 | -7.583 | -29.123 | 22.902 | 3.725 | -7.522 | -1.147 | -3.144 | | -0.935 |
| | 25.184 | 20.754 | 7.827 | 4.226 | 26.047 | -6.419 | 13.176 | 16.661 | 22.046 | 3.555 | 2.035 | | -2.364 |
| | 16.190 | -2.514 | 20.532 | -19.506 | -8.371 | 8.040 | 5.840 | -12.055 | 10.371 | 5.854 | 6.077 | | 19.034 |
| | 3.757 | -18.080 | 8.699 | 23.758 | 23.995 | 26.718 | 7.221 | -8.399 | -8.966 | 3.012 | 7.127 | | 22.182 |
| | -17.077 | 12.674 | 11.494 | -0.764 | 15.892 | 0.160 | 19.943 | 5.395 | 20.185 | 14.360 | | | 21.587 |
| | -14.771 | -17.471 | 20.120 | -15.525 | -5.924 | 11.355 | -8.597 | 20.782 | 13.840 | -8.918 | | | 13.464 |
| | -21.234 | 19.911 | -16.490 | 15.872 | 29.710 | 24.218 | 4.065 | 15.206 | -17.598 | 20.488 | | | -25.416 |
| | 17.635 | 23.400 | 9.088 | -14.472 | -13.917 | 19.511 | -2.664 | -11.465 | 9.256 | 16.899 | | | 19.719 |
| | -12.030 | -11.390 | 13.310 | 28.187 | -20.398 | 8.049 | 11.456 | 13.544 | 21.297 | -1.021 | | | 7.836 |
| | 4.871 | 13.134 | 13.010 | 20.091 | 26.866 | -10.806 | -3.694 | -3.799 | -4.201 | 23.172 | | | 23.188 |
| | 7.940 | -3.892 | -3.669 | 18.180 | -4.038 | -18.479 | -14.070 | 6.170 | 22.430 | 12.132 | | | 18.057 |
| | 23.889 | -5.342 | -15.027 | -11.248 | -12.580 | 25.324 | 1.850 | 10.049 | -24.443 | 22.064 | | | 8.624 |
| | 4.070 | 16.278 | 0.969 | -3.314 | 21.425 | -9.643 | 17.794 | -5.108 | 5.335 | 28.029 | | | 23.535 |
| | -5.419 | 12.957 | -2.679 | 11.630 | 27.997 | 28.768 | 24.601 | 26.408 | -5.061 | 21.150 | | | -24.290 |
| | -29.541 | 6.595 | -2.425 | -14.767 | 24.364 | 10.608 | -5.600 | 15.901 | 15.435 | 16.854 | | | -15.622 |
| | -24.612 | -22.193 | 16.747 | -10.085 | -10.193 | 18.767 | -10.509 | 13.223 | -17.317 | 30.093 | | | -22.208 |
| | -22.886 | -23.779 | 26.125 | -25.297 | -25.715 | 22.021 | 16.724 | 18.317 | 6.292 | 24.121 | | | |
| | 6.788 | -23.575 | -7.040 | -23.989 | -28.958 | -17.188 | 9.920 | 16.683 | 7.978 | 10.941 | | | |
| | 0.702 | -12.597 | -1.335 | 26.895 | 10.498 | 29.379 | -7.503 | 17.562 | 23.278 | 15.408 | | | |
| | -0.595 | -17.681 | -15.003 | 18.727 | 24.477 | 8.346 | -6.820 | -10.114 | 13.437 | 27.225 | | | |
| | -1.851 | -21.983 | 7.938 | 13.517 | 17.339 | 6.085 | 19.849 | 11.412 | -8.558 | 12.476 | | | |

| | | | | | | | | | | |
|---|---|---|---|---|---|---|---|---|---|---|
| | -18.761 | -4.913 | 10.287 | -11.664 | -27.706 | -30.727 | -15.698 | -0.028 | 20.546 | 9.906 |
| | 22.600 | 17.277 | 18.035 | -29.493 | -21.364 | -16.355 | 28.810 | 12.875 | 18.440 | -25.873 |
| | 11.138 | 8.520 | 24.093 | 23.909 | -13.635 | -26.025 | 22.557 | 18.042 | 15.711 | 28.952 |
| | -21.486 | -1.648 | 25.921 | -22.396 | 19.694 | 22.579 | 0.994 | 22.287 | 9.143 | 21.674 |
| | -22.290 | -22.628 | -24.430 | 24.481 | -16.161 | -18.700 | 27.503 | 10.146 | -3.646 | 4.438 |
| | 10.008 | 17.963 | -8.542 | -12.537 | -27.272 | 17.588 | 20.972 | 10.648 | 29.277 | 18.495 |
| | -20.908 | 19.785 | -23.103 | -28.304 | -5.296 | 0.757 | 24.906 | 27.928 | 26.085 | 30.800 |
| | 25.042 | -6.691 | -22.837 | -11.726 | -15.041 | -27.625 | -1.562 | 9.051 | 0.691 | 27.291 |
| | 5.942 | -11.143 | -10.469 | -3.483 | -12.258 | 7.785 | 1.681 | 15.771 | 21.236 | 19.251 |
| | 8.209 | 15.507 | 1.261 | 22.152 | 9.140 | 28.959 | 23.135 | 14.664 | 15.376 | -14.456 |
| | -14.501 | 14.384 | -9.693 | -29.552 | 12.330 | 13.255 | -7.439 | 5.607 | 10.640 | 25.149 |
| | 19.812 | -13.211 | -22.001 | -20.037 | -29.865 | -10.852 | 14.384 | 21.014 | -12.590 | 13.959 |
| | -0.125 | 11.178 | 4.857 | -22.279 | 23.623 | -16.666 | 11.826 | -14.369 | 16.967 | 11.401 |
| | 18.707 | -25.995 | 8.682 | -26.601 | -7.608 | -21.830 | 6.056 | 13.811 | 5.280 | 22.156 |
| | 12.385 | -4.000 | -13.067 | 6.279 | 21.417 | 4.673 | 15.294 | 18.399 | 9.275 | -2.150 |
| | 30.198 | 4.751 | -13.912 | 1.039 | -9.790 | 26.777 | -10.977 | 20.708 | 18.395 | 12.516 |
| | -30.064 | 21.447 | 10.250 | -19.271 | -3.021 | 23.691 | -19.784 | 11.851 | 3.562 | 16.658 |
| | 1.766 | 16.017 | -0.106 | -6.036 | -15.642 | 22.274 | -1.191 | 7.249 | 7.201 | 22.304 |
| | -4.987 | 4.108 | -25.888 | -7.739 | 11.875 | -3.356 | -13.618 | -3.409 | 8.047 | 0.250 |
| | 6.217 | -1.054 | -5.815 | 20.777 | 26.533 | -17.509 | -14.550 | 23.431 | 5.706 | -0.583 |
| | 3.559 | 5.608 | -4.917 | -23.463 | 4.625 | 25.699 | 16.909 | 22.166 | 28.857 | 18.024 |
| | -0.282 | 12.973 | -7.391 | -11.370 | -6.515 | 18.314 | -21.698 | 4.403 | 30.264 | 3.435 |
| | -27.164 | 8.842 | 12.488 | -5.347 | 19.818 | 24.860 | 3.407 | 17.069 | 12.270 | -5.222 |
| | 24.355 | -11.247 | 16.923 | -7.596 | -28.812 | -8.744 | 24.864 | -16.981 | -5.475 | 27.061 |
| | 20.899 | -21.103 | -19.262 | 7.302 | -25.055 | 27.984 | 12.676 | -3.721 | -25.134 | 21.576 |
| | 9.349 | 8.090 | -20.610 | 23.231 | -18.957 | -29.304 | -5.066 | -20.091 | -3.530 | -3.363 |
| | 28.673 | -7.923 | 1.552 | 1.183 | -27.126 | 21.811 | 14.299 | -22.573 | 17.681 | 10.063 |
| | 17.499 | 13.774 | 8.088 | 21.472 | 3.301 | -15.227 | 6.980 | 27.604 | 24.253 | 24.897 |
| | 27.859 | 22.350 | 25.440 | 27.664 | 5.305 | 1.835 | -7.449 | 24.718 | -2.471 | 23.428 |
| | -17.597 | 17.124 | -16.183 | 26.785 | 13.661 | 1.201 | 8.073 | 13.458 | 10.158 | 11.629 |
| | 13.880 | 6.773 | 20.219 | 18.276 | 20.199 | -28.737 | 11.890 | 2.528 | 13.674 | 16.437 |
| | 27.523 | 9.514 | 11.534 | 9.056 | -12.258 | 22.431 | 20.943 | 19.754 | 16.854 | 15.456 |
| | 12.373 | -2.537 | 5.900 | 11.915 | 6.721 | -13.950 | 1.473 | 12.593 | 15.150 | 11.103 |
| | 16.514 | -3.590 | -22.260 | 15.029 | -2.081 | -22.118 | 5.584 | 5.800 | 18.066 | 25.319 |
| | 25.806 | 23.817 | 6.985 | 18.615 | -9.199 | -18.516 | 26.190 | -1.079 | 2.774 | 17.845 |
| | 8.196 | -0.563 | -18.872 | 25.440 | -0.063 | -19.604 | 29.361 | 10.813 | 29.009 | 27.429 |

| | | | | | | | | | | |
|---|---|---|---|---|---|---|---|---|---|---|
| | -17.311 | -24.875 | 28.898 | 9.138 | -28.373 | 3.393 | -0.850 | 24.970 | 25.050 | 20.304 |
| | -13.232 | 18.848 | 26.989 | 3.432 | -20.121 | 10.092 | -8.201 | 5.218 | -24.363 | 21.135 |
| | 17.655 | -22.443 | -14.031 | 22.577 | 17.633 | 13.247 | -16.677 | -5.434 | 29.955 | 24.117 |
| | -14.776 | 14.107 | -14.908 | 23.824 | 6.344 | 18.040 | 21.712 | -0.550 | 16.545 | 30.194 |
| | -0.355 | -10.732 | -25.592 | -20.413 | -13.518 | -11.611 | -2.608 | -3.042 | 5.278 | 22.066 |
| | -2.347 | 6.730 | -16.273 | 7.652 | 9.421 | 4.540 | -3.990 | 19.964 | 7.397 | 14.351 |
| | -15.342 | 5.494 | 21.569 | 4.320 | -16.153 | -17.724 | 8.890 | -20.114 | -10.050 | 29.567 |
| | 20.554 | 11.852 | -2.087 | 21.060 | -22.193 | -27.401 | 3.407 | 22.407 | 24.071 | 16.491 |
| | -14.320 | 8.501 | 10.143 | 12.851 | 3.161 | -7.642 | -1.779 | -9.651 | -7.772 | 25.012 |
| | 28.743 | -11.589 | -25.129 | 18.000 | -8.812 | -26.423 | 17.398 | -6.909 | -3.873 | 20.836 |
| | 23.617 | -9.279 | -26.573 | -25.001 | 11.434 | 21.899 | 0.225 | 16.943 | -21.483 | 26.760 |
| | 31.512 | 2.703 | 16.688 | 24.834 | 0.122 | 8.600 | -6.154 | 14.252 | 4.070 | 18.480 |
| | -21.560 | 13.117 | 15.058 | 16.393 | 12.710 | 6.264 | 13.123 | 24.681 | -1.028 | 9.019 |
| | 29.758 | 14.852 | -3.955 | 3.931 | 23.048 | 1.150 | -7.727 | 8.051 | -4.602 | 24.759 |
| | 8.420 | 13.965 | -22.975 | -21.169 | 19.054 | -22.199 | -9.375 | 5.566 | 8.317 | 0.142 |
| | -11.149 | -7.200 | -21.172 | -22.095 | 15.463 | 20.808 | -21.901 | -4.693 | -14.534 | 19.656 |
| | -22.229 | -5.647 | -25.526 | 5.125 | -4.389 | 5.518 | 26.230 | 25.884 | 15.597 | 15.953 |
| | -26.560 | -13.325 | 27.594 | 3.922 | 24.214 | -15.675 | 18.183 | 18.605 | 6.661 | 11.514 |
| | 30.328 | 25.440 | -11.339 | 3.006 | -25.104 | -20.838 | 23.296 | 14.115 | 4.757 | 23.479 |
| | 17.370 | -14.224 | -7.388 | 1.638 | -17.173 | 10.649 | 12.618 | 15.634 | 13.729 | 15.180 |
| | 10.721 | 15.720 | 18.278 | -8.078 | -26.964 | -22.985 | 8.842 | 18.125 | 27.879 | 24.780 |
| | -3.441 | -10.690 | -8.827 | 0.210 | 14.413 | -28.318 | -3.541 | 29.335 | 9.999 | 12.379 |
| | -1.989 | -12.665 | -11.595 | -6.304 | 6.606 | 11.586 | 14.249 | 13.133 | 29.723 | 9.570 |
| | 23.000 | -11.994 | -23.135 | -10.339 | 4.513 | 13.174 | 20.421 | -5.630 | 16.214 | -0.049 |
| | 8.220 | 18.933 | -2.999 | -15.714 | 3.019 | 22.794 | 1.047 | -5.925 | -0.245 | 29.121 |
| | -22.010 | 8.375 | 8.688 | -11.067 | -21.393 | -2.174 | -2.117 | 7.032 | 22.479 | 14.990 |
| | -1.507 | 16.770 | -9.201 | -5.132 | 8.187 | 24.206 | -19.712 | 2.626 | 12.605 | 25.032 |
| | 21.794 | 10.809 | 6.114 | 21.656 | -9.605 | -10.486 | 22.275 | 4.783 | -7.004 | 24.193 |
| | 28.593 | -2.876 | 9.701 | -12.812 | 5.204 | 21.192 | 15.180 | 7.796 | 15.544 | 17.909 |
| | 9.740 | -27.777 | 21.723 | -14.509 | -6.812 | -24.783 | 27.540 | 27.734 | 11.746 | 20.654 |
| | 1.765 | -26.273 | 2.874 | -18.065 | 23.972 | 6.863 | 4.429 | 22.709 | -23.243 | 4.281 |
| | -19.404 | 27.379 | 16.072 | 18.342 | -25.664 | 9.780 | 3.516 | 20.587 | 1.750 | 10.207 |
| | 10.193 | 2.297 | -9.611 | 20.169 | -10.883 | -19.924 | 26.268 | 29.055 | -4.254 | 21.428 |
| | 12.639 | 0.538 | 4.921 | -8.293 | -17.445 | 10.198 | 19.483 | -14.704 | 25.064 | 26.481 |
| | 26.451 | -14.661 | -13.740 | -24.012 | -20.004 | 22.273 | 7.122 | 12.056 | 12.103 | 21.877 |
| | 5.615 | -21.081 | 16.485 | 18.926 | -7.743 | -16.007 | -8.920 | -4.972 | 19.622 | 8.779 |

| | | | | | | | | | | |
|---|---|---|---|---|---|---|---|---|---|---|
| | 25.201 | -22.200 | -16.419 | -13.720 | -0.955 | -11.896 | 23.345 | 23.671 | 14.119 | 24.160 |
| | 16.900 | 14.062 | -12.665 | 7.223 | -16.404 | -5.998 | -0.955 | 15.183 | 20.335 | 22.854 |
| | -13.820 | 10.310 | 9.471 | 1.539 | -6.054 | -26.744 | -21.816 | -0.935 | 15.026 | 7.810 |
| | -4.090 | -24.855 | -22.661 | -0.279 | 23.691 | 14.674 | 17.468 | -10.421 | -11.791 | 12.699 |
| | -14.577 | -26.896 | 18.774 | -2.295 | 15.856 | 16.864 | -15.140 | 6.002 | 10.745 | 19.199 |
| | -4.697 | 21.685 | -3.722 | 23.344 | -22.139 | -24.939 | -5.655 | 14.692 | 9.802 | 3.257 |
| | -9.402 | 3.726 | -20.901 | 7.384 | -2.151 | 24.453 | -1.914 | 2.884 | 26.741 | 26.294 |
| | 23.441 | 20.851 | 10.929 | -14.354 | -27.465 | 9.618 | 12.806 | 28.011 | -14.892 | 15.875 |
| | -31.415 | -7.729 | -26.543 | -18.121 | 9.145 | 8.339 | -4.066 | -3.658 | 29.889 | 16.562 |
| | -21.595 | 11.688 | -19.700 | -5.123 | 15.370 | -27.269 | 10.393 | 3.500 | -28.684 | 4.604 |
| | 11.939 | 26.537 | -7.048 | -7.443 | 23.089 | -1.654 | 22.118 | -18.198 | 8.838 | 12.360 |
| | -18.743 | 6.909 | -29.273 | -11.352 | -14.067 | -24.315 | -2.543 | -11.842 | -1.753 | -0.132 |
| | 4.005 | -6.255 | -11.885 | 9.912 | -16.071 | 26.207 | 24.789 | -3.148 | 18.826 | 20.193 |
| | 24.753 | 9.656 | 25.531 | -4.089 | -7.662 | 21.183 | 6.462 | -8.696 | 20.879 | 9.731 |
| | 13.918 | 0.935 | 2.613 | 13.769 | -2.142 | -15.373 | -5.211 | 24.156 | -0.238 | 28.410 |
| | 8.448 | -9.734 | 26.366 | 1.294 | -21.103 | -21.691 | 18.401 | 28.849 | 4.590 | 0.747 |
| | 27.419 | -12.902 | 20.107 | 0.500 | 4.778 | -8.547 | -13.589 | 12.949 | 11.719 | 17.727 |
| | 3.448 | -15.144 | -20.586 | -21.619 | 20.076 | 7.489 | 2.569 | -0.066 | -6.826 | 24.152 |
| | -29.272 | -28.408 | 15.975 | 17.527 | -22.119 | -4.720 | 23.914 | -1.329 | -0.636 | 27.469 |
| | 20.032 | 12.316 | -30.207 | -6.773 | -25.961 | -14.189 | 21.224 | -17.618 | 27.240 | 6.604 |
| | 8.949 | 19.597 | -3.113 | 8.703 | -8.911 | 21.740 | 20.112 | 18.205 | 8.055 | 23.151 |
| | 5.209 | 20.323 | -18.825 | -8.656 | 7.296 | -7.339 | 15.200 | -13.333 | 5.946 | 16.690 |
| | 5.726 | -17.296 | 17.717 | 11.431 | -15.222 | 14.305 | 11.130 | 21.420 | 6.922 | 14.474 |
| | -14.609 | -20.974 | 11.568 | -16.545 | -13.763 | -15.961 | 7.263 | 13.661 | 28.313 | 21.051 |
| | -16.474 | 2.166 | -27.820 | 12.944 | 17.373 | 0.331 | 27.505 | 30.322 | -2.696 | 4.239 |
| | -27.184 | 21.000 | -23.693 | 21.858 | -22.998 | 1.805 | 5.871 | 17.589 | 21.587 | 24.184 |
| | -9.017 | 22.256 | 5.687 | -14.664 | -19.232 | 14.777 | -19.269 | 11.382 | -1.573 | 26.264 |
| | -8.523 | 9.627 | -24.770 | -24.861 | -16.047 | -21.291 | 3.654 | -2.498 | -5.995 | 4.790 |
| | -5.549 | 13.615 | 26.292 | 2.327 | 4.066 | -5.197 | 14.598 | 10.237 | 9.487 | 15.719 |
| | 27.516 | 7.866 | 3.734 | -10.305 | -20.583 | 23.339 | -2.603 | -4.376 | 17.593 | 21.758 |
| | 19.375 | 3.798 | -29.848 | -17.780 | 19.116 | 4.788 | 18.600 | -19.047 | 25.184 | 26.284 |
| | 17.048 | 26.641 | -4.028 | 21.340 | 2.184 | -6.537 | 16.800 | 29.730 | -17.734 | 19.869 |
| | 1.409 | -8.342 | -28.530 | 15.498 | -3.377 | 7.286 | 2.803 | 28.476 | 23.536 | 8.328 |
| | -11.872 | 16.721 | 2.469 | 7.294 | 19.574 | -13.563 | 6.513 | 25.267 | 11.991 | 25.055 |
| | 21.064 | 10.801 | 9.564 | 19.527 | -25.068 | -19.218 | 11.743 | -6.272 | 26.105 | 26.839 |
| | -24.295 | 7.929 | -30.640 | 8.846 | 18.551 | -17.463 | 13.410 | 1.222 | -4.137 | 9.374 |

| | | | | | | | | | | |
|---|---|---|---|---|---|---|---|---|---|---|
| | 24.593 | 1.069 | 0.751 | -0.914 | 20.773 | 20.988 | 9.508 | 22.403 | 19.361 | 12.956 |
| | -23.844 | 28.543 | 17.209 | 13.620 | -9.711 | -2.163 | 22.734 | -5.394 | 20.325 | 3.283 |
| | 26.812 | 3.179 | 6.471 | -9.988 | 10.218 | 21.782 | -17.285 | -12.937 | -13.242 | 26.210 |
| | 9.673 | -10.000 | 3.545 | 18.430 | -17.816 | -4.622 | 15.566 | 18.801 | 0.171 | 25.832 |
| | -14.464 | -4.065 | 19.479 | 12.225 | -8.180 | -21.226 | 0.944 | 2.939 | 9.025 | 20.244 |
| | 16.435 | -15.612 | -5.948 | -8.985 | 18.186 | 2.162 | -0.891 | -10.741 | -6.275 | 16.290 |
| | 10.821 | -26.736 | -20.744 | 10.347 | 2.550 | 4.513 | 4.129 | 17.061 | 24.881 | 17.156 |
| | 7.167 | 15.689 | 20.552 | 6.205 | 3.268 | -19.640 | -11.474 | 17.972 | 23.816 | 14.401 |
| | -15.632 | -10.542 | -19.900 | -20.127 | 16.012 | -24.099 | 5.643 | -3.514 | 21.179 | -6.809 |
| | -25.153 | -1.444 | 27.238 | -7.847 | -10.790 | 22.486 | -3.018 | 21.551 | 23.495 | 14.792 |
| | -29.215 | 1.720 | 10.761 | 22.105 | 8.254 | -9.365 | 19.575 | -8.909 | 12.622 | 12.390 |
| | -7.645 | 22.790 | 29.450 | 17.246 | 4.877 | 1.145 | -14.831 | 23.487 | -0.812 | 23.255 |
| | -21.995 | 14.881 | -1.030 | 10.991 | 22.351 | -1.458 | 27.307 | 28.070 | 21.751 | 19.833 |
| | -18.437 | 18.705 | 20.996 | -8.665 | -0.132 | -17.243 | 24.828 | 24.497 | 19.060 | 13.203 |
| | 28.536 | -4.489 | -25.122 | 3.908 | 24.062 | 21.566 | 6.735 | 10.986 | 15.789 | |
| | 3.652 | -5.527 | 25.622 | -19.958 | -2.528 | 19.396 | -4.784 | 9.176 | 6.342 | |
| | 0.826 | 14.101 | -30.101 | 0.059 | 11.180 | 13.562 | -6.118 | 29.173 | 16.300 | |
| | -8.213 | 9.803 | -22.271 | 15.993 | 15.453 | 17.504 | 16.351 | -17.274 | -5.727 | |
| | 24.720 | 9.031 | -21.376 | -16.329 | -6.394 | 15.767 | 22.026 | 14.708 | 28.114 | |
| | -24.296 | -10.579 | 16.661 | -2.087 | 2.533 | 11.767 | -13.863 | 27.357 | -2.367 | |
| | -15.337 | 15.063 | -13.315 | -3.961 | -1.015 | 18.613 | 1.316 | 7.627 | -14.493 | |
| | -25.769 | 6.083 | -25.618 | 0.811 | -11.939 | 23.042 | -10.269 | 26.202 | 7.857 | |
| | 26.962 | 26.041 | 28.383 | 24.861 | 16.442 | 24.334 | -0.462 | -4.200 | 7.436 | |
| | 14.997 | -28.198 | 19.338 | 1.173 | 18.724 | 17.308 | 20.248 | 15.460 | 22.461 | |
| | 2.244 | 29.620 | 21.084 | -11.534 | 7.217 | 20.950 | 4.006 | -2.319 | 12.749 | |
| | 8.508 | -26.204 | -23.833 | 19.250 | 3.739 | 0.244 | 5.912 | -10.161 | -4.041 | |
| | -25.546 | 13.057 | -6.048 | -26.183 | 11.138 | 24.635 | 17.126 | -12.757 | 3.198 | |
| | 26.954 | 18.331 | 2.428 | -7.291 | -9.174 | 7.057 | -1.904 | 14.552 | 15.310 | |
| | -19.181 | 12.074 | 4.691 | -18.925 | -9.344 | -8.293 | 7.471 | 20.708 | -12.203 | |
| | 21.068 | 22.670 | -29.421 | 15.036 | 14.059 | 22.526 | 23.618 | 13.234 | -7.757 | |
| | 25.861 | 29.056 | 21.550 | -24.612 | 12.675 | -11.964 | 0.430 | 3.181 | 8.769 | |
| | -23.310 | 17.236 | -8.674 | 23.151 | -15.674 | -14.265 | 26.728 | 5.139 | -3.102 | |
| | -27.813 | -19.001 | 8.065 | 3.407 | -12.743 | -4.161 | 1.801 | -4.975 | 25.425 | |
| | 27.529 | -3.186 | 15.856 | 5.506 | 0.297 | 24.843 | 18.995 | 7.531 | 20.885 | |
| | -4.135 | 13.386 | -14.179 | -19.995 | 18.375 | 20.241 | -9.840 | 17.311 | 6.156 | |
| | -16.946 | -23.393 | -26.624 | -24.006 | -2.064 | -19.376 | -14.152 | -3.459 | 2.314 | |

| | | | | | | | | | |
|---|---|---|---|---|---|---|---|---|---|
| | -25.893 | 24.910 | 12.035 | -23.087 | -10.524 | -14.931 | -8.981 | 11.298 | -0.340 |
| | -1.140 | -28.481 | -3.766 | 10.347 | -19.205 | 14.791 | 8.884 | 27.153 | 22.659 |
| | -10.534 | -11.890 | -21.123 | -6.227 | -21.008 | 6.899 | 17.193 | 24.327 | 26.803 |
| | 12.027 | 27.621 | 18.058 | 14.721 | -6.991 | -16.142 | 6.115 | -23.319 | 14.000 |
| | -20.569 | -0.982 | -14.425 | 2.017 | -25.959 | -22.291 | 16.071 | 22.648 | 16.467 |
| | -8.946 | -5.392 | 25.027 | -8.390 | -12.968 | -18.336 | 2.821 | 21.177 | 16.953 |
| | 2.118 | 9.116 | -24.337 | -13.003 | 27.196 | -20.090 | 10.139 | 28.323 | 1.587 |
| | 25.315 | 5.007 | 6.248 | -16.483 | 25.964 | 26.346 | 0.631 | 9.979 | 11.990 |
| | -19.286 | -2.698 | -5.360 | 4.502 | 7.505 | -11.788 | 3.483 | -27.936 | 24.071 |
| | -22.321 | 15.453 | 1.169 | 24.775 | 3.852 | -7.538 | -1.028 | 6.996 | 22.900 |
| | -24.160 | -28.035 | -22.674 | -10.042 | 17.040 | 15.399 | 7.024 | -19.906 | 2.752 |
| | -18.512 | 18.622 | 1.622 | 13.112 | 7.923 | 14.569 | 9.200 | -1.894 | -3.352 |
| | -19.755 | 6.576 | 3.705 | -2.092 | -11.511 | 12.666 | 25.963 | -3.727 | -4.230 |
| | 19.526 | 25.384 | -13.047 | 20.330 | -17.838 | -20.845 | 22.346 | 9.151 | 0.564 |
| | -28.313 | 15.594 | -8.045 | 17.930 | -0.237 | 19.742 | 24.645 | -16.106 | 20.705 |
| | -27.445 | 7.232 | 18.919 | 2.268 | -21.825 | -8.462 | -1.718 | 16.651 | 21.361 |
| | -14.686 | -2.100 | 15.109 | -15.729 | -8.607 | -17.404 | 1.708 | 0.798 | 7.680 |
| | -12.503 | -20.939 | -21.344 | 17.051 | -8.089 | -23.565 | 12.463 | 7.799 | -2.779 |
| | -6.746 | 12.041 | 28.828 | -10.563 | -6.833 | 22.713 | 8.769 | 18.822 | 19.175 |
| | -27.687 | 26.922 | 20.502 | 0.085 | 21.882 | 26.755 | -2.600 | 28.906 | 23.582 |
| | 27.205 | 17.428 | 9.706 | 14.691 | 6.878 | 3.042 | -17.366 | 20.198 | 20.085 |
| | 3.746 | -7.952 | 6.342 | -11.369 | -20.562 | 27.503 | 11.614 | -2.840 | 13.766 |
| | -11.809 | 21.551 | 12.037 | -21.735 | 17.363 | -4.644 | -0.735 | 12.461 | 4.673 |
| | -25.144 | -22.395 | -7.506 | -4.420 | 1.778 | 20.973 | -5.150 | -8.711 | 6.260 |
| | 24.737 | 26.214 | -16.635 | 12.661 | 26.309 | 10.035 | 8.675 | 19.570 | 7.202 |
| | 23.921 | 16.205 | -20.781 | -6.792 | -9.567 | 1.605 | 7.903 | -2.035 | 3.958 |
| | -7.289 | -10.269 | 0.504 | -2.966 | -27.032 | -7.634 | -10.648 | 6.870 | 12.786 |
| | -10.684 | 22.258 | 15.639 | 21.979 | 8.865 | 25.179 | -3.259 | -5.740 | 22.537 |
| | 22.476 | -6.493 | -3.012 | 20.497 | 15.373 | 22.021 | 9.154 | -4.601 | 26.950 |
| | 5.271 | 23.878 | 2.164 | 17.448 | 6.368 | -7.195 | -16.163 | 2.867 | 10.703 |
| | 14.837 | 2.089 | 27.630 | -14.605 | -7.683 | -22.959 | 17.964 | 10.652 | 21.503 |
| | -16.129 | 25.217 | 1.016 | 18.383 | -11.344 | -16.761 | 16.310 | -3.626 | 16.130 |
| | 20.133 | 17.208 | -22.077 | -5.709 | 5.000 | -15.374 | 0.337 | 9.383 | 18.614 |
| | 25.148 | -4.649 | 22.512 | 0.164 | -3.098 | 27.432 | 18.692 | 14.769 | 1.977 |
| | 26.688 | 1.279 | 16.990 | -13.955 | -0.130 | 12.678 | 15.824 | 18.792 | 4.741 |
| | -21.790 | 22.474 | -20.412 | 19.051 | 13.296 | -24.781 | 9.930 | -2.570 | -13.814 |

| | | | | | | | | | |
|---|---|---|---|---|---|---|---|---|---|
| | 25.863 | 5.176 | 1.680 | -9.849 | 10.872 | -27.307 | 6.352 | -10.252 | 19.620 |
| | -26.130 | -18.465 | 18.404 | -8.226 | 2.865 | -23.556 | 10.112 | 21.609 | 0.468 |
| | -3.795 | -21.297 | -27.085 | 9.417 | 27.927 | 14.210 | -4.030 | 19.829 | 9.812 |
| | -24.450 | -6.991 | -12.086 | 27.048 | -18.239 | 22.457 | 0.420 | 22.002 | 20.812 |
| | -1.562 | -11.990 | -23.597 | 12.665 | -5.365 | -14.271 | 6.730 | 15.415 | 26.024 |
| | 9.652 | 26.757 | 0.568 | 11.320 | -12.197 | 21.035 | 13.416 | 8.243 | 7.098 |
| | -22.166 | 14.545 | 7.683 | -17.715 | 19.414 | 4.799 | 2.591 | 25.731 | 18.598 |
| | 19.257 | 14.838 | 22.640 | -11.900 | 25.253 | 17.030 | 15.819 | 7.004 | -5.641 |
| | -14.928 | 23.011 | 25.363 | 5.666 | -16.706 | -2.632 | -18.355 | 8.509 | 15.002 |
| | -3.007 | 15.952 | -26.684 | -1.384 | 18.051 | -17.250 | 20.154 | 5.955 | -10.322 |
| | -6.872 | 6.861 | -25.586 | 3.430 | -7.287 | 28.225 | 8.470 | -16.833 | 19.924 |
| | -13.311 | -5.008 | 19.668 | -11.473 | 24.508 | 3.661 | -4.133 | 24.828 | 23.305 |
| | -16.043 | 29.240 | 22.416 | 27.378 | 13.480 | -15.948 | 11.275 | 19.711 | 14.268 |
| | 23.302 | -13.366 | -22.081 | 17.322 | -20.535 | 27.439 | 10.758 | 6.477 | 6.119 |
| | 22.890 | 0.939 | 21.680 | -1.340 | 24.065 | -1.068 | -2.213 | -4.431 | 16.072 |
| | 12.907 | 21.246 | -16.860 | -9.166 | 22.718 | -8.386 | 4.446 | 7.508 | 21.122 |
| | -13.816 | -3.189 | -11.335 | 8.659 | 0.942 | 26.193 | 2.751 | -1.629 | 19.244 |
| | -26.015 | -8.182 | -22.590 | -10.933 | 11.337 | 12.863 | -9.512 | 11.459 | -0.984 |
| | 0.540 | -15.992 | 10.362 | -20.114 | -0.911 | 10.432 | 18.016 | 15.692 | 20.563 |
| | -24.161 | -26.175 | -3.295 | -9.992 | -12.864 | 8.725 | 18.855 | 4.612 | 7.610 |
| | -25.174 | 25.003 | -14.002 | 23.493 | 4.789 | -2.164 | 8.651 | -13.136 | 3.317 |
| | -12.833 | -6.344 | -17.833 | -16.821 | -7.782 | 22.646 | -2.369 | 21.029 | 11.123 |
| | -14.617 | 20.249 | -18.560 | 0.921 | -27.679 | 0.918 | 6.218 | 17.653 | 5.053 |
| | -17.460 | 3.622 | 26.137 | 26.513 | 11.471 | 0.461 | -1.702 | 30.067 | 8.149 |
| | -22.943 | 13.072 | -12.728 | 3.274 | 7.791 | -8.135 | -6.649 | 7.257 | 14.090 |
| | -16.762 | 23.076 | -23.177 | 15.492 | -11.895 | 23.560 | -8.713 | 0.292 | 25.115 |
| | -24.420 | 19.657 | -9.170 | -5.559 | -10.437 | 0.273 | 9.650 | 23.829 | -1.432 |
| | -23.120 | 4.446 | 13.281 | 19.717 | -1.187 | -18.534 | -19.795 | 10.813 | 16.865 |
| | 8.668 | -1.543 | 21.189 | -6.733 | 20.614 | -14.150 | 20.340 | 25.430 | 18.047 |
| | 18.642 | 15.948 | 14.737 | -22.174 | 10.272 | -11.407 | 3.103 | -22.350 | 1.780 |
| | 8.132 | -12.667 | 2.691 | 7.061 | -1.892 | 11.669 | 1.142 | -12.347 | 8.939 |
| | -17.166 | -7.899 | -4.842 | 24.528 | 5.589 | -13.748 | 14.358 | 2.239 | 7.018 |
| | -18.230 | -4.781 | -10.197 | 2.786 | -9.097 | -25.044 | -2.688 | 22.636 | 15.457 |
| | -10.272 | 5.208 | -16.576 | 1.910 | 1.629 | -29.337 | -7.733 | 16.405 | 9.832 |
| | 2.018 | 26.582 | -1.144 | 11.509 | -14.053 | 27.401 | 4.481 | 18.886 | 5.034 |
| | -17.261 | -14.862 | -23.461 | 17.918 | -16.286 | 12.631 | 7.795 | 19.711 | -11.009 |

| | | | | | | | | | |
|---|---|---|---|---|---|---|---|---|---|
| | -8.028 | -9.419 | -8.554 | 0.560 | 24.971 | -27.353 | 16.462 | 9.836 | 21.653 |
| | 5.687 | -21.915 | 19.314 | -2.164 | 22.534 | 20.322 | 12.259 | 24.707 | 5.700 |
| | 2.513 | 8.414 | -3.361 | 10.697 | 17.530 | -5.192 | 8.409 | 11.240 | 18.403 |
| | 9.000 | 10.102 | -24.622 | -22.652 | -19.389 | -21.380 | -10.087 | 22.373 | 20.671 |
| | -13.674 | 22.744 | 14.222 | -23.538 | -8.641 | 22.198 | -7.124 | -28.210 | 23.791 |
| | -21.998 | -2.507 | -11.136 | -3.637 | 8.077 | 3.120 | 10.120 | 11.993 | 0.300 |
| | 21.598 | -29.750 | 5.470 | 12.195 | -27.657 | 24.355 | -3.184 | 3.089 | 15.529 |
| | -15.829 | -4.519 | -23.690 | -0.347 | -15.905 | 30.337 | -21.532 | 15.804 | -5.090 |
| | -12.120 | -19.547 | 0.432 | -6.012 | 16.210 | -12.236 | -12.800 | 12.993 | 2.444 |
| | 18.748 | -25.001 | 12.818 | 16.684 | -0.878 | 14.329 | -16.221 | 8.987 | -0.656 |
| | -8.162 | 19.595 | -9.176 | 21.273 | -12.960 | 26.288 | 0.141 | 7.326 | 9.320 |
| | 17.507 | 25.980 | -23.259 | 20.265 | 24.853 | -28.740 | 5.243 | -9.233 | -2.807 |
| | -20.751 | -11.868 | -14.424 | 18.734 | 15.187 | 19.394 | 21.502 | 14.429 | 11.203 |
| | 11.010 | -5.865 | -16.680 | 23.398 | -10.893 | -13.305 | -9.974 | 1.426 | 18.432 |
| | -4.293 | -13.086 | -2.196 | 5.793 | 20.808 | 19.650 | 17.103 | 24.167 | 17.000 |
| | -19.178 | 30.151 | -7.945 | -15.174 | 15.415 | 21.014 | -2.671 | -4.950 | -5.392 |
| | 23.861 | -18.286 | -23.218 | 0.059 | -23.780 | -29.812 | -4.325 | 26.041 | -1.068 |
| | -12.472 | -8.101 | 2.680 | 25.415 | -9.815 | -7.280 | 8.975 | 6.768 | 17.510 |
| | -19.805 | 12.865 | -8.393 | -21.214 | 19.285 | 22.851 | -7.942 | 19.839 | -5.209 |
| | -17.286 | 15.175 | 24.018 | -10.976 | 3.880 | 23.735 | 4.188 | 5.132 | 12.258 |
| | -15.590 | -0.256 | -17.303 | -9.755 | -28.533 | -10.567 | 22.134 | 28.164 | -6.551 |
| | -0.530 | 18.048 | -4.572 | 8.289 | -0.148 | 11.957 | 2.906 | -9.399 | 16.874 |
| | 24.973 | -11.147 | -19.275 | 15.219 | -7.573 | 10.000 | -17.919 | 16.782 | 6.847 |
| | 21.207 | 29.135 | -0.156 | -22.606 | 23.449 | 11.336 | 1.036 | 23.115 | 1.434 |
| | 17.688 | 16.148 | 18.574 | 12.718 | 13.213 | 14.373 | 14.987 | 27.226 | 8.953 |
| | 0.582 | -2.326 | -8.055 | 25.138 | -5.169 | -18.939 | 17.409 | 19.161 | 12.448 |
| | 26.727 | 26.798 | -24.863 | 1.753 | 29.857 | 30.909 | -6.255 | 22.102 | -20.972 |
| | -25.644 | -12.177 | -13.846 | 17.952 | 28.021 | -19.794 | -7.846 | -1.353 | 1.334 |
| | -3.575 | 22.931 | -24.238 | 7.598 | -22.905 | -25.984 | 13.001 | 8.817 | 11.989 |
| | -27.014 | 9.777 | -10.085 | 10.355 | -10.656 | 25.985 | -3.100 | 3.361 | -12.191 |
| | 3.182 | -3.166 | 4.777 | -12.463 | -4.232 | -12.888 | 11.757 | -2.217 | 13.509 |
| | -24.264 | 17.491 | 7.110 | -20.102 | -14.485 | -27.183 | -1.777 | 27.409 | 7.571 |
| | 6.450 | -21.005 | -5.905 | 17.537 | 21.542 | 25.034 | 13.983 | 5.793 | 10.053 |
| | -7.021 | -27.178 | -6.813 | -16.044 | 1.627 | 11.486 | 25.311 | 27.055 | 16.179 |
| | 2.015 | 28.372 | -12.490 | 19.686 | -21.989 | -9.383 | -0.553 | 12.225 | 12.442 |
| | 19.587 | 10.234 | -18.161 | -3.146 | -12.892 | 25.569 | 20.173 | 24.793 | 4.914 |

| | | | | | | | | | |
|---|---|---|---|---|---|---|---|---|---|
| | -6.871 | -24.596 | -23.141 | -23.280 | 8.023 | 4.788 | 3.915 | 28.768 | -1.479 |
| | -16.540 | -23.380 | -18.580 | -17.269 | 8.658 | -22.959 | -24.571 | -6.548 | 17.793 |
| | -12.698 | -22.240 | -22.241 | 2.699 | 9.648 | -19.175 | 17.941 | 20.873 | -1.763 |
| | -13.260 | 27.053 | 5.827 | 28.119 | -30.650 | 29.636 | -9.970 | 10.224 | 8.828 |
| | 22.254 | 22.340 | -9.085 | 9.564 | 17.639 | -25.169 | -4.615 | 3.208 | 3.076 |
| | 3.123 | 13.497 | 0.812 | 6.541 | 0.827 | 16.207 | 2.118 | 6.309 | 22.880 |
| | -22.826 | -22.909 | 26.685 | 15.438 | 10.013 | 27.744 | 19.099 | 2.369 | 10.552 |
| | 14.780 | -19.602 | 23.390 | 14.893 | -22.524 | 22.625 | 5.531 | -0.627 | 2.391 |
| | 10.639 | 17.201 | 8.265 | -1.673 | 6.557 | 17.920 | 7.154 | 19.485 | 20.303 |
| | -2.917 | 15.150 | -25.005 | -1.245 | 26.971 | -6.334 | 12.421 | 23.275 | -3.884 |
| | 19.987 | 26.324 | -15.408 | 22.229 | 22.709 | -30.428 | 9.942 | 18.049 | 15.023 |
| | -17.109 | -7.557 | -12.955 | -2.526 | -9.548 | -16.780 | 8.778 | 13.618 | 26.834 |
| | 26.930 | 6.230 | -2.025 | 1.483 | 8.434 | -27.613 | 8.387 | -5.357 | 18.063 |
| | -14.169 | 11.652 | -5.826 | 23.381 | -12.702 | 24.377 | -9.000 | 2.091 | 16.597 |
| | -26.693 | 10.614 | -3.723 | 5.726 | -7.138 | 6.157 | 13.190 | 9.747 | 22.083 |
| | 17.801 | 24.663 | -9.053 | -22.378 | -31.111 | -5.321 | 7.347 | -2.678 | 8.235 |
| | -21.448 | 20.608 | -6.624 | -24.693 | 0.233 | 27.415 | 26.516 | 3.959 | 10.253 |
| | -3.863 | 14.418 | -24.480 | -6.152 | -21.309 | 6.306 | -4.343 | 7.841 | -6.100 |
| | -15.262 | -14.805 | 5.981 | 19.173 | -24.769 | 23.649 | 11.426 | -15.138 | 6.075 |
| | -20.742 | 12.335 | 7.224 | 0.297 | 14.325 | -1.302 | -14.450 | 12.269 | 25.839 |
| | -10.478 | 5.054 | -18.375 | -11.164 | -11.059 | 0.575 | 24.070 | 14.524 | 14.269 |
| | 8.243 | 3.138 | 3.423 | 26.450 | 6.137 | 25.612 | 22.398 | 25.597 | -10.522 |
| | -8.973 | 23.222 | 19.751 | 4.444 | -3.927 | -13.523 | 6.800 | -20.792 | 13.435 |
| | 6.558 | -15.443 | 25.584 | 29.331 | -22.826 | 24.706 | 12.879 | 15.674 | 1.507 |
| | 21.021 | -20.646 | -16.635 | -24.038 | 30.710 | 6.908 | -9.082 | 3.721 | 11.631 |
| | -24.777 | 22.357 | -5.949 | -7.792 | -4.516 | -23.682 | 2.773 | 0.270 | -4.853 |
| | -2.863 | -6.996 | -1.211 | -17.514 | 17.397 | 28.453 | 25.450 | 22.500 | 12.582 |
| | -6.408 | 5.533 | 27.414 | 26.846 | -5.596 | -26.292 | 5.992 | 15.583 | 18.862 |
| | -8.521 | -24.937 | 8.299 | -11.023 | -19.014 | 22.337 | 13.362 | 1.733 | -10.001 |
| | -5.472 | -4.562 | -15.362 | 1.513 | -14.735 | 28.480 | 17.772 | 9.862 | -1.703 |
| | 7.415 | 11.064 | -9.554 | 24.562 | -0.506 | 20.542 | 8.932 | -4.007 | -12.117 |
| | 23.015 | -2.176 | -22.307 | -4.069 | -31.709 | -24.537 | 21.835 | 18.969 | 10.975 |
| | -26.700 | 25.991 | -0.076 | 16.010 | 13.495 | 14.571 | 8.082 | -0.944 | -16.645 |
| | 11.688 | -12.554 | -5.435 | 8.716 | -10.454 | -21.579 | -7.872 | -3.118 | 24.064 |
| | -3.538 | 21.367 | 16.959 | 17.813 | 27.136 | -26.523 | -10.786 | 21.553 | 9.078 |
| | 4.496 | -8.417 | 18.217 | 27.116 | 14.529 | 26.646 | -23.442 | 7.482 | 17.727 |

| | | | | | | | | | |
|---|---|---|---|---|---|---|---|---|---|
| | -4.695 | -21.766 | -11.470 | -17.165 | -17.649 | -7.990 | 13.838 | -7.440 | 12.546 |
| | 27.421 | 26.496 | -25.006 | 3.948 | 10.676 | -7.448 | -12.457 | 19.960 | 4.819 |
| | 16.714 | 3.806 | -25.809 | -26.649 | 2.736 | -32.094 | 13.068 | 7.840 | -21.463 |
| | 11.896 | 7.767 | -5.250 | -23.043 | 29.623 | 22.989 | 15.951 | 24.929 | 18.937 |
| | -19.564 | 5.917 | 18.430 | -24.172 | -16.365 | -12.528 | 9.689 | 24.413 | 11.672 |
| | -7.099 | 20.091 | 10.850 | 13.287 | -8.284 | 30.668 | 22.529 | 24.975 | 22.755 |
| | -6.459 | 15.265 | -11.501 | 16.531 | 16.890 | 1.489 | 6.960 | 10.413 | 2.612 |
| | -22.197 | -13.613 | 26.936 | -7.307 | 4.472 | 13.265 | 23.735 | 1.040 | 27.793 |
| | 19.564 | -3.505 | 1.612 | -16.357 | -0.843 | 24.959 | 21.668 | 25.201 | 18.221 |
| | -18.381 | -15.052 | 15.254 | 14.407 | 30.855 | 19.965 | -25.372 | 8.820 | 12.681 |
| | -23.144 | -10.521 | -18.283 | 30.941 | 24.851 | 26.075 | 6.253 | 22.183 | 20.061 |
| | -25.336 | -22.468 | -23.522 | 8.812 | -29.947 | 5.217 | -15.038 | 0.287 | 8.007 |
| | 23.105 | 5.448 | 27.913 | -2.702 | 13.046 | -30.118 | 11.665 | 23.612 | 23.025 |
| | -20.525 | 13.078 | -13.131 | 0.379 | -19.928 | 3.551 | 9.030 | -2.200 | -1.108 |
| | -10.399 | 4.348 | 7.961 | -9.615 | -5.637 | -11.871 | 24.017 | 22.669 | 2.118 |
| | 23.846 | -4.920 | -15.336 | 21.930 | 1.921 | 9.873 | 16.920 | -6.052 | 22.369 |
| | 4.215 | 22.726 | 19.147 | 4.702 | -17.294 | 25.439 | 7.775 | 13.655 | 8.779 |
| | -28.377 | 21.057 | 18.902 | -23.506 | -26.745 | 19.471 | 12.571 | 3.621 | 15.483 |
| | 10.343 | -23.649 | -16.671 | -14.428 | 23.160 | 15.243 | 26.956 | 21.830 | 11.686 |
| | 14.663 | 14.198 | 5.963 | 15.899 | 26.986 | 21.545 | 19.774 | -4.050 | 20.220 |
| | -11.848 | -7.270 | 4.049 | -27.793 | -3.366 | 17.206 | 25.830 | -4.393 | 24.337 |
| | -5.883 | 2.452 | -12.267 | 25.405 | 15.459 | -26.160 | -3.190 | 26.518 | 10.140 |
| | -19.808 | 0.675 | 19.263 | 28.761 | -7.800 | 22.550 | 21.697 | 18.154 | 26.164 |
| | -16.522 | 18.908 | -5.757 | 0.213 | -28.160 | 20.878 | 15.587 | -13.395 | 6.203 |
| | 10.602 | 20.281 | -10.355 | -21.877 | -12.938 | 8.922 | 17.277 | 14.210 | 23.403 |
| | 1.077 | -12.662 | -18.546 | -28.552 | -9.161 | -20.879 | 13.629 | 6.525 | 8.992 |
| | 0.233 | -3.260 | -27.770 | -22.737 | 28.177 | -24.514 | 29.197 | -25.079 | 14.272 |
| | -24.584 | 21.498 | 20.183 | -3.140 | -6.788 | -27.971 | -3.383 | -0.219 | -16.432 |
| | -27.986 | -8.010 | -20.726 | 23.485 | -15.361 | 27.592 | -13.164 | 19.913 | 26.049 |
| | -15.887 | -0.865 | 16.860 | 11.150 | 13.024 | -11.406 | -1.469 | -12.489 | 15.815 |
| | -19.068 | 18.869 | -14.283 | 24.732 | -1.213 | -5.708 | 27.113 | 11.029 | 11.631 |
| | 7.895 | 11.109 | -4.667 | 16.369 | -21.133 | 20.765 | 28.260 | 22.165 | 20.027 |
| | -11.476 | -15.617 | 17.934 | 0.319 | 30.096 | -21.561 | 19.854 | 20.908 | 12.934 |
| | 28.809 | 24.745 | 22.948 | -3.585 | -26.766 | -16.517 | 14.674 | 25.175 | 10.677 |
| | 23.398 | -21.144 | -17.108 | 8.731 | -23.521 | -30.847 | 10.321 | 3.939 | 22.161 |
| | 19.523 | -2.008 | 22.050 | -18.863 | -18.076 | 25.488 | -4.934 | -4.356 | 9.426 |

| | | | | | | | | | |
|---|---|---|---|---|---|---|---|---|---|
| | -15.164 | -10.683 | -12.141 | 28.022 | 11.629 | 19.922 | 19.180 | 9.605 | -8.342 |
| | -7.520 | 5.903 | 27.059 | -21.623 | -24.301 | 24.782 | -2.338 | 11.368 | 28.219 |
| | -10.217 | 2.034 | -11.165 | 0.823 | 27.680 | 4.466 | 26.367 | 8.119 | 13.744 |
| | 10.437 | 4.970 | 24.665 | -2.273 | 21.929 | 26.301 | 7.039 | 0.347 | 19.058 |
| | -17.854 | -4.065 | -16.006 | -16.919 | 0.093 | 16.944 | -1.840 | -8.980 | 14.491 |
| | 26.310 | 14.864 | -29.848 | -11.081 | -12.411 | -10.363 | -27.465 | 18.940 | 17.499 |
| | -21.469 | 16.106 | -13.356 | -14.469 | -5.159 | -30.218 | 13.405 | 5.069 | -14.224 |
| | -12.164 | 7.271 | 23.648 | -13.087 | -25.630 | 10.373 | -0.689 | -2.711 | 25.789 |
| | -19.818 | -7.524 | 10.473 | 23.420 | -11.700 | -14.079 | 8.738 | -3.442 | 15.562 |
| | 20.057 | -9.013 | -30.373 | -28.059 | 28.325 | -13.400 | -29.424 | 6.946 | -3.450 |
| | -26.214 | 23.100 | -7.126 | -25.400 | -17.800 | -24.887 | 4.687 | 18.189 | 13.621 |
| | 3.613 | 12.630 | -9.191 | 26.903 | -7.388 | -8.236 | 12.891 | -4.290 | 18.395 |
| | -27.744 | -0.072 | 11.161 | 14.797 | -9.703 | -19.993 | 21.545 | 21.420 | -0.063 |
| | 23.793 | -20.850 | -16.373 | -21.757 | 19.352 | 13.878 | -6.252 | 11.439 | 12.836 |
| | 9.404 | -2.941 | -1.911 | 5.825 | -14.274 | -29.866 | 7.467 | 22.827 | 1.825 |
| | 5.036 | -15.883 | 8.717 | -11.323 | -10.364 | -0.075 | 10.762 | 14.785 | 24.970 |
| | 18.490 | -12.426 | -22.257 | 9.068 | 4.308 | 27.361 | -17.088 | 19.403 | 22.716 |
| | -10.273 | 13.804 | 23.915 | 8.682 | 4.907 | -11.526 | 24.639 | 4.152 | 14.225 |
| | -5.658 | 4.865 | -26.241 | -26.579 | 0.706 | 28.057 | -0.453 | -4.600 | 13.437 |
| | -2.821 | 24.679 | 26.506 | -7.719 | -2.419 | -20.533 | 24.100 | 2.238 | -7.787 |
| | 24.307 | 11.622 | -14.027 | -20.890 | -6.228 | 4.652 | -2.828 | -0.378 | -13.229 |
| | -7.666 | 8.592 | 17.293 | 26.229 | -9.269 | -19.911 | 6.490 | 3.291 | 24.512 |
| | 21.827 | -23.286 | 3.619 | 22.984 | -11.804 | 14.085 | 16.704 | 7.522 | 20.812 |
| | -14.336 | -25.382 | 24.106 | -23.688 | -3.735 | -27.587 | 17.738 | 6.211 | 17.748 |
| | 27.129 | -21.570 | -8.428 | 26.743 | 17.374 | 23.074 | 12.242 | 9.921 | 21.962 |
| | -1.255 | 24.014 | 12.437 | -9.918 | 18.518 | -5.128 | 13.954 | 2.015 | 26.099 |
| | 7.327 | -5.653 | 21.829 | -7.883 | -7.210 | -10.466 | 0.741 | -4.841 | -22.591 |
| | -22.356 | -11.489 | -0.301 | 21.567 | 25.420 | 3.129 | 30.312 | | -19.314 |
| | -30.844 | -2.555 | -22.536 | 10.224 | 3.442 | 8.170 | -15.617 | | 16.329 |
| | 25.650 | 15.419 | -28.313 | -25.215 | -21.928 | 4.969 | 22.405 | | 11.175 |
| | 15.749 | 16.091 | -5.949 | 6.743 | 15.855 | 27.957 | 25.170 | | -0.939 |
| | -4.604 | -20.622 | -14.858 | 25.305 | -27.356 | 27.038 | 27.165 | | 0.187 |
| | -9.212 | -4.866 | -3.620 | -15.017 | -12.170 | -24.844 | 20.480 | | 8.821 |
| | 23.144 | -17.702 | -2.637 | 7.314 | 7.694 | 4.785 | 8.597 | | 24.435 |
| | 19.771 | 17.880 | 7.263 | -4.888 | 18.955 | -19.082 | 15.627 | | 23.292 |
| | 9.405 | 4.789 | -1.886 | 1.403 | 10.882 | 24.249 | 2.965 | | 14.998 |

| | | | | | | | | | |
|---|---|---|---|---|---|---|---|---|---|
| | 26.492 | -22.872 | -29.154 | 1.979 | -4.444 | 15.194 | 18.478 | | 9.663 |
| | 26.067 | 4.859 | 10.570 | -5.796 | -26.100 | -19.425 | 23.626 | | 10.575 |
| | -5.488 | -0.933 | -21.259 | -0.103 | 6.803 | 12.858 | 26.661 | | -0.460 |
| | -7.385 | 19.142 | -6.935 | 18.886 | -27.892 | 8.434 | -5.493 | | 19.686 |
| | -23.886 | -26.113 | -13.851 | -7.149 | 19.426 | -20.376 | 11.375 | | 27.955 |
| | -26.278 | -1.659 | 21.348 | 21.280 | -5.724 | -20.916 | 9.527 | | 2.912 |
| | 18.883 | -17.218 | -0.303 | -0.826 | -11.054 | 25.959 | 21.461 | | -4.530 |
| | 28.949 | -15.446 | 26.921 | 14.312 | 27.381 | 1.971 | 16.764 | | 18.491 |
| | 25.768 | -8.579 | 24.791 | -2.478 | 9.979 | -17.083 | -2.572 | | 27.121 |
| | 9.723 | 27.068 | -10.435 | 5.197 | 22.256 | -7.149 | -8.925 | | 24.983 |
| | -23.954 | 23.425 | -4.244 | 2.502 | -14.669 | 4.044 | -5.941 | | 14.371 |
| | -17.044 | -18.099 | -6.191 | 1.793 | -14.083 | -26.308 | 25.373 | | 12.134 |
| | 5.904 | -2.374 | 8.465 | | -24.952 | 5.005 | -15.851 | | 24.017 |
| | -7.219 | -3.696 | 23.928 | | -9.169 | 9.850 | 4.710 | | -2.237 |
| | -30.157 | 15.038 | -28.073 | | 17.257 | -8.227 | 10.832 | | 27.463 |
| | -17.725 | 3.404 | 19.608 | | -20.149 | 19.860 | -2.930 | | 9.112 |
| | -11.044 | 24.766 | 3.548 | | -2.495 | 10.942 | -11.936 | | 0.406 |
| | 29.130 | -12.114 | -14.430 | | 1.815 | -18.838 | 0.835 | | 13.073 |
| | 21.113 | -11.101 | 13.472 | | 18.335 | 22.087 | 2.373 | | 15.688 |
| | -31.730 | -3.085 | -19.148 | | 21.768 | -22.774 | 1.306 | | 11.508 |
| | -21.212 | -9.044 | 6.962 | | -19.461 | -24.109 | -1.296 | | -4.439 |
| | -20.005 | -6.375 | 11.338 | | -22.480 | -3.737 | -4.719 | | 10.451 |
| | 28.691 | -19.217 | -20.889 | | -2.281 | -26.793 | 17.952 | | 8.898 |
| | 13.926 | -18.736 | 2.752 | | -23.243 | -10.596 | 5.691 | | 5.189 |
| | -26.736 | -14.548 | -27.940 | | 0.175 | -4.479 | -22.106 | | 6.862 |
| | 15.374 | -9.776 | 22.029 | | -26.325 | -24.995 | -10.044 | | 0.034 |
| | -28.013 | -22.216 | -3.159 | | -21.018 | 18.224 | 22.460 | | -25.315 |
| | 27.603 | -24.527 | -7.271 | | -22.440 | 23.889 | 2.998 | | 22.650 |
| | 14.485 | -4.633 | -30.026 | | -4.017 | 6.690 | -23.110 | | 1.827 |
| | -13.266 | 25.744 | -5.713 | | -14.813 | -19.731 | 14.316 | | 7.587 |
| | 25.534 | 17.933 | 0.965 | | -7.377 | 4.497 | -0.673 | | 30.318 |
| | -5.008 | 12.553 | -19.898 | | 9.148 | 3.676 | 28.842 | | 18.028 |
| | 4.978 | -27.671 | 22.633 | | 1.523 | 23.690 | 2.501 | | 3.340 |
| | -20.485 | -17.983 | 5.850 | | -2.289 | 2.720 | 9.974 | | -19.983 |
| | -31.701 | -9.354 | 18.269 | | -12.655 | 0.514 | 20.330 | | 6.638 |
| | 16.929 | 9.806 | 17.178 | | -3.550 | -7.019 | 6.337 | | 26.845 |

| | | | | | | | | | |
|---|---|---|---|---|---|---|---|---|---|
| | -30.386 | -10.617 | -8.334 | | 26.136 | -3.249 | 16.750 | | -8.516 |
| | -23.084 | -7.957 | 14.287 | | 17.051 | -14.319 | -15.941 | | 28.991 |
| | 0.295 | -23.180 | -2.043 | | -21.066 | -21.466 | -13.787 | | 9.531 |
| | -6.986 | 27.492 | 25.371 | | 23.289 | -16.570 | -22.871 | | 17.473 |
| | 30.235 | -6.416 | -8.558 | | 25.838 | -0.355 | 27.974 | | 20.063 |
| | -29.120 | 11.748 | 7.222 | | -25.688 | -5.810 | -4.535 | | 1.929 |
| | 31.223 | 25.177 | -4.963 | | -19.952 | -25.674 | -10.973 | | 26.348 |
| | 22.141 | 13.450 | 11.213 | | 4.950 | 1.249 | 25.246 | | 27.839 |
| | -5.850 | 26.457 | 24.854 | | 13.744 | -2.356 | 18.773 | | 11.554 |
| | -22.378 | -1.034 | -24.584 | | 21.543 | -20.955 | -6.506 | | 22.092 |
| | 25.177 | 17.663 | -16.304 | | -12.478 | -10.408 | 28.224 | | 16.114 |
| | 4.107 | -28.512 | 8.810 | | 13.649 | -17.745 | 4.966 | | 18.297 |
| | 14.948 | 23.011 | 11.937 | | -22.832 | -14.536 | -2.683 | | 25.114 |
| | 1.387 | -25.058 | 28.529 | | -18.208 | -21.203 | -8.953 | | 5.620 |
| | -3.995 | -25.967 | -11.453 | | -14.190 | -23.309 | -5.866 | | -2.780 |
| | -7.983 | -7.443 | -14.152 | | -19.805 | -23.943 | -7.533 | | 28.920 |
| | -13.534 | -22.812 | 3.224 | | -8.106 | -24.370 | 16.210 | | 1.598 |
| | -15.033 | 7.814 | -21.148 | | 13.416 | | 11.741 | | -4.047 |
| | -30.802 | 7.315 | 24.303 | | 5.418 | | -19.341 | | 9.173 |
| | 1.878 | -23.612 | 27.909 | | -8.789 | | 19.296 | | 7.944 |
| | -16.536 | 26.949 | 23.404 | | -25.828 | | 30.279 | | 28.253 |
| | 29.507 | -8.051 | 7.712 | | 19.253 | | 8.387 | | 11.031 |
| | -19.398 | -16.608 | 26.965 | | -0.286 | | -13.990 | | 9.280 |
| | 20.206 | -5.858 | -22.635 | | 13.775 | | 20.690 | | 14.610 |
| | 30.869 | -6.626 | -19.386 | | -8.489 | | -27.587 | | -5.993 |
| | -29.794 | 9.814 | -16.453 | | 6.855 | | 16.034 | | 5.933 |
| | -18.964 | 3.338 | 8.243 | | 25.490 | | 22.465 | | 24.431 |
| | -17.369 | -12.547 | -7.023 | | -17.605 | | 15.328 | | 17.852 |
| | 24.352 | -5.088 | 28.826 | | 0.865 | | -3.447 | | -19.599 |
| | -10.024 | -18.079 | -27.280 | | -11.010 | | 14.189 | | -30.114 |
| | -13.343 | -7.277 | -20.058 | | 16.292 | | 4.190 | | 15.687 |
| | 5.095 | 13.336 | 30.057 | | -10.319 | | 10.924 | | -2.279 |
| | 0.073 | -3.983 | -17.005 | | -23.252 | | 12.168 | | 17.193 |
| | 5.956 | 6.530 | 17.461 | | -1.715 | | 5.169 | | 25.785 |
| | -11.887 | -6.067 | 8.773 | | -2.429 | | 13.274 | | 24.224 |
| | -16.795 | -11.622 | 11.711 | | -14.932 | | -5.644 | | 10.242 |

| | | | | | | | | | |
|---|---|---|---|---|---|---|---|---|---|
| | -28.518 | -13.344 | -14.133 | | 13.176 | | -8.374 | | 1.269 |
| | 13.731 | -9.602 | -22.993 | | 13.881 | | 16.778 | | 18.758 |
| | 10.173 | 26.526 | 11.284 | | 12.324 | | -7.215 | | 9.511 |
| | -17.890 | 5.779 | 9.853 | | 10.139 | | 24.550 | | 27.397 |
| | 27.102 | 18.643 | 25.911 | | 22.052 | | 16.123 | | 5.410 |
| | 18.466 | -26.012 | -12.149 | | -0.406 | | 7.984 | | 14.713 |
| | 3.883 | 25.227 | 6.548 | | -6.280 | | 21.615 | | -4.675 |
| | -1.691 | 23.652 | -25.413 | | -16.358 | | 6.486 | | 8.263 |
| | -9.432 | -13.744 | -8.556 | | 5.465 | | -4.820 | | 22.206 |
| | -26.518 | -4.652 | 3.500 | | 20.225 | | 26.549 | | 4.945 |
| | 23.137 | -7.306 | -18.122 | | 19.377 | | 11.264 | | 24.068 |
| | 19.597 | 3.641 | 21.477 | | 21.582 | | 8.076 | | 27.977 |
| | -21.940 | -15.515 | -1.638 | | 7.872 | | 26.832 | | 21.554 |
| | 26.672 | -16.657 | -22.536 | | -2.593 | | 27.860 | | 23.391 |
| | -23.716 | -28.083 | -6.814 | | -25.452 | | 21.693 | | 7.706 |
| | -27.713 | -18.551 | -8.032 | | -7.374 | | 10.358 | | 21.096 |
| | -26.533 | -26.730 | -9.868 | | -24.034 | | -10.201 | | 16.626 |
| | 29.565 | 29.048 | -22.941 | | 11.964 | | 17.204 | | 2.319 |
| | 21.197 | 4.918 | 19.316 | | -22.008 | | -2.452 | | 19.371 |
| | 10.453 | 14.588 | -17.023 | | -26.773 | | -23.276 | | 15.958 |
| | 2.262 | 4.478 | 12.946 | | -1.976 | | -16.670 | | 13.295 |
| | -24.593 | 6.843 | 8.654 | | 11.679 | | 19.018 | | 27.218 |
| | -5.282 | -20.373 | -23.960 | | 25.094 | | -8.934 | | 23.625 |
| | 26.779 | -3.121 | -4.572 | | 19.522 | | 28.372 | | 22.084 |
| | -11.622 | -4.143 | -20.908 | | 3.112 | | -6.726 | | -3.192 |
| | 7.889 | 21.921 | -3.897 | | 4.093 | | 12.987 | | 27.828 |
| | 26.984 | -12.843 | -19.045 | | -12.940 | | 13.669 | | -0.235 |
| | 13.143 | -14.049 | -2.511 | | -16.651 | | 10.753 | | 24.309 |
| | 4.295 | -14.394 | 3.066 | | 2.032 | | 2.561 | | 26.731 |
| | 20.795 | -19.142 | -13.641 | | -26.382 | | 1.091 | | 14.964 |
| | -3.110 | 8.935 | 9.387 | | 13.066 | | -7.628 | | 19.240 |
| | -28.894 | 27.920 | 21.679 | | -22.056 | | 0.623 | | 21.406 |
| | -29.895 | -18.089 | -10.359 | | 18.161 | | 9.015 | | 7.398 |
| | -20.214 | 14.373 | 4.316 | | -9.359 | | 1.442 | | 26.743 |
| | 24.936 | 20.871 | -27.309 | | 26.156 | | -4.338 | | 5.551 |
| | 25.982 | 15.748 | 0.622 | | 24.133 | | -20.429 | | 26.976 |

| | | | | | | | | | |
|---|---|---|---|---|---|---|---|---|---|
| | -13.967 | 1.324 | 7.321 | | 20.991 | | -0.141 | | 9.065 |
| | 22.083 | 22.950 | 8.742 | | 16.245 | | 23.376 | | -2.680 |
| | -11.853 | -29.815 | -9.654 | | -4.760 | | 24.816 | | 21.232 |
| | 0.980 | 18.789 | 23.922 | | -24.263 | | 26.900 | | 17.831 |
| | -2.580 | 0.074 | -15.389 | | 20.260 | | 18.766 | | 20.519 |
| | -20.023 | -10.897 | -4.404 | | -17.506 | | 2.010 | | 22.816 |
| | -18.150 | -20.853 | -21.796 | | 1.366 | | 7.326 | | 4.583 |
| | 25.108 | -21.921 | -5.433 | | -6.052 | | 3.104 | | 5.063 |
| | -4.850 | 19.923 | -14.251 | | -10.985 | | -2.768 | | 12.131 |
| | 16.427 | 18.239 | -0.504 | | 11.749 | | 16.551 | | 19.049 |
| | -16.965 | -2.780 | -6.971 | | -1.347 | | 9.595 | | 13.346 |
| | 16.042 | 16.524 | 4.687 | | 12.670 | | 25.644 | | 22.168 |
| | 20.561 | 17.851 | -16.468 | | 8.196 | | 24.089 | | 21.294 |
| | -8.896 | 31.242 | -22.152 | | 0.461 | | -5.861 | | -7.743 |
| | -25.273 | 16.940 | -5.624 | | -0.826 | | -1.956 | | 10.980 |
| | -18.844 | -30.605 | 9.128 | | -15.810 | | 5.947 | | 11.946 |
| | 19.640 | -25.436 | 21.633 | | 24.177 | | -23.763 | | 15.045 |
| | -22.129 | 0.248 | -11.002 | | 17.800 | | -0.960 | | 18.988 |
| | -14.648 | 9.251 | -3.852 | | -6.927 | | -4.159 | | 21.544 |
| | 25.330 | 3.525 | -24.012 | | 15.004 | | 8.263 | | 8.368 |
| | -23.553 | -19.575 | -25.599 | | 19.089 | | 26.121 | | 25.898 |
| | 16.790 | 24.224 | -7.697 | | 4.147 | | -3.876 | | 14.460 |
| | 16.392 | 27.500 | 22.247 | | -5.613 | | 1.241 | | 5.024 |
| | -15.784 | 18.603 | 16.386 | | -22.042 | | 3.375 | | 1.779 |
| | 11.565 | -22.543 | -17.792 | | -15.037 | | 21.448 | | -0.346 |
| | -5.213 | 12.996 | 17.113 | | 9.984 | | -16.960 | | 21.568 |
| | -26.502 | -13.197 | 11.201 | | 6.295 | | 11.717 | | 3.443 |
| | 20.270 | -17.093 | -0.761 | | -4.060 | | -2.864 | | -7.581 |
| | -3.508 | -14.706 | -3.480 | | 25.474 | | 12.479 | | 10.392 |
| | 0.684 | -16.414 | -7.147 | | 17.534 | | 16.382 | | 10.065 |
| | -14.197 | 26.281 | -4.403 | | 11.150 | | 5.429 | | 14.044 |
| | -15.011 | 11.931 | -12.291 | | 15.589 | | 10.248 | | 11.551 |
| | -19.066 | -8.388 | 9.221 | | 26.283 | | 2.529 | | -0.991 |
| | 20.249 | -21.489 | -15.095 | | 13.840 | | 24.534 | | 19.903 |
| | 12.171 | 19.926 | 23.042 | | 9.290 | | 9.124 | | 16.450 |
| | 21.672 | -4.757 | -4.079 | | 18.398 | | 12.927 | | 15.031 |

| | | | | | | | | | |
|---|---|---|---|---|---|---|---|---|---|
| | -13.685 | -27.215 | 24.929 | | 8.115 | | 15.417 | | 1.176 |
| | 12.076 | 4.526 | -17.812 | | 6.560 | | 0.834 | | 21.894 |
| | -25.064 | -21.345 | 1.538 | | -12.791 | | -1.795 | | 20.598 |
| | -18.962 | -20.391 | 0.915 | | 2.683 | | 2.864 | | -7.776 |
| | -3.019 | 19.446 | -20.337 | | -27.638 | | 20.510 | | -1.238 |
| | 27.144 | 28.306 | -10.222 | | 20.911 | | 18.789 | | |
| | -26.631 | -6.932 | 8.103 | | 19.912 | | -4.410 | | |
| | 22.558 | 18.203 | -12.655 | | -13.070 | | 4.884 | | |
| | -5.722 | -10.428 | 2.594 | | -1.385 | | 5.786 | | |
| | -18.758 | -17.118 | -9.672 | | -12.106 | | 16.907 | | |
| | -9.172 | -22.189 | 12.516 | | 14.775 | | -12.107 | | |
| | -19.928 | 8.267 | 1.475 | | 16.007 | | 15.462 | | |
| | -16.567 | -3.296 | -3.011 | | -7.492 | | -10.885 | | |
| | -0.554 | 30.310 | 14.402 | | -8.203 | | 23.313 | | |
| | -5.129 | 18.116 | -9.430 | | 5.597 | | 13.250 | | |
| | -12.177 | 13.407 | -5.404 | | 3.614 | | 7.816 | | |
| | 11.120 | -15.995 | 5.978 | | 8.889 | | 12.380 | | |
| | 20.591 | -21.333 | -0.367 | | 7.075 | | 10.356 | | |
| | -23.179 | -9.919 | | | -13.488 | | 17.326 | | |
| | 14.799 | 22.201 | | | 20.600 | | 25.878 | | |
| | 1.758 | -23.959 | | | 23.407 | | 2.711 | | |
| | 4.821 | -8.278 | | | 16.212 | | 19.157 | | |
| | -1.995 | 7.552 | | | -3.023 | | 25.012 | | |
| | -9.404 | 9.876 | | | -18.423 | | -1.941 | | |
| | -8.783 | 15.640 | | | -2.143 | | 18.463 | | |
| | -6.611 | -17.763 | | | -22.662 | | 21.306 | | |
| | -22.013 | 12.072 | | | 17.414 | | 12.707 | | |
| | 9.397 | 12.733 | | | 14.106 | | 0.399 | | |
| | -5.650 | 7.137 | | | 7.608 | | -0.533 | | |
| | -11.858 | -14.227 | | | -6.510 | | 14.699 | | |
| | -12.842 | 18.418 | | | -26.955 | | -7.400 | | |
| | 20.659 | 11.544 | | | 27.480 | | -5.806 | | |
| | 5.580 | 19.116 | | | 18.883 | | -10.925 | | |
| | -4.710 | -8.277 | | | 2.644 | | -2.994 | | |
| | -20.326 | 8.743 | | | -24.545 | | -6.607 | | |
| | -17.902 | -19.565 | | | -10.491 | | -4.725 | | |

| | | | | | | | |
|---|---|---|---|---|---|---|---|
| | -0.286 | -29.617 | | | 25.571 | | -7.006 |
| | 12.630 | 19.349 | | | -0.517 | | -0.712 |
| | 11.591 | -28.263 | | | -17.217 | | 24.369 |
| | 6.810 | -9.690 | | | -15.728 | | 0.031 |
| | -15.441 | 17.145 | | | -7.109 | | 2.137 |
| | -1.564 | 11.502 | | | 12.524 | | 19.825 |
| | 8.541 | -16.102 | | | -19.257 | | -22.591 |
| | 21.568 | 24.235 | | | -3.301 | | -16.003 |
| | 16.072 | 23.729 | | | 23.558 | | 5.538 |
| | | 25.293 | | | 18.899 | | 3.758 |
| | | -3.060 | | | 18.104 | | -3.742 |
| | | -6.479 | | | 26.762 | | -0.978 |
| | | -1.176 | | | 0.158 | | -12.764 |
| | | -15.833 | | | -9.079 | | 20.465 |
| | | -10.089 | | | -12.508 | | 9.902 |
| | | 28.194 | | | 11.018 | | 23.524 |
| | | 21.629 | | | -5.896 | | |
| | | 23.809 | | | 15.179 | | |
| | | 24.568 | | | 9.024 | | |
| | | -24.911 | | | 22.167 | | |
| | | -13.663 | | | -7.195 | | |
| | | 19.516 | | | -12.750 | | |
| | | -6.439 | | | 9.828 | | |
| | | -20.716 | | | 28.606 | | |
| | | -19.847 | | | -10.686 | | |
| | | 5.756 | | | -16.914 | | |
| | | -18.834 | | | -27.581 | | |
| | | -28.630 | | | 5.483 | | |
| | | 15.157 | | | 3.634 | | |
| | | 13.611 | | | -25.497 | | |
| | | -13.333 | | | 23.019 | | |
| | | -14.189 | | | 26.772 | | |
| | | -1.293 | | | 28.155 | | |
| | | -22.336 | | | -24.450 | | |
| | | 28.663 | | | 11.462 | | |
| | | 9.257 | | | 25.207 | | |

| | | | | | |
|---|---|---|---|---|---|
| | | 18.320 | | | 22.620 |
| | | -10.392 | | | 12.545 |
| | | 8.899 | | | -25.141 |
| | | 10.909 | | | 19.169 |
| | | -16.468 | | | -4.662 |
| | | -23.141 | | | 24.570 |
| | | -20.756 | | | 16.900 |
| | | -11.278 | | | -0.548 |
| | | 25.049 | | | -18.011 |
| | | 16.423 | | | 20.360 |
| | | -13.898 | | | 16.408 |
| | | -19.006 | | | 27.492 |
| | | 26.075 | | | -25.627 |
| | | -8.747 | | | -7.687 |
| | | 1.402 | | | -14.185 |
| | | 19.179 | | | 19.314 |
| | | -24.649 | | | 8.611 |
| | | -16.648 | | | 29.644 |
| | | -22.262 | | | 25.678 |
| | | 27.053 | | | 14.733 |
| | | 15.004 | | | 7.349 |
| | | 20.383 | | | -18.202 |
| | | 21.715 | | | 13.642 |
| | | -5.695 | | | -21.464 |
| | | 13.682 | | | 2.326 |
| | | -23.113 | | | -10.016 |
| | | 6.874 | | | -20.084 |
| | | -13.988 | | | 21.073 |
| | | -13.251 | | | 23.104 |
| | | 28.033 | | | 20.653 |
| | | -16.834 | | | 13.596 |
| | | 3.471 | | | 7.897 |
| | | 14.850 | | | 19.101 |
| | | -24.635 | | | -0.940 |
| | | 5.613 | | | 27.911 |
| | | -2.917 | | | 27.088 |

| | | | | | |
|---|---|---|---|---|---|
| | | 6.746 | | | -17.448 |
| | | -13.024 | | | 20.063 |
| | | -23.283 | | | 19.707 |
| | | 23.903 | | | 7.616 |
| | | -19.170 | | | -14.827 |
| | | 12.770 | | | 0.582 |
| | | -23.589 | | | 27.921 |
| | | -21.817 | | | -21.533 |
| | | 2.330 | | | -4.565 |
| | | -17.429 | | | -18.802 |
| | | 18.701 | | | 7.049 |
| | | -12.080 | | | 28.503 |
| | | -20.603 | | | 17.654 |
| | | 21.286 | | | -26.096 |
| | | -11.813 | | | -4.846 |
| | | 0.116 | | | -16.644 |
| | | -22.430 | | | 29.396 |
| | | 7.135 | | | 9.197 |
| | | 24.344 | | | 2.161 |
| | | -9.028 | | | 30.730 |
| | | -3.470 | | | -25.708 |
| | | -1.414 | | | 4.192 |
| | | -7.504 | | | 26.091 |
| | | -25.894 | | | -6.552 |
| | | 20.190 | | | 10.954 |
| | | 25.301 | | | 12.743 |
| | | 26.352 | | | 27.451 |
| | | -2.369 | | | -20.099 |
| | | 3.746 | | | 22.619 |
| | | -15.920 | | | -15.133 |
| | | -20.504 | | | 22.259 |
| | | -26.436 | | | -9.165 |
| | | -24.880 | | | 27.973 |
| | | 2.359 | | | 22.555 |
| | | 18.182 | | | 9.614 |
| | | -21.756 | | | -12.745 |

| | | | | | |
|---|---|---|---|---|---|
| | | -21.283 | | | -22.780 |
| | | -8.683 | | | -17.358 |
| | | -16.532 | | | -22.367 |
| | | 9.528 | | | -24.575 |
| | | 8.913 | | | -20.181 |
| | | 16.637 | | | -1.721 |
| | | -0.748 | | | 5.452 |
| | | 22.567 | | | -7.194 |
| | | -22.565 | | | 6.706 |
| | | 18.976 | | | -18.627 |
| | | 2.793 | | | 11.279 |
| | | 24.413 | | | -21.937 |
| | | -10.278 | | | -10.353 |
| | | 13.693 | | | -1.203 |
| | | 16.236 | | | 29.113 |
| | | 12.552 | | | -5.389 |
| | | 5.617 | | | -1.888 |
| | | -14.826 | | | -23.387 |
| | | -0.091 | | | 9.249 |
| | | -13.826 | | | -4.220 |
| | | -23.865 | | | 20.233 |
| | | -8.137 | | | -15.315 |
| | | 4.632 | | | 21.457 |
| | | -22.038 | | | 22.915 |
| | | -15.228 | | | 5.825 |
| | | | | | 27.152 |
| | | | | | -6.660 |
| | | | | | -18.658 |
| | | | | | 9.282 |
| | | | | | -10.097 |
| | | | | | 27.625 |
| | | | | | -15.109 |
| | | | | | 12.397 |
| | | | | | -18.068 |
| | | | | | 10.307 |
| | | | | | 22.604 |

| | | | | | |
|---|---|---|---|---|---|
| | | | | | -2.443 |
| | | | | | 26.399 |
| | | | | | 7.789 |
| | | | | | 8.296 |
| | | | | | -21.782 |
| | | | | | -13.533 |
| | | | | | -22.899 |
| | | | | | 13.938 |
| | | | | | 1.445 |
| | | | | | 27.606 |
| | | | | | 7.368 |
| | | | | | 6.249 |
| | | | | | 11.045 |
| | | | | | 2.852 |
| | | | | | -6.261 |
| | | | | | 4.644 |
| | | | | | -26.146 |
| | | | | | -27.607 |
| | | | | | 8.159 |
| | | | | | 0.589 |
| | | | | | 13.207 |
| | | | | | 3.350 |
| | | | | | 21.058 |
| | | | | | -6.968 |
| | | | | | 14.156 |
| | | | | | -26.736 |
| | | | | | -19.650 |
| | | | | | 14.896 |
| | | | | | -9.872 |
| | | | | | -16.421 |
| | | | | | 23.242 |
| | | | | | 21.709 |
| | | | | | -26.768 |
| | | | | | 19.003 |
| | | | | | -18.539 |
| | | | | | 4.597 |

| | | | | | |
|---|---|---|---|---|---|
| | | | | | -4.270 |
| | | | | | -22.247 |
| | | | | | -6.581 |
| | | | | | 14.182 |
| | | | | | 3.375 |
| | | | | | 7.602 |
| | | | | | -19.144 |
| | | | | | -6.444 |
| | | | | | -23.796 |
| | | | | | 14.159 |
| | | | | | -11.585 |
| | | | | | -16.787 |
| | | | | | 18.898 |
| | | | | | -3.072 |
| | | | | | 2.051 |
| | | | | | -4.242 |
| | | | | | 6.097 |
| | | | | | 22.511 |
| | | | | | 2.065 |
| | | | | | 20.987 |
| | | | | | 17.264 |
| | | | | | -13.165 |
| | | | | | -12.208 |
| | | | | | 1.929 |
| | | | | | -7.008 |

**FIGURE-3A-2M1B-STARVED-WORM-POSITIONS (units = mm)**

| DMSO | MB D01 | MB D02 | MB D03 | MB D04 | MB D05 | MB D06 | MB D07 | MB D08 | MB D09 | MB D10 | MB D11 | MB D12 | MB D13 |
|---|---|---|---|---|---|---|---|---|---|---|---|---|---|
| -19.092 | 18.241 | -2.516 | 10.286 | 10.776 | -0.034 | 3.842 | 15.930 | 23.365 | -18.074 | -7.615 | 15.025 | 9.405 | 21.305 |
| 14.577 | -2.795 | 5.453 | 5.213 | 22.282 | 6.865 | 10.299 | 5.483 | 13.919 | -2.365 | 13.604 | 19.455 | 22.869 | 11.474 |
| 12.848 | -13.293 | -9.921 | 8.464 | -23.854 | 19.017 | 2.742 | 2.525 | -10.222 | 6.851 | 21.164 | 15.231 | -20.186 | 21.914 |
| -7.029 | 17.052 | 25.053 | -19.025 | 21.205 | 18.257 | -4.028 | 24.077 | -12.292 | 22.580 | -10.494 | 3.238 | 14.340 | 9.862 |
| -11.313 | -9.796 | 21.812 | -12.659 | 4.755 | -11.268 | 10.971 | 9.392 | 2.010 | 20.899 | 15.173 | 24.055 | -15.876 | 12.810 |
| 11.675 | 4.597 | 19.682 | -5.706 | 4.216 | 2.304 | -2.514 | 13.796 | 23.922 | 24.086 | -2.342 | -0.106 | -2.379 | 21.010 |
| 22.327 | 7.332 | 3.691 | 20.953 | 13.158 | 20.240 | -19.935 | -18.294 | 12.109 | -12.509 | 23.460 | -3.880 | 26.229 | -1.756 |
| -24.643 | -5.741 | 16.924 | 23.388 | -12.148 | 5.648 | -13.103 | 4.384 | -21.119 | 21.321 | 1.555 | 14.701 | 24.278 | -13.967 |
| 8.469 | -2.216 | 8.692 | -0.564 | 1.219 | 17.300 | 7.041 | -23.417 | -21.588 | -0.031 | 24.004 | 13.500 | 22.431 | 17.503 |
| 7.852 | 24.773 | -13.376 | -16.857 | 22.200 | -9.758 | -5.839 | 3.290 | -14.098 | -25.137 | 0.194 | -9.043 | -26.141 | 14.633 |
| -15.983 | -2.737 | 9.529 | -22.523 | -22.022 | 12.425 | -16.232 | 6.772 | 2.895 | 8.267 | 14.315 | 6.090 | 6.404 | 10.535 |
| 3.296 | -3.458 | -24.231 | -25.761 | -19.686 | 20.946 | 11.898 | -11.009 | 25.331 | -1.981 | 19.100 | 10.364 | 23.380 | 9.201 |
| -22.535 | 15.412 | 23.829 | -14.111 | 25.284 | 6.952 | -11.701 | -21.757 | -6.018 | -13.862 | 18.221 | 5.431 | 3.699 | 19.930 |
| -23.268 | 4.451 | -13.453 | 18.432 | 12.123 | -0.069 | 18.432 | -21.066 | 23.984 | 19.873 | 22.551 | -19.936 | 24.476 | 15.510 |
| 13.555 | 21.405 | 25.823 | 6.836 | -24.532 | 18.160 | -24.884 | -12.144 | -4.449 | 18.667 | 10.317 | 13.781 | 12.146 | -8.486 |
| -8.153 | 3.092 | 16.780 | 4.858 | 5.688 | 9.410 | 23.302 | 21.990 | -3.381 | 6.342 | -17.395 | 21.250 | 21.379 | 3.781 |
| 19.462 | -18.830 | 10.433 | 19.672 | 5.012 | -19.462 | 22.373 | 14.536 | 21.940 | 0.665 | -5.697 | 22.663 | -3.093 | -17.663 |
| -8.729 | 7.425 | -22.936 | 16.775 | 22.354 | -9.226 | -18.985 | -7.998 | 22.952 | -17.680 | -11.238 | 18.655 | 3.391 | -0.245 |
| 1.628 | 18.260 | -11.522 | 2.170 | 15.798 | 0.714 | -5.636 | 14.498 | 23.794 | -1.418 | 15.781 | 5.916 | 24.940 | 11.318 |
| 21.445 | 26.917 | -0.568 | 21.532 | -3.363 | 7.449 | 25.038 | -4.897 | 21.410 | 22.678 | 6.187 | 16.306 | 12.957 | 25.334 |
| -21.294 | -5.833 | -13.290 | 17.874 | -21.998 | -20.248 | 25.597 | -13.467 | 25.513 | -3.954 | 20.964 | -0.473 | 7.568 | 7.105 |
| 24.206 | 8.738 | 24.690 | 8.762 | 18.129 | -1.886 | 20.329 | 22.654 | 14.004 | 10.736 | -19.368 | -7.876 | 23.237 | 23.182 |
| 6.370 | -11.285 | 20.425 | 9.557 | -14.262 | 0.140 | -8.972 | 17.702 | 9.452 | 15.813 | 9.261 | -7.069 | 16.058 | -19.262 |
| -7.783 | -14.483 | 10.772 | 6.855 | 15.517 | 7.298 | -3.295 | 5.663 | 19.159 | -17.716 | -10.493 | 25.089 | 12.038 | -7.401 |
| -6.268 | 18.478 | -25.419 | -14.178 | 13.445 | -23.825 | -9.309 | 11.934 | 9.827 | 24.691 | -8.700 | 20.335 | 13.704 | 9.501 |
| -15.415 | 12.803 | 21.872 | 26.885 | 10.014 | -14.710 | 2.832 | -2.433 | 6.796 | -11.487 | -17.911 | 8.175 | 4.912 | 24.466 |
| -22.038 | -23.465 | -1.030 | -7.214 | 12.861 | -3.896 | 9.639 | -4.332 | -26.314 | 20.401 | -0.647 | 2.346 | 17.903 | 22.161 |
| 14.301 | -2.851 | 3.838 | 23.767 | 20.960 | 3.019 | -4.931 | 9.260 | -8.958 | -10.041 | -2.373 | -18.219 | 14.922 | -4.044 |
| 15.403 | 21.593 | -26.628 | 4.865 | -25.506 | 8.009 | -14.781 | -7.964 | -22.277 | -7.509 | 18.149 | -20.343 | -3.464 | 18.813 |
| 22.176 | -10.237 | 24.679 | 17.461 | -13.760 | 7.188 | -3.508 | 22.842 | 13.235 | -19.776 | -23.704 | 1.000 | 26.894 | 0.971 |
| 17.708 | 24.901 | -17.693 | -8.308 | -15.721 | -21.030 | -22.583 | -16.664 | -25.925 | -19.211 | -3.003 | 18.855 | 4.080 | -25.454 |
| -3.415 | 16.700 | 22.957 | -9.136 | -5.469 | 5.037 | 18.151 | -25.704 | -0.839 | -22.860 | 12.844 | -10.185 | 12.418 | 19.395 |
| 25.340 | -24.556 | -26.616 | 8.897 | 25.075 | -11.682 | -18.836 | 5.220 | 23.572 | -22.347 | -9.267 | 22.790 | -18.842 | 0.327 |
| -2.759 | -5.226 | 27.152 | -18.980 | 19.540 | 18.895 | 11.120 | -20.541 | 20.809 | -11.861 | -25.256 | 17.845 | 14.361 | 12.254 |
| 8.504 | 8.151 | 3.055 | 5.753 | 16.554 | -14.973 | -22.695 | 13.023 | 19.204 | -5.976 | 25.336 | -20.099 | 23.456 | 22.893 |

| | | | | | | | | | | | | | |
|---|---|---|---|---|---|---|---|---|---|---|---|---|---|
| 23.146 | -11.726 | -17.920 | 27.531 | -11.146 | 23.993 | 21.411 | -0.799 | -2.691 | -8.669 | 20.844 | 20.321 | 22.959 | -10.228 |
| 1.937 | 25.760 | -20.267 | 28.107 | 19.875 | -3.841 | -16.971 | -14.044 | 7.608 | -3.159 | 19.855 | -11.291 | -10.282 | 25.515 |
| -10.608 | 5.780 | 1.266 | 4.735 | 17.693 | -9.467 | -20.927 | -22.156 | -15.633 | -3.772 | -1.736 | 22.403 | 12.928 | 21.229 |
| -3.563 | 26.787 | 10.247 | -5.810 | 22.824 | 24.572 | 12.881 | 3.994 | 7.906 | -7.280 | -15.734 | -16.505 | 22.548 | 18.362 |
| 0.773 | 5.371 | 25.729 | -7.490 | 8.053 | 20.129 | 19.043 | -27.209 | -2.198 | 12.218 | 12.223 | -1.515 | -9.112 | 0.155 |
| -18.583 | 22.829 | -24.495 | 17.612 | 6.147 | -12.215 | -23.420 | 25.438 | 1.883 | 18.182 | -13.129 | -4.085 | 25.676 | 26.076 |
| -19.597 | -1.843 | -15.243 | -13.306 | 21.833 | -8.684 | 23.708 | 22.536 | -3.720 | 16.569 | -13.609 | 4.047 | 18.193 | 17.929 |
| 8.085 | -6.400 | -23.643 | -16.549 | 6.739 | -12.921 | -1.699 | -22.442 | 25.916 | -9.308 | -7.470 | 25.456 | 15.845 | 11.486 |
| 22.216 | -19.643 | -25.372 | 8.517 | 0.594 | -11.411 | 21.083 | -20.558 | -7.586 | 26.816 | -18.040 | -18.151 | 21.076 | 21.265 |
| -23.817 | -9.089 | -3.986 | -23.177 | -25.894 | 24.888 | 10.686 | 20.957 | 2.547 | 1.376 | -5.586 | -1.243 | 20.355 | -6.480 |
| 18.145 | 21.187 | 11.384 | -21.251 | 20.632 | 21.300 | 13.945 | -23.683 | -26.148 | 27.447 | 22.168 | 19.412 | 15.408 | 19.210 |
| -8.755 | 25.276 | 18.797 | 25.995 | -2.434 | -18.966 | -13.334 | 16.518 | 21.979 | 0.349 | 17.940 | 15.325 | 5.712 | 13.998 |
| -1.020 | 13.718 | 14.430 | 14.910 | 14.521 | -1.199 | 12.563 | -10.484 | -23.835 | -11.433 | 21.568 | -8.663 | 26.104 | -16.773 |
| -1.720 | -6.974 | 3.307 | 19.758 | -1.034 | -16.143 | 26.589 | 27.239 | 17.916 | 22.069 | -20.802 | 9.881 | 19.517 | 19.835 |
| -5.347 | 27.324 | -13.124 | -9.345 | -5.990 | -24.242 | -4.151 | -26.814 | 22.567 | -19.035 | 1.832 | -5.120 | -24.684 | -24.973 |
| -26.037 | 14.396 | -21.694 | 16.751 | -20.079 | 22.549 | -12.611 | -27.658 | 3.119 | -12.795 | -2.809 | -7.127 | -13.912 | 0.638 |
| -25.618 | 7.738 | 18.291 | 7.300 | 26.608 | -21.629 | 14.505 | -20.574 | 16.368 | -8.698 | -17.615 | 0.168 | 16.014 | -3.569 |
| 23.009 | -23.892 | -21.105 | 24.760 | 27.556 | -25.450 | 21.770 | -6.937 | -9.965 | -5.209 | -26.467 | 13.161 | 28.114 | 23.432 |
| 21.066 | -8.775 | 0.107 | 20.094 | 27.075 | 13.741 | -25.331 | -18.441 | 4.347 | 2.168 | 20.074 | 1.730 | 19.581 | -6.604 |
| -10.148 | 3.914 | -24.272 | 5.945 | 1.012 | -19.173 | 7.675 | 14.205 | 24.516 | -0.934 | -4.354 | 10.209 | 20.843 | 22.782 |
| -21.218 | -25.351 | -20.658 | 28.875 | -6.943 | 25.043 | 3.312 | -26.134 | 4.817 | 26.896 | -19.501 | -15.058 | -4.875 | 20.693 |
| -18.017 | -0.369 | 9.189 | 27.016 | 23.701 | 19.038 | -9.605 | 25.690 | 10.726 | 21.233 | -3.274 | 26.118 | -26.657 | 13.569 |
| 24.239 | 4.540 | 2.200 | -0.693 | 15.238 | 25.567 | -23.699 | -23.722 | 4.116 | 24.024 | -6.169 | 6.017 | 20.298 | 1.866 |
| 4.226 | 25.222 | 1.513 | -16.097 | 8.664 | -13.830 | 9.861 | 18.112 | -0.981 | 9.819 | 6.945 | 12.146 | 4.582 | -9.846 |
| -13.439 | -6.839 | -5.706 | -28.117 | -25.307 | -25.825 | 19.191 | 25.167 | 11.391 | -3.660 | -7.662 | -24.750 | 21.587 | 16.288 |
| 4.809 | -16.818 | 8.478 | -10.356 | 23.061 | -15.281 | 5.223 | -14.080 | 8.450 | 14.577 | 25.683 | -20.607 | 17.472 | 21.488 |
| 3.170 | 13.391 | -18.270 | 24.543 | -13.925 | 0.260 | -17.230 | -9.325 | 0.106 | -24.554 | -25.787 | -2.224 | -23.595 | 15.119 |
| -12.232 | -1.234 | -0.104 | 21.369 | -12.894 | -26.401 | 29.038 | -3.874 | 10.920 | 18.782 | -3.950 | -15.107 | 22.602 | 13.109 |
| -5.944 | -21.612 | 28.139 | 16.132 | 4.921 | 4.772 | -4.394 | 22.441 | -6.093 | 19.819 | 1.918 | -3.619 | 24.835 | 24.668 |
| -6.565 | -27.739 | 8.036 | 12.584 | -4.280 | 16.641 | -29.163 | 10.244 | 10.926 | 4.722 | -14.881 | 5.928 | 16.552 | -3.385 |
| -23.534 | 26.415 | -28.821 | 9.659 | -4.350 | 11.600 | 18.265 | -17.990 | -3.699 | -3.182 | 5.041 | -21.187 | 29.113 | 20.306 |
| -1.181 | -6.816 | 21.714 | -27.352 | 25.475 | -2.410 | 20.717 | 25.624 | 26.533 | -19.727 | 1.881 | 27.732 | -10.802 | 22.355 |
| 16.755 | 7.281 | -6.633 | 2.823 | -23.025 | -7.289 | 27.968 | 15.893 | 25.640 | -22.902 | 24.029 | 14.679 | 21.124 | 17.568 |
| 22.437 | 27.937 | 27.484 | 25.126 | -20.270 | 15.892 | -23.455 | 13.606 | -6.194 | -8.931 | -5.085 | 21.174 | 17.744 | -8.238 |
| 0.475 | 23.802 | 22.760 | -11.735 | -2.698 | 2.085 | -13.014 | -29.220 | 14.949 | 21.226 | -26.592 | 4.917 | 26.781 | 9.296 |
| 21.186 | -22.193 | 5.076 | -14.014 | 26.522 | 22.759 | -28.019 | 28.910 | 8.221 | -1.389 | -0.609 | -12.781 | 28.236 | 23.276 |

| | | | | | | | | | | | | | |
|---|---|---|---|---|---|---|---|---|---|---|---|---|---|
| -20.374 | -23.955 | 28.225 | 4.849 | 7.084 | -9.337 | -15.144 | 14.468 | -26.589 | 10.403 | -4.315 | 0.944 | 19.307 | 4.682 |
| 7.213 | 14.893 | -4.079 | 14.798 | 9.882 | -3.072 | -25.319 | -16.871 | 15.930 | 28.289 | 7.322 | -10.125 | 9.496 | 17.789 |
| 8.081 | 8.537 | 13.909 | -24.766 | -0.862 | 23.893 | 15.822 | 15.057 | -15.405 | 22.265 | 0.421 | -19.249 | 30.256 | 26.455 |
| -21.307 | 27.732 | 19.575 | 7.995 | 5.926 | -10.867 | 23.246 | -5.597 | 13.916 | 26.144 | 12.633 | -2.363 | 22.065 | 3.882 |
| -23.825 | 4.038 | -8.585 | -25.066 | -15.099 | 5.395 | 25.500 | -23.858 | 9.916 | 1.754 | -1.626 | 23.599 | -1.554 | -11.315 |
| 20.721 | 0.713 | -13.407 | -28.042 | 22.402 | -8.057 | 29.759 | -6.324 | 3.905 | 0.064 | -6.467 | 26.059 | 24.426 | 5.039 |
| 27.794 | -3.582 | 8.954 | 17.587 | 13.719 | 24.636 | 18.226 | 2.366 | 2.309 | 3.896 | 18.910 | -24.667 | 15.960 | -4.889 |
| 19.834 | 7.548 | 7.553 | 0.996 | 8.632 | -1.436 | -6.162 | 21.104 | 20.009 | -16.659 | -5.248 | -19.550 | 25.466 | 13.249 |
| -7.524 | -9.050 | 1.763 | -13.180 | 29.536 | 6.773 | 22.618 | -12.089 | 25.315 | -26.596 | 12.147 | 7.747 | 23.616 | 10.888 |
| -19.908 | -19.059 | 10.334 | -1.508 | 23.856 | -13.834 | 20.104 | 25.085 | 24.030 | 15.181 | -17.537 | 16.798 | -27.107 | 16.838 |
| 15.225 | -23.167 | -0.878 | 2.000 | -15.722 | 17.794 | -7.504 | 9.038 | 26.040 | 4.266 | 10.889 | -26.802 | 24.771 | -3.545 |
| 14.588 | -18.487 | -12.176 | -26.117 | -23.764 | 20.523 | 25.805 | 18.717 | -3.787 | 27.042 | -19.212 | 2.271 | 20.768 | -1.590 |
| 25.401 | 13.419 | 23.114 | -27.248 | 27.165 | -23.575 | -28.655 | 11.662 | -19.442 | 19.387 | -16.183 | 17.237 | 21.557 | 15.884 |
| 18.898 | -27.666 | 19.029 | 12.918 | -26.188 | 15.195 | -12.401 | 8.856 | 20.477 | 24.738 | -11.330 | 6.348 | 28.726 | 17.639 |
| 18.171 | 8.637 | 1.685 | 22.518 | -11.946 | 19.729 | -15.927 | -15.426 | 24.633 | 20.313 | -10.550 | 0.771 | -25.924 | 15.045 |
| -18.780 | 15.487 | -4.534 | 24.914 | 19.753 | 17.244 | -17.586 | 26.151 | -10.766 | 2.709 | -26.519 | 24.922 | 28.890 | 23.541 |
| -3.570 | -28.590 | 18.624 | 17.498 | -6.704 | -7.786 | -21.969 | 18.434 | 8.297 | 27.270 | 15.486 | 15.415 | 25.119 | 20.150 |
| 0.485 | -1.425 | 18.252 | 3.674 | -22.649 | 18.760 | -29.897 | -5.996 | 0.181 | 12.978 | -4.696 | 19.576 | 16.310 | -1.063 |
| -0.419 | -8.232 | 0.277 | 6.303 | 17.283 | 12.873 | -15.161 | 22.579 | 9.476 | 25.790 | 23.644 | -0.133 | 2.144 | -3.091 |
| -5.162 | -21.797 | 6.190 | -27.867 | 7.480 | 0.521 | 27.749 | 24.838 | -12.518 | 16.652 | -8.072 | -22.673 | 26.616 | 6.813 |
| -18.230 | 24.420 | 4.031 | -14.685 | -27.263 | -3.653 | -29.257 | -16.763 | 1.150 | -29.140 | -6.736 | -28.452 | 20.190 | 7.401 |
| -24.258 | -14.386 | 2.875 | 13.985 | -21.039 | -25.577 | 0.945 | 5.391 | -0.100 | 23.940 | -22.580 | 24.499 | 30.191 | 21.965 |
| -23.176 | 18.099 | 22.812 | -30.328 | 0.455 | 5.059 | -4.369 | -19.304 | 19.453 | -16.031 | 16.768 | 1.210 | 24.004 | 25.722 |
| 17.262 | -7.612 | 28.317 | -11.283 | -10.897 | 0.977 | 20.354 | 0.505 | 27.819 | -12.126 | -15.284 | -4.772 | 27.545 | 13.622 |
| 5.864 | -18.190 | 25.427 | 7.620 | -13.951 | -16.681 | 11.128 | 18.145 | 25.442 | -27.624 | -3.214 | 15.883 | 16.923 | 19.391 |
| 4.776 | 12.325 | -26.409 | 22.052 | -3.621 | 26.221 | 7.881 | -15.587 | -8.706 | 27.372 | 22.422 | -17.719 | 25.960 | 10.522 |
| -12.751 | -20.427 | 0.923 | 16.958 | 27.455 | 23.537 | -22.559 | 25.095 | 8.779 | 18.878 | 1.213 | 25.815 | 23.000 | 4.321 |
| 24.805 | -5.770 | -10.034 | -22.194 | 25.928 | 0.281 | 20.247 | 10.100 | 14.086 | 10.528 | 17.456 | -2.404 | 25.545 | 16.864 |
| 4.178 | 13.971 | -3.096 | -0.263 | -22.753 | 6.252 | 20.911 | -6.214 | 12.917 | -26.661 | 18.755 | 14.314 | 13.333 | 16.067 |
| 1.001 | -29.193 | -23.490 | 25.570 | -17.347 | -26.119 | -4.343 | 1.088 | -1.640 | 26.383 | -2.079 | 2.653 | 4.299 | 22.536 |
| 14.093 | 28.208 | -24.625 | -16.388 | 8.304 | -27.660 | -8.491 | 20.091 | -2.197 | -3.545 | -7.276 | 4.414 | 26.938 | 10.977 |
| 12.778 | 6.291 | 19.421 | 5.952 | -23.221 | -1.300 | -19.051 | -24.494 | 10.037 | -5.644 | -10.049 | -10.936 | 21.753 | 9.142 |
| 7.518 | 21.188 | 23.984 | -18.725 | -9.027 | -6.687 | -19.670 | -26.623 | 4.415 | 1.264 | -5.946 | 13.295 | 21.517 | 21.024 |
| 27.935 | 19.739 | -3.692 | -13.571 | -1.909 | 14.360 | -5.708 | -13.860 | 26.866 | -20.280 | -10.983 | 25.033 | 19.051 | 18.298 |
| 2.645 | 4.043 | -5.805 | 8.440 | 15.816 | -2.044 | 28.173 | -19.603 | 2.200 | 24.082 | -23.933 | 27.128 | 14.490 | 4.744 |
| 0.307 | -18.672 | 2.066 | -29.633 | -0.657 | -15.813 | -26.705 | 30.476 | -12.111 | -11.455 | 5.288 | 8.772 | 24.837 | -17.601 |

| | | | | | | | | | | | | | |
|---|---|---|---|---|---|---|---|---|---|---|---|---|---|
| 22.680 | 18.569 | -15.389 | -15.583 | 15.447 | -21.275 | 27.241 | 0.475 | 21.706 | 24.847 | -12.418 | 21.941 | 18.236 | -24.201 |
| 15.994 | -5.085 | 6.002 | 17.863 | 5.057 | 5.922 | -30.760 | -3.139 | 9.321 | 22.520 | -8.335 | -4.480 | 26.298 | 6.385 |
| -17.593 | -27.363 | 14.517 | -10.856 | -1.105 | 19.349 | -21.173 | 20.393 | -0.836 | -3.882 | 20.829 | -0.520 | 29.579 | 15.807 |
| -6.626 | -9.995 | 10.195 | 25.823 | -17.448 | 5.102 | -7.316 | -10.622 | 25.762 | -8.192 | -10.341 | 12.076 | 20.629 | 18.484 |
| -26.561 | 22.091 | 5.673 | -20.444 | -26.028 | 26.539 | -5.170 | -25.655 | 14.193 | -22.310 | -17.717 | -7.150 | 27.635 | -11.961 |
| -21.208 | -0.906 | 8.988 | -20.712 | -9.497 | -22.287 | -15.433 | -17.630 | -19.570 | -25.848 | -24.897 | -28.304 | 22.854 | 27.738 |
| -25.373 | 27.807 | 4.367 | 1.221 | 15.039 | 3.249 | -17.541 | 15.980 | -2.484 | 15.529 | 2.002 | 23.534 | 13.347 | 26.495 |
| 16.465 | 8.654 | -5.378 | -8.621 | 11.902 | -17.582 | 15.487 | 25.730 | 2.713 | 12.817 | 7.193 | 8.205 | 30.176 | 14.651 |
| 12.128 | 26.867 | 22.107 | 11.323 | -22.831 | 1.269 | 1.002 | -26.746 | 23.603 | 0.561 | 25.041 | -29.594 | 16.266 | 13.412 |
| 18.869 | -7.541 | 21.240 | -0.601 | 19.832 | 29.382 | 12.595 | -30.587 | -1.356 | -17.417 | -2.944 | 18.545 | 24.710 | -0.848 |
| -5.026 | 17.616 | -9.462 | -23.803 | -22.454 | -9.256 | 2.170 | -20.320 | -5.164 | 14.641 | -7.778 | -5.062 | 23.612 | 20.577 |
| -13.471 | -24.545 | -10.730 | -15.575 | -7.189 | -10.394 | 19.794 | -29.717 | 1.413 | 17.424 | 13.984 | -21.636 | 21.547 | 2.513 |
| 0.010 | 8.840 | 25.268 | 7.122 | -16.387 | 20.761 | -19.205 | 11.599 | 15.153 | -11.570 | -6.792 | 28.601 | 27.315 | 22.787 |
| -19.253 | 22.784 | 6.704 | -13.289 | -3.351 | -2.141 | 27.485 | -23.383 | -8.894 | -3.959 | 18.736 | -11.419 | -20.414 | -28.158 |
| -27.257 | 25.520 | -16.120 | -2.901 | -24.298 | -18.276 | -13.696 | 19.209 | 29.392 | -24.404 | -22.208 | -18.370 | 29.653 | 18.889 |
| 18.171 | 12.590 | 30.427 | 14.024 | 6.086 | -12.671 | -18.593 | -30.564 | -23.868 | 3.431 | -21.645 | 26.206 | 16.662 | 12.265 |
| 19.423 | -7.624 | -19.388 | -22.329 | -5.367 | -6.380 | 7.398 | 29.849 | -19.880 | -20.908 | 23.368 | -10.172 | 12.337 | 22.354 |
| 2.089 | -27.270 | -11.766 | 9.789 | -18.279 | -0.416 | -2.357 | 12.349 | -4.776 | 19.697 | 19.968 | 9.351 | 25.098 | 9.880 |
| 20.564 | 29.608 | -24.104 | 28.693 | 10.619 | -24.216 | 23.437 | 3.003 | 6.301 | 20.974 | 15.239 | -16.568 | 20.147 | 3.672 |
| 13.363 | 22.169 | -9.610 | 24.973 | -27.887 | 28.664 | 16.534 | 30.070 | 27.708 | -9.288 | -16.256 | -22.655 | 23.169 | 29.675 |
| -0.579 | 8.364 | 1.194 | -2.645 | -25.145 | 17.774 | 14.152 | 26.853 | -9.618 | 28.157 | -1.046 | 5.740 | 6.798 | 21.243 |
| 14.932 | -13.465 | -4.528 | 13.798 | 30.041 | 12.353 | 19.376 | 16.437 | 11.772 | 29.521 | -16.905 | 29.344 | -31.026 | 8.743 |
| 12.795 | -18.387 | 20.231 | 5.664 | 24.809 | -10.551 | -27.315 | -15.328 | -9.840 | -3.363 | 4.822 | 17.123 | 21.360 | 19.411 |
| 1.175 | 7.136 | -7.333 | -21.651 | -8.119 | 29.758 | 25.200 | -21.601 | -25.275 | -16.835 | -5.832 | 5.076 | 29.957 | 16.160 |
| 24.180 | 13.673 | 21.955 | 7.828 | 21.922 | 27.291 | 7.252 | -11.449 | -18.697 | 6.973 | 1.863 | -4.433 | 24.691 | -2.255 |
| -15.223 | -1.980 | -1.596 | -18.256 | 30.467 | 6.154 | 14.459 | 24.360 | 23.646 | -22.365 | -1.572 | 22.792 | 17.188 | -3.047 |
| 20.855 | -25.546 | -17.530 | -19.334 | 3.908 | 23.280 | 18.847 | 2.659 | 19.525 | 20.695 | 25.773 | 21.759 | -6.457 | 21.986 |
| 2.593 | 17.425 | -28.793 | 9.314 | 20.888 | -29.143 | 7.956 | -3.449 | 15.736 | -28.428 | -10.511 | -1.201 | 3.701 | 24.985 |
| -1.155 | -9.217 | 1.520 | 6.445 | 18.625 | 8.469 | 27.465 | -10.874 | -21.750 | -14.764 | 0.503 | 20.682 | 22.643 | 26.118 |
| -2.539 | -2.684 | -26.455 | 28.015 | -28.236 | 21.683 | 9.392 | -31.152 | -25.649 | -0.904 | 21.432 | 1.846 | 29.862 | 17.872 |
| -25.203 | -24.137 | -13.717 | -20.847 | 16.282 | -9.240 | 17.513 | 6.008 | -28.727 | -2.429 | -0.377 | 17.468 | 27.499 | 22.995 |
| 20.031 | -5.662 | 10.047 | -6.287 | 8.194 | 26.672 | 20.890 | 16.773 | -27.792 | -13.019 | 23.034 | -18.830 | 18.109 | 22.471 |
| 26.822 | 17.746 | 19.942 | -10.543 | 31.807 | 11.437 | 16.477 | 8.661 | -6.000 | 12.282 | 26.499 | -20.164 | 18.904 | 27.099 |
| 0.285 | -23.766 | -11.304 | -8.205 | 17.976 | -23.646 | 18.939 | 9.758 | -4.062 | -19.445 | -14.551 | 12.037 | 22.168 | 12.642 |
| 29.260 | 0.920 | 15.084 | -12.876 | 16.493 | -13.143 | -16.232 | 4.189 | 4.568 | 22.004 | 14.850 | -27.970 | 17.877 | 18.684 |
| -22.874 | 14.443 | -3.703 | -11.744 | -17.311 | 6.519 | 28.183 | 24.563 | -26.084 | 20.959 | -25.874 | 24.157 | -20.404 | 17.327 |

| | | | | | | | | | | | | | |
|---|---|---|---|---|---|---|---|---|---|---|---|---|---|
| 16.629 | -22.887 | 9.756 | -25.192 | -10.945 | -11.648 | 19.964 | 16.226 | 18.881 | -10.393 | 22.630 | -7.966 | 9.304 | 13.158 |
| -17.462 | 17.785 | -10.675 | 3.764 | 12.987 | 20.595 | -11.626 | -16.944 | 30.094 | -8.454 | 18.532 | -17.649 | 20.491 | 19.900 |
| -3.627 | -3.200 | 18.106 | -17.123 | 0.640 | -26.411 | -4.223 | 17.893 | 28.041 | -22.494 | -9.437 | -26.297 | -30.985 | 21.878 |
| -19.045 | 23.307 | 11.109 | 28.557 | 6.456 | -3.368 | -5.626 | 17.081 | 25.233 | 27.350 | 16.153 | 19.516 | 24.132 | 20.427 |
| 19.115 | -29.122 | -1.772 | -14.540 | -25.083 | -19.584 | 29.713 | 13.943 | -8.464 | -25.245 | 10.512 | -29.891 | 7.900 | 18.653 |
| -28.516 | -8.877 | 14.445 | -19.517 | 11.303 | -9.653 | 7.917 | -15.586 | -12.374 | 17.600 | -7.508 | 13.091 | 11.749 | 4.510 |
| 11.869 | 25.758 | -29.714 | -21.657 | -12.903 | 27.125 | -7.793 | 27.770 | -30.730 | -20.996 | 24.174 | 26.186 | 4.425 | -1.538 |
| -5.516 | -6.008 | 30.425 | -9.730 | 1.875 | 16.447 | 22.021 | -2.536 | -22.936 | -7.177 | -15.371 | 21.444 | 25.483 | -3.529 |
| -20.809 | -10.795 | 6.128 | 15.172 | 24.916 | 1.355 | -20.895 | -31.674 | 10.357 | -0.227 | -9.780 | 12.600 | 26.648 | 22.830 |
| -21.624 | -17.391 | -2.242 | -8.091 | 5.497 | -2.983 | -14.277 | -4.678 | -14.027 | 16.600 | 19.423 | 13.534 | 24.559 | 27.806 |
| 25.280 | 18.256 | 10.439 | 18.751 | -18.973 | 4.828 | -24.067 | 9.365 | 22.176 | -2.954 | -12.245 | 1.497 | 23.087 | 25.036 |
| 21.186 | 7.553 | -19.297 | 8.238 | 19.516 | -0.627 | 22.401 | -14.953 | -5.088 | 17.033 | -4.030 | 11.841 | 21.010 | 12.915 |
| -23.656 | -8.941 | -13.836 | -11.068 | -7.846 | -23.894 | -10.708 | 19.517 | -14.673 | 1.895 | -17.133 | 14.584 | 8.504 | 30.780 |
| -29.090 | 19.895 | -21.247 | -26.589 | -18.459 | 14.382 | -19.043 | 18.085 | -0.241 | -0.551 | 23.390 | -2.352 | 24.218 | 25.756 |
| -17.835 | 0.813 | 5.167 | -19.080 | -28.011 | -2.567 | 15.896 | 6.952 | -4.026 | -11.840 | -2.177 | -7.350 | 0.556 | 23.081 |
| 8.227 | -28.287 | 18.386 | 1.735 | -22.367 | 26.277 | 26.769 | 21.462 | 20.192 | -2.112 | -1.651 | -4.816 | 15.480 | 15.081 |
| 1.904 | -7.886 | 16.785 | 12.784 | -17.881 | 2.737 | 24.138 | -4.983 | -5.354 | 16.364 | -9.127 | -22.904 | -23.626 | 19.392 |
| -0.527 | 4.644 | -4.976 | -8.047 | 23.135 | -24.987 | -10.006 | -9.525 | 22.658 | 15.775 | 18.884 | 29.313 | 12.889 | 13.508 |
| 4.770 | 4.280 | 0.911 | 19.487 | -7.319 | 0.662 | -5.218 | -15.733 | 9.539 | 10.255 | -21.100 | 18.826 | 23.465 | -30.000 |
| 27.921 | -25.233 | 11.118 | -8.539 | -16.435 | -21.860 | -1.246 | 13.912 | 14.109 | -18.357 | -2.944 | 30.540 | 22.543 | 21.609 |
| 0.590 | -9.035 | 4.431 | 28.833 | -21.827 | 23.189 | 19.210 | 15.412 | 2.143 | 22.832 | -12.952 | 28.659 | 27.272 | 29.510 |
| -1.928 | 16.761 | 17.661 | -20.717 | -12.566 | 10.729 | -12.896 | 21.961 | 1.158 | -11.068 | -14.357 | -7.667 | 4.741 | 20.228 |
| 14.507 | 20.481 | -26.319 | -23.389 | 6.256 | -19.805 | 12.851 | 28.087 | -16.914 | 25.511 | 12.821 | -17.429 | 19.065 | -2.801 |
| -19.660 | 17.057 | 3.666 | -7.348 | 16.024 | 7.313 | -29.553 | 19.199 | 30.239 | 7.780 | -24.964 | 21.999 | 2.707 | 0.496 |
| -4.079 | 27.825 | -3.283 | -22.835 | -0.723 | 6.116 | -1.558 | 13.141 | -29.981 | 8.887 | -0.885 | -28.873 | -12.940 | 17.006 |
| 3.782 | -16.400 | 2.345 | 14.832 | -14.126 | 3.210 | -3.634 | -13.926 | -1.248 | 5.238 | 28.657 | 15.954 | 20.538 | 15.338 |
| -25.933 | 7.869 | -7.957 | -13.205 | 15.309 | 17.426 | -7.120 | 18.849 | -0.578 | -12.484 | -10.786 | 25.123 | 15.685 | -22.236 |
| 26.962 | 31.149 | 14.929 | 19.818 | -24.631 | -1.743 | -27.975 | 2.559 | -7.463 | 4.003 | 5.295 | 2.171 | -21.700 | 25.057 |
| 0.974 | 27.372 | -9.105 | 5.554 | 22.275 | -11.084 | -4.162 | -24.158 | 18.249 | 24.961 | -28.275 | -20.800 | -10.649 | 16.339 |
| -17.758 | -2.327 | -5.831 | -25.995 | -14.566 | 18.396 | -0.819 | -22.792 | 9.493 | -29.106 | 22.256 | 4.656 | -14.079 | 26.443 |
| 28.560 | 21.144 | -25.236 | 14.122 | 19.064 | 30.615 | 25.227 | -9.103 | 0.955 | 8.392 | 21.905 | 3.485 | 3.508 | 13.823 |
| 12.148 | 16.207 | 1.741 | 9.987 | -15.228 | -25.465 | 2.553 | -11.422 | 21.425 | 27.636 | 15.551 | 27.366 | 16.536 | 19.513 |
| 4.080 | 21.606 | -14.830 | 4.141 | 16.894 | 14.676 | -4.258 | -21.955 | 30.843 | 5.897 | -2.158 | 23.865 | 15.143 | -3.219 |
| -1.442 | -8.394 | -12.539 | -14.369 | -19.365 | 29.856 | -17.342 | 7.355 | -6.147 | -0.852 | -8.041 | 20.425 | 13.045 | 27.358 |
| 1.669 | -0.284 | -26.131 | -12.784 | -12.013 | -10.036 | 7.803 | 29.954 | -25.334 | 3.354 | -24.122 | -25.451 | 24.486 | 11.910 |
| -27.155 | 15.027 | 1.210 | 12.621 | 3.108 | -19.519 | -23.375 | 20.986 | -23.774 | -24.004 | 14.489 | 21.178 | 14.741 | 21.705 |

| | | | | | | | | | | | | | |
|---|---|---|---|---|---|---|---|---|---|---|---|---|---|
| 24.055 | -12.138 | -6.382 | 7.487 | 23.907 | -5.485 | 9.404 | -6.866 | 24.209 | 17.948 | -22.760 | 22.340 | 17.919 | -15.791 |
| 5.453 | -6.364 | 18.027 | 0.863 | 22.904 | 11.639 | -7.547 | 25.720 | -12.153 | 12.636 | 6.404 | 7.303 | 26.589 | 15.619 |
| -21.173 | 17.957 | -21.441 | 3.859 | 18.328 | 6.897 | 7.610 | 27.051 | 17.779 | 30.266 | 29.312 | 21.117 | 22.633 | -29.502 |
| 27.643 | -5.240 | -3.358 | 14.044 | 9.606 | -29.041 | 4.867 | 18.023 | -2.631 | -10.723 | 13.428 | 23.271 | 19.137 | 21.115 |
| 0.519 | 15.716 | 26.751 | 10.229 | 15.827 | 13.055 | -28.801 | -5.406 | 15.733 | -16.391 | -19.998 | 31.319 | -7.372 | -27.441 |
| -2.442 | -26.136 | -0.742 | 18.527 | 0.700 | -12.337 | -19.952 | -26.196 | 1.474 | 17.539 | 11.464 | 29.545 | -4.415 | 10.564 |
| 29.399 | -9.985 | 17.258 | -21.553 | -1.142 | 6.191 | 9.002 | 28.633 | 8.866 | 9.323 | -14.411 | 16.640 | 16.577 | 0.598 |
| -12.170 | 14.590 | -8.703 | -13.321 | -24.652 | -29.805 | -5.871 | 18.335 | 10.885 | 6.911 | -0.477 | -12.120 | 2.476 | 24.295 |
| 25.495 | 19.443 | -25.006 | 1.679 | -29.161 | 22.622 | 22.009 | -3.757 | -31.677 | 2.627 | -8.937 | 23.629 | 16.793 | 18.379 |
| -0.663 | 27.126 | 14.242 | -1.472 | 22.038 | 15.396 | 26.523 | 5.594 | 28.894 | 28.045 | -2.544 | -1.389 | 9.650 | 14.583 |
| 26.961 | 9.112 | 7.421 | 12.866 | 20.517 | 24.399 | -4.903 | 18.920 | -5.787 | -18.772 | 2.803 | 3.712 | 24.032 | -2.244 |
| -29.691 | -4.365 | -20.167 | 27.572 | 31.093 | 30.974 | 28.299 | -21.015 | 10.359 | -19.852 | -3.912 | 31.518 | 6.714 | 17.661 |
| -3.749 | -16.063 | 0.007 | -3.602 | 28.119 | 23.568 | 3.840 | 1.552 | 21.614 | -5.702 | 26.402 | 14.483 | 20.467 | -1.298 |
| 0.050 | -26.645 | 4.657 | 21.489 | 20.073 | 14.619 | -19.105 | 6.660 | -1.108 | 1.082 | 23.512 | 25.920 | 22.487 | -4.141 |
| -28.845 | -14.582 | 1.670 | -17.093 | 4.821 | 0.237 | -1.137 | 24.749 | 9.887 | 13.730 | -6.812 | 10.228 | -1.399 | 20.399 |
| -25.473 | -4.552 | -17.326 | -20.411 | 15.024 | 10.907 | 6.344 | 17.924 | 25.921 | 16.863 | -8.555 | 27.417 | -8.203 | 30.797 |
| -18.128 | -29.921 | 12.882 | 18.761 | -9.249 | -6.053 | -16.923 | 11.408 | -26.461 | -1.959 | 7.820 | 18.431 | 27.202 | 15.251 |
| -7.259 | 18.660 | 27.877 | -13.265 | -12.180 | -9.801 | 16.479 | 8.120 | 24.097 | -24.728 | 30.240 | 19.337 | 5.534 | -3.052 |
| 6.175 | -7.597 | 2.474 | -19.985 | 10.515 | -6.686 | -4.705 | 1.747 | 3.113 | 1.429 | 1.787 | -29.988 | 26.694 | 28.093 |
| 4.498 | -15.365 | -10.892 | 25.753 | 21.744 | 16.448 | -22.330 | -3.741 | 23.475 | -21.218 | 1.259 | 30.065 | 21.875 | 16.283 |
| 9.207 | -9.276 | 1.650 | -28.051 | -23.465 | 24.741 | -25.462 | -2.191 | 8.638 | -12.717 | -3.509 | -24.580 | 25.856 | 9.313 |
| -24.007 | -27.123 | -4.528 | -15.789 | 11.820 | 6.111 | 3.349 | 5.452 | -14.497 | -22.374 | -4.612 | -0.676 | -23.064 | -2.487 |
| 27.606 | -3.517 | 21.958 | 2.609 | 9.224 | -21.610 | -1.247 | -1.016 | -4.688 | 11.846 | 12.906 | -28.780 | -12.480 | 30.277 |
| -26.682 | 5.568 | -20.522 | 26.659 | -21.835 | 29.395 | -19.241 | -13.898 | -6.800 | 8.138 | 16.708 | 21.996 | -12.740 | 24.104 |
| -8.573 | -12.254 | 24.827 | 16.543 | -18.457 | -18.907 | 19.478 | -25.672 | -7.068 | 1.259 | 15.629 | 18.026 | 15.046 | 10.491 |
| 3.614 | 20.237 | -6.603 | -3.231 | 4.067 | -2.321 | -1.549 | -22.639 | 8.155 | 2.859 | 7.028 | -4.695 | 20.016 | 7.291 |
| -1.489 | 3.935 | -14.459 | -23.794 | 4.693 | -11.574 | 9.440 | 24.156 | 6.723 | -29.929 | -18.633 | 24.274 | 25.036 | 23.154 |
| -5.268 | -6.499 | -1.943 | 6.947 | 2.746 | -20.903 | -12.071 | 26.177 | -25.421 | 19.945 | -26.350 | 8.803 | 4.311 | 31.781 |
| 16.216 | 7.176 | 15.003 | 7.525 | 13.956 | 27.565 | -8.578 | 15.159 | 18.729 | -6.277 | 21.658 | -23.644 | 2.984 | 27.610 |
| -9.114 | 9.497 | -23.402 | 19.080 | -24.941 | -15.011 | 15.092 | -21.648 | 5.206 | 5.921 | 9.242 | 28.391 | -27.100 | 18.982 |
| 17.474 | -6.111 | 12.288 | -26.785 | 13.067 | -0.602 | -7.430 | 5.406 | -7.593 | -19.573 | 18.205 | 6.270 | -11.050 | 5.349 |
| -18.221 | -9.346 | 10.531 | 1.223 | 16.501 | 17.676 | 11.217 | -24.739 | 24.221 | -1.365 | 25.352 | 16.357 | 4.093 | 31.162 |
| 25.711 | -26.180 | 9.472 | -6.584 | 1.987 | 15.667 | -10.835 | 24.588 | 7.650 | -16.467 | -10.259 | 24.793 | -1.367 | 12.002 |
| 20.922 | -15.614 | 3.885 | 27.523 | -7.004 | 2.348 | -3.000 | 25.390 | 0.653 | -13.883 | -23.088 | 1.346 | 19.579 | 9.533 |
| -14.311 | 15.451 | -25.114 | 5.229 | 21.521 | 0.875 | -5.834 | 8.344 | -8.583 | 14.553 | 8.339 | 2.666 | -25.909 | -3.630 |
| -20.347 | -24.810 | 17.348 | 10.825 | 12.526 | -24.515 | -6.532 | 3.794 | 22.287 | -3.906 | 1.205 | 30.709 | 1.231 | 26.315 |

| | | | | | | | | | | | | | |
|---|---|---|---|---|---|---|---|---|---|---|---|---|---|
| 23.022 | 25.290 | -8.477 | -1.789 | 11.148 | -19.218 | 12.433 | -27.835 | 13.386 | 18.563 | -0.127 | 7.022 | 15.393 | 17.400 |
| -13.371 | 19.560 | -0.087 | -4.645 | -7.076 | -29.406 | 8.781 | 1.894 | 14.576 | -32.012 | -14.660 | -29.716 | 16.127 | -14.665 |
| -27.457 | 16.329 | 29.158 | -12.726 | 6.084 | 23.452 | -15.809 | -4.839 | 9.296 | 17.190 | 24.435 | 15.921 | 2.846 | 23.739 |
| 14.756 | 22.401 | 20.336 | -26.735 | -26.292 | 25.491 | 12.924 | -8.188 | 20.906 | 12.380 | -2.911 | -6.711 | -4.837 | 18.228 |
| 13.873 | -27.728 | 8.624 | -2.931 | 20.583 | -20.297 | 22.152 | 11.783 | -24.388 | -11.996 | -18.413 | 23.721 | 9.946 | 16.140 |
| -3.381 | 23.270 | 4.828 | 11.702 | -12.706 | -25.334 | 18.410 | -19.822 | -31.598 | 22.206 | 5.538 | -21.645 | 18.649 | 6.033 |
| -25.727 | -2.927 | 21.951 | -20.912 | 12.012 | -2.254 | 3.743 | -1.282 | 4.244 | -15.556 | 15.125 | -5.479 | 15.547 | -4.821 |
| -19.240 | 4.176 | -22.987 | -26.005 | 3.194 | -29.804 | 0.777 | -7.450 | -28.340 | -29.617 | -2.285 | 29.826 | -8.298 | -2.841 |
| 28.764 | -7.392 | 6.631 | 10.596 | 27.949 | -3.658 | 20.882 | -17.214 | 1.418 | -3.289 | 14.065 | 15.265 | -16.185 | 29.241 |
| 29.844 | 15.651 | -18.453 | 4.106 | 17.434 | -14.146 | 14.633 | 25.428 | 6.971 | -20.870 | 11.978 | -1.750 | 18.402 | 19.991 |
| -29.705 | 12.717 | -20.719 | -6.472 | -28.042 | 27.689 | 13.469 | 11.446 | 30.367 | -6.237 | 21.461 | -2.230 | 2.696 | 19.303 |
| 15.866 | 12.004 | -2.280 | -11.403 | -0.291 | 13.052 | 20.005 | 21.298 | -5.690 | 28.750 | -8.667 | 18.110 | -6.584 | 12.959 |
| 30.527 | -10.553 | -19.140 | -18.246 | 14.330 | 5.790 | 10.300 | 9.787 | 3.467 | -14.479 | -29.974 | -27.730 | 23.209 | 8.952 |
| -23.047 | 26.793 | -10.075 | -2.936 | 0.519 | -6.900 | -5.981 | 20.187 | 13.390 | -10.951 | 29.379 | 21.800 | 6.999 | 21.022 |
| 23.397 | 9.348 | -16.804 | -10.688 | -16.968 | 14.561 | 11.727 | 13.917 | -7.325 | -17.055 | 28.812 | -1.588 | 16.903 | 23.137 |
| 3.857 | -26.982 | -29.722 | -22.075 | -21.162 | -5.207 | 7.653 | 14.265 | -0.256 | 3.276 | 12.413 | 23.819 | 25.200 | 26.677 |
| -17.854 | -27.991 | -13.663 | -15.251 | 18.473 | 4.675 | -12.443 | -16.043 | 18.333 | -30.902 | -20.769 | 8.174 | -23.271 | 17.298 |
| -6.069 | 21.185 | -20.155 | -5.146 | 22.684 | -28.058 | 18.624 | 28.571 | 16.910 | -6.755 | 23.731 | 2.600 | -2.651 | 23.789 |
| 26.823 | -4.592 | -14.961 | -8.139 | -5.097 | 8.168 | -4.883 | 22.303 | 6.217 | 13.058 | -29.349 | -28.393 | 16.317 | 17.676 |
| 1.978 | 11.902 | 0.091 | -27.657 | 2.651 | -4.490 | -0.640 | 26.353 | -5.452 | -15.836 | -28.797 | 11.292 | 22.005 | 14.665 |
| -1.677 | -23.547 | 8.769 | 25.555 | -24.559 | -2.305 | 8.566 | 16.709 | -27.360 | 4.494 | 16.982 | 26.792 | 22.489 | 10.701 |
| -26.062 | -21.860 | 2.496 | 3.935 | 4.983 | -30.516 | 10.958 | -21.166 | 30.772 | 2.503 | -3.212 | -0.954 | 26.398 | 25.808 |
| -9.345 | -28.823 | -5.015 | -15.323 | -10.283 | 19.806 | 3.904 | -22.227 | 26.354 | 5.144 | 26.245 | 27.367 | 18.580 | 1.776 |
| 12.829 | 9.229 | -11.965 | -3.653 | -15.059 | 9.077 | 0.927 | -15.273 | 9.854 | -20.125 | -5.634 | 6.281 | -6.051 | 18.839 |
| -2.917 | 17.704 | 6.308 | 12.437 | -27.042 | 26.244 | -20.360 | -3.612 | 15.999 | -21.766 | -9.158 | 28.340 | 13.789 | 29.057 |
| 29.158 | -23.417 | -0.648 | 19.016 | 21.123 | -10.545 | 8.125 | 19.053 | -1.886 | 0.843 | 27.696 | 20.243 | 10.602 | 10.373 |
| -21.971 | -12.247 | -17.677 | 8.637 | -25.083 | 26.806 | 19.469 | -17.704 | 4.887 | 16.229 | -10.019 | 9.788 | -27.376 | 27.954 |
| -4.774 | -3.836 | 10.099 | -25.662 | -2.584 | 15.674 | 15.175 | -6.403 | -20.411 | -19.990 | -11.063 | 4.350 | -25.632 | -16.218 |
| 21.719 | -6.565 | -29.237 | -3.022 | -18.434 | -15.592 | 8.115 | 16.143 | 11.846 | 0.955 | 23.327 | -18.059 | 24.835 | -21.762 |
| 21.044 | 5.707 | 13.910 | -5.605 | 26.380 | 10.209 | 14.810 | -2.272 | 8.530 | 18.566 | -16.504 | 24.001 | 25.203 | 13.761 |
| -17.091 | -18.706 | 4.121 | 7.244 | -13.737 | -19.297 | 0.719 | -24.258 | 7.892 | -12.732 | -20.153 | 22.609 | 11.428 | 3.536 |
| 1.945 | -2.992 | -3.608 | -23.911 | -8.904 | -3.207 | -2.767 | 22.860 | -31.063 | 0.491 | 28.515 | -21.761 | 17.776 | 6.490 |
| 16.629 | -28.920 | -8.143 | -3.709 | 18.947 | 21.110 | -18.230 | 5.536 | 0.011 | 26.517 | 25.111 | -20.302 | 23.540 | 15.127 |
| -18.775 | -13.820 | -21.491 | 25.140 | -4.148 | 28.764 | -6.284 | -23.200 | 27.844 | 22.542 | 22.778 | -8.300 | 18.940 | 27.283 |
| 23.176 | -25.578 | 20.075 | -9.686 | 19.632 | 18.396 | 11.576 | -16.351 | -15.612 | 5.025 | -1.490 | -2.838 | 25.325 | -5.181 |
| 10.010 | 8.646 | 20.583 | -15.617 | -19.382 | 19.385 | -4.948 | -24.977 | -11.909 | -12.382 | 20.406 | -2.417 | 15.143 | 20.623 |

| | | | | | | | | | | | | | |
|---|---|---|---|---|---|---|---|---|---|---|---|---|---|
| -24.614 | -8.482 | 22.367 | 3.485 | -22.688 | -13.224 | -1.735 | -0.578 | 10.151 | 29.571 | -24.697 | -11.934 | 25.816 | -7.985 |
| -25.975 | -13.057 | -19.893 | 22.282 | 24.435 | 9.403 | 18.326 | 8.218 | -7.089 | 19.001 | 20.982 | -16.203 | -17.358 | 12.675 |
| 9.274 | -0.768 | 10.736 | -3.930 | -25.529 | -22.643 | -2.064 | -4.709 | -16.831 | 10.524 | 21.845 | 22.152 | 16.901 | 28.137 |
| 24.443 | 24.487 | 15.330 | -2.811 | 1.518 | 5.302 | 21.841 | 7.578 | 10.514 | 11.570 | 29.099 | -18.542 | 27.728 | 24.201 |
| -20.817 | 13.502 | 0.023 | 11.960 | 11.424 | -11.830 | -15.851 | -8.343 | 5.719 | -4.992 | 25.766 | -29.416 | -13.537 | 13.497 |
| 20.783 | -15.674 | 6.297 | -9.105 | -5.021 | 26.164 | 25.558 | -21.034 | 9.617 | -7.859 | 0.874 | 7.499 | 15.970 | 20.898 |
| 6.990 | 25.075 | 22.586 | -14.467 | -12.340 | 15.504 | 24.397 | 17.407 | -10.095 | -21.215 | 16.403 | 5.500 | 15.386 | 9.946 |
| 18.718 | 17.366 | -6.237 | -12.077 | 29.431 | 13.160 | -8.235 | 16.478 | 11.210 | 5.870 | 7.009 | -16.848 | 18.897 | 15.099 |
| -14.762 | -13.026 | -16.828 | 7.571 | 23.322 | -2.841 | 0.299 | 6.152 | 0.161 | -20.117 | 30.291 | 25.648 | 19.265 | 19.361 |
| 29.379 | 1.844 | 29.416 | -21.038 | 5.600 | -7.060 | -3.239 | -22.441 | -30.810 | 15.431 | 26.470 | -20.953 | 27.559 | 22.424 |
| 21.332 | -2.186 | -11.861 | -0.289 | 10.554 | 20.939 | 4.891 | 1.105 | -17.371 | -26.054 | 29.743 | 19.056 | 17.250 | 20.560 |
| 16.061 | 4.491 | -17.856 | 10.307 | 16.435 | -21.108 | 1.501 | 8.626 | 26.123 | -27.972 | 19.393 | 23.562 | 25.397 | -13.669 |
| -7.296 | 26.762 | -29.562 | 11.188 | 0.640 | 14.526 | -2.676 | -18.920 | 18.979 | 27.807 | 17.277 | 27.059 | -22.597 | -28.366 |
| -29.111 | -27.007 | -8.013 | 6.429 | 7.961 | -10.324 | -4.059 | 0.325 | -21.632 | 8.636 | 30.893 | -6.839 | -4.533 | 19.669 |
| 14.142 | 6.063 | 13.132 | 2.310 | -9.358 | 2.001 | 16.734 | -2.687 | 21.872 | -0.519 | 3.926 | 19.583 | 21.612 | -21.607 |
| -18.268 | -4.186 | -19.137 | 4.701 | -2.807 | -7.559 | 3.772 | -23.565 | 2.466 | 9.453 | 3.250 | -4.776 | 18.968 | 15.496 |
| 14.578 | 16.120 | 27.238 | -10.617 | 21.529 | -18.643 | -5.246 | 11.754 | -2.000 | -22.926 | 5.188 | 11.463 | 24.576 | 14.723 |
| -9.255 | -3.256 | 14.613 | -17.060 | 19.577 | -9.962 | -12.436 | -7.425 | -24.296 | -14.954 | 21.911 | 27.492 | 20.372 | 26.903 |
| 0.222 | -11.900 | -12.331 | -13.242 | 9.298 | -6.182 | -8.575 | 12.473 | -16.460 | -1.533 | 12.554 | 14.365 | 13.862 | 11.922 |
| -28.240 | 25.949 | 10.325 | 3.552 | 10.012 | 24.351 | 13.073 | -12.294 | 12.940 | 3.655 | 22.725 | -23.066 | 25.635 | -5.827 |
| -0.356 | 3.283 | -25.214 | -11.360 | 21.747 | 14.969 | 8.520 | -1.288 | 21.842 | 18.442 | -30.752 | -0.897 | 28.225 | 16.902 |
| 31.348 | -21.610 | 7.787 | -24.485 | 3.425 | -19.508 | 12.347 | -10.625 | 26.156 | -9.833 | 26.260 | 1.785 | 15.021 | 18.017 |
| -2.157 | -5.336 | -22.463 | -4.945 | -24.733 | -2.590 | -1.426 | -3.790 | 16.739 | 9.141 | 8.718 | -9.743 | -9.080 | 29.420 |
| 23.867 | 6.930 | 4.635 | -12.330 | 22.596 | -16.769 | -15.792 | 13.748 | 9.881 | 23.252 | 24.924 | 5.169 | 18.196 | -23.291 |
| 10.367 | -17.354 | -15.965 | -17.207 | 5.908 | -8.615 | -5.787 | 10.597 | 24.769 | 4.725 | 23.359 | -19.506 | 9.575 | 21.819 |
| 5.354 | 6.360 | -1.573 | -21.643 | 0.346 | 8.782 | -20.441 | 16.917 | 2.167 | -14.381 | -28.370 | 12.792 | 23.755 | -1.581 |
| -29.194 | -0.210 | 15.111 | 9.105 | -4.636 | 25.158 | -11.794 | 5.055 | -5.404 | 1.638 | 18.362 | -22.226 | 28.716 | 22.358 |
| 28.602 | 25.044 | 25.150 | -16.869 | -6.260 | -21.568 | -18.513 | 18.901 | 17.925 | 17.586 | -12.522 | -8.368 | 26.374 | 25.818 |
| 0.415 | 2.200 | 16.211 | -14.789 | -11.854 | 23.002 | 17.306 | 12.605 | -6.527 | -12.832 | -2.643 | -28.232 | 17.113 | 18.921 |
| -21.345 | 11.900 | -5.686 | 22.568 | 2.894 | -4.660 | -7.314 | -2.822 | -11.866 | 1.195 | 19.658 | 25.538 | 22.701 | 13.806 |
| 15.327 | -0.731 | -19.263 | 24.868 | -9.580 | -27.894 | 11.693 | -11.083 | 10.607 | -6.971 | -27.952 | -22.278 | 29.730 | 0.594 |
| 7.742 | -28.089 | 13.413 | 1.803 | 5.885 | 19.505 | 11.140 | 5.782 | 0.552 | -4.567 | 12.955 | -12.195 | 19.146 | -21.533 |
| -4.885 | 8.406 | 28.123 | 2.628 | -8.609 | 20.798 | 17.029 | 7.689 | -20.354 | -2.340 | -11.725 | 4.449 | 27.228 | -7.813 |
| 23.305 | -2.472 | 10.549 | -1.524 | -23.155 | 14.369 | 1.178 | -5.772 | 2.873 | -5.324 | -1.454 | 19.166 | 20.577 | -11.946 |
| -13.061 | 24.408 | 10.069 | 11.245 | 4.523 | 16.192 | -2.882 | 9.837 | 23.753 | -25.501 | 31.031 | -27.199 | 26.103 | 17.685 |
| -14.863 | -5.233 | 24.960 | 20.715 | -2.610 | 10.092 | -1.359 | -10.494 | 27.770 | 22.475 | 22.171 | 16.104 | 18.351 | -20.657 |

| | | | | | | | | | | | | | |
|---|---|---|---|---|---|---|---|---|---|---|---|---|---|
| -27.140 | 12.877 | 14.167 | 1.057 | 1.021 | -29.465 | -8.185 | 5.656 | 13.888 | 20.633 | -29.359 | 0.048 | 17.842 | 21.482 |
| -12.643 | 11.089 | -7.926 | -14.411 | 11.142 | -22.332 | -11.814 | 6.521 | -5.642 | 25.685 | 26.295 | 21.678 | -9.160 | 16.921 |
| -24.708 | -19.249 | 2.098 | 0.783 | 26.432 | 26.401 | 2.190 | 2.960 | -15.557 | -18.188 | 29.810 | 6.408 | 6.781 | -14.983 |
| -2.419 | -4.579 | 0.750 | -11.762 | 0.344 | 24.456 | -19.384 | 8.200 | -3.595 | -19.201 | -11.428 | 23.914 | 20.264 | 20.866 |
| 0.898 | 26.501 | 25.920 | 21.538 | -7.311 | -12.081 | -14.845 | 13.641 | 27.028 | 10.401 | 27.969 | 3.923 | 26.885 | -25.118 |
| 13.803 | 5.914 | -26.038 | -5.261 | -8.961 | -25.092 | -7.456 | -15.785 | 5.158 | -0.211 | 0.010 | -4.550 | -6.416 | 26.332 |
| -6.947 | 17.731 | -0.254 | 24.176 | -25.311 | 24.773 | -6.533 | 23.048 | 21.457 | 4.821 | -28.125 | 13.733 | 25.878 | 25.597 |
| -3.640 | 14.566 | 12.217 | -16.787 | 14.457 | 6.180 | -9.419 | -8.938 | 9.437 | 6.413 | 29.353 | -27.488 | 23.663 | 22.043 |
| 14.253 | -3.757 | 9.265 | -19.628 | -21.099 | -12.781 | 4.035 | -16.415 | -18.405 | 17.621 | -29.895 | 20.471 | 26.992 | -14.188 |
| 30.541 | -1.241 | -3.866 | 2.314 | -13.494 | -1.040 | 5.667 | 4.828 | 7.361 | -0.624 | 13.622 | 11.422 | 19.344 | -0.251 |
| 21.570 | -7.220 | -17.967 | -16.514 | -8.433 | 27.459 | 1.594 | 2.266 | 15.563 | 27.709 | -0.520 | -8.022 | 14.325 | -5.603 |
| -4.518 | -7.706 | -20.384 | 6.452 | -10.950 | 14.899 | -25.427 | -24.731 | 12.159 | 11.275 | 7.273 | 26.174 | 22.602 | 10.432 |
| -1.113 | -4.205 | -7.061 | -21.268 | 4.886 | -6.430 | -10.338 | 8.922 | 20.599 | -4.729 | -22.071 | 4.958 | -30.519 | 7.132 |
| -29.106 | -2.862 | -14.112 | 9.007 | 15.374 | 4.310 | 11.142 | -0.440 | -6.146 | 0.333 | -4.793 | 16.795 | 23.192 | -29.508 |
| 23.910 | -12.154 | 3.148 | -18.591 | 1.945 | -24.225 | 10.313 | 21.579 | -5.071 | -1.166 | -30.618 | 1.331 | 19.874 | 24.080 |
| -0.649 | 6.243 | -23.652 | 21.328 | -4.127 | -13.337 | 12.729 | 1.267 | 16.607 | 1.725 | -0.203 | 18.975 | 21.066 | 27.374 |
| 1.187 | 20.173 | 26.870 | 27.213 | -5.456 | 16.849 | 22.330 | 0.749 | -10.455 | -5.886 | -17.093 | -9.230 | 28.529 | 23.021 |
| 17.953 | 4.796 | 16.348 | -5.595 | -27.394 | 19.718 | -26.251 | -13.504 | 0.525 | 17.410 | -14.271 | 6.042 | 7.154 | 20.300 |
| 5.709 | 10.998 | -1.523 | -4.972 | 0.367 | 10.021 | 7.965 | -20.605 | -0.453 | -23.396 | 4.230 | -25.551 | 29.496 | 16.483 |
| -4.948 | 10.059 | 5.967 | -25.157 | -12.382 | -11.588 | 16.721 | 7.696 | 28.726 | -6.677 | -3.727 | 12.909 | 26.493 | 12.138 |
| -20.133 | 1.741 | -2.245 | 28.434 | -14.199 | -2.538 | 2.028 | 8.375 | 7.165 | 7.667 | 6.319 | 2.226 | 19.026 | 17.582 |
| 26.586 | -20.831 | 18.759 | 7.132 | -14.915 | -1.779 | -23.445 | -25.750 | 23.168 | -1.939 | -14.798 | -16.494 | 17.999 | 3.164 |
| -9.208 | 24.782 | 24.276 | -13.022 | 3.075 | 26.981 | 5.643 | 19.689 | 17.950 | -20.805 | 19.594 | 0.844 | 30.490 | 28.207 |
| 9.165 | -24.976 | -0.770 | 20.179 | 26.778 | -25.116 | 7.981 | 4.566 | 18.956 | -22.863 | -19.993 | 7.601 | 26.926 | 16.269 |
| 0.054 | -19.969 | 23.277 | -9.691 | 19.367 | 8.315 | -25.922 | -6.077 | -13.197 | -17.756 | 23.741 | 23.999 | 20.531 | 18.909 |
| -19.435 | -22.504 | 22.245 | 23.984 | 11.986 | 5.630 | 15.264 | -18.697 | -27.710 | 0.887 | -9.027 | -11.861 | 31.727 | 14.207 |
| 6.668 | 5.261 | 2.275 | -5.106 | -3.796 | -22.068 | -5.615 | -6.463 | 4.493 | 2.657 | 12.535 | -17.548 | -20.228 | 0.239 |
| -27.963 | 0.970 | 7.702 | -26.171 | 3.901 | -9.546 | -6.389 | 7.386 | 21.011 | 7.074 | 9.998 | 12.912 | 21.957 | -6.582 |
| 25.855 | 20.854 | -23.472 | -5.829 | 1.055 | -10.921 | 13.952 | -24.567 | -3.725 | -21.216 | 16.611 | 20.526 | 22.826 | 15.312 |
| -10.689 | -24.200 | -8.321 | -24.306 | 12.413 | 15.571 | -5.897 | -11.235 | 9.427 | 10.616 | 1.244 | -21.927 | 28.469 | 24.788 |
| 21.885 | 0.794 | -18.445 | 18.389 | 9.319 | -3.764 | -21.884 | -5.508 | -23.820 | 25.153 | 16.890 | 25.986 | 25.131 | 19.367 |
| 29.190 | -0.322 | 8.719 | -11.565 | -13.877 | 16.924 | -4.065 | -19.914 | 21.913 | 18.872 | 11.632 | -16.746 | 19.112 | 20.602 |
| 3.420 | -5.025 | -9.292 | -12.654 | 5.944 | -28.027 | -23.411 | 12.954 | 8.396 | 5.125 | 15.272 | 17.597 | 21.600 | 19.735 |
| -24.938 | 3.811 | -27.644 | 3.648 | -3.297 | 26.636 | 16.219 | 24.509 | -2.813 | 14.189 | -4.245 | 15.097 | 20.187 | -9.020 |
| -2.551 | 11.774 | -11.268 | 19.560 | -8.978 | 17.198 | -26.383 | 11.019 | 19.767 | -2.605 | 16.043 | 25.001 | 27.919 | 11.781 |
| -17.471 | -8.430 | 18.199 | -14.103 | -26.470 | 25.167 | 10.123 | -5.127 | -4.194 | -23.319 | -5.018 | 9.381 | 23.281 | -23.637 |

| | | | | | | | | | | | | | |
|---|---|---|---|---|---|---|---|---|---|---|---|---|---|
| -27.161 | -14.834 | 21.674 | -18.186 | 5.164 | 8.518 | -17.860 | -8.372 | -17.858 | 1.490 | 11.450 | -8.024 | 24.327 | 10.642 |
| 12.938 | -11.471 | -8.630 | -11.951 | 2.068 | -1.208 | 25.627 | 18.492 | 17.726 | -7.605 | 2.866 | -11.290 | 23.006 | 6.796 |
| -22.125 | 24.289 | 2.937 | -19.205 | 23.609 | -16.794 | 10.460 | 3.742 | 3.153 | -3.759 | 25.465 | 1.971 | 27.416 | 22.324 |
| -18.945 | 14.905 | 7.656 | 7.878 | -4.915 | -24.672 | -24.030 | -12.228 | 7.150 | 9.276 | -25.719 | -5.573 | 22.056 | 21.681 |
| -26.506 | 14.471 | 27.522 | 28.284 | 14.816 | 24.810 | 22.023 | -18.330 | -7.993 | 27.396 | 11.940 | -3.908 | -18.819 | 14.983 |
| 19.759 | 26.223 | 10.836 | -23.459 | -5.817 | -5.522 | 18.824 | -15.062 | 21.553 | -4.814 | -10.070 | 21.951 | 14.270 | -4.342 |
| 18.324 | 7.211 | -23.337 | -2.524 | 10.158 | 10.085 | -22.855 | -16.644 | 11.939 | -19.804 | 17.734 | 13.431 | -6.351 | -6.703 |
| -21.789 | -18.899 | -24.304 | 11.811 | -22.571 | -15.147 | 19.883 | -26.120 | -4.439 | 15.245 | -29.586 | 19.124 | 23.463 | 19.294 |
| -5.959 | -21.894 | -26.907 | -23.024 | 14.465 | 19.199 | 8.098 | 18.980 | -17.815 | 26.635 | 9.949 | -0.098 | 8.361 | 19.891 |
| 5.369 | -7.993 | 18.760 | 21.321 | 4.855 | 4.330 | 25.163 | 22.824 | -19.383 | 25.892 | 13.186 | 18.773 | 22.031 | -4.143 |
| -22.840 | 9.757 | -1.954 | -5.945 | 22.879 | 23.761 | 11.409 | -13.508 | 15.815 | 6.068 | 2.201 | 6.272 | 28.578 | -2.267 |
| 20.891 | 4.343 | -4.235 | -14.951 | 17.863 | 2.723 | -17.428 | 4.246 | 4.140 | -21.867 | -26.747 | -21.675 | -12.421 | 14.028 |
| -20.180 | -6.284 | -9.745 | -28.628 | 8.778 | -15.647 | -21.702 | 15.034 | 20.655 | -11.140 | 16.199 | 1.044 | 19.888 | -1.499 |
| -18.363 | -12.388 | 15.952 | 25.712 | 19.561 | 2.413 | 13.967 | 1.500 | 9.442 | 5.539 | 26.454 | -8.180 | 7.511 | -26.364 |
| 22.104 | 6.042 | -26.363 | -21.941 | 4.623 | -6.929 | -29.123 | 17.634 | 8.599 | -24.511 | -25.479 | 13.561 | 20.710 | 3.876 |
| 16.292 | 1.405 | -21.888 | -21.345 | 1.140 | -9.247 | -3.136 | -12.724 | 6.627 | 9.646 | 27.581 | -20.687 | 16.527 | -15.751 |
| -23.670 | 25.343 | 9.277 | 26.906 | -6.391 | 6.061 | 28.240 | 21.013 | 12.195 | 12.513 | 29.052 | 21.280 | 29.227 | 13.301 |
| -24.896 | 24.085 | 26.619 | -12.928 | 11.804 | 8.566 | 17.410 | -1.622 | 11.386 | 18.678 | 8.058 | 18.516 | 25.726 | 3.320 |
| 3.005 | -26.472 | 1.822 | -15.330 | 7.466 | -21.296 | -8.940 | -0.653 | -8.682 | -10.175 | -4.834 | 8.949 | 26.835 | 26.642 |
| -26.938 | 23.549 | -15.912 | 29.079 | -13.528 | -25.743 | -19.368 | -25.050 | -17.550 | -27.566 | -28.923 | 7.974 | 23.818 | -5.308 |
| 24.735 | 11.691 | -13.987 | 17.946 | -18.430 | 11.609 | -29.451 | -8.238 | 11.072 | -2.837 | 18.617 | 20.804 | 24.269 | 5.925 |
| -1.639 | 8.452 | 23.029 | -11.773 | -23.738 | 14.108 | 14.349 | -19.935 | 15.286 | -12.208 | -2.668 | 0.966 | 23.117 | 27.461 |
| -16.902 | 13.442 | -2.364 | -23.365 | 3.847 | -7.624 | 6.965 | 25.123 | 1.099 | 7.615 | -6.726 | -12.182 | 2.753 | 16.149 |
| -5.245 | -21.948 | 15.278 | 30.586 | 18.985 | 3.566 | -15.994 | -20.438 | -0.250 | -13.333 | 23.578 | 1.992 | 6.428 | 6.528 |
| -9.650 | 24.384 | -21.439 | -26.032 | 10.556 | -20.729 | -1.977 | 11.382 | -28.135 | -8.187 | -0.378 | -2.789 | -4.660 | 21.643 |
| -12.709 | 9.158 | -9.241 | 2.698 | -9.262 | 27.184 | -22.179 | 0.129 | 22.187 | 23.492 | -3.302 | 16.333 | 10.108 | 23.141 |
| -24.096 | 2.318 | -21.919 | 20.824 | 13.480 | 26.705 | -18.377 | 10.309 | -7.307 | 21.882 | 1.574 | 7.920 | -16.233 | -1.467 |
| 5.373 | -3.898 | 16.075 | 23.606 | 2.318 | -8.104 | -27.672 | -6.787 | 20.900 | -17.907 | -7.476 | 24.591 | 25.174 | -3.931 |
| -21.538 | 15.092 | 5.508 | -6.356 | 8.123 | -25.225 | -25.108 | -3.789 | -3.071 | 6.591 | 21.809 | -12.181 | 18.493 | 25.676 |
| 23.845 | 11.310 | 24.302 | -17.257 | 10.998 | -13.188 | 26.852 | 23.146 | 20.241 | -7.248 | 24.128 | 3.350 | 27.663 | 6.937 |
| -12.063 | -16.121 | -26.224 | 15.757 | -0.831 | 11.131 | -8.590 | -23.711 | -14.909 | 4.760 | -14.076 | 11.421 | 17.896 | 13.888 |
| -24.658 | 19.197 | -13.058 | -16.318 | -2.473 | 6.292 | -19.295 | 13.513 | 8.365 | 14.772 | 27.993 | -20.585 | 23.789 | 16.645 |
| 28.767 | 19.779 | 14.652 | 19.946 | -10.683 | 14.763 | -15.517 | -24.721 | 22.493 | -1.352 | -15.936 | 25.256 | 20.411 | -0.754 |
| -23.438 | 26.145 | 24.478 | -25.838 | -22.600 | -3.755 | 22.193 | 19.454 | -8.210 | 3.951 | -5.294 | 13.508 | 23.415 | 23.751 |
| 8.689 | -9.876 | 18.487 | -6.031 | 20.853 | -14.103 | 14.498 | 16.632 | 13.921 | -18.683 | -10.302 | -6.938 | 26.886 | 11.430 |
| 22.959 | -1.549 | 11.211 | -15.870 | 27.337 | -27.118 | -27.989 | 4.945 | -24.991 | 15.938 | -26.159 | -17.526 | 18.583 | 24.546 |

| | | | | | | | | | | | | | |
|---|---|---|---|---|---|---|---|---|---|---|---|---|---|
| 17.422 | 12.263 | -5.535 | -24.726 | 18.279 | 22.049 | -22.363 | -4.486 | 9.053 | 2.130 | -24.056 | 25.857 | 17.129 | 18.433 |
| -22.065 | -1.187 | -25.145 | -13.959 | 0.634 | -16.016 | -18.976 | -2.313 | -10.103 | -4.064 | 19.974 | 6.151 | 24.066 | 27.830 |
| -9.738 | 22.629 | 15.707 | -19.388 | -0.247 | -24.593 | 20.089 | 19.593 | -11.960 | -5.950 | 29.444 | -0.543 | 22.167 | 21.290 |
| 1.043 | -0.757 | -9.987 | 22.808 | -20.189 | -22.075 | 21.361 | 0.131 | -0.569 | -21.710 | -18.267 | 7.771 | 22.854 | 9.377 |
| -2.645 | -2.816 | -24.730 | -6.691 | 25.990 | 9.919 | -12.421 | -0.446 | 25.903 | 5.181 | 18.947 | 23.822 | 25.397 | 16.258 |
| -25.181 | 13.622 | 23.640 | -14.594 | -4.496 | 2.699 | 20.020 | -25.778 | -27.797 | -3.098 | -15.094 | 12.473 | 17.620 | 10.313 |
| 17.874 | 6.653 | -12.494 | 18.942 | 21.272 | 9.285 | 12.591 | 17.901 | 10.304 | -8.798 | 21.494 | 2.174 | -14.348 | -25.904 |
| 13.112 | -2.136 | -0.232 | -18.799 | 18.624 | 7.328 | -17.635 | 7.725 | -15.145 | 3.062 | 26.189 | 10.727 | 21.797 | -14.676 |
| -19.467 | -20.370 | -25.434 | -20.750 | -21.640 | -10.666 | -21.338 | -19.530 | 1.357 | -21.308 | -0.680 | -26.813 | 16.804 | 21.363 |
| 21.725 | -4.818 | -3.597 | -17.726 | 5.206 | -1.041 | -16.325 | -18.835 | -13.371 | 6.527 | -8.120 | 20.450 | 18.384 | 19.271 |
| -6.370 | -9.327 | 17.468 | 29.050 | 28.414 | -11.719 | -8.170 | -2.662 | 14.236 | -20.278 | -14.749 | 18.817 | 27.864 | 23.029 |
| -26.273 | -7.481 | 21.047 | -1.167 | 15.778 | -25.633 | -10.452 | -6.023 | 20.448 | 19.098 | 24.485 | 22.330 | 24.520 | 22.410 |
| 11.340 | 7.877 | 22.583 | 18.816 | 5.671 | 13.122 | -22.562 | -0.747 | 2.214 | 21.462 | 10.793 | 13.217 | 23.047 | 23.804 |
| -14.543 | 4.182 | 9.911 | 14.268 | -11.089 | 21.023 | 2.329 | -2.055 | 27.952 | 26.961 | 6.368 | -2.566 | -22.370 | -0.882 |
| 1.898 | 2.107 | 13.135 | -23.619 | -24.572 | -13.189 | -4.292 | 26.783 | -4.296 | 11.181 | -5.905 | 3.083 | 24.160 | -26.228 |
| 6.613 | 5.980 | 6.128 | -30.955 | -18.865 | 6.916 | -15.744 | -24.629 | -3.482 | -16.730 | -8.841 | -7.853 | 15.646 | 15.877 |
| -25.409 | -1.493 | 11.738 | 14.099 | -26.039 | 16.402 | -18.534 | 5.782 | -14.095 | -23.889 | 28.658 | -25.617 | 18.180 | -4.457 |
| -25.888 | 20.604 | 14.613 | -4.989 | 4.162 | -4.560 | -25.946 | 7.921 | 4.750 | 9.236 | -3.038 | 9.764 | 3.268 | 21.937 |
| 27.709 | -12.321 | -6.267 | -25.547 | -3.079 | 16.870 | -9.338 | -26.008 | -25.636 | -11.028 | 25.283 | 26.842 | 12.595 | 18.242 |
| 2.501 | 12.883 | -10.086 | 23.892 | 25.820 | -7.079 | -8.685 | 15.143 | -6.223 | -24.030 | -27.751 | -26.416 | 22.159 | -1.168 |
| 14.475 | 10.152 | -16.427 | 19.002 | 19.814 | 15.697 | -9.877 | 13.029 | 20.878 | 21.284 | -3.817 | 23.000 | -3.521 | 14.949 |
| 7.713 | 13.607 | 16.475 | -7.637 | 0.087 | -5.552 | 28.637 | 6.458 | 13.367 | -4.266 | -19.244 | 4.362 | 11.811 | 8.402 |
| 3.110 | 2.313 | -7.837 | -8.699 | -13.889 | -18.907 | -1.023 | 4.212 | 23.573 | -13.155 | 28.196 | -5.583 | 18.984 | -19.803 |
| 15.234 | 6.473 | 12.974 | -28.311 | 22.385 | 21.675 | -14.780 | -20.056 | -0.162 | -22.248 | -21.171 | 27.327 | 22.296 | -21.666 |
| -13.487 | 0.355 | 14.071 | -25.551 | 8.863 | 8.280 | 27.957 | 5.874 | 5.911 | 13.664 | -23.778 | -23.574 | 19.342 | 27.105 |
| 20.958 | -11.705 | -11.037 | 20.972 | -3.706 | 22.781 | 21.444 | -23.354 | 27.072 | -10.670 | 25.963 | 24.291 | -1.734 | -3.659 |
| -20.806 | -5.101 | -12.210 | -16.439 | 6.050 | 17.625 | 23.039 | -0.490 | -10.430 | 7.854 | 12.355 | -11.340 | 17.562 | 20.729 |
| -24.438 | -2.471 | 19.429 | -18.819 | 1.039 | -17.646 | 20.663 | -27.682 | -11.013 | -7.126 | 23.212 | 10.995 | 25.181 | 12.855 |
| 25.968 | -4.274 | 24.687 | 10.914 | 27.468 | -20.667 | 19.251 | 12.287 | 10.975 | -25.566 | 0.457 | 15.033 | 19.994 | 5.491 |
| 20.369 | 15.210 | -17.998 | -10.751 | 18.959 | -13.065 | -18.631 | -0.403 | -17.978 | -5.177 | -26.886 | -24.237 | 26.685 | 1.526 |
| 8.390 | -14.958 | -14.601 | 18.165 | -1.299 | -8.394 | -23.737 | -7.610 | 9.652 | 21.510 | 8.102 | -6.271 | 18.043 | 10.226 |
| 0.791 | -9.215 | 8.509 | -19.478 | -6.376 | 0.656 | 30.420 | 16.897 | 8.091 | 6.016 | -16.090 | -22.438 | 2.782 | -2.228 |
| 9.024 | -5.616 | -22.903 | -29.944 | 25.152 | 3.931 | -19.846 | 26.091 | 25.892 | 17.465 | -13.919 | 24.897 | 21.193 | 24.424 |
| 23.650 | 14.863 | -21.747 | -21.190 | 15.416 | -6.704 | -25.669 | -0.187 | 18.607 | 4.480 | -26.440 | 3.916 | -26.725 | 6.369 |
| -10.483 | -14.015 | 7.838 | 1.973 | 1.377 | -14.600 | -9.438 | -4.850 | -25.073 | -8.929 | -13.211 | -25.984 | -7.238 | -5.431 |
| 12.622 | 1.705 | 16.621 | 6.223 | -16.601 | -23.597 | -30.536 | -17.874 | 7.542 | -2.324 | 27.007 | 26.052 | 24.093 | -16.531 |

| | | | | | | | | | | | | | |
|---|---|---|---|---|---|---|---|---|---|---|---|---|---|
| 27.066 | -20.678 | 19.811 | 24.430 | -10.509 | -2.216 | 27.934 | -15.617 | -9.399 | 11.327 | 16.551 | -18.795 | 25.290 | 7.123 |
| -11.549 | -0.585 | -22.282 | -20.616 | 23.795 | 13.766 | 21.550 | 18.040 | 24.463 | -1.251 | 14.258 | 11.892 | 22.941 | 18.262 |
| -25.513 | -7.253 | 9.832 | -18.942 | 17.212 | -20.603 | -28.431 | 23.219 | -7.976 | -16.036 | 3.206 | 3.477 | 15.731 | 21.751 |
| 25.903 | 1.434 | -27.276 | 25.729 | 4.863 | 15.705 | -2.945 | 9.886 | 26.472 | 0.424 | 15.478 | -5.670 | 5.485 | 3.885 |
| -17.015 | 0.829 | 14.318 | -29.857 | -18.182 | 20.372 | -13.893 | -23.740 | -19.150 | -4.686 | 24.582 | 27.363 | 6.282 | 8.471 |
| 5.611 | -6.573 | -16.375 | -19.083 | 13.876 | 12.561 | 17.009 | -13.001 | -26.551 | 13.243 | 23.402 | 9.330 | -0.810 | 25.035 |
| 3.583 | -16.527 | -15.323 | -17.817 | -9.821 | -0.635 | -19.424 | -21.429 | 6.614 | -21.601 | 7.605 | 21.119 | -6.689 | 2.403 |
| 7.776 | -12.632 | -26.170 | -12.552 | 11.079 | -18.530 | -10.219 | -7.571 | -10.730 | 9.688 | -2.068 | 21.731 | -9.175 | 16.648 |
| 6.746 | -7.945 | -1.419 | -24.248 | -25.348 | -6.148 | 10.914 | 29.124 | -16.475 | -3.988 | 7.002 | 19.918 | -23.430 | 17.846 |
| 22.700 | -9.023 | -14.821 | 22.734 | -15.489 | -5.393 | -24.173 | -0.287 | 16.378 | 6.337 | -3.192 | 7.631 | -2.070 | 20.009 |
| -14.887 | -16.382 | -21.504 | -3.854 | -7.377 | 6.696 | -11.185 | -19.250 | 19.734 | 12.056 | 22.015 | 12.906 | -5.168 | -22.381 |
| -19.069 | -21.685 | 11.128 | 21.068 | 13.105 | 2.054 | 18.605 | -17.464 | 22.165 | 16.761 | -13.622 | 22.916 | 5.684 | 6.820 |
| 4.544 | 19.928 | 2.000 | -6.115 | -23.792 | 20.233 | -30.895 | 11.252 | 12.385 | -0.279 | 20.540 | 20.510 | -1.067 | 23.417 |
| -8.215 | 1.135 | 24.019 | -8.339 | 1.509 | 25.905 | 26.244 | -25.416 | -3.389 | -27.976 | -17.960 | -1.249 | 0.334 | 20.730 |
| 1.680 | -10.739 | 22.858 | -4.059 | -0.017 | -17.109 | 1.968 | -8.352 | -1.925 | 1.093 | 22.992 | 13.216 | -1.208 | 25.238 |
| -26.368 | -15.311 | 21.532 | -22.916 | -0.636 | 5.682 | 16.506 | -7.311 | 7.996 | -9.251 | 26.822 | 20.094 | 24.527 | 18.225 |
| 22.642 | -2.193 | 14.979 | -19.364 | -13.194 | 24.649 | -8.443 | -9.514 | 24.827 | -18.118 | 1.183 | 29.421 | 0.912 | 7.368 |
| -22.362 | -19.729 | -9.186 | -13.248 | 22.090 | -18.983 | 22.387 | -26.400 | -4.378 | 5.373 | 24.137 | 18.360 | 5.716 | -0.523 |
| -7.663 | 1.488 | 11.889 | 18.865 | 7.690 | 8.848 | -29.941 | -8.932 | 2.081 | 21.223 | -6.152 | 3.117 | -27.203 | -3.896 |
| 14.743 | -20.811 | 24.543 | -5.405 | -6.373 | 15.745 | 19.154 | 22.882 | 16.494 | -6.936 | -17.109 | -11.740 | 14.708 | 13.026 |
| 20.717 | 9.472 | 20.177 | -17.557 | 0.817 | -23.250 | -12.037 | 9.343 | -26.918 | 26.824 | 25.339 | -8.549 | 25.119 | 24.406 |
| -3.353 | -15.698 | 9.684 | -25.461 | -8.076 | 9.303 | 5.114 | -18.707 | -5.034 | 0.715 | -3.754 | -19.822 | -9.631 | 9.257 |
| 23.431 | -16.415 | 0.181 | -10.713 | 18.579 | 23.808 | -25.134 | 28.434 | 15.035 | -9.841 | 18.725 | 1.798 | 17.596 | -12.203 |
| 6.772 | -22.157 | 22.194 | 15.122 | 6.304 | 21.095 | 16.815 | 4.668 | 5.372 | -1.577 | -17.940 | -12.352 | 23.592 | 8.823 |
| 4.177 | 20.103 | -4.743 | 27.494 | 2.400 | -25.160 | 6.781 | -13.635 | 3.002 | 5.762 | 24.937 | 3.398 | 23.711 | 23.876 |
| 18.684 | -8.215 | -22.538 | 19.414 | -22.216 | 20.606 | -29.121 | -30.047 | 11.318 | 2.285 | 17.343 | -0.571 | 11.374 | -2.131 |
| -14.731 | 7.862 | 8.731 | 8.546 | -2.811 | -23.072 | 30.433 | 9.577 | 2.511 | -17.040 | 22.822 | 29.117 | 24.044 | 12.538 |
| 22.159 | 15.993 | 12.862 | -12.563 | -5.636 | 13.270 | 18.339 | -8.895 | -4.034 | -2.964 | -10.661 | 11.588 | 26.083 | 0.040 |
| 19.937 | 4.206 | 16.097 | -0.942 | 4.767 | 12.350 | -5.880 | 4.210 | -17.090 | 14.893 | 19.285 | 29.807 | 22.768 | -4.284 |
| -24.784 | 7.410 | 4.255 | -17.108 | 1.674 | 26.119 | -27.936 | -24.581 | -26.231 | 6.708 | 16.586 | -1.402 | 22.102 | -13.618 |
| 5.034 | 3.159 | 6.532 | -19.643 | 8.175 | 27.117 | 8.934 | -16.804 | 8.319 | 16.018 | -6.602 | 26.187 | 18.897 | 9.882 |
| 20.671 | 1.533 | 18.596 | -1.865 | 25.750 | -20.495 | 21.222 | 28.255 | 20.113 | 3.317 | -10.107 | 24.651 | 14.678 | 5.810 |
| 22.147 | 11.084 | 3.685 | -0.423 | 22.736 | -17.495 | -14.271 | -0.591 | 9.502 | -21.826 | 21.613 | 2.552 | 23.262 | 4.046 |
| -16.388 | -17.837 | -8.853 | 9.937 | -1.094 | 23.171 | 6.666 | -13.708 | 18.789 | 8.130 | 14.535 | -19.602 | 15.992 | -9.375 |
| 25.664 | -19.226 | 21.479 | -19.771 | -1.621 | -24.256 | 24.586 | 3.041 | 6.872 | 2.628 | -18.767 | -26.146 | 21.457 | -7.412 |
| 24.929 | -6.537 | -17.539 | -7.243 | 20.649 | -18.853 | 6.375 | 19.400 | 1.363 | -4.050 | 9.843 | -1.960 | 24.709 | 0.528 |

| | | | | | | | | | | | | | |
|---|---|---|---|---|---|---|---|---|---|---|---|---|---|
| -13.921 | 16.633 | -2.472 | 27.797 | 11.791 | -20.995 | 5.404 | 6.706 | -8.089 | 24.412 | 7.673 | -13.975 | 27.928 | 22.169 |
| -17.238 | 22.040 | 0.776 | -0.680 | 20.167 | 8.901 | 29.818 | 14.344 | -10.707 | -25.769 | 0.795 | 8.762 | -5.604 | 21.568 |
| -22.711 | -4.601 | -14.720 | -4.001 | -4.494 | -25.961 | 30.781 | -11.589 | -23.870 | -13.477 | -11.334 | 23.758 | 19.825 | 14.550 |
| 12.098 | 22.967 | -6.068 | -22.583 | -13.768 | 20.886 | -12.658 | 30.980 | -11.943 | -10.788 | 19.070 | 28.466 | 22.752 | 23.037 |
| 4.547 | 2.644 | 28.050 | 15.519 | 15.115 | 17.231 | -8.051 | 19.190 | 20.751 | -18.690 | 10.706 | -28.050 | 28.082 | 13.135 |
| -12.205 | -2.723 | 19.855 | -11.786 | -22.942 | 5.131 | -6.290 | -6.882 | -24.307 | 4.125 | 26.098 | -10.357 | -26.058 | -7.676 |
| 0.968 | 1.399 | -12.320 | -26.261 | 7.858 | 24.101 | 22.391 | -25.880 | 3.985 | -9.395 | -1.671 | -0.966 | 22.366 | -14.110 |
| -19.620 | -17.433 | -0.286 | -2.768 | 6.876 | 19.321 | 10.564 | -17.514 | 19.789 | 0.041 | 6.503 | 29.055 | 19.315 | -21.784 |
| -18.818 | -5.627 | -17.450 | -18.094 | 10.196 | -22.034 | 8.262 | 13.961 | 11.457 | 11.033 | 22.352 | -20.422 | 23.633 | -2.073 |
| 3.308 | 4.691 | 14.644 | -12.884 | -6.972 | 2.320 | 16.705 | 22.565 | 15.881 | 4.632 | 0.883 | 19.993 | 14.807 | -6.273 |
| 25.703 | -7.605 | 9.011 | 17.461 | 4.730 | -10.368 | -1.878 | -2.030 | 7.486 | 13.303 | -25.467 | 11.034 | 7.498 | -4.475 |
| 7.680 | 7.300 | 1.177 | -3.668 | -21.314 | 24.894 | 17.940 | -31.375 | -5.570 | -2.757 | -5.543 | -26.877 | -2.506 | -11.117 |
| 8.719 | -25.892 | 18.592 | -14.086 | 3.647 | -26.591 | -14.281 | -9.320 | -9.413 | 1.692 | -1.038 | -6.213 | 22.498 | 4.883 |
| -21.691 | 24.959 | 4.149 | 12.143 | 21.225 | 19.885 | 12.105 | -6.349 | -3.228 | -5.895 | -22.228 | 12.434 | 12.685 | 18.021 |
| 15.013 | 5.824 | -15.336 | -6.939 | 12.796 | 14.109 | -12.063 | -15.314 | 6.786 | -4.906 | -7.210 | -15.809 | -8.638 | -2.936 |
| 10.267 | -20.144 | 23.815 | 4.214 | 10.642 | -24.465 | 4.425 | -1.134 | 0.772 | -6.715 | -17.478 | 22.215 | 20.317 | -21.264 |
| 12.106 | 18.300 | 14.039 | -27.214 | -3.170 | -23.927 | 5.635 | -30.548 | -7.203 | 12.517 | 2.205 | -30.060 | 13.913 | -2.554 |
| 0.240 | 9.582 | 2.330 | -0.801 | -17.334 | 27.478 | 2.097 | -16.881 | 4.853 | 6.959 | -4.428 | 17.124 | 26.368 | 9.313 |
| -20.477 | 24.252 | -3.603 | 20.359 | 26.553 | 26.592 | -19.334 | -27.036 | 2.230 | 3.258 | 3.978 | -17.907 | -6.952 | -7.648 |
| -2.111 | -0.564 | 17.773 | 0.572 | 9.670 | 14.658 | -26.286 | 25.869 | 19.426 | 7.858 | -12.750 | -5.639 | -13.397 | 24.148 |
| 5.455 | -17.832 | 8.256 | 11.675 | 8.974 | 7.069 | 14.591 | 3.053 | 17.067 | -7.472 | 23.494 | -0.024 | 9.743 | -13.507 |
| 23.696 | 2.333 | -7.280 | 9.504 | -12.757 | -4.568 | -20.220 | -10.972 | 14.920 | -11.259 | 8.303 | -15.740 | 19.524 | -1.349 |
| 15.746 | 12.552 | -7.731 | 24.438 | 5.549 | 22.273 | 9.103 | -21.139 | -1.127 | 5.316 | -1.828 | -30.478 | 18.171 | 6.145 |
| -21.405 | 12.145 | 28.455 | -8.312 | 23.262 | 15.245 | 18.838 | 24.171 | -1.741 | -23.784 | -18.883 | 21.135 | 24.994 | -0.052 |
| -10.306 | 19.375 | 18.542 | -5.587 | 16.810 | 22.989 | -17.387 | -14.536 | 1.612 | 11.141 | 9.278 | 29.091 | 21.591 | 4.536 |
| -2.950 | -21.758 | 16.724 | -3.271 | -14.913 | -18.004 | 20.677 | -8.149 | 9.415 | 20.480 | 24.147 | 25.983 | 20.926 | 3.947 |
| -23.707 | 0.632 | 15.312 | -4.561 | -9.507 | 3.818 | -1.696 | -9.624 | 0.195 | -4.610 | -10.280 | 27.531 | 26.446 | 17.045 |
| 1.140 | -6.134 | 0.732 | -20.615 | 17.602 | 16.395 | 20.130 | 29.237 | -6.893 | -16.083 | 10.857 | 31.271 | 23.699 | 6.524 |
| 5.019 | 16.329 | 26.997 | -26.237 | 13.486 | -14.972 | -14.812 | 9.302 | -7.890 | 25.155 | 1.511 | 13.129 | 12.418 | 5.220 |
| 0.640 | -5.041 | -1.137 | -0.896 | 11.930 | 27.989 | 21.011 | -14.971 | 22.463 | 9.792 | -6.623 | 1.373 | 27.965 | 20.104 |
| -22.453 | 22.571 | -12.545 | 25.850 | 3.006 | 23.619 | -20.618 | -21.083 | 4.628 | -9.056 | 2.212 | 30.409 | 22.820 | 2.220 |
| 9.139 | 14.333 | -3.606 | -21.331 | 11.239 | 0.114 | -6.295 | 17.483 | 13.589 | -14.853 | 5.537 | 5.245 | 19.504 | -3.603 |
| 14.409 | 18.278 | -29.658 | -2.824 | 1.307 | 26.105 | 12.028 | 9.799 | 5.691 | 24.195 | -20.204 | -3.268 | 27.336 | 8.443 |
| 0.061 | 10.074 | 4.745 | 15.607 | -19.654 | 17.373 | -9.528 | 13.962 | -8.479 | -6.002 | 26.026 | -19.726 | -23.371 | -5.623 |
| 15.899 | 13.710 | 23.696 | -15.913 | 22.210 | -23.985 | 24.590 | 7.456 | -18.167 | 22.331 | 16.061 | 28.904 | 6.864 | -24.954 |
| -21.000 | 13.271 | -4.940 | -5.109 | 19.793 | 20.184 | 19.251 | -13.615 | 17.293 | -2.846 | 21.758 | -10.727 | 20.784 | 15.275 |

| | | | | | | | | | | | | | |
|---|---|---|---|---|---|---|---|---|---|---|---|---|---|
| 4.301 | 2.358 | 28.975 | 25.966 | 17.331 | 27.535 | -4.761 | 22.544 | 7.701 | 11.859 | -1.567 | 22.265 | 26.437 | 6.891 |
| -8.883 | 8.573 | -0.282 | -20.560 | -21.505 | -19.752 | -23.786 | -6.955 | -14.465 | 5.764 | 20.718 | -2.691 | 19.807 | -0.014 |
| -18.760 | 0.629 | -28.582 | 2.992 | 16.643 | 24.370 | -5.569 | 17.952 | 21.599 | -5.368 | -0.298 | -4.209 | 13.406 | -6.878 |
| 25.182 | -22.265 | 13.994 | 16.741 | -7.489 | 2.541 | -24.949 | -11.424 | 14.379 | -9.670 | 7.505 | 18.334 | 22.839 | -2.713 |
| 19.336 | -22.905 | -26.089 | -8.466 | -11.236 | 15.822 | -14.794 | 17.519 | 2.645 | -22.869 | 19.197 | -26.442 | 14.807 | 16.453 |
| -20.185 | -2.851 | -8.340 | 10.345 | 24.264 | 16.817 | -6.817 | 15.713 | -13.199 | 5.280 | 17.977 | 20.872 | 20.413 | -5.022 |
| 15.315 | -1.172 | 6.778 | -8.050 | -12.947 | 5.638 | 20.843 | 18.610 | 19.708 | -10.620 | -23.452 | 19.027 | 24.964 | -8.138 |
| 2.972 | 17.047 | -1.558 | -4.994 | 2.503 | -17.016 | 2.037 | 16.136 | 10.482 | 0.636 | 5.772 | 24.865 | 24.048 | -10.159 |
| -12.754 | 23.154 | -6.536 | 17.231 | 8.212 | -24.105 | -5.519 | -28.058 | -5.027 | 9.025 | 17.241 | 13.351 | 24.520 | 20.328 |
| -22.335 | -3.787 | 3.817 | 14.969 | 5.722 | 18.030 | -22.998 | -21.017 | -4.531 | 7.103 | 25.575 | -0.810 | 22.125 | 3.852 |
| 23.654 | 26.064 | -21.500 | 3.513 | -3.891 | -27.477 | 25.843 | -29.880 | -11.851 | 15.886 | -6.912 | 5.512 | 27.806 | -24.970 |
| 22.774 | 8.025 | 29.664 | -14.636 | -16.782 | -19.173 | 5.810 | -16.808 | -2.392 | 9.544 | 9.126 | 4.131 | 10.266 | 13.017 |
| 14.018 | -24.036 | -12.516 | -24.804 | 7.248 | 27.726 | 1.629 | 26.687 | -16.097 | 1.213 | -3.286 | 28.108 | 22.898 | -13.357 |
| -21.674 | -24.886 | 3.483 | 21.977 | -15.167 | 19.560 | 22.038 | -13.068 | 15.383 | 23.273 | 21.427 | -1.739 | 20.812 | 11.086 |
| 16.071 | -3.121 | -12.096 | -9.349 | -21.014 | -27.242 | 16.347 | 14.599 | -12.135 | 2.046 | 4.754 | -30.897 | 28.896 | 19.067 |
| -24.191 | 10.260 | -14.052 | -23.460 | -8.623 | -26.128 | -10.279 | -12.758 | 5.895 | -5.975 | -22.357 | -28.317 | 23.796 | 18.179 |
| 25.268 | -8.269 | 16.163 | 4.919 | 25.594 | -15.260 | -14.122 | -22.315 | -0.692 | 18.098 | 15.492 | 31.776 | 26.282 | 10.151 |
| 8.122 | -14.549 | -10.718 | 4.278 | -19.238 | -21.293 | -16.497 | -4.152 | 5.441 | 6.465 | 11.241 | 23.595 | 8.054 | 17.642 |
| -4.031 | 20.155 | -0.866 | -5.560 | 16.066 | 6.396 | -12.210 | 20.192 | -22.309 | -3.471 | 20.837 | 15.894 | -8.449 | 12.128 |
| 22.786 | 19.470 | 16.580 | -8.277 | 8.806 | 20.810 | 1.896 | 13.238 | -5.804 | 19.264 | 6.091 | -5.160 | 27.372 | 21.254 |
| -1.949 | 3.820 | -6.482 | 18.743 | -2.393 | 10.638 | 11.155 | 9.901 | -8.095 | 22.185 | -11.756 | 1.637 | 25.563 | 0.554 |
| 2.504 | -6.800 | -17.508 | 4.640 | -11.470 | -10.855 | -5.568 | 5.389 | 4.078 | -26.226 | 0.561 | 24.807 | 18.304 | 12.977 |
| 16.417 | -10.954 | -18.906 | -9.721 | 11.074 | 23.312 | 13.777 | 12.363 | 14.305 | 1.348 | -9.094 | -19.094 | 22.404 | 18.144 |
| -6.551 | 22.852 | 5.857 | -25.051 | 8.152 | -4.420 | -23.867 | -22.414 | 8.892 | -11.272 | 4.691 | 30.270 | 23.918 | 2.900 |
| -8.322 | 1.409 | 24.161 | -5.598 | -7.942 | -19.240 | 22.384 | 10.767 | -2.814 | -18.880 | 14.310 | 27.303 | 19.927 | 15.825 |
| -19.587 | -20.185 | 13.593 | -5.016 | -2.753 | 17.617 | -6.302 | -9.195 | -10.567 | 9.017 | -1.786 | 0.979 | 29.093 | -0.899 |
| -20.150 | 10.405 | 21.613 | 7.143 | -10.585 | 24.718 | 8.547 | -28.681 | -11.268 | 7.609 | -5.432 | 8.989 | 26.139 | 1.489 |
| 19.310 | 6.356 | 28.806 | 9.438 | -0.719 | 9.349 | 2.352 | 14.069 | 9.343 | -7.012 | 16.286 | 26.450 | 25.232 | 14.332 |
| -26.073 | -0.079 | 3.577 | -7.473 | 17.104 | 29.124 | 10.111 | 0.251 | -23.575 | -4.619 | 3.842 | 21.669 | 12.460 | 15.221 |
| 21.934 | 26.249 | -14.510 | -20.653 | 9.472 | -22.285 | 21.248 | 1.991 | 3.496 | 1.771 | 11.162 | -31.528 | 0.143 | 19.826 |
| 10.638 | -10.487 | -3.355 | -17.863 | 17.127 | 20.773 | 9.634 | 25.031 | 17.352 | 2.331 | -24.428 | 3.983 | 23.159 | 16.545 |
| 3.499 | 21.056 | -0.325 | -20.781 | 14.894 | 27.199 | 14.171 | 14.582 | -4.697 | -21.286 | 20.260 | -26.187 | 29.219 | 21.113 |
| -13.359 | -23.620 | -1.637 | -22.523 | -8.346 | 19.666 | 4.487 | 17.366 | 19.483 | 15.756 | 5.439 | -8.410 | 18.243 | 13.522 |
| 23.400 | 19.916 | 29.446 | -4.830 | 3.386 | 17.665 | 18.482 | -23.329 | 0.498 | 19.658 | 12.926 | 11.418 | 12.862 | 17.472 |
| -0.911 | -7.319 | 28.239 | -1.683 | -9.541 | -7.777 | -9.190 | -25.601 | -10.854 | -3.739 | 10.250 | 0.149 | 26.984 | 4.785 |
| -9.480 | 3.310 | 26.506 | 1.818 | 12.707 | 30.088 | -3.282 | -10.235 | -3.432 | -9.353 | 9.510 | 26.192 | 24.254 | -12.098 |

| | | | | | | | | | | | | | |
|---|---|---|---|---|---|---|---|---|---|---|---|---|---|
| 15.444 | 7.450 | 24.333 | 1.093 | -10.549 | 28.368 | 8.763 | 1.527 | -7.332 | 13.976 | 7.064 | -23.449 | 15.887 | 15.894 |
| 10.301 | 18.617 | 20.737 | -0.738 | 0.760 | 24.921 | -6.210 | -11.096 | -3.960 | -1.679 | -17.115 | 29.333 | 27.693 | -15.479 |
| -22.604 | -1.676 | -1.252 | 15.190 | -5.633 | -20.728 | -20.406 | -17.265 | -18.103 | -16.908 | -0.666 | -20.910 | 20.495 | 22.201 |
| -6.460 | 25.277 | 1.585 | 17.588 | 11.956 | 21.632 | -7.816 | -14.657 | 11.992 | -12.532 | -0.036 | 28.067 | 26.164 | 18.375 |
| 0.584 | 21.978 | -26.161 | -6.828 | -8.361 | -10.209 | -11.379 | 27.275 | 5.007 | -14.369 | -25.383 | -0.919 | 14.690 | 11.503 |
| 24.640 | -24.873 | -25.391 | -2.101 | -16.034 | 18.941 | -26.133 | 15.236 | -17.387 | 9.932 | 25.242 | 24.880 | 27.313 | 10.029 |
| -21.888 | -20.285 | -6.113 | -3.367 | 13.140 | 28.099 | -20.961 | 21.790 | -1.686 | 5.173 | -6.176 | 26.613 | 7.168 | 14.487 |
| 23.397 | 28.412 | 25.107 | 21.207 | 14.263 | 29.358 | 11.701 | -14.329 | 14.394 | -8.405 | 24.918 | 13.617 | 12.253 | 0.528 |
| 20.711 | 8.129 | 15.682 | -22.758 | -10.612 | 25.976 | 18.175 | -2.241 | 18.947 | -16.273 | 10.365 | -11.299 | 29.834 | 23.832 |
| 3.364 | 13.530 | 18.632 | 2.182 | -4.901 | 17.993 | 9.521 | 7.009 | -8.675 | 12.231 | -7.705 | -17.809 | 21.537 | 10.664 |
| -20.506 | -7.933 | 12.898 | 7.351 | 3.789 | -9.304 | -16.559 | 4.938 | 10.471 | -10.656 | -24.233 | -9.328 | 31.134 | -2.012 |
| -25.683 | -26.008 | 6.988 | 26.033 | -22.772 | 10.538 | 18.449 | -6.798 | -11.182 | -13.594 | -19.268 | 6.473 | 10.906 | 17.904 |
| 19.341 | -22.014 | 4.866 | -11.152 | 24.287 | 0.098 | 7.337 | 15.618 | 12.323 | 6.232 | -14.516 | 25.870 | -15.352 | 25.821 |
| -3.925 | -19.745 | -15.858 | 15.848 | -24.561 | -3.649 | 2.914 | 6.470 | 19.423 | -13.210 | 26.509 | 18.004 | 19.172 | 17.016 |
| -10.316 | -27.129 | 10.909 | 5.760 | -3.935 | -28.953 | 10.070 | -20.052 | -14.465 | 6.917 | -25.072 | 21.000 | 27.017 | -16.503 |
| -24.773 | 2.435 | 26.519 | 16.162 | 3.322 | 23.088 | 1.209 | 10.578 | -21.539 | -22.236 | 2.183 | 12.215 | 22.951 | 19.909 |
| -23.439 | -24.200 | 16.562 | -25.942 | 2.474 | -14.625 | 6.291 | 17.441 | 10.073 | -19.251 | -20.025 | 0.755 | 22.012 | 2.080 |
| 5.426 | 25.639 | -8.269 | 7.238 | -8.834 | -22.021 | 11.999 | -18.045 | 22.022 | 10.502 | -20.854 | 27.852 | 25.153 | 0.672 |
| 4.909 | 7.606 | -25.588 | 17.572 | 1.799 | 25.566 | -4.720 | 3.664 | -7.541 | 1.240 | -6.835 | 26.442 | 21.170 | 21.166 |
| 27.350 | 4.076 | -6.785 | -24.229 | -7.933 | 28.624 | 21.327 | -13.167 | 14.714 | -12.375 | -22.486 | 3.680 | 23.719 | 20.360 |
| -26.767 | -8.339 | -24.590 | -22.516 | -6.798 | -6.401 | 3.216 | 23.611 | -16.640 | -15.891 | 26.435 | 19.498 | 6.935 | 23.822 |
| 24.578 | 27.165 | 7.622 | 5.252 | -17.409 | 14.890 | 5.012 | 6.801 | -8.074 | -18.638 | 20.054 | 28.690 | 22.261 | 23.260 |
| 4.559 | -4.466 | 27.667 | -10.125 | 1.059 | 1.815 | 11.244 | -27.265 | -17.889 | 7.101 | 1.509 | -25.344 | 14.547 | 8.019 |
| 20.565 | -10.965 | 5.829 | -7.765 | 22.531 | 9.861 | -8.135 | -17.981 | 25.124 | 18.492 | -0.378 | 16.456 | 19.078 | 24.988 |
| -24.248 | -26.454 | 16.968 | -12.957 | -14.987 | 26.083 | -7.756 | 21.891 | 15.579 | -10.150 | -26.798 | 23.051 | -12.756 | 5.495 |
| -18.930 | 15.336 | 28.846 | 3.468 | -25.392 | 11.214 | -20.810 | -9.266 | -8.812 | -11.483 | -23.822 | 14.452 | 29.952 | 20.607 |
| 27.533 | 2.771 | -18.126 | -9.196 | 24.945 | 4.059 | 7.159 | 19.887 | 23.933 | 6.521 | -16.569 | 4.414 | 7.659 | 23.560 |
| 11.383 | 14.374 | -21.259 | 20.780 | -19.616 | 13.728 | -24.615 | -21.017 | 7.100 | -25.951 | -6.144 | -13.818 | 20.460 | 11.199 |
| 23.998 | 19.535 | 18.930 | 18.965 | 7.232 | 19.104 | 5.264 | 23.616 | 3.434 | 14.817 | -26.009 | 4.887 | 25.808 | -0.300 |
| -9.987 | -18.485 | 31.243 | -22.745 | 20.766 | -22.553 | 9.871 | -17.679 | 2.016 | 4.172 | 24.370 | -27.830 | 23.892 | 18.331 |
| -8.872 | 6.491 | -19.340 | -6.539 | 23.655 | 23.401 | 1.238 | 21.471 | 13.653 | -1.392 | 24.225 | 29.517 | 29.013 | 26.040 |
| -27.450 | 8.640 | -15.257 | -3.693 | 19.974 | -11.861 | -9.597 | -0.251 | 10.178 | -4.359 | -20.403 | -5.250 | -0.256 | 26.921 |
| 0.737 | 20.442 | -25.878 | -10.184 | -4.326 | -17.658 | -25.300 | -0.932 | -24.430 | -6.429 | 22.431 | -28.273 | 9.335 | 26.433 |
| 21.560 | 23.699 | 20.305 | -24.219 | 22.143 | 22.901 | -6.547 | -3.626 | 14.977 | 15.592 | 14.301 | -19.953 | 20.920 | 17.231 |
| 11.973 | -16.288 | 21.477 | 27.449 | 22.531 | 29.611 | -10.742 | -16.179 | -25.092 | 23.357 | -25.755 | 22.676 | 24.628 | 21.715 |
| -5.504 | 0.487 | -3.846 | -5.214 | 6.498 | 11.772 | 20.051 | 3.665 | 17.468 | 16.583 | -11.705 | 1.883 | 16.721 | 24.916 |

| | | | | | | | | | | | | | |
|---|---|---|---|---|---|---|---|---|---|---|---|---|---|
| 14.640 | 4.247 | -2.915 | -27.381 | 19.081 | -19.136 | -14.524 | 4.032 | -2.809 | -0.232 | -6.693 | 19.032 | 5.331 | 19.971 |
| -2.503 | -17.933 | 19.485 | -6.072 | -20.481 | -20.440 | 23.809 | -3.740 | 16.520 | 6.076 | 9.644 | 27.629 | -7.306 | -25.576 |
| -26.457 | 22.031 | 5.188 | -21.555 | -24.421 | 28.177 | 19.414 | 12.388 | 4.295 | 4.904 | 27.499 | 3.217 | 27.710 | -9.849 |
| -4.168 | 6.571 | -9.569 | -13.157 | 2.459 | -1.485 | -8.146 | -21.340 | 23.171 | -5.806 | 14.677 | 29.203 | 15.465 | 23.741 |
| -0.784 | 27.457 | -10.444 | -19.259 | -6.850 | 16.954 | 12.215 | 17.681 | -14.264 | -3.228 | -16.560 | 27.238 | -26.524 | 16.718 |
| 13.863 | -0.888 | 9.579 | 18.758 | 5.305 | -22.091 | -4.460 | 13.267 | 10.433 | -4.805 | 25.503 | 22.886 | 28.402 | 20.297 |
| -24.245 | -8.443 | 6.290 | 15.726 | -16.719 | 6.136 | 2.517 | 11.714 | 4.622 | -1.369 | 22.931 | 7.183 | 20.667 | 14.637 |
| -6.351 | -25.878 | -13.415 | -24.981 | -7.309 | -16.252 | 9.338 | -10.941 | -16.119 | 5.686 | 4.356 | 15.953 | 23.010 | 18.777 |
| 27.440 | 17.571 | 25.707 | -6.227 | -0.852 | 12.931 | 7.357 | 10.194 | -21.957 | 6.966 | 1.853 | 14.664 | 25.962 | 27.226 |
| 24.527 | -9.154 | 7.062 | -19.497 | 10.956 | 0.163 | -20.220 | 2.812 | -23.630 | 22.250 | -19.147 | 9.971 | 22.757 | 16.433 |
| 25.728 | 16.934 | 0.487 | -17.537 | -2.930 | -15.792 | -16.273 | 4.883 | -12.725 | -4.220 | -8.208 | -3.955 | 23.083 | -6.896 |
| 18.730 | 8.757 | -12.093 | 14.859 | -5.153 | -16.731 | 20.556 | 3.956 | 12.520 | 14.793 | -1.831 | 0.318 | 18.231 | 22.415 |
| 5.423 | 13.916 | 27.736 | -5.044 | 18.315 | 17.745 | 14.767 | -3.863 | 16.170 | -0.495 | 26.594 | -21.113 | 26.386 | -0.705 |
| 24.133 | 7.712 | -8.254 | 15.485 | 21.793 | 21.114 | -14.600 | -5.312 | -8.785 | 19.604 | 2.726 | 25.524 | 22.117 | 9.934 |
| -17.787 | 24.396 | -20.374 | 14.412 | 22.794 | -14.205 | -8.444 | 11.473 | 20.536 | -7.035 | -16.050 | 20.466 | 25.254 | 24.974 |
| 23.434 | 21.107 | -26.832 | 4.591 | 2.764 | -25.388 | 24.120 | -6.351 | 10.066 | -2.407 | -23.834 | 7.815 | 18.946 | 21.503 |
| -1.212 | -20.447 | 18.597 | -8.771 | -10.436 | -26.558 | 18.584 | -14.700 | -9.321 | 24.963 | 21.655 | 17.402 | 7.294 | -20.289 |
| -25.105 | -24.201 | 4.957 | 8.575 | -15.356 | -13.719 | -15.521 | 9.350 | 18.618 | -20.320 | -2.415 | 15.047 | 23.161 | 23.978 |
| -25.848 | 15.052 | 28.532 | -17.878 | -2.366 | 22.190 | -10.322 | 5.866 | -18.734 | 7.685 | 29.022 | 11.440 | 27.774 | 27.777 |
| 2.447 | 25.289 | 8.532 | -19.962 | -25.510 | 16.180 | 3.445 | 2.014 | -6.410 | -22.502 | -6.406 | -11.436 | -20.114 | -3.349 |
| -9.454 | -4.620 | -10.212 | -15.613 | 19.309 | -18.464 | -9.426 | -1.429 | 7.241 | -10.500 | -27.220 | -4.935 | 21.808 | -15.515 |
| 18.703 | -9.152 | -3.856 | 24.213 | 14.587 | -28.823 | -25.540 | 3.402 | 2.855 | 5.020 | 5.161 | 24.058 | 18.138 | 25.854 |
| -3.930 | 19.852 | -18.957 | -17.606 | -4.588 | 12.535 | 27.328 | 6.358 | 15.943 | -17.268 | -20.233 | 9.006 | 29.181 | 27.940 |
| 22.944 | -14.655 | 0.970 | -5.157 | 0.180 | 6.322 | -21.725 | -0.588 | 24.706 | -11.507 | -23.960 | 13.194 | 16.640 | 12.664 |
| 1.918 | 29.318 | -0.911 | 6.453 | 17.900 | -16.010 | 13.494 | -7.947 | 11.939 | 8.096 | 25.828 | -2.443 | 4.120 | 26.782 |
| 4.213 | 20.545 | 30.861 | -18.015 | 6.676 | -28.264 | 11.081 | 4.429 | 20.108 | -3.183 | -8.109 | 10.417 | 19.932 | 17.387 |
| 26.402 | 7.594 | -26.251 | 21.009 | 8.895 | 28.393 | -9.143 | 2.934 | -10.684 | -7.419 | 3.551 | -1.127 | 16.255 | 28.320 |
| 2.230 | -28.909 | 27.244 | 9.778 | -0.547 | -22.866 | 5.587 | 19.337 | 14.260 | -19.221 | 8.581 | -16.446 | 21.228 | 21.130 |
| -28.078 | 28.736 | 23.199 | -19.369 | 27.165 | -23.797 | -5.874 | -4.582 | 24.301 | -24.583 | 0.282 | -0.562 | -10.105 | 14.319 |
| 25.695 | -21.826 | 1.879 | -28.233 | 1.798 | 27.309 | -25.637 | 20.358 | 1.604 | 8.763 | 13.780 | -5.521 | -26.450 | -4.018 |
| 0.787 | 27.729 | 2.896 | -18.190 | -26.820 | -27.331 | 15.637 | 8.511 | 9.238 | -8.010 | -6.944 | -18.552 | 23.963 | 27.341 |
| -16.752 | -22.910 | -3.154 | -20.910 | -0.135 | -25.377 | -23.905 | 23.348 | 7.303 | -14.251 | 21.540 | 25.711 | 27.080 | 4.388 |
| 7.366 | -7.090 | -6.613 | -7.267 | 26.431 | -30.261 | -12.411 | -21.573 | -3.354 | -2.866 | 11.222 | 22.151 | 6.607 | 9.894 |
| -24.155 | -20.819 | 9.221 | -15.469 | 8.270 | 28.861 | -21.034 | 25.207 | -2.364 | 25.265 | -13.214 | -25.691 | 19.744 | 23.021 |
| -24.955 | 16.584 | -16.709 | 12.434 | 4.480 | 23.662 | 13.951 | -26.907 | -25.252 | -2.235 | -17.211 | -0.414 | 24.963 | 21.615 |
| -1.021 | 25.621 | -22.728 | 28.306 | -20.090 | 13.815 | 17.941 | 2.949 | 6.624 | -0.848 | 11.754 | -7.231 | 26.427 | -12.099 |

| | | | | | | | | | | | | | |
|---|---|---|---|---|---|---|---|---|---|---|---|---|---|
| -26.313 | -14.593 | 9.902 | -21.134 | 1.554 | -8.311 | 0.206 | -19.005 | -18.264 | -22.133 | 26.341 | 11.471 | 2.627 | 13.153 |
| 27.473 | 8.267 | 11.811 | -9.264 | 9.468 | -11.405 | -16.174 | 15.651 | 18.440 | -4.483 | -27.919 | 24.209 | 21.725 | 7.409 |
| -23.548 | 12.086 | -10.582 | 28.803 | -1.164 | 26.842 | -16.706 | -25.088 | 7.886 | 25.240 | 24.188 | -10.946 | 11.731 | 14.026 |
| -9.740 | 14.625 | -20.183 | -27.201 | 11.940 | -23.719 | 20.492 | -13.730 | 12.582 | -2.490 | -6.944 | 22.409 | 24.372 | 17.039 |
| 16.181 | 13.533 | 15.851 | 10.148 | -8.522 | 12.929 | -18.680 | -15.885 | 2.177 | 26.284 | 5.897 | 6.698 | 16.160 | 20.846 |
| -15.989 | -26.935 | -4.887 | 24.155 | 0.661 | -16.207 | 11.411 | 3.623 | 23.493 | 8.461 | 21.252 | -24.019 | 23.817 | 24.559 |
| -22.713 | -29.921 | -18.445 | -14.620 | -25.480 | 29.845 | 3.132 | -6.295 | 0.809 | -0.685 | 27.523 | -26.824 | 27.298 | -3.660 |
| -11.690 | -22.515 | 5.103 | -22.095 | -14.497 | 3.091 | -13.387 | 0.862 | -4.453 | -23.948 | 22.926 | 23.149 | 21.377 | -27.353 |
| -4.513 | 28.537 | 3.997 | -27.928 | 2.467 | 9.134 | -18.035 | 5.702 | 23.265 | 23.391 | 7.420 | -3.544 | 27.386 | 18.765 |
| 1.770 | 24.103 | 17.527 | -17.576 | -13.005 | 8.156 | 6.548 | -14.219 | 1.292 | -1.699 | -9.542 | 2.700 | 25.541 | 9.949 |
| 7.143 | -18.246 | 29.018 | 14.135 | 13.550 | -1.447 | -8.654 | 9.119 | 25.965 | -17.691 | 18.783 | -2.772 | 21.473 | 1.078 |
| 24.971 | -28.152 | -16.278 | 11.157 | 25.242 | -5.916 | -6.153 | 18.879 | 3.149 | -10.217 | -10.909 | -27.498 | 15.935 | 17.318 |
| 2.203 | 5.091 | 0.383 | -11.851 | -12.452 | 1.536 | -26.381 | 4.578 | -9.810 | -16.979 | -19.929 | 12.109 | 18.709 | 2.079 |
| 3.827 | 2.383 | 12.957 | -27.693 | 15.920 | -25.697 | 27.299 | 3.939 | 16.034 | 22.534 | 26.865 | 25.990 | 4.398 | 23.985 |
| 0.090 | 19.726 | -30.883 | 30.626 | -23.221 | -30.701 | -20.993 | -8.874 | 0.139 | -3.269 | 28.847 | 25.231 | 17.933 | 21.546 |
| -14.853 | 8.824 | 26.713 | 16.894 | -19.855 | 22.178 | 23.457 | -3.802 | 9.287 | -19.150 | 17.643 | 5.241 | 25.125 | 0.465 |
| -17.102 | 4.162 | -28.429 | -14.717 | 0.428 | -27.214 | -20.823 | 22.075 | -4.254 | 25.300 | 5.476 | 14.644 | 14.905 | 22.673 |
| 5.849 | -29.966 | 8.574 | 7.061 | 27.743 | -9.795 | 19.601 | -6.055 | -4.667 | 19.354 | 19.923 | 23.371 | 0.486 | 9.566 |
| 22.023 | 12.180 | 23.813 | -0.659 | -25.724 | 27.970 | -28.667 | 3.143 | 20.125 | -26.720 | -16.566 | 1.512 | -7.497 | 25.482 |
| 12.732 | 24.846 | -8.779 | -28.527 | 3.158 | 26.232 | 11.715 | 16.022 | 6.389 | -10.229 | 29.444 | -3.844 | 14.964 | 16.470 |
| 26.797 | -16.961 | 20.991 | 14.094 | -0.322 | -9.523 | -3.223 | -19.577 | 0.985 | -9.848 | 26.333 | -16.645 | 0.451 | 27.467 |
| -6.884 | -12.335 | 19.953 | 6.328 | 13.148 | -8.775 | 18.013 | -25.148 | 6.879 | -3.727 | 16.381 | 21.429 | 21.547 | 21.629 |
| -26.940 | -5.854 | 30.148 | -14.208 | -16.803 | 22.865 | 18.798 | -6.753 | 12.578 | -25.549 | 11.226 | -13.196 | 8.317 | -14.968 |
| -8.280 | -22.226 | -29.553 | -24.043 | -11.274 | 26.751 | -20.421 | 12.006 | 23.737 | -22.807 | -24.693 | 26.422 | 18.569 | 12.356 |
| -23.874 | 18.287 | 29.911 | 23.639 | -24.687 | -20.498 | -22.920 | 7.924 | 14.023 | 21.999 | -22.849 | 18.340 | 2.361 | -18.553 |
| -24.706 | -26.709 | -1.442 | 29.642 | 0.786 | 28.741 | 14.323 | -18.724 | -15.039 | 8.047 | 28.417 | 17.665 | -22.467 | 20.932 |
| -27.652 | 13.942 | -7.753 | -17.667 | 1.508 | 17.970 | 6.102 | -11.404 | -1.744 | 5.440 | 22.140 | -9.700 | 9.064 | -23.103 |
| 18.662 | 4.673 | 13.852 | 15.085 | 22.116 | 29.838 | -15.992 | -10.287 | -10.992 | -23.542 | -8.854 | -27.011 | -18.188 | 24.635 |
| 26.083 | -3.885 | -3.912 | 6.365 | -5.219 | 7.219 | 22.799 | 2.300 | 22.258 | -4.484 | -27.422 | -14.882 | -2.720 | 16.333 |
| -20.546 | 8.511 | -23.469 | 16.741 | -13.265 | 27.007 | 28.097 | 25.845 | 13.017 | -6.534 | 23.597 | -6.156 | -10.352 | 8.693 |
| 14.436 | -16.665 | 25.010 | -13.137 | -9.959 | 21.964 | -13.206 | 15.701 | 11.023 | -6.899 | -1.330 | 23.638 | 22.602 | 18.229 |
| 24.770 | 29.123 | 18.380 | 19.796 | -13.994 | 20.348 | 20.528 | 12.087 | 20.522 | 4.827 | -24.040 | -25.435 | 21.393 | 29.145 |
| 20.031 | 30.402 | 22.989 | -9.978 | -14.680 | 24.280 | 18.019 | -13.232 | -3.653 | 3.779 | -10.695 | -23.215 | -7.873 | 9.525 |
| 10.452 | -22.580 | -15.011 | -18.530 | 20.547 | -30.168 | 27.348 | -25.790 | -20.996 | 22.838 | 19.390 | 15.900 | 23.080 | 8.915 |
| -15.162 | 6.595 | 25.753 | 11.414 | 21.623 | -23.451 | -7.090 | -23.397 | 0.354 | -18.778 | 4.620 | 14.451 | 6.723 | 27.295 |
| -16.113 | 3.884 | 12.836 | -30.205 | -18.983 | 17.645 | 28.113 | -26.529 | 10.353 | -0.866 | 18.136 | 23.173 | 3.885 | 25.157 |

| | | | | | | | | | | | | | |
|---|---|---|---|---|---|---|---|---|---|---|---|---|---|
| 1.123 | 5.538 | -5.273 | 24.301 | 0.238 | 19.311 | -25.313 | 26.929 | 19.377 | 19.750 | 19.888 | -0.561 | 18.426 | 14.646 |
| 0.188 | -19.054 | -3.047 | 3.550 | 22.462 | 0.053 | -12.259 | 23.906 | -6.931 | -4.633 | -7.676 | -26.186 | -4.911 | 21.005 |
| -19.980 | -13.614 | -0.773 | -3.413 | 25.238 | 13.271 | -30.363 | 25.090 | -6.020 | -5.726 | -27.698 | 15.503 | 21.056 | 20.293 |
| -2.148 | 17.442 | -1.878 | -23.908 | -13.363 | 20.621 | 21.408 | -15.204 | 3.174 | -15.640 | 26.945 | -13.120 | 23.065 | 17.260 |
| 19.031 | -27.624 | 12.398 | -29.746 | -14.496 | -15.389 | -10.192 | 13.104 | -4.860 | -22.736 | 23.205 | -23.874 | 11.171 | 21.881 |
| -28.910 | -24.864 | -2.805 | -23.228 | -11.083 | -3.400 | 19.582 | 10.436 | -21.439 | 12.645 | 30.276 | 25.700 | 6.854 | 29.539 |
| -24.927 | 6.999 | 15.450 | -24.890 | -7.211 | 17.780 | 22.402 | -1.308 | -4.470 | -19.829 | 17.026 | -18.523 | 20.409 | 16.207 |
| 23.155 | -11.467 | 7.348 | -9.922 | 11.851 | -10.526 | 8.247 | 20.910 | 24.146 | 18.665 | -22.765 | -1.308 | 18.315 | -10.737 |
| 12.020 | -28.133 | 28.805 | 9.338 | -19.891 | 22.496 | 6.218 | 9.029 | 3.806 | -8.325 | -30.254 | 10.979 | 25.554 | 0.801 |
| 2.724 | 9.654 | 20.890 | 18.508 | 19.301 | -21.803 | -29.771 | 0.703 | 17.339 | -19.180 | -26.967 | 1.478 | 5.974 | -19.671 |
| -14.311 | 25.330 | 25.723 | -26.292 | 7.395 | 14.748 | -10.150 | 3.242 | -0.713 | 20.484 | 21.768 | 18.694 | -9.002 | 26.766 |
| 15.369 | 20.066 | -19.577 | 11.704 | -23.739 | -5.356 | -12.393 | 8.534 | -5.813 | 21.295 | 3.987 | 11.943 | 16.619 | 20.917 |
| -0.804 | 1.787 | 16.972 | -8.083 | 23.359 | 25.667 | -23.311 | 18.574 | -24.733 | 15.096 | 22.865 | 22.487 | 10.479 | 24.807 |
| 3.200 | -6.734 | 17.407 | -7.443 | 4.728 | 6.537 | 9.652 | 15.649 | -17.181 | 1.725 | 24.024 | 19.300 | -18.471 | 18.207 |
| 5.409 | 12.919 | 26.231 | 10.048 | 0.372 | -2.244 | 28.947 | -4.951 | 24.854 | -2.839 | -1.438 | -21.410 | -4.778 | 11.613 |
| -17.720 | -6.286 | 0.833 | 19.419 | -12.059 | 12.357 | -26.660 | -20.631 | -8.017 | -3.522 | 27.537 | 23.371 | 11.724 | 25.612 |
| 8.053 | -23.569 | 15.090 | 7.723 | -20.644 | 14.041 | 26.644 | -12.074 | 17.703 | -22.452 | 11.877 | 13.341 | -23.132 | 14.825 |
| -26.395 | 0.828 | 9.360 | -5.483 | -14.040 | 21.779 | 29.367 | -18.644 | 18.142 | 24.368 | -8.440 | 5.472 | 20.402 | 28.834 |
| 25.828 | 4.477 | -6.906 | -8.320 | -18.643 | 5.558 | -18.552 | -5.152 | -8.318 | 13.024 | 21.815 | 8.216 | 17.575 | 17.587 |
| 1.735 | -22.055 | -20.649 | -10.268 | 2.721 | -7.509 | -1.364 | -11.295 | -7.213 | -2.462 | 25.000 | 19.097 | 4.600 | -1.139 |
| -9.467 | 5.502 | -1.825 | 20.081 | 16.326 | -24.366 | -8.485 | 22.937 | 15.372 | 11.016 | 8.700 | -1.741 | 10.537 | 21.910 |
| -22.113 | -10.755 | -29.852 | 3.342 | 4.217 | -11.656 | 12.078 | -15.112 | 3.519 | 25.789 | -22.729 | -17.560 | 14.560 | 20.384 |
| -7.228 | -23.949 | -11.113 | 22.283 | 28.032 | 14.997 | -24.312 | 17.469 | 10.711 | -4.983 | -26.988 | -6.990 | 23.028 | 3.578 |
| 3.782 | 13.138 | -12.737 | 11.717 | -23.280 | 20.410 | -30.753 | 17.092 | -13.809 | 7.400 | -29.378 | -24.784 | 26.597 | 7.113 |
| 6.797 | 14.142 | 8.654 | 8.753 | 17.365 | 27.326 | -17.854 | -26.870 | 2.128 | -24.231 | 26.636 | 4.387 | 23.879 | -8.821 |
| -19.026 | -28.878 | -27.893 | -21.617 | 29.873 | -29.605 | 23.484 | 19.907 | 2.967 | 27.510 | -9.205 | -16.973 | -19.821 | 28.122 |
| -1.866 | 18.005 | 28.914 | -11.120 | 3.024 | 4.513 | 16.715 | 7.322 | -6.696 | 24.448 | 15.738 | -14.211 | -13.473 | 16.357 |
| -6.113 | 27.364 | 6.863 | 12.681 | -1.324 | -27.356 | -26.557 | -8.232 | 0.223 | 4.859 | 30.223 | 3.795 | 2.609 | -25.724 |
| -28.427 | 1.165 | -3.457 | 24.158 | 7.447 | 23.116 | 20.920 | 22.798 | -0.693 | 18.471 | -29.750 | 16.695 | -0.190 | 27.283 |
| 26.611 | 28.501 | -24.213 | 19.451 | -9.156 | 28.926 | -21.212 | 18.329 | 14.694 | -9.460 | -8.588 | -19.805 | 19.736 | 23.757 |
| 1.328 | 30.257 | 23.209 | 11.358 | -20.186 | -3.422 | 10.337 | 7.535 | -21.809 | -8.789 | 26.651 | -25.041 | -21.259 | 27.668 |
| -12.110 | 21.269 | -18.117 | -3.825 | 16.685 | 9.740 | -1.053 | -30.267 | 20.959 | 1.888 | -19.949 | -15.548 | 13.295 | 12.966 |
| 28.390 | -30.529 | 4.606 | 7.075 | 8.271 | -0.517 | 23.865 | 28.196 | -5.803 | -2.552 | -30.183 | 9.533 | 21.085 | -10.767 |
| -0.231 | 4.595 | -7.221 | -24.938 | -0.700 | 13.180 | 3.338 | 20.034 | -23.477 | -11.172 | 4.329 | -14.894 | 0.557 | 15.270 |
| -15.308 | -31.071 | -14.299 | 10.584 | 21.118 | 0.446 | -19.935 | -19.683 | -8.630 | -7.530 | -18.189 | -20.414 | 13.022 | 26.913 |
| 5.644 | 1.800 | -19.937 | -18.126 | 26.415 | -10.184 | 25.847 | -7.225 | 8.960 | -9.533 | 17.856 | -6.760 | 24.118 | 22.149 |

| | | | | | | | | | | | | | |
|---|---|---|---|---|---|---|---|---|---|---|---|---|---|
| 17.066 | 29.794 | -29.523 | -15.518 | 17.905 | -24.603 | 27.868 | -10.551 | 5.189 | 8.073 | -8.842 | -10.917 | -12.029 | -8.078 |
| 14.834 | 23.906 | -21.753 | -14.849 | 13.873 | -2.357 | -14.932 | 6.132 | 22.562 | 20.137 | -2.454 | 9.490 | 25.863 | 10.135 |
| 4.663 | 25.456 | 24.122 | -27.014 | -7.196 | 20.216 | -27.902 | -22.478 | -26.554 | -17.151 | 15.950 | 25.210 | 15.682 | 18.542 |
| -11.574 | 4.894 | -10.353 | 25.019 | 27.653 | -12.567 | -10.525 | -19.694 | -7.519 | -2.283 | 17.158 | -7.636 | 6.248 | 30.540 |
| -8.797 | -11.470 | 27.809 | 9.936 | 22.089 | 23.465 | -5.676 | 26.287 | -11.424 | -1.945 | 29.859 | 0.982 | 22.461 | 29.077 |
| -19.486 | -28.698 | 8.286 | 28.191 | -14.795 | 6.105 | 25.479 | 20.399 | 24.693 | 1.202 | -21.539 | -19.127 | 7.361 | 16.153 |
| 20.756 | -19.122 | 13.226 | 2.226 | 28.988 | 1.397 | -5.538 | -16.972 | -10.532 | 9.877 | -29.913 | -20.034 | -21.804 | -7.462 |
| -11.163 | 27.945 | 10.774 | -10.061 | 8.972 | 20.890 | 19.501 | 7.393 | 10.469 | 25.586 | 19.079 | 4.176 | 18.756 | 9.417 |
| -28.180 | -20.743 | 20.469 | 25.491 | 1.958 | 8.844 | -25.405 | -20.080 | 18.212 | 14.656 | 29.462 | 20.261 | 25.992 | 14.545 |
| 22.058 | 12.915 | 6.561 | 25.963 | -28.921 | 3.741 | 23.207 | 29.845 | -25.904 | 28.994 | -16.769 | -3.492 | 25.564 | 23.507 |
| 5.102 | 5.941 | -5.667 | 1.740 | 24.370 | -18.073 | 20.462 | -6.786 | -27.173 | -27.984 | -17.799 | -6.930 | 19.176 | 21.317 |
| 24.895 | 7.012 | 27.253 | -22.993 | -21.332 | -28.413 | -13.455 | 22.773 | 20.161 | 27.514 | 20.801 | -16.152 | 22.377 | 26.656 |
| 21.549 | 30.578 | 26.256 | -8.380 | -7.112 | 16.970 | -28.269 | 25.304 | 21.237 | 0.037 | 25.961 | 24.751 | 27.727 | 18.196 |
| -25.673 | 18.017 | 25.469 | 19.671 | 11.550 | -2.774 | 12.947 | -28.573 | -17.718 | -28.562 | 24.791 | -22.887 | 21.578 | 20.096 |
| -4.371 | -27.063 | -0.867 | 26.332 | 7.145 | 25.341 | -8.441 | -27.802 | 26.361 | -18.002 | -0.896 | 20.794 | 9.222 | 11.609 |
| 8.560 | 9.607 | -16.634 | -19.405 | 29.885 | 18.218 | -24.667 | 19.652 | -12.733 | -11.139 | 3.195 | 11.573 | 16.886 | 5.622 |
| 5.910 | -20.002 | -18.122 | 5.905 | 23.689 | 21.914 | 21.679 | -29.921 | 15.824 | 7.203 | 17.140 | 5.436 | 10.115 | 30.231 |
| -9.367 | 18.649 | 28.525 | 13.004 | -1.924 | -10.567 | 20.258 | -18.376 | -12.198 | 11.090 | -26.521 | -18.856 | -2.015 | 29.148 |
| 23.909 | -25.181 | 23.073 | -10.149 | 14.097 | 23.888 | 15.238 | 23.867 | -9.602 | 23.634 | 2.313 | 25.902 | 21.770 | 21.730 |
| -0.254 | 16.906 | 5.843 | -5.897 | 26.432 | 7.981 | -19.040 | 16.725 | -22.996 | -1.725 | 20.257 | -10.921 | 23.266 | 26.337 |
| -23.245 | 13.921 | 24.230 | 9.103 | 17.516 | 11.800 | -23.239 | -26.916 | -4.182 | -29.921 | 13.914 | 21.714 | 15.916 | -4.416 |
| 18.319 | 29.501 | -22.943 | -20.957 | -24.632 | 27.265 | 29.060 | -6.178 | 27.923 | -7.130 | -26.198 | 12.162 | 28.751 | -2.871 |
| -2.346 | 19.144 | 2.876 | 2.402 | 19.842 | 19.737 | 24.675 | 30.320 | 20.334 | 1.273 | 22.371 | 6.417 | -6.542 | 14.069 |
| 20.221 | 25.455 | 5.208 | 17.022 | -14.516 | -13.267 | -24.192 | 27.574 | 7.504 | 9.003 | 0.426 | 3.244 | 22.837 | 2.273 |
| -2.939 | 24.638 | 3.683 | 8.199 | 29.948 | 26.047 | -10.588 | -25.522 | -24.446 | -11.375 | -9.004 | -4.821 | 1.704 | -25.535 |
| -24.317 | 21.452 | 19.611 | -13.435 | -8.582 | 16.310 | 22.127 | 19.599 | 23.654 | 28.555 | -2.301 | -26.344 | 18.955 | 24.006 |
| -23.491 | -6.686 | 9.571 | 22.299 | 30.655 | -26.678 | 28.224 | -1.599 | 5.288 | 24.923 | -29.595 | -16.254 | 20.430 | 20.227 |
| 16.228 | 12.573 | -9.375 | -14.301 | 0.258 | -20.606 | 20.405 | -5.898 | 25.011 | -19.361 | -28.131 | 17.528 | 24.885 | 7.883 |
| 19.550 | 20.847 | -8.694 | -21.224 | 10.347 | 13.949 | -22.221 | 23.962 | 8.857 | 22.469 | 3.147 | 4.122 | 16.747 | 30.755 |
| 17.976 | 1.415 | 17.704 | 7.430 | -10.002 | 19.222 | 25.373 | 14.425 | 18.852 | -26.177 | -8.016 | 14.666 | 9.542 | 17.195 |
| 11.223 | 16.274 | -6.218 | -16.299 | -30.451 | 12.713 | 20.017 | 9.894 | 5.753 | -28.647 | 20.733 | 9.392 | 8.305 | 15.937 |
| -20.352 | 27.780 | 10.478 | 20.511 | -2.035 | -3.871 | 5.342 | 6.869 | 11.063 | 2.794 | 27.972 | -3.853 | -3.900 | 15.357 |
| -7.199 | -20.978 | 8.171 | -11.040 | 13.309 | 3.044 | -7.064 | -19.012 | -16.343 | 17.510 | -19.249 | -26.744 | 23.646 | 30.199 |
| 22.268 | 28.444 | -15.568 | 21.177 | 31.535 | 27.103 | 15.338 | 28.168 | -20.027 | 4.058 | 11.870 | -21.450 | 25.749 | 19.419 |
| -30.407 | 7.438 | -20.619 | 13.664 | 19.145 | 21.395 | 21.479 | -6.079 | -24.988 | -20.573 | 19.399 | 9.196 | 27.770 | 22.787 |
| -18.850 | -6.350 | -9.724 | -17.621 | 19.822 | 24.491 | -17.852 | -19.833 | 25.583 | 27.934 | -18.138 | -23.548 | 24.897 | 18.142 |

| | | | | | | | | | | | | | |
|---|---|---|---|---|---|---|---|---|---|---|---|---|---|
| 24.167 | -14.184 | -13.892 | -7.779 | -8.444 | -6.670 | 5.848 | -8.950 | -23.350 | 26.718 | 15.684 | 5.195 | 28.919 | 25.981 |
| 20.977 | -26.359 | 17.245 | -9.062 | 29.310 | -9.638 | -27.930 | 19.846 | 20.526 | 23.655 | 25.126 | 18.362 | 22.219 | 21.457 |
| -25.187 | 30.258 | -16.719 | -7.918 | 18.709 | 11.656 | 23.496 | -18.738 | -18.664 | -0.919 | 24.364 | 13.213 | 28.247 | -6.959 |
| -29.285 | -9.325 | 20.118 | 21.360 | 18.577 | 19.400 | -26.063 | -13.502 | 10.204 | 5.246 | 22.409 | -19.563 | 9.875 | -24.550 |
| 3.655 | -16.260 | 11.656 | 10.967 | -21.764 | -8.515 | 27.480 | -8.206 | -26.485 | -10.646 | -1.340 | 25.486 | 5.422 | -32.187 |
| 16.123 | 14.587 | -12.213 | -20.073 | -7.589 | -22.741 | 22.454 | 14.521 | 17.743 | 28.830 | -3.036 | 3.694 | 18.707 | 10.852 |
| 5.342 | -1.144 | 4.458 | -24.947 | 17.254 | -19.438 | -9.638 | 24.781 | 25.046 | -18.006 | 13.129 | 20.314 | 14.899 | 11.802 |
| -3.181 | -5.794 | 21.580 | -23.617 | 5.030 | 7.438 | -16.218 | 15.046 | -7.894 | -22.764 | 9.524 | 27.595 | 27.053 | 28.571 |
| 8.733 | -11.769 | 2.466 | -21.657 | 7.685 | -20.277 | -20.973 | 11.916 | -27.234 | 30.234 | 26.822 | 24.606 | -15.236 | 26.182 |
| 29.110 | -20.956 | 25.665 | -5.873 | -27.114 | -15.970 | -0.718 | -15.238 | 28.073 | -27.015 | 20.109 | -19.109 | 24.562 | 14.607 |
| 29.687 | 8.630 | -13.495 | -8.838 | 29.730 | 20.350 | 12.438 | 22.736 | 9.752 | 13.233 | 3.667 | -26.640 | 22.243 | 20.218 |
| 2.510 | 5.933 | -20.001 | -0.035 | -0.601 | 26.509 | 0.662 | -21.875 | 15.203 | -1.387 | -7.367 | 17.324 | 23.295 | 18.581 |
| -28.291 | -12.870 | -22.569 | 10.486 | -27.826 | -4.989 | -8.344 | -14.148 | -25.195 | 25.739 | -3.415 | 9.582 | 19.928 | 15.114 |
| 27.817 | 26.300 | -18.621 | 3.661 | 7.127 | -9.419 | -17.676 | -6.677 | -8.179 | 20.731 | -5.424 | 6.457 | -3.771 | 10.577 |
| 26.149 | 16.964 | 5.063 | 23.114 | -12.887 | 13.213 | -7.956 | -18.447 | 22.578 | -11.917 | -6.474 | 19.874 | 15.073 | 2.097 |
| -0.759 | -29.444 | -24.234 | -18.608 | 29.907 | 19.293 | -15.372 | 19.250 | 7.563 | -22.175 | -11.215 | -21.548 | 24.389 | -29.945 |
| -27.183 | 10.010 | -4.990 | -1.477 | 6.764 | 13.623 | -22.171 | 5.295 | 22.355 | -13.994 | 29.720 | 3.067 | 18.978 | 13.228 |
| 24.324 | 22.533 | -27.461 | 16.333 | -19.230 | 25.300 | 6.304 | -10.799 | 5.395 | 10.704 | 23.512 | -15.989 | 22.412 | 13.886 |
| -19.716 | 27.277 | -14.936 | 19.660 | 21.238 | -7.689 | -7.388 | -30.117 | -16.406 | -1.674 | 21.624 | 17.818 | 9.623 | 14.775 |
| -29.762 | 25.102 | 16.447 | -0.958 | -30.434 | 4.541 | -20.926 | -18.452 | -26.661 | 29.120 | 15.092 | -2.651 | -1.157 | -1.399 |
| -31.296 | -22.087 | 15.174 | -16.922 | -11.279 | -21.473 | -20.263 | -7.219 | -11.289 | -0.693 | -8.002 | -28.419 | -7.302 | 5.594 |
| -3.992 | 0.237 | -24.820 | 23.877 | -7.823 | -1.191 | 18.838 | 21.106 | 9.034 | -6.537 | -0.754 | 6.779 | 17.981 | 24.251 |
| 26.837 | -13.686 | -14.390 | 9.213 | 15.595 | 16.477 | 22.877 | 15.283 | -6.847 | 30.237 | 12.752 | 23.379 | 10.321 | 16.834 |
| 16.768 | -29.843 | 24.135 | 22.058 | 11.631 | 22.748 | 25.156 | -28.976 | 3.690 | 7.110 | 21.021 | -0.499 | -21.780 | -1.806 |
| 4.671 | -18.065 | 26.572 | -24.733 | 16.508 | -16.920 | -25.870 | -7.110 | 16.420 | 25.878 | -16.015 | 22.500 | -11.146 | -2.691 |
| 3.098 | 15.765 | 9.871 | -10.590 | 21.611 | 9.867 | -13.641 | 9.161 | -20.211 | 18.246 | -22.849 | -11.079 | -8.346 | -17.734 |
| 1.636 | -8.287 | 8.259 | -8.718 | 4.779 | 25.955 | -20.923 | 2.380 | 11.422 | 23.060 | -29.238 | 11.199 | 16.930 | 17.183 |
| -0.458 | 28.416 | 5.224 | 19.040 | -3.117 | 23.590 | 9.931 | -10.294 | 4.096 | -30.955 | 27.197 | 19.148 | 26.911 | -30.696 |
| 5.863 | -25.883 | 23.404 | 24.383 | 19.371 | -25.684 | 14.959 | -23.237 | 24.368 | -19.076 | -4.332 | 7.293 | 25.313 | -16.373 |
| -5.937 | 30.241 | -25.359 | 4.742 | 3.431 | 14.220 | 6.338 | 13.496 | 6.471 | 9.823 | 2.557 | 18.265 | 14.175 | -7.867 |
| 10.130 | 23.438 | 24.148 | 21.247 | -6.861 | -26.374 | -14.662 | 17.816 | 1.455 | 30.899 | 23.369 | 13.507 | 28.706 | 16.729 |
| 14.685 | -11.834 | -10.290 | -3.603 | 12.874 | 24.406 | 19.090 | -18.335 | -28.265 | -4.040 | 4.504 | 27.536 | 15.006 | 5.242 |
| 20.987 | 0.604 | 2.945 | -6.766 | 8.258 | -8.351 | -22.099 | 22.403 | -7.982 | -23.216 | -17.283 | 24.099 | 19.016 | -4.974 |
| -19.515 | 24.447 | -5.903 | -6.401 | -25.666 | 20.213 | 18.333 | -28.038 | -7.589 | -21.785 | 22.674 | -19.193 | 28.164 | 11.641 |
| 12.888 | 27.303 | -16.666 | -17.146 | -29.215 | 15.878 | 8.902 | 5.070 | 16.942 | -6.074 | 1.137 | 5.352 | 30.896 | 20.545 |
| 30.811 | 7.022 | 17.881 | -9.630 | -27.241 | -17.514 | -2.590 | -21.061 | -28.823 | 30.496 | -28.240 | -19.809 | 12.264 | 8.744 |

| | | | | | | | | | | | | | |
|---|---|---|---|---|---|---|---|---|---|---|---|---|---|
| -1.290 | -8.705 | -24.807 | -8.974 | -15.425 | 1.343 | 23.991 | -7.848 | -9.800 | -5.536 | 27.657 | 25.190 | 12.430 | 17.833 |
| -28.957 | -11.252 | 25.520 | 13.341 | -26.336 | -5.841 | -6.861 | 20.009 | 29.092 | -29.843 | -3.357 | 22.936 | 16.140 | 13.046 |
| -2.276 | 21.649 | -18.815 | 3.586 | 30.030 | -2.185 | 15.664 | 1.391 | 21.143 | -6.623 | 15.417 | -2.393 | 19.973 | 24.662 |
| -16.056 | 29.324 | -27.083 | -21.062 | 21.920 | -23.974 | -23.608 | -24.270 | 1.738 | 7.795 | -13.731 | -13.517 | -30.316 | 17.117 |
| 24.580 | 23.487 | -5.276 | 3.970 | 21.264 | 11.379 | -9.684 | 27.144 | -5.170 | -9.874 | 28.903 | 24.653 | 29.043 | 26.833 |
| 4.126 | -17.496 | 22.016 | 20.241 | 11.579 | -7.870 | 19.648 | 17.318 | 24.993 | 30.429 | 23.967 | 13.457 | 27.531 | -9.501 |
| 3.147 | 19.425 | -3.186 | -4.663 | 6.673 | -7.075 | -11.792 | 1.207 | -11.308 | 15.829 | -7.016 | -7.567 | 18.325 | -25.975 |
| 19.830 | -19.972 | 0.766 | -0.875 | 18.208 | 5.745 | 9.265 | 21.812 | 24.356 | -2.180 | 13.576 | 27.395 | 11.429 | 19.609 |
| -13.237 | 6.371 | 21.389 | 23.516 | -25.176 | 25.123 | 5.612 | 26.564 | 5.981 | -12.739 | 27.197 | -17.739 | 22.637 | 9.664 |
| 30.251 | 14.076 | 15.220 | -19.335 | -22.382 | 5.955 | 0.593 | 7.654 | -4.751 | 24.562 | -23.454 | -26.356 | 22.238 | 7.541 |
| 6.210 | 9.169 | 22.177 | 22.654 | 26.823 | 10.558 | -18.802 | 19.913 | -9.339 | -25.161 | 20.045 | 28.244 | 26.256 | 21.063 |
| 1.691 | -12.923 | -14.605 | 12.125 | 4.206 | -19.002 | 21.981 | -11.747 | 16.278 | 31.885 | 21.733 | 29.042 | 23.699 | 23.855 |
| -15.439 | 7.739 | -17.365 | -22.059 | -13.858 | -24.148 | 13.585 | -17.283 | 29.698 | 9.919 | -9.689 | 9.005 | 10.142 | 18.476 |
| -17.614 | 21.992 | 19.138 | 7.732 | -1.140 | 24.735 | 6.177 | 11.579 | 22.797 | 29.794 | -16.851 | 5.704 | 11.920 | 5.750 |
| -22.476 | 21.021 | -22.972 | 6.678 | 22.046 | 13.309 | 15.704 | -5.838 | 17.073 | 22.848 | 28.576 | -25.215 | 24.686 | 25.977 |
| -28.465 | 2.711 | 17.111 | -19.066 | 4.711 | 23.679 | -3.845 | -23.035 | 8.083 | -13.041 | -4.900 | -18.339 | 26.087 | 16.455 |
| 18.048 | 14.457 | 20.657 | -8.714 | -21.911 | 24.307 | -18.575 | 17.250 | -18.563 | 28.792 | -19.615 | 26.217 | 5.546 | 30.244 |
| -26.057 | 8.598 | 17.380 | -25.784 | 22.845 | -15.326 | -20.974 | 12.217 | -4.406 | 6.219 | 24.976 | 29.355 | 28.090 | 25.137 |
| -23.600 | -19.180 | -0.090 | -10.728 | 29.195 | 8.105 | -11.491 | -1.365 | -20.511 | -14.532 | 25.780 | 21.394 | 26.861 | 5.325 |
| 23.342 | 6.189 | 13.748 | 5.559 | 15.794 | -17.567 | -19.612 | -4.004 | 2.287 | 23.749 | -0.923 | 27.455 | 22.959 | 10.845 |
| 2.280 | -5.698 | -12.914 | -20.426 | 20.537 | -0.674 | 10.033 | -18.978 | 17.992 | 3.625 | 4.326 | 24.760 | 19.834 | 6.920 |
| -21.526 | -6.139 | 15.125 | -17.778 | 15.406 | -11.420 | -12.578 | 6.590 | 10.380 | -25.693 | -25.970 | 29.169 | 20.430 | 8.692 |
| 4.421 | -14.666 | -10.683 | 23.098 | 27.311 | 7.728 | 1.806 | -23.205 | 6.463 | -5.734 | 12.395 | 27.069 | 15.558 | 18.759 |
| 3.833 | 12.230 | 5.093 | -21.008 | 28.454 | 22.490 | 6.854 | -13.672 | -29.120 | -16.354 | 5.768 | 20.543 | 26.064 | 17.328 |
| -20.641 | 10.843 | 16.639 | -22.692 | 10.502 | 25.163 | 8.120 | 18.971 | 15.055 | -13.584 | 27.931 | 1.761 | 21.581 | -8.741 |
| 1.328 | 20.099 | 10.715 | 22.341 | -21.975 | 20.435 | 24.158 | -3.494 | -28.226 | 6.981 | 15.006 | -25.943 | 1.928 | 24.945 |
| -9.637 | 26.188 | -13.876 | 24.867 | 26.456 | 15.947 | 15.384 | 18.719 | 7.104 | 7.883 | 25.583 | 23.746 | 9.046 | 20.761 |
| 19.243 | -3.087 | 5.396 | -13.811 | 20.230 | 11.931 | -18.819 | 15.262 | 14.272 | 29.073 | 8.653 | 17.786 | 26.984 | 5.836 |
| -5.568 | 18.214 | -17.011 | -17.756 | 18.399 | -13.881 | -7.420 | -0.964 | 5.647 | 13.599 | 26.450 | 25.808 | 22.404 | 6.816 |
| 17.653 | 16.875 | -22.589 | 17.797 | -1.624 | -16.428 | 2.989 | 16.534 | 28.416 | 29.330 | -8.662 | 28.607 | 13.789 | 1.343 |
| -19.857 | -17.131 | -26.010 | 5.947 | -15.096 | -6.867 | -23.298 | -24.959 | 17.462 | -9.127 | 23.982 | 18.582 | 23.660 | 18.756 |
| 16.432 | -2.080 | 25.295 | -5.170 | 7.099 | -16.368 | -7.848 | -16.541 | 10.141 | -25.221 | -17.593 | 12.530 | 18.676 | 29.205 |
| 8.267 | -8.867 | 15.604 | 5.015 | 20.311 | -5.083 | -21.873 | -18.963 | 7.550 | 28.799 | -18.955 | 24.955 | 26.579 | 15.685 |
| -3.414 | 24.741 | -14.246 | -19.637 | 21.735 | -6.633 | 2.929 | 19.156 | -27.310 | 21.536 | -15.468 | 5.304 | 15.886 | 14.937 |
| 22.715 | 4.721 | -17.854 | 19.951 | -23.353 | 0.979 | -16.760 | 10.689 | 15.743 | -18.119 | -16.368 | -30.875 | 28.203 | 10.137 |
| -29.149 | 21.707 | -21.168 | 25.321 | 25.156 | -13.351 | -7.349 | 12.228 | -19.917 | -13.633 | 3.647 | 28.239 | 20.851 | -11.423 |

| | | | | | | | | | | | | | |
|---|---|---|---|---|---|---|---|---|---|---|---|---|---|
| 27.788 | 12.420 | 5.284 | -21.692 | -10.442 | 8.202 | -5.226 | 11.794 | 20.950 | 2.438 | 14.656 | -29.891 | 22.901 | -27.846 |
| 20.031 | -23.997 | 10.113 | 16.741 | -5.226 | -10.650 | 1.084 | 16.777 | -0.575 | -1.671 | 23.460 | 21.143 | 12.976 | 25.587 |
| 10.176 | 16.800 | -11.947 | -7.736 | 24.083 | 9.217 | -20.330 | 18.080 | 13.495 | 30.174 | -24.407 | 10.720 | 7.125 | -9.097 |
| -13.627 | -15.944 | 12.002 | -18.361 | 0.805 | 16.052 | -27.155 | 19.149 | 7.312 | -31.667 | -25.828 | 24.022 | 20.156 | 23.359 |
| -22.297 | 2.309 | 7.693 | 7.477 | 28.905 | -19.237 | -24.580 | 4.391 | 23.098 | 22.575 | 1.741 | 26.039 | 19.597 | 27.906 |
| -28.304 | -16.265 | 6.917 | 3.885 | 4.414 | -23.170 | 16.544 | -3.187 | 12.405 | 27.913 | 24.775 | -30.272 | 17.498 | 6.141 |
| -25.525 | 17.774 | -11.411 | 27.878 | 3.752 | -18.468 | -26.655 | -19.075 | 5.565 | -2.293 | 12.262 | 2.234 | 15.766 | 1.011 |
| -1.236 | -12.163 | -1.189 | 8.453 | -28.207 | -21.921 | 12.881 | -5.309 | 28.736 | -22.247 | -2.959 | 18.407 | 10.224 | 8.881 |
| -24.210 | -25.895 | -16.182 | -22.395 | 18.797 | 5.733 | -0.203 | 5.028 | 21.399 | 27.313 | -14.672 | 12.494 | 19.666 | 26.633 |
| 9.016 | 4.297 | -2.263 | -16.854 | 0.943 | -25.596 | -22.130 | -7.479 | 18.110 | -9.381 | 27.423 | 25.145 | 17.920 | -19.089 |
| 4.900 | 28.117 | 17.796 | -14.786 | -19.653 | 9.217 | -12.278 | -2.202 | 1.084 | -19.461 | 3.612 | 3.920 | 23.955 | 23.663 |
| 2.313 | -6.981 | -5.230 | -20.039 | 6.409 | 22.198 | 9.445 | 11.380 | 15.141 | -12.949 | -19.427 | -1.538 | 12.264 | 19.118 |
| -10.671 | 15.082 | -3.072 | 27.025 | -24.761 | 23.993 | -4.502 | -8.491 | -24.443 | 20.262 | 21.974 | -13.593 | 26.618 | 19.710 |
| 17.368 | 15.443 | -3.411 | -21.386 | 0.237 | 26.367 | -18.970 | 21.306 | -16.079 | -24.358 | -18.852 | 19.559 | 28.845 | 20.532 |
| 26.861 | 14.666 | 6.244 | -28.643 | -25.564 | -10.950 | -21.024 | 6.699 | -17.700 | 20.954 | -15.252 | 13.105 | 21.470 | 15.386 |
| -24.404 | -13.016 | 20.471 | -26.061 | 1.346 | 10.877 | -11.185 | -3.468 | 11.340 | -21.575 | 15.162 | 24.069 | 26.148 | 7.650 |
| 8.563 | -17.411 | 16.048 | -5.697 | 16.453 | -24.129 | 24.592 | 7.634 | -3.370 | 6.384 | -12.825 | -13.013 | 17.178 | 21.841 |
| 19.859 | -26.554 | -22.358 | 21.975 | -1.777 | 22.241 | -15.686 | -23.338 | 28.237 | 16.562 | -6.688 | 30.307 | 24.059 | -12.318 |
| 0.225 | -24.051 | 2.980 | -27.787 | 15.088 | 17.147 | -7.035 | 23.698 | 19.019 | 22.638 | -8.407 | 19.791 | -28.721 | 26.044 |
| 10.077 | -16.300 | -20.133 | 28.796 | 3.855 | 5.492 | 3.933 | 3.975 | 13.108 | 1.906 | 11.357 | 0.637 | 9.079 | -26.165 |
| 16.145 | 19.209 | 13.874 | 26.364 | 3.255 | -26.298 | 2.736 | -6.085 | 14.814 | -29.002 | 26.277 | 14.154 | 28.367 | 13.504 |
| -8.777 | 21.446 | 22.855 | 22.717 | 23.086 | 23.081 | 15.020 | -12.507 | 3.054 | 23.193 | 25.059 | 12.773 | 23.318 | 24.646 |
| 17.965 | -4.171 | 18.890 | -22.923 | 24.666 | 18.110 | -5.733 | -0.534 | 19.948 | 19.823 | 9.927 | -15.410 | 18.402 | 28.597 |
| 22.008 | 27.141 | 16.950 | -22.500 | -9.728 | -6.168 | 8.303 | 2.711 | 12.802 | -2.554 | 12.952 | 31.166 | 14.667 | 10.946 |
| 20.369 | 6.998 | -11.565 | -4.814 | 11.655 | 8.511 | 3.282 | -11.633 | 1.191 | 21.813 | 21.034 | 13.040 | 27.661 | 5.296 |
| -7.522 | -9.736 | 8.277 | -19.317 | 5.897 | 7.582 | -19.963 | -26.283 | 0.213 | -21.618 | 18.112 | 28.143 | 22.837 | 16.101 |
| -28.241 | -27.724 | 17.444 | 18.975 | 16.131 | -12.440 | 1.311 | 24.369 | -22.824 | 5.468 | -11.438 | 26.981 | 19.241 | 27.954 |
| 17.028 | 24.870 | 4.935 | -1.728 | -18.095 | 14.744 | 13.890 | -0.519 | -27.378 | 26.200 | -17.506 | 26.362 | 26.379 | 27.441 |
| -6.081 | -20.787 | 3.471 | 12.805 | 11.180 | 19.180 | -13.476 | 10.543 | 29.356 | 23.736 | 0.593 | -26.446 | 25.668 | -14.285 |
| -10.206 | -15.034 | 6.305 | -19.112 | 3.382 | -2.920 | -10.666 | -9.425 | 27.765 | 30.055 | -1.858 | 27.901 | 28.239 | 17.364 |
| 8.103 | -17.010 | -0.550 | 20.293 | -10.407 | -15.107 | -23.336 | 7.289 | 30.209 | -17.575 | 24.426 | 16.990 | 3.568 | 8.826 |
| 6.448 | 25.359 | -10.783 | 7.533 | 27.937 | 13.147 | 17.873 | -18.527 | 18.676 | 17.853 | -20.368 | -13.993 | 25.314 | 25.377 |
| -21.132 | 22.924 | -15.951 | -15.077 | 0.254 | -4.566 | -8.726 | -5.513 | 5.539 | 9.104 | 26.189 | 29.251 | 15.272 | -4.346 |
| -16.514 | 1.689 | -9.721 | 26.886 | -13.844 | 22.414 | 17.298 | 6.088 | 11.851 | -28.278 | -14.617 | -27.929 | 21.067 | 24.289 |
| -6.903 | -3.539 | -14.443 | 28.838 | 18.646 | 4.934 | 9.686 | 1.814 | -30.639 | 8.215 | 1.490 | -20.097 | 16.201 | 21.838 |
| -20.569 | -13.213 | -17.114 | 22.172 | -7.005 | 26.789 | 21.102 | -4.949 | -23.148 | -26.706 | 11.718 | 14.317 | 3.733 | 25.962 |

| | | | | | | | | | | | | | |
|---|---|---|---|---|---|---|---|---|---|---|---|---|---|
| -28.888 | -24.525 | 11.412 | 18.523 | 15.627 | 16.256 | 0.709 | 13.897 | 13.329 | 4.743 | -3.380 | -9.916 | -18.296 | -2.084 |
| 29.064 | 6.515 | 23.640 | -13.852 | 23.423 | 0.999 | 4.921 | 3.976 | 4.947 | -27.080 | 10.730 | -15.748 | 24.114 | -8.577 |
| 7.162 | -18.298 | 17.602 | 26.444 | 26.796 | 12.555 | -25.812 | -4.503 | -12.409 | 29.531 | 11.088 | -25.130 | 25.578 | 20.069 |
| -14.683 | 23.968 | 12.946 | 17.849 | 22.130 | 1.776 | -20.764 | 9.724 | 25.498 | -24.502 | 10.386 | -25.507 | 24.289 | 9.718 |
| 19.516 | 19.553 | 22.057 | -4.768 | -5.216 | -14.971 | 10.835 | 18.317 | -13.140 | 24.723 | -24.374 | 29.639 | 26.776 | 20.516 |
| -18.961 | -26.642 | -25.443 | -28.616 | -10.975 | -23.422 | -10.744 | -21.650 | -20.354 | -28.757 | -20.634 | -12.981 | 16.679 | -6.688 |
| 22.686 | -19.956 | 6.287 | 23.192 | 10.195 | 24.837 | 1.430 | -8.247 | 14.776 | 27.719 | 13.813 | 26.357 | 11.561 | -11.575 |
| -25.624 | 10.269 | 5.406 | 7.421 | 8.835 | 5.754 | 7.952 | 4.801 | 22.402 | 6.637 | 9.992 | 15.775 | -13.629 | -4.049 |
| 26.326 | 9.024 | -10.266 | -7.931 | -8.675 | -10.924 | 19.616 | -22.237 | 29.010 | 2.881 | -21.377 | 18.889 | 23.566 | -26.982 |
| -3.487 | -14.045 | -9.097 | -27.031 | -2.108 | 24.267 | 6.984 | -7.492 | -0.879 | 26.642 | -8.035 | 27.007 | 21.025 | 0.140 |
| -8.124 | 7.879 | 13.854 | -25.408 | -8.077 | -5.246 | -5.167 | -1.131 | 9.541 | -8.956 | -13.703 | 11.470 | 20.241 | -24.427 |
| 1.141 | -4.583 | -15.795 | 22.124 | -20.754 | -19.599 | 11.317 | -20.484 | -9.533 | -22.452 | -12.224 | 27.200 | 10.016 | 6.343 |
| -26.633 | -6.442 | 26.470 | 23.270 | -23.878 | 10.142 | -23.869 | -5.557 | 13.465 | 5.423 | 19.220 | 30.358 | 5.539 | 27.832 |
| 4.162 | 26.633 | -20.598 | -28.361 | 16.406 | -7.697 | -6.042 | 3.831 | 3.609 | 26.359 | -4.800 | 5.676 | -4.529 | 17.284 |
| -5.623 | 5.732 | 22.842 | 19.496 | 20.779 | 18.979 | -7.267 | 2.240 | -0.208 | -2.762 | 6.841 | 24.484 | 26.167 | 26.842 |
| 4.891 | 2.900 | -1.021 | -27.918 | 2.766 | -13.220 | 5.870 | 10.847 | 1.516 | -8.379 | 22.146 | 25.965 | -9.493 | -6.411 |
| -4.027 | -5.137 | -6.262 | 7.590 | -6.468 | 23.120 | 0.297 | -1.655 | -26.334 | 3.575 | -16.920 | 29.348 | 9.304 | -7.505 |
| 9.025 | -24.959 | -3.025 | -11.947 | 10.412 | -9.924 | -0.594 | 4.271 | -2.760 | 27.188 | -1.835 | 10.905 | 8.502 | 16.901 |
| 18.260 | -26.415 | -23.889 | 14.772 | -22.635 | 12.097 | 23.098 | -5.999 | 27.569 | -14.503 | 18.622 | -14.187 | 26.947 | -5.979 |
| -2.409 | -9.659 | 8.923 | -26.371 | -0.586 | 6.579 | 20.246 | 7.157 | 8.236 | 28.324 | 16.399 | 28.798 | 22.715 | 15.608 |
| -11.347 | 17.285 | 26.219 | 28.061 | 4.203 | 21.660 | -12.000 | 4.900 | 25.919 | -23.439 | -3.872 | -30.184 | -2.291 | -18.628 |
| -18.082 | 11.583 | -6.867 | 11.328 | 1.294 | -25.228 | 12.756 | 8.892 | 31.846 | -14.832 | -17.668 | 16.428 | 13.747 | 4.472 |
| 28.564 | -21.493 | -13.430 | -12.496 | 11.390 | 26.790 | 7.094 | -4.233 | 18.017 | 25.350 | -25.263 | 19.895 | 5.759 | 27.390 |
| 27.201 | 23.211 | -24.924 | -9.521 | 3.452 | -2.295 | -2.102 | -9.577 | -12.563 | -5.161 | 26.104 | 1.212 | 24.497 | 6.766 |
| -0.755 | -7.477 | 23.844 | 30.580 | -18.097 | 15.457 | 5.609 | 17.957 | 22.581 | -14.277 | -15.617 | -10.029 | 5.367 | 20.863 |
| 23.877 | -6.060 | 19.042 | 28.900 | -4.475 | -22.814 | -27.907 | -25.702 | 14.890 | -4.203 | 10.242 | -8.317 | -2.828 | -12.840 |
| 6.824 | -14.533 | -5.110 | 24.156 | -11.769 | 28.388 | 6.896 | 23.914 | 31.015 | 23.571 | 13.676 | -19.710 | 6.671 | 5.238 |
| 27.890 | 21.915 | -19.003 | 15.226 | -12.444 | 22.118 | -21.438 | 4.387 | 19.390 | -1.340 | 20.244 | 13.669 | 16.161 | -2.461 |
| 25.509 | -0.513 | -26.529 | 6.368 | -15.781 | -26.139 | 9.333 | -4.951 | 17.615 | 5.337 | -13.250 | 25.438 | 24.284 | 12.506 |
| 18.696 | -25.536 | -26.077 | -11.000 | 10.238 | -5.818 | 18.301 | 6.345 | 26.721 | -23.150 | -15.781 | 26.368 | 22.762 | 11.400 |
| 29.082 | 10.217 | -20.583 | -27.553 | -5.626 | 21.978 | -27.270 | -8.998 | 16.369 | 7.721 | -5.782 | -20.891 | 13.377 | 4.621 |
| -28.346 | -4.375 | 22.924 | 18.063 | 21.132 | -24.116 | -29.615 | 18.622 | 13.007 | 4.933 | -18.124 | 24.853 | 12.879 | 3.453 |
| 14.702 | 13.004 | -17.699 | 1.181 | 9.429 | 20.433 | 20.335 | 10.363 | 23.156 | 9.061 | -2.452 | -28.452 | 19.824 | -0.526 |
| 24.108 | -16.898 | -6.338 | 21.589 | 10.727 | 24.302 | 17.465 | 2.588 | -31.026 | 1.644 | -1.416 | 25.923 | -11.726 | 26.083 |
| 22.203 | -22.610 | -22.954 | 17.163 | 0.566 | -25.879 | 5.446 | -6.058 | -15.396 | 0.210 | -21.093 | -30.404 | -16.410 | 12.544 |
| 9.775 | -23.134 | 15.566 | 27.910 | -9.050 | 15.024 | -17.119 | -3.520 | 30.220 | -16.692 | 1.045 | 27.552 | 22.250 | 17.081 |

| | | | | | | | | | | | | | |
|---|---|---|---|---|---|---|---|---|---|---|---|---|---|
| 6.494 | -12.258 | -20.022 | 10.751 | 12.788 | 1.534 | -9.252 | -6.673 | 10.279 | -22.554 | -17.253 | -6.272 | 12.134 | 0.558 |
| 3.361 | -19.217 | 11.233 | -29.968 | 7.377 | -27.482 | -21.665 | 8.400 | -28.795 | 24.348 | -6.387 | 29.734 | 17.652 | 19.439 |
| 21.058 | 23.544 | 27.006 | -18.261 | -6.831 | 7.776 | 15.149 | 0.837 | -14.226 | 6.382 | -8.641 | 1.129 | -14.335 | 8.762 |
| -11.202 | 22.639 | -4.314 | -24.435 | 3.225 | 1.003 | 26.992 | 19.228 | -19.387 | -6.318 | -14.035 | 23.249 | 7.837 | 9.761 |
| -26.993 | 18.456 | 0.315 | 22.734 | 4.960 | -10.765 | -24.380 | -14.756 | 13.925 | -4.453 | -10.173 | 22.327 | 9.938 | 6.317 |
| -27.903 | -4.307 | -18.253 | -19.820 | 8.933 | 26.712 | 14.305 | 1.941 | 4.271 | 23.410 | 0.607 | -24.727 | 3.871 | 20.147 |
| 22.921 | 17.771 | 25.159 | 11.515 | 4.701 | 2.335 | -8.408 | 4.539 | -8.753 | 24.777 | -4.264 | 29.393 | 6.869 | -9.031 |
| -0.451 | 22.345 | 8.264 | -18.568 | 21.875 | 10.845 | 9.902 | -6.600 | 27.605 | 3.483 | -20.058 | 23.729 | -14.120 | 20.848 |
| -27.536 | 2.018 | -12.036 | -20.825 | -13.955 | 18.567 | -7.482 | 18.169 | 26.221 | 19.619 | -2.891 | 27.789 | 4.085 | 24.796 |
| 24.582 | -17.803 | 26.645 | 24.632 | -16.686 | 5.030 | -16.319 | 21.800 | -31.528 | 8.789 | -5.716 | -3.843 | 12.463 | 1.229 |
| -4.309 | 15.910 | -26.233 | -26.014 | 2.628 | -1.701 | 12.819 | -14.083 | 5.460 | 27.733 | -7.809 | -12.797 | 17.167 | 25.973 |
| -20.650 | -5.760 | -16.410 | -9.371 | 23.481 | 26.045 | -15.745 | -16.364 | -24.044 | -13.496 | -14.097 | -14.687 | 12.054 | 20.216 |
| -26.201 | 17.548 | -20.015 | -18.738 | -0.777 | -17.935 | -19.324 | -11.146 | 6.496 | 13.369 | 10.060 | -26.704 | -9.841 | -24.022 |
| -18.890 | 3.999 | -22.480 | -11.951 | 20.049 | -7.641 | -30.219 | -11.621 | 14.832 | -11.069 | 0.299 | -25.561 | -21.238 | 5.570 |
| 0.854 | 23.634 | 23.851 | -8.109 | 4.435 | -17.298 | 8.225 | 26.317 | 16.639 | 1.744 | 9.058 | -7.737 | 24.789 | -19.120 |
| 25.900 | -21.246 | -8.869 | -30.746 | -1.630 | 4.566 | 20.403 | -25.073 | -5.782 | -5.430 | -7.797 | 22.894 | 26.022 | 24.214 |
| -24.260 | 17.369 | -11.695 | 12.489 | 0.890 | 17.472 | 29.256 | -7.837 | -30.318 | 24.388 | -4.828 | 8.802 | 25.334 | 23.341 |
| -23.032 | -3.705 | 24.391 | 21.047 | -10.372 | -12.859 | -16.651 | 19.161 | 21.356 | 2.043 | -3.403 | 24.347 | -6.209 | 8.014 |
| -9.405 | -22.192 | -12.430 | -23.328 | -3.577 | -13.245 | 6.825 | 9.254 | -18.796 | 1.140 | -14.932 | 10.872 | 6.548 | -7.907 |
| -1.096 | -9.869 | -16.929 | 6.297 | -25.232 | 16.543 | -25.250 | 20.848 | 17.726 | 22.156 | -8.598 | 26.598 | -4.974 | 18.658 |
| -15.727 | 4.285 | 2.404 | 24.172 | 21.599 | 7.735 | -21.001 | -0.643 | 26.614 | 6.498 | 1.723 | -29.182 | 12.641 | 11.930 |
| -19.498 | -5.254 | 27.731 | -8.079 | 1.946 | 29.904 | -22.245 | 19.575 | -4.405 | 4.578 | 19.600 | -26.277 | 24.019 | 22.216 |
| 21.700 | -5.678 | 18.453 | -30.278 | -0.131 | 22.578 | -2.492 | -10.923 | 14.367 | -20.095 | 19.897 | 19.414 | -12.678 | -7.411 |
| -3.055 | 20.028 | -5.693 | -26.891 | 11.895 | -7.074 | 23.820 | -7.374 | 3.208 | -0.935 | -16.871 | 15.917 | 20.744 | -8.298 |
| -6.599 | -19.546 | -11.137 | -28.692 | -14.096 | 21.443 | 12.688 | 22.936 | -9.807 | -8.520 | 18.395 | -19.137 | 2.041 | -7.010 |
| 4.580 | 5.809 | -17.529 | 17.299 | -13.635 | -20.582 | -15.842 | -25.005 | 13.656 | 11.512 | 19.050 | 8.887 | 14.143 | 20.700 |
| -25.221 | -6.142 | -20.193 | 13.436 | 3.167 | -28.351 | -28.365 | 24.876 | 7.269 | 26.867 | 3.301 | 22.383 | -22.981 | 16.445 |
| 20.452 | 22.985 | 20.857 | -6.411 | 22.002 | 18.007 | -16.945 | 13.560 | -3.785 | -14.301 | 8.178 | 25.783 | 26.385 | -25.870 |
| -10.273 | 12.628 | -27.334 | -19.899 | 24.979 | -0.110 | -20.976 | -26.749 | 29.756 | 22.824 | 12.297 | 25.058 | 20.124 | 17.859 |
| 27.283 | -23.226 | 16.882 | -20.898 | -6.017 | -4.182 | 0.873 | 4.917 | -2.100 | 4.213 | 24.075 | 5.795 | -10.597 | -2.510 |
| 28.284 | 2.970 | 19.545 | -22.837 | 3.592 | 24.347 | 18.726 | 22.298 | 28.856 | -20.831 | -8.358 | 4.443 | -6.113 | 16.914 |
| -2.384 | -26.325 | 1.361 | -17.598 | 24.111 | 23.270 | -12.155 | -0.782 | 16.150 | -2.262 | 20.625 | 21.134 | 23.270 | 8.221 |
| 10.771 | 1.544 | -15.754 | -18.007 | -12.151 | 19.287 | -27.554 | 14.815 | -27.050 | 15.122 | 22.580 | 14.315 | 2.975 | 4.925 |
| -11.847 | -16.348 | 10.984 | -9.252 | -1.233 | -21.926 | 5.557 | -6.523 | 8.338 | 16.652 | 15.503 | -28.189 | 22.061 | -14.574 |
| -26.287 | 7.281 | -28.476 | 26.085 | -4.579 | -13.154 | -8.360 | -12.256 | 15.332 | 3.714 | -5.233 | -19.900 | 19.122 | -5.446 |
| 14.322 | -10.461 | -23.063 | 21.964 | -7.583 | 1.144 | -19.821 | -22.917 | 23.892 | -3.764 | -12.585 | 20.251 | -9.501 | 3.261 |

| | | | | | | | | | | | | | |
|---|---|---|---|---|---|---|---|---|---|---|---|---|---|
| 26.435 | -5.334 | 19.864 | 14.869 | 2.252 | -2.760 | -22.467 | 12.288 | 21.412 | -26.485 | 16.884 | -15.120 | -19.504 | 21.745 |
| 18.223 | -7.044 | -3.195 | -16.912 | -0.293 | 29.195 | -17.824 | -10.140 | 26.805 | -13.893 | 2.566 | 13.305 | 4.682 | 9.692 |
| 29.127 | 26.556 | 2.559 | 29.015 | 3.241 | 29.674 | 28.245 | -18.546 | 25.487 | -13.323 | 24.364 | -7.703 | 12.927 | 5.964 |
| 9.184 | -20.068 | -26.965 | 30.094 | -5.561 | -7.219 | 27.994 | 10.042 | 4.550 | 20.049 | 1.946 | -4.894 | 11.663 | 14.464 |
| 9.935 | 24.138 | -19.661 | -19.403 | -2.853 | 21.375 | -11.274 | 19.113 | 3.661 | -4.228 | 15.394 | 26.435 | 22.554 | 15.116 |
| -13.959 | -24.426 | 12.518 | -27.418 | -0.964 | 26.078 | 18.764 | -1.324 | -31.342 | -27.347 | -11.252 | -26.709 | -23.604 | 6.342 |
| 6.440 | -16.396 | -20.406 | -15.707 | -13.928 | 27.896 | -23.837 | 21.009 | -21.601 | -5.367 | -25.271 | -27.981 | 18.588 | 8.220 |
| 14.579 | 8.036 | 11.562 | -21.377 | 3.508 | -5.086 | 30.342 | 16.736 | -23.201 | 7.282 | 2.569 | 1.551 | -14.074 | 3.247 |
| 5.577 | 0.607 | 20.709 | 27.094 | 9.445 | 28.941 | 19.909 | -22.257 | 18.936 | -26.311 | -19.749 | 19.995 | 27.124 | -16.170 |
| -6.855 | -17.133 | -11.639 | 23.858 | 11.946 | 15.245 | 15.840 | -17.987 | 30.842 | -10.079 | -11.958 | -5.833 | 16.168 | -3.530 |
| 7.144 | -6.562 | -28.249 | -24.274 | -0.118 | -29.418 | 10.472 | 17.731 | 9.025 | 18.629 | 26.562 | 25.670 | -6.855 | 12.973 |
| 4.827 | -23.340 | -1.258 | -18.235 | -9.918 | -21.460 | -17.160 | 18.292 | -28.805 | -18.919 | -21.416 | 0.073 | 8.001 | -9.571 |
| -25.468 | 18.170 | -13.180 | 28.522 | 12.592 | 2.752 | -18.740 | 0.416 | 17.443 | 11.042 | -16.299 | -27.221 | 21.103 | -22.898 |
| -26.413 | -3.854 | 0.158 | -24.709 | 17.834 | -4.125 | -20.342 | 23.495 | 10.563 | 6.062 | 13.354 | -5.167 | -21.258 | 7.396 |
| -13.756 | 22.642 | -10.080 | -8.264 | 1.847 | -14.791 | -23.672 | -2.591 | -22.509 | -2.436 | -18.526 | 17.407 | 23.844 | 16.018 |
| 2.650 | 12.894 | -14.693 | 9.779 | 19.240 | 6.792 | 1.558 | -30.435 | -20.776 | -1.921 | -10.382 | 24.277 | 10.439 | 9.846 |
| 0.577 | -0.905 | -20.275 | 29.373 | 5.071 | -4.648 | -8.591 | 19.905 | 6.445 | -15.292 | 24.441 | 10.696 | 0.599 | 11.425 |
| -4.546 | 2.615 | 4.079 | 7.298 | 0.216 | 11.816 | -16.484 | 11.205 | 0.144 | -20.128 | -27.676 | 16.216 | 25.440 | 20.092 |
| 27.910 | -4.759 | -21.587 | -29.172 | -0.446 | 17.921 | 19.629 | 24.620 | 7.128 | -22.796 | 18.905 | 14.125 | 18.170 | -24.480 |
| -10.007 | -16.128 | 19.243 | -17.842 | -8.463 | -12.071 | 4.686 | 12.674 | 5.915 | 6.872 | 27.084 | 11.356 | 24.853 | 15.434 |
| 25.651 | 19.963 | -22.098 | -17.206 | -4.188 | 19.881 | -14.588 | -6.949 | 21.795 | 23.748 | 8.620 | 3.115 | -18.189 | 8.549 |
| 17.615 | 19.086 | 25.035 | -11.373 | 1.476 | -24.552 | 16.928 | 17.039 | 9.559 | 5.759 | -1.083 | -22.187 | -22.012 | 17.340 |
| 8.811 | -2.923 | 21.437 | -21.032 | 4.630 | 8.214 | 23.298 | 11.337 | -27.453 | -0.810 | -17.573 | 16.952 | 26.913 | 2.475 |
| 21.774 | -6.465 | -16.629 | -19.928 | -6.441 | 3.671 | 28.232 | 7.455 | 3.137 | 8.685 | 19.640 | -14.110 | 23.012 | -2.426 |
| -7.907 | -1.583 | 21.943 | -19.356 | -2.828 | 5.523 | 3.960 | -21.268 | -8.063 | 7.770 | 1.235 | 17.956 | 24.392 | 21.294 |
| -16.152 | 18.370 | -17.500 | -24.828 | 2.861 | 20.820 | 20.580 | -0.531 | 12.409 | 16.522 | -11.331 | 22.805 | -3.699 | 25.395 |
| -17.156 | 15.543 | -18.907 | -30.284 | 0.692 | 28.780 | -0.655 | 13.502 | 15.540 | -24.635 | 6.583 | 7.400 | 16.352 | 19.528 |
| 14.691 | -7.417 | 18.041 | -20.492 | 11.742 | -19.699 | -22.860 | -31.332 | -30.326 | -26.294 | -21.891 | 20.927 | 13.122 | 1.281 |
| -5.146 | -11.284 | 28.846 | 15.254 | 8.833 | -29.022 | -22.185 | 4.437 | 29.852 | 15.773 | -7.016 | 21.876 | 13.716 | 10.000 |
| -13.295 | 21.964 | 11.793 | 6.409 | 18.389 | -15.445 | -19.662 | 8.857 | 14.996 | -0.006 | 26.488 | 9.971 | -27.090 | 8.104 |
| 23.818 | -4.857 | 0.996 | 18.293 | 20.590 | 21.645 | -16.566 | -18.893 | -7.934 | -7.489 | 22.677 | -25.714 | -0.950 | 13.370 |
| -8.745 | -6.990 | 14.465 | -8.327 | 5.477 | 30.094 | 12.852 | 17.889 | 17.834 | 13.278 | 5.771 | -1.550 | 20.879 | 1.063 |
| 5.717 | -22.901 | -14.268 | 5.963 | -3.210 | 17.389 | -20.361 | 30.773 | 19.042 | 2.061 | -9.706 | 2.642 | -15.750 | 13.833 |
| -19.611 | 22.517 | 18.170 | -26.103 | -0.407 | -4.594 | 1.061 | -11.443 | -0.520 | 5.933 | 22.653 | 23.880 | 21.472 | 22.542 |
| -27.311 | 4.108 | -2.370 | -30.211 | -9.902 | -9.686 | 28.268 | -2.920 | -26.783 | 0.227 | -3.530 | 2.196 | -25.443 | -2.675 |
| 22.860 | -2.173 | 19.481 | -25.602 | 23.720 | 26.096 | 4.837 | -14.232 | 11.708 | 5.119 | -21.537 | -10.903 | -28.112 | -10.020 |

| | | | | | | | | | | | | | |
|---|---|---|---|---|---|---|---|---|---|---|---|---|---|
| 17.515 | 9.665 | 21.521 | -28.829 | 12.804 | 26.814 | 1.701 | -30.327 | -9.887 | -1.992 | 24.063 | 24.678 | 26.153 | 11.207 |
| -1.010 | 14.008 | 9.145 | 21.940 | -10.271 | 8.480 | 7.697 | 28.943 | 20.813 | 4.168 | -20.861 | 15.678 | 23.145 | 9.485 |
| -4.222 | 5.341 | -18.635 | 14.060 | 22.958 | 16.323 | 0.190 | 8.469 | 12.494 | -4.306 | -27.717 | 14.298 | -8.349 | 18.956 |
| -5.841 | -18.514 | 23.367 | 8.083 | 6.304 | 19.675 | -15.730 | 5.988 | -4.808 | 12.655 | -2.438 | 1.367 | 22.389 | 4.828 |
| -23.745 | 14.610 | 12.195 | -11.960 | 0.831 | 31.021 | 16.165 | 24.833 | 9.522 | 8.120 | -1.159 | 6.433 | 2.820 | 25.784 |
| 11.881 | 12.663 | 5.441 | -25.010 | -4.883 | 23.429 | -27.369 | -7.899 | 13.731 | -2.177 | 27.099 | 3.734 | 5.626 | 19.344 |
| -1.864 | 23.750 | -13.308 | 28.076 | -8.754 | -24.946 | -19.950 | -26.472 | -25.044 | 0.398 | 11.774 | -4.216 | -0.024 | 7.798 |
| 2.449 | 6.338 | -6.062 | 18.853 | 0.066 | 28.966 | -8.663 | 25.473 | 8.822 | 10.055 | 26.198 | 12.899 | -25.293 | 24.755 |
| -15.844 | -24.014 | -6.795 | -26.842 | 1.397 | -16.454 | 2.329 | -28.681 | 16.522 | -8.271 | 20.963 | 8.726 | -19.251 | 23.748 |
| 18.516 | -5.384 | -1.616 | 12.056 | 15.932 | 12.783 | 5.191 | -22.053 | 18.468 | 17.359 | 14.329 | -23.568 | 24.505 | 21.337 |
| -25.735 | 1.831 | 16.113 | 7.217 | 4.508 | -5.956 | -26.436 | -16.134 | 3.926 | -9.587 | -9.070 | 24.497 | 5.083 | 19.696 |
| -6.164 | -0.517 | -11.795 | -17.870 | 1.890 | 3.673 | 8.769 | -23.579 | 29.585 | -5.774 | 24.490 | -2.501 | -24.807 | -10.574 |
| 16.955 | 16.945 | -15.531 | -15.878 | 22.575 | 22.860 | 6.007 | -17.829 | 13.177 | -18.707 | 11.863 | 13.751 | 28.768 | 25.229 |
| -18.767 | 7.880 | -9.578 | 13.694 | 2.514 | 5.860 | 4.623 | 10.920 | -6.319 | -7.434 | -7.517 | 1.987 | 20.142 | 18.287 |
| 27.818 | -6.409 | -14.037 | 4.475 | 6.626 | -13.737 | -14.097 | -4.888 | -20.151 | -6.099 | 19.349 | 17.518 | -0.857 | 14.032 |
| -26.534 | -23.357 | 28.635 | 5.542 | 21.842 | -1.823 | -26.771 | -24.716 | -24.182 | -15.965 | -4.329 | -14.208 | -18.278 | 13.207 |
| -5.247 | 9.491 | 11.146 | -23.673 | 7.185 | -23.186 | -10.367 | -13.917 | -11.138 | -8.625 | 4.496 | -8.668 | 25.983 | 4.683 |
| 19.366 | -9.936 | -18.651 | 27.883 | 19.385 | 26.172 | -3.992 | 23.363 | 28.222 | -1.962 | 3.867 | -0.414 | 17.068 | -0.439 |
| 10.816 | 18.533 | 23.560 | 26.024 | -10.983 | 22.471 | -24.773 | -20.081 | 23.697 | 16.690 | -21.582 | 15.329 | 11.357 | 20.273 |
| 27.401 | 17.093 | 17.437 | -14.460 | -15.905 | 24.214 | 22.273 | -20.890 | -1.987 | 0.493 | 5.702 | -7.342 | -5.574 | 12.003 |
| -6.437 | 8.251 | 22.487 | -15.478 | 23.284 | -19.690 | 9.608 | 24.005 | -17.319 | -3.513 | -1.296 | -12.708 | -23.626 | 26.816 |
| -9.938 | -1.690 | 20.856 | -17.638 | -5.449 | 12.203 | -2.565 | -11.852 | -16.808 | 22.718 | 18.334 | -1.516 | 22.121 | 22.351 |
| 24.347 | -12.108 | 15.384 | 10.496 | -6.265 | 7.030 | -1.277 | -7.049 | 8.297 | -2.742 | -17.045 | -5.068 | -22.349 | 16.464 |
| -14.051 | -19.044 | -21.186 | -17.179 | 26.233 | 14.703 | -0.001 | -13.919 | -25.207 | -1.820 | -25.285 | 11.613 | 21.419 | -15.451 |
| 26.469 | -14.440 | -1.357 | -25.544 | -16.506 | -10.543 | 19.109 | 4.001 | 21.683 | -9.576 | 28.243 | -9.592 | -28.119 | -20.576 |
| 22.114 | 1.093 | 20.200 | 6.792 | 2.384 | -18.139 | 4.931 | -0.304 | -7.178 | -21.465 | 27.632 | -12.880 | 23.476 | 20.910 |
| -24.548 | 6.229 | -7.881 | -23.375 | -23.743 | -21.265 | 2.992 | 16.741 | 27.581 | -0.233 | 25.829 | -26.061 | 27.122 | 11.510 |
| 25.335 | -2.920 | 26.315 | -6.062 | 14.243 | -24.757 | 25.732 | -18.925 | 3.771 | -3.998 | 6.619 | -10.648 | -25.623 | 1.492 |
| 26.069 | -8.083 | 9.825 | 10.437 | 15.912 | 16.908 | -7.232 | -17.392 | -23.671 | -13.997 | -20.671 | -14.417 | -19.781 | 7.374 |
| 20.656 | -13.289 | 3.835 | 20.531 | 9.066 | 22.272 | -21.925 | -14.327 | 3.411 | -8.732 | 17.022 | 8.843 | 20.267 | 16.987 |
| 14.222 | -16.583 | -27.970 | 16.079 | 3.170 | -8.662 | 1.456 | -25.446 | 22.311 | -3.518 | -2.905 | 24.346 | 3.143 | -4.896 |
| -1.340 | -18.422 | 24.798 | 19.661 | -15.085 | 14.896 | 2.747 | -27.954 | 20.782 | 6.557 | 17.416 | 16.817 | -25.305 | -24.928 |
| -3.579 | -25.017 | -12.109 | 24.141 | -19.107 | 27.346 | -9.329 | -15.421 | 20.043 | -4.379 | 21.101 | 10.418 | 27.794 | 3.587 |
| -25.196 | -20.536 | -15.631 | -5.231 | 10.881 | 8.580 | -2.288 | -3.093 | 5.126 | -13.618 | 22.106 | -0.264 | 24.304 | 0.243 |
| 16.850 | 5.006 | -20.897 | 25.499 | -24.616 | 28.278 | 7.880 | 5.189 | 4.802 | -1.304 | 3.950 | -3.705 | 26.711 | 13.717 |
| 23.318 | 11.673 | -4.172 | -3.595 | -12.716 | 12.873 | 23.071 | -18.995 | -19.186 | 17.320 | 24.793 | 21.131 | 21.542 | 21.564 |

| | | | | | | | | | | | | | |
|---|---|---|---|---|---|---|---|---|---|---|---|---|---|
| 25.423 | -0.993 | 12.507 | -27.564 | 0.656 | 7.522 | 2.131 | -23.357 | 19.180 | 23.066 | 9.964 | -18.475 | -19.814 | 6.626 |
| -9.302 | -11.072 | -25.600 | 24.416 | -14.281 | -17.243 | 9.271 | -24.010 | -22.181 | 1.368 | 11.243 | -8.319 | 19.463 | 4.838 |
| 12.981 | 6.107 | 3.072 | -13.174 | 10.180 | 28.720 | 19.661 | -5.403 | -27.936 | -3.519 | 25.779 | 18.254 | 30.027 | -1.666 |
| 2.102 | 2.641 | 18.759 | -21.177 | 19.687 | 20.986 | -9.929 | 0.365 | 26.486 | -6.601 | -20.161 | -4.135 | 23.095 | 12.202 |
| -4.017 | -2.347 | 16.194 | -13.714 | 0.281 | 8.064 | 8.686 | -4.812 | 13.241 | 4.180 | -29.934 | -25.166 | 15.037 | 23.361 |
| 5.860 | -22.600 | -16.576 | -23.364 | -3.691 | 25.655 | -6.901 | -3.426 | -6.446 | 10.495 | 25.355 | 5.541 | 20.867 | 10.235 |
| 8.717 | 12.716 | 14.090 | 7.728 | 7.552 | 19.068 | 20.635 | 23.009 | 21.426 | 0.564 | -1.963 | 12.102 | 24.539 | 14.419 |
| -18.881 | 17.134 | -20.151 | 21.497 | 6.169 | -0.698 | -8.012 | -16.317 | -7.372 | -19.651 | 21.165 | 10.721 | 25.798 | 13.957 |
| -20.180 | -4.148 | 30.734 | 9.124 | 13.960 | -22.084 | -19.478 | -12.305 | -4.721 | -20.488 | 12.255 | 7.357 | -25.288 | 15.817 |
| 26.495 | -13.882 | 23.625 | 5.957 | -5.462 | 30.293 | -10.060 | -19.076 | 9.279 | 15.334 | -16.032 | -20.943 | -23.276 | -10.205 |
| -16.140 | -8.012 | 20.751 | -20.938 | 18.674 | -28.061 | 10.736 | 15.962 | 6.190 | 6.949 | 14.945 | 7.366 | -28.593 | 11.877 |
| 18.761 | 19.596 | -13.463 | 20.100 | 0.557 | -6.148 | 17.801 | -18.540 | -20.482 | 2.386 | -28.812 | -16.438 | 25.149 | 0.334 |
| 22.792 | 1.921 | 8.943 | 18.190 | 6.472 | -18.162 | 10.116 | 19.436 | -25.270 | -1.730 | -10.270 | 10.613 | 4.420 | 4.533 |
| 20.000 | 1.136 | 7.347 | 12.763 | 5.220 | 29.860 | -2.925 | 8.882 | -21.629 | -8.690 | -0.202 | -9.992 | 28.985 | 1.734 |
| 5.016 | -8.446 | 10.741 | 3.613 | -4.620 | 10.612 | 18.259 | -3.085 | -4.133 | 16.259 | -3.872 | -21.979 | 23.577 | 18.543 |
| 15.859 | 13.238 | 10.228 | 1.249 | 23.512 | 13.757 | 8.541 | -0.218 | 26.837 | 9.015 | 23.256 | -23.764 | 23.060 | 26.485 |
| 17.736 | -5.736 | -8.291 | 24.640 | -1.616 | 2.511 | -0.640 | -22.773 | 4.209 | 8.162 | 9.335 | 22.412 | 29.545 | -1.597 |
| 16.843 | -7.587 | -10.595 | 5.020 | -22.510 | 16.046 | -9.046 | 7.483 | 19.816 | -14.321 | -20.518 | 21.660 | 27.899 | 13.506 |
| 7.177 | -11.055 | -23.709 | -24.200 | -11.800 | 17.845 | 17.559 | 13.615 | 21.944 | -12.135 | -29.962 | 20.539 | -0.860 | 0.722 |
| -24.494 | 21.333 | 7.144 | -27.011 | -16.893 | -30.259 | 11.100 | 14.664 | 15.650 | -9.518 | -6.900 | -5.704 | 30.382 | 16.969 |
| 9.617 | 3.878 | -9.790 | -3.250 | -12.975 | -26.094 | -16.804 | 4.194 | -2.154 | 5.451 | 19.313 | 17.723 | 19.819 | 19.223 |
| -15.707 | 18.550 | -19.230 | 27.041 | 28.039 | -2.254 | -25.294 | 0.841 | 20.287 | -1.957 | 25.805 | 27.041 | 11.671 | 6.383 |
| -6.280 | 8.860 | 28.986 | 24.027 | -20.447 | -29.361 | 0.540 | -4.847 | 6.543 | -4.096 | 27.336 | -25.601 | 25.998 | 8.692 |
| 10.491 | -25.093 | -24.921 | -8.868 | -7.750 | 26.023 | 25.085 | -10.467 | 11.158 | -4.477 | 27.726 | 6.682 | 21.719 | -8.395 |
| -4.285 | -0.007 | -12.887 | -14.160 | 2.065 | 23.114 | 7.174 | 10.930 | -12.631 | -10.439 | 13.166 | -0.703 | 27.714 | 20.773 |
| -25.619 | 23.460 | 14.105 | -5.393 | 7.143 | 18.738 | 20.476 | 25.682 | -18.171 | -21.104 | 21.668 | -18.429 | -2.484 | 2.977 |
| 8.185 | -14.727 | -22.204 | 14.775 | -24.003 | 9.923 | -19.039 | 0.531 | 4.577 | -17.172 | -22.108 | -27.894 | 26.461 | 17.838 |
| -23.091 | 22.087 | 6.623 | -17.546 | 11.757 | 13.389 | -24.660 | 18.968 | -7.407 | 0.436 | -8.973 | 23.983 | 0.498 | 1.618 |
| -25.835 | -21.586 | -24.692 | 19.629 | 18.601 | -24.687 | 11.149 | 19.672 | -1.372 | -8.030 | 28.925 | 13.086 | 29.852 | 22.225 |
| 1.682 | 1.973 | -25.528 | -16.624 | 8.094 | 30.460 | 16.507 | -12.882 | -2.917 | 1.665 | 4.014 | 14.894 | -29.251 | 23.200 |
| 0.955 | -2.762 | 6.033 | -0.549 | -18.189 | -23.997 | -0.141 | 20.332 | -3.620 | -5.730 | -27.611 | 9.798 | 4.745 | 11.078 |
| 9.296 | 12.849 | -26.849 | 25.555 | 21.268 | -27.782 | 24.223 | -1.217 | 24.339 | -13.862 | 19.163 | 16.606 | 23.650 | -7.171 |
| 20.704 | 11.171 | -0.248 | -22.174 | 23.168 | 1.246 | -2.150 | 6.396 | 5.160 | 4.417 | -29.865 | 13.974 | -30.373 | 9.099 |
| 14.062 | 4.646 | -5.325 | 20.840 | 19.595 | 1.646 | -21.280 | 4.392 | 7.488 | 1.141 | -0.180 | 23.087 | 22.531 | 14.358 |
| 10.122 | 20.500 | 27.799 | -5.120 | 7.772 | 5.693 | -19.627 | -25.341 | 21.529 | -15.923 | -14.009 | 19.200 | -20.859 | 15.190 |
| 22.230 | 7.383 | 23.527 | 3.393 | -4.621 | 20.958 | -22.516 | 24.651 | 20.422 | -0.435 | 28.381 | 21.984 | 21.245 | 6.603 |

| | | | | | | | | | | | | | |
|---|---|---|---|---|---|---|---|---|---|---|---|---|---|
| 6.857 | -3.866 | -1.877 | -7.048 | 24.814 | -20.593 | -23.999 | -16.327 | -0.499 | 24.529 | 17.300 | -4.684 | -0.846 | 4.715 |
| -12.127 | -3.243 | -15.057 | 7.030 | -26.825 | 29.268 | -8.769 | -0.260 | -20.585 | -12.367 | -28.227 | 27.624 | 23.799 | 13.424 |
| -23.738 | 21.806 | 16.915 | 23.841 | -2.614 | 25.732 | -23.005 | 8.038 | -11.376 | -15.330 | 16.025 | -24.512 | 9.372 | 4.129 |
| 21.068 | 11.530 | 20.035 | 24.918 | 5.096 | 1.030 | -14.758 | -7.371 | -28.466 | -18.964 | 8.772 | -24.927 | 30.873 | -16.516 |
| -19.667 | 8.326 | 0.625 | -19.168 | 20.539 | 19.204 | -18.551 | 18.929 | -1.712 | -6.634 | -4.445 | 26.604 | 17.559 | -18.440 |
| 18.247 | -5.660 | 12.148 | 16.773 | 15.358 | 6.317 | -19.569 | 12.206 | 18.551 | 14.873 | -14.436 | -11.247 | 19.129 | 25.759 |
| 14.992 | -21.695 | -17.271 | 8.975 | 7.371 | -21.826 | 17.710 | -3.636 | 7.949 | 14.387 | -24.822 | 20.081 | 28.261 | 7.175 |
| 23.779 | -8.256 | -3.507 | 19.320 | 16.157 | 27.107 | -6.285 | 14.461 | 2.654 | 8.856 | -10.810 | 23.984 | 20.065 | 13.778 |
| 17.418 | -12.198 | -10.733 | -10.623 | -3.667 | 23.882 | -26.852 | 3.273 | -8.151 | 20.953 | -31.368 | 11.793 | -19.161 | 27.395 |
| 5.168 | -2.045 | -11.942 | 24.658 | 6.872 | 21.458 | 21.375 | -13.165 | -17.510 | -8.398 | -13.785 | -8.595 | -26.412 | 3.269 |
| 16.162 | 9.210 | 24.893 | -8.930 | 11.853 | 15.376 | 8.475 | -17.579 | 25.105 | -2.629 | 16.593 | 9.936 | -0.279 | 1.905 |
| 10.654 | 5.937 | 3.096 | 12.991 | -1.890 | 2.101 | -4.891 | -15.377 | -22.228 | -23.357 | 14.837 | -16.867 | 19.839 | 8.391 |
| 15.862 | -3.187 | 2.100 | 20.270 | -15.929 | -18.658 | -14.888 | -7.142 | 13.600 | 2.284 | -15.003 | -0.586 | -1.004 | -2.620 |
| 8.348 | -7.204 | 24.203 | 3.868 | -28.915 | -7.618 | 9.710 | 8.280 | 2.896 | 7.412 | -27.617 | 1.976 | -21.402 | -24.482 |
| -7.701 | -17.021 | 14.864 | -23.668 | 28.076 | -28.761 | 23.817 | -21.323 | -0.883 | 27.564 | 10.539 | 10.076 | 28.967 | 14.073 |
| 16.683 | -6.880 | 0.140 | 17.901 | 14.273 | -29.341 | 15.769 | -26.802 | -15.064 | -1.549 | 30.449 | 25.780 | 25.671 | 22.552 |
| -13.029 | 3.005 | 30.240 | -8.283 | -10.656 | 4.657 | -26.158 | 27.637 | 21.809 | -25.317 | 21.969 | -19.690 | 18.819 | 15.996 |
| 12.685 | -10.279 | 20.931 | 0.577 | 7.567 | 9.459 | 24.692 | 7.395 | 20.066 | -4.316 | 26.404 | -28.394 | 26.366 | 10.170 |
| 20.127 | 25.892 | 12.008 | -16.735 | 16.023 | -15.887 | 18.685 | -25.789 | 2.396 | 3.930 | 29.452 | 22.982 | 22.381 | 4.640 |
| 0.754 | 21.299 | 23.061 | -9.861 | 4.701 | -4.545 | 22.917 | 23.560 | 7.379 | 8.745 | -0.310 | 6.929 | 11.582 | 25.901 |
| -23.325 | 0.008 | 28.055 | 15.215 | 19.406 | -5.142 | 20.560 | -2.615 | 16.758 | 6.211 | 24.375 | 13.399 | -26.932 | 19.775 |
| -2.493 | 0.654 | 21.386 | -19.824 | 4.902 | 18.330 | -0.485 | -26.502 | 14.978 | -19.857 | -11.505 | -22.066 | 19.812 | 25.017 |
| 2.209 | 4.802 | 19.511 | 8.761 | -11.137 | -0.789 | 5.332 | -4.135 | 24.066 | -15.300 | -12.930 | 21.718 | 21.083 | -13.245 |
| 13.540 | -1.750 | -27.894 | -0.783 | 13.550 | -4.156 | -18.413 | 12.212 | 10.920 | -17.463 | 25.125 | 18.924 | -0.598 | 17.731 |
| 8.554 | 10.037 | -19.842 | -1.031 | 7.238 | -13.515 | -24.846 | -17.837 | 3.930 | 10.262 | -1.570 | 27.124 | 23.277 | 12.202 |
| 24.436 | 0.037 | 10.329 | -22.253 | 27.127 | -11.597 | 12.901 | 21.413 | -7.794 | 13.130 | -4.282 | 0.031 | -19.479 | 6.432 |
| -21.354 | -4.286 | -11.453 | -10.906 | 24.114 | -29.817 | 25.242 | -8.105 | -13.143 | -5.371 | -0.052 | 20.589 | 27.773 | 22.376 |
| 20.220 | -5.231 | -22.683 | 25.109 | -19.412 | -24.540 | 24.629 | 10.327 | -1.241 | -4.685 | 6.921 | 26.367 | 17.208 | 21.139 |
| 19.290 | 9.682 | 23.445 | -24.628 | 19.865 | 27.796 | -26.886 | -2.910 | 0.748 | -1.843 | 9.644 | -7.610 | 5.076 | -7.906 |
| -13.944 | 25.195 | 10.945 | 21.550 | 0.699 | 26.178 | -0.768 | 1.123 | -15.970 | -0.908 | -3.710 | -27.565 | 22.759 | 15.524 |
| -18.850 | -9.166 | -17.289 | 19.178 | -24.880 | 14.494 | -2.276 | -5.569 | 8.879 | -7.710 | -25.640 | 0.375 | -12.501 | 26.466 |
| -5.167 | 23.810 | -30.702 | -22.777 | 9.196 | 28.618 | -4.801 | 27.389 | -2.751 | -16.651 | -2.635 | -13.580 | 26.858 | 18.317 |
| -14.595 | 6.658 | 7.438 | -9.864 | 17.358 | 7.908 | 23.458 | -26.315 | 17.979 | -20.656 | -30.744 | 25.184 | 24.791 | 1.107 |
| -3.034 | 23.225 | 3.380 | -19.747 | 21.914 | 3.808 | -23.520 | 19.579 | 1.552 | 0.941 | -30.199 | 19.284 | -20.607 | 19.187 |
| 6.771 | 22.506 | -7.989 | 0.027 | -13.564 | -4.338 | 27.537 | 16.468 | -25.658 | -5.082 | -14.139 | 24.311 | -5.099 | 13.037 |
| 5.218 | -12.773 | -17.744 | -2.706 | -10.087 | -29.220 | 11.432 | 2.883 | 10.926 | 1.456 | 23.064 | 7.171 | 28.442 | 13.858 |

| | | | | | | | | | | | | | |
|---|---|---|---|---|---|---|---|---|---|---|---|---|---|
| 14.496 | 21.014 | -13.066 | -7.544 | -5.702 | 19.038 | -15.926 | -16.260 | -18.915 | -13.214 | 12.537 | -1.381 | -4.711 | 24.641 |
| 7.411 | -18.971 | -23.529 | -16.617 | -0.786 | 11.686 | -22.765 | 20.226 | 24.757 | -10.827 | 28.555 | -11.097 | 19.978 | 14.942 |
| -0.944 | -3.527 | -25.794 | 19.854 | -23.163 | -2.281 | -19.331 | -1.921 | 5.330 | -17.935 | 25.232 | -24.106 | 13.881 | 17.283 |
| -4.995 | -6.893 | 18.095 | 3.465 | 4.647 | -12.200 | 5.512 | -10.189 | -5.537 | 2.382 | -20.247 | -10.337 | 0.701 | 26.853 |
| 15.563 | -7.313 | 14.003 | 25.454 | 28.367 | -28.694 | -20.007 | 0.990 | 3.239 | 19.662 | 23.526 | -26.615 | -7.320 | 8.277 |
| 11.978 | -15.985 | -18.841 | 10.290 | 21.664 | 27.642 | 21.527 | -2.564 | 16.286 | -11.932 | 26.173 | -28.696 | -29.543 | 12.246 |
| 7.632 | 13.237 | 13.424 | -12.709 | -18.266 | -1.711 | -28.007 | -27.829 | 8.412 | -19.970 | -29.778 | 17.184 | 20.588 | -26.460 |
| 16.559 | 26.906 | 4.169 | -24.123 | 6.726 | -18.773 | -22.372 | 19.861 | -9.928 | 24.883 | 26.530 | -28.358 | 5.440 | 11.693 |
| 11.437 | -6.549 | -5.107 | 3.718 | -0.429 | 27.085 | -21.637 | -3.226 | -8.373 | 4.926 | 1.551 | 8.941 | 2.067 | 5.414 |
| -17.455 | 25.985 | -30.898 | -1.812 | -29.542 | 12.426 | 18.328 | 27.783 | -15.679 | -28.252 | -19.540 | -7.782 | 26.803 | 13.830 |
| -11.970 | 16.549 | 30.970 | -15.561 | 27.163 | 1.916 | 2.042 | -4.747 | 25.092 | 7.720 | 30.606 | 1.194 | 15.866 | 28.319 |
| -23.148 | -4.054 | -2.704 | 15.702 | 9.111 | -26.116 | -10.730 | 24.363 | 0.487 | -22.576 | -26.706 | 28.819 | -8.562 | 6.891 |
| 11.436 | -12.874 | 29.951 | -21.398 | -4.722 | 16.751 | -9.714 | 29.203 | 5.782 | -13.244 | 8.629 | 26.133 | 3.008 | -29.311 |
| 10.788 | -24.161 | 29.298 | -6.820 | 9.731 | -6.160 | 3.828 | 19.681 | 26.114 | -5.415 | -20.009 | 25.469 | -1.251 | 22.483 |
| 9.973 | -9.727 | -16.524 | -23.607 | 5.632 | 3.970 | -15.891 | -24.771 | 24.102 | -7.423 | -21.119 | 11.479 | -31.047 | 20.823 |
| 9.439 | 26.376 | 24.426 | 17.653 | 27.541 | 21.177 | 6.887 | -20.590 | -2.195 | -8.159 | -17.524 | -27.343 | 17.603 | -12.597 |
| -5.667 | -21.856 | 3.307 | 18.431 | -15.573 | -19.342 | -17.157 | -15.050 | 3.576 | -6.281 | 27.667 | 20.160 | -3.301 | 23.600 |
| 8.009 | 27.234 | 27.904 | -11.459 | 11.752 | -27.435 | -4.291 | -4.533 | -19.872 | 20.173 | 25.760 | 2.712 | 30.039 | -2.564 |
| -3.532 | -1.535 | -9.372 | -25.320 | -12.034 | -18.249 | 20.425 | 19.788 | 10.978 | -1.929 | -24.011 | 21.134 | 29.177 | 17.759 |
| 21.316 | 19.228 | -7.058 | -22.252 | 30.514 | 17.990 | 21.430 | 5.913 | 2.873 | -19.356 | -27.302 | 7.265 | 25.851 | 9.431 |
| 21.036 | -5.855 | 20.114 | 11.365 | 2.652 | 11.525 | 27.770 | 29.210 | 1.539 | -21.531 | 18.847 | 22.180 | -2.905 | 15.324 |
| 15.459 | 9.419 | -16.776 | 15.619 | 23.772 | 9.651 | 24.910 | 20.355 | -9.338 | -26.959 | 23.781 | 21.857 | -22.350 | 20.581 |
| -4.213 | 25.881 | -10.100 | 5.840 | 15.427 | -26.704 | 6.413 | -26.193 | 15.960 | 2.097 | -28.413 | -5.204 | -12.138 | -11.090 |
| -11.882 | 0.578 | -21.514 | 10.628 | -7.121 | 18.477 | -13.998 | -8.122 | -6.210 | -16.602 | 14.128 | -10.416 | 0.344 | 6.240 |
| -18.856 | -14.216 | 19.938 | -19.515 | -9.717 | -8.957 | 5.592 | -5.343 | 5.745 | -20.428 | 2.189 | -24.248 | 1.755 | -5.129 |
| 8.961 | -22.316 | 16.439 | 26.496 | -22.580 | -24.887 | -29.076 | 14.689 | -22.146 | 3.744 | 16.015 | -25.480 | -24.259 | 21.915 |
| -14.421 | 3.755 | 4.947 | -14.770 | 9.351 | -5.141 | -19.312 | 25.849 | -27.395 | -1.354 | -18.355 | 24.123 | 13.336 | 14.065 |
| -7.956 | 12.232 | 18.477 | -11.854 | -10.549 | 20.792 | -18.724 | 13.193 | -4.438 | 9.689 | -24.046 | -12.446 | 12.761 | 21.377 |
| -20.236 | -0.828 | -19.255 | -17.039 | -26.349 | -22.581 | -12.243 | -29.786 | 20.603 | 7.013 | 2.818 | 1.142 | -7.278 | 9.487 |
| -26.122 | -18.538 | 6.982 | 23.864 | 19.245 | -14.577 | -18.002 | 5.444 | -5.828 | -19.374 | 20.630 | 24.869 | -1.022 | -24.114 |
| 20.570 | 8.509 | 29.685 | 24.516 | -10.246 | 12.449 | 28.253 | 27.659 | 22.252 | -25.431 | 11.766 | -22.301 | -16.117 | 13.465 |
| 19.783 | 6.266 | 5.659 | -0.923 | 20.837 | 2.860 | 17.351 | 4.528 | -13.796 | 9.854 | 0.292 | 29.465 | 21.377 | 23.027 |
| 25.893 | 18.423 | -2.644 | -21.249 | 7.447 | 19.340 | 12.632 | -21.794 | 15.192 | -28.021 | -22.829 | -28.965 | -12.039 | -15.431 |
| 3.337 | -6.194 | -19.544 | 2.405 | 15.529 | 22.314 | -14.428 | 23.711 | 13.033 | -9.643 | 25.932 | -8.995 | -20.427 | 18.664 |
| 24.300 | 1.682 | 16.275 | -4.985 | 4.361 | 20.901 | 27.444 | 24.633 | 0.277 | -4.555 | -21.183 | 27.131 | 1.879 | 16.571 |
| -18.043 | -14.682 | 24.050 | 9.098 | -1.368 | -6.343 | -25.147 | -8.659 | -2.234 | -16.229 | -29.546 | 2.470 | -9.898 | 24.641 |

| | | | | | | | | | | | | | |
|---|---|---|---|---|---|---|---|---|---|---|---|---|---|
| 11.013 | 23.180 | -13.344 | -14.422 | -17.677 | -16.545 | -27.664 | 2.160 | 2.499 | 14.499 | -0.670 | 17.900 | 1.237 | 27.854 |
| 6.350 | 11.710 | 19.588 | -24.051 | 4.095 | -25.972 | 18.286 | -6.483 | -26.772 | 26.067 | -13.356 | 24.595 | -7.859 | 21.517 |
| -13.116 | -7.863 | 25.007 | -8.045 | -14.460 | -17.960 | 28.879 | -29.322 | 6.314 | -18.355 | 26.703 | 22.629 | 19.032 | 15.116 |
| -23.528 | 23.903 | 22.254 | -8.646 | -16.913 | 6.234 | 23.594 | 13.003 | -10.803 | 5.142 | 22.301 | 21.671 | -22.774 | 5.687 |
| 7.631 | -6.587 | 17.367 | -7.369 | 27.659 | 19.591 | 4.161 | -12.166 | -3.674 | -5.176 | -22.201 | -21.816 | -24.878 | 11.232 |
| 2.314 | -16.457 | 23.289 | 24.196 | 4.817 | 10.332 | -0.564 | -22.858 | 0.631 | -10.041 | -0.089 | 29.674 | 0.107 | 23.254 |
| 21.148 | 14.344 | 10.791 | 13.811 | -1.996 | -16.699 | -23.671 | 29.838 | -5.325 | 18.200 | -26.916 | 14.488 | -15.349 | 20.607 |
| 25.484 | 8.410 | 20.443 | -0.490 | 3.292 | 7.346 | -13.953 | 28.484 | -1.594 | -14.616 | 23.217 | -19.678 | -2.060 | 19.814 |
| -21.753 | 4.109 | 13.876 | -21.820 | 4.648 | 24.227 | -26.095 | -13.498 | -0.399 | -9.453 | 8.999 | -16.198 | 24.374 | 11.804 |
| 23.832 | -22.342 | -1.955 | -22.860 | 28.755 | -6.120 | 0.423 | -14.482 | 8.138 | 27.402 | 10.806 | 14.043 | 4.112 | 23.820 |
| -26.180 | -1.527 | -16.766 | -7.547 | 9.293 | -7.374 | 7.545 | -17.565 | -15.873 | 0.302 | -17.464 | 25.750 | 16.231 | -2.928 |
| 18.533 | 22.233 | 27.629 | 1.116 | -20.041 | -10.570 | -26.559 | 1.093 | 16.930 | 28.041 | -11.730 | 20.506 | 2.671 | -15.109 |
| 6.603 | -5.223 | 18.297 | -18.451 | -14.108 | -24.543 | -24.897 | 17.604 | 1.565 | -19.281 | -27.976 | -29.587 | 23.237 | 12.263 |
| 25.760 | -24.802 | -17.758 | -24.545 | 14.735 | 21.725 | -9.185 | -10.954 | 25.973 | 18.178 | 21.271 | 16.571 | 21.744 | 12.878 |
| -5.492 | -2.449 | 29.152 | -15.900 | -10.014 | -20.214 | -0.144 | 22.102 | -26.089 | -18.235 | -0.097 | 6.598 | -29.945 | 7.803 |
| 22.289 | 26.403 | 12.291 | 23.211 | -7.635 | -13.992 | -12.487 | 1.311 | -12.517 | -3.630 | -3.065 | -13.759 | -28.248 | 18.029 |
| 14.833 | -3.623 | 2.736 | 27.555 | 8.404 | 20.981 | -29.056 | 8.589 | 10.944 | 8.879 | 5.293 | -17.514 | 26.258 | 19.528 |
| 8.675 | -19.293 | 19.466 | 9.493 | 7.477 | -8.763 | 28.781 | -1.684 | -8.123 | -15.819 | -24.893 | -8.073 | 18.054 | 29.584 |
| 5.804 | 27.760 | -13.095 | -16.323 | -25.030 | -11.955 | 21.047 | -5.361 | 6.905 | 18.659 | 24.845 | -14.277 | 20.678 | 28.079 |
| -2.952 | 27.523 | 13.000 | 11.064 | 6.490 | -17.484 | 1.758 | 3.539 | -1.088 | 17.565 | 7.538 | 11.564 | 23.936 | 21.264 |
| 13.075 | 7.144 | -27.304 | -20.533 | 23.300 | 19.091 | -2.286 | 30.098 | 19.280 | 0.049 | 10.602 | -28.007 | -2.680 | 2.254 |
| -10.383 | 19.007 | 17.982 | 4.015 | 4.686 | 0.458 | 25.502 | -8.483 | 1.576 | 5.347 | -10.817 | 17.287 | -20.015 | 27.390 |
| 20.617 | 21.035 | -14.555 | -17.368 | 24.279 | -2.443 | -15.293 | 27.177 | 8.253 | -7.617 | 23.988 | -18.537 | 16.642 | 13.166 |
| 21.585 | -18.524 | -6.166 | 1.547 | 2.644 | 14.685 | 19.989 | 24.392 | -13.750 | -11.967 | 22.061 | 14.764 | 8.581 | 14.447 |
| -24.615 | 24.813 | 15.259 | -27.100 | 25.002 | 25.058 | -30.072 | -2.431 | -4.912 | 23.058 | 16.334 | 22.994 | -1.094 | -16.956 |
| -7.419 | 23.516 | 14.549 | -4.624 | 28.083 | -0.604 | -29.585 | -22.413 | -5.868 | -18.982 | -6.450 | 24.328 | -21.109 | 24.208 |
| 15.532 | 27.285 | -19.710 | 7.392 | -19.235 | 21.756 | 25.713 | 5.069 | 18.296 | -22.990 | 27.049 | -28.505 | 13.810 | 20.117 |
| 0.982 | 4.473 | -24.363 | -3.343 | -7.416 | 3.377 | 18.229 | -8.783 | 24.744 | -3.589 | -23.290 | 23.355 | 23.353 | -28.592 |
| 14.150 | 22.059 | -8.314 | -24.134 | -30.315 | 9.573 | 30.013 | 14.864 | -16.805 | 3.441 | 0.817 | -24.797 | 25.241 | 22.918 |
| -22.427 | 14.333 | -22.362 | 11.265 | -6.045 | -10.406 | -28.292 | 20.998 | 20.951 | 3.003 | 9.683 | -30.536 | 7.238 | 16.078 |
| 18.651 | -6.167 | 23.121 | 23.808 | -26.680 | -26.202 | 2.536 | 9.563 | -6.653 | 17.971 | 27.497 | -14.327 | -6.537 | -6.641 |
| 23.577 | -2.095 | -13.774 | -12.783 | -4.283 | -19.394 | -19.022 | -17.433 | -16.211 | -20.312 | -24.937 | 26.662 | -23.024 | 13.206 |
| -17.549 | 5.596 | -9.572 | 6.021 | 20.240 | -15.202 | 21.926 | -31.425 | -25.788 | 6.786 | 19.230 | -19.894 | 22.724 | -22.921 |
| -3.471 | -4.334 | 28.938 | -21.250 | 10.880 | 17.889 | 29.118 | 28.236 | 9.276 | 4.200 | -23.855 | 25.307 | 20.291 | 19.163 |
| -27.560 | 19.972 | -10.608 | -7.850 | 28.091 | 0.359 | -2.881 | 20.324 | 14.771 | -12.721 | -14.179 | -22.560 | -18.204 | 1.985 |
| -26.035 | -3.861 | 29.353 | -26.942 | -30.428 | -9.605 | -24.964 | -24.382 | 6.060 | 20.318 | -23.091 | 17.796 | -23.382 | 3.207 |

| | | | | | | | | | | | | | |
|---|---|---|---|---|---|---|---|---|---|---|---|---|---|
| 9.494 | 23.445 | 27.394 | 19.113 | -27.013 | -5.480 | 10.918 | 10.979 | 4.328 | 17.128 | 12.022 | 10.834 | 20.587 | 11.259 |
| -18.941 | -20.206 | 13.886 | -6.815 | -7.366 | -7.010 | 13.112 | -6.520 | -11.990 | -12.415 | 2.341 | -7.771 | 29.274 | 31.088 |
| 19.834 | 9.461 | 1.373 | 2.601 | 13.095 | -0.891 | -29.231 | -12.704 | 12.427 | 15.699 | 26.761 | 27.389 | 1.473 | 20.628 |
| -26.354 | -12.673 | -5.547 | -2.026 | 5.274 | -25.224 | 25.516 | -7.074 | -12.763 | 1.585 | 9.483 | 8.320 | 27.121 | 14.114 |
| 22.459 | 17.889 | -24.976 | -27.932 | 14.763 | -6.043 | -15.801 | -20.542 | 3.600 | -30.685 | 19.996 | 23.914 | 15.299 | 26.256 |
| -15.578 | -28.133 | 23.111 | 13.974 | -10.652 | 16.807 | -31.058 | -22.075 | -2.675 | 14.741 | -16.821 | 21.560 | 0.144 | 24.902 |
| 6.184 | 22.052 | 4.438 | -13.896 | -26.226 | 8.884 | 12.571 | 0.140 | 13.325 | 3.204 | 1.372 | 21.033 | -0.930 | 27.741 |
| -21.570 | -8.343 | 21.235 | 8.744 | 24.787 | 24.394 | -3.823 | -11.771 | 17.911 | -6.799 | 7.305 | 20.089 | -2.399 | 14.619 |
| 16.968 | -26.300 | -0.787 | -21.989 | 20.239 | 21.139 | 22.308 | -32.480 | 24.165 | -5.610 | -7.970 | -27.309 | 26.140 | 1.587 |
| -27.577 | -5.729 | 20.774 | 6.849 | 7.166 | 5.282 | -3.378 | 22.001 | 21.961 | 25.014 | 28.573 | -17.205 | 22.664 | 25.805 |
| 14.024 | 16.650 | 0.569 | 29.092 | 30.650 | 25.950 | -26.547 | 18.681 | 15.605 | -22.261 | 18.724 | -11.225 | -18.849 | 17.278 |
| -1.779 | 0.404 | 25.720 | -4.345 | -18.043 | -22.459 | 26.021 | 2.080 | -13.559 | 10.612 | 29.616 | -13.885 | 20.667 | 19.856 |
| 10.318 | -19.994 | -23.181 | 15.918 | 8.559 | 25.046 | 25.013 | -12.485 | 13.949 | -1.512 | 4.947 | 0.387 | -25.296 | 30.484 |
| -5.444 | -0.503 | -13.436 | 0.711 | 23.254 | 19.638 | -30.507 | -13.681 | 4.427 | -10.720 | 1.805 | 15.233 | 3.358 | 11.152 |
| -18.838 | -17.944 | -17.206 | 17.584 | -19.900 | 10.791 | 19.749 | -31.753 | 10.223 | 28.897 | 22.240 | 2.444 | 1.633 | 17.949 |
| 20.619 | 27.712 | 19.475 | 26.465 | 8.300 | 23.560 | 22.816 | -6.721 | -3.854 | -8.783 | 13.050 | -6.981 | -29.514 | 22.327 |
| -11.921 | 19.651 | -2.393 | 22.052 | -23.563 | -6.738 | 3.963 | -10.393 | -9.616 | -14.724 | 24.243 | -4.524 | -3.887 | 8.767 |
| -10.745 | 2.334 | 27.653 | -0.810 | -9.005 | 19.040 | -15.267 | 6.488 | 7.942 | 8.066 | -27.163 | 30.000 | 9.223 | 21.308 |
| 27.838 | -3.012 | -21.169 | -14.930 | 7.209 | -17.213 | 5.579 | 29.624 | 3.293 | -1.758 | -12.488 | 23.963 | 18.690 | 15.241 |
| -26.005 | -0.105 | 24.539 | 6.046 | 8.607 | -25.919 | 29.358 | 23.896 | 2.695 | -6.891 | -9.814 | 14.780 | 17.391 | 0.388 |
| -16.562 | 19.930 | -1.468 | -28.053 | -22.618 | 13.382 | -30.901 | -10.997 | -0.506 | -15.541 | -23.444 | 4.567 | -0.539 | 23.390 |
| 25.365 | -5.207 | 28.321 | -7.908 | 7.786 | -12.740 | 10.163 | -27.173 | -1.003 | -17.126 | 26.184 | -14.059 | -0.041 | -17.545 |
| -21.530 | -10.439 | -25.375 | 15.474 | -15.898 | -17.607 | 6.858 | 0.412 | 4.905 | -21.113 | -26.781 | 19.432 | 27.512 | 25.595 |
| 8.978 | -22.872 | -4.954 | 1.839 | -8.343 | -10.294 | -31.490 | -21.218 | -2.162 | 5.558 | 17.184 | -26.463 | 15.938 | 14.398 |
| 21.930 | -29.544 | -13.279 | 14.266 | 18.938 | -13.260 | 31.475 | 3.344 | 13.498 | -24.433 | -11.692 | 9.718 | 23.995 | 21.515 |
| -7.060 | -11.656 | 20.569 | -16.310 | 12.005 | -25.673 | 20.879 | -5.044 | 12.317 | -18.218 | -14.534 | -24.479 | 7.031 | -3.081 |
| -27.623 | 27.184 | 25.138 | 23.633 | 22.043 | 15.578 | 19.614 | 13.069 | -15.814 | 28.482 | -5.677 | -31.334 | 23.496 | -29.067 |
| 25.236 | 5.077 | -22.798 | 18.545 | -0.720 | -21.885 | 21.733 | -18.948 | -18.565 | -27.477 | 27.922 | 28.174 | 20.322 | 19.193 |
| -3.089 | -19.730 | 7.117 | -25.232 | -22.409 | 4.853 | -6.961 | 18.334 | -1.547 | 0.263 | 22.318 | 26.265 | -20.719 | 17.135 |
| -9.267 | -0.927 | 13.532 | -29.244 | -6.095 | 19.657 | 3.639 | -1.066 | 9.496 | 24.187 | 11.775 | 11.108 | 21.028 | 18.494 |
| -2.255 | 23.552 | 11.516 | 21.456 | 6.599 | -20.958 | -6.439 | -16.795 | 5.024 | -20.150 | 17.822 | 30.796 | 10.549 | -25.841 |
| -16.174 | 27.527 | 10.305 | -14.683 | 29.756 | 25.276 | -21.279 | 16.738 | 20.876 | -13.076 | 29.557 | 11.733 | 9.916 | -29.903 |
| -20.248 | 21.454 | 26.862 | 10.481 | 4.171 | 21.295 | 28.655 | 7.774 | 0.249 | -5.862 | 28.710 | 3.379 | -9.026 | 28.635 |
| -3.872 | 9.199 | -26.367 | -7.804 | -4.475 | 24.420 | -7.497 | -12.075 | 22.671 | -18.412 | -18.871 | 27.485 | 18.849 | 27.327 |
| 18.864 | 6.144 | 22.465 | -8.737 | 26.062 | -3.344 | -17.415 | -14.249 | -5.537 | 3.066 | 26.529 | 17.106 | -28.398 | -6.153 |
| 21.381 | 3.932 | 4.729 | 3.556 | -2.669 | 17.189 | -30.332 | -23.906 | -7.615 | -9.933 | -25.462 | 12.649 | 21.965 | 30.883 |

| | | | | | | | | | | | | | |
|---|---|---|---|---|---|---|---|---|---|---|---|---|---|
| -20.372 | 13.857 | 6.306 | 1.796 | 24.462 | -10.004 | 21.515 | -16.416 | -14.536 | -4.824 | -10.337 | -29.667 | -21.921 | 16.289 |
| -27.198 | -1.437 | 17.100 | -2.981 | 9.211 | 22.727 | 6.629 | 11.882 | -2.658 | -21.354 | -2.185 | 26.344 | 25.110 | -8.046 |
| -18.883 | 6.264 | 19.276 | -26.764 | 3.801 | -6.012 | -17.998 | 20.542 | 11.159 | 20.828 | 23.184 | 24.325 | 25.548 | 20.855 |
| -24.667 | -8.834 | -4.924 | -13.198 | -21.578 | 8.066 | -19.182 | -21.080 | 5.020 | 8.129 | 27.231 | -30.687 | -0.141 | -2.754 |
| -15.384 | -17.170 | -5.907 | 3.207 | 24.921 | 6.364 | 5.900 | -30.178 | -19.444 | -10.403 | 25.245 | 27.024 | -5.797 | -6.678 |
| 28.265 | 11.764 | -9.851 | -21.376 | 1.750 | -2.445 | 16.995 | -27.734 | -23.179 | -16.681 | -23.307 | -5.252 | -3.291 | 28.408 |
| -8.460 | 1.157 | 24.632 | 13.527 | -8.256 | -10.437 | -1.757 | 9.696 | -7.134 | 24.928 | 23.889 | 18.789 | -0.948 | 25.555 |
| -26.519 | 16.558 | -15.414 | -18.663 | 3.507 | -1.063 | -29.540 | -10.171 | -12.709 | 15.946 | -6.120 | 30.337 | -16.205 | 12.185 |
| -0.514 | 24.863 | -24.997 | -0.224 | 10.073 | -21.709 | 18.341 | -8.357 | 7.122 | -27.165 | 22.934 | 11.196 | -18.102 | 15.289 |
| -15.820 | -6.337 | -0.490 | -25.314 | 27.883 | -15.328 | -8.798 | 16.788 | -0.912 | -14.769 | -9.995 | -11.953 | 22.789 | 16.383 |
| -10.734 | -28.063 | -24.401 | 28.920 | -24.371 | 23.382 | 3.290 | 3.974 | -1.898 | 17.250 | 14.169 | -21.767 | 2.021 | 24.683 |
| 22.916 | 18.274 | -5.089 | 7.777 | -7.762 | -6.853 | -8.358 | 29.348 | 2.995 | -13.736 | 14.927 | 6.643 | -1.659 | 20.444 |
| 20.533 | 29.567 | -25.395 | 11.455 | -22.390 | -21.255 | 0.375 | 21.791 | -13.566 | 6.997 | -17.412 | 28.550 | -25.875 | 29.902 |
| -5.006 | 20.439 | -1.456 | -7.351 | 3.491 | 14.679 | 20.845 | 17.216 | 2.161 | -20.981 | 26.363 | 18.330 | 15.988 | 18.180 |
| -21.820 | 4.924 | -19.597 | 10.443 | -28.497 | -1.500 | -6.759 | -1.885 | 7.752 | -7.811 | -4.521 | 19.953 | 5.700 | 13.412 |
| -13.668 | 8.066 | -2.474 | -0.497 | 25.873 | -16.133 | -28.998 | -22.289 | -20.064 | -11.053 | -12.076 | 17.919 | -27.596 | -1.533 |
| 27.736 | 8.738 | -26.954 | 5.585 | 22.476 | 10.046 | -31.415 | 25.301 | -24.583 | -16.792 | -26.695 | -28.160 | -2.545 | 14.862 |
| 25.471 | 23.903 | 27.140 | 4.111 | 4.286 | 15.490 | -23.785 | 1.604 | 7.672 | 29.047 | 26.885 | 3.243 | -24.627 | 21.419 |
| -1.511 | 5.264 | 22.019 | 20.431 | -3.073 | -4.174 | -24.355 | 27.857 | 23.840 | 7.976 | -1.842 | 25.523 | 26.317 | -6.032 |
| 19.412 | -10.501 | 9.869 | -26.801 | 3.842 | 0.141 | 28.868 | 26.643 | 4.754 | -20.423 | -20.956 | -3.372 | 20.293 | 19.779 |
| 15.411 | -8.220 | 26.103 | -2.609 | -26.768 | -5.604 | 25.919 | 1.248 | 5.410 | -26.321 | 9.643 | 7.849 | -4.478 | -9.806 |
| 12.748 | 6.949 | 15.649 | -15.624 | 7.770 | -24.502 | 26.370 | 8.672 | 9.538 | -0.507 | -24.077 | 27.012 | -3.387 | 24.174 |
| 4.005 | -0.436 | 23.116 | 8.522 | -3.616 | -20.703 | 15.128 | 21.971 | 16.531 | 23.731 | 25.826 | -16.336 | 4.068 | 21.611 |
| -20.629 | -29.434 | 14.082 | 18.839 | -1.169 | -11.481 | 19.937 | -21.091 | 2.608 | 17.682 | 7.537 | 22.010 | 24.789 | 20.978 |
| 8.854 | 10.833 | 6.124 | -28.974 | -5.778 | 4.682 | 22.076 | -3.659 | -17.876 | -2.702 | -13.132 | 29.039 | -25.530 | 19.437 |
| 17.951 | -26.454 | 12.785 | 0.073 | 28.501 | 8.274 | 18.889 | -8.050 | -6.747 | -15.238 | 17.614 | 23.338 | -5.531 | 14.858 |
| 9.961 | 22.869 | -1.309 | 14.455 | -23.821 | 17.895 | 3.185 | -19.607 | 12.232 | 22.127 | 0.762 | 1.573 | 7.604 | 24.793 |
| -7.577 | -15.774 | 11.303 | 18.343 | 27.517 | -7.187 | -4.561 | 13.484 | 1.581 | 19.070 | -21.863 | -29.524 | -6.343 | 16.477 |
| 23.778 | 2.412 | 1.686 | -21.224 | 6.395 | -8.561 | -26.259 | 3.706 | -19.010 | -25.636 | 16.700 | 27.900 | 20.688 | 10.372 |
| -19.961 | -18.030 | -25.729 | 11.503 | 7.926 | 1.120 | 22.337 | -26.827 | 25.302 | -19.700 | -9.005 | -22.726 | 1.290 | -9.257 |
| 2.516 | -7.640 | 22.106 | 24.596 | -26.819 | 14.157 | 24.135 | 1.096 | -16.705 | 24.974 | 7.079 | 10.427 | 21.333 | 26.552 |
| -15.348 | -23.934 | -5.345 | 29.481 | -5.470 | -2.323 | 8.665 | -4.426 | 5.513 | 6.853 | -17.704 | -17.875 | -8.250 | -21.263 |
| 25.979 | -7.276 | -21.572 | -15.024 | 8.514 | 24.776 | -20.156 | -6.383 | -22.688 | -14.512 | -22.130 | 26.586 | 18.542 | 16.983 |
| -8.106 | -22.803 | 24.667 | 19.231 | 14.107 | 4.036 | 28.682 | -8.915 | 11.431 | -21.976 | -24.920 | 25.027 | -27.554 | 19.623 |
| -9.571 | -10.299 | -12.445 | -2.513 | 17.366 | 9.707 | 28.503 | -28.216 | -11.231 | 27.811 | -10.423 | 22.892 | 19.389 | 7.587 |
| 22.901 | -12.565 | -23.136 | 6.334 | 25.099 | -11.210 | 6.164 | -20.811 | -2.476 | -11.271 | 21.098 | 25.829 | 16.045 | -16.703 |

| | | | | | | | | | | | | | |
|---|---|---|---|---|---|---|---|---|---|---|---|---|---|
| 1.839 | -18.598 | 15.358 | -18.538 | -5.053 | 21.281 | 18.147 | -1.472 | 22.105 | 29.618 | -23.255 | -10.701 | 12.293 | -16.989 |
| -28.023 | -5.201 | -20.606 | -24.084 | 9.883 | 23.112 | -8.730 | 23.933 | 18.920 | 30.507 | 23.918 | -13.104 | -15.448 | 20.784 |
| -14.497 | -15.669 | -13.781 | 2.463 | 1.390 | 16.291 | 21.330 | -22.712 | 1.030 | 12.170 | 19.834 | -30.049 | 12.704 | 22.176 |
| -29.636 | 25.561 | -11.121 | -16.822 | -26.314 | 13.238 | 4.118 | 5.205 | -4.215 | 11.294 | 14.041 | 28.116 | 19.076 | 25.993 |
| 27.971 | -28.873 | -19.410 | 5.545 | -4.748 | 10.518 | 3.046 | -16.372 | -24.777 | 9.976 | -9.641 | -26.263 | -1.182 | 23.235 |
| 24.416 | 22.067 | 23.412 | -12.692 | -22.111 | -2.150 | 23.658 | 4.941 | -8.811 | -16.176 | -21.857 | 9.420 | 24.777 | -10.952 |
| -8.728 | 29.428 | -24.948 | 25.074 | 3.342 | -4.541 | 5.432 | -5.545 | 22.793 | 19.733 | 26.327 | 29.186 | -4.260 | 26.380 |
| 30.541 | 9.118 | -26.479 | -9.963 | 9.695 | -17.648 | 7.382 | 18.799 | 17.101 | 23.575 | -6.712 | -29.396 | 0.498 | 18.770 |
| -11.112 | 21.436 | -12.056 | 1.207 | -24.543 | 8.873 | -3.185 | -12.429 | -0.892 | 28.126 | 14.990 | 12.356 | -27.009 | 6.011 |
| -3.926 | -4.084 | -15.981 | 18.327 | -1.370 | 19.102 | -24.205 | 21.308 | 21.615 | 23.056 | 10.675 | 8.395 | 16.082 | -10.481 |
| -4.966 | -9.100 | 21.002 | -5.811 | 17.724 | 20.360 | -1.710 | 4.626 | -3.706 | -4.180 | 17.673 | -1.933 | -7.517 | 27.296 |
| 16.294 | 5.936 | 6.973 | -28.598 | 13.083 | 17.640 | -4.359 | -4.817 | -6.240 | 8.318 | 23.923 | 14.997 | -2.539 | 22.581 |
| -29.076 | -12.183 | 17.760 | 25.845 | -3.555 | 12.538 | -18.456 | -22.655 | 6.767 | 1.749 | -21.035 | 23.915 | -25.139 | 24.637 |
| 19.630 | 23.996 | -8.663 | 0.133 | 25.908 | -3.321 | 23.265 | 20.123 | -10.422 | 12.687 | -12.764 | -10.226 | 25.523 | 17.701 |
| 21.380 | 12.620 | -19.314 | -2.907 | -13.180 | -11.359 | -17.517 | 26.700 | -0.200 | -3.988 | 13.861 | 15.656 | 22.696 | 14.965 |
| 26.911 | 7.549 | 1.553 | -8.802 | 23.947 | 7.871 | -21.368 | -9.625 | 9.785 | 1.265 | -10.216 | 17.229 | -23.211 | 28.431 |
| 22.294 | -14.721 | -11.769 | 25.290 | -7.876 | 23.955 | -19.990 | 16.144 | 5.601 | -24.781 | 24.891 | 22.124 | 4.731 | -23.379 |
| -21.947 | -30.949 | -26.138 | 14.752 | -16.354 | -6.158 | 17.911 | -4.240 | 10.669 | 7.313 | -8.833 | 25.246 | -23.493 | 18.294 |
| -30.688 | -24.775 | 21.766 | -20.489 | -6.739 | 2.354 | -19.457 | -6.199 | 8.268 | 27.647 | -21.151 | -11.539 | 5.836 | 6.951 |
| -23.802 | -2.445 | -22.802 | 23.667 | 13.817 | 9.572 | 26.293 | 19.985 | 1.594 | 21.624 | 16.217 | 10.396 | -27.980 | 18.940 |
| 9.374 | 30.493 | 5.108 | -1.215 | -2.581 | -7.816 | 27.867 | -25.829 | -0.862 | -18.414 | -14.891 | 1.132 | 12.331 | 14.409 |
| -7.787 | -3.815 | 12.617 | 27.320 | 2.628 | 18.453 | 8.303 | 9.786 | 11.081 | -17.782 | 23.311 | -6.113 | -0.808 | 22.169 |
| -16.669 | 2.866 | 8.494 | -29.688 | 3.948 | 1.581 | -20.713 | -7.385 | 7.458 | -19.485 | 9.535 | 26.066 | -22.167 | 21.788 |
| -26.368 | -11.309 | 16.816 | 26.127 | -9.288 | -20.801 | -30.868 | -0.242 | 12.184 | -26.143 | 12.372 | 27.546 | 24.023 | 7.858 |
| -2.598 | 24.406 | 9.577 | -26.625 | -13.713 | -25.103 | -2.673 | 15.533 | -17.196 | 7.730 | -8.120 | 17.855 | -8.296 | 10.067 |
| -18.703 | -22.098 | -0.941 | 9.908 | 4.779 | 13.599 | 19.000 | -8.914 | -12.740 | -6.332 | -11.115 | 13.369 | 25.663 | 8.515 |
| 6.605 | 21.830 | 22.919 | 19.919 | 1.706 | 25.433 | 4.717 | -2.382 | -6.766 | -14.509 | -0.490 | 4.287 | 5.524 | 23.038 |
| -7.417 | -25.397 | 24.813 | 3.773 | 9.012 | 12.510 | 23.758 | -9.560 | 0.622 | -18.327 | 12.576 | 27.022 | -26.808 | -21.607 |
| 8.155 | -6.064 | -16.573 | 14.924 | -0.515 | 24.779 | 18.379 | 26.823 | -7.871 | -23.733 | 8.615 | -7.353 | 23.461 | 13.360 |
| 24.602 | -0.770 | 13.925 | 1.154 | -22.173 | -16.683 | -15.512 | -8.776 | -1.794 | 12.115 | 22.952 | -21.899 | 6.642 | 19.954 |
| 10.616 | 16.873 | 3.783 | -1.723 | -3.031 | 23.773 | -28.337 | -25.596 | 7.071 | -8.313 | 19.740 | -27.332 | 2.631 | 0.194 |
| -27.146 | -13.142 | 9.566 | 21.709 | -12.960 | 6.642 | 21.785 | 25.900 | -20.077 | 5.765 | -4.823 | -12.324 | 9.366 | 9.046 |
| 17.589 | 24.948 | 21.827 | 15.679 | 11.555 | 20.844 | 28.253 | -4.121 | 21.926 | 19.106 | -2.986 | -17.781 | -1.686 | 28.286 |
| 29.166 | 26.145 | -3.434 | 10.759 | -26.449 | 15.288 | -7.608 | -17.171 | 13.675 | -16.787 | 6.306 | 27.916 | 22.688 | 23.024 |
| 14.594 | 13.653 | -6.590 | 29.293 | 22.366 | 19.507 | -0.523 | 23.945 | -3.577 | 3.397 | 16.284 | -8.219 | 14.560 | 21.070 |
| -21.378 | -26.909 | -8.945 | 8.109 | -4.916 | 15.498 | -7.090 | -27.682 | 5.443 | -7.473 | 6.809 | -24.438 | 10.289 | 12.370 |

| | | | | | | | | | | | | | |
|---|---|---|---|---|---|---|---|---|---|---|---|---|---|
| 21.706 | 21.754 | 4.133 | -6.596 | -10.411 | 7.849 | -12.764 | -13.544 | 4.817 | 28.657 | 24.175 | 28.521 | -5.748 | -5.681 |
| -24.019 | -8.575 | -1.077 | 6.248 | 25.466 | -26.582 | -26.646 | -22.634 | -16.133 | -3.534 | 15.970 | 29.393 | 24.853 | 21.392 |
| -5.477 | 15.051 | -1.522 | -18.765 | 17.858 | 27.168 | -5.667 | -3.324 | 8.381 | -18.918 | -13.876 | 23.026 | 24.518 | -7.662 |
| 22.106 | -20.497 | -23.946 | -3.207 | -6.004 | 13.780 | -29.149 | 22.766 | 19.978 | 6.162 | -20.191 | 24.753 | 1.144 | 1.573 |
| -6.762 | -3.318 | 22.205 | -29.745 | -6.806 | -2.106 | 22.806 | -0.866 | -0.659 | 12.095 | -24.808 | 28.824 | 26.992 | 26.373 |
| 0.252 | -27.490 | -16.775 | -24.750 | -1.123 | 26.380 | -18.698 | -4.460 | 17.501 | -24.601 | -3.810 | 12.369 | 5.064 | -21.829 |
| -4.528 | -7.397 | 24.243 | -25.541 | 3.638 | 18.759 | 16.602 | -11.102 | -5.141 | 11.414 | -8.754 | -4.708 | -5.911 | 26.825 |
| -2.985 | 17.153 | -22.956 | 16.546 | 20.151 | 10.016 | 25.671 | 8.349 | -9.055 | -10.363 | 3.339 | -5.553 | 6.444 | 21.600 |
| 28.486 | 14.060 | -21.429 | 0.680 | 14.645 | 24.450 | -21.921 | -6.698 | 19.027 | 4.638 | -21.416 | -27.274 | -4.936 | -2.981 |
| -19.050 | 18.069 | 3.065 | 1.514 | 15.342 | -25.567 | -22.204 | 10.567 | 1.713 | -15.773 | 13.300 | 24.245 | 13.865 | -7.378 |
| 4.023 | 30.978 | -0.790 | -2.683 | -12.832 | 7.054 | 5.072 | -7.344 | -4.574 | -14.191 | -12.552 | 22.549 | -23.649 | -23.983 |
| -1.664 | 4.560 | 18.159 | -13.549 | -7.935 | -12.076 | -29.839 | -6.264 | -10.561 | 26.844 | -9.113 | 21.717 | -0.257 | 12.706 |
| 0.527 | 15.984 | 21.571 | -26.814 | 5.340 | 16.005 | -14.821 | -2.718 | 20.369 | -9.787 | -17.881 | 26.569 | -23.131 | 27.497 |
| 12.799 | 20.029 | 22.624 | 4.019 | 12.187 | 3.539 | 20.915 | 13.291 | -12.496 | 27.682 | -26.068 | 25.850 | 2.413 | -3.842 |
| 16.149 | -26.038 | -1.574 | 24.075 | 21.395 | 21.573 | 18.563 | -24.077 | -11.745 | 20.653 | 13.788 | 1.865 | 3.079 | 25.420 |
| 17.479 | 17.835 | 13.000 | -2.137 | 15.414 | 25.484 | 9.034 | -2.416 | 9.275 | -1.438 | 22.103 | -7.648 | 14.196 | 1.343 |
| 24.938 | 30.245 | 0.637 | 12.159 | 8.188 | 17.301 | 25.355 | -14.581 | 14.777 | 9.306 | 19.527 | 0.995 | 6.402 | 4.006 |
| 17.808 | 31.491 | 16.000 | 6.023 | -13.001 | -25.882 | 7.810 | 2.691 | 22.832 | -15.314 | 3.300 | 16.320 | -7.630 | -3.615 |
| 28.854 | 13.263 | 15.162 | -7.345 | 23.986 | -8.708 | -10.448 | -27.148 | -1.630 | 27.956 | 16.766 | -18.452 | 13.589 | -6.169 |
| 2.350 | 28.677 | -17.718 | 21.021 | 18.344 | -24.003 | 16.245 | -15.847 | -2.222 | -20.262 | -8.468 | -18.831 | -3.651 | 26.421 |
| -25.553 | 24.575 | 0.734 | -15.353 | -22.127 | 6.517 | -20.364 | -20.411 | 18.297 | -16.218 | -6.599 | 18.850 | -26.365 | 26.780 |
| -27.868 | -29.598 | -14.298 | 16.732 | 25.426 | -17.236 | -0.405 | -0.055 | -2.845 | -7.609 | 6.067 | -6.636 | -6.256 | 1.851 |
| -28.971 | 16.363 | -21.083 | 3.599 | 24.464 | -11.131 | 5.492 | 15.818 | 24.244 | 7.715 | 15.256 | 28.140 | -12.476 | 22.232 |
| 19.103 | 23.834 | 12.092 | -1.365 | 7.506 | 11.947 | 10.147 | 12.603 | -8.567 | -12.955 | 7.093 | 11.137 | -26.025 | 16.048 |
| 10.839 | 4.022 | -11.323 | -26.050 | -9.023 | -19.496 | 29.315 | -0.331 | 24.873 | 24.299 | 21.506 | 23.152 | -22.679 | -26.737 |
| -11.903 | -26.347 | 17.877 | -2.526 | -4.741 | 10.504 | 3.748 | -4.183 | -26.453 | -23.014 | 12.864 | -22.164 | 17.580 | -16.473 |
| -1.048 | -11.014 | -18.985 | -18.088 | -13.367 | -22.481 | 0.347 | -5.177 | 26.406 | 5.381 | 24.997 | 21.367 | 6.797 | 11.414 |
| -22.718 | -18.471 | -22.276 | -6.874 | 22.570 | 14.290 | -25.363 | -13.750 | -12.443 | -9.414 | 14.765 | -10.883 | -24.809 | 20.668 |
| -24.363 | 19.712 | -16.535 | -3.942 | -16.855 | -23.127 | 4.256 | -7.591 | -1.075 | 3.095 | 5.051 | 27.384 | 25.204 | 3.426 |
| 12.160 | 29.868 | -25.445 | -25.496 | -5.719 | 6.154 | 5.847 | 22.594 | -11.304 | 3.607 | -11.525 | 14.709 | -6.988 | -6.756 |
| 27.540 | 17.339 | -12.233 | 12.946 | 8.849 | -21.642 | 28.022 | 2.764 | 23.578 | -14.392 | -3.757 | 17.624 | -19.703 | -4.223 |
| 11.805 | 27.805 | -12.731 | -24.320 | -0.356 | -2.330 | 7.395 | -25.282 | 11.726 | -19.178 | 15.861 | 0.968 | -2.844 | 2.362 |
| 21.435 | -9.529 | -13.423 | -2.562 | 4.447 | -2.779 | -4.409 | -23.181 | -5.544 | -18.404 | -26.347 | 19.053 | 0.601 | -0.093 |
| 23.989 | 3.156 | 15.428 | 26.352 | 21.178 | 9.897 | -28.030 | -11.731 | -10.012 | 7.214 | 7.414 | 2.725 | -7.391 | 24.267 |
| 15.593 | 14.300 | 25.104 | -20.905 | 5.369 | -16.269 | -6.606 | 0.526 | 12.089 | -13.631 | 16.589 | 15.471 | 26.021 | 23.103 |
| -7.011 | 13.711 | -6.278 | 2.119 | 3.857 | -26.441 | -21.980 | 4.030 | -21.611 | -4.450 | -20.518 | 21.883 | -12.736 | 9.912 |

| | | | | | | | | | | | | | |
|---|---|---|---|---|---|---|---|---|---|---|---|---|---|
| 22.128 | -20.916 | -24.946 | 3.063 | -2.885 | 19.228 | -29.432 | -4.854 | 1.637 | -5.701 | -14.340 | 20.241 | -2.084 | 14.286 |
| -18.577 | -21.490 | -23.043 | 12.209 | 11.534 | -26.814 | -28.563 | -15.279 | -0.559 | -28.548 | -9.116 | -6.589 | 4.131 | 24.830 |
| -21.711 | 30.684 | 18.761 | -7.851 | 10.094 | 8.670 | 21.719 | -7.235 | -21.212 | 22.942 | 18.455 | -7.645 | -15.505 | 17.567 |
| -18.985 | -13.772 | 16.084 | -24.619 | 14.067 | -23.288 | -2.199 | 1.576 | 8.910 | -19.604 | -1.638 | 24.066 | -0.759 | 25.726 |
| -22.793 | 5.401 | -22.136 | -23.769 | 2.802 | 22.636 | -23.125 | 10.251 | -4.597 | -26.512 | -3.833 | 10.568 | 21.839 | -0.155 |
| 9.010 | 12.999 | -14.893 | -28.297 | -15.830 | -15.282 | 19.001 | -3.413 | -4.067 | -0.195 | 7.639 | 5.291 | 17.719 | 7.567 |
| 23.210 | 12.556 | -24.637 | 29.129 | -4.222 | 26.697 | -23.493 | -16.641 | 2.959 | -6.812 | -14.930 | 21.172 | 3.402 | 19.646 |
| -14.477 | -28.676 | -26.840 | -20.465 | 15.380 | 21.867 | 25.849 | -16.364 | -27.815 | 16.001 | -21.501 | -18.365 | -10.838 | 15.896 |
| -6.942 | 15.415 | 22.526 | 4.484 | -1.552 | 17.779 | -26.322 | 24.984 | 15.692 | 26.859 | 26.882 | 25.193 | 8.409 | 21.286 |
| -13.126 | 29.994 | -19.735 | -16.439 | -14.213 | 9.427 | 4.064 | 11.157 | 4.531 | -2.752 | 21.595 | 25.919 | 1.674 | 13.462 |
| -19.097 | -23.571 | -17.311 | 25.619 | 20.658 | -10.865 | 23.394 | 24.291 | -6.688 | 5.624 | 0.301 | 24.200 | 13.802 | 23.059 |
| 19.551 | 5.034 | -25.598 | 5.565 | 3.057 | 16.260 | 11.527 | 6.252 | 16.409 | 24.307 | 26.063 | -1.996 | 6.968 | 16.081 |
| -14.341 | 3.575 | 13.634 | 0.861 | -12.137 | 12.833 | 2.460 | 1.150 | -23.684 | 1.137 | 17.072 | 14.706 | 22.707 | 2.920 |
| -24.796 | 13.150 | -23.320 | -7.914 | 25.509 | 22.228 | 10.493 | 25.031 | 2.453 | -21.219 | 23.919 | 22.287 | -2.182 | -1.076 |
| -6.698 | 19.100 | -19.532 | -25.543 | -25.059 | 1.813 | -6.888 | -0.264 | 0.782 | -3.268 | 21.297 | -26.152 | -8.812 | -11.266 |
| 29.500 | 17.074 | 22.820 | -24.608 | 7.993 | -3.006 | 11.033 | 12.204 | 1.900 | -26.490 | 8.401 | -21.204 | -10.602 | -24.754 |
| 28.091 | 11.932 | 20.147 | -4.799 | 13.475 | 26.390 | 25.383 | -4.523 | 25.648 | -22.661 | -15.044 | 26.616 | -14.001 | 22.226 |
| 24.237 | 17.704 | 17.456 | -0.600 | -0.155 | 3.498 | -25.118 | -7.080 | 4.778 | 3.480 | -23.477 | 19.184 | 8.772 | -22.097 |
| -25.811 | -27.310 | 25.198 | -22.265 | 4.305 | 28.157 | -3.512 | 20.969 | 25.985 | -27.461 | 20.391 | -20.971 | -19.997 | 0.773 |
| -1.800 | -29.217 | -14.136 | -3.563 | 9.458 | 19.593 | -0.716 | 25.290 | 24.335 | 8.452 | -11.854 | -17.902 | 18.459 | -23.564 |
| -10.243 | 19.045 | -15.681 | 17.591 | 6.981 | 23.459 | -17.862 | 23.335 | 4.080 | 12.220 | -15.730 | 13.293 | -1.051 | 23.999 |
| 26.363 | -15.791 | 2.480 | 19.718 | -11.912 | -1.873 | -5.903 | -0.314 | 24.006 | -19.782 | 23.377 | 19.929 | -8.414 | 17.346 |
| 3.723 | -19.706 | -20.757 | 28.146 | 12.843 | -17.914 | 3.140 | 9.892 | -26.902 | 10.006 | 25.117 | 20.643 | 15.633 | -0.622 |
| -0.682 | -17.657 | -7.742 | -27.456 | 14.181 | 24.291 | -24.167 | -7.943 | -23.665 | -22.603 | 15.587 | 10.336 | 22.902 | 6.161 |
| 9.639 | -10.889 | -4.329 | -28.493 | 13.755 | 5.454 | 8.725 | 26.008 | 13.821 | -6.484 | 16.145 | -22.961 | 2.938 | -22.500 |
| -2.686 | 23.083 | 7.745 | -8.732 | 8.533 | 18.695 | 18.307 | -13.793 | -17.635 | -13.273 | -10.726 | -5.120 | 21.027 | -1.672 |
| 15.656 | -20.162 | -21.857 | -7.380 | 17.855 | 6.255 | -1.354 | -6.364 | 23.521 | -25.656 | -4.693 | -21.661 | 3.171 | 22.503 |
| -12.712 | 10.340 | 19.740 | 0.115 | 9.753 | -26.445 | -2.348 | 5.865 | 15.031 | 15.544 | -18.921 | 22.836 | 17.682 | -4.537 |
| -28.080 | -28.229 | 24.779 | 4.205 | 6.077 | 16.666 | 7.451 | 10.799 | 18.104 | 6.137 | -21.397 | 15.310 | 21.976 | 15.753 |
| 11.081 | 17.736 | -13.575 | 3.416 | 8.198 | -3.664 | 5.019 | -25.136 | 8.335 | -1.956 | 18.800 | -19.593 | 1.713 | 13.548 |
| 8.408 | 22.392 | -5.812 | -4.172 | -7.889 | 29.024 | 21.187 | 10.273 | -7.821 | -12.799 | 9.041 | -24.452 | -12.183 | 16.555 |
| 29.051 | -0.106 | -8.945 | 9.656 | -21.519 | 22.962 | -7.735 | -6.564 | -19.321 | 5.389 | 6.942 | 25.085 | 25.852 | -6.741 |
| 0.182 | 18.515 | 6.814 | 23.537 | -17.248 | 13.532 | -22.950 | -5.622 | -28.151 | 1.903 | -25.555 | 4.331 | 22.757 | 0.955 |
| 9.175 | -12.613 | -6.405 | 22.946 | 16.188 | 4.149 | -0.835 | -8.893 | -16.152 | -18.226 | 26.229 | 6.351 | -5.818 | 18.507 |
| -0.655 | 29.627 | 14.386 | -26.852 | 3.255 | -27.986 | 8.100 | 26.605 | -18.824 | -2.962 | -17.214 | 0.276 | -11.018 | 2.235 |
| -4.281 | 3.511 | -14.390 | 5.146 | -6.138 | 8.275 | -5.401 | -17.946 | 26.333 | -3.649 | -14.250 | 16.447 | -24.318 | -0.159 |

| | | | | | | | | | | | | | |
|---|---|---|---|---|---|---|---|---|---|---|---|---|---|
| -22.298 | 27.994 | 24.009 | -9.515 | 16.860 | -0.965 | -7.210 | -19.916 | -26.237 | -16.941 | 15.665 | 7.288 | -10.267 | -10.049 |
| -13.380 | 6.494 | -25.391 | 27.765 | -10.230 | 22.531 | -27.714 | 25.791 | -3.632 | -17.490 | -22.692 | 8.064 | -16.429 | 25.061 |
| 27.676 | -8.120 | 13.058 | 9.963 | 19.941 | -23.177 | -4.190 | -16.765 | -4.095 | -5.351 | -28.177 | -26.579 | 4.427 | 2.944 |
| -19.680 | 18.740 | -28.291 | 7.196 | 19.343 | 27.052 | -9.540 | -6.171 | -6.372 | -1.200 | 16.765 | 20.414 | -22.559 | -0.161 |
| 19.882 | -5.764 | 19.032 | 0.684 | -2.831 | -17.905 | -22.389 | -8.449 | 21.935 | -12.030 | 0.751 | 24.570 | -24.923 | 6.331 |
| 3.606 | -11.092 | -22.955 | -3.296 | 17.464 | 2.502 | 17.918 | -20.619 | 0.367 | 1.998 | 7.919 | -19.750 | 18.478 | 10.820 |
| -10.467 | -22.579 | 7.329 | -2.501 | -3.613 | 23.995 | -25.738 | -19.852 | -19.442 | -18.891 | 21.540 | 12.076 | 17.538 | 20.869 |
| -1.702 | -25.091 | 20.713 | 9.010 | 10.330 | -2.784 | 9.690 | -19.404 | 19.600 | 5.332 | -17.819 | 12.828 | 1.213 | 4.184 |
| -20.787 | -8.649 | 9.583 | -6.848 | 15.477 | -5.820 | 3.157 | 18.154 | 9.856 | -21.003 | -3.571 | 21.946 | 19.855 | -9.208 |
| 2.720 | -2.649 | -2.708 | -12.109 | 11.687 | 28.076 | -12.438 | -3.077 | 13.135 | 8.867 | 24.004 | 17.219 | 0.261 | 12.400 |
| 11.883 | -6.701 | 15.062 | -13.850 | -8.139 | -24.296 | 2.424 | -0.738 | -25.129 | 4.660 | -1.435 | -10.819 | 25.202 | -10.995 |
| -3.623 | -26.154 | -0.376 | -17.861 | -0.642 | -26.654 | -11.194 | 27.405 | 18.659 | 24.839 | -25.083 | -16.416 | 23.963 | -6.416 |
| 8.627 | 15.599 | -15.413 | 1.008 | 22.986 | -14.970 | -11.950 | -17.132 | 6.921 | 19.110 | 2.300 | -7.724 | 22.975 | 26.044 |
| 13.914 | 25.422 | -0.967 | 14.401 | 23.591 | -2.253 | -20.134 | 9.518 | -16.995 | 7.120 | -16.802 | 2.370 | -21.786 | -8.331 |
| 22.462 | 5.040 | -6.299 | -8.354 | 0.741 | 10.974 | 21.268 | -8.316 | 5.177 | -13.402 | -20.758 | -8.854 | -24.884 | -24.678 |
| -19.501 | -10.225 | 8.121 | -27.546 | 3.647 | -18.277 | -3.753 | -26.007 | 12.561 | 0.414 | 14.676 | -11.943 | 17.918 | 8.165 |
| -28.139 | 16.687 | 18.762 | -27.023 | 5.234 | 21.967 | -21.787 | 15.703 | -13.763 | -18.951 | 18.618 | -14.281 | 12.830 | -26.135 |
| -2.139 | 9.082 | -4.820 | 6.199 | -17.420 | 20.242 | -23.827 | 23.413 | -14.886 | 12.139 | -8.413 | -20.065 | 16.223 | 21.306 |
| -3.939 | 19.047 | 17.646 | 4.649 | 8.858 | 19.287 | 22.059 | 15.022 | -25.473 | -2.073 | 5.386 | -1.966 | 23.121 | 1.307 |
| -23.423 | 4.044 | -10.366 | 3.136 | 12.710 | 10.079 | 8.009 | -0.446 | 27.134 | -10.206 | 8.944 | -20.489 | -25.058 | -4.808 |
| -26.257 | -16.200 | 8.267 | 24.020 | -4.101 | 2.059 | -17.328 | 0.947 | -14.382 | 1.020 | 0.672 | -25.550 | -7.855 | -3.491 |
| -26.679 | 9.641 | 21.530 | 22.449 | 16.203 | 12.284 | 5.493 | 5.999 | 15.328 | -0.225 | 4.516 | -24.749 | -4.398 | 26.261 |
| 11.441 | 10.530 | -12.733 | 5.506 | 0.409 | -7.421 | 3.457 | -26.815 | -8.657 | 22.678 | -19.664 | -12.552 | 0.121 | 1.878 |
| 23.376 | -1.998 | 28.106 | -2.756 | -10.447 | 17.726 | -11.247 | -26.050 | 3.583 | 18.046 | -27.843 | 21.400 | -17.817 | -8.560 |
| 18.295 | 18.380 | -24.929 | 1.896 | -12.674 | 25.625 | -0.472 | -7.103 | 21.144 | 18.632 | -20.043 | 6.192 | 15.451 | 8.987 |
| 30.526 | 20.601 | 19.223 | 14.793 | -20.891 | 20.887 | -6.444 | 27.959 | -28.093 | -21.728 | 25.410 | 23.528 | -2.430 | 12.989 |
| -24.985 | 16.494 | 14.777 | -15.401 | 15.297 | -9.214 | -15.318 | -14.713 | 24.089 | 4.708 | -26.933 | 7.296 | 21.408 | 21.920 |
| -5.453 | -16.967 | -14.791 | 26.962 | -13.946 | -29.299 | 2.803 | 10.215 | -11.002 | 22.971 | 28.953 | -6.601 | 17.162 | 27.422 |
| 5.217 | 27.228 | -29.296 | 27.582 | 19.793 | -11.953 | -26.591 | 15.091 | 8.276 | 18.013 | -21.131 | 14.735 | -26.279 | -22.520 |
| 20.059 | 14.913 | 25.967 | -0.786 | -0.809 | -2.014 | -7.746 | 0.500 | -1.682 | 16.218 | -23.226 | -20.170 | 27.683 | 2.623 |
| 13.877 | -23.696 | 24.526 | 13.230 | -5.011 | -1.105 | -3.251 | -17.611 | -5.555 | 3.411 | 12.573 | -19.142 | 28.378 | 23.293 |
| -13.976 | -19.747 | 22.944 | -13.933 | -2.401 | 27.337 | 0.901 | 28.820 | -6.545 | 25.877 | 14.540 | 3.406 | 22.268 | 20.777 |
| -20.767 | 22.520 | -26.252 | 14.399 | -16.670 | 14.225 | -21.454 | 21.965 | 20.562 | -23.160 | 5.843 | -9.442 | -12.402 | 1.698 |
| 25.951 | -12.633 | -28.645 | -8.373 | 6.047 | -17.194 | 8.227 | 20.356 | 27.166 | 6.031 | 20.213 | -6.630 | 8.497 | 0.178 |
| 15.999 | -18.714 | -27.894 | 25.998 | 5.101 | -21.786 | 4.778 | 28.165 | 27.651 | 8.266 | 21.972 | -12.852 | 25.659 | -10.117 |
| -13.249 | 18.793 | -6.614 | -7.181 | 19.973 | -0.462 | -1.743 | 27.310 | -12.032 | -22.428 | 3.019 | -7.504 | 24.962 | 27.747 |

| | | | | | | | | | | | | | |
|---|---|---|---|---|---|---|---|---|---|---|---|---|---|
| -15.636 | -2.434 | 25.451 | -24.516 | 21.956 | 23.407 | -18.433 | 25.273 | 4.932 | -8.726 | 17.513 | 4.432 | 10.313 | 7.926 |
| 24.591 | 21.342 | -23.950 | -10.085 | -1.714 | 14.918 | -10.826 | -1.164 | 0.811 | -11.137 | 16.499 | 8.214 | 21.484 | 9.601 |
| 28.918 | 2.864 | 30.358 | 5.191 | 7.143 | -26.406 | 0.731 | 29.217 | -14.118 | -9.724 | -3.137 | 5.010 | -24.557 | 12.605 |
| -27.748 | -15.123 | 2.589 | 0.602 | -10.957 | 9.616 | -4.084 | -14.586 | 25.540 | 2.055 | -7.346 | 21.073 | -27.733 | 11.007 |
| 7.344 | -5.116 | 14.206 | -4.295 | 2.871 | -22.782 | 15.155 | -8.569 | 25.939 | -14.363 | 15.456 | 13.035 | 24.084 | 4.338 |
| 21.508 | 20.178 | -3.668 | -17.868 | -0.704 | 22.845 | -17.263 | -7.361 | 7.642 | -26.452 | 7.906 | -3.181 | -14.543 | -5.474 |
| 9.232 | -10.550 | -16.020 | 8.909 | 24.753 | -18.526 | 11.547 | -6.961 | -19.689 | 5.112 | -18.600 | 22.658 | -16.461 | -21.901 |
| -8.584 | 17.569 | 5.747 | 5.971 | 26.732 | 23.859 | -19.250 | -20.814 | -23.429 | -13.563 | 26.061 | 8.980 | -24.426 | -27.799 |
| 18.295 | 1.095 | 2.928 | -14.490 | 5.745 | -7.897 | -26.166 | 22.361 | -29.252 | -24.400 | 23.536 | -16.760 | 20.266 | 13.981 |
| 17.164 | -18.725 | 1.141 | -21.488 | -4.651 | -5.212 | 7.177 | 29.780 | -29.819 | -27.701 | -28.371 | -18.890 | -26.481 | 6.672 |
| -11.504 | 25.537 | 10.530 | -0.609 | -19.862 | -16.928 | -5.096 | -28.362 | 19.866 | 13.075 | 6.429 | -2.148 | 16.287 | -17.232 |
| -2.312 | 8.522 | -26.488 | -1.719 | -3.812 | -10.170 | -6.112 | 3.353 | 8.045 | 7.285 | -13.274 | -4.859 | 27.804 | -3.253 |
| -21.891 | -9.642 | 9.864 | 4.096 | 9.977 | 8.389 | 9.840 | 22.424 | -8.722 | -10.758 | -21.889 | -1.412 | 25.822 | 2.346 |
| 21.539 | -24.932 | 22.822 | 1.577 | 8.444 | -6.793 | -8.531 | 15.413 | 12.244 | -4.768 | -2.988 | -17.422 | 24.682 | -6.361 |
| -19.715 | -4.465 | -12.996 | -0.923 | 7.118 | -10.935 | 24.436 | 4.412 | 26.802 | -12.043 | 11.715 | 3.205 | 2.092 | 21.494 |
| 8.743 | 22.235 | -21.395 | 10.547 | -1.025 | -20.325 | 1.589 | -1.813 | -27.749 | 27.668 | 25.462 | -20.693 | -7.427 | 11.063 |
| -20.375 | -26.061 | -11.455 | -12.801 | -15.404 | -28.833 | 6.007 | -23.831 | -24.158 | -0.907 | -11.306 | -4.791 | 21.090 | -4.650 |
| -24.396 | -22.304 | -12.461 | -11.166 | 6.303 | 30.465 | -9.921 | -28.809 | 2.550 | -5.228 | -29.898 | -5.609 | 5.041 | 4.818 |
| 2.447 | 11.905 | -18.691 | 11.410 | 6.058 | 21.161 | 3.739 | -15.644 | -22.586 | -7.704 | 28.916 | 0.563 | -20.980 | 3.858 |
| -0.210 | 2.026 | 7.662 | -6.153 | 1.754 | 29.223 | -20.341 | -26.009 | 6.539 | -22.519 | -30.243 | -6.560 | -29.129 | 5.586 |
| 23.766 | 0.718 | -10.319 | 2.831 | 10.771 | 4.881 | -25.180 | -18.122 | 11.362 | 8.009 | 9.303 | 2.991 | 22.176 | 16.192 |
| 29.678 | -27.754 | -19.512 | 0.035 | 5.613 | -11.400 | -11.403 | 8.257 | 7.685 | -3.728 | -16.450 | 7.706 | 9.859 | -19.188 |
| 13.174 | 13.143 | 19.530 | 0.735 | -13.674 | 27.195 | -12.610 | 4.887 | 7.051 | 26.420 | 13.105 | 4.328 | -24.525 | 23.281 |
| 7.570 | 16.940 | 12.876 | -2.052 | 13.603 | 23.514 | 2.836 | -24.342 | 8.996 | -7.264 | 11.014 | -4.021 | 14.932 | 18.129 |
| 4.243 | 0.129 | -0.808 | -9.759 | 23.364 | 28.657 | -2.019 | -27.764 | 5.245 | -2.699 | 14.549 | 21.743 | 28.678 | -18.764 |
| -7.142 | -2.236 | -4.307 | 26.043 | -6.654 | 20.819 | -26.666 | 19.531 | -1.508 | 19.862 | -4.670 | 8.996 | 23.737 | 8.736 |
| 1.385 | 26.887 | -17.079 | 23.551 | 22.834 | 19.308 | 0.463 | -15.675 | -16.973 | -15.506 | 27.247 | 7.015 | 27.388 | -5.358 |
| -3.326 | -3.006 | -24.874 | 3.823 | 25.987 | 24.571 | -5.236 | -14.538 | -24.624 | -19.707 | -0.852 | -3.287 | -20.936 | -3.163 |
| 1.648 | 9.096 | -8.243 | -6.164 | 19.538 | -14.943 | -25.602 | 14.947 | -28.413 | 14.752 | 15.606 | 2.465 | 25.749 | 3.066 |
| 8.907 | 11.843 | 22.051 | -25.284 | -15.508 | 6.518 | 4.440 | -2.148 | 25.331 | 15.703 | 11.303 | 5.058 | -3.748 | 2.166 |
| 19.213 | 25.509 | 14.793 | -3.918 | 24.826 | -0.098 | 13.576 | -5.921 | 24.360 | 24.732 | 9.940 | -6.998 | -7.979 | -14.853 |
| 26.516 | 6.484 | 12.305 | 10.082 | 28.396 | 12.632 | 11.474 | -19.287 | 22.828 | 8.349 | 28.392 | 6.336 | 19.509 | -7.150 |
| -24.058 | -25.748 | 6.181 | 11.413 | -13.592 | -21.004 | -27.344 | 26.691 | -3.144 | 3.489 | 22.586 | -1.334 | -26.579 | 29.119 |
| 22.255 | -24.489 | -29.475 | 0.457 | 9.854 | -26.122 | 9.972 | 18.486 | 28.742 | 0.886 | -17.184 | -4.807 | 4.291 | 15.829 |
| 0.095 | -12.801 | -22.907 | 5.600 | -9.114 | -4.560 | -1.733 | -28.798 | 23.341 | -20.567 | -21.133 | 22.518 | 9.119 | -3.641 |
| 24.971 | 21.392 | 14.882 | -11.001 | 15.904 | 21.610 | -8.595 | -11.377 | 29.965 | 19.343 | 1.994 | 7.364 | 24.379 | -10.701 |

| | | | | | | | | | | | | | |
|---|---|---|---|---|---|---|---|---|---|---|---|---|---|
| 10.163 | 12.718 | -5.194 | 24.837 | 11.086 | 17.801 | -24.717 | -27.296 | -14.959 | -14.787 | -12.770 | 6.102 | -1.666 | -4.344 |
| -16.472 | 0.785 | -14.703 | 19.274 | -12.058 | -14.351 | 6.240 | 17.362 | 10.977 | -27.381 | -28.186 | -24.411 | 6.973 | 10.084 |
| 21.127 | -10.717 | -17.359 | -13.677 | 13.898 | 9.604 | -19.571 | -22.030 | -27.279 | 28.943 | -0.669 | -4.730 | 26.724 | 21.955 |
| -1.526 | 19.172 | 17.559 | 0.187 | 25.461 | -18.575 | 12.735 | 21.747 | 7.116 | -20.516 | 18.286 | -0.248 | 27.655 | 0.412 |
| 5.608 | 9.625 | -18.735 | 6.146 | 20.057 | -27.138 | 6.919 | -29.597 | 20.150 | 10.831 | -29.452 | -1.916 | -13.281 | -12.153 |
| 19.394 | 13.661 | -21.756 | 0.787 | 7.399 | -24.185 | 10.378 | -23.871 | 17.159 | 25.130 | 23.975 | -8.054 | 23.284 | 14.181 |
| -1.057 | -22.903 | 9.671 | -1.786 | -2.643 | 16.362 | -1.216 | 0.181 | 3.954 | 18.144 | -19.138 | -3.390 | 0.843 | -1.012 |
| -13.310 | -10.954 | 6.954 | -7.116 | 7.738 | -29.984 | -6.216 | 26.243 | -21.567 | -25.052 | 8.064 | -2.751 | 23.639 | 24.467 |
| -6.873 | -17.402 | -11.961 | 17.321 | -1.112 | 27.425 | -14.100 | -3.637 | 10.690 | 17.737 | 23.032 | -15.047 | 25.477 | 1.507 |
| 15.885 | -24.770 | -18.096 | 2.258 | -15.079 | -13.030 | 0.578 | -8.592 | 26.945 | -5.337 | 2.357 | -22.535 | 22.707 | -28.208 |
| -6.326 | -0.612 | 5.258 | 4.707 | 9.452 | 28.295 | -22.933 | 12.151 | 30.234 | -1.503 | 11.630 | -4.405 | 20.845 | 22.731 |
| 16.552 | 0.874 | 13.483 | 16.486 | 8.152 | 18.990 | -7.326 | -28.815 | 1.017 | -19.186 | 25.606 | -0.161 | 20.415 | -3.487 |
| -19.242 | 22.198 | -2.957 | 9.957 | -9.261 | 13.529 | 3.495 | -15.845 | -5.021 | -24.363 | 4.478 | -1.570 | 26.093 | -11.070 |
| 7.421 | 10.765 | 10.309 | -8.470 | 1.140 | 22.369 | -24.118 | 24.410 | -0.014 | 19.390 | -11.800 | -12.606 | 22.091 | -8.554 |
| 12.242 | 4.567 | 24.068 | 25.401 | -7.337 | 14.839 | -2.185 | 16.351 | 4.756 | 14.746 | -22.866 | 25.149 | -28.246 | 13.160 |
| -22.685 | 19.621 | -15.718 | 1.158 | 7.067 | -0.009 | -12.539 | 10.056 | -25.485 | -19.972 | -17.507 | 4.456 | 16.099 | 7.905 |
| 27.973 | -4.132 | 17.655 | 16.710 | 26.428 | 11.581 | 9.498 | -18.338 | -25.925 | 9.705 | 0.924 | -8.826 | 29.729 | -1.530 |
| -3.395 | -14.628 | -1.049 | 18.457 | 10.587 | -19.423 | -6.203 | 31.994 | 18.516 | 7.555 | 4.439 | 10.216 | 17.495 | 11.421 |
| 11.367 | 6.378 | -23.786 | 16.270 | 6.344 | 15.353 | 1.651 | -21.955 | 25.211 | -7.261 | 21.724 | 3.452 | 23.496 | 9.017 |
| 10.417 | -24.433 | 19.774 | 23.943 | -26.483 | -22.116 | -0.479 | 10.821 | 5.471 | -11.463 | 29.701 | -20.430 | 30.980 | -21.864 |
| -13.300 | 22.695 | -14.409 | 10.362 | 24.576 | -30.842 | -4.943 | -24.462 | 26.467 | 12.329 | -30.096 | -19.529 | 21.880 | 20.477 |
| -17.549 | -26.067 | 30.513 | -11.137 | -25.063 | -24.693 | 3.648 | 26.505 | 8.620 | 1.575 | 14.898 | -11.207 | 19.397 | -7.574 |
| -26.770 | 23.498 | -22.483 | 24.155 | -24.190 | 7.017 | 1.152 | -2.619 | 15.443 | 25.207 | 1.288 | -5.121 | 27.282 | 27.948 |
| 23.341 | -10.105 | -16.510 | -21.451 | 29.421 | 2.675 | -13.302 | 3.733 | -5.620 | 22.200 | -6.167 | 7.464 | -21.067 | 23.397 |
| 14.907 | -2.373 | 25.196 | -9.926 | 4.702 | 24.931 | 15.042 | -30.576 | -23.015 | 1.080 | 21.122 | 25.697 | 28.764 | 24.571 |
| 20.039 | -23.890 | 20.625 | 21.540 | 24.104 | 29.558 | 13.589 | -0.560 | 21.476 | -20.562 | -3.669 | -5.231 | -15.134 | 19.135 |
| -9.463 | 26.041 | 0.009 | -7.843 | 11.855 | 17.313 | 7.810 | -28.099 | 9.592 | 23.804 | 13.153 | 5.159 | 20.965 | -7.151 |
| -7.565 | 5.700 | 21.478 | -0.715 | -0.919 | 18.324 | 4.309 | 1.022 | 18.981 | -26.726 | 19.111 | -14.994 | -29.331 | 3.608 |
| 23.014 | 5.369 | -2.292 | 24.918 | -27.007 | 6.494 | 9.766 | -4.275 | -24.346 | -8.807 | 28.426 | -10.746 | -30.000 | 21.044 |
| 19.249 | 7.237 | -12.654 | 18.688 | 21.486 | -22.901 | 4.983 | -20.892 | 1.645 | 28.431 | 26.274 | 24.699 | 20.499 | -13.669 |
| -24.813 | -23.970 | -15.750 | 20.359 | 10.418 | 20.858 | -6.929 | -30.995 | -25.368 | -15.729 | 1.746 | -5.335 | 23.898 | -6.157 |
| -3.549 | 20.652 | 27.720 | -20.478 | 7.416 | -18.777 | 8.185 | 14.263 | 12.264 | -25.893 | -5.847 | -12.652 | 24.371 | 5.639 |
| 11.392 | -9.269 | 15.241 | -23.054 | 6.249 | -28.228 | -0.261 | 18.713 | 28.033 | 22.636 | -15.337 | 6.075 | 22.453 | 5.095 |
| -9.409 | 24.286 | 9.865 | 18.896 | 9.905 | 25.885 | -12.842 | -21.482 | 30.121 | 16.913 | -26.418 | 27.695 | 29.953 | 22.833 |
| 25.250 | -10.184 | 3.126 | -15.116 | -14.489 | 13.824 | -4.910 | 27.228 | -6.142 | 27.511 | 17.073 | 20.794 | 25.539 | 9.564 |
| 21.127 | 12.498 | -25.117 | -26.045 | -16.213 | 0.362 | -2.720 | -25.635 | 29.013 | -3.574 | 23.484 | -5.830 | 13.812 | 10.698 |

| | | | | | | | | | | | | | |
|---|---|---|---|---|---|---|---|---|---|---|---|---|---|
| 21.698 | 5.758 | 18.801 | 25.108 | -17.677 | 27.229 | 19.883 | 29.284 | -9.510 | 25.089 | 8.206 | -6.894 | 27.950 | 10.262 |
| 13.582 | 13.809 | 13.742 | 15.439 | -8.570 | 23.422 | -21.620 | -19.446 | -14.537 | 24.392 | -22.109 | 3.634 | -29.067 | -1.413 |
| -16.919 | -2.290 | -14.396 | -21.766 | 0.775 | 12.883 | -20.678 | 24.873 | 15.530 | -0.791 | -25.706 | -29.614 | 23.986 | -12.985 |
| 18.499 | 21.440 | -8.133 | -22.972 | 22.845 | 1.519 | -23.021 | -29.218 | -22.449 | 24.020 | 26.909 | 2.913 | 22.189 | -13.911 |
| -18.680 | -1.435 | 24.553 | 25.328 | 19.356 | 19.417 | 9.559 | 0.141 | -27.768 | -23.134 | -4.926 | -22.477 | -21.189 | 26.764 |
| 9.307 | 3.450 | -8.630 | -21.689 | 27.526 | 5.777 | 21.002 | 18.663 | 20.226 | 10.382 | -17.144 | 23.052 | 17.138 | -9.680 |
| 14.691 | 16.445 | 12.259 | -17.476 | 12.024 | -1.498 | -16.295 | -15.976 | -20.692 | -21.857 | 13.435 | -4.197 | 26.222 | 8.070 |
| -21.920 | 23.169 | 2.567 | 9.452 | 20.180 | -8.791 | 1.172 | -30.123 | -31.141 | -5.869 | -16.791 | 25.874 | 8.488 | 13.715 |
| 25.316 | 22.058 | -5.544 | -15.632 | 0.184 | -9.565 | -10.271 | -15.105 | 14.226 | -24.327 | 26.494 | -6.450 | 28.709 | -27.507 |
| -11.217 | 1.637 | -9.541 | -19.819 | 17.074 | 30.827 | 18.228 | 30.136 | 7.418 | -15.016 | 3.968 | -7.612 | -30.072 | -7.868 |
| -4.536 | -11.288 | -1.911 | -24.525 | -2.347 | 14.487 | -12.147 | 23.758 | 13.857 | -19.388 | -28.220 | 3.833 | 24.858 | 11.842 |
| -13.736 | -9.961 | -7.814 | 13.578 | 10.315 | -5.669 | 9.770 | 1.122 | -11.004 | -27.157 | 27.966 | -12.611 | -19.763 | 0.995 |
| -8.104 | -4.819 | -20.879 | 18.557 | -14.762 | -20.104 | 0.835 | -1.092 | -6.730 | -12.954 | -1.130 | -4.582 | -24.505 | -15.085 |
| 8.610 | -0.101 | 6.493 | -22.144 | 17.996 | -12.363 | -19.921 | 10.545 | -10.425 | 30.801 | 23.135 | 31.193 | 16.341 | 14.559 |
| -17.661 | -6.105 | 21.460 | 26.218 | -4.122 | -14.807 | -14.398 | -21.823 | 31.626 | 6.706 | -13.586 | 0.579 | -9.068 | 31.843 |
| -15.442 | 1.181 | -6.879 | 3.685 | -14.283 | -30.245 | -0.315 | -15.210 | -25.905 | 21.053 | -25.024 | 31.068 | 17.987 | 18.654 |
| -4.138 | 19.787 | -15.170 | 25.581 | 6.631 | -31.091 | -23.343 | -24.129 | 24.880 | 2.845 | 29.598 | 2.533 | 17.052 | 28.812 |
| 4.931 | 4.109 | 15.795 | -3.922 | 24.086 | 25.881 | -26.387 | 29.717 | 14.810 | 1.409 | -3.262 | 23.163 | 20.950 | 11.870 |
| 23.822 | 20.236 | -25.021 | 8.504 | -12.175 | 23.886 | 10.730 | -21.341 | -24.699 | -25.240 | 18.751 | -25.633 | 24.411 | 5.988 |
| 20.219 | 9.080 | 24.782 | -26.236 | 20.752 | -7.873 | -14.244 | -0.469 | 30.396 | -12.136 | -7.133 | -6.882 | 22.819 | -8.433 |
| 24.929 | -23.726 | -0.312 | 4.043 | -6.426 | -14.269 | -15.968 | -14.900 | 4.819 | 23.838 | -27.159 | 25.395 | 15.708 | 13.046 |
| 19.776 | -8.781 | -12.018 | 20.716 | -19.330 | 27.488 | 6.401 | -8.470 | -7.140 | -16.931 | 28.939 | -6.418 | 14.875 | -6.621 |
| -6.159 | 19.747 | -2.788 | -0.063 | 27.378 | -21.367 | -24.930 | -30.121 | -26.572 | 29.687 | 30.552 | 18.248 | 28.207 | -6.983 |
| -8.981 | -20.009 | -10.701 | -26.811 | 25.892 | 26.347 | -26.022 | 21.743 | -19.018 | -21.718 | 31.121 | -1.355 | 25.236 | 29.565 |
| 1.011 | 12.358 | 14.049 | -25.598 | -4.331 | -31.191 | 24.434 | 18.791 | 21.598 | 19.314 | 9.250 | 5.765 | 28.029 | 2.469 |
| 3.883 | -2.951 | 16.182 | 11.305 | -7.655 | 18.960 | -2.974 | 25.052 | 23.656 | 27.293 | 4.573 | 29.490 | -28.433 | -0.782 |
| 26.431 | -15.533 | -16.923 | 25.645 | 27.904 | -4.787 | -2.684 | -2.198 | -3.282 | -19.502 | -22.773 | -28.690 | 23.814 | -5.790 |
| -21.191 | -17.348 | 21.155 | 24.540 | -24.559 | 22.883 | -18.648 | 5.400 | -10.952 | 25.122 | 11.216 | -6.206 | 20.616 | 28.768 |
| 14.818 | -2.689 | 7.633 | 7.877 | 5.030 | 8.536 | 26.285 | 0.077 | 7.601 | 23.964 | 9.528 | 0.466 | 0.618 | -30.971 |
| -24.398 | 4.573 | -1.210 | 27.497 | 2.321 | -10.045 | 22.850 | -13.343 | -9.913 | 20.889 | 10.468 | -27.402 | -27.199 | 20.157 |
| 11.598 | 19.835 | 31.389 | 17.969 | -22.059 | 12.350 | -19.788 | -8.558 | -20.920 | 19.923 | 2.699 | 1.727 | 27.722 | 2.514 |
| 19.203 | -6.435 | 13.443 | -24.587 | -5.905 | -10.734 | 7.543 | 10.435 | 2.315 | -21.193 | 1.335 | -1.104 | 18.151 | 0.503 |
| 6.862 | -7.794 | 8.416 | 28.062 | 11.957 | 13.850 | -26.925 | -28.360 | 5.147 | -25.378 | 30.438 | -28.164 | 29.818 | 3.577 |
| -16.991 | 4.168 | -30.657 | -5.430 | -16.833 | 10.053 | -17.059 | 5.932 | 23.417 | -6.914 | -23.537 | 17.558 | -8.283 | -7.884 |
| 2.435 | -5.431 | 19.276 | -6.430 | 19.389 | -25.084 | -22.723 | -20.616 | 16.720 | -9.873 | -5.262 | -0.677 | 26.812 | -1.194 |
| 13.580 | 1.240 | 15.294 | 14.672 | 19.875 | -4.094 | -22.044 | 9.906 | 29.365 | 8.204 | 1.505 | 0.985 | 25.997 | 20.857 |

| | | | | | | | | | | | | | |
|---|---|---|---|---|---|---|---|---|---|---|---|---|---|
| 13.828 | -1.854 | -19.811 | 21.980 | -11.942 | 17.109 | -27.131 | 14.885 | 10.003 | -23.416 | 18.317 | -16.592 | 11.533 | 6.042 |
| 1.316 | -7.876 | 2.592 | 26.519 | -22.187 | -27.919 | 20.059 | 8.343 | 24.802 | -27.014 | 4.833 | -28.881 | 22.220 | 6.553 |
| -22.226 | 18.278 | 6.559 | -0.482 | 9.655 | -11.243 | 26.063 | 1.765 | 22.057 | -3.590 | 24.291 | 4.701 | 21.365 | -8.654 |
| 24.403 | 10.061 | -24.880 | -1.957 | -3.726 | -23.292 | 22.550 | -30.679 | 12.218 | 28.839 | -18.070 | 1.523 | 26.195 | -2.852 |
| 4.412 | 8.986 | -14.072 | -22.636 | 13.753 | -25.664 | 13.553 | -25.926 | -26.234 | -20.189 | 21.287 | 11.279 | 19.081 | 17.158 |
| -11.365 | -18.612 | 20.876 | -3.132 | -7.477 | -1.796 | 27.293 | 24.315 | 18.173 | -20.521 | 23.532 | -4.420 | 27.696 | 4.468 |
| -4.910 | -1.500 | 12.167 | 13.807 | 18.648 | 27.105 | -24.301 | 12.559 | -15.360 | 20.840 | -8.543 | -23.805 | -6.825 | -4.304 |
| -21.323 | 5.519 | 22.184 | 23.062 | 5.380 | 13.208 | -17.310 | -23.826 | 7.969 | 4.565 | 9.721 | 3.864 | -22.141 | -22.940 |
| 0.645 | -2.675 | -0.237 | 12.600 | -6.362 | -6.739 | 3.714 | 7.963 | -13.606 | -12.096 | 15.933 | 7.267 | -20.159 | 6.455 |
| 2.888 | 3.049 | 5.433 | 8.213 | -26.960 | -26.574 | -17.799 | -31.045 | -29.895 | -23.935 | 2.513 | 19.511 | -26.012 | 2.989 |
| 8.471 | -7.464 | -10.719 | 5.302 | 26.408 | 19.569 | -27.061 | 23.414 | 27.091 | -22.983 | -7.288 | -9.835 | 9.358 | -26.519 |
| -23.289 | 17.154 | 27.584 | -24.902 | 7.364 | 5.901 | -21.143 | -0.806 | 21.663 | -6.348 | 18.492 | -12.396 | -24.928 | -12.902 |
| 17.646 | -9.116 | -12.901 | -4.843 | 4.500 | 27.680 | -27.610 | -23.562 | 15.958 | 19.034 | 8.139 | 3.599 | -25.533 | 7.685 |
| -24.546 | 11.692 | -5.634 | -11.682 | -11.409 | 1.816 | -24.923 | 17.727 | -19.161 | -21.468 | -12.529 | 20.609 | 15.326 | 25.499 |
| 23.779 | 0.352 | 21.663 | 24.987 | -13.550 | 17.834 | -23.622 | -16.840 | -3.377 | 9.345 | 13.334 | -3.428 | 28.384 | 5.144 |
| -10.660 | 11.249 | -25.663 | 15.430 | 19.995 | 14.908 | 14.372 | -10.846 | 30.322 | -26.123 | 5.903 | 10.258 | -29.040 | -2.100 |
| -3.608 | 18.627 | -30.615 | 28.136 | -2.877 | 30.914 | -21.709 | -27.735 | 5.972 | -28.359 | 10.260 | -3.527 | 29.253 | 8.913 |
| -9.608 | -4.078 | 11.656 | 24.302 | 23.451 | -23.433 | 20.946 | -26.437 | 19.214 | -24.137 | 21.423 | -4.147 | -22.922 | 1.497 |
| 5.244 | 3.598 | 12.780 | 26.513 | 15.261 | -29.984 | -6.406 | 28.094 | -11.811 | -19.786 | 11.610 | -5.289 | 24.399 | 12.651 |
| 14.495 | -11.783 | 0.468 | 7.723 | 19.349 | 25.155 | -13.969 | -3.336 | -8.044 | 12.171 | -1.736 | 1.841 | 21.790 | -3.436 |
| -18.820 | 0.269 | 3.674 | -23.670 | -18.540 | -29.336 | -11.426 | -20.131 | 7.044 | -0.969 | 20.016 | 26.687 | 20.109 | -30.565 |
| 19.264 | -4.782 | -28.452 | -26.458 | 17.518 | 7.435 | -19.370 | -28.150 | -19.290 | -30.935 | -28.454 | -5.447 | 27.713 | 11.997 |
| 2.580 | -9.429 | -17.500 | -1.723 | 9.918 | 24.377 | 3.015 | -19.317 | 26.296 | 19.598 | 21.905 | -20.859 | 17.550 | 0.899 |
| 18.478 | -8.583 | -16.893 | -5.325 | 8.140 | -30.348 | -5.030 | -6.063 | -26.204 | 25.819 | 5.975 | 20.861 | 27.086 | 6.762 |
| 6.030 | 5.644 | 1.981 | -2.901 | -21.782 | -16.023 | -23.197 | 12.517 | 27.511 | 14.531 | 14.664 | -22.164 | -27.403 | -27.292 |
| -3.544 | -1.797 | -12.974 | 27.743 | -8.678 | 14.361 | 18.527 | -26.096 | -7.047 | 21.973 | 28.134 | -3.699 | 12.864 | 2.576 |
| 3.102 | 8.281 | -2.032 | 10.440 | 12.373 | -3.230 | 27.398 | 27.279 | 26.646 | -23.777 | 25.683 | -0.551 | 26.610 | -1.588 |
| 16.253 | -0.259 | 22.665 | -28.815 | 20.931 | -2.191 | -3.348 | 11.183 | -19.317 | 16.106 | 8.663 | 10.367 | 22.660 | -11.512 |
| 9.347 | 9.049 | 20.424 | -8.200 | 22.556 | 6.492 | 5.157 | -19.949 | -23.303 | -27.788 | 0.817 | 6.054 | 14.860 | 3.405 |
| -11.672 | -2.655 | -1.429 | 2.467 | -27.572 | 26.873 | -27.445 | -23.646 | 28.216 | -17.885 | 24.907 | -12.175 | -23.424 | -29.695 |
| -4.738 | -3.179 | -3.013 | -23.963 | 10.979 | -13.049 | -17.901 | 1.123 | 25.044 | 23.057 | 19.356 | -10.401 | 19.384 | -14.254 |
| 17.956 | 7.547 | -18.562 | -4.410 | 2.652 | 18.325 | 22.406 | -21.850 | -28.346 | 15.582 | -23.351 | 23.180 | 20.622 | 24.601 |
| 10.283 | -25.608 | 14.240 | -12.511 | -5.535 | -5.288 | 28.670 | 25.095 | 6.559 | -25.049 | 23.036 | -2.448 | 21.697 | 7.272 |
| -4.427 | -13.741 | -19.598 | 14.025 | -3.999 | 26.426 | 7.939 | 11.080 | -10.731 | 6.530 | -0.622 | -0.058 | 27.806 | 5.048 |
| -0.932 | 4.513 | -4.011 | -27.195 | 15.777 | 15.189 | -23.627 | -21.168 | 24.321 | 5.729 | 29.284 | 1.784 | 1.487 | -7.652 |
| 14.618 | 3.639 | -6.021 | -27.897 | 14.376 | 23.812 | -20.791 | -22.318 | 29.273 | -13.510 | 0.745 | 9.962 | -26.926 | -5.590 |

| | | | | | | | | | | | | | |
|---|---|---|---|---|---|---|---|---|---|---|---|---|---|
| 5.783 | -10.409 | -10.178 | -24.888 | 28.577 | 20.442 | 0.058 | 12.817 | 5.524 | -18.649 | 23.970 | 5.423 | 26.125 | -10.141 |
| 5.390 | 1.940 | 8.272 | -24.028 | -13.325 | 8.256 | -22.983 | -23.262 | -0.658 | 23.930 | -15.376 | -22.292 | -26.108 | 9.438 |
| -3.730 | -14.870 | 0.769 | 25.808 | -23.299 | -11.850 | -23.407 | 4.949 | -23.700 | -10.886 | 18.290 | 25.999 | 21.164 | 0.412 |
| 0.009 | 0.604 | -6.392 | 18.823 | 31.475 | -1.390 | 5.898 | 28.037 | 4.404 | -22.003 | -16.729 | -18.507 | -19.152 | -6.884 |
| -7.741 | 11.872 | -20.737 | 5.799 | 30.267 | 25.426 | 21.339 | 16.364 | -1.940 | -25.915 | 25.262 | -3.258 | 24.660 | 3.682 |
| -10.332 | 18.731 | -22.100 | -0.291 | -7.602 | -13.394 | -17.639 | -27.891 | -20.707 | 20.444 | 21.107 | 1.146 | 22.216 | -1.557 |
| -6.908 | 4.862 | 11.689 | -1.683 | -28.775 | -21.817 | 16.497 | 10.557 | -16.483 | 14.781 | -27.890 | 26.915 | -28.164 | 5.569 |
| 21.685 | -4.122 | -16.470 | 28.921 | 14.055 | 14.242 | 25.336 | 14.363 | -8.649 | 26.586 | 22.261 | -2.784 | 22.471 | -2.869 |
| 8.100 | 22.559 | -20.201 | 16.671 | -22.900 | -29.719 | 7.599 | 21.258 | -27.530 | -30.435 | 5.338 | -8.924 | -28.150 | -6.576 |
| 20.737 | 7.860 | 6.194 | 23.089 | -6.968 | -7.969 | 1.133 | 1.836 | 17.697 | -23.203 | -21.800 | 5.296 | -27.555 | 17.892 |
| 9.090 | -19.837 | -7.109 | -25.554 | 2.495 | -27.618 | 17.637 | -19.793 | -4.162 | 12.709 | 22.724 | -4.053 | 25.625 | -9.804 |
| 10.360 | -10.261 | -16.017 | 10.887 | 15.394 | 24.239 | -2.248 | -7.981 | 24.358 | -23.905 | 8.761 | 4.932 | 2.995 | 2.075 |
| -22.581 | 2.638 | -22.966 | -2.928 | -19.331 | -23.043 | 6.576 | -16.676 | -22.761 | -7.478 | -9.613 | -3.710 | 0.511 | 6.070 |
| -8.121 | -4.070 | -8.826 | 26.043 | -24.782 | -14.051 | -25.996 | 26.519 | 20.353 | 22.686 | -24.998 | -0.367 | -22.602 | 24.016 |
| 6.944 | 0.047 | 2.692 | 29.231 | -24.340 | 4.451 | 8.843 | -17.955 | -25.364 | 10.203 | -10.348 | 2.739 | 27.501 | 14.279 |
| 17.396 | -6.253 | -23.798 | 6.343 | 22.573 | -12.100 | -29.199 | 2.553 | -0.481 | 14.534 | -14.351 | 2.662 | 18.624 | 0.682 |
| 24.531 | 10.908 | 28.373 | -26.713 | 9.703 | 8.973 | 2.601 | 27.271 | -2.195 | -21.005 | -5.635 | 4.214 | -5.248 | -4.097 |
| -4.181 | 6.491 | 16.804 | -10.352 | -17.010 | 6.119 | 21.899 | -16.784 | 16.030 | -12.832 | -2.796 | -0.827 | 28.303 | -28.305 |
| 13.885 | -6.414 | 25.173 | 15.986 | 5.509 | 22.289 | 23.016 | -20.918 | 24.686 | -21.732 | 7.359 | -3.977 | 19.039 | 29.856 |
| 19.401 | 1.627 | 24.603 | 9.127 | 6.107 | -3.450 | -3.307 | -24.656 | 23.077 | -4.650 | 18.633 | 7.072 | 9.580 | -15.621 |
| -5.135 | 11.670 | 12.625 | 13.129 | 1.869 | 15.484 | 25.399 | 26.084 | -3.867 | -30.457 | 20.292 | 4.295 | -22.979 | -8.432 |
| 1.613 | 20.561 | 5.983 | -5.592 | -10.860 | 25.698 | 20.911 | -3.415 | 29.487 | 23.718 | -3.817 | 8.458 | -23.490 | 8.524 |
| -19.271 | 8.091 | -4.548 | -17.968 | 0.511 | -7.931 | -25.342 | -24.174 | 14.584 | -7.526 | -15.391 | -2.996 | 19.473 | -7.049 |
| -12.656 | -2.795 | 21.149 | -16.367 | 22.193 | -11.567 | -5.038 | -26.777 | -7.790 | -11.225 | -4.760 | -8.474 | 11.862 | 7.870 |
| -23.338 | 3.244 | 7.058 | 12.003 | 9.251 | -17.859 | -29.383 | -19.871 | -25.542 | -15.475 | -8.854 | 9.447 | 26.537 | 14.623 |
| -6.558 | -11.969 | 22.373 | 22.689 | -10.028 | -8.917 | 8.899 | 14.996 | 19.225 | -18.623 | 4.358 | 1.900 | 25.218 | -8.109 |
| 25.026 | 4.416 | -5.862 | 11.285 | 13.418 | -22.609 | -26.256 | -22.226 | 25.776 | -8.806 | 19.795 | -3.171 | -21.853 | -2.011 |
| 16.358 | 5.851 | -19.352 | 21.807 | 4.630 | -25.011 | 26.082 | 21.037 | 5.137 | -16.455 | 21.092 | -1.235 | -20.389 | 10.551 |
| 18.299 | -4.231 | 26.370 | -9.467 | -8.528 | 21.124 | -4.471 | -3.631 | 29.168 | 14.363 | 14.849 | 25.433 | 23.906 | 10.851 |
| 12.474 | 5.149 | 12.298 | 27.060 | -18.083 | 25.090 | 26.752 | -19.263 | 27.153 | -10.375 | 0.289 | -12.035 | 25.710 | -3.945 |
| -24.445 | -0.729 | 10.760 | 28.818 | -7.579 | -9.705 | 25.014 | -22.881 | 22.116 | -13.670 | 22.784 | -1.016 | -3.579 | 20.797 |
| 20.018 | -3.524 | 26.693 | 18.257 | -12.891 | 24.539 | -27.256 | -16.289 | 10.041 | -19.311 | 16.216 | 15.955 | 20.754 | 3.269 |
| -3.223 | 22.399 | 1.145 | 7.586 | -21.929 | -0.316 | -29.410 | 24.846 | -5.594 | -23.099 | 18.979 | 23.679 | 11.607 | -4.852 |
| -19.910 | 20.419 | -21.905 | 24.030 | 28.195 | 15.259 | 27.862 | -9.699 | -14.303 | -28.499 | -19.880 | -3.884 | 3.680 | -6.049 |
| -2.269 | -8.894 | -3.061 | 29.551 | 22.216 | 2.995 | 2.712 | -18.736 | 18.354 | 19.670 | -6.493 | 11.651 | -23.135 | 12.710 |
| -23.972 | 1.860 | 10.241 | -0.188 | -6.647 | 27.965 | -0.986 | 5.011 | 4.292 | -16.024 | 22.148 | 13.897 | 3.332 | -3.233 |

| | | | | | | | | | | | | | |
|---|---|---|---|---|---|---|---|---|---|---|---|---|---|
| -3.909 | 24.701 | -28.996 | -21.702 | 8.176 | 11.012 | 26.037 | -27.423 | 26.237 | -17.965 | 4.160 | 0.053 | 22.732 | -19.448 |
| 23.418 | 8.980 | 23.994 | -27.005 | -0.078 | -12.750 | -8.507 | -20.624 | 25.398 | 24.924 | -4.244 | 1.562 | -11.975 | 4.165 |
| 2.103 | -8.989 | 9.506 | 23.127 | -20.096 | 10.207 | 26.729 | 1.166 | -6.620 | -28.993 | 16.923 | -2.228 | -24.509 | 24.743 |
| 22.739 | -12.602 | 21.935 | -7.293 | -22.234 | 4.733 | -2.174 | 25.486 | 0.712 | -5.550 | 26.701 | -7.630 | 19.811 | -30.153 |
| -5.719 | 6.043 | 16.328 | 15.134 | 22.978 | 13.301 | -6.179 | -8.075 | -10.386 | -24.927 | -24.397 | -23.642 | 24.825 | 7.600 |
| 26.636 | -13.827 | 3.436 | 5.149 | -18.422 | 7.907 | -17.507 | 10.456 | -24.218 | 1.845 | -1.531 | 6.650 | -13.819 | -0.300 |
| -13.511 | -4.720 | -13.166 | -29.177 | -3.583 | -4.391 | 23.073 | -25.504 | 12.051 | -23.389 | 23.862 | 25.013 | -17.180 | -26.207 |
| -15.185 | -18.024 | -16.666 | 20.818 | -20.719 | -17.316 | 22.282 | 18.819 | 3.069 | -13.998 | -17.828 | -12.498 | 24.191 | -6.269 |
| 14.122 | -18.737 | 0.025 | -19.935 | 9.177 | -27.394 | -22.745 | 0.375 | 4.244 | 24.445 | -25.733 | 20.090 | -24.340 | 2.040 |
| -9.603 | 2.703 | -15.791 | -11.368 | 26.871 | -27.800 | -28.990 | -22.573 | 6.202 | -25.982 | 22.591 | -5.692 | 18.228 | -1.729 |
| -19.895 | -1.187 | -20.529 | -18.302 | 11.618 | 15.022 | 21.264 | -24.523 | -18.052 | -11.250 | -1.294 | -21.489 | 22.771 | -23.292 |
| 11.649 | -19.080 | -11.109 | -29.420 | 0.124 | 11.960 | -22.300 | -26.170 | 19.765 | -18.583 | -9.361 | 1.406 | -10.755 | 2.894 |
| -13.808 | 7.856 | 3.490 | 23.462 | 28.831 | 19.853 | -9.571 | 11.758 | -28.339 | -17.251 | -16.537 | 23.242 | -0.258 | 0.883 |
| -3.410 | -9.426 | -12.770 | 26.546 | 16.732 | -10.707 | -25.281 | -3.520 | -15.809 | -26.448 | 17.305 | -1.349 | -15.728 | 3.504 |
| -25.210 | -5.854 | 14.062 | 22.112 | 12.230 | -20.499 | -29.829 | -8.029 | -2.042 | 28.072 | 18.838 | -25.849 | -10.534 | -3.441 |
| -20.267 | -20.686 | 11.244 | 19.304 | 3.920 | 5.426 | 24.838 | -14.341 | 11.317 | -24.646 | -23.795 | -12.256 | -13.556 | -9.938 |
| 3.001 | 18.044 | -27.709 | 25.883 | -19.542 | 18.970 | -25.944 | 8.048 | 21.117 | -27.275 | 26.524 | -1.743 | -25.130 | 15.109 |
| -8.034 | 10.762 | 18.450 | 0.839 | -3.847 | 18.053 | 30.841 | -4.026 | -9.261 | 25.213 | -11.376 | -6.100 | 23.651 | -9.486 |
| -10.131 | -7.126 | -9.988 | -29.258 | -25.803 | -1.230 | 25.365 | 1.218 | 14.641 | -4.667 | 25.432 | 2.809 | 8.229 | 5.062 |
| 21.427 | -2.611 | -15.265 | -24.763 | -13.470 | 25.632 | 2.536 | -9.310 | 6.764 | -8.422 | 15.319 | -2.788 | -16.496 | -3.110 |
| 9.992 | -7.967 | 12.846 | -20.065 | -22.343 | -12.140 | -2.262 | -14.000 | 0.887 | -21.804 | 16.720 | 10.760 | 16.844 | -7.077 |
| -24.084 | 19.885 | 28.085 | 29.828 | -2.397 | -8.418 | 6.638 | -18.292 | 6.400 | 15.248 | -14.646 | 3.800 | -3.777 | 11.838 |
| 24.137 | 0.526 | 25.276 | 17.778 | -18.134 | -8.023 | -26.538 | -24.949 | -5.392 | 8.892 | 0.721 | -18.403 | -2.631 | 7.176 |
| -7.408 | -22.089 | 4.694 | 28.751 | -6.731 | 23.784 | 28.155 | -2.998 | 27.439 | -7.800 | -14.001 | -2.433 | -7.643 | -10.272 |
| 6.843 | 13.787 | -19.440 | -19.140 | 2.325 | 16.262 | -3.850 | 5.926 | 25.584 | -0.687 | -17.463 | -5.578 | -2.161 | -0.990 |
| 23.491 | 9.885 | 11.487 | 30.928 | -12.532 | 26.855 | 23.229 | -23.707 | 23.625 | -23.867 | 14.320 | -2.228 | 10.197 | 11.275 |
| -21.287 | -10.610 | 0.721 | 24.539 | -7.105 | 11.539 | -0.649 | -20.322 | 8.800 | -10.107 | 9.700 | -7.738 | -7.647 | 9.395 |
| -20.732 | 6.020 | -12.600 | -8.572 | 19.574 | -28.598 | -4.588 | -14.907 | -9.912 | -7.256 | 4.273 | 6.832 | 9.906 | -25.048 |
| 26.285 | 9.474 | -24.057 | -30.120 | 5.420 | 1.529 | -19.261 | 6.713 | 26.278 | -13.335 | 8.450 | 1.617 | 17.059 | -12.034 |
| -20.287 | -13.296 | -26.095 | 18.625 | -25.362 | -6.750 | 30.944 | -20.922 | 27.102 | -15.861 | 1.100 | -6.559 | 8.127 | -1.619 |
| -25.607 | 15.480 | -0.481 | 5.295 | 23.581 | 20.532 | -27.984 | 15.265 | -13.832 | -3.590 | 19.602 | 8.229 | 9.070 | -23.147 |
| 9.122 | -1.719 | 2.553 | 0.166 | -10.893 | 8.452 | 25.935 | 19.774 | -22.630 | 9.193 | -25.327 | -4.456 | 2.280 | -5.700 |
| -8.404 | -19.990 | 19.218 | 25.034 | 1.320 | 27.364 | -18.784 | 23.039 | -1.743 | -4.151 | 24.373 | -0.209 | -23.148 | 13.227 |
| -0.028 | 16.654 | -15.692 | -10.028 | -22.118 | -25.040 | -7.421 | 19.033 | 22.612 | 24.103 | -2.837 | -6.074 | 19.927 | -2.960 |
| -24.901 | -5.242 | -21.121 | 24.265 | -27.166 | 2.950 | -6.417 | -19.065 | 3.064 | 29.370 | -21.563 | 11.928 | -9.729 | 23.157 |
| 20.884 | 22.944 | 24.146 | 27.087 | -21.362 | 22.550 | 3.519 | 0.065 | 23.011 | -6.530 | 11.601 | 2.773 | 21.568 | 5.979 |

| | | | | | | | | | | | | | |
|---|---|---|---|---|---|---|---|---|---|---|---|---|---|
| 14.000 | -4.373 | 16.376 | 22.109 | 11.108 | -19.005 | 17.361 | -20.183 | 5.481 | 19.142 | 24.680 | 17.329 | 8.853 | -5.303 |
| -22.686 | 29.684 | -17.916 | 18.170 | 10.316 | -4.709 | 2.265 | -18.020 | -2.179 | -5.111 | -7.697 | -6.349 | 1.322 | 0.997 |
| -26.518 | 5.884 | -21.992 | -11.288 | -3.131 | -20.316 | 29.835 | 0.917 | -24.969 | -12.310 | 13.626 | 7.561 | -4.793 | -1.305 |
| 1.171 | 14.338 | 25.968 | -0.964 | 28.875 | 23.488 | -2.943 | 21.776 | -22.899 | -27.600 | 10.257 | 7.078 | -12.833 | -4.939 |
| 19.530 | -3.617 | -9.685 | -6.188 | 0.895 | 8.682 | 24.282 | -3.972 | -23.493 | 6.274 | -20.123 | -0.219 | -7.116 | 28.370 |
| -7.176 | 3.050 | -27.770 | 20.893 | -9.480 | -14.025 | 11.757 | 24.980 | -8.951 | -22.306 | -9.988 | -0.791 | -5.973 | 12.093 |
| -15.692 | -2.141 | -0.004 | -31.412 | 14.629 | -27.901 | -21.209 | -19.203 | 0.745 | -7.187 | -10.583 | 28.132 | -24.960 | 3.962 |
| -17.294 | 1.722 | 7.099 | 29.069 | 9.677 | -9.061 | -0.226 | -8.225 | 19.625 | -1.162 | -6.789 | -22.288 | 24.207 | 7.861 |
| 5.515 | 1.078 | 27.303 | 31.448 | -4.718 | 4.248 | -7.871 | -10.633 | -13.504 | 19.576 | 6.973 | -0.905 | 25.173 | 1.958 |
| -20.605 | 12.132 | 8.322 | 28.078 | 26.056 | -16.565 | 26.779 | -12.891 | -3.996 | -12.211 | 25.680 | -5.512 | -24.471 | -3.514 |
| -21.910 | -24.273 | -19.390 | 1.536 | -13.448 | -18.485 | -12.150 | 25.799 | -3.414 | 14.115 | 13.987 | -4.697 | 22.519 | -5.196 |
| -19.756 | -16.311 | -14.678 | -1.716 | 13.196 | 25.021 | -18.049 | -21.063 | -24.590 | -23.579 | -21.435 | -14.898 | -0.297 | 0.151 |
| -2.907 | -20.421 | 15.990 | -21.864 | -0.794 | 14.398 | 6.340 | 24.185 | 23.700 | -2.811 | -17.906 | -23.254 | 21.609 | 5.061 |
| -11.383 | 21.438 | 5.647 | 6.780 | 23.725 | 16.111 | -20.027 | 5.983 | -19.814 | -24.335 | 13.138 | -18.188 | 25.703 | 7.314 |
| 27.028 | 10.255 | 2.965 | 6.292 | -20.107 | 10.285 | 16.059 | -1.419 | -23.801 | -14.713 | -21.350 | -8.693 | 11.762 | 20.202 |
| 15.241 | 6.222 | 18.220 | 4.610 | 5.436 | 27.965 | 7.992 | -7.049 | 4.933 | 2.986 | 18.269 | 21.539 | -21.204 | 24.797 |
| -12.644 | 7.388 | 25.350 | -4.189 | 23.301 | -23.845 | -1.999 | 4.174 | -5.492 | 27.281 | -4.044 | 26.466 | 4.012 | -6.919 |
| 28.347 | -9.760 | -7.010 | -2.391 | 2.368 | -13.541 | 20.843 | -25.455 | -10.277 | -10.418 | -7.463 | -11.135 | 26.793 | 22.491 |
| -23.427 | 5.706 | 22.477 | -6.790 | 22.638 | 13.732 | 30.443 | 12.541 | 23.031 | -17.384 | -22.207 | -6.525 | 21.435 | 21.729 |
| -0.007 | 4.720 | -11.684 | 18.668 | 10.620 | 25.319 | -22.377 | -3.589 | 10.402 | -8.658 | 19.686 | -6.938 | 16.580 | -1.078 |
| -8.108 | -0.773 | 3.345 | 22.944 | 10.377 | -6.034 | -19.851 | 7.046 | -27.565 | 17.677 | 7.283 | -5.869 | 19.502 | 10.402 |
| -14.366 | 10.804 | 26.490 | -0.199 | 8.499 | -20.672 | -18.933 | 1.113 | 4.336 | -7.488 | 17.329 | -19.163 | -0.454 | 3.154 |
| 23.714 | 9.042 | -0.980 | -5.194 | 6.181 | 26.131 | 17.297 | -18.004 | -13.111 | -26.806 | -9.896 | -0.792 | 13.749 | -14.153 |
| 13.122 | 24.258 | 16.917 | 5.262 | 1.195 | 20.544 | -3.596 | -10.308 | -7.117 | 6.581 | -14.884 | -26.029 | -26.502 | 8.947 |
| -18.848 | 20.305 | 7.862 | -8.014 | 13.540 | -22.932 | -11.699 | 17.371 | -9.576 | -28.646 | -12.999 | -14.535 | -2.869 | 0.697 |
| 18.726 | 5.377 | -5.262 | -20.559 | 5.053 | 2.500 | -26.390 | 1.422 | 8.004 | -20.196 | 7.984 | -17.290 | 23.065 | 8.255 |
| -7.026 | 30.914 | -23.403 | 23.641 | 0.520 | 27.161 | -2.409 | 23.434 | 5.814 | -6.223 | 15.602 | -23.326 | 27.193 | -4.007 |
| 5.318 | 7.913 | 21.458 | -0.921 | 3.814 | 24.067 | -1.221 | 25.906 | -13.432 | -22.875 | -8.986 | -2.243 | 19.832 | 6.916 |
| 2.496 | -11.667 | -16.316 | -21.626 | 14.978 | -25.989 | -20.131 | -24.244 | 6.300 | -2.253 | 9.339 | -6.737 | 23.640 | 21.087 |
| 8.061 | 2.192 | 20.613 | 20.682 | 11.888 | 6.095 | 0.435 | -3.920 | -17.710 | -12.771 | -7.407 | 3.549 | 25.247 | 2.686 |
| -1.261 | 8.571 | -22.248 | 0.406 | -19.153 | 27.510 | -27.958 | -6.171 | -23.204 | 10.198 | -8.284 | 2.767 | -5.141 | -8.778 |
| 22.875 | -14.651 | 2.653 | 8.839 | -20.419 | 11.054 | 28.777 | 24.540 | -4.736 | 4.023 | 12.241 | -21.512 | -23.598 | 4.131 |
| 13.634 | 19.306 | -13.293 | -5.497 | -3.234 | 14.367 | -30.711 | -8.316 | 24.338 | -3.965 | -0.682 | 7.637 | -23.245 | 9.525 |
| -9.687 | 17.703 | -6.141 | -21.117 | -3.953 | 22.363 | -4.462 | -10.827 | -21.069 | -25.282 | 1.591 | -13.828 | -20.118 | -0.703 |
| -25.867 | 16.084 | -13.820 | 22.760 | -4.608 | -18.985 | 5.660 | -25.799 | 15.629 | 8.942 | 17.616 | -7.473 | -24.960 | -9.373 |
| -16.515 | 6.200 | 19.743 | -27.433 | 15.716 | 11.827 | -2.081 | -18.185 | -11.904 | -14.014 | 8.995 | -22.118 | -25.854 | -27.934 |

| | | | | | | | | | | | | | |
|---|---|---|---|---|---|---|---|---|---|---|---|---|---|
| 28.987 | 25.852 | 0.247 | 15.503 | -14.491 | -7.779 | 4.096 | 8.141 | 7.022 | 9.676 | 12.869 | -26.957 | 18.707 | 10.985 |
| 26.146 | -5.108 | 21.938 | -18.984 | -21.270 | -13.364 | -5.390 | 16.389 | -3.527 | -12.214 | 9.822 | -9.243 | 24.155 | 23.466 |
| 13.034 | -6.353 | 24.045 | 9.198 | 5.501 | -11.682 | 21.140 | -23.862 | 20.884 | 3.497 | 10.753 | -10.882 | 24.767 | 25.228 |
| -3.894 | 7.620 | -17.509 | 12.476 | -26.984 | 21.124 | 29.316 | 22.551 | 0.096 | -7.006 | 2.309 | 1.374 | 20.678 | 4.777 |
| -8.145 | -27.794 | -19.719 | 6.584 | -7.955 | -20.409 | 24.758 | 21.678 | -6.547 | 15.286 | 23.151 | 20.318 | 25.845 | 5.263 |
| 10.756 | -7.018 | -8.286 | 13.568 | 5.110 | -19.503 | 30.808 | -26.175 | 12.895 | -28.906 | -2.793 | -11.537 | -7.817 | -6.482 |
| -24.000 | 3.080 | -6.986 | -15.093 | 14.133 | 10.475 | -0.861 | -2.048 | 17.454 | -23.359 | -24.907 | -0.626 | -22.288 | -2.479 |
| 24.962 | 7.924 | 18.072 | -0.413 | 4.598 | 5.385 | -20.437 | -0.492 | 24.540 | -26.986 | -3.788 | -7.289 | -25.424 | -10.700 |
| 7.496 | 6.973 | -22.056 | 25.500 | -17.494 | 18.938 | 16.226 | 1.566 | 9.331 | 6.813 | 7.341 | 4.401 | 10.102 | 3.348 |
| 24.028 | -3.148 | -25.428 | -23.521 | -18.104 | -16.333 | -23.753 | -18.074 | 19.402 | -16.333 | 8.061 | 29.583 | -28.123 | 25.481 |
| 18.741 | 28.351 | 5.716 | -21.826 | -18.815 | 23.677 | 21.277 | 20.353 | 18.032 | -20.879 | 3.104 | -26.193 | 27.916 | -9.229 |
| -24.863 | 9.891 | -3.068 | -20.598 | -6.905 | 21.877 | 30.905 | 24.421 | 4.461 | -20.364 | 0.023 | 4.729 | -4.928 | -7.693 |
| 15.381 | -9.641 | 16.251 | -24.914 | -19.887 | 0.598 | 24.635 | -16.018 | -10.206 | 4.599 | 1.572 | 4.437 | -0.464 | -4.276 |
| 2.019 | 17.158 | 15.035 | 2.696 | -11.982 | -17.608 | 0.696 | 8.520 | -21.846 | -19.094 | 14.748 | -28.929 | -27.111 | -0.983 |
| -22.462 | 15.390 | -6.745 | 14.185 | 26.638 | 10.074 | -4.027 | 7.666 | -24.582 | -17.169 | -3.023 | -21.803 | 25.384 | -16.555 |
| -0.501 | 10.725 | -26.853 | -23.396 | 2.812 | -2.071 | -14.022 | -17.156 | 10.258 | -9.213 | -9.858 | -2.250 | 26.683 | -6.360 |
| -19.893 | 1.220 | 22.823 | 1.101 | -21.867 | -6.109 | -22.383 | 25.507 | -13.073 | -18.586 | -2.889 | 8.198 | 25.640 | 8.521 |
| 26.225 | -16.107 | 9.476 | 23.446 | 21.052 | -22.720 | 24.403 | -22.351 | 13.320 | 26.690 | -5.895 | 28.786 | 13.030 | 11.637 |
| -2.625 | 0.037 | 5.643 | -23.014 | -16.232 | 24.640 | 8.153 | -18.559 | -7.856 | -12.719 | 3.768 | 30.816 | 29.248 | -12.560 |
| -9.306 | -10.668 | 12.650 | -21.562 | 17.529 | -11.835 | -6.905 | 17.813 | -0.102 | -5.695 | -6.315 | 1.145 | 12.541 | 4.503 |
| -26.843 | -2.460 | -8.970 | -25.182 | -20.315 | 9.500 | -19.467 | -24.211 | 10.031 | 14.233 | 2.158 | -29.216 | -25.132 | 6.055 |
| 21.540 | -8.130 | 19.832 | 19.265 | -1.491 | -21.742 | -12.748 | 28.935 | 1.292 | -2.135 | 10.302 | -13.752 | 14.679 | -5.190 |
| 1.774 | 25.466 | 17.916 | 28.025 | -5.109 | 21.780 | 28.635 | 19.090 | 25.783 | -2.747 | 23.650 | 26.762 | -1.208 | -14.754 |
| -13.092 | 11.296 | 18.994 | -3.856 | 4.670 | 14.577 | -14.819 | -21.856 | -13.560 | -8.279 | 1.037 | 12.248 | 27.846 | -1.949 |
| -14.752 | -3.695 | -2.277 | 20.673 | 6.087 | 5.171 | -28.904 | -1.970 | 2.177 | 9.583 | 6.472 | -5.331 | 26.470 | -26.739 |
| 16.315 | -5.353 | -9.733 | -9.189 | -2.987 | 22.496 | -2.487 | 6.685 | 24.233 | -6.677 | -1.464 | 25.150 | 26.988 | 4.173 |
| -20.852 | 17.392 | 15.989 | 1.505 | 2.079 | 2.694 | -26.056 | -28.363 | 7.912 | 19.286 | -5.132 | 7.971 | 24.092 | 0.375 |
| -11.913 | 2.145 | 10.912 | 5.145 | -12.218 | 23.513 | 26.231 | 18.311 | 21.766 | -15.252 | 19.328 | 2.333 | -28.803 | 12.624 |
| 14.144 | 7.901 | 16.527 | 2.502 | 26.036 | 15.968 | -1.098 | -27.841 | 6.472 | 17.416 | 25.856 | 12.260 | 23.095 | 11.000 |
| -3.334 | 10.410 | -18.007 | 18.797 | -7.759 | -14.388 | 5.541 | -13.854 | 1.593 | -7.577 | -7.122 | 4.194 | -26.780 | 2.458 |
| 26.034 | 1.142 | 18.635 | 17.220 | 16.880 | -20.942 | 27.418 | 4.736 | 11.396 | 8.573 | 24.967 | -7.320 | 15.732 | -2.215 |
| 23.057 | 18.052 | 14.039 | -4.434 | 10.610 | 3.394 | -23.113 | 7.496 | 8.139 | 7.483 | -5.137 | 18.882 | 19.603 | 16.755 |
| 10.206 | 11.113 | -2.617 | 19.293 | -13.010 | -10.396 | -0.442 | -24.709 | -2.458 | -25.002 | -6.451 | 29.836 | -15.358 | -4.755 |
| 29.443 | -19.071 | 9.028 | 25.553 | -1.410 | -23.958 | 30.212 | 27.542 | -24.468 | -19.494 | -27.349 | 23.684 | 16.726 | -8.124 |
| 10.923 | -17.330 | -6.329 | -24.423 | 16.277 | 19.189 | -23.959 | 23.415 | 20.586 | 11.957 | -5.771 | 4.349 | 17.763 | 5.730 |
| 3.253 | 22.176 | 6.269 | -2.968 | 24.963 | 25.540 | 7.768 | -19.967 | 13.689 | -8.594 | 25.333 | -11.851 | 28.164 | -3.573 |

| | | | | | | | | | | | | | |
|---|---|---|---|---|---|---|---|---|---|---|---|---|---|
| -10.021 | -12.043 | -15.783 | -4.996 | 6.423 | -22.331 | -29.693 | 8.279 | -7.070 | 2.638 | 24.377 | 6.093 | 23.835 | 6.055 |
| -5.384 | 8.719 | -24.755 | 21.489 | 22.414 | -24.802 | 17.480 | 29.007 | -3.140 | -16.976 | 28.092 | 9.364 | 19.009 | 6.959 |
| 17.779 | -10.525 | -18.190 | -21.907 | 7.392 | 18.003 | 6.443 | -10.154 | -0.091 | 23.192 | 24.119 | -27.062 | 19.757 | -20.547 |
| 20.169 | 7.120 | -10.352 | -27.686 | 11.944 | -18.306 | 22.243 | 20.523 | 22.546 | -22.094 | 4.040 | 24.060 | 27.605 | -7.387 |
| -22.830 | -1.776 | 23.666 | 27.566 | -0.723 | -23.423 | -25.003 | 14.857 | 3.209 | 4.769 | -5.032 | -6.732 | 21.287 | -11.101 |
| 26.774 | -11.390 | -22.945 | -5.004 | 25.739 | -25.614 | -28.268 | -0.426 | -2.812 | 1.690 | 25.155 | -3.408 | 28.196 | -0.760 |
| 16.775 | -12.435 | -19.760 | 2.733 | 21.725 | 8.673 | 0.713 | -5.156 | 7.321 | 8.776 | 8.213 | 7.701 | 4.301 | -5.603 |
| 21.644 | -0.028 | 20.826 | -24.132 | -5.381 | 16.041 | -1.026 | -23.458 | 4.540 | 14.501 | 0.396 | -3.921 | -4.076 | -3.952 |
| 28.650 | 13.059 | 4.356 | -21.961 | 24.531 | 20.880 | 1.754 | 0.307 | -16.596 | -16.371 | 24.961 | -1.864 | -17.037 | -6.243 |
| 24.201 | 10.704 | 3.464 | -9.442 | -8.659 | -7.620 | -29.031 | -22.854 | 1.391 | 7.950 | -9.029 | 0.223 | 21.647 | 3.504 |
| -17.329 | -7.879 | 25.161 | 8.691 | 5.399 | -9.307 | -11.816 | -25.277 | -1.932 | 19.306 | 23.453 | -6.506 | 17.977 | -9.906 |
| 10.029 | 12.668 | -5.689 | 5.970 | 20.976 | 5.395 | -18.117 | 9.248 | 0.807 | -0.436 | 11.924 | -2.974 | 14.100 | -10.454 |
| -24.138 | 17.723 | 16.106 | -25.614 | 16.090 | 25.526 | 5.989 | -8.606 | 3.528 | -2.293 | -6.293 | -0.156 | 22.297 | 1.526 |
| 0.188 | 10.434 | 6.331 | -24.558 | -1.146 | 24.172 | 4.526 | 11.013 | -17.854 | -4.411 | -27.848 | -21.557 | -24.706 | 0.291 |
| -24.795 | 4.555 | 17.467 | 7.557 | -25.231 | -18.184 | 27.131 | -28.240 | 4.948 | -24.783 | 5.356 | -4.058 | 24.145 | -20.819 |
| 11.600 | -9.381 | -12.425 | -21.108 | 20.824 | 21.796 | 9.072 | 5.263 | -0.681 | -1.364 | 9.821 | -6.021 | -2.557 | 10.185 |
| -4.080 | 6.718 | -16.802 | 19.894 | 10.488 | -18.924 | -0.735 | -6.842 | -7.979 | -7.026 | 26.673 | -0.978 | 26.729 | 6.184 |
| -27.246 | 1.553 | -15.073 | 10.122 | -6.906 | -0.364 | -23.124 | -17.524 | -5.132 | 1.472 | 27.297 | -0.821 | -28.203 | 0.678 |
| 10.412 | -20.949 | 0.131 | 26.492 | 16.675 | -12.418 | 4.301 | -3.984 | -10.212 | 15.006 | -6.283 | -16.055 | 11.019 | -18.593 |
| -7.021 | 18.482 | -26.008 | -23.934 | 6.629 | 4.656 | -23.719 | -24.481 | -25.023 | -21.951 | -9.147 | -4.302 | 28.545 | -0.772 |
| -28.034 | 9.325 | 9.518 | -9.207 | 19.642 | -16.162 | -15.491 | 5.868 | 5.296 | 19.239 | -6.987 | -4.992 | 27.499 | 9.104 |
| -15.297 | 15.389 | 15.139 | -25.389 | 22.698 | -21.302 | 20.122 | 25.002 | 18.341 | 2.299 | -6.765 | 0.335 | -29.572 | -6.661 |
| 1.415 | -5.219 | 14.408 | -8.508 | 20.038 | -1.887 | 4.669 | 22.950 | 10.015 | 12.983 | -19.138 | -7.597 | 30.006 | -9.200 |
| -2.167 | -2.151 | 2.999 | 8.371 | -12.263 | -9.008 | 18.145 | 12.330 | -1.477 | -9.133 | 17.841 | -2.047 | -21.414 | 7.279 |
| 14.581 | -17.154 | -6.236 | -26.236 | -12.982 | 25.796 | -22.353 | 3.097 | 23.954 | -11.769 | 26.410 | -5.352 | 15.724 | -12.215 |
| 11.089 | -14.045 | 6.500 | -23.190 | 0.257 | 10.500 | 23.259 | 24.122 | 8.267 | -13.435 | -6.163 | -13.504 | -2.107 | 0.921 |
| -10.272 | -16.003 | -17.341 | 24.166 | 11.975 | 9.556 | -0.734 | 21.944 | 0.804 | 9.816 | -13.287 | 5.554 | 25.285 | -1.490 |
| -17.840 | -3.841 | 9.338 | 17.825 | 14.583 | -23.940 | 2.013 | -14.937 | -12.076 | 17.252 | 21.003 | -23.283 | -3.181 | -8.414 |
| 17.420 | -14.863 | 12.474 | 5.215 | -7.384 | -17.188 | 22.900 | 19.427 | -3.009 | -5.855 | -1.051 | -7.047 | 20.443 | 12.641 |
| 20.722 | -15.161 | -14.228 | 3.787 | -6.614 | 4.271 | -24.455 | 8.461 | -5.748 | -8.393 | -7.046 | -8.674 | 22.569 | -19.045 |
| -17.027 | -15.552 | -21.976 | 26.931 | -22.490 | -20.719 | -17.007 | -18.509 | -7.160 | -5.007 | 29.534 | -3.594 | 18.720 | -25.961 |
| 29.678 | -7.867 | -25.000 | 2.399 | 25.704 | 7.321 | -28.080 | 26.832 | 14.711 | 5.880 | 25.698 | -12.820 | -22.545 | 4.943 |
| 18.175 | -8.897 | 16.228 | -26.609 | 10.760 | 22.160 | 4.214 | -0.822 | 4.704 | -7.588 | 4.626 | 5.524 | 26.039 | -3.634 |
| 15.426 | -19.765 | -4.644 | 25.435 | -3.949 | -4.551 | -2.326 | -12.710 | 14.079 | -26.157 | 30.189 | -0.462 | 30.864 | -20.609 |
| -25.946 | 20.860 | 23.976 | 23.276 | -15.492 | -7.649 | -27.189 | -7.702 | -14.442 | -16.363 | 28.849 | -9.545 | 28.632 | 10.901 |
| -8.165 | 8.556 | 18.695 | 0.076 | -17.293 | 20.030 | 24.621 | 23.253 | 8.793 | -22.963 | -31.287 | -18.712 | 21.777 | -6.700 |

| | | | | | | | | | | | | | |
|---|---|---|---|---|---|---|---|---|---|---|---|---|---|
| 25.155 | -4.047 | 1.842 | 4.764 | 15.028 | 6.697 | 1.399 | 27.848 | 23.337 | -20.945 | 17.572 | 11.539 | -23.713 | 16.106 |
| 18.700 | -5.306 | -11.622 | -24.663 | 10.405 | -14.945 | -20.780 | 1.874 | 1.512 | -9.020 | 1.342 | 23.826 | 30.144 | 12.299 |
| -11.788 | -20.036 | 14.203 | -4.244 | 0.685 | 24.870 | -22.874 | 27.567 | -5.323 | -0.065 | 19.172 | -2.887 | 19.786 | -13.262 |
| -14.609 | 14.739 | 1.408 | -1.943 | -11.518 | 22.571 | 0.308 | -17.986 | -23.824 | 8.653 | 20.038 | -5.687 | 29.228 | 0.623 |
| 20.690 | -6.066 | 4.256 | -3.781 | -0.952 | 16.547 | -17.037 | -22.875 | 18.267 | 7.430 | 28.886 | -7.155 | -30.832 | -8.040 |
| 17.242 | -21.623 | 5.355 | -6.664 | -4.051 | 7.048 | 23.481 | -23.420 | 12.549 | 0.748 | -5.030 | 13.011 | 17.533 | 14.223 |
| -22.454 | 18.190 | -2.635 | 19.962 | 10.051 | -13.834 | -9.744 | -24.186 | 9.766 | 4.670 | 5.453 | -10.396 | -28.223 | 1.519 |
| -4.298 | 8.819 | 15.013 | -20.793 | -4.939 | 5.608 | -4.293 | -29.596 | -4.532 | 3.568 | -6.061 | 1.184 | 26.062 | 1.829 |
| -7.159 | -0.163 | 6.122 | 22.567 | -2.488 | 6.889 | 8.576 | -29.293 | 15.300 | 6.851 | -2.392 | 19.763 | 11.251 | 3.544 |
| 28.973 | -3.378 | 18.077 | 3.754 | 7.029 | 20.997 | -20.444 | 25.567 | -0.056 | -0.783 | 1.522 | -4.447 | 20.877 | 8.587 |
| 11.029 | -13.436 | -14.085 | -3.465 | -0.388 | 2.992 | 4.649 | 2.408 | -24.439 | -10.930 | 0.420 | 24.661 | -27.224 | -1.815 |
| -5.538 | 3.882 | -1.369 | 26.207 | -17.670 | -4.459 | 5.767 | 30.474 | -25.614 | -12.196 | 26.217 | -3.674 | 18.861 | 11.178 |
| 12.782 | 9.634 | -21.011 | -4.545 | -22.480 | 21.914 | -25.484 | -14.265 | 4.027 | -12.730 | -2.201 | -6.215 | -14.853 | -1.435 |
| 1.318 | 7.213 | 24.447 | 24.341 | -12.943 | -0.115 | -8.865 | 22.957 | -18.289 | -3.224 | 18.478 | -13.268 | 28.401 | 9.391 |
| 21.572 | 15.213 | -2.291 | 25.403 | -4.775 | 12.363 | -24.191 | -23.744 | -1.500 | -24.240 | 31.106 | -6.285 | 27.205 | -8.034 |
| -17.424 | 13.524 | 8.301 | 1.817 | 1.012 | 22.242 | 23.010 | -0.302 | -0.704 | 26.420 | 4.946 | 23.566 | 2.793 | -15.010 |
| -4.665 | -1.715 | -0.217 | -23.142 | 7.533 | 9.315 | -1.374 | -3.403 | 20.712 | 3.387 | -1.427 | -22.979 | -30.347 | -3.131 |
| -29.054 | -17.930 | 1.807 | -3.838 | 21.491 | 8.711 | 26.537 | -23.305 | 12.922 | -4.195 | 23.000 | 4.112 | 30.511 | 0.104 |
| 9.599 | -11.804 | -8.475 | 7.191 | 16.484 | 3.020 | -27.517 | -17.749 | 3.566 | -5.221 | -5.284 | 1.166 | 26.435 | 6.193 |
| 23.025 | 4.588 | -7.909 | -1.327 | 6.066 | 1.903 | 27.140 | -19.306 | 25.190 | -25.456 | 18.984 | -18.406 | 22.481 | 11.290 |
| -27.782 | 0.324 | 1.340 | -0.090 | -0.721 | 4.227 | 19.909 | 21.969 | -7.934 | 7.339 | -30.787 | 3.760 | -1.929 | -11.070 |
| 30.333 | -11.973 | -6.140 | 22.471 | 13.789 | 21.692 | -15.825 | 7.353 | 10.449 | -26.806 | 2.961 | 0.744 | 24.864 | 2.308 |
| 6.203 | -20.248 | -10.669 | 1.483 | 26.115 | -2.109 | -27.967 | -18.457 | -9.652 | 2.380 | 25.741 | -6.385 | -19.297 | -18.894 |
| -8.943 | 15.794 | 2.954 | -21.382 | 13.148 | 8.586 | 21.225 | -9.642 | -5.577 | -24.218 | -4.628 | -4.957 | 29.406 | -13.159 |
| -20.805 | 21.893 | 6.329 | -13.592 | 1.986 | -3.102 | -0.587 | 17.963 | -8.887 | 5.901 | 24.170 | 0.396 | 17.101 | -14.835 |
| -18.810 | -7.412 | -1.184 | 5.593 | 5.597 | -9.487 | -15.355 | 3.316 | -0.853 | -9.124 | 24.278 | -2.586 | -18.044 | -14.219 |
| 16.714 | 23.977 | -5.800 | 6.133 | 9.051 | -19.541 | -18.272 | -29.198 | 5.315 | 23.634 | 22.919 | 2.585 | 31.156 | -18.686 |
| 21.153 | -10.484 | -2.874 | -7.924 | 22.259 | 24.007 | -25.831 | -14.765 | 13.802 | 8.633 | -7.122 | -5.356 | -22.564 | -3.875 |
| -21.520 | -2.620 | 2.526 | -0.517 | 20.642 | 13.016 | 2.412 | 25.218 | 21.649 | 3.832 | -1.924 | 3.074 | 15.722 | 6.962 |
| 5.203 | 18.654 | -20.537 | 13.336 | -18.669 | 0.986 | -2.011 | -13.998 | -18.279 | -1.783 | -6.010 | 0.701 | 19.796 | -2.725 |
| -14.742 | 13.642 | -3.110 | -22.372 | -10.973 | 0.203 | -4.271 | -6.508 | 23.966 | -0.114 | 22.963 | -18.768 | 28.406 | 10.355 |
| 13.607 | 7.723 | -8.080 | -21.820 | -11.437 | 24.218 | 3.068 | -18.933 | 7.575 | -17.824 | 24.708 | -21.620 | -28.082 | -3.219 |
| -29.725 | -8.772 | 3.458 | 17.501 | 15.391 | -4.987 | 22.039 | -28.471 | -2.654 | -6.219 | -4.935 | -2.133 | 0.538 | 1.417 |
| -6.220 | -16.537 | 14.073 | -3.381 | 1.823 | 0.746 | -26.703 | 11.277 | 0.332 | -12.148 | 22.417 | 0.484 | -11.393 | 4.357 |
| 23.887 | 2.611 | 4.780 | -19.102 | 10.259 | -7.817 | -1.331 | 26.347 | -15.298 | 3.184 | 2.566 | -0.727 | -22.793 | 2.397 |
| -12.040 | 26.820 | -0.301 | 14.327 | -20.869 | 0.286 | -20.707 | -19.734 | 3.646 | -24.620 | -4.518 | -0.189 | 7.589 | 0.169 |

| | | | | | | | | | | | | | |
|---|---|---|---|---|---|---|---|---|---|---|---|---|---|
| -15.279 | 6.166 | 4.189 | -5.630 | -10.049 | -5.086 | 9.159 | 9.887 | -17.304 | 0.132 | 26.959 | 2.523 | 11.195 | -3.345 |
| -16.345 | -6.197 | 1.760 | 1.376 | -19.514 | 4.864 | 8.771 | 28.773 | -8.532 | -0.279 | -4.308 | 9.545 | 27.076 | 16.563 |
| 27.522 | -3.886 | -21.025 | -16.087 | 23.565 | 2.011 | -3.024 | -2.172 | -6.093 | -2.248 | 28.934 | -7.670 | 25.759 | 14.704 |
| 19.849 | -9.566 | -16.401 | 4.019 | -0.422 | -16.220 | 2.066 | -20.665 | -22.018 | -13.670 | 28.132 | -5.553 | -10.799 | -4.822 |
| 29.195 | 4.809 | 7.680 | -18.260 | 21.299 | 10.110 | 3.368 | -21.989 | 0.582 | 2.528 | 24.471 | 5.295 | 27.921 | 10.799 |
| -18.572 | 19.286 | 5.697 | -2.262 | 20.201 | -8.276 | -26.799 | 31.594 | -5.073 | -1.247 | 20.153 | 6.132 | 24.500 | -15.759 |
| 30.648 | 17.159 | -1.680 | -0.451 | 22.328 | -15.221 | 24.967 | -12.814 | 16.960 | 16.913 | 28.693 | -3.328 | 15.612 | -2.688 |
| 7.269 | 22.296 | 1.983 | 11.161 | -24.382 | 0.539 | -22.929 | -5.459 | -22.834 | -4.594 | 5.079 | 22.383 | -29.010 | 5.486 |
| -13.691 | -19.688 | -5.385 | -2.100 | 21.949 | -2.957 | -9.712 | -13.515 | 4.640 | -15.871 | 3.342 | 9.248 | -1.328 | -18.248 |
| 4.068 | -21.209 | -9.263 | 13.589 | 2.151 | 6.433 | 7.947 | -27.312 | -25.452 | -10.096 | 20.893 | 24.477 | -21.337 | 7.015 |
| 17.976 | -1.809 | 4.207 | 0.428 | 27.154 | 24.178 | -26.829 | -30.385 | 26.268 | 20.481 | 27.252 | -11.759 | 17.897 | -5.619 |
| -4.593 | 13.113 | 24.365 | 8.920 | -0.794 | 23.168 | -25.286 | 29.662 | 22.270 | -11.047 | 21.448 | -22.391 | 21.065 | -8.503 |
| 25.784 | -17.893 | 18.767 | -8.125 | -24.346 | 17.652 | -2.686 | 27.604 | 10.555 | 6.144 | 3.566 | -16.090 | 14.590 | 17.939 |
| -27.976 | -25.851 | 7.593 | 8.108 | 25.428 | 26.868 | 3.577 | 3.353 | 9.707 | -7.251 | -1.913 | -0.444 | -2.732 | -7.195 |
| -28.875 | 20.719 | -2.719 | -2.441 | 15.794 | 0.844 | 2.937 | 30.852 | 13.810 | 0.509 | -2.817 | 3.398 | 26.067 | -1.284 |
| -15.945 | 8.537 | -4.382 | -9.117 | -14.588 | 6.520 | -26.606 | 1.651 | 3.835 | 4.923 | 23.016 | -23.865 | 30.032 | 12.522 |
| 6.344 | -16.000 | -12.504 | 14.718 | -17.955 | -16.651 | -22.036 | 20.599 | 6.476 | -6.508 | 25.351 | -4.493 | 15.028 | 19.323 |
| 27.486 | 4.436 | -0.564 | -0.013 | 18.325 | 18.620 | -3.823 | 3.092 | 1.185 | 17.764 | 11.952 | 25.842 | 29.501 | 0.435 |
| -30.886 | 11.807 | -6.279 | -18.053 | 16.447 | -0.989 | -6.489 | -4.256 | -0.616 | -20.819 | 3.234 | -12.953 | 26.905 | -0.840 |
| -3.755 | 10.413 | -19.063 | -7.360 | 4.017 | 19.398 | -18.215 | -8.590 | -2.014 | -5.451 | 9.771 | -27.291 | 16.730 | -19.122 |
| -14.517 | -6.053 | 17.260 | 2.834 | -25.142 | 27.590 | -7.752 | 23.232 | 9.058 | -19.572 | 25.138 | -24.853 | 25.022 | -24.455 |
| -2.594 | 1.851 | -3.133 | -19.012 | 21.919 | -19.020 | 6.074 | 11.984 | -14.826 | -16.468 | 23.510 | -16.718 | 1.976 | 17.194 |
| 18.398 | -9.175 | 6.384 | 22.645 | -12.178 | -1.383 | -23.526 | -26.575 | -1.708 | -15.822 | 7.824 | 3.701 | 27.708 | 15.035 |
| 3.452 | 16.240 | 2.554 | -18.471 | -20.448 | -21.925 | 18.966 | -5.207 | -24.154 | 1.625 | 6.710 | 0.379 | 22.126 | -3.053 |
| -4.406 | 21.107 | 4.294 | 12.087 | 5.014 | 0.084 | 3.604 | 9.720 | -27.988 | -13.982 | 25.003 | 3.544 | 20.443 | -9.810 |
| -29.677 | 14.411 | 9.121 | 22.392 | 8.643 | -2.332 | 16.217 | -12.051 | -20.143 | -1.537 | -3.655 | -17.442 | 18.125 | -12.153 |
| -12.163 | 9.881 | -0.127 | 21.323 | -21.334 | -5.372 | 16.887 | -3.335 | 0.498 | 0.107 | -6.532 | -10.393 | -27.733 | -4.600 |
| 28.899 | 18.580 | 6.884 | 6.410 | -26.790 | 28.538 | -25.068 | -20.278 | 17.988 | -0.783 | 0.754 | 3.146 | 23.970 | 14.239 |
| 15.126 | -10.787 | -4.659 | 7.243 | 15.717 | 20.800 | -2.184 | 24.876 | -3.802 | -15.497 | -0.002 | -6.274 | -1.698 | -1.926 |
| 24.693 | -13.382 | 17.430 | 5.187 | -1.811 | 22.987 | -7.362 | -0.471 | -5.995 | -6.131 | 2.703 | 5.852 | -23.354 | 3.963 |
| 17.408 | 21.542 | 4.420 | 10.835 | -8.780 | -22.788 | 5.091 | 8.672 | 21.966 | -8.050 | -2.745 | 3.705 | 0.996 | -3.503 |
| 5.647 | 7.984 | 5.829 | -19.965 | -5.370 | -11.233 | -8.188 | 23.882 | -7.725 | -1.062 | 4.063 | -12.850 | 11.958 | -8.852 |
| 18.817 | 19.570 | -1.510 | -3.091 | -18.821 | 25.989 | 3.171 | 21.109 | 5.097 | -7.798 | -0.661 | 3.292 | -30.166 | -1.154 |
| 12.092 | 5.440 | -1.924 | 13.675 | 21.004 | -24.296 | 7.096 | 11.413 | -18.073 | 2.196 | 23.481 | -5.696 | -26.719 | -5.962 |
| -26.153 | -0.589 | -21.928 | 12.793 | 26.283 | -26.926 | -12.823 | 27.283 | 11.363 | 5.259 | -2.501 | 6.747 | 18.833 | 16.349 |
| 20.685 | 11.937 | -6.584 | 6.102 | 10.683 | -24.997 | 25.110 | 25.646 | -22.069 | 6.341 | 2.970 | 5.484 | 21.959 | 24.156 |

| | | | | | | | | | | | | | |
|---|---|---|---|---|---|---|---|---|---|---|---|---|---|
| -13.763 | 15.300 | -0.938 | -6.355 | 21.881 | 0.218 | -9.020 | 17.364 | 27.019 | -10.785 | 23.514 | -4.995 | 19.404 | -5.150 |
| 11.032 | 7.312 | 5.765 | -3.922 | 18.989 | 5.084 | -20.364 | -17.232 | 9.273 | -14.552 | 13.277 | -23.676 | -27.418 | 2.440 |
| 0.712 | 16.987 | 1.072 | -18.996 | 5.945 | 1.589 | -6.844 | 21.824 | -10.543 | -16.723 | 19.533 | 0.696 | 28.041 | 10.271 |
| -19.334 | -20.315 | 23.807 | -23.268 | 0.886 | -17.063 | -14.016 | -12.389 | 7.914 | -12.357 | 8.029 | -8.177 | 18.001 | 12.819 |
| -4.388 | -14.601 | -3.332 | -9.774 | 25.052 | 15.308 | -17.885 | -6.365 | 16.803 | 10.818 | 16.512 | -18.250 | -12.561 | 3.689 |
| -9.365 | 22.820 | -12.617 | 3.381 | 19.742 | 27.952 | -9.377 | 17.936 | 0.666 | 8.100 | 17.105 | 0.961 | 23.657 | 9.000 |
| 6.253 | 6.606 | -5.853 | -3.727 | -22.423 | 29.772 | 11.425 | 3.260 | -14.214 | 0.638 | 0.230 | -27.489 | 13.275 | -1.884 |
| -8.110 | 20.738 | 27.393 | 13.835 | 13.570 | 29.087 | 4.375 | -22.827 | 5.579 | 19.987 | -24.930 | -25.172 | 29.117 | -9.316 |
| 2.209 | 9.324 | 3.411 | -1.193 | 9.216 | 2.840 | 18.602 | 23.425 | -8.799 | 24.659 | -18.184 | -0.554 | 28.432 | 27.050 |
| -5.821 | -7.888 | 4.225 | 16.495 | 4.227 | -3.903 | -10.987 | 7.999 | -23.483 | 0.355 | 1.243 | 7.592 | 22.534 | 6.452 |
| -3.851 | -13.013 | -5.231 | 23.835 | -19.261 | 4.099 | -3.395 | 22.743 | 10.134 | -9.242 | 8.717 | 28.600 | -4.290 | -8.033 |
| 26.797 | 4.305 | 14.939 | 26.686 | 11.720 | -7.068 | -15.715 | -19.578 | 16.499 | 22.324 | 4.402 | -6.034 | 14.231 | 8.474 |
| -20.917 | 22.392 | 21.952 | 21.786 | -1.755 | -19.129 | -14.946 | -31.082 | -9.836 | -1.758 | -1.236 | -9.960 | 27.021 | 3.365 |
| -7.225 | -5.605 | -17.344 | -16.045 | 17.289 | -15.617 | 9.597 | 2.753 | 9.744 | 9.348 | -0.795 | -22.137 | 20.921 | 4.004 |
| -29.771 | 6.192 | 5.046 | -22.681 | -4.959 | 21.214 | -21.591 | 30.197 | -11.452 | -21.556 | 23.660 | 4.219 | 25.916 | 6.484 |
| 11.560 | -1.711 | -8.051 | 2.635 | 27.068 | 1.280 | -2.303 | -28.792 | -22.671 | -1.232 | -24.872 | 29.222 | 11.809 | 21.401 |
| 14.552 | -2.684 | -0.061 | 8.436 | -14.397 | 5.078 | 2.186 | 1.266 | 3.702 | -8.765 | -0.587 | 25.748 | 21.221 | 25.415 |
| 22.880 | 21.406 | 26.023 | -3.386 | -4.513 | 21.602 | 11.140 | -3.892 | -12.786 | -15.438 | 11.371 | -28.903 | 0.178 | 10.718 |
| 2.875 | 6.602 | -5.718 | 5.333 | -18.569 | -5.390 | -17.396 | 22.523 | 27.504 | 3.460 | 14.540 | -8.194 | 20.270 | 4.557 |
| -24.923 | 24.829 | -22.387 | 6.449 | -12.008 | 5.303 | -7.388 | 21.061 | -25.221 | 9.977 | -22.789 | 16.322 | -17.863 | -6.902 |
| 29.473 | -13.426 | -17.446 | -8.581 | -19.622 | 28.211 | 24.206 | -15.456 | 6.177 | 3.981 | -1.130 | 6.229 | 4.851 | -0.196 |
| 17.976 | -25.615 | -27.879 | 21.690 | 15.230 | -8.438 | -19.961 | -20.469 | 14.591 | 4.433 | 24.317 | 4.095 | 28.990 | -0.940 |
| 1.927 | 10.147 | 24.994 | -19.614 | 22.260 | -21.665 | -9.235 | 26.375 | -25.719 | 2.496 | -9.032 | -6.265 | 1.786 | -2.941 |
| -16.512 | 0.975 | 24.125 | 19.799 | 27.780 | -3.733 | 25.973 | -16.777 | 6.398 | 0.012 | 20.997 | 9.571 | -1.106 | 24.005 |
| 21.320 | -10.583 | 27.904 | -24.976 | -7.473 | 10.295 | 21.316 | 17.082 | 12.453 | -4.257 | -1.329 | 13.788 | 12.110 | 18.388 |
| 4.372 | 21.851 | 15.909 | -26.514 | -5.871 | -0.602 | 7.957 | 0.460 | -4.426 | -4.615 | 23.477 | 7.155 | -1.720 | 8.783 |
| 17.284 | 2.832 | -7.779 | 25.547 | 25.421 | -22.141 | 5.499 | -19.185 | -3.323 | -15.079 | 0.193 | -6.727 | 14.517 | -4.187 |
| 2.332 | -5.066 | 26.517 | 24.115 | 4.475 | 9.891 | 17.757 | 21.732 | 5.362 | 3.085 | -2.766 | 19.675 | 24.294 | -9.063 |
| -9.140 | 13.352 | -6.793 | 1.147 | -21.854 | 12.568 | -24.806 | 15.146 | 28.420 | 2.431 | -13.719 | 8.248 | -28.897 | 5.698 |
| -14.976 | 1.737 | 16.274 | 29.158 | 5.904 | -19.649 | -19.168 | 5.366 | -7.655 | -8.793 | -15.680 | 23.367 | 8.749 | 0.530 |
| -19.314 | -14.629 | 17.372 | 2.503 | 14.423 | -26.780 | 17.632 | 23.255 | -22.380 | -9.872 | 4.245 | 31.239 | -19.006 | -5.219 |
| 13.998 | -1.633 | -15.857 | 14.053 | 6.927 | 26.790 | 2.067 | -10.262 | 15.141 | -16.216 | -1.860 | -2.885 | 25.591 | 13.269 |
| -13.570 | -6.598 | 14.433 | -27.934 | 21.635 | 5.267 | 0.616 | -23.335 | -6.864 | -10.737 | -1.616 | 11.122 | -23.191 | 7.539 |
| 1.606 | 22.416 | 3.628 | -6.018 | 10.518 | 31.793 | -25.430 | 22.332 | -25.678 | -6.425 | -2.834 | -2.035 | 4.298 | 18.672 |
| 24.624 | 15.057 | 22.874 | -26.022 | -21.734 | -29.780 | 3.955 | 13.008 | 6.443 | 15.136 | 2.169 | -2.301 | 22.824 | -19.529 |
| -17.390 | -9.907 | 1.259 | 19.322 | -7.002 | 10.753 | 19.759 | -5.074 | 3.706 | 7.531 | 9.395 | 15.898 | 21.720 | 8.808 |

| | | | | | | | | | | | | | |
|---|---|---|---|---|---|---|---|---|---|---|---|---|---|
| -19.030 | 8.352 | 17.409 | -19.343 | -4.313 | -8.159 | -2.709 | 12.352 | -26.561 | -17.853 | 1.706 | -6.993 | 18.715 | -8.184 |
| 3.601 | -1.127 | 24.814 | 19.893 | -5.105 | -6.861 | -0.951 | 29.086 | -14.934 | 20.205 | 20.217 | -20.319 | 9.546 | 23.353 |
| -18.735 | -2.233 | 21.617 | 0.365 | 10.813 | -7.492 | -4.876 | -29.149 | 17.324 | -13.461 | 7.560 | 23.340 | 11.278 | 4.960 |
| -19.828 | 11.679 | 26.381 | -5.877 | -22.643 | 29.644 | -20.736 | 25.068 | -1.390 | -14.926 | 2.244 | -2.059 | 24.073 | -3.434 |
| -4.398 | -7.348 | 24.614 | -10.615 | 12.097 | 4.850 | 25.022 | 10.576 | 16.674 | -9.621 | 21.262 | 5.235 | 19.555 | -0.656 |
| -8.340 | 17.429 | -4.780 | -21.174 | 23.300 | -7.047 | -5.776 | 21.986 | 5.092 | -0.193 | 4.863 | -7.485 | 22.009 | -13.222 |
| -21.860 | -8.264 | -6.910 | 19.850 | 18.752 | 28.051 | -23.348 | 0.350 | 19.461 | -18.952 | -3.353 | 4.705 | -27.567 | 10.463 |
| -29.629 | 7.018 | 5.293 | -3.030 | -26.843 | 3.149 | 24.536 | 3.797 | 14.837 | 12.341 | 1.143 | -7.704 | -26.319 | -21.410 |
| 0.776 | -21.365 | -1.547 | 23.473 | 20.252 | -4.658 | -25.971 | -3.791 | -26.945 | 6.246 | -0.336 | -11.786 | -16.569 | 3.350 |
| 19.772 | 21.423 | -5.104 | -9.724 | 19.227 | -19.527 | -1.368 | -3.200 | 10.356 | -6.705 | 24.116 | 8.665 | 25.026 | 11.573 |
| -17.021 | -8.547 | 4.549 | 3.720 | 26.203 | 23.503 | -25.479 | 23.670 | 21.099 | 27.084 | 23.367 | 27.219 | 27.942 | 23.444 |
| 17.853 | -11.299 | 25.023 | 16.043 | 27.699 | -11.510 | 0.104 | 12.133 | 9.739 | 12.721 | 3.865 | 3.805 | 26.213 | -24.775 |
| -5.864 | -5.133 | -23.749 | -8.398 | -9.841 | -2.655 | -6.363 | 29.589 | -13.443 | 13.865 | -1.641 | 9.804 | -23.513 | 22.234 |
| -7.828 | 6.026 | 24.076 | -10.524 | -28.056 | -6.463 | 23.236 | 4.916 | -17.911 | -12.976 | 5.973 | 4.954 | 3.384 | -13.874 |
| 28.572 | 12.770 | 1.153 | 11.592 | -21.634 | 7.366 | 20.965 | -22.365 | -9.237 | -19.426 | 16.623 | -10.635 | -12.643 | -1.394 |
| -7.172 | -7.875 | 22.424 | 22.612 | 9.108 | -6.765 | 2.093 | 24.702 | 13.597 | -10.080 | -1.037 | 22.576 | -21.692 | 25.186 |
| 29.474 | -3.871 | -30.495 | -27.854 | 28.993 | 30.767 | 3.672 | 7.149 | -12.070 | 19.761 | 25.616 | 7.067 | 15.193 | 2.928 |
| 21.886 | -0.349 | 31.728 | 27.640 | -3.577 | 26.045 | -2.798 | -2.763 | 3.851 | 7.982 | 25.010 | 5.714 | -18.232 | -2.333 |
| -19.585 | -8.920 | 3.801 | 21.537 | 23.660 | 22.573 | -9.564 | 9.827 | 12.862 | 2.813 | -22.481 | -18.146 | 20.812 | 2.423 |
| -28.467 | -7.094 | -4.752 | 25.357 | -18.437 | 4.506 | 25.944 | -21.374 | -1.154 | 10.532 | 2.318 | 6.318 | 25.393 | 26.941 |
| 19.286 | 9.988 | 24.064 | 18.252 | -10.088 | -3.233 | -3.775 | -25.237 | -23.773 | -6.417 | 23.284 | 1.759 | 22.605 | -22.712 |
| -21.405 | 0.708 | 18.880 | -4.628 | 20.994 | 29.079 | 0.897 | 9.300 | 4.976 | -5.636 | -11.815 | 6.755 | -26.095 | 5.357 |
| -25.353 | -1.956 | 3.030 | 24.316 | 17.821 | 24.288 | -4.425 | -6.475 | -5.027 | 9.227 | 21.942 | 10.435 | -12.375 | 4.113 |
| -26.992 | 20.798 | -31.642 | -1.051 | -17.607 | 29.055 | 24.806 | 24.059 | -2.202 | -22.091 | -1.575 | -5.043 | 21.341 | 10.279 |
| -5.730 | 19.321 | -8.678 | -9.634 | -25.691 | 26.172 | 17.681 | 3.106 | -1.883 | -13.215 | -4.487 | 10.303 | 6.612 | 11.421 |
| -22.878 | -4.110 | 17.092 | 15.130 | -8.488 | 2.728 | 20.274 | -27.142 | -7.723 | 11.652 | 2.576 | -3.830 | 1.151 | -15.185 |
| 5.897 | 11.731 | -1.480 | 3.562 | 21.278 | -26.391 | -13.656 | -20.808 | 23.764 | -20.296 | 13.858 | -23.787 | -5.485 | 3.097 |
| 24.466 | 4.203 | 5.889 | -5.129 | -15.275 | -3.363 | -18.784 | 26.791 | -3.513 | 26.638 | -3.260 | 22.557 | 14.930 | 20.758 |
| 28.325 | 18.726 | 2.715 | -24.390 | -23.264 | 0.614 | 1.452 | 23.620 | 1.881 | 18.047 | 27.442 | 3.277 | -13.361 | 5.685 |
| 17.310 | 6.518 | 21.742 | 3.125 | -14.164 | 28.229 | -10.302 | -8.624 | -20.637 | 5.842 | -0.523 | 28.171 | 26.566 | -6.041 |
| -3.011 | 4.781 | 3.512 | -29.349 | 8.164 | -3.385 | -3.226 | 14.905 | 12.350 | 0.043 | 6.366 | 4.051 | 25.823 | 17.764 |
| -14.554 | -3.661 | 13.149 | 25.353 | 23.245 | 16.017 | -22.051 | -10.737 | -0.632 | 23.752 | 24.372 | -7.512 | 13.166 | -18.471 |
| 20.287 | 0.176 | -4.798 | 23.500 | 16.959 | 2.791 | -0.069 | 24.575 | 25.026 | 12.694 | 8.029 | 22.554 | -10.010 | 17.319 |
| 14.786 | -4.978 | -7.693 | 22.784 | 18.267 | -5.433 | -19.905 | 28.748 | 11.991 | 18.430 | 1.791 | -24.295 | 24.399 | 24.479 |
| 11.220 | -4.333 | 29.821 | 27.855 | -29.456 | -24.452 | -26.688 | 22.870 | 17.600 | 9.790 | -2.456 | 5.500 | 13.747 | -1.264 |
| -4.796 | -11.636 | -5.061 | 14.437 | 15.094 | -3.468 | 23.609 | 2.540 | 17.954 | -4.065 | -10.844 | -24.593 | 6.038 | 28.453 |

| | | | | | | | | | | | | | |
|---|---|---|---|---|---|---|---|---|---|---|---|---|---|
| -3.339 | 3.304 | -25.901 | -23.077 | 16.257 | -1.436 | -4.653 | -13.921 | -20.842 | 13.777 | -12.636 | -21.639 | 23.608 | 20.158 |
| -25.214 | 2.662 | -3.227 | 28.334 | -21.349 | -2.305 | 28.030 | -23.218 | -13.463 | -8.459 | -16.736 | 24.638 | 22.061 | 11.626 |
| 13.507 | -5.883 | 22.359 | -2.810 | -4.170 | -2.850 | 18.666 | 25.768 | 19.585 | -24.153 | 1.187 | 21.998 | -1.759 | 5.279 |
| -6.761 | 1.534 | 19.537 | -27.041 | -8.821 | 0.451 | -8.388 | -2.881 | 21.709 | -26.028 | 0.876 | 0.531 | -15.329 | -6.821 |
| -11.044 | -3.990 | 23.209 | 1.725 | 15.504 | -6.624 | -22.093 | 15.951 | -17.616 | -13.873 | -28.278 | -1.251 | 27.235 | 6.963 |
| 1.971 | 15.855 | -29.417 | -19.965 | -8.044 | 24.018 | -18.939 | -11.616 | -27.538 | 5.541 | 28.199 | -2.042 | 19.451 | -5.257 |
| -23.720 | -3.117 | -8.492 | 3.677 | -14.716 | -26.677 | -25.522 | 26.127 | -14.057 | -15.604 | -6.105 | 3.990 | 20.881 | -6.431 |
| 26.184 | -3.554 | 25.969 | -0.142 | -24.029 | -9.448 | -3.112 | 27.244 | 24.117 | 11.472 | 5.946 | 10.101 | -3.776 | 21.803 |
| 8.573 | -7.017 | -4.512 | -13.568 | 21.904 | -1.507 | 6.112 | -29.916 | 12.950 | -14.307 | -10.682 | 21.425 | -6.753 | -21.707 |
| -18.981 | 4.129 | 27.354 | 24.926 | -6.828 | 21.465 | -5.277 | 23.940 | 6.073 | 7.659 | 2.773 | -11.470 | -8.801 | -7.693 |
| -20.510 | -0.437 | -2.904 | -9.657 | 19.408 | -3.323 | 3.561 | 24.755 | -21.482 | 21.079 | 8.398 | -1.535 | 26.048 | 1.917 |
| -0.312 | 6.465 | 10.659 | -29.220 | 9.344 | -3.513 | -27.100 | 14.036 | 22.494 | 21.308 | 27.940 | 14.303 | -16.818 | -28.272 |
| -16.696 | 14.015 | 3.750 | -7.880 | -18.999 | -0.716 | -10.481 | -11.950 | 11.542 | -10.221 | -4.906 | 24.246 | 8.546 | 23.080 |
| -10.457 | -5.774 | 6.391 | 16.431 | -17.923 | 2.290 | 1.522 | -17.105 | -7.974 | -5.840 | 6.128 | -7.779 | -5.078 | 20.261 |
| 27.831 | 8.630 | 10.369 | -30.730 | -25.044 | 15.741 | -17.482 | 12.724 | -24.910 | -17.649 | 9.480 | -1.060 | 17.222 | 10.280 |
| 17.476 | 10.858 | -2.636 | 28.333 | 26.381 | -28.447 | -6.258 | 2.777 | 13.227 | -22.151 | 12.150 | 22.723 | 22.260 | 11.503 |
| 6.728 | -17.851 | 26.060 | 27.064 | -20.157 | -1.560 | -13.317 | 1.676 | -8.702 | 9.287 | 22.406 | 1.979 | 9.451 | 5.703 |
| -28.190 | 5.128 | 17.949 | 20.272 | -27.489 | 1.759 | 0.625 | 14.287 | 30.201 | -4.882 | 5.445 | -6.964 | 24.064 | -15.125 |
| -10.895 | -1.306 | 4.558 | 6.207 | -13.489 | -9.368 | -4.030 | 28.020 | -10.402 | -23.178 | 28.096 | -1.371 | 7.852 | 4.221 |
| -28.760 | 0.782 | -1.052 | 23.808 | 24.996 | 22.481 | -12.639 | -20.458 | -22.408 | 6.320 | 23.820 | -0.561 | 19.878 | 15.914 |
| -21.433 | -4.622 | -18.584 | 6.918 | 11.192 | 26.354 | -27.370 | -26.374 | -13.848 | 28.291 | 2.841 | -8.560 | -2.494 | -13.345 |
| -22.674 | 5.710 | 10.573 | -28.692 | 26.159 | -1.691 | 2.442 | 22.991 | -28.231 | 19.250 | 20.171 | -8.000 | -12.653 | 26.142 |
| 9.544 | -0.035 | 0.484 | -21.685 | 14.527 | -0.931 | -26.845 | 19.246 | -29.441 | 14.513 | 22.398 | -17.790 | 21.784 | 17.952 |
| 18.137 | 11.453 | 20.135 | -5.737 | -17.880 | 2.321 | 8.334 | 25.949 | 12.219 | -11.107 | 19.516 | 4.294 | 17.215 | 17.341 |
| 5.806 | -6.040 | 14.530 | 19.823 | -14.528 | 0.535 | -8.322 | -6.045 | -12.069 | -9.437 | 27.631 | 21.541 | 24.651 | -3.569 |
| -20.664 | -4.960 | -2.667 | 0.320 | 27.952 | 19.981 | 28.333 | 13.953 | -27.973 | -17.380 | 24.328 | -5.781 | 23.924 | 3.695 |
| 26.838 | -7.736 | -4.572 | -1.190 | -17.898 | 6.164 | -24.238 | -26.949 | -26.159 | 6.289 | -14.312 | -25.709 | 25.705 | -1.165 |
| 7.392 | -9.904 | 13.755 | -22.910 | 21.898 | -21.768 | -0.284 | -0.550 | -19.908 | 7.850 | 21.432 | -23.102 | 3.524 | -4.412 |
| -4.780 | -2.788 | 22.580 | 8.269 | 10.721 | 17.271 | -2.437 | -23.490 | 23.709 | 5.767 | 3.631 | 15.162 | 3.922 | 26.906 |
| 1.546 | 1.432 | 23.249 | -19.483 | -25.253 | -1.517 | -1.161 | -18.131 | 29.661 | -2.275 | 0.881 | 5.791 | 2.174 | 4.264 |
| -0.034 | 0.593 | -0.066 | 18.914 | 19.500 | 1.419 | -28.428 | 24.167 | 21.366 | -20.377 | 18.235 | 3.545 | -23.183 | -24.412 |
| -18.418 | 12.627 | -0.376 | 2.749 | -23.820 | -24.194 | 25.142 | 27.710 | 18.225 | -1.029 | 23.875 | 25.512 | -24.154 | 5.769 |
| 8.717 | 3.712 | -17.731 | 5.915 | 22.504 | 0.816 | -1.895 | 17.277 | -10.947 | -9.967 | 26.874 | -16.908 | -9.050 | -16.512 |
| -9.009 | 9.748 | -13.279 | 0.125 | -14.514 | 17.856 | -24.538 | 24.794 | -23.991 | 21.567 | 23.053 | 25.937 | 9.861 | 27.751 |
| -9.964 | -5.473 | -4.564 | -10.670 | -22.100 | -25.229 | 28.616 | 20.786 | 11.760 | 20.509 | -5.198 | 16.817 | 22.757 | 17.636 |
| -17.868 | 2.443 | 2.106 | -29.335 | -18.968 | 2.522 | -25.761 | 15.161 | 6.142 | -25.466 | 24.048 | -0.835 | -13.252 | 14.193 |

| | | | | | | | | | | | | | |
|---|---|---|---|---|---|---|---|---|---|---|---|---|---|
| -27.618 | -10.178 | 26.040 | -3.728 | -28.602 | -5.112 | 26.047 | 22.399 | 10.224 | -14.704 | 6.405 | -8.615 | 18.536 | 23.392 |
| -12.997 | -2.939 | 2.938 | -30.730 | 7.344 | -1.055 | 17.782 | -18.770 | -7.832 | 18.809 | 19.146 | -5.990 | -8.030 | -1.305 |
| -7.895 | -6.610 | 16.608 | -19.578 | -21.030 | -2.701 | 1.042 | 0.343 | 8.922 | -19.705 | 30.950 | -7.206 | -25.780 | -9.992 |
| -16.152 | -9.113 | 18.298 | 9.175 | 11.555 | -25.114 | 21.472 | -4.442 | 6.735 | 6.076 | 16.973 | -24.959 | 23.293 | -27.323 |
| 25.154 | 1.868 | 13.390 | -27.423 | 9.511 | -6.501 | -5.248 | 0.668 | 2.861 | -8.348 | 6.518 | 15.674 | 14.824 | -7.027 |
| 16.303 | 7.621 | -2.218 | 2.570 | 20.363 | 4.487 | 14.861 | -25.932 | -14.714 | -25.825 | 3.099 | -14.399 | -2.656 | 10.481 |
| -22.151 | -1.051 | 1.185 | -30.516 | 21.490 | 7.796 | -22.053 | -15.481 | 22.168 | -4.970 | 24.129 | -17.766 | -10.045 | -18.312 |
| 23.639 | 4.335 | 5.500 | 21.412 | -20.324 | -6.072 | -28.697 | -20.118 | -1.233 | -21.624 | 14.353 | 1.614 | -18.868 | 8.581 |
| -10.744 | 17.945 | -1.387 | 17.257 | -25.770 | 24.057 | -18.119 | -3.650 | 29.141 | 19.253 | 0.973 | -15.503 | -24.428 | 3.598 |
| -23.979 | -2.661 | 4.610 | -17.440 | 16.070 | 26.909 | 5.162 | 9.136 | 10.973 | -28.487 | 26.907 | 21.562 | 2.916 | -28.642 |
| 9.713 | -4.209 | -10.615 | 1.778 | -9.982 | 1.417 | 5.849 | -21.444 | -9.388 | -20.395 | -4.073 | -4.962 | 22.116 | -11.611 |
| 21.581 | 1.142 | 13.633 | 0.358 | -23.873 | 16.198 | 20.276 | -6.713 | -11.045 | -21.246 | -17.201 | -10.944 | 20.168 | 27.134 |
| -0.425 | -0.331 | 1.632 | -15.342 | -23.081 | -13.960 | 26.446 | 25.453 | 17.181 | 5.621 | -6.272 | -2.689 | -11.772 | 20.157 |
| 21.310 | -8.339 | -1.351 | 5.173 | -24.259 | 17.567 | -4.790 | -21.811 | -23.005 | -13.990 | 5.666 | -2.115 | -21.209 | 4.955 |
| -11.488 | -6.032 | -5.951 | 27.909 | 26.252 | 1.622 | -26.616 | 2.192 | -12.805 | -12.418 | 26.558 | -0.359 | 2.558 | -21.925 |
| 10.349 | -5.054 | 9.952 | 18.146 | -11.827 | -0.858 | -29.366 | -18.266 | 17.981 | 25.868 | 6.776 | 11.379 | 8.369 | -13.105 |
| 8.220 | 15.929 | -22.318 | -11.643 | 6.447 | 20.596 | 29.343 | -27.532 | 4.309 | 8.048 | 0.532 | 18.468 | 16.856 | 7.170 |
| -24.886 | -9.746 | -16.550 | -1.759 | -27.958 | -2.058 | 28.497 | -14.428 | -32.265 | -13.213 | 28.582 | 15.737 | 18.383 | 11.230 |
| 2.219 | 15.209 | -20.098 | 25.308 | -20.563 | 26.639 | -19.140 | -22.937 | -16.360 | 28.995 | -0.892 | 24.433 | 16.969 | 30.166 |
| 24.404 | -2.287 | -14.608 | -5.574 | -18.990 | 23.008 | -6.560 | -7.322 | 10.856 | -7.799 | -3.617 | -5.786 | -11.357 | -7.215 |
| -7.207 | 11.987 | -17.873 | 17.567 | 13.003 | 19.646 | 23.174 | -26.328 | 7.093 | -11.481 | -7.376 | 19.019 | -12.135 | -8.694 |
| 18.724 | 2.033 | -4.061 | 1.924 | -15.916 | 11.622 | 25.933 | 12.709 | 22.426 | -28.112 | 2.477 | -16.296 | -2.508 | -0.835 |
| 17.703 | 0.519 | 2.125 | 8.110 | -19.813 | 1.031 | -24.219 | -28.085 | -29.843 | 17.582 | 25.559 | 17.719 | 13.608 | 21.539 |
| 4.559 | 5.029 | -0.471 | -29.899 | 8.646 | 3.041 | -0.192 | -13.025 | -30.502 | 8.454 | 14.498 | 0.347 | 4.928 | -4.582 |
| 19.335 | -1.555 | -1.325 | -1.962 | 17.244 | 20.521 | 27.032 | -22.234 | -6.964 | 26.108 | 9.783 | 2.088 | -10.843 | 5.174 |
| 27.860 | -5.933 | 1.566 | -15.681 | 20.786 | 10.974 | 0.466 | 23.471 | -2.289 | 23.193 | 2.617 | -16.555 | 23.147 | 15.100 |
| -27.760 | -4.128 | 1.936 | -27.048 | 7.180 | -1.219 | -28.121 | -15.207 | -28.007 | 11.297 | 7.696 | 25.476 | 15.571 | -3.370 |
| 23.880 | -2.757 | -20.172 | 21.665 | -18.748 | 6.914 | -5.087 | -18.199 | 18.976 | 13.664 | 6.734 | 19.598 | 4.334 | -5.996 |
| 1.327 | -2.083 | 2.586 | 23.696 | -17.992 | 13.463 | -26.075 | -16.161 | 11.278 | 3.287 | 18.065 | 7.947 | 20.845 | 20.642 |
| -2.990 | -2.543 | 16.420 | 3.297 | 11.364 | 25.021 | -29.948 | -12.741 | -28.613 | 7.132 | 6.176 | 2.684 | -15.349 | 17.827 |
| -25.892 | -1.337 | 15.776 | 20.445 | 21.650 | 21.457 | 27.570 | 18.417 | -29.036 | 5.944 | -0.861 | 22.003 | 20.182 | -23.874 |
| -1.699 | -5.721 | -0.280 | -28.975 | 12.324 | 15.889 | -2.308 | 8.239 | -3.301 | -27.641 | 21.799 | -21.284 | 25.905 | -30.190 |
| -25.682 | 3.055 | -1.645 | 6.691 | -13.603 | 3.124 | 25.040 | -19.862 | 7.172 | -19.915 | 7.672 | 18.404 | -17.093 | -29.798 |
| 9.253 | 13.823 | 16.924 | -7.771 | 19.538 | -3.629 | 20.334 | 24.454 | -1.219 | -3.953 | 22.750 | -13.793 | -20.250 | 25.328 |
| -11.796 | -11.364 | 7.217 | 9.189 | 10.000 | -22.537 | 1.683 | -14.241 | 28.508 | -21.560 | -23.634 | -11.661 | -15.238 | -3.750 |
| -19.344 | -14.644 | 1.425 | 4.217 | 15.261 | -23.352 | -19.006 | -9.592 | -18.846 | 10.223 | 21.974 | 10.769 | -21.800 | 4.107 |

| | | | | | | | | | | | | | |
|---|---|---|---|---|---|---|---|---|---|---|---|---|---|
| 3.680 | 5.901 | 22.686 | -1.436 | 12.452 | 23.203 | 1.302 | 22.429 | 27.561 | -6.704 | 17.226 | 0.896 | 2.285 | 15.860 |
| -15.925 | -0.905 | 5.608 | 25.561 | 17.859 | -6.671 | -0.143 | 19.866 | 20.741 | 27.161 | 6.853 | -23.275 | 23.859 | 3.194 |
| 5.758 | -3.150 | -2.824 | -21.098 | 5.356 | 15.152 | -3.198 | 21.705 | -9.042 | -0.403 | -0.141 | -7.130 | -25.887 | 12.574 |
| 3.212 | -3.802 | 23.076 | 27.485 | -29.298 | 0.259 | -29.243 | 25.130 | 22.635 | 2.797 | 24.910 | -20.709 | 27.013 | -27.446 |
| -27.069 | -4.227 | 23.825 | 16.165 | -27.004 | 18.445 | -21.253 | -26.431 | -16.703 | 29.599 | -1.094 | 7.808 | -7.261 | 31.175 |
| 19.946 | -10.953 | -7.807 | -18.890 | 20.863 | 17.057 | 1.938 | -27.184 | 26.618 | 9.546 | 28.118 | 19.729 | 17.792 | 18.709 |
| 0.796 | 23.047 | 6.417 | 17.859 | 11.470 | -10.280 | 11.542 | -24.081 | 28.781 | -9.326 | -2.935 | 23.898 | 21.741 | 7.966 |
| 7.067 | 9.037 | -0.613 | 3.681 | -3.449 | 10.204 | -22.258 | -23.471 | -23.202 | 20.320 | 14.708 | 21.092 | 22.091 | 28.890 |
| 27.355 | -11.589 | -16.870 | -27.541 | 0.594 | -11.719 | -16.296 | -15.657 | 28.080 | -19.506 | -1.206 | 9.017 | -13.148 | 26.180 |
| -26.450 | -14.041 | 6.033 | -5.513 | -22.161 | 19.905 | -14.849 | 20.283 | 15.934 | -23.741 | -1.942 | -24.838 | 26.086 | 9.726 |
| 7.900 | 0.838 | 22.206 | -3.043 | 23.856 | -8.418 | 24.537 | -11.946 | -15.872 | 24.546 | 3.744 | -7.503 | -10.849 | 30.328 |
| -9.916 | -22.312 | 2.551 | -22.192 | -6.927 | -5.834 | -0.981 | -14.810 | 19.620 | 13.743 | 22.593 | 26.257 | 23.016 | -8.015 |
| 22.183 | 3.549 | 9.639 | -20.194 | -6.472 | 4.672 | 1.312 | 5.604 | -26.943 | -10.406 | 1.253 | -25.108 | 24.340 | 7.522 |
| -11.173 | -15.821 | 2.398 | 21.887 | -21.121 | 1.494 | 26.745 | -21.723 | -29.673 | -12.043 | 20.748 | 23.206 | -0.278 | 5.815 |
| -2.929 | 8.248 | 3.492 | 0.884 | -24.287 | -23.438 | 4.014 | -26.379 | -2.343 | -14.001 | 6.041 | -3.928 | 27.507 | 3.048 |
| 9.383 | -0.708 | 1.765 | -6.661 | 2.833 | 5.527 | -20.063 | -11.115 | 15.035 | -3.356 | -2.897 | 26.608 | -25.369 | 8.267 |
| -14.014 | 18.514 | -23.409 | 22.341 | -21.752 | -19.047 | 3.050 | 18.706 | 23.740 | -4.770 | 0.866 | 17.207 | 23.659 | 23.014 |
| -20.646 | 7.216 | -26.836 | 5.001 | 7.214 | -21.748 | -5.874 | 23.651 | 7.194 | -8.982 | 6.448 | 21.183 | -23.739 | -13.985 |
| -5.872 | 2.389 | 4.318 | 28.041 | 14.496 | 13.433 | -8.879 | 25.666 | 18.280 | -15.885 | 25.188 | 20.092 | -26.308 | 9.137 |
| -22.811 | 6.170 | -1.476 | -7.688 | -16.544 | -7.100 | 14.435 | -20.395 | 1.765 | 21.676 | 0.693 | 12.174 | 1.558 | 4.096 |
| 9.989 | 3.499 | 26.786 | 4.536 | -18.956 | -4.442 | 13.071 | 26.675 | 19.422 | 4.788 | 21.574 | -17.416 | 24.967 | 3.519 |
| 24.138 | -9.256 | 0.551 | -20.211 | 16.740 | 5.973 | -21.682 | -17.464 | 16.823 | 12.716 | -0.103 | 22.445 | 7.672 | -0.578 |
| 0.367 | -0.198 | -19.143 | 13.817 | -5.706 | -3.808 | -19.223 | -24.793 | 26.945 | -18.966 | 26.237 | -27.514 | 10.279 | 21.253 |
| -10.230 | -7.681 | 4.013 | -21.615 | -13.399 | -21.716 | -28.926 | 6.902 | 12.475 | -20.776 | -1.717 | 7.742 | 22.379 | -0.003 |
| -24.390 | 8.245 | -13.531 | -27.177 | -8.263 | -18.530 | 22.517 | 8.838 | -2.744 | 27.416 | -0.798 | 26.850 | -25.160 | 19.619 |
| 4.307 | 10.585 | -5.573 | -18.915 | 23.264 | 7.393 | 31.029 | 15.303 | -5.961 | 18.124 | -2.367 | 26.252 | 23.546 | 14.157 |
| -6.591 | -3.364 | -19.531 | -5.950 | 19.131 | -1.629 | -1.922 | 12.014 | 9.981 | -16.340 | 21.194 | 12.299 | 19.150 | 30.737 |
| -17.192 | -7.118 | -22.961 | -7.014 | 4.070 | -15.679 | -7.747 | 4.607 | 1.648 | -15.227 | 0.904 | 27.969 | -26.336 | 2.291 |
| 21.836 | -10.005 | 3.379 | -19.440 | 17.690 | -0.194 | -20.167 | -19.300 | -19.636 | 14.122 | -16.326 | 23.765 | -11.889 | 19.114 |
| 14.376 | -5.931 | -2.353 | -27.298 | 10.005 | 21.406 | 29.975 | 5.781 | -13.042 | -24.566 | 2.362 | -11.781 | 25.003 | 5.398 |
| -25.328 | 6.100 | -8.198 | -17.478 | 23.913 | -4.498 | -5.559 | -15.719 | 19.158 | 21.527 | 9.354 | 27.382 | -27.194 | -5.170 |
| -12.006 | -2.095 | -16.820 | 26.859 | -20.128 | -11.384 | -23.764 | -0.505 | 24.975 | 9.463 | 3.099 | 12.423 | 26.012 | 7.579 |
| -16.321 | 2.311 | 5.757 | -0.127 | -26.243 | -25.419 | 23.738 | -25.580 | 19.248 | -21.604 | -8.024 | 22.606 | -3.515 | -23.969 |
| -19.627 | -9.983 | 17.654 | -24.315 | -22.565 | 19.627 | -26.713 | 3.661 | 2.801 | -19.271 | 5.680 | 23.866 | 27.841 | 4.294 |
| 12.864 | 4.888 | -5.436 | 11.237 | 7.497 | 20.584 | 24.608 | -2.538 | 13.272 | 6.934 | 7.674 | 15.710 | 19.425 | -4.567 |
| -14.454 | -12.576 | 1.070 | -12.758 | 23.224 | -1.366 | -27.460 | -1.217 | 10.532 | -0.449 | 12.311 | 28.774 | 13.291 | 8.696 |

| | | | | | | | | | | | | | |
|---|---|---|---|---|---|---|---|---|---|---|---|---|---|
| -3.819 | 1.291 | 20.857 | 19.318 | -8.374 | 26.501 | 1.063 | -17.871 | 8.492 | -18.317 | -2.751 | 27.588 | 17.889 | 8.298 |
| -13.096 | -21.545 | -5.300 | 24.243 | 22.440 | -18.461 | -4.714 | -6.854 | 24.602 | -11.834 | 10.243 | 26.922 | -24.731 | -19.353 |
| -19.250 | -1.426 | 6.703 | -2.467 | -10.134 | 14.419 | -22.687 | 18.773 | 0.483 | 11.728 | 0.369 | -26.525 | 23.740 | -24.346 |
| -9.799 | 0.129 | -12.062 | 6.064 | 20.846 | -15.311 | 10.685 | -9.702 | -12.873 | -10.344 | -18.017 | 24.954 | 26.554 | 22.715 |
| -19.878 | -1.063 | -2.002 | -3.340 | 17.815 | 0.370 | 9.289 | -13.349 | 28.284 | 17.712 | -20.641 | 27.936 | 20.921 | 16.598 |
| 15.873 | -14.271 | 16.405 | -9.921 | -21.462 | -26.447 | 22.297 | -22.750 | -26.962 | 14.906 | 20.052 | 26.463 | 7.532 | -1.623 |
| 17.318 | -12.116 | 25.175 | -24.280 | 22.885 | -14.408 | 14.913 | -19.924 | 27.849 | 12.546 | -9.532 | 25.226 | 22.085 | 12.594 |
| 1.954 | 1.789 | 20.453 | 25.625 | 19.363 | -5.344 | 8.804 | -20.926 | 1.686 | -6.845 | 6.408 | 25.903 | 19.980 | 16.017 |
| 24.271 | 5.457 | 16.935 | 11.066 | 12.516 | 18.189 | 16.933 | 25.720 | 11.836 | -17.898 | -19.876 | 29.344 | 20.547 | 29.682 |
| 25.007 | 4.305 | 5.588 | 1.897 | -4.929 | 28.096 | 19.405 | -6.122 | -28.666 | 22.024 | 25.989 | 20.706 | -19.917 | -16.240 |
| -7.448 | -5.211 | -6.188 | -27.978 | 20.097 | -1.289 | 27.794 | 22.371 | -4.541 | -16.090 | -23.539 | 18.438 | -24.102 | -7.592 |
| -24.062 | -6.621 | -14.539 | -5.014 | -27.813 | 19.818 | 1.156 | 22.111 | -3.841 | 9.999 | 19.503 | 28.415 | -22.510 | 8.472 |
| 26.865 | 10.189 | 0.922 | 14.900 | 9.799 | 12.975 | -27.801 | -23.118 | 9.929 | 11.725 | 19.038 | 29.137 | 26.965 | 0.298 |
| 5.389 | 1.223 | 4.720 | -23.482 | -19.749 | 22.114 | 10.184 | -24.951 | -1.989 | -21.509 | 4.888 | 26.815 | 20.923 | -10.893 |
| -19.129 | 16.018 | 8.637 | 6.957 | -11.872 | 20.802 | 20.355 | 23.224 | 1.638 | -11.437 | -19.476 | -16.215 | -28.170 | 3.822 |
| -23.086 | -8.650 | -5.319 | 1.734 | -20.737 | 24.776 | -29.457 | 16.907 | -5.820 | 21.470 | 22.834 | 24.872 | 22.698 | 5.195 |
| -3.760 | -4.593 | 8.835 | -12.727 | -4.498 | 16.398 | -13.235 | 15.580 | 24.751 | -24.122 | -18.054 | 26.855 | 18.462 | -11.437 |
| -3.159 | -8.071 | -15.386 | 23.399 | -25.966 | -9.407 | -0.038 | -2.739 | 20.509 | 1.065 | 5.542 | 29.893 | 24.111 | -7.095 |
| -18.734 | -4.108 | -5.525 | -5.296 | 1.605 | -10.295 | -19.327 | -16.727 | -7.293 | -14.827 | -9.592 | -28.955 | 25.300 | 14.165 |
| 18.987 | -19.542 | 19.217 | 21.832 | -12.394 | -6.831 | 14.347 | -10.675 | 1.638 | -22.034 | -24.393 | 29.040 | 13.254 | 11.048 |
| 4.144 | -3.656 | 2.317 | -22.430 | -6.872 | -2.342 | -4.797 | 17.102 | -22.140 | 23.849 | 17.304 | 20.487 | -22.816 | 17.692 |
| 14.106 | -2.579 | -19.891 | -4.905 | 21.472 | -25.936 | 16.003 | 22.355 | 18.077 | 15.550 | -17.654 | 29.337 | -9.375 | -0.142 |
| -11.651 | 6.202 | 16.521 | -25.017 | -18.618 | 10.420 | -31.377 | 0.347 | 10.656 | 7.795 | -1.104 | -30.282 | 25.846 | 1.126 |
| -20.228 | -8.966 | 5.677 | -14.678 | -5.785 | -18.517 | 22.321 | 8.564 | -24.745 | 23.443 | 10.282 | 17.893 | -3.175 | 16.726 |
| 6.386 | 8.732 | -19.453 | -13.394 | -28.576 | 6.104 | -18.067 | -0.889 | 24.723 | 22.699 | -19.001 | -26.730 | 14.206 | 13.787 |
| -2.212 | -9.919 | 26.284 | -0.223 | -2.314 | -11.214 | 2.744 | 10.431 | 21.700 | -23.317 | 0.631 | 28.393 | 12.799 | 3.474 |
| 8.596 | 6.927 | -8.991 | -25.921 | 0.066 | 8.695 | -6.618 | 7.562 | 6.818 | 20.824 | -8.322 | 29.809 | -27.877 | -11.290 |
| 25.428 | -0.079 | -24.831 | -14.060 | -23.484 | -9.839 | -2.444 | -8.568 | 4.632 | 21.909 | -17.062 | 25.674 | 20.866 | -29.454 |
| 9.906 | -7.538 | 21.095 | 15.724 | -12.939 | -23.856 | -28.289 | -20.092 | 10.688 | -5.183 | 11.229 | 22.646 | 28.616 | 27.513 |
| 11.857 | -14.552 | 4.862 | -17.582 | 27.069 | -27.658 | -2.677 | 19.142 | 1.375 | -3.053 | 9.677 | 12.440 | 29.430 | 9.330 |
| -10.434 | 17.804 | 9.758 | -5.315 | 19.258 | 12.606 | -29.335 | 19.451 | 7.421 | 4.318 | -12.812 | 21.302 | 26.904 | -0.931 |
| 7.204 | 13.216 | -7.677 | 14.474 | 21.792 | 24.915 | 23.580 | -7.040 | 2.023 | -8.109 | -23.045 | 30.506 | 22.360 | -5.251 |
| 4.682 | -5.092 | 4.181 | -9.627 | 20.801 | -2.807 | 15.848 | -19.232 | -23.654 | 26.370 | 23.387 | -7.289 | 26.072 | 2.014 |
| -25.426 | -0.051 | 4.912 | 26.015 | 18.209 | -9.434 | -11.799 | -21.005 | -13.725 | 6.747 | -19.114 | 7.546 | -23.050 | 6.554 |
| 21.058 | 14.357 | -2.536 | -15.128 | -22.267 | -12.556 | 27.053 | 14.373 | 5.074 | 13.006 | -9.706 | 25.448 | -5.001 | -10.259 |
| 20.277 | 3.666 | 24.116 | 0.083 | -18.194 | 20.791 | -7.916 | -13.144 | -3.143 | 10.120 | 23.667 | 25.613 | 10.994 | -5.641 |

| | | | | | | | | | | | | | |
|---|---|---|---|---|---|---|---|---|---|---|---|---|---|
| 5.518 | 7.185 | -4.177 | 0.793 | -21.421 | 18.958 | -9.551 | 15.819 | 3.526 | -17.475 | -15.506 | 18.852 | 24.380 | 5.929 |
| 18.063 | -15.541 | 1.451 | 24.673 | -6.739 | -17.711 | 18.012 | 22.583 | 11.004 | 17.153 | -21.355 | 19.201 | 23.414 | -4.903 |
| -12.628 | 15.651 | 7.720 | 1.352 | -27.958 | -3.501 | 10.939 | -5.515 | -0.723 | -17.846 | -22.175 | 28.011 | 25.592 | -29.021 |
| 22.324 | 21.412 | -25.407 | 5.155 | 10.147 | -27.071 | 26.329 | -17.643 | 2.258 | 14.369 | 19.183 | 30.476 | 26.800 | 28.245 |
| -6.719 | -19.638 | 0.415 | -11.904 | -16.750 | -1.950 | -15.603 | -4.000 | 17.172 | 28.629 | 24.931 | 29.108 | 21.273 | 17.412 |
| 18.652 | 5.567 | -18.673 | -22.434 | 3.688 | 13.706 | 1.438 | -6.060 | 19.692 | -6.632 | 14.306 | 17.777 | -20.653 | 15.768 |
| 9.936 | -6.275 | -14.788 | -1.166 | 17.988 | -7.497 | -16.834 | 4.105 | 18.567 | 8.268 | -23.822 | 26.909 | 27.910 | -3.748 |
| 6.055 | 4.777 | 0.332 | -19.146 | 9.723 | 27.965 | 27.025 | -5.561 | 7.037 | 0.395 | -16.032 | 31.078 | 20.106 | -18.146 |
| 16.867 | 1.403 | 2.121 | -25.867 | 12.628 | 28.802 | -3.402 | 10.075 | 23.848 | -24.314 | -26.243 | 3.348 | 18.512 | -9.146 |
| 14.611 | 19.527 | -2.754 | 8.156 | 13.732 | -14.817 | 24.878 | 15.670 | 13.236 | 22.327 | 22.193 | 29.615 | -6.397 | -16.712 |
| -21.393 | -20.061 | 26.025 | -23.061 | 7.042 | 23.230 | -6.711 | 25.782 | -8.815 | 19.812 | 25.005 | 31.427 | 20.872 | 12.038 |
| 11.720 | 3.070 | 17.152 | -0.056 | -8.251 | -26.495 | -11.341 | 24.872 | 10.172 | 5.460 | 8.639 | 30.872 | -30.169 | 5.488 |
| -24.618 | 8.419 | 2.852 | -13.748 | 22.993 | 11.947 | 25.674 | -18.372 | -11.362 | -12.280 | 17.670 | -30.000 | 16.866 | 25.707 |
| 22.548 | 10.500 | -0.376 | 1.148 | -18.259 | -11.534 | 19.063 | -25.177 | 15.104 | 7.141 | 16.187 | 5.237 | 0.628 | 14.216 |
| 0.083 | 12.956 | 10.658 | 6.363 | 16.546 | -8.863 | -6.257 | 8.056 | -10.605 | 22.988 | -13.772 | 24.173 | 22.026 | -25.108 |
| 9.117 | 9.776 | 17.333 | -4.738 | 8.563 | -3.221 | 29.959 | -11.657 | 24.127 | 24.050 | -21.237 | -29.239 | -19.875 | -21.107 |
| -16.342 | 20.341 | 3.155 | -18.957 | -13.717 | 10.170 | -3.976 | -26.713 | 0.572 | 3.245 | 27.030 | 21.120 | 18.416 | 16.933 |
| 1.323 | 9.254 | 0.929 | -16.258 | 10.979 | 9.616 | -9.510 | 23.485 | -5.091 | -16.729 | -3.014 | 5.994 | 26.702 | -0.467 |
| 5.096 | -8.655 | -0.853 | 4.700 | -22.745 | -16.044 | -30.561 | -26.233 | 24.930 | -0.185 | 5.404 | 9.491 | 29.293 | 8.142 |
| -19.350 | -6.717 | -12.284 | 1.996 | -24.537 | -17.568 | 27.602 | -7.031 | 6.661 | 5.867 | 12.628 | 27.264 | 23.284 | 1.827 |
| 25.586 | -9.891 | -2.310 | -7.016 | -19.266 | 24.280 | 20.759 | -21.166 | -12.061 | 23.396 | 26.030 | 7.118 | 28.628 | 22.179 |
| 6.046 | -12.235 | -1.326 | -21.868 | 21.188 | 10.985 | 1.393 | -14.271 | -1.577 | -9.596 | 18.001 | 17.460 | -2.687 | -1.983 |
| -1.524 | 7.873 | 13.111 | 1.981 | 10.355 | -16.494 | 28.494 | -16.600 | 18.649 | -6.263 | 25.032 | -6.855 | -30.997 | -14.656 |
| 22.369 | -3.498 | -10.708 | -13.754 | 20.132 | -4.572 | 30.635 | 3.886 | 14.910 | -5.381 | 10.746 | 29.449 | -20.011 | 11.149 |
| -21.843 | 13.285 | 3.545 | -25.484 | 6.308 | 22.589 | 22.050 | 19.733 | 1.314 | -13.276 | -24.155 | 21.031 | 20.642 | -18.971 |
| -13.513 | 10.749 | 25.946 | -23.964 | 5.677 | 19.384 | -1.975 | -24.765 | 8.714 | -20.384 | -25.691 | -19.499 | 31.133 | 10.685 |
| -17.462 | 18.872 | 4.460 | -18.381 | -27.049 | -9.852 | 24.050 | 9.948 | -3.454 | -6.841 | -24.825 | 19.684 | -9.212 | -11.368 |
| -19.628 | -11.359 | 24.054 | -25.051 | -15.023 | 5.038 | 4.663 | -27.532 | -9.431 | -1.563 | -1.713 | 25.580 | -23.429 | 1.066 |
| 21.760 | 1.103 | 25.633 | 11.619 | 24.435 | -10.579 | 12.178 | 14.708 | -1.806 | -22.512 | 12.239 | 25.132 | 1.767 | -10.671 |
| -12.061 | 10.069 | 5.406 | -6.267 | -29.310 | -12.830 | 2.505 | 4.435 | -6.314 | 20.961 | -26.889 | 26.548 | 17.448 | -7.866 |
| -10.946 | 8.558 | 3.669 | -24.249 | 22.602 | -7.665 | -21.526 | 24.955 | -19.470 | -4.096 | -11.449 | 21.425 | -15.505 | 24.218 |
| 16.815 | 4.521 | 4.528 | -2.669 | -16.253 | -6.717 | -27.513 | -11.079 | 8.511 | -19.027 | 22.862 | -7.631 | 31.813 | -4.657 |
| -15.351 | 11.276 | -2.631 | -8.649 | 6.559 | -26.909 | 22.097 | -6.572 | 3.026 | -7.946 | 27.011 | 29.062 | -31.533 | -1.067 |
| 9.152 | 5.554 | 8.510 | -9.852 | 9.961 | -20.884 | -29.204 | -23.393 | -0.506 | -16.007 | 21.694 | 22.054 | 21.843 | -9.722 |
| -0.339 | 11.657 | -2.993 | -13.844 | 30.097 | 9.977 | 1.122 | -18.547 | 9.364 | -9.194 | -7.089 | 30.571 | 13.207 | 13.502 |
| 23.923 | 6.963 | -7.390 | -23.318 | -3.391 | -1.944 | 1.787 | 13.370 | -1.287 | 28.576 | -25.485 | -6.832 | 3.675 | 12.530 |

| | | | | | | | | | | | | | |
|---|---|---|---|---|---|---|---|---|---|---|---|---|---|
| 14.126 | 5.941 | -0.173 | -4.314 | -18.489 | 18.952 | -9.482 | 8.732 | 5.389 | 21.941 | -22.455 | 23.041 | 23.695 | 3.470 |
| 1.358 | -14.032 | -3.025 | -12.006 | 26.805 | -23.036 | 19.182 | 11.691 | 13.346 | 6.730 | 2.068 | 28.370 | 25.784 | 1.698 |
| 19.089 | 19.150 | -27.717 | -22.410 | 15.668 | -9.481 | 21.057 | -23.663 | -4.218 | 18.962 | -26.016 | 14.522 | -11.093 | -27.091 |
| -7.681 | 8.820 | 27.969 | -23.987 | -25.635 | -11.162 | 13.351 | 1.388 | -20.381 | -4.580 | 23.025 | -4.951 | 23.293 | -11.866 |
| 4.372 | -5.743 | 6.497 | -12.141 | 0.541 | -6.030 | 8.675 | 24.929 | 7.953 | -27.402 | -8.509 | 29.406 | 25.030 | -12.576 |
| 15.662 | -4.215 | -6.574 | 6.378 | -0.367 | 20.833 | 6.756 | -6.686 | -20.849 | 30.227 | 28.552 | 29.932 | -28.057 | -3.027 |
| -3.904 | -9.066 | -10.203 | 7.049 | -19.558 | -29.190 | 29.937 | -10.536 | 14.237 | -20.870 | -2.303 | -5.941 | -21.991 | -7.002 |
| -19.178 | 13.383 | 10.813 | 1.388 | 23.210 | 30.855 | 22.806 | -7.758 | 4.171 | 17.360 | 27.872 | 22.323 | 12.852 | 11.594 |
| -20.058 | 4.203 | -2.577 | -21.578 | 19.574 | -16.483 | 28.944 | 3.589 | -3.243 | -17.170 | 19.698 | 27.909 | -29.501 | -9.170 |
| -9.303 | -11.302 | 3.970 | 5.905 | -9.112 | -28.379 | 25.386 | -19.950 | 2.921 | 8.067 | -24.534 | 19.669 | 26.511 | 7.611 |
| -11.350 | -19.568 | -6.543 | 9.135 | 18.853 | 8.594 | -8.557 | -11.242 | 20.219 | 3.379 | 23.895 | 19.158 | 24.400 | 4.711 |
| 25.489 | -9.724 | -18.354 | -7.516 | -22.507 | -29.683 | -19.602 | -18.929 | -21.758 | -25.315 | -18.374 | 15.711 | -27.683 | -6.293 |
| -4.671 | -3.293 | -25.396 | -19.610 | 5.089 | 28.102 | -27.085 | -8.171 | -9.101 | 22.499 | 18.499 | 22.098 | 28.992 | 23.665 |
| 22.769 | -8.345 | -13.792 | -1.177 | 22.484 | 16.832 | 2.998 | -21.645 | 0.098 | 0.240 | -4.234 | 26.074 | 23.023 | -18.741 |
| 0.843 | 20.437 | -4.257 | -6.403 | -24.447 | 1.516 | 16.458 | -5.628 | 15.144 | 21.176 | 10.610 | -10.265 | -23.013 | 6.681 |
| -8.258 | 19.670 | 0.344 | -24.148 | 8.212 | -12.563 | -23.057 | 2.240 | 27.091 | 11.605 | -1.723 | 16.969 | 22.804 | -7.844 |
| 19.996 | 6.646 | -1.052 | -26.271 | 23.663 | -15.049 | 21.384 | 17.376 | 9.657 | -8.328 | -17.146 | 7.205 | -26.542 | -16.569 |
| -7.498 | -16.735 | -11.739 | 22.053 | 18.225 | -19.754 | 23.437 | -19.242 | 3.196 | 23.807 | -3.650 | 2.884 | 12.016 | 8.276 |
| 8.845 | -11.683 | -9.699 | 5.923 | -21.758 | -10.203 | -5.200 | -26.880 | 1.399 | 10.819 | 2.293 | -1.593 | 14.588 | 22.137 |
| 18.315 | 23.891 | -15.761 | 7.582 | -20.586 | 21.248 | -6.020 | 27.239 | -25.042 | -24.419 | -12.888 | -21.696 | 1.790 | -28.335 |
| 9.507 | 3.868 | -10.737 | 0.330 | 2.451 | -29.730 | -22.124 | 4.701 | -15.417 | -7.586 | -20.041 | 23.707 | 29.162 | 2.191 |
| -7.433 | 22.112 | 9.603 | -5.591 | -10.170 | -2.821 | -25.141 | -16.309 | 21.689 | 3.053 | -2.471 | 23.289 | -29.679 | -5.436 |
| -1.454 | 16.978 | 8.947 | -12.685 | 6.083 | 26.781 | -24.190 | -3.004 | -13.255 | -18.312 | 27.700 | 25.053 | 27.374 | 0.197 |
| -18.441 | -7.380 | -0.424 | 20.828 | -16.403 | -12.699 | 13.275 | -25.871 | 9.399 | -2.864 | 27.121 | 5.485 | -22.683 | -24.311 |
| 14.862 | 15.286 | -8.571 | 4.701 | 9.685 | 3.867 | 24.202 | 24.234 | -17.854 | 4.056 | 26.036 | -8.546 | 24.578 | 27.991 |
| -0.464 | 13.429 | -12.238 | 0.917 | -7.511 | -5.651 | 4.606 | -23.399 | -5.151 | 5.194 | -5.298 | -7.802 | -10.158 | -3.429 |
| -23.761 | 8.834 | -1.432 | -1.084 | -12.919 | -6.848 | -18.975 | -25.473 | -6.583 | 6.089 | -7.180 | -3.028 | -2.503 | -26.108 |
| -11.785 | -8.656 | -1.102 | -3.716 | -21.387 | 17.524 | -4.026 | -6.128 | -15.042 | -10.470 | 29.600 | -10.975 | -22.326 | 10.636 |
| -21.688 | 7.626 | 21.530 | 26.121 | 20.083 | 16.228 | 2.567 | -7.154 | -23.719 | -16.061 | 7.881 | 26.686 | 3.624 | 16.811 |
| 23.796 | 22.733 | -17.842 | 3.371 | -17.378 | 19.051 | 21.442 | 6.303 | 12.338 | -29.329 | -20.427 | -1.029 | 12.972 | -10.645 |
| 12.546 | 17.010 | -19.555 | -7.619 | -5.908 | 10.199 | 5.657 | 9.584 | -12.096 | 8.964 | -14.865 | -4.802 | 29.524 | 7.285 |
| -20.180 | -20.359 | 2.347 | 14.157 | 14.159 | -14.672 | -8.126 | -27.796 | 9.569 | 9.866 | -25.913 | 24.293 | 28.169 | 21.293 |
| -1.814 | -19.538 | -14.588 | -6.004 | -13.449 | -8.182 | -21.099 | 22.322 | -22.513 | -5.927 | -21.366 | -9.776 | -27.156 | -7.358 |
| -9.968 | -10.642 | 14.713 | 2.047 | -22.879 | -7.350 | -4.571 | -0.407 | 8.144 | -0.118 | -27.882 | 3.665 | 1.292 | -8.823 |
| -8.643 | 17.360 | -16.607 | -17.106 | 17.338 | 0.156 | -25.679 | 28.622 | 25.951 | -6.988 | 15.357 | 9.412 | -30.632 | -23.549 |
| -17.951 | 23.414 | -22.223 | 19.378 | 21.272 | -16.864 | 7.916 | -27.385 | 5.942 | 12.480 | -9.057 | 22.476 | 23.491 | 7.088 |

| | | | | | | | | | | | | | |
|---|---|---|---|---|---|---|---|---|---|---|---|---|---|
| 0.227 | -5.986 | -25.321 | -19.288 | -15.205 | 20.724 | 1.205 | 18.767 | -9.367 | -19.871 | 22.345 | -6.159 | 24.797 | 27.320 |
| -16.681 | 15.267 | 11.978 | 0.030 | -18.586 | 15.402 | 27.771 | 19.948 | 11.993 | 23.448 | -20.642 | 5.434 | -28.531 | 6.387 |
| -19.368 | 13.606 | 23.679 | -18.707 | -6.976 | -13.560 | 26.399 | 15.915 | -6.746 | -4.881 | 14.851 | 0.865 | 18.427 | 17.432 |
| -18.814 | 16.833 | 17.718 | -0.428 | 16.015 | -0.911 | -6.076 | 13.849 | 11.210 | -4.491 | -28.173 | -3.851 | 14.217 | -3.314 |
| -10.989 | -15.029 | 15.878 | 1.287 | 12.012 | -2.095 | -27.624 | -11.652 | -8.660 | 0.289 | 12.016 | -0.061 | 19.624 | -3.663 |
| 2.130 | -5.664 | 1.221 | -22.872 | 5.299 | 8.164 | 24.181 | -20.115 | 23.901 | -12.885 | 21.933 | -9.648 | -23.159 | 22.591 |
| 0.986 | 16.205 | -4.595 | 0.659 | -2.503 | 2.152 | 20.816 | 23.056 | -19.406 | -8.603 | 18.262 | -23.166 | -30.176 | 10.726 |
| 3.636 | 19.377 | 9.693 | -1.743 | 8.066 | -12.718 | -0.280 | 5.980 | -22.764 | 14.802 | 26.772 | -19.691 | 26.366 | -0.911 |
| 1.444 | 18.310 | 10.053 | 4.855 | -28.304 | 5.965 | 3.460 | -3.755 | -9.565 | 10.716 | -29.711 | 1.276 | 22.437 | 28.192 |
| 4.528 | -18.820 | 3.898 | -7.824 | 12.938 | 5.175 | 25.793 | -20.799 | -24.732 | 18.663 | 2.405 | -22.795 | -27.803 | 24.466 |
| -1.752 | 9.871 | 23.998 | -26.596 | -12.625 | 31.298 | -0.789 | 0.981 | -22.106 | 17.394 | -2.853 | -11.966 | -19.378 | -12.632 |
| 22.204 | 5.920 | 4.937 | 22.131 | -22.966 | 3.073 | -17.791 | 25.652 | 20.063 | -1.474 | -20.411 | -22.009 | -26.854 | 16.923 |
| 18.139 | 7.846 | 11.416 | -1.280 | -18.286 | -5.944 | -7.017 | -7.875 | 22.030 | -4.264 | 30.240 | 20.514 | -25.169 | 1.067 |
| -24.252 | -19.414 | 5.746 | 8.918 | 10.775 | 19.300 | -23.817 | 5.343 | 5.447 | 28.390 | 26.262 | -20.382 | 23.577 | 12.019 |
| 9.949 | 14.889 | 20.603 | 21.847 | -0.510 | 0.437 | -24.757 | -10.941 | -17.784 | 19.581 | -26.012 | 26.522 | 21.179 | 4.031 |
| 9.463 | 11.473 | 6.441 | -23.786 | -2.224 | 3.779 | -2.197 | -11.416 | -20.984 | -6.860 | 21.154 | 8.740 | -22.189 | -3.146 |
| -0.749 | -8.793 | 14.726 | -19.587 | 25.084 | -9.837 | 26.573 | 7.786 | 13.013 | -24.182 | -7.349 | -28.467 | -11.655 | -10.541 |
| -19.307 | -11.070 | 22.954 | 0.410 | -5.543 | -14.771 | -4.250 | 0.226 | -21.660 | 24.768 | -18.298 | -18.930 | -3.257 | 5.134 |
| 7.194 | 0.896 | 20.126 | 8.451 | -17.127 | 22.059 | 18.191 | -1.527 | -27.684 | 21.866 | -12.744 | 13.239 | 16.337 | -22.047 |
| 23.436 | -19.046 | 16.455 | -12.109 | 16.450 | -13.154 | -22.106 | 6.994 | 20.982 | 14.295 | 31.119 | 6.191 | 1.250 | 2.800 |
| 14.527 | 7.042 | -3.991 | -13.056 | 23.193 | -30.941 | 21.415 | 3.962 | 8.917 | 0.849 | -16.897 | -13.873 | 28.303 | -25.541 |
| -5.942 | -10.542 | 12.766 | 3.133 | -14.857 | -1.959 | 3.945 | -0.877 | -13.496 | -0.295 | 21.728 | 3.810 | -11.403 | 8.608 |
| 7.344 | -5.463 | 7.045 | -25.808 | -3.833 | 2.415 | -8.883 | -2.719 | 20.087 | -13.019 | -5.790 | -2.871 | 22.484 | 1.767 |
| -7.422 | 19.998 | -7.708 | -0.720 | 6.018 | -20.290 | 16.480 | 11.867 | 17.252 | 2.121 | -5.157 | 14.971 | -27.214 | -4.852 |
| 19.582 | 9.267 | 19.659 | -21.467 | 3.724 | 28.472 | 11.216 | -13.523 | 8.632 | -21.967 | 13.703 | 22.763 | -22.356 | -5.756 |
| -4.549 | -19.472 | 22.485 | 0.906 | -20.062 | 4.683 | -18.464 | 21.446 | -6.247 | 7.295 | 5.373 | -7.817 | -26.462 | 17.575 |
| -15.632 | 17.471 | 6.395 | -10.679 | 10.282 | -8.641 | 19.563 | -27.279 | -24.850 | -6.378 | 22.999 | 5.745 | -21.567 | 7.354 |
| -10.457 | -7.710 | 11.927 | -23.578 | 17.063 | 23.755 | -21.841 | -24.295 | -10.123 | -8.359 | 20.110 | 1.451 | 25.340 | 2.400 |
| -0.150 | 21.298 | 4.768 | -2.171 | 26.280 | 29.167 | 11.679 | 24.196 | 22.692 | 5.087 | -30.057 | -12.941 | 18.043 | 0.780 |
| 15.856 | 14.084 | 17.396 | 22.510 | 15.821 | -17.786 | 10.125 | 12.982 | -27.206 | -27.336 | -6.579 | -18.774 | -23.145 | -2.196 |
| -25.268 | -18.611 | -5.856 | -5.496 | -14.287 | 14.727 | -23.122 | -9.497 | 29.966 | 25.020 | -21.240 | -24.796 | -28.287 | -26.881 |
| 24.630 | 27.512 | -11.516 | -1.304 | 18.943 | -7.726 | -4.354 | -14.862 | -2.133 | -4.522 | -7.910 | -21.792 | 25.072 | 3.491 |
| 11.673 | -18.603 | 17.572 | -3.113 | 28.890 | -6.401 | 6.519 | -28.082 | -22.278 | -11.924 | -0.810 | -8.295 | 20.892 | -5.339 |
| 7.474 | 10.070 | -21.977 | 11.152 | 20.884 | 1.271 | 25.519 | -16.624 | 21.409 | -29.002 | -17.668 | 0.574 | 6.769 | -1.306 |
| 18.556 | -14.534 | -15.773 | -27.540 | -16.654 | -15.477 | -5.935 | 20.251 | 5.393 | 11.558 | 26.830 | -19.801 | -29.988 | 16.525 |
| 12.347 | 8.515 | -20.658 | -20.785 | 5.651 | 30.588 | -0.277 | 8.766 | 13.722 | 18.501 | 8.773 | 24.658 | 1.190 | -7.897 |

| | | | | | | | | | | | | | |
|---|---|---|---|---|---|---|---|---|---|---|---|---|---|
| 13.978 | -8.671 | -22.445 | -4.696 | -20.770 | -25.059 | 23.027 | 22.868 | 20.357 | 0.264 | -3.710 | -0.930 | -0.391 | 1.761 |
| -19.408 | 11.682 | -24.583 | 5.437 | 10.013 | -7.110 | 26.688 | -1.974 | 7.115 | -21.498 | 17.809 | 22.266 | 5.234 | -23.360 |
| -22.793 | 16.050 | 19.214 | -24.248 | -18.241 | -19.600 | -10.593 | -15.081 | 21.827 | 4.754 | 28.003 | 20.618 | -23.488 | 23.272 |
| 12.885 | 16.959 | 7.211 | 2.634 | -21.424 | 0.886 | 23.790 | 25.921 | -23.022 | 27.709 | 10.116 | 9.749 | 27.531 | 20.030 |
| -21.684 | -17.929 | -5.178 | 2.970 | 21.427 | -13.556 | 24.965 | 12.621 | -20.248 | 12.171 | -18.897 | 19.456 | 18.017 | 27.956 |
| 11.593 | 20.707 | 17.985 | -26.218 | 13.176 | 23.839 | 5.118 | -17.769 | 23.864 | -3.782 | 27.068 | -21.138 | 21.114 | -13.764 |
| -0.259 | -15.335 | -13.257 | 4.866 | -4.245 | -17.754 | 3.653 | -16.635 | 5.020 | 8.065 | 8.007 | 3.176 | -13.552 | -10.654 |
| 21.662 | 12.749 | 10.431 | -1.129 | 14.927 | 29.164 | -2.990 | -13.495 | 28.060 | 7.128 | -9.017 | -4.012 | 18.464 | -6.403 |
| 25.370 | 9.760 | -24.595 | -23.373 | 2.536 | 28.832 | 18.430 | 3.960 | 7.261 | 4.642 | -7.407 | 20.546 | -17.783 | -22.610 |
| -20.963 | -1.301 | -12.440 | -14.975 | 7.451 | 7.212 | 12.229 | -20.699 | 4.682 | 21.429 | 21.520 | 6.161 | 29.578 | -4.203 |
| 27.922 | -8.837 | 4.053 | 20.217 | -3.095 | 4.400 | -1.057 | 8.913 | -12.881 | 0.406 | 25.507 | 4.247 | 25.632 | 16.779 |
| -21.914 | 17.622 | 26.852 | 16.691 | -19.433 | -21.370 | 19.868 | 2.323 | -16.863 | -2.741 | 29.686 | -19.980 | -16.816 | 1.778 |
| 8.055 | 8.418 | -0.629 | -19.073 | -22.188 | 2.574 | -27.923 | 21.597 | 11.980 | 18.228 | -4.619 | 21.076 | -25.478 | -3.704 |
| -23.675 | 7.592 | -15.559 | -9.746 | -12.160 | -22.130 | -24.570 | 0.798 | 4.395 | 19.570 | -26.848 | -8.699 | -16.371 | -10.792 |
| -11.279 | -7.687 | 11.485 | -27.300 | -23.131 | 26.007 | 22.191 | -19.368 | 30.638 | -0.315 | -26.446 | -14.748 | 27.079 | -2.033 |
| 10.387 | 6.117 | -15.284 | -3.948 | 4.931 | -30.591 | -9.638 | -18.887 | -19.059 | 2.756 | -9.822 | 21.946 | 23.214 | -5.644 |
| 9.813 | -13.143 | 19.009 | 26.153 | -5.959 | 4.426 | 19.158 | 11.884 | 23.630 | 6.719 | -15.285 | 7.602 | 24.540 | -8.642 |
| 22.555 | -27.250 | -14.204 | 0.788 | -12.652 | -6.784 | -15.670 | 29.090 | -17.632 | 19.106 | 29.486 | 16.109 | -28.545 | 17.255 |
| -9.161 | -6.728 | -20.738 | -12.561 | 25.350 | -4.936 | 1.982 | 26.782 | 19.628 | 1.100 | -16.726 | -4.331 | 18.297 | 25.081 |
| 25.298 | -8.067 | -26.572 | 11.025 | 16.965 | 9.972 | 7.711 | 2.932 | 17.619 | -6.119 | -9.371 | 11.222 | 22.103 | 16.062 |
| -25.820 | -21.609 | -11.352 | -3.061 | 29.295 | -17.177 | -21.495 | -13.843 | 14.741 | -12.521 | -8.082 | 13.115 | 16.919 | 11.377 |
| -26.435 | 18.822 | 23.396 | 1.271 | 0.887 | -4.372 | -23.639 | -14.891 | -12.476 | 6.337 | 16.200 | 0.304 | -10.665 | -5.237 |
| 2.792 | 16.727 | 8.917 | -27.092 | -17.930 | 3.830 | 26.470 | 13.432 | -29.998 | 14.874 | -24.718 | -2.816 | 25.525 | -1.094 |
| -3.094 | -12.302 | -7.647 | 28.580 | 23.328 | -23.039 | 5.082 | 19.889 | -24.524 | -4.403 | -4.822 | -6.989 | -17.858 | 9.317 |
| -10.621 | 7.205 | -29.363 | 0.477 | -2.793 | 10.771 | 3.959 | -7.723 | 13.000 | -5.681 | 27.282 | 25.964 | 20.813 | 5.942 |
| -25.338 | -7.507 | 17.768 | -14.843 | -10.598 | -20.157 | -25.325 | -23.298 | 20.270 | 24.444 | 1.727 | 18.451 | 2.781 | 21.564 |
| 9.567 | 10.562 | 28.188 | 24.010 | 17.714 | -16.579 | 22.271 | 8.894 | 12.123 | 7.409 | -26.404 | -0.698 | -9.844 | -24.390 |
| 24.075 | -10.807 | -30.285 | -5.346 | -0.687 | 29.007 | 9.259 | 27.222 | -18.340 | -21.847 | -7.395 | -5.083 | 24.384 | 13.774 |
| 23.448 | -19.688 | -24.687 | -29.584 | -3.581 | 7.395 | -17.979 | -5.065 | -21.951 | -13.612 | 21.888 | -10.069 | 26.043 | 25.816 |
| 21.290 | 2.958 | -31.037 | -0.797 | 1.139 | 1.284 | 25.035 | -13.518 | 5.924 | -19.362 | -1.395 | 25.182 | -26.182 | -6.473 |
| 0.018 | -15.806 | 30.886 | 18.747 | -11.417 | 16.310 | 2.322 | 4.931 | 21.618 | 17.715 | 14.123 | -15.580 | 1.680 | 23.029 |
| -15.659 | 16.279 | -29.118 | -21.820 | 12.836 | 29.419 | 7.079 | 26.120 | 17.612 | -1.002 | 0.849 | 6.659 | -6.064 | 19.279 |
| -9.032 | 21.293 | -30.240 | 0.721 | -5.147 | -3.415 | 20.418 | 28.725 | 29.937 | -5.144 | -3.065 | -8.263 | -12.930 | 0.477 |
| 3.843 | 23.988 | -1.908 | 20.013 | -14.977 | 5.330 | 23.924 | 18.026 | -5.365 | 21.381 | -7.712 | 15.314 | -22.649 | -1.647 |
| -16.068 | -5.887 | 17.918 | -6.941 | 13.569 | 4.321 | 5.607 | 10.454 | -28.847 | -1.960 | -5.463 | -3.414 | 25.544 | -19.119 |
| 13.469 | 18.113 | -31.125 | -10.287 | -21.169 | -13.483 | 26.909 | -14.287 | 23.821 | 3.859 | -6.412 | -4.033 | -25.691 | 2.610 |

| | | | | | | | | | | | | | |
|---|---|---|---|---|---|---|---|---|---|---|---|---|---|
| 4.918 | -1.856 | -1.474 | 7.784 | 4.999 | -1.436 | 3.363 | 3.439 | -20.598 | 16.186 | -11.993 | -5.481 | 20.737 | 1.492 |
| 22.778 | 11.338 | 19.262 | 27.821 | -9.377 | 27.210 | -5.809 | -4.350 | -26.089 | -19.869 | 12.584 | 21.662 | -3.765 | 3.986 |
| 27.409 | 21.891 | -30.006 | -7.585 | 24.847 | 3.919 | 23.393 | 26.525 | -7.963 | -17.389 | 25.328 | -7.167 | -1.218 | 14.082 |
| -16.466 | -14.022 | 20.108 | -1.554 | -6.766 | -5.264 | 27.665 | -27.975 | -23.092 | 22.951 | -6.633 | 9.808 | -26.873 | 6.761 |
| 8.618 | -11.790 | -0.223 | 23.215 | 10.991 | -7.795 | -23.109 | -22.272 | 29.410 | -16.265 | -9.294 | -0.203 | -19.342 | 23.218 |
| 23.871 | 1.514 | 26.399 | 3.843 | -3.072 | 6.689 | 1.086 | 23.764 | 23.258 | 11.746 | 24.527 | 14.472 | -23.040 | -8.482 |
| 0.331 | -28.918 | 22.996 | 4.846 | -13.365 | 11.223 | 20.713 | 11.589 | 13.436 | 1.248 | 6.915 | 16.729 | 18.737 | -1.277 |
| 5.839 | 20.614 | 4.218 | -3.752 | 5.641 | -3.082 | 26.156 | 6.677 | -10.924 | 5.027 | 0.626 | 4.313 | 6.206 | 17.741 |
| -21.924 | 11.017 | 16.504 | -10.876 | -5.249 | -21.354 | 2.160 | 21.337 | -9.276 | -19.308 | -16.971 | 3.663 | 14.690 | 16.177 |
| -12.850 | 6.201 | 28.997 | 25.007 | 23.920 | -13.672 | -2.259 | -3.146 | -21.996 | 4.670 | 20.469 | 6.150 | -23.896 | -15.241 |
| 2.020 | 8.793 | 7.925 | 18.215 | 14.035 | -26.464 | 10.505 | 10.777 | -20.289 | -8.034 | -1.923 | -8.476 | -25.292 | 10.455 |
| -20.594 | 9.284 | 12.946 | -5.889 | 11.716 | 22.255 | -19.000 | -16.180 | 20.554 | 17.644 | 27.215 | -10.905 | -14.804 | -0.397 |
| 15.944 | 15.589 | -21.393 | -10.460 | -16.376 | 16.315 | 5.995 | 21.643 | 22.019 | 2.935 | -2.939 | -1.290 | 22.556 | 16.587 |
| -2.653 | 25.231 | -0.862 | -13.990 | 8.262 | -12.856 | 27.218 | -2.673 | -22.406 | 3.569 | -5.083 | 2.027 | 23.049 | -21.014 |
| -8.249 | 14.004 | -29.360 | 6.946 | 4.595 | 27.728 | 22.203 | 27.397 | 3.762 | -19.206 | 5.452 | -4.274 | -15.828 | 1.865 |
| -6.067 | -5.853 | 6.184 | -11.304 | -0.931 | -8.541 | -24.528 | 26.956 | 16.577 | 26.782 | 16.013 | 21.817 | 27.027 | 9.334 |
| 22.592 | 9.186 | -3.126 | 17.204 | -11.921 | -18.911 | -25.063 | 9.537 | 5.509 | 7.395 | -8.996 | 2.137 | -27.261 | 9.814 |
| 18.741 | 20.815 | 9.423 | 25.016 | 3.753 | -11.145 | 15.611 | 29.484 | -10.497 | -6.877 | 8.226 | -9.990 | 21.805 | 5.766 |
| -18.665 | -8.354 | 8.377 | -0.407 | 6.198 | 23.361 | 1.813 | 1.904 | 28.522 | 0.210 | -23.279 | 3.281 | 15.809 | 7.167 |
| 17.246 | 16.423 | 10.585 | -8.804 | -6.882 | -4.536 | 17.939 | -20.041 | -19.932 | 14.959 | -10.407 | -9.347 | -2.341 | -3.568 |
| 13.198 | -11.122 | -5.686 | -21.802 | 0.396 | -28.406 | 24.230 | -29.513 | -23.232 | -2.094 | -19.066 | 19.741 | 25.881 | -20.288 |
| -19.236 | 10.047 | 23.727 | 19.047 | -19.926 | -14.611 | -10.485 | -11.554 | 9.904 | 12.919 | 7.038 | 23.099 | 2.169 | -25.456 |
| -25.239 | -9.547 | -9.608 | -17.988 | -22.687 | 24.085 | -2.489 | 18.833 | 0.410 | 10.550 | 6.082 | 19.389 | 26.291 | -7.531 |
| 18.462 | 24.271 | 3.121 | -7.324 | -1.362 | -22.101 | -21.168 | 30.542 | 24.353 | -23.853 | -11.244 | 16.904 | 20.369 | -12.879 |
| -2.925 | -8.121 | 1.858 | -3.243 | 1.766 | 9.884 | -26.500 | -9.460 | -11.948 | 27.115 | -7.538 | 0.647 | -21.094 | 8.617 |
| -0.690 | 5.668 | -0.443 | 4.009 | -20.549 | -1.074 | -20.797 | 28.485 | 12.733 | -11.685 | -14.974 | 5.483 | 8.368 | 11.737 |
| 14.387 | 11.255 | 17.204 | 1.004 | -8.964 | -5.812 | 17.498 | 3.401 | -5.816 | 2.197 | -4.604 | 9.138 | -22.439 | 2.996 |
| -27.147 | 7.689 | 1.432 | 23.208 | -4.256 | 14.976 | 5.148 | -15.693 | 1.164 | -13.411 | -2.265 | -11.536 | 21.900 | -0.676 |
| -17.581 | -1.963 | -20.667 | 5.610 | -9.654 | 11.540 | -23.996 | -17.365 | -20.683 | 22.507 | 8.058 | -8.342 | -27.242 | 5.362 |
| 9.555 | -6.780 | -21.444 | 16.576 | -22.119 | -2.833 | -24.921 | 18.707 | -11.777 | 5.706 | -5.346 | -19.541 | -24.071 | -4.592 |
| 2.749 | -20.306 | 0.493 | 12.073 | -3.211 | 16.346 | -17.165 | -7.918 | -2.452 | 5.170 | -6.396 | | 13.651 | 3.854 |
| 4.481 | 21.856 | 24.870 | -27.863 | -10.861 | -25.962 | -1.274 | -30.430 | -2.047 | 8.333 | -7.944 | | 15.830 | 6.058 |
| 21.781 | -22.962 | -17.139 | -8.821 | 20.266 | 14.144 | -25.883 | 20.445 | -18.042 | -6.216 | -9.169 | | -18.287 | -26.308 |
| 8.885 | 4.698 | -0.113 | 3.474 | -0.241 | 2.513 | 0.488 | 25.926 | 12.116 | 1.144 | -11.576 | | -19.567 | -17.572 |
| -17.012 | 7.125 | 14.983 | -29.247 | -2.717 | 0.084 | -3.964 | 11.032 | 22.457 | -0.354 | -0.359 | | 21.064 | 13.043 |
| -3.848 | -8.897 | -8.516 | -30.368 | -7.830 | -1.638 | -8.477 | 25.297 | -6.168 | -1.307 | -3.474 | | 24.057 | -6.267 |

| | | | | | | | | | | | | | |
|---|---|---|---|---|---|---|---|---|---|---|---|---|---|
| -13.155 | -5.308 | -24.228 | 26.021 | -18.921 | 28.249 | -24.509 | 8.398 | 1.079 | -2.693 | 0.388 | | 13.002 | 4.568 |
| -14.820 | -12.640 | 1.432 | 11.800 | 6.443 | 24.093 | 21.635 | -8.492 | 14.184 | 11.338 | -4.530 | | 26.889 | 8.765 |
| 22.630 | 20.497 | -1.368 | -9.763 | -14.564 | 26.041 | 17.640 | 20.655 | 11.000 | -8.574 | -14.289 | | -23.471 | 5.428 |
| 28.756 | -24.692 | 8.357 | -25.344 | -1.411 | -1.315 | 4.589 | 0.957 | -26.741 | 20.567 | -20.617 | | -6.239 | 3.566 |
| 23.265 | -5.246 | -4.148 | -1.630 | 7.553 | -26.412 | 5.165 | -5.804 | -17.134 | -10.768 | 4.322 | | 15.704 | -1.524 |
| 26.770 | 0.803 | 5.035 | -4.120 | -24.774 | 10.735 | -25.528 | 16.314 | -23.491 | -3.754 | 7.634 | | -25.064 | -8.790 |
| 13.790 | 24.552 | 20.937 | -2.661 | 10.846 | -15.775 | 7.525 | -24.471 | 27.452 | -12.171 | -4.914 | | 3.799 | -0.154 |
| 1.473 | -6.087 | -0.894 | 6.898 | -7.200 | 0.229 | -20.971 | 11.982 | -10.848 | -20.729 | -18.831 | | 18.797 | 6.185 |
| 25.541 | 4.677 | -3.848 | -8.136 | -22.303 | 3.291 | -24.033 | -2.608 | -19.193 | 25.754 | -22.827 | | 12.035 | -3.507 |
| 12.544 | -5.000 | -0.156 | -3.387 | -3.244 | -22.467 | -2.154 | -8.881 | -7.763 | -1.768 | -19.746 | | 22.560 | -12.754 |
| -1.146 | -3.706 | -6.762 | -5.053 | -2.013 | -3.814 | -11.686 | -20.268 | 4.192 | 8.414 | -0.599 | | 20.299 | 4.420 |
| -29.406 | 6.504 | 11.162 | -18.324 | 27.463 | 10.106 | -16.365 | -19.599 | 17.116 | 5.961 | -12.578 | | 2.693 | 2.488 |
| 7.666 | -6.286 | 19.456 | 31.422 | -22.615 | -20.325 | 1.252 | -18.590 | 7.917 | -6.580 | 5.348 | | -2.611 | -4.602 |
| -11.776 | 7.863 | 3.058 | -4.475 | 10.653 | -7.140 | -12.658 | 24.857 | 24.516 | 7.024 | -3.163 | | -2.004 | 1.739 |
| -6.447 | -2.825 | 12.971 | 5.404 | -13.864 | 0.877 | -23.350 | -31.653 | 15.786 | -5.484 | 3.635 | | 25.572 | 0.043 |
| -19.962 | -8.620 | -23.303 | -14.545 | -14.875 | -4.152 | 16.153 | -7.926 | -9.916 | 10.183 | 0.871 | | -3.327 | 10.572 |
| 6.615 | -18.723 | 10.473 | 8.109 | 3.455 | 22.518 | -25.363 | 2.773 | 19.701 | -12.546 | -1.580 | | 25.944 | 9.812 |
| 4.730 | -20.716 | -4.645 | -31.267 | -11.107 | 4.300 | -11.334 | -12.218 | -4.458 | -6.910 | -18.395 | | -21.426 | -4.944 |
| 17.464 | -3.313 | 16.511 | -11.378 | 11.624 | 2.190 | -20.822 | 16.596 | -6.658 | -28.564 | 11.231 | | 18.156 | 16.724 |
| 22.071 | -4.489 | 8.467 | -20.953 | -1.309 | -8.813 | 0.060 | 8.583 | -5.760 | 27.332 | -10.818 | | -25.670 | 8.366 |
| -28.322 | -16.791 | -2.451 | -2.136 | -9.365 | -1.233 | 7.079 | 13.845 | -17.496 | 2.896 | 2.680 | | 17.210 | -13.955 |
| 14.393 | -6.209 | 5.057 | -18.833 | 15.822 | -27.044 | -17.155 | -5.662 | -23.433 | -4.976 | -2.638 | | 10.722 | 6.736 |
| 16.621 | -10.524 | 12.044 | 16.251 | 19.971 | 17.113 | -16.687 | 24.142 | 12.268 | 17.438 | 26.426 | | -25.553 | -3.428 |
| -19.048 | 1.860 | -23.918 | -12.136 | -26.081 | -5.540 | -18.206 | 30.709 | 8.818 | 4.549 | 6.168 | | -24.162 | -7.362 |
| -23.937 | -0.934 | 14.144 | -0.117 | 9.467 | -26.300 | 20.084 | 17.957 | -9.170 | -2.385 | 5.354 | | 18.562 | -22.985 |
| 22.876 | 5.533 | 13.436 | -0.566 | 13.648 | -8.574 | 1.756 | -15.270 | -28.980 | -3.247 | -3.828 | | 21.430 | -1.783 |
| -15.411 | 15.466 | -0.490 | -21.718 | 3.997 | 27.555 | -19.704 | 6.929 | -10.744 | -8.015 | -4.545 | | -6.470 | -10.347 |
| 19.119 | 4.819 | -8.476 | 6.910 | -8.821 | -12.352 | 16.177 | -19.202 | 25.916 | -24.324 | 3.820 | | -11.199 | 16.604 |
| -16.804 | -24.294 | 2.144 | 3.293 | 6.242 | -5.847 | 13.870 | 25.747 | 25.030 | 14.462 | -1.590 | | 20.935 | 14.643 |
| 20.823 | -20.059 | 4.653 | -25.797 | -9.170 | 1.640 | 3.181 | 18.995 | 7.299 | 6.558 | -2.863 | | -15.135 | 22.727 |
| -2.266 | -21.250 | -13.019 | 15.733 | 2.400 | -19.327 | -11.377 | -21.072 | 21.217 | 18.065 | -14.018 | | 19.749 | 4.677 |
| -21.844 | -9.183 | 3.868 | 23.475 | 4.429 | -9.403 | 1.121 | 25.427 | 11.472 | 12.095 | 25.044 | | 22.079 | 22.028 |
| 18.574 | -14.452 | -0.931 | -1.112 | -0.176 | -7.067 | -4.977 | 23.428 | 10.607 | -7.589 | -2.336 | | -24.984 | 10.954 |
| -10.954 | 11.218 | -5.405 | -19.876 | -7.979 | 9.050 | -7.203 | 1.205 | -12.604 | 20.404 | -0.973 | | -4.941 | 1.549 |
| 8.277 | 0.839 | 9.706 | 19.594 | 16.966 | 12.885 | -6.665 | 13.537 | -0.743 | 3.093 | 1.423 | | -9.073 | 21.461 |
| 22.400 | -8.489 | 7.385 | -8.189 | 7.762 | 14.659 | 15.580 | 9.202 | 14.499 | 6.075 | -4.679 | | 1.524 | 12.086 |

| | | | | | | | | | | | | | |
|---|---|---|---|---|---|---|---|---|---|---|---|---|---|
| 3.860 | -3.352 | -6.715 | 8.806 | -12.959 | -9.197 | -9.220 | -11.865 | -23.604 | -27.701 | -12.372 | | -2.244 | -8.605 |
| 8.826 | 5.963 | -9.337 | -23.462 | -14.422 | 20.118 | -11.103 | 28.542 | -6.284 | -11.286 | -16.761 | | 13.176 | -15.359 |
| -14.273 | -14.945 | 1.154 | -27.812 | -4.227 | 23.826 | 5.249 | 1.460 | -13.108 | 8.703 | -15.452 | | -20.816 | 14.547 |
| -19.602 | -10.701 | -1.670 | -11.730 | 22.086 | 15.398 | -9.706 | 3.356 | -12.062 | 3.951 | 13.487 | | 15.393 | 1.620 |
| 6.213 | -5.908 | 8.351 | -5.026 | 7.248 | 8.566 | -18.652 | -32.032 | 10.117 | 2.332 | -11.085 | | -0.485 | 23.233 |
| 29.417 | 4.571 | 2.688 | 21.319 | -6.692 | 2.739 | 4.524 | -28.938 | -25.702 | -17.911 | -22.880 | | 16.154 | 17.555 |
| 2.551 | 1.480 | -7.384 | -9.115 | 5.876 | 0.541 | -13.908 | 26.332 | -10.761 | 7.633 | 4.712 | | -5.153 | 4.328 |
| -17.389 | -8.213 | 14.804 | 4.082 | -5.946 | -7.888 | 0.331 | -11.420 | -29.788 | -20.104 | 2.483 | | -17.020 | 3.098 |
| 19.435 | -7.432 | -3.779 | -22.896 | 3.837 | 2.120 | -20.823 | 16.582 | 4.442 | 27.263 | 8.017 | | 1.771 | -14.272 |
| -28.922 | -4.555 | -5.136 | 10.272 | -13.125 | -2.214 | -5.405 | -12.045 | -2.122 | -10.304 | -2.155 | | | 9.496 |
| -1.521 | -0.734 | -18.635 | -25.311 | 10.572 | 4.591 | 11.056 | -28.469 | 20.345 | 1.380 | 5.398 | | | 10.189 |
| -15.846 | 1.254 | 5.382 | 5.584 | -12.683 | -8.782 | 5.687 | 6.571 | -8.433 | 18.859 | -4.370 | | | -15.462 |
| 0.535 | -3.633 | 21.796 | 23.256 | -16.347 | -19.164 | 18.398 | -27.648 | -28.228 | -5.285 | 7.349 | | | -13.258 |
| -31.359 | 2.645 | -2.455 | 6.169 | 9.346 | -0.180 | 0.101 | 14.247 | 6.509 | 9.930 | 12.162 | | | 21.201 |
| 9.879 | -6.473 | -24.865 | -31.166 | 8.998 | 1.232 | 23.419 | -22.581 | -9.667 | -3.161 | -3.524 | | | 11.905 |
| 9.325 | 3.166 | -17.485 | -19.731 | 4.660 | -4.350 | 21.803 | 18.144 | 11.441 | -27.942 | -22.733 | | | 10.632 |
| -11.545 | -1.216 | -4.398 | -21.123 | -1.185 | -0.973 | -5.873 | -1.344 | 22.401 | 18.279 | -21.516 | | | -6.028 |
| 29.973 | 3.303 | 8.847 | -28.720 | -6.482 | 10.788 | 1.682 | 25.610 | -11.347 | 9.453 | 1.293 | | | 9.930 |
| -19.383 | 6.449 | 0.633 | 3.096 | -11.055 | -3.626 | 26.024 | 0.689 | -22.490 | 14.364 | -5.074 | | | 24.410 |
| -20.640 | -2.565 | -26.072 | 17.753 | -3.378 | -8.055 | 6.936 | 20.837 | 12.559 | -1.339 | 16.761 | | | -4.767 |
| -12.374 | -5.541 | 10.390 | -10.003 | -6.008 | -3.071 | -12.120 | 12.554 | 10.841 | -20.962 | -3.292 | | | -10.730 |
| 12.888 | 0.623 | 4.489 | -12.681 | 24.601 | 2.242 | -23.908 | -20.291 | 9.277 | 20.377 | 24.395 | | | 18.241 |
| 11.954 | -25.378 | 4.089 | -25.481 | -2.211 | -12.874 | 20.726 | 29.945 | 26.052 | 1.507 | 1.449 | | | -3.601 |
| -9.273 | -23.107 | -11.890 | 25.530 | -7.079 | 8.246 | -14.326 | -8.599 | 17.712 | -6.927 | 2.858 | | | 1.218 |
| -21.579 | -3.463 | -21.257 | 20.139 | -9.053 | 25.442 | 17.273 | -19.499 | 25.008 | -8.166 | -0.544 | | | 7.806 |
| -18.380 | 5.249 | -14.377 | 0.361 | -19.759 | -2.195 | -5.377 | -7.743 | 19.033 | 20.058 | 5.087 | | | 11.221 |
| 9.586 | 8.457 | -0.746 | 20.657 | 0.204 | -6.440 | 24.985 | 27.104 | 16.923 | -18.748 | 9.128 | | | -1.842 |
| 8.830 | -8.300 | 6.134 | -2.015 | 1.464 | -15.479 | -27.309 | 15.373 | -7.959 | -1.470 | 3.174 | | | 13.548 |
| 11.419 | -6.719 | -3.964 | 10.283 | -17.628 | -21.977 | -18.540 | -2.843 | -8.665 | 4.471 | -19.462 | | | 8.289 |
| 2.356 | -15.154 | -26.220 | -9.081 | 7.724 | 4.986 | -25.938 | -18.392 | 8.587 | 6.102 | -2.352 | | | 5.551 |
| 1.897 | 1.466 | 1.401 | 12.525 | -18.493 | -10.841 | 23.253 | 26.140 | -6.001 | 4.147 | -16.411 | | | -5.520 |
| -11.472 | 3.192 | 2.742 | 13.921 | 2.022 | -25.622 | 5.115 | 21.576 | -12.573 | -0.862 | -24.998 | | | -11.983 |
| -5.048 | 2.579 | 4.932 | 3.970 | -8.399 | 23.680 | 12.270 | -5.347 | -24.068 | 12.635 | 9.343 | | | -13.241 |
| 10.790 | -23.754 | -9.411 | 13.696 | 26.100 | -23.923 | 18.154 | -1.650 | 5.950 | -10.631 | 15.856 | | | 8.851 |
| 18.536 | -6.360 | 7.031 | 19.516 | 0.692 | 3.374 | -17.685 | -27.535 | -0.234 | 5.376 | 2.217 | | | 21.943 |
| 4.850 | -5.261 | -7.350 | 2.688 | -20.345 | 20.313 | -21.438 | 27.595 | -19.818 | -18.238 | -20.034 | | | -9.888 |

| | | | | | | | | | | | | | |
|---|---|---|---|---|---|---|---|---|---|---|---|---|---|
| 0.463 | -0.112 | 16.528 | 25.264 | -1.266 | -4.775 | 8.486 | 14.800 | -10.113 | 2.701 | -7.481 | | | 13.050 |
| -20.023 | -7.304 | 11.117 | -2.637 | 4.312 | -1.369 | -18.474 | 4.220 | 21.515 | 9.013 | 3.860 | | | 9.649 |
| -6.556 | -12.992 | -11.179 | -13.696 | -7.346 | -0.336 | 23.073 | 10.503 | 15.701 | -6.218 | 7.492 | | | -10.662 |
| 11.845 | -19.995 | 7.513 | -22.948 | -3.977 | 14.937 | -22.965 | -21.932 | -17.501 | 16.509 | 5.043 | | | 20.542 |
| 1.572 | -8.653 | -13.145 | -27.102 | 3.794 | -11.012 | -26.203 | -27.982 | 23.768 | -4.064 | 1.571 | | | -2.350 |
| 17.688 | -9.611 | -2.147 | 18.515 | 7.922 | -14.157 | 19.009 | 29.771 | 13.208 | 7.429 | 10.801 | | | -5.737 |
| -16.239 | -0.931 | -4.804 | -27.627 | -3.436 | 8.714 | -0.948 | 20.522 | -6.606 | -2.320 | 24.941 | | | -3.933 |
| -10.506 | 9.359 | 0.150 | 19.614 | -22.637 | -5.486 | -22.696 | 16.776 | 27.674 | 7.711 | 19.553 | | | 3.871 |
| 28.033 | -1.893 | -23.013 | -3.745 | -21.144 | -4.207 | 25.557 | -15.386 | 22.209 | 6.041 | -0.045 | | | -4.892 |
| 28.380 | -12.066 | -2.743 | -1.017 | 4.818 | 22.980 | 24.711 | -20.290 | -13.574 | 2.887 | 25.645 | | | 11.008 |
| -8.958 | 0.914 | -5.309 | 2.873 | -5.409 | 19.086 | -1.752 | -18.295 | -29.341 | -10.720 | 25.292 | | | 25.759 |
| -28.650 | -0.521 | -10.463 | 23.302 | -20.127 | 24.695 | 27.864 | 6.038 | 19.037 | 12.698 | 22.565 | | | 3.007 |
| -20.385 | -3.674 | 10.350 | 0.522 | -16.059 | 17.567 | -2.316 | -29.682 | -7.995 | -16.042 | 24.306 | | | 11.777 |
| -27.194 | -0.072 | 9.790 | 3.748 | 3.848 | 5.029 | 20.084 | 28.932 | 21.010 | 19.233 | 20.036 | | | 4.943 |
| 21.435 | -6.538 | -1.319 | -29.673 | -4.072 | 23.671 | -14.444 | 23.458 | 10.214 | -16.728 | -15.505 | | | -9.447 |
| -11.669 | -4.812 | -0.387 | -25.339 | -23.169 | -3.419 | -16.929 | 17.767 | 8.477 | -9.405 | -25.671 | | | -10.351 |
| -17.974 | 6.533 | -3.882 | -21.592 | 23.166 | 19.210 | -26.596 | 10.375 | -19.213 | -10.143 | 4.166 | | | 26.238 |
| 7.342 | -0.971 | -8.308 | 2.026 | -2.660 | 16.358 | 11.448 | -2.371 | 23.972 | -3.407 | -18.740 | | | 8.213 |
| -2.374 | -22.397 | 1.362 | 17.768 | -8.728 | 13.723 | 22.608 | -18.862 | 3.731 | 4.519 | 9.361 | | | 1.482 |
| 16.838 | 0.866 | -1.942 | 11.862 | 11.320 | -0.123 | -10.331 | 23.917 | 7.871 | -2.846 | -1.451 | | | 0.293 |
| 10.106 | -10.967 | -17.398 | -3.114 | 16.324 | 25.869 | 20.629 | 20.167 | 6.835 | 2.525 | -3.939 | | | 22.993 |
| -25.958 | -5.626 | -1.274 | -10.523 | -11.355 | -18.933 | -16.940 | 6.883 | -5.201 | 22.992 | -10.235 | | | 20.717 |
| 22.528 | -6.479 | 4.581 | 10.739 | -4.517 | -1.107 | -23.528 | 3.006 | 9.533 | 12.577 | 2.483 | | | -3.342 |
| -24.121 | -24.723 | -10.350 | 5.037 | -6.355 | 2.678 | 22.898 | -25.707 | -23.788 | -6.653 | -22.061 | | | -16.350 |
| -29.472 | 13.041 | -2.902 | 15.680 | 13.259 | -6.025 | 4.020 | 13.731 | -27.385 | -10.087 | -4.532 | | | -6.240 |
| 14.618 | 9.788 | -12.226 | -22.022 | -20.667 | -17.905 | -0.778 | -30.087 | -6.573 | 22.793 | -15.554 | | | 19.735 |
| -15.337 | -3.354 | 22.153 | -24.549 | 25.108 | -5.139 | -25.638 | -18.908 | -13.710 | 16.192 | 16.338 | | | 10.391 |
| 17.379 | 10.705 | 9.694 | -29.311 | 8.314 | 19.561 | -4.443 | -26.715 | -9.671 | 9.217 | 20.241 | | | 13.974 |
| -6.514 | 8.584 | 12.929 | 1.100 | -5.997 | -10.441 | 4.966 | 0.893 | 22.088 | 1.873 | -0.505 | | | 18.566 |
| -17.249 | -12.465 | 18.939 | -10.504 | -19.140 | -21.243 | 26.580 | 1.551 | -4.923 | 23.575 | -20.753 | | | 8.801 |
| -26.249 | -20.334 | 16.333 | -0.724 | 25.758 | 1.917 | 20.756 | -16.600 | -10.274 | 7.453 | 25.324 | | | -5.834 |
| -18.675 | 2.014 | 19.893 | -12.420 | 17.263 | 24.842 | -28.332 | -28.818 | -11.932 | 9.542 | -25.578 | | | -25.597 |
| -14.676 | -11.843 | -8.573 | -5.932 | 13.284 | -21.425 | 6.896 | 5.106 | -11.051 | 2.067 | 22.114 | | | 17.041 |
| 15.277 | -12.158 | -7.701 | -14.481 | 7.870 | 14.689 | -22.660 | 28.645 | 16.754 | 1.549 | 21.445 | | | 15.832 |
| 14.345 | 1.182 | 21.980 | -3.035 | -6.679 | 11.457 | 23.286 | 26.669 | 12.427 | 11.826 | 0.807 | | | 7.774 |
| -16.443 | 5.357 | -21.130 | -0.243 | -10.860 | -2.754 | 25.057 | 17.782 | 6.662 | -1.910 | -5.984 | | | -10.102 |

| | | | | | | | | | | | | | |
|---|---|---|---|---|---|---|---|---|---|---|---|---|---|
| 27.249 | 3.598 | -22.347 | -4.341 | -17.730 | 9.115 | -20.668 | -29.351 | -21.212 | 5.532 | 21.946 | | | -25.046 |
| 12.825 | 21.053 | 23.964 | 18.630 | -10.063 | 19.486 | 19.836 | 24.460 | -15.058 | 1.023 | -25.430 | | | -0.991 |
| 13.647 | -21.931 | 23.470 | 27.149 | -19.065 | -0.198 | -7.226 | 18.519 | -15.650 | -1.125 | 14.054 | | | -22.275 |
| 0.882 | 0.380 | 23.067 | -28.743 | 24.050 | -24.700 | 22.652 | -17.318 | -18.669 | 7.442 | 9.150 | | | 6.274 |
| 16.657 | -15.815 | 9.045 | -5.633 | 17.062 | 13.023 | -2.115 | 13.265 | 16.709 | -18.143 | 1.535 | | | 3.649 |
| -1.609 | 15.017 | -7.928 | -27.465 | -0.377 | 18.687 | 12.041 | 18.887 | 13.678 | 13.061 | 5.073 | | | -19.457 |
| 23.031 | -6.668 | 12.761 | 27.956 | 8.625 | -8.088 | 17.054 | -2.217 | 3.817 | -2.613 | 26.102 | | | -1.848 |
| -20.894 | 6.080 | 26.375 | 26.357 | -15.396 | 5.400 | -23.147 | -16.322 | -7.068 | -9.315 | -7.693 | | | -5.617 |
| 12.448 | 7.632 | 5.990 | 12.114 | -5.606 | -9.042 | -24.218 | -19.529 | 11.374 | -3.398 | -15.100 | | | -7.150 |
| 23.536 | -1.607 | 4.361 | 3.667 | 1.527 | -0.810 | 25.076 | -29.255 | -17.444 | -1.844 | 8.855 | | | 1.788 |
| -29.081 | -2.022 | 3.461 | 6.735 | -8.257 | 25.317 | 21.962 | -20.457 | -23.657 | -9.109 | 26.027 | | | -3.551 |
| 27.593 | -4.622 | -8.623 | -1.180 | 7.953 | 13.740 | 10.119 | -10.324 | 20.821 | -12.847 | 24.549 | | | 4.067 |
| 2.800 | 7.496 | -1.472 | -13.005 | 1.229 | 0.878 | -29.546 | 21.068 | 22.023 | 24.786 | -10.028 | | | -2.303 |
| 9.172 | 0.715 | -4.590 | -26.013 | 19.044 | -24.447 | -8.564 | -22.377 | 14.118 | -0.825 | -12.973 | | | -11.093 |
| -22.809 | -1.203 | 25.905 | 22.468 | -18.929 | -0.062 | 2.473 | 20.522 | 4.881 | 2.404 | 25.426 | | | 2.941 |
| 20.341 | 10.176 | 15.151 | -23.062 | 2.378 | 12.260 | 22.392 | -17.922 | -10.411 | 14.803 | 26.979 | | | -4.357 |
| 8.470 | 11.097 | -22.838 | -9.576 | -16.177 | -10.728 | -19.395 | -28.201 | 18.489 | 6.548 | 27.606 | | | -5.816 |
| 6.549 | -5.143 | 2.385 | -3.339 | 5.095 | -9.910 | 17.017 | 24.722 | 6.734 | -4.254 | -1.335 | | | 10.140 |
| 5.995 | 2.291 | 6.050 | 8.472 | -17.419 | 15.838 | -23.899 | -8.334 | -3.357 | -3.463 | 16.584 | | | 12.093 |
| -27.018 | -23.885 | 27.263 | -20.883 | 0.545 | 20.800 | 29.183 | -20.257 | -9.842 | 1.982 | 0.691 | | | 13.288 |
| 10.692 | 8.252 | 24.232 | 18.686 | 19.255 | 20.653 | -0.966 | -21.425 | 17.210 | 0.486 | -26.081 | | | -21.290 |
| 14.444 | 11.614 | -19.292 | 0.537 | -4.101 | -17.320 | 27.033 | -27.568 | -21.766 | 7.309 | -20.000 | | | -9.687 |
| -25.190 | -5.568 | 13.856 | -6.928 | -2.275 | 26.501 | 25.439 | 26.697 | 26.294 | 12.829 | 21.307 | | | 7.077 |
| 26.086 | -18.040 | 20.057 | -27.111 | -4.462 | 3.439 | 12.304 | 19.690 | 21.204 | -1.387 | -26.810 | | | -6.842 |
| 18.946 | 8.704 | -0.945 | 2.639 | -11.982 | -7.452 | -8.180 | -1.214 | 16.750 | 9.941 | -9.742 | | | 1.186 |
| -6.780 | 0.549 | 3.131 | -19.308 | -16.777 | 21.507 | 15.673 | 23.061 | 0.740 | -4.592 | -10.733 | | | -9.024 |
| -17.684 | 6.622 | 24.283 | -17.975 | 8.591 | 20.212 | 21.267 | 17.885 | -6.624 | 5.596 | 26.987 | | | -1.972 |
| 15.808 | -7.705 | -11.704 | -26.592 | 16.551 | -18.391 | 11.035 | -26.475 | -2.028 | 8.087 | -1.133 | | | -16.739 |
| 10.058 | -4.114 | -26.377 | 27.875 | 4.118 | -20.862 | -19.899 | 18.746 | -16.167 | -5.790 | 21.935 | | | 3.432 |
| -16.892 | -10.858 | 17.396 | 17.138 | 12.695 | -8.591 | 22.113 | 11.454 | -18.130 | 11.393 | -7.605 | | | 6.528 |
| 27.360 | 1.544 | -2.555 | 16.540 | -0.223 | -9.656 | -21.633 | -15.741 | -22.316 | -19.864 | 27.926 | | | 18.903 |
| -2.537 | -0.908 | -1.897 | 20.494 | 17.571 | -26.115 | 20.338 | -23.493 | 27.873 | -0.325 | 26.318 | | | 16.611 |
| -19.301 | 5.182 | 25.080 | 27.598 | 24.469 | 27.189 | 25.090 | -20.586 | 7.250 | -25.300 | 9.418 | | | 14.984 |
| 22.378 | -2.367 | -21.143 | 24.497 | 5.887 | 6.061 | -25.162 | -22.877 | 25.366 | -8.263 | 4.339 | | | -2.139 |
| 21.969 | 3.295 | 23.534 | -3.761 | 20.339 | 3.155 | -29.150 | -28.029 | 21.968 | 6.130 | 13.950 | | | 11.679 |
| 24.332 | -15.252 | -16.081 | -11.562 | -2.623 | 0.858 | 17.006 | 26.181 | 12.623 | 3.081 | 25.845 | | | -6.960 |

| | | | | | | | | | | | | | |
|---|---|---|---|---|---|---|---|---|---|---|---|---|---|
| -3.584 | -7.397 | -14.706 | -2.000 | 15.778 | -0.375 | -1.975 | -9.894 | 16.166 | -3.758 | 1.734 | | | 20.935 |
| 26.469 | -11.711 | 7.372 | -2.507 | -9.802 | 19.451 | -23.807 | -7.826 | 9.697 | -4.424 | -2.022 | | | 23.810 |
| 28.858 | 5.664 | 4.952 | 5.695 | -17.907 | -23.693 | 19.273 | 5.027 | -9.907 | 4.706 | 28.279 | | | 1.968 |
| 21.659 | 1.423 | 26.800 | -25.150 | -0.258 | -1.342 | -19.982 | 25.190 | 7.436 | -1.830 | -22.001 | | | -11.421 |
| -8.757 | -23.572 | 1.387 | -23.530 | -14.721 | -2.085 | -25.422 | 2.681 | -14.150 | 21.303 | 0.313 | | | 21.958 |
| 4.656 | -0.418 | 9.080 | 10.303 | 2.869 | -20.291 | -4.604 | -18.917 | 3.724 | 27.613 | 25.211 | | | 6.168 |
| -27.995 | 12.324 | 15.672 | -8.599 | 23.318 | -8.320 | 28.847 | 27.176 | 2.901 | -1.503 | 15.685 | | | 5.622 |
| -7.268 | 7.315 | -27.737 | 2.489 | 1.160 | 15.217 | 27.871 | -8.460 | 22.746 | -18.700 | 27.643 | | | 20.369 |
| -20.232 | 3.011 | 2.501 | 23.859 | -7.344 | -2.855 | 2.578 | -23.335 | 21.504 | -6.477 | 4.177 | | | 11.067 |
| 18.350 | -4.998 | 13.601 | 23.230 | 4.265 | -6.794 | 1.027 | 21.364 | -9.293 | 19.100 | 28.551 | | | -3.664 |
| 7.948 | 6.266 | 3.561 | -24.986 | -11.772 | -14.193 | -21.522 | -20.737 | 6.842 | 3.346 | 0.724 | | | -4.357 |
| 12.268 | -10.391 | 28.684 | 26.154 | 15.310 | 3.573 | 22.899 | 18.990 | 1.827 | 0.174 | -7.025 | | | 13.931 |
| 10.261 | 10.109 | 25.341 | 8.240 | 19.865 | -13.162 | 6.905 | -13.188 | 18.980 | -0.569 | 22.259 | | | -2.527 |
| -11.266 | -0.414 | -22.073 | 7.011 | 6.992 | -23.508 | 11.318 | 8.560 | 9.370 | -1.980 | 26.193 | | | -9.016 |
| -18.650 | -4.430 | -26.311 | 22.463 | 18.265 | 25.381 | -28.417 | -24.756 | -22.161 | -10.340 | -13.121 | | | -23.301 |
| -21.927 | -13.024 | 18.833 | 5.760 | 1.959 | 18.925 | -14.036 | -11.171 | -24.975 | -2.687 | 13.174 | | | 3.871 |
| -24.703 | -2.264 | 6.810 | 25.541 | 4.816 | 1.665 | -0.233 | -15.573 | 20.274 | 12.014 | -2.760 | | | -28.008 |
| -14.445 | -5.621 | -1.821 | -3.521 | 15.272 | 9.489 | 27.290 | 22.456 | 16.352 | -8.223 | 27.481 | | | 24.186 |
| 16.105 | -24.140 | -22.855 | -10.720 | -2.438 | -21.379 | 25.080 | 3.153 | -7.828 | 10.437 | -3.417 | | | 11.379 |
| 2.607 | -16.982 | -25.277 | 16.043 | -4.600 | -25.118 | 6.981 | 18.819 | 17.680 | -9.215 | -20.088 | | | 1.666 |
| -10.996 | -13.819 | -26.688 | -23.277 | -14.984 | 4.258 | -14.743 | -27.228 | -9.320 | -24.559 | -6.690 | | | 3.118 |
| -19.766 | -20.793 | -11.799 | 2.540 | 22.337 | -4.539 | 28.968 | 24.975 | -16.432 | -2.712 | 29.331 | | | -13.856 |
| -25.673 | -20.316 | -20.425 | 22.438 | 11.291 | 5.710 | -5.695 | 26.238 | -4.985 | -5.909 | 23.610 | | | 17.783 |
| 28.775 | 12.175 | 25.044 | -25.737 | -13.959 | 10.294 | 30.297 | -12.081 | -6.154 | 23.341 | -7.423 | | | -6.160 |
| -23.833 | -17.409 | -22.488 | 6.839 | -15.917 | -5.569 | -17.020 | 22.239 | 25.215 | -14.809 | 16.265 | | | 0.942 |
| -25.109 | -26.326 | 22.306 | 9.719 | -16.193 | 27.113 | -25.386 | -21.037 | 4.681 | -3.756 | 2.727 | | | -17.187 |
| 12.114 | -12.682 | -14.073 | -2.002 | 17.172 | -2.397 | 16.608 | 21.461 | -7.400 | 19.727 | -3.567 | | | -26.831 |
| -24.378 | -9.280 | 1.559 | -2.549 | 26.803 | -21.680 | 29.842 | 18.375 | -4.299 | -26.544 | 26.312 | | | 4.191 |
| 21.759 | -16.689 | 28.255 | 18.579 | -3.198 | -23.006 | 22.181 | -20.588 | 19.910 | 21.852 | -2.540 | | | 10.091 |
| 11.391 | 7.685 | 21.057 | 23.248 | -6.775 | 1.053 | 24.215 | -23.632 | -20.094 | -8.342 | -14.623 | | | 7.547 |
| -19.551 | 21.950 | -21.733 | 4.823 | 2.041 | 23.330 | 23.570 | -13.112 | -13.649 | 7.481 | 29.213 | | | -8.704 |
| -13.463 | -2.503 | 9.934 | -23.018 | -10.231 | 20.264 | -15.227 | -4.981 | 14.166 | 11.610 | 28.023 | | | 29.283 |
| -20.251 | -0.012 | 13.202 | 16.715 | -20.222 | 2.624 | 17.079 | -7.016 | -17.376 | -12.968 | 18.541 | | | 6.260 |
| 13.694 | -15.888 | -7.913 | 1.128 | 6.620 | 12.548 | -29.627 | -15.203 | 5.503 | 10.921 | 29.705 | | | 27.594 |
| -24.754 | -4.217 | 30.266 | 24.031 | 0.471 | 3.909 | -16.551 | -24.413 | 20.354 | -2.394 | 24.618 | | | 25.093 |
| -9.628 | 5.611 | 27.500 | 14.241 | -4.897 | -26.359 | -27.712 | 19.707 | -25.884 | -3.220 | 21.848 | | | -27.727 |

| | | | | | | | | | | | | | |
|---|---|---|---|---|---|---|---|---|---|---|---|---|---|
| -23.028 | -13.664 | 22.434 | 11.053 | -24.671 | 7.009 | -25.364 | -18.915 | -27.488 | 15.792 | 30.138 | | | -27.028 |
| -6.150 | -20.437 | 24.216 | -7.975 | 21.119 | -6.207 | -31.843 | -20.904 | -1.547 | -11.286 | 20.795 | | | -25.422 |
| 8.441 | 8.508 | -29.611 | -22.280 | 4.990 | -20.036 | 21.038 | 23.238 | 21.183 | -18.270 | -12.579 | | | -21.616 |
| 19.180 | -11.046 | -24.044 | -17.915 | -16.795 | -22.261 | -18.904 | 18.157 | -6.145 | 0.243 | -6.336 | | | 3.924 |
| 0.625 | -5.276 | 28.199 | -3.120 | 17.874 | 29.768 | -25.192 | -22.931 | 17.532 | -1.673 | 2.816 | | | -9.906 |
| -7.501 | -6.547 | 24.199 | -7.244 | -10.338 | 7.920 | -13.781 | 20.169 | 9.720 | -4.213 | 26.755 | | | 17.655 |
| 7.349 | -8.339 | 7.259 | -9.882 | -25.727 | -27.796 | -1.926 | -25.745 | -5.086 | 11.854 | 25.856 | | | 14.504 |
| 23.194 | -9.805 | -21.578 | -23.972 | -8.908 | 16.080 | -28.750 | -1.305 | 5.842 | -14.895 | -28.680 | | | -18.490 |
| -8.727 | -16.174 | 23.025 | -7.991 | -18.383 | -5.249 | 22.667 | -23.666 | 9.308 | -13.925 | -26.008 | | | -7.802 |
| 14.071 | 22.083 | 30.663 | -16.832 | 14.873 | -20.703 | 17.638 | -24.836 | -4.165 | 24.187 | -4.373 | | | 17.992 |
| -10.270 | -0.079 | 29.778 | 21.071 | 21.577 | -12.592 | 31.523 | 21.946 | -0.169 | -8.710 | -5.937 | | | -15.405 |
| 9.112 | -11.300 | 28.707 | -6.612 | -13.660 | -9.410 | -18.208 | 15.490 | -5.647 | 5.229 | -21.605 | | | 22.447 |
| 23.782 | 11.569 | 12.629 | -11.961 | -16.587 | -18.444 | -28.142 | -9.757 | -15.193 | 2.179 | -22.293 | | | 30.763 |
| -24.666 | -12.645 | -27.122 | 24.140 | 12.834 | 8.596 | -27.178 | 18.968 | -1.540 | -6.550 | 11.060 | | | 5.314 |
| -25.710 | 20.960 | 29.178 | 18.240 | 24.391 | 9.912 | -26.205 | -19.345 | -8.544 | -3.646 | -17.230 | | | -30.237 |
| 4.820 | -5.786 | 14.379 | 8.831 | -11.335 | 21.946 | 25.112 | 22.500 | -9.429 | -7.362 | 29.078 | | | -20.850 |
| -1.118 | -21.963 | 5.150 | -14.036 | 3.967 | -21.651 | -1.684 | 11.874 | -14.409 | -9.348 | -15.599 | | | 4.161 |
| -0.720 | 7.167 | 16.510 | 20.760 | -16.852 | -5.109 | 14.477 | -12.186 | 8.123 | -5.034 | 30.267 | | | -2.336 |
| -3.255 | 17.477 | 24.954 | 14.487 | 4.689 | -13.442 | -2.418 | -14.865 | 8.545 | -8.454 | 6.937 | | | 8.352 |
| 20.903 | -7.714 | 19.250 | -9.503 | -7.619 | -19.871 | 24.857 | 1.893 | -16.350 | -12.806 | 28.711 | | | 22.390 |
| -15.398 | 5.531 | -28.762 | 6.790 | -21.736 | 30.288 | 25.300 | 11.359 | -13.767 | 21.628 | 24.783 | | | -20.269 |
| 8.243 | -19.097 | 17.911 | -22.683 | -23.181 | -20.462 | 6.728 | -13.120 | -11.813 | 14.326 | 26.871 | | | -13.571 |
| 0.683 | -3.898 | -30.608 | -7.421 | 25.309 | -14.277 | 23.497 | -8.321 | 17.839 | 6.317 | -20.455 | | | -19.222 |
| 26.769 | -3.294 | 2.827 | -21.406 | -10.752 | -26.018 | -24.342 | | -6.574 | -2.742 | -0.519 | | | 6.929 |
| 10.206 | -6.236 | 19.328 | 1.936 | -9.877 | 14.762 | 21.970 | | -11.214 | 7.915 | -9.877 | | | 30.622 |
| -10.132 | 16.005 | 26.449 | 23.095 | -11.570 | 10.299 | 30.325 | | 4.257 | 25.215 | -29.311 | | | 6.253 |
| -21.566 | 2.179 | 1.469 | 18.815 | 2.880 | -4.418 | -24.892 | | -0.863 | 24.789 | -19.734 | | | 25.368 |
| 16.980 | -5.714 | -4.522 | -3.190 | 14.324 | 4.590 | 11.922 | | -7.481 | 8.600 | 29.749 | | | 17.783 |
| 25.903 | 14.726 | 31.009 | 10.169 | 16.246 | -13.418 | 22.488 | | -12.575 | 27.216 | -30.735 | | | 3.389 |
| 15.175 | -22.309 | 19.603 | -4.120 | 12.402 | 26.729 | 27.682 | | 14.315 | 2.533 | -27.183 | | | -16.193 |
| -13.912 | -20.152 | -15.184 | -14.778 | -15.885 | 19.468 | 20.321 | | -2.189 | 24.394 | -22.434 | | | 8.101 |
| 6.079 | -8.566 | -12.018 | -8.318 | -17.020 | -21.831 | -23.122 | | -5.743 | 17.639 | -18.341 | | | 15.207 |
| -27.906 | -7.015 | 28.587 | -16.034 | -22.133 | 29.307 | 25.541 | | 4.404 | -4.008 | -30.106 | | | 19.155 |
| -5.254 | -5.188 | 26.884 | 1.390 | -22.582 | 22.226 | -27.296 | | -0.305 | 1.305 | 20.431 | | | 1.883 |
| -15.047 | 8.747 | 28.195 | 19.469 | -11.556 | -19.594 | -7.692 | | 5.017 | -4.617 | -7.685 | | | -8.917 |
| -22.352 | -18.165 | -29.165 | 3.726 | -12.692 | -24.040 | -21.174 | | -7.715 | 16.782 | -27.855 | | | 18.403 |

| | | | | | | | | | | | | | |
|---|---|---|---|---|---|---|---|---|---|---|---|---|---|
| 2.544 | 20.983 | 28.580 | 11.764 | 6.102 | -7.908 | -29.278 | | -12.288 | -27.239 | -0.771 | | | 17.095 |
| 4.986 | -17.511 | -13.848 | 18.411 | 11.275 | -0.991 | 21.376 | | 2.685 | 21.373 | -4.166 | | | 11.406 |
| 23.991 | 5.356 | 21.733 | 17.284 | -11.944 | 26.090 | -28.140 | | -9.566 | 10.393 | -2.800 | | | 13.400 |
| -16.092 | -4.489 | 4.533 | 11.660 | -23.588 | -13.223 | -0.845 | | -22.106 | -0.357 | -8.196 | | | 10.611 |
| -17.143 | -22.978 | -11.072 | | 19.715 | 23.515 | -9.923 | | 7.394 | 22.379 | -6.861 | | | -1.377 |
| -21.900 | -12.452 | -14.193 | | -26.854 | 14.246 | -31.297 | | 23.882 | 12.940 | 30.781 | | | 23.184 |
| 3.946 | 24.903 | -15.227 | | -22.816 | 19.117 | -6.786 | | -0.660 | 11.002 | -9.907 | | | -11.516 |
| -20.228 | 13.869 | 27.560 | | 13.149 | -8.511 | 1.335 | | -2.733 | -10.328 | 25.539 | | | -21.578 |
| 8.236 | -9.262 | 5.649 | | 11.994 | 6.742 | -16.722 | | -6.736 | -7.044 | 6.526 | | | 11.790 |
| 27.411 | 6.420 | 29.675 | | 18.882 | -6.710 | 29.163 | | -15.380 | -5.804 | -13.514 | | | 5.864 |
| -5.860 | 23.386 | 26.309 | | 21.059 | -26.583 | 14.132 | | -3.075 | 27.075 | 14.510 | | | 20.885 |
| -11.652 | 22.719 | -19.178 | | 9.495 | -29.035 | 3.343 | | -18.332 | -28.725 | 5.671 | | | -11.834 |
| 12.804 | -16.812 | 28.181 | | -28.460 | -5.904 | 19.782 | | 4.585 | -4.691 | -4.294 | | | 17.622 |
| 1.346 | 2.861 | -28.328 | | 10.578 | -17.338 | -25.025 | | 25.391 | -9.596 | 16.635 | | | 5.298 |
| -20.652 | 5.752 | -17.995 | | -19.351 | 22.396 | -13.114 | | 3.727 | 22.595 | 5.191 | | | 15.095 |
| 10.878 | -24.240 | 0.849 | | 14.483 | -13.450 | -22.702 | | -9.639 | 10.119 | 18.076 | | | 19.691 |
| -14.473 | 8.641 | 23.707 | | -17.547 | -25.177 | 30.203 | | -10.528 | -22.407 | 23.008 | | | -22.981 |
| 13.887 | -9.256 | 25.189 | | -21.001 | -23.780 | 25.274 | | -11.003 | 20.780 | 26.532 | | | 26.493 |
| -2.207 | 13.203 | -29.657 | | 16.416 | -10.748 | 4.867 | | -9.246 | 21.901 | 7.414 | | | -9.953 |
| -3.228 | -6.230 | -12.145 | | 0.066 | 28.720 | 21.209 | | -21.285 | 17.226 | -5.327 | | | -22.293 |
| 20.254 | -17.500 | -17.019 | | 22.144 | -15.905 | 23.069 | | 18.189 | 25.429 | -26.366 | | | -31.023 |
| 22.372 | -11.109 | -5.267 | | -5.527 | -18.713 | -24.650 | | 8.184 | 23.031 | -6.535 | | | 20.844 |
| 6.673 | -7.420 | 8.931 | | -24.683 | -1.514 | -26.364 | | -22.791 | -3.887 | -3.054 | | | 14.156 |
| -23.921 | 5.425 | -28.948 | | -29.944 | -20.091 | 29.451 | | 22.229 | -19.068 | 30.074 | | | -16.742 |
| 9.140 | -12.764 | 20.194 | | -7.989 | -23.137 | -30.175 | | 0.227 | -2.444 | 26.081 | | | -23.892 |
| 0.561 | 15.116 | 21.430 | | 28.066 | 29.625 | 26.779 | | 3.184 | -15.940 | 25.625 | | | 11.333 |
| -25.382 | -8.593 | 26.311 | | 7.555 | 14.991 | 16.623 | | 1.061 | 19.925 | -17.789 | | | 10.845 |
| 19.880 | -13.456 | 28.104 | | -4.052 | 20.508 | -15.584 | | 1.795 | 9.510 | -1.219 | | | -24.317 |
| 16.467 | -21.403 | 11.241 | | -5.035 | 27.027 | -6.624 | | -2.300 | -14.673 | 23.142 | | | 20.687 |
| -6.342 | 20.788 | -0.485 | | -12.353 | -29.247 | -27.318 | | -6.384 | -11.721 | 20.289 | | | 7.864 |
| 12.673 | 0.603 | -23.662 | | -4.669 | 16.035 | 3.894 | | 7.055 | 14.675 | -12.240 | | | 29.759 |
| -17.022 | 6.095 | -22.030 | | 22.531 | -25.939 | 28.078 | | 22.803 | 1.713 | 9.019 | | | 11.249 |
| -22.066 | -15.351 | -13.065 | | -18.545 | -17.941 | -25.691 | | 8.605 | -29.610 | 3.829 | | | -13.578 |
| -27.804 | 9.593 | -14.518 | | 12.574 | -5.207 | -20.688 | | -5.758 | 12.777 | -1.695 | | | -9.264 |
| -12.850 | -23.962 | 1.672 | | -3.377 | 22.068 | 23.081 | | -8.986 | -20.060 | -3.510 | | | -0.003 |
| 18.476 | -6.412 | 18.222 | | -16.499 | -22.180 | 19.564 | | -19.235 | -4.522 | 27.763 | | | -29.596 |

| | | | | | | | | | | | | | |
|---|---|---|---|---|---|---|---|---|---|---|---|---|---|
| 4.270 | 23.016 | 0.677 | | 19.646 | 27.047 | -5.790 | | -10.485 | -16.903 | 10.495 | | | -6.473 |
| -1.438 | -14.382 | -1.685 | | 18.643 | -31.333 | 24.652 | | -3.191 | 10.260 | -17.848 | | | 12.068 |
| -7.274 | -29.342 | 25.780 | | -13.839 | 9.221 | -3.735 | | -5.276 | -7.748 | 27.356 | | | -6.987 |
| 21.686 | 10.806 | 8.889 | | -14.329 | -17.169 | -14.879 | | 7.841 | 1.224 | 22.452 | | | -12.110 |
| -22.454 | 10.433 | -22.040 | | -6.038 | -20.365 | -18.038 | | -4.092 | -7.474 | 20.850 | | | -20.159 |
| 0.434 | -14.085 | 12.050 | | -21.236 | -29.525 | -0.628 | | 5.131 | 21.131 | -22.201 | | | 19.843 |
| -6.861 | -2.348 | -12.711 | | -17.496 | 28.142 | -22.472 | | 10.461 | -21.675 | -26.943 | | | -28.094 |
| -24.644 | -21.585 | -24.081 | | 24.751 | -30.309 | -24.780 | | -6.992 | 14.656 | 13.625 | | | 29.115 |
| -1.955 | -7.356 | -28.412 | | 15.730 | 18.172 | -19.122 | | -0.614 | 22.194 | -1.694 | | | 13.207 |
| 16.220 | -8.768 | -6.223 | | -11.659 | -7.746 | 12.930 | | -10.050 | -8.624 | -0.143 | | | 20.282 |
| -27.057 | -11.828 | 22.701 | | -13.964 | 25.257 | 0.523 | | 3.316 | 22.606 | -8.607 | | | -17.034 |
| 5.397 | -13.257 | -5.319 | | -30.556 | -8.935 | 21.690 | | 0.126 | -4.043 | -25.013 | | | -30.422 |
| -6.838 | 10.339 | -8.812 | | -26.434 | -30.954 | 27.636 | | 9.708 | 5.589 | 2.594 | | | -11.832 |
| 9.170 | -5.761 | 25.300 | | 12.081 | -14.090 | 22.155 | | -3.126 | -18.222 | 0.490 | | | 0.876 |
| 7.311 | -27.878 | 4.513 | | -8.384 | 28.901 | -12.103 | | -1.163 | 8.297 | -17.610 | | | 28.300 |
| -8.158 | 3.634 | 4.016 | | 27.383 | 20.244 | -29.508 | | 16.612 | -28.353 | 20.192 | | | 7.520 |
| -23.292 | 2.019 | 27.030 | | 10.750 | 8.957 | -23.956 | | -1.644 | 28.675 | 8.653 | | | 12.779 |
| 19.955 | -10.183 | -7.626 | | 6.276 | 18.223 | 26.399 | | 15.163 | -13.461 | 25.801 | | | -12.292 |
| -11.127 | 11.339 | -27.096 | | -5.710 | 28.309 | 24.689 | | -17.017 | -8.764 | 18.826 | | | 27.271 |
| -15.119 | -13.648 | 12.566 | | 20.352 | 17.118 | 5.561 | | -0.469 | 5.133 | 27.571 | | | -19.046 |
| 18.300 | -15.754 | -3.060 | | 16.139 | 15.467 | 23.636 | | -6.473 | -1.530 | 3.377 | | | -23.031 |
| 1.559 | -11.437 | -25.389 | | -13.434 | -27.730 | -22.905 | | -14.321 | -21.030 | 28.851 | | | 20.169 |
| 22.829 | -13.273 | 1.606 | | -24.743 | 19.807 | 12.423 | | 22.665 | 8.461 | 26.505 | | | -23.683 |
| 11.219 | 13.604 | -14.319 | | -20.954 | -13.482 | 28.190 | | -7.541 | 19.296 | 5.314 | | | 13.507 |
| -12.165 | -19.549 | 26.659 | | -22.166 | 29.135 | 21.937 | | 17.507 | 18.037 | -25.985 | | | 16.011 |
| -12.548 | 19.953 | 3.086 | | 1.701 | -4.195 | 24.964 | | 14.043 | -2.724 | -13.020 | | | -16.374 |
| 18.735 | -21.777 | 2.711 | | 15.699 | 2.618 | 21.286 | | 5.731 | -7.506 | -14.736 | | | -24.914 |
| 15.884 | 17.381 | -4.521 | | -16.177 | 14.474 | -21.653 | | 24.826 | 10.499 | -18.226 | | | -20.421 |
| -3.922 | -10.545 | -26.658 | | 10.005 | 25.284 | -18.450 | | -11.536 | 14.091 | 27.208 | | | 0.353 |
| 4.427 | 9.887 | 24.402 | | 22.626 | -5.576 | -20.589 | | -7.009 | -18.084 | -1.924 | | | -9.005 |
| 3.613 | -5.327 | 6.408 | | 21.967 | -26.374 | 22.430 | | 4.108 | -24.704 | 2.290 | | | -14.983 |
| -15.630 | 25.404 | 11.788 | | -21.833 | -18.764 | 23.421 | | -7.755 | 8.355 | -7.264 | | | 27.015 |
| 24.596 | 22.579 | 7.401 | | -23.438 | -9.224 | 19.850 | | 16.931 | -15.122 | -11.577 | | | -22.928 |
| -7.898 | 7.680 | -2.531 | | -7.469 | 6.137 | 14.812 | | 16.162 | -3.440 | -4.123 | | | 8.733 |
| 8.642 | -19.126 | 19.435 | | -19.324 | -3.567 | 9.157 | | 1.393 | 21.557 | 13.858 | | | 12.634 |
| -9.707 | -23.772 | 5.269 | | 14.654 | 29.254 | -18.794 | | 2.819 | -10.179 | 13.261 | | | 21.158 |

| | | | | | | | | | | | | | |
|---|---|---|---|---|---|---|---|---|---|---|---|---|---|
| 10.617 | -2.862 | -0.268 | | -6.871 | -25.346 | -1.354 | | -8.266 | 3.691 | -6.222 | | | 29.229 |
| 26.037 | -6.943 | 1.472 | | -23.601 | 24.684 | 23.884 | | -0.678 | -21.006 | -26.230 | | | -10.247 |
| 26.820 | -25.677 | 20.968 | | 11.178 | -1.486 | 16.245 | | -1.937 | 22.746 | -8.946 | | | -12.042 |
| 10.308 | 21.934 | 10.681 | | 10.654 | 2.483 | 17.267 | | 20.561 | -16.558 | 20.685 | | | 26.708 |
| 0.997 | -6.085 | 4.146 | | -8.112 | -29.223 | 4.527 | | 25.377 | 20.189 | -5.794 | | | -21.825 |
| -3.009 | -16.877 | 19.468 | | 27.522 | 28.785 | -4.503 | | -16.565 | 9.428 | 0.067 | | | 7.503 |
| 0.473 | -7.615 | 2.225 | | 9.721 | 10.569 | -26.355 | | -7.765 | 27.673 | -15.325 | | | 11.436 |
| 18.791 | -17.581 | 10.668 | | 19.818 | -21.912 | 22.151 | | 11.381 | -14.432 | -10.663 | | | 0.586 |
| 14.506 | -21.338 | 26.013 | | -30.167 | 27.885 | -12.456 | | 16.348 | -4.305 | -19.690 | | | -3.135 |
| -9.260 | -11.640 | -1.726 | | 21.347 | -13.523 | 26.849 | | 10.364 | 9.024 | 27.634 | | | -11.138 |
| -24.501 | -10.951 | -25.814 | | 0.737 | 5.641 | -12.087 | | 0.122 | -27.572 | 22.794 | | | -13.532 |
| 3.440 | -0.906 | 24.880 | | -22.621 | 23.129 | -21.228 | | -5.834 | 10.092 | -1.522 | | | 14.592 |
| -4.392 | -24.383 | -0.668 | | 18.749 | -27.778 | 27.875 | | -24.696 | 7.875 | -0.890 | | | 4.866 |
| -13.843 | -7.821 | 14.926 | | -8.577 | 14.880 | 3.075 | | 12.090 | -22.867 | 7.226 | | | 13.990 |
| 3.015 | 9.237 | 0.936 | | 8.913 | 8.102 | -28.570 | | -3.137 | 17.876 | 10.418 | | | -24.046 |
| 8.120 | 7.049 | 24.552 | | 18.193 | 28.125 | 6.698 | | -7.383 | 18.335 | 1.497 | | | -10.665 |
| 2.617 | -15.407 | -1.228 | | -6.019 | -5.001 | 26.254 | | 10.079 | 3.716 | -20.304 | | | 19.062 |
| -23.442 | -10.026 | -6.904 | | -7.912 | -19.107 | -25.398 | | 20.295 | -11.279 | 5.590 | | | 26.164 |
| 10.254 | -10.304 | -4.055 | | -20.281 | 21.083 | 18.759 | | -20.990 | 27.840 | 2.538 | | | 21.658 |
| -5.110 | -9.720 | -21.967 | | -25.019 | 22.874 | 5.323 | | -26.078 | 11.855 | -4.550 | | | 3.470 |
| 13.687 | -19.618 | -16.941 | | 19.615 | -23.323 | 20.040 | | -1.440 | -24.314 | 15.712 | | | -27.144 |
| 1.844 | 25.173 | 5.019 | | 5.202 | -19.505 | -19.740 | | 26.874 | 14.058 | -7.171 | | | 8.116 |
| 5.110 | -6.759 | -6.227 | | 27.829 | 13.967 | 11.672 | | 25.047 | 9.568 | -5.145 | | | -6.882 |
| -16.273 | 10.315 | 22.893 | | 20.629 | 25.242 | -24.124 | | 3.793 | -19.664 | 4.912 | | | -20.357 |
| -22.414 | 8.557 | -6.600 | | -16.130 | 22.316 | 7.645 | | 11.802 | 22.303 | -14.812 | | | 7.254 |
| -3.640 | 17.598 | 3.627 | | 17.448 | 26.062 | 24.003 | | -10.438 | -12.555 | 0.938 | | | -20.735 |
| -10.324 | -8.561 | 21.811 | | -13.297 | -14.081 | 27.306 | | 5.686 | 0.885 | -15.834 | | | -10.735 |
| 5.890 | 8.967 | 10.626 | | 19.000 | -16.624 | 19.442 | | -12.887 | -15.586 | -26.203 | | | -7.457 |
| 1.608 | 0.840 | -1.645 | | 6.885 | -25.410 | -21.183 | | -20.521 | 8.881 | -10.213 | | | -19.165 |
| -10.795 | -10.901 | 11.117 | | -15.634 | -4.888 | -15.645 | | 15.249 | 12.686 | -1.142 | | | -3.502 |
| 23.072 | -5.930 | -4.291 | | 23.474 | -9.242 | 25.834 | | -9.052 | -3.205 | -8.514 | | | 3.582 |
| -1.077 | -9.429 | -24.724 | | 7.941 | -6.150 | -18.679 | | 4.288 | -21.464 | 27.867 | | | 20.157 |
| -11.644 | -20.736 | 16.501 | | -20.752 | 14.066 | -20.629 | | 19.037 | -7.255 | -11.226 | | | 5.151 |
| -17.313 | 8.322 | 12.470 | | 19.678 | -9.518 | 26.177 | | -24.743 | -18.347 | -23.255 | | | -1.654 |
| 10.429 | -20.144 | -11.668 | | -29.948 | -26.434 | 15.026 | | 21.558 | -11.786 | -22.299 | | | 20.928 |
| 20.236 | -15.660 | 3.288 | | -9.367 | 18.266 | -0.111 | | 0.404 | 20.700 | -3.897 | | | -20.033 |

| | | | | | | | | | | | | | |
|---|---|---|---|---|---|---|---|---|---|---|---|---|---|
| 14.489 | 10.519 | -13.432 | | -29.287 | -7.526 | -26.361 | | 11.028 | -6.575 | 16.643 | | | 26.714 |
| -9.471 | -3.170 | -15.114 | | -16.368 | 26.876 | 27.805 | | -6.319 | -4.314 | -7.974 | | | -0.015 |
| -8.046 | -16.968 | -8.594 | | 17.194 | 3.302 | 11.201 | | 15.263 | -13.381 | 20.206 | | | 1.913 |
| 8.941 | -6.716 | -5.998 | | 12.551 | -2.102 | 25.712 | | -1.819 | 7.874 | 17.923 | | | -7.929 |
| -5.311 | 20.536 | -14.420 | | 7.369 | -8.583 | -25.350 | | -6.972 | 15.914 | 26.213 | | | -18.810 |
| 0.020 | -0.222 | | | -19.971 | -25.526 | 14.588 | | -10.836 | -8.237 | -7.561 | | | -19.393 |
| 11.424 | 21.032 | | | 14.965 | -24.169 | -27.159 | | 4.025 | 21.237 | 22.133 | | | 25.076 |
| 10.880 | 13.185 | | | 15.352 | 17.389 | 23.717 | | -24.265 | 29.145 | -6.247 | | | 22.534 |
| -24.667 | 27.149 | | | 14.550 | -25.858 | -14.264 | | -2.552 | -10.176 | -1.960 | | | -5.957 |
| -1.401 | 16.502 | | | 13.471 | -16.989 | -17.102 | | 19.263 | 1.091 | -22.725 | | | -4.385 |
| -6.820 | 25.223 | | | -14.053 | 15.220 | -23.748 | | -21.547 | -18.886 | -5.697 | | | -7.727 |
| -13.615 | 19.972 | | | -22.432 | -24.953 | -4.236 | | 25.601 | -21.184 | -11.116 | | | 3.498 |
| -8.763 | -13.921 | | | 26.834 | 1.139 | 24.682 | | 1.684 | 30.582 | 25.114 | | | -19.017 |
| -17.810 | -27.776 | | | -27.063 | -9.164 | 6.005 | | 26.238 | 4.950 | 1.257 | | | -9.429 |
| -17.085 | -11.680 | | | 10.751 | -10.159 | -20.588 | | 4.289 | -16.880 | 22.480 | | | 7.372 |
| -4.682 | -0.232 | | | -12.775 | 2.380 | 26.577 | | 8.309 | 13.758 | -4.509 | | | -18.280 |
| 12.237 | 21.934 | | | -20.703 | 3.457 | -1.406 | | 7.564 | -8.280 | -13.480 | | | 17.914 |
| 2.299 | 18.762 | | | 21.809 | -24.446 | -22.182 | | -0.838 | -1.107 | -14.292 | | | -2.767 |
| -15.178 | -20.520 | | | 13.836 | -22.681 | -19.407 | | -7.120 | -22.376 | -25.944 | | | -25.980 |
| -20.790 | -18.787 | | | -7.384 | -2.906 | -24.663 | | -4.853 | 14.156 | -10.887 | | | 0.742 |
| 20.638 | 6.351 | | | -25.218 | 23.268 | 16.622 | | -22.265 | -11.299 | -15.224 | | | 3.515 |
| 19.157 | -0.538 | | | 17.968 | 17.497 | -16.639 | | -16.519 | -18.317 | -6.413 | | | 21.842 |
| -10.590 | 13.178 | | | 7.213 | 6.853 | -11.311 | | 18.636 | -1.552 | 23.176 | | | -8.139 |
| 5.413 | 11.834 | | | -18.280 | -19.847 | 8.948 | | -6.257 | -15.850 | 26.407 | | | 6.515 |
| 13.930 | 9.493 | | | -6.156 | 21.758 | -4.938 | | 12.214 | -13.909 | 2.837 | | | -0.124 |
| -12.413 | 3.121 | | | -14.496 | -20.448 | -25.689 | | -9.322 | 24.652 | 1.236 | | | 18.337 |
| -18.388 | -7.224 | | | 2.065 | 19.867 | 8.024 | | -7.508 | -16.415 | -5.675 | | | -3.812 |
| -4.006 | -11.049 | | | 12.177 | 6.406 | 19.362 | | -19.883 | -3.228 | -17.904 | | | -21.494 |
| 7.721 | 0.096 | | | -2.374 | -14.097 | 24.301 | | 8.316 | -27.983 | 25.250 | | | 0.133 |
| 1.601 | -15.748 | | | 30.929 | -25.219 | 7.240 | | -13.097 | -12.387 | 15.548 | | | -24.944 |
| 0.738 | -5.596 | | | -14.836 | -9.600 | 25.353 | | -28.291 | -4.069 | 26.804 | | | 12.164 |
| -0.473 | 10.065 | | | -21.920 | -7.768 | -3.686 | | -9.874 | -12.051 | 11.338 | | | -1.917 |
| 21.067 | 27.856 | | | 19.222 | -1.396 | 21.435 | | -21.628 | 10.391 | 1.842 | | | -21.831 |
| -19.421 | 15.076 | | | -13.554 | 23.245 | -24.913 | | -16.890 | 11.495 | 16.247 | | | 20.869 |
| -21.772 | 5.312 | | | 5.754 | 1.879 | 16.745 | | 6.692 | -2.605 | 12.786 | | | 25.077 |
| 3.003 | -3.990 | | | 9.471 | -5.991 | 18.902 | | 26.616 | 30.013 | 8.300 | | | 2.609 |

| | | | | | | | | | | | | | |
|---|---|---|---|---|---|---|---|---|---|---|---|---|---|
| 0.120 | 7.420 | | | 11.041 | -23.450 | -5.810 | | -4.543 | -6.953 | -25.714 | | | 4.305 |
| -6.202 | 17.334 | | | 4.407 | 12.750 | 20.023 | | -29.008 | -17.673 | -18.690 | | | 10.158 |
| -10.303 | 14.570 | | | -19.226 | 2.963 | -4.901 | | -18.991 | 7.510 | 23.583 | | | 6.584 |
| -19.602 | -21.470 | | | 6.633 | -17.800 | 6.904 | | -27.184 | -2.368 | 3.911 | | | -2.281 |
| -24.664 | -4.984 | | | 15.042 | 18.268 | -20.446 | | -17.863 | -23.048 | -25.374 | | | -11.530 |
| -24.310 | 11.238 | | | 30.575 | -3.855 | 21.842 | | -8.983 | -25.412 | -12.025 | | | 10.644 |
| 22.866 | -11.133 | | | -24.319 | 4.555 | -2.738 | | -10.453 | 22.661 | -7.610 | | | 7.223 |
| 18.649 | 20.670 | | | -16.910 | -11.803 | -16.564 | | -23.164 | -12.696 | -24.123 | | | 1.884 |
| 1.568 | 12.129 | | | -9.188 | -12.896 | -19.862 | | 20.592 | -8.709 | -18.582 | | | 13.808 |
| -9.923 | -4.122 | | | -16.480 | 20.008 | 17.850 | | -6.765 | 26.167 | 0.225 | | | 11.720 |
| 3.492 | -14.913 | | | -27.824 | -21.635 | 23.068 | | -15.384 | -18.366 | 26.797 | | | 9.353 |
| 0.214 | -20.810 | | | 5.804 | -19.177 | 18.578 | | -7.716 | -11.065 | -17.110 | | | -12.961 |
| -0.361 | 20.164 | | | 15.979 | -2.924 | 26.083 | | 29.155 | 3.881 | -13.036 | | | -23.537 |
| 4.762 | -9.398 | | | -9.671 | -19.914 | 24.751 | | -9.013 | 21.888 | -1.577 | | | -24.178 |
| -6.750 | 22.038 | | | 11.394 | -4.507 | -4.711 | | -26.787 | 4.700 | -2.264 | | | -17.329 |
| 18.679 | 20.904 | | | 20.286 | 0.540 | -26.359 | | -21.182 | 8.891 | -20.449 | | | 16.109 |
| 23.184 | -18.052 | | | 18.538 | -3.698 | -9.064 | | -4.961 | 19.614 | 10.470 | | | -3.180 |
| -23.346 | -13.201 | | | -25.514 | -17.166 | -23.270 | | 0.044 | -3.059 | -9.074 | | | 8.396 |
| 25.149 | -21.976 | | | -20.401 | -23.086 | -0.271 | | 21.797 | -2.076 | -12.332 | | | 2.906 |
| -3.682 | -16.109 | | | 21.260 | -21.059 | -7.079 | | 24.474 | 6.268 | 13.079 | | | -20.347 |
| 20.407 | 23.047 | | | -18.593 | 4.002 | -24.109 | | -6.390 | 4.671 | 25.551 | | | -11.880 |
| 22.950 | 17.326 | | | -1.007 | 7.286 | 10.548 | | -20.574 | -6.849 | -3.963 | | | -5.644 |
| 21.653 | 10.249 | | | -13.880 | -3.650 | -21.115 | | 16.299 | -4.783 | -0.056 | | | 6.410 |
| 5.970 | 20.091 | | | 7.244 | 7.582 | -13.336 | | 6.762 | 7.180 | -21.764 | | | 16.087 |
| -1.856 | 15.781 | | | -19.451 | 17.515 | -24.858 | | 20.279 | -7.843 | -2.722 | | | -7.704 |
| 13.539 | 13.783 | | | -25.010 | -19.552 | 4.959 | | -8.058 | -3.376 | -19.867 | | | -6.271 |
| 21.714 | -18.277 | | | 24.023 | -0.428 | 16.009 | | 26.183 | 0.252 | 7.399 | | | -19.943 |
| 0.485 | 18.999 | | | 26.428 | 3.642 | 18.898 | | 21.214 | -1.196 | 17.627 | | | -22.270 |
| 9.453 | 28.734 | | | -19.981 | -14.441 | -21.564 | | -19.630 | -15.674 | -19.373 | | | 10.718 |
| -20.844 | 6.711 | | | -16.602 | -21.416 | 17.909 | | 29.286 | 26.034 | -2.967 | | | -12.863 |
| 20.885 | 3.208 | | | 1.642 | -17.674 | -24.051 | | -22.088 | 1.056 | -0.547 | | | -0.126 |
| -1.014 | -21.269 | | | -8.804 | -22.127 | -1.741 | | 26.472 | -6.261 | -4.043 | | | 12.785 |
| -22.891 | -2.258 | | | -25.768 | -21.661 | 21.652 | | -14.790 | 21.423 | 16.533 | | | -3.918 |
| -0.043 | 14.502 | | | -14.157 | -2.670 | -2.499 | | -21.640 | -0.261 | | | | -15.920 |
| -11.814 | -3.885 | | | 9.694 | -25.267 | 20.578 | | -24.272 | 16.402 | | | | 0.618 |
| 24.567 | -20.668 | | | -23.773 | 6.360 | -18.717 | | 22.989 | -19.508 | | | | -10.162 |

| | | | | | | | | | | | | | |
|---|---|---|---|---|---|---|---|---|---|---|---|---|---|
| 1.426 | -10.593 | | | -20.915 | 20.326 | 10.438 | | 29.743 | 4.958 | | | | 1.785 |
| -16.827 | -12.681 | | | -24.553 | -9.126 | 13.290 | | 27.678 | 22.178 | | | | 4.799 |
| -19.535 | 10.691 | | | -1.456 | -4.947 | 24.041 | | -25.642 | -18.530 | | | | 18.828 |
| 7.244 | -8.428 | | | 18.650 | -14.607 | -15.838 | | -18.232 | 8.765 | | | | -7.836 |
| 8.549 | -21.926 | | | 20.512 | 22.477 | -23.003 | | -26.380 | 0.904 | | | | 9.638 |
| 14.011 | -1.456 | | | 17.511 | -13.045 | -2.060 | | 13.231 | 7.235 | | | | -12.829 |
| 1.046 | -11.500 | | | -7.227 | -11.947 | -14.114 | | 16.827 | -18.292 | | | | 5.483 |
| -25.290 | 1.317 | | | 18.312 | 24.750 | 7.546 | | -3.387 | 3.828 | | | | 2.289 |
| -4.596 | -7.805 | | | 16.365 | -2.829 | -18.831 | | -20.812 | -28.925 | | | | -8.146 |
| -8.896 | 18.934 | | | -13.989 | 25.200 | 22.837 | | 23.029 | -17.187 | | | | 10.314 |
| 23.056 | -12.743 | | | -15.790 | 21.768 | -23.928 | | 7.604 | 0.212 | | | | 6.114 |
| -1.266 | -14.380 | | | -25.882 | 18.265 | -16.217 | | -29.520 | 15.701 | | | | -6.241 |
| -16.760 | 10.081 | | | -4.905 | -14.333 | 4.828 | | -11.187 | -6.399 | | | | 9.156 |
| 22.964 | 8.093 | | | -26.647 | -13.202 | 14.593 | | -18.807 | -4.679 | | | | 17.032 |
| 5.923 | 29.287 | | | 15.472 | -5.332 | 6.087 | | -21.880 | -1.020 | | | | -7.058 |
| 17.920 | -17.683 | | | -12.913 | 4.599 | -12.994 | | -24.608 | 19.956 | | | | 8.680 |
| 3.425 | 24.349 | | | -17.408 | 8.840 | -23.015 | | -26.736 | -14.311 | | | | -25.136 |
| 26.980 | 21.221 | | | -11.460 | -4.187 | | | -12.570 | 13.664 | | | | 0.886 |
| -26.252 | -9.898 | | | -16.145 | 12.588 | | | -9.143 | -5.951 | | | | -16.768 |
| 9.060 | -21.000 | | | 17.647 | -23.475 | | | -7.441 | -8.094 | | | | 2.004 |
| 4.051 | -11.116 | | | 14.666 | -7.967 | | | 17.530 | 2.952 | | | | -9.371 |
| -24.446 | -6.555 | | | -8.078 | 4.476 | | | -6.862 | -2.932 | | | | 2.853 |
| -10.353 | 27.914 | | | -26.920 | -16.716 | | | -14.329 | 8.951 | | | | -10.249 |
| -11.734 | 16.791 | | | -8.483 | 16.975 | | | -22.043 | 7.529 | | | | 0.269 |
| 22.526 | -4.143 | | | 13.285 | 8.041 | | | 20.298 | -14.972 | | | | -2.309 |
| -25.834 | -18.486 | | | -21.388 | -1.354 | | | 26.945 | -7.129 | | | | 6.758 |
| 6.051 | -10.841 | | | -24.183 | 23.733 | | | 23.205 | 0.556 | | | | 1.132 |
| -7.837 | -2.700 | | | -10.281 | -21.805 | | | -20.041 | -11.277 | | | | -17.268 |
| 21.608 | -19.487 | | | -16.742 | -7.293 | | | -21.680 | -23.341 | | | | 18.472 |
| 6.834 | -23.634 | | | 11.048 | 26.104 | | | -25.964 | 18.415 | | | | -15.644 |
| -16.901 | -21.514 | | | -9.454 | -10.657 | | | 3.723 | 16.742 | | | | 7.739 |
| -16.318 | 5.685 | | | -6.283 | -22.783 | | | -25.604 | -21.755 | | | | -12.466 |
| -18.837 | 14.347 | | | -15.059 | -27.102 | | | 28.420 | 6.152 | | | | -1.717 |
| 27.290 | -7.714 | | | 20.987 | 15.622 | | | -10.195 | -19.345 | | | | -0.945 |
| 24.180 | 17.040 | | | -0.228 | 4.259 | | | -22.414 | 19.846 | | | | -6.320 |
| 3.553 | 3.312 | | | -14.688 | 25.637 | | | -20.202 | -2.307 | | | | -0.301 |

| | | | | | | | | | | | | | |
|---|---|---|---|---|---|---|---|---|---|---|---|---|---|
| 22.982 | -19.866 | | | -7.667 | 5.078 | | | 23.817 | -8.605 | | | | 5.507 |
| 12.061 | 8.853 | | | 7.484 | 13.106 | | | -3.246 | -5.454 | | | | 3.197 |
| 1.580 | 23.371 | | | -16.513 | 17.474 | | | 20.973 | -17.448 | | | | -8.068 |
| 18.897 | 11.907 | | | -11.980 | -11.544 | | | -9.691 | -14.471 | | | | -4.556 |
| -0.495 | 16.004 | | | -24.116 | -12.693 | | | 2.004 | -9.406 | | | | 11.641 |
| 13.873 | -11.224 | | | -25.233 | -19.591 | | | -6.212 | 11.808 | | | | 9.308 |
| 11.134 | -7.245 | | | 14.148 | -0.745 | | | 12.081 | -14.190 | | | | -2.689 |
| -2.855 | 21.039 | | | 20.184 | 11.711 | | | 9.877 | -25.313 | | | | -3.693 |
| -21.896 | -12.063 | | | 6.121 | -24.144 | | | -31.138 | -21.256 | | | | -7.251 |
| 20.134 | 6.913 | | | -22.818 | -3.734 | | | 17.428 | 10.204 | | | | 2.055 |
| -27.073 | 4.427 | | | 8.704 | 20.208 | | | -21.849 | 14.206 | | | | 22.619 |
| 15.309 | -3.028 | | | 7.993 | -1.700 | | | 25.692 | 0.351 | | | | 20.662 |
| -9.333 | 26.192 | | | 5.648 | 3.773 | | | 20.301 | -12.077 | | | | 5.898 |
| -8.652 | -16.230 | | | -27.274 | -19.255 | | | 19.485 | 6.695 | | | | 5.251 |
| 17.812 | -14.346 | | | -5.994 | 25.479 | | | 11.631 | -2.769 | | | | 17.205 |
| 2.687 | -15.711 | | | -19.058 | 27.397 | | | -11.972 | -7.297 | | | | 8.441 |
| -7.817 | 17.871 | | | 23.041 | 2.843 | | | 30.791 | -14.171 | | | | -3.211 |
| 10.002 | -21.964 | | | -10.642 | -15.406 | | | 21.033 | 19.155 | | | | 14.172 |
| -23.620 | -6.508 | | | 9.197 | -7.303 | | | -25.801 | 2.326 | | | | 4.338 |
| 7.579 | 9.137 | | | 11.250 | 24.773 | | | -12.790 | 3.256 | | | | -9.531 |
| -27.819 | 5.196 | | | -4.957 | -21.962 | | | 13.955 | 22.181 | | | | -1.119 |
| -1.047 | -4.761 | | | -23.835 | 25.695 | | | 13.269 | -16.597 | | | | -2.302 |
| 29.997 | -5.325 | | | -8.243 | 14.100 | | | -4.816 | -20.242 | | | | -11.065 |
| -23.215 | 24.550 | | | 14.256 | -2.003 | | | 22.601 | 1.092 | | | | -8.342 |
| 29.147 | -8.632 | | | 6.377 | -23.799 | | | -9.181 | 0.026 | | | | 7.421 |
| 5.656 | -21.318 | | | 13.621 | 4.357 | | | -24.825 | -12.686 | | | | 0.523 |
| 7.907 | 7.505 | | | 5.432 | 10.405 | | | -24.483 | -1.503 | | | | -20.930 |
| 23.067 | -18.676 | | | -17.093 | -27.362 | | | 31.059 | -12.231 | | | | 12.333 |
| 18.889 | 15.773 | | | -23.591 | -20.182 | | | -10.332 | 5.526 | | | | 4.892 |
| -27.193 | -0.793 | | | -18.067 | 7.315 | | | -7.823 | -24.168 | | | | -12.927 |
| 20.302 | 13.969 | | | -21.524 | 3.289 | | | 25.138 | 14.688 | | | | -17.174 |
| 11.642 | 22.194 | | | 18.091 | -12.411 | | | 16.592 | -6.170 | | | | -12.496 |
| -28.239 | 13.540 | | | -9.018 | -25.344 | | | -3.022 | 2.021 | | | | -6.733 |
| 5.385 | -3.253 | | | -5.840 | 1.799 | | | 23.343 | 21.961 | | | | -1.591 |
| 20.780 | -7.977 | | | -19.192 | -24.718 | | | -2.364 | -9.240 | | | | 11.054 |
| 16.134 | -20.582 | | | -13.677 | -23.339 | | | -7.547 | 14.686 | | | | 6.049 |

| | | | | | | | | | | | | | |
|---|---|---|---|---|---|---|---|---|---|---|---|---|---|
| -0.126 | -4.654 | | | 15.993 | -7.492 | | | -27.210 | -12.998 | | | | 16.910 |
| -26.552 | -6.955 | | | -19.492 | 19.130 | | | -20.246 | -14.608 | | | | -18.203 |
| -23.370 | -14.190 | | | -3.942 | -5.716 | | | 20.552 | -8.943 | | | | -0.861 |
| 29.821 | -3.878 | | | 22.581 | -7.238 | | | 26.529 | 3.707 | | | | -4.423 |
| -2.301 | 3.322 | | | 7.570 | -25.687 | | | 8.989 | 19.430 | | | | 5.370 |
| -4.915 | -0.641 | | | -6.357 | 22.376 | | | 5.122 | 7.450 | | | | 27.613 |
| 17.225 | 7.124 | | | -10.832 | -18.745 | | | -12.471 | -6.282 | | | | 8.493 |
| 30.193 | 21.051 | | | 12.081 | 14.867 | | | -29.554 | -6.900 | | | | -3.840 |
| -3.220 | -2.131 | | | -16.113 | -8.653 | | | 19.510 | -24.869 | | | | -7.858 |
| 4.276 | -9.835 | | | 17.732 | -28.586 | | | 22.556 | -19.884 | | | | 2.222 |
| 17.405 | -18.201 | | | 1.277 | 15.819 | | | 19.731 | -3.525 | | | | -5.762 |
| 21.720 | -5.716 | | | 13.975 | -17.373 | | | -10.351 | 1.283 | | | | -23.394 |
| 19.437 | 14.438 | | | 3.569 | 19.390 | | | -15.647 | 21.227 | | | | 16.832 |
| -1.307 | -4.012 | | | 7.379 | 1.809 | | | -4.676 | 21.983 | | | | 21.163 |
| -25.075 | -5.954 | | | 14.556 | -10.625 | | | -13.417 | 9.787 | | | | 18.856 |
| 5.542 | -6.318 | | | -19.416 | 9.833 | | | -15.922 | 11.325 | | | | -0.229 |
| 29.080 | -3.387 | | | -10.877 | 3.558 | | | -20.279 | 13.812 | | | | -3.430 |
| 6.049 | 2.305 | | | -7.889 | -27.333 | | | -30.096 | -10.002 | | | | -10.028 |
| -1.097 | 5.247 | | | -20.654 | 8.923 | | | 17.416 | 19.367 | | | | -25.613 |
| 27.759 | 3.418 | | | 3.226 | -9.029 | | | 4.043 | 19.709 | | | | 0.930 |
| 24.523 | -1.405 | | | -11.870 | -21.388 | | | -16.897 | 14.335 | | | | 8.930 |
| 22.454 | -2.857 | | | -4.846 | 12.083 | | | 12.650 | 1.816 | | | | -7.228 |
| -4.098 | -13.828 | | | -5.355 | -1.855 | | | 5.715 | -21.809 | | | | 11.333 |
| -7.938 | 27.398 | | | -12.892 | 16.478 | | | -9.452 | 9.053 | | | | 21.744 |
| 21.392 | 5.926 | | | -14.389 | 23.126 | | | 21.276 | -9.858 | | | | 14.874 |
| -30.407 | -0.958 | | | -2.471 | -24.592 | | | -22.545 | -0.817 | | | | -4.645 |
| -19.592 | -7.133 | | | 0.379 | 28.610 | | | 8.777 | -16.446 | | | | -6.129 |
| 30.804 | -4.930 | | | -16.039 | 27.894 | | | -14.510 | 0.025 | | | | 6.191 |
| 20.103 | -0.365 | | | -23.834 | 24.878 | | | -4.464 | -2.460 | | | | 24.597 |
| -20.241 | -8.587 | | | -22.920 | -12.180 | | | -24.381 | 7.157 | | | | 0.242 |
| -1.828 | -4.195 | | | 25.963 | -29.506 | | | -6.670 | -8.688 | | | | 26.580 |
| 18.311 | -18.754 | | | 5.070 | -1.884 | | | -21.254 | -3.291 | | | | 21.805 |
| 4.575 | 3.060 | | | 14.517 | 7.443 | | | 23.514 | -3.858 | | | | 10.149 |
| -20.983 | -1.348 | | | 23.438 | 13.081 | | | 8.317 | -18.892 | | | | 4.907 |
| -5.874 | 12.531 | | | 2.990 | -24.219 | | | -23.766 | 10.051 | | | | -26.554 |
| 30.683 | -4.858 | | | -26.813 | 10.775 | | | -6.216 | -20.195 | | | | 11.076 |

| | | | | | | | | | | | | | |
|---|---|---|---|---|---|---|---|---|---|---|---|---|---|
| -30.293 | 6.616 | | | -7.927 | 19.969 | | | -12.257 | 8.419 | | | | -3.739 |
| -2.184 | -12.330 | | | -9.691 | -9.154 | | | 23.102 | 5.236 | | | | -12.666 |
| 19.993 | 0.383 | | | -17.653 | -26.158 | | | 20.139 | 6.107 | | | | -8.249 |
| -24.387 | 14.327 | | | 15.671 | 9.928 | | | -10.691 | -19.317 | | | | -6.986 |
| 29.787 | 22.170 | | | -11.171 | 6.028 | | | 10.634 | -0.590 | | | | 10.774 |
| -8.901 | -6.314 | | | 25.875 | 16.223 | | | -7.464 | -6.338 | | | | -3.602 |
| 0.759 | -4.055 | | | 12.894 | 4.803 | | | -12.787 | 5.456 | | | | 20.950 |
| 26.956 | 16.669 | | | -24.349 | -28.520 | | | 13.811 | 13.624 | | | | -12.078 |
| 13.874 | -9.833 | | | 17.516 | 17.497 | | | 17.502 | -3.079 | | | | -5.650 |
| -1.842 | -1.114 | | | 10.117 | -15.517 | | | 16.479 | -11.591 | | | | 23.714 |
| -17.961 | -1.938 | | | -22.562 | 7.988 | | | 13.281 | -4.152 | | | | -26.254 |
| 11.728 | 4.988 | | | -7.861 | -9.972 | | | 1.975 | 1.879 | | | | 19.615 |
| -25.547 | 0.282 | | | -25.351 | -19.891 | | | -7.822 | -14.498 | | | | 3.931 |
| -22.881 | 2.337 | | | -14.421 | 1.252 | | | -17.721 | -13.091 | | | | -17.325 |
| 23.861 | -4.778 | | | -10.447 | 26.446 | | | -29.529 | 26.044 | | | | 3.462 |
| 21.416 | -7.203 | | | 6.635 | 27.211 | | | 24.245 | 7.111 | | | | 4.506 |
| -3.169 | -4.103 | | | -19.094 | 27.670 | | | -7.038 | 5.945 | | | | -10.142 |
| 19.482 | -4.301 | | | -18.486 | -7.015 | | | 19.253 | 0.032 | | | | -0.523 |
| 28.421 | 18.400 | | | 8.308 | -7.423 | | | 6.696 | 12.198 | | | | -6.138 |
| 30.323 | 0.196 | | | 3.589 | -27.302 | | | -11.645 | -9.402 | | | | -20.229 |
| -18.879 | -2.441 | | | -2.865 | 26.889 | | | -14.079 | -13.901 | | | | 2.322 |
| -12.403 | 7.737 | | | -11.114 | 15.158 | | | -21.669 | -22.035 | | | | 1.460 |
| -23.246 | 15.687 | | | 4.682 | 16.284 | | | 25.052 | 22.603 | | | | -21.958 |
| -7.688 | -19.478 | | | -20.148 | -29.180 | | | -8.286 | 6.667 | | | | 8.813 |
| 19.921 | -22.490 | | | -3.790 | -2.864 | | | -7.653 | -4.810 | | | | 26.629 |
| -12.959 | 23.229 | | | -16.683 | -21.312 | | | -12.995 | -1.586 | | | | 0.242 |
| -29.533 | -23.205 | | | 11.686 | 30.291 | | | -1.379 | 11.558 | | | | -7.974 |
| -14.304 | 5.549 | | | -1.922 | 0.861 | | | 27.027 | -7.094 | | | | -1.803 |
| 21.459 | -8.353 | | | -9.458 | -15.369 | | | 15.366 | 21.225 | | | | 19.153 |
| 7.796 | -2.034 | | | 0.604 | 8.728 | | | -9.236 | 20.424 | | | | 11.173 |
| -9.023 | -5.635 | | | 0.228 | 21.452 | | | -28.214 | 2.614 | | | | 2.110 |
| 11.758 | 1.120 | | | -15.949 | -16.727 | | | 17.988 | -16.343 | | | | -3.418 |
| -5.684 | -2.601 | | | 7.133 | 5.822 | | | -12.335 | -18.436 | | | | 5.778 |
| -3.932 | -25.366 | | | 0.972 | 10.104 | | | -4.878 | -23.504 | | | | -16.928 |
| 11.273 | -3.047 | | | -11.829 | 11.700 | | | -5.460 | 19.660 | | | | 15.961 |
| 8.153 | -0.913 | | | -3.558 | 17.439 | | | -13.449 | -7.845 | | | | -9.443 |

| | | | | | | | | | | | | | |
|---|---|---|---|---|---|---|---|---|---|---|---|---|---|
| -26.989 | -1.566 | | | -0.643 | 7.449 | | | 18.408 | 4.813 | | | | -5.463 |
| 26.200 | -7.556 | | | -0.259 | -24.635 | | | -26.659 | 15.628 | | | | -6.132 |
| -15.849 | -2.263 | | | -3.253 | 6.947 | | | -3.183 | -14.185 | | | | -9.017 |
| -18.329 | -21.512 | | | -7.035 | 23.266 | | | -18.096 | 22.640 | | | | 9.690 |
| -21.737 | 1.973 | | | 25.334 | -16.514 | | | -20.955 | 6.037 | | | | -14.739 |
| -5.002 | 4.903 | | | 0.787 | 21.093 | | | 18.784 | 0.715 | | | | -2.575 |
| -24.164 | -10.654 | | | 6.980 | 12.414 | | | -1.111 | 18.297 | | | | 7.649 |
| 1.350 | -24.105 | | | -17.621 | 30.477 | | | -28.955 | -17.154 | | | | -0.258 |
| 0.549 | -13.877 | | | -5.629 | 8.910 | | | 5.668 | -13.105 | | | | -5.076 |
| 26.603 | 0.963 | | | -6.541 | 19.554 | | | -4.489 | -1.227 | | | | 15.511 |
| 18.885 | -11.060 | | | -10.449 | 9.798 | | | -6.956 | 8.700 | | | | 11.031 |
| 22.444 | 16.383 | | | 3.168 | 6.073 | | | 13.356 | 7.700 | | | | -17.932 |
| -11.191 | -0.545 | | | -1.402 | -29.645 | | | 22.383 | -16.422 | | | | -9.701 |
| 10.037 | 7.322 | | | -0.463 | -27.275 | | | 4.307 | -7.615 | | | | -9.459 |
| 27.101 | -5.473 | | | 4.669 | 10.899 | | | 12.489 | 12.450 | | | | 6.657 |
| -6.029 | -8.132 | | | -17.065 | 26.612 | | | 1.816 | 7.450 | | | | -7.002 |
| -1.916 | -4.633 | | | -18.493 | -21.807 | | | 20.272 | 24.731 | | | | -24.929 |
| 28.690 | 15.973 | | | 22.104 | 28.321 | | | 13.755 | 9.377 | | | | 1.310 |
| 20.975 | 5.630 | | | -8.156 | 21.104 | | | 6.537 | -19.370 | | | | -5.808 |
| -22.495 | -4.048 | | | -14.882 | 9.285 | | | -14.666 | -4.593 | | | | 20.072 |
| -22.017 | -20.348 | | | 10.075 | -9.531 | | | -10.255 | -24.770 | | | | -19.689 |
| -10.257 | -9.916 | | | 13.484 | -18.564 | | | -24.729 | -23.995 | | | | -28.390 |
| 22.487 | -5.161 | | | -22.312 | 22.297 | | | -7.670 | 18.383 | | | | -0.657 |
| 25.988 | 11.268 | | | 13.305 | 18.858 | | | -19.769 | 17.488 | | | | 11.130 |
| 11.867 | -19.983 | | | 8.505 | -15.117 | | | 17.415 | -22.620 | | | | 17.692 |
| -1.670 | -17.855 | | | 20.278 | 27.972 | | | -21.135 | -3.071 | | | | 16.218 |
| -12.928 | 2.685 | | | 2.132 | -29.585 | | | 20.772 | 26.498 | | | | -11.435 |
| -20.312 | -4.533 | | | 1.599 | 17.163 | | | 10.255 | -18.577 | | | | 9.861 |
| -25.117 | -6.803 | | | -23.869 | -20.609 | | | 7.470 | -4.995 | | | | 2.328 |
| -6.200 | -3.749 | | | -22.938 | -14.622 | | | -5.224 | 6.246 | | | | 27.047 |
| 10.946 | -13.727 | | | 11.692 | 24.090 | | | 23.322 | 5.981 | | | | 18.857 |
| -22.679 | 25.843 | | | -9.092 | 28.067 | | | 11.132 | -3.773 | | | | 3.758 |
| 4.172 | 22.134 | | | -17.129 | -15.110 | | | -3.620 | -7.983 | | | | -0.711 |
| 24.434 | 16.729 | | | 0.588 | -17.206 | | | 19.376 | -23.913 | | | | -1.279 |
| 9.492 | -24.364 | | | -15.687 | 21.919 | | | 13.928 | -4.435 | | | | -9.073 |
| -3.437 | -10.248 | | | 9.347 | -23.477 | | | 3.265 | -5.448 | | | | 9.414 |

| | | | | | | | | | | | | | |
|---|---|---|---|---|---|---|---|---|---|---|---|---|---|
| -4.197 | -21.329 | | | -0.304 | -28.361 | | | 25.335 | -6.073 | | | | -5.244 |
| 6.353 | 1.600 | | | -5.367 | -3.068 | | | 22.972 | 6.642 | | | | -25.424 |
| 21.125 | -9.840 | | | 25.018 | -19.069 | | | 10.453 | 9.072 | | | | 7.212 |
| -25.621 | -4.832 | | | 10.537 | -29.234 | | | 9.521 | 17.237 | | | | 25.301 |
| -13.196 | -23.433 | | | 1.378 | 27.678 | | | -29.119 | -19.549 | | | | -11.000 |
| -12.075 | -12.364 | | | -7.001 | -21.653 | | | 1.861 | -21.610 | | | | -5.986 |
| -5.082 | 13.407 | | | 19.926 | -23.814 | | | -23.226 | 8.464 | | | | 22.005 |
| 18.662 | 19.915 | | | 24.497 | -8.861 | | | 9.722 | 26.060 | | | | 1.153 |
| 14.633 | -8.803 | | | -19.233 | 17.509 | | | -8.364 | 5.744 | | | | -28.166 |
| 26.016 | 23.599 | | | 4.621 | -13.748 | | | -16.216 | -13.264 | | | | -1.792 |
| -5.814 | 6.413 | | | 9.184 | 19.035 | | | 11.587 | -18.143 | | | | -0.454 |
| 1.610 | 13.699 | | | 2.009 | -25.157 | | | -10.673 | -6.856 | | | | -12.471 |
| 6.915 | 5.478 | | | -9.058 | 2.672 | | | 17.497 | 7.345 | | | | 2.178 |
| 11.723 | -2.095 | | | -2.297 | 25.944 | | | 8.406 | -7.591 | | | | -26.375 |
| -9.508 | -5.690 | | | -5.742 | 24.284 | | | 24.727 | -9.791 | | | | -27.684 |
| -4.940 | -27.896 | | | 22.747 | -29.740 | | | 21.646 | -19.089 | | | | -8.859 |
| -10.968 | 6.010 | | | -19.467 | 29.656 | | | -5.040 | 5.528 | | | | 28.256 |
| 27.431 | 15.104 | | | 7.371 | -3.525 | | | -13.762 | -5.956 | | | | 13.704 |
| 6.756 | -18.049 | | | 14.969 | 29.251 | | | -25.870 | -11.112 | | | | 14.997 |
| -24.836 | -28.763 | | | -11.041 | 18.028 | | | 19.539 | -23.394 | | | | -16.404 |
| 4.101 | -5.294 | | | -17.847 | -29.269 | | | 18.181 | -5.062 | | | | 17.670 |
| 2.322 | 25.990 | | | 13.890 | 22.707 | | | -22.972 | 3.242 | | | | 0.358 |
| -14.604 | 19.210 | | | 5.022 | -23.820 | | | 11.460 | -22.499 | | | | 5.060 |
| -9.664 | -8.719 | | | -1.761 | -7.431 | | | 9.088 | 24.280 | | | | -6.552 |
| 12.548 | 9.245 | | | 1.059 | 26.228 | | | -18.822 | 11.105 | | | | 25.824 |
| -8.340 | 3.719 | | | -2.954 | -21.651 | | | 3.186 | 0.310 | | | | 17.915 |
| -4.350 | 6.971 | | | 11.036 | -0.223 | | | 2.133 | 19.973 | | | | -3.297 |
| 23.035 | 10.370 | | | -3.707 | -1.291 | | | 9.184 | -14.353 | | | | 8.993 |
| 21.896 | 8.042 | | | -16.124 | 29.236 | | | 4.121 | 17.632 | | | | 10.655 |
| 4.496 | -29.116 | | | 6.602 | 26.718 | | | -22.648 | -1.014 | | | | 6.668 |
| -15.447 | -24.943 | | | 5.973 | 7.201 | | | -1.457 | -4.553 | | | | 19.922 |
| 0.133 | -22.967 | | | -10.338 | 14.778 | | | 10.863 | 8.663 | | | | 29.303 |
| -25.403 | -11.452 | | | -13.323 | -17.831 | | | 11.827 | -7.096 | | | | -13.316 |
| 0.745 | 23.453 | | | -4.590 | 16.604 | | | 2.671 | -24.862 | | | | 16.179 |
| 5.343 | -11.989 | | | -9.917 | -21.446 | | | 23.429 | 25.622 | | | | 26.032 |
| 26.570 | -15.647 | | | -13.785 | 8.723 | | | 8.433 | 3.317 | | | | -12.929 |

| | | | | | | | | | | | | | |
|---|---|---|---|---|---|---|---|---|---|---|---|---|---|
| 8.970 | -6.035 | | | 15.629 | 22.633 | | | 3.167 | 1.588 | | | | -11.452 |
| 2.743 | 19.146 | | | -22.759 | 10.501 | | | -17.218 | -29.237 | | | | 24.921 |
| -2.508 | 6.733 | | | 2.538 | 28.415 | | | 22.034 | -23.294 | | | | -1.002 |
| -10.430 | 10.508 | | | -2.590 | -23.356 | | | -10.285 | -25.922 | | | | 10.883 |
| -4.797 | -13.921 | | | -19.035 | -22.720 | | | -12.134 | -20.535 | | | | -9.181 |
| 12.508 | -21.806 | | | -21.483 | 25.409 | | | -19.829 | -19.004 | | | | -23.455 |
| 2.384 | -6.613 | | | 6.199 | 22.291 | | | 10.361 | -22.926 | | | | -22.657 |
| -3.249 | 5.338 | | | 4.109 | -24.326 | | | 7.166 | 20.654 | | | | 20.709 |
| -18.535 | 14.398 | | | -0.003 | 8.513 | | | 16.288 | 19.030 | | | | -5.797 |
| 0.397 | -23.502 | | | -0.951 | -11.828 | | | -3.004 | -19.577 | | | | -15.609 |
| -7.953 | 25.744 | | | 8.975 | -22.332 | | | -12.436 | 17.548 | | | | 0.590 |
| 10.082 | -22.139 | | | -12.568 | 7.633 | | | 24.162 | -18.358 | | | | -6.382 |
| -16.923 | 21.508 | | | -0.602 | -1.695 | | | 27.059 | 8.773 | | | | 29.659 |
| 23.950 | 8.484 | | | 9.578 | -17.144 | | | 12.169 | 22.293 | | | | -8.348 |
| 16.057 | -27.801 | | | 3.535 | 13.060 | | | 7.842 | 11.733 | | | | 28.525 |
| -6.875 | 18.786 | | | 2.767 | -8.740 | | | -21.736 | 9.079 | | | | 9.200 |
| 11.287 | 14.325 | | | -7.475 | -25.604 | | | 6.490 | 26.716 | | | | -15.245 |
| 5.635 | -17.305 | | | -16.710 | -20.552 | | | -2.254 | 24.948 | | | | 1.985 |
| 6.108 | -21.118 | | | -23.172 | 22.404 | | | -23.779 | 7.944 | | | | 4.289 |
| 6.846 | 22.377 | | | -18.662 | -20.229 | | | 11.775 | 6.826 | | | | -17.140 |
| -9.941 | 11.340 | | | -25.824 | 1.298 | | | -16.515 | 23.720 | | | | -8.454 |
| 15.022 | -14.429 | | | 15.868 | -8.432 | | | -0.353 | 19.822 | | | | -9.594 |
| 8.085 | -5.299 | | | -4.294 | 26.618 | | | -15.912 | 10.590 | | | | -21.518 |
| -14.702 | 20.954 | | | 3.376 | -18.388 | | | 22.885 | -17.003 | | | | -10.268 |
| -24.881 | -20.010 | | | -7.037 | 24.125 | | | 1.396 | -17.653 | | | | -12.568 |
| 13.397 | 23.058 | | | 5.516 | -2.452 | | | -4.195 | -1.469 | | | | -19.725 |
| 1.696 | -9.641 | | | -22.757 | 22.681 | | | -7.431 | 19.685 | | | | -18.763 |
| -3.455 | 0.562 | | | 17.784 | -14.424 | | | -25.832 | 21.462 | | | | 31.055 |
| 8.967 | -7.609 | | | -0.363 | 12.523 | | | 3.635 | -16.004 | | | | 0.033 |
| -12.324 | -10.896 | | | -14.295 | -22.008 | | | -6.218 | 15.559 | | | | -6.864 |
| 3.961 | 20.018 | | | -21.200 | 8.912 | | | -20.474 | 21.789 | | | | 14.591 |
| -22.791 | -8.741 | | | -7.529 | -23.891 | | | 5.652 | 17.969 | | | | 21.164 |
| -6.558 | 21.551 | | | -8.690 | -8.448 | | | 26.709 | -0.816 | | | | 3.020 |
| 19.233 | -5.461 | | | -10.606 | 0.509 | | | 27.026 | 23.189 | | | | -22.377 |
| -6.285 | 23.696 | | | -23.706 | 25.248 | | | 17.010 | -9.327 | | | | -17.178 |
| 7.019 | -1.603 | | | 8.747 | 24.119 | | | -0.892 | 23.914 | | | | -4.963 |

| | | | | | | | | | | | | | |
|---|---|---|---|---|---|---|---|---|---|---|---|---|---|
| -23.477 | -11.996 | | | 24.190 | -8.553 | | | -12.526 | -19.062 | | | | 23.569 |
| 12.193 | 13.041 | | | -5.922 | -12.714 | | | 1.522 | 10.741 | | | | 7.734 |
| 4.691 | 11.070 | | | 4.772 | -16.827 | | | 3.934 | 24.431 | | | | -18.591 |
| 18.655 | 19.991 | | | 16.145 | -20.585 | | | -9.405 | -21.161 | | | | -5.841 |
| 6.530 | -19.703 | | | -25.198 | -1.244 | | | -0.247 | 10.164 | | | | 17.220 |
| -7.510 | 9.126 | | | 21.637 | -20.971 | | | 4.782 | -29.878 | | | | 22.362 |
| 22.067 | 5.053 | | | 5.727 | -7.141 | | | -7.827 | 4.318 | | | | -6.664 |
| -4.064 | 18.588 | | | 3.437 | 21.723 | | | 21.576 | 25.919 | | | | -16.323 |
| -4.666 | 8.720 | | | -13.099 | -25.077 | | | -1.872 | -22.476 | | | | 20.908 |
| 2.338 | -10.283 | | | -10.055 | 19.226 | | | -10.660 | -6.903 | | | | 15.690 |
| 3.753 | -5.904 | | | -12.783 | 0.398 | | | 13.704 | -29.032 | | | | 21.645 |
| -15.631 | 16.413 | | | -18.597 | 14.195 | | | -19.577 | -18.693 | | | | 11.559 |
| -7.904 | 15.415 | | | -16.438 | -16.646 | | | -3.458 | 21.772 | | | | 14.116 |
| 13.183 | -7.821 | | | -7.946 | -2.778 | | | 5.788 | -3.985 | | | | 18.146 |
| 7.852 | 7.011 | | | -14.241 | -4.496 | | | 11.352 | 16.470 | | | | -7.051 |
| -22.614 | 16.220 | | | -24.058 | 3.988 | | | -12.994 | -14.839 | | | | 19.025 |
| -15.098 | -8.983 | | | -22.580 | 24.857 | | | 6.859 | -17.271 | | | | 28.856 |
| 1.357 | -18.790 | | | -0.664 | 5.155 | | | 17.271 | 22.678 | | | | 10.122 |
| -3.957 | -22.879 | | | 12.683 | -19.365 | | | -14.150 | 15.211 | | | | -23.908 |
| -1.448 | 8.239 | | | 15.078 | -16.006 | | | 0.862 | -0.743 | | | | 15.663 |
| -25.776 | -8.829 | | | -2.461 | 22.352 | | | -4.120 | 29.601 | | | | 12.799 |
| 2.586 | -20.422 | | | 7.400 | -14.457 | | | -26.727 | 16.556 | | | | -6.027 |
| 7.567 | -23.646 | | | -23.207 | -16.744 | | | -21.699 | -25.713 | | | | 9.264 |
| -18.681 | -25.172 | | | 7.949 | 10.169 | | | 10.332 | -18.314 | | | | 24.459 |
| 8.503 | 24.135 | | | 21.961 | 2.535 | | | 8.838 | 11.192 | | | | -26.778 |
| 5.278 | -10.334 | | | -18.773 | -9.164 | | | 1.872 | -5.223 | | | | 3.702 |
| -10.360 | -14.813 | | | 21.570 | -8.183 | | | -5.112 | -15.497 | | | | -7.350 |
| 19.738 | -7.437 | | | -18.368 | -22.298 | | | -7.960 | -24.320 | | | | 28.442 |
| 20.941 | -24.407 | | | -19.169 | 10.988 | | | 20.383 | 24.446 | | | | -22.427 |
| -1.401 | 18.560 | | | -10.944 | 18.143 | | | -3.654 | 27.753 | | | | 16.289 |
| -10.590 | -27.136 | | | -13.695 | 10.076 | | | 22.929 | -17.544 | | | | -20.973 |
| -9.435 | 27.290 | | | -4.064 | 13.681 | | | -7.160 | 22.998 | | | | 5.832 |
| -0.242 | 17.274 | | | 18.740 | 8.889 | | | 7.892 | 22.347 | | | | 22.034 |
| -14.373 | 10.230 | | | 14.747 | -1.694 | | | 4.299 | -8.994 | | | | -6.992 |
| -19.857 | -13.005 | | | -25.840 | | | | 9.036 | -5.771 | | | | -8.705 |
| 8.881 | 20.541 | | | 3.847 | | | | -15.611 | 4.243 | | | | -9.319 |

| | | | | | | | | | | | | | |
|---|---|---|---|---|---|---|---|---|---|---|---|---|---|
| 22.089 | 7.899 | | | -15.630 | | | | -4.647 | -27.453 | | | | 13.095 |
| 2.646 | -12.042 | | | -20.516 | | | | 13.484 | 7.261 | | | | 9.967 |
| -9.134 | 22.727 | | | 21.597 | | | | 2.654 | 10.627 | | | | 20.856 |
| -13.007 | -19.615 | | | -20.739 | | | | -1.233 | 21.112 | | | | 12.106 |
| 4.846 | 26.104 | | | 15.534 | | | | -14.881 | 5.556 | | | | -7.623 |
| 25.468 | 22.041 | | | 13.257 | | | | 8.469 | -7.939 | | | | 14.661 |
| -17.645 | 11.449 | | | 9.260 | | | | 6.991 | -19.485 | | | | 15.504 |
| 9.992 | -11.711 | | | -10.524 | | | | 11.221 | -24.605 | | | | 2.870 |
| 7.686 | 6.712 | | | -1.455 | | | | -11.206 | 20.441 | | | | -10.530 |
| -5.970 | 4.418 | | | -17.578 | | | | -10.021 | 16.145 | | | | 7.225 |
| 3.757 | -16.794 | | | 16.264 | | | | 11.853 | -6.972 | | | | 1.630 |
| 20.340 | 5.736 | | | -14.056 | | | | -22.689 | 22.176 | | | | -26.858 |
| 20.632 | -25.857 | | | 21.255 | | | | -22.257 | -29.455 | | | | -12.518 |
| -7.014 | -15.676 | | | 19.207 | | | | -15.911 | -13.202 | | | | -29.011 |
| -2.177 | -25.536 | | | 24.189 | | | | 20.572 | 7.434 | | | | -9.507 |
| -10.555 | 4.311 | | | -17.883 | | | | 5.615 | 17.737 | | | | -26.053 |
| -25.580 | 20.226 | | | -12.412 | | | | 2.395 | -5.096 | | | | -18.824 |
| 9.229 | 22.533 | | | -16.391 | | | | 1.136 | 15.954 | | | | -22.384 |
| 14.153 | -21.996 | | | -14.905 | | | | -19.603 | 5.135 | | | | -4.510 |
| -24.524 | 12.467 | | | 15.607 | | | | 18.963 | 26.106 | | | | 10.296 |
| -26.616 | 7.852 | | | -23.233 | | | | 12.353 | 20.752 | | | | -10.479 |
| 4.503 | 20.973 | | | -22.611 | | | | -2.094 | 10.294 | | | | 23.091 |
| -8.129 | 18.759 | | | 19.502 | | | | 0.116 | 29.854 | | | | -11.985 |
| -14.843 | 17.373 | | | -15.379 | | | | 11.908 | -21.155 | | | | 6.284 |
| -11.907 | -19.708 | | | -20.832 | | | | 1.347 | -6.051 | | | | -15.234 |
| -3.960 | 19.884 | | | 18.601 | | | | -7.464 | 20.032 | | | | 4.760 |
| 20.012 | 16.238 | | | -11.781 | | | | 2.012 | -1.811 | | | | 9.597 |
| 3.094 | -10.549 | | | 24.147 | | | | -17.687 | 25.397 | | | | 10.808 |
| -18.162 | -2.929 | | | -14.707 | | | | -0.494 | 23.907 | | | | -17.205 |
| -13.118 | -8.479 | | | 17.052 | | | | 7.112 | -15.060 | | | | -4.084 |
| -3.258 | 0.599 | | | -19.280 | | | | 5.231 | -20.518 | | | | 6.730 |
| -1.440 | -0.397 | | | 7.122 | | | | 7.960 | 19.346 | | | | 21.576 |
| -5.177 | 14.699 | | | 10.419 | | | | 19.623 | 29.477 | | | | -9.883 |
| -18.587 | -9.816 | | | 8.213 | | | | 5.605 | 0.965 | | | | -16.220 |
| 5.795 | 24.631 | | | 9.050 | | | | 22.614 | -16.041 | | | | 8.463 |
| 2.020 | -21.838 | | | 6.689 | | | | -2.263 | 5.863 | | | | 14.585 |

| | | | | | | | | | | | | | |
|---|---|---|---|---|---|---|---|---|---|---|---|---|---|
| -25.141 | 25.343 | | | -13.797 | | | | 8.737 | -28.317 | | | | 5.601 |
| -15.831 | 21.588 | | | 21.224 | | | | 10.087 | -24.172 | | | | -12.548 |
| 8.699 | 19.805 | | | 7.262 | | | | 1.995 | -18.108 | | | | 19.983 |
| 9.971 | 19.050 | | | -19.786 | | | | -5.917 | -5.429 | | | | -22.819 |
| 6.588 | 8.262 | | | -18.735 | | | | 3.495 | -1.636 | | | | 27.844 |
| -1.501 | -2.663 | | | 16.106 | | | | 16.720 | 12.205 | | | | -6.310 |
| 11.921 | -8.206 | | | 22.821 | | | | 0.129 | -14.657 | | | | -16.612 |
| -24.391 | -7.839 | | | 15.194 | | | | -3.499 | 28.868 | | | | -7.063 |
| 17.593 | 17.987 | | | 14.124 | | | | -8.788 | -26.552 | | | | 4.561 |
| -6.193 | 11.914 | | | 20.014 | | | | 3.938 | 23.519 | | | | -8.895 |
| -20.775 | -24.205 | | | 5.118 | | | | -6.987 | -20.515 | | | | 24.022 |
| -28.512 | -0.739 | | | -8.860 | | | | -0.649 | 23.226 | | | | -27.370 |
| -0.075 | 6.778 | | | -21.373 | | | | -24.988 | -28.476 | | | | -18.216 |
| 21.620 | 19.510 | | | -19.937 | | | | 9.690 | 16.965 | | | | -24.370 |
| -21.758 | 21.573 | | | -16.326 | | | | -2.446 | 25.876 | | | | 13.780 |
| -15.087 | 0.368 | | | 19.103 | | | | -19.161 | 11.133 | | | | 21.422 |
| 13.030 | 19.782 | | | -22.569 | | | | 3.421 | 1.183 | | | | -25.867 |
| 11.188 | 10.599 | | | -24.836 | | | | 18.143 | -5.207 | | | | -9.433 |
| -4.678 | 11.810 | | | 15.707 | | | | -21.145 | 29.049 | | | | 25.544 |
| -21.359 | 13.147 | | | -20.054 | | | | -6.125 | 25.448 | | | | 23.244 |
| -28.463 | 8.364 | | | 8.404 | | | | 15.932 | -17.094 | | | | -17.193 |
| 17.561 | -8.422 | | | -7.472 | | | | -1.492 | -19.995 | | | | -3.918 |
| 4.542 | 6.256 | | | -13.392 | | | | -4.632 | 29.359 | | | | -6.544 |
| -20.771 | -24.159 | | | -19.261 | | | | 2.396 | -1.796 | | | | -20.016 |
| -1.459 | 15.384 | | | -18.495 | | | | -7.893 | 9.001 | | | | 2.063 |
| -20.218 | 18.139 | | | -14.946 | | | | 1.858 | -26.399 | | | | 0.082 |
| 8.627 | -25.975 | | | 14.830 | | | | -2.649 | -28.897 | | | | 2.323 |
| -29.657 | 22.530 | | | -21.870 | | | | 7.916 | 23.610 | | | | 23.238 |
| 5.548 | 21.055 | | | 9.314 | | | | 0.072 | -11.780 | | | | 17.965 |
| 2.046 | 16.732 | | | -11.005 | | | | -26.252 | 2.120 | | | | 21.472 |
| 24.660 | -14.638 | | | -20.911 | | | | -16.566 | 3.252 | | | | 17.006 |
| 7.966 | 8.777 | | | -19.972 | | | | 5.562 | -7.895 | | | | 12.906 |
| 4.352 | -6.309 | | | -26.804 | | | | -8.942 | -29.443 | | | | 21.606 |
| -27.087 | -7.420 | | | 5.866 | | | | 8.636 | 5.745 | | | | -24.171 |
| -7.550 | 19.913 | | | -10.691 | | | | 22.692 | 23.889 | | | | -3.972 |
| 20.334 | 23.537 | | | 20.642 | | | | -2.647 | -15.770 | | | | -5.306 |

| | | | | | | | | | | | | | |
|---|---|---|---|---|---|---|---|---|---|---|---|---|---|
| 21.858 | 15.757 | | | 20.099 | | | | -10.199 | 4.337 | | | | 7.390 |
| -15.227 | -16.826 | | | -14.917 | | | | -19.397 | -3.445 | | | | -2.684 |
| -28.015 | -15.294 | | | 31.953 | | | | 15.417 | 22.365 | | | | -27.868 |
| -0.883 | 20.926 | | | -16.540 | | | | 0.297 | 6.057 | | | | 0.648 |
| -8.919 | -11.876 | | | -22.813 | | | | -5.726 | -17.469 | | | | -1.601 |
| -11.440 | 13.043 | | | -25.218 | | | | 1.659 | 22.358 | | | | 8.148 |
| -13.886 | -18.561 | | | 22.799 | | | | 9.764 | 12.851 | | | | 16.591 |
| -21.361 | 17.240 | | | 18.667 | | | | -10.695 | -21.349 | | | | -4.214 |
| -13.333 | 14.497 | | | 17.737 | | | | 13.230 | 6.965 | | | | -20.962 |
| -6.307 | -6.754 | | | 7.102 | | | | -17.107 | -4.627 | | | | -0.334 |
| 23.397 | 10.749 | | | -18.548 | | | | -5.100 | 24.647 | | | | 9.611 |
| 20.058 | 21.630 | | | -20.762 | | | | -2.103 | 5.072 | | | | 4.442 |
| 0.389 | -20.282 | | | 6.286 | | | | 20.840 | 9.124 | | | | -24.537 |
| -2.374 | -10.629 | | | -24.360 | | | | 4.910 | 22.487 | | | | 10.486 |
| 14.662 | -7.343 | | | 22.005 | | | | -11.392 | -20.849 | | | | -1.057 |
| -21.828 | -1.489 | | | 19.728 | | | | 3.638 | -4.983 | | | | 20.844 |
| -20.626 | -12.189 | | | 10.136 | | | | 12.619 | -6.484 | | | | 9.030 |
| 0.536 | 9.751 | | | 21.006 | | | | -8.363 | -19.006 | | | | 4.058 |
| -29.243 | -13.939 | | | 5.892 | | | | 16.021 | 21.083 | | | | 7.941 |
| 25.449 | 7.950 | | | -29.712 | | | | 21.657 | 5.292 | | | | -16.908 |
| -11.668 | 11.737 | | | 13.100 | | | | 14.467 | -16.517 | | | | 1.987 |
| -7.897 | -25.404 | | | -17.713 | | | | 24.056 | -20.077 | | | | -15.968 |
| 20.584 | 3.246 | | | -19.506 | | | | -19.457 | 14.726 | | | | 23.877 |
| 13.482 | -25.882 | | | -16.797 | | | | 5.536 | 19.287 | | | | 10.792 |
| -10.413 | -13.535 | | | 18.785 | | | | 4.616 | -16.877 | | | | -22.732 |
| -31.173 | -1.843 | | | 22.658 | | | | -16.801 | -11.361 | | | | -5.317 |
| 7.802 | -18.366 | | | -16.076 | | | | 8.511 | -27.618 | | | | 3.203 |
| -29.980 | -7.870 | | | -22.132 | | | | -27.691 | -2.165 | | | | -9.281 |
| -20.808 | 23.272 | | | 9.603 | | | | -10.954 | 2.448 | | | | 20.246 |
| -13.464 | 23.751 | | | -11.642 | | | | 19.799 | -28.130 | | | | 19.656 |
| -30.567 | -9.574 | | | -17.505 | | | | 10.823 | 13.079 | | | | -10.928 |
| -3.337 | 11.334 | | | -24.760 | | | | -11.462 | -9.336 | | | | -3.551 |
| 29.893 | -1.938 | | | -17.016 | | | | -12.024 | -19.251 | | | | -24.302 |
| 13.578 | -15.176 | | | -19.786 | | | | -6.589 | -26.471 | | | | 18.842 |
| 10.304 | -3.418 | | | 21.421 | | | | -23.696 | 24.743 | | | | 25.614 |
| -20.321 | 10.756 | | | 16.201 | | | | -12.867 | -9.048 | | | | 19.324 |

| | | | | | | | | | | | | | |
|---|---|---|---|---|---|---|---|---|---|---|---|---|---|
| 19.447 | 8.133 | | | -8.088 | | | | 10.101 | 12.411 | | | | -5.400 |
| 12.165 | -20.351 | | | 6.309 | | | | -7.297 | -26.490 | | | | 22.178 |
| 28.073 | -19.493 | | | -21.518 | | | | -6.936 | 14.376 | | | | -3.495 |
| -21.379 | 14.939 | | | 12.587 | | | | -14.937 | -10.377 | | | | -4.061 |
| 1.241 | 6.299 | | | 4.486 | | | | -9.721 | -13.760 | | | | -7.884 |
| -6.635 | -3.379 | | | 0.250 | | | | 1.220 | -16.030 | | | | 2.904 |
| -4.691 | 4.350 | | | -15.265 | | | | -28.261 | 8.738 | | | | -18.010 |
| 25.180 | -15.314 | | | 6.910 | | | | 22.799 | 20.972 | | | | 6.548 |
| -13.980 | 0.441 | | | -17.242 | | | | -21.491 | -0.718 | | | | -14.981 |
| 18.923 | 2.603 | | | -20.385 | | | | 14.888 | -9.961 | | | | 11.680 |
| 16.949 | -5.075 | | | -13.373 | | | | -7.857 | -6.460 | | | | -12.078 |
| 14.418 | 8.421 | | | 9.080 | | | | 12.475 | 7.461 | | | | -23.846 |
| -18.038 | 0.688 | | | 17.307 | | | | -11.485 | 15.906 | | | | -10.400 |
| -30.127 | 6.924 | | | -23.038 | | | | -11.856 | -12.626 | | | | 21.393 |
| 17.880 | -10.119 | | | -27.455 | | | | 3.522 | 16.679 | | | | 10.693 |
| 21.883 | 1.815 | | | 5.548 | | | | -26.053 | 22.590 | | | | 1.550 |
| -10.587 | -4.899 | | | -19.469 | | | | 24.670 | -20.361 | | | | 14.264 |
| 1.853 | -14.590 | | | 19.896 | | | | -9.006 | 13.361 | | | | 13.439 |
| -28.923 | 3.646 | | | -11.204 | | | | 8.614 | -9.700 | | | | 6.929 |
| -14.412 | 11.540 | | | -9.987 | | | | 22.450 | -10.905 | | | | 0.432 |
| -29.372 | -9.927 | | | -22.443 | | | | 18.055 | 2.340 | | | | -7.016 |
| -4.463 | -10.175 | | | -25.262 | | | | 4.100 | -25.553 | | | | 21.280 |
| 11.289 | -6.058 | | | 10.515 | | | | 7.023 | -18.360 | | | | 7.139 |
| -20.112 | -1.074 | | | 13.595 | | | | -7.827 | 0.790 | | | | 12.080 |
| -7.962 | -4.539 | | | 15.928 | | | | -21.133 | 15.224 | | | | 11.343 |
| -9.910 | -1.300 | | | 18.004 | | | | 4.772 | -21.917 | | | | -16.159 |
| 30.831 | 22.148 | | | 20.788 | | | | 10.459 | 1.703 | | | | 1.487 |
| -11.432 | 24.827 | | | -17.737 | | | | -4.297 | -21.642 | | | | 3.026 |
| -14.864 | -7.898 | | | 9.815 | | | | 19.427 | -3.363 | | | | -8.332 |
| 9.474 | -23.852 | | | 25.427 | | | | -8.863 | -24.592 | | | | 13.165 |
| -25.788 | 21.503 | | | 4.024 | | | | -9.989 | -12.833 | | | | -19.365 |
| 25.573 | 22.763 | | | -6.284 | | | | 7.614 | 9.466 | | | | -2.686 |
| -18.020 | 14.580 | | | -23.338 | | | | -4.555 | -6.168 | | | | -16.397 |
| 0.183 | -4.417 | | | 6.241 | | | | -21.923 | 3.027 | | | | 7.873 |
| -0.731 | 21.659 | | | 16.044 | | | | 23.331 | -0.296 | | | | 10.720 |
| -8.208 | 9.413 | | | 22.044 | | | | 22.160 | -4.983 | | | | 13.442 |

| | | | | | | | | | | | | | |
|---|---|---|---|---|---|---|---|---|---|---|---|---|---|
| 1.593 | 2.590 | | | 7.661 | | | | 5.414 | 3.921 | | | | -6.603 |
| -22.379 | -2.929 | | | -20.810 | | | | -6.552 | 2.027 | | | | 14.513 |
| -10.018 | 13.361 | | | -24.913 | | | | -7.891 | -17.893 | | | | -15.151 |
| 24.442 | 1.604 | | | 21.111 | | | | 6.398 | 17.532 | | | | 23.888 |
| 30.370 | -1.440 | | | 19.503 | | | | -0.055 | 19.612 | | | | -25.362 |
| -21.708 | 14.234 | | | -19.668 | | | | -18.850 | -20.788 | | | | 13.959 |
| 30.812 | -17.491 | | | 20.031 | | | | 13.862 | -2.168 | | | | 13.649 |
| -11.132 | -3.964 | | | 5.640 | | | | -13.491 | 25.340 | | | | -1.637 |
| -12.994 | 10.896 | | | 20.759 | | | | 0.751 | 1.409 | | | | 0.242 |
| -14.351 | 24.973 | | | 19.250 | | | | -19.315 | -24.704 | | | | 19.063 |
| -13.585 | 9.939 | | | -7.806 | | | | -27.620 | 14.363 | | | | 23.886 |
| 14.315 | -1.740 | | | -14.199 | | | | 7.482 | -0.442 | | | | -18.775 |
| -6.865 | -18.845 | | | 10.073 | | | | 13.043 | 21.445 | | | | -23.890 |
| 18.119 | -0.936 | | | -13.470 | | | | -7.609 | -8.165 | | | | 18.455 |
| -2.835 | 5.876 | | | -17.543 | | | | 3.206 | -21.779 | | | | 2.491 |
| -8.554 | 23.562 | | | 12.753 | | | | -22.317 | 0.940 | | | | 17.114 |
| 21.309 | 0.123 | | | -24.842 | | | | -8.696 | 23.553 | | | | -11.167 |
| -18.010 | 1.977 | | | 16.897 | | | | 16.727 | -17.764 | | | | -7.484 |
| 29.703 | 3.935 | | | 1.582 | | | | -5.169 | -0.693 | | | | 24.611 |
| -22.670 | -12.397 | | | -24.166 | | | | -19.353 | -4.986 | | | | 21.683 |
| -15.362 | -13.934 | | | -25.191 | | | | 19.961 | -16.687 | | | | 8.823 |
| 15.327 | 19.953 | | | -3.885 | | | | 15.279 | -15.311 | | | | -10.693 |
| 21.973 | 9.173 | | | -18.260 | | | | -8.226 | 23.191 | | | | 26.422 |
| -12.136 | 11.128 | | | -12.342 | | | | 3.053 | -10.962 | | | | -2.691 |
| -16.784 | -9.454 | | | 10.696 | | | | 6.008 | -18.509 | | | | -11.909 |
| -23.800 | -25.973 | | | -20.362 | | | | 13.640 | -0.222 | | | | -21.428 |
| 19.384 | 6.799 | | | -13.451 | | | | -18.003 | 1.209 | | | | 20.335 |
| 29.109 | -0.884 | | | -19.161 | | | | 7.154 | -20.471 | | | | 19.583 |
| -4.761 | 18.291 | | | 18.826 | | | | -16.604 | 15.470 | | | | 9.395 |
| -29.639 | -9.450 | | | 1.819 | | | | -2.138 | 22.673 | | | | 17.179 |
| -28.591 | -3.382 | | | -15.966 | | | | -0.004 | 4.459 | | | | 7.174 |
| 22.700 | 12.207 | | | -22.087 | | | | -17.217 | -16.041 | | | | -18.493 |
| 7.064 | -12.177 | | | 14.373 | | | | -23.027 | 23.789 | | | | -21.384 |
| -11.275 | -4.256 | | | 11.138 | | | | 5.891 | -2.089 | | | | -27.054 |
| -12.324 | 5.304 | | | -24.127 | | | | 13.589 | 3.893 | | | | -25.139 |
| 30.216 | -22.766 | | | -10.056 | | | | 19.992 | 12.151 | | | | -10.216 |

| | | | | | | | | | | | | | |
|---|---|---|---|---|---|---|---|---|---|---|---|---|---|
| -0.537 | 26.055 | | | 25.351 | | | | 28.896 | 9.665 | | | | 13.866 |
| -13.585 | 9.285 | | | -20.918 | | | | 23.173 | 27.174 | | | | -22.841 |
| -9.106 | 7.505 | | | -3.488 | | | | -9.613 | -13.830 | | | | -26.092 |
| -5.483 | -19.311 | | | 21.475 | | | | -17.613 | 25.965 | | | | 1.619 |
| -18.472 | 3.166 | | | -19.522 | | | | 1.489 | 25.311 | | | | 16.702 |
| -3.980 | 3.486 | | | -22.945 | | | | -10.088 | -8.113 | | | | -8.018 |
| -9.627 | -25.926 | | | 12.575 | | | | -13.731 | -15.644 | | | | -13.713 |
| 0.168 | 8.678 | | | -7.608 | | | | -9.035 | 7.210 | | | | -0.910 |
| 22.352 | -5.304 | | | 17.437 | | | | -19.548 | -20.394 | | | | -25.549 |
| 16.400 | 5.116 | | | 7.363 | | | | -11.131 | -17.310 | | | | 21.018 |
| -15.735 | -10.638 | | | -6.711 | | | | 25.752 | -0.888 | | | | -5.972 |
| -11.835 | -19.660 | | | -25.652 | | | | 20.291 | 24.238 | | | | -14.723 |
| -22.305 | -14.019 | | | -17.423 | | | | 17.466 | -22.773 | | | | 17.642 |
| 28.673 | 11.015 | | | 20.152 | | | | 12.792 | -25.286 | | | | -4.257 |
| -6.832 | -0.722 | | | 2.931 | | | | 11.083 | -23.095 | | | | -8.211 |
| -6.269 | 0.703 | | | 26.458 | | | | 8.805 | 26.060 | | | | -10.305 |
| -19.478 | -26.117 | | | 10.200 | | | | 6.742 | -21.534 | | | | -18.990 |
| 10.352 | -5.204 | | | -9.112 | | | | -23.512 | -18.188 | | | | 19.027 |
| 28.004 | -11.860 | | | 18.556 | | | | 5.162 | -19.986 | | | | 23.627 |
| -0.731 | 1.050 | | | 8.355 | | | | -14.337 | 17.886 | | | | -22.058 |
| -7.873 | -6.666 | | | -16.993 | | | | -16.582 | 28.052 | | | | 19.767 |
| -10.246 | -24.988 | | | -20.196 | | | | -22.636 | -22.873 | | | | -24.293 |
| -5.597 | -9.705 | | | -20.640 | | | | -20.843 | -19.649 | | | | -17.917 |
| -27.983 | -13.307 | | | -25.786 | | | | -9.533 | 23.603 | | | | -4.964 |
| -9.388 | -11.109 | | | -5.186 | | | | 27.282 | 21.402 | | | | 25.206 |
| -24.250 | -6.310 | | | -24.546 | | | | 3.945 | -17.036 | | | | -23.059 |
| -4.556 | -19.006 | | | -24.989 | | | | -32.231 | 27.304 | | | | 20.304 |
| -13.971 | 4.729 | | | -13.244 | | | | -23.047 | -21.442 | | | | 7.745 |
| 26.952 | -24.415 | | | 14.899 | | | | 30.807 | -19.424 | | | | -11.667 |
| 6.803 | -1.272 | | | 3.089 | | | | 19.582 | 28.019 | | | | 22.885 |
| -27.096 | -26.023 | | | -18.133 | | | | 10.055 | 26.534 | | | | -13.726 |
| -5.637 | -23.586 | | | -4.764 | | | | 5.966 | -26.813 | | | | 13.379 |
| 23.222 | -19.382 | | | 19.171 | | | | 13.727 | 28.745 | | | | 5.163 |
| -22.055 | -6.871 | | | 13.448 | | | | 21.055 | -25.270 | | | | 22.829 |
| -6.685 | 6.714 | | | 22.541 | | | | 26.088 | -19.118 | | | | -6.807 |
| -27.717 | -17.182 | | | 6.468 | | | | -6.859 | 27.133 | | | | -22.350 |

| | | | | | | | | | | | | | |
|---|---|---|---|---|---|---|---|---|---|---|---|---|---|
| -20.520 | 23.491 | | | 16.047 | | | | 22.957 | -11.540 | | | | 5.626 |
| -16.602 | -9.458 | | | -16.825 | | | | -12.809 | 5.234 | | | | -14.979 |
| -2.807 | 4.613 | | | -23.563 | | | | -26.272 | -15.577 | | | | -3.533 |
| 26.019 | -28.488 | | | 5.004 | | | | -10.516 | 28.055 | | | | -0.979 |
| -7.980 | 18.726 | | | -14.628 | | | | -3.733 | 26.767 | | | | -16.445 |
| -18.327 | -3.325 | | | -21.397 | | | | 25.187 | 24.619 | | | | 3.368 |
| -9.455 | -8.136 | | | -6.642 | | | | -17.403 | -2.494 | | | | 7.184 |
| -23.423 | -0.772 | | | -1.313 | | | | 8.690 | -28.114 | | | | 17.092 |
| -6.095 | -10.122 | | | 18.603 | | | | -18.362 | 28.536 | | | | -4.601 |
| -1.549 | 29.210 | | | -5.123 | | | | 7.575 | -0.492 | | | | 10.533 |
| -6.843 | -23.660 | | | 11.703 | | | | 0.809 | -22.002 | | | | -8.115 |
| -25.362 | 18.711 | | | -13.167 | | | | -5.445 | 20.801 | | | | 8.317 |
| 10.130 | -13.962 | | | -0.688 | | | | -17.010 | 27.191 | | | | -28.069 |
| 2.308 | 9.521 | | | 20.726 | | | | -30.512 | 14.184 | | | | 28.802 |
| -21.054 | -4.514 | | | -7.458 | | | | 24.746 | 0.101 | | | | -23.189 |
| -10.390 | 0.298 | | | -15.447 | | | | -12.812 | -9.079 | | | | -7.539 |
| 7.791 | -22.409 | | | -18.199 | | | | 13.897 | -10.108 | | | | -6.625 |
| 0.834 | -0.387 | | | -20.577 | | | | 11.305 | 17.267 | | | | 8.686 |
| 5.252 | 28.807 | | | -21.597 | | | | -8.036 | 27.643 | | | | -2.005 |
| -13.748 | 8.033 | | | -16.364 | | | | -23.721 | -2.017 | | | | -11.253 |
| -5.570 | 7.052 | | | 10.335 | | | | 23.531 | 28.077 | | | | -16.858 |
| -26.734 | 13.207 | | | 7.297 | | | | 18.775 | -20.030 | | | | -8.774 |
| -7.167 | -22.252 | | | 2.337 | | | | -21.803 | -5.962 | | | | -29.095 |
| -1.272 | 18.194 | | | -14.204 | | | | 29.454 | -29.597 | | | | 28.598 |
| -9.925 | 0.724 | | | -13.313 | | | | -20.681 | -16.726 | | | | 29.316 |
| -16.439 | -6.173 | | | -22.692 | | | | -21.212 | -28.983 | | | | 6.471 |
| -20.749 | -3.554 | | | -20.964 | | | | 7.299 | 15.649 | | | | 25.660 |
| 9.705 | -25.699 | | | 22.037 | | | | 20.166 | -11.164 | | | | 24.047 |
| 7.115 | -1.867 | | | 15.618 | | | | 1.759 | -10.482 | | | | -12.599 |
| 13.732 | 24.856 | | | -5.915 | | | | 9.879 | 20.867 | | | | -0.165 |
| -6.768 | -5.396 | | | -17.821 | | | | -8.257 | -9.088 | | | | -16.262 |
| -9.473 | -11.253 | | | 15.063 | | | | -13.144 | 7.547 | | | | -6.288 |
| -5.978 | -21.296 | | | -23.960 | | | | -10.196 | 20.331 | | | | 2.664 |
| -15.123 | 26.447 | | | 17.628 | | | | 4.779 | -7.249 | | | | 6.445 |
| -23.181 | -30.185 | | | -10.063 | | | | -18.380 | -1.886 | | | | 14.503 |
| -19.595 | 8.188 | | | 18.065 | | | | 18.088 | -18.316 | | | | 2.352 |

| | | | | | | | | | | | | | |
|---|---|---|---|---|---|---|---|---|---|---|---|---|---|
| -2.473 | 15.246 | | | 10.930 | | | | -8.821 | -24.972 | | | | 4.861 |
| -11.393 | 6.027 | | | -4.335 | | | | 25.236 | -11.715 | | | | 21.713 |
| -8.355 | -9.151 | | | -16.036 | | | | 16.297 | 28.335 | | | | -5.769 |
| 5.076 | -11.695 | | | -19.782 | | | | 3.974 | 28.908 | | | | -24.348 |
| -25.570 | -5.145 | | | 17.661 | | | | 13.067 | -7.049 | | | | -8.619 |
| -15.528 | -20.537 | | | -15.102 | | | | 22.946 | -16.324 | | | | -16.623 |
| 14.071 | 29.011 | | | 14.601 | | | | 20.313 | -27.411 | | | | 7.247 |
| 5.989 | 19.216 | | | 16.691 | | | | 19.381 | -8.364 | | | | -9.605 |
| 15.979 | -9.701 | | | -13.605 | | | | -30.751 | -26.356 | | | | -12.619 |
| 11.480 | 30.482 | | | 19.888 | | | | 24.322 | 18.223 | | | | -28.446 |
| 1.639 | -28.790 | | | 11.952 | | | | 9.808 | 28.193 | | | | 14.392 |
| -7.606 | 1.318 | | | 0.460 | | | | 23.466 | -18.432 | | | | -27.127 |
| -22.507 | 15.825 | | | -1.065 | | | | -14.164 | 21.227 | | | | 8.732 |
| 18.925 | 31.329 | | | -22.016 | | | | 6.139 | -27.795 | | | | 7.446 |
| -5.486 | 4.930 | | | 0.917 | | | | -29.433 | -29.229 | | | | -8.585 |
| -23.380 | -15.482 | | | -7.951 | | | | 18.892 | 20.459 | | | | 19.427 |
| 25.030 | 17.338 | | | -19.962 | | | | 8.095 | 1.482 | | | | -9.810 |
| 6.694 | 28.515 | | | -10.589 | | | | -18.206 | -8.376 | | | | 7.877 |
| 12.002 | 24.490 | | | -19.263 | | | | 5.350 | -5.993 | | | | -1.692 |
| 12.638 | 26.637 | | | 12.855 | | | | 9.804 | -30.293 | | | | 21.942 |
| -9.373 | -20.303 | | | -0.211 | | | | 8.924 | -22.055 | | | | -5.523 |
| 0.701 | 21.681 | | | 15.107 | | | | -11.668 | -30.787 | | | | -19.367 |
| -4.923 | 28.736 | | | -3.408 | | | | -19.860 | 30.671 | | | | 27.018 |
| 5.140 | -29.402 | | | 15.995 | | | | -7.579 | 22.562 | | | | -3.464 |
| -2.000 | 21.683 | | | -19.828 | | | | 5.851 | 30.876 | | | | -9.214 |
| -3.643 | 29.157 | | | -8.157 | | | | -8.217 | 13.712 | | | | -20.344 |
| -7.458 | 27.880 | | | 16.490 | | | | -22.273 | -23.848 | | | | -18.826 |
| 24.743 | 23.262 | | | 12.456 | | | | -24.833 | 27.451 | | | | -30.962 |
| 10.667 | -13.757 | | | 2.936 | | | | 16.075 | 10.605 | | | | 13.541 |
| 23.343 | 24.053 | | | 21.355 | | | | -17.958 | 31.409 | | | | -29.824 |
| -6.017 | -20.251 | | | -19.778 | | | | -6.863 | -13.290 | | | | -6.242 |
| 14.967 | -4.293 | | | 6.902 | | | | -4.784 | -19.527 | | | | -24.822 |
| 0.350 | -4.776 | | | -14.350 | | | | 12.553 | 15.695 | | | | -27.467 |
| 5.598 | 19.522 | | | -7.402 | | | | -5.624 | -6.974 | | | | -8.876 |
| 20.978 | -28.349 | | | -13.447 | | | | 13.721 | -27.188 | | | | -20.154 |
| 16.368 | -22.902 | | | 22.961 | | | | 29.636 | 28.666 | | | | 4.848 |

| | | | | | | | | | | | | | |
|---|---|---|---|---|---|---|---|---|---|---|---|---|---|
| 9.847 | -7.441 | | | -21.807 | | | | 10.544 | 17.712 | | | | -19.668 |
| 7.318 | -22.318 | | | -22.899 | | | | -26.006 | -29.911 | | | | -0.362 |
| 8.150 | 21.883 | | | 9.334 | | | | 15.269 | 30.788 | | | | -5.847 |
| 17.262 | -29.157 | | | -2.396 | | | | 9.721 | -15.665 | | | | -5.220 |
| -3.046 | 12.960 | | | -7.493 | | | | 3.197 | -20.073 | | | | -10.712 |
| 20.282 | -21.261 | | | -17.964 | | | | -18.961 | -4.683 | | | | 21.610 |
| 7.851 | -24.541 | | | 16.158 | | | | -23.372 | -18.498 | | | | 21.225 |
| -1.565 | -8.239 | | | -10.065 | | | | 19.153 | -28.521 | | | | -2.442 |
| 12.566 | -7.131 | | | 8.302 | | | | -13.882 | 30.192 | | | | -18.910 |
| 8.429 | -5.214 | | | -5.442 | | | | 6.008 | 22.158 | | | | -8.291 |
| 4.045 | 1.498 | | | -18.369 | | | | -7.004 | -18.886 | | | | -1.231 |
| 9.215 | -27.363 | | | -18.722 | | | | -8.361 | -7.005 | | | | -7.250 |
| -2.839 | -16.898 | | | -12.034 | | | | -24.567 | -30.816 | | | | 11.721 |
| -15.210 | 19.689 | | | 15.700 | | | | -22.803 | -28.914 | | | | 19.784 |
| -11.418 | 21.014 | | | 17.688 | | | | -25.707 | 28.728 | | | | 17.909 |
| 7.385 | -27.792 | | | 2.346 | | | | -14.245 | 24.530 | | | | -19.295 |
| -10.528 | -19.223 | | | 14.621 | | | | -7.718 | 19.401 | | | | -2.323 |
| 15.579 | 27.434 | | | 16.688 | | | | 24.481 | -27.462 | | | | 1.318 |
| -9.672 | 27.872 | | | 20.732 | | | | -4.876 | 28.394 | | | | 3.157 |
| 1.324 | -25.713 | | | 18.592 | | | | -17.858 | -25.409 | | | | -4.407 |
| -3.977 | 11.074 | | | 19.172 | | | | -8.898 | -28.682 | | | | 29.567 |
| 5.934 | -1.077 | | | 18.218 | | | | 9.276 | -30.048 | | | | -13.280 |
| -3.285 | 27.122 | | | 19.823 | | | | 25.743 | -1.188 | | | | -18.399 |
| 4.648 | -3.136 | | | 3.306 | | | | -7.133 | -8.136 | | | | -7.475 |
| 18.903 | 0.512 | | | 7.307 | | | | -2.826 | -1.477 | | | | -8.847 |
| 23.633 | 20.257 | | | 19.616 | | | | -20.005 | -22.383 | | | | 1.638 |
| -19.543 | -8.102 | | | -8.521 | | | | -18.185 | -7.154 | | | | -21.207 |
| -7.509 | 24.217 | | | 23.695 | | | | 14.119 | 17.975 | | | | 19.234 |
| -1.369 | -16.779 | | | -11.450 | | | | 5.657 | -22.776 | | | | 2.975 |
| 2.484 | 26.525 | | | -13.875 | | | | 7.927 | 1.671 | | | | -14.577 |
| 14.267 | 8.476 | | | -6.162 | | | | -6.657 | 30.074 | | | | 7.647 |
| 4.390 | 18.029 | | | -17.228 | | | | 20.485 | 29.027 | | | | 8.444 |
| -19.684 | 12.470 | | | 2.465 | | | | 3.103 | -19.190 | | | | 6.690 |
| 3.802 | -3.729 | | | -14.687 | | | | 9.731 | -28.518 | | | | -1.291 |
| 2.174 | 14.416 | | | 1.422 | | | | -12.909 | 20.459 | | | | 29.542 |
| 22.410 | -26.407 | | | 14.304 | | | | -4.647 | 27.672 | | | | 19.996 |

| | | | | | | | | | | | | | |
|---|---|---|---|---|---|---|---|---|---|---|---|---|---|
| 5.944 | -27.542 | | | 8.660 | | | | 6.726 | -6.640 | | | | -8.807 |
| -3.850 | -11.576 | | | -5.903 | | | | 19.580 | -16.964 | | | | -4.463 |
| 6.904 | -10.363 | | | -20.297 | | | | 21.051 | 12.542 | | | | -5.118 |
| -2.324 | 23.678 | | | 20.762 | | | | -0.585 | -29.363 | | | | 13.303 |
| -0.720 | 6.938 | | | -10.132 | | | | -6.055 | 27.094 | | | | -12.519 |
| -11.050 | 26.555 | | | 23.186 | | | | 22.259 | -10.833 | | | | 2.773 |
| -11.414 | -24.227 | | | -12.822 | | | | 23.926 | 14.207 | | | | -24.913 |
| -25.590 | -7.891 | | | 0.084 | | | | 23.046 | -15.560 | | | | 0.366 |
| 0.680 | 1.975 | | | -3.199 | | | | 8.322 | 21.774 | | | | -2.408 |
| -23.846 | -26.949 | | | -5.166 | | | | 5.385 | -10.837 | | | | -12.725 |
| 9.594 | 4.213 | | | -11.900 | | | | -6.810 | -6.291 | | | | 9.441 |
| 3.943 | 27.178 | | | -17.726 | | | | -14.360 | -3.821 | | | | -5.283 |
| -0.972 | -2.488 | | | 0.736 | | | | -0.731 | -5.607 | | | | -23.699 |
| -3.579 | 26.623 | | | 4.797 | | | | 7.862 | 1.487 | | | | -1.699 |
| -26.035 | 25.173 | | | -3.825 | | | | -8.606 | 22.697 | | | | -7.836 |
| 10.776 | -16.642 | | | -25.214 | | | | -28.677 | 14.119 | | | | -6.493 |
| -2.929 | 12.085 | | | 18.121 | | | | -16.172 | -13.679 | | | | 19.958 |
| 20.124 | -6.305 | | | -5.758 | | | | -5.741 | -29.597 | | | | 14.632 |
| -8.234 | -19.173 | | | -5.102 | | | | 5.891 | -4.975 | | | | -10.514 |
| 8.892 | -18.782 | | | 19.927 | | | | 10.847 | 27.986 | | | | 8.865 |
| -15.971 | -8.260 | | | 7.710 | | | | 1.692 | 17.770 | | | | -21.900 |
| -13.692 | -20.258 | | | -23.743 | | | | -6.547 | -5.855 | | | | -21.517 |
| -5.036 | 14.726 | | | 0.064 | | | | 9.480 | -13.234 | | | | 12.554 |
| -6.964 | -4.935 | | | -2.772 | | | | -23.533 | 21.557 | | | | 16.812 |
| -18.994 | -27.041 | | | -6.605 | | | | 10.290 | 27.362 | | | | -4.954 |
| -10.111 | -24.396 | | | 2.155 | | | | 6.345 | 15.004 | | | | 9.267 |
| -10.891 | -26.616 | | | -0.775 | | | | 4.992 | -23.248 | | | | 0.195 |
| -18.313 | -24.904 | | | -9.955 | | | | -16.370 | 20.838 | | | | -7.973 |
| 24.981 | -25.205 | | | -2.819 | | | | -20.702 | -4.540 | | | | -26.890 |
| -26.927 | -22.829 | | | -7.003 | | | | 1.821 | 17.365 | | | | 9.505 |
| -23.830 | -4.702 | | | -21.272 | | | | -10.566 | -1.300 | | | | -6.145 |
| 1.649 | 20.937 | | | 6.835 | | | | 26.888 | 12.842 | | | | 14.039 |
| 10.574 | 12.897 | | | 2.419 | | | | 1.014 | 11.178 | | | | 3.313 |
| 26.411 | -5.500 | | | -1.185 | | | | -20.027 | -14.226 | | | | -25.511 |
| 6.902 | -10.985 | | | -14.373 | | | | -21.818 | -18.151 | | | | 9.265 |
| -10.427 | 16.944 | | | 20.832 | | | | 12.162 | -11.118 | | | | 19.917 |

| | | | | | | | | | | | | | |
|---|---|---|---|---|---|---|---|---|---|---|---|---|---|
| 3.201 | -6.776 | | | 4.883 | | | | -5.831 | -28.263 | | | | 5.739 |
| 25.232 | -9.215 | | | 8.463 | | | | -2.530 | -23.989 | | | | -29.489 |
| -7.206 | -23.782 | | | -3.348 | | | | 8.959 | -27.827 | | | | -8.995 |
| -1.724 | -1.271 | | | 18.478 | | | | 1.070 | 22.570 | | | | -21.520 |
| -21.067 | -5.708 | | | 9.719 | | | | 11.123 | -26.934 | | | | -0.210 |
| 5.994 | -0.824 | | | 6.124 | | | | 0.343 | 26.732 | | | | -4.208 |
| 0.210 | -21.544 | | | -7.297 | | | | 15.603 | 28.132 | | | | 1.696 |
| -17.008 | 9.462 | | | 8.938 | | | | -5.605 | 1.774 | | | | -3.389 |
| -13.760 | -13.122 | | | -1.009 | | | | -16.899 | 21.394 | | | | 22.439 |
| -20.321 | 4.374 | | | -9.456 | | | | 7.211 | -6.148 | | | | 18.429 |
| -16.371 | -25.151 | | | -25.426 | | | | 1.299 | -28.242 | | | | -1.664 |
| 1.305 | -15.880 | | | -12.009 | | | | -21.777 | 24.288 | | | | -17.244 |
| -6.336 | -20.331 | | | 7.425 | | | | 6.304 | -1.742 | | | | -27.022 |
| -11.593 | -7.044 | | | 4.720 | | | | -0.956 | -22.192 | | | | -6.976 |
| 0.997 | 11.136 | | | 13.067 | | | | 12.417 | -4.242 | | | | -7.720 |
| -4.642 | -13.524 | | | -5.824 | | | | 0.738 | -10.141 | | | | 6.268 |
| -22.417 | -4.562 | | | -10.647 | | | | 10.411 | 25.384 | | | | -5.373 |
| 1.359 | -9.640 | | | -1.485 | | | | 5.792 | -1.212 | | | | -27.565 |
| 0.195 | 0.805 | | | 14.517 | | | | 6.358 | 10.489 | | | | -2.619 |
| 20.532 | 20.459 | | | -23.603 | | | | 7.018 | 21.460 | | | | -4.502 |
| 17.195 | 13.741 | | | -7.446 | | | | -6.030 | -11.300 | | | | -9.402 |
| -3.117 | 24.203 | | | -22.323 | | | | -11.496 | -23.972 | | | | 18.413 |
| 12.166 | -3.404 | | | 1.662 | | | | -8.058 | -18.410 | | | | -3.863 |
| -5.686 | 1.915 | | | 2.242 | | | | 22.254 | -12.268 | | | | 2.645 |
| 14.076 | -11.083 | | | -12.144 | | | | 10.780 | -3.740 | | | | -1.541 |
| -7.749 | -9.073 | | | -8.844 | | | | -16.980 | -9.301 | | | | 2.084 |
| 12.954 | 8.642 | | | 18.153 | | | | 18.247 | -0.776 | | | | -11.550 |
| -14.188 | 0.060 | | | 4.670 | | | | -5.168 | 15.469 | | | | 26.470 |
| -15.756 | -21.161 | | | -10.523 | | | | -24.900 | -17.064 | | | | -19.632 |
| -5.351 | | | | -26.075 | | | | 4.390 | 1.401 | | | | -7.966 |
| -8.646 | | | | 3.404 | | | | -4.448 | -27.373 | | | | 0.814 |
| -20.929 | | | | -22.120 | | | | 25.323 | -15.242 | | | | -20.743 |
| -27.753 | | | | -27.663 | | | | -10.832 | 16.884 | | | | -12.473 |
| -23.358 | | | | -12.047 | | | | 4.572 | 22.080 | | | | -7.538 |
| -1.182 | | | | -17.502 | | | | -11.384 | 15.766 | | | | 19.733 |
| 17.954 | | | | -4.044 | | | | -1.781 | -11.206 | | | | 4.537 |

| | | | | | | | | | | | | | |
|---|---|---|---|---|---|---|---|---|---|---|---|---|---|
| 25.428 | | | | 6.399 | | | | -25.834 | -7.806 | | | | 2.194 |
| 22.440 | | | | -20.540 | | | | 8.816 | 26.152 | | | | -3.362 |
| -22.996 | | | | 3.292 | | | | -14.525 | -3.309 | | | | -3.494 |
| -5.440 | | | | 1.432 | | | | 24.588 | 4.192 | | | | 24.261 |
| 27.330 | | | | -5.294 | | | | 17.006 | -6.218 | | | | 13.810 |
| 18.401 | | | | -12.585 | | | | 15.280 | -3.998 | | | | 9.266 |
| 19.226 | | | | 8.737 | | | | 3.175 | -0.523 | | | | 22.262 |
| -12.932 | | | | -6.105 | | | | 23.692 | -26.842 | | | | 1.014 |
| -9.015 | | | | 27.924 | | | | 11.142 | -13.990 | | | | -6.458 |
| 18.672 | | | | 19.900 | | | | -2.240 | -24.616 | | | | 6.576 |
| -9.698 | | | | -22.498 | | | | -11.330 | -26.197 | | | | -4.528 |
| -20.697 | | | | -1.043 | | | | 7.799 | 7.784 | | | | -16.903 |
| 0.488 | | | | -13.342 | | | | 4.263 | -23.833 | | | | -8.171 |
| -29.077 | | | | 11.760 | | | | 13.930 | -9.772 | | | | -19.665 |
| 24.747 | | | | -12.162 | | | | 4.983 | -13.180 | | | | 25.979 |
| -19.117 | | | | -20.266 | | | | -12.567 | -16.672 | | | | 17.669 |
| -26.580 | | | | 21.190 | | | | -6.669 | 22.260 | | | | 4.205 |
| -16.427 | | | | -13.726 | | | | -7.822 | -9.276 | | | | 7.359 |
| 21.545 | | | | 9.978 | | | | 3.888 | 15.368 | | | | -15.356 |
| 20.417 | | | | 22.771 | | | | 0.785 | -10.974 | | | | -17.854 |
| 19.834 | | | | -0.033 | | | | -12.916 | 22.408 | | | | -4.951 |
| 22.312 | | | | -3.596 | | | | -2.716 | -4.608 | | | | -18.277 |
| 26.691 | | | | -2.352 | | | | -4.544 | -24.154 | | | | 16.336 |
| 27.074 | | | | 5.544 | | | | 7.426 | -9.186 | | | | 1.183 |
| 23.602 | | | | -4.772 | | | | -11.451 | 24.349 | | | | 1.759 |
| -7.107 | | | | 21.918 | | | | 13.560 | 3.327 | | | | -27.438 |
| 20.175 | | | | 19.432 | | | | -0.760 | -19.517 | | | | -22.662 |
| 25.137 | | | | -19.257 | | | | -3.481 | 7.540 | | | | 5.978 |
| -18.024 | | | | 6.271 | | | | 4.212 | -22.409 | | | | -9.591 |
| 28.786 | | | | 3.386 | | | | -25.761 | 9.265 | | | | 2.539 |
| -5.936 | | | | -2.772 | | | | -11.042 | 16.189 | | | | -5.586 |
| -3.701 | | | | -0.689 | | | | 9.030 | -2.156 | | | | -0.528 |
| -21.916 | | | | 7.651 | | | | 1.994 | -7.248 | | | | -6.478 |
| -7.391 | | | | -4.429 | | | | -0.411 | -3.816 | | | | 24.628 |
| 20.656 | | | | -20.647 | | | | -26.479 | | | | | -4.693 |
| 29.203 | | | | 7.298 | | | | -6.695 | | | | | 5.013 |

| | | | | | | | | | | | | | |
|---|---|---|---|---|---|---|---|---|---|---|---|---|---|
| -22.272 | | | | -23.714 | | | | 4.891 | | | | | 26.710 |
| 9.523 | | | | -27.481 | | | | 10.315 | | | | | -0.346 |
| 12.435 | | | | -25.729 | | | | -8.490 | | | | | -11.125 |
| 20.475 | | | | 13.983 | | | | 1.046 | | | | | -15.463 |
| 11.629 | | | | 11.610 | | | | 11.627 | | | | | -2.519 |
| -18.451 | | | | -19.002 | | | | -5.172 | | | | | 0.631 |
| 10.187 | | | | -14.602 | | | | -20.098 | | | | | -6.222 |
| 4.138 | | | | 18.884 | | | | 13.925 | | | | | 9.008 |
| -8.549 | | | | 14.450 | | | | -22.163 | | | | | -6.648 |
| 7.417 | | | | -24.357 | | | | 14.215 | | | | | 11.945 |
| 17.585 | | | | -7.758 | | | | 0.334 | | | | | 3.445 |
| 14.396 | | | | 28.717 | | | | -13.930 | | | | | 10.458 |
| 2.793 | | | | 4.443 | | | | -23.740 | | | | | 19.752 |
| 30.870 | | | | -15.798 | | | | -0.337 | | | | | 18.255 |
| -21.665 | | | | 5.579 | | | | -20.101 | | | | | -5.764 |
| 5.935 | | | | 3.824 | | | | -3.729 | | | | | -9.797 |
| 1.974 | | | | 22.553 | | | | -13.320 | | | | | 18.897 |
| 23.461 | | | | 13.959 | | | | 10.286 | | | | | 25.175 |
| -11.213 | | | | 4.749 | | | | 18.102 | | | | | 4.254 |
| -11.778 | | | | -21.459 | | | | -6.832 | | | | | -4.100 |
| 30.715 | | | | -27.509 | | | | 12.019 | | | | | -7.478 |
| -16.881 | | | | -16.972 | | | | -9.992 | | | | | -1.792 |
| 23.651 | | | | -22.330 | | | | -13.556 | | | | | -20.732 |
| 1.911 | | | | 13.672 | | | | -10.293 | | | | | 5.459 |
| 16.943 | | | | -9.996 | | | | 3.304 | | | | | 1.745 |
| -20.018 | | | | 20.003 | | | | -0.896 | | | | | -3.414 |
| 17.388 | | | | -13.254 | | | | -7.281 | | | | | 7.846 |
| -19.607 | | | | -22.918 | | | | -26.484 | | | | | -16.460 |
| 30.292 | | | | 9.142 | | | | -23.990 | | | | | 1.222 |
| 6.705 | | | | -23.994 | | | | -5.058 | | | | | -5.083 |
| 28.414 | | | | -16.545 | | | | -2.984 | | | | | 19.072 |
| 24.217 | | | | -10.655 | | | | -14.307 | | | | | -17.003 |
| -20.776 | | | | -19.858 | | | | 7.796 | | | | | -7.948 |
| -12.168 | | | | 9.302 | | | | -17.000 | | | | | 11.226 |
| -16.205 | | | | 2.003 | | | | 14.250 | | | | | 3.139 |
| 22.060 | | | | -24.965 | | | | 1.345 | | | | | -18.465 |

| | | | | | | | | | | | | | |
|---|---|---|---|---|---|---|---|---|---|---|---|---|---|
| -31.067 | | | | -8.938 | | | | -19.687 | | | | | -1.348 |
| 30.540 | | | | -2.216 | | | | 6.250 | | | | | -6.108 |
| 18.775 | | | | 3.810 | | | | -7.840 | | | | | 19.592 |
| 12.073 | | | | -21.009 | | | | -6.864 | | | | | 12.595 |
| 14.130 | | | | -22.475 | | | | -17.289 | | | | | 3.606 |
| -11.748 | | | | -13.640 | | | | 24.009 | | | | | -2.580 |
| 25.745 | | | | 4.446 | | | | 2.823 | | | | | -21.199 |
| 14.776 | | | | -23.319 | | | | -5.357 | | | | | -26.350 |
| 29.848 | | | | -19.218 | | | | -0.590 | | | | | 0.883 |
| -6.742 | | | | 9.522 | | | | -2.642 | | | | | -12.415 |
| -2.942 | | | | -21.302 | | | | 13.455 | | | | | 15.986 |
| 19.016 | | | | -25.378 | | | | 9.706 | | | | | -11.726 |
| 9.814 | | | | -17.868 | | | | -23.064 | | | | | -23.137 |
| 9.176 | | | | 20.877 | | | | -8.606 | | | | | 4.909 |
| 3.570 | | | | -12.821 | | | | -20.423 | | | | | -22.576 |
| 22.652 | | | | 5.497 | | | | 18.106 | | | | | 3.496 |
| 5.030 | | | | -4.107 | | | | -14.272 | | | | | -1.108 |
| 28.607 | | | | -15.079 | | | | 2.416 | | | | | 7.464 |
| 17.105 | | | | -16.769 | | | | 20.008 | | | | | 6.888 |
| 10.705 | | | | 19.661 | | | | 21.573 | | | | | -19.904 |
| -23.361 | | | | 16.074 | | | | -11.060 | | | | | 5.075 |
| -21.752 | | | | -8.309 | | | | 11.419 | | | | | -9.860 |
| -5.448 | | | | 16.860 | | | | 8.155 | | | | | 2.907 |
| -1.396 | | | | 18.107 | | | | -4.639 | | | | | -10.542 |
| -18.701 | | | | -26.536 | | | | 6.481 | | | | | -14.895 |
| -27.287 | | | | 17.180 | | | | -3.050 | | | | | -1.827 |
| -13.704 | | | | 21.740 | | | | 12.562 | | | | | -17.457 |
| 21.416 | | | | -24.815 | | | | -12.046 | | | | | 19.795 |
| -2.255 | | | | 5.315 | | | | 5.761 | | | | | 10.445 |
| -23.345 | | | | -30.042 | | | | -15.716 | | | | | 0.385 |
| 8.741 | | | | -28.900 | | | | 0.448 | | | | | -12.945 |
| 21.683 | | | | -20.884 | | | | -1.958 | | | | | -13.735 |
| -28.064 | | | | -13.785 | | | | -8.445 | | | | | -5.444 |
| -10.919 | | | | -18.929 | | | | 7.781 | | | | | 16.587 |
| 23.115 | | | | -19.961 | | | | 23.380 | | | | | -0.990 |
| 0.367 | | | | -17.458 | | | | 10.118 | | | | | 4.152 |

| | | | | | | | | | | | | | |
|---|---|---|---|---|---|---|---|---|---|---|---|---|---|
| -27.248 | | | | -23.980 | | | | 6.921 | | | | | -7.139 |
| -29.049 | | | | -22.215 | | | | -4.630 | | | | | -12.226 |
| 25.616 | | | | -5.267 | | | | 4.666 | | | | | 0.772 |
| 24.965 | | | | -5.858 | | | | -9.774 | | | | | -15.228 |
| -3.983 | | | | 3.113 | | | | -7.270 | | | | | -5.112 |
| 17.918 | | | | -9.125 | | | | 4.176 | | | | | |
| -20.003 | | | | -27.469 | | | | -4.200 | | | | | |
| 23.153 | | | | -26.754 | | | | 16.003 | | | | | |
| -19.638 | | | | -3.800 | | | | 17.243 | | | | | |
| 14.443 | | | | -24.804 | | | | -16.142 | | | | | |
| -21.930 | | | | -14.579 | | | | 24.874 | | | | | |
| -18.613 | | | | 14.888 | | | | -1.402 | | | | | |
| 10.593 | | | | -12.435 | | | | -2.510 | | | | | |
| 8.290 | | | | -22.996 | | | | 22.642 | | | | | |
| 6.390 | | | | -10.478 | | | | -10.224 | | | | | |
| 8.870 | | | | -20.795 | | | | 26.392 | | | | | |
| -15.386 | | | | 15.266 | | | | -14.107 | | | | | |
| 4.541 | | | | -11.640 | | | | 10.465 | | | | | |
| 20.202 | | | | -12.873 | | | | 18.200 | | | | | |
| 18.634 | | | | -16.693 | | | | 6.407 | | | | | |
| -20.544 | | | | -22.757 | | | | -9.928 | | | | | |
| -25.111 | | | | -15.391 | | | | 2.415 | | | | | |
| -0.174 | | | | -5.843 | | | | -2.854 | | | | | |
| -8.562 | | | | -29.631 | | | | -17.709 | | | | | |
| -27.099 | | | | 12.302 | | | | 12.584 | | | | | |
| -3.643 | | | | -0.082 | | | | -19.490 | | | | | |
| -25.852 | | | | -8.356 | | | | 5.856 | | | | | |
| -2.079 | | | | -23.395 | | | | 0.627 | | | | | |
| 13.829 | | | | 16.094 | | | | -11.153 | | | | | |
| -4.191 | | | | -14.398 | | | | 9.519 | | | | | |
| -15.579 | | | | 14.470 | | | | -3.576 | | | | | |
| 0.596 | | | | 21.464 | | | | -13.752 | | | | | |
| -23.481 | | | | -13.549 | | | | 13.736 | | | | | |
| 8.826 | | | | 16.819 | | | | -24.609 | | | | | |
| -0.226 | | | | -5.477 | | | | -3.140 | | | | | |
| 1.218 | | | | -12.810 | | | | -23.728 | | | | | |

| | | | | | | | | |
|---|---|---|---|---|---|---|---|---|
| -15.284 | | | | -16.103 | | | | -21.909 |
| 25.504 | | | | -9.152 | | | | 27.747 |
| 12.900 | | | | -21.171 | | | | 22.774 |
| -5.992 | | | | 11.942 | | | | -6.643 |
| -10.869 | | | | -25.767 | | | | -5.365 |
| -8.454 | | | | -10.582 | | | | 8.733 |
| -23.567 | | | | 15.094 | | | | -4.019 |
| -24.695 | | | | 30.003 | | | | -19.613 |
| 17.608 | | | | 22.340 | | | | 8.965 |
| 15.128 | | | | 5.547 | | | | 1.907 |
| 9.489 | | | | -16.318 | | | | -4.694 |
| 21.987 | | | | 12.456 | | | | 5.550 |
| -20.902 | | | | -24.063 | | | | -8.755 |
| 3.522 | | | | -22.279 | | | | 17.569 |
| -3.291 | | | | -26.764 | | | | -11.499 |
| 9.827 | | | | -20.857 | | | | 12.633 |
| 17.691 | | | | -21.563 | | | | 0.609 |
| 23.535 | | | | 19.084 | | | | 2.778 |
| -4.154 | | | | -27.599 | | | | -5.830 |
| -9.285 | | | | -23.097 | | | | -16.351 |
| -1.925 | | | | 2.630 | | | | -4.300 |
| -13.015 | | | | -6.840 | | | | -25.204 |
| -20.378 | | | | -13.861 | | | | 12.257 |
| -25.857 | | | | -20.046 | | | | -13.397 |
| -8.621 | | | | 25.002 | | | | -6.626 |
| -19.812 | | | | 15.476 | | | | -21.718 |
| 4.094 | | | | 8.794 | | | | 21.963 |
| -17.100 | | | | 6.345 | | | | -7.632 |
| -11.097 | | | | 20.710 | | | | -18.337 |
| -18.261 | | | | -2.009 | | | | 19.355 |
| -22.465 | | | | -21.174 | | | | 6.791 |
| -12.443 | | | | 20.234 | | | | -5.396 |
| -6.297 | | | | 19.074 | | | | -8.348 |
| -11.250 | | | | 23.035 | | | | -21.282 |
| -20.348 | | | | 21.171 | | | | -9.331 |
| -19.809 | | | | 17.347 | | | | 19.642 |

| | | | | | | | | |
|---|---|---|---|---|---|---|---|---|
| -21.445 | | | | -0.035 | | | | -19.758 |
| | | | | 16.107 | | | | 7.477 |
| | | | | -21.622 | | | | -7.089 |
| | | | | 18.570 | | | | -8.062 |
| | | | | -24.635 | | | | -2.182 |
| | | | | 11.837 | | | | -4.134 |
| | | | | 12.761 | | | | -22.743 |
| | | | | -5.552 | | | | -9.529 |
| | | | | 10.401 | | | | -5.057 |
| | | | | 3.758 | | | | -15.709 |
| | | | | 12.164 | | | | -4.150 |
| | | | | 1.278 | | | | -8.784 |
| | | | | -16.936 | | | | 3.135 |
| | | | | 0.808 | | | | -3.439 |
| | | | | 8.871 | | | | 28.188 |
| | | | | 15.601 | | | | 13.529 |
| | | | | -21.230 | | | | -6.646 |
| | | | | -25.419 | | | | 10.566 |
| | | | | 9.493 | | | | -9.131 |
| | | | | 19.274 | | | | -14.998 |
| | | | | -20.083 | | | | -16.110 |
| | | | | 24.970 | | | | -18.044 |
| | | | | 12.573 | | | | 18.696 |
| | | | | 4.470 | | | | 15.711 |
| | | | | -23.192 | | | | -9.433 |
| | | | | 15.470 | | | | 6.322 |
| | | | | 3.802 | | | | -15.510 |
| | | | | 18.162 | | | | 16.827 |
| | | | | -7.665 | | | | 11.336 |
| | | | | -12.758 | | | | -7.967 |
| | | | | -20.266 | | | | 22.570 |
| | | | | 0.285 | | | | -4.105 |
| | | | | -26.392 | | | | 21.747 |
| | | | | 7.266 | | | | -13.782 |
| | | | | 6.756 | | | | 5.368 |
| | | | | -23.333 | | | | -26.065 |

| | | | | | | | | |
|---|---|---|---|---|---|---|---|---|
| | | | | 21.553 | | | | -22.977 |
| | | | | -22.289 | | | | -24.609 |
| | | | | 9.484 | | | | 9.514 |
| | | | | -24.104 | | | | -17.552 |
| | | | | -17.852 | | | | 7.265 |
| | | | | 16.527 | | | | -15.328 |
| | | | | 8.198 | | | | -7.601 |
| | | | | 16.263 | | | | 18.810 |
| | | | | 9.290 | | | | -11.299 |
| | | | | -23.178 | | | | 20.735 |
| | | | | -12.931 | | | | -4.770 |
| | | | | 23.257 | | | | 28.368 |
| | | | | 18.058 | | | | -5.567 |
| | | | | 0.651 | | | | -18.274 |
| | | | | 19.547 | | | | -16.465 |
| | | | | 21.963 | | | | 22.210 |
| | | | | -24.452 | | | | 17.875 |
| | | | | 16.113 | | | | -13.645 |
| | | | | -26.569 | | | | 3.755 |
| | | | | -22.428 | | | | 16.460 |
| | | | | -8.976 | | | | -12.292 |
| | | | | -25.261 | | | | -18.870 |
| | | | | 10.807 | | | | -16.865 |
| | | | | 10.237 | | | | 23.313 |
| | | | | 14.518 | | | | -18.409 |
| | | | | 7.329 | | | | -6.904 |
| | | | | -7.795 | | | | 22.600 |
| | | | | -23.215 | | | | 2.782 |
| | | | | 12.698 | | | | -7.618 |
| | | | | 2.329 | | | | -23.515 |
| | | | | 15.757 | | | | 26.968 |
| | | | | -25.358 | | | | -17.000 |
| | | | | 6.989 | | | | 19.062 |
| | | | | -17.673 | | | | -11.983 |
| | | | | 10.258 | | | | -11.376 |
| | | | | 18.810 | | | | -9.671 |

| | | | | | | | | |
|---|---|---|---|---|---|---|---|---|
| | | | | 7.733 | | | | -1.843 |
| | | | | 4.857 | | | | -25.267 |
| | | | | 5.378 | | | | -22.761 |
| | | | | 15.918 | | | | 1.034 |
| | | | | 9.555 | | | | -4.901 |
| | | | | -4.055 | | | | 20.509 |
| | | | | -11.665 | | | | -7.530 |
| | | | | -21.352 | | | | 16.003 |
| | | | | 10.538 | | | | -19.227 |
| | | | | -24.790 | | | | 10.841 |
| | | | | 6.691 | | | | -2.424 |
| | | | | -16.896 | | | | -3.794 |
| | | | | 9.580 | | | | 0.589 |
| | | | | 5.530 | | | | -6.448 |
| | | | | -8.573 | | | | -22.297 |
| | | | | 11.558 | | | | 23.766 |
| | | | | -11.909 | | | | -25.185 |
| | | | | -8.099 | | | | -5.873 |
| | | | | 14.799 | | | | -23.008 |
| | | | | 27.773 | | | | 18.911 |
| | | | | -5.723 | | | | -7.447 |
| | | | | -7.820 | | | | 22.646 |
| | | | | -4.511 | | | | 12.080 |
| | | | | -9.099 | | | | -8.276 |
| | | | | -9.838 | | | | -11.876 |
| | | | | 2.494 | | | | -9.394 |
| | | | | 6.741 | | | | -9.777 |
| | | | | -22.012 | | | | 11.544 |
| | | | | 23.478 | | | | -10.592 |
| | | | | 10.173 | | | | -12.802 |
| | | | | 8.289 | | | | -19.107 |
| | | | | -0.078 | | | | 19.728 |
| | | | | 24.106 | | | | -4.050 |
| | | | | 12.265 | | | | -14.420 |
| | | | | 2.459 | | | | -18.245 |
| | | | | -6.868 | | | | -12.196 |

| | | | | | | | | |
|---|---|---|---|---|---|---|---|---|
| | | | | -26.302 | | | | -21.081 |
| | | | | -11.823 | | | | 6.491 |
| | | | | 5.787 | | | | -4.992 |
| | | | | -23.101 | | | | -11.590 |
| | | | | 7.379 | | | | -14.968 |
| | | | | 11.596 | | | | -19.882 |
| | | | | -21.121 | | | | -10.593 |
| | | | | 21.763 | | | | -13.462 |
| | | | | 7.971 | | | | -16.915 |
| | | | | 19.556 | | | | 11.753 |
| | | | | 3.945 | | | | 14.158 |
| | | | | -20.181 | | | | -14.489 |
| | | | | 13.510 | | | | -9.877 |
| | | | | 4.216 | | | | -22.608 |
| | | | | -1.833 | | | | 18.662 |
| | | | | -6.403 | | | | -9.178 |
| | | | | 1.101 | | | | -20.236 |
| | | | | -23.940 | | | | -1.612 |
| | | | | -10.192 | | | | -6.172 |
| | | | | 5.535 | | | | 0.260 |
| | | | | 0.808 | | | | -25.767 |
| | | | | 14.472 | | | | -10.966 |
| | | | | 19.561 | | | | -19.274 |
| | | | | -11.652 | | | | -23.509 |
| | | | | -2.115 | | | | -9.985 |
| | | | | -10.692 | | | | -11.644 |
| | | | | -6.853 | | | | -12.184 |
| | | | | 20.970 | | | | -14.235 |
| | | | | -0.834 | | | | -2.145 |
| | | | | 11.041 | | | | -5.508 |
| | | | | -3.037 | | | | -7.541 |
| | | | | 2.102 | | | | -16.661 |
| | | | | 13.380 | | | | 7.067 |
| | | | | 8.253 | | | | -3.601 |
| | | | | -6.997 | | | | -17.038 |
| | | | | 20.701 | | | | -21.008 |

| | | | | | | | | |
|---|---|---|---|---|---|---|---|---|
| | | | | 2.988 | | | | 8.649 |
| | | | | -12.895 | | | | -23.511 |
| | | | | | | | | 26.621 |
| | | | | | | | | 19.008 |
| | | | | | | | | -9.489 |
| | | | | | | | | 11.557 |
| | | | | | | | | 24.292 |
| | | | | | | | | 21.388 |
| | | | | | | | | -13.481 |
| | | | | | | | | -16.373 |
| | | | | | | | | -20.048 |
| | | | | | | | | -12.440 |
| | | | | | | | | -18.919 |
| | | | | | | | | -22.868 |
| | | | | | | | | -1.829 |
| | | | | | | | | -20.822 |
| | | | | | | | | -21.974 |
| | | | | | | | | -17.843 |
| | | | | | | | | 2.958 |
| | | | | | | | | -24.213 |
| | | | | | | | | -1.050 |
| | | | | | | | | 0.424 |
| | | | | | | | | -17.018 |
| | | | | | | | | -21.426 |
| | | | | | | | | -8.273 |
| | | | | | | | | -11.413 |
| | | | | | | | | -30.292 |
| | | | | | | | | -15.732 |
| | | | | | | | | 5.372 |
| | | | | | | | | 3.744 |
| | | | | | | | | -11.297 |
| | | | | | | | | -13.817 |
| | | | | | | | | -8.888 |
| | | | | | | | | -10.898 |
| | | | | | | | | -12.313 |
| | | | | | | | | -6.205 |

| | | | | | | | | |
|---|---|---|---|---|---|---|---|---|
| | | | | | | | | -22.093 |
| | | | | | | | | 15.251 |
| | | | | | | | | -15.335 |
| | | | | | | | | 17.792 |
| | | | | | | | | -5.478 |
| | | | | | | | | -9.336 |
| | | | | | | | | -13.237 |
| | | | | | | | | 21.821 |
| | | | | | | | | -15.885 |
| | | | | | | | | -19.815 |
| | | | | | | | | 10.837 |
| | | | | | | | | -6.027 |
| | | | | | | | | -12.610 |
| | | | | | | | | -26.125 |
| | | | | | | | | -11.589 |
| | | | | | | | | -28.626 |
| | | | | | | | | -6.969 |
| | | | | | | | | 12.412 |
| | | | | | | | | -5.814 |
| | | | | | | | | -10.623 |
| | | | | | | | | 27.644 |
| | | | | | | | | 25.622 |
| | | | | | | | | -6.567 |
| | | | | | | | | -8.069 |
| | | | | | | | | -16.234 |
| | | | | | | | | -25.014 |
| | | | | | | | | -10.069 |
| | | | | | | | | -11.040 |
| | | | | | | | | -11.404 |
| | | | | | | | | -24.304 |
| | | | | | | | | -14.060 |
| | | | | | | | | -19.024 |
| | | | | | | | | 20.112 |
| | | | | | | | | -11.724 |
| | | | | | | | | -10.385 |
| | | | | | | | | -18.764 |

| | | | | | | | | |
|---|---|---|---|---|---|---|---|---|
| | | | | | | | | 10.964 |
| | | | | | | | | -9.197 |
| | | | | | | | | 3.276 |
| | | | | | | | | 4.549 |
| | | | | | | | | 0.345 |
| | | | | | | | | -1.925 |
| | | | | | | | | -7.713 |
| | | | | | | | | 2.227 |
| | | | | | | | | 21.265 |
| | | | | | | | | 1.399 |
| | | | | | | | | -9.972 |
| | | | | | | | | 7.007 |
| | | | | | | | | 23.102 |
| | | | | | | | | -8.073 |
| | | | | | | | | -3.247 |
| | | | | | | | | -9.188 |
| | | | | | | | | -9.866 |
| | | | | | | | | 4.901 |
| | | | | | | | | 15.028 |
| | | | | | | | | 4.336 |
| | | | | | | | | 8.065 |
| | | | | | | | | -7.306 |
| | | | | | | | | 1.825 |
| | | | | | | | | -8.907 |
| | | | | | | | | 0.801 |
| | | | | | | | | 8.394 |
| | | | | | | | | -21.536 |
| | | | | | | | | 9.455 |
| | | | | | | | | 4.407 |
| | | | | | | | | -0.548 |
| | | | | | | | | -13.684 |
| | | | | | | | | -14.525 |
| | | | | | | | | -16.107 |
| | | | | | | | | -8.235 |

**FIGURE-3B-DOSE-RESPONSE-CURVE (units = mm)**

| Test compound | Concentration (mM) | Physiological condition | Mean difference (mm) | CI 5 | CI 95 | Error high | Error low |
|---|---|---|---|---|---|---|---|
| 2M1B | 0.003 | Fed | 0.119 | -0.960 | 1.190 | 1.071 | 1.079 |
| 2M1B | 0.01 | Fed | -0.973 | -2.099 | 0.095 | 1.069 | 1.126 |
| 2M1B | 0.03 | Fed | -0.551 | -1.657 | 0.534 | 1.085 | 1.106 |
| 2M1B | 0.1 | Fed | 1.536 | 0.436 | 2.635 | 1.099 | 1.100 |
| 2M1B | 0.3 | Fed | 0.275 | -0.768 | 1.343 | 1.068 | 1.043 |
| 2M1B | 1 | Fed | 0.374 | -0.771 | 1.492 | 1.118 | 1.146 |
| 2M1B | 3 | Fed | 4.872 | 3.881 | 5.913 | 1.041 | 0.991 |
| 2M1B | 10 | Fed | 8.133 | 7.144 | 9.230 | 1.096 | 0.989 |
| 2M1B | 30 | Fed | 8.674 | 7.627 | 9.648 | 0.974 | 1.048 |
| 2M1B | 100 | Fed | 12.190 | 11.211 | 13.151 | 0.960 | 0.980 |
| 2M1B | 300 | Fed | 13.181 | 12.205 | 14.135 | 0.954 | 0.976 |
| 2M1B | 1000 | Fed | 12.018 | 11.039 | 13.038 | 1.019 | 0.979 |
| 2M1B | 3000 | Fed | 9.086 | 8.001 | 10.107 | 1.021 | 1.085 |
| 2M1B | 0.003 | Starved | 0.280 | -0.492 | 1.069 | 0.789 | 0.772 |
| 2M1B | 0.01 | Starved | 1.023 | 0.164 | 1.884 | 0.861 | 0.859 |
| 2M1B | 0.03 | Starved | -1.094 | -1.972 | -0.222 | 0.872 | 0.877 |
| 2M1B | 0.1 | Starved | -0.066 | -0.823 | 0.725 | 0.792 | 0.757 |
| 2M1B | 0.3 | Starved | 1.286 | 0.434 | 2.166 | 0.879 | 0.852 |
| 2M1B | 1 | Starved | -0.651 | -1.536 | 0.284 | 0.935 | 0.885 |
| 2M1B | 3 | Starved | -0.488 | -1.390 | 0.453 | 0.941 | 0.903 |
| 2M1B | 10 | Starved | 0.587 | -0.160 | 1.365 | 0.778 | 0.747 |
| 2M1B | 30 | Starved | -0.650 | -1.438 | 0.142 | 0.792 | 0.788 |
| 2M1B | 100 | Starved | 2.548 | 1.660 | 3.418 | 0.870 | 0.888 |
| 2M1B | 300 | Starved | 3.421 | 2.476 | 4.362 | 0.940 | 0.945 |
| 2M1B | 1000 | Starved | 8.438 | 7.455 | 9.347 | 0.908 | 0.983 |
| 2M1B | 3000 | Starved | 4.109 | 3.343 | 4.887 | 0.778 | 0.766 |

**FIGURE-4-MEAN-DIFFERENCE-CONFIDENCE INTERVALS (units = mm)**

| Strain | Physiological condition | Test compound | Mean difference (mm) | CI 5 | CI 95 |
|---|---|---|---|---|---|
| N2 | Fed | I1 | 0.836 | 0.012 | 1.640 |
| N2 | Fed | I2 | -4.450 | -5.288 | -3.627 |
| N2 | Fed | I3 | 2.283 | 1.371 | 3.117 |
| N2 | Fed | I4 | 12.984 | 12.139 | 13.809 |
| N2 | Fed | M1 | 0.156 | -0.752 | 1.061 |
| N2 | Fed | M2 | 1.030 | 0.176 | 1.892 |
| N2 | Fed | M3 | 0.989 | 0.141 | 1.908 |
| N2 | Fed | M4 | 11.607 | 10.798 | 12.418 |
| CX10 | Fed | I1 | 4.664 | 3.909 | 5.430 |
| CX10 | Fed | I2 | 4.554 | 3.773 | 5.350 |
| CX10 | Fed | I3 | 8.845 | 8.055 | 9.622 |
| CX10 | Fed | I4 | 16.219 | 15.538 | 16.901 |
| CX10 | Fed | M1 | 4.032 | 3.253 | 4.762 |
| CX10 | Fed | M2 | 6.873 | 6.120 | 7.652 |
| CX10 | Fed | M3 | 7.065 | 6.320 | 7.803 |
| CX10 | Fed | M4 | 16.372 | 15.740 | 17.018 |
| PR678 | Fed | I1 | -0.019 | -0.660 | 0.641 |
| PR678 | Fed | I2 | -0.077 | -0.742 | 0.587 |
| PR678 | Fed | I3 | -0.348 | -0.965 | 0.345 |
| PR678 | Fed | I4 | 0.225 | -0.432 | 0.876 |
| PR678 | Fed | M1 | 0.458 | -0.193 | 1.114 |
| PR678 | Fed | M2 | 0.856 | 0.168 | 1.554 |
| PR678 | Fed | M3 | -0.274 | -0.923 | 0.384 |
| PR678 | Fed | M4 | -0.561 | -1.178 | 0.097 |
| GN1077 | Fed | I1 | 0.155 | -0.694 | 0.997 |
| GN1077 | Fed | I2 | 0.070 | -0.811 | 0.984 |
| GN1077 | Fed | I3 | -0.565 | -1.417 | 0.343 |
| GN1077 | Fed | I4 | -0.263 | -1.179 | 0.654 |
| GN1077 | Fed | M1 | -0.250 | -1.168 | 0.653 |
| GN1077 | Fed | M2 | 0.083 | -0.804 | 0.958 |
| GN1077 | Fed | M3 | 0.086 | -0.751 | 1.011 |
| GN1077 | Fed | M4 | -0.381 | -1.217 | 0.498 |
| N2 | Starved | I1 | -0.907 | -1.720 | -0.104 |
| N2 | Starved | I2 | -7.828 | -8.601 | -7.070 |
| N2 | Starved | I3 | -0.754 | -1.516 | 0.004 |
| N2 | Starved | I4 | 2.436 | 1.660 | 3.217 |
| N2 | Starved | M1 | -0.202 | -1.015 | 0.619 |
| N2 | Starved | M2 | 1.542 | 0.732 | 2.362 |
| N2 | Starved | M3 | -0.804 | -1.596 | 0.017 |
| N2 | Starved | M4 | -1.257 | -2.059 | -0.463 |
| CX10 | Starved | I1 | -0.341 | -1.323 | 0.729 |
| CX10 | Starved | I2 | 0.041 | -1.002 | 1.085 |
| CX10 | Starved | I3 | 1.666 | 0.708 | 2.659 |
| CX10 | Starved | I4 | 8.396 | 7.385 | 9.461 |

| | | | | | |
|---|---|---|---|---|---|
| CX10 | Starved | M1 | 0.269 | -0.738 | 1.213 |
| CX10 | Starved | M2 | 2.764 | 1.764 | 3.737 |
| CX10 | Starved | M3 | 1.299 | 0.291 | 2.285 |
| CX10 | Starved | M4 | 3.249 | 2.188 | 4.241 |
| PR678 | Starved | I1 | -1.245 | -2.404 | -0.148 |
| PR678 | Starved | I2 | -4.158 | -5.186 | -3.148 |
| PR678 | Starved | I3 | -0.710 | -1.782 | 0.359 |
| PR678 | Starved | I4 | 0.910 | -0.188 | 2.036 |
| PR678 | Starved | M1 | -0.412 | -1.557 | 0.710 |
| PR678 | Starved | M2 | 0.946 | -0.233 | 2.048 |
| PR678 | Starved | M3 | -0.935 | -2.001 | 0.124 |
| PR678 | Starved | M4 | -1.164 | -2.211 | -0.018 |
| GN1077 | Starved | I1 | -0.774 | -1.482 | -0.078 |
| GN1077 | Starved | I2 | 0.846 | 0.166 | 1.571 |
| GN1077 | Starved | I3 | -0.565 | -1.316 | 0.148 |
| GN1077 | Starved | I4 | 0.011 | -0.738 | 0.728 |
| GN1077 | Starved | M1 | 0.232 | -0.511 | 0.976 |
| GN1077 | Starved | M2 | -0.462 | -1.176 | 0.291 |
| GN1077 | Starved | M3 | -0.564 | -1.273 | 0.197 |
| GN1077 | Starved | M4 | -0.635 | -1.380 | 0.072 |
